# Prototypic SNARE proteins are encoded in the genomes of Heimdallarchaeota, potentially bridging the gap between the prokaryotes and eukaryotes

**DOI:** 10.1101/810531

**Authors:** Emilie Neveu, Dany Khalifeh, Nicolas Salamin, Dirk Fasshauer

## Abstract

A defining feature of eukaryotic cells is the presence of numerous membrane-bound organelles that subdivide the intracellular space into distinct compartments. How the eukaryotic cell acquired its internal complexity is still poorly understood. Material exchange among most organelles occurs via vesicles that bud off from a source and specifically fuse with a target compartment. Central players in the vesicle fusion process are the <u>S</u>oluble <u>*N*</u>-ethylmaleimide-sensitive factor <u>A</u>ttachment protein <u>RE</u>ceptor (SNARE) proteins. These small tail-anchored (TA) membrane proteins zipper into elongated four-helix bundles that pull membranes together^1–3^. SNARE proteins are highly conserved among eukaryotes but are thought to be absent in prokaryotes. Here, we identified SNARE-like factors in the genomes of uncultured organisms of Asgard archaea of the *Heimdallarchaeota* clade^4,5^, which are thought to be the closest living relatives of eukaryotes. Biochemical experiments show that the archaeal SNARE-like proteins can interact with eukaryotic SNARE proteins. We did not detect SNAREs in α-proteobacteria, the closest relatives of mitochondria, but identified several genes encoding for SNARE proteins in γ-proteobacteria of the order Legionellales, pathogens that live inside eukaryotic cells. Very probably, their SNAREs stem from lateral gene transfer from eukaryotes. Together, this suggests that the diverse set of eukaryotic SNAREs evolved from an archaeal precursor. However, whether *Heimdallarchaeota* actually have a simplified endomembrane system will only be seen when we succeed studying these organisms under the microscope.

---

All SNARE proteins share an evolutionary conserved α-helical stretch ~ 60 amino acids long – the so-called SNARE-motif – whereas their *N*-terminal regulatory domains can have different folds. In most SNAREs, the SNARE motif is *C*-terminally anchored via a single-pass transmembrane domain (TMD) and is facing the cytosol. *N*- to *C*-terminal assembly into a tight parallel four-helix bundle SNARE complex^6^ between apposing membranes thus pulls the membranes together^1–3^. In the core of the SNARE bundle, 16 layers of mostly hydrophobic residues interact tightly^6^. The central ionic “0-layer”, consisting of three glutamine (Q) residues and one arginine (R) residue, is almost unchanged throughout the SNARE protein family^7^. Four basic types, namely Qa-, Qb-, Qc-, and R-SNAREs, can be distinguished by their sequence profiles, reflecting their position in the heterologous four-helix bundle^8^. The structures of different SNARE complexes reveal that the four helices have distinctive features, particularly Qa- and R-SNAREs, which interlock^6,9^. This feature renders QabcR-SNARE complexes stable and may prevent other combinations. Among the four basic SNARE types, about 20 subtypes can be distinguished^8^. They assemble into distinct QabcR units that work in different trafficking steps and probably represent the original repertoire of the Last Eukaryotic Common Ancestor (LECA)^8^, which was a fairly sophisticated cell with a nucleus, peroxisomes, mitochondria, and probably all compartments of the endomembrane system^10,11^. The diverse SNARE set of the LECA and, by extension, of all present-day eukaryotes can be traced back to a single QabcR complex that was multiplied about 2 billion years ago. The different vesicle fusion machines then adapted to different intracellular trafficking steps^1,8,10^. Moreover, as all four basic SNARE types are related, it is conceivable that the first QabcR unit arose by gene duplication from one common SNARE ancestor as a prototypic SNARE protein assembled into homomeric bundles. But do such prototypic SNARE proteins still exist?

In order to search for prototypic SNARE proteins, we scanned the NCBI protein databases for archaea and bacteria (collectively known as prokaryotes). For this, we took advantage of Hidden Markov Model (HMM) profiles trained previously^8^ to classify the SNARE motifs of eukaryotic SNARE proteins. We implemented a 1E^−4^ expectation value cutoff and kept only the sequences for which the target motif was at least 40 amino acids long to minimize false positive results. Around 5,000 prokaryotic sequences met these criteria (see Methods and Supplementary Information, Section 1 for details). For an overview of the relationships among the collected prokaryotic sequences, we clustered them with the Basic Local Alignment Search Tool (BLAST) to construct groups of similar factors for further inspection (Methods). The sequences split into 96 different clusters of different sizes from a large cluster with ~ 4,200 sequences to clusters with only two sequences; 178 sequences remained isolated (Supplementary Information, Section 1). We then restricted the search to tail-anchored (TA) proteins, which are a subset of membrane proteins (~ 5%) that play important membrane-active roles in eukaryotes, these include the SNARE proteins^12^. TA proteins have been reported in all domains of life, although their number is usually restricted to about a dozen in most prokaryotes^13,14^, while eukaryotes can have several hundreds^15,16^.

Altogether, the collection of prokaryotic SNARE-like sequences found by our bioinformatic screen contained only 20 candidate sequences for TA protein (Methods). The sequences were contained in smaller clusters and as singletons and had only moderate e-values (Supplementary Information, Section 1). Cluster 8 contained sequences from several closely related species of the genus *Variovorax* (γ-proteobacteria). These sequences were about 10 residues too short for a SNARE motif. Cluster 40, containing TA proteins from the genus *Methylobacterium* (α-proteobacteria), was even less enticing, as it was too short and lacked a central glutamine. By contrast, Cluster 28 containing a pair of related sequences (OLS22354.1 and PWI47941.1) from two metagenomes of the Asgard group, *Heimdallarchaeota archaeon LC_2* ^5^ and *B3-JM-08* ^17^, looked like promising candidates. Both sequences possess an entire SNARE motif and a central glutamine residue within the motif and the motif was connected to a TMD via a short linker. Most strikingly, both sequence’s *N*-terminal regions are predicted to contain α-helices that may fold into a three-helix bundle, a structural feature found in the vast majority of eukaryotic Q-SNARE subtypes (Extended Data Fig. 1; Supplementary Information, Section 2). Overall, these two Asgard sequences posed the good prototypic prokaryotic SNARE protein candidates uncovered. This observation was intriguing, as this archaeal lineages is considered to be the closest extant relative of eukaryotes^5,17,18^ and the existence of functional SNARE proteins within this lineage would provide fascinating insights into the origin of intracellular trafficking and eukaryotes in general. However, do these Asgard genes really encode SNARE proteins?

In order to validate this initial bioinformatic assessment, we tested the ability of these Asgard proteins to interact with eukaryotic SNARE proteins. We recombinantly expressed the SNARE-like region of PWI47941.1 from *H. B3-JM-08* and mixed it in different combinations with the neuronal SNARE proteins Syx1 (Qa-), SNAP-25 (Qbc-), or Syb2 (R-SNARE). These well-characterized neuronal SNARE proteins assemble during synaptic secretion^1^. Assembly can be monitored by native gel electrophoresis^19,20^, a technique that allows for the separation of the individual proteins from complexes, provided that the latter are of sufficient stability (Methods). Another reason for choosing this particular SNARE unit was because it has been shown that its subunits can also interact non-specifically with SNARE proteins working in other trafficking steps^21^, thus optimizing our chances of observing a binding event with evolutionary distant SNARE proteins.

When the *H. B3-JM-08* protein, termed HPS1 for <u>H</u>eimdallarchaeota <u>P</u>rototypic <u>S</u>NARE, was mixed with individual neuronal SNARE proteins, no new band indicative of a stable interaction appeared in native gel electrophoresis. However, when HPS1 was mixed with Syx1 and SNAP-25, a new band appeared on top of the separating gel, suggesting that they formed a ternary complex (Extended Data Fig. 2), in which HPS1 may have taken over the position of the R-SNARE. Additional titration experiments indicated, that this HPS1 complex was not very stable (Supplementary Information, Section 2, Fig. S2.3.).

In the next set of binding experiments, the two helices of the Qbc-SNARE SNAP-25 were used as independent constructs, a Qb-(SN1) and a Qc-helix (SN2). We mixed HPS1 with each of the individual SNARE proteins and in different combinations. Supporting our previous observation, new bands appeared on top of the gel when HPS1 was mixed with (i) Syx1 and SN1 (ii) Syx1, SN1, and SN2 (iii) Syx1, SN1, and Syb2; and (iv) Syx1, SN2, and Syb2, although the latter was less prominent (Fig. 1). As the new bands containing HPS1 could not be distinguished on the native gel, further experiments were carried out in which we preformed stable complexes of the neuronal SNAREs Syx1, SN1, and Syb2 and Syx1, SN2, and Syb2. Upon addition of HPS1, both complexes were transformed into complexes containing HPS1 (Extended Data Fig. 3). These observations were supported by size exclusion experiments (Methods, Supplementary Information, Section 2, Fig. S2.5). All in all, these experiments suggested that HPS1 can take over different helix positions when it interacts with neuronal SNARE proteins: apparently, it can act as a R-, Qb-, or Qc-helix. Not surprisingly, the different HPS1-complex were less stable than the quaternary neuronal SNARE complex, which we observed by addition of the matching neuronal SNARE.

**Fig. 1:**
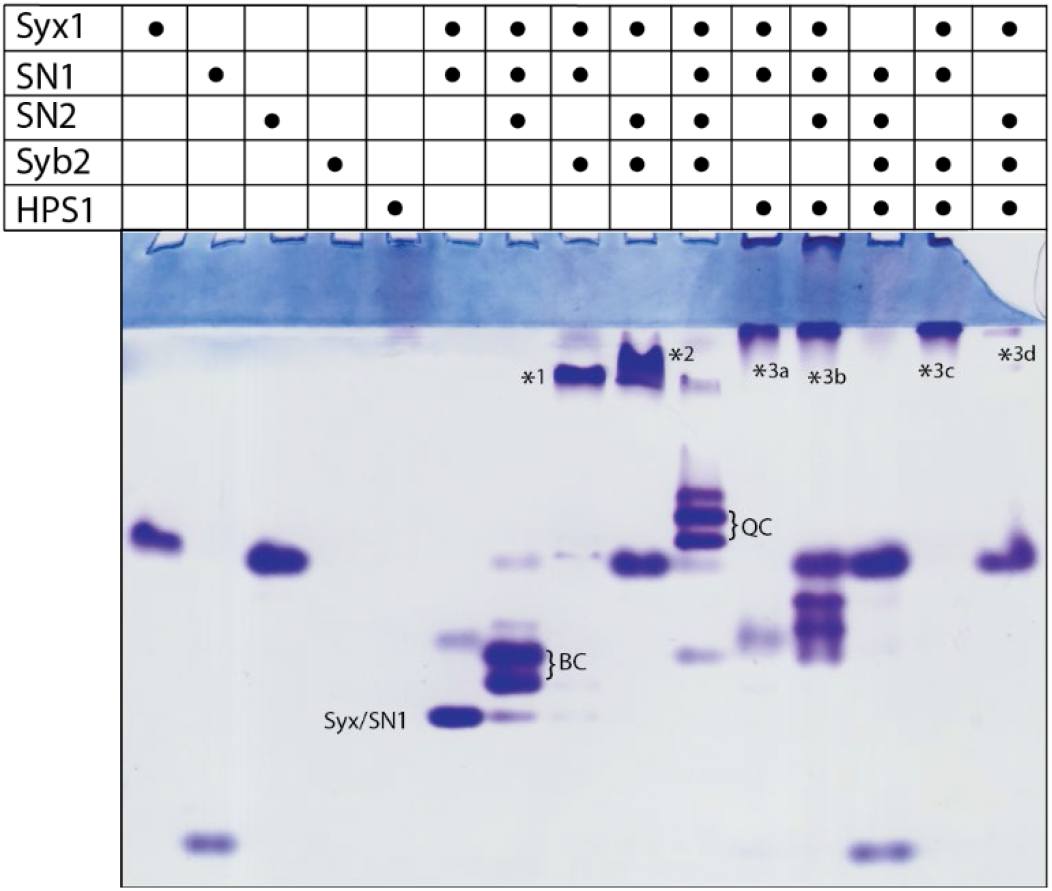
Interaction pattern of HPS1 with neuronal SNARE proteins by nondenaturing gel electrophoresis. The SNARE-like region of PWI47941.1 from *Heimdallarchaeota B3-JM-08*, termed HPS1 (aa 159–211), was mixed in different combinations with the neuronal SNARE proteins Syx1 (Qa-), Syb2 (R-), and the two SNARE motifs of SNAP-25, SN1 (Qb-) and SN2 (Qc-SNARE), which were used as independent proteins. The proteins were incubated overnight at 4°C with equimolar ratios at ~15 µM concentration prior to nondenaturing gel electrophoresis, which separates native proteins and stable interactions by their charge only. Proteins were visualized by Coomassie Blue staining. Note that Syb2 and HPS1 are not detectable in the non-denaturing gel (Lanes 4 & 5) because of their isoelectric points of 8.5 and 8.1, respectively. As shown before, the neuronal SNAREs assemble into a Syx1–SN1 complex, a complex consisting of Syx– SN1–SN2 (BC), a Syx1–SN1–Syb2 complex (*1), a Syx1–SN2–Syb2 complex (*2), and a quaternary complex (QC)^20^. New complex bands appeared when HPS1 was added to (i) Syx1 and SN1 (*3a); (ii) Syx1, SN1, and SN2 (*3b); (iii) Syx1, SN1, and Syb2 (*3c); and (iv) Syx1, SN2, and Syb2 (*3d). All new bands containing HPS1 ran on top of the gel. Further analyses of these HPS1 complexes are shown in Extended Data Fig. 3.

We performed similar mixing experiments with the SNARE-like region of OLS22354.1 (HPS2) from *H. LC_2*. HPS2 was able to interact as well but, in contrast to the somewhat promiscuous HPS1, HPS2 formed a stable complex only in combination with two SNAREs, Syx1 and Syb2 (Extended Data Fig. 4). The biochemical experiments suggest that the Asgard SNARE-like proteins are able to interact with a set of eukaryotic SNARE proteins that normally assemble into a stable complex to drive secretion of synaptic vesicles in animals. Given the fact that the Asgard proteins lack the distinctive sequence features of the four different eukaryotic SNARE types and in line with the idea that they might be ancestral, it is not surprising that the Asgard proteins are less selective than eukaryotic proteins for the position in the bundle. The lack of distinctive sequence features provides an explanation why the non-cognate complexes containing Asgard proteins are less stable than the cognate neuronal SNARE complex.

As we found only one prototypic SNARE protein each in the two *Heimdallarchaeota* genomes, we scrutinized their genomes for other TA proteins that might constitute other SNARE protein candidates but which had evaded our initial HMM screen. We uncovered 11 TA protein candidates in *H. B3-JM-08* and we found 33 in *H. LC_2*, including the already described SNARE-like sequences (Supplementary Information, Section 3). However, none of the other putative TA proteins from these metagenomes resembled a SNARE protein. We also searched for TA proteins in other relevant genomes. In *H. LC_3* ^5^, we found a larger repertoire of 41 TA proteins, one of which (OLS23218.1) turned out to be homologous to HPS1 and HPS2 but had been less well recognized by our HMMs (Extended Data Table 1). The relatively large number of different TA proteins in the genomes *H. LC_3* and *LC_2* compared with the genomes of other prokaryotes was surprising. It seems that TA proteins, which play membrane-active roles in eukaryotes, may have moved to the fore in *Heimdallarchaeota*. It is conceivable that the expanded repertoire of TA proteins in deep-branching Asgard archaea might have served as evolutionary breeding ground for the development of the endomembrane system in eukaryotes.

Further sequence searches uncovered an additional archaeal SNARE-like sequence in the NCBI nr database: RMG36020.1 from an unclassified *Euryarchaeota* archaeon ^22^ (Extended Data Table 1). The latter sequence is most closely related to the one from *H. LC_2* (48% identity), suggesting that the sequence belongs to a species belonging to the *Heimdallarchaeota* clade. Overall, the four prototypic SNARE proteins display ~ 27– 48% identity, whereas their SNARE motifs show a higher degree (~ 36 – 66% identity). The SNARE motifs of Asgard proteins displayed only low-level identity to eukaryotic SNAREs. For example, the highest identity in human SNAREs was achieved by the Qc-SNARE syntaxin 6 (20 – 28%). We did not find comparable SNARE-like sequences in other Asgard archaea, suggesting that prototypic SNARE proteins only emerged within the *Heimdallarchaeota* lineage or have been lost in other Asgard lineages. However, we cannot exclude that such sequences may exist in other Asgard archaea but are too divergent for us to detect via this analysis. Another possibility is that they are still missing because the metagenomes of Asgard archaea may not be fully reconstructed. It should also be noted that other factors have been reported to have a dispersed phylogenetic distribution within the Asgard clade. For example, the ATPase Get3, which is at the heart of the "Guided Entry of TA proteins" (GET) pathway and inserts TA proteins into the membrane through a conformational change^23,24^, is conserved across eukaryotes and is also present in archaea^25–27^, including *Heimdallarchaeota* (Supplementary Information, Section 3.3).

In a phylogenetic tree (Methods), the SNARE-like sequences from *Heimdallarchaeota* branched outside the four major types of eukaryotic SNAREs (Fig. 2), supporting the idea that the Asgard proteins are likely to constitute prototypic SNARE proteins that are related to the last common ancestor of SNARE proteins, from which the four basic eukaryotic SNARE types arose.

**Fig. 2:**
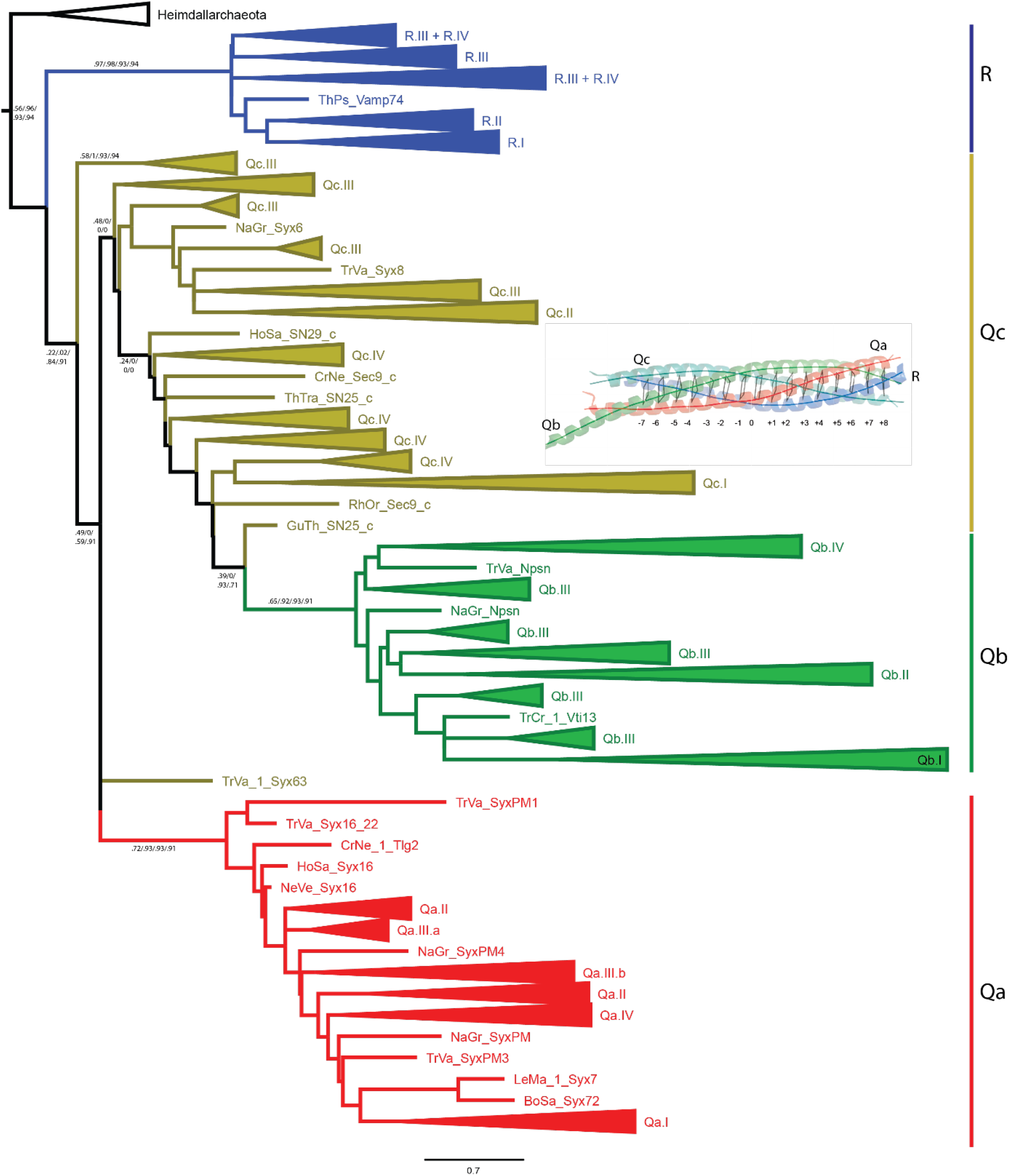
Phylogenetic analysis of eukaryotic and archaeal SNARE proteins. To gain insights into the evolutionary relationship between eukaryotic SNAREs and Asgard SNARE-like proteins, we constructed a maximum likelihood tree from typical SNAREs of 24 representative eukaryotic species, which cover the diversity of the eukaryotic domain (Extended Data Table 3), and from four SNARE-like sequences from the Heimdallarchaeota clade. In the tree, the four Asgard SNARE-like sequences form a clade that is separate from eukaryotic SNAREs, indicating that they are genuinely derived from Asgard metagenomes and are not the result of eukaryotic contamination. The eukaryotic SNAREs split, as shown previously, into four main groups, Qa-, Qb-, Qc-, and R-SNAREs, which correspond to the four different positions in the four-helix bundle SNARE complexes^8^ (shown schematically as an inset). This suggests that eukaryotic SNAREs co-evolved into four interlocking types from a prototypic common ancestor. Despite the relatively short sequence alignment (53 amino acids), the eukaryotic Qa- and R-SNARE groups are relatively well supported, whereas the Qb- and the Qc-groups are more divergent, since their sequences features are less conserved, as can be seen in a Weblogo representation in Extended Data Figure 1. Among the four main SNARE types, conserved subtypes can be distinguished. Generally, SNARE subtypes involved in ER- (Type I) and Golgi-trafficking (Type II) are more conserved across eukaryotes than SNAREs involved in endosomal trafficking (Type III) and secretion (Type IV). Conserved SNAREs are shown as collapsed clades. The remaining branches are labeled by a four letter species identifier (the first two letters from the genus and species names)^8^. Abbreviations of the species names are given in Extended Data Table 3; see also our public <u>SNARE database</u>. The entire tree, without collapsed clades, is given in Supplementary Information, Section 6. Statistical support values (likelihood-mapping, IQ-TREE support, RAxML support, PhyML support) are given at selected inner edges. The scale bar indicates the number of substitutions per site. Note that we constructed additional trees for the complete SNARE set of each of the 24 representative species (Extended Data Table 1) together with the four archaeal sequences (Supplementary Information, Section 7). All individual phylogenetic trees and the corresponding sequence alignments are available at DOI 10.5281/zenodo.3478558.

In the next step, we turned our attention to the other best-scoring sequences we identified with our bioinformatical analysis. From previous studies^8^, we inferred that the separation of true and false positives were of high confidence at expectation values below about 1E^−10^ (see also Supplementary Information, Section 1, Fig. S1.2). Most of these high scoring sequences were found in distinct small clusters, though others stayed isolated. Usually, these sequences possessed an entire SNARE motif. A closer inspection revealed that more than 30 sequences were detected with high confidence by different HMMs (i.e., they represent different types of SNARE proteins (Extended Data Table 2)). For example, three clusters (and one isolated sequence) contained R-SNAREs, one cluster contained a Qc-SNARE, and two clusters contained SNAP-25-like proteins. In phylogenetic analyses, the bacterial SNARE proteins were located on different branches in the SNARE tree (Extended Data Fig. 5), corroborating the idea that they represent different types of SNARE proteins.

We then tested the ability of bacterial SNARE proteins to form a SNARE complex. For this, we recombinantly expressed and purified two representative bacterial proteins, a SNAP-25 like and an R-SNARE, for binding experiments with Syb2, Syx1, and SNAP-25 as described in the previous section. When we mixed stoichiometric amounts of a SNAP-25 like SNARE from *Legionella cherrii* with constructs encompassing the SNARE domains of Syx1 and Syb2, a new band corresponding to a stable ternary complex appeared, confirming that the protein is able to form a SNARE complex. Comparable results were obtained for the bacterial R-SNARE (Fig. 3).

**Fig. 3:**
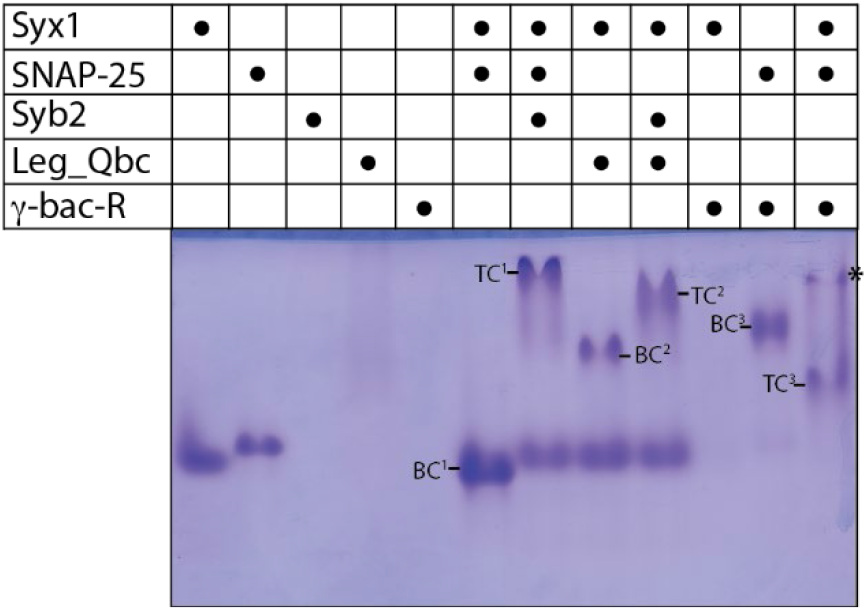
Formation of stable complexes between neuronal SNARE proteins and SNARE proteins from different γ-proteobacteria of the order Legionellales analyzed by non-denaturing gel electrophoresis. A SNAP-25 like SNARE protein from *Legionella cherrii* (Leg_Qbc, aa 1–209, WP_028380397.1) or an R-SNARE from *Gammaproteobacteria bacterium RIFCSPHIGHO2_12_FULL_42_13* (μ-bac-R, aa 116–176, OGT53257.1) was mixed with the neuronal SNARE proteins synaptobrevin (Syb2, R-SNARE), SNAP-25 (Qbc-SNARE), and the SNARE motif of syntaxin (Syx1, Qa-SNARE) as indicated. All incubations were performed as described in the legend of Fig. 1. Proteins were visualized by Coomassie Blue staining. Note that the R-SNAREs Syb2 and μ-bac-R are not detectable in the nondenaturing gel (Lanes 3 & 5) because their isoelectric points are 8.5 and 9.4, respectively. As shown before, the neuronal SNAREs Syx1 and SNAP-25 assemble into a binary complex (BC^2^) and the mixture of all three forms a very stable ternary SNARE complex (TC^1^) ^73^. Comparably, Leg_Qbc forms a binary complex with Syx1 (BC^2^) and a ternary complex with Syb2 and Syx1 (TC^2^). μ-bac-R forms a binary complex with SNAP-25 (BC^3^) and a ternary complex with SNAP-25 and Syx (TC^3^). Note that the corresponding SDS-PAGE is shown in (Supplementary Information, Section 2, Fig. S4.2).

The overwhelming majority of high scoring sequences were from γ-proteobacteria of the order Legionellales, bacterial pathogens that live inside eukaryotic cells ^28,29^. Note that the sequences described here do not mimic SNARE proteins, as has been described for some intracellular bacteria in previous studies^30,31^, but are *bona fide* SNARE proteins. One of the *bona fide* SNARE sequences, a Qc-SNARE protein, was expressed by *Legionella pneumophila* and has been described before^32^. The authors of that study noted that this Qc-SNARE protein, termed LseA, does not have a transmembrane domain but instead features a *C*-terminal CAAX motif^33^, which is farnesylated after it has been translocated by the bacterial Dot/Icm Type IV secretion system into the host cell’s cytosol. Remarkably, most of the bacterial SNAREs discovered by us, although they are of different types, also possess a *C*-terminal CAAX motif, suggesting that they also obtain a post-translational membrane anchor. In fact, only three of the *bona fide* SNAREs from Legionellales matched the general domain layout of eukaryotic SNARE proteins, as a *C*-terminal TMD was found directly adjacent to the SNARE motif. However, the TMD is followed directly, in a hairpin-like fashion, by another TMD, exposing the *C* terminus to the cytosol; this explains why these proteins were not found in our screen for TA proteins.

Most of the Legionellales bacteria have only one SNARE protein. A notable exception is *Berkiella cookevillensis*^34^, whose metagenome codes for three different SNARE proteins, an R-, a Qbc-, and a Qa-SNARE, which could, in principle, assemble into one SNARE complex. A more detailed description of the *bona fide* SNARE proteins in the genomes of Legionellales is given in Supplementary Information, Section 4. Note that other, less-well scoring sequences found are described in Supplementary Information, Section 5. The role the bacterial SNARE proteins play in the life cycle of these pathogenic intracellular bacteriaIt needs to be further investigated. Taken together, our data strongly suggest that the SNARE proteins of Legionellales have been acquired by lateral gene transfer of different SNARE genes from their eukaryotic hosts and thus do not represent the source of the eukaryotic SNARE repertoire.

The chimeric nature of the eukaryotic genome indicates that eukaryotes originated from a merger of an archaeal host with at least one bacterial lineage^35^. A readily traceable event is the entry of an α-proteobacterium that evolved into power-producing organelles, the mitochondria, inside the host cell^36–38^. Whether the α-proteobacterium entered the archaeal host early or late is currently being intensely debated (e.g. ^39,40^). During this process many bacterial genes were transferred to the host’s genome. Nevertheless, it is clear that the eukaryotic genome is more than the sum of two prokaryotic genomes, since many subsequent key inventions facilitated the organizational and structural complexity of eukaryote cells ^41,42^. Until recently, it seemed that their cellular complexity had emerged suddenly, as no life form has been discovered that represents an intermediate between eukaryotes and their more simply built prokaryotic ancestors. Evidently, the discovery of Asgard archaea marks a breakthrough in the quest for the elusive intermediate, as their genomes contain some key building blocks formerly considered to be eukaryotic inventions, suggesting that this lineage gave rise to eukaryotes^4,10,43,44^. The eukaryote signature proteins in the Asgard clade include factors that are involved in membrane trafficking such as an expanded range of small GTPases, ESCRT complexes, putative vesicle coat protein homologs and TRAPP proteins^5,18^. Together, these points indicate that an ancestral endomembrane system probably exists in Asgard archaea.

Currently, no cellular investigations can corroborate this bold idea of an endomembrane system within Asgard archaea, as all the information stems from samples unearthed from deep-sea marine sediments and other forbidding places. Initial steps towards culturing these organisms have been taken. In one study, different *Lokiarchaeota* and *Heimdallarchaeota* have been visualized from environmental samples by fluorescence *in situ* hybridization. It has been observed that though *Lokiarchaeota* can have very different shapes, *Heimdallarchaeota* displayed a more uniform cellular morphology with an interesting central DNA localization^45^. In another study, a *Lokiarchaeota*-related Asgard archaeon, termed *Candidatus Prometheoarchaeum syntrophicum* strain MK-D1, was isolated in a decade-long effort from deep marine sediments and studied in culture^46^. Electron microscopy revealed that *Candidatus Prometheoarchaeum syntrophicum* strain MK-D1 does not possess intracellular organelles, but can produce outer membrane vesicles as well as several long extensions of the cellular membrane that might serve in the exchange with symbiotic partners^46^. Probably, it will take less time to isolate and study the next Asgard archaeons *in vivo*.

As no archaeon of the *Heimdallarchaeota* clade has been cultured yet, we still have to rely on functional and structural investigations of their eukaryote signature proteins. For example, a recent study has elegantly shown that eukaryotic-like profilins of Asgard archaea can regulate actin polymerization, suggesting that they possess a dynamic cytoskeleton^47^. Likewise, our biochemical investigations indicate that eukaryotic-like SNARE proteins of Asgard archaeons can assemble into SNARE complexes, suggesting that these proteins could serve as a simple mechanism for fusing membranes by assembling them into a homomeric complex between two membranes. However, this finding does not rule out that the archaeal SNARE-like proteins might have a somewhat different cellular role, such as tethering and bending membranes. Indeed, it needs to be borne in mind that homology alone does not necessarily mean that eukaryote signature proteins, including prototypic SNAREs, carry out exactly the same function in the archaeon as they do in eukaryotic cells. For example, ESCRT proteins may play a role in archaeal cell division instead of vesicle trafficking^48,49^.

It is certain that homologous factors, possibly in a closely related but yet undiscovered Asgard lineage, served as the building blocks for the machinery used in eukaryotic membrane trafficking. It is apparent that prototypic trafficking factors were present in the archaeal genomes before they hosted mitochondria and it is conceivable that the entry of an α-proteobacterium sparked the evolution of a true eukaryotic membrane trafficking system, which was then refined by duplications of a primordial set of vesicle formation and fusion proteins^8,10,50^.

## Methods

### Sequences search and classification

In earlier studies, we identified 23 basic subtypes within the eukaryotic SNARE protein family and had developed specific as well as sensitive HMMs for each subtype^8,51,52^. By using hmmscan from <u>HMMER</u> v3.2.1 ^53^, we now searched for SNAREs with these HMMs in prokaryotes from the nr-database at NCBI as of 21 November 2018. As it can be challenging to find homologous sequences across domains of life^54^, particularly for coiled-coil sequences^55^, we used a 10^−4^ expectation value cutoff and kept only the sequences for which the target motif was at least 40 amino acids long to minimize false positive results. Next, we identified groups of similar sequence types by using the pairwise sequence similarity between all protein sequences as described in CLANS ^56^, which is based on the Basic Local Alignment Search Tool (BLAST)^57^. Only e-values of ≤ 10^−15^ were used for clustering and visualization via a method from the Python package <u>networkX</u>. With this, we were able to define 97 similarity groups of proteins with SNARE-like motifs. To better assess the presence of conserved domains and their arrangement within the similarity groups, we used SMART^58^, and PFAM^59^. For a more detailed analysis, we made use of secondary structure predictions ^60–62^.

### Search for TA proteins

The prokaryotic proteins with SNARE-like motifs were scanned for the presence of a single TMD by TMHMM 2.0 ^63^. We kept only the sequences whose TMD was at a distance of less than 30 residues from the *C*-terminus, as established in a previous study^13^. Similar scans were performed for individual proteomes of *Thalassospira australica* (txid1528106), *Lokiarchaeum sp. GC14_75* (txid1538547), *Candidatus Heimdallarchaeota archaeon LC_2* (txid1841597), *Candidatus Heimdallarchaeota archaeon LC_3* (txid1841598), and *Candidatus Heimdallarchaeota archaeon B3-JM-08* (txid2012493).

### Phylogenetic reconstruction

Phylogenetic reconstructions were carried out essentially as described ^8,64^. The alignments were based on the SNARE motifs 53 amino acid long stretch detected by HMMER v3.2.1 and refined globally with HMMER v2^53^. We used a combination of three different maximum likelihood programs (IQ-TREE^65^, Randomized Accelerated Maximum Likelihood (RAxML)^66^, and Phylogenetic estimation using Maximum Likelihood (PhyML)^67^). To be able to calculate the best trees, we first used IQ-TREE to estimate the best model and the model parameters. For all trees, the LG matrix^68^ with a Γ-distribution for rate heterogeneity was found to be the most appropriate model. We executed IQ-TREE with 1000 rapid bootstrap replicates^69^. PhyML was set to start with 20 random start trees and 1000 standard non-parametric bootstrap replicates. Additionally, we used Subtree Pruning and Regrafting transformations and a random seed of 9. For RAxML, we again chose a random seed of 9 and 1000 standard non-parametric bootstrap replicates. We then used RAxML to estimate site-wise log-likelihoods for all calculated trees and used Consel ^70^ to estimate an Approximately Unbiased (AU) ranking^71^. The highest-ranking tree was taken as a reference. Again by making use of Consel, we adjusted the bias of the support values of the different bootstrap replicates from RAxML and PhyMl via the AU test^71^. IQ-TREE has a built-in correction and no further adjustment was necessary. Finally, as an additional and more independent confidence estimator, we used IQ-TREE to run likelihood mapping^72^ on the best tree. The main edges in all trees are annotated in the following order: likelihood-mapping, IQ-TREE support, RAxML support, PhyML support (the two latter as AU p-values). All phylogenetic trees and corresponding sequence alignments are available at DOI 10.5281/zenodo.3478558.

### Protein expression and purification

All recombinant proteins were cloned iton the *pET28a* vector, which contains an *N*-terminal, thrombin-cleavable His_6_-tag. The constructs for neuronal SNARE proteins from *Rattus norvegicus* have been described before: the SNARE domain of syntaxin 1a (aa 180 –262, Syx), a cysteine-free variant of SNAP-25b (aa 1–206, SN25), the first helix of SNAP-25 (aa 1–83, SN1), the second helix of SNAP-25 (aa 120–206, SN2), and synaptobrevin 2 (aa 1–96, Syb2)^20,73^. Codon-optimized versions of the following sequences were synthesized and subcloned into the *pET28a* vector (GenScript): OGT53257.1 (*Gammaproteobacteria bacterium RIFCSPHIGHO2_12_FULL_42_13*, aa 116–176, μ-bac-R), a SNAP-25-like SNARE from *Legionella cherrii* (WP_028380397.1, aa 1–209, Leg_Qbc) and the SNARE-like regions of WP_081944311.1 (*Thalassospira australica*, aa 616–686, ThAu_MCP), PWI47941.1 (*Candidatus Heimdallarchaeota archaeon B3-JM-08*, aa 150–216, HPS1), OLS22354.1 (*Candidatus Heimdallarchaeota archaeon LC_2*, aa 135–217, HPS2). All proteins were expressed in the *Escherichia coli* strain BL21 (DE3) and purified by Ni^2+^-chromatography. After cleavage of the His_6_-tags by thrombin, the proteins were further purified by ion exchange chromatography on an Äkta system (GE Healthcare). The proteins were eluted with a linear gradient of NaCl in a standard buffer (20 mM Tris pH 7.4, 1 mM EDTA) as previously described ^20,73^. The eluted proteins were 95% pure, as determined by gel electrophoresis. Protein concentrations were determined by absorption at 280 nm and the Bradford assay.

### Size exclusion chromatography

Size exclusion chromatography was performed on a HR-10/300 Superdex-200 column (GE Healthcare) in standard buffer containing 150 mM NaCl at a flow rate of 0.5 mL min^−1^. The elution profiles were monitored by UV absorption at 230 and 280 nm.

### Electrophoretic procedures

Routinely, SDS-PAGE was carried out as described by Laemmli. Non-denaturing gels were prepared and run in an identical manner to the SDS-polyacrylamide gels, except that SDS was omitted from all buffers^20,73^.

## Author contributions

E.N., D.K., N.S., and D.F. designed the study; E.N. and D.K. performed the experiments; E.N. and D.F. analysed the data; D.F. wrote the paper; and all authors reviewed, commented on, and edited the manuscript.

## Acknowledgements

This work was supported by the Swiss National Science Foundation (grant 31003A_182732 to D.F.. We thank the Division de Calcul et Soutien à la Recherche of the UNIL for access to the university computer infrastructure. We thank all members of the Fasshauer Laboratory for helpful discussions. We thank Tobias Klöpper for support with the analysis and comments on the manuscript.

## Competing interests

The authors declare no competing interests.

## List of abbreviations

SNARE: Soluble *N*-ethylmaleimide-sensitive factor attachment protein receptor
TMD: transmembrane domain
MCP: methyl-accepting chemotaxis protein
HMM: Hidden Markov Model
TA: tail anchored
GET: Guided Entry of Tail-anchored proteins
HPS: Heimdallarchaeota Prototypic SNARE
AU: Approximately Unbiased

## Additional Information

Supplementary Information is available for this paper. The file contains Supplementary Information Sections 1–7. Supplementary Figures are included within the respective sections.

Correspondence and requests for materials should be addressed to D.F.

---

**Extended Data Figure 1:**
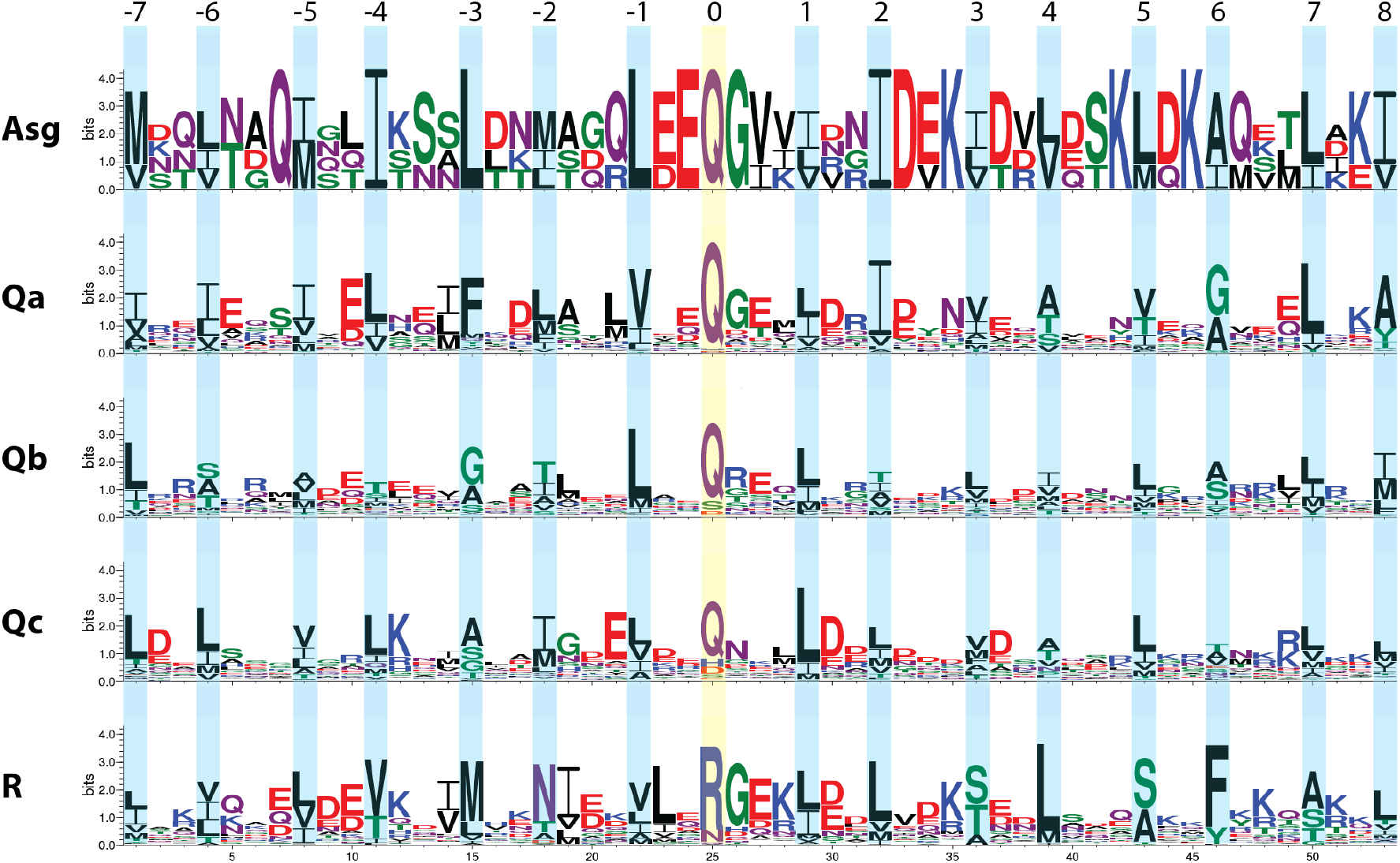
<u>WebLogo</u>^74^ representation of the SNARE motif region of SNARE-like proteins from Asgard archaea and of the four basic types of eukaryotic SNARE proteins. The positions of the layer residues (−7 to +8) forming the hydrophobic core of the four-helix bundle SNARE complex are indicated. The representation of the Asgard proteins is based on four different sequences only (Extended Data Table 1), whereas the eukaryotic types are based on SNARE sets from 24 representative eukaryotic species (Extended Data Table 3). In detail, the weblogos for eukaryotic SNAREs are based on alignments of 116 Qa-, 129 Qb-, 120 Qc-, and 96 R-SNAREs, respectively.

**Extended Data Fig. 2:**
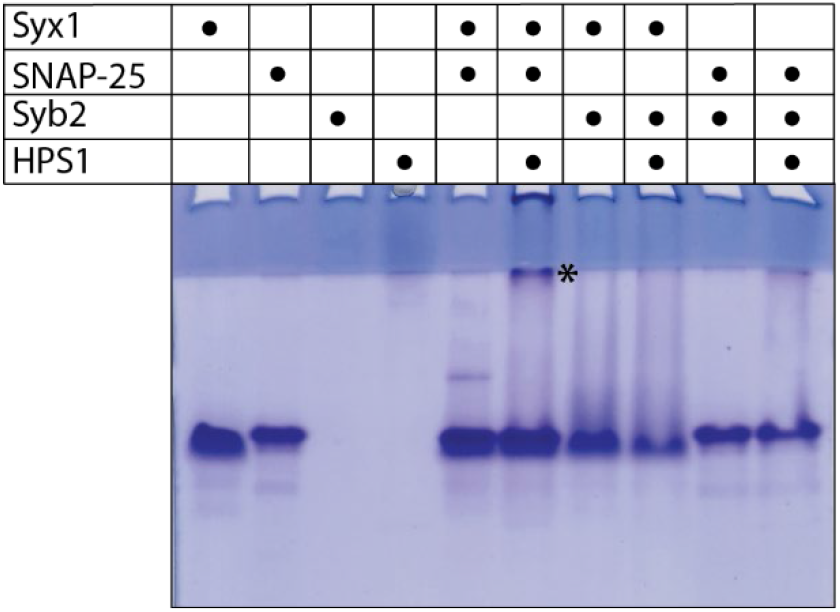
Interaction of HPS1 with the neuronal SNARE proteins SNAP-25 and Syx1 tested by non-denaturing gel electrophoresis. The SNARE-like region of PWI47941.1 from *Heimdallarchaeota B3-JM-08*, termed HPS1 (aa 150–209), was mixed with the neuronal SNARE proteins synaptobrevin (Syb2, R-SNARE), SNAP-25 (Qbc-SNARE), and the SNARE motif of syntaxin (Syx1, Qa-SNARE) as indicated. All incubations were performed as described in the legend of Fig. 1. Proteins were visualized by Coomassie Blue staining. A new band appears (*) upon mixing of HPS1 with SNAP-25 and Syx1. Additional titration experiments are shown in (Supplementary Information, Section 2, Fig. S2.3).

**Extended Data Fig. 3:**
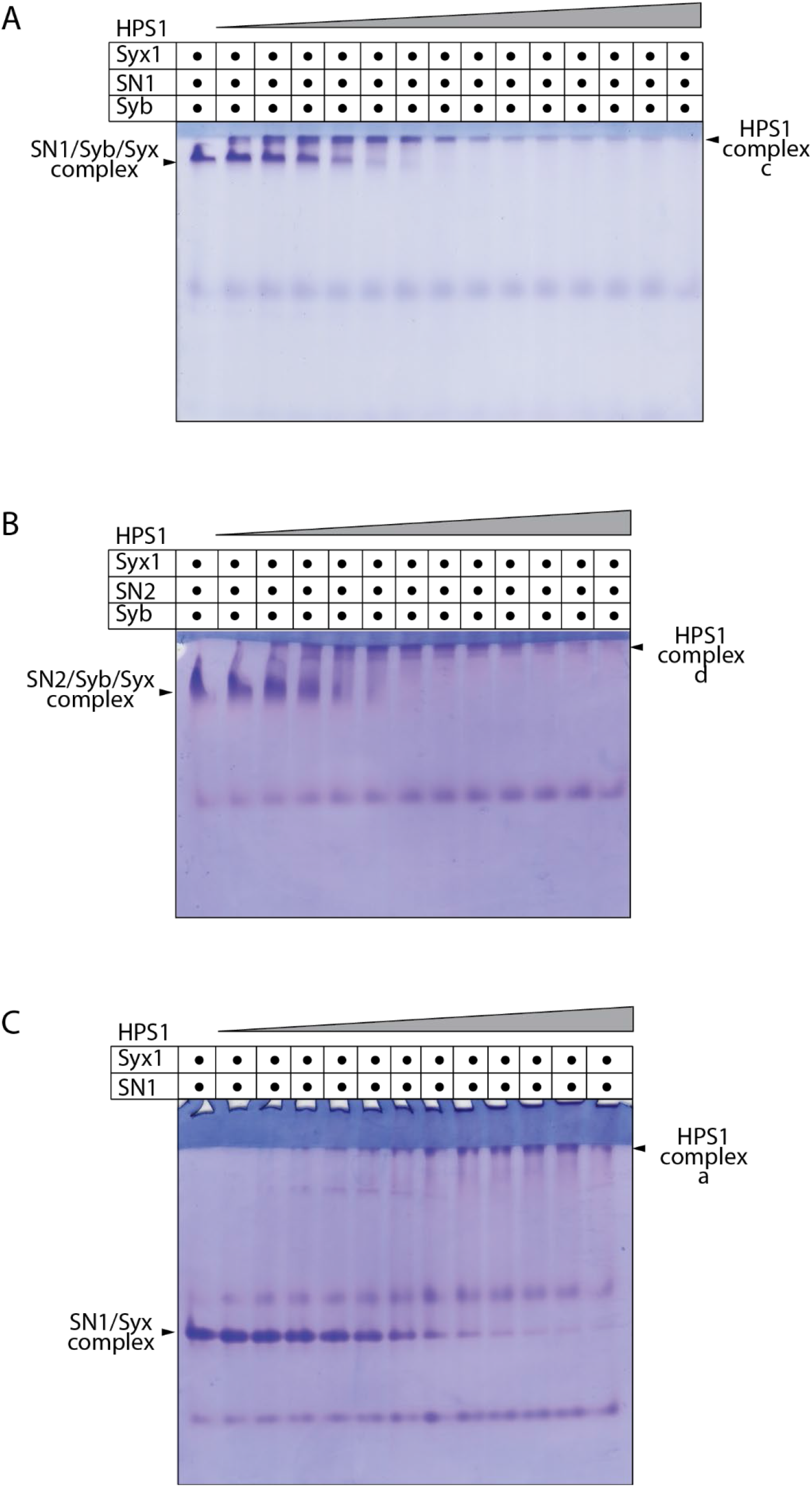
HPS1 interacts with different subcomplexes of neuronal SNARE proteins. The four SNARE motifs of neuronal SNARE proteins can assemble into different stable subcomplexes^20^. HPS1 can transform these subcomplexes by forming new stable complexes containing HPS1. However, these HPS1 complexes cannot be separated by non-denaturing gel electrophoresis. In three different experiments, constant amounts of preassembled neuronal subcomplexes, (A) the SN1–Syb2–Syx, (B) the SN2–Syb2–Syx (see Supplementary Information, Section 2, Fig. S2.4) for the enrichment of the subcomplexes) and (C) the SN1–Syx complex were mixed with increasing amounts of HPS1. In each gel, the respective neuronal subcomplex disappeared upon the addition of increasing amounts of HPS1; a new complex band appeared as indicated. All incubations were performed as described in the legend of Fig. 1. Proteins were visualized by Coomassie Blue staining.

**Extended Data Fig. 4:**
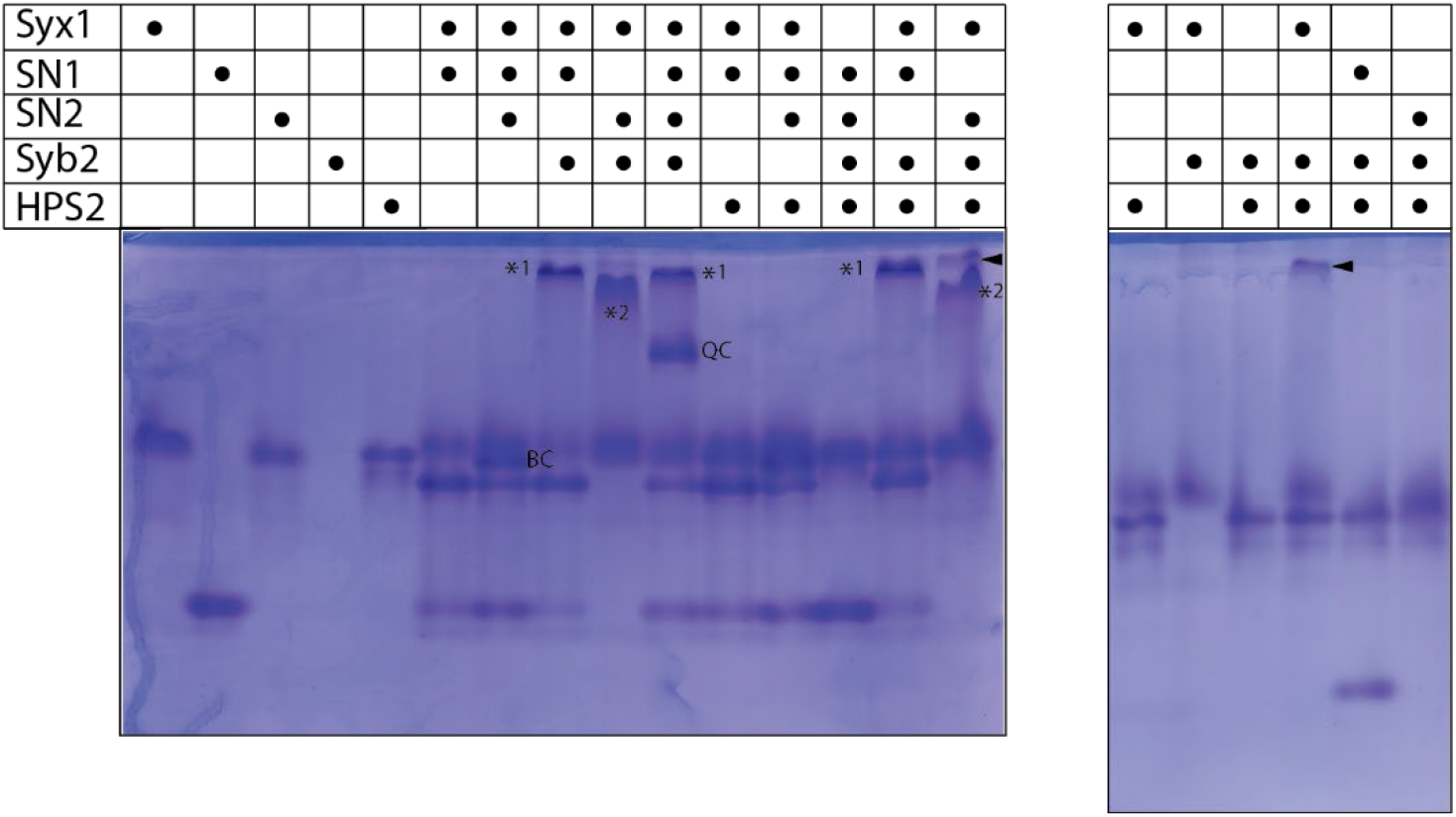
Interaction pattern of HPS2 with neuronal SNARE proteins, visualized by non-denaturing gel electrophoresis. The SNARE-like region of OLS22354.1 (HPS2) from *Heimdallarchaeota LC_2*, termed HPS2 (aa 135–217), was mixed in different combinations with the neuronal SNARE proteins Syx1 (Qa-), Syb2 (R-), and the two SNARE motifs of SNAP-25, SN1 (Qb-), and SN2 (Qc-SNARE), which were used as independent proteins. All incubations were performed as described in the legend of Figure 1. Proteins were visualized by Coomassie Blue staining. HPS2 formed a stable complex in combination with the two SNAREs Syx1 and Syb2 as indicated by the arrows. Stable complexes of neuronal proteins are indicated as in Figure 1: binary complex consisting of Syx– SN1–SN2 (BC), a Syx1–SN1–Syb2 complex; *1), a Syx1–SN2–Syb2 complex (*2), and a quaternary complex (QC)^20^. Note that Syb2 is not detectable in the non-denaturing gel (Lane 4) because of its isoelectric point of 8.5.

**Extended Data Fig. 5:**
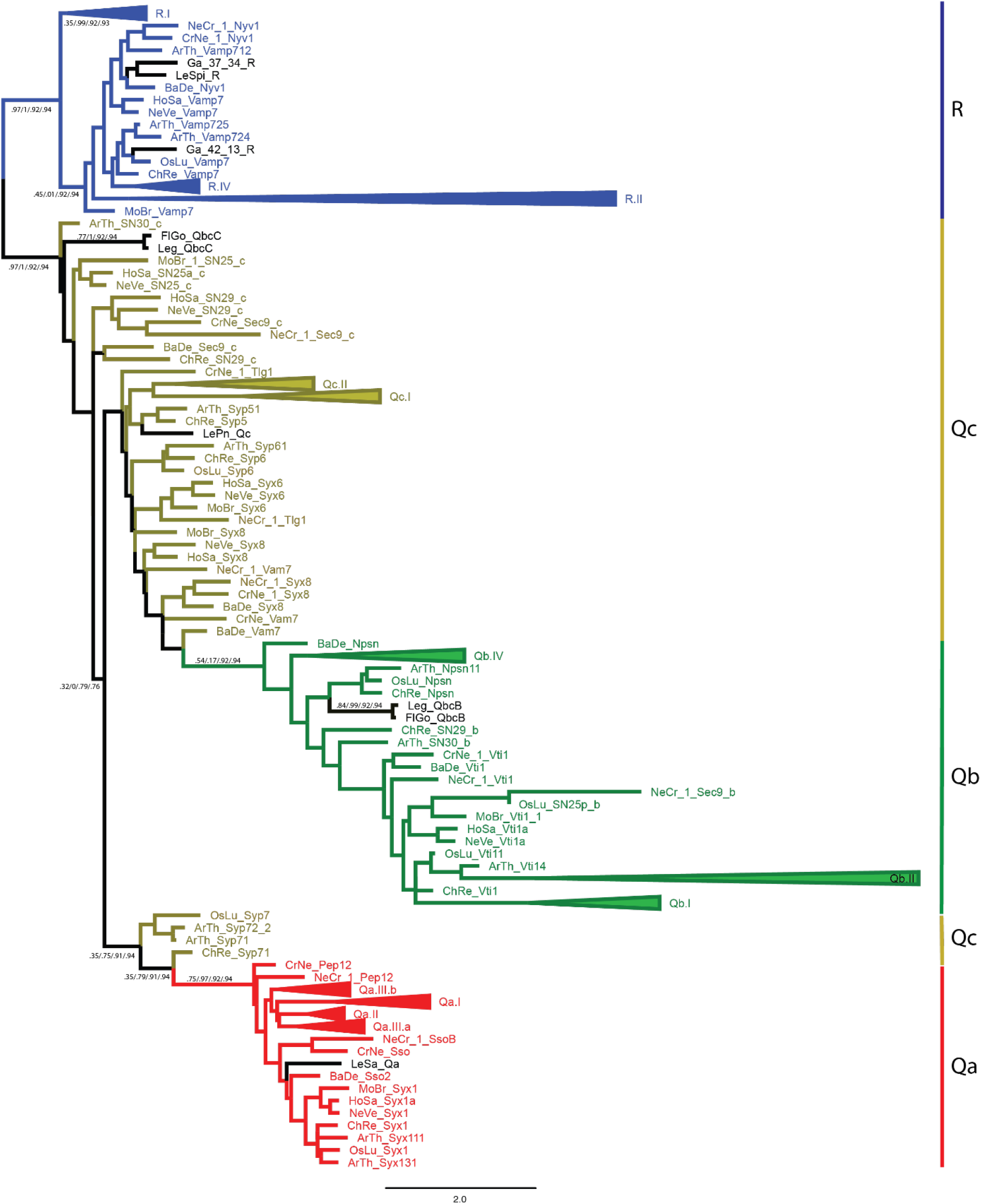
Phylogenetic analysis of eukaryotic and bacterial SNARE proteins. The maximum likelihood tree was constructed from six SNARE proteins from different γ-proteobacteria of the order Legionellales (Extended Data Table 2) and typical SNARE sets of nine representative eukaryotic species (Extended Data Table 3). Note that the two SNARE motifs of Qbc-SNAREs are indicated by a B or C for the Qb- or the Qc-helix, respectively. Qa-SNAREs are in red, Qb-SNAREs in khaki, Qc-SNAREs in moss green, and R-SNAREs in blue. As in Fig. 2, the eukaryotic SNAREs split into four main groups that correspond to the four different positions in the four-helix bundle SNARE complexes. The different γ-proteobacterial SNAREs, labeled in black, are nested within different branches of the eukaryotic SNARE types, suggesting that different SNARE genes were transferred independently from eukaryotes to parasitic intracellular bacteria. Some SNARE types are shown as collapsed clades. The entire tree, without collapsed clades, is given in the Supplementary Information, Section 6. Statistical support values (likelihood-mapping, IQ-TREE support, RAxML support, PhyML support) are given at selected inner edges. The scale bar indicates the number of substitutions per site.

**Extended Data Table 1:**
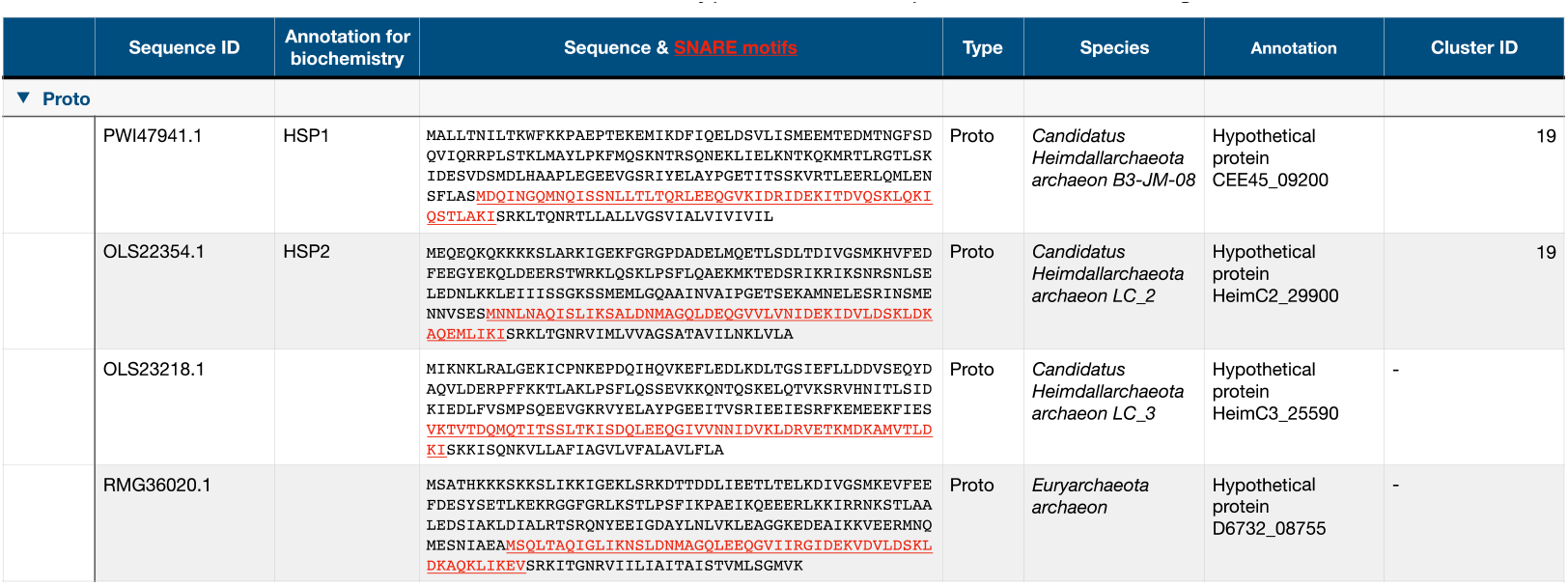
List of Prototypic SNARE Sequences Found in Asgard Archaea

**Extended Data Table 2:**
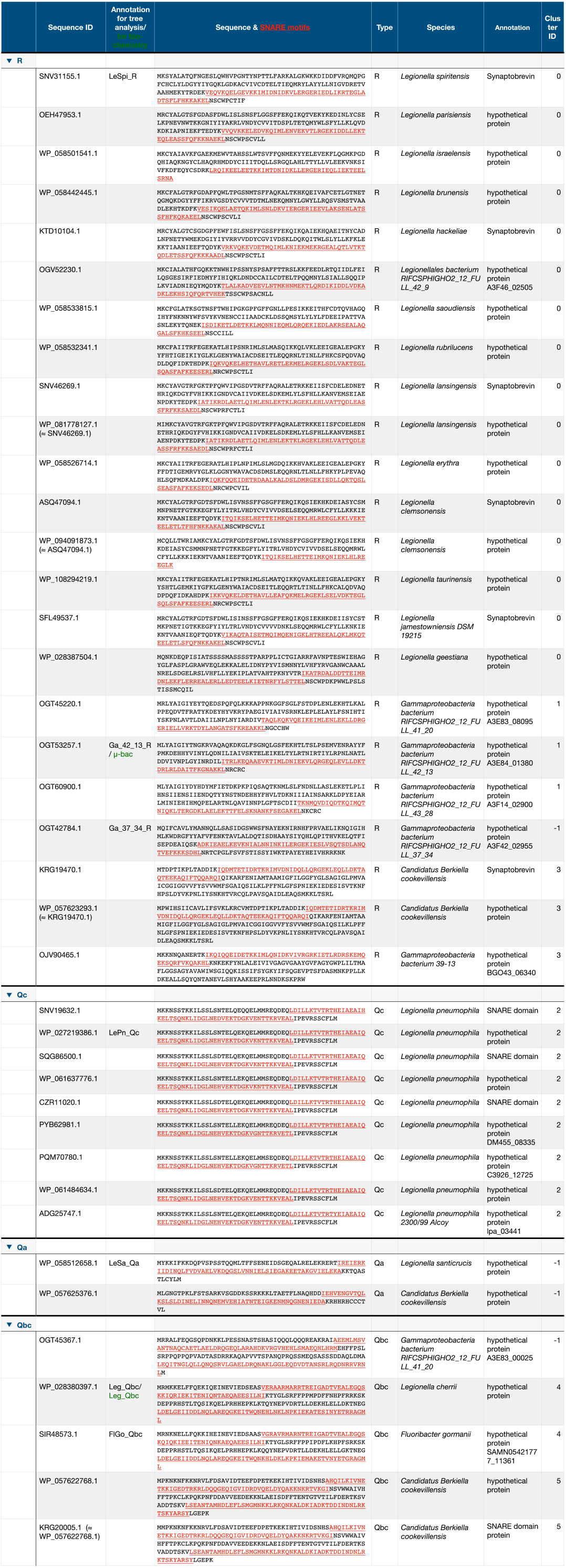
List of fide SNARE in y-proteobacteria of the order Legionellales

**Extended Data Table 3:**
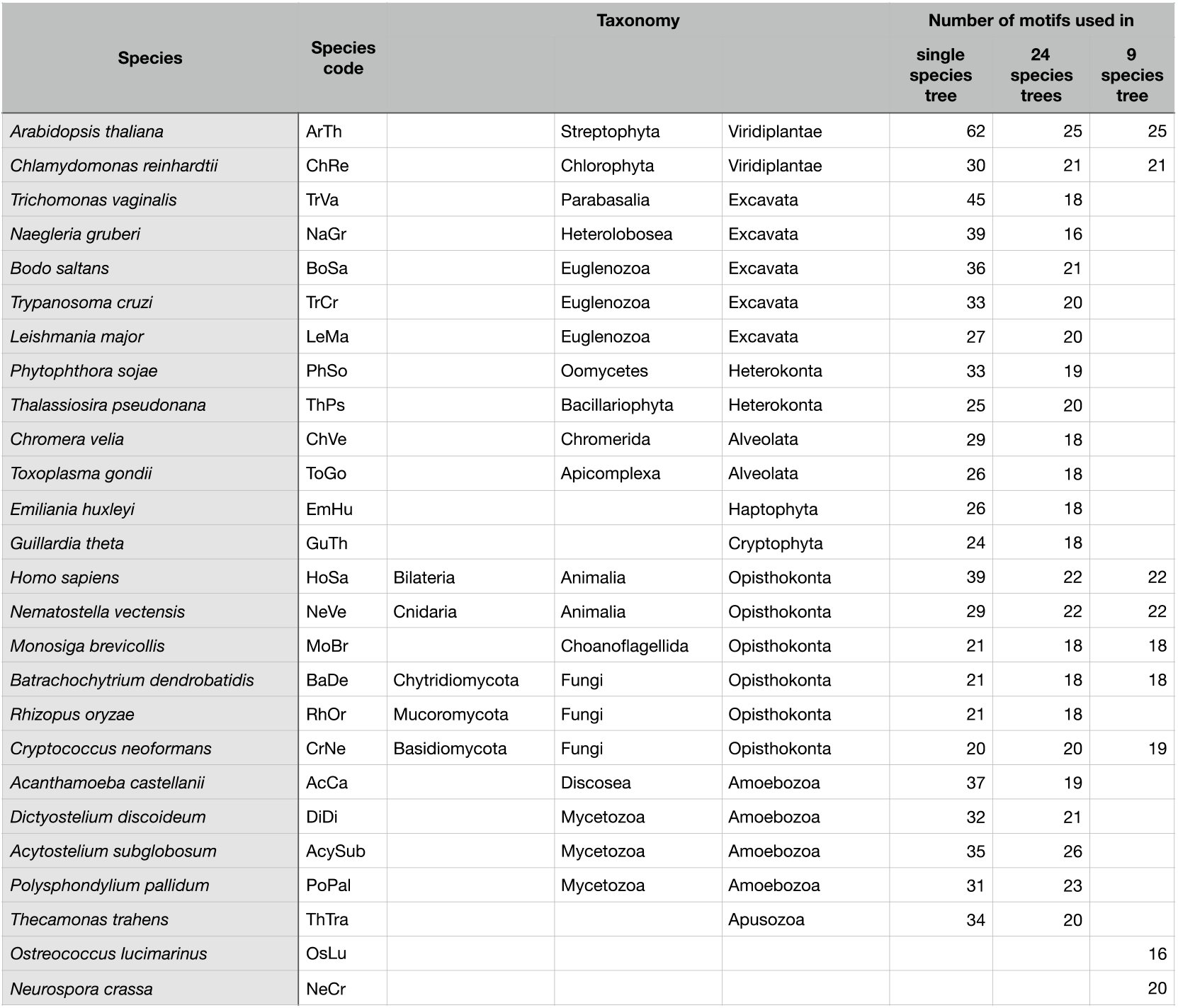
Number Of Snare Sequences Per Species Used For Tree Calculations

## Notes

#### Summary of Updates

In the abstract of the first version, the special characters for α- and γ-proteobacteria were placed incorrectly.

https://zenodo.org/record/3478559#.XamhZC2B0Uw

