## Supplemental Data 1 for "Prototypic SNARE proteins are encoded in the genomes of Heimdallarchaeota, potentially bridging the gap between the prokaryotes and eukaryotes"

#### Section 1

As described in the main text, we searched for archaea and bacteria (collectively, prokaryotes) the NCBI protein databases, which contained about 528 million sequences in Nov. 2018, for sequences containing SNARE-like motifs. For the scan, we used the Hidden Markov Model (HMM) profiles trained previously[1-3] to classify the SNARE motifs of eukaryotic SNARE proteins into about 20 different subtypes. We implemented a $10^{-4}$ expectation value cutoff and kept only the sequences for which the target motif was at least 40 amino acids long to minimize false positive results (see Methods). Around 5000 prokaryotic sequences met these criteria (Supplementary Information, Section 8, Supplementary Table 1).

For an overview of the relationships between the collected prokaryotic sequences, we clustered them by using the Basic Local Alignment Search Tool (BLAST) to construct groups of similar factors for further inspection (Methods). The sequences split into 96 different clusters of different sizes. A cluster map is depicted in Fig. S1.1. The distribution of the e-values of different clusters is shown in Fig. S1.2. A more detailed description of the sequences found in different clusters is given in the main text. Larger clusters with lower scoring sequences are described in Section 5.

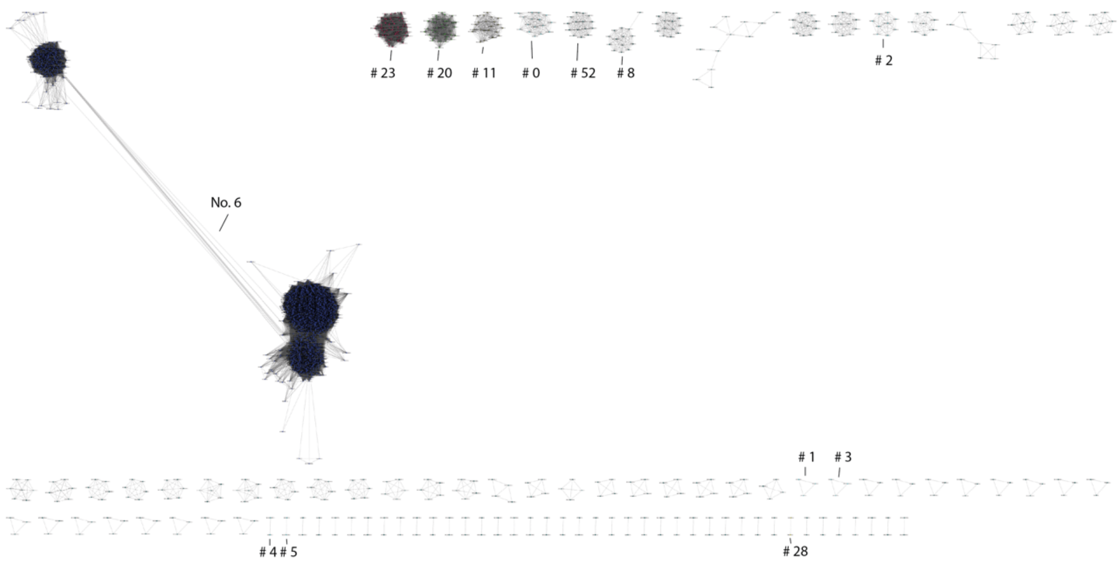

**Fig. S1.1. Cluster map of prokaryotic sequences with SNARE-like motifs established by a sequence similarity network approach.**

The clusters have been visualized with the default prefuse force directed layout of Cytoscape 3.5.1 (https://cytoscape.org). Homology is depicted by lines between nodes. Clusters are ordered according to the sequence with the lowest e-value within a cluster (Supplementary Table 1). Clusters 0–5 comprise the clusters with *bona fide* SNARE sequences found in γ-proteobacteria of the order Legionellales. Cluster 6 contains different types of prokaryotic chemotaxis proteins (see Section 5 for details).

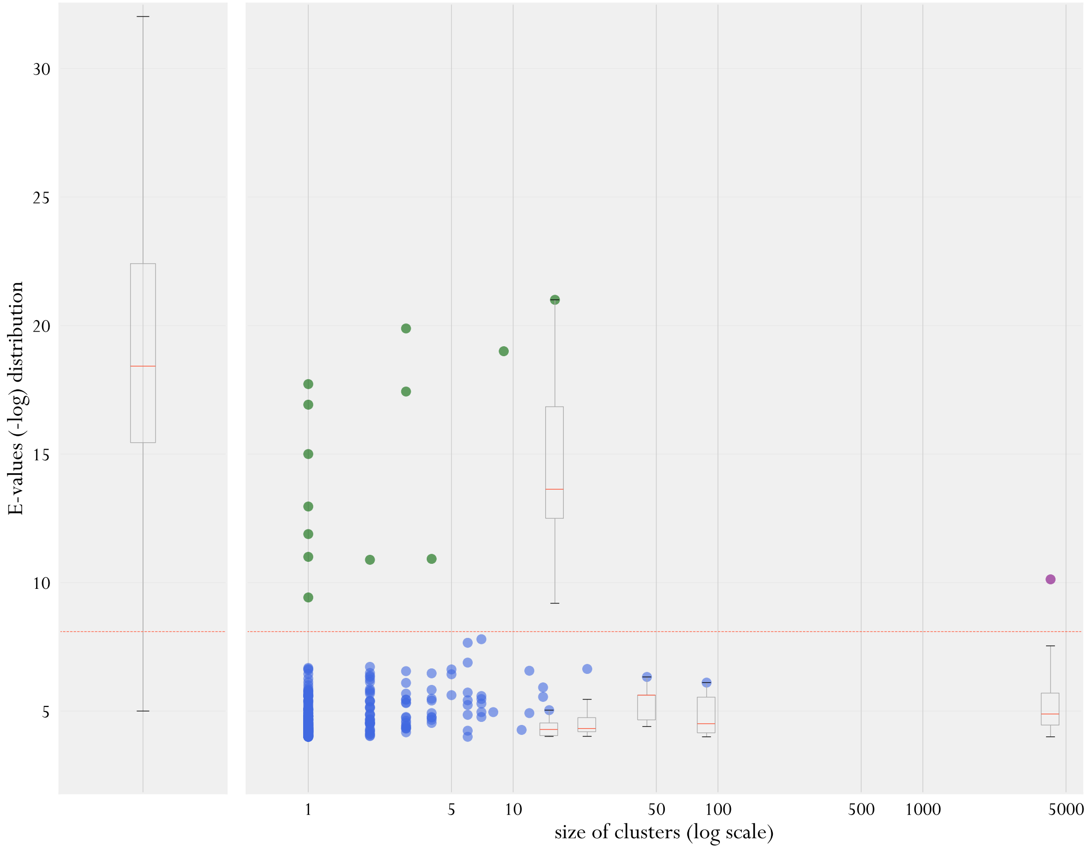

**Fig. S1.2. Box plot showing the distribution of e-values of different clusters of prokaryotic sequences with SNARE protein-like regions.**

On the left, the e-value distribution (see methods) of 19533 eukaryotic SNARE domains from our database is shown as box-and-whisker plots; the box length is the interquartile range, the upper whisker marks the smaller of the maximum value and Quartile 3 + 1.5 interquartile range (IQR); the lower whisker marks the larger of the smallest value and Quartile 1 – 1.5 IQR. The solid horizontal red line inside the box represents the median e-value. On the right, the e-value distributions for prokaryotic sequences grouped into homology clusters (see Methods), including singletons, are given. Each cluster, ordered by size, is represented with its lowest e-value as a colored circle (violet: methyl-accepting chemotaxis proteins (MCPs); green: prokaryotic sequences from Legionellales that encode very probable bona fide SNARE proteins; blue: other prokaryotic sequences). For clusters with more than 20 sequences, box-and-whisker plots are also shown. The dashed horizontal red line represents the value of the median – 2 σ of the eukaryotic SNARE e-values. Note that the e-values for eukaryotic SNAREs and for prokaryotic sequences with SNARE-like motifs were computed with the hmmscan tool from HMMER v3.2.1[4].

#### References – Section 1

1 Kienle, N., Kloepper, T. H. & Fasshauer, D. Phylogeny of the SNARE vesicle fusion machinery yields insights into the conservation of the secretory pathway in fungi. *Bmc Evolutionary Biology* **9**, doi:10.1186/1471-2148-9-19 (2009).

2 Kloepper, T. H., Kienle, C. N. & Fasshauer, D. An elaborate classification of SNARE proteins sheds light on the conservation of the eukaryotic endomembrane system. *Molecular Biology of the Cell* **18**, 3463-3471, doi:10.1091/mbc.E07-03-0193 (2007).

3 Kloepper, T. H., Kienle, C. N. & Fasshauer, D. SNAREing the basis of multicellularity: Consequences of protein family expansion during evolution. *Molecular Biology and Evolution* **25**, 2055-2068, doi:10.1093/molbev/msn151 (2008).

4 Eddy, S. R. Profile hidden Markov models. *Bioinformatics* **14**, 755-763 (1998).

### Section 2: SNARE-like proteins from Asgard archaea

#### 2.1 Domain organization of SNARE-like proteins from Asgard archaea

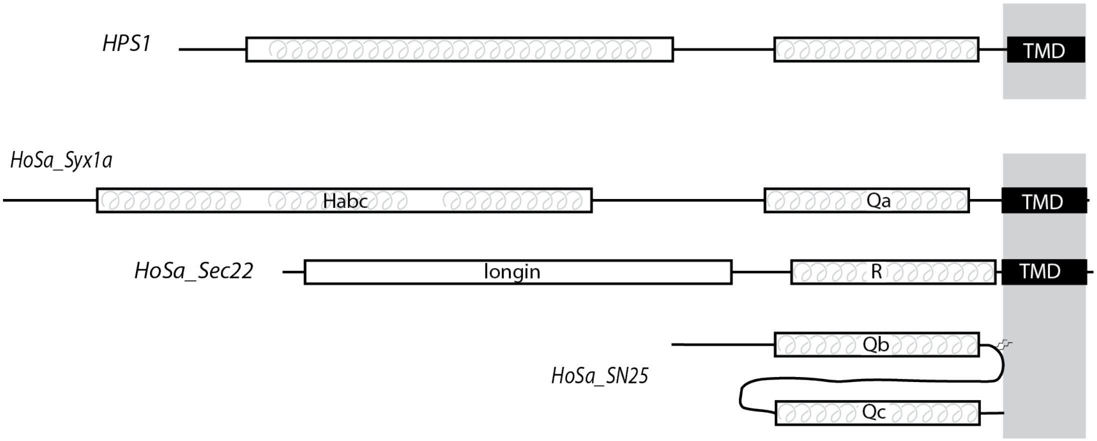

**Fig. S2.1. Schematic representation of the domain organization of SNARE-like proteins from Asgard archaea.** On top, the domain organization of the SNARE-like protein HPS1 from *Candidatus Heimdallarchaeota archaeon B3_Heim 179* is shown. Its SNARE-like region is connected to a *C*-terminal transmembrane domain (TMD) by a short linker region. The *N*-terminal region is predicted[1] to fold into a three-helix bundle structure, as found in several eukaryotic Q-SNARE types. The other SNARE-like proteins from Asgard archaea (Extended Data Table 1) have a very similar domain arrangement. The domain architecture of the most important types of SNARE proteins are depicted below (Syx1a, Sec22, and SNAP-25 from *Homo sapiens* are shown). In most SNAREs proteins, the SNARE motifs are connected by a short linker to a *C*-terminal TMD. The Qa-SNARE Syx1a carries an *N*-terminal Habc-domain that folds into a three-helix bundle structure[2,3]. This domain arrangement can be found in several other types of Q-SNAREs[4-7] but not in R-SNAREs, which usually carry an *N*-terminal longin domain, a globular fold with an α-β-α sandwich architecture[8]. In SNAP-25, two SNARE motifs of different types (Qb and Qc) are connected by a linker that is palmitoylated, whereas a TMD is lacking.

#### 2.2 Genomic neighborhood of the SNARE-like genes in Asgard archaea

*Candidatus Heimdallarchaeota archaeon B3_Heim 179*

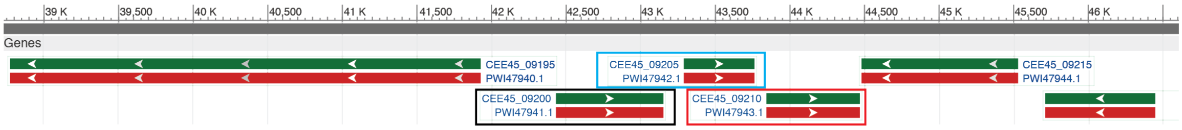

*Candidatus Heimdallarchaeota archaeon LC_3*

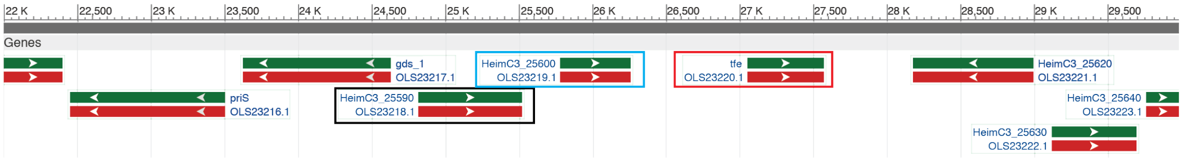

*Candidatus Heimdallarchaeota archaeon LC_2*

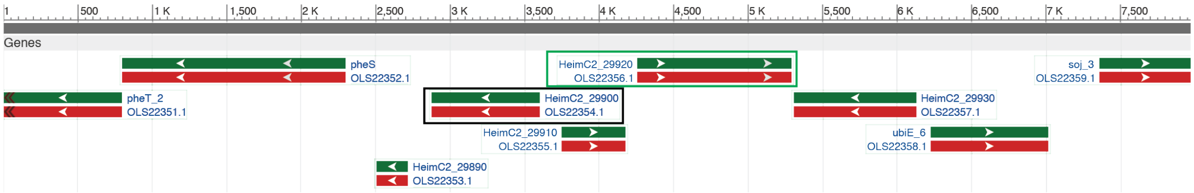

*Euryarchaeota archaeon isolate J059*

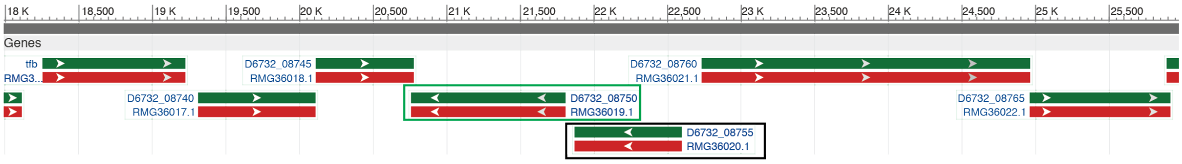

**Fig. S2.2. View of the region (~ 8000 nucleotides) encompassing the SNARE-like factors in the four Asgard metagenomes.**

The SNARE-like factors are indicated by black boxes. Note that the gene organization for two genes downstream of the SNARE-like factor is shared between *Candidatus Heimdallarchaeota archaeon B3_Heim 179* (NJBF01000020.1) and *Candidatus Heimdallarchaeota archaeon LC_3* (MDVS01000044.1). The gene next to the SNARE-like factor encodes for a protein with a roadblock domain (blue box), which often serves as binding platform for small GTPases[9]. The following gene encodes for Transcription Factor E (red box). A gene encoding for a protein of unknown function (green box) is found next to the SNARE-like factor in Euryarchaeota archaeon isolate J059 (RFHV01000376.1) and *Candidatus Heimdallarchaeota archaeon LC_2* (MDVR01000079.1), although in reversed orientation.

#### 2.3 Native gel titration of HPS1 with neuronal SNAREs

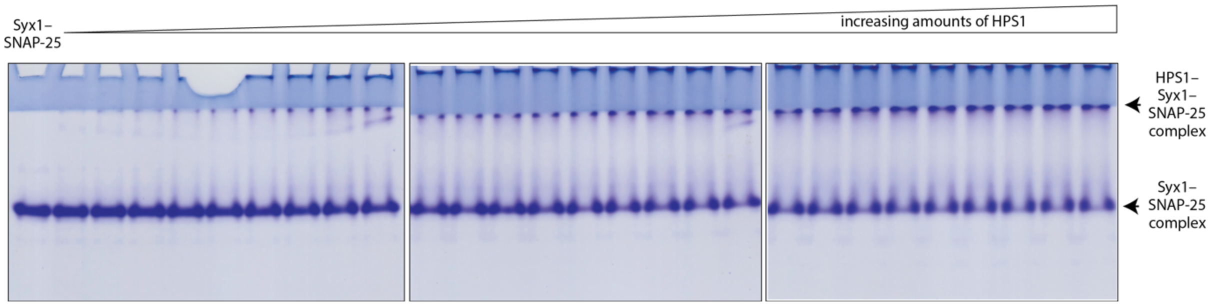

**Fig. S2.3. Interaction of HPS1 with the binary Syx1–SNAP-25 complex.**
Approximate stoichiometric amounts of Syx1 and SNAP-25 were mixed to preform the binary Syx1–SNAP-25 complex[10]. Increasing amounts of HPS1 were added and incubated overnight in standard buffer (20 mM Tris, pH 7.4, 1 mM EDTA) containing 100 mM NaCl prior to separation by non-denaturing gel electrophoresis. The gel was stained with Coomassie Blue.

#### 2.4 Enrichment of stable neuronal subcomplexes for binding experiments

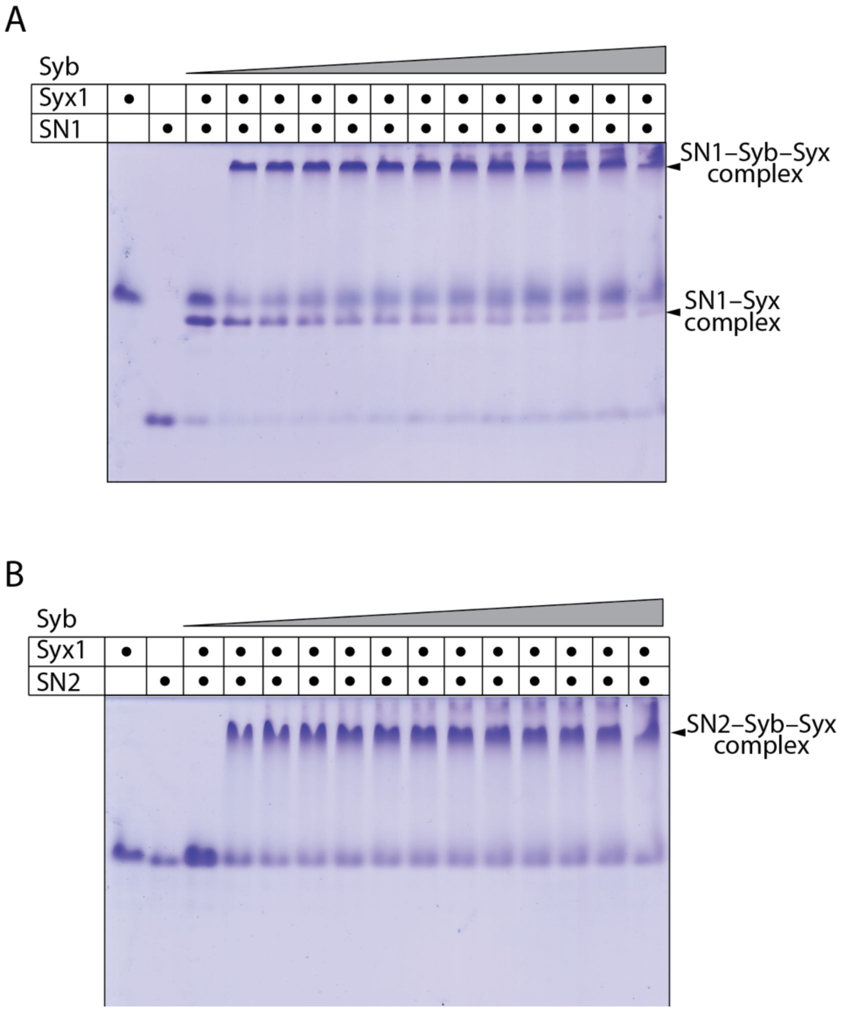

**Fig. S2.4. Enrichment of SN1–Syb–Syx and SN2–Syb–Syx complexes monitored by nondenaturing gel electrophoresis.**

Approximate stoichiometric amounts of (A) Syx1 and SN1 or (B) Syx1 and SN2 were mixed. Note that Syx1 and SN1 assemble into a stable Syx1–SN1 complex, whereas Syx1 and SN2 do not interact. Upon addition of increasing amounts of Syb to the two different premixes, a new complex band appears. The Syx–Syb–SN1- and Syx–Syb–SN2 complexes can be isolated, but their precise stoichiometry is not known. Addition of the missing SNARE domain is transforming them into the quaternary SNARE complex. The gel was stained with Coomassie Blue. Extended Data Figure 3 shows that a new complex band appeared upon mixing of the enriched Syx–Syb–SN1 and Syx–Syb–SN2 complexes with HPS1.

#### 2.5 Size exclusion chromatography experiments

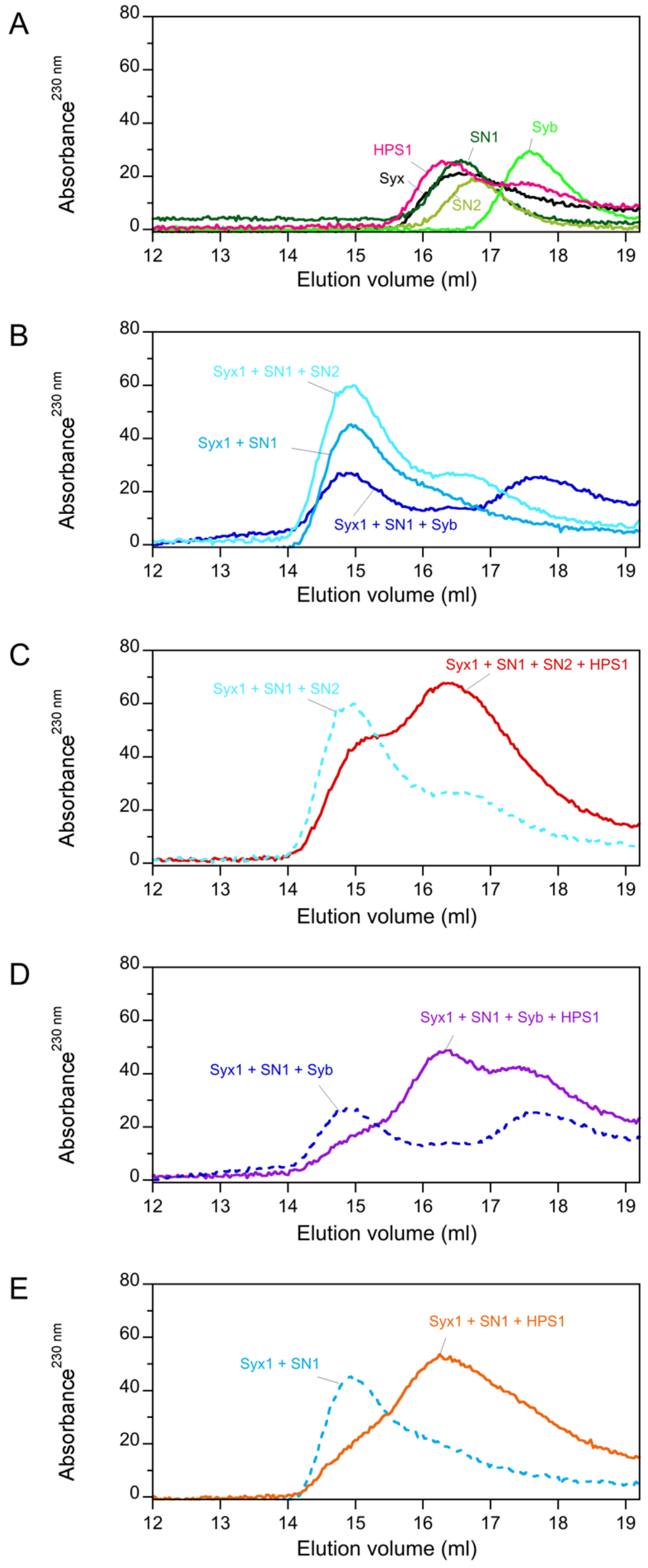

**Fig. S2.5. Size exclusion chromatographic elution profiles for the SNARE motifs of individual neuronal SNARE proteins, HPS1 and different mixes as indicated.**

Prior to separation on a Superdex 200 column, equimolar protein mixes (about 15 µM) were incubated overnight in standard buffer (20 mM Tris pH 7.4, 1 mM EDTA) containing 150 mM NaCl. Note that the individual proteins, although their molecular mass is very similar, elute at different volumes.

#### References – Section 2

1 Lobley, A., Sadowski, M. I. & Jones, D. T. pGenTHREADER and pDomTHREADER: new methods for improved protein fold recognition and superfamily discrimination. *Bioinformatics (Oxford, England)* **25**, 1761-1767, doi:10.1093/bioinformatics/btp302 (2009).

2 Fernandez, I. *et al.* Three-dimensional structure of an evolutionarily conserved N-terminal domain of syntaxin 1A. *Cell* **94**, 841-849 (1998).

3 Lerman, J. C., Robblee, J., Fairman, R. & Hughson, F. M. Structural analysis of the neuronal SNARE protein syntaxin-1A. *Biochemistry* **39**, 8470-8479 (2000).

4 Kloepper, T. H., Kienle, C. N. & Fasshauer, D. An elaborate classification of SNARE proteins sheds light on the conservation of the eukaryotic endomembrane system. *Molecular biology of the cell* **18**, 3463-3471, doi:10.1091/mbc.E07-03-0193 (2007).

5 Jahn, R. & Fasshauer, D. Molecular machines governing exocytosis of synaptic vesicles. *Nature* **490**, 201-207, doi:10.1038/nature11320 (2012).

6 Baker, R. W. & Hughson, F. M. Chaperoning SNARE assembly and disassembly. *Nature reviews* **17**, 465-479, doi:10.1038/nrm.2016.65 (2016).

7 Wang, T., Li, L. & Hong, W. SNARE proteins in membrane trafficking. *Traffic (Copenhagen, Denmark)* **18**, 767-775, doi:10.1111/tra.12524 (2017).

8 Daste, F., Galli, T. & Tareste, D. Structure and function of longin SNAREs. *J Cell Sci* **128**, 4263-4272, doi:10.1242/jcs.178574 (2015).

9 Levine, T. P. *et al.* Discovery of new Longin and Roadblock domains that form platforms for small GTPases in Ragulator and TRAPP-II. *Small GTPases* **4**, 62-69, doi:10.4161/sgtp.24262 (2013).

10 Fasshauer, D., Otto, H., Eliason, W., Jahn, R. & Brunger, A. Structural changes are associated with soluble N-ethylmaleimide-sensitive fusion protein attachment protein receptor complex formation. *Journal of Biological Chemistry* **272**, 28036-28041, doi:10.1074/jbc.272.44.28036 (1997).

#### Section 3: Putative tail-anchored proteins

As described in the main text, tail-anchored membrane proteins (TA proteins) are a subset of membrane proteins that are anchored via a *C*-terminal single-pass TMD facing the cytosol[1-3]. They are not inserted co-translationally into the membrane via the signal recognition particle and the Sec61 translocation channel, but use the "Guided Entry of Tail-anchored proteins" (GET) pathway[4,5], which is driven by a conformational change of the ATPase Get3 that binds directly to a TA protein[6]. We screened the NCBI database for TA proteins for the prokaryotes that were relevant to our study. The repertoire of TA proteins, usually contained the $\gamma$- (SecE) and β-subunits (SecG) of the Sec61/SecYEG protein secretory system[7]. It should be noted that the bacterial β-subunits have an additional TMD that precedes the *C*-terminal TMD and are therefore usually not found as TA proteins in bacteria[7].

##### 3.1 TA proteins found in the HMM screen for SNARE proteins

Cluster 8:
```
>OJZ14246.1 hypothetical protein BGP22_06020 [Variovorax sp. 67-131]
MSEPHARTQELLLLGQIHGLVQALKDGQDRQNRRMDGFDTRFDALDGRLRTVEQRAAVFGAASGGAMAIGTALLAEAVKQWFR
NGPGGN
>SEJ95423.1 hypothetical protein SAMN05518853_105110 [Variovorax sp. OK202]
MSEPHVRTQELLLLGQIHGLVQALKDGQDQQNRRMDSFDTRFDALDGRLRSVEQRAAMFGAASGGAMAIGTALLAEAIKQWFR
SGPGIN
>WP_093343527.1 hypothetical protein [Variovorax sp. PDC80]
MNDPEIRAQELFLLGQIHGLVSALKEGQDRQSRRMDAVDARFDSLDSRLRAVEQRAAAFGAASGGAMAIATALLAESIKQWFR
SNASGS
>PBI84102.1 hypothetical protein BKP43_55320 [Variovorax boronicumulans]
MSDSHARAQELLLLGQIHGLVQALKEGQDQQNRRMDSFDARFDALDGRLRAVEQRAAVFGAASGGAMAIGTALLIEAIKQWFR
GGGPGAN
>SFB93767.1 hypothetical protein SAMN03159379_01087 [Variovorax sp. NFACC26]
MSDLHARTQELLLLGQIHGLVQALKDGQDRQTRRMDGFDARFDALDGRLRTVEQRAAVFGAASGGAMAVGTALLAEAVKQWFR
NGPGIN
>SCX70901.1 hypothetical protein SAMN03159363_3693 [Variovorax sp. EL159]
MSDLHERTQELLLLGQIHGLVQALKDGQDRQNRRMDGFDTRFDALDGRLRSVEQRAAVFGAASGGAMAIGTALLAEAVKQWFR
NGPGVN
```

Cluster 28:
```
>PWI47941.1 hypothetical protein CEE45_09200 [Candidatus Heimdallarchaeota archaeon
B3-JM-08]
MALLTNILTKWFKKPAEPTEKEMIKDFIQELDSVLISMEEMTEDMTNGFSDQVIQRRPLSTKLMAYLPKFMQSKNTRSQNEKL
IELKNTKQKMRTLRGTLSKIDESVDSMDLHAAPLEGEEVGSRIYELAYPGETITSSKVRTLEERLQMLENSFLASMDQINGQM
NQISSNLLTLTQRLEEQGVKIDRIDEKITDVQSKLQKIQSTLAKISRKLTQNRTLLALLVGSVIALVIVIVIL
>OLS22354.1 hypothetical protein HeimC2_29900 [Candidatus Heimdallarchaeota
archaeon LC_2]
MEQEQKQKKKKSLARKIGEKFGRGPDADELMQETLSDLTDIVGSMKHVFEDFEEGYEKQLDEERSTWRKLQSKLPSFLQAEKM
KTEDSRIKRIKSNRSNLSELEDNLKKLEIIISSGKSSMEMLGQAAINVAIPGETSEKAMNELESRINSMENNVSESMNNLNAQ
ISLIKSALDNMAGQLDEQGVVLVNIDEKIDVLDSKLDKAQEMLIKISRKLTGNRVIMLVVAGSATAVILNKLVLA
```

Cluster 40:
```
>WP_109962236.1 hypothetical protein [Methylobacterium sp. 17Sr1-28]
MLQGQTPTAAPPNAATAVPAPVPPNPVPPGGPAQPDAPARVAVALPSQVADQLARIEDKASRIEDKYARSEALLGRVEDRVEA
AGARMNEAARQSDLAALRSEVRALSERTRGLPGLGALMLTALVTALLTVALTLVAQRLNLQGLIPLR
>WP_099956808.1 hypothetical protein [Methylobacterium currus]
MPTSAPQGTTPAAPVQVAPAQMSPAQASPAHVAPGHVPPGPIPPNAVAPPSGTPARIAVALPSQVADQLARIEDKASRIEDKY
ARSEALLTRVEDRVDAAGARMNEAARQADLAALRNEVRALTERTRRLPGLGALVLTAIITAVLTVALTLVAQRLNLQGLIPLR
>OAS17275.1 hypothetical protein A5481_27510 [Methylobacterium platani]
MTDAVPSARPATAPLPQGQMAPAQALQGQASPAQAAPAQVAPVQVSAAPASHAPTPPLPSQVADQLARIEDKASRIEDKYARS
EALLGRVEDRVEAAGARMNEAARQADLAALRNEVRALSERTRGLPGLGALMLTALVTAVLTVALTLVAQRLNLQGLIPLR
>WP_091950135.1 hypothetical protein [Methylorubrum salsuginis]
MTETSPTKSSGTDFIPTTPPPAATVTPAPAPAPAIGKADLPAASAVNHADQLARIEDKAARIEDKYARSEALLSRVEDKVENA
SSRMNEAARQADLASLRGEVRGMADRINRLPGTGALVLTALITAVLTVALMVAVQRANLDKLLPPSLGGQPQTTSNP
```

>SFL60932.1 hypothetical protein SAMN04488125_1199 [Methylorubrum salsuginis]
MPAASRMNGASASLQNNGVHAMTETSPTKSSGTDFIPTTPPPAATVTPAPAPAPAIGKADLPAASAVNHADQLARIEDKAARI
EDKYARSEALLSRVEDKVENASSRMNEAARQADLASLRGEVRGMADRINRLPGTGALVLTALITAVLTVALMVAVQRANLDKL
LPPSLGGQPQTTSNP
>WP_056244371.1 hypothetical protein [Methylobacterium sp. Leaf456]
MTETSPTKSSGTDFIPTTPPVPAAPAPAPAPSLAKADLSAAAPVNHSDQLARIEDKAARIEDKYARSEALLSRVEDKVENASS
RMNEAARQADLASLRSEVRGMADRINRLPGTGALVLTALITAVLTVALMVAVQRANLDKLLPPSLGGQPQTTTTNP

Cluster 87:
>WP_042403592.1 hypothetical protein [Geomicrobium sp. JCM 19037]
MTWDKFKKERMAISQEIIQGDAATERKMVDQFKRITDATNRAAENQEKLLTRIADTTDSNTAAIRELTKAQEHLNSSINQIQN
EHGRRLENIESSRAQEHEIKTISLKGWFGVGAAALGGTGIITVIVTFLLNYLPNY
>GAK05654.1 hypothetical protein JCM19037_4173 [Geomicrobium sp. JCM 19037]
MDQANTRSGTDPVTWDKFKKERMAISQEIIQGDAATERKMVDQFKRITDATNRAAENQEKLLTRIADTTDSNTAAIRELTKAQ
EHLNSSINQIQNEHGRRLENIESSRAQEHEIKTISLKGWFGVGAAALGGTGIITVIVTFLLNYLPNY

Singletons:
>WP_062266217.1 hypothetical protein [Endozoicomonas arenosclerae]
MMFYTDLAQSYGKRFSKAVKSAGEGAMKTASDKVSHSKWNPFTGKGKPQVRTESHGQNEPQKTPVKTRPTQTSDKAEKKQALQ
TFKAKLSSLQKQFAASLTGSDENAASTALDAKELLRTAPGLLKGHASEKLLNGHIDRAQRDLETPQDMSEKGSQDVMTVPLEM
AMDQLKGDMDTQEAKARQTLTAMAEVRHQRNYSSPRFTPLEDEVDFMQVRAEAELEEEQAMLQLERDAQATITRAEELRKLEH
EIRDLNEIFKSMSQVVYDQGQQFETLESLIDSAAFSVQEGVKTLDTCVKVDKSNHKLVAAVAASLAVLGLVALALIIFV
>WP_055737291.1 MULTISPECIES: hypothetical protein [Bacillus]
MTEPSAYDLLRKLTEVEIALAKVTQQLESQTSVLEEVRKELSMTREKADAAEEKAEIAYRKAKQALGEIKQTREETLRTQNER
KVDRRWFIGTVLTVVSLIMPMIITFYFQ
>WP_066112547.1 MULTISPECIES: hypothetical protein [Blastomonas]
MTIDADLTRKAMLDSINRLAEDMREMRTDMKKELGELRQISERLVKIETRLEDIQELKNENLAIKAELALLKADLNKRDGERG
MLATILKSPLSYLIAAGAGLIAWLKGGV
>WP_050640989.1 hypothetical protein [Eubacterium sp. SB2]
MDLEHEKRLTSVEERSKSNTHRLDKLEPIIEEIHNMSAALVEMTAELKHTNKSVEEIKDKVEDIEKEPARKWKDSTKALFNAF
LGAIGTAVAGGMIYLLTNMK

#### 3.2 Repertoire of TA proteins in selected prokaryotes

We have searched for TA proteins in selected prokaryotes. In a previous report[8], the number of TA proteins in the Euryarchaeota *Methanococcus maripaludis* was given as 12, which approximately matches that of *E. coli*[8,9] and probably of many other prokaryotes[10]. We found a comparable number of TA proteins for the aforementioned $\alpha$-proteobacterium *Thalassospira australica*. We found 19 TA proteins in *Lokiarchaeum sp. GC14_75*[11], but none featured a SNARE-like region, corroborating an earlier survey[12]. However, we uncovered 11 TA protein candidates in *H. B3-JM-08,* 33 in *H. LC_2*, and 41 TA proteins in *H. LC_3*[13]. Note that many putative TA proteins from one *Heimdallarchaeota* species were not recognized as homologs from any other inspected *Heimdallarchaeota* species, revealing that the set of TA proteins in *Heimdallarchaeota* is diverse. In addition, we noted that the repertoire of *H. LC_3* contained small subgroups of homologous TA proteins that may stem from internal duplications; some were even found in the direct genetic neighborhoods.

*Thalassospira australica*

```
>WP_033067262.1 hypothetical protein [Thalassospira australica]
MMYDENGQDDRQPVYLPETRHHWASNDEVAKPVQDLSRGPVRPRLKSKRDTGPRPSPSEWMKQNLPKIGATAIVAIIILCVVV
AIYTDVAGIVVD
>WP_033067263.1 hypothetical protein [Thalassospira australica]
MAENTPSRRGRAPGRFLPTRTVREAVKRQKPVRETPIKRFDRKDMMERDLERYRAEARARLLKNGFDIPAFLMPKPKTETNPE
DNEDEYDPAHDLDVQPPALIGPKPTPLISNANLASAKQERGQGAEKVKPRRHVHVQNKPGGLMSMFLPQDEQQALQSSQGGPA
AQRQDPTVGADAHAAAQKAAIAKEKTAYEEYIDSLDGGRTLLDHARGEDDGAGHSKNDISGRVDDLKEADARKSRIDLPDNDS
NWASQEDFAAKRKAILEKQGRAAGGRSGKDQKGSGRSKSWNKAVARFAAYVITAVAVVYVLILTVNEFSLIIGK
>WP_033068685.1 preprotein translocase subunit SecE [Thalassospira australica]
MAKTSPAQFVQQVRSEAKKVSWPTRKETTVSTIMVFVMVALASIFFFVVDQLLSWGVKLVFGVGG
>WP_033068763.1 hypothetical protein [Thalassospira australica]
MQNEPVAFTQRASNMAFAMEQQPERRKRVKVDLPENRHKWASNDEVEKPTVILRGPNYLVDDPAKLKKSPRKRYLPTIMAYAF
SGAVLITGFMYFYTEMSAILR
>WP_033069518.1 DUF393 domain-containing protein [Thalassospira australica]
MITVFYDGKCGLCSREINHYRKIASEGIFDWRDVTEHANDLEKQGVSVADGLQQLHAKDASGQLHVGVDAFILIWRQLRRWRL
LGAIVALPIIHQIAHVVYRAFGKWRFKRLTHCQLASTRGA
>WP_033069586.1 hypothetical protein [Thalassospira australica]
MSDAIMKKSYTVMCVRSTGQQMELETVSDYAAAVNSAKEALTNAKNTEIRVIENKYDQATRKDKKQVVKILTRESTQGASGSG
SSGARRGPKKPAVTREVANQATKGIINIVIAITVLVLLAGIMVPLIMR
>WP_052065538.1 DNA-3-methyladenine glycosylase I [Thalassospira australica]
MHNDNHCRCGWAEIHAAKPFYIDYHDTEWGVPVHDDRLFFEMLILEGAQAGLSWLTILARRDTYRAAYDNFDVTKIAAYDDAK
KEALLADPGIIRNKLKVAASIQNAQTFIAIQNEFGSFDSYIWDFVGGTSVINHWKTMSDVPVSTELSDRISKDLKKRGMKFVG
TTIIYSFLQATGIVMDHTTDCYRYAELS
>WP_081944293.1 aa3-type cytochrome c oxidase subunit IV [Thalassospira australica]
MRQPDVPCDDEPRAFRHLPALWLQIQHPINPWDQTISRSDVKSVKGPTLRLIDILHCTNLVDIHLQNSKPKEARLQGNDYELT
ETHVDIYNGYTSFVKWGTISVIAILVLMAIFLL
```

*Lokiarchaeum sp. GC14_75*

```
>KKK46064.1 Carboxymuconolactone decarboxylase family protein [Lokiarchaeum sp.
GC14_75]
MNEENKKDRVERNLNNDGLLRFGSSRVSPLNELEYVQGLISSRQAVNHIKFDEKSDDQSEWNDAVLNIIKELKGSPSFYMNPI
SQYLKLAEKGLDLNSINLEDFEEPIKDFIKQREYIFNLRATLMRHKDLFLRWGLFANQVFVHSDIPPREKEIIILRIAWLSQS
EYEWEHHVAGAKQAKLFSDEVIKRIKQGPEAEGWDFFDASLVRAVDELYAKNCMSDSTWKILSERYKTNQLIEILFIVGYYNL
LALTMNSLGAQVEGMYKTNQE
>KKK45694.1 hypothetical protein Lokiarch_07710 [Lokiarchaeum sp. GC14_75]
MKISLITVLKTGDLFSSNIPLVLTDLLRACHTSEPSLTAPDFACRICTFLLDTSFNFYQNSAFNLSLTVFIALKISFTIFLRE
FKIVKI
>KKK45607.1 hypothetical protein Lokiarch_08350 [Lokiarchaeum sp. GC14_75]
MILKLKAKSNKNKTVTAWIQKHKDFNDDVQQIFTFFKDKITFSKLSKITKYYVVTSTNPAIIFSLFSAIQDLIPEAYYSQLDS
MDI
>KKK45281.1 hypothetical protein Lokiarch_10990 [Lokiarchaeum sp. GC14_75]
MIIYQIKVSIISLRLLLKAENIKDISVEILFLSYILSDLSSFIWIWIL
>KKK44899.1 hypothetical protein Lokiarch_14320 [Lokiarchaeum sp. GC14_75]
MSEKDNSNKSEQKEIDKKPVLKGWNSFLKDGFDKFQKSLENQTRKNKDFWVENKEKTNKFFTGLKQEWNNKIKEWDTDIKKRK
IETKEQLDAYKKKVGEDINNWQEKTQKDWKDGFSLVRKGFFKAYFWFLILTIPILIIVVVVFAFIVKL
```

```
>KKK44804.1 hypothetical protein Lokiarch_15150 [Lokiarchaeum sp. GC14_75]
MKELKEQKAESANKSKFIPSNNLGTRIKDLSYPRFLGSIHLRKEEMNPMYNQVKNLIANENELKLNKSLKNTLFNPVTKILII
LAALSNIIWVLYSYIF
>KKK44250.1 hypothetical protein Lokiarch_19930 [Lokiarchaeum sp. GC14_75]
MNPINTETYNRTRTFIGYTPTKGKHSTSMEFVQFNEPVILSQKFLILGEIAEEMANIYKASKNGDLMELSYSYNTLAEEIKSI
SSLVKEQLFEYDWQILESSAVLADYHDPGNGGGGGNFLTCIICIAGCELAVLGGCIIACLVFIPFCPVCVLVVDLWELFDLGC
GYVCEWIGAC
>KKK44150.1 hypothetical protein Lokiarch_20780 [Lokiarchaeum sp. GC14_75]
MLKSDNEDDEIDEEEEDSRKRNKYVIICIIIILVMMLPSTMAFLGIF
>KKK42570.1 hypothetical protein Lokiarch_34060 [Lokiarchaeum sp. GC14_75]
MSQSSWKDQIKKKWGPHSHCSVCGKAMPTDKKYCSQGCKDNYMQHERKQKKKGRFQCIFLIVMLVAMMLLLFIPSLLGG
>KKK42521.1 hypothetical protein Lokiarch_34490 [Lokiarchaeum sp. GC14_75]
MKKIDFKCTLCGYKVIYNTKLESLTCSNCNTEYDKGIFRFSKKRKLGIYFKKYGFLLAVLSSIYLLYLVYRILIY
>KKK42450.1 hypothetical protein Lokiarch_34680 [Lokiarchaeum sp. GC14_75]
MTENEIQDDLKNIEKKLDDVKKINTAIRSLLLVIVFILVCFFIMSFIYFSGHFPPI
>KKK42394.1 hypothetical protein Lokiarch_35320 [Lokiarchaeum sp. GC14_75]
MSNCPNCNNPIREDQVICLNYGSQIKPLKIQRRSYKKICTIFIIFWVVMIVAMIVLLSIPSFLGG
>KKK41581.1 hypothetical protein Lokiarch_41850 [Lokiarchaeum sp. GC14_75]
MRKKRILLIGILALMVFSGIPKNVNAETEEVYSLEGSITYDKDNWRDFNGTKEEWEILYNISIPIPLKSGRDRLYWYNRFEDG
CDDLTIIFYFRPIPEGGSGGDAFYIYPDDNDVCDQRKYHDAGNFSPDNRNNTFVLVKRRGFLNPKGYLYHLTWKFWIERDIVE
NQPNDINDIIGINLMPFLLGFICIGVIIVYTKKIKN
>KKK41352.1 preprotein translocase subunit SecG [Lokiarchaeum sp. GC14_75]
MSPRSNQSRRKKRRSGNAPMPMGGAGLMRFFEDSSIGIKIGPISAVLLSTVLIVLVILAHVGVFNWLFTAVGGG
>KKK41324.1 hypothetical protein Lokiarch_43830 [Lokiarchaeum sp. GC14_75]
MKDIVNPAIIIYLPLEDGIFGNTPPGFNISITEVNLDSTWYTIQGNVTQYPLTGTIGTIDQDAWNAALEGQITIIFYAQDRAG
NIGTETVIVIKRIPSQPSIPGYNMYLLYGIVFIGLIITLQKKHKS
>KKK40990.1 hypothetical protein Lokiarch_46670 [Lokiarchaeum sp. GC14_75]
MESCAYCEKESVSSCAICGKHLCQDHILHGLSKRNTAPAVNCANCQKKFPRKLRKNIIIMTTVFIIASIFLIYYLSTIIPF
>KKK40788.1 Toxin ParE2 [Lokiarchaeum sp. GC14_75]
MNKHYILRFHPDVEGDILKIYSWYEEKLVGLGNDFLQIFYSSTKLIMENPLQYSKIYKNYHRYLFRKFPYALYYIIEDNLIIV
MGVFHHARDPRMIKSKLGNRTD
>KKK40646.1 hypothetical protein Lokiarch_49600 [Lokiarchaeum sp. GC14_75]
MSEKYNSNKSEQKETDKKPVLKGWNSFLKDGFDKFQKSLENQTKKNKDFWTENKEKTNKFFTGLNQDWDNKIKEWDADIKKRK
IETIEQFDAHKKKIGEDFNNWKEKTKKDWNDGVASVRKGFFKAYFWFLVLTIPILIIVIVVFVIINSLIG
>KKK40168.1 hypothetical protein Lokiarch_53620 [Lokiarchaeum sp. GC14_75]
MRFNGSIFEERMTTIRQTSQMDINSLREYVSSCKSDYYSELEKRVLRSYAYSKGLLGFQPMFIYKLLYILISIYITKFKLK
```

*Candidatus Heimdallarchaeota archaeon B3-JM-08*

```
>PWI47941.1 hypothetical protein CEE45_09200 [Candidatus Heimdallarchaeota archaeon
B3-JM-08]
MALLTNILTKWFKKPAEPTEKEMIKDFIQELDSVLISMEEMTEDMTNGFSDQVIQRRPLSTKLMAYLPKFMQSKNTRSQNEKL
IELKNTKQKMRTLRGTLSKIDESVDSMDLHAAPLEGEEVGSRIYELAYPGETITSSKVRTLEERLQMLENSFLASMDQINGQM
NQISSNLLTLTQRLEEQGVKIDRIDEKITDVQSKLQKIQSTLAKISRKLTQNRTLLALLVGSVIALVIVIVIL
>PWI49187.1 protein translocase SEC61 complex subunit gamma [Candidatus
Heimdallarchaeota archaeon B3-JM-08]
MGIDNSRMGSASFYKRALRIMRLAKKPDRSEVSLVIKITIAGMAILGFIAFVIRFIIFVLLGREL
>PWI48048.1 hypothetical protein CEE45_08460 [Candidatus Heimdallarchaeota archaeon
B3-JM-08]
MNNANRISQWVDEILNDTGADKIDLVGHSMGGLSSRYYIKFLDGLDKVEDYVSLGSPHHGENIATCGAQGVNALAVILNEGDE
TPGGILNDTLGNRTDPVSGIIYNSTHVPGNISYTSIYSLDDAIPSISPPLDGANNTAVEGLAHIELLSSERVYKRIKTAVKDD
FPDPTTTTTTQTTTTETSGVGILPILVIMMLVTWSKRKK
>PWI47578.1 hypothetical protein CEE45_11030 [Candidatus Heimdallarchaeota archaeon
B3-JM-08]
MKRWFEEDGRPKTALYAYDFDDKFNCSYQANINNAHKIKQWVDDILNETGAEKIDLVGHSMGGMSSRYYIKFLDGIDTVDDYV
SLGSPHHGNPNYHCGREGVKEIALILNVGDETPGGVLNDTLGIRFDPIWIEENITYNGAHILGNISYSSIYSPDDGLCPDTSS
PLDGAHNISVPNVPHSFLTEDWSVYKHVRVAVNDLKDIAFRTTPLTSTTMTTTDDTPLTLIPVLCTISVMALIVIRRRRLKTL
ANIKSNHK
>PWI48027.1 hypothetical protein CEE45_09000 [Candidatus Heimdallarchaeota archaeon
B3-JM-08]
MMSDYCVNCGQKLVQSDIFCSNCGSEVEKDKTSYSPRSSPISEYSTRRLTTQPKLYRSRGDRWIAGVCGGLGKHFSIDPILIR
IGFIIAFFGYGAGLILYIILAIFIEEEPLNSEFAPPKITDAPY
>PWI46340.1 hypothetical protein CEE45_17280, partial [Candidatus Heimdallarchaeota
archaeon B3-JM-08]
MAQEKHAYLLQGLSCTDCAMKIERTLKQEGYSSVQLNFATRRLHIGSDDVTDINSIISKIEPGAIAIPERESQEHSSAKDQEN
WLLRQIIVSTFLLLVGVLITSLS
>PWI48992.1 hypothetical protein CEE45_03495 [Candidatus Heimdallarchaeota archaeon
B3-JM-08]
MSSTEDNQDEQQPEVTMEELLDAIANFLKKTKIDQMYERMSKESSSLEFKKFAIQAFIVIIIILGTLYLVAIQILQGDQFLLI
MSLVLGYILAKADFKSV
```

>PWI47839.1 hypothetical protein CEE45_10020 [Candidatus Heimdallarchaeota archaeon B3-JM-08]
MNQVNRDIVLDKEVIWKIKKGIDFNSDAETFLEYLKKEKEMFDTFRPMSTLAPVIGILVCLTLLFLYTSMIKFV
>PWI47749.1 hypothetical protein CEE45_10225 [Candidatus Heimdallarchaeota archaeon B3-JM-08]
MKGFSSTFSRFIVGDWGTTGIMLGEQPVLSMIQRLFPASIELVLFSMTLALGMGIPLGIFLLVTK
>PWI47434.1 hypothetical protein CEE45_11500, partial [Candidatus Heimdallarchaeota archaeon B3-JM-08]
MRELEDTKFDSLAKSVGLDQEDITRIKSSKKKGSLNKLNTVYWFGQLLIFLIFLFYSVTIGI
>PWI47199.1 hypothetical protein CEE45_12925 [Candidatus Heimdallarchaeota archaeon B3-JM-08]
MLEVRYKVIQLLVLSETLMSDYTNILNAEITKEKYQWLLTRMEPFLLMFQKEIHNSNDKSHGITTSTIPNLLGIGEGSTPQSD
DLYLGIIATIKCKEPSLSEVLQNLSLIKYESYTTRKSSSLIRSFLRYNFPDEIKPIIELLKIDFPSTAHIMKFKMEIQKIKLI
GASSGYYFLVGVLWQLEYYENQKQNLIVKK

*Candidatus Heimdallarchaeota archaeon LC_2*

>OLS22354.1 hypothetical protein HeimC2_29900 [Candidatus Heimdallarchaeota archaeon LC_2]
MEQEQKQKKKKSLARKIGEKFGRGPDADELMQETLSDLTDIVGSMKHVFEDFEEGYEKQLDEERSTWRKLQSKLPSFLQAEKM
KTEDSRIKRIKSNRSNLSELEDNLKKLEIIISSGKSSMEMLGQAAINVAIPGETSEKAMNELESRINSMENNVSESMNNLNAQ
ISLIKSALDNMAGQLDEQGVVLVNIDEKIDVLDSKLDKAQEMLIKISRKLTGNRVIMLVVAGSATAVILNKLVLA
>OLS28442.1 hypothetical protein HeimC2_06610 [Candidatus Heimdallarchaeota archaeon LC_2]
MSSNDSEVYISLSAMAQMIKFDISNPGKETAGLLIGEEINNVVHVDEIRVGKQKGNAVHVEISDEELTMAAIEVSTREDGKVI
VGWWHTHPGLTSFMSGTDVKTQSMYQALMPSAIAIVIDDVKYEKTGSINDLDFGVYRLIDGKSQRLEYRIKDSVEFGLNSYVL
SDTGFDPTKSKSVTSKYYVPSMNKDKLRRLRINVDKNKNQLDPIDADSISLWLDLAESVEDGAVNEVPVDVNTLLGALDASVG
EVAIELKEMNSRMYNSQAQKTLFFIIFGLVLEFLAIFVLT
>OLS28177.1 hypothetical protein HeimC2_08320 [Candidatus Heimdallarchaeota archaeon LC_2]
MSRKEVYGMLLHDSDSLIKDYCLFKQMQNTKASTPINYTALESGYQKVKFDCSACGSVEVTVKSDNIYRRLYEFFPVKGLGYI
IGLILATLFFGSVNLLLDC
>OLS26590.1 hypothetical protein HeimC2_14290 [Candidatus Heimdallarchaeota archaeon LC_2]
MVKGDILKWYLDMDIFDELGPDISDEKYDNWSTYNDILFRALQLEYEGKDSIQLMDEFLEAVNEFCSTACTLAIENIIEDTIA
TELYLKGEKLKKEINQIYEKERSNYDILYSEYKSFITNFEEWRLELKAEKHSHKKQNYITLKQGIIIGAPILFVSIIAIIMSF
ISIIK
>OLS25240.1 hypothetical protein HeimC2_19160 [Candidatus Heimdallarchaeota archaeon LC_2]
MTNLCINCSQITTNDSDFCNSCMDKMEAGVEMSVHSTNANIIDTKSHGLFQTRNVTGKILSDDEDLIANLHRNNDQGNYYFWM
FILVIIFFMVLIAVIN
>OLS24987.1 hypothetical protein HeimC2_20420 [Candidatus Heimdallarchaeota archaeon LC_2]
MCREVLSEDFRDELKPKSEQERKRLINIMRTLDNLSDSVYRLETGKKHGKRIQKMRIDEMGDSRGPSLLKPIATVVIFLIWIV
LVIAIFA
>OLS24956.1 hypothetical protein HeimC2_20110 [Candidatus Heimdallarchaeota archaeon LC_2]
MNCGNCGNANEQGSEFCSSCGTKLNLANDSLPNSQTSTSPPPYASRGKLYRSRDDRWVGGVSGGIGEHYDIDPNMIRIIWLIL
IFAGGTGVLAYIIAWFIIPENPIVKPRSY
>OLS24138.1 hypothetical protein HeimC2_23580 [Candidatus Heimdallarchaeota archaeon LC_2]
METAAEVRDSVGDVVEDVKDKSFSTIQFATNMAAEKIEAVKHTADAAKDQTLGAFAAAKQKASNFGADVKNKIADTIWKIKDG
AISIAGKVKGALTGGLNKILMIVGIALVALIVLFVVFSKMGMMPTP
>OLS17105.1 hypothetical protein HeimC2_44940 [Candidatus Heimdallarchaeota archaeon LC_2]
MCKKKYSDTYRYNLIEENLKYHDRDLLKEFQTLAQIERKHEKRERRHDEHRLRKSQGIKEPNGLALEILLGIIVFVIGALFVI
GGDTN
>OLS29425.1 hypothetical protein HeimC2_00550 [Candidatus Heimdallarchaeota archaeon LC_2]
MFSSDEILKQLNPKILEDQVYEIQVNCLKMKRQKKFKNKKGYQQGAKYYKTKRPLEGKMNIQMFCIYCENDYSLLIRSKGETK
FNYLWHSFVLIVFLLPFFSWGLNYGYVGFPALAYYLNKMRLKWNIYVKPSNSHSSRVIREYYPIHVNFG
>OLS29184.1 hypothetical protein HeimC2_01760 [Candidatus Heimdallarchaeota archaeon LC_2]
MIDMKMSLRKPKSKKDNAPMPSTGAGLMRYFDEELPGFKISPRGVGFLTSALIFTVLLLNSTVNPFA
>OLS29024.1 hypothetical protein HeimC2_02190 [Candidatus Heimdallarchaeota archaeon LC_2]
MNTSNTETIDNSSNSIYSKMKNKFNIHKPGIWKGFCITCASTLGIVMALTVGTVLVYYTLFYLDWLGR
>OLS19481.1 hypothetical protein HeimC2_40880 [Candidatus Heimdallarchaeota archaeon LC_2] = OLS21170.1 hypothetical protein HeimC2_34680 [Candidatus Heimdallarchaeota archaeon LC_2]

MSTIDSTSLDTNSMCNKCGKLMPSYAKFCYECGLQVTSKSFGFENMDITAPKSTVGKLFDFISIAAMATFGVILAATVGFILL
YYTLFYLDYIG
>OLS28804.1 hypothetical protein HeimC2_04050 [Candidatus Heimdallarchaeota archaeon LC_2]
MDSVTTRNLVKKNQKRLESTSTLFLPEFSKKDVIKMILPLLPIHIQKSIKNLNVKFETLFEPLKSEELIKQFNDKNIYNLLIL
SELIIHNIGYMRFYVDKYRLNKND
>OLS28177.1 hypothetical protein HeimC2_08320 [Candidatus Heimdallarchaeota archaeon LC_2]
MSRKEVYGMLLHDSDSLIKDYCLFKQMQNTKASTPINYTALESGYQKVKFDCSACGSVEVTVKSDNIYRRLYEFFPVKGLGYI
IGLILATLFFGSVNLLLDC
>OLS27208.1 hypothetical protein HeimC2_13180 [Candidatus Heimdallarchaeota archaeon LC_2]
MITELTNCYNHSNVKTKWVCDRCFIAICELDHKTRGSPTNYELYCFGCYDKMKYRNTVLTFFIFVFLFGIIFGWATSDFMKWL
>OLS25712.1 hypothetical protein HeimC2_17670 [Candidatus Heimdallarchaeota archaeon LC_2]
MEDKNDEIITISKKEYEAIKKELLAFSDYQRLSSLLIWIGIAMVILHIAADVFFHIV
>OLS25240.1 hypothetical protein HeimC2_19160 [Candidatus Heimdallarchaeota archaeon LC_2]
MTNLCINCSQITTNDSDFCNSCMDKMEAGVEMSVHSTNANIIDTKSHGLFQTRNVTGKILSDDEDLIANLHRNNDQGNYYFWM
FILVIIFFMVLIAVIN
>OLS25097.1 hypothetical protein HeimC2_19690 [Candidatus Heimdallarchaeota archaeon LC_2]
MLNAKKILSARVKKLVSWKTKSIMALRSARIMGKKLQSSLAFNARIMVTSMKSKLPYSFKANRVNTAGLAVQSFKLNSYAAKG
ATPYQYNVNGPVVQSYNGGKAIQSFNGGKAILVKSGLAIAGIAITTVLISGIAVTGFAWLSRVSLLTRGFAFM
>OLS24678.1 hypothetical protein HeimC2_21590 [Candidatus Heimdallarchaeota archaeon LC_2]
MSTISQKEIQQAVNKDDLSIKPIDKKHKKENQAHQNNYGFAKNKWYSDRINFGILLYNFVRLVIVMVTTITFLELERAL
>OLS24101.1 hypothetical protein HeimC2_23750 [Candidatus Heimdallarchaeota archaeon LC_2]
MIQIDAVGWSEEYLTKTCSKCDSDNSLQFVRTQKRYKIGNFKLINSGSRHFGRECFNCFNLTPISKSDLRTVTKLDYYFKRPT
EENINRKSYYARNVKFPVLDKKKRDEIRSENIKEGLMSMVIGSIVGVIIALFYGPAIWIVPIMLLFGIYAALEDPETKFKGLI
EKSKGVRPKRRDSKRRLQ
>OLS23068.1 hypothetical protein HeimC2_28080 [Candidatus Heimdallarchaeota archaeon LC_2]
MVNHSTEENSSITLAKEHRSSARIEGEPRYTKLKKIGLVVLALLFAGPLELLIILYLA
>OLS22062.1 hypothetical protein HeimC2_31160 [Candidatus Heimdallarchaeota archaeon LC_2]
MNESFNDLIDQENDNKVDWDEVVGKSPKYIEINKFQFFLVILILVFGFLVIQNSILGMWKDGDQ
>OLS22060.1 hypothetical protein HeimC2_31140 [Candidatus Heimdallarchaeota archaeon LC_2]
MSKASIICESCGRDNELGATFCIECGTSISISKLGKSLNLEKQDPPNPNSTNIRRSEDQNDETFQSESKASSPFFMRCWHMMR
MNMNPMFIMMPIMIIVMVFIVFRRFDY
>OLS21991.1 hypothetical protein HeimC2_31390 [Candidatus Heimdallarchaeota archaeon LC_2]
MSKKEIQQATAKDGLSIKPLNTQRLIEQNIKNQGHQDNYGFGNKGWYSNKFNFEILSYNFIRFVIVVVTTISLMELERAL
>OLS21773.1 Protein translocase subunit SecE [Candidatus Heimdallarchaeota archaeon LC_2]
MGFVDTASRYYIKSKRLLFKSTRRPSRREVVTTSRIVALGLLFIGAIGFVVGLLVDFIVDSTSA
>OLS21589.1 hypothetical protein HeimC2_32930 [Candidatus Heimdallarchaeota archaeon LC_2]
MYEDQYLEDCAGLNSNFTRSEKALFIVLVLMALTGFFLAVTSILFAIL
>OLS19696.1 hypothetical protein HeimC2_40020 [Candidatus Heimdallarchaeota archaeon LC_2]
MTHSKIKIVMISLILIFSLGSIHPSNSQNIENSIYDLEITIVKFRIDGNPDENITDDNGFLEFEFYYELLDMTILDPERLEEN
LELNSSVELTDIVPFQNNTLVTKLKSIKYIQVDGRRANFNLRFWMDEHTTDNEGDEKTTKWNYNSVKIPLSNENKTISWETPY
QVLVDLMLEYKLSQSNKDDSQDGLSFNYLFSLSSLIFVALILKITKKQ
>OLS19211.1 hypothetical protein HeimC2_42180 [Candidatus Heimdallarchaeota archaeon LC_2]
MIKDSSVGRIAIIAAMVPEAINIANRSVNAESWIDRSWLLPLALINLWILIVVATYSNKFEPDSKQIISPNNLLTN
>OLS26333.1 hypothetical protein HeimC2_15770 [Candidatus Heimdallarchaeota archaeon LC_2]
MRKSPTDLTYLIGTTGNVLIWAFDANESGDEPSKYFITLDGVVIPEHDFVNWQDNVDIIVNVDGLDLGSYVVAIVANDTGTDN
NQASSTTDAAIVSVVLEIIDTDQPSDTTSDSITSTPSDDSTSSDTTLDTTTSTTGTTNTVDTQSSNSGTSDVNAFPITILSLI
FALAIGIPILKYNKY
>OLS28811.1 hypothetical protein HeimC2_04120 [Candidatus Heimdallarchaeota archaeon LC_2]
MVNKRKIKSKLFIILFLSLFNQFASAQFDLNSIHEFDLIDLDIEGKIVGGLRDEYLFQDLESVQFNVSQKILIKGLYETFLRM
KLDESRSGIIVTPENGYKDNSTDLYGETLLFNITINRIIHGDSETFTSGVAYVNPKLTRFSDFAQGFGLLIAIVLTTVLLKAV
FDYLKLRNQSQLIIR

>OLS24287.1 hypothetical protein HeimC2_23520 [Candidatus Heimdallarchaeota archaeon LC_2]
MNNIISSKGDYRNYVLDFNKLCGVCNSKIVSKDELSKWIKCESCTQGYHGNHILEVGKDGFCPICYGPLLVGVNLYLYEITGI
ESGQLQFQARKSTKTPSIQILFLKIFPFLLLFLGLGVMVSFPVSS
>OLS22046.1 hypothetical protein HeimC2_31000 [Candidatus Heimdallarchaeota archaeon LC_2]
MRNLRTYALIILFLGTNSLSQINFVHGCSFSTEVDNYITVFDLNGVKVQEKKLDYLGVGADCSITNSIFPSTSDRIYIQDEKN
KVDFVDISNDKWEVETQFQIGNIYRPLNIYQNKVILNPKIDDLLNKIKFEIVLYDLLTQEETIIKFDLIEDISQTFDNFTIIH
EQISVTSNNNWIAFFAQLQPNGCNDSCEEDVRYWLYRYNMQNSSNNTYKGILSQVYDHGFGPLDQFQISRDGRFVFFVEDYPY
NNFVSFDLDLNLSKEMTNKIPIRDKWIKEDSIYYINSENSIYSLYYLNLTNNVLETLIEFDNIYPNKIAVTDNYILIADTRPT
NNNFLAGNNTFFMIISTLFMILIIKRKLNLKSSQ
>OLS21807.1 hypothetical protein HeimC2_32180 [Candidatus Heimdallarchaeota archaeon LC_2]
MIRKSLSKSKSSSNKKKKLHKRTSKKNQKSYTCNLHPVKERYSSTSAKDFEKHLSKAHSRKRGKGVDLTMFFEVLRVIGVISI
VGFLGAIAYNSTDMVANETEAGKTIKQDNPLINGDIE
>OLS19419.1 hypothetical protein HeimC2_41460 [Candidatus Heimdallarchaeota archaeon LC_2]
MCDKHIVHKDIKDLPEHVSVFIDVQNFDLLRAELSMLGFRNTPFQIMNPGQIYAKILPTTFNNNDVELHVRCYHDGKLEAEYE
PKRLGNIIQHLSRRSYSAHEFLIESLNHLNIEHVVDQQIRERYNSKFPKEFPKQKWKFFHWFFISSMIIYPLGILWRIQWEIK
KKLRSRTSSDKSEEKGSPVTGN

*Candidatus Heimdallarchaeota archaeon LC_3*
>OLS23218.1 hypothetical protein HeimC3_25590 [Candidatus Heimdallarchaeota archaeon LC_3]
MIKNKLRALGEKICPNKEPDQIHQVKEFLEDLKDLTGSIEFLLDDVSEQYDAQVLDERPFFKKTLAKLPSFLQSSEVKKQNTQ
SKELQTVKSRVHNITLSIDKIEDLFVSMPSQEEVGKRVYELAYPGEEITVSRIEEIESRFKEMEEKFIESVKTVTDQMQTITS
SLTKISDQLEEQGIVVNNIDVKLDRVETKMDKAMVTLDKISKKISQNKVLLAFIAGVLVFALAVLFLA
>OLS27510.1 5'-nucleotidase SurE [Candidatus Heimdallarchaeota archaeon LC_3]
MQKTILLTNDDGIQSPGLKALKIELKTDYDVKIVASYTAQSAQGVSHSHGDRWVKYEKFGDGIAVHGSPATCVSVALRELGIK
PDLVVSGINFGEKLGLNIFILEQLVQLGNLQCRDIYLWQLHWNYLQIYTMSSMNLLTLAMLPILLKKL
>OLS22530.1 hypothetical protein HeimC3_30710 [Candidatus Heimdallarchaeota archaeon LC_3]
MSHTHQDKEIAERIKGLIETSGAKANLLTYEVDPSQTIIEKVKNGIKKCDLGIILWTKNSEKKEWIIQEAGALAITEKPIIVL
MESSINPPGAMLEGIHYVRFGDIEGMKSLVEWLKQRVQNEELWKIILILGGGLFLIWYLFSK
>OLS22216.1 hypothetical protein HeimC3_32200 [Candidatus Heimdallarchaeota archaeon LC_3]
MQNDVLDLLNSNKKKLEELLQEDEDWEYFFDDLIELLNNIEIIIDNLDNRIKKNYKIPPIHPDLNYDVWKKFYGKKQNEPNNN
FNKITNKLLVLTNKLLIEVKLQLSSHTIINKKTKSNLDTSFSKFGLTFPIKIKENPKEAFTQGLIWGFGATIISIILLAIPVI
LSK
>OLS22208.1 hypothetical protein HeimC3_32120 [Candidatus Heimdallarchaeota archaeon LC_3]
MEMDFFEDNFTKLFFNTETGEVNHESLSDYVQTAINVAEDVIWELALEKTSLKELNQARDELRKYILRFFDFMRWMREYDGKT
HSLRDKINIIDSAVVVIKKEMDLIDSEVATLKQNAIAEKTLKATRRAITIAYIALVISISFNLLFIIDRIFFGSN
>OLS20691.1 hypothetical protein HeimC3_39570 [Candidatus Heimdallarchaeota archaeon LC_3]
MKSNFLEDNFDELLLEEGTTDVNIELFNEYTLNVRMEITTLLFVQLRLTLDEESELERNKQKIQKYVMRVFEYVNWINNNVEN
EKERELKGEIFRDILYNHLIKDIDLLNNQILTKKQNEIAEKTLKSTNTAVKMAKYAIGLTLAVSIINLLISILGK
>OLS21438.1 hypothetical protein HeimC3_35710 [Candidatus Heimdallarchaeota archaeon LC_3]
MILVVKQHIKRWWKFQREYKKQDPELYGKKTLCRNVSHPYHDSKFEALTCRVLHKFYPAEKILVPEQHGCKTKFDFVIPGVAV
IEPHGVWENKKGEIGEYTRKRVWLANQEEMTKGLQVIVVGSMRDLKILDKKLKEFSNKRKEKRRGNLIWRMVIIYEILLIMII
VL
>OLS20559.1 hypothetical protein HeimC3_40170 [Candidatus Heimdallarchaeota archaeon LC_3]
MGSKKDMIKLYSYPKRIVRWWNFHQNFKRKNLHRWNYGRPGLCRNINHPYHLSQFEISCCNILSEHFDQSEILVPSQHKCKTK
FDFIIKERIIVEPHASWRWNNNKEYMNYYYRRKHLALKEEKTKNLPVIVIPSIIDLRIMRIFLHKYHDPIKALKQLGLNLINK
YSVEPLIKVDYQKRTKKSKILIPLALYELIIGIVGIATNLSYSY
>OLS20562.1 hypothetical protein HeimC3_40200 [Candidatus Heimdallarchaeota archaeon LC_3]
MDYYYKRKHLAQREEKTRNLPVLVLSSISDVRSMKDYLEKYIDPIHALKRLRLNLINKYSIEPINKIEYHKKQNLSKILIPLA
LYELILGIAGVITFF
>OLS25611.1 hypothetical protein HeimC3_13590 [Candidatus Heimdallarchaeota archaeon LC_3]
MGSKNILSNLYSYPKRIVRWWNFHLIFKRKNRHRWKFSRPGLCENINHPYHLSQFEIICCKILSQYFNQSEILTPSQHKFKTK
FDFIIKDNIILEPHASWKWSKTNEYMDYYYKRKHLALREEKTRNLPVLVLSSISDVRSMKNYLEKYNDPIHALKRLRLNLINK
YSIEPIKKIDYHKKQKLSKILIPLALYELILGIAGEIIGVYVSDKVMEEIIDGKNVQHYIP
>OLS16874.1 hypothetical protein HeimC3_51040 [Candidatus Heimdallarchaeota archaeon LC_3]

MVFSLKFVKRIIRWLKFHLFIKNKFKKRFGNPKLCNNLEHRYHDSNFEKETCLILEKYFHQKSIIVPSQHKLKTKFDFIIKGI
AIIEPHGIWNGNGFYSYYKKRKILAEKFDLDLPVIIIPNYSDLTQFDTYLQKENNPVLAMKQFQLLMLNRYQDSNIVHSPIIA
TRISEKWKSFSLLNLSIIIIQFVYIFYLS
>OLS16648.1 hypothetical protein HeimC3_52320 [Candidatus Heimdallarchaeota archaeon LC_3]
MLTKVFTYPKRIVRWWNFYQNFKRKNHHRWNYGRPGLCRNINHPYHLSQFEISCCNILSKHFDQSEILVPSQHKCKTKFDFII
KERIIIEPHASWKWNNNNEYMNYYYKRRDLALKERRTSNLPVLVLSSITDFKKMIMFFNNYQDPINALKHLRLNLINKYSVEV
IKKVDYKKKLKKSKIVIPLVLYEAILGIAGILTFL
>OLS20991.1 hypothetical protein HeimC3_37400 [Candidatus Heimdallarchaeota archaeon LC_3]
MEKDQKLKAEKPPYHKLESNENYTPSYDGLKSNENYTPSYDGLKSNENFTPSYDGLKSKKFIPSYDSLKA
DNKFKPTYGRPRRSILTLPFRHYKTLLILVVLMVIVYLLITFL
>OLS16932.1 hypothetical protein HeimC3_50780 [Candidatus Heimdallarchaeota archaeon LC_3] = OLS21012.1 hypothetical protein HeimC3_37610 [Candidatus Heimdallarchaeota archaeon LC_3]
MEKDQKLKAEKPPYHKLESNENYTPSYTGLKSNENFTPFYDGLKSNENFIPSYDGLKSKKFIPSYDSLKADNKFKPTYGRPRR
SILTLLFRHYKTLLILVVLMVIVYLLTTFL
>OLS16942.1 hypothetical protein HeimC3_50880 [Candidatus Heimdallarchaeota archaeon LC_3]
MKTRIKLESNENFTPSYDGLKSNENFTPSYDGLKSKKFIPSYDSLKADNKFKPTYGRPRRSILTLPFRHYKTLLILVVLIVIV
YIITTFL
>OLS19958.1 hypothetical protein HeimC3_42840 [Candidatus Heimdallarchaeota archaeon LC_3]
MNKTNIVNEKKILKEVQKSYCVLHTKRRAYRLCDNCYLPYCKEDIVESWSHNFLSYAYLGSKKEFKKQSLCKSCERRKRRNSV
GFAVFLLIVFGIFILGFAFNP
>OLS19959.1 hypothetical protein HeimC3_42850 [Candidatus Heimdallarchaeota archaeon LC_3]
MVESYSNEDDIKTKDQLDLEYLNQDIIGNSLCYRLSCFDKAEKYCEKCNEVFCSSHLNVYWSQNFLQHAFVGQKRQFISETLC
HKCERSNRFLGVFLAFFLLFPFLFTPLLILFG
>OLS20352.1 hypothetical protein HeimC3_40910 [Candidatus Heimdallarchaeota archaeon LC_3]
MNQTNGTPKLDALRTTLIDAISDPEIVPPYELKLLYISTFHLPTDTPEMMQLHEIQFGYKKEQAEMKREMKFMRYMIIGLAIG
SGLLKIITVDVLEAFLRGWGLF
>OLS19858.1 hypothetical protein HeimC3_42970 [Candidatus Heimdallarchaeota archaeon LC_3]
MVSISRSSKKKDLHRRPSIQKTRIINRFQRYDAFSAQTGKTLKQIDLTEKKYQSFIETLISEGILKTSSASGTAKYWLKAEKV
NPKEKDNKFLITSVVFAVTFVIIMFIFFAFS
>OLS28094.1 hypothetical protein HeimC3_00540 [Candidatus Heimdallarchaeota archaeon LC_3]
MDKIIIPDVETAIGTIAEKPFTTQRRNVQLDEIGLLFESKKFEWTSLNDVKVKFFLGNPYIQMNFENDQKISLWFPTKWYKLR
RNPWFASSEQLTMDFIEKVSSFQSDPSLKTKIQSQISESQNSLTLVGRIYQTSTNAVFLLLPLITLYLLISFILWVMDNPGGV
FEL
>OLS27280.1 hypothetical protein HeimC3_04510 [Candidatus Heimdallarchaeota archaeon LC_3]
MNEREFSQSKEEINNKEMFSDDRMDALLKESLQYFQISVVIILLMVLVTLIHMGWSHVLHIQDLIF
>OLS27279.1 hypothetical protein HeimC3_04500 [Candidatus Heimdallarchaeota archaeon LC_3]
MSENRINDKPSIVDEILDKKNNFERNQKIIFVFSIIIFLITLSHVIMDVFFH
>OLS27278.1 hypothetical protein HeimC3_04490 [Candidatus Heimdallarchaeota archaeon LC_3]
MDEEKNLAKSAPSFPMPKESVMSDKYTDPKEKRLANWIEDVVTMQRVNLVLLIVTLMMLVLHLVLDFFFHV
>OLS26012.1 hypothetical protein HeimC3_12860 [Candidatus Heimdallarchaeota archaeon LC_3]
MKIDPLLQCYELFDQYKFKKALLLAKQKINHIELESETRRNLVGLIIWCYNFLGNFEKCLENIKVLEKNLGKTMPEWHLGLLN
YNKALIIKRKENKEKASAIFRKAASHYKAHRENEIESFPDNQQLVLKNLNMLVLILIFMVIVLKEQD
>OLS25649.1 hypothetical protein HeimC3_13970 [Candidatus Heimdallarchaeota archaeon LC_3]
MDNYCYIHTNRPKTTYCDKSKKNICEEDTRRISSGIGNEETVMVYCPNCYSSVKKENLLNWILYLVAVITFGIIVILISIVGG
IGFF
>OLS26791.1 hypothetical protein HeimC3_06830 [Candidatus Heimdallarchaeota archaeon LC_3]
MRRFFHTTSPLFAKSLAVLFFKEKLIVKIILGTLFLIFGIWILFT
>OLS25039.1 hypothetical protein HeimC3_16690 [Candidatus Heimdallarchaeota archaeon LC_3]
MEEDLELKGLNESRIISRLEELITLIKVILFVNILILATLVITAIF
>OLS24889.1 hypothetical protein HeimC3_17650 [Candidatus Heimdallarchaeota archaeon LC_3]
MSSRRSKKSKSDNPMPATGAGLIRFYSESDSPGIKVGPRLTVFFAIFLIIFILAVNILIKPGA
>OLS16506.1 Protein translocase subunit SecE [Candidatus Heimdallarchaeota archaeon LC_3]

MGFLNDSVRLIKIAKKPSLKEIWLISKVTIVGMSILGLIGFIIYVVGRLIFESIR
>OLS24865.1 hypothetical protein HeimC3_17400 [Candidatus Heimdallarchaeota archaeon LC_3]
MFDVSLDPQRELVIFHMNNPNENNELLFKHKKKQKPNFPTDNRMKLIIFFCLVMVLSFLFALLLASPLRFWF
>OLS24327.1 hypothetical protein HeimC3_20450 [Candidatus Heimdallarchaeota archaeon LC_3]
MIVSQSGDCSEYVKFNLDISIGYFRKIDKEEFPITEITNDLVKELIYIQGNYGMLLERLFYVISLSQNLLLISLFASILLIET
YYLKF
>OLS23965.1 hypothetical protein HeimC3_21900 [Candidatus Heimdallarchaeota archaeon LC_3]
MSFCNVCGEEIGNANFCSNCGTAQDNEPPQSKNLPQPPTFQTQAPSQQGMQGMMQNISPMMIIMIIAMIIMMVFMFIFIFPTI
FNFMNR
>OLS23915.1 hypothetical protein HeimC3_22580 [Candidatus Heimdallarchaeota archaeon LC_3]
MILIFFVRINNTSTNFFNFEGSNMIKNNLSYADFEEVSLKDLGIVDETWVEIKIKENLSMLKLGNLTFTIIIIIFPFLNHF
>OLS23255.1 hypothetical protein HeimC3_25960 [Candidatus Heimdallarchaeota archaeon LC_3]
MKASIDPPNSKDFFLKIREDYNEYKRQTNGHLTGIHALVLVEEAEDLYYNIATLPTVYDLKEADQIERYLKEYTQMMTEFHWL
YSRVKLSTTNTSEDDFTDRIFALNKVEGLINIIWNNLNHLKQSKNELFVKDQLELAEIQNDLVRESLKRIDKQIKVAWITFGV
ASIAVLISIFMYIIA
>OLS22919.1 hypothetical protein HeimC3_28620 [Candidatus Heimdallarchaeota archaeon LC_3]
MELFSEQIIGLRQVFYNKKDDGSEILSKFKVFIFYALFLVMFLFS
>OLS22622.1 hypothetical protein HeimC3_28960 [Candidatus Heimdallarchaeota archaeon LC_3]
MTQKVCPNCGKEISESQFCLHCGMPLTDLARKSPTQRAIKNETKEMSNLWIFLIMCLCLTPIGGIIFLFLIPTFFF
>OLS21976.1 hypothetical protein HeimC3_33220 [Candidatus Heimdallarchaeota archaeon LC_3]
MITCPAIRIKKPRNPSNDIPTTVTLITTHNSFRDGFLAILNKRAESEKNPKNSNQTFICLFLITLFIKLILYY
>OLS21688.1 hypothetical protein HeimC3_34420 [Candidatus Heimdallarchaeota archaeon LC_3]
MSVRNLMIEKQKFSLYSSHLEHADLIELTINQNNLKRLKDMDKFPNSSMLAVIPIITLISSIITAIFGFIF
>OLS20677.1 hypothetical protein HeimC3_39430 [Candidatus Heimdallarchaeota archaeon LC_3]
MQIKKKIDGRQLIIDWSVWSSKLIIEYDNETIHETWEITFWVTVAGLFTLLIGIFFGIYIGMLYF
>OLS17029.1 hypothetical protein HeimC3_50030 [Candidatus Heimdallarchaeota archaeon LC_3]
MTENYNKFYLSETDYNLLSEYKNLSNNEKKILEDFRYKPKGYIKIHKDDKERRIKIVNSFITPVLVFSLT
YLRGMLS
>OLS16588.1 hypothetical protein HeimC3_53340 [Candidatus Heimdallarchaeota archaeon LC_3]
MSYSADQMVIFLLWVGHDVRNEIGINQKYIGETVETVKNINDYWRIFFTTFKAVSIYSFISFSVTVFVSF
DTI

##### 3.3 Putative Get3 homologs in selected Asgard archaea

The ATPase Get3, which is at the heart of the GET pathway, is conserved across eukaryotes and is also present in archaea[14-16]. Through a Blast search, we found putative Get3 homologs in *Lokiarchaea* and some *Heimdallarchaeota,* but not in *H. LC_2* and *H. AB_125*.

*Lokiarchaeum sp. GC14_75*

```
>KKK42590.1 putative arsenical pump-driving ATPase [Lokiarchaeum sp. GC14_75]
MELKEQLLKLKIIMFGGKGGVGKTSCAASSAIWAADHGRNTLIISTDPAHSLGDSLGINLLPGIPTPIEGIENLTALEINPKV
NMAEYQGLTNINPMEEMNIPGIMENMSLFGDLEELSSMSPPGIDEALAFGKILEFIETEHDYDLVIFDTAPTGHTLRFLSLPE
TLSGWIGKLIKMRVSFGKMFGAVKRLFTRQEKKEDNSLEILEKLNNSIINARDDLMNPVKNSFIIVMIAEEMAITETGRLLNE
LIKQNIPVHTIVVNQLYLDREELCKFCKARREMQQKNLLKVIEIFSENFHKNIIQVPLFKDEIREYDKLKEMSEFLIEKV
```

*Candidatus Heimdallarchaeota archaeon B3-JM-08*

```
>PWI49592.1 arsenic-transporting ATPase [Candidatus Heimdallarchaeota archaeon B3-
JM-08]
MSNSIFSKHFLFFGGKGGVGKTTMAAATAIRAADLGHNTLLVSTDPAHSVSDSLDQQIGGDDYVKVNNVDNLWAIELSTDKAM
STYSEMISQQDPTGAFNELLGDGDANSLSPPGTDETVAFIQLLEFIQNPEYDIVVFDTAPTGHTLKLLQLPELTQNWLFRLIK
MRRRIGGLMSGFKSLIGGGTDLDEQDAFDKLEELRDQVEIARTHLSNNEETEFVAVTIPTVMAIWETERLIRTLFEVAFPIKR
IYINQLQPDNPDCTYCMNRYTDQLKNLGKIKDLYDEFDLQEIPSFEYEIRGIVHLRELANLLYGVRT
```

*Candidatus Heimdallarchaeota archaeon LC_3*

```
>OLS20412.1 putative arsenical pump-driving ATPase [Candidatus Heimdallarchaeota
archaeon LC_3]
MIRLKTKLIGGKGGVGKTTIASSIAIYHALKGLKTLVISTDPAHSLADCLDQEIGSEIVPVRKIKNLWALEIDSEKATQEYGN
LLVQQGFDQSSIFSQFLGGDDISSLTPPGADETVAFLKLLEFIENPLEYEVIIYDTAPTGHTLKLLSLPELTSNWLFKIAMLR
QKLSSTLGGIKKIFGGGGKNVNTADMKQSIDVLRKRIESAREHLQNHEETEFIPITIPTLMSIWETERLLQALRQYGISAKTI
IVNQVNPENDKCDFCKLKHKQHNSIIDQLKDLYSDEYRIHTIEMFKDEIRGIDNLIEFNSKISSIFENN
```

#### References – Section 3

1 Kutay, U., Hartmann, E. & Rapoport, T. A. A class of membrane proteins with a C-terminal anchor. *Trends Cell Biol* **3**, 72-75 (1993).

2 Beilharz, T., Egan, B., Silver, P. A., Hofmann, K. & Lithgow, T. Bipartite signals mediate subcellular targeting of tail-anchored membrane proteins in Saccharomyces cerevisiae. *The Journal of biological chemistry* **278**, 8219-8223, doi:10.1074/jbc.M212725200 (2003).

3 Kalbfleisch, T., Cambon, A. & Wattenberg, B. W. A bioinformatics approach to identifying tail-anchored proteins in the human genome. *Traffic (Copenhagen, Denmark)* **8**, 1687-1694, doi:10.1111/j.1600-0854.2007.00661.x (2007).

4 Chio, U. S., Cho, H. & Shan, S. O. Mechanisms of Tail-Anchored Membrane Protein Targeting and Insertion. *Annu Rev Cell Dev Biol* **33**, 417-438, doi:10.1146/annurev-cellbio-100616-060839 (2017).

5 Mateja, A. & Keenan, R. J. A structural perspective on tail-anchored protein biogenesis by the GET pathway. *Current opinion in structural biology* **51**, 195-202, doi:10.1016/j.sbi.2018.07.009 (2018).

6 Mateja, A. *et al.* Protein targeting. Structure of the Get3 targeting factor in complex with its membrane protein cargo. *Science (New York, N.Y* **347**, 1152-1155, doi:10.1126/science.1261671 (2015).

7 Zimmer, J., Nam, Y. & Rapoport, T. A. Structure of a complex of the ATPase SecA and the protein-translocation channel. *Nature* **455**, 936-943, doi:10.1038/nature07335 (2008).
8 Borgese, N. & Righi, M. Remote origins of tail-anchored proteins. *Traffic (Copenhagen, Denmark)* **11**, 877-885, doi:10.1111/j.1600-0854.2010.01068.x (2010).
9 Lutfullahoglu-Bal, G., Seferoglu, A. B., Keskin, A., Akdogan, E. & Dunn, C. D. A bacteria-derived tail anchor localizes to peroxisomes in yeast and mammalian cells. *Scientific reports* **8**, 16374, doi:10.1038/s41598-018-34646-7 (2018).
10 Craney, A., Tahlan, K., Andrews, D. & Nodwell, J. Bacterial transmembrane proteins that lack N-terminal signal sequences. *PLoS ONE* **6**, e19421, doi:10.1371/journal.pone.0019421 (2011).
11 Spang, A. *et al.* Complex archaea that bridge the gap between prokaryotes and eukaryotes. *Nature* **521**, 173-179, doi:10.1038/nature14447 (2015).
12 Klinger, C. M., Spang, A., Dacks, J. B. & Ettema, T. J. Tracing the Archaeal Origins of Eukaryotic Membrane-Trafficking System Building Blocks. *Mol Biol Evol* **33**, 1528-1541, doi:10.1093/molbev/msw034 (2016).
13 Zaremba-Niedzwiedzka, K. *et al.* Asgard archaea illuminate the origin of eukaryotic cellular complexity. *Nature* **541**, 353-358, doi:10.1038/nature21031 (2017).
14 Farkas, A., De Laurentiis, E. I. & Schwappach, B. The natural history of Get3-like chaperones. *Traffic (Copenhagen, Denmark)* **20**, 311-324, doi:10.1111/tra.12643 (2019).
15 Sherrill, J., Mariappan, M., Dominik, P., Hegde, R. S. & Keenan, R. J. A conserved archaeal pathway for tail-anchored membrane protein insertion. *Traffic (Copenhagen, Denmark)* **12**, 1119-1123, doi:10.1111/j.1600-0854.2011.01229.x (2011).
16 Suloway, C. J., Rome, M. E. & Clemons, W. M., Jr. Tail-anchor targeting by a Get3 tetramer: the structure of an archaeal homologue. *EMBO J* **31**, 707-719, doi:10.1038/emboj.2011.433 (2012).

#### Section 4: Description of *bona fide* SNARE proteins from γ-proteobacteria of the order Legionellales

##### 4.1 Different types of SNARE proteins were found in γ-proteobacteria

The best-scoring sequences of our HMM screen for prokaryotic SNARE sequences came from γ-proteobacteria of the order Legionellales, bacterial pathogens that live inside eukaryotic cells[1]. SNARE proteins operate via a fundamental mechanism in eukaryotic vesicle trafficking: their sequential assembly into stable, heterologous membrane-bridging complexes pulls membranes together[2-6]. Our analysis demonstrates that the bacterial sequences belong to different types of SNARE proteins[7]. Several clusters contained R-SNAREs, one cluster contained a Qc-SNARE, and two clusters contained Qbc-SNARE proteins, which have two different SNARE domains connected by a longer linker. Moreover, we found Qa.IV-SNAREs. Most of the *bona fide* SNARE sequences grouped into six different clusters (Cluster 0–5), whereas six sequences were found as singletons and are listed in Cluster -1 (Supplementary Table 1). An overview of the different domain organizations is given in Fig. S4.1.

Three clusters (i.e. Clusters 0, 1, and 3) contained R-SNARE sequences (Extended Data Table 2). The sequences in Cluster 0 are from genomes of various *Legionella* species, whereas the three R-SNARE sequences in Cluster 1 originate from metagenomes of different unclassified γ-proteobacteria[8]. A related R-SNARE sequence (WP_114833497.1) was found in *Aquicella lusitana* (Legionellales, Coxiellaceae), suggesting that the unclassified γ-proteobacteria might be from a related lineage. We noted that the R-SNARE sequences in Clusters 0 and 1 carry a *C*-terminal CAAX motif. A *C*-terminal CAAX motif is also found in Ykt6, whereas all other eukaryotic R-SNAREs usually have a *C*-terminal transmembrane domain. It is therefore possible that the R-SNARE sequences of Clusters 0 and 1 are related, although they are very divergent and are found in different lineages of γ-proteobacteria. Another R-SNARE sequence found in the metagenome of γ-proteobacteria bacterium RIFCSPHIGHO2_12_FULL_37_34[8], which did not cluster with the other R-SNARE does not carry a *C*-terminal CAAX domain. The above-mentioned bacterial R-SNAREs have a longer *N*-terminal region, which, according to secondary structure predictions[9,10], folds into a longin domain, which can be found in three different R-SNARE types, Sec22 (R.I-type), Ykt6 (R.II-type) and Vamp7 (R.III-type)[11]. Cluster 3 contained R-SNARE sequences from *Berkiella cookevillensis* (Legionellales, Coxiellaceae)[12] and an unclassified γ-proteobacterium, γ-proteobacteria bacterium 39-13[13]. An R-SNARE sequence from *Berkiella aquae* (KRG20970.1) is closely related. By contrast, the R-SNARE sequences contained in Cluster 3 do not have a longin domain and appear to have two consecutive transmembrane domains following their SNARE domain. It is likely that only the short linker between the two TMDs faces the intravesicular (= extracellular) side, whereas the SNARE motif and the longer *C*-terminal extension face the cytoplasmic side. The R-SNAREs of Cluster 3 are probably not directly related to the other bacterial R-SNAREs.

Cluster 2 contains highly similar Qc-SNARE protein sequences, which are expressed by different *Legionella pneumophila* strains. This protein has been described earlier[14].

Clusters 4 and 5 contained Qbc-SNARE proteins (i.e. SNAP-25-like sequences), which have two different consecutive SNARE domains, a Qb- and Qc-motif, connected by a longer linker. Some eukaryotic Qbc-SNAREs (e.g. SNAP-25, which plays a role in neuronal secretion) are known to be attached to membranes by palmitoylation of cysteines in the linker region, whereas others do not have cysteines for post-translational modifications. It remains unclear whether the bacterial Qbc-SNAREs undergo post-translational modifications. The sequences from Cluster 4 are from *Fluoribacter gormanii* and *Legionella cherrii*, and seem to be related to a SNAP-25-like protein from the amoebae *Acanthamoeba castellanii* (XP_004368292.1), which is an established host for intracellular bacterial pathogens. The two almost identical sequences in Cluster 5 are from *Berkiella cookevillensis*. Another divergent Qbc-SNARE sequence was found in the metagenome of *γ-proteobacterium bacterium RIFCSPHIGHO2_12_FULL_41_20*. It is unclear whether these different SNAP-25-like SNARE sequences ended up in bacterial genomes through only one lateral gene transfer event or through many.

The metagenome of *Berkiella cookevillensis* also codes for a Qa.IV-SNARE (WP_057625376.1). Usually, Qa.IV-SNARE proteins are involved in secretion. We noted that the other two SNARE sequences from *Berkiella cookevillensis* mentioned above belong to secretory types of SNARE proteins as well. As mentioned in the main text, *Berkiella cookevillensis* encodes for three different secretory SNARE proteins, an R-, a Qbc-, and a Qa-SNARE, which could in principle assemble into one SNARE complex. *Berkiella cookevillensis* has only recently been described as a intranuclear bacterium[15] of amoebae[12,16,17] and not much is known about its life cycle and the role of the unusual set of SNARE proteins is unclear. A different Qa.IV-SNARE sequence (WP_058512658.1) is present in the genome of *Legionella santicrucis*. Both bacterial Qa.IV SNAREs have a *C*-terminal CAAX motif but do not possess a longer *N*-terminal domain, which, in eukaryotic Qa-SNAREs, consists of an independently folded three-helix bundle domain, termed the Habc domain.

We also found a sequence (OUW42374.1) with a Qb.III-type SNARE motif from a metagenome of the unclassified bacterium TMED181[18]. A BLAST search revealed that the sequence is very similar to a putative Vti1 sequence (XP_002505281.1) from the green alga *Micromonas commoda*. It is therefore unclear whether the sequence indeed represents a bacterial sequence or a sequence from green algae closely related to *Micromonas commoda*.

A SNARE sequence of the Qa.III-type (WP_062266217.1) was found in the genome of *Endozoicomonas arenosclerae* (γ-proteobacteria, Oceanospirillales)[19]. A BLAST search showed that the sequence is related to a syntaxin-7-like protein from the demosponge *Amphimedon queenslandica*. As the bacterial sequence was obtained from the flora of the marine sponge *Arenosclera brasiliensis*, we concluded that this sequence instead represents a syntaxin-7-like sequence from the demosponge *Arenosclera brasiliensis* than from a bacterium.

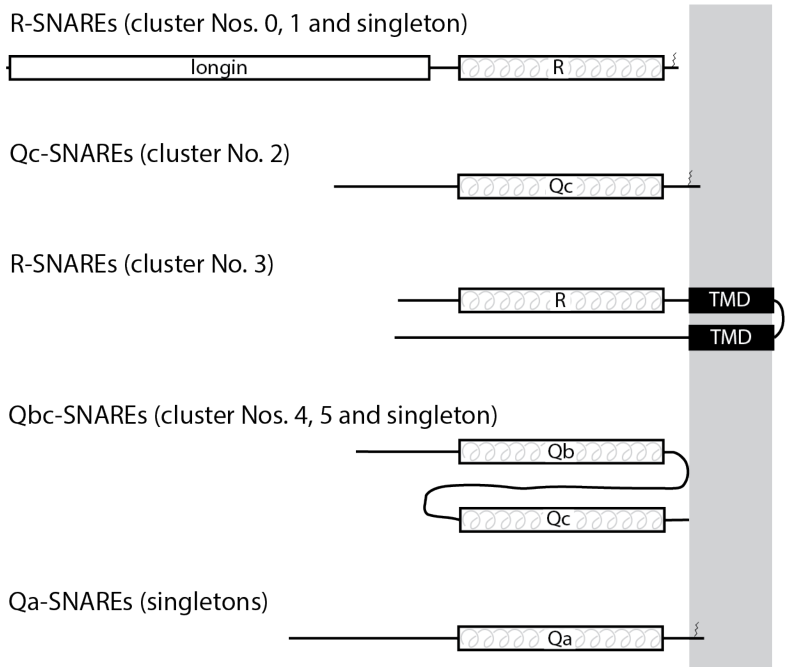

**Fig. S4.1. Domain organization of the different SNARE protein types found in γ-proteobacteria of the order Legionellales.**

The different types of SNARE motifs are indicated. Note that most R-SNAREs of Legionellales were predicted[10 9] to possess an *N*-terminal longin domain[11]. They also harbor a *C*-terminal CAAX motif (C = cysteine, A = aliphatic amino acid, X = terminal residue)[20] which is farnesylated for membrane anchoring. CAAX motifs are also present in bacterial Qc- and Qa-SNAREs. These proteins do not seem to have an independently folded *N*-terminal domain. The membrane is shaded in gray.

#### 4.2 Formation of stable complexes between neuronal SNARE proteins and those from different γ-proteobacteria of the order Legionellales

As described in the main text, we demonstrated through non denaturing gel electrophoresis that two different SNARE proteins from different γ-proteobacteria can form a stable ternary SNARE complex with neuronal SNARE proteins (Fig. 3). In contrast to the ternary complex formed by the three neuronal SNARE proteins, the ternary complexes containing bacterial SNARE proteins were not SDS-resistant (Fig. S4.2).

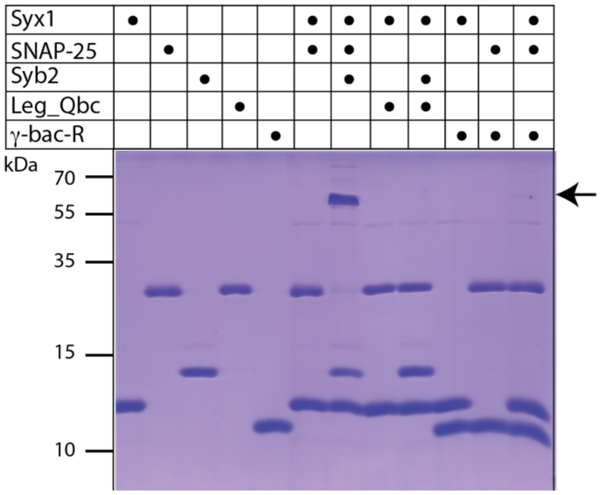

**Fig. S4.2. Combinations of neuronal SNARE proteins and two SNARE proteins from γ-proteobacteria of the order Legionellales analyzed by SDS-PAGE.**

As described in the legend of Fig. 3, a SNAP-25 like SNARE protein from *Legionella cherrii* (Leg_Qbc, WP_028380397.1) or an R-SNARE from *Gammaproteobacteria bacterium RIFCSPHIGHO2_12_FULL_42_13* (µ-bac-R, aa 116-176, OGT53257.1) were mixed with the neuronal SNARE proteins synaptobrevin (Syb2, R-SNARE), SNAP-25 (Qbc-SNARE) and the SNARE motif of syntaxin (Syx1, Qa-SNARE). The proteins were incubated overnight at 4°C with about equimolar ratios at about 15 µM concentration prior to separation by SDS-PAGE. Note that the samples were not boiled before SDS-PAGE to separate the SDS-resistant ternary complex formed by the three neuronal SNARE proteins from monomers. The 65-kDa band corresponding to the SDS-resistant ternary complex is indicated by an arrow. Note that the ternary complexes formed by the bacterial SNAREs and neuronal proteins are not SDS-resistant[21,22].

#### References – Section 4

1 Duron, O., Doublet, P., Vavre, F. & Bouchon, D. The Importance of Revisiting Legionellales Diversity. *Trends Parasitol* **34**, 1027-1037, doi:10.1016/j.pt.2018.09.008 (2018).

2 Jahn, R. & Scheller, R. H. SNAREs — engines for membrane fusion. *Nature Publishing Group* **7**, 631-643, doi:10.1038/nrm2002 (2006).

3 Sudhof, T. C. & Rothman, J. E. Membrane fusion: grappling with SNARE and SM proteins. *Science (New York, N.Y* **323**, 474-477 (2009).

4 Jahn, R. & Fasshauer, D. Molecular machines governing exocytosis of synaptic vesicles. *Nature* **490**, 201-207, doi:10.1038/nature11320 (2012).

5 Hong, W. & Lev, S. Tethering the assembly of SNARE complexes. *Trends Cell Biol* **24**, 35-43, doi:10.1016/j.tcb.2013.09.006 (2014).

6 Wang, T., Li, L. & Hong, W. SNARE proteins in membrane trafficking. *Traffic (Copenhagen, Denmark)* **18**, 767-775, doi:10.1111/tra.12524 (2017).

7 Kloepper, T. H., Kienle, C. N. & Fasshauer, D. An elaborate classification of SNARE proteins sheds light on the conservation of the eukaryotic endomembrane system. *Molecular biology of the cell* **18**, 3463-3471, doi:10.1091/mbc.E07-03-0193 (2007).

8 Anantharaman, K. *et al.* Thousands of microbial genomes shed light on interconnected biogeochemical processes in an aquifer system. *Nat Commun* **7**, 13219, doi:10.1038/ncomms13219 (2016).

9 Kelley, L. A., Mezulis, S., Yates, C. M., Wass, M. N. & Sternberg, M. J. The Phyre2 web portal for protein modeling, prediction and analysis. *Nat Protoc* **10**, 845-858, doi:10.1038/nprot.2015.053 (2015).

10 Lobley, A., Sadowski, M. I. & Jones, D. T. pGenTHREADER and pDomTHREADER: new methods for improved protein fold recognition and superfamily discrimination. *Bioinformatics (Oxford, England)* **25**, 1761-1767, doi:10.1093/bioinformatics/btp302 (2009).

11 Daste, F., Galli, T. & Tareste, D. Structure and function of longin SNAREs. *J Cell Sci* **128**, 4263-4272, doi:10.1242/jcs.178574 (2015).

12 Mehari, Y. T., Arivett, B. A., Farone, A. L., Gunderson, J. H. & Farone, M. B. Draft Genome Sequences of Two Novel Amoeba-Resistant Intranuclear Bacteria, "Candidatus Berkiella cookevillensis" and "Candidatus Berkiella aquae". *Genome Announc* **4**, doi:10.1128/genomeA.01732-15 (2016).

13 Kantor, R. S. *et al.* Genome-Resolved Meta-Omics Ties Microbial Dynamics to Process Performance in Biotechnology for Thiocyanate Degradation. *Environ Sci Technol* **51**, 2944-2953, doi:10.1021/acs.est.6b04477 (2017).

14 King, N. P. *et al.* Soluble NSF attachment protein receptor molecular mimicry by a Legionella pneumophila Dot/Icm effector. *Cell Microbiol* **17**, 767-784, doi:10.1111/cmi.12405 (2015).

15 Schulz, F. & Horn, M. Intranuclear bacteria: inside the cellular control center of eukaryotes. *Trends Cell Biol* **25**, 339-346, doi:10.1016/j.tcb.2015.01.002 (2015).

16 Mehari, Y. T. *et al.* Description of 'Candidatus Berkiella aquae' and 'Candidatus Berkiella cookevillensis', two intranuclear bacteria of freshwater amoebae. *Int J Syst Evol Microbiol* **66**, 536-541, doi:10.1099/ijsem.0.000750 (2016).

17 Chamberlain, N. B. *et al.* Infection and nuclear interaction in mammalian cells by 'Candidatus Berkiella cookevillensis', a novel bacterium isolated from amoebae. *BMC Microbiol* **19**, 91, doi:10.1186/s12866-019-1457-z (2019).
18 Tully, B. J., Sachdeva, R., Graham, E. D. & Heidelberg, J. F. 290 metagenome-assembled genomes from the Mediterranean Sea: a resource for marine microbiology. *PeerJ* **5**, e3558, doi:10.7717/peerj.3558 (2017).
19 Appolinario, L. R. *et al.* Description of Endozoicomonas arenosclerae sp. nov. using a genomic taxonomy approach. *Antonie Van Leeuwenhoek* **109**, 431-438, doi:10.1007/s10482-016-0649-x (2016).
20 Wang, M. & Casey, P. J. Protein prenylation: unique fats make their mark on biology. *Nature reviews* **17**, 110-122, doi:10.1038/nrm.2015.11 (2016).
21 Fasshauer, D., Eliason, W., Brunger, A. & Jahn, R. Identification of a minimal core of the synaptic SNARE complex sufficient for reversible assembly and disassembly. *Biochemistry* **37**, 10354-10362, doi:10.1021/bi980542h (1998).
22 Fasshauer, D., Otto, H., Eliason, W., Jahn, R. & Brunger, A. Structural changes are associated with soluble N-ethylmaleimide-sensitive fusion protein attachment protein receptor complex formation. *Journal of Biological Chemistry* **272**, 28036-28041, doi:10.1074/jbc.272.44.28036 (1997).

#### Section 5: Larger clusters of proteins with SNARE-like regions

##### 5.1 Prokaryotic signaling proteins contained in a large cluster

Finally, we inspected sequences that scored less well in our initial screen (i.e. those with e-values of ~ $1E^{-10}$ or more). Very probably, these moderately scoring sequences include false-positives. As the original HMMs were trained on the coiled-coil stretch of eukaryotic SNARE proteins only, it was likely that our screen had also detected unrelated prokaryotic coiled-coil proteins[1,2], which may be challenging to tell apart from a prototypical SNARE protein. Indeed, notable clusters contained distinct prokaryotic proteins with extended coiled-coil segments that have remote similarity to the coiled-coil pattern found in SNARE proteins.

Most of moderately scoring sequences were contained in the largest cluster, Cluster 6 (Supplementary Table 1) with about 4200 different prokaryotic signaling proteins. This cluster consists of different types of chemoreceptors, a large group of proteins that help bacteria and archaea to sense environmental and intracellular cues and relay them to intracellular signaling pathways. This cluster consists mostly of two subclusters, one containing mostly classical membrane-bound methyl-accepting chemotaxis proteins (MCPs)[3,4] and the other subcluster containing soluble chemotaxis proteins[5,6]. A typical MCP has a periplasmic ligand-sensing domain. Ligand binding is transmitted to the cytoplasmic region by a conformational change via the HAMP domain onto a long four-helix bundle of the homodimeric receptor. The tip of the bundle serves as binding platform for the histidine kinase cheA and the receptor coupling protein cheW, which ultimately control the rotation of the flagellar motor. In the center of the bundle, a flexible glycine hinge separates the hairpin tip from a region that can be reversibly methylated at specific glutamate residues by a methyltransferase and a methylesterase. Methylation leads to a subtle conformational change that counteracts the effect of ligand binding. Glutamine can be irreversibly deamidated to glutamate by the methylesterase CheB[7]. The *C*-terminal portion of numerous MCPs was found by our screen to contain a SNARE-like sequence region. Indeed, the methylation region somewhat resembles a SNARE bundle, although the MCP bundle has an antiparallel orientation[8,9]. Intriguingly, the *C*-terminal helices display buried polar glutamine (Q) residues in the center, although the two glutamines do not form a hydrophilic layer with residues from the *N*-terminal helices comparable to the setting in the 0-layer of eukaryotic SNARE complexes.

The most abundant soluble chemotaxis factor in the second subcluster, often referred to as PAS domain S-box protein, was recognized by a specific HMM for a more divergent SNARE type (Qb.I, Sec20) that is involved in ER-Golgi trafficking in eukaryotic cells[10]. In many eukaryotes, Sec20 features a serine residue in the 0-layer position, which is also present in the SNARE-like region of this chemotaxis factor. In the soluble chemotaxis factor, this region may form an extended coiled coil as a dimerization interface[11-13] between the Per-Arnt-Sim (PAS) domains[14]. Next to the PAS domains, methyltransferase (CheR), methylesterase (CheB), histidine kinase domains and cheY-homologous receiver domains were present. These soluble chemotaxis factors are present mostly in β-, γ- and δ- proteobacteria, some firmicutes and also in some archaea[15,16]. Typical domain architectures are given below (Fig. S5.1).

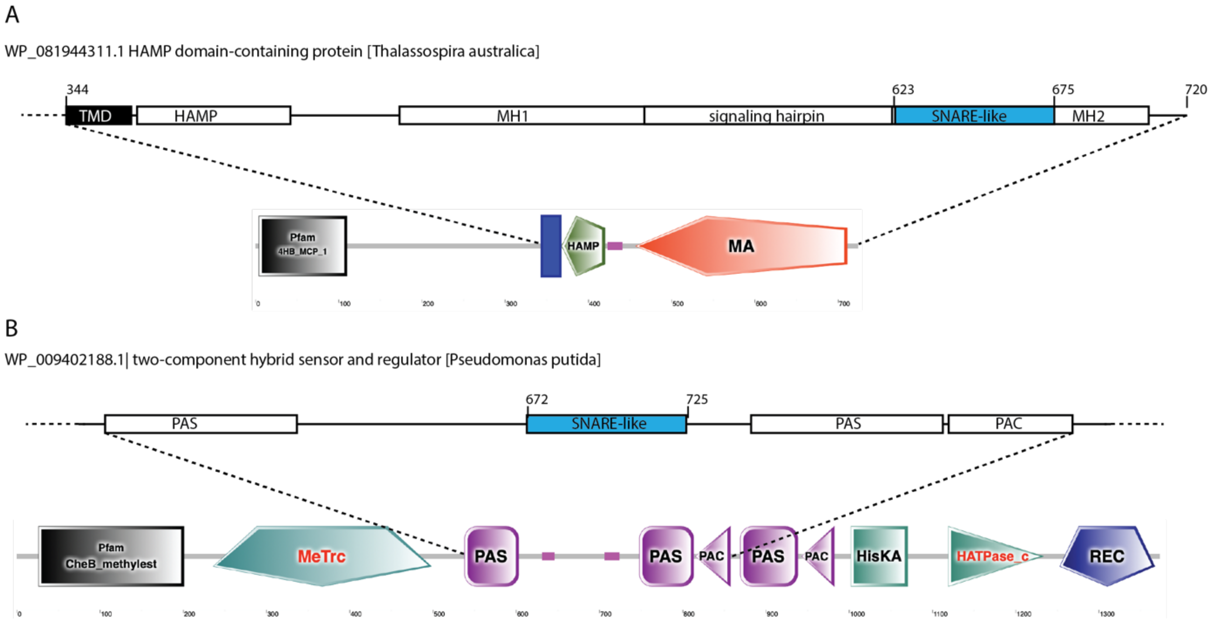

**Fig. S5.1. Domain organization of typical prokaryotic signaling molecules with SNARE-like regions.**
The domain arrangement of a membrane-bound methyl-accepting chemotaxis protein from *Thalassospira australica* (WP_081944311.1) (A) and a soluble chemotaxis protein from *Pseudomonas putida CSV86* (WP_009402188.1) (B) are shown. For both, the SMART annotation is shown below[17]. A more detailed scheme domain arrangement including the SNARE-like region is shown on top.

#### 5.2 The SNARE-like region of an MCP from the $\alpha$-proteobacterium *Thalassospira australica* does not interact with neuronal SNARE proteins

MCPs with SNARE-like regions were found in different lineages of bacteria as well as in archaea. Sequences from $\alpha$-proteobacteria were among the best-scoring MCPs, which was intriguing, because this lineage of bacteria includes the closest bacterial relatives of mitochondria. To test whether the SNARE-like region of an MCP can interact with eukaryotic SNARE proteins, we expressed this region of an MCP from the $\alpha$-proteobacterium *Thalassospira australica* as recombinant protein, as it was among the best-scoring sequences. For the biochemical test, we again used the two helices of neuronal SNAP-25 as independent constructs[18]. We then mixed all the different combinations of neuronal SNAREs with the MCP from *T. australica* but no stable complex was formed (Fig. S5.2). We are aware that this negative finding does not entirely rule out that MCPs could act as a SNARE-like protein. However, given the complex domain architecture of MCPs and the lack of the *C*-terminal membrane anchor that is present in most SNARE proteins, an evolutionary scenario in which a small portion of an MCP was co-opted for a new function does not seem likely.

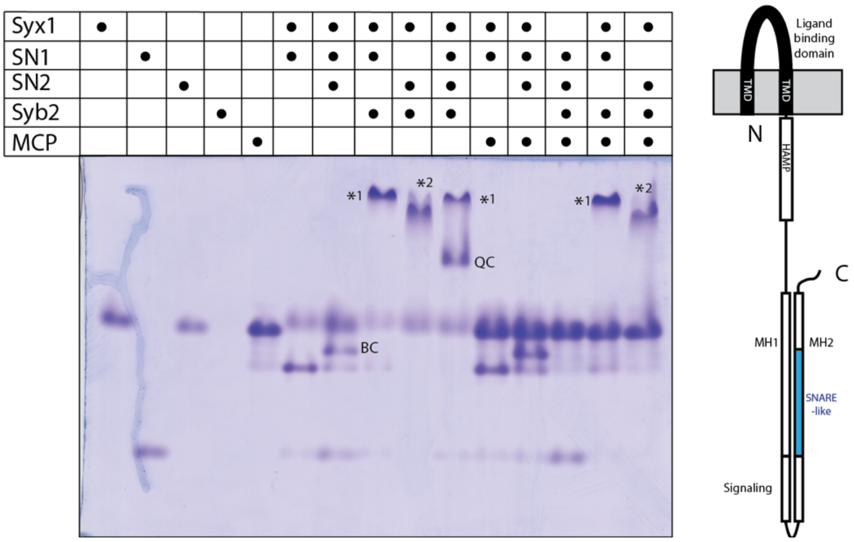

**Fig. S5.2. The SNARE-like region of an MCP from the $\alpha$-proteobacterium *Thalassospira australica* does not form a stable complex with neuronal SNARE proteins**.

On the left, a non-denaturing gel is shown, in which the SNARE-like region of the MCP from *T. australica* (aa 616–685) is mixed with neuronal SNARE proteins. All incubations were performed as described in the legend of Figure 1. The neuronal SNAREs assemble into a Syx1–SN1 complex, a complex consisting of Syx–SN1–SN2 (BC), a Syx1– SN1–Syb2 complex (*1), a Syx1–SN2–Syb2 complex (*2), and a quaternary complex (QC)[18], but no additional complex band appears in the presence of MCP. On the right, the domain architecture of a typical MCP is shown. The sensory (ligand-binding) domain is extracellular and is anchored in the membrane by two transmembrane regions. The HAMP domain and and the methyl-accepting domain comprising two methylation regions (MH1 and MH2) and the signaling subdomain are in the cytoplasm. The putative SNARE-like region is shown in blue.

##### 5.3 Other prokaryotic factors in sizable clusters

Overall, our screen for SNARE-like sequences only detected a subset of the large repertoire of different prokaryotic signal proteins, namely the ones with extended coiled-coil segments that show some similarity to the specific coiled-coil pattern found in SNARE proteins. Comparable subsets of proteins with extended coiled-coil segments, sometimes from a distinct taxonomic subgroup of bacteria, were also present in other larger clusters of our sequence collection. Among these are factors with known functions and structures such as MCE, flagellin, murein lipoprotein, and HlyD/EmrA. Similar to chemotaxis proteins, most of these factors found by our screen for SNARE-like regions are also unlikely to constitute a SNARE ancestor (see Fig. S5.3 for a domain overview).

- Cluster 23 contains a group of actinobacterial proteins with a mammalian cell entry (MCE) domain[19-21]. These proteins form a hexameric membrane channel that transports lipids and steroids. The MCE proteins found by our screen also possess a coiled-coil stretch that bears some similarity to a SNARE domain. We noted, however, that other homologous MCEs found by BLAST searches had a coiled-coil stretch as well but were less well detected by our HMMs.
- Cluster 20 contains closely related proteins of spirochetes with unknown function. It comprises a signal sequence and therefore is likely to be secreted by the bacteria.
- Cluster 11 comprises flagellin, the principal component of the bacterial flagellum[22,23]. Different HMMs for SNARE motifs recognized the longer *C*-terminal helix region of flagellins from different bacterial lineages.
- Cluster 52 consists of murein lipoproteins, which is one of the most abundant membrane proteins of in some Gram-negative bacteria. It is a small protein that is covalently attached to the peptidoglycan layer via a *C*-terminal lysine residue and forms homotrimeric coiled-coil structures[24]. Our HMMs detected murein lipoproteins mainly from bacteria of the order Methylococcales.
- Cluster 36 comprises the periplasmic adaptor protein EmrA of bacterial tripartite efflux pumps. The protein forms an extended alpha-helical coiled coil that is attached to the inner membrane by a lipoyl domain, which may form a channel that connects the inner and outer membranes[25,26].

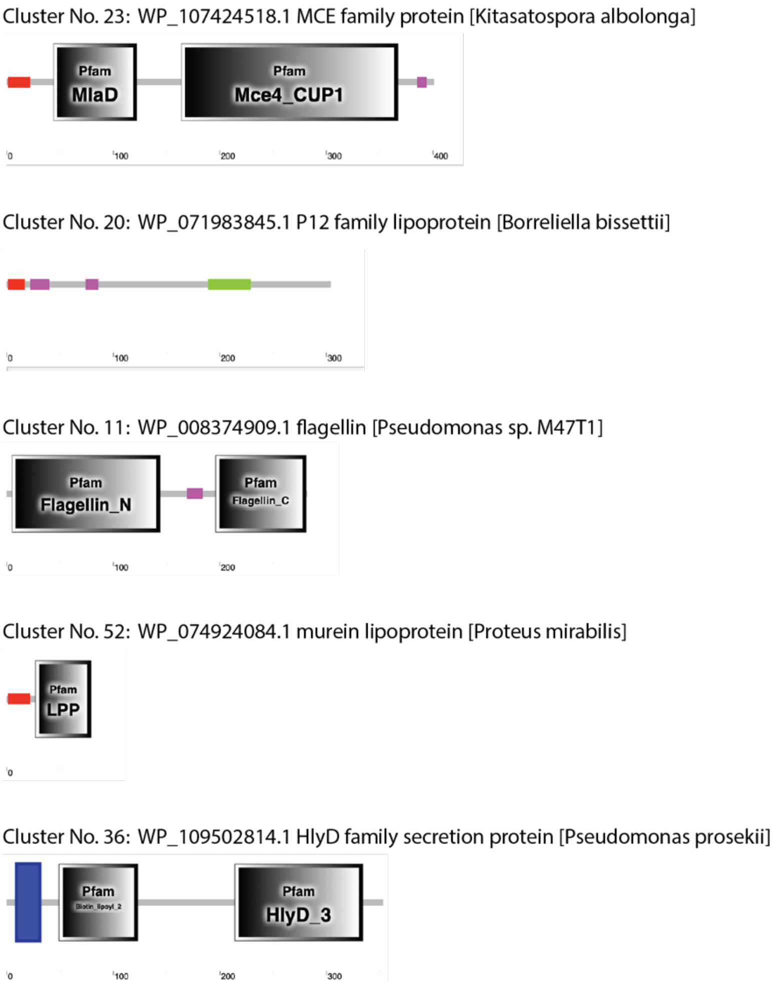

**Fig. S5.3. Domain organization of factors found in other sizable clusters.**
For the factors listed, the SMART annotation[17] is shown below. Note that three factors possess signal peptides (red) at the amino terminus that targets the proteins into, or across, membranes [27].

#### References – Section 5

1 Mistry, J., Finn, R. D., Eddy, S. R., Bateman, A. & Punta, M. Challenges in homology search: HMMER3 and convergent evolution of coiled-coil regions. *Nucleic Acids Res* **41**, e121, doi:10.1093/nar/gkt263 (2013).

2 Surkont, J. & Pereira-Leal, J. B. Evolutionary patterns in coiled-coils. *Genome Biol Evol* **7**, 545-556, doi:10.1093/gbe/evv007 (2015).

3 Parkinson, J. S. Signaling mechanisms of HAMP domains in chemoreceptors and sensor kinases. *Annu Rev Microbiol* **64**, 101-122, doi:10.1146/annurev.micro.112408.134215 (2010).

4 Salah Ud-Din, A. I. M. & Roujeinikova, A. Methyl-accepting chemotaxis proteins: a core sensing element in prokaryotes and archaea. *Cell Mol Life Sci* **74**, 3293-3303, doi:10.1007/s00018-017-2514-0 (2017).

5 Alexander, R. P. & Zhulin, I. B. Evolutionary genomics reveals conserved structural determinants of signaling and adaptation in microbial chemoreceptors. *Proceedings*

*of the National Academy of Sciences of the United States of America* **104**, 2885-2890, doi:10.1073/pnas.0609359104 (2007).
6 Zhulin, I. B. The superfamily of chemotaxis transducers: from physiology to genomics and back. *Adv Microb Physiol* **45**, 157-198 (2001).
7 Parkinson, J. S., Hazelbauer, G. L. & Falke, J. J. Signaling and sensory adaptation in Escherichia coli chemoreceptors: 2015 update. *Trends Microbiol* **23**, 257-266, doi:10.1016/j.tim.2015.03.003 (2015).
8 Kim, K. K., Yokota, H. & Kim, S. H. Four-helical-bundle structure of the cytoplasmic domain of a serine chemotaxis receptor. *Nature* **400**, 787-792, doi:10.1038/23512 (1999).
9 Park, S. Y. *et al.* Reconstruction of the chemotaxis receptor-kinase assembly. *Nature structural & molecular biology* **13**, 400-407, doi:10.1038/nsmb1085 (2006).
10 Kloepper, T. H., Kienle, C. N. & Fasshauer, D. An elaborate classification of SNARE proteins sheds light on the conservation of the eukaryotic endomembrane system. *Molecular biology of the cell* **18**, 3463-3471, doi:10.1091/mbc.E07-03-0193 (2007).
11 Anantharaman, V., Balaji, S. & Aravind, L. The signaling helix: a common functional theme in diverse signaling proteins. *Biol Direct* **1**, 25, doi:10.1186/1745-6150-1-25 (2006).
12 Singh, M., Berger, B., Kim, P. S., Berger, J. M. & Cochran, A. G. Computational learning reveals coiled coil-like motifs in histidine kinase linker domains. *Proceedings of the National Academy of Sciences of the United States of America* **95**, 2738-2743, doi:10.1073/pnas.95.6.2738 (1998).
13 Collins, K. D., Lacal, J. & Ottemann, K. M. Internal sense of direction: sensing and signaling from cytoplasmic chemoreceptors. *Microbiol Mol Biol Rev* **78**, 672-684, doi:10.1128/MMBR.00033-14 (2014).
14 Henry, J. T. & Crosson, S. Ligand-binding PAS domains in a genomic, cellular, and structural context. *Annu Rev Microbiol* **65**, 261-286, doi:10.1146/annurev-micro-121809-151631 (2011).
15 Taylor, B. L. & Zhulin, I. B. PAS domains: internal sensors of oxygen, redox potential, and light. *Microbiol Mol Biol Rev* **63**, 479-506 (1999).
16 Zhulin, I. B., Taylor, B. L. & Dixon, R. PAS domain S-boxes in Archaea, Bacteria and sensors for oxygen and redox. *Trends Biochem Sci* **22**, 331-333 (1997).
17 Letunic, I. & Bork, P. 20 years of the SMART protein domain annotation resource. *Nucleic Acids Res* **46**, D493-D496, doi:10.1093/nar/gkx922 (2018).
18 Fasshauer, D., Eliason, W., Brunger, A. & Jahn, R. Identification of a minimal core of the synaptic SNARE complex sufficient for reversible assembly and disassembly. *Biochemistry* **37**, 10354-10362, doi:10.1021/bi980542h (1998).
19 Ekiert, D. C. *et al.* Architectures of Lipid Transport Systems for the Bacterial Outer Membrane. *Cell* **169**, 273-285 e217, doi:10.1016/j.cell.2017.03.019 (2017).
20 Mohn, W. W. *et al.* The actinobacterial mce4 locus encodes a steroid transporter. *The Journal of biological chemistry* **283**, 35368-35374, doi:10.1074/jbc.M805496200 (2008).
21 Pandey, A. K. & Sassetti, C. M. Mycobacterial persistence requires the utilization of host cholesterol. *Proceedings of the National Academy of Sciences of the United States of America* **105**, 4376-4380, doi:10.1073/pnas.0711159105 (2008).
22 Wang, F. *et al.* A structural model of flagellar filament switching across multiple bacterial species. *Nat Commun* **8**, 960, doi:10.1038/s41467-017-01075-5 (2017).

23 Yonekura, K., Maki-Yonekura, S. & Namba, K. Complete atomic model of the bacterial flagellar filament by electron cryomicroscopy. *Nature* **424**, 643-650, doi:10.1038/nature01830 (2003).
24 Shu, W., Liu, J., Ji, H. & Lu, M. Core structure of the outer membrane lipoprotein from Escherichia coli at 1.9 A resolution. *Journal of molecular biology* **299**, 1101-1112, doi:10.1006/jmbi.2000.3776 (2000).
25 Hinchliffe, P. *et al.* Structure of the periplasmic adaptor protein from a major facilitator superfamily (MFS) multidrug efflux pump. *FEBS letters* **588**, 3147-3153, doi:10.1016/j.febslet.2014.06.055 (2014).
26 Kim, J. S. *et al.* Crystal Structure of a Soluble Fragment of the Membrane Fusion Protein HlyD in a Type I Secretion System of Gram-Negative Bacteria. *Structure* **24**, 477-485, doi:10.1016/j.str.2015.12.012 (2016).
27 Almagro Armenteros, J. J. *et al.* SignalP 5.0 improves signal peptide predictions using deep neural networks. *Nat Biotechnol* **37**, 420-423, doi:10.1038/s41587-019-0036-z (2019).

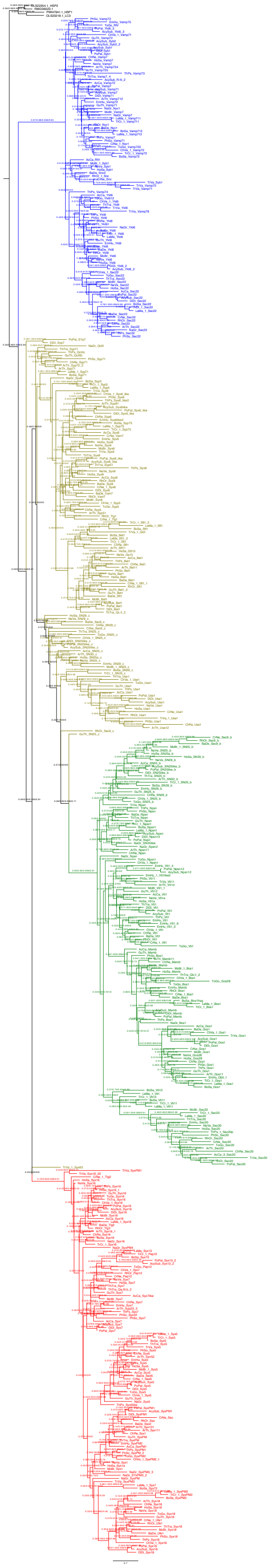

#### Section 6

**Fig. S6.1: Maximum likelihood tree from typical SNAREs of 24 representative eukaryotic species.**

Statistical branch support values (likelihood-mapping, IQ-TREE support, RAxML support, PhyML support) are given. Qa-SNAREs are in red, Qb-SNAREs in khaki, Qc-SNAREs in moss green, and R-SNAREs in blue. A version with collapsed clades is shown in Fig. 2.

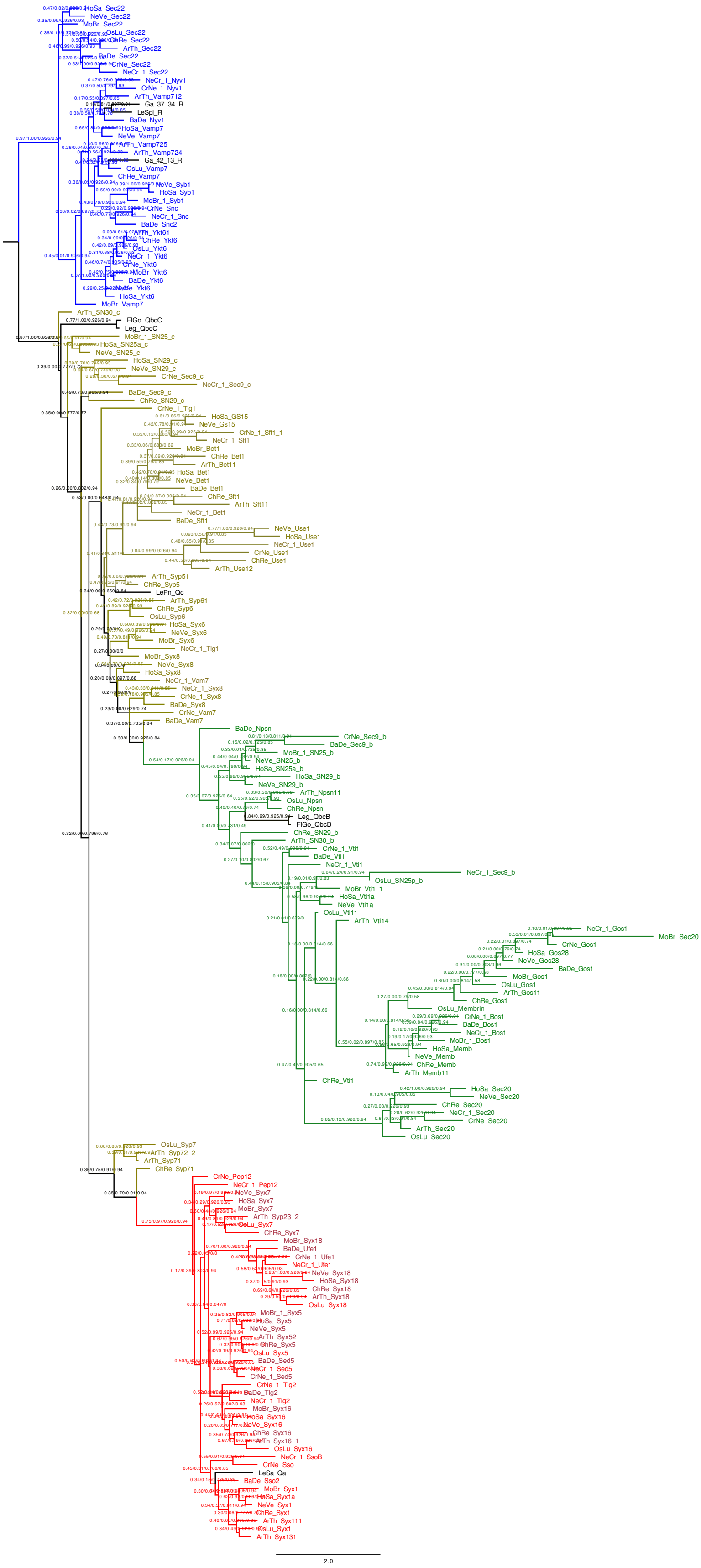

**Fig. S6.2. Maximum-likelihood tree from six SNARE proteins from different γ-proteobacteria of the order Legionellales and typical SNARE sets from nine representative eukaryotic species.** Statistical branch support values (likelihood-mapping, IQ-TREE support, RAxML support, PhyML support) are given. A version with collapsed clades is shown in Extended Data Fig. 5.

A

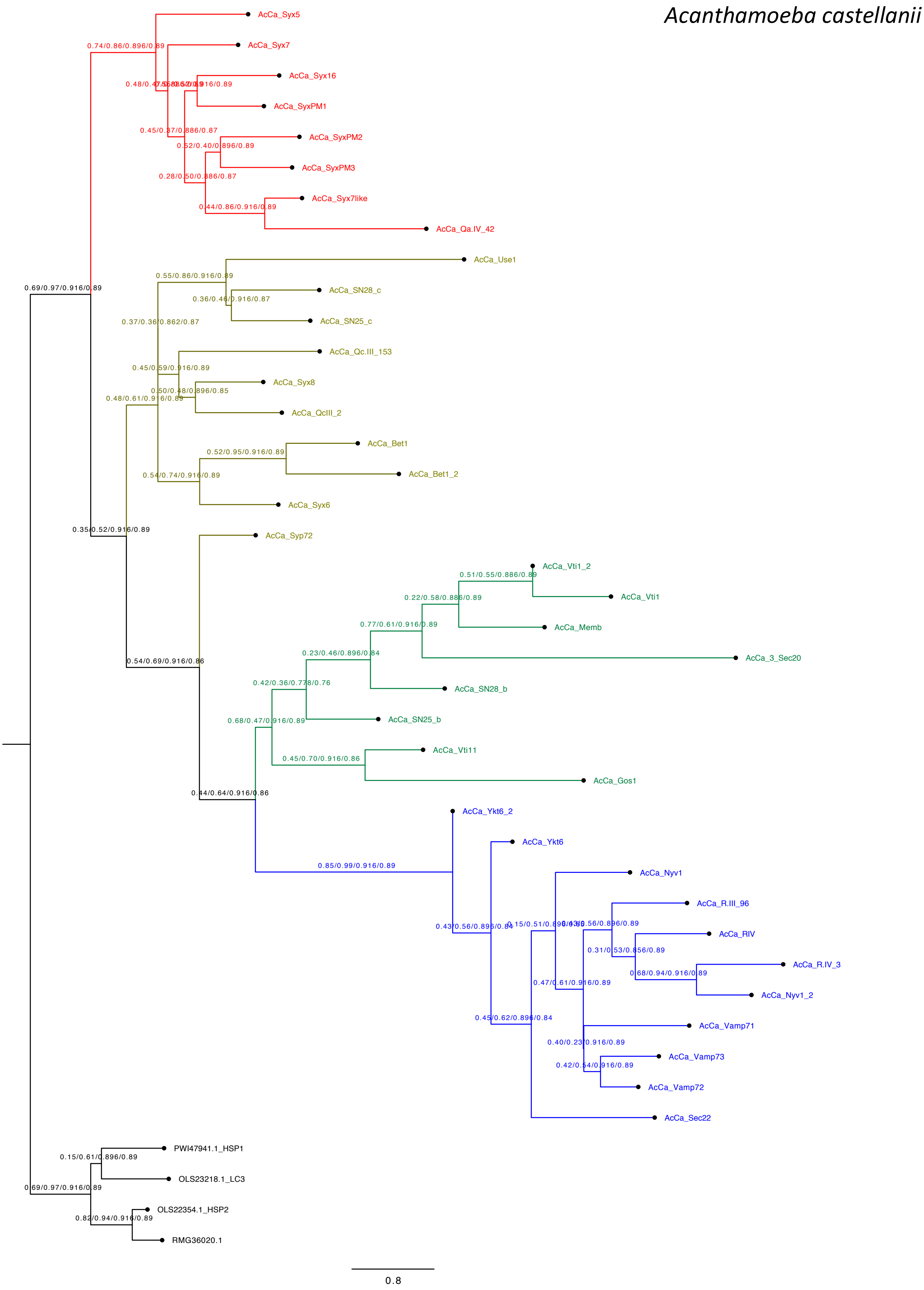
Acanthamoeba castellanii
AcCa_Syx5
0.74/0.86/0.896/0.89
AcCa_Syx7
AcCa_Syx16
AcCa_SyxPM1
0.45/0.37/0.886/0.87
AcCa_SyxPM2
0.52/0.40/0.896/0.89
AcCa_SyxPM3
0.28/0.50/0.886/0.87
AcCa_Syx7like
0.44/0.86/0.916/0.89
AcCa_Qa.IV_42
AcCa_Use1
0.55/0.86/0.916/0.89
AcCa_SN28_c
0.36/0.46/0.916/0.87
AcCa_SN25_c
0.37/0.36/0.862/0.87
AcCa_Qc.III_153
0.45/0.59/0.916/0.89
AcCa_Syx8
0.50/0.48/0.896/0.85
0.48/0.61/0.916/0.89
AcCa_QcIII_2
AcCa_Bet1
0.52/0.95/0.916/0.89
AcCa_Bet1_2
0.54/0.74/0.916/0.89
AcCa_Syx6
0.69/0.97/0.916/0.89
0.35/0.52/0.916/0.89
AcCa_Syp72
AcCa_Vti1_2
0.51/0.55/0.886/0.89
AcCa_Vti1
0.22/0.58/0.886/0.89
AcCa_Memb
0.77/0.61/0.916/0.89
AcCa_3_Sec20
0.23/0.46/0.896/0.84
0.54/0.69/0.916/0.86
AcCa_SN28_b
0.42/0.36/0.778/0.76
AcCa_SN25_b
0.68/0.47/0.916/0.89
AcCa_Vti11
0.45/0.70/0.916/0.86
AcCa_Gos1
0.44/0.64/0.916/0.86
AcCa_Ykt6_2
AcCa_Ykt6
0.85/0.99/0.916/0.89
AcCa_Nyv1
AcCa_R.III_96
0.43/0.56/0.896/0.84
AcCa_RIV
0.31/0.53/0.856/0.89
AcCa_R.IV_3
0.68/0.94/0.916/0.89
0.47/0.61/0.916/0.89
AcCa_Nyv1_2
0.45/0.62/0.896/0.84
AcCa_Vamp71
0.40/0.23/0.916/0.89
AcCa_Vamp73
0.42/0.54/0.916/0.89
AcCa_Vamp72
AcCa_Sec22
PWI47941.1_HSP1
0.15/0.61/0.896/0.89
OLS23218.1_LC3
0.69/0.97/0.916/0.89
OLS22354.1_HSP2
0.82/0.94/0.916/0.89
RMG36020.1
0.8

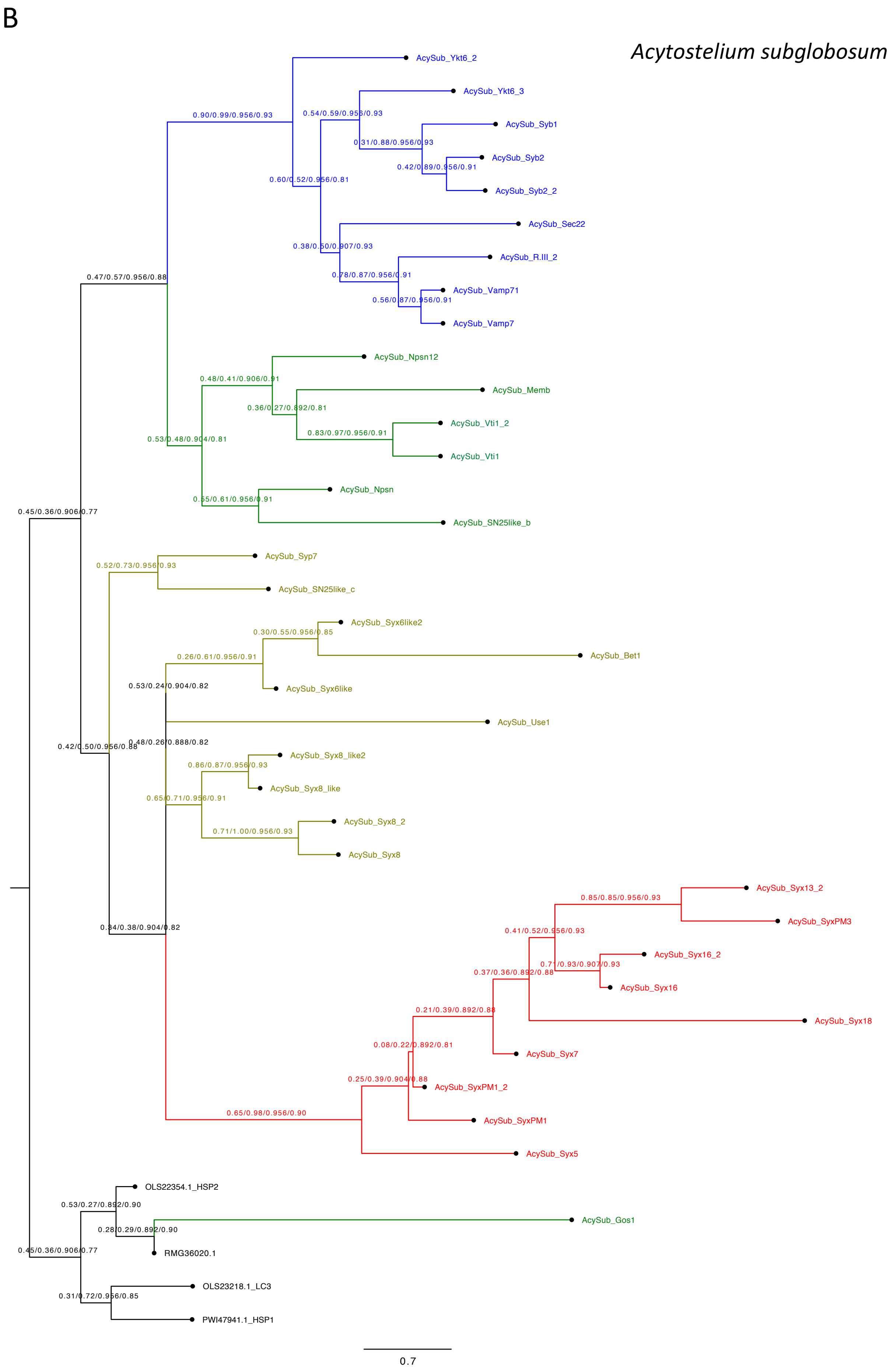

B
Acytostelium subglobosum
AcySub_Ykt6_2
0.90/0.99/0.956/0.93
AcySub_Ykt6_3
0.54/0.59/0.956/0.93
AcySub_Syb1
0.31/0.88/0.956/0.93
AcySub_Syb2
0.42/0.89/0.956/0.91
0.60/0.52/0.956/0.81
AcySub_Syb2_2
AcySub_Sec22
0.38/0.50/0.907/0.93
AcySub_R.III_2
0.78/0.87/0.956/0.91
0.47/0.57/0.956/0.88
AcySub_Vamp71
0.56/0.87/0.956/0.91
AcySub_Vamp7
AcySub_Npsn12
0.48/0.41/0.906/0.91
AcySub_Memb
0.36/0.27/0.892/0.81
AcySub_Vti1_2
0.83/0.97/0.956/0.91
0.53/0.48/0.904/0.81
AcySub_Vti1
AcySub_Npsn
0.55/0.61/0.956/0.91
0.45/0.36/0.906/0.77
AcySub_SN25like_b
AcySub_Syp7
0.52/0.73/0.956/0.93
AcySub_SN25like_c
AcySub_Syx6like2
0.30/0.55/0.956/0.85
0.26/0.61/0.956/0.91
AcySub_Bet1
0.53/0.24/0.904/0.82
AcySub_Syx6like
AcySub_Use1
0.48/0.26/0.888/0.82
0.42/0.50/0.956/0.88
AcySub_Syx8_like2
0.86/0.87/0.956/0.93
AcySub_Syx8_like
0.65/0.71/0.956/0.91
AcySub_Syx8_2
0.71/1.00/0.956/0.93
AcySub_Syx8
AcySub_Syx13_2
0.85/0.85/0.956/0.93
AcySub_SyxPM3
0.41/0.52/0.956/0.93
0.34/0.38/0.904/0.82
AcySub_Syx16_2
0.71/0.93/0.907/0.93
0.37/0.36/0.892/0.88
AcySub_Syx16
AcySub_Syx18
0.21/0.39/0.892/0.88
AcySub_Syx7
0.08/0.22/0.892/0.81
0.25/0.39/0.904/0.88
AcySub_SyxPM1_2
0.65/0.98/0.956/0.90
AcySub_SyxPM1
AcySub_Syx5
OLS22354.1_HSP2
0.53/0.27/0.892/0.90
AcySub_Gos1
0.28/0.29/0.892/0.90
0.45/0.36/0.906/0.77
RMG36020.1
OLS23218.1_LC3
0.31/0.72/0.956/0.85
PWI47941.1_HSP1
0.7

C

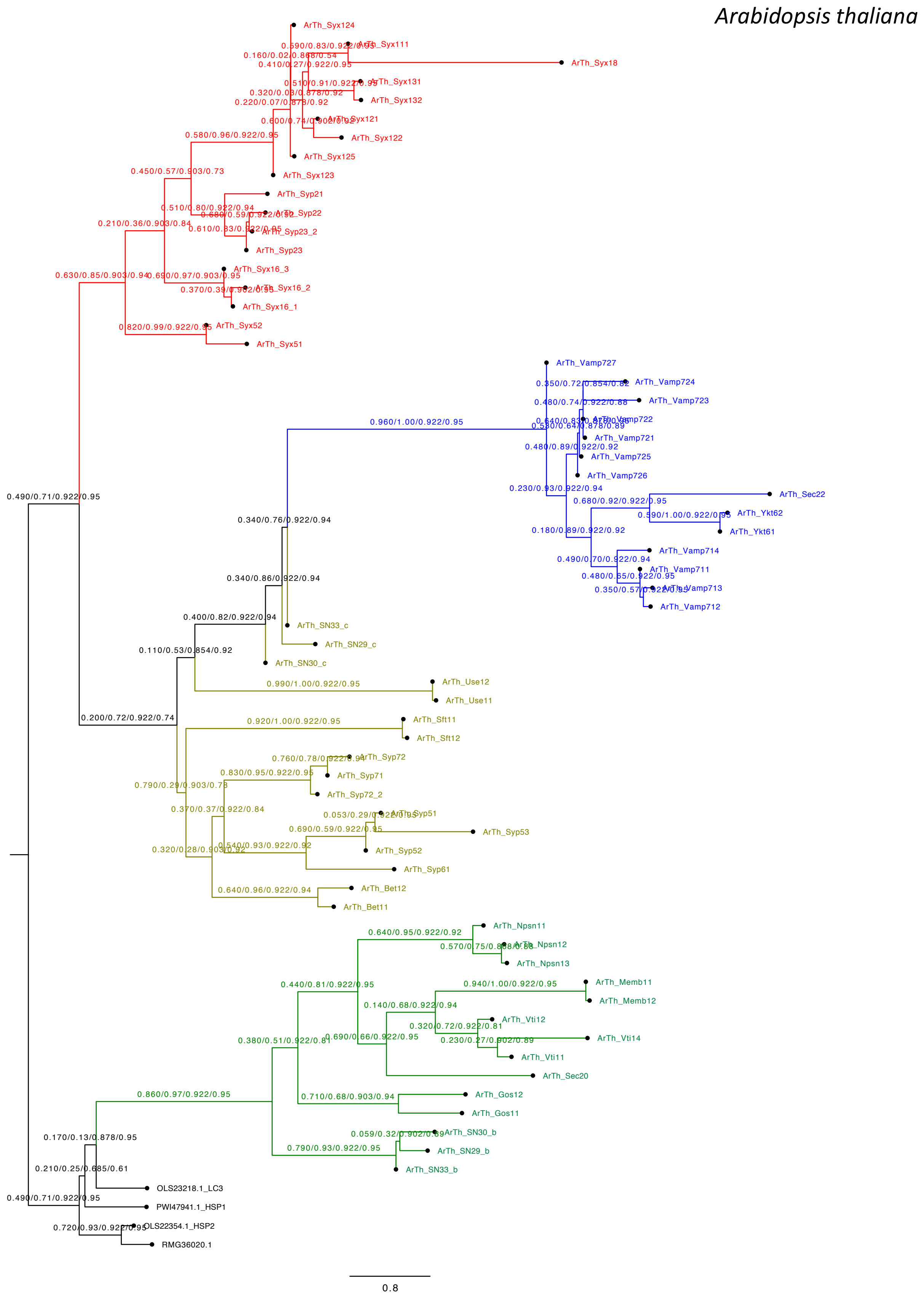
Arabidopsis thaliana
ArTh_Syx124
ArTh_Syx111
ArTh_Syx18
ArTh_Syx131
ArTh_Syx132
ArTh_Syx121
ArTh_Syx122
ArTh_Syx125
ArTh_Syx123
ArTh_Syp21
ArTh_Syp22
ArTh_Syp23_2
ArTh_Syp23
ArTh_Syx16_3
ArTh_Syx16_2
ArTh_Syx16_1
ArTh_Syx52
ArTh_Syx51
ArTh_Vamp727
ArTh_Vamp724
ArTh_Vamp723
ArTh_Vamp722
ArTh_Vamp721
ArTh_Vamp725
ArTh_Vamp726
ArTh_Sec22
ArTh_Ykt62
ArTh_Ykt61
ArTh_Vamp714
ArTh_Vamp711
ArTh_Vamp713
ArTh_Vamp712
ArTh_SN33_c
ArTh_SN29_c
ArTh_SN30_c
ArTh_Use12
ArTh_Use11
ArTh_Sft11
ArTh_Sft12
ArTh_Syp72
ArTh_Syp71
ArTh_Syp72_2
ArTh_Syp51
ArTh_Syp53
ArTh_Syp52
ArTh_Syp61
ArTh_Bet12
ArTh_Bet11
ArTh_Npsn11
ArTh_Npsn12
ArTh_Npsn13
ArTh_Memb11
ArTh_Memb12
ArTh_Vti12
ArTh_Vti14
ArTh_Vti11
ArTh_Sec20
ArTh_Gos12
ArTh_Gos11
ArTh_SN30_b
ArTh_SN29_b
ArTh_SN33_b
OLS23218.1_LC3
PWI47941.1_HSP1
OLS22354.1_HSP2
RMG36020.1
0.490/0.71/0.922/0.95
0.200/0.72/0.922/0.74
0.630/0.85/0.903/0.94
0.960/1.00/0.922/0.95
0.340/0.76/0.922/0.94
0.340/0.86/0.922/0.94
0.400/0.82/0.922/0.94
0.110/0.53/0.854/0.92
0.990/1.00/0.922/0.95
0.920/1.00/0.922/0.95
0.860/0.97/0.922/0.95
0.170/0.13/0.878/0.95
0.210/0.25/0.685/0.61
0.720/0.93/0.922/0.95
0.8

D

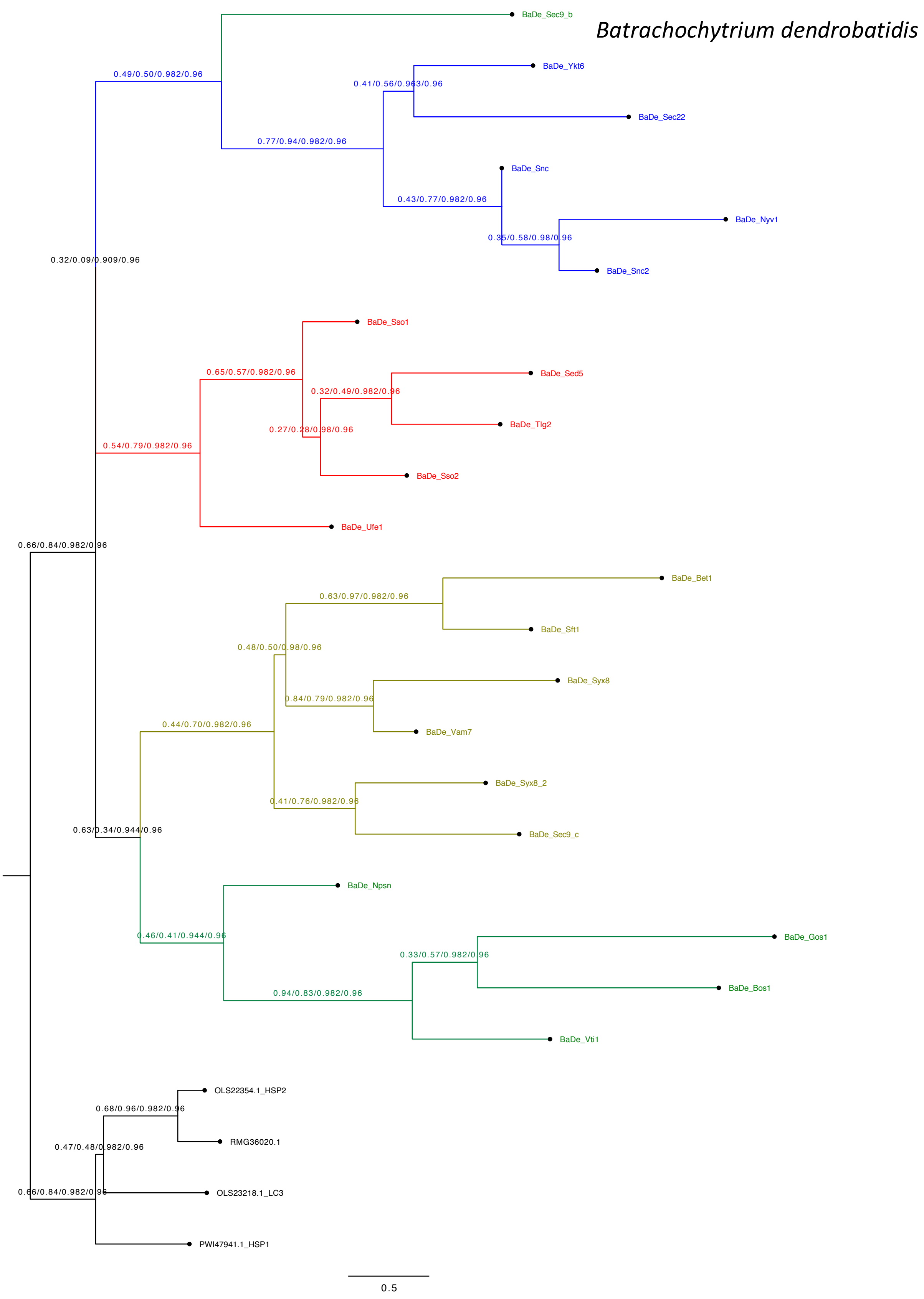
Batrachochytrium dendrobatidis
BaDe_Sec9_b
0.49/0.50/0.982/0.96
BaDe_Ykt6
0.41/0.56/0.963/0.96
BaDe_Sec22
0.77/0.94/0.982/0.96
BaDe_Snc
0.43/0.77/0.982/0.96
BaDe_Nyv1
0.35/0.58/0.98/0.96
BaDe_Snc2
0.32/0.09/0.909/0.96
BaDe_Sso1
0.65/0.57/0.982/0.96
BaDe_Sed5
0.32/0.49/0.982/0.96
0.27/0.28/0.98/0.96
BaDe_Tlg2
0.54/0.79/0.982/0.96
BaDe_Sso2
BaDe_Ufe1
0.66/0.84/0.982/0.96
BaDe_Bet1
0.63/0.97/0.982/0.96
BaDe_Sft1
0.48/0.50/0.98/0.96
BaDe_Syx8
0.84/0.79/0.982/0.96
0.44/0.70/0.982/0.96
BaDe_Vam7
BaDe_Syx8_2
0.41/0.76/0.982/0.96
0.63/0.34/0.944/0.96
BaDe_Sec9_c
BaDe_Npsn
0.46/0.41/0.944/0.96
BaDe_Gos1
0.33/0.57/0.982/0.96
0.94/0.83/0.982/0.96
BaDe_Bos1
BaDe_Vti1
OLS22354.1_HSP2
0.68/0.96/0.982/0.96
RMG36020.1
0.47/0.48/0.982/0.96
0.66/0.84/0.982/0.96
OLS23218.1_LC3
PWI47941.1_HSP1
0.5

E

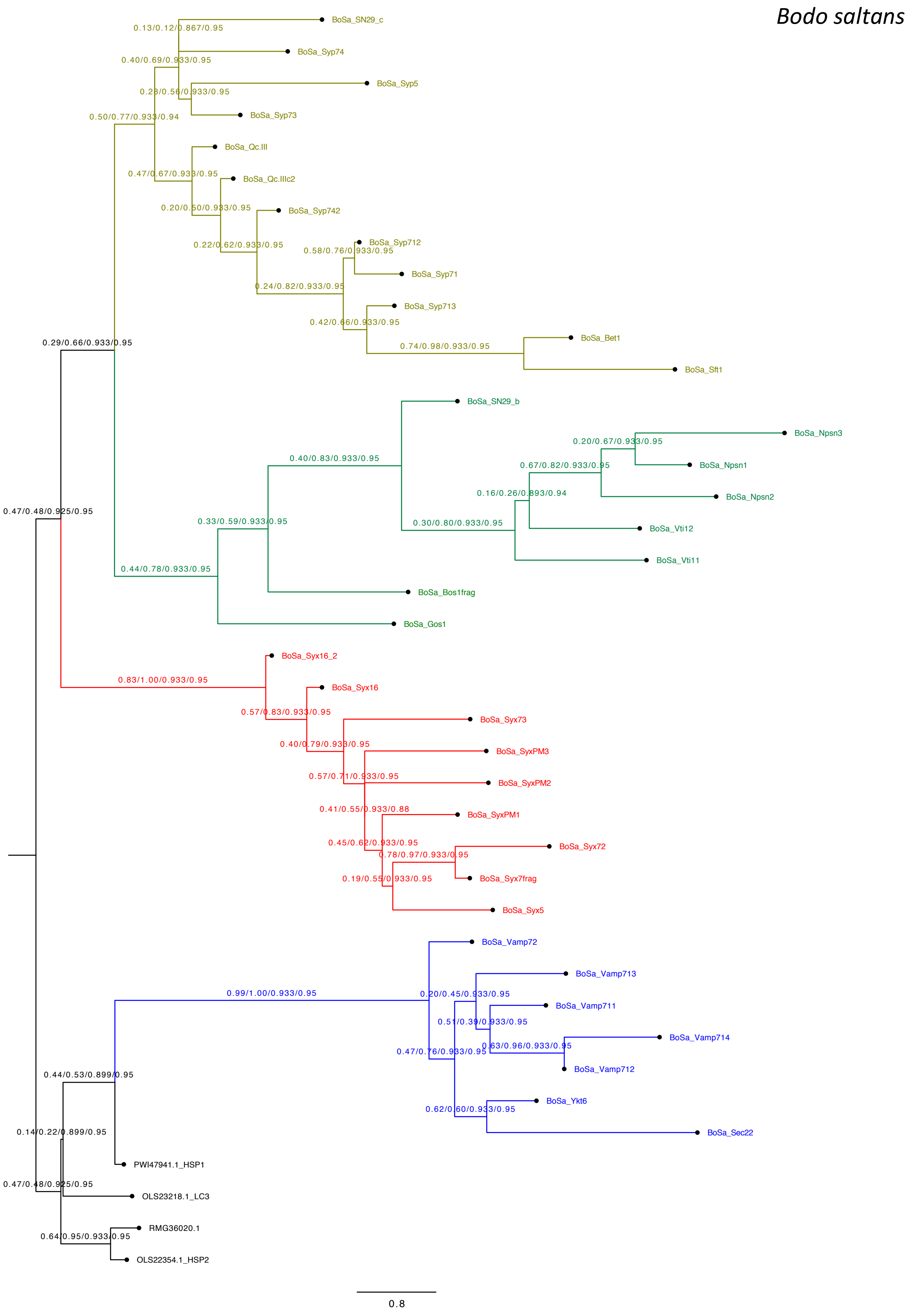
Bodo saltans
BoSa_SN29_c
0.13/0.12/0.867/0.95
BoSa_Syp74
0.40/0.69/0.933/0.95
BoSa_Syp5
0.23/0.56/0.933/0.95
BoSa_Syp73
0.50/0.77/0.933/0.94
BoSa_Qc.III
0.47/0.67/0.933/0.95
BoSa_Qc.IIIc2
0.20/0.50/0.933/0.95
BoSa_Syp742
0.22/0.62/0.933/0.95
BoSa_Syp712
0.58/0.76/0.933/0.95
BoSa_Syp71
0.24/0.82/0.933/0.95
BoSa_Syp713
0.42/0.66/0.933/0.95
BoSa_Bet1
0.74/0.98/0.933/0.95
BoSa_Sft1
0.29/0.66/0.933/0.95
BoSa_SN29_b
0.20/0.67/0.933/0.95
BoSa_Npsn3
0.67/0.82/0.933/0.95
BoSa_Npsn1
0.40/0.83/0.933/0.95
0.16/0.26/0.893/0.94
BoSa_Npsn2
0.47/0.48/0.925/0.95
0.33/0.59/0.933/0.95
0.30/0.80/0.933/0.95
BoSa_Vti12
BoSa_Vti11
0.44/0.78/0.933/0.95
BoSa_Bos1frag
BoSa_Gos1
BoSa_Syx16_2
0.83/1.00/0.933/0.95
BoSa_Syx16
0.57/0.83/0.933/0.95
BoSa_Syx73
0.40/0.79/0.933/0.95
BoSa_SyxPM3
0.57/0.71/0.933/0.95
BoSa_SyxPM2
0.41/0.55/0.933/0.88
BoSa_SyxPM1
0.45/0.62/0.933/0.95
BoSa_Syx72
0.78/0.97/0.933/0.95
0.19/0.55/0.933/0.95
BoSa_Syx7frag
BoSa_Syx5
BoSa_Vamp72
BoSa_Vamp713
0.99/1.00/0.933/0.95
0.20/0.45/0.933/0.95
BoSa_Vamp711
0.51/0.39/0.933/0.95
BoSa_Vamp714
0.47/0.76/0.933/0.95
0.63/0.96/0.933/0.95
BoSa_Vamp712
0.44/0.53/0.899/0.95
BoSa_Ykt6
0.62/0.60/0.933/0.95
BoSa_Sec22
0.14/0.22/0.899/0.95
PWI47941.1_HSP1
0.47/0.48/0.925/0.95
OLS23218.1_LC3
RMG36020.1
0.64/0.95/0.933/0.95
OLS22354.1_HSP2
0.8

F

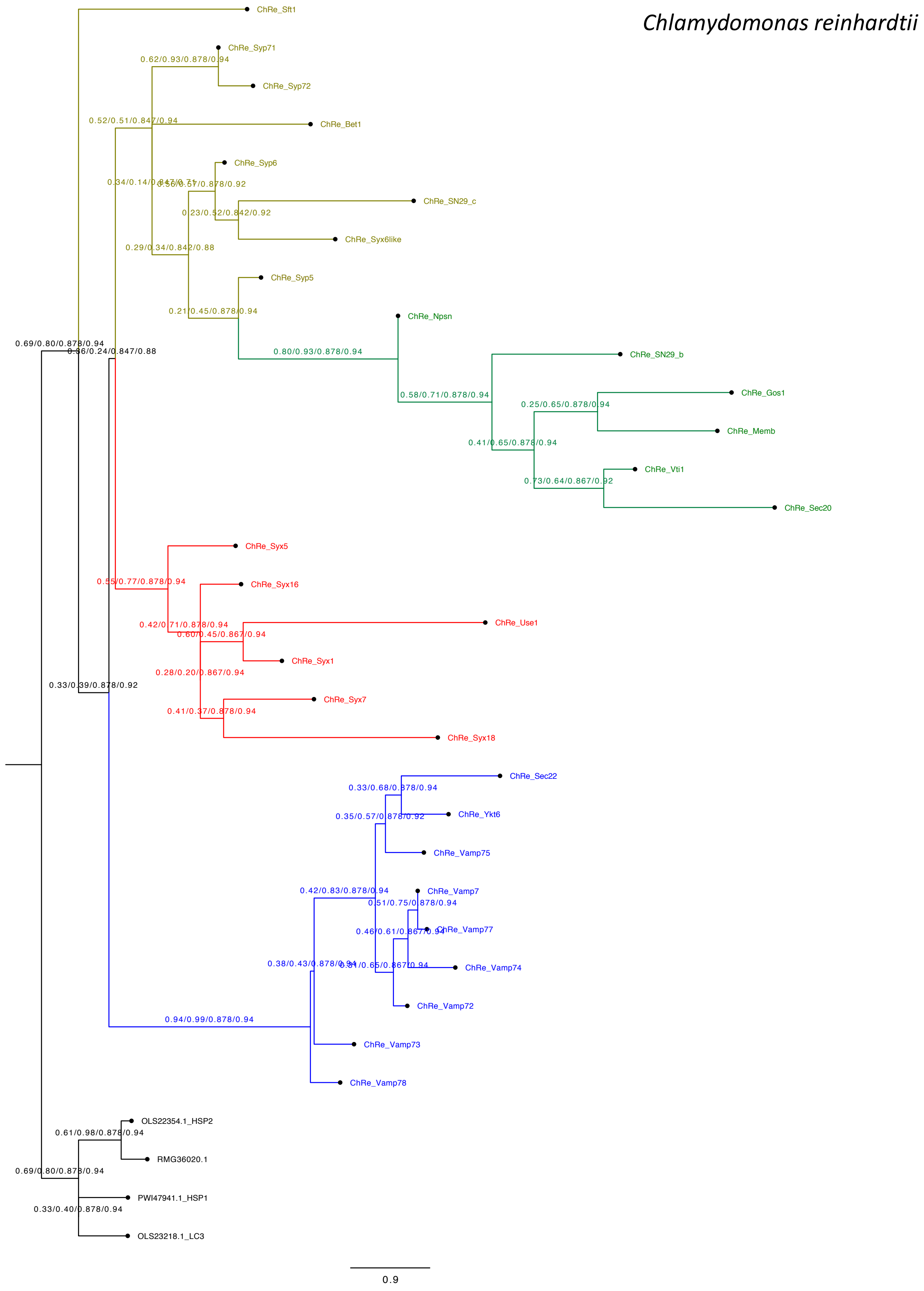

*Chlamydomonas reinhardtii*
ChRe_Sft1
ChRe_Syp71
ChRe_Syp72
ChRe_Bet1
ChRe_Syp6
ChRe_SN29_c
ChRe_Syx6like
ChRe_Syp5
ChRe_Npsn
ChRe_SN29_b
ChRe_Gos1
ChRe_Memb
ChRe_Vti1
ChRe_Sec20
ChRe_Syx5
ChRe_Syx16
ChRe_Use1
ChRe_Syx1
ChRe_Syx7
ChRe_Syx18
ChRe_Sec22
ChRe_Ykt6
ChRe_Vamp75
ChRe_Vamp7
ChRe_Vamp77
ChRe_Vamp74
ChRe_Vamp72
ChRe_Vamp73
ChRe_Vamp78
OLS22354.1_HSP2
RMG36020.1
PWI47941.1_HSP1
OLS23218.1_LC3
0.9

G

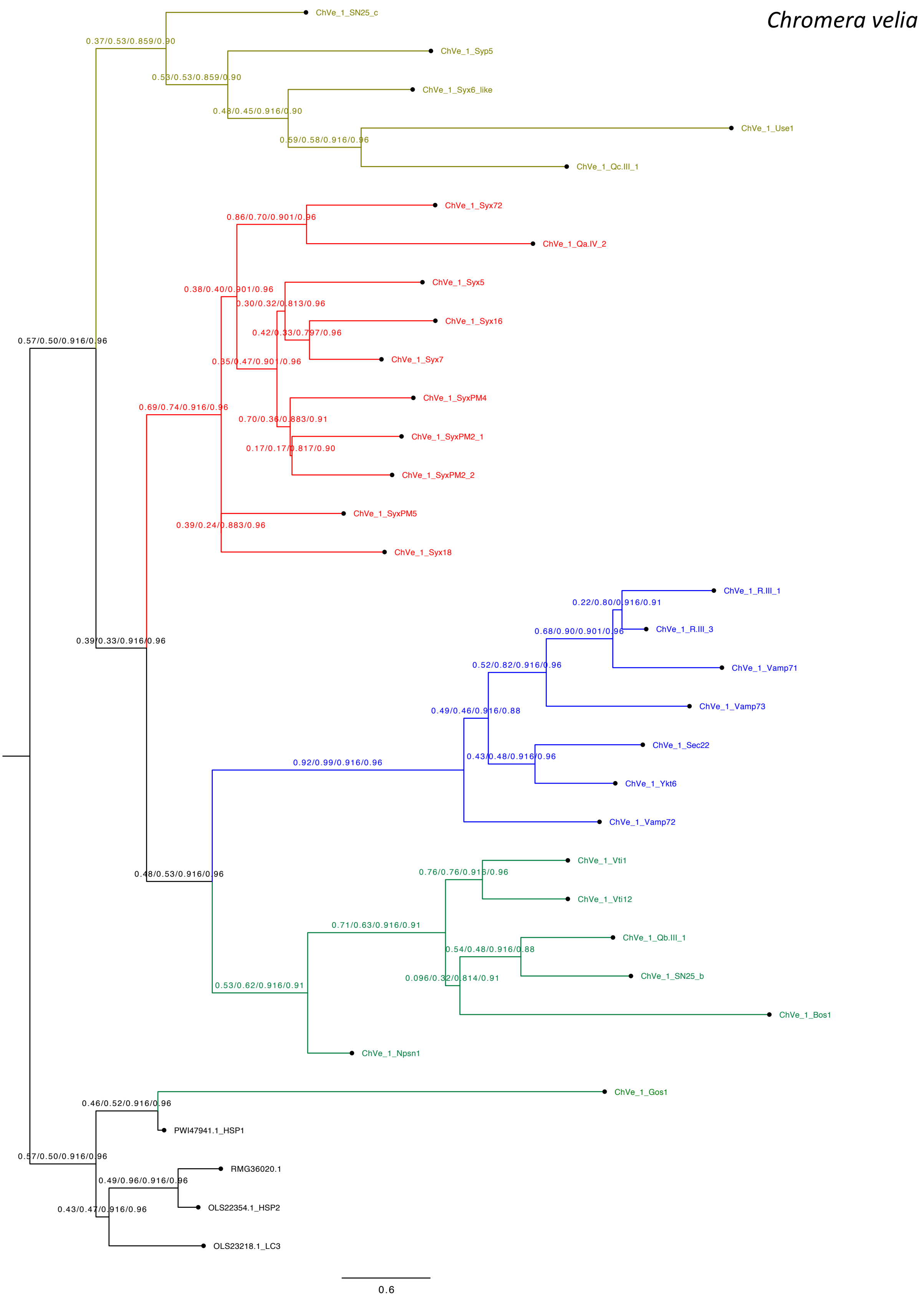
Chromera velia
ChVe_1_SN25_c
0.37/0.53/0.859/0.90
ChVe_1_Syp5
0.53/0.53/0.859/0.90
ChVe_1_Syx6_like
0.48/0.45/0.916/0.90
ChVe_1_Use1
0.59/0.58/0.916/0.96
ChVe_1_Qc.III_1
ChVe_1_Syx72
0.86/0.70/0.901/0.96
ChVe_1_Qa.IV_2
ChVe_1_Syx5
0.38/0.40/0.901/0.96
0.30/0.32/0.813/0.96
ChVe_1_Syx16
0.42/0.33/0.797/0.96
0.57/0.50/0.916/0.96
ChVe_1_Syx7
0.35/0.47/0.901/0.96
ChVe_1_SyxPM4
0.69/0.74/0.916/0.96
0.70/0.36/0.883/0.91
ChVe_1_SyxPM2_1
0.17/0.17/0.817/0.90
ChVe_1_SyxPM2_2
ChVe_1_SyxPM5
0.39/0.24/0.883/0.96
ChVe_1_Syx18
ChVe_1_R.III_1
0.22/0.80/0.916/0.91
ChVe_1_R.III_3
0.68/0.90/0.901/0.96
0.39/0.33/0.916/0.96
0.52/0.82/0.916/0.96
ChVe_1_Vamp71
ChVe_1_Vamp73
0.49/0.46/0.916/0.88
ChVe_1_Sec22
0.43/0.48/0.916/0.96
0.92/0.99/0.916/0.96
ChVe_1_Ykt6
ChVe_1_Vamp72
ChVe_1_Vti1
0.76/0.76/0.916/0.96
0.48/0.53/0.916/0.96
ChVe_1_Vti12
0.71/0.63/0.916/0.91
ChVe_1_Qb.III_1
0.54/0.48/0.916/0.88
0.096/0.32/0.814/0.91
ChVe_1_SN25_b
0.53/0.62/0.916/0.91
ChVe_1_Bos1
ChVe_1_Npsn1
ChVe_1_Gos1
0.46/0.52/0.916/0.96
PWI47941.1_HSP1
0.57/0.50/0.916/0.96
RMG36020.1
0.49/0.96/0.916/0.96
OLS22354.1_HSP2
0.43/0.47/0.916/0.96
OLS23218.1_LC3
0.6

H

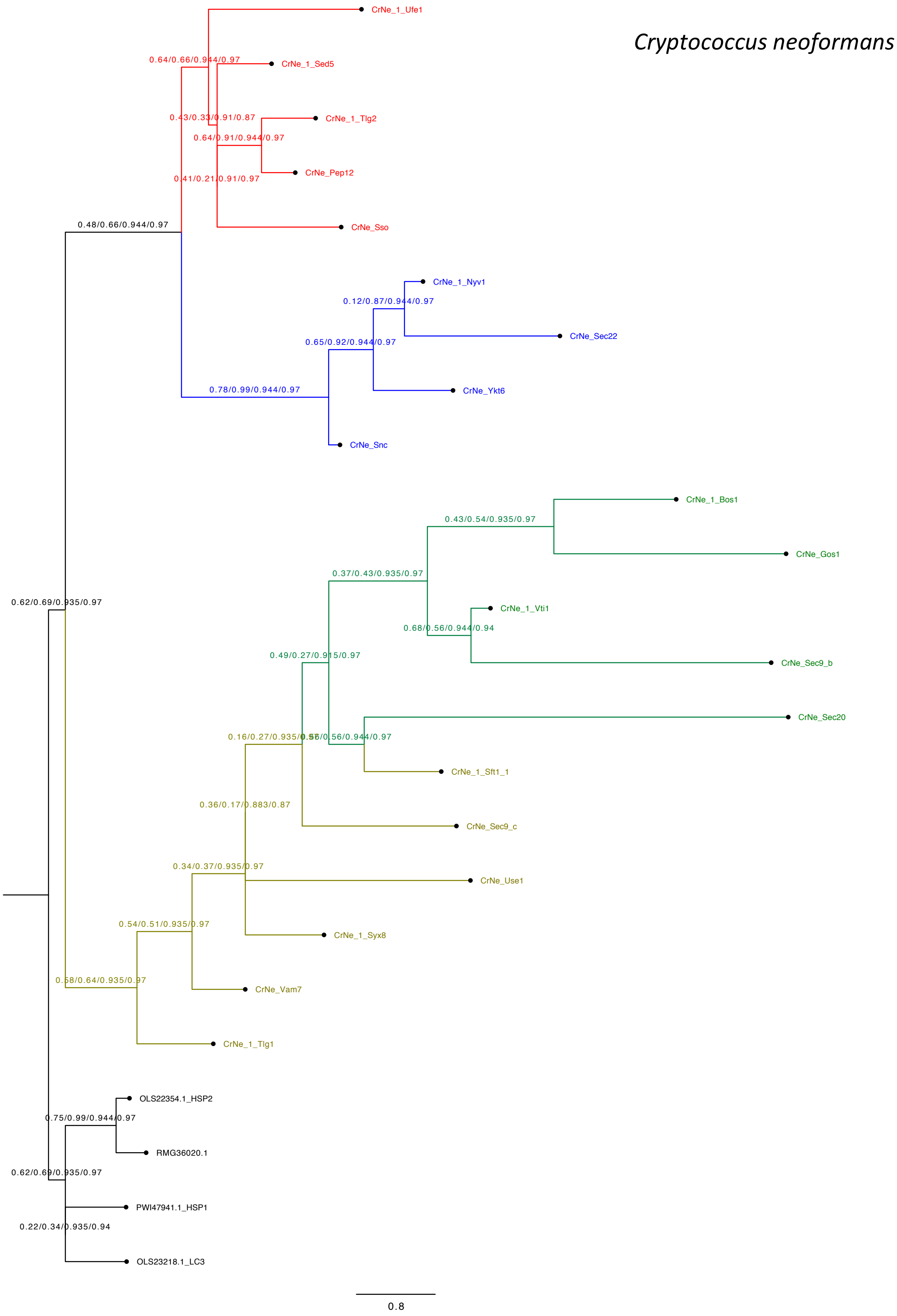

Cryptococcus neoformans
CrNe_1_Ufe1
0.64/0.66/0.944/0.97
CrNe_1_Sed5
0.43/0.33/0.91/0.87
CrNe_1_Tlg2
0.64/0.91/0.944/0.97
CrNe_Pep12
0.41/0.21/0.91/0.97
CrNe_Sso
0.48/0.66/0.944/0.97
CrNe_1_Nyv1
0.12/0.87/0.944/0.97
CrNe_Sec22
0.65/0.92/0.944/0.97
CrNe_Ykt6
0.78/0.99/0.944/0.97
CrNe_Snc
CrNe_1_Bos1
0.43/0.54/0.935/0.97
CrNe_Gos1
0.37/0.43/0.935/0.97
0.62/0.69/0.935/0.97
CrNe_1_Vti1
0.68/0.56/0.944/0.94
0.49/0.27/0.915/0.97
CrNe_Sec9_b
CrNe_Sec20
0.16/0.27/0.935/0.97
0.56/0.56/0.944/0.97
CrNe_1_Sft1_1
0.36/0.17/0.883/0.87
CrNe_Sec9_c
0.34/0.37/0.935/0.97
CrNe_Use1
0.54/0.51/0.935/0.97
CrNe_1_Syx8
0.58/0.64/0.935/0.97
CrNe_Vam7
CrNe_1_Tlg1
OLS22354.1_HSP2
0.75/0.99/0.944/0.97
RMG36020.1
0.62/0.69/0.935/0.97
PWI47941.1_HSP1
0.22/0.34/0.935/0.94
OLS23218.1_LC3
0.8

I

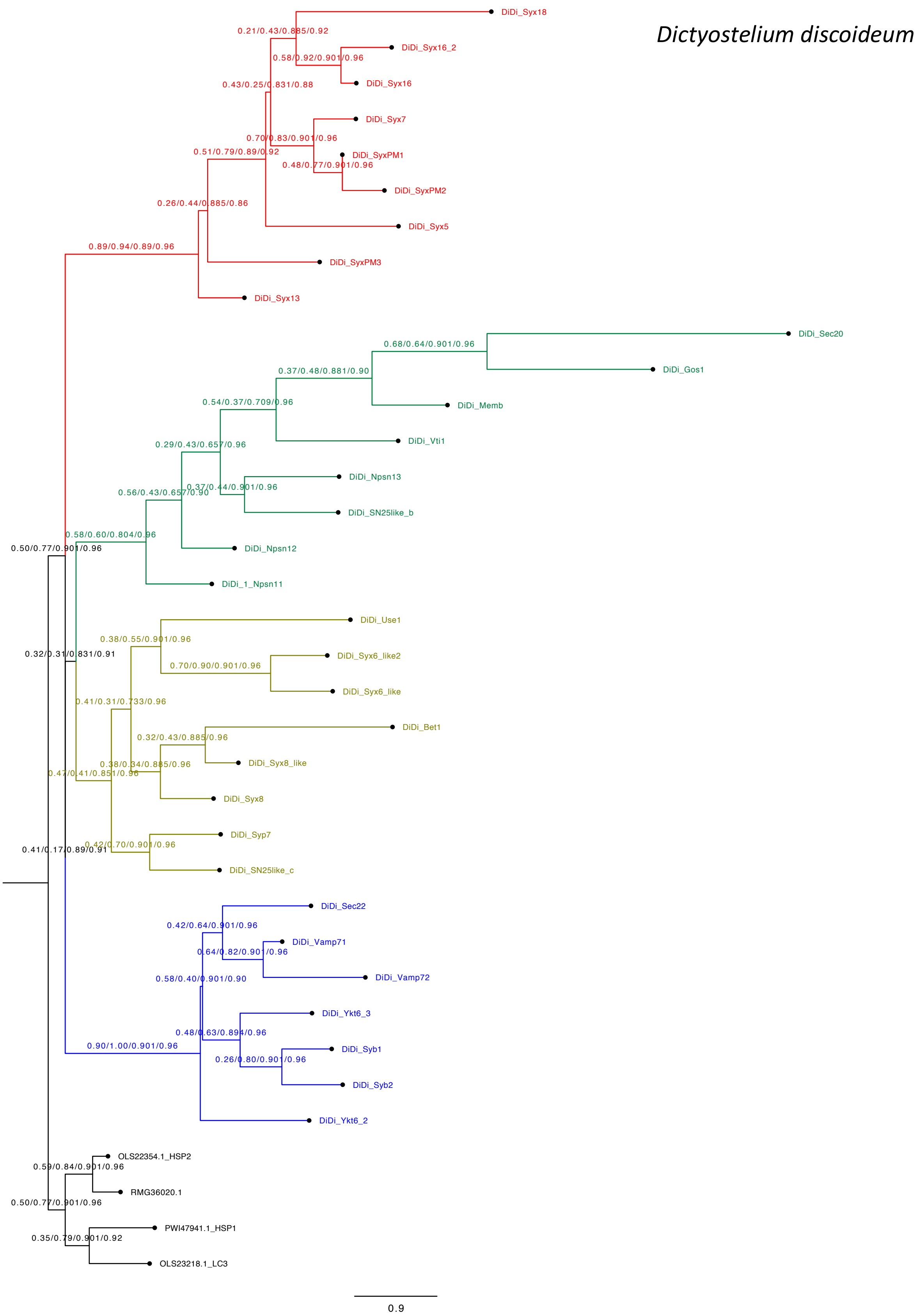
Dictyostelium discoideum
DiDi_Syx18
DiDi_Syx16_2
DiDi_Syx16
DiDi_Syx7
DiDi_SyxPM1
DiDi_SyxPM2
DiDi_Syx5
DiDi_SyxPM3
DiDi_Syx13
DiDi_Sec20
DiDi_Gos1
DiDi_Memb
DiDi_Vti1
DiDi_Npsn13
DiDi_SN25like_b
DiDi_Npsn12
DiDi_1_Npsn11
DiDi_Use1
DiDi_Syx6_like2
DiDi_Syx6_like
DiDi_Bet1
DiDi_Syx8_like
DiDi_Syx8
DiDi_Syp7
DiDi_SN25like_c
DiDi_Sec22
DiDi_Vamp71
DiDi_Vamp72
DiDi_Ykt6_3
DiDi_Syb1
DiDi_Syb2
DiDi_Ykt6_2
OLS22354.1_HSP2
RMG36020.1
PWI47941.1_HSP1
OLS23218.1_LC3
0.21/0.43/0.885/0.92
0.58/0.92/0.901/0.96
0.43/0.25/0.831/0.88
0.70/0.83/0.901/0.96
0.51/0.79/0.89/0.92
0.48/0.77/0.901/0.96
0.26/0.44/0.885/0.86
0.89/0.94/0.89/0.96
0.68/0.64/0.901/0.96
0.37/0.48/0.881/0.90
0.54/0.37/0.709/0.96
0.29/0.43/0.657/0.96
0.37/0.44/0.901/0.96
0.56/0.43/0.657/0.90
0.58/0.60/0.804/0.96
0.50/0.77/0.901/0.96
0.38/0.55/0.901/0.96
0.32/0.31/0.831/0.91
0.70/0.90/0.901/0.96
0.41/0.31/0.733/0.96
0.32/0.43/0.885/0.96
0.38/0.34/0.885/0.96
0.47/0.41/0.851/0.96
0.42/0.70/0.901/0.96
0.41/0.17/0.89/0.91
0.42/0.64/0.901/0.96
0.64/0.82/0.901/0.96
0.58/0.40/0.901/0.90
0.48/0.63/0.894/0.96
0.90/1.00/0.901/0.96
0.26/0.80/0.901/0.96
0.59/0.84/0.901/0.96
0.50/0.77/0.901/0.96
0.35/0.79/0.901/0.92
0.9

J

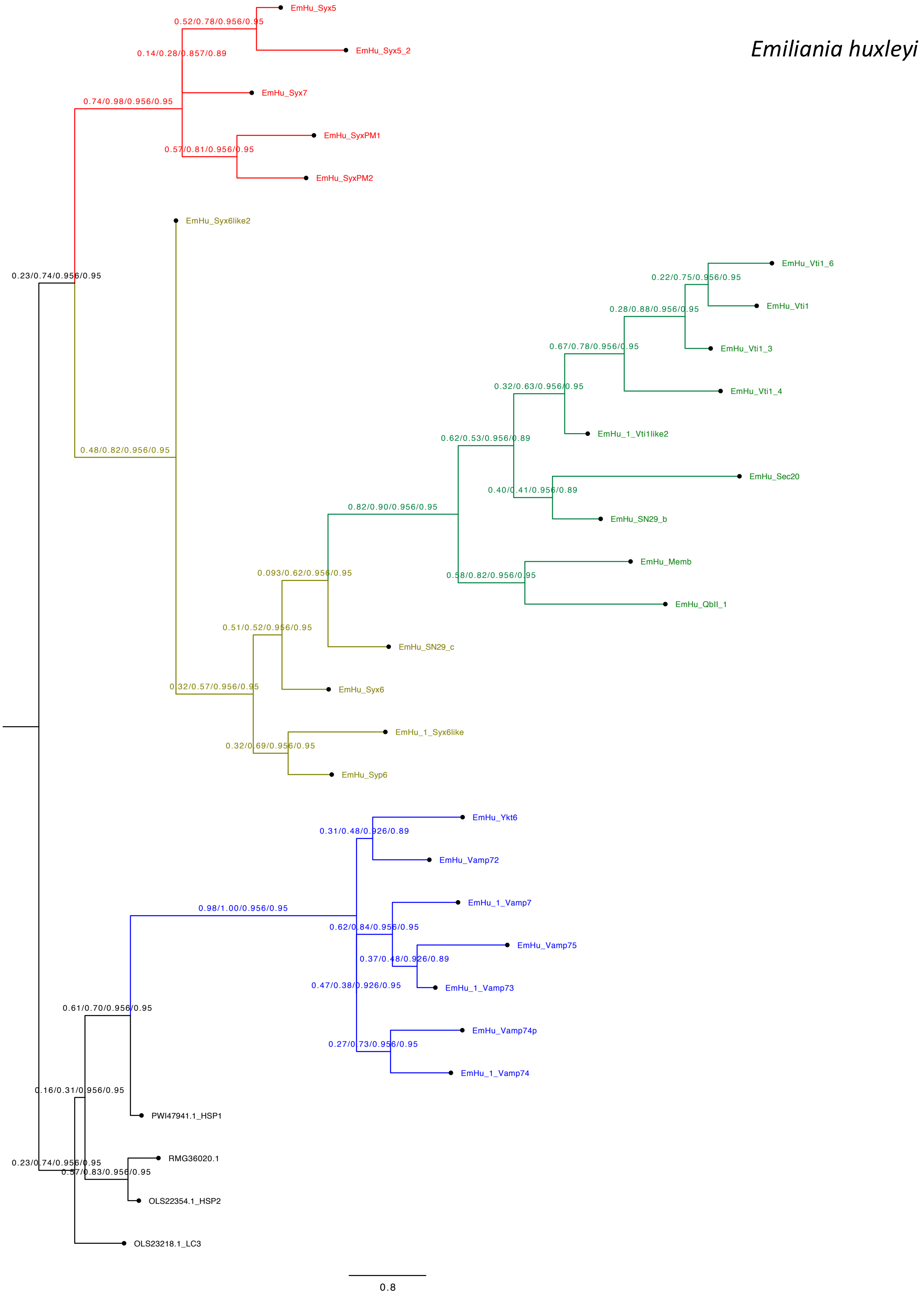
Emiliania huxleyi
EmHu_Syx5
0.52/0.78/0.956/0.95
EmHu_Syx5_2
0.14/0.28/0.857/0.89
EmHu_Syx7
0.74/0.98/0.956/0.95
EmHu_SyxPM1
0.57/0.81/0.956/0.95
EmHu_SyxPM2
EmHu_Syx6like2
EmHu_Vti1_6
0.23/0.74/0.956/0.95
0.22/0.75/0.956/0.95
EmHu_Vti1
0.28/0.88/0.956/0.95
EmHu_Vti1_3
0.67/0.78/0.956/0.95
EmHu_Vti1_4
0.32/0.63/0.956/0.95
EmHu_1_Vti1like2
0.62/0.53/0.956/0.89
0.48/0.82/0.956/0.95
EmHu_Sec20
0.40/0.41/0.956/0.89
EmHu_SN29_b
0.82/0.90/0.956/0.95
EmHu_Memb
0.093/0.62/0.956/0.95
0.58/0.82/0.956/0.95
EmHu_QbII_1
0.51/0.52/0.956/0.95
EmHu_SN29_c
EmHu_Syx6
0.32/0.57/0.956/0.95
EmHu_1_Syx6like
0.32/0.69/0.956/0.95
EmHu_Syp6
EmHu_Ykt6
0.31/0.48/0.926/0.89
EmHu_Vamp72
EmHu_1_Vamp7
0.98/1.00/0.956/0.95
0.62/0.84/0.956/0.95
EmHu_Vamp75
0.37/0.48/0.926/0.89
0.47/0.38/0.926/0.95
EmHu_1_Vamp73
0.61/0.70/0.956/0.95
EmHu_Vamp74p
0.27/0.73/0.956/0.95
EmHu_1_Vamp74
0.16/0.31/0.956/0.95
PWI47941.1_HSP1
RMG36020.1
0.23/0.74/0.956/0.95
0.57/0.83/0.956/0.95
OLS22354.1_HSP2
OLS23218.1_LC3
0.8

K

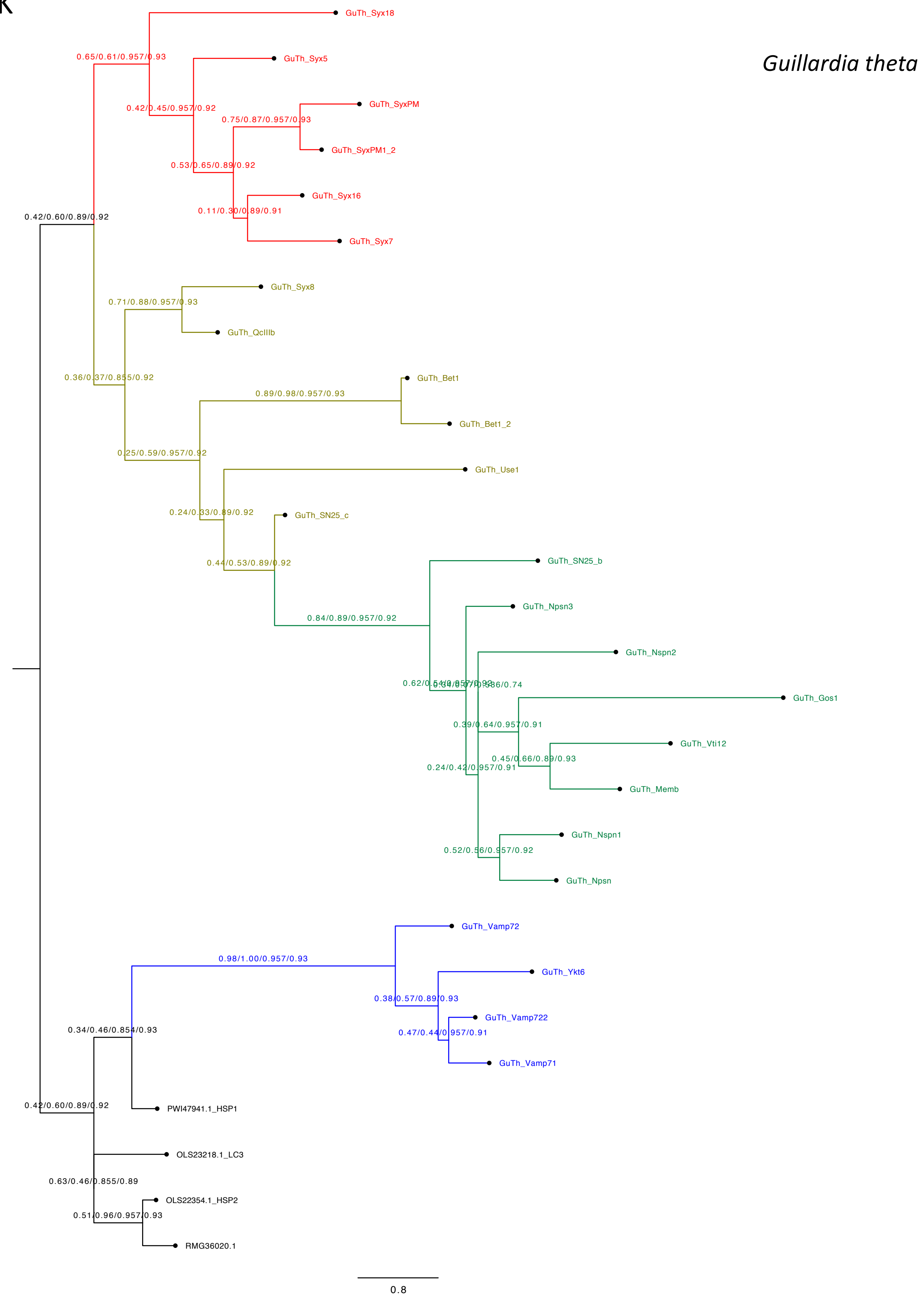

Guillardia theta
GuTh_Syx18
0.65/0.61/0.957/0.93
GuTh_Syx5
0.42/0.45/0.957/0.92
GuTh_SyxPM
0.75/0.87/0.957/0.93
GuTh_SyxPM1_2
0.53/0.65/0.89/0.92
GuTh_Syx16
0.11/0.30/0.89/0.91
GuTh_Syx7
0.42/0.60/0.89/0.92
GuTh_Syx8
0.71/0.88/0.957/0.93
GuTh_QcIIIb
0.36/0.37/0.855/0.92
GuTh_Bet1
0.89/0.98/0.957/0.93
GuTh_Bet1_2
0.25/0.59/0.957/0.92
GuTh_Use1
0.24/0.33/0.89/0.92
GuTh_SN25_c
0.44/0.53/0.89/0.92
GuTh_SN25_b
GuTh_Npsn3
0.84/0.89/0.957/0.92
GuTh_Nspn2
GuTh_Gos1
0.39/0.64/0.957/0.91
GuTh_Vti12
0.45/0.66/0.89/0.93
0.24/0.42/0.957/0.91
GuTh_Memb
GuTh_Nspn1
0.52/0.56/0.957/0.92
GuTh_Npsn
GuTh_Vamp72
0.98/1.00/0.957/0.93
GuTh_Ykt6
0.38/0.57/0.89/0.93
GuTh_Vamp722
0.47/0.44/0.957/0.91
GuTh_Vamp71
0.34/0.46/0.854/0.93
PWI47941.1_HSP1
0.42/0.60/0.89/0.92
OLS23218.1_LC3
0.63/0.46/0.855/0.89
OLS22354.1_HSP2
0.51/0.96/0.957/0.93
RMG36020.1
0.8

L

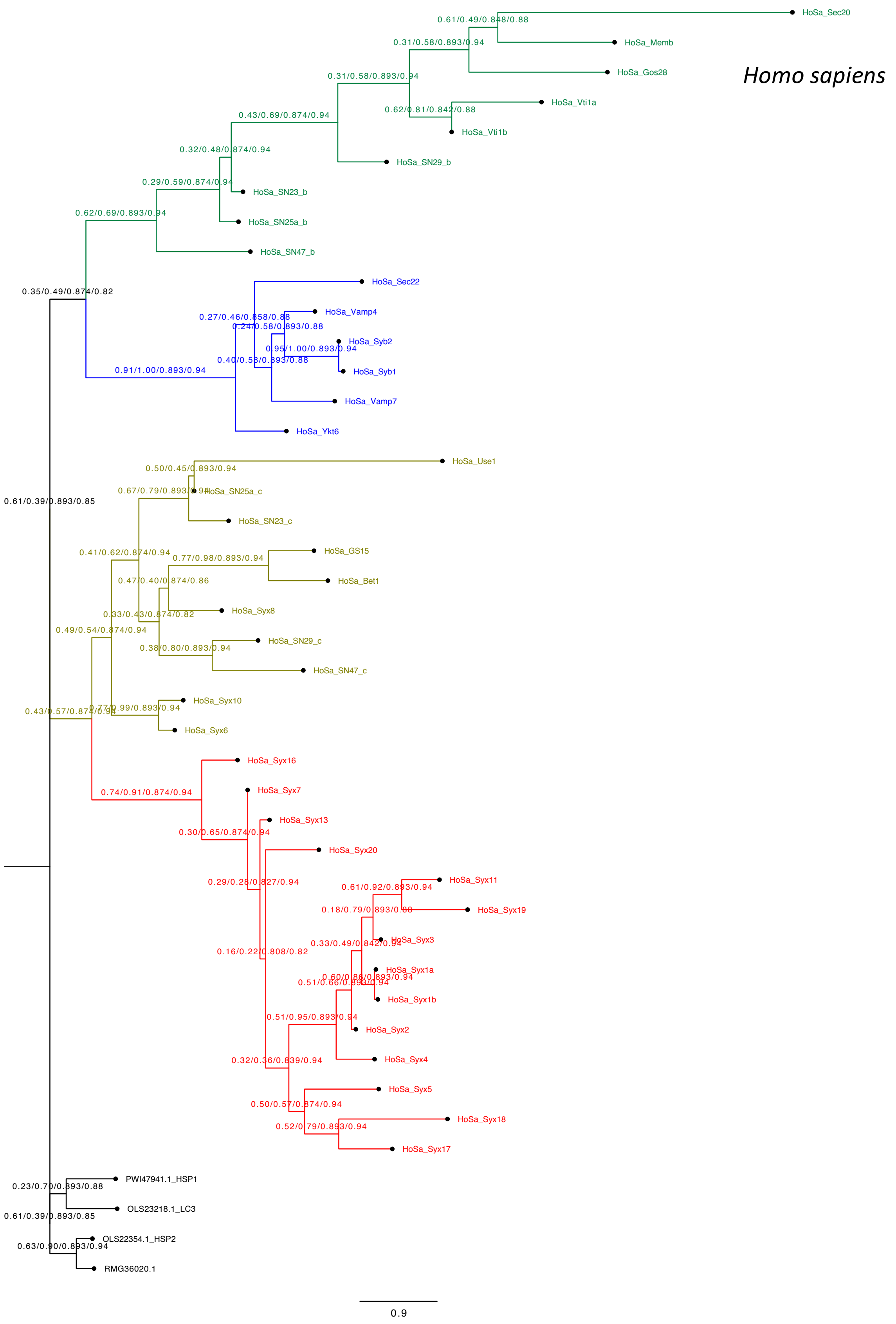

Homo sapiens
HoSa_Sec20
HoSa_Memb
HoSa_Gos28
HoSa_Vti1a
HoSa_Vti1b
HoSa_SN29_b
HoSa_SN23_b
HoSa_SN25a_b
HoSa_SN47_b
HoSa_Sec22
HoSa_Vamp4
HoSa_Syb2
HoSa_Syb1
HoSa_Vamp7
HoSa_Ykt6
HoSa_Use1
HoSa_SN25a_c
HoSa_SN23_c
HoSa_GS15
HoSa_Bet1
HoSa_Syx8
HoSa_SN29_c
HoSa_SN47_c
HoSa_Syx10
HoSa_Syx6
HoSa_Syx16
HoSa_Syx7
HoSa_Syx13
HoSa_Syx20
HoSa_Syx11
HoSa_Syx19
HoSa_Syx3
HoSa_Syx1a
HoSa_Syx1b
HoSa_Syx2
HoSa_Syx4
HoSa_Syx5
HoSa_Syx18
HoSa_Syx17
PWI47941.1_HSP1
OLS23218.1_LC3
OLS22354.1_HSP2
RMG36020.1
0.9

M

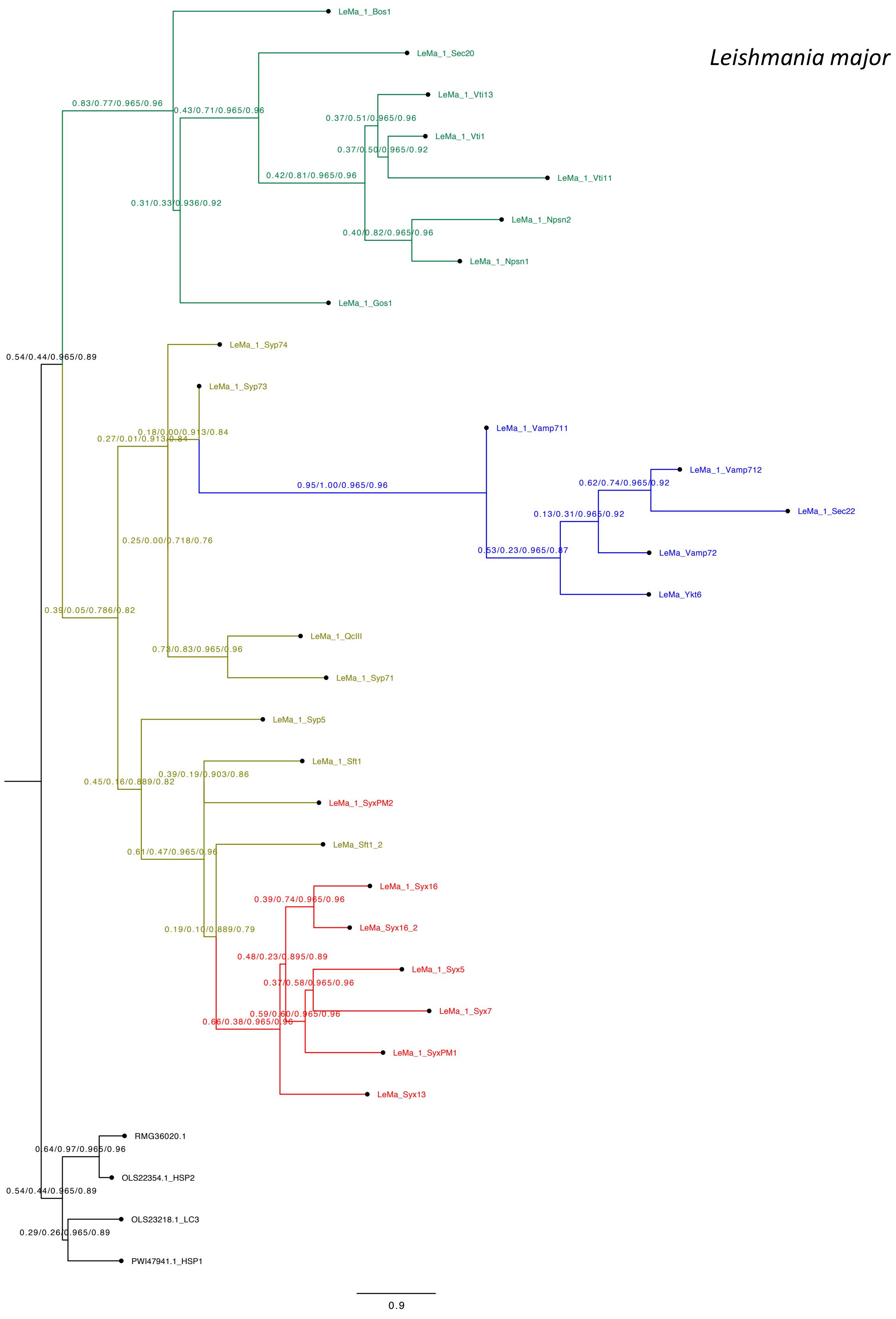
Leishmania major
LeMa_1_Bos1
LeMa_1_Sec20
LeMa_1_Vti13
LeMa_1_Vti1
LeMa_1_Vti11
LeMa_1_Npsn2
LeMa_1_Npsn1
LeMa_1_Gos1
LeMa_1_Syp74
LeMa_1_Syp73
LeMa_1_Vamp711
LeMa_1_Vamp712
LeMa_1_Sec22
LeMa_Vamp72
LeMa_Ykt6
LeMa_1_QcIII
LeMa_1_Syp71
LeMa_1_Syp5
LeMa_1_Sft1
LeMa_1_SyxPM2
LeMa_Sft1_2
LeMa_1_Syx16
LeMa_Syx16_2
LeMa_1_Syx5
LeMa_1_Syx7
LeMa_1_SyxPM1
LeMa_Syx13
RMG36020.1
OLS22354.1_HSP2
OLS23218.1_LC3
PWI47941.1_HSP1
0.83/0.77/0.965/0.96
0.43/0.71/0.965/0.96
0.37/0.51/0.965/0.96
0.37/0.50/0.965/0.92
0.42/0.81/0.965/0.96
0.31/0.33/0.936/0.92
0.40/0.82/0.965/0.96
0.54/0.44/0.965/0.89
0.18/0.00/0.913/0.84
0.27/0.01/0.913/0.84
0.95/1.00/0.965/0.96
0.62/0.74/0.965/0.92
0.13/0.31/0.965/0.92
0.53/0.23/0.965/0.87
0.25/0.00/0.718/0.76
0.39/0.05/0.786/0.82
0.73/0.83/0.965/0.96
0.39/0.19/0.903/0.86
0.45/0.16/0.889/0.82
0.61/0.47/0.965/0.96
0.39/0.74/0.965/0.96
0.19/0.10/0.889/0.79
0.48/0.23/0.895/0.89
0.37/0.58/0.965/0.96
0.59/0.60/0.965/0.96
0.66/0.38/0.965/0.96
0.64/0.97/0.965/0.96
0.54/0.44/0.965/0.89
0.29/0.26/0.965/0.89
0.9

N

*Monosiga brevicollis*

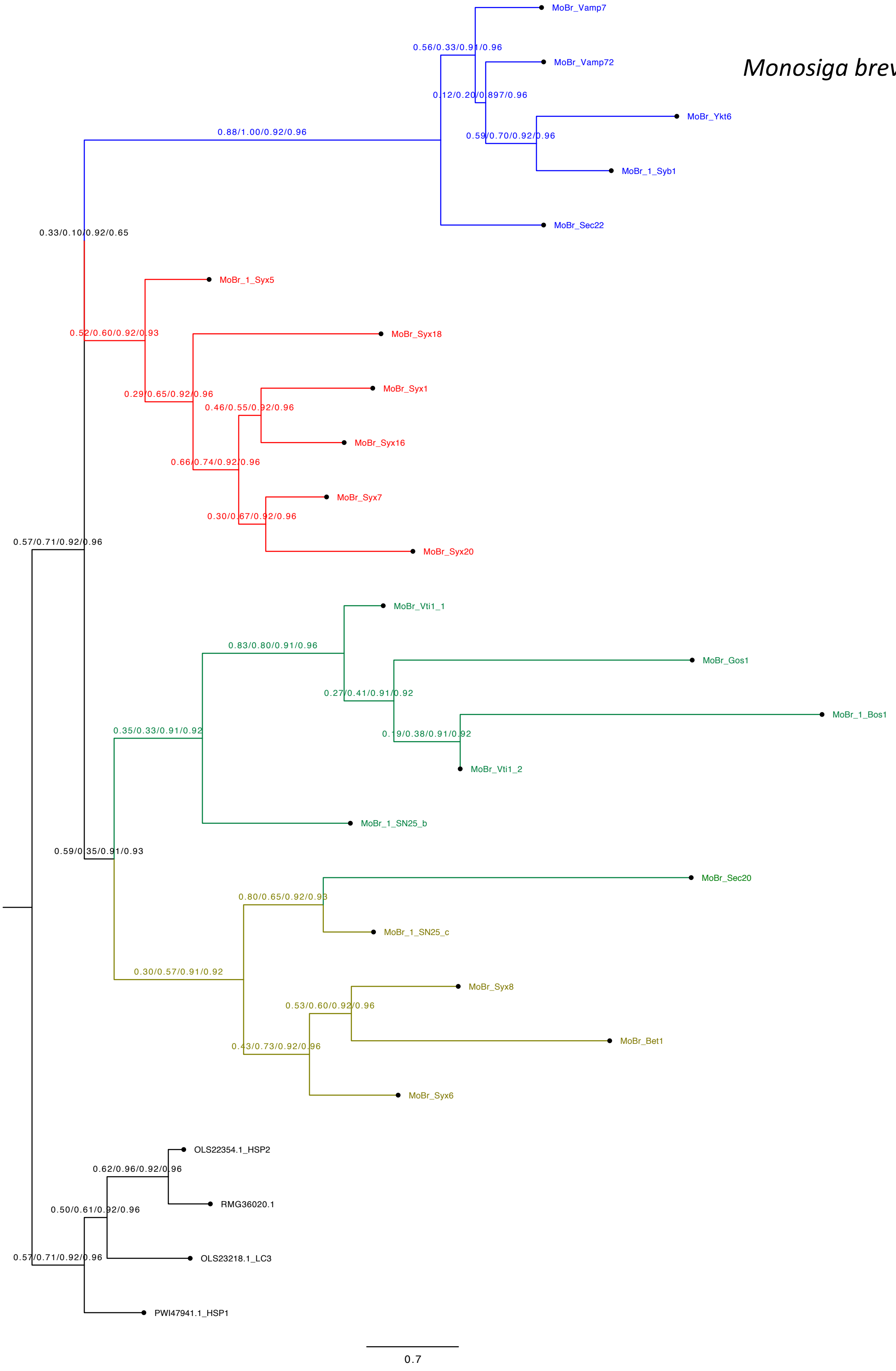

O

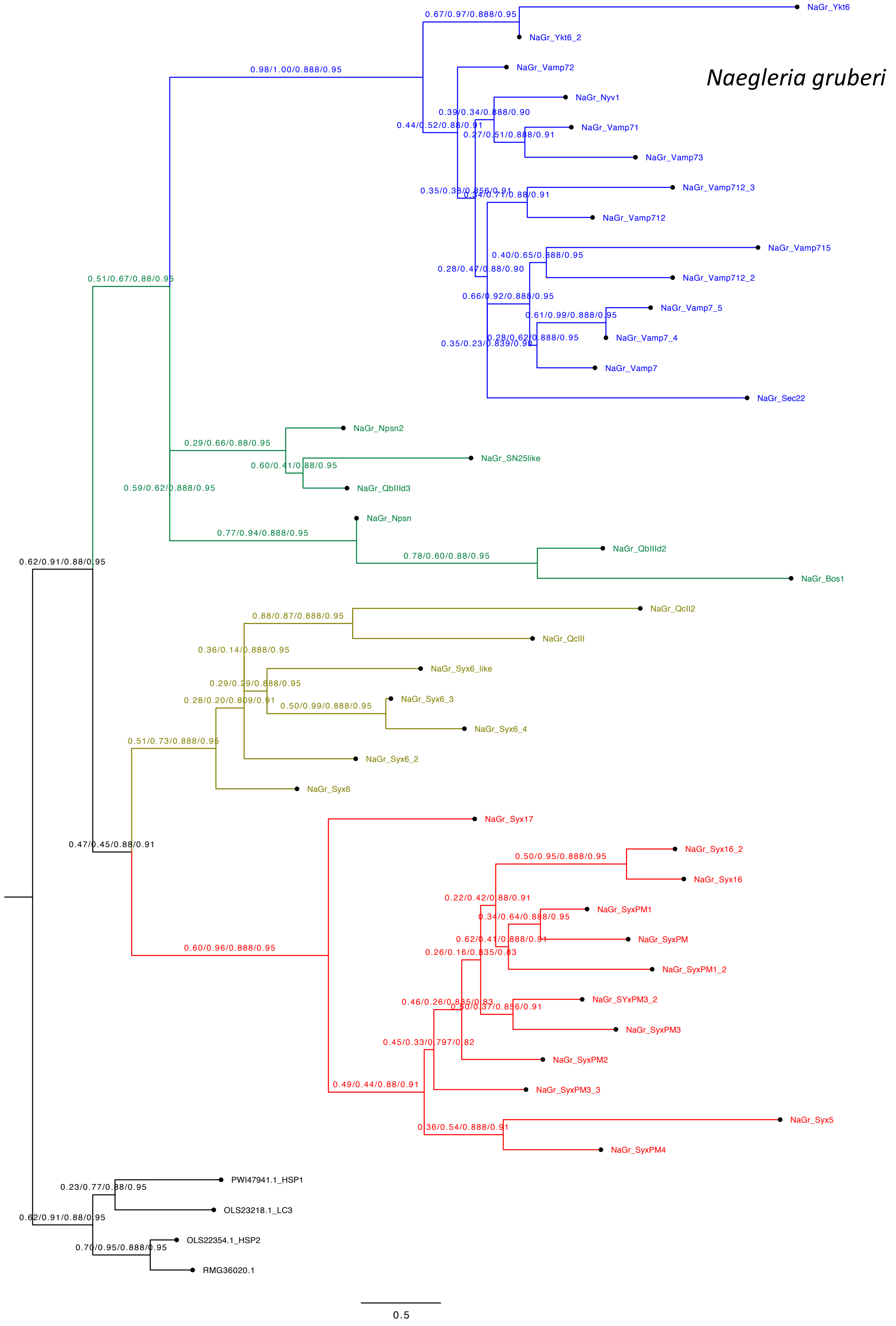

Naegleria gruberi
NaGr_Ykt6
NaGr_Ykt6_2
NaGr_Vamp72
NaGr_Nyv1
NaGr_Vamp71
NaGr_Vamp73
NaGr_Vamp712_3
NaGr_Vamp712
NaGr_Vamp715
NaGr_Vamp712_2
NaGr_Vamp7_5
NaGr_Vamp7_4
NaGr_Vamp7
NaGr_Sec22
NaGr_Npsn2
NaGr_SN25like
NaGr_QbIIId3
NaGr_Npsn
NaGr_QbIIId2
NaGr_Bos1
NaGr_QcII2
NaGr_QcIII
NaGr_Syx6_like
NaGr_Syx6_3
NaGr_Syx6_4
NaGr_Syx6_2
NaGr_Syx6
NaGr_Syx17
NaGr_Syx16_2
NaGr_Syx16
NaGr_SyxPM1
NaGr_SyxPM
NaGr_SyxPM1_2
NaGr_SYxPM3_2
NaGr_SyxPM3
NaGr_SyxPM2
NaGr_SyxPM3_3
NaGr_Syx5
NaGr_SyxPM4
PWI47941.1_HSP1
OLS23218.1_LC3
OLS22354.1_HSP2
RMG36020.1
0.67/0.97/0.888/0.95
0.98/1.00/0.888/0.95
0.39/0.34/0.888/0.90
0.44/0.52/0.88/0.91
0.27/0.51/0.888/0.91
0.35/0.38/0.856/0.91
0.34/0.71/0.88/0.91
0.40/0.65/0.888/0.95
0.28/0.47/0.88/0.90
0.66/0.92/0.888/0.95
0.61/0.99/0.888/0.95
0.28/0.62/0.888/0.95
0.35/0.23/0.839/0.90
0.51/0.67/0.88/0.95
0.29/0.66/0.88/0.95
0.60/0.41/0.88/0.95
0.59/0.62/0.888/0.95
0.77/0.94/0.888/0.95
0.78/0.60/0.88/0.95
0.62/0.91/0.88/0.95
0.88/0.87/0.888/0.95
0.36/0.14/0.888/0.95
0.29/0.29/0.888/0.95
0.28/0.20/0.809/0.91
0.50/0.99/0.888/0.95
0.51/0.73/0.888/0.95
0.47/0.45/0.88/0.91
0.50/0.95/0.888/0.95
0.22/0.42/0.88/0.91
0.34/0.64/0.888/0.95
0.62/0.41/0.888/0.91
0.26/0.16/0.835/0.83
0.60/0.96/0.888/0.95
0.46/0.26/0.835/0.83
0.50/0.37/0.856/0.91
0.45/0.33/0.797/0.82
0.49/0.44/0.88/0.91
0.36/0.54/0.888/0.91
0.23/0.77/0.88/0.95
0.62/0.91/0.88/0.95
0.70/0.95/0.888/0.95
0.5

P

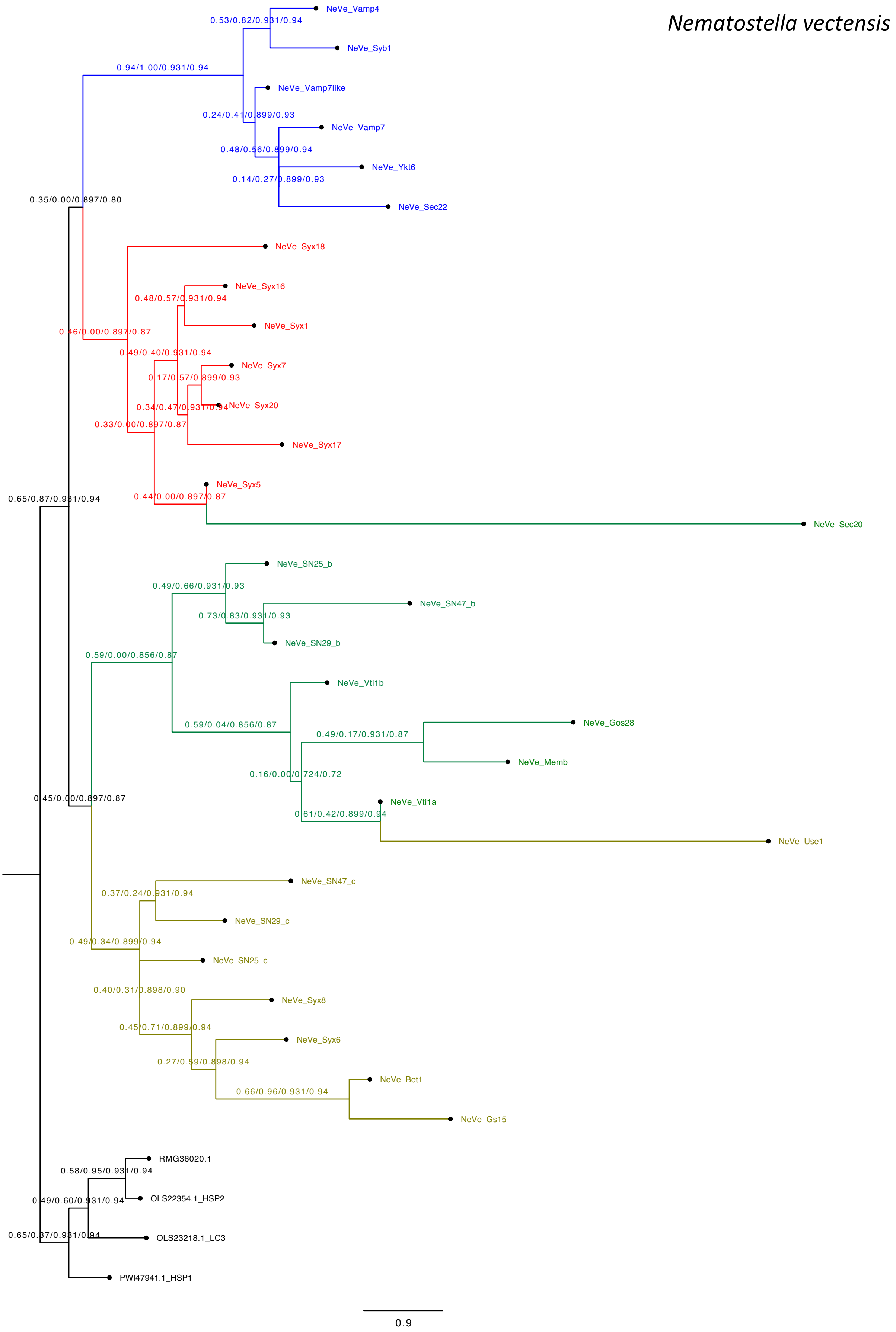
Nematostella vectensis
NeVe_Vamp4
0.53/0.82/0.931/0.94
NeVe_Syb1
0.94/1.00/0.931/0.94
NeVe_Vamp7like
0.24/0.41/0.899/0.93
NeVe_Vamp7
0.48/0.56/0.899/0.94
NeVe_Ykt6
0.14/0.27/0.899/0.93
NeVe_Sec22
0.35/0.00/0.897/0.80
NeVe_Syx18
NeVe_Syx16
0.48/0.57/0.931/0.94
NeVe_Syx1
0.46/0.00/0.897/0.87
0.49/0.40/0.931/0.94
NeVe_Syx7
0.17/0.57/0.899/0.93
0.34/0.47/0.931/0.94
NeVe_Syx20
0.33/0.00/0.897/0.87
NeVe_Syx17
NeVe_Syx5
0.65/0.87/0.931/0.94
0.44/0.00/0.897/0.87
NeVe_Sec20
NeVe_SN25_b
0.49/0.66/0.931/0.93
NeVe_SN47_b
0.73/0.83/0.931/0.93
NeVe_SN29_b
0.59/0.00/0.856/0.87
NeVe_Vti1b
NeVe_Gos28
0.59/0.04/0.856/0.87
0.49/0.17/0.931/0.87
NeVe_Memb
0.16/0.00/0.724/0.72
0.45/0.00/0.897/0.87
NeVe_Vti1a
0.61/0.42/0.899/0.94
NeVe_Use1
NeVe_SN47_c
0.37/0.24/0.931/0.94
NeVe_SN29_c
0.49/0.34/0.899/0.94
NeVe_SN25_c
0.40/0.31/0.898/0.90
NeVe_Syx8
0.45/0.71/0.899/0.94
NeVe_Syx6
0.27/0.59/0.898/0.94
NeVe_Bet1
0.66/0.96/0.931/0.94
NeVe_Gs15
RMG36020.1
0.58/0.95/0.931/0.94
OLS22354.1_HSP2
0.49/0.60/0.931/0.94
0.65/0.87/0.931/0.94
OLS23218.1_LC3
PWI47941.1_HSP1
0.9

Q

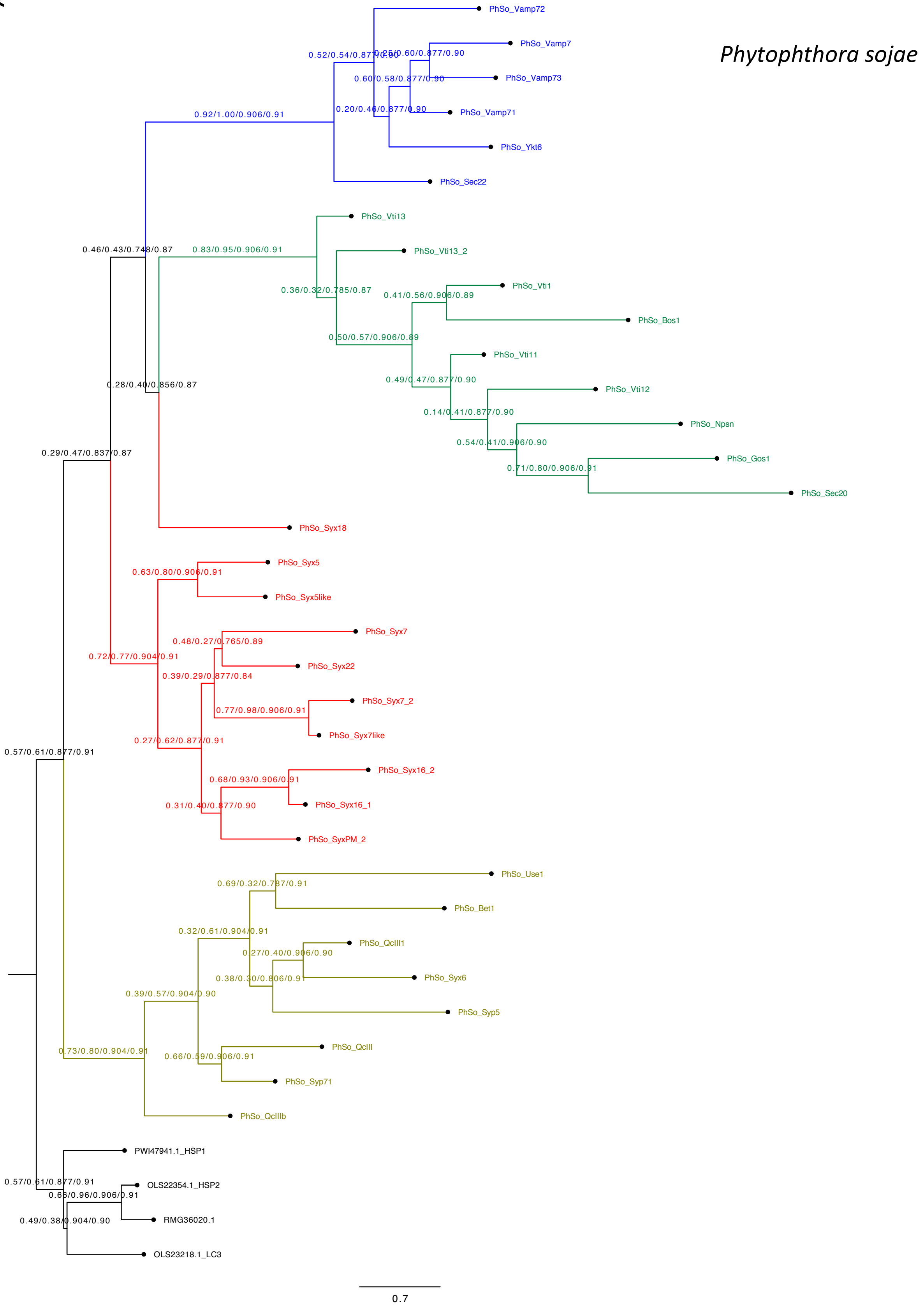
Phytophthora sojae
PhSo_Vamp72
PhSo_Vamp7
PhSo_Vamp73
PhSo_Vamp71
PhSo_Ykt6
PhSo_Sec22
PhSo_Vti13
PhSo_Vti13_2
PhSo_Vti1
PhSo_Bos1
PhSo_Vti11
PhSo_Vti12
PhSo_Npsn
PhSo_Gos1
PhSo_Sec20
PhSo_Syx18
PhSo_Syx5
PhSo_Syx5like
PhSo_Syx7
PhSo_Syx22
PhSo_Syx7_2
PhSo_Syx7like
PhSo_Syx16_2
PhSo_Syx16_1
PhSo_SyxPM_2
PhSo_Use1
PhSo_Bet1
PhSo_QcIII1
PhSo_Syx6
PhSo_Syp5
PhSo_QcIII
PhSo_Syp71
PhSo_QcIIIb
PWI47941.1_HSP1
OLS22354.1_HSP2
RMG36020.1
OLS23218.1_LC3
0.92/1.00/0.906/0.91
0.52/0.54/0.877/0.90
0.25/0.60/0.877/0.90
0.60/0.58/0.877/0.90
0.20/0.46/0.877/0.90
0.46/0.43/0.748/0.87
0.83/0.95/0.906/0.91
0.36/0.32/0.785/0.87
0.41/0.56/0.906/0.89
0.50/0.57/0.906/0.89
0.49/0.47/0.877/0.90
0.14/0.41/0.877/0.90
0.54/0.41/0.906/0.90
0.71/0.80/0.906/0.91
0.28/0.40/0.856/0.87
0.29/0.47/0.837/0.87
0.63/0.80/0.906/0.91
0.48/0.27/0.765/0.89
0.72/0.77/0.904/0.91
0.39/0.29/0.877/0.84
0.77/0.98/0.906/0.91
0.27/0.62/0.877/0.91
0.57/0.61/0.877/0.91
0.68/0.93/0.906/0.91
0.31/0.40/0.877/0.90
0.69/0.32/0.787/0.91
0.32/0.61/0.904/0.91
0.27/0.40/0.906/0.90
0.38/0.30/0.806/0.91
0.39/0.57/0.904/0.90
0.73/0.80/0.904/0.91
0.66/0.59/0.906/0.91
0.57/0.61/0.877/0.91
0.66/0.96/0.906/0.91
0.49/0.38/0.904/0.90
0.7

R

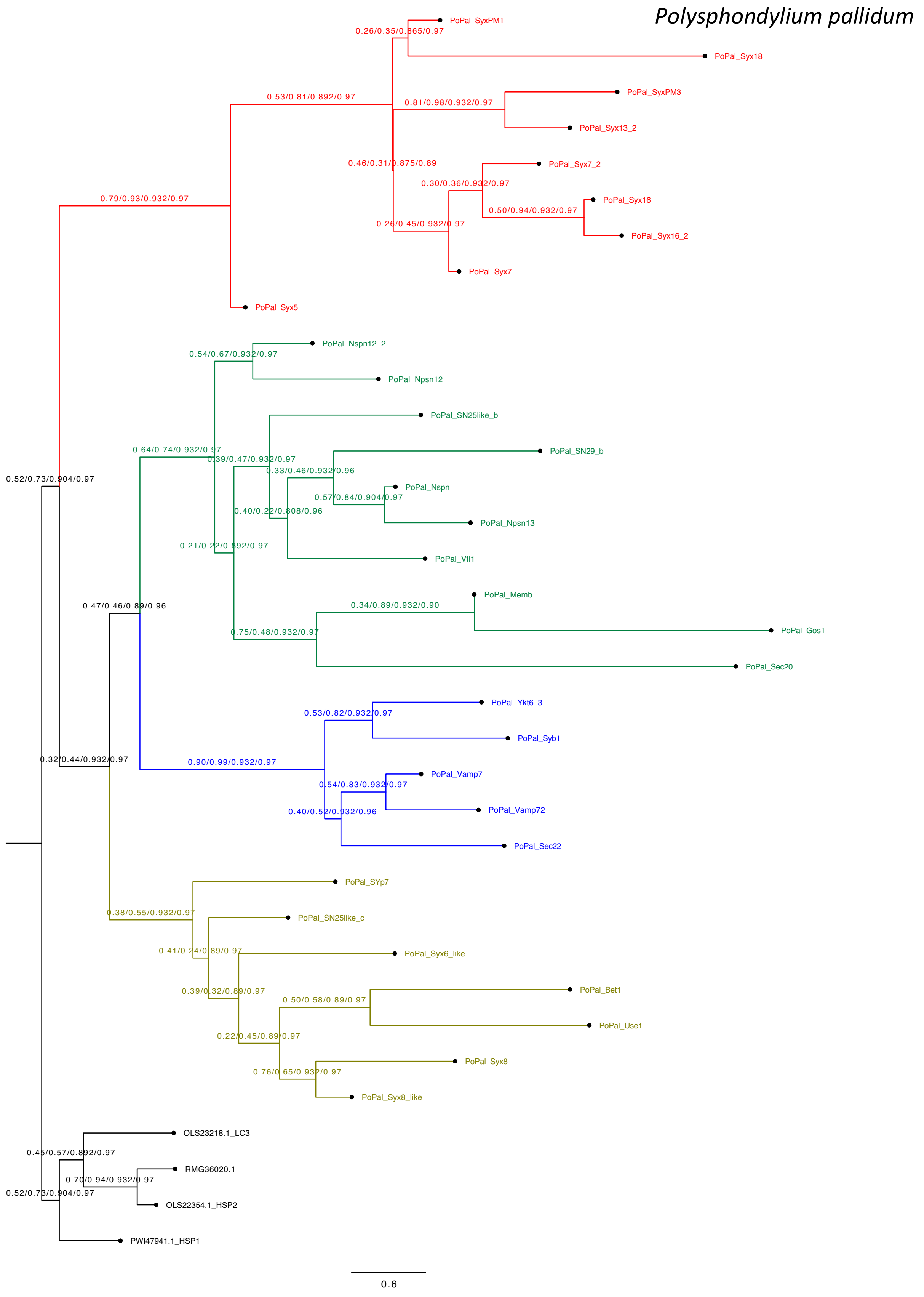

Polysphondylium pallidum
PoPal_SyxPM1
0.26/0.35/0.865/0.97
PoPal_Syx18
0.53/0.81/0.892/0.97
PoPal_SyxPM3
0.81/0.98/0.932/0.97
PoPal_Syx13_2
0.46/0.31/0.875/0.89
PoPal_Syx7_2
0.30/0.36/0.932/0.97
0.79/0.93/0.932/0.97
PoPal_Syx16
0.50/0.94/0.932/0.97
0.26/0.45/0.932/0.97
PoPal_Syx16_2
PoPal_Syx7
PoPal_Syx5
PoPal_Nspn12_2
0.54/0.67/0.932/0.97
PoPal_Npsn12
PoPal_SN25like_b
0.64/0.74/0.932/0.97
PoPal_SN29_b
0.52/0.73/0.904/0.97
0.39/0.47/0.932/0.97
0.33/0.46/0.932/0.96
PoPal_Nspn
0.57/0.84/0.904/0.97
0.40/0.22/0.808/0.96
PoPal_Npsn13
0.21/0.22/0.892/0.97
PoPal_Vti1
PoPal_Memb
0.47/0.46/0.89/0.96
0.34/0.89/0.932/0.90
PoPal_Gos1
0.75/0.48/0.932/0.97
PoPal_Sec20
PoPal_Ykt6_3
0.53/0.82/0.932/0.97
PoPal_Syb1
0.32/0.44/0.932/0.97
0.90/0.99/0.932/0.97
PoPal_Vamp7
0.54/0.83/0.932/0.97
0.40/0.52/0.932/0.96
PoPal_Vamp72
PoPal_Sec22
PoPal_SYp7
0.38/0.55/0.932/0.97
PoPal_SN25like_c
0.41/0.24/0.89/0.97
PoPal_Syx6_like
0.39/0.32/0.89/0.97
PoPal_Bet1
0.50/0.58/0.89/0.97
PoPal_Use1
0.22/0.45/0.89/0.97
PoPal_Syx8
0.76/0.65/0.932/0.97
PoPal_Syx8_like
OLS23218.1_LC3
0.45/0.57/0.892/0.97
RMG36020.1
0.70/0.94/0.932/0.97
0.52/0.73/0.904/0.97
OLS22354.1_HSP2
PWI47941.1_HSP1
0.6

S

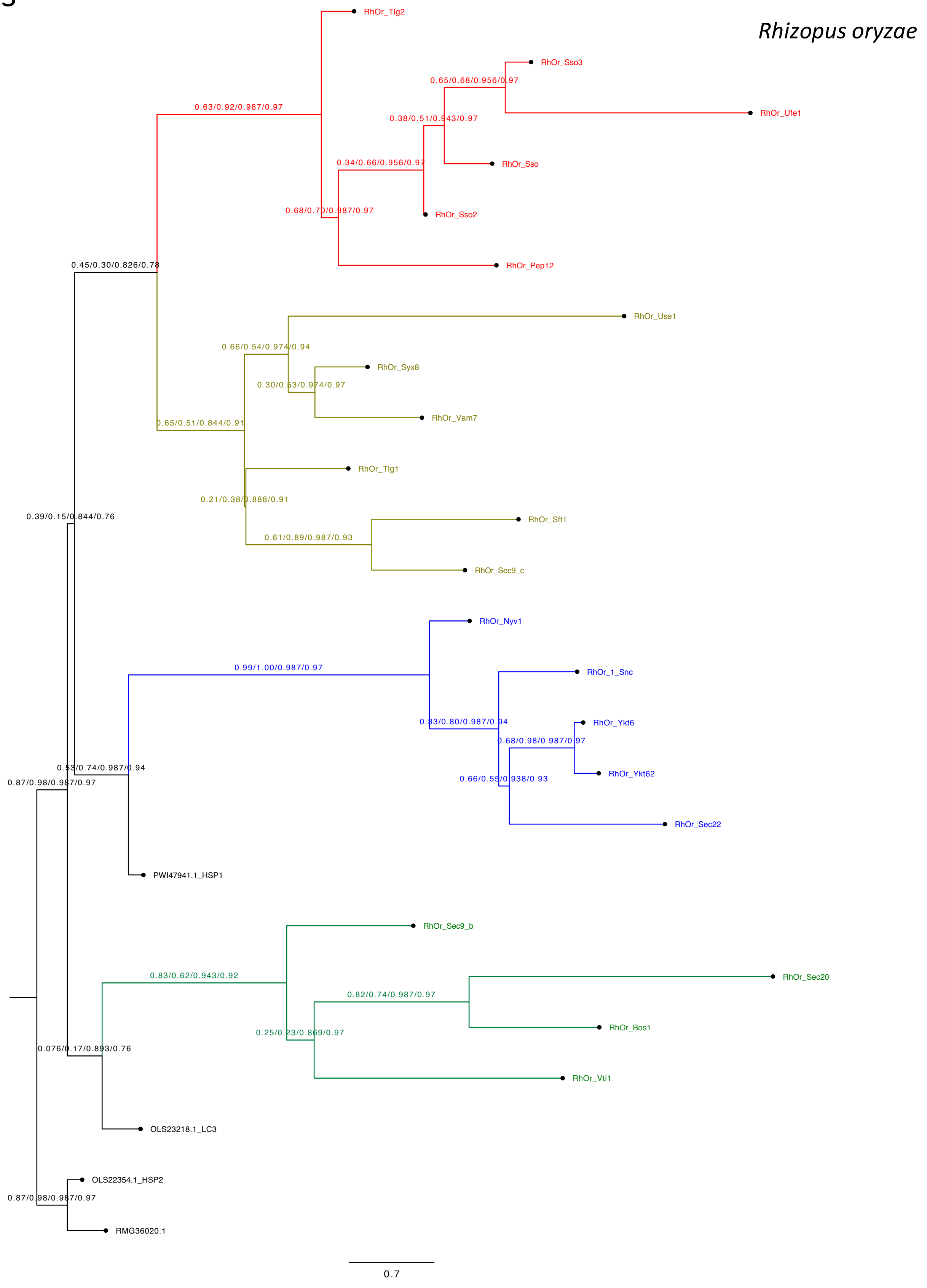
Rhizopus oryzae
RhOr_Tlg2
0.63/0.92/0.987/0.97
RhOr_Sso3
0.65/0.68/0.956/0.97
RhOr_Ufe1
0.38/0.51/0.943/0.97
0.34/0.66/0.956/0.97
RhOr_Sso
0.68/0.70/0.987/0.97
RhOr_Sso2
0.45/0.30/0.826/0.78
RhOr_Pep12
RhOr_Use1
0.66/0.54/0.974/0.94
RhOr_Syx8
0.30/0.53/0.974/0.97
RhOr_Vam7
0.65/0.51/0.844/0.91
RhOr_Tlg1
0.21/0.38/0.888/0.91
0.39/0.15/0.844/0.76
RhOr_Sft1
0.61/0.89/0.987/0.93
RhOr_Sec9_c
RhOr_Nyv1
0.99/1.00/0.987/0.97
RhOr_1_Snc
0.83/0.80/0.987/0.94
RhOr_Ykt6
0.68/0.98/0.987/0.97
0.53/0.74/0.987/0.94
RhOr_Ykt62
0.87/0.98/0.987/0.97
0.66/0.55/0.938/0.93
RhOr_Sec22
PWI47941.1_HSP1
RhOr_Sec9_b
0.83/0.62/0.943/0.92
RhOr_Sec20
0.82/0.74/0.987/0.97
RhOr_Bos1
0.25/0.23/0.869/0.97
0.076/0.17/0.893/0.76
RhOr_Vti1
OLS23218.1_LC3
OLS22354.1_HSP2
0.87/0.98/0.987/0.97
RMG36020.1
0.7

T

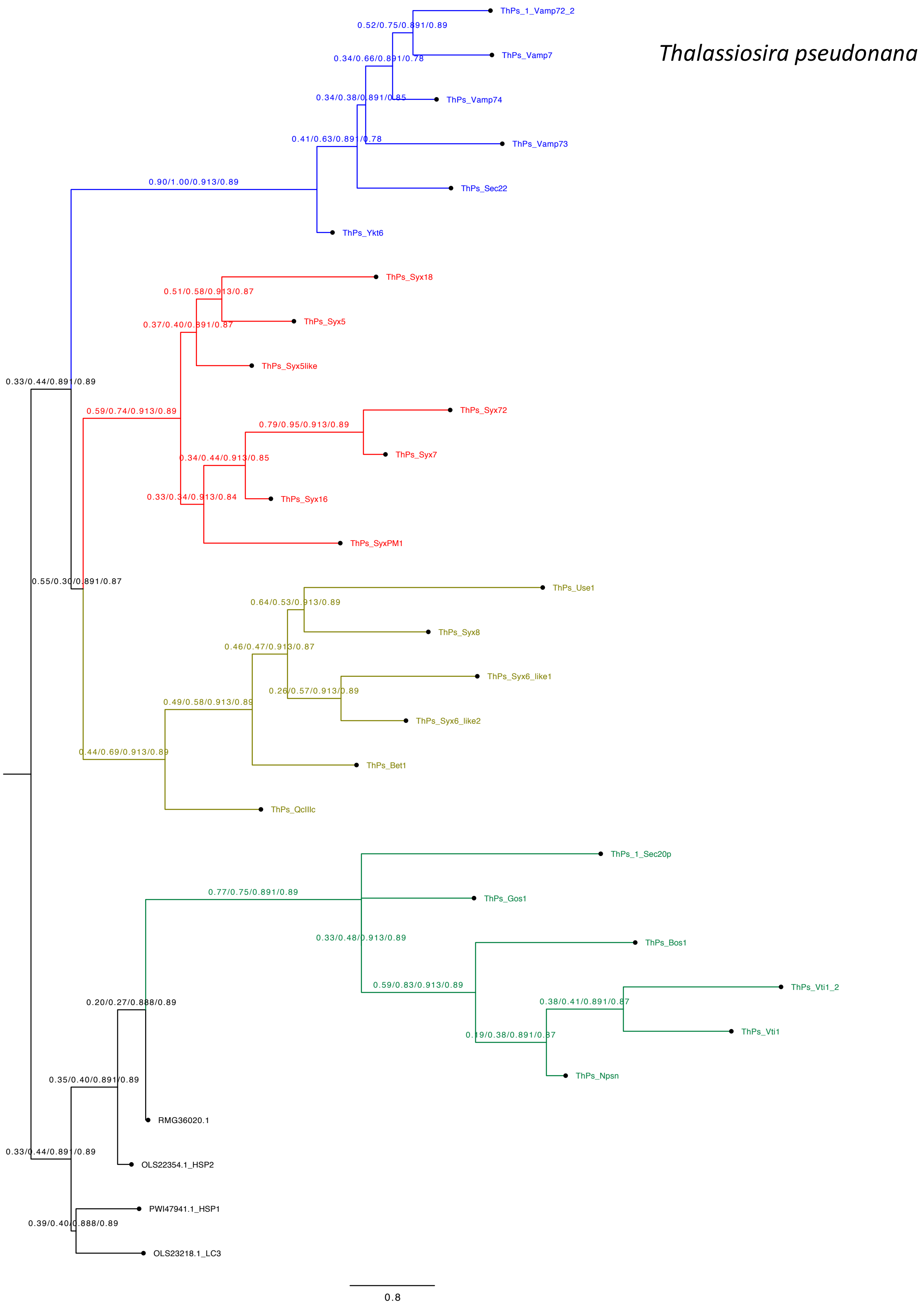

*Thalassiosira pseudonana*
ThPs_1_Vamp72_2
0.52/0.75/0.891/0.89
ThPs_Vamp7
0.34/0.66/0.891/0.78
0.34/0.38/0.891/0.85
ThPs_Vamp74
0.41/0.63/0.891/0.78
ThPs_Vamp73
0.90/1.00/0.913/0.89
ThPs_Sec22
ThPs_Ykt6
ThPs_Syx18
0.51/0.58/0.913/0.87
ThPs_Syx5
0.37/0.40/0.891/0.87
ThPs_Syx5like
0.33/0.44/0.891/0.89
0.59/0.74/0.913/0.89
ThPs_Syx72
0.79/0.95/0.913/0.89
ThPs_Syx7
0.34/0.44/0.913/0.85
0.33/0.34/0.913/0.84
ThPs_Syx16
ThPs_SyxPM1
0.55/0.30/0.891/0.87
ThPs_Use1
0.64/0.53/0.913/0.89
ThPs_Syx8
0.46/0.47/0.913/0.87
ThPs_Syx6_like1
0.26/0.57/0.913/0.89
0.49/0.58/0.913/0.89
ThPs_Syx6_like2
0.44/0.69/0.913/0.89
ThPs_Bet1
ThPs_QcIIIc
ThPs_1_Sec20p
0.77/0.75/0.891/0.89
ThPs_Gos1
0.33/0.48/0.913/0.89
ThPs_Bos1
0.59/0.83/0.913/0.89
ThPs_Vti1_2
0.20/0.27/0.888/0.89
0.38/0.41/0.891/0.87
ThPs_Vti1
0.19/0.38/0.891/0.87
ThPs_Npsn
0.35/0.40/0.891/0.89
RMG36020.1
0.33/0.44/0.891/0.89
OLS22354.1_HSP2
PWI47941.1_HSP1
0.39/0.40/0.888/0.89
OLS23218.1_LC3
0.8

U

*Thecamonas trahens*

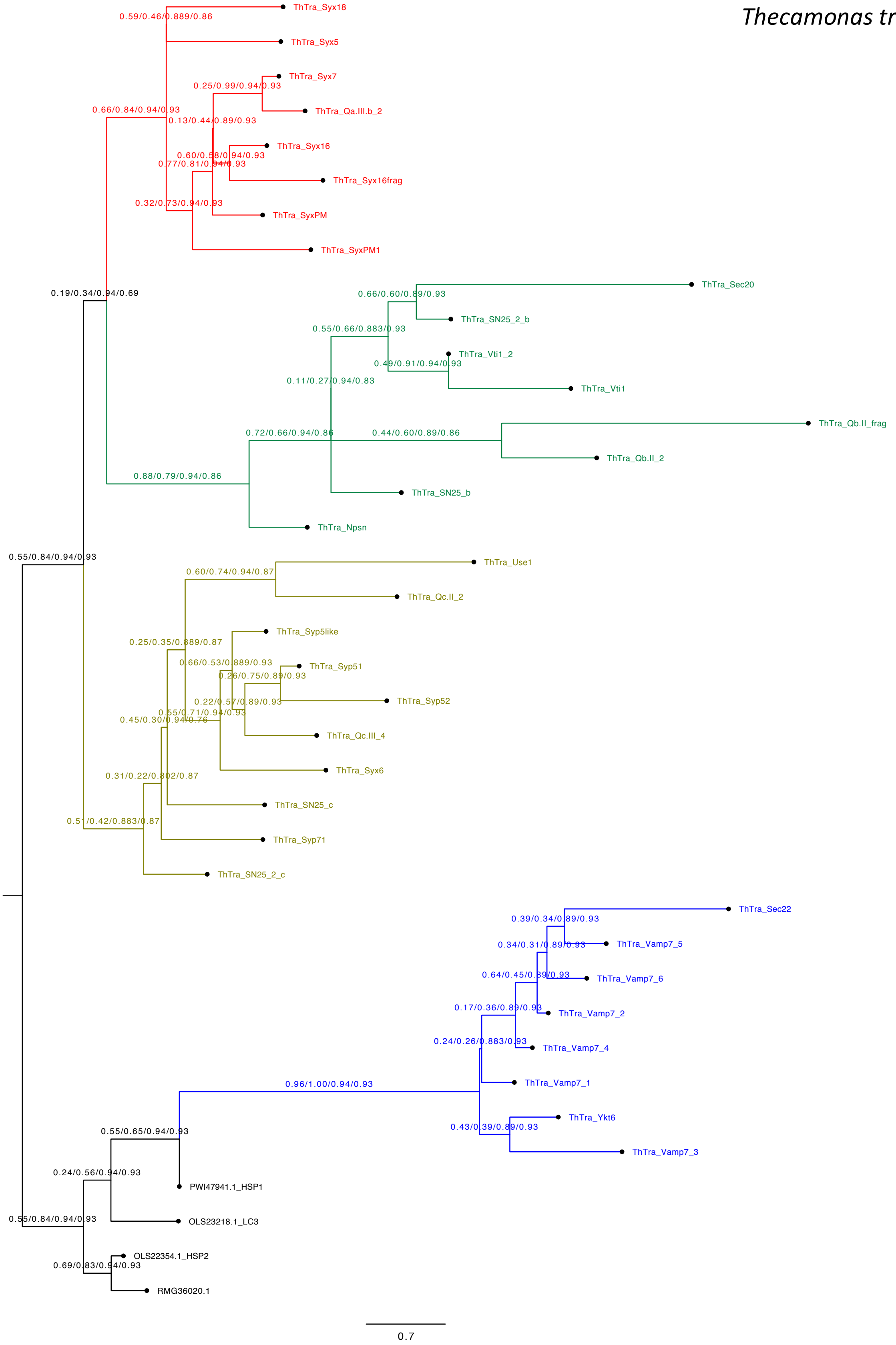

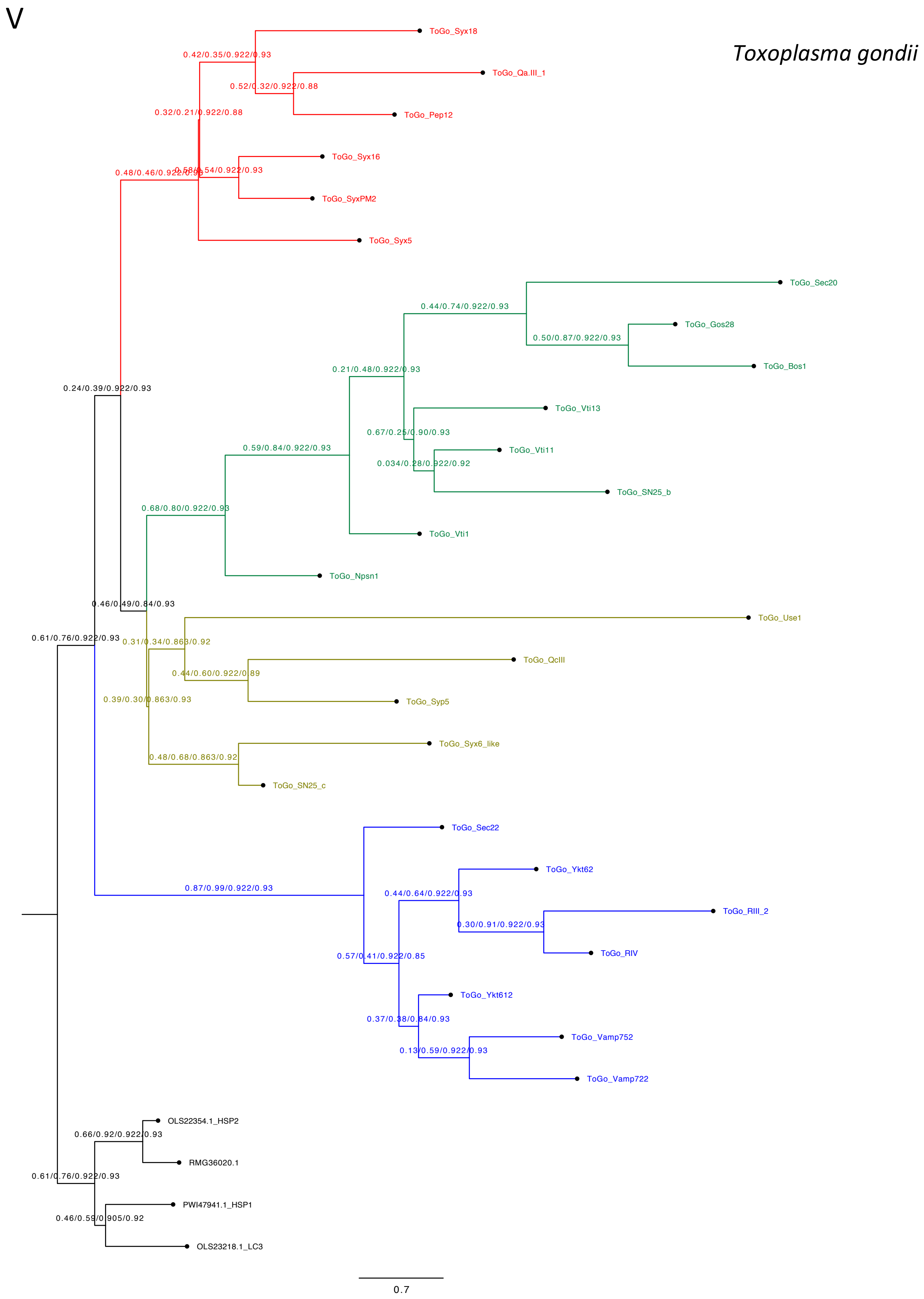
V
Toxoplasma gondii
ToGo_Syx18
0.42/0.35/0.922/0.93
ToGo_Qa.III_1
0.52/0.32/0.922/0.88
ToGo_Pep12
0.32/0.21/0.922/0.88
ToGo_Syx16
0.48/0.46/0.922/0.93
0.58/0.54/0.922/0.93
ToGo_SyxPM2
ToGo_Syx5
ToGo_Sec20
0.44/0.74/0.922/0.93
ToGo_Gos28
0.50/0.87/0.922/0.93
ToGo_Bos1
0.21/0.48/0.922/0.93
0.24/0.39/0.922/0.93
ToGo_Vti13
0.67/0.25/0.90/0.93
ToGo_Vti11
0.59/0.84/0.922/0.93
0.034/0.28/0.922/0.92
ToGo_SN25_b
0.68/0.80/0.922/0.93
ToGo_Vti1
ToGo_Npsn1
0.46/0.49/0.84/0.93
ToGo_Use1
0.61/0.76/0.922/0.93
0.31/0.34/0.863/0.92
ToGo_QcIII
0.44/0.60/0.922/0.89
ToGo_Syp5
0.39/0.30/0.863/0.93
ToGo_Syx6_like
0.48/0.68/0.863/0.92
ToGo_SN25_c
ToGo_Sec22
ToGo_Ykt62
0.87/0.99/0.922/0.93
0.44/0.64/0.922/0.93
ToGo_RIII_2
0.30/0.91/0.922/0.93
ToGo_RIV
0.57/0.41/0.922/0.85
ToGo_Ykt612
0.37/0.38/0.84/0.93
ToGo_Vamp752
0.13/0.59/0.922/0.93
ToGo_Vamp722
OLS22354.1_HSP2
0.66/0.92/0.922/0.93
RMG36020.1
0.61/0.76/0.922/0.93
PWI47941.1_HSP1
0.46/0.59/0.905/0.92
OLS23218.1_LC3
0.7

W

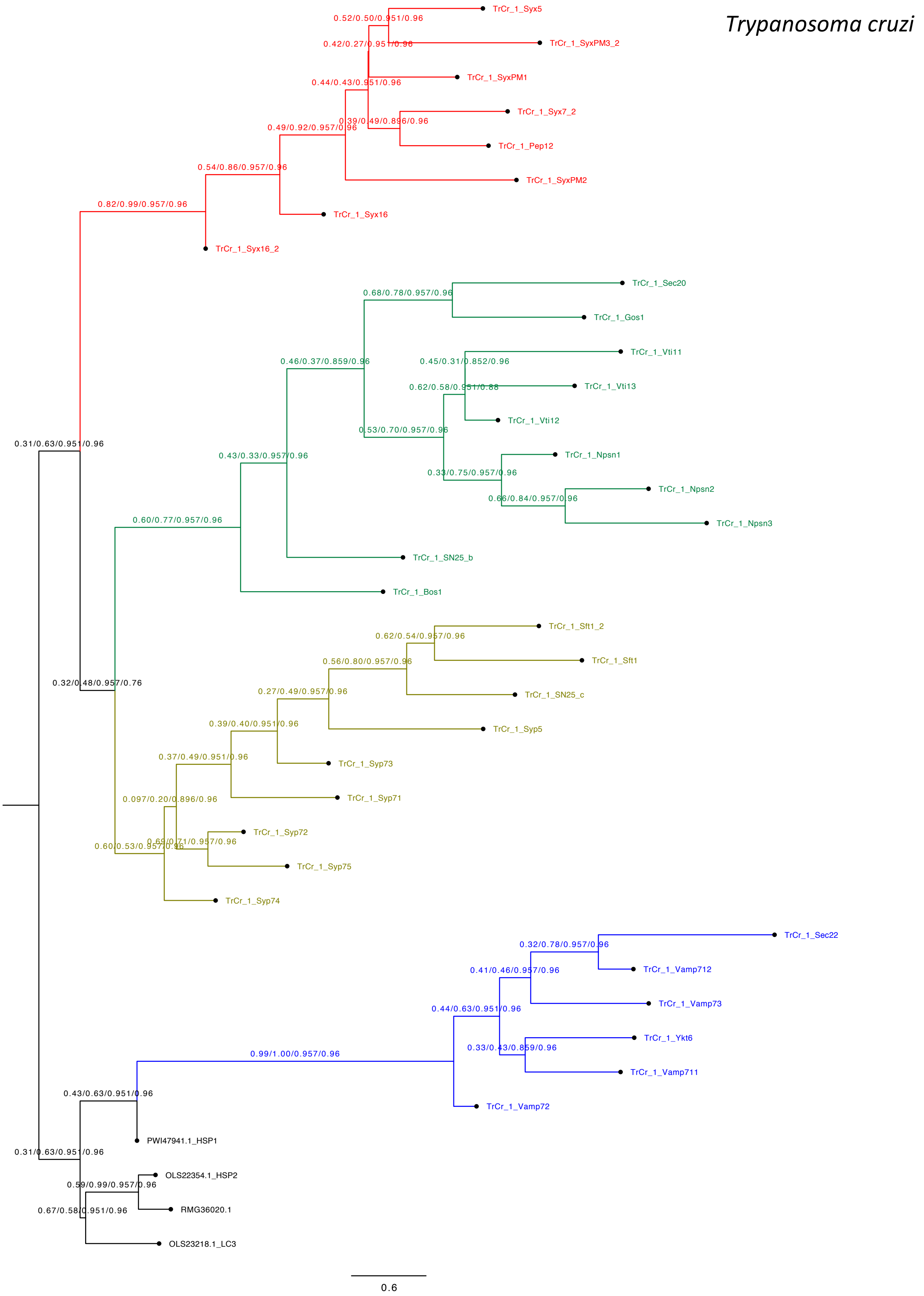
Trypanosoma cruzi
TrCr_1_Syx5
TrCr_1_SyxPM3_2
TrCr_1_SyxPM1
TrCr_1_Syx7_2
TrCr_1_Pep12
TrCr_1_SyxPM2
TrCr_1_Syx16
TrCr_1_Syx16_2
TrCr_1_Sec20
TrCr_1_Gos1
TrCr_1_Vti11
TrCr_1_Vti13
TrCr_1_Vti12
TrCr_1_Npsn1
TrCr_1_Npsn2
TrCr_1_Npsn3
TrCr_1_SN25_b
TrCr_1_Bos1
TrCr_1_Sft1_2
TrCr_1_Sft1
TrCr_1_SN25_c
TrCr_1_Syp5
TrCr_1_Syp73
TrCr_1_Syp71
TrCr_1_Syp72
TrCr_1_Syp75
TrCr_1_Syp74
TrCr_1_Sec22
TrCr_1_Vamp712
TrCr_1_Vamp73
TrCr_1_Ykt6
TrCr_1_Vamp711
TrCr_1_Vamp72
PWI47941.1_HSP1
OLS22354.1_HSP2
RMG36020.1
OLS23218.1_LC3
0.6

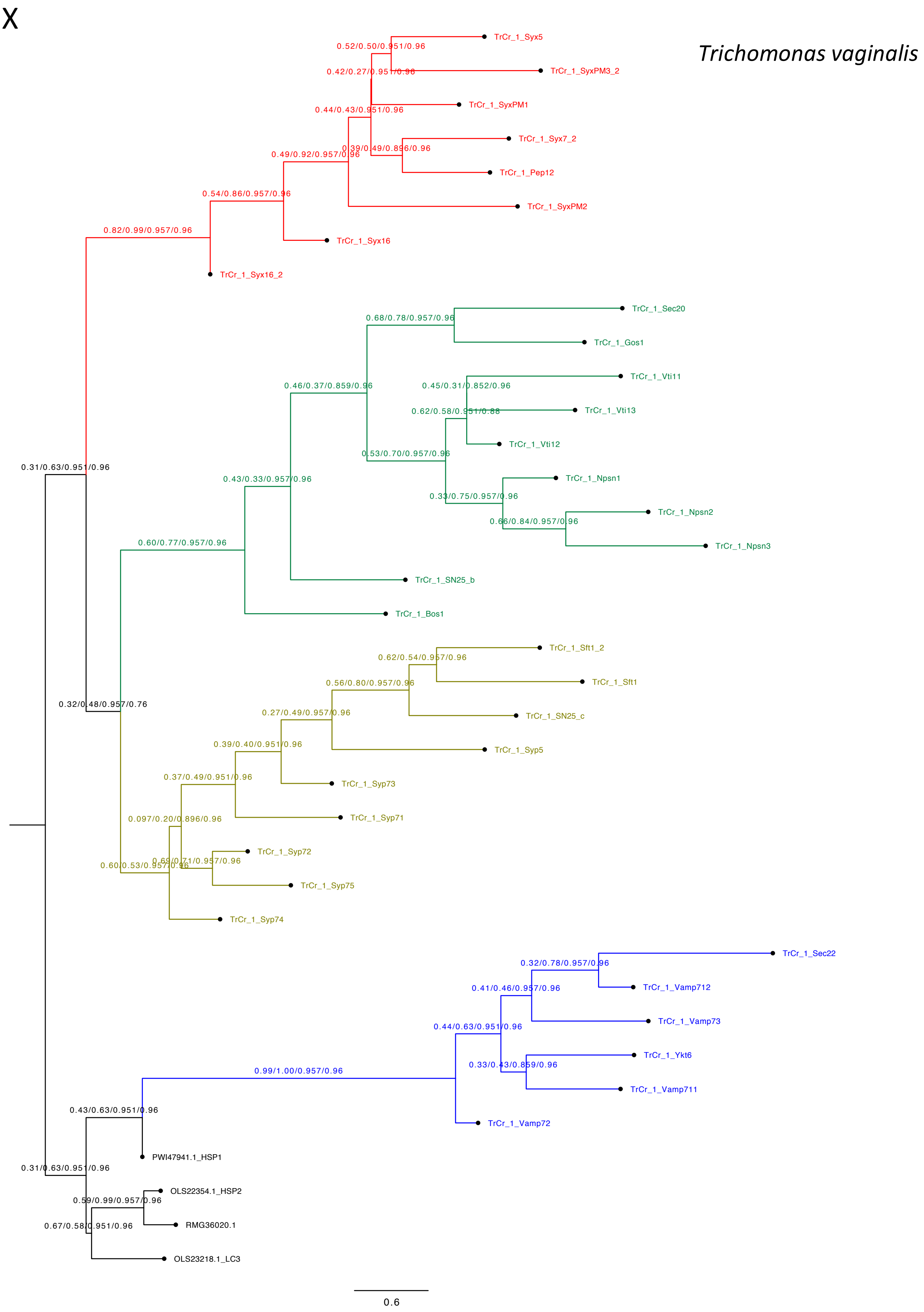
X
Trichomonas vaginalis
TrCr_1_Syx5
TrCr_1_SyxPM3_2
TrCr_1_SyxPM1
TrCr_1_Syx7_2
TrCr_1_Pep12
TrCr_1_SyxPM2
TrCr_1_Syx16
TrCr_1_Syx16_2
TrCr_1_Sec20
TrCr_1_Gos1
TrCr_1_Vti11
TrCr_1_Vti13
TrCr_1_Vti12
TrCr_1_Npsn1
TrCr_1_Npsn2
TrCr_1_Npsn3
TrCr_1_SN25_b
TrCr_1_Bos1
TrCr_1_Sft1_2
TrCr_1_Sft1
TrCr_1_SN25_c
TrCr_1_Syp5
TrCr_1_Syp73
TrCr_1_Syp71
TrCr_1_Syp72
TrCr_1_Syp75
TrCr_1_Syp74
TrCr_1_Sec22
TrCr_1_Vamp712
TrCr_1_Vamp73
TrCr_1_Ykt6
TrCr_1_Vamp711
TrCr_1_Vamp72
PWI47941.1_HSP1
OLS22354.1_HSP2
RMG36020.1
OLS23218.1_LC3
0.52/0.50/0.951/0.96
0.42/0.27/0.951/0.96
0.44/0.43/0.951/0.96
0.39/0.49/0.896/0.96
0.49/0.92/0.957/0.96
0.54/0.86/0.957/0.96
0.82/0.99/0.957/0.96
0.68/0.78/0.957/0.96
0.46/0.37/0.859/0.96
0.45/0.31/0.852/0.96
0.62/0.58/0.951/0.88
0.53/0.70/0.957/0.96
0.43/0.33/0.957/0.96
0.33/0.75/0.957/0.96
0.66/0.84/0.957/0.96
0.60/0.77/0.957/0.96
0.31/0.63/0.951/0.96
0.62/0.54/0.957/0.96
0.56/0.80/0.957/0.96
0.27/0.49/0.957/0.96
0.32/0.48/0.957/0.76
0.39/0.40/0.951/0.96
0.37/0.49/0.951/0.96
0.097/0.20/0.896/0.96
0.69/0.71/0.957/0.96
0.60/0.53/0.957/0.96
0.32/0.78/0.957/0.96
0.41/0.46/0.957/0.96
0.44/0.63/0.951/0.96
0.33/0.43/0.859/0.96
0.99/1.00/0.957/0.96
0.43/0.63/0.951/0.96
0.31/0.63/0.951/0.96
0.59/0.99/0.957/0.96
0.67/0.58/0.951/0.96
0.6

**Fig. S7. Maximum likelihood trees for the complete SNARE set of each of the 24 representative species used for the overview tree shown in Fig. 2 and the four archaeal SNARE-like sequences from the Heimdallarchaeota clade.**

The different eukaryotic species were selected to cover the diversity of the entire eukaryotic domain. Although the SNARE repertoire of each eukaryotic species usually includes all conserved SNARE types, different sets of SNARE genes were multiplied in different species. Thus different species have different numbers of SNARE genes. The SNARE repertoire of the selected 24 species varies from 20 to 62 individual SNARE motifs. Note that Qbc-SNAREs have two SNARE motifs, which are indicated by a B or C for the Qb- or the Qc-helix, respectively. Statistical branch support values (likelihood-mapping, IQ-TREE support, RAxML support, PhyML support) are given. Qa-SNAREs are in red, Qb-SNAREs in khaki, Qc-SNAREs in moss green, and R-SNAREs in blue.

#### Section 8: List of Cluster containing sequences with a SNARE-like region

Supplementary Table 1: List of Clusters Containing Prokaryotic Sequences With a SNARE-Like Region

| | sequence ID (NCBI) | HMM e-value | HMM SNARE type | Motif detected by HMMER 3.2.1 | Species name | Taxonomy | annotation | Cluster ID |
|---|---|---|---|---|---|---|---|---|
| ▼ 0 | | | | | | | | |
| | SNV31155.1 | 1E-21 | R.III | VEQVKQELGEVKKIMIDNIDKVLERGERIEDLIKRTEGLADTSFLFHKKAK-- | *Legionella spiritensis* | Bacteria; Proteobacteria; Gammaproteobacteria; Legionellales; Legionellaceae; Legionella | Synaptobrevin | 0 |
| | OEH47953.1 | 1.7E-20 | R.III | -VQVKKELEDVKQIMLENVEKVTLRGEKIDDLLEKTEQLEASSFQFKKNAEKL | *Legionella parisiensis* | Bacteria; Proteobacteria; Gammaproteobacteria; Legionellales; Legionellaceae; Legionella | hypothetical protein lpari_01141 | 0 |
|  | WP_058501541.1 | 3.6E-20 | R.III | LRQIKEELEETKKIMTDNIDKLLERGERIEQLIEKTEELSRNAKQFKHE---- | *Legionella israelensis* | Bacteria; Proteobacteria; Gammaproteobacteria; Legionellales; Legionellaceae; Legionella | hypothetical protein | 0 |
| | WP_058442445.1 | 8.8E-18 | R.III | VESIKQELAETQKIMLSNLDKVIERGERIEEVLAKSENLATSSFHFKQKA--- | *Legionella brunensis* | Bacteria; Proteobacteria; Gammaproteobacteria; Legionellales; Legionellaceae; Legionella | hypothetical protein | 0 |
| | KTD10104.1 | 1.7E-17 | R.III | VRKVQKEVDETMQIMLKNIEKMEKRGEALQTLVTKTQDLETSSFQFKKKAA-- | *Legionella hackeliae* | Bacteria; Proteobacteria; Gammaproteobacteria; Legionellales; Legionellaceae; Legionella | Synaptobrevin | 0 |
|  | OGV52230.1 | 6E-16 | R.III | --ALKADVEEVLNTMKHNMEKTLQRDIKIDDLVDKADKLEKHSIQFQRT---- | *Legionellales bacterium RIFCSPHIGHO2_12_FULL_42_9* | Bacteria; Proteobacteria; Gammaproteobacteria; Legionellales | hypothetical protein A3F46_02505 | 0 |
| | WP_058533815.1 | 1.4E-15 | R.III | -SDIKETLDETKKLMQNNIEQMLQRQEKIEDLAKRSEALAQGALSFKHKS--- | *Legionella saoudiensis* | Bacteria; Proteobacteria; Gammaproteobacteria; Legionellales; Legionellaceae; Legionella | hypothetical protein | 0 |
| | WP_058532341.1 | 8.7E-15 | R.II | IQKVQKELHETHAVLRETLEKMELRGEKLSDLVAKTEGLSQASFAFKEES--- | *Legionella rubrilucens* | Bacteria; Proteobacteria; Gammaproteobacteria; Legionellales; Legionellaceae; Legionella | hypothetical protein | 0 |
| | SNV46269.1 | 6.3E-14 | R.III | IATIKRDLAETLQIMLENLEKTKLRGEKLEHLVATTQDLEASSFRFKKSAED- | *Legionella lansingensis* | Bacteria; Proteobacteria; Gammaproteobacteria; Legionellales; Legionellaceae; Legionella | Synaptobrevin | 0 |
| | WP_081778127.1 | 6.4E-14 | R.III | IATIKRDLAETLQIMLENLEKTKLRGEKLEHLVATTQDLEASSFRFKKSAED- | *Legionella lansingensis* | Bacteria; Proteobacteria; Gammaproteobacteria; Legionellales; Legionellaceae; Legionella | hypothetical protein | 0 |
|  | WP_058526714.1 | 1.3E-13 | R.III | ----QQEIDETRDAALKALDSLDMRGEKISDLLQKTQSLSEASFAFKEKSE-- | *Legionella erythra* | Bacteria; Proteobacteria; Gammaproteobacteria; Legionellales; Legionellaceae; Legionella | hypothetical protein | 0 |
| | ASQ47094.1 | 3.1E-13 | R.I | ITQIKSELHETTEIMKQNIEKLHLREEGLKKLVEKTEELETLTFHFNKKAKAL | *Legionella clemsonensis* | Bacteria; Proteobacteria; Gammaproteobacteria; Legionellales; Legionellaceae; Legionella | Synaptobrevin | 0 |
| | WP_094091873.1 | 3.4E-13 | R.I | ITQIKSELHETTEIMKQNIEKLHLREEGLKKLVEKTEELETLTFHFNKKAKAL | *Legionella clemsonensis* | Bacteria; Proteobacteria; Gammaproteobacteria; Legionellales; Legionellaceae; Legionella | hypothetical protein | 0 |
| | WP_108294219.1 | 6.2E-13 | R.III | IKKVQKELDETHAVLLEAFQKMELRGEKLSELVDKTEGLSQLSFAFKEESERL | *Legionella taurinensis* | Bacteria; Proteobacteria; Gammaproteobacteria; Legionellales; Legionellaceae; Legionella | hypothetical protein | 0 |
| | SFL49537.1 | 2.9E-10 | R.III | --KAQTAISETMQIMQENIGKLHTREEALQKLMKQTEELETLSFQFNKKAK-- | *Legionella jamestowniensis DSM 19215* | Bacteria; Proteobacteria; Gammaproteobacteria; Legionellales; Legionellaceae; Legionella | Synaptobrevin | 0 |
|  | WP_028387504.1 | 6.4E-10 | R.III | -KATRDALDDTTEIMRDNLEKFLERREALERLLEDTEELKIETNRF------- | *Legionella geestiana* | Bacteria; Proteobacteria; Gammaproteobacteria; Legionellales; Legionellaceae; Legionella | hypothetical protein | 0 |
| ▼ 1 | | | | | | | | |
| | OGT45220.1 | 1.3E-20 | R.III | -AQLKQKVQEIKEIMLENLEKLLDRGERIELLVRKTDYLANGATSFKREAKKL | *Gammaproteobacteria bacterium RIFCSPHIGHO2_12_FULL_41_20* | Bacteria; Proteobacteria; Gammaproteobacteria | hypothetical protein A3E83_08095 | 1 |
| | OGT53257.1 | 7.4E-20 | R.III | ITRLKEQAAEVKTIMLDNIEKVLQRGEQLEVLLDKTDRLRLDAITFKGNAKKL | *Gammaproteobacteria bacterium RIFCSPHIGHO2_12_FULL_42_13* | Bacteria; Proteobacteria; Gammaproteobacteria | hypothetical protein A3E84_01380 | 1 |
| | OGT60900.1 | 2.2E-14 | R.IV | ---MQVDIQDTKQIMQTNIQKLTERGDKLAELEKTTFELSKNANKFSEGAKEL | *Gammaproteobacteria bacterium RIFCSPHIGHO2_12_FULL_43_28* | Bacteria; Proteobacteria; Gammaproteobacteria | hypothetical protein A3F14_02900 | 1 |
| ▼ 2 | | | | | | | | |
| | SNV19632.1 | 1E-19 | Qc.III.c | LDILLKTVTRTHEIAEAIHKELTSQNKLIDGLNEDVEKTDGKVENTTKRV--- | *Legionella pneumophila* | Bacteria; Proteobacteria; Gammaproteobacteria; Legionellales; Legionellaceae; Legionella | SNARE domain | 2 |
| | CZR11020.1 | 4.8E-19 | Qc.III.c | LDILLKTVTRTHEIAEAIQEELTSQNKLIDGLNEHVEKTDGKVKNTTKKV--- | *Legionella pneumophila* | Bacteria; Proteobacteria; Gammaproteobacteria; Legionellales; Legionellaceae; Legionella | SNARE domain | 2 |
| | SQG86500.1 | 4.8E-19 | Qc.III.c | LDILLKTVTRTHEIAEAIQEELTSQNKLIDGLNEHVEKTDGKVKNTTKKV--- | *Legionella pneumophila* | Bacteria; Proteobacteria; Gammaproteobacteria; Legionellales; Legionellaceae; Legionella | SNARE domain | 2 |
| | WP_027219386.1 | 4.8E-19 | Qc.III.c | LDILLKTVTRTHEIAEAIQEELTSQNKLIDGLNEHVEKTDGKVKNTTKKV--- | *Legionella pneumophila* | Bacteria; Proteobacteria; Gammaproteobacteria; Legionellales; Legionellaceae; Legionella | hypothetical protein | 2 |
| | WP_061637776.1 | 4.8E-19 | Qc.III.c | LDILLKTVTRTHEIAEAIQEELTSQNKLIDGLNEHVEKTDGKVKNTTKKV--- | *Legionella pneumophila* | Bacteria; Proteobacteria; Gammaproteobacteria; Legionellales; Legionellaceae; Legionella | hypothetical protein | 2 |
| | PQM70780.1 | 1.2E-18 | Qc.III.c | LDILLKTVTRTHEIAEAIQEELTSQNKLIDGLNEHVEKTDGKVGNTTKRVET- | *Legionella pneumophila* | Bacteria; Proteobacteria; Gammaproteobacteria; Legionellales; Legionellaceae; Legionella | hypothetical protein C3926_12725 | 2 |
| | PYB62981.1 | 1.2E-18 | Qc.III.c | LDILLKTVTRTHEIAEAIQEELTSQNKLIDGLNEHVEKTDGKVGNTTKRVET- | *Legionella pneumophila* | Bacteria; Proteobacteria; Gammaproteobacteria; Legionellales; Legionellaceae; Legionella | hypothetical protein DM455_08335 | 2 |
| | WP_061484634.1 | 2.2E-18 | Qc.III.c | LDILLKTVTRTHEIAEAIQEELTSQNKLIDGLNEHVEKTDGKVENTTKKV--- | *Legionella pneumophila* | Bacteria; Proteobacteria; Gammaproteobacteria; Legionellales; Legionellaceae; Legionella | hypothetical protein | 2 |
| | ADG25747.1 | 2.9E-17 | Qc.III.c | LDILLKTVTRTYEIAEAIQEELTSQNKLIDGLNEHVEKTDGKVENTTKKV--- | *Legionella pneumophila 2300/99 Alcoy* | Bacteria; Proteobacteria; Gammaproteobacteria; Legionellales; Legionellaceae; Legionella | hypothetical protein lpa_03441 | 2 |
| ▼ -1 | | | | | | | | |
| | OGT42784.1 | 1.9E-18 | R.III | --KIEAELKEVKNIALNNINKILERGEKIESLVSQTSDLANQTVEFKKKS--- | *Gammaproteobacteria bacterium RIFCSPHIGHO2_12_FULL_37_34* | Bacteria; Proteobacteria; Gammaproteobacteria | hypothetical protein A3F42_02955 | -1 |
| | WP_058512658.1 | 1.2E-17 | Qa.IV | IREIERKIIDINQLFVDVAELVKDQGSLVNNIELSIEGAKEETAKGVIELEKA | *Legionella santicrucis* | Bacteria; Proteobacteria; Gammaproteobacteria; Legionellales; Legionellaceae; Legionella | hypothetical protein | -1 |
| | WP_062266217.1 | 1E-15 | Qa.III | LRKLEHEIRDLNEIFKSMSQVVYDQGQQFETLESLIDSAAFSVQEGVKTL--- | *Endozoicomonas arenosclerae* | Bacteria; Proteobacteria; Gammaproteobacteria; Oceanospirillales; Hahellaceae; Endozoicomonas | hypothetical protein | -1 |
| | OUW42374.1 | 1.1E-13 | Qb.III.b | LTDAQRAAADANETGDAILGTLREQRDALIRSRAGMEGVKEDMKHNERLVNNM | *bacterium TMED181* | Bacteria | hypothetical protein CBD41_09135 | -1 |
| | OGT45367.1 | 1.3E-12 | SNAP.c | LEQITNGLQLLQNQSRVLGAELDRQNAKLGGLEQDVDTANSRLRQDNRRVRNL | *Gammaproteobacteria bacterium RIFCSPHIGHO2_12_FULL_41_20* | Bacteria; Proteobacteria; Gammaproteobacteria | hypothetical protein A3E83_00025 | -1 |
| | OGT45367.1 | 1E-11 | SNAP.b | --EMLMSVANTNAQCAETLAELDRQGEQLARAHDKVRGVHEHLSMAEQHLHRM | *Gammaproteobacteria bacterium RIFCSPHIGHO2_12_FULL_41_20* | Bacteria; Proteobacteria; Gammaproteobacteria | hypothetical protein A3E83_00025 | -1 |
| | WP_057625376.1 | 3.8E-10 | Qa.IV | ---VENGVTQLKSLSLDINELINNQNEMVEHIATNTEIGKENMNQGNENIEDA | *Candidatus Berkiella cookevillensis* | Bacteria; Proteobacteria; Gammaproteobacteria; Legionellales; Coxiellaceae; Candidatus Berkiella | hypothetical protein | -1 |
|  | WP_089081872.1 | 2.1E-07 | Qc.III | ---------VTNNVSDKLNKALEDQIELLKELNNNLKTENFRLNNLVIEL--- | *Flavobacterium branchiophilum* | Bacteria; Bacteroidetes; Flavobacteriia; Flavobacteriales; Flavobacteriaceae; Flavobacterium | XRE family transcriptional regulator | -1 |
| | OGY39434.1 | 2.3E-07 | Qc.III | LELIADDVSTIKENVSHLTDRMDHMTERMDHMTERMDHMTERMDHMTEKVD-- | *Candidatus Andersenbacteria bacterium RIFCSPLOWO2_12_FULL_45_8* | Bacteria; Candidatus Andersenbacteria | hypothetical protein A3G57_00530 | -1 |
| | WP_078921088.1 | 2.5E-07 | Qc.III | IDRLTTQIDHQQQQSDNLHDEVNQMEKKISDIDKTVYSNEQKIEALLNRLQ-- | *Thiothrix eikelboomii* | Bacteria; Proteobacteria; Gammaproteobacteria; Thiotrichales; Thiotrichaceae; Thiothrix | hypothetical protein | -1 |
|  | WP_056089750.1 | 3.6E-07 | Qb.II | IQHNSQQTDRYVQLSNQVTNALKTQNSLIRKINTNNNAVPN------------ | *Pedobacter sp. Leaf176* | Bacteria; Bacteroidetes; Sphingobacteriia; Sphingobacteriales; Sphingobacteriaceae; Pedobacter | serine protease | -1 |
|  | PTC01180.1 | 5.5E-07 | Qa.III | -------------LYEEVSDELDEQDDKVERLEEAVTTAKENLVNTRKALEEA | *Thalassospira xiamenensis* | Bacteria; Proteobacteria; Alphaproteobacteria; Rhodospirillales; Rhodospirillaceae; Thalassospira | hypothetical protein C9975_03495 | -1 |
|  | OYT30919.1 | 7.6E-07 | Qb.III | --QTASVVNQAEQQANQVDQELNQDIEELDKTIAEIDQIDSDLDTTD------ | *Candidatus Woesearchaeota archaeon ex4484_78* | Archaea; Candidatus Woesearchaeota | hypothetical protein B6U93_04480 partial | -1 |
|  | WP_065592534.1 | 1E-06 | Qa.IV | --KLASSTKQLGDLFRKISLQLEQDKHAIMEIAENAKSASTN----------- | *Aliivibrio* | Bacteria; Proteobacteria; Gammaproteobacteria; Vibrionales; Vibrionaceae | MULTISPECIES: hypothetical protein | -1 |
| | WP_107348248.1 | 1.3E-06 | Qa.III | --RISEAIQQINDMYLQIASAAEEQSLVAEEINTNTLNIKELSVQVADAAV-- | *Photobacterium lutimaris* | Bacteria; Proteobacteria; Gammaproteobacteria; Vibrionales; Vibrionaceae; Photobacterium | hypothetical protein | -1 |
|  | WP_102074318.1 | 1.5E-06 | SNAP.b | --RSTSMANQAAEQARQALSMLEGQTAQLDALRSSLDESREQ----------- | *Pusillimonas sp. JR1/69-3-13* | Bacteria; Proteobacteria; Betaproteobacteria; Burkholderiales; Alcaligenaceae; Pusillimonas | hypothetical protein | -1 |
| | WP_066064391.1 | 1.6E-06 | Qc.III | ---IDQKVSSLDQKVNSMDQELKIHGKMINHLIETVAATNVKISETNERLDRL | *Bacillus soli* | Bacteria; Firmicutes; Bacilli; Bacillales; Bacillaceae; Bacillus | hypothetical protein | -1 |
|  | PZX66526.1 | 1.7E-06 | Qc.II | -----GKVKQLRMKTGasaEIKEEIRKENKALNRLQKDLDSLRSKITSAIN----- | *Salegentibacter mishustinae* | Bacteria; Bacteroidetes; Flavobacteriia; Flavobacteriales; Flavobacteriaceae; Salegentibacter | phage integrase family protein | -1 |
|  | WP_006565661.1 | 1.8E-06 | SNAP.b | ------------ERFNAVEEHLGEQDERFNTIEKHLGEVDEKFHIVDRRMDTM | *Anaerostipes caccae* | Bacteria; Firmicutes; Clostridia; Clostridiales; Lachnospiraceae; Anaerostipes | hypothetical protein | -1 |
|  | WP_065592426.1 | 2E-06 | R.III | -----ENIVKVNDLAKDTVDDVQDNACIVEELNTMALYLRKSLRIFK------ | *Aliivibrio* | Bacteria; Proteobacteria; Gammaproteobacteria; Vibrionales; Vibrionaceae | MULTISPECIES: hypothetical protein | -1 |
|  | OKZ80410.1 | 2.2E-06 | Qa.IV | ---IENTIDSLQETLSDLNSLLEKHEDEISEIADKVKNADKELSD-------- | *Clostridium sp. CAG:12237_41* | Bacteria; Firmicutes; Clostridia; Clostridiales; Clostridiaceae; Clostridium | hypothetical protein BHW08_05950 | -1 |
|  | WP_057507700.1 | 2.3E-06 | Qc.III.b | ---LDTTTGTLVQRTNSINKEIKALEKQLDDLDARMEKVSDRYTKQ------- | *Stenotrophomonas chelatiphaga* | Bacteria; Proteobacteria; Gammaproteobacteria; Xanthomonadales; Xanthomonadaceae; Stenotrophomonas | flagellar protein | -1 |
| | PZP38966.1 | 2.4E-06 | Qc.III.b | --ELFGSVNNLHNLINGINGTLTNHGNRLNVLDNQVNNINSRINTINNQISTL | *Pseudomonas fluorescens* | Bacteria; Proteobacteria; Gammaproteobacteria; Pseudomonadales; Pseudomonadaceae; Pseudomonas | hypothetical protein DI585_05715 | -1 |
|  | WP_053747023.1 | 2.5E-06 | Qa.III | ---------KLNELITQVNAILKQQGELFGEIEDDLKETLLKMFKTQDE---- | *Streptomyces sp. NRRL WC-3618* | Bacteria; Actinobacteria; Streptomycetales; Streptomycetaceae; Streptomyces | type VII secretion system-associated protein | -1 |
|  | AEV95415.1 | 2.6E-06 | Qc.III | --QLSQSTKNLRQNSNRLQNELKKTQKTVDEISNSVDHFEFKL---------- | *Pediococcus claussenii ATCC BAA-344* | Bacteria; Firmicutes; Bacilli; Lactobacillales; Lactobacillaceae; Pediococcus | hypothetical protein PECL_1153 | -1 |
|  | WP_104471415.1 | 2.6E-06 | R.III | ----RTKTNEFREQLAKNIDpMVAKRAERLEAIQSQAETFRAVAQEF------- | *Acinetobacter indicus* | Bacteria; Proteobacteria; Gammaproteobacteria; Pseudomonadales; Moraxellaceae; Acinetobacter | DUF4102 domain-containing protein | -1 |
|  | WP_107440037.1 | 2.7E-06 | R.IV | -----AKVKQVKGKMKETVGKAvgnrsLKREGQAEMLGGKAQEMAEKA-------- | *Streptomyces sp. 142MFCol3.1* | Bacteria; Actinobacteria; Streptomycetales; Streptomycetaceae; Streptomyces | CsbD family protein | -1 |
|  | WP_104724683.1 | 2.8E-06 | Qb.I | ----------LVELSKKSLDELSHQNLSLKESLEIYKQGLAYLNEAQRLLEE- | *Helicobacter cholecystus* | Bacteria; Proteobacteria; Epsilonproteobacteria; Campylobacterales; Helicobacteraceae; Helicobacter | exodeoxyribonuclease VII small subunit | -1 |
|  | RDD62576.1 | 3.8E-06 | Qa.IV | -QRLETQVMQARELSDDMNFLVEKATSLADRLEGQIGQART------------ | *Rhodospirillaceae bacterium* | Bacteria; Proteobacteria; Alphaproteobacteria; Rhodospirillales; Rhodospirillaceae | hypothetical protein DRB17_08015 | -1 |

| | | | | | | | | |
|---|---|---|---|---|---|---|---|---|
|  | **WP_025790170.1** | 3.8E-06 | SNAP.c | -----AGAKRLAEIAEEMGQEVGEQAKRMHDLLTDIPTVNIDVVS-------- | *Prevotella loescheii* | Bacteria; Bacteroidetes; Bacteroidia; Bacteroidales; Prevotellaceae; Prevotella | hypothetical protein | -1 |
|  | **OQY41163.1** | 4.1E-06 | Qc.III | IDKIISEVKRTEHTADKIIKEANDQVEIiIDNAHEKAKKLIERIKE-------- | *Candidatus Atribacteria bacterium 4572_76* | Bacteria; Candidatus Atribacteria | ATP synthase archaeal subunit H | -1 |
|  | **PKO29673.1** | 4.4E-06 | Qb.III.b | -----DQANTTEELMINLGEQLRSQRLRM---NLTLEEVAQSAGVSINAVRR- | *Betaproteobacteria bacterium HGW-Betaproteobacteria-9* | Bacteria; Proteobacteria; Betaproteobacteria | XRE family transcriptional regulator | -1 |
|  | **WP_097079058.1** | 4.6E-06 | Qb.III | -------ENQTIDLSDGSQIQLNTQSKVVVKMRLHSREIQLQQGEATFHV--- | *Acinetobacter puyangensis* | Bacteria; Proteobacteria; Gammaproteobacteria; Pseudomonadales; Moraxellaceae; Acinetobacter | DUF4974 domain-containing protein | -1 |
|  | **OFV68491.1** | 5.4E-06 | Qa.III.b | -------INDVKQLMKEIADIAQKHSLVIQIVDGDLVAGKEHVRSAV------ | *Candidatus Syntrophoarchaeum caldarius* | Archaea; Euryarchaeota; Methanomicrobia; Methanosarcinales; ANME-2 cluster; Candidatus Syntrophoarchaeum | Metal-dependent phosphohydrolase | -1 |
|  | **WP_084428681.1** | 6.1E-06 | Qb.I | -----QNLNDTVKQLAAAADTLSGSREDLFATVDSLGKFTEALASSDKAV--- | *Kibdelosporangium aridum* | Bacteria; Actinobacteria; Pseudonocardiales; Pseudonocardiaceae; Kibdelosporangium | MCE family protein | -1 |
|  | **PHV62861.1** | 7.1E-06 | Qc.I | --EITELSEEMKGGVKSASGAVQKVDRTLINAKKQVESAQTKSK--------- | *Cyanobacterium aponinum IPPAS B-1201* | Bacteria; Cyanobacteria; Oscillatoriophycideae; Chroococcales; Cyanobacteriaceae; Cyanobacterium | hypothetical protein CSQ80_08270 | -1 |
| | **GBD26362.1** | 8E-06 | Qc.III | -ERLAASVTELASHVDQLGSRVDQLGARVDQIGSHVDRVGSLMDRLAARVDE- | *bacterium HR30* | Bacteria | Chromosome partition protein Smc | -1 |
|  | **OHA26315.1** | 8.1E-06 | Qc.III | LQNLMETLGTIAREGYTIRQRIEHHEKMASLLKEKLELQGGSLD--------- | *Candidatus Taylorbacteria bacterium RIFCSPHIGHO2_02_FULL_46_13* | Bacteria; Candidatus Taylorbacteria | hypothetical protein A3C06_04725 | -1 |
|  | **WP_070779095.1** | 8.5E-06 | Qc.III.b | --ALAAQVSSLADQVAHIEATLEDHGQSLNAVQTRVDRLGEELH--------- | *Actinomyces sp. HMSC062G12* | Bacteria; Actinobacteria; Actinomycetales; Actinomycetaceae; Actinomyces | DUF2746 domain-containing protein | -1 |
|  | **RDB36013.1** | 8.6E-06 | Qb.III.d | -----AIADDIKKISSSLAQMLSSQETggSLKRIIANIDGITSDLK--------- | *Spirobacillus cienkowskii* | Bacteria; Proteobacteria; Oligoflexia; Silvanigrellales; Spirobacillus | MCE family protein | -1 |
|  | **OYT74226.1** | 8.8E-06 | Qb.I | -RRIRQQLQQQIEAVRAQIAALQQQQEQLRRQNDEFAEQGAALAS-------- | *Armatimonadetes bacterium JP3_11* | Bacteria; Armatimonadetes | hypothetical protein CFK49_09500 | -1 |
|  | **WP_112052017.1** | 8.9E-06 | R.IV | ----QEEMKKTLTTMTESLNKMTESNEKLAEFQEKmSETLVMLS---------- | *Helicobacter sp. 15-1451* | Bacteria; Proteobacteria; Epsilonproteobacteria; Campylobacterales; Helicobacteraceae; Helicobacter | hypothetical protein | -1 |
| | **WP_068606002.1** | 9.3E-06 | Qc.III | --ELSEGAKRILHTAGDISNDVKGKIRKVEPIVDTVQDVGEMLHNVTSTVKE- | *Paenibacillus swuensis* | Bacteria; Firmicutes; Bacilli; Bacillales; Paenibacillaceae; Paenibacillus | DUF948 domain-containing protein | -1 |
|  | **OGW37336.1** | 1E-05 | Qc.III | ---------VIKDKSEKIGEEADKLKEKTGDIKDVTKKVTEGIKEQGEKVKE- | *Nitrospirae bacterium RBG_13_39_12* | Bacteria; Nitrospirae | hypothetical protein A2Y97_05345 | -1 |
|  | **PKP52188.1** | 1.1E-05 | Qc.III | ----GDKVDAVGDKVDAVGDEVKAVGDKVDALGDKVDAVGDEVKAVGSKVDNL | *Candidatus Altiarchaeales archaeon HGW-Altiarchaeales-3* | Archaea; Euryarchaeota; Candidatus Altiarchaeales | hypothetical protein CVT90_02735 | -1 |
|  | **KKT45151.1** | 1.2E-05 | Qc.III.b | LDAIEKTLGPLVHVSSAMQRALREHERRLSLVEHKVGLTNK------------ | *Parcubacteria group bacterium GW2011_GWA2_44_15* | Bacteria; unclassified Parcubacteria group | hypothetical protein UW34_C0004G0014 | -1 |
|  | **RCK13034.1** | 1.2E-05 | R.IV | ---IKINSDAGVNNLNQIIEKLMKNGAEIRSLEEKAPNLVPHS---------- | *Bacillus licheniformis* | Bacteria; Firmicutes; Bacilli; Bacillales; Bacillaceae; Bacillus | ATP-binding cassette domain-containing protein | -1 |
|  | **WP_034560074.1** | 1.2E-05 | Qc.III | ------------EQAELMQEKINEMQAALDYLVHKVDNYEDHMRVAEKRLSK- | *Carnobacterium gallinarum* | Bacteria; Firmicutes; Bacilli; Lactobacillales; Carnobacteriaceae; Carnobacterium | MerR family transcriptional regulator | -1 |
|  | **WP_016281978.1** | 1.3E-05 | Qa.III | ----EAMIMQMNETMEILKSYISVQGLMILQIKDSMERLLEKVEKGQA----- | *Lachnospiraceae bacterium A4* | Bacteria; Firmicutes; Clostridia; Clostridiales; Lachnospiraceae | hypothetical protein | -1 |
|  | **WP_101354561.1** | 1.4E-05 | Qc.III | -----------RELAIQITTELKKLTEKLDNIHVLAVYGGQDVEQQMRKLK-- | *Bacillus camelliae* | Bacteria; Firmicutes; Bacilli; Bacillales; Bacillaceae; Bacillus | DEAD/DEAH box helicase | -1 |
|  | **WP_020531895.1** | 1.5E-05 | Qb.I | -------LAQQVQSGIENLETLKETKETISETANTLDDMRSAITKVNDVVKQI | *Flexithrix dorotheae* | Bacteria; Bacteroidetes; Cytophagia; Cytophagales; Flammeovirgaceae; Flexithrix | hypothetical protein | -1 |
| | **WP_111701224.1** | 1.5E-05 | SNAP.c | IDQMQRQVDELKEQARELTERVGRQKESVRILGESWQATVDEVDRAQQALD-- | *Microbispora sp. NEAU-HRDPA2-9* | Bacteria; Actinobacteria; Streptosporangiales; Streptosporangiaceae; Microbispora | hypothetical protein | -1 |
|  | **KIF36445.1** | 1.6E-05 | Qc.II | ----EERLSRLENIFVNIGETVVAQSDTIDVLVANINQLTENVNAVTNRVD-- | *Hassallia byssoidea VB512170* | Bacteria; Cyanobacteria; Nostocales; Tolypothrichaceae; Hassallia | hypothetical protein PI95_10830 | -1 |
|  | **KPK75711.1** | 1.6E-05 | Qc.III | --QIGELMGQLADMSRQINGQIDTRIKKLEILLRDADQAAAKLEKLTGK---- | *Phycisphaerae bacterium SM23_30* | Bacteria; Planctomycetes; Phycisphaerae | hypothetical protein AMJ79_10120 | -1 |
|  | **SCJ50933.1** | 1.6E-05 | Qa.III | -----SEVNELKSGFTELKSEVNELKSNVFRLESNVSVLNDNVFEIDQSL--- | *uncultured Clostridium sp.* | Bacteria; Firmicutes; Clostridia; Clostridiales; Clostridiaceae; Clostridium; environmental samples | Uncharacterized protein containing TOPRIM domain%2C potential nuclease | -1 |
|  | **WP_006915017.1** | 1.6E-05 | Qa.IV | ----EKTLESVRALTDNVNTQLESRDQQLTRLLNNAEATSANLEQ-------- | *Salinisphaera shabanensis* | Bacteria; Proteobacteria; Gammaproteobacteria; Salinisphaerales; Salinisphaeraceae; Salinisphaera | MCE family protein | -1 |
|  | **WP_057942896.1** | 1.6E-05 | Qc.I | ------LAHLLRDQASVLGDILRKSTQQVDRIEDRLLANRISTDRT------- | *Lysobacter gummosus* | Bacteria; Proteobacteria; Gammaproteobacteria; Xanthomonadales; Xanthomonadaceae; Lysobacter | magnesium transporter CorA | -1 |
|  | **WP_062187726.1** | 1.6E-05 | Qb.III.d | -------------ISVLAAITLISQAEALEKGTTEMEQVESKLaqlKKAENLLK-- | *Bacillales* | Bacteria; Firmicutes; Bacilli | MULTISPECIES: cell division protein DIVIC | -1 |
|  | **KUO91040.1** | 1.7E-05 | Qb.III.d | --------TTSDSLSSSLMQTLNKSDELINSIGKYVNEVQSQLDKVKDTLSS- | *Thermocladium sp. ECH_B* | Archaea; Crenarchaeota; Thermoprotei; Thermoproteales; Thermoproteaceae; Thermocladium | hypothetical protein AT710_07665 | -1 |
|  | **OEU75218.1** | 1.9E-05 | Qa.I | --DVDEKIVEISNGTEFFLTGLSEHEKKVSNIVDDANSTIKS----------- | *Desulfuromonadales bacterium C00003107* | Bacteria; Proteobacteria; Deltaproteobacteria; Desulfuromonadales | hypothetical protein BA869_04635 | -1 |
|  | **WP_009744881.1** | 2E-05 | Qb.I | ------RAQREMRKAEKTNKELASSDKKIQKLQKEISKVQEKIAKTG------ | *Capnocytophaga sp. oral taxon 338* | Bacteria; Bacteroidetes; Flavobacteriia; Flavobacteriales; Flavobacteriaceae; Capnocytophaga | hypothetical protein | -1 |
|  | **WP_057822268.1** | 2E-05 | Qc.III | -----------QEKGARLNDELTNTQDSLNETNHKMDANTKDSRQVVQRLE-- | *Lactobacillus perolens* | Bacteria; Firmicutes; Bacilli; Lactobacillales; Lactobacillaceae; Lactobacillus | hypothetical protein | -1 |
|  | **WP_099463716.1** | 2E-05 | Qc.III.c | ------------GQSYTLNAQMKVQTNLMDEVVVSVSRYEQKLSDVTVSMEL- | *Parabacteroides sp. Marseille-P3668* | Bacteria; Bacteroidetes; Bacteroidia; Bacteroidales; Tannerellaceae; Parabacteroides | TonB-dependent receptor | -1 |
|  | **WP_101635542.1** | 2E-05 | Qc.III | ----------INECSIRVHRKIDEKNKLIEELDKEIERVHKELKALKIKDD-- | *Bacillus sp. V5-8f* | Bacteria; Firmicutes; Bacilli; Bacillales; Bacillaceae; Bacillus | hypothetical protein | -1 |
|  | **WP_099435211.1** | 2.1E-05 | Qb.III | --TSEKIESLTEESGEKIMISLEKNNDSLEEMNNKLSTAQQL----------- | *Cyanobacterium aponinum* | Bacteria; Cyanobacteria; Oscillatoriophycideae; Chroococcales; Cyanobacteriaceae; Cyanobacterium | hypothetical protein | -1 |
| | **OGJ46455.1** | 2.2E-05 | Qc.III | -DETVEAVDNLKNEAVDIKEGVEEKIDQVNEAKEDVGEAVDAVNEAAGSLK-- | *Candidatus Peregrinibacteria bacterium RIFCSPLOWO2_02_FULL_48_14* | Bacteria; Candidatus Peregrinibacteria | hypothetical protein A3J23_00605 | -1 |
|  | **WP_006682114.1** | 2.2E-05 | SNAP.b | ----RQDAQNAEAALAVRRAELAAQGAQLKRARVEVEKARADLDYT------- | *Candidatus Glomeribacter gigasporarum* | Bacteria; Proteobacteria; Betaproteobacteria; Burkholderiales; Burkholderiaceae; Candidatus Glomeribacter | efflux RND transporter periplasmic adaptor subunit | -1 |
|  | **WP_023013633.1** | 2.2E-05 | Qb.I | --NLRALLNESIEKFDLVSkQELERQRTALNSANQRLNELQQQMKS-------- | *Acinetobacter sp. COS3* | Bacteria; Proteobacteria; Gammaproteobacteria; Pseudomonadales; Moraxellaceae; Acinetobacter | membrane fusogenic activity | -1 |
|  | **WP_095907158.1** | 2.2E-05 | Qc.II | ---------SLKSINQQLGETAKYEREQISGIKIKFEQLKANLRNAMEIMI-- | *Enterobacter cloacae* | Bacteria; Proteobacteria; Gammaproteobacteria; Enterobacterales; Enterobacteriaceae; Enterobacter; Enterobacter cloacae complex | hypothetical protein | -1 |
|  | **WP_053612689.1** | 2.3E-05 | Qc.I | --------RELRARVQQLARSVAADGETIRQIGRRADENAMGVARVADSL--- | *Streptomyces sp. XY533* | Bacteria; Actinobacteria; Streptomycetales; Streptomycetaceae; Streptomyces | hypothetical protein | -1 |
|  | **WP_091207672.1** | 2.3E-05 | Qa.IV | -----NTLKEVRSLTDHFNNQLIIQKEQIDILSHDIKSHELYLESAAH----- | *Flavobacterium xueshanense* | Bacteria; Bacteroidetes; Flavobacteriia; Flavobacteriales; Flavobacteriaceae; Flavobacterium | hypothetical protein | -1 |
|  | **WP_078672419.1** | 2.4E-05 | Qb.I | LKHTRDRMAVQKQKSAAAAQELETSRTNLQSLYQ--QAQADILT--------- | *Chitinophaga eiseniae* | Bacteria; Bacteroidetes; Chitinophagia; Chitinophagales; Chitinophagaceae; Chitinophaga | TolC family protein | -1 |
|  | **WP_020222899.1** | 2.6E-05 | Qc.III | ---------NLRDTVSSLNDRVDELNDRVDSLNDRIDRLQSELEEARSQ---- | *Halarchaeum acidiphilum* | Archaea; Euryarchaeota; Halobacteria; Halobacteriales; Halobacteriaceae | hypothetical protein | -1 |
|  | **OGK29302.1** | 2.7E-05 | Qc.III | IKDLDDSVHALTESSGMMGDELNYENKKIQSIKDDLEKLHENLR--------- | *Candidatus Roizmanbacteria bacterium RIFCSPHIGHO2_02_FULL_40_9* | Bacteria; Candidatus Roizmanbacteria | hypothetical protein A3D06_01585 partial | -1 |
|  | **WP_056247610.1** | 2.7E-05 | Qc.III | ----------ILQKANQIKDEINGHNDKLKKINTTINDAELVVKKVKKSIQ-- | *Flavobacterium sp. Leaf82* | Bacteria; Bacteroidetes; Flavobacteriia; Flavobacteriales; Flavobacteriaceae; Flavobacterium | hypothetical protein | -1 |
| | **WP_010097852.1** | 2.8E-05 | Qa.I | --ELSGKMSDLANKQDIMVEKVATQSDNHDNVIQHLENHEALMEKSNRQL--- | *Ornithinibacillus scapharcae* | Bacteria; Firmicutes; Bacilli; Bacillales; Bacillaceae; Ornithinibacillus | hypothetical protein | -1 |
|  | **OHB03277.1** | 2.9E-05 | Qa.III | ------SVASVYEATKGLFDEVNEQGQAIDF-EAKYQDIKTQVESGEIQL--- | *Candidatus Zambryskibacteria bacterium RIFCSPLOWO2_01_FULL_43_17* | Bacteria; Candidatus Zambryskibacteria | hypothetical protein A2920_00175 | -1 |
|  | **WP_006863774.1** | 2.9E-05 | Qc.III | ---------QLAEQGKKLKDKVTRQEQQISRLTDKVMEQSNTIDRLQEKVADL | *Marvinbryantia formatexigens* | Bacteria; Firmicutes; Clostridia; Clostridiales; Lachnospiraceae; Marvinbryantia | recombinase | -1 |
|  | **WP_031598337.1** | 2.9E-05 | Qb.I | --NARALLSSLLAKADLvTREEFDRQTQVLARTREKLNELEAKVA--------- | *Ferrovum* | Bacteria; Proteobacteria; Betaproteobacteria; Ferrovales; Ferrovaceae | MULTISPECIES: phosphoheptose isomerase | -1 |
|  | **WP_068325682.1** | 3E-05 | Qb.I | --ETKSLITRELDMSLAAAVELDASTQALLMTTQDHTEFHAAFN--------- | *Janibacter terrae* | Bacteria; Actinobacteria; Micrococcales; Intrasporangiaceae; Janibacter | enoyl-CoA hydratase family protein | -1 |
|  | **WP_096189578.1** | 3E-05 | SNAP.b | LRETREKLRETREKLRETREKLRVTREKLRETREKLRETREKLRRNEE----- | *Bacillus sp. FJAT-44876* | Bacteria; Firmicutes; Bacilli; Bacillales; Bacillaceae; Bacillus | hypothetical protein | -1 |
| | **WP_096187236.1** | 3.1E-05 | SNAP.b | LRESMEKLRQSAEKLRQSTEKLRQSKEKLRQSKEKLRESTEKLRQSKEKL--- | *Bacillus sp. FJAT-44876* | Bacteria; Firmicutes; Bacilli; Bacillales; Bacillaceae; Bacillus | hypothetical protein | -1 |
|  | **PKL78332.1** | 3.2E-05 | Qa.III.b | --AINSSIANLSELSRELNSAVKEQKPNIERIIADMKQVSEK----------- | *Ignavibacteriae bacterium HGW-Ignavibacteriae-4* | Bacteria; Ignavibacteriae | hypothetical protein CVV25_12060 | -1 |
|  | **WP_074894735.1** | 3.3E-05 | R.IV | ------QIDEIKDSTKHpNIRKLIKRMDNALANEDYSGVLHASASVFETLAKDL | *Bacillus megaterium* | Bacteria; Firmicutes; Bacilli; Bacillales; Bacillaceae; Bacillus | hypothetical protein | -1 |
| | **OFY60351.1** | 3.4E-05 | Qc.III | ---LQKQVADLKDNVSKLNSKVDDLNGKITKLEEEAEKKEKEIKELKSRIK-- | *Bacteroidetes bacterium RBG_19FT_COMBO_42_10* | Bacteria; Bacteroidetes | hypothetical protein A2V50_04890 | -1 |
|  | **WP_084176478.1** | 3.4E-05 | Qa.IV | ----KQTMGGINELTTNLNSLVVENRQTLVGTLDSITRTSDQMQNAIASLDTA | *Limnothrix rosea* | Bacteria; Cyanobacteria; Synechococcales; Pseudanabaenaceae; Limnothrix | MCE family protein | -1 |
|  | **WP_089291378.1** | 3.4E-05 | SNAP.b | ----------SDEQATATRKELAAQHEKIAQAGRELEAAQAKVAEAERRL--- | *Actinoplanes regularis* | Bacteria; Actinobacteria; Micromonosporales; Micromonosporaceae; Actinoplanes | Laminin subunit beta-1 | -1 |
|  | **WP_091625378.1** | 3.4E-05 | SNAP.b | ------MASQYEKMLEDALGAYQRQRERLTDAKERMDSLTSTATSARR----- | *Amycolatopsis saalfeldensis* | Bacteria; Actinobacteria; Pseudonocardiales; Pseudonocardiaceae; Amycolatopsis | YbaB/EbfC family DNA-binding protein | -1 |
|  | **WP_026817911.1** | 3.5E-05 | SNAP.b | ---------QSVARPFQTEQQLADARQKLENGEQKLSDAKAQLQDGTRKL--- | *Arthrobacter castelli* | Bacteria; Actinobacteria; Micrococcales; Micrococcaceae; Arthrobacter | RND transporter | -1 |
|  | **RAI14819.1** | 3.6E-05 | Qa.IV | --DIDKAMVDLSNAMTLVNKMTPEQKTQIECIVSDTAETTRNLKK-------- | *Candidatus Melainabacteria bacterium* | Bacteria; Candidatus Melainabacteria; unclassified Candidatus Melainabacteria | hypothetical protein DKM22_07250 | -1 |
|  | **SCN20291.1** | 3.6E-05 | Qc.III | -----------KELAKQIYEKVCEMEKKVDKLDNRIDSLEGKVDSLEGSFK-- | *Bacillus cereus* | Bacteria; Firmicutes; Bacilli; Bacillales; Bacillaceae; Bacillus; Bacillus cereus group | SMC1-family ATPase involved in DNA repair | -1 |

| | | | | | | | | |
|---|---|---|---|---|---|---|---|---|
|  | AEM69790.1 | 3.7E-05 | Qb.III | -EQIISLQQESLQAMQKVLDGLRNQNERLERSNQKLRKRISE----------- | *Muricauda ruestringensis DSM 13258* | Bacteria; Bacteroidetes; Flavobacteriia; Flavobacteriales; Flavobacteriaceae; Muricauda | hypothetical protein Murru_0740 | -1 |
|  | WP_095132019.1 | 3.7E-05 | Qa.III.b | MKQVEEIIYELEELRRELNSLISEKTKLI----------ESQVIETSQKL--- | *Anaeromicrobium sediminis* | Bacteria; Firmicutes; Clostridia; Clostridiales; Clostridiaceae; Anaeromicrobium | aspartyl-phosphate phosphatase Spo0E family protein | -1 |
|  | WP_104809085.1 | 3.8E-05 | Qb.III.b | ------------ETGKAEILNLYNQKLEELNINYRYQEIDTTFGKTNIIITG- | *Polaribacter filamentus* | Bacteria; Bacteroidetes; Flavobacteriia; Flavobacteriales; Flavobacteriaceae; Polaribacter | hypothetical protein | -1 |
| | WP_092694614.1 | 3.9E-05 | Qb.III | -DQQEAQLAKAHELGHARLDEMVAQRDAL---NEAIDELRGQLDWVGRTLNSM | *Roseivivax roseus* | Bacteria; Proteobacteria; Alphaproteobacteria; Rhodobacterales; Rhodobacteraceae; Roseivivax | MerR family DNA-binding transcriptional regulator | -1 |
| | WP_020621256.1 | 4E-05 | Qc.III.b | LDSLYADIAALRRLVETLNSTIQDQNKIIQELRSDTHEQTAVSQNQTKKVE-- | *Succinimonas amylolytica* | Bacteria; Proteobacteria; Gammaproteobacteria; Aeromonadales; Succinivibrionaceae; Succinimonas | hypothetical protein | -1 |
|  | OGR17086.1 | 4.1E-05 | Qa.III.b | -------MRNVNAAVKDIGRILKENQEALSRAIPNIAIASENVAVITQDL--- | *Desulfobacterales bacterium GWB2_56_26* | Bacteria; Proteobacteria; Deltaproteobacteria; Desulfobacterales | hypothetical protein A2X81_15350 | -1 |
|  | WP_008993578.1 | 4.2E-05 | R.III | --ALESQAAETTGVASEALENKADAVEKAGEA--KSDALENQADAIKK----- | *Novosphingobium sp. Rr 2-17* | Bacteria; Proteobacteria; Alphaproteobacteria; Sphingomonadales; Sphingomonadaceae; Novosphingobium | hypothetical protein | -1 |
|  | WP_059407064.1 | 4.2E-05 | Qb.III | ------MVIETVKISDRTLAVLAEQIKNLSGANKTADSVDEAFAKTR------ | *Pseudomonas sp. RIT-PI-q* | Bacteria; Proteobacteria; Gammaproteobacteria; Pseudomonadales; Pseudomonadaceae; Pseudomonas | hypothetical protein | -1 |
|  | WP_063571525.1 | 4.2E-05 | SNAP.c | LDKVSEDIVDLKVQTVAIGARLTGVEERLDRTDERVDRAEER----------- | *Luteibacter rhizovicinus* | Bacteria; Proteobacteria; Gammaproteobacteria; Xanthomonadales; Rhodanobacteraceae; Luteibacter | hypothetical protein | -1 |
|  | KGP75063.1 | 4.3E-05 | R.IV | -DSAMETLDETKKKTEKLIDKA---EEKAKELKDKAEDMKDMAE--------- | *Desulfosporosinus sp. Tol-M* | Bacteria; Firmicutes; Clostridia; Clostridiales; Peptococcaceae; Desulfosporosinus | hypothetical protein JT05_12480 | -1 |
|  | WP_016131477.1 | 4.3E-05 | Qa.III | --ELHEKIADLHEMVEKVLSLLNVQQSIIFLLKDVFQYSLSDIAK-------- | *Bacillus cereus* | Bacteria; Firmicutes; Bacilli; Bacillales; Bacillaceae; Bacillus; Bacillus cereus group | RNA polymerase subunit sigma-70 | -1 |
|  | AKT90549.1 | 4.5E-05 | Qc.III | ----ENAINRTNSKLDNIDNKLSNTNSKLDSINNKLDRTNSKLDDINKNIS-- | *Campylobacter ureolyticus RIGS 9880* | Bacteria; Proteobacteria; Epsilonproteobacteria; Campylobacterales; Campylobacteraceae; Campylobacter | hypothetical protein CUREO_0685 | -1 |
|  | WP_018930791.1 | 4.5E-05 | Qb.III | LKEAYAQLEEAKLTGSEINSTLLDQINQLEKQVQNVDIID------------- | *Gracilibacillus lacisalsi* | Bacteria; Firmicutes; Bacilli; Bacillales; Bacillaceae; Gracilibacillus | low molecular weight protein arginine phosphatase | -1 |
| | WP_022592437.1 | 4.5E-05 | Qb.III.d | ---SEKEVEKTAHLARAAAEEAKRAMEQAAKAEEAANLARSKLKKAKKELKQ- | *Vibrio nigripulchritudo* | Bacteria; Proteobacteria; Gammaproteobacteria; Vibrionales; Vibrionaceae; Vibrio | hypothetical protein | -1 |
|  | WP_066112547.1 | 4.6E-05 | SNAP.b | -NRLAEDMREMRTDMKKELGELRQISERLVKIETRLEDIQE------------ | *Blastomonas* | Bacteria; Proteobacteria; Alphaproteobacteria; Sphingomonadales; Sphingomonadaceae | MULTISPECIES: hypothetical protein | -1 |
|  | PCH56635.1 | 4.7E-05 | Qc.III | ---IESDLTHLEQLVASLNHTITEHDKMIQQLQTQVNRVATGLE--------- | *Verrucomicrobiales bacterium* | Bacteria; Verrucomicrobia; Verrucomicrobiae; Verrucomicrobiales | hypothetical protein COC21_00295 | -1 |
|  | PSH01042.1 | 4.8E-05 | Qa.IV | --ALEKEVTELLADFSDLSDRIEGSEETVEGLESDIESVEERV---------- | *Nanohaloarchaea archaeon SW_7_46_7* | Archaea; Candidatus Nanohaloarchaeota; Nanohaloarchaea | hypothetical protein BRC30_00350 partial | -1 |
| | WP_069447388.1 | 4.8E-05 | Qc.III.c | LQKLQEKLRTMKEPSAALTQQIVKQKEKLDSLSVGNDAYVKKLKSLGKELN-- | *Vibrio scophthalmi* | Bacteria; Proteobacteria; Gammaproteobacteria; Vibrionales; Vibrionaceae; Vibrio | hypothetical protein | -1 |
|  | WP_095480797.1 | 4.8E-05 | Qc.III | --------NILADNGEMIQQDIRIQEGKIVEMGDQLSANDEAVFEANGQL--- | *Bacillaceae bacterium SAOS 7* | Bacteria; Firmicutes; Bacilli; Bacillales; Bacillaceae; unclassified Bacillaceae | dihydroorotase | -1 |
| | PHQ55371.1 | 4.9E-05 | Qb.III | INALHKELDDIKEHSKQVIEDLNADGDSVTTADRVVSGLDKILSKISKII--- | *Sulfurimonas sp.* | Bacteria; Proteobacteria; Epsilonproteobacteria; Campylobacterales; Helicobacteraceae; Sulfurimonas | hypothetical protein COA30_06905 | -1 |
|  | PZU47420.1 | 4.9E-05 | Qc.III | ---------KVAQTGNMLKGEVELLDEQLRAFEDDISKLEAKLREARSR---- | *Sphingomonas sp.* | Bacteria; Proteobacteria; Alphaproteobacteria; Sphingomonadales; Sphingomonadaceae; Sphingomonas | phage shock protein PspA | -1 |
|  | WP_073246164.1 | 4.9E-05 | R.IV | -KKIKDKTKELKDTMKNDFDKAIQKiEDKYQNIAQNTNDLVQE----------- | *Flavobacterium flevense* | Bacteria; Bacteroidetes; Flavobacteriia; Flavobacteriales; Flavobacteriaceae; Flavobacterium | YtxH domain-containing protein | -1 |
|  | APS82305.1 | 5E-05 | Qa.IV | -----VAMNNIKNMNSHVATTTDEQNNMTKRIEEQVSSTLLSVRKA------- | *Piscirickettsia salmonis* | Bacteria; Proteobacteria; Gammaproteobacteria; Thiotrichales; Piscirickettsiaceae; Piscirickettsia | hypothetical protein AVM71_02530 | -1 |
| | OYX45924.1 | 5.1E-05 | SNAP.c | -RQLENDAKNLTSLGRTFAPELMSQLDQLDQLINAARGLALKVNETRTAL--- | *Alphaproteobacteria bacterium 32-64-14* | Bacteria; Proteobacteria; Alphaproteobacteria | P-type conjugative transfer protein TrbJ | -1 |
|  | WP_007694330.1 | 5.2E-05 | SNAP.b | -----------SQSSSTIIDALSHQTEDLCQVHNHITQVIADLDVSQEMLS-- | *Halococcus hamelinensis* | Archaea; Euryarchaeota; Halobacteria; Halobacteriales; Halococcaceae; Halococcus | transposase | -1 |
|  | WP_080706143.1 | 5.2E-05 | Qc.II | LDRMANDIYQSKNATNEILGRINASNDLLQSINSKMGSSGS------------ | *Aeromonas hydrophila* | Bacteria; Proteobacteria; Gammaproteobacteria; Aeromonadales; Aeromonadaceae; Aeromonas | hypothetical protein | -1 |
|  | OUX02977.1 | 5.3E-05 | Qc.II | IDELGQEVQTLEQAFVEINNELTKTDTLLDGTRASFEKQIRTLK--------- | *Flavobacteriaceae bacterium TMED238* | Bacteria; Bacteroidetes; Flavobacteriia; Flavobacteriales; Flavobacteriaceae | hypothetical protein CBD98_00580 partial | -1 |
|  | WP_014404862.1 | 5.3E-05 | Qc.II | ----------LRKLAEQMEKEIEGGNASLVNLSRVFDKSPTFLKGVAKRSR-- | *Methanocella conradii* | Archaea; Euryarchaeota; Stenosarchaea group; Methanomicrobia; Methanocellales; Methanocellaceae; Methanocella | hypothetical protein | -1 |
|  | WP_012580465.1 | 5.4E-05 | SNAP.b | ----------ATTTLLNIKEQLEKNKKRLDEIEEMINKIKVDAKVVARKI--- | *Thermosipho africanus* | Bacteria; Thermotogae; Thermotogales; Fervidobacteriaceae; Thermosipho | DUF342 domain-containing protein | -1 |
|  | WP_018575683.1 | 5.4E-05 | Qb.III | ----ERTADLVERVEAIAQKELKHQAQQLTRVEYHLSDINSETR--------- | *Oligella ureolytica* | Bacteria; Proteobacteria; Betaproteobacteria; Burkholderiales; Alcaligenaceae; Oligella | hypothetical protein | -1 |
|  | WP_106898922.1 | 5.4E-05 | Qc.I | -SNLVSYTRNMRAGFINLKEIIKKDTKSKEKILKIIEQNYIKAKK-------- | *Clostridium botulinum* | Bacteria; Firmicutes; Clostridia; Clostridiales; Clostridiaceae; Clostridium | hypothetical protein | -1 |
|  | OGT90947.1 | 5.5E-05 | Qb.I | --------AQTADRFSRAAQQIEATSQAVKQSHDQLSEVARTIRDGAQLVQ-- | *Gammaproteobacteria bacterium RIFOXYD12_FULL_61_37* | Bacteria; Proteobacteria; Gammaproteobacteria | hypothetical protein A2514_07490 | -1 |
|  | WP_087844761.1 | 5.6E-05 | SNAP.b | ------LATSTVAHAESALKQLHQVNNQLSHIENAVNGTQKKYHNTQRTIKEI | *Photobacterium sp. CECT 9190* | Bacteria; Proteobacteria; Gammaproteobacteria; Vibrionales; Vibrionaceae; Photobacterium | hypothetical protein | -1 |
|  | WP_113475159.1 | 5.6E-05 | SNAP.b | -----------AEAARANAGVLEELNEKIARLVSENEKLTADLDEADKKISAL | *Rhizobiales bacterium* | Bacteria; Proteobacteria; Alphaproteobacteria; Rhizobiales | hypothetical protein | -1 |
|  | KPB68883.1 | 5.7E-05 | Qc.I | -------TRELRDHAEQMLRAIAADLQTDQSSQEQVDKSQGQVESQE------ | *Pseudomonas syringae pv. maculicola* | Bacteria; Proteobacteria; Gammaproteobacteria; Pseudomonadales; Pseudomonadaceae; Pseudomonas | Signal transduction histidine kinase | -1 |
|  | KKR01288.1 | 5.8E-05 | SNAP.b | -------------RLDRIVTMLDKQSGNLQSIEKELTLIRASLDTNDTNQSSL | *Microgenomates group bacterium GW2011_GWC1_39_12* | Bacteria | hypothetical protein UT26_C0034G0023 | -1 |
| | SHE13204.1 | 5.9E-05 | Qa.IV | MIQIQTSAEDLNRTVTELHTQLGEHKESLAQLQSQFDHVQSDLEAVKEELSLA | *Chlamydia abortus* | Bacteria; Chlamydiae; Chlamydiales; Chlamydiaceae; Chlamydia/ Chlamydophila group; Chlamydia | Membrane-bound metallopeptidase | -1 |
|  | WP_075346056.1 | 5.9E-05 | Qc.I | -----NLAQKLQDEAVIYVASIGEDISVVDNSQKAAVNQSSSSQTA------- | *Streptococcus sp. 'caviae'* | Bacteria; Firmicutes; Bacilli; Lactobacillales; Streptococcaceae; Streptococcus | competence protein CelA | -1 |
|  | WP_101844228.1 | 5.9E-05 | Qc.III | ------TLDELRDQVESLDEEIDRHDQVLDLVNNQLHSVDRKINRPGRELD-- | *Halobacillus sp. Marseille-P3879* | Bacteria; Firmicutes; Bacilli; Bacillales; Bacillaceae; Halobacillus | hypothetical protein | -1 |
|  | CAP12996.1 | 6E-05 | Qa.III | -REIRKELEDVTSAIEDVSDELSEQGSGI-----------RETEAVKKELQK- | *Halobacterium salinarum R1* | Archaea; Euryarchaeota; Halobacteria; Halobacteriales; Halobacteriaceae; Halobacterium | homolog to pHK2-ORF10 | -1 |
|  | WP_093824606.1 | 6E-05 | Qb.I | LENTASSASQTVGNMNKASETLTKTSENLNKASEKLNNTNS------------ | *Spirosoma endophyticum* | Bacteria; Bacteroidetes; Cytophagia; Cytophagales; Cytophagaceae; Spirosoma | MCE family protein | -1 |
| | AFS48962.1 | 6.1E-05 | R.IV | ---LDQKIEEIKSNINSSLKSFSDAENKLKEAEKQTADLDNKIQELLNSAK-- | *alpha proteobacterium HIMB59* | Bacteria; Proteobacteria; Alphaproteobacteria; Pelagibacterales; Pelagibacteraceae | ATP synthase B/B' CF(0) | -1 |
|  | ETO92760.1 | 6.1E-05 | Qb.I | -------LSLLVNMSSPALKRLHASTSTLKAMVKEYQELRAIMLRNQQKL--- | *Legionella oakridgensis RV-2-2007* | Bacteria; Proteobacteria; Gammaproteobacteria; Legionellales; Legionellaceae; Legionella | hypothetical protein LOR_74c21250 | -1 |
|  | WP_076643895.1 | 6.1E-05 | R.IV | TDKIKGAANEVAGNVKQSVGNVTGNDKlqaegKAQELEGKTQ------------- | *Methylorubrum extorquens* | Bacteria; Proteobacteria; Alphaproteobacteria; Rhizobiales; Methylobacteriaceae; Methylorubrum | CsbD family protein | -1 |
| | WP_090303857.1 | 6.1E-05 | SNAP.c | -DALESQLRSLTERVERLETELERKDERIATLESKLDEHDARLEQANTQLK-- | *Natronorubrum texcoconense* | Archaea; Euryarchaeota; Halobacteria; Natrialbales; Natrialbaceae; Natronorubrum | hypothetical protein | -1 |
|  | WP_103776862.1 | 6.2E-05 | Qb.II | LQRVRQGGNRALDGIQQALDTISDQRAQMAVQQEKARQLI------------- | *Citrobacter amalonaticus* | Bacteria; Proteobacteria; Gammaproteobacteria; Enterobacterales; Enterobacteriaceae; Citrobacter | flagellar hook-associated protein | -1 |
|  | WP_053071213.1 | 6.3E-05 | R.IV | -ESLKSAIQEIVGEMNQTLAELEEKFSNLENIETKTGAQAKADKA-------- | *Bacillus glycinifermentans* | Bacteria; Firmicutes; Bacilli; Bacillales; Bacillaceae; Bacillus | phage baseplate upper protein | -1 |
|  | PLY12952.1 | 6.5E-05 | Qc.III | -EEVLQTLSQQNETGKNLHSELQQMGSQLETISDSIDKGNG------------ | *Desulfuromonas sp.* | Bacteria; Proteobacteria; Deltaproteobacteria; Desulfuromonadales; Desulfuromonadaceae; Desulfuromonas | hypothetical protein C0624_00250 | -1 |
|  | WP_037062938.1 | 6.6E-05 | Qb.I | -------MSDETSRVTQLEETLAHQAKAIEELSDQLTEQWKVVEQTRAKL--- | *Rhizobium leguminosarum* | Bacteria; Proteobacteria; Alphaproteobacteria; Rhizobiales; Rhizobiaceae; Rhizobium/Agrobacterium group; Rhizobium | hypothetical protein | -1 |
|  | WP_057005010.1 | 6.6E-05 | Qc.I | -------TRGLQDNIQRLSNDIEETERRLKESVPGSSKAISNLKVVSDR---- | *Pseudomonas* | Bacteria; Proteobacteria; Gammaproteobacteria; Pseudomonadales; Pseudomonadaceae | MULTISPECIES: hypothetical protein | -1 |
| | ABL87258.1 | 6.7E-05 | SNAP.c | MAQITSDVSSLKNTAAALQNEVSAVKNAVSQVSTKVDAVGGQVAALSQKIDQL | *Pyrobaculum islandicum DSM 4184* | Archaea; Crenarchaeota; Thermoprotei; Thermoproteales; Thermoproteaceae; Pyrobaculum | conserved hypothetical protein | -1 |
|  | CCG55689.1 | 6.7E-05 | SNAP.c | LDNIESISEKVNNIALFVESEADRTKKSLSDVHDAIDNVTS------------ | *Brachyspira pilosicoli WesB* | Bacteria; Spirochaetes; Brachyspirales; Brachyspiraceae; Brachyspira | hypothetical protein WESB_0217 | -1 |
|  | OHA43422.1 | 6.7E-05 | Qc.I | -----EMAGKLKESYAGLEQKVRDRTRDLDAANQQLKASNQQIRASNQQLNA- | *Candidatus Taylorbacteria bacterium RIFCSPLOWO2_12_FULL_44_15c* | Bacteria; Candidatus Taylorbacteria | hypothetical protein A3G03_01240 | -1 |
|  | WP_084888362.1 | 6.7E-05 | Qc.I | ---------RLKVDIENQNRILDERQQKLETIQTSIAKNISELTKQQNQLS-- | *Streptococcus mitis* | Bacteria; Firmicutes; Bacilli; Lactobacillales; Streptococcaceae; Streptococcus | hypothetical protein | -1 |
|  | GBC91993.1 | 7E-05 | SNAP.b | ------IVRELVEVARQGMERLGRVEERLERVEERTERMEERLERVEERTERM | *bacterium HR15* | Bacteria | Chromosome partition protein Smc | -1 |
|  | PIR42132.1 | 7E-05 | Qc.I | ---------GLKNKVGEARQSLRNDLKVVQDTIKNAHEAVRNIIIALSR---- | *Candidatus Yanofskybacteria bacterium CG10_big_fil_rev_8_21_14_0_10_3 7_15* | Bacteria; Candidatus Yanofskybacteria | hypothetical protein COV30_00050 | -1 |
| | WP_061324945.1 | 7E-05 | SNAP.c | LGVVEQQIERMDQRAERMEQRVEQMDLRVERMEQRVEQMDLRVERMEQRVERM | *Serratia rubidaea* | Bacteria; Proteobacteria; Gammaproteobacteria; Enterobacterales; Yersiniaceae; Serratia | hypothetical protein | -1 |
|  | WP_064631228.1 | 7.2E-05 | Qc.III | ------AITRFGENANNVQNRLGLQENALSQAGDKMARVTELTVQAN------ | *Xanthomonas nasturtii* | Bacteria; Proteobacteria; Gammaproteobacteria; Xanthomonadales; Xanthomonadaceae; Xanthomonas | flagellar hook-associated protein FlgL | -1 |
|  | PIV38794.1 | 7.3E-05 | Qc.III | -------VQTTKTQGTTLKAEITRMQNQINKLEAEIRLTQTKISQTSLKIQG- | *Candidatus Portnoybacteria bacterium CG02_land_8_20_14_3_00_45_8* | Bacteria; Candidatus Portnoybacteria | hypothetical protein COS30_00205 partial | -1 |
|  | WP_042072886.1 | 7.5E-05 | Qb.I | ----SKQSHQTINQTDANEVSLTESSLVVLKINP--NDVAKITRSGNAAVITL | *Acinetobacter gerneri* | Bacteria; Proteobacteria; Gammaproteobacteria; Pseudomonadales; Moraxellaceae; Acinetobacter | hypothetical protein partial | -1 |
|  | WP_091746365.1 | 7.5E-05 | Qc.III | --ELKTVLTRIEEEQKNIGNRLARTEESLKSVHKRVDKLEGKVH--------- | *Propionispora vibrioides* | Bacteria; Firmicutes; Negativicutes; Selenomonadales; Sporomusaceae; Propionispora | hypothetical protein | -1 |

| | | | | | | | | |
|---|---|---|---|---|---|---|---|---|
| | **WP_021661040.1** | 7.6E-05 | SNAP.b | ---LSRRMDAMETRLNQRMDTLESQlTERLDGIDERLDGIDERLDRVEIRLD-- | *Clostridium* | Bacteria; Firmicutes; Clostridia; Clostridiales; Clostridiaceae | MULTISPECIES: hypothetical protein | -1 |
|  | **WP_094424925.1** | 7.8E-05 | Qa.I | IEKAKENMEQIQQLLRDYQALVLDNASDIEALVDQFVESEANVAKA------- | *Bacillus* | Bacteria; Firmicutes; Bacilli; Bacillales; Bacillaceae | MULTISPECIES: hypothetical protein | -1 |
|  | **CDE96600.1** | 7.9E-05 | Qb.III.d | -------------VSVSSGTDIARDSAQVVIINDNLERINDLLDISSKTVRNI | *Clostridium sp. CAG:567* | Bacteria; Firmicutes; Clostridia; Clostridiales; Clostridiaceae; Clostridium; environmental samples | heavy metal translocating P-type ATPase | -1 |
|  | **PKL15878.1** | 8.2E-05 | Qc.II | LDELSGQISSMKsmisdtlNMTNSISQEAKTGETSIKQMNDNMESMNR-------- | *Spirochaetae bacterium HGW-Spirochaetae-5* | Bacteria; Spirochaetes | hypothetical protein CVV49_19145 partial | -1 |
|  | **WP_019189831.1** | 8.5E-05 | SNAP.c | ---LSNITKQLNEVVKKNNEELDATIKSLPVLTEKADHTITKVN--------- | *Levyella massiliensis* | Bacteria; Firmicutes; Clostridia; Clostridiales; Levyella | hypothetical protein | -1 |
|  | **WP_035162960.1** | 8.5E-05 | Qc.III | -REILNEIGSLKNQVTSMNDRITSMEGRINSIETEVKATRQDIKEL------- | *Caloranaerobacter azorensis* | Bacteria; Firmicutes; Clostridia; Clostridiales; Clostridiaceae; Caloranaerobacter | hypothetical protein | -1 |
|  | **WP_108018099.1** | 8.5E-05 | Qb.III.b | IKRALDVTVEALSVASTIAMGLARQKKQL-DALQTTNETASDL---------- | *Marinobacter sp. PT19DW* | Bacteria; Proteobacteria; Gammaproteobacteria; Alteromonadales; Alteromonadaceae; Marinobacter | toxic anion resistance protein | -1 |
|  | **PIT98396.1** | 8.6E-05 | SNAP.b | -----------KEDLKNIETRLNGVDGRLDSVDKRLDGIDQRLDKIERVQHS- | *Candidatus Andersenbacteria bacterium CG10_big_fil_rev_8_21_14_0_10_54_11* | Bacteria; Candidatus Andersenbacteria | hypothetical protein COT71_00940 | -1 |
|  | **WP_097445825.1** | 8.6E-05 | SNAP.b | ----VGALTESLQNTQNNIQALDRQSKVFERAAGNLSKYREQIARAEQRLAGL | *Escherichia coli* | Bacteria; Proteobacteria; Gammaproteobacteria; Enterobacterales; Enterobacteriaceae; Escherichia | phage tail tape measure protein partial | -1 |
|  | **WP_050640989.1** | 8.7E-05 | Qc.III | LDKLEPIIEEIHNMSaalVEMTAELKHTNKSVEEIKDKVED-------------- | *Eubacterium sp. SB2* | Bacteria; Firmicutes; Clostridia; Clostridiales; Eubacteriaceae; Eubacterium | hypothetical protein | -1 |
|  | **WP_106706705.1** | 8.7E-05 | Qc.I | -------------KAKKLQDRLEQDNTIMEESERrNLERNLANLSREYQRAQK- | *Nitrosomonas sp. APG5* | Bacteria; Proteobacteria; Betaproteobacteria; Nitrosomonadales; Nitrosomonadaceae; Nitrosomonas | OmpH family outer membrane protein | -1 |
|  | **AIW54858.1** | 8.8E-05 | Qa.IV | -----KSLEELKELSKKLAETNKENKSIVDIIQDKKDELNNYTENAIKNI--- | *Clostridium botulinum* | Bacteria; Firmicutes; Clostridia; Clostridiales; Clostridiaceae; Clostridium | hypothetical protein (plasmid) | -1 |
|  | **WP_033827909.1** | 8.9E-05 | SNAP.c | ---LKEILSALKNHSENMNKRFDQLDSRMDKLEERVDRLEEKVD--------- | *Bacillus andreraoultii* | Bacteria; Firmicutes; Bacilli; Bacillales; Bacillaceae; Bacillus | hypothetical protein | -1 |
|  | **WP_081724007.1** | 8.9E-05 | Qb.I | ------FLKEEIEKMKTDIKSLKEQTTSLKTVVDALNAGKVITN-----VK-- | *Bacteroides pyogenes* | Bacteria; Bacteroidetes; Bacteroidia; Bacteroidales; Bacteroidaceae; Bacteroides | hypothetical protein | -1 |
|  | **WP_015134488.1** | 9E-05 | Qc.I | --------RNMKNHLMVYQDKLLARKRALIETVIDQLKNISQIEHSRH----- | *Leptolyngbya sp. PCC 7376* | Bacteria; Cyanobacteria; Synechococcales; Leptolyngbyaceae; Leptolyngbya | IS982 family transposase | -1 |
|  | **WP_097074175.1** | 9E-05 | Qc.I | ---------ELKDSISSFKQAIEPDATKLKEEIEGLQKSITEIENNLSELN-- | *Lysinibacillus xyleni* | Bacteria; Firmicutes; Bacilli; Bacillales; Bacillaceae; Lysinibacillus | gas vesicle protein | -1 |
|  | **WP_010304401.1** | 9.4E-05 | Qc.I | -SELSRLGRSLKETLNTIESIVQEKQARLILIKQGLDINPHNTNNLTN----- | *Candidatus Odyssella thessalonicensis* | Bacteria; Proteobacteria; Alphaproteobacteria; Holosporales; Candidatus Paracaedibacteraceae; Candidatus Odyssella | resolvase | -1 |
|  | **PIQ10023.1** | 9.5E-05 | Qb.II | -----QAMQSLISEASTLLEGFERQNRQIMNVVEAVYARLNQLPG-------- | *Hydrogenophilales bacterium CG18_big_fil_WC_8_21_14_2_50_58_12* | Bacteria; Proteobacteria; Hydrogenophilalia; Hydrogenophilales | hypothetical protein COW70_14295 | -1 |
|  | **OHE82596.1** | 9.7E-05 | Qa.IV | AQQILKQLQELRQNQQQLQQTIIQQQRTIEALEKRVAAAD------------ | *Verrucomicrobia bacterium GWF2_62_7* | Bacteria; Verrucomicrobia; unclassified Verrucomicrobia | hypothetical protein A2107_08025 | -1 |
|  | **PWM48277.1** | 9.9E-05 | SNAP.b | ----HNLAHQAEDAGFETA------GEKLHAAMSALDDIRALLDDAKEAL--- | *Coriobacteriia bacterium* | Bacteria; Actinobacteria; Coriobacteriia | dynein gamma chain protein | -1 |
|  | **ANW59524.1** | 0.0001 | Qa.III | ------NVNQLNTLSTSMSQSLSNQQTQLNNLGSQLNQTQQQLQQTDTM---- | *Burkholderia pseudomallei* | Bacteria; Proteobacteria; Betaproteobacteria; Burkholderiales; Burkholderiaceae; Burkholderia; pseudomallei group | hypothetical protein A7U59_26260 | -1 |
| | **KUO86143.1** | 0.0001 | SNAP.c | ---VEDGLTRLENAVEELTRAVKSHDERLTRLENAVEELTKAIKAHEERLTRL | *Caldivirga sp. CIS_19* | Archaea; Crenarchaeota; Thermoprotei; Thermoproteales; Thermoproteaceae; Caldivirga | hypothetical protein AT712_04990 | -1 |
|  | **WP_002771781.1** | 0.0001 | Qb.III.b | -----------ESLAENTLTELREIRLGLQEIKLGLQETNQGLKETRAIANS- | *Microcystis aeruginosa* | Bacteria; Cyanobacteria; Oscillatoriophycideae; Chroococcales; Microcystaceae; Microcystis | hypothetical protein | -1 |
|  | **WP_038264096.1** | 0.0001 | Qc.I | ------------NNVEEIGEKLVFSIQLLNEDQIGVVNTLANLKATLNKLVSY | *Zhouia amylolytica* | Bacteria; Bacteroidetes; Flavobacteriia; Flavobacteriales; Flavobacteriaceae; Zhouia | DNA repair protein RecN | -1 |
|  | **WP_070633353.1** | 0.0001 | R.IV | ---IEAVVEADAGAMRQVLHNILKNGAEAAAADKQPEVLIQTA---------- | *Eikenella* | Bacteria; Proteobacteria; Betaproteobacteria; Neisseriales; Neisseriaceae | MULTISPECIES: PAS domain-containing sensor histidine kinase | -1 |
|  | **WP_083558381.1** | 0.0001 | Qc.I | -----------RARASELARAVEADRATVDALRTAVEEAETALRAAHARLEE- | *Olsenella phocaeensis* | Bacteria; Actinobacteria; Coriobacteriia; Coriobacteriales; Atopobiaceae; Olsenella | hypothetical protein | -1 |
|  | **WP_113634993.1** | 0.0001 | Qa.IV | --------RDLHDGIGQMLSVVKMNVSTLNSADKTVDNTLNLVDKTIDELR-- | *Nubsella zeaxanthinifaciens* | Bacteria; Bacteroidetes; Sphingobacteriia; Sphingobacteriales; Sphingobacteriaceae; Nubsella | hypothetical protein | -1 |
| **▼ 3** | | | | | | | | |
| | **OJV90465.1** | 3.7E-18 | R.IV | IKQIQQEIDETKKIMLQNIDKVIVRGRKIETLRDRSKEMQEKSQRFVKQAKHL | *Gammaproteobacteria bacterium 39-13* | Bacteria; Proteobacteria; Gammaproteobacteria | hypothetical protein BGO43_06340 | 3 |
|  | **KRG19470.1** | 2E-14 | R.III | -----ETIDRTKRIMVDNIDQLLQRGEKLEQLLDKTAQTEEKAQIFTQQARQ- | *Candidatus Berkiella cookevillensis* | Bacteria; Proteobacteria; Gammaproteobacteria; Legionellales; Coxiellaceae; Candidatus Berkiella | Synaptobrevin | 3 |
|  | **WP_057623293.1** | 2.4E-14 | R.III | -----ETIDRTKRIMVDNIDQLLQRGEKLEQLLDKTAQTEEKAQIFTQQARQ- | *Candidatus Berkiella cookevillensis* | Bacteria; Proteobacteria; Gammaproteobacteria; Legionellales; Coxiellaceae; Candidatus Berkiella | hypothetical protein | 3 |
| **▼ 4** | | | | | | | | |
|  | **SIR48573.1** | 1.2E-11 | SNAP.b | --RAVRMARNTREIGADTVEALEGQSKQIQKIEEITENIQNKAEQAEE----- | *Fluoribacter gormanii* | Bacteria; Proteobacteria; Gammaproteobacteria; Legionellales; Legionellaceae; Fluoribacter | hypothetical protein SAMN05421777_11361 | 4 |
|  | **WP_028380397.1** | 5.4E-11 | SNAP.c | LDELGEIIDDLNQLAREQGKEITWQNEHLNKLNPKIEKATESINYE------- | *Legionella cherrii* | Bacteria; Proteobacteria; Gammaproteobacteria; Legionellales; Legionellaceae; Legionella | hypothetical protein | 4 |
|  | **SIR48573.1** | 3.4E-10 | SNAP.c | LDELGEIIDDLNQLAREQGKEITWQNKHLDKLKPKIEMAKETVN--------- | *Fluoribacter gormanii* | Bacteria; Proteobacteria; Gammaproteobacteria; Legionellales; Legionellaceae; Fluoribacter | hypothetical protein SAMN05421777_11361 | 4 |
|  | **WP_028380397.1** | 1.3E-09 | SNAP.b | --RAARMARRTREIGADTVEALEGQSKKIQRIEKITENIQNTAEQAEE----- | *Legionella cherrii* | Bacteria; Proteobacteria; Gammaproteobacteria; Legionellales; Legionellaceae; Legionella | hypothetical protein | 4 |
| **▼ 5** | | | | | | | | |
| | **WP_057622768.1** | 1.3E-11 | Qb.III.d | --QILKIVNETKKIGEDTRKRLDQQGEQIGVIDRDVQELDYQAKKNKRTVKG- | *Candidatus Berkiella cookevillensis* | Bacteria; Proteobacteria; Gammaproteobacteria; Legionellales; Coxiellaceae; Candidatus Berkiella | hypothetical protein | 5 |
| | **KRG20005.1** | 1.4E-11 | Qb.III.d | --QILKIVNETKKIGEDTRKRLDQQGEQIGVIDRDVQELDYQAKKNKRTVKG- | *Candidatus Berkiella cookevillensis* | Bacteria; Proteobacteria; Gammaproteobacteria; Legionellales; Coxiellaceae; Candidatus Berkiella | SNARE domain protein | 5 |
| **▼ 6** | | | | | | | | |
| | **OSQ28999.1** | 7.5E-11 | Qc.III | IEDIGRIIENINEIATTISAAVEEQGAATDEITRNVRSAADRTRTVSSSIND- | *Thalassospira sp. MCCC 1A03138* | Bacteria; Proteobacteria; Alphaproteobacteria; Rhodospirillales; Rhodospirillaceae; Thalassospira | hypothetical protein TH468_15855 partial | 6 |
|  | **KEO50828.1** | 1E-09 | Qc.III.c | -EDIGRIIENINEIATTISAAVEEQGAATDEITRNVRSAAD------------ | *Thalassospira permensis NBRC 106175* | Bacteria; Proteobacteria; Alphaproteobacteria; Rhodospirillales; Rhodospirillaceae; Thalassospira | hypothetical protein SMB34_21865 partial | 6 |
| | **SEE46404.1** | 1.4E-09 | Qc.III | IEDIGRIIENINEIATTISAAVEEQGAATDEITRNVRSAADRTRTVSSSIND- | *Thalassospira sp. KO164* | Bacteria; Proteobacteria; Alphaproteobacteria; Rhodospirillales; Rhodospirillaceae; Thalassospira | Methyl-accepting chemotaxis protein | 6 |
| | **WP_065701939.1** | 1.4E-09 | Qc.III | IEDIGRIIENINEIATTISAAVEEQGAATDEITRNVRSAADRTRTVSSSIND- | *Thalassospira* | Bacteria; Proteobacteria; Alphaproteobacteria; Rhodospirillales; Rhodospirillaceae | MULTISPECIES: HAMP domain-containing protein | 6 |
| | **WP_071239546.1** | 1.4E-09 | Qc.III | IEDIGRIIENINEIATTISAAVEEQGAATDEITRNVRSAADRTRTVSSSIND- | *Thalassospira sp. MIT1004* | Bacteria; Proteobacteria; Alphaproteobacteria; Rhodospirillales; Rhodospirillaceae; Thalassospira | HAMP domain-containing protein | 6 |
|  | **WP_110252300.1** | 1.7E-09 | Qc.III | -----RIIENINEIATTISAAVEEQGAATDEITRNVRSAADRTRTVSSSIND- | *Thalassospira sp. 11-3* | Bacteria; Proteobacteria; Alphaproteobacteria; Rhodospirillales; Rhodospirillaceae; Thalassospira | chemotaxis protein partial | 6 |
|  | **WP_089672391.1** | 2E-09 | Qc.III | -EELSAAASQIATGATQINEEVGDQNRSLRQVSDEMQNLSATM---------- | *Halohasta litchfieldiae* | Archaea; Euryarchaeota; Halobacteria; Haloferacales; Halorubraceae; Halohasta | PAS domain-containing protein | 6 |
| | **WP_085618428.1** | 2.4E-09 | Qc.III | IEDIGKIIENINEIATTISAAVEEQGAATDEITRNVRGAADRTRAVSSSIND- | *Thalassospira alkalitolerans* | Bacteria; Proteobacteria; Alphaproteobacteria; Rhodospirillales; Rhodospirillaceae; Thalassospira | HAMP domain-containing protein | 6 |
| | **WP_085618430.1** | 2.4E-09 | Qc.III | IEDIGKIIENINEIATTISAAVEEQGAATDEITRNVRGAADRTRAVSSSIND- | *Thalassospira alkalitolerans* | Bacteria; Proteobacteria; Alphaproteobacteria; Rhodospirillales; Rhodospirillaceae; Thalassospira | HAMP domain-containing protein | 6 |
| | **RCK50379.1** | 3.2E-09 | Qc.III | IEMIGTIIESINEIATTISAAVEEQGAATNEITRNVRNAAERTRMVSTSIS-- | *Thalassospira profundimaris* | Bacteria; Proteobacteria; Alphaproteobacteria; Rhodospirillales; Rhodospirillaceae; Thalassospira | hypothetical protein TH25_12405 partial | 6 |
|  | **WP_063318529.1** | 3.4E-09 | Qa.IV | INAITKTMSDVNDITATIAAAVEQQGAATTEISENVQQAA------------ | *Pseudovibrio sp. Ad13* | Bacteria; Proteobacteria; Alphaproteobacteria; Rhodobacterales; Rhodobacteraceae; Pseudovibrio | methyl-accepting chemotaxis protein | 6 |
|  | **PKP78959.1** | 3.8E-09 | Qa.IV | IKSIEKTITEMNQIATSTAAAVDEQDAATNEIARNVQEAA------------ | *Alphaproteobacteria bacterium HGW-Alphaproteobacteria-3* | Bacteria; Proteobacteria; Alphaproteobacteria | chemotaxis protein | 6 |
| | **WP_052268357.1** | 4.4E-09 | Qc.III | IEDIGRIIENINEIATTISAAVEEQGAATDEITRNVRSAADRTRTVSASIN-- | *Thalassospira xiamenensis* | Bacteria; Proteobacteria; Alphaproteobacteria; Rhodospirillales; Rhodospirillaceae; Thalassospira | HAMP domain-containing protein | 6 |
| | **WP_097050601.1** | 4.4E-09 | Qc.III | IEDIGRIIENINEIATTISAAVEEQGAATDEITRNVRSAADRTRTVSASIN-- | *Thalassospira xiamenensis* | Bacteria; Proteobacteria; Alphaproteobacteria; Rhodospirillales; Rhodospirillaceae; Thalassospira | HAMP domain-containing protein | 6 |
|  | **WP_008974387.1** | 4.5E-09 | Qa.III | IQAIGGTIAEIDEISKEIATAVEQQGLATGEISGNVQQAAEGTRQ-------- | *Bradyrhizobium sp. STM 3843* | Bacteria; Proteobacteria; Alphaproteobacteria; Rhizobiales; Bradyrhizobiaceae; Bradyrhizobium | methyl-accepting chemotaxis protein | 6 |
| | **WP_071239545.1** | 4.5E-09 | Qc.III | IEDIGRIIENINEIATTISAAVEEQGAATDEITRNVRSAADRTRTVSASIN-- | *Thalassospira sp. MIT1004* | Bacteria; Proteobacteria; Alphaproteobacteria; Rhodospirillales; Rhodospirillaceae; Thalassospira | HAMP domain-containing protein | 6 |
| | **WP_085645604.1** | 4.5E-09 | Qc.III | IEDIGRIIENINEIATTISAAVEEQGAATDEITRNVRSAADRTRTVSASIN-- | *Thalassospira sp. MCCC 1A03138* | Bacteria; Proteobacteria; Alphaproteobacteria; Rhodospirillales; Rhodospirillaceae; Thalassospira | HAMP domain-containing protein | 6 |
| | **WP_097050602.1** | 4.5E-09 | Qc.III | IEDIGRIIENINEIATTISAAVEEQGAATDEITRNVRSAADRTRTVSASIN-- | *Thalassospira xiamenensis* | Bacteria; Proteobacteria; Alphaproteobacteria; Rhodospirillales; Rhodospirillaceae; Thalassospira | HAMP domain-containing protein | 6 |
| | **WP_114124035.1** | 4.5E-09 | Qc.III | IEDIGRIIENINEIATTISAAVEEQGAATDEITRNVRSAADRTRTVSASIN-- | *Thalassospira xianhensis* | Bacteria; Proteobacteria; Alphaproteobacteria; Rhodospirillales; Rhodospirillaceae; Thalassospira | HAMP domain-containing protein | 6 |
|  | **WP_008615362.1** | 5.4E-09 | Qa.IV | IRDILSTISELSQVTSSIASAVQEQQAATAEIARNVEQAAQGTE--------- | *Magnetospirillum caucaseum* | Bacteria; Proteobacteria; Alphaproteobacteria; Rhodospirillales; Rhodospirillaceae; Magnetospirillum | methyl-accepting chemotaxis protein | 6 |
|  | **WP_085374982.1** | 6.5E-09 | Qa.IV | IQDILSTISELSQVTSSIASAVQEQQAATAEIARNVEQAAQGTE--------- | *Magnetospirillum sp. ME-1* | Bacteria; Proteobacteria; Alphaproteobacteria; Rhodospirillales; Rhodospirillaceae; Magnetospirillum | methyl-accepting chemotaxis protein | 6 |
|  | **WP_073956994.1** | 7.2E-09 | Qa.III | IDGITRIIHDLNEISAEVAAAVQQQGSATNEISNSVQQAVSSTQE-------- | *Thalassospira sp. TSL5-1* | Bacteria; Proteobacteria; Alphaproteobacteria; Rhodospirillales; Rhodospirillaceae; Thalassospira | methyl-accepting chemotaxis protein | 6 |

| | | | | | | | | |
|---|---|---|---|---|---|---|---|---|
|  | **PDT85964.1** | 7.4E-09 | Qa.IV | IKEISSTIERLSEISSTIAAAVEEQGAATQEIARNVQQAAQ----------- | *Bradyrhizobium sp. Y36* | Bacteria; Proteobacteria; Alphaproteobacteria; Rhizobiales; Bradyrhizobiaceae; Bradyrhizobium | methyl-accepting chemotaxis protein partial | 6 |
| | **WP_085581341.1** | 8.5E-09 | Qc.III | IEMIGHIIENINEIATTISAAVEEQGAATNEITRNVRNAADRTRMVSTSIT-- | *Thalassospira mesophila* | Bacteria; Proteobacteria; Alphaproteobacteria; Rhodospirillales; Rhodospirillaceae; Thalassospira | HAMP domain-containing protein | 6 |
| | **WP_108174471.1** | 8.5E-09 | Qc.III.c | -GEISATILRINEISTTIASAVEEQNAVTTEMSSNMRVAAEGVRAISGRM--- | *Phreatobacter oligotrophus* | Bacteria; Proteobacteria; Alphaproteobacteria; Phreatobacter | PAS domain S-box protein | 6 |
| | **WP_081944311.1** | 1.1E-08 | Qc.III | -GDIGRIIENINEIATTISAAVEEQGAATDEITRNVRGAADRTRSVSSSIT-- | *Thalassospira australica* | Bacteria; Proteobacteria; Alphaproteobacteria; Rhodospirillales; Rhodospirillaceae; Thalassospira | HAMP domain-containing protein | 6 |
| | **WP_022730340.1** | 1.2E-08 | Qc.III | -GDIGRIIENLNEIATTISAAVEEQGAATDEITRNVRGAADRTRSVSSSIT-- | *Thalassospira lucentensis* | Bacteria; Proteobacteria; Alphaproteobacteria; Rhodospirillales; Rhodospirillaceae; Thalassospira | HAMP domain-containing protein | 6 |
| | **WP_022730341.1** | 1.2E-08 | Qc.III | -GDIGRIIENLNEIATTISAAVEEQGAATDEITRNVRGAADRTRSVSSSIT-- | *Thalassospira lucentensis* | Bacteria; Proteobacteria; Alphaproteobacteria; Rhodospirillales; Rhodospirillaceae; Thalassospira | HAMP domain-containing protein | 6 |
|  | **WP_094454577.1** | 1.2E-08 | Qa.IV | --TIDRTIREMNQISGAIAAAVEQQGAATAEIARNVQMAAMAT---------- | *Niveispirillum lacus* | Bacteria; Proteobacteria; Alphaproteobacteria; Rhodospirillales; Rhodospirillaceae; Niveispirillum | hypothetical protein | 6 |
|  | **WP_054362215.1** | 1.3E-08 | Qa.IV | IQEIERTIAEISQISTSMAAAVEEQSAATGDIARTVQETATNT---------- | *Prosthecomicrobium hirschii* | Bacteria; Proteobacteria; Alphaproteobacteria; Rhizobiales; Hyphomicrobiaceae; Prosthecomicrobium | chemotaxis protein | 6 |
| | **WP_101267987.1** | 1.3E-08 | Qc.III | IEMIGAIIESINEIATTISAAVEEQGAATNEITRNVRNAAERTRMVSTSIS-- | *Thalassospira marina* | Bacteria; Proteobacteria; Alphaproteobacteria; Rhodospirillales; Rhodospirillaceae; Thalassospira | HAMP domain-containing protein | 6 |
| | **ONH87533.1** | 1.5E-08 | Qc.III | -GDIGRIIENINEIATTISAAVEEQGAATDEITRNVRGAADRTRSVSASIN-- | *Thalassospira sp. MCCC 1A02803* | Bacteria; Proteobacteria; Alphaproteobacteria; Rhodospirillales; Rhodospirillaceae; Thalassospira | chemotaxis protein | 6 |
|  | **WP_067559934.1** | 1.6E-08 | Qa.IV | -----NTIREINQISTTIASAVEEQGAATQEIARNVQQASVGTQQVNQNI--- | *Oceanibaculum pacificum* | Bacteria; Proteobacteria; Alphaproteobacteria; Rhodospirillales; Rhodospirillaceae; Oceanibaculum | methyl-accepting chemotaxis protein | 6 |
|  | **EKD37181.1** | 1.8E-08 | Qa.III | IDQIAKIIHSVNEIVSTIAVAVGEQSSATEEIARNISQAS------------ | *uncultured bacterium* | Bacteria; environmental samples | hypothetical protein ACD_75C01226G0003 | 6 |
|  | **SDQ94461.1** | 1.9E-08 | Qa.IV | INAITKTMADVNDITATIAAAVEQQGAATTEISENVQQAA------------ | *Pseudovibrio sp. Tun.PSC04-5.I4* | Bacteria; Proteobacteria; Alphaproteobacteria; Rhodobacterales; Rhodobacteraceae; Pseudovibrio | methyl-accepting chemotaxis sensory transducer with Cache sensor | 6 |
|  | **WP_028138419.1** | 2.2E-08 | Qa.IV | IKTIGSTIKSLSEISSTIAAAVEEQGAATQEIARNVQQAAQ----------- | *Bradyrhizobium japonicum* | Bacteria; Proteobacteria; Alphaproteobacteria; Rhizobiales; Bradyrhizobiaceae; Bradyrhizobium | PAS domain S-box protein | 6 |
|  | **WP_052268356.1** | 2.4E-08 | Qa.III | IEDIGRIIENINEIATTISAAVEEQGSATDEITRNVRSAADRT---------- | *Thalassospira xiamenensis* | Bacteria; Proteobacteria; Alphaproteobacteria; Rhodospirillales; Rhodospirillaceae; Thalassospira | HAMP domain-containing protein | 6 |
| | **KZD00728.1** | 2.6E-08 | Qc.III | -GDIGRIIENINEIATTISAAVEEQGAATDEITRNVRGAADRTRSVSASIN-- | *Thalassospira sp. MCCC 1A02898* | Bacteria; Proteobacteria; Alphaproteobacteria; Rhodospirillales; Rhodospirillaceae; Thalassospira | hypothetical protein AUQ41_04950 | 6 |
| | **KXJ57125.1** | 2.7E-08 | Qc.III | -GDIGRIIENINEIATTISAAVEEQGAATDEITRNVRGAADRTRSVSASIN-- | *Thalassospira sp. Nap_22* | Bacteria; Proteobacteria; Alphaproteobacteria; Rhodospirillales; Rhodospirillaceae; Thalassospira | hypothetical protein AXW12_06915 | 6 |
| | **WP_008891372.1** | 2.7E-08 | Qc.III | -GDIGRIIENINEIATTISAAVEEQGAATDEITRNVRGAADRTRSVSASIN-- | *Thalassospira* | Bacteria; Proteobacteria; Alphaproteobacteria; Rhodospirillales; Rhodospirillaceae | MULTISPECIES: HAMP domain-containing protein | 6 |
| | **WP_064790150.1** | 2.7E-08 | Qc.III | -GDIGRIIENINEIATTISAAVEEQGAATDEITRNVRGAADRTRSVSASIN-- | *Thalassospira profundimaris* | Bacteria; Proteobacteria; Alphaproteobacteria; Rhodospirillales; Rhodospirillaceae; Thalassospira | HAMP domain-containing protein | 6 |
| | **WP_082049696.1** | 2.7E-08 | Qc.III | -GDIGRIIENINEIATTISAAVEEQGAATDEITRNVRGAADRTRSVSASIN-- | *Thalassospira sp. HJ* | Bacteria; Proteobacteria; Alphaproteobacteria; Rhodospirillales; Rhodospirillaceae; Thalassospira | HAMP domain-containing protein | 6 |
| | **WP_083997020.1** | 2.7E-08 | Qc.III | -GDIGRIIENINEIATTISAAVEEQGAATDEITRNVRGAADRTRSVSASIN-- | *Thalassospira tepidiphila* | Bacteria; Proteobacteria; Alphaproteobacteria; Rhodospirillales; Rhodospirillaceae; Thalassospira | HAMP domain-containing protein | 6 |
| | **CAM77113.1** | 2.9E-08 | Qc.III | IEGMGGTISNINQIASAIAAAVEEQNAATQEISRNVQLAAEGTNSVSRNIG-- | *Magnetospirillum gryphiswaldense MSR-1* | Bacteria; Proteobacteria; Alphaproteobacteria; Rhodospirillales; Rhodospirillaceae; Magnetospirillum | Histidine kinase HAMP region:Bacterial chemotaxis sensory transducer | 6 |
| | **KZB70159.1** | 2.9E-08 | Qc.III | -GDIGRIIENINEIATTISAAVEEQGAATDEITRNVRGAADRTRSVSASI--- | *Thalassospira sp. MCCC 1A01148* | Bacteria; Proteobacteria; Alphaproteobacteria; Rhodospirillales; Rhodospirillaceae; Thalassospira | hypothetical protein AUQ43_14900 | 6 |
|  | **WP_029058275.1** | 2.9E-08 | Qa.IV | IREISRTIEQINQISGDIATSVGQQGEATTEIAMNVQEAT------------ | *Stappia stellulata* | Bacteria; Proteobacteria; Alphaproteobacteria; Rhodobacterales; Rhodobacteraceae; Stappia | methyl-accepting chemotaxis protein | 6 |
|  | **WP_109514941.1** | 2.9E-08 | Qa.III | IQSIANTINEVNEIAAAIAAAVQEQGSATNEIARNVQQAA------------ | *Roseomonas aestuarii* | Bacteria; Proteobacteria; Alphaproteobacteria; Rhodospirillales; Acetobacteraceae; Roseomonas | methyl-accepting chemotaxis protein | 6 |
|  | **WP_028482171.1** | 3E-08 | Qa.III | -GEIAKTMGNVNDITQSIAAAVEQQGAATTEISDNVQQAAE----------- | *Nesiotobacter exalbescens* | Bacteria; Proteobacteria; Alphaproteobacteria; Rhodobacterales; Rhodobacteraceae; Nesiotobacter | methyl-accepting chemotaxis protein | 6 |
| | **WP_068518121.1** | 3E-08 | Qc.III | -GDIGRIIENINEIATTISAAVEEQGAATDEITRNVRGAADRTRSVSASI--- | *Thalassospira* | Bacteria; Proteobacteria; Alphaproteobacteria; Rhodospirillales; Rhodospirillaceae | MULTISPECIES: methyl-accepting chemotaxis protein | 6 |
| | **WP_084272160.1** | 3E-08 | Qc.III | -GDIGRIIENINEIATTISAAVEEQGAATDEITRNVRGAADRTRSVSASI--- | *Thalassospira* | Bacteria; Proteobacteria; Alphaproteobacteria; Rhodospirillales; Rhodospirillaceae | MULTISPECIES: HAMP domain-containing protein | 6 |
| | **WP_114099838.1** | 3E-08 | Qc.III | -GDIGRIIENINEIATTISAAVEEQGAATDEITRNVRGAADRTRSVSASI--- | *Thalassospira profundimaris* | Bacteria; Proteobacteria; Alphaproteobacteria; Rhodospirillales; Rhodospirillaceae; Thalassospira | HAMP domain-containing protein | 6 |
| | **WP_114102360.1** | 3E-08 | Qc.III | -GDIGRIIENINEIATTISAAVEEQGAATDEITRNVRGAADRTRSVSASI--- | *Thalassospira profundimaris* | Bacteria; Proteobacteria; Alphaproteobacteria; Rhodospirillales; Rhodospirillaceae; Thalassospira | HAMP domain-containing protein | 6 |
| | **WP_114112606.1** | 3E-08 | Qc.III | -GDIGRIIENINEIATTISAAVEEQGAATDEITRNVRGAADRTRSVSASI--- | *Thalassospira profundimaris* | Bacteria; Proteobacteria; Alphaproteobacteria; Rhodospirillales; Rhodospirillaceae; Thalassospira | HAMP domain-containing protein | 6 |
|  | **EJN13705.1** | 3.1E-08 | Qa.IV | IREISSTIERLSEISSTVAAAVEEQGAATQEISRNVQQAAQ----------- | *Bradyrhizobium sp. YR681* | Bacteria; Proteobacteria; Alphaproteobacteria; Rhizobiales; Bradyrhizobiaceae; Bradyrhizobium | methyl-accepting chemotaxis protein partial | 6 |
|  | **WP_042340398.1** | 3.1E-08 | Qa.III | IQAIGGTIAEIDDISKEIATAVEQQGLATGEISGNVQQAAE----------- | *Bradyrhizobium oligotrophicum* | Bacteria; Proteobacteria; Alphaproteobacteria; Rhizobiales; Bradyrhizobiaceae; Bradyrhizobium | chemotaxis protein | 6 |
|  | **WP_097173782.1** | 3.1E-08 | Qa.IV | IRSITETIREMNIVTTAIASAVEQQGAATQEIARNVQSAA------------ | *Stappia indica* | Bacteria; Proteobacteria; Alphaproteobacteria; Rhodobacterales; Rhodobacteraceae; Stappia | methyl-accepting chemotaxis protein | 6 |
| | **WP_114102359.1** | 3.1E-08 | Qc.III | -GEIGAIIDNLNEIATTISAAVEEQGAATDEISRNVRGAADRTRAVSASI--- | *Thalassospira profundimaris* | Bacteria; Proteobacteria; Alphaproteobacteria; Rhodospirillales; Rhodospirillaceae; Thalassospira | HAMP domain-containing protein | 6 |
|  | **BAM92825.1** | 3.2E-08 | Qa.III | IQAIGGTIAEIDDISKEIATAVEQQGLATGEISGNVQQAAE----------- | *Bradyrhizobium oligotrophicum S58* | Bacteria; Proteobacteria; Alphaproteobacteria; Rhizobiales; Bradyrhizobiaceae; Bradyrhizobium | conserved membrane hypothetical protein | 6 |
|  | **PDT63924.1** | 3.2E-08 | Qa.IV | IKEISSTIERLSEISSAIAAAVEEQGAATQEIARNVQQAAQ----------- | *Bradyrhizobium ottawaense* | Bacteria; Proteobacteria; Alphaproteobacteria; Rhizobiales; Bradyrhizobiaceae; Bradyrhizobium | methyl-accepting chemotaxis protein partial | 6 |
|  | **SFJ85833.1** | 3.3E-08 | Qa.IV | IRDISATIERLSEVSSTIAAAVEEQGAATQEISRNVQQAA------------ | *Bradyrhizobium sp. cf659* | Bacteria; Proteobacteria; Alphaproteobacteria; Rhizobiales; Bradyrhizobiaceae; Bradyrhizobium | hypothetical protein SAMN04487925_112139 | 6 |
|  | **WP_035899298.1** | 3.3E-08 | Qa.IV | IQEITDTMEEVNEYTSTIAAAVEQQGAATNEISVNVQRAA------------ | *Labrenzia sp. DG1229* | Bacteria; Proteobacteria; Alphaproteobacteria; Rhodobacterales; Rhodobacteraceae; Labrenzia | HAMP domain-containing protein | 6 |
|  | **WP_104705408.1** | 3.3E-08 | Qa.IV | IQEITDTMEEVNEYTSTIAAAVEQQGAATNEISVNVQRAA------------ | *Labrenzia sp. EL143* | Bacteria; Proteobacteria; Alphaproteobacteria; Rhodobacterales; Rhodobacteraceae; Labrenzia | HAMP domain-containing protein | 6 |
|  | **WP_108875991.1** | 3.3E-08 | Qa.IV | IQEITDTMEEVNEYTSTIAAAVEQQGAATNEISVNVQRAA------------ | *Labrenzia sp. Alg231-36* | Bacteria; Proteobacteria; Alphaproteobacteria; Rhodobacterales; Rhodobacteraceae; Labrenzia | HAMP domain-containing protein | 6 |
| | **WP_085590579.1** | 3.4E-08 | Qc.III | IEMIGSIIESINEIATTISAAVEEQGAATNEITRNVRNAADRTRMVSTSIT-- | *Thalassospira sp. MCCC 1A01428* | Bacteria; Proteobacteria; Alphaproteobacteria; Rhodospirillales; Rhodospirillaceae; Thalassospira | HAMP domain-containing protein | 6 |
| | **SMF39594.1** | 3.8E-08 | Qc.III | -RSIGEVIGQMSEIATSISAAVEEQGAATDEIAQTMSRAADGTQTVAGNI--- | *Tistlia consotensis USBA 355* | Bacteria; Proteobacteria; Alphaproteobacteria; Rhodospirillales; Rhodospirillaceae; Tistlia | methyl-accepting chemotaxis protein partial | 6 |
| | **WP_073954017.1** | 4E-08 | Qc.III | IETIGSIIESINEIATTISAAVEEQGAATNEITRNVRNAAERTRMVSTSIS-- | *Thalassospira sp. TSL5-1* | Bacteria; Proteobacteria; Alphaproteobacteria; Rhodospirillales; Rhodospirillaceae; Thalassospira | HAMP domain-containing protein | 6 |
| | **WP_073009798.1** | 4.2E-08 | Qa.III | IEAIMKIIQNVNEISSGIAAAITEQSAVTRDISDNMQRAARETMAASSSL--- | *Labrenzia suaedae* | Bacteria; Proteobacteria; Alphaproteobacteria; Rhodobacterales; Rhodobacteraceae; Labrenzia | methyl-accepting chemotaxis protein | 6 |
|  | **WP_028319247.1** | 4.4E-08 | Qa.III | INQITKIINNVNEIVATISTAVGEQSSATEEIANNIAQAS------------ | *Desulfobulbus elongatus* | Bacteria; Proteobacteria; Deltaproteobacteria; Desulfobacterales; Desulfobulbaceae; Desulfobulbus | methyl-accepting chemotaxis protein | 6 |
| | **OYX01872.1** | 4.6E-08 | Qa.III | ITGIASTIGQMNEISNTIASAVSQQASATDEIANSIARASEGTRNANSNLEQ- | *Rhizobiales bacterium 32-66-8* | Bacteria; Proteobacteria; Alphaproteobacteria; Rhizobiales | hypothetical protein B7Z15_19310 partial | 6 |
|  | **WP_093809603.1** | 4.6E-08 | Qa.IV | IREISRTIEQINQISGDIAASVGQQGEATTEIAMNVQEAT------------ | *Stappia sp. ES.058* | Bacteria; Proteobacteria; Alphaproteobacteria; Rhodobacterales; Rhodobacteraceae; Stappia | methyl-accepting chemotaxis protein | 6 |
|  | **ACL60785.1** | 4.8E-08 | Qa.III | IQEIVRVIGEVSTIATTIAAAVEEQTAVTSEIAQNVQQTATNTQTAV------ | *Methylobacterium nodulans ORS 2060* | Bacteria; Proteobacteria; Alphaproteobacteria; Rhizobiales; Methylobacteriaceae; Methylobacterium | methyl-accepting chemotaxis sensory transducer | 6 |
|  | **OUX59732.1** | 4.8E-08 | Qa.IV | IKEISSTINLISEISSTIAAAVEEQGAATQEIARNVQQAAEL---------- | *Afipia sp. TMED4* | Bacteria; Proteobacteria; Alphaproteobacteria; Rhizobiales; Bradyrhizobiaceae; Afipia | methyl-accepting chemotaxis protein | 6 |
|  | **WP_006019103.1** | 4.8E-08 | Qa.IV | IKEISSTINLISEISSTIAAAVEEQGAATQEIARNVQQAAEL---------- | *Afipia broomeae* | Bacteria; Proteobacteria; Alphaproteobacteria; Rhizobiales; Bradyrhizobiaceae; Afipia | hypothetical protein | 6 |
|  | **WP_079566556.1** | 5.1E-08 | Qa.IV | IRSITNTISEINDVTTAIAAAVEEQGAATREIARNIQHAA------------ | *Bradyrhizobium erythrophlei* | Bacteria; Proteobacteria; Alphaproteobacteria; Rhizobiales; Bradyrhizobiaceae; Bradyrhizobium | HAMP domain-containing protein | 6 |
| | **WP_085909374.1** | 5.2E-08 | Qc.III | -GSISGTIGQINEIATAISTAVEEQSAATGEISSNVQQAHTGTRNVTSNIG-- | *Kiloniella majae* | Bacteria; Proteobacteria; Alphaproteobacteria; Kiloniellales; Kiloniellaceae; Kiloniella | methyl-accepting chemotaxis protein | 6 |
| | **WP_082049671.1** | 5.3E-08 | Qc.III | IQEISETINRISSIATAISSAVEEQGAATQEIARNVSESAGRSKEVTSGIQ-- | *Thalassospira sp. HJ* | Bacteria; Proteobacteria; Alphaproteobacteria; Rhodospirillales; Rhodospirillaceae; Thalassospira | methyl-accepting chemotaxis protein | 6 |
|  | **PIW26536.1** | 5.4E-08 | Qa.IV | IQSITTTISEMSEIAGTIAAAVEQQGAATSEIARNVEQAS------------ | *Rhodospirillales bacterium CG15_BIG_FIL_POST_REV_8_21_14_020_66_15* | Bacteria; Proteobacteria; Alphaproteobacteria; Rhodospirillales; unclassified Rhodospirillales | methyl-accepting chemotaxis protein | 6 |
| | **RCK38639.1** | 5.5E-08 | Qc.III | IEAIGSIIENINEIATTISAAIEQQGSATNEITRNVRGAADRTRAVSKSIND- | *Thalassospira profundimaris* | Bacteria; Proteobacteria; Alphaproteobacteria; Rhodospirillales; Rhodospirillaceae; Thalassospira | hypothetical protein TH25_25485 partial | 6 |
|  | **KPA19254.1** | 5.6E-08 | Qa.III.b | -------VEEINMIMNSIATAVEQQTETTDGISNNLTDSASSARSVSDNLQKA | *Candidatus Magnetomorum sp. HK-1* | Bacteria; Proteobacteria; Deltaproteobacteria; Desulfobacterales; Desulfobacteraceae; Candidatus Magnetomorum | methyl-accepting chemotaxis protein | 6 |

| | | | | | | | |
|---|---|---|---|---|---|---|---|
| WP_029065797.1 | 5.8E-08 | Qc.III | INSIMGVIKNIAEIATSISTAVEEQTAVTNDISQNMQSAAQGVDVVTTNLQ-- | *Labrenzia sp. DG1229* | Bacteria; Proteobacteria; Alphaproteobacteria; Rhodobacterales; Rhodobacteraceae; Labrenzia | methyl-accepting chemotaxis protein | 6 |
| OQW55538.1 | 5.9E-08 | Qa.IV | IKEISTTITLISEISSTIAAAVEEQGAATQEIARNVQHAAQRSTAVAE----- | *Proteobacteria bacterium SG_bin9* | Bacteria; Proteobacteria | hypothetical protein A4S14_12925 | 6 |
| WP_114090499.1 | 6.2E-08 | Qa.III | IDGITRIIHDLNEISAEVAAAVQQQGSATSEISNSVQQAVTSTQE-------- | *Thalassospira profundimaris* | Bacteria; Proteobacteria; Alphaproteobacteria; Rhodospirillales; Rhodospirillaceae; Thalassospira | HAMP domain-containing protein | 6 |
| WP_008970719.1 | 6.7E-08 | Qa.IV | IRAITGTIERLSEISSTIAAAVEQQGAATQEISRNVQQAAE------------ | *Bradyrhizobium sp. STM 3843* | Bacteria; Proteobacteria; Alphaproteobacteria; Rhizobiales; Bradyrhizobiaceae; Bradyrhizobium | methyl-accepting chemotaxis protein | 6 |
| WP_044538644.1 | 6.8E-08 | Qa.IV | IRDISSTIERLSEISSTIAAAVEEQGAATQEISRNVQQAAE------------ | *Bradyrhizobium sp. LTSP885* | Bacteria; Proteobacteria; Alphaproteobacteria; Rhizobiales; Bradyrhizobiaceae; Bradyrhizobium | methyl-accepting chemotaxis protein | 6 |
| WP_109327219.1 | 6.8E-08 | Qa.III | IVGIGRTIANINSIAGDIGVAIGQQSSAIREIVANMEQAAA------------ | *Azospirillum sp. CFH 70021* | Bacteria; Proteobacteria; Alphaproteobacteria; Rhodospirillales; Rhodospirillaceae; Azospirillum | chemotaxis protein | 6 |
| WP_052065390.1 | 7.2E-08 | Qc.III | -GEIGSIIDSLNEIATTISAAVEEQGAATDEITRNVRGAADRTRAVSASI--- | *Thalassospira australica* | Bacteria; Proteobacteria; Alphaproteobacteria; Rhodospirillales; Rhodospirillaceae; Thalassospira | HAMP domain-containing protein | 6 |
| WP_062957707.1 | 7.2E-08 | Qc.III | -GEIGSIIDSLNEIATTISAAVEEQGAATDEITRNVRGAADRTRAVSASI--- | *Thalassospira* | Bacteria; Proteobacteria; Alphaproteobacteria; Rhodospirillales; Rhodospirillaceae | MULTISPECIES: HAMP domain-containing protein | 6 |
| WP_044538645.1 | 7.6E-08 | Qa.IV | IKEISSTIERLSEISSTIAAAVEEQGAATQEISRNVQQAAQ------------ | *Bradyrhizobium sp. LTSP885* | Bacteria; Proteobacteria; Alphaproteobacteria; Rhizobiales; Bradyrhizobiaceae; Bradyrhizobium | methyl-accepting chemotaxis protein | 6 |
| WP_044588899.1 | 7.6E-08 | Qa.IV | IKEISSTIERLSEISSTIAAAVEEQGAATQEISRNVQQAAQ------------ | *Bradyrhizobium sp. LTSPM299* | Bacteria; Proteobacteria; Alphaproteobacteria; Rhizobiales; Bradyrhizobiaceae; Bradyrhizobium | methyl-accepting chemotaxis protein | 6 |
| WP_002714792.1 | 8E-08 | Qa.IV | IKEIGKTITLISEISQTIAAAVEEQGAATQEISRNVQQAA------------- | *Afipia clevelandensis* | Bacteria; Proteobacteria; Alphaproteobacteria; Rhizobiales; Bradyrhizobiaceae; Afipia | methyl-accepting chemotaxis protein | 6 |
| WP_009735413.1 | 8E-08 | Qa.IV | IKEIGKTITLISEISQTIAAAVEEQGAATQEISRNVQQAA------------- | *Bradyrhizobiaceae bacterium SG-6C* | Bacteria; Proteobacteria; Alphaproteobacteria; Rhizobiales; Bradyrhizobiaceae | methyl-accepting chemotaxis protein | 6 |
| WP_049974523.1 | 8E-08 | Qa.IV | -RRIGATIADVNELTVTIASAVEQQSAATQEIARNVQQAAQ------------ | *Azospirillum sp. B4* | Bacteria; Proteobacteria; Alphaproteobacteria; Rhodospirillales; Rhodospirillaceae; Azospirillum | methyl-accepting chemotaxis protein | 6 |
| WP_065730696.1 | 8.3E-08 | Qc.III | IKEISGTIQRLAEIASAIAAAVEEQGAATQEISRNVQQAAEGTHQVSANIT-- | *Bradyrhizobium icense* | Bacteria; Proteobacteria; Alphaproteobacteria; Rhizobiales; Bradyrhizobiaceae; Bradyrhizobium | methyl-accepting chemotaxis protein | 6 |
| PYE99017.1 | 8.5E-08 | Qa.IV | IKEIGQTISRMSEIASTIASAVEEQGAATQEISRNVQEAA------------- | *Rhodopseudomonas faecalis* | Bacteria; Proteobacteria; Alphaproteobacteria; Rhizobiales; Bradyrhizobiaceae; Rhodopseudomonas | hypothetical protein BJ122_1451 partial | 6 |
| WP_024576125.1 | 8.7E-08 | Qa.IV | IKEISSTINLISEISSTIAAAVEEQGAATQEIARNVQQAAEL----------- | *Afipia* | Bacteria; Proteobacteria; Alphaproteobacteria; Rhizobiales; Bradyrhizobiaceae | MULTISPECIES: chemotaxis protein | 6 |
| WP_097670513.1 | 8.7E-08 | Qa.IV | IKEISSTIERLSEISSTIAAAVEEQGAATQEISRNVQQAAQ------------ | *Bradyrhizobium sp. C9* | Bacteria; Proteobacteria; Alphaproteobacteria; Rhizobiales; Bradyrhizobiaceae; Bradyrhizobium | methyl-accepting chemotaxis protein | 6 |
| WP_085581345.1 | 8.8E-08 | Qc.III | IEMIGRIIENINEIATTISAAIEEQGAATNEITRNVRGAADRTRAVSSSIT-- | *Thalassospira mesophila* | Bacteria; Proteobacteria; Alphaproteobacteria; Rhodospirillales; Rhodospirillaceae; Thalassospira | HAMP domain-containing protein | 6 |
| PIT06569.1 | 9.1E-08 | Qa.IV | IKEISSTIERLSEISSAIAAAVEEQGAATQEIARNVQQAAQ------------ | *Bradyrhizobium sp. TSA1* | Bacteria; Proteobacteria; Alphaproteobacteria; Rhizobiales; Bradyrhizobiaceae; Bradyrhizobium | chemotaxis protein partial | 6 |
| WP_028164954.1 | 9.1E-08 | Qa.IV | IKEISNTIERLSEISSTIAAAVEEQGAATQEISRNVQQAAQ------------ | *Bradyrhizobium elkanii* | Bacteria; Proteobacteria; Alphaproteobacteria; Rhizobiales; Bradyrhizobiaceae; Bradyrhizobium | methyl-accepting chemotaxis protein | 6 |
| WP_073617116.1 | 9.1E-08 | Qa.III | ISEIKTIINDVNDIVTTIATAVEEQSVSTNEIATNVSQAA------------- | *Desulfopila aestuarii* | Bacteria; Proteobacteria; Deltaproteobacteria; Desulfobacterales; Desulfobulbaceae; Desulfopila | methyl-accepting chemotaxis protein | 6 |
| WP_029012845.1 | 9.3E-08 | Qa.III | --AIEGTIREIDGISSTIATAVGEQGEATREIARNVQNAATATAEVADN---- | *Niveispirillum irakense* | Bacteria; Proteobacteria; Alphaproteobacteria; Rhodospirillales; Rhodospirillaceae; Niveispirillum | hypothetical protein | 6 |
| WP_104701320.1 | 9.3E-08 | Qc.III | --SILGVIKNIAEIATSISSAVEEQTAVTNDISQNMQSAAQGVDVVTTNLQ-- | *Labrenzia sp. EL143* | Bacteria; Proteobacteria; Alphaproteobacteria; Rhodobacterales; Rhodobacteraceae; Labrenzia | methyl-accepting chemotaxis protein | 6 |
| OSM02529.1 | 9.5E-08 | Qc.III | --EVSASIARINEVSDQINGAVDQQAITAKAIGESVTRVTEATDTVTRNSKE- | *Magnetofaba australis IT-1* | Bacteria; Proteobacteria; Alphaproteobacteria; Magnetococcales; Magnetococcaceae; Magnetofaba | putative methyl-accepting chemotaxis sensory transducer | 6 |
| WP_066504387.1 | 9.5E-08 | Qa.IV | IKEISSTIERLSEISSTIAAAVEEQGAATQEISRNVQQAA------------- | *Bradyrhizobium sp. BR 10303* | Bacteria; Proteobacteria; Alphaproteobacteria; Rhizobiales; Bradyrhizobiaceae; Bradyrhizobium | methyl-accepting chemotaxis protein | 6 |
| WP_084627821.1 | 9.5E-08 | Qa.III | IHGIGRTISQLNEVASTISAAVEEQGAATNEIARNVQQAATSSATVN------ | *Elstera litoralis* | Bacteria; Proteobacteria; Alphaproteobacteria; Rhodospirillales; Rhodospirillaceae; Elstera | hypothetical protein | 6 |
| ASR44020.1 | 9.6E-08 | Qb.I | LQRTRDRLQATIEQSEASNEELKASNEELQSINEELRSATEELETS------- | *Xanthomonas citri pv. mangiferaeindicae* | Bacteria; Proteobacteria; Gammaproteobacteria; Xanthomonadales; Xanthomonadaceae; Xanthomonas | hypothetical protein BEN78_12200 | 6 |
| WP_020592951.1 | 9.6E-08 | Qa.IV | --KITGTIKQINEITASVAAAVEEQGAATQEISRNVQEAAAAT---------- | *Kiloniella laminariae* | Bacteria; Proteobacteria; Alphaproteobacteria; Kiloniellales; Kiloniellaceae; Kiloniella | HAMP domain-containing protein | 6 |
| KJV08874.1 | 9.7E-08 | Qa.III | IHGIGRTISQLNEVASTISAAVEEQGAATNEIARNVQQAATSSATVN------ | *Elstera litoralis* | Bacteria; Proteobacteria; Alphaproteobacteria; Rhodospirillales; Rhodospirillaceae; Elstera | hypothetical protein VZ95_14940 | 6 |
| KRQ00031.1 | 9.7E-08 | Qa.III | IRSICATIQQVDEISAAIAAAVGQQGTATQEIAQNVQQAA------------- | *Bradyrhizobium manausense* | Bacteria; Proteobacteria; Alphaproteobacteria; Rhizobiales; Bradyrhizobiaceae; Bradyrhizobium | chemotaxis protein | 6 |
| WP_025037116.1 | 9.9E-08 | Qc.III.c | --AISQTINRISEISTAIASAVEEQAAATKEVTTNISSTNQAVGQ-------- | *Bradyrhizobium sp. DOA9* | Bacteria; Proteobacteria; Alphaproteobacteria; Rhizobiales; Bradyrhizobiaceae; Bradyrhizobium | hypothetical protein | 6 |
| WP_020587626.1 | 1E-07 | Qa.IV | IQNITDVINQVNDIVSTIAMTVEEQTETIAEIDENVSQGANALQEVNANVAK- | *Desulfobacter curvatus* | Bacteria; Proteobacteria; Deltaproteobacteria; Desulfobacterales; Desulfobacteraceae; Desulfobacter | methyl-accepting chemotaxis protein | 6 |
| WP_024870635.1 | 1E-07 | Qa.III | LEQITKSIGDIQNMTHQIATAAEEQTSVIEDISSNLIDIKNT----------- | *Tolumonas lignilytica* | Bacteria; Proteobacteria; Gammaproteobacteria; Aeromonadales; Aeromonadaceae; Tolumonas | methyl-accepting chemotaxis protein | 6 |
| WP_027520028.1 | 1E-07 | Qa.III | IQSICATIQEVDEISAAIAAAVGQQGTATQEIAQNVQQAAARTSEVSQ----- | *Bradyrhizobium sp. WSM1417* | Bacteria; Proteobacteria; Alphaproteobacteria; Rhizobiales; Bradyrhizobiaceae; Bradyrhizobium | methyl-accepting chemotaxis protein | 6 |
| WP_085445383.1 | 1E-07 | Qc.III | --EVSASIARINEVSDQINGAVDQQAITAKAIGESVTRVTEATDTVTRNSKE- | *Magnetofaba australis* | Bacteria; Proteobacteria; Alphaproteobacteria; Magnetococcales; Magnetococcaceae; Magnetofaba | methyl-accepting chemotaxis protein | 6 |
| WP_092191809.1 | 1E-07 | Qa.III | -GQIVKVISDVNEIVTTIAAAIEEQSAVTREVATNINQATSGVQDA------- | *Desulfomicrobium norvegicum* | Bacteria; Proteobacteria; Deltaproteobacteria; Desulfovibrionales; Desulfomicrobiaceae; Desulfomicrobium | methyl-accepting chemotaxis protein | 6 |
| BAM90860.1 | 1.1E-07 | Qc.III | IKEIGSTIGRLSEIAAAVSAAVEEQGAATQEISRNVQHAANGTQRVSRNIDD- | *Bradyrhizobium oligotrophicum S58* | Bacteria; Proteobacteria; Alphaproteobacteria; Rhizobiales; Bradyrhizobiaceae; Bradyrhizobium | putative methyl-accepting chemotaxis protein | 6 |
| PYF04213.1 | 1.1E-07 | Qa.IV | IKEIGQTISRMSEIASTIASAVEEQGAATQEIARNVQEAA------------- | *Rhodopseudomonas faecalis* | Bacteria; Proteobacteria; Alphaproteobacteria; Rhizobiales; Bradyrhizobiaceae; Rhodopseudomonas | hypothetical protein BJ122_104193 partial | 6 |
| WP_042340181.1 | 1.1E-07 | Qc.III | IKEIGSTIGRLSEIAAAVSAAVEEQGAATQEISRNVQHAANGTQRVSRNIDD- | *Bradyrhizobium oligotrophicum* | Bacteria; Proteobacteria; Alphaproteobacteria; Rhizobiales; Bradyrhizobiaceae; Bradyrhizobium | HAMP domain-containing protein | 6 |
| WP_095410375.1 | 1.1E-07 | Qb.I | LQRTKEQLQSTIEQSETSTEELKASNEELQAINEELQAINEELRS-------- | *Burkholderia ubonensis* | Bacteria; Proteobacteria; Betaproteobacteria; Burkholderiales; Burkholderiaceae; Burkholderia; Burkholderia cepacia complex | PAS domain S-box protein | 6 |
| EDM64614.1 | 1.2E-07 | Qc.III | LDSMTNSVMQINDLGIQIATAAEEQSSVTEEINRNMTAIQSMVNKLTENGDK- | *Moritella sp. PE36* | Bacteria; Proteobacteria; Gammaproteobacteria; Alteromonadales; Moritellaceae; Moritella | hypothetical methyl-accepting chemotaxis protein | 6 |
| OIM93438.1 | 1.2E-07 | Qa.IV | IKEISSTIERLSEISSTIAAAVEEQGAATQEISRNVQQAAQ------------ | *Bradyrhizobium elkanii* | Bacteria; Proteobacteria; Alphaproteobacteria; Rhizobiales; Bradyrhizobiaceae; Bradyrhizobium | methyl-accepting chemotaxis protein | 6 |
| OUX59733.1 | 1.2E-07 | Qa.IV | IKEIGKTITLISEISSTIAAAVEEQGAATQEIARNVQQAA------------- | *Afipia sp. TMED4* | Bacteria; Proteobacteria; Alphaproteobacteria; Rhizobiales; Bradyrhizobiaceae; Afipia | methyl-accepting chemotaxis protein | 6 |
| PCJ56858.1 | 1.2E-07 | Qa.III | IKGIGTTIKEINEIAASIASAVEEQGAATSEISRNVQEAA------------- | *Rhodospirillaceae bacterium* | Bacteria; Proteobacteria; Alphaproteobacteria; Rhodospirillales; Rhodospirillaceae | hypothetical protein COA65_10130 partial | 6 |
| WP_018081677.1 | 1.2E-07 | Qa.IV | IEKIVTTISDIRSISTILASAVEQQGTATQEIAGNTAQAA------------- | *Asticcacaulis benevestitus* | Bacteria; Proteobacteria; Alphaproteobacteria; Caulobacterales; Caulobacteraceae; Asticcacaulis | methyl-accepting chemotaxis protein | 6 |
| WP_018270531.1 | 1.2E-07 | Qa.IV | IKEISSTIERLSEISSTIAAAVEEQGAATQEISRNVQQAAQ------------ | *Bradyrhizobium elkanii* | Bacteria; Proteobacteria; Alphaproteobacteria; Rhizobiales; Bradyrhizobiaceae; Bradyrhizobium | methyl-accepting chemotaxis protein | 6 |
| WP_021076189.1 | 1.2E-07 | Qa.IV | IKEISSTIERLSEISSTIAAAVEEQGAATQEISRNVQQAAQ------------ | *Bradyrhizobium sp. MOS004* | Bacteria; Proteobacteria; Alphaproteobacteria; Rhizobiales; Bradyrhizobiaceae; Bradyrhizobium | methyl-accepting chemotaxis protein | 6 |
| WP_022694479.1 | 1.2E-07 | Qc.III | -SEITGVIERINDISMSISGAVEEQAVVTQDISRNMSEATSAVNNI------- | *Ponticaulis koreensis* | Bacteria; Proteobacteria; Alphaproteobacteria; Rhodobacterales; Hyphomonadaceae; Ponticaulis | methyl-accepting chemotaxis protein | 6 |
| WP_024576124.1 | 1.2E-07 | Qa.IV | IKEIGKTITLISEISSTIAAAVEEQGAATQEIARNVQQAA------------- | *Afipia* | Bacteria; Proteobacteria; Alphaproteobacteria; Rhizobiales; Bradyrhizobiaceae | MULTISPECIES: methyl-accepting chemotaxis protein | 6 |
| WP_024921920.1 | 1.2E-07 | Qa.IV | IKEIGKTITLISEISSTIAAAVEEQGAATQEIARNVQQAA------------- | *Afipia* | Bacteria; Proteobacteria; Alphaproteobacteria; Rhizobiales; Bradyrhizobiaceae | MULTISPECIES: methyl-accepting chemotaxis protein | 6 |
| WP_028331780.1 | 1.2E-07 | Qa.IV | IKEISSTIERLSEISSTIAAAVEEQGAATQEISRNVQQAAQ------------ | *Bradyrhizobium elkanii* | Bacteria; Proteobacteria; Alphaproteobacteria; Rhizobiales; Bradyrhizobiaceae; Bradyrhizobium | methyl-accepting chemotaxis protein | 6 |
| WP_028338756.1 | 1.2E-07 | Qa.IV | IKEISSTIERLSEISSTIAAAVEEQGAATQEISRNVQQAAQ------------ | *Bradyrhizobium elkanii* | Bacteria; Proteobacteria; Alphaproteobacteria; Rhizobiales; Bradyrhizobiaceae; Bradyrhizobium | methyl-accepting chemotaxis protein | 6 |
| WP_029061297.1 | 1.2E-07 | Qc.III | --SIMGVIKNIAEIATSISTAVEEQTAVTNDISQNMQSAAQGVDVVTTNLQ-- | *Labrenzia sp. DG1229* | Bacteria; Proteobacteria; Alphaproteobacteria; Rhodobacterales; Rhodobacteraceae; Labrenzia | methyl-accepting chemotaxis protein | 6 |
| WP_038379440.1 | 1.2E-07 | Qa.IV | IKEISSTIERLSEISSTIAAAVEEQGAATQEISRNVQQAAQ------------ | *Bradyrhizobium elkanii* | Bacteria; Proteobacteria; Alphaproteobacteria; Rhizobiales; Bradyrhizobiaceae; Bradyrhizobium | methyl-accepting chemotaxis protein | 6 |
| WP_038387995.1 | 1.2E-07 | Qa.IV | IKEISSTIERLSEISSTIAAAVEEQGAATQEISRNVQQAAQ------------ | *Bradyrhizobium elkanii* | Bacteria; Proteobacteria; Alphaproteobacteria; Rhizobiales; Bradyrhizobiaceae; Bradyrhizobium | methyl-accepting chemotaxis protein | 6 |
| WP_050406773.1 | 1.2E-07 | Qa.IV | IKEISSTIERLSEISSTIAAAVEEQGAATQEISRNVQQAAQ------------ | *Bradyrhizobium embrapense* | Bacteria; Proteobacteria; Alphaproteobacteria; Rhizobiales; Bradyrhizobiaceae; Bradyrhizobium | methyl-accepting chemotaxis protein | 6 |
| WP_055114211.1 | 1.2E-07 | Qc.III | --SIMGVIKNIAEIATSISTAVEEQTAVTNDISQNMQSAAQGVDVVTTNLQ-- | *Labrenzia* | Bacteria; Proteobacteria; Alphaproteobacteria; Rhodobacterales; Rhodobacteraceae | MULTISPECIES: methyl-accepting chemotaxis protein | 6 |
| WP_057017651.1 | 1.2E-07 | Qa.IV | IKEISSTIERLSEISSTIAAAVEEQGAATQEISRNVQQAAQ------------ | *Bradyrhizobium* | Bacteria; Proteobacteria; Alphaproteobacteria; Rhizobiales; Bradyrhizobiaceae | MULTISPECIES: methyl-accepting chemotaxis protein | 6 |
| WP_069276194.1 | 1.2E-07 | Qa.IV | IKEISSTIERLSEISSTIAAAVEEQGAATQEISRNVQQAAQ------------ | *Bradyrhizobium elkanii* | Bacteria; Proteobacteria; Alphaproteobacteria; Rhizobiales; Bradyrhizobiaceae; Bradyrhizobium | methyl-accepting chemotaxis protein | 6 |

| | | | | | | | | |
|---|---|---|---|---|---|---|---|---|
|  | WP_076862468.1 | 1.2E-07 | Qa.IV | IKEISSTIERLSEISSTIAAAVEEQGAATQEISRNVQQAAQ------------ | *Bradyrhizobium mercantei* | Bacteria; Proteobacteria; Alphaproteobacteria; Rhizobiales; Bradyrhizobiaceae; Bradyrhizobium | methyl-accepting chemotaxis protein | 6 |
| | WP_085899826.1 | 1.2E-07 | Qc.III | -GSISGTIGQINEIATAISTAVEEQSAATGEISSNVQQAHTGTQNVTSNIG-- | *Kiloniella majae* | Bacteria; Proteobacteria; Alphaproteobacteria; Kiloniellales; Kiloniellaceae; Kiloniella | methyl-accepting chemotaxis protein | 6 |
| | WP_085905337.1 | 1.2E-07 | Qc.III | -GSISGTIGQINEIATAISTAVEEQSAATGEISSNVQQAHTGTQNVTSNIG-- | *Kiloniella majae* | Bacteria; Proteobacteria; Alphaproteobacteria; Kiloniellales; Kiloniellaceae; Kiloniella | methyl-accepting chemotaxis protein | 6 |
|  | WP_092083291.1 | 1.2E-07 | Qa.IV | IKEISSTIERLSEISSTIAAAVEEQGAATQEISRNVQQAAQ------------ | *Bradyrhizobium sp. R5* | Bacteria; Proteobacteria; Alphaproteobacteria; Rhizobiales; Bradyrhizobiaceae; Bradyrhizobium | methyl-accepting chemotaxis protein | 6 |
|  | WP_092125055.1 | 1.2E-07 | Qa.IV | IKEISSTIERLSEISSTIAAAVEEQGAATQEISRNVQQAAQ------------ | *Bradyrhizobium erythrophlei* | Bacteria; Proteobacteria; Alphaproteobacteria; Rhizobiales; Bradyrhizobiaceae; Bradyrhizobium | methyl-accepting chemotaxis protein | 6 |
|  | WP_092681190.1 | 1.2E-07 | Qa.IV | IKEISATINGISEISSTIASAVEEQGAATQEISRNVQQAAQ------------ | *Rhodopseudomonas pseudopalustris* | Bacteria; Proteobacteria; Alphaproteobacteria; Rhizobiales; Bradyrhizobiaceae; Rhodopseudomonas | methyl-accepting chemotaxis protein | 6 |
|  | WP_106329021.1 | 1.2E-07 | Qa.IV | IKEISSTIERLSEISSTIAAAVEEQGAATQEISRNVQQAAQ------------ | *Rhizobium tropici* | Bacteria; Proteobacteria; Alphaproteobacteria; Rhizobiales; Rhizobiaceae; Rhizobium/Agrobacterium group; Rhizobium | methyl-accepting chemotaxis protein | 6 |
| | WP_114102358.1 | 1.2E-07 | Qc.III | -GEIGSIIDNLNEIATSISAAVEQQGAATDEISRNVRGAADRTRAVSASI--- | *Thalassospira profundimaris* | Bacteria; Proteobacteria; Alphaproteobacteria; Rhodospirillales; Rhodospirillaceae; Thalassospira | HAMP domain-containing protein | 6 |
|  | SDH86654.1 | 1.3E-07 | Qa.IV | IKAISSTIERLSEISSTIAAAVEEQGAATQEISRNVQQAAQ------------ | *Bradyrhizobium ottawaense* | Bacteria; Proteobacteria; Alphaproteobacteria; Rhizobiales; Bradyrhizobiaceae; Bradyrhizobium | methyl-accepting chemotaxis sensory transducer | 6 |
|  | WP_006019102.1 | 1.3E-07 | Qa.IV | IKEIGKTITLISEISSTIAAAVEEQGAATQEIARNVQQAA------------- | *Afipia broomeae* | Bacteria; Proteobacteria; Alphaproteobacteria; Rhizobiales; Bradyrhizobiaceae; Afipia | methyl-accepting chemotaxis protein | 6 |
| | WP_009027997.1 | 1.3E-07 | Qc.III | IKEIGSTIGRLSEISAAVSAAVEEQGAATQEISRNVQHAANGTHRVSRNIDD- | *Bradyrhizobium sp. ORS 375* | Bacteria; Proteobacteria; Alphaproteobacteria; Rhizobiales; Bradyrhizobiaceae; Bradyrhizobium | methyl-accepting chemotaxis protein | 6 |
|  | WP_074831849.1 | 1.3E-07 | Qa.IV | IKAISSTIERLSEISSTIAAAVEEQGAATQEISRNVQQAAQ------------ | *Bradyrhizobium* | Bacteria; Proteobacteria; Alphaproteobacteria; Rhizobiales; Bradyrhizobiaceae | MULTISPECIES: methyl-accepting chemotaxis protein | 6 |
|  | WP_075710850.1 | 1.3E-07 | Qa.III | LDKITHSIQEISEMASQISAAASEQRTVTEEVSKNIQQAN------------- | *Vibrio panuliri* | Bacteria; Proteobacteria; Gammaproteobacteria; Vibrionales; Vibrionaceae; Vibrio | methyl-accepting chemotaxis protein | 6 |
|  | WP_075715123.1 | 1.3E-07 | Qa.III | LDKITHSIQEISEMASQISAAASEQRTVTEEVSKNIQQAN------------- | *Vibrio panuliri* | Bacteria; Proteobacteria; Gammaproteobacteria; Vibrionales; Vibrionaceae; Vibrio | methyl-accepting chemotaxis protein | 6 |
|  | WP_081628435.1 | 1.3E-07 | Qa.III.b | IREIVGVIGEVNALAGDISDAVSQQGEATREIARNVTEAAQGTQEVAES---- | *Novispirillum itersonii* | Bacteria; Proteobacteria; Alphaproteobacteria; Rhodospirillales; Rhodospirillaceae; Novispirillum | methyl-accepting chemotaxis protein | 6 |
|  | WP_090070721.1 | 1.3E-07 | Qa.IV | IRAITQTMTEVGSTSNTIAAAVEEQGSATENISENVQRAA------------- | *Cohaesibacter marisflavi* | Bacteria; Proteobacteria; Alphaproteobacteria; Rhizobiales; Cohaesibacteraceae | methyl-accepting chemotaxis protein | 6 |
|  | AUC95903.1 | 1.4E-07 | Qa.IV | IKEISTTIERLSEISSTIAAAVEEQGAATQEISRNVQQAAQ------------ | *Bradyrhizobium sp. SK17* | Bacteria; Proteobacteria; Alphaproteobacteria; Rhizobiales; Bradyrhizobiaceae; Bradyrhizobium | methyl-accepting chemotaxis protein | 6 |
|  | BAR58224.1 | 1.4E-07 | Qa.III | IQSICATIQQVDEISAAIAAAVGQQGTATQEIAQNVQQAA------------- | *Bradyrhizobium diazoefficiens* | Bacteria; Proteobacteria; Alphaproteobacteria; Rhizobiales; Bradyrhizobiaceae; Bradyrhizobium | putative methyl accepting chemotaxis protein | 6 |
|  | KGJ63535.1 | 1.4E-07 | Qa.III | IQSICATIQQVDEISAAIAAAVGQQGTATQEIAQNVQQAA------------- | *Bradyrhizobium diazoefficiens SEMIA 5080* | Bacteria; Proteobacteria; Alphaproteobacteria; Rhizobiales; Bradyrhizobiaceae; Bradyrhizobium | putative methyl accepting chemotaxis protein | 6 |
|  | SCB29332.1 | 1.4E-07 | Qa.III | IQSICATIQQVDEISAAIAAAVGQQGTATQEIAQNVQQAA------------- | *Bradyrhizobium shewense* | Bacteria; Proteobacteria; Alphaproteobacteria; Rhizobiales; Bradyrhizobiaceae; Bradyrhizobium | methyl-accepting chemotaxis protein | 6 |
|  | SFI69764.1 | 1.4E-07 | Qa.III | IQSICATIQQVDEISAAIAAAVGQQGTATQEIAQNVQQAA------------- | *Bradyrhizobium sp. cf659* | Bacteria; Proteobacteria; Alphaproteobacteria; Rhizobiales; Bradyrhizobiaceae; Bradyrhizobium | methyl-accepting chemotaxis protein | 6 |
|  | SFM80674.1 | 1.4E-07 | Qa.III | IQSICATIQQVDEISAAIAAAVGQQGTATQEIAQNVQQAA------------- | *Bradyrhizobium sp. Rc3b* | Bacteria; Proteobacteria; Alphaproteobacteria; Rhizobiales; Bradyrhizobiaceae; Bradyrhizobium | methyl-accepting chemotaxis sensory transducer | 6 |
|  | WP_014492382.1 | 1.4E-07 | Qa.III | IQSICATIQQVDEISAAIAAAVGQQGTATQEIAQNVQQAA------------- | *Bradyrhizobium japonicum* | Bacteria; Proteobacteria; Alphaproteobacteria; Rhizobiales; Bradyrhizobiaceae; Bradyrhizobium | methyl-accepting chemotaxis protein | 6 |
|  | WP_019195974.1 | 1.4E-07 | Qa.IV | IKEIGKTITLISEISSTIAAAVEEQGAATQEIARNVQQAA------------- | *Bradyrhizobiaceae* | Bacteria; Proteobacteria; Alphaproteobacteria; Rhizobiales | MULTISPECIES: methyl-accepting chemotaxis protein | 6 |
|  | WP_028140341.1 | 1.4E-07 | Qa.III | IQSICATIQQVDEISAAIAAAVGQQGTATQEIAQNVQQAA------------- | *Bradyrhizobium japonicum* | Bacteria; Proteobacteria; Alphaproteobacteria; Rhizobiales; Bradyrhizobiaceae; Bradyrhizobium | methyl-accepting chemotaxis protein | 6 |
|  | WP_028161098.1 | 1.4E-07 | Qa.III | IQSICATIQQVDEISAAIAAAVGQQGTATQEIAQNVQQAA------------- | *Bradyrhizobium japonicum* | Bacteria; Proteobacteria; Alphaproteobacteria; Rhizobiales; Bradyrhizobiaceae; Bradyrhizobium | methyl-accepting chemotaxis protein | 6 |
|  | WP_038932323.1 | 1.4E-07 | Qa.III | IQSICATIQQVDEISAAIAAAVGQQGTATQEIAQNVQQAA------------- | *Bradyrhizobium japonicum* | Bacteria; Proteobacteria; Alphaproteobacteria; Rhizobiales; Bradyrhizobiaceae; Bradyrhizobium | methyl-accepting chemotaxis protein | 6 |
|  | WP_041953837.1 | 1.4E-07 | Qa.III | IQSICATIQQVDEISAAIAAAVGQQGTATQEIAQNVQQAA------------- | *Bradyrhizobium japonicum* | Bacteria; Proteobacteria; Alphaproteobacteria; Rhizobiales; Bradyrhizobiaceae; Bradyrhizobium | methyl-accepting chemotaxis protein | 6 |
|  | WP_044562653.1 | 1.4E-07 | Qa.III | ITSIASTIVRINEIASTIASAVEEQGAATREIASNVQQAA------------- | *Azospirillum sp. B4* | Bacteria; Proteobacteria; Alphaproteobacteria; Rhodospirillales; Rhodospirillaceae; Azospirillum | hypothetical protein | 6 |
|  | WP_045002695.1 | 1.4E-07 | Qa.III | IQSICATIQQVDEISAAIAAAVGQQGTATQEIAQNVQQAA------------- | *Bradyrhizobium sp. LTSP857* | Bacteria; Proteobacteria; Alphaproteobacteria; Rhizobiales; Bradyrhizobiaceae; Bradyrhizobium | methyl-accepting chemotaxis protein | 6 |
|  | WP_046829031.1 | 1.4E-07 | Qa.IV | IKEIGKTITLISEISSTIAAAVEEQGAATQEISRNVQQAA------------- | *Afipia massiliensis* | Bacteria; Proteobacteria; Alphaproteobacteria; Rhizobiales; Bradyrhizobiaceae; Afipia | methyl-accepting chemotaxis protein | 6 |
|  | WP_046829032.1 | 1.4E-07 | Qa.IV | IKEIGKTITLISEISSTIAAAVEEQGAATQEISRNVQQAA------------- | *Afipia massiliensis* | Bacteria; Proteobacteria; Alphaproteobacteria; Rhizobiales; Bradyrhizobiaceae; Afipia | methyl-accepting chemotaxis protein | 6 |
|  | WP_073956858.1 | 1.4E-07 | Qa.III | IQSISATITEINEIASSIASAIEEQGSATEEITRNVQQAS------------- | *Thalassospira sp. TSL5-1* | Bacteria; Proteobacteria; Alphaproteobacteria; Rhodospirillales; Rhodospirillaceae; Thalassospira | methyl-accepting chemotaxis protein | 6 |
|  | WP_092034571.1 | 1.4E-07 | Qa.III | IQSICATIQQVDEISAAIAAAVGQQGTATQEIAQNVQQAA------------- | *Bradyrhizobium sp. OK095* | Bacteria; Proteobacteria; Alphaproteobacteria; Rhizobiales; Bradyrhizobiaceae; Bradyrhizobium | methyl-accepting chemotaxis protein | 6 |
|  | WP_092376632.1 | 1.4E-07 | Qa.III | -GQIVKVISDVNEIVTTIAAAIEEQSAVTREVATNINQATTGVQDA------- | *Desulfomicrobium apsheronum* | Bacteria; Proteobacteria; Deltaproteobacteria; Desulfovibrionales; Desulfomicrobiaceae; Desulfomicrobium | methyl-accepting chemotaxis protein | 6 |
|  | NP_773813.1 | 1.5E-07 | Qa.III | IQSICATIQQVDEISAAIAAAVGQQGTATQEIAQNVQQAA------------- | *Bradyrhizobium diazoefficiens USDA 110* | Bacteria; Proteobacteria; Alphaproteobacteria; Rhizobiales; Bradyrhizobiaceae; Bradyrhizobium | methyl accepting chemotaxis protein | 6 |
| | OYY58313.1 | 1.5E-07 | Qc.III | IERVGTQIRKMSDVANEISAAVEEQGMATKEIVRNVDQAATGTNSVTSHISD- | *Rhizobiales bacterium 35-66-30* | Bacteria; Proteobacteria; Alphaproteobacteria; Rhizobiales | hypothetical protein B7Y61_24810 | 6 |
| | PIW26489.1 | 1.5E-07 | Qc.III | --GISETIHRISEIAAAISAAVEEQGAATSEIARNVEQAATGTQEVSENI--- | *Rhodospirillales bacterium CG15_BIG_FIL_POST_REV_8_21_14_020_66_15* | Bacteria; Proteobacteria; Alphaproteobacteria; Rhodospirillales; unclassified Rhodospirillales | methyl-accepting chemotaxis protein | 6 |
|  | PKN41917.1 | 1.5E-07 | Qa.III | -GQIVKVISDVNEIVTTIAAAIEEQSAVTREVATNINQATTGVQDA------- | *Deltaproteobacteria bacterium HGW-Deltaproteobacteria-18* | Bacteria; Proteobacteria; Deltaproteobacteria | chemotaxis protein | 6 |
|  | WP_028480039.1 | 1.5E-07 | Qa.IV | --AITEKIEEVSQVSSTIASSVEQQGAATNEISMNIQSASQY----------- | *Nesiotobacter exalbescens* | Bacteria; Proteobacteria; Alphaproteobacteria; Rhodobacterales; Rhodobacteraceae; Nesiotobacter | chemotaxis protein | 6 |
|  | WP_047190158.1 | 1.5E-07 | Qa.IV | IQEIAQTITEMAQISTGIAAAVEEQGAATREISRNVQEAAR------------ | *Microvirga vignae* | Bacteria; Proteobacteria; Alphaproteobacteria; Rhizobiales; Methylobacteriaceae; Microvirga | methyl-accepting chemotaxis protein | 6 |
|  | WP_084251805.1 | 1.5E-07 | Qa.IV | IKEISSTIERLSEISSTIAAAVEEQGAATQEIARNVQQAAQ------------ | *Bradyrhizobium sp. AT1* | Bacteria; Proteobacteria; Alphaproteobacteria; Rhizobiales; Bradyrhizobiaceae; Bradyrhizobium | methyl-accepting chemotaxis protein partial | 6 |
|  | WP_099554299.1 | 1.5E-07 | Qa.IV | IQSIATTIGDINAISSSIASAVEQQNASTDEISRNVEEAS------------- | *Hartmannibacter diazotrophicus* | Bacteria; Proteobacteria; Alphaproteobacteria; Rhizobiales; Hartmannibacter | methyl-accepting chemotaxis protein | 6 |
|  | OFA09263.1 | 1.6E-07 | Qb.I | LRRTKAQLQDTVEHSETSNEELKASNEELQAINEELRSATEELETS------- | *Duganella phyllosphaerae* | Bacteria; Proteobacteria; Betaproteobacteria; Burkholderiales; Oxalobacteraceae; Duganella | aerobic respiration control sensor protein ArcB | 6 |
|  | OSJ26377.1 | 1.6E-07 | Qa.III | IQSICATIQEVDEISAAIAAAVGQQGTATQEIAQNVQQAA------------- | *Bradyrhizobium canariense* | Bacteria; Proteobacteria; Alphaproteobacteria; Rhizobiales; Bradyrhizobiaceae; Bradyrhizobium | methyl-accepting chemotaxis protein | 6 |
|  | OSQ31397.1 | 1.6E-07 | Qa.III | --DIVGTIEKVNEIASSIAAAVEEQQVATGEIARNVEEAARGTQD-------- | *Thalassospira sp. MCCC 1A01428* | Bacteria; Proteobacteria; Alphaproteobacteria; Rhodospirillales; Rhodospirillaceae; Thalassospira | chemotaxis protein partial | 6 |
|  | PDT64136.1 | 1.6E-07 | Qa.IV | IKEISSTIERLSEISSAIAAAVEEQGAATQEIARNVQQAAQ------------ | *Bradyrhizobium ottawaense* | Bacteria; Proteobacteria; Alphaproteobacteria; Rhizobiales; Bradyrhizobiaceae; Bradyrhizobium | methyl-accepting chemotaxis protein partial | 6 |
|  | SFO45274.1 | 1.6E-07 | Qa.III | IQSICATIREVDEISAAIAAAVGQQGTATQEIAQNVQQAA------------- | *Bradyrhizobium sp. Ghvi* | Bacteria; Proteobacteria; Alphaproteobacteria; Rhizobiales; Bradyrhizobiaceae; Bradyrhizobium | methyl-accepting chemotaxis sensory transducer | 6 |
|  | WP_015668127.1 | 1.6E-07 | Qa.IV | IREISSTIERLSEISSTIAAAVEEQGAATQEISRNVQQAA------------- | *Bradyrhizobium oligotrophicum* | Bacteria; Proteobacteria; Alphaproteobacteria; Rhizobiales; Bradyrhizobiaceae; Bradyrhizobium | HAMP domain-containing protein | 6 |
| | WP_024082201.1 | 1.6E-07 | Qc.III | IEGMGGTISNINQIASAIAAAVEEQNAATQEISRNVQLAAEGTNSVSRNIG-- | *Magnetospirillum gryphiswaldense* | Bacteria; Proteobacteria; Alphaproteobacteria; Rhodospirillales; Rhodospirillaceae; Magnetospirillum | methyl-accepting chemotaxis protein | 6 |
|  | WP_027570321.1 | 1.6E-07 | Qa.III | IQSICATIQEVDEISAAIAAAVGQQGTATQEIAQNVQQAA------------- | *Bradyrhizobium sp. URHA0013* | Bacteria; Proteobacteria; Alphaproteobacteria; Rhizobiales; Bradyrhizobiaceae; Bradyrhizobium | methyl-accepting chemotaxis protein | 6 |
| | WP_046504776.1 | 1.6E-07 | Qc.III | IQSISGTIKSIHEVATAIASAVEEQGAATSEITMSVQQAASGTDQVTSNI--- | *Kiloniella litopenaei* | Bacteria; Proteobacteria; Alphaproteobacteria; Kiloniellales; Kiloniellaceae; Kiloniella | hypothetical protein | 6 |
|  | WP_061029167.1 | 1.6E-07 | Qa.IV | IREISNTIERLSEISSTVAAAVEEQGAATQEISRNVQQAA------------- | *Bradyrhizobium sp. CCH5-F6* | Bacteria; Proteobacteria; Alphaproteobacteria; Rhizobiales; Bradyrhizobiaceae; Bradyrhizobium | methyl-accepting chemotaxis protein | 6 |
|  | WP_063994759.1 | 1.6E-07 | Qa.III | IQSICATIQEVDEISAAIAAAVGQQGTATQEIAQNVQQAA------------- | *Bradyrhizobium sp.* | Bacteria; Proteobacteria; Alphaproteobacteria; Rhizobiales; Bradyrhizobiaceae; Bradyrhizobium | methyl-accepting chemotaxis protein | 6 |
|  | WP_070245720.1 | 1.6E-07 | Qb.I | LRRTKAQLQDTVEHSETSNEELKASNEELQAINEELRSATEELETS------- | *Duganella phyllosphaerae* | Bacteria; Proteobacteria; Betaproteobacteria; Burkholderiales; Oxalobacteraceae; Duganella | PAS domain S-box protein | 6 |
| | WP_106002580.1 | 1.6E-07 | Qc.III | IEGMGGTISNINQIASAIAAAVEEQNAATQEISRNVQLAAEGTNSVSRNIG-- | *Magnetospirillum gryphiswaldense* | Bacteria; Proteobacteria; Alphaproteobacteria; Rhodospirillales; Rhodospirillaceae; Magnetospirillum | methyl-accepting chemotaxis protein | 6 |
| | EEA92923.1 | 1.7E-07 | Qc.III.c | ---IGETIERINSLSTGINEAVSEQRQATDDISRNLNSAHQNVSQVASNMQ-- | *Pseudovibrio sp. JE062* | Bacteria; Proteobacteria; Alphaproteobacteria; Rhodobacterales; Rhodobacteraceae; Pseudovibrio | methyl-accepting chemotaxis receptor/ sensory transducer | 6 |
|  | PHR62999.1 | 1.7E-07 | Qc.III | IEGISKTIGQINEYSAAISAAMEEQTAVTSDMSANMQTAASNVDNITH----- | *Robiginitomaculum sp.* | Bacteria; Proteobacteria; Alphaproteobacteria; Rhodobacterales; Hyphomonadaceae; Robiginitomaculum | chemotaxis protein | 6 |

| | | | | | | | | |
|---|---|---|---|---|---|---|---|---|
| | WP_026987121.1 | 1.7E-07 | Qc.III | IQGISNVISEMNEISTSISAAVEEQGAATDEIANTMNRVSDGTQKVNQNIEE- | *Fodinicurvata fenggangensis* | Bacteria; Proteobacteria; Alphaproteobacteria; Rhodospirillales; Rhodospirillaceae; Fodinicurvata | HAMP domain-containing protein | 6 |
| | WP_083417429.1 | 1.7E-07 | Qc.III.c | ---IGETIERINSLSTGINEAVSEQRQATDDISRNLNSAHQNVSQVASNMQ-- | *Pseudovibrio denitrificans* | Bacteria; Proteobacteria; Alphaproteobacteria; Rhodobacterales; Rhodobacteraceae; Pseudovibrio | methyl-accepting chemotaxis protein | 6 |
| | WP_083646136.1 | 1.7E-07 | Qc.III.c | ---IGETIERINSLSTGINEAVSEQRQATDDISRNLNSAHQNVSQVASNMQ-- | *Pseudovibrio sp. Ab134* | Bacteria; Proteobacteria; Alphaproteobacteria; Rhodobacterales; Rhodobacteraceae; Pseudovibrio | methyl-accepting chemotaxis protein | 6 |
| | AEV39294.1 | 1.8E-07 | Qc.III.c | ---IGETIERINSLSTGINEAVSEQRQATDDISRNLNSAHQNVSQVASNMQ-- | *Pseudovibrio sp. FO-BEG1* | Bacteria; Proteobacteria; Alphaproteobacteria; Rhodobacterales; Rhodobacteraceae; Pseudovibrio | methyl-accepting chemotaxis receptor/ sensory transducer | 6 |
| | KXJ57126.1 | 1.8E-07 | Qc.III | -GEIGSIIDSLNEIATTISAAVEQQGAATDEITRNVRGAADRTRAVSASI--- | *Thalassospira sp. Nap_22* | Bacteria; Proteobacteria; Alphaproteobacteria; Rhodospirillales; Rhodospirillaceae; Thalassospira | hypothetical protein AXW12_06920 | 6 |
|  | KYG97161.1 | 1.8E-07 | Qa.IV | IKEISSTIERLSEISSAIAAAVEEQGAATQEIARNVQQAAQ------------ | *Bradyrhizobium sp. DOA1* | Bacteria; Proteobacteria; Alphaproteobacteria; Rhizobiales; Bradyrhizobiaceae; Bradyrhizobium | chemotaxis protein partial | 6 |
|  | KYH00694.1 | 1.8E-07 | Qa.IV | IKEISSTIERLSEISSAIAAAVEEQGAATQEIARNVQQAAQ------------ | *Bradyrhizobium sp. DOA1* | Bacteria; Proteobacteria; Alphaproteobacteria; Rhizobiales; Bradyrhizobiaceae; Bradyrhizobium | chemotaxis protein partial | 6 |
| | OYW42554.1 | 1.8E-07 | Qc.III | IERVGTQIRKMSDVANEISAAVEEQGMATKEIVRNVDQAATGTNSVTSHISD- | *Azorhizobium sp. 12-66-6* | Bacteria; Proteobacteria; Alphaproteobacteria; Rhizobiales; Xanthobacteraceae; Azorhizobium | hypothetical protein B7Z45_01925 partial | 6 |
|  | PZP23743.1 | 1.8E-07 | Qb.I | LQRTKQQLQATIEQSETSTEELKASNEELQAINEELRSTTEELETSK------ | *Pseudomonas citronellolis* | Bacteria; Proteobacteria; Gammaproteobacteria; Pseudomonadales; Pseudomonadaceae; Pseudomonas | chemotaxis protein | 6 |
| | SFU12850.1 | 1.8E-07 | Qc.III.c | ---IGETIERINSLSTGINEAVSEQRQATDDISRNLNSAHQNVSQVASNMQ-- | *Pseudovibrio denitrificans* | Bacteria; Proteobacteria; Alphaproteobacteria; Rhodobacterales; Rhodobacteraceae; Pseudovibrio | methyl-accepting chemotaxis protein | 6 |
|  | WP_046508769.1 | 1.8E-07 | Qa.III.b | -EQIDNVIKEINEIASGISTAVEEQGGATREISQTVNQSANN----------- | *Kiloniella litopenaei* | Bacteria; Proteobacteria; Alphaproteobacteria; Kiloniellales; Kiloniellaceae; Kiloniella | HAMP domain-containing protein | 6 |
| | WP_050773957.1 | 1.8E-07 | Qc.III.c | ---IGETIERINSLSTGINEAVSEQRQATDDISRNLNSAHQNVSQVASNMQ-- | *Pseudovibrio sp. JE062* | Bacteria; Proteobacteria; Alphaproteobacteria; Rhodobacterales; Rhodobacteraceae; Pseudovibrio | methyl-accepting chemotaxis protein | 6 |
|  | WP_056296723.1 | 1.8E-07 | Qa.IV | IREISTTINLISEISSTIAAAVEEQGAATQEIARNVQQAAGLS---------- | *Afipia sp. Root123D2* | Bacteria; Proteobacteria; Alphaproteobacteria; Rhizobiales; Bradyrhizobiaceae; Afipia | HAMP domain-containing protein | 6 |
|  | WP_085584554.1 | 1.8E-07 | Qa.III | IDGITRIINDLNSISAEVASAVHQQGSATSEISHSVQQAVSSTRQVS------ | *Thalassospira mesophila* | Bacteria; Proteobacteria; Alphaproteobacteria; Rhodospirillales; Rhodospirillaceae; Thalassospira | methyl-accepting chemotaxis protein | 6 |
|  | OQX62800.1 | 1.9E-07 | Qa.IV | -SQITRTINDINEVVSAIATAVEQQTTTTREIANNVAKASNGI---------- | *Desulfococcus sp. 4484_241* | Bacteria; Proteobacteria; Deltaproteobacteria; Desulfobacterales; Desulfobacteraceae; Desulfococcus | hypothetical protein B5M56_05235 | 6 |
| | WP_008891371.1 | 1.9E-07 | Qc.III | -GEIGSIIDSLNEIATTISAAVEQQGAATDEITRNVRGAADRTRAVSASI--- | *Thalassospira profundimaris* | Bacteria; Proteobacteria; Alphaproteobacteria; Rhodospirillales; Rhodospirillaceae; Thalassospira | HAMP domain-containing protein | 6 |
| | WP_022727453.1 | 1.9E-07 | Qc.III | IQGISSVISEMNEISTSISAAVEEQGAATDEIASTMNRVSDGTQKVNQNIEE- | *Fodinicurvata sediminis* | Bacteria; Proteobacteria; Alphaproteobacteria; Rhodospirillales; Rhodospirillaceae; Fodinicurvata | HAMP domain-containing protein | 6 |
| | WP_026888755.1 | 1.9E-07 | Qb.I | ---INKLVSDTVKQSHDSAEYINESSYLISGTVDTFDNIFNKINSANDLIQE- | *Clostridium beijerinckii* | Bacteria; Firmicutes; Clostridia; Clostridiales; Clostridiaceae; Clostridium | methyl-accepting chemotaxis protein | 6 |
| | WP_052657577.1 | 1.9E-07 | Qc.III | -GEIGSIIDSLNEIATTISAAVEQQGAATDEITRNVRGAADRTRAVSASI--- | *Thalassospira sp. HJ* | Bacteria; Proteobacteria; Alphaproteobacteria; Rhodospirillales; Rhodospirillaceae; Thalassospira | HAMP domain-containing protein | 6 |
| | WP_057854071.1 | 1.9E-07 | Qc.III | IKEISGTIERLAEIASTIAAAVEEQGAATQEISRNVQQAAEGTQQVSSNITD- | *Bradyrhizobium valentinum* | Bacteria; Proteobacteria; Alphaproteobacteria; Rhizobiales; Bradyrhizobiaceae; Bradyrhizobium | methyl-accepting chemotaxis protein | 6 |
| | WP_063087180.1 | 1.9E-07 | Qc.III | -GEIGSIIDSLNEIATTISAAVEQQGAATDEITRNVRGAADRTRAVSASI--- | *Thalassospira* | Bacteria; Proteobacteria; Alphaproteobacteria; Rhodospirillales; Rhodospirillaceae | MULTISPECIES: HAMP domain-containing protein | 6 |
| | WP_064780631.1 | 1.9E-07 | Qc.III | -GEIGSIIDSLNEIATTISAAVEQQGAATDEITRNVRGAADRTRAVSASI--- | *Thalassospira tepidiphila* | Bacteria; Proteobacteria; Alphaproteobacteria; Rhodospirillales; Rhodospirillaceae; Thalassospira | HAMP domain-containing protein | 6 |
| | WP_064790147.1 | 1.9E-07 | Qc.III | -GEIGSIIDSLNEIATTISAAVEQQGAATDEITRNVRGAADRTRAVSASI--- | *Thalassospira profundimaris* | Bacteria; Proteobacteria; Alphaproteobacteria; Rhodospirillales; Rhodospirillaceae; Thalassospira | HAMP domain-containing protein | 6 |
| | WP_077839644.1 | 1.9E-07 | Qb.I | ---INKLVSDTVKQSHDSAEYINESSYLISGTVDTFDNIFNKINSANDLIQE- | *Clostridium beijerinckii* | Bacteria; Firmicutes; Clostridia; Clostridiales; Clostridiaceae; Clostridium | methyl-accepting chemotaxis protein | 6 |
| | WP_078117474.1 | 1.9E-07 | Qb.I | ---INKLVSDTVKQSHDSAEYINESSYLISGTVDTFDNIFNKINSANDLIQE- | *Clostridium* | Bacteria; Firmicutes; Clostridia; Clostridiales; Clostridiaceae | MULTISPECIES: methyl-accepting chemotaxis protein | 6 |
| | WP_085900419.1 | 1.9E-07 | Qc.III | IKSISGTIKSIHEVATAIASAVEEQGAATSEITMSVQQAANGTDQVTSNIRQ- | *Kiloniella majae* | Bacteria; Proteobacteria; Alphaproteobacteria; Kiloniellales; Kiloniellaceae; Kiloniella | hypothetical protein | 6 |
| | WP_085904628.1 | 1.9E-07 | Qc.III | IKSISGTIKSIHEVATAIASAVEEQGAATSEITMSVQQAANGTDQVTSNIRQ- | *Kiloniella majae* | Bacteria; Proteobacteria; Alphaproteobacteria; Kiloniellales; Kiloniellaceae; Kiloniella | hypothetical protein | 6 |
| | WP_085908390.1 | 1.9E-07 | Qc.III | IKSISGTIKSIHEVATAIASAVEEQGAATSEITMSVQQAANGTDQVTSNIRQ- | *Kiloniella majae* | Bacteria; Proteobacteria; Alphaproteobacteria; Kiloniellales; Kiloniellaceae; Kiloniella | hypothetical protein | 6 |
| | OAV38365.1 | 2E-07 | Qb.I | ---IKKLVSDTVKQSHESADYINESSYLISGTVDTFDDIFNKINGTNNLI--- | *Clostridium saccharobutylicum DSM 13864* | Bacteria; Firmicutes; Clostridia; Clostridiales; Clostridiaceae; Clostridium | methyl-accepting chemotaxis protein | 6 |
|  | WP_020585217.1 | 2E-07 | Qa.III | IKDISGIIQNVNEIVSDIAAAVDEQSATTGEIVENMNRTSSKMQEVND----- | *Desulfobacter curvatus* | Bacteria; Proteobacteria; Deltaproteobacteria; Desulfobacterales; Desulfobacteraceae; Desulfobacter | methyl-accepting chemotaxis protein | 6 |
|  | WP_109906860.1 | 2E-07 | Qa.IV | -----RTIASINDITSNIAGAVEQQGAATSEIARNVAHAADDTRQ-------- | *Zavarzinia sp. HR-AS* | Bacteria; Proteobacteria; Alphaproteobacteria; Rhodospirillales; Acetobacteraceae; Zavarzinia | HAMP domain-containing protein | 6 |
|  | EJN14795.1 | 2.1E-07 | Qa.III | IQSICATIQQVDEISAAIAAAVGQQGTATQEIAQNVQQAA------------- | *Bradyrhizobium sp. YR681* | Bacteria; Proteobacteria; Alphaproteobacteria; Rhizobiales; Bradyrhizobiaceae; Bradyrhizobium | methyl-accepting chemotaxis protein | 6 |
|  | PZM95177.1 | 2.1E-07 | Qc.III | IQEIGKIIERINETQIAIASAVEEQTAATNDIGRSADAAAE------------ | *Actinobacteria bacterium* | Bacteria; Actinobacteria | methyl-accepting chemotaxis protein | 6 |
|  | SFV08122.1 | 2.1E-07 | Qa.III | IQSICATIQQVDEISAAIAAAVGQQGTATQEIAQNVQQAA------------- | *Bradyrhizobium arachidis* | Bacteria; Proteobacteria; Alphaproteobacteria; Rhizobiales; Bradyrhizobiaceae; Bradyrhizobium | methyl-accepting chemotaxis sensory transducer | 6 |
|  | WP_018641617.1 | 2.1E-07 | Qa.III | IQSICATIQQVDEISAAIAAAVGQQGTATQEIAQNVQQAA------------- | *Bradyrhizobium japonicum* | Bacteria; Proteobacteria; Alphaproteobacteria; Rhizobiales; Bradyrhizobiaceae; Bradyrhizobium | methyl-accepting chemotaxis protein | 6 |
|  | WP_027561081.1 | 2.1E-07 | Qa.III | IQSICATIQQVDEISAAIAAAVGQQGTATQEIAQNVQQAA------------- | *Bradyrhizobium genosp. SA-4* | Bacteria; Proteobacteria; Alphaproteobacteria; Rhizobiales; Bradyrhizobiaceae; Bradyrhizobium | methyl-accepting chemotaxis protein | 6 |
|  | WP_028138416.1 | 2.1E-07 | Qa.III | IQSICATIQQVDEISAAIAAAVGQQGTATQEIAQNVQQAA------------- | *Bradyrhizobium japonicum* | Bacteria; Proteobacteria; Alphaproteobacteria; Rhizobiales; Bradyrhizobiaceae; Bradyrhizobium | methyl-accepting chemotaxis protein | 6 |
|  | WP_063201038.1 | 2.1E-07 | Qa.III | IQSICATIQQVDEISAAIAAAVGQQGTATQEIAQNVQQAA------------- | *Bradyrhizobium sp. AT1* | Bacteria; Proteobacteria; Alphaproteobacteria; Rhizobiales; Bradyrhizobiaceae; Bradyrhizobium | methyl-accepting chemotaxis protein | 6 |
|  | WP_063694119.1 | 2.1E-07 | Qa.III | IQSICATIQQVDEISAAIAAAVGQQGTATQEIAQNVQQAA------------- | *Bradyrhizobium stylosanthis* | Bacteria; Proteobacteria; Alphaproteobacteria; Rhizobiales; Bradyrhizobiaceae; Bradyrhizobium | methyl-accepting chemotaxis protein | 6 |
|  | WP_096700859.1 | 2.1E-07 | Qa.III | IQSIAGTIDQLNEIATNIASAIEQQGAATSTIAGNIQEAA------------- | *Magnetospirillum sp. 15-1* | Bacteria; Proteobacteria; Alphaproteobacteria; Rhodospirillales; Rhodospirillaceae; Magnetospirillum | methyl-accepting chemotaxis protein | 6 |
|  | AUC95904.1 | 2.2E-07 | Qa.IV | IREIGATIERLSEISSTIAAAVEEQGAATQEISRNVQHAAE------------ | *Bradyrhizobium sp. SK17* | Bacteria; Proteobacteria; Alphaproteobacteria; Rhizobiales; Bradyrhizobiaceae; Bradyrhizobium | methyl-accepting chemotaxis protein | 6 |
| | CAC84419.1 | 2.2E-07 | Qc.III | --DVSKVIGEIHEISASISAAVEEQGAATREIARNVDEAATGTNRVSSAIGR- | *Rhodospirillum rubrum ATCC 11170* | Bacteria; Proteobacteria; Alphaproteobacteria; Rhodospirillales; Rhodospirillaceae; Rhodospirillum | hypothetical protein partial | 6 |
|  | WP_024579376.1 | 2.2E-07 | Qa.IV | IREIGATIERLSEISSTIAAAVEEQGAATQEISRNVQHAAE------------ | *Bradyrhizobium* | Bacteria; Proteobacteria; Alphaproteobacteria; Rhizobiales; Bradyrhizobiaceae | MULTISPECIES: methyl-accepting chemotaxis protein | 6 |
|  | WP_041757297.1 | 2.2E-07 | Qa.III | IQMIGGTIAEIDGISKEIATAVEQQGLATGEISGNVQQASE------------ | *Bradyrhizobium sp. ORS 278* | Bacteria; Proteobacteria; Alphaproteobacteria; Rhizobiales; Bradyrhizobiaceae; Bradyrhizobium | chemotaxis protein | 6 |
|  | WP_054029474.1 | 2.2E-07 | Qa.III | IEEITRIIQEVNDVVSGIATAVEEQSTATSEIAGNVSQAARGIQ--------- | *Desulfatitalea tepidiphila* | Bacteria; Proteobacteria; Deltaproteobacteria; Desulfobacterales; Desulfobacteraceae; Desulfatitalea | methyl-accepting chemotaxis protein | 6 |
| | WP_068518122.1 | 2.2E-07 | Qc.III | -GEIGTIIDNLNQIATTISAAVEEQGAATDEISRNVRGAADRTRAVSASI--- | *Thalassospira* | Bacteria; Proteobacteria; Alphaproteobacteria; Rhodospirillales; Rhodospirillaceae | MULTISPECIES: methyl-accepting chemotaxis protein | 6 |
|  | WP_104462836.1 | 2.2E-07 | Qa.IV | IREISATIERLSEISSTVAAAVEQQGAATQEISRNVQQAA------------- | *Bradyrhizobium sp. AC87j1* | Bacteria; Proteobacteria; Alphaproteobacteria; Rhizobiales; Bradyrhizobiaceae; Bradyrhizobium | methyl-accepting chemotaxis protein | 6 |
| | WP_114112607.1 | 2.2E-07 | Qc.III | -GEIGTIIDNLNQIATTISAAVEEQGAATDEISRNVRGAADRTRAVSASI--- | *Thalassospira profundimaris* | Bacteria; Proteobacteria; Alphaproteobacteria; Rhodospirillales; Rhodospirillaceae; Thalassospira | HAMP domain-containing protein | 6 |
|  | SDI67046.1 | 2.4E-07 | Qa.IV | -RDISDTIEKLSEISSTIAAAVEEQGAATQEISRNVQQAAE------------ | *Bradyrhizobium sp. Rc2d* | Bacteria; Proteobacteria; Alphaproteobacteria; Rhizobiales; Bradyrhizobiaceae; Bradyrhizobium | methyl-accepting chemotaxis protein | 6 |
|  | WP_036348336.1 | 2.4E-07 | Qc.III | IEAITSTVQRLNEISVAISAAVEEQSAVTQEMSSNMQGAS------------- | *Microvirga sp. BSC39* | Bacteria; Proteobacteria; Alphaproteobacteria; Rhizobiales; Methylobacteriaceae; Microvirga | PAS domain S-box protein | 6 |
|  | WP_092063678.1 | 2.4E-07 | Qa.III | -NAITESIKEVNDLANTIASATEEQTVTMNDITSNVKRTTEAAEK-------- | *Desulfonauticus submarinus* | Bacteria; Proteobacteria; Deltaproteobacteria; Desulfovibrionales; Desulfohalobiaceae; Desulfonauticus | bacteriohemerythrin | 6 |
|  | CEG10403.1 | 2.5E-07 | Qa.III | IDEITQTIGRVNEIASAIASAVEEQGAATREIANNVSQAAE------------ | *Afipia felis* | Bacteria; Proteobacteria; Alphaproteobacteria; Rhizobiales; Bradyrhizobiaceae; Afipia | Methyl-accepting chemotaxis protein 4 | 6 |
|  | OYX14924.1 | 2.5E-07 | Qa.IV | --KIVGTISDIRQATSTIASAIEEQGAATQEIATNTQHAAHGTG--------- | *Rhizobiales bacterium 32-66-8* | Bacteria; Proteobacteria; Alphaproteobacteria; Rhizobiales | hypothetical protein B7Z15_02295 | 6 |
|  | WP_015773350.1 | 2.5E-07 | Qa.III | -GQIVKVIGDVNEIVTTIAAAIEEQSAVTREVATNINQATTGVQDA------- | *Desulfomicrobium baculatum* | Bacteria; Proteobacteria; Deltaproteobacteria; Desulfovibrionales; Desulfomicrobiaceae; Desulfomicrobium | methyl-accepting chemotaxis protein | 6 |
|  | WP_093556408.1 | 2.5E-07 | Qb.I | LQRTRAQLQETVEHSETSNEELKASNEELQAINEELRSATEELETS------- | *Massilia namucuonensis* | Bacteria; Proteobacteria; Betaproteobacteria; Burkholderiales; Oxalobacteraceae; Massilia | PAS domain S-box protein | 6 |
|  | WP_094375875.1 | 2.5E-07 | Qb.I | ------LVNNTVEQSKESAEYINSSTQKISAATDTFDSIFDAINQTGQMI--- | *Lachnotalea glycerini* | Bacteria; Firmicutes; Clostridia; Clostridiales; Lachnospiraceae; Lachnotalea | methyl-accepting chemotaxis protein | 6 |
|  | WP_103012690.1 | 2.5E-07 | Qa.IV | IRKIGETISRMSEISSTIASAVEQQGAATQEISRNVQQAA------------- | *Rhodopseudomonas palustris* | Bacteria; Proteobacteria; Alphaproteobacteria; Rhizobiales; Bradyrhizobiaceae; Rhodopseudomonas | methyl-accepting chemotaxis protein | 6 |
|  | WP_110291165.1 | 2.5E-07 | Qb.I | ------LVNNTVEQSKESAEYINSSTQKISAATDTFDSIFDAINQTGQMI--- | *Lachnotalea glycerini* | Bacteria; Firmicutes; Clostridia; Clostridiales; Lachnospiraceae; Lachnotalea | methyl-accepting chemotaxis protein | 6 |
|  | WP_110779578.1 | 2.5E-07 | Qa.IV | IRKIGETISRMSEISSTIASAVEQQGAATQEISRNVQQAA------------- | *Rhodopseudomonas faecalis* | Bacteria; Proteobacteria; Alphaproteobacteria; Rhizobiales; Bradyrhizobiaceae; Rhodopseudomonas | methyl-accepting chemotaxis protein | 6 |
|  | KMO22875.1 | 2.6E-07 | Qa.IV | ILQIGRTIASINEISGMIAATVVQQTEATTEISRNAAEAA------------- | *Methylobacterium platani* | Bacteria; Proteobacteria; Alphaproteobacteria; Rhizobiales; Methylobacteriaceae; Methylobacterium | chemotaxis protein partial | 6 |

| | | | | | | | |
|---|---|---|---|---|---|---|---|
| KMO25974.1 | 2.6E-07 | Qa.IV | ILQIGRTIASINEISGMIAATVVQQTEATTEISRNAAEAA------------- | *Methylobacterium platani* | Bacteria; Proteobacteria; Alphaproteobacteria; Rhizobiales; Methylobacteriaceae; Methylobacterium | chemotaxis protein partial | 6 |
| PXV90333.1 | 2.6E-07 | Qb.I | ------LVNNTVEQSKESAEYINSSTQKISAATDTFDSIFDAINQTGQMI--- | *Lachnotalea glycerini* | Bacteria; Firmicutes; Clostridia; Clostridiales; Lachnospiraceae; Lachnotalea | methyl-accepting chemotaxis protein | 6 |
| WP_009029505.1 | 2.6E-07 | Qa.III | IQMIGGTIAEIDGISKEIATAVEQQGLATGEISGNVQQASE------------ | *Bradyrhizobium sp. ORS 375* | Bacteria; Proteobacteria; Alphaproteobacteria; Rhizobiales; Bradyrhizobiaceae; Bradyrhizobium | methyl-accepting chemotaxis protein | 6 |
| WP_026653399.1 | 2.6E-07 | Qb.I | ---ITNLIKDLAEKSQANMEAITNSSGAVSKAGDSFSKINQELNSTA------ | *Butyrivibrio proteoclasticus* | Bacteria; Firmicutes; Clostridia; Clostridiales; Lachnospiraceae; Butyrivibrio | methyl-accepting chemotaxis protein | 6 |
| WP_027277864.1 | 2.6E-07 | Qa.IV | IRKIGETISRMSEISSTIASAVEQQGAATQEISRNVQQAA------------- | *Rhodopseudomonas palustris* | Bacteria; Proteobacteria; Alphaproteobacteria; Rhizobiales; Bradyrhizobiaceae; Rhodopseudomonas | methyl-accepting chemotaxis protein | 6 |
| CAL74683.1 | 2.7E-07 | Qa.III | IQMIGGTIAEIDGISKEIATAVEQQGLATGEISGNVQQASE------------ | *Bradyrhizobium sp. ORS 278* | Bacteria; Proteobacteria; Alphaproteobacteria; Rhizobiales; Bradyrhizobiaceae; Bradyrhizobium | Hypothetical protein; Putative Methyl-accepting chemotaxis protein (fragment) | 6 |
| WP_006612433.1 | 2.7E-07 | Qc.III | IKEIGGTINRIAEISAAIAAAIEQQGAATQEIARNVDQAAQGTTQVASNIT-- | *Bradyrhizobium sp. ORS 285* | Bacteria; Proteobacteria; Alphaproteobacteria; Rhizobiales; Bradyrhizobiaceae; Bradyrhizobium | methyl-accepting chemotaxis protein | 6 |
| WP_071178934.1 | 2.7E-07 | Qa.III | IKQIVAMIDQLNDMNTQIATATDEQREVTEEINSNMTSISDASKKVAQQ---- | *Vibrio cholerae* | Bacteria; Proteobacteria; Gammaproteobacteria; Vibrionales; Vibrionaceae; Vibrio | methyl-accepting chemotaxis protein | 6 |
| WP_084988670.1 | 2.7E-07 | Qa.III | IKQIVAMIDQLNDMNTQIATATDEQREVTEEINSNMTSISDASKKVAQQ---- | *Vibrio cholerae* | Bacteria; Proteobacteria; Gammaproteobacteria; Vibrionales; Vibrionaceae; Vibrio | methyl-accepting chemotaxis protein | 6 |
| WP_108315083.1 | 2.7E-07 | Qa.III | IKQIVAMIDQLNDMNTQIATATDEQREVTEEINSNMTSISDASKKVAQQ---- | *Vibrio cholerae* | Bacteria; Proteobacteria; Gammaproteobacteria; Vibrionales; Vibrionaceae; Vibrio | methyl-accepting chemotaxis protein | 6 |
| WP_113593663.1 | 2.7E-07 | Qa.III | IKQIVAMIDQLNDMNTQIATATDEQREVTEEINSNMTSISDASKKVAQQ---- | *Vibrio* | Bacteria; Proteobacteria; Gammaproteobacteria; Vibrionales; Vibrionaceae | MULTISPECIES: methyl-accepting chemotaxis protein | 6 |
| RCK50404.1 | 2.8E-07 | Qa.III | IDGITSIIANLNEIASEVAAAVQEQGQATSEISNSVQQAVQSTQE-------- | *Thalassospira xiamenensis* | Bacteria; Proteobacteria; Alphaproteobacteria; Rhodospirillales; Rhodospirillaceae; Thalassospira | chemotaxis protein | 6 |
| SMX59454.1 | 2.8E-07 | Qc.III | IKEIGSTIGRLSEISAAVSAAVEEQGAATQEISRNVQHAANGTQRVSRNIDD- | *Bradyrhizobium sp. ORS 285* | Bacteria; Proteobacteria; Alphaproteobacteria; Rhizobiales; Bradyrhizobiaceae; Bradyrhizobium | putative methyl-accepting chemotaxis protein | 6 |
| WP_013501517.1 | 2.8E-07 | Qa.IV | IKEIGTTISQMSEISSAIASAVEQQGSATQEISRNVQQAA------------- | *Rhodopseudomonas palustris* | Bacteria; Proteobacteria; Alphaproteobacteria; Rhizobiales; Bradyrhizobiaceae; Rhodopseudomonas | methyl-accepting chemotaxis protein | 6 |
| WP_062661193.1 | 2.8E-07 | Qa.III | --QIEGMMVDINDMNTQIATAATEQSSVTEEISMNVNNIARSS---------- | *Grimontia celer* | Bacteria; Proteobacteria; Gammaproteobacteria; Vibrionales; Vibrionaceae; Grimontia | methyl-accepting chemotaxis protein | 6 |
| WP_097053227.1 | 2.8E-07 | Qa.III | IDGITSIIANLNEIASEVAAAVQEQGQATSEISNSVQQAVQSTQE-------- | *Thalassospira xiamenensis* | Bacteria; Proteobacteria; Alphaproteobacteria; Rhodospirillales; Rhodospirillaceae; Thalassospira | methyl-accepting chemotaxis protein | 6 |
| WP_114109093.1 | 2.8E-07 | Qa.III | IDGITSIIANLNEIASEVAAAVQEQGQATSEISNSVQQAVQSTQE-------- | *Thalassospira xiamenensis* | Bacteria; Proteobacteria; Alphaproteobacteria; Rhodospirillales; Rhodospirillaceae; Thalassospira | HAMP domain-containing protein | 6 |
| WP_027539080.1 | 2.9E-07 | Qa.IV | IKEISSTIERLSEISSTIAAAVEEQGAATQEISRNVQQAAQ------------ | *Bradyrhizobium sp. URHA0002* | Bacteria; Proteobacteria; Alphaproteobacteria; Rhizobiales; Bradyrhizobiaceae; Bradyrhizobium | methyl-accepting chemotaxis protein partial | 6 |
| WP_068499254.1 | 2.9E-07 | Qa.III.b | IDSISKVIREVNEISTTVASAVQEQTAATDEIARNVEQAASGTQEVSA----- | *Magnetospirillum moscoviense* | Bacteria; Proteobacteria; Alphaproteobacteria; Rhodospirillales; Rhodospirillaceae; Magnetospirillum | HAMP domain-containing protein | 6 |
| WP_077577972.1 | 2.9E-07 | Qc.III | -DEISGNLSNIKNLSSDMNEEIEQLGPIVSNLKENVDSLSGAIR--------- | *Salinivibrio* | Bacteria; Proteobacteria; Gammaproteobacteria; Vibrionales; Vibrionaceae | MULTISPECIES: methyl-accepting chemotaxis protein | 6 |
| EFO30724.1 | 3E-07 | Qa.III | IKSIASTISQINEISSGISEAVEQQSLATSEIAQNIEQASSGTQ--------- | *Roseibium sp. TrichSKD4* | Bacteria; Proteobacteria; Alphaproteobacteria; Rhodobacterales; Rhodobacteraceae; Roseibium | methyl-accepting chemotaxis protein | 6 |
| EKD27094.1 | 3E-07 | Qa.III | IKEIVNIIEKVNDITHTIASAVEEQSATVSEIAQTVNNTKTSVQETASNIK-- | *uncultured bacterium* | Bacteria; environmental samples | hypothetical protein ACD_79C00882G0002 partial | 6 |
| KPG01045.1 | 3E-07 | Qa.IV | IKQITGTIEQLSEIASTIAAAVEEQGAATQEISRNVQQAAQ------------ | *Rhodopseudomonas sp. AAP120* | Bacteria; Proteobacteria; Alphaproteobacteria; Rhizobiales; Bradyrhizobiaceae; Rhodopseudomonas | chemotaxis protein | 6 |
| OAN52456.1 | 3E-07 | Qa.III.b | IDSISKVIREVNEISTTVASAVQEQTAATDEIARNVEQAASGTQEVSA----- | *Magnetospirillum moscoviense* | Bacteria; Proteobacteria; Alphaproteobacteria; Rhodospirillales; Rhodospirillaceae; Magnetospirillum | hypothetical protein A6A05_10770 | 6 |
| SMX58010.1 | 3E-07 | Qa.IV | IREISTTIEKLSEISSTIAAAVEQQGAATQEISRNVQHAA------------- | *Bradyrhizobium sp. ORS 285* | Bacteria; Proteobacteria; Alphaproteobacteria; Rhizobiales; Bradyrhizobiaceae; Bradyrhizobium | putative methyl-accepting chemotaxis receptor/sensory transducer | 6 |
| WP_044537065.1 | 3E-07 | Qa.IV | IQSIRNTIMEVNEISSSIAVAVEQQGAATREIASSVQKAA------------- | *Bradyrhizobium sp. LTSP885* | Bacteria; Proteobacteria; Alphaproteobacteria; Rhizobiales; Bradyrhizobiaceae; Bradyrhizobium | methyl-accepting chemotaxis protein | 6 |
| WP_050763477.1 | 3E-07 | Qa.III | IKSIASTISQINEISSGISEAVEQQSLATSEIAQNIEQASSGTQ--------- | *Roseibium sp. TrichSKD4* | Bacteria; Proteobacteria; Alphaproteobacteria; Rhodobacterales; Rhodobacteraceae; Roseibium | methyl-accepting chemotaxis protein | 6 |
| WP_074121346.1 | 3E-07 | Qa.III | IQSICATIQQVDEISAAIAAAVSQQGTATQEIAQNVQQAA------------- | *Bradyrhizobium sp. AS23.2* | Bacteria; Proteobacteria; Alphaproteobacteria; Rhizobiales; Bradyrhizobiaceae; Bradyrhizobium | methyl-accepting chemotaxis protein | 6 |
| WP_082910799.1 | 3E-07 | Qa.III.b | IDSISKVIREVNEISTTVASAVQEQTAATDEIARNVEQAASGTQEVSA----- | *Magnetospirillum moscoviense* | Bacteria; Proteobacteria; Alphaproteobacteria; Rhodospirillales; Rhodospirillaceae; Magnetospirillum | HAMP domain-containing protein | 6 |
| AEO47118.1 | 3.1E-07 | Qc.III | --DVSKVIGEIHEISASISAAVEEQGAATREIARNVDEAATGTNRVSSAIGR- | *Rhodospirillum rubrum F11* | Bacteria; Proteobacteria; Alphaproteobacteria; Rhodospirillales; Rhodospirillaceae; Rhodospirillum | chemotaxis sensory transducer | 6 |
| PIE74191.1 | 3.1E-07 | Qa.III | IRDIAETIDKVNEIVASIATAVEDQATTTAEIADNITQAS------------- | *Deltaproteobacteria bacterium* | Bacteria; Proteobacteria; Deltaproteobacteria | chemotaxis protein | 6 |
| WP_108514433.1 | 3.1E-07 | Qa.IV | IRDISNTISEINEVTTAIAAAVEEQGAATREIARNIQHAA------------- | *Bradyrhizobium algeriense* | Bacteria; Proteobacteria; Alphaproteobacteria; Rhizobiales; Bradyrhizobiaceae; Bradyrhizobium | HAMP domain-containing protein | 6 |
| WP_011388390.1 | 3.2E-07 | Qc.III | --DVSKVIGEIHEISASISAAVEEQGAATREIARNVDEAATGTNRVSSAIGR- | *Rhodospirillum rubrum* | Bacteria; Proteobacteria; Alphaproteobacteria; Rhodospirillales; Rhodospirillaceae; Rhodospirillum | HAMP domain-containing protein | 6 |
| WP_046829033.1 | 3.2E-07 | Qa.IV | IKEISSTINLMSEISSTIAAAVEEQGAATQEIARNVQQAAE------------ | *Afipia massiliensis* | Bacteria; Proteobacteria; Alphaproteobacteria; Rhizobiales; Bradyrhizobiaceae; Afipia | methyl-accepting chemotaxis protein | 6 |
| WP_074831850.1 | 3.2E-07 | Qa.IV | IKAISSTIERLSEISSTIAAAVEEQGAATQEISRNVQQAAQ------------ | *Bradyrhizobium* | Bacteria; Proteobacteria; Alphaproteobacteria; Rhizobiales; Bradyrhizobiaceae | MULTISPECIES: methyl-accepting chemotaxis protein | 6 |
| WP_079587968.1 | 3.2E-07 | Qa.IV | IKAISSTIERLSEISSTIAAAVEEQGAATQEISRNVQQAAQ------------ | *Bradyrhizobium lablabi* | Bacteria; Proteobacteria; Alphaproteobacteria; Rhizobiales; Bradyrhizobiaceae; Bradyrhizobium | methyl-accepting chemotaxis protein | 6 |
| WP_079587969.1 | 3.2E-07 | Qa.IV | IKAISSTIERLSEISSTIAAAVEEQGAATQEISRNVQQAAQ------------ | *Bradyrhizobium lablabi* | Bacteria; Proteobacteria; Alphaproteobacteria; Rhizobiales; Bradyrhizobiaceae; Bradyrhizobium | methyl-accepting chemotaxis protein | 6 |
| OOF27716.1 | 3.3E-07 | Qc.III | -DEISSNLSNIKTLSGDMNEAIDQLGPIVSDLKNNVDSLSSAIKH-------- | *Salinivibrio sp. IB872* | Bacteria; Proteobacteria; Gammaproteobacteria; Vibrionales; Vibrionaceae; Salinivibrio | methyl-accepting chemotaxis protein | 6 |
| SDH86623.1 | 3.3E-07 | Qa.IV | IKAISSTIERLSEISSTIAAAVEEQGAATQEISRNVQQAAQ------------ | *Bradyrhizobium ottawaense* | Bacteria; Proteobacteria; Alphaproteobacteria; Rhizobiales; Bradyrhizobiaceae; Bradyrhizobium | Methyl-accepting chemotaxis protein | 6 |
| SHL96171.1 | 3.3E-07 | Qa.IV | IKAISSTIERLSEISSTIAAAVEEQGAATQEISRNVQQAAQ------------ | *Bradyrhizobium lablabi* | Bacteria; Proteobacteria; Alphaproteobacteria; Rhizobiales; Bradyrhizobiaceae; Bradyrhizobium | Methyl-accepting chemotaxis protein | 6 |
| SHL96198.1 | 3.3E-07 | Qa.IV | IKAISSTIERLSEISSTIAAAVEEQGAATQEISRNVQQAAQ------------ | *Bradyrhizobium lablabi* | Bacteria; Proteobacteria; Alphaproteobacteria; Rhizobiales; Bradyrhizobiaceae; Bradyrhizobium | methyl-accepting chemotaxis sensory transducer | 6 |
| WP_044408004.1 | 3.3E-07 | Qa.IV | IKEIGTTISQMSEISSAIASAVEQQGAATQEISRNVQQAA------------- | *Rhodopseudomonas palustris* | Bacteria; Proteobacteria; Alphaproteobacteria; Rhizobiales; Bradyrhizobiaceae; Rhodopseudomonas | chemotaxis protein | 6 |
| WP_057850125.1 | 3.3E-07 | Qa.IV | IRNISNTISEINEVTTAIAAAVEEQGAATREIARNIQHAA------------- | *Bradyrhizobium valentinum* | Bacteria; Proteobacteria; Alphaproteobacteria; Rhizobiales; Bradyrhizobiaceae; Bradyrhizobium | HAMP domain-containing protein | 6 |
| WP_063236911.1 | 3.3E-07 | Qb.I | LHRTKEQLQATIEQSETSTEELKASNEELQAINEELRSTTEELETSK------ | *Cupriavidus oxalaticus* | Bacteria; Proteobacteria; Betaproteobacteria; Burkholderiales; Burkholderiaceae; Cupriavidus | PAS domain S-box protein | 6 |
| WP_082363863.1 | 3.3E-07 | Qa.IV | IKQITGTIEQLSEIASTIAAAVEEQGAATQEISRNVQQAAQ------------ | *Rhodopseudomonas sp. AAP120* | Bacteria; Proteobacteria; Alphaproteobacteria; Rhizobiales; Bradyrhizobiaceae; Rhodopseudomonas | HAMP domain-containing protein | 6 |
| WP_108743130.1 | 3.3E-07 | Qc.III | -DEISSNLSNIKTLSGDMNEAIDQLGPIVSDLKNNVDSLSSAIKH-------- | *Salinivibrio sp. IB872* | Bacteria; Proteobacteria; Gammaproteobacteria; Vibrionales; Vibrionaceae; Salinivibrio | methyl-accepting chemotaxis protein | 6 |
| WP_114359794.1 | 3.3E-07 | Qa.IV | IKQITGTIEQLSEIASTIAAAVEEQGAATQEISRNVQQAAQ------------ | *Rhodopseudomonas pentothenatexigens* | Bacteria; Proteobacteria; Alphaproteobacteria; Rhizobiales; Bradyrhizobiaceae; Rhodopseudomonas | HAMP domain-containing protein | 6 |
| PHR89509.1 | 3.4E-07 | Qc.III | LDSMTNSVMQINDLGIQIATAAEEQSSVTEEINRNMTAIQSMVNKLTENGDK- | *Moritella sp.* | Bacteria; Proteobacteria; Gammaproteobacteria; Alteromonadales; Moritellaceae; Moritella | methyl-accepting chemotaxis protein | 6 |
| SMF86095.1 | 3.4E-07 | Qc.III | -RSIGEVIGQMSEIATSISAAVEEQGAATDEIAQTMSRAADGTQTVAGNI--- | *Tistlia consotensis USBA 355* | Bacteria; Proteobacteria; Alphaproteobacteria; Rhodospirillales; Rhodospirillaceae; Tistlia | Methyl-accepting chemotaxis protein partial | 6 |
| WP_063691908.1 | 3.5E-07 | Qa.IV | IKDISNTIEKLSEISSTIAAAVEEQGAATQEISRNVQQAAE------------ | *Bradyrhizobium stylosanthis* | Bacteria; Proteobacteria; Alphaproteobacteria; Rhizobiales; Bradyrhizobiaceae; Bradyrhizobium | methyl-accepting chemotaxis protein | 6 |
| AIB13512.1 | 3.6E-07 | Qa.IV | --NIVGTIQEVHETSSSIAAAIHEQHAAIAEISRNVQQTAAGTADINRHI--- | *Azospirillum brasilense* | Bacteria; Proteobacteria; Alphaproteobacteria; Rhodospirillales; Rhodospirillaceae; Azospirillum | hypothetical protein ABAZ39_16345 (plasmid) | 6 |
| ANK81588.1 | 3.6E-07 | Qa.IV | IQKVTSAVKELNGIASTIASAVEEQTAVTQDIARNVQQAA------------- | *Rhizobiales bacterium NRL2* | Bacteria; Proteobacteria; Alphaproteobacteria; Rhizobiales | hypothetical protein TEF_12885 | 6 |
| WP_006614844.1 | 3.6E-07 | Qa.IV | IKEISTTIERLSEISSTIAAAVEQQGAATQEISRNIQEASR------------ | *Bradyrhizobium sp. ORS 285* | Bacteria; Proteobacteria; Alphaproteobacteria; Rhizobiales; Bradyrhizobiaceae; Bradyrhizobium | PAS domain S-box protein | 6 |
| WP_008145161.1 | 3.6E-07 | Qa.IV | IKDISNTIEKLSEISSTIAAAVEEQGAATQEISRNVQQAAE------------ | *Bradyrhizobium sp. YR681* | Bacteria; Proteobacteria; Alphaproteobacteria; Rhizobiales; Bradyrhizobiaceae; Bradyrhizobium | methyl-accepting chemotaxis protein | 6 |
| WP_109794641.1 | 3.6E-07 | Qa.IV | IQKVTSAVKELNGIASTIASAVEEQTAVTQDIARNVQQAA------------- | *Rhizobiales bacterium* | Bacteria; Proteobacteria; Alphaproteobacteria; Rhizobiales | HAMP domain-containing protein | 6 |
| AFK52068.1 | 3.7E-07 | Qa.IV | ---VTVSIREMHEIALTIASAIDEQGAATAEIARNVQEAATATDR-------- | *Tistrella mobilis KA081020-065* | Bacteria; Proteobacteria; Alphaproteobacteria; Rhodospirillales; Rhodospirillaceae; Tistrella | chemotaxis sensory transducer | 6 |
| WP_028348059.1 | 3.7E-07 | Qa.IV | IRNISTTISEINEVTTAIAAAVEEQGAATREIARNIQHAA------------- | *Bradyrhizobium elkanii* | Bacteria; Proteobacteria; Alphaproteobacteria; Rhizobiales; Bradyrhizobiaceae; Bradyrhizobium | HAMP domain-containing protein | 6 |
| WP_065726616.1 | 3.7E-07 | Qa.IV | IRNISTTISEINEVTTAIAAAVEEQGAATREIARNIQHAA------------- | *Bradyrhizobium icense* | Bacteria; Proteobacteria; Alphaproteobacteria; Rhizobiales; Bradyrhizobiaceae; Bradyrhizobium | methyl-accepting chemotaxis protein | 6 |

| | | | | | | | |
|---|---|---|---|---|---|---|---|
| **AWJ93740.1** | 3.8E-07 | Qa.IV | --NIVGTIQEVHETSSSIAAAIHEQHAAIAEISRNVQQTAAGTADINRHI--- | *Azospirillum brasilense Sp245* | Bacteria; Proteobacteria; Alphaproteobacteria; Rhodospirillales; Rhodospirillaceae; Azospirillum | methyl-accepting chemotaxis sensory transducer (plasmid) | 6 |
| **EIM62400.1** | 3.8E-07 | Qa.III | IEEISAVIADVNDIVSTIATAVEEQSSATREIAQNIAQASSGIEEVN------ | *Desulfobacter postgatei 2ac9* | Bacteria; Proteobacteria; Deltaproteobacteria; Desulfobacterales; Desulfobacteraceae; Desulfobacter | methyl-accepting chemotaxis protein | 6 |
| **PIE67358.1** | 3.8E-07 | Qa.III | --EISNVIGNVDEIVSSIATAVEEQSVTTREISDNIAQASSGIQE-------- | *Deltaproteobacteria bacterium* | Bacteria; Proteobacteria; Deltaproteobacteria | chemotaxis protein | 6 |
| **PNQ95376.1** | 3.8E-07 | Qa.IV | --NIVGTIQEVHETSSSIAAAIHEQHAAIAEISRNVQQTAAGTADINRHI--- | *Azospirillum brasilense* | Bacteria; Proteobacteria; Alphaproteobacteria; Rhodospirillales; Rhodospirillaceae; Azospirillum | methyl-accepting chemotaxis protein (plasmid) | 6 |
| **WP_008972784.1** | 3.8E-07 | Qc.III | -REIGSTIGRLSEIAAAVSAAVEEQGAATQEISRNVQHAASGTQRVSTNIDN- | *Bradyrhizobium sp. STM 3843* | Bacteria; Proteobacteria; Alphaproteobacteria; Rhizobiales; Bradyrhizobiaceae; Bradyrhizobium | HAMP domain-containing protein | 6 |
| **WP_081598718.1** | 3.8E-07 | Qa.IV | ---VTVSIREMHEIALTIASAIDEQGAATAEIARNVQEAATATDR-------- | *Tistrella mobilis* | Bacteria; Proteobacteria; Alphaproteobacteria; Rhodospirillales; Rhodospirillaceae; Tistrella | methyl-accepting chemotaxis protein | 6 |
| **WP_044616392.1** | 3.9E-07 | Qa.III | --EIRSAISTINEMATQIATAAEEQTSVTAEINQNTT----NIHTASQEL--- | *Gynuella sunshinyii* | Bacteria; Proteobacteria; Gammaproteobacteria; Oceanospirillales; Saccharospirillaceae; Gynuella | methyl-accepting chemotaxis protein | 6 |
| **WP_092516015.1** | 3.9E-07 | Qa.IV | -KEISGTIERLSEISSTIAAAVEEQGAATQEISRNVQQAAQ------------ | *Afipia sp. GAS231* | Bacteria; Proteobacteria; Alphaproteobacteria; Rhizobiales; Bradyrhizobiaceae; Afipia | methyl-accepting chemotaxis protein | 6 |
| **WP_018323368.1** | 4E-07 | Qa.IV | IRDISNTIEKLSEISSTIAAAVEEQGAATQEISRNVQQAAE------------ | *Bradyrhizobium sp. WSM2793* | Bacteria; Proteobacteria; Alphaproteobacteria; Rhizobiales; Bradyrhizobiaceae; Bradyrhizobium | HAMP domain-containing protein | 6 |
| **WP_074125513.1** | 4E-07 | Qa.IV | IKEISSTIEKLSEISSTIAAAVEEQGAATQEISRNVQQAA------------- | *Bradyrhizobium sp. NAS96.2* | Bacteria; Proteobacteria; Alphaproteobacteria; Rhizobiales; Bradyrhizobiaceae; Bradyrhizobium | methyl-accepting chemotaxis protein | 6 |
| **CUW41714.1** | 4.1E-07 | Qa.IV | ------TIRDINGISTAIASAVEQQSAATSEISRNVQQAAKAVDDVNDNIV-- | *Magnetospirillum sp. XM-1* | Bacteria; Proteobacteria; Alphaproteobacteria; Rhodospirillales; Rhodospirillaceae; Magnetospirillum | putative methyl-accepting chemotaxis protein | 6 |
| **WP_073954342.1** | 4.1E-07 | Qa.III | IQNISRTINEVDEISTAIASAIEQQGSVTHEISNSVYQAVKSTESAQAN---- | *Thalassospira sp. TSL5-1* | Bacteria; Proteobacteria; Alphaproteobacteria; Rhodospirillales; Rhodospirillaceae; Thalassospira | chemotaxis protein | 6 |
| **WP_084809158.1** | 4.1E-07 | Qa.III | IQSICATIRQVDEISAAIAAAVSQQGTATQEIAQNVQQAA------------- | *Bradyrhizobium sp. NAS80.1* | Bacteria; Proteobacteria; Alphaproteobacteria; Rhizobiales; Bradyrhizobiaceae; Bradyrhizobium | methyl-accepting chemotaxis protein | 6 |
| **OJU25415.1** | 4.2E-07 | Qa.IV | IREIGTTIKTISEISSAIAAAVEEQGAATQEIARNVQQAAQR----------- | *Nitrobacter sp. 62-13* | Bacteria; Proteobacteria; Alphaproteobacteria; Rhizobiales; Bradyrhizobiaceae; Nitrobacter | methyl-accepting chemotaxis protein | 6 |
| **WP_008969109.1** | 4.3E-07 | Qa.IV | IRDISSTIEKLSEISSTIAAAVEEQGAATQEISRNVQQAAQ------------ | *Bradyrhizobium sp. STM 3843* | Bacteria; Proteobacteria; Alphaproteobacteria; Rhizobiales; Bradyrhizobiaceae; Bradyrhizobium | methyl-accepting chemotaxis protein | 6 |
| **WP_074271194.1** | 4.3E-07 | Qa.IV | IRNISNTISEINDVTTAIAAAVEEQGAATREIARNIQHAA------------- | *Bradyrhizobium erythrophlei* | Bacteria; Proteobacteria; Alphaproteobacteria; Rhizobiales; Bradyrhizobiaceae; Bradyrhizobium | HAMP domain-containing protein | 6 |
| **WP_082700656.1** | 4.3E-07 | Qa.IV | ------TIRDINGISTAIASAVEQQSAATSEISRNVQQAAKAVDDVNDNIV-- | *Magnetospirillum sp. XM-1* | Bacteria; Proteobacteria; Alphaproteobacteria; Rhodospirillales; Rhodospirillaceae; Magnetospirillum | HAMP domain-containing protein | 6 |
| **PKL59707.1** | 4.4E-07 | Qb.I | ---TKENLQATVEKMQAANEELKSTNEELQSTNEELQSTNEELETSR------ | *Methanomicrobiales archaeon HGW-Methanomicrobiales-4* | Archaea; Euryarchaeota; Methanomicrobia; Methanomicrobiales | SAM-dependent methyltransferase partial | 6 |
| **WP_025034936.1** | 4.4E-07 | Qa.IV | IKEISSTIERLSEISSAIAAAVEEQGAATQEIARNVQQAAQ------------ | *Bradyrhizobium sp. DOA9* | Bacteria; Proteobacteria; Alphaproteobacteria; Rhizobiales; Bradyrhizobiaceae; Bradyrhizobium | methyl-accepting chemotaxis protein | 6 |
| **WP_091132009.1** | 4.4E-07 | Qa.IV | IQDIARTITEMSQISTSIAAAMEEQGAATAEIARNVQEAARVTETASENIA-- | *Microvirga guangxiensis* | Bacteria; Proteobacteria; Alphaproteobacteria; Rhizobiales; Methylobacteriaceae; Microvirga | methyl-accepting chemotaxis protein | 6 |
| **WP_097667111.1** | 4.4E-07 | Qa.IV | IKEISSTIERLSEISSAIAAAVEEQGAATQEIARNVQQAAQ------------ | *Bradyrhizobium sp. Y36* | Bacteria; Proteobacteria; Alphaproteobacteria; Rhizobiales; Bradyrhizobiaceae; Bradyrhizobium | HAMP domain-containing protein | 6 |
| **CAL76374.1** | 4.5E-07 | Qa.IV | IKEISSTIEKLSEISSTIAAAVEQQGAATQEISRNVQQAA------------- | *Bradyrhizobium sp. ORS 278* | Bacteria; Proteobacteria; Alphaproteobacteria; Rhizobiales; Bradyrhizobiaceae; Bradyrhizobium | putative methyl-accepting chemotaxis receptor/sensory transducer (Chemoreceptor) | 6 |
| **OQX11309.1** | 4.5E-07 | Qa.III | IEDILRVINEIDHIVQTIATAVESQTSTTDSIADNIAQAS------------- | *Desulfobulbaceae bacterium A2* | Bacteria; Proteobacteria; Deltaproteobacteria; Desulfobacterales; Desulfobulbaceae | hypothetical protein BWK76_19355 | 6 |
| **PZN41387.1** | 4.5E-07 | Qa.I | IEQIARVVEEVSRIATAIAASVEEQSAVTEGIATNVAQTTEH----------- | *Proteobacteria bacterium* | Bacteria; Proteobacteria | chemotaxis protein partial | 6 |
| **WP_008133441.1** | 4.5E-07 | Qa.IV | IKEISSSIERLSEISSAIAAAVEEQGAATQEIARNVQQAA------------- | *Bradyrhizobium sp. YR681* | Bacteria; Proteobacteria; Alphaproteobacteria; Rhizobiales; Bradyrhizobiaceae; Bradyrhizobium | methyl-accepting chemotaxis protein partial | 6 |
| **WP_008969917.1** | 4.5E-07 | Qc.III | -REISGTIGRLSEISATIASAVEQQGAATQEISRNVQQTAEGTHRVSSNITE- | *Bradyrhizobium sp. STM 3843* | Bacteria; Proteobacteria; Alphaproteobacteria; Rhizobiales; Bradyrhizobiaceae; Bradyrhizobium | methyl-accepting chemotaxis protein | 6 |
| **AMA61232.1** | 4.6E-07 | Qa.IV | IKEISSTIERLSEISSAIAAAVEEQGAATQEIARNVQQAAQ------------ | *Bradyrhizobium sp. CCGE-LA001* | Bacteria; Proteobacteria; Alphaproteobacteria; Rhizobiales; Bradyrhizobiaceae; Bradyrhizobium | chemotaxis protein | 6 |
| **PCJ60477.1** | 4.6E-07 | Qa.III | IKGIGSTIKEINEIAASIASAVEEQGAATSEISRNVQEAA------------- | *Rhodospirillaceae bacterium* | Bacteria; Proteobacteria; Alphaproteobacteria; Rhodospirillales; Rhodospirillaceae | chemotaxis protein | 6 |
| **RAI19904.1** | 4.6E-07 | Qa.III | IESITGTIKQIDEISSTIASSVEEQGSATGEIAQNVQQAAEGTQEVSTSIVS- | *Afifella marina DSM 2698* | Bacteria; Proteobacteria; Alphaproteobacteria; Rhizobiales; Rhodobiaceae; Afifella | methyl-accepting chemotaxis protein | 6 |
| **WP_035717735.1** | 4.6E-07 | Qa.IV | IKEISSTIERLSEISSAIAAAVEEQGAATQEIARNVQQAAQ------------ | *Bradyrhizobium japonicum* | Bacteria; Proteobacteria; Alphaproteobacteria; Rhizobiales; Bradyrhizobiaceae; Bradyrhizobium | methyl-accepting chemotaxis protein | 6 |
| **WP_061020301.1** | 4.6E-07 | Qa.IV | IKEISSTIERLSEISSAIAAAVEEQGAATQEIARNVQQAAQ------------ | *Bradyrhizobium sp. CCH5-F6* | Bacteria; Proteobacteria; Alphaproteobacteria; Rhizobiales; Bradyrhizobiaceae; Bradyrhizobium | methyl-accepting chemotaxis protein | 6 |
| **SCY20531.1** | 4.7E-07 | Qa.IV | IQDIARTITEMSQISTSIAAAMEEQGAATAEIARNVQEAARVTETASENIA-- | *Microvirga guangxiensis* | Bacteria; Proteobacteria; Alphaproteobacteria; Rhizobiales; Methylobacteriaceae; Microvirga | methyl-accepting chemotaxis sensory transducer with Cache sensor | 6 |
| **WP_008967355.1** | 4.7E-07 | Qa.IV | IREIGTTIEKLSEISSTIAAAVEQQGAATQEISRNVQHAA------------- | *Bradyrhizobium sp. STM 3843* | Bacteria; Proteobacteria; Alphaproteobacteria; Rhizobiales; Bradyrhizobiaceae; Bradyrhizobium | methyl-accepting chemotaxis protein | 6 |
| **WP_011509355.1** | 4.7E-07 | Qa.IV | IREIGTTIKLISEISSAIAAAVEEQGAATQEIARNVQHAAQR----------- | *Nitrobacter hamburgensis* | Bacteria; Proteobacteria; Alphaproteobacteria; Rhizobiales; Bradyrhizobiaceae; Nitrobacter | methyl-accepting chemotaxis protein | 6 |
| **WP_017981122.1** | 4.7E-07 | Qa.IV | ---------EVNNAFLKIKEAIDKANETIVRIQESAKTSKENVETALRHIQ-- | *Methanocaldococcus villosus* | Archaea; Euryarchaeota; Methanococci; Methanococcales; Methanocaldococcaceae; Methanocaldococcus | hypothetical protein | 6 |
| **WP_017981148.1** | 4.7E-07 | Qa.IV | ---------EVNNAFLKIKEAIDKANETIVRIQESAKNSKENVETALRHIQ-- | *Methanocaldococcus villosus* | Archaea; Euryarchaeota; Methanococci; Methanococcales; Methanocaldococcaceae; Methanocaldococcus | hypothetical protein | 6 |
| **WP_028146888.1** | 4.7E-07 | Qa.IV | IKEISSTIERLSEISSAIAAAVEEQGAATQEIARNVQQAAQ------------ | *Bradyrhizobium japonicum* | Bacteria; Proteobacteria; Alphaproteobacteria; Rhizobiales; Bradyrhizobiaceae; Bradyrhizobium | methyl-accepting chemotaxis protein | 6 |
| **WP_041756412.1** | 4.7E-07 | Qa.IV | IKEISSTIEKLSEISSTIAAAVEQQGAATQEISRNVQQAA------------- | *Bradyrhizobium sp. ORS 278* | Bacteria; Proteobacteria; Alphaproteobacteria; Rhizobiales; Bradyrhizobiaceae; Bradyrhizobium | HAMP domain-containing protein | 6 |
| **AHB49398.1** | 4.8E-07 | Qa.IV | IKEITATINRMSEISGTIAAAVEEQGATTQEISRNVMEAA------------- | *Hyphomicrobium nitrativorans NL23* | Bacteria; Proteobacteria; Alphaproteobacteria; Rhizobiales; Hyphomicrobiaceae; Hyphomicrobium | methyl-accepting chemotaxis protein | 6 |
| **PZQ10881.1** | 4.8E-07 | Qc.III | IESIAGTIARINEISVAIASAVEQQSGVTREISVSMQVASQGVSAISGRMKD- | *Starkeya novella* | Bacteria; Proteobacteria; Alphaproteobacteria; Rhizobiales; Xanthobacteraceae; Starkeya | chemotaxis protein | 6 |
| **WP_009338269.1** | 4.8E-07 | Qa.III | IDEITQTIGRVNEIASAIASAVEEQGAATREIANNVSQAAE------------ | *Afipia sp. 1NLS2* | Bacteria; Proteobacteria; Alphaproteobacteria; Rhizobiales; Bradyrhizobiaceae; Afipia | methyl-accepting chemotaxis protein | 6 |
| **WP_057845453.1** | 4.8E-07 | Qa.IV | IRNISTTISEINDVTTAIAAAVEEQGAATREIARNIQHAA------------- | *Bradyrhizobium retamae* | Bacteria; Proteobacteria; Alphaproteobacteria; Rhizobiales; Bradyrhizobiaceae; Bradyrhizobium | HAMP domain-containing protein | 6 |
| **AWM00179.1** | 4.9E-07 | Qa.IV | IKEISSTIERLSEISSAIAAAVEEQGAATQEIARNVQQAAQ------------ | *Bradyrhizobium sp. 2 39S1MB* | Bacteria; Proteobacteria; Alphaproteobacteria; Rhizobiales; Bradyrhizobiaceae; Bradyrhizobium | methyl-accepting chemotaxis protein | 6 |
| **WP_008133438.1** | 4.9E-07 | Qa.IV | IKEISSTIERLSEISSAIAAAVEEQGAATQEIARNVQQAAQ------------ | *Bradyrhizobium sp. YR681* | Bacteria; Proteobacteria; Alphaproteobacteria; Rhizobiales; Bradyrhizobiaceae; Bradyrhizobium | methyl-accepting chemotaxis protein | 6 |
| **WP_014496890.1** | 4.9E-07 | Qa.IV | IKEISSTIERLSEISSAIAAAVEEQGAATQEIARNVQQAAQ------------ | *Bradyrhizobium japonicum* | Bacteria; Proteobacteria; Alphaproteobacteria; Rhizobiales; Bradyrhizobiaceae; Bradyrhizobium | methyl-accepting chemotaxis protein | 6 |
| **WP_018315627.1** | 4.9E-07 | Qa.IV | IKEISSTIERLSEISSAIAAAVEEQGAATQEIARNVQQAAQ------------ | *Bradyrhizobium sp. WSM2793* | Bacteria; Proteobacteria; Alphaproteobacteria; Rhizobiales; Bradyrhizobiaceae; Bradyrhizobium | methyl-accepting chemotaxis protein | 6 |
| **WP_018456982.1** | 4.9E-07 | Qa.IV | IKEISSTIERLSEISSAIAAAVEEQGAATQEIARNVQQAAQ------------ | *Bradyrhizobium sp. WSM4349* | Bacteria; Proteobacteria; Alphaproteobacteria; Rhizobiales; Bradyrhizobiaceae; Bradyrhizobium | methyl-accepting chemotaxis protein | 6 |
| **WP_018648080.1** | 4.9E-07 | Qa.IV | IKEISSTIERLSEISSAIAAAVEEQGAATQEIARNVQQAAQ------------ | *Bradyrhizobium japonicum* | Bacteria; Proteobacteria; Alphaproteobacteria; Rhizobiales; Bradyrhizobiaceae; Bradyrhizobium | methyl-accepting chemotaxis protein | 6 |
| **WP_024339676.1** | 4.9E-07 | Qa.IV | IKEISSTIERLSEISSAIAAAVEEQGAATQEIARNVQQAAQ------------ | *Bradyrhizobium japonicum* | Bacteria; Proteobacteria; Alphaproteobacteria; Rhizobiales; Bradyrhizobiaceae; Bradyrhizobium | methyl-accepting chemotaxis protein | 6 |
| **WP_027516790.1** | 4.9E-07 | Qa.IV | IKEISSTIERLSEISSAIAAAVEEQGAATQEIARNVQQAAQ------------ | *Bradyrhizobium sp. WSM1417* | Bacteria; Proteobacteria; Alphaproteobacteria; Rhizobiales; Bradyrhizobiaceae; Bradyrhizobium | chemotaxis protein | 6 |
| **WP_027550344.1** | 4.9E-07 | Qa.IV | IKEISSTIERLSEISSAIAAAVEEQGAATQEIARNVQQAAQ------------ | *Bradyrhizobium sp. Cp5.3* | Bacteria; Proteobacteria; Alphaproteobacteria; Rhizobiales; Bradyrhizobiaceae; Bradyrhizobium | methyl-accepting chemotaxis protein | 6 |
| **WP_027558624.1** | 4.9E-07 | Qa.IV | IKEISSTIERLSEISSAIAAAVEEQGAATQEIARNVQQAAQ------------ | *Bradyrhizobium* | Bacteria; Proteobacteria; Alphaproteobacteria; Rhizobiales; Bradyrhizobiaceae | MULTISPECIES: methyl-accepting chemotaxis protein | 6 |
| **WP_027568945.1** | 4.9E-07 | Qa.IV | IKEISSTIERLSEISSAIAAAVEEQGAATQEIARNVQQAAQ------------ | *Bradyrhizobium sp. URHA0013* | Bacteria; Proteobacteria; Alphaproteobacteria; Rhizobiales; Bradyrhizobiaceae; Bradyrhizobium | methyl-accepting chemotaxis protein | 6 |
| **WP_028135258.1** | 4.9E-07 | Qa.IV | IKEISSTIERLSEISSAIAAAVEEQGAATQEIARNVQQAAQ------------ | *Bradyrhizobium japonicum* | Bacteria; Proteobacteria; Alphaproteobacteria; Rhizobiales; Bradyrhizobiaceae; Bradyrhizobium | methyl-accepting chemotaxis protein | 6 |
| **WP_028143383.1** | 4.9E-07 | Qa.IV | IKEISSTIERLSEISSAIAAAVEEQGAATQEIARNVQQAAQ------------ | *Bradyrhizobium* | Bacteria; Proteobacteria; Alphaproteobacteria; Rhizobiales; Bradyrhizobiaceae | MULTISPECIES: methyl-accepting chemotaxis protein | 6 |
| **WP_028157151.1** | 4.9E-07 | Qa.IV | IKEISSTIERLSEISSAIAAAVEEQGAATQEIARNVQQAAQ------------ | *Bradyrhizobium japonicum* | Bacteria; Proteobacteria; Alphaproteobacteria; Rhizobiales; Bradyrhizobiaceae; Bradyrhizobium | methyl-accepting chemotaxis protein | 6 |
| **WP_028178349.1** | 4.9E-07 | Qa.IV | IKEISSTIERLSEISSAIAAAVEEQGAATQEIARNVQQAAQ------------ | *Bradyrhizobium* | Bacteria; Proteobacteria; Alphaproteobacteria; Rhizobiales; Bradyrhizobiaceae | MULTISPECIES: methyl-accepting chemotaxis protein | 6 |
| **WP_038942077.1** | 4.9E-07 | Qa.IV | IKEISSTIERLSEISSAIAAAVEEQGAATQEIARNVQQAAQ------------ | *Bradyrhizobium japonicum* | Bacteria; Proteobacteria; Alphaproteobacteria; Rhizobiales; Bradyrhizobiaceae; Bradyrhizobium | methyl-accepting chemotaxis protein | 6 |

| | | | | | | | |
|---|---|---|---|---|---|---|---|
| **WP_038957124.1** | 4.9E-07 | Qa.IV | IKEISSTIERLSEISSAIAAAVEEQGAATQEIARNVQQAAQ------------ | *Bradyrhizobium japonicum* | Bacteria; Proteobacteria; Alphaproteobacteria; Rhizobiales; Bradyrhizobiaceae; Bradyrhizobium | methyl-accepting chemotaxis protein | 6 |
| **WP_039151333.1** | 4.9E-07 | Qa.IV | IKEISSTIERLSEISSAIAAAVEEQGAATQEIARNVQQAAQ------------ | *Bradyrhizobium japonicum* | Bacteria; Proteobacteria; Alphaproteobacteria; Rhizobiales; Bradyrhizobiaceae; Bradyrhizobium | methyl-accepting chemotaxis protein | 6 |
| **WP_045009005.1** | 4.9E-07 | Qa.IV | IKEISSTIERLSEISSAIAAAVEEQGAATQEIARNVQQAAQ------------ | *Bradyrhizobium sp. LTSP857* | Bacteria; Proteobacteria; Alphaproteobacteria; Rhizobiales; Bradyrhizobiaceae; Bradyrhizobium | methyl-accepting chemotaxis protein | 6 |
| **WP_045010096.1** | 4.9E-07 | Qa.IV | IKEISSTIERLSEISSAIAAAVEEQGAATQEIARNVQQAAQ------------ | *Bradyrhizobium sp. LTSP849* | Bacteria; Proteobacteria; Alphaproteobacteria; Rhizobiales; Bradyrhizobiaceae; Bradyrhizobium | methyl-accepting chemotaxis protein | 6 |
| **WP_049820253.1** | 4.9E-07 | Qa.IV | IKEISSTIERLSEISSAIAAAVEEQGAATQEIARNVQQAAQ------------ | *Bradyrhizobium japonicum* | Bacteria; Proteobacteria; Alphaproteobacteria; Rhizobiales; Bradyrhizobiaceae; Bradyrhizobium | methyl-accepting chemotaxis protein | 6 |
| **WP_057027090.1** | 4.9E-07 | Qa.IV | IKEISSTIERLSEISSAIAAAVEEQGAATQEIARNVQQAAQ------------ | *Bradyrhizobium yuanmingense* | Bacteria; Proteobacteria; Alphaproteobacteria; Rhizobiales; Bradyrhizobiaceae; Bradyrhizobium | methyl-accepting chemotaxis protein | 6 |
| **WP_057753145.1** | 4.9E-07 | Qa.IV | IKEISSTIERLSEISSAIAAAVEEQGAATQEIARNVQQAAQ------------ | *Bradyrhizobium manausense* | Bacteria; Proteobacteria; Alphaproteobacteria; Rhizobiales; Bradyrhizobiaceae; Bradyrhizobium | methyl-accepting chemotaxis protein | 6 |
| **WP_060735483.1** | 4.9E-07 | Qa.IV | IKEISSTIERLSEISSAIAAAVEEQGAATQEIARNVQQAAQ------------ | *Bradyrhizobium sp. CCGE-LA001* | Bacteria; Proteobacteria; Alphaproteobacteria; Rhizobiales; Bradyrhizobiaceae; Bradyrhizobium | methyl-accepting chemotaxis protein | 6 |
| **WP_061878144.1** | 4.9E-07 | Qa.IV | IKEISSTIERLSEISSAIAAAVEEQGAATQEIARNVQQAAQ------------ | *Bradyrhizobium liaoningense* | Bacteria; Proteobacteria; Alphaproteobacteria; Rhizobiales; Bradyrhizobiaceae; Bradyrhizobium | methyl-accepting chemotaxis protein | 6 |
| **WP_063196145.1** | 4.9E-07 | Qa.IV | IKEISSTIERLSEISSAIAAAVEEQGAATQEIARNVQQAAQ------------ | *Bradyrhizobium sp. AT1* | Bacteria; Proteobacteria; Alphaproteobacteria; Rhizobiales; Bradyrhizobiaceae; Bradyrhizobium | methyl-accepting chemotaxis protein | 6 |
| **WP_063680813.1** | 4.9E-07 | Qa.IV | IKEISSTIERLSEISSAIAAAVEEQGAATQEIARNVQQAAQ------------ | *Bradyrhizobium neotropicale* | Bacteria; Proteobacteria; Alphaproteobacteria; Rhizobiales; Bradyrhizobiaceae; Bradyrhizobium | methyl-accepting chemotaxis protein | 6 |
| **WP_063691177.1** | 4.9E-07 | Qa.IV | IKEISSTIERLSEISSAIAAAVEEQGAATQEIARNVQQAAQ------------ | *Bradyrhizobium stylosanthis* | Bacteria; Proteobacteria; Alphaproteobacteria; Rhizobiales; Bradyrhizobiaceae; Bradyrhizobium | methyl-accepting chemotaxis protein | 6 |
| **WP_063709183.1** | 4.9E-07 | Qa.IV | IKEISSTIERLSEISSAIAAAVEEQGAATQEIARNVQQAAQ------------ | *Bradyrhizobium sp. BR 10245* | Bacteria; Proteobacteria; Alphaproteobacteria; Rhizobiales; Bradyrhizobiaceae; Bradyrhizobium | methyl-accepting chemotaxis protein | 6 |
| **WP_063984694.1** | 4.9E-07 | Qa.IV | IKEISSTIERLSEISSAIAAAVEEQGAATQEIARNVQQAAQ------------ | *Bradyrhizobium* | Bacteria; Proteobacteria; Alphaproteobacteria; Rhizobiales; Bradyrhizobiaceae | MULTISPECIES: methyl-accepting chemotaxis protein | 6 |
| **WP_063993163.1** | 4.9E-07 | Qa.IV | IKEISSTIERLSEISSAIAAAVEEQGAATQEIARNVQQAAQ------------ | *Bradyrhizobium sp.* | Bacteria; Proteobacteria; Alphaproteobacteria; Rhizobiales; Bradyrhizobiaceae; Bradyrhizobium | methyl-accepting chemotaxis protein | 6 |
| **WP_074119872.1** | 4.9E-07 | Qa.IV | IKEISSTIERLSEISSAIAAAVEEQGAATQEIARNVQQAAQ------------ | *Bradyrhizobium sp. AS23.2* | Bacteria; Proteobacteria; Alphaproteobacteria; Rhizobiales; Bradyrhizobiaceae; Bradyrhizobium | methyl-accepting chemotaxis protein | 6 |
| **WP_080136674.1** | 4.9E-07 | Qa.IV | IKEISSTIERLSEISSAIAAAVEEQGAATQEIARNVQQAAQ------------ | *Bradyrhizobium sp. BR10280* | Bacteria; Proteobacteria; Alphaproteobacteria; Rhizobiales; Bradyrhizobiaceae; Bradyrhizobium | methyl-accepting chemotaxis protein | 6 |
| **WP_083543340.1** | 4.9E-07 | Qa.IV | IKEISSTIERLSEISSAIAAAVEEQGAATQEIARNVQQAAQ------------ | *Bradyrhizobium sp. CCGE-LA001* | Bacteria; Proteobacteria; Alphaproteobacteria; Rhizobiales; Bradyrhizobiaceae; Bradyrhizobium | methyl-accepting chemotaxis protein | 6 |
| **WP_084803148.1** | 4.9E-07 | Qa.IV | IKEISSTIERLSEISSAIAAAVEEQGAATQEIARNVQQAAQ------------ | *Bradyrhizobium sp. NAS80.1* | Bacteria; Proteobacteria; Alphaproteobacteria; Rhizobiales; Bradyrhizobiaceae; Bradyrhizobium | methyl-accepting chemotaxis protein | 6 |
| **WP_085352178.1** | 4.9E-07 | Qa.IV | IKEISSTIERLSEISSAIAAAVEEQGAATQEIARNVQQAAQ------------ | *Bradyrhizobium canariense* | Bacteria; Proteobacteria; Alphaproteobacteria; Rhizobiales; Bradyrhizobiaceae; Bradyrhizobium | methyl-accepting chemotaxis protein | 6 |
| **WP_085362315.1** | 4.9E-07 | Qa.IV | IKEISSTIERLSEISSAIAAAVEEQGAATQEIARNVQQAAQ------------ | *Bradyrhizobium canariense* | Bacteria; Proteobacteria; Alphaproteobacteria; Rhizobiales; Bradyrhizobiaceae; Bradyrhizobium | methyl-accepting chemotaxis protein | 6 |
| **WP_085966299.1** | 4.9E-07 | Qa.IV | IKEISSTIERLSEISSAIAAAVEEQGAATQEIARNVQQAAQ------------ | *Bradyrhizobium liaoningense* | Bacteria; Proteobacteria; Alphaproteobacteria; Rhizobiales; Bradyrhizobiaceae; Bradyrhizobium | methyl-accepting chemotaxis protein | 6 |
| **WP_085967432.1** | 4.9E-07 | Qa.IV | IKEISSTIERLSEISSAIAAAVEEQGAATQEIARNVQQAAQ------------ | *Bradyrhizobium liaoningense* | Bacteria; Proteobacteria; Alphaproteobacteria; Rhizobiales; Bradyrhizobiaceae; Bradyrhizobium | methyl-accepting chemotaxis protein | 6 |
| **WP_090073454.1** | 4.9E-07 | Qa.I | IRSIAAKMSEISQYTTAISAAVQEQNASTNEISLNIQQAA------------- | *Cohaesibacter marisflavi* | Bacteria; Proteobacteria; Alphaproteobacteria; Rhizobiales; Cohaesibacteraceae | HAMP domain-containing protein | 6 |
| **WP_091895281.1** | 4.9E-07 | Qa.IV | IKEISSTIERLSEISSAIAAAVEEQGAATQEIARNVQQAAQ------------ | *Bradyrhizobium sp. Rc2d* | Bacteria; Proteobacteria; Alphaproteobacteria; Rhizobiales; Bradyrhizobiaceae; Bradyrhizobium | methyl-accepting chemotaxis protein | 6 |
| **WP_091953519.1** | 4.9E-07 | Qa.IV | IKEISSTIERLSEISSAIAAAVEEQGAATQEIARNVQQAAQ------------ | *Bradyrhizobium shewense* | Bacteria; Proteobacteria; Alphaproteobacteria; Rhizobiales; Bradyrhizobiaceae; Bradyrhizobium | methyl-accepting chemotaxis protein | 6 |
| **WP_092024365.1** | 4.9E-07 | Qa.IV | IKEISSTIERLSEISSAIAAAVEEQGAATQEIARNVQQAAQ------------ | *Bradyrhizobium sp. OK095* | Bacteria; Proteobacteria; Alphaproteobacteria; Rhizobiales; Bradyrhizobiaceae; Bradyrhizobium | methyl-accepting chemotaxis protein | 6 |
| **WP_092029119.1** | 4.9E-07 | Qa.IV | IKDISNTIERLSEISSAIAAAVEEQGAATQEISRNVQQAAE------------ | *Bradyrhizobium sp. OK095* | Bacteria; Proteobacteria; Alphaproteobacteria; Rhizobiales; Bradyrhizobiaceae; Bradyrhizobium | methyl-accepting chemotaxis protein | 6 |
| **WP_092186098.1** | 4.9E-07 | Qa.IV | IKEISSTIERLSEISSAIAAAVEEQGAATQEIARNVQQAAQ------------ | *Bradyrhizobium sp. cf659* | Bacteria; Proteobacteria; Alphaproteobacteria; Rhizobiales; Bradyrhizobiaceae; Bradyrhizobium | methyl-accepting chemotaxis protein | 6 |
| **WP_092216926.1** | 4.9E-07 | Qa.IV | IKEISSTIERLSEISSAIAAAVEEQGAATQEIARNVQQAAQ------------ | *Bradyrhizobium arachidis* | Bacteria; Proteobacteria; Alphaproteobacteria; Rhizobiales; Bradyrhizobiaceae; Bradyrhizobium | methyl-accepting chemotaxis protein | 6 |
| **WP_092259484.1** | 4.9E-07 | Qa.IV | IKEISSTIERLSEISSAIAAAVEEQGAATQEIARNVQQAAQ------------ | *Bradyrhizobium sp. Rc3b* | Bacteria; Proteobacteria; Alphaproteobacteria; Rhizobiales; Bradyrhizobiaceae; Bradyrhizobium | methyl-accepting chemotaxis protein | 6 |
| **WP_100178967.1** | 4.9E-07 | Qa.IV | IKEISSTIERLSEISSAIAAAVEEQGAATQEIARNVQQAAQ------------ | *Bradyrhizobium sp. TSA1* | Bacteria; Proteobacteria; Alphaproteobacteria; Rhizobiales; Bradyrhizobiaceae; Bradyrhizobium | methyl-accepting chemotaxis protein | 6 |
| **WP_100230110.1** | 4.9E-07 | Qa.IV | IKEISSTIERLSEISSAIAAAVEEQGAATQEIARNVQQAAQ------------ | *Bradyrhizobium sp. INPA54B* | Bacteria; Proteobacteria; Alphaproteobacteria; Rhizobiales; Bradyrhizobiaceae; Bradyrhizobium | methyl-accepting chemotaxis protein | 6 |
| **WP_104464683.1** | 4.9E-07 | Qa.IV | IKEISSTIERLSEISSAIAAAVEEQGAATQEIARNVQQAAQ------------ | *Bradyrhizobium sp. AC87j1* | Bacteria; Proteobacteria; Alphaproteobacteria; Rhizobiales; Bradyrhizobiaceae; Bradyrhizobium | methyl-accepting chemotaxis protein | 6 |
| **WP_106942588.1** | 4.9E-07 | Qa.IV | IKEISSTIERLSEISSAIAAAVEEQGAATQEIARNVQQAAQ------------ | *Bradyrhizobium sp. MOS002* | Bacteria; Proteobacteria; Alphaproteobacteria; Rhizobiales; Bradyrhizobiaceae; Bradyrhizobium | methyl-accepting chemotaxis protein | 6 |
| **WP_109117286.1** | 4.9E-07 | Qa.IV | IVDIVQTIAEINQIATTIAAAVEQQDAATREIARNVQQAA------------- | *Azospirillum sp. TSO22-1* | Bacteria; Proteobacteria; Alphaproteobacteria; Rhodospirillales; Rhodospirillaceae; Azospirillum | methyl-accepting chemotaxis protein | 6 |
| **AWM06437.1** | 5E-07 | Qa.IV | IKEISSTIERLSEISSAIAAAVEEQGAATQEIARNVQQAAQ------------ | *Bradyrhizobium sp. 3 85S1MB* | Bacteria; Proteobacteria; Alphaproteobacteria; Rhizobiales; Bradyrhizobiaceae; Bradyrhizobium | methyl-accepting chemotaxis protein | 6 |
| **AWM06438.1** | 5E-07 | Qa.IV | IKEISSTIERLSEISSAIAAAVEEQGAATQEIARNVQQAAQ------------ | *Bradyrhizobium sp. 3 85S1MB* | Bacteria; Proteobacteria; Alphaproteobacteria; Rhizobiales; Bradyrhizobiaceae; Bradyrhizobium | methyl-accepting chemotaxis protein | 6 |
| **WP_028172386.1** | 5E-07 | Qa.IV | IKEISSTIERLSEISSAIAAAVEEQGAATQEIARNVQQAAQ------------ | *Bradyrhizobium* | Bacteria; Proteobacteria; Alphaproteobacteria; Rhizobiales; Bradyrhizobiaceae | MULTISPECIES: methyl-accepting chemotaxis protein | 6 |
| **WP_085899985.1** | 5E-07 | Qa.III | IRSITGTISEVNDIAQTIGDSIEQQDLATREISQNVQETAINTTEASNE---- | *Kiloniella majae* | Bacteria; Proteobacteria; Alphaproteobacteria; Kiloniellales; Kiloniellaceae; Kiloniella | methyl-accepting chemotaxis protein | 6 |
| **WP_085905195.1** | 5E-07 | Qa.III | IRSITGTISEVNDIAQTIGDSIEQQDLATREISQNVQETAINTTEASNE---- | *Kiloniella majae* | Bacteria; Proteobacteria; Alphaproteobacteria; Kiloniellales; Kiloniellaceae; Kiloniella | methyl-accepting chemotaxis protein | 6 |
| **WP_085909487.1** | 5E-07 | Qa.III | IRSITGTISEVNDIAQTIGDSIEQQDLATREISQNVQETAINTTEASNE---- | *Kiloniella majae* | Bacteria; Proteobacteria; Alphaproteobacteria; Kiloniellales; Kiloniellaceae; Kiloniella | methyl-accepting chemotaxis protein | 6 |
| **WP_106949977.1** | 5E-07 | Qa.IV | IKEISSTIERLSEISSAIAAAVEEQGAATQEIARNVQQAAQ------------ | *Bradyrhizobium sp. MOS003* | Bacteria; Proteobacteria; Alphaproteobacteria; Rhizobiales; Bradyrhizobiaceae; Bradyrhizobium | methyl-accepting chemotaxis protein | 6 |
| **WP_106949978.1** | 5E-07 | Qa.IV | IKEISSTIERLSEISSAIAAAVEEQGAATQEIARNVQQAAQ------------ | *Bradyrhizobium sp. MOS003* | Bacteria; Proteobacteria; Alphaproteobacteria; Rhizobiales; Bradyrhizobiaceae; Bradyrhizobium | methyl-accepting chemotaxis protein | 6 |
| **OYU88815.1** | 5.1E-07 | Qa.IV | IRNIGQTITEINEVSTAIATAVEQQGAATREIARNIQHAA------------- | *Bradyrhizobiaceae bacterium PARB1* | Bacteria; Proteobacteria; Alphaproteobacteria; Rhizobiales; Bradyrhizobiaceae | methyl-accepting chemotaxis protein | 6 |
| **PLX53986.1** | 5.1E-07 | Qa.III | IDQISKIINDVNEIVTTIASAVEEQSVTTKEIAGNVAQASQGIQE-------- | *Desulfobacteraceae bacterium* | Bacteria; Proteobacteria; Deltaproteobacteria; Desulfobacterales; Desulfobacteraceae | hypothetical protein C0611_02360 | 6 |
| **WP_008138054.1** | 5.1E-07 | Qa.IV | IRDISATIERLSEVSSTIAAAVEEQGAATQEISRNVQQAA------------- | *Bradyrhizobium sp. YR681* | Bacteria; Proteobacteria; Alphaproteobacteria; Rhizobiales; Bradyrhizobiaceae; Bradyrhizobium | methyl-accepting chemotaxis protein | 6 |
| **WP_014494370.1** | 5.1E-07 | Qa.IV | IRDISATIERLSEVSSTIAAAVEEQGAATQEISRNVQQAA------------- | *Bradyrhizobium japonicum* | Bacteria; Proteobacteria; Alphaproteobacteria; Rhizobiales; Bradyrhizobiaceae; Bradyrhizobium | methyl-accepting chemotaxis protein | 6 |
| **WP_022774009.1** | 5.1E-07 | Qb.I | ---ITNLIKDLAEKSQANMEAITNSSGAVSKAGDSFSKINQELNSTA------ | *Butyrivibrio sp. AE2015* | Bacteria; Firmicutes; Clostridia; Clostridiales; Lachnospiraceae; Butyrivibrio | methyl-accepting chemotaxis protein | 6 |
| **WP_022779070.1** | 5.1E-07 | Qb.I | ---ITNLIKDLAEKSQANMEAITNSSGAVSKAGDSFSKINQELNSTA------ | *Butyrivibrio sp. AE3009* | Bacteria; Firmicutes; Clostridia; Clostridiales; Lachnospiraceae; Butyrivibrio | methyl-accepting chemotaxis protein | 6 |
| **WP_026490097.1** | 5.1E-07 | Qb.I | ---ITNLIKDLAEKSQANMEAITNSSGAVSKAGDSFSKINQELNSTA------ | *Butyrivibrio sp. XBB1001* | Bacteria; Firmicutes; Clostridia; Clostridiales; Lachnospiraceae; Butyrivibrio | methyl-accepting chemotaxis protein | 6 |
| **WP_026516404.1** | 5.1E-07 | Qb.I | ---ITNLIKDLAEKSQANMEAITNSSGAVSKAGDSFSKINQELNSTA------ | *Butyrivibrio sp. MC2021* | Bacteria; Firmicutes; Clostridia; Clostridiales; Lachnospiraceae; Butyrivibrio | methyl-accepting chemotaxis protein | 6 |
| **WP_027533849.1** | 5.1E-07 | Qa.IV | IRDISATIERLSEVSSTIAAAVEEQGAATQEISRNVQQAA------------- | *Bradyrhizobium sp. WSM3983* | Bacteria; Proteobacteria; Alphaproteobacteria; Rhizobiales; Bradyrhizobiaceae; Bradyrhizobium | methyl-accepting chemotaxis protein | 6 |
| **WP_028150215.1** | 5.1E-07 | Qa.IV | IRDISATIERLSEVSSTIAAAVEEQGAATQEISRNVQQAA------------- | *Bradyrhizobium japonicum* | Bacteria; Proteobacteria; Alphaproteobacteria; Rhizobiales; Bradyrhizobiaceae; Bradyrhizobium | methyl-accepting chemotaxis protein | 6 |
| **WP_028154791.1** | 5.1E-07 | Qa.IV | IRDISATIERLSEVSSTIAAAVEEQGAATQEISRNVQQAA------------- | *Bradyrhizobium japonicum* | Bacteria; Proteobacteria; Alphaproteobacteria; Rhizobiales; Bradyrhizobiaceae; Bradyrhizobium | methyl-accepting chemotaxis protein | 6 |
| **WP_038935981.1** | 5.1E-07 | Qa.IV | IRDISATIERLSEVSSTIAAAVEEQGAATQEISRNVQQAA------------- | *Bradyrhizobium japonicum* | Bacteria; Proteobacteria; Alphaproteobacteria; Rhizobiales; Bradyrhizobiaceae; Bradyrhizobium | methyl-accepting chemotaxis protein | 6 |
| **WP_039147377.1** | 5.1E-07 | Qa.IV | IRDISATIERLSEVSSTIAAAVEEQGAATQEISRNVQQAA------------- | *Bradyrhizobium japonicum* | Bacteria; Proteobacteria; Alphaproteobacteria; Rhizobiales; Bradyrhizobiaceae; Bradyrhizobium | methyl-accepting chemotaxis protein | 6 |
| **WP_041958445.1** | 5.1E-07 | Qa.IV | IRDISATIERLSEVSSTIAAAVEEQGAATQEISRNVQQAA------------- | *Bradyrhizobium japonicum* | Bacteria; Proteobacteria; Alphaproteobacteria; Rhizobiales; Bradyrhizobiaceae; Bradyrhizobium | methyl-accepting chemotaxis protein | 6 |
| **WP_063982528.1** | 5.1E-07 | Qa.IV | IRDISATIERLSEVSSTIAAAVEEQGAATQEISRNVQQAA------------- | *Bradyrhizobium sp.* | Bacteria; Proteobacteria; Alphaproteobacteria; Rhizobiales; Bradyrhizobiaceae; Bradyrhizobium | methyl-accepting chemotaxis protein | 6 |
| **WP_071912709.1** | 5.1E-07 | Qa.IV | IRDISATIERLSEVSSTIAAAVEEQGAATQEISRNVQQAA------------- | *Bradyrhizobium japonicum* | Bacteria; Proteobacteria; Alphaproteobacteria; Rhizobiales; Bradyrhizobiaceae; Bradyrhizobium | methyl-accepting chemotaxis protein | 6 |
| **WP_074883783.1** | 5.1E-07 | Qb.I | ---ITNLIKDLAEKSQANMEAITNSSGAVSKAGDSFSKINQELNSTA------ | *Butyrivibrio proteoclasticus* | Bacteria; Firmicutes; Clostridia; Clostridiales; Lachnospiraceae; Butyrivibrio | methyl-accepting chemotaxis protein | 6 |

| | | | | | | | | |
|---|---|---|---|---|---|---|---|---|
|  | **WP_085405356.1** | 5.1E-07 | Qa.IV | IRDISATIERLSEVSSTIAAAVEEQGAATQEISRNVQQAA------------- | *Bradyrhizobium japonicum* | Bacteria; Proteobacteria; Alphaproteobacteria; Rhizobiales; Bradyrhizobiaceae; Bradyrhizobium | methyl-accepting chemotaxis protein | 6 |
|  | **WP_092028411.1** | 5.1E-07 | Qa.IV | IRDISATIERLSEVSSTIAAAVEEQGAATQEISRNVQQAA------------- | *Bradyrhizobium sp. OK095* | Bacteria; Proteobacteria; Alphaproteobacteria; Rhizobiales; Bradyrhizobiaceae; Bradyrhizobium | methyl-accepting chemotaxis protein | 6 |
|  | **WP_092228646.1** | 5.1E-07 | Qa.IV | IRDISATIERLSEVSSTIAAAVEEQGAATQEISRNVQQAA------------- | *Bradyrhizobium sp. Gha* | Bacteria; Proteobacteria; Alphaproteobacteria; Rhizobiales; Bradyrhizobiaceae; Bradyrhizobium | methyl-accepting chemotaxis protein | 6 |
|  | **WP_092294954.1** | 5.1E-07 | Qa.IV | IRDISATIERLSEVSSTIAAAVEEQGAATQEISRNVQQAA------------- | *Bradyrhizobium sp. Ghvi* | Bacteria; Proteobacteria; Alphaproteobacteria; Rhizobiales; Bradyrhizobiaceae; Bradyrhizobium | methyl-accepting chemotaxis protein | 6 |
|  | **WP_092329976.1** | 5.1E-07 | Qb.I | ---ITNLIKDLAEKSQANMEAITNSSGAVSKAGDSFSKINQELNSTA------ | *Butyrivibrio sp. YAB3001* | Bacteria; Firmicutes; Clostridia; Clostridiales; Lachnospiraceae; Butyrivibrio | methyl-accepting chemotaxis protein | 6 |
|  | **WP_100418893.1** | 5.1E-07 | Qa.IV | -KEISGTIERLSEISSTIAAAVEEQGAATQEISRNVQQAAQ------------ | *Bradyrhizobium lablabi* | Bacteria; Proteobacteria; Alphaproteobacteria; Rhizobiales; Bradyrhizobiaceae; Bradyrhizobium | HAMP domain-containing protein | 6 |
|  | **WP_105645351.1** | 5.1E-07 | Qa.III | LDGIRQAIERVNDMAGQIASAAEEQSAVVDEINQNITNIA------------- | *Pseudomonas sp. MYb185* | Bacteria; Proteobacteria; Gammaproteobacteria; Pseudomonadales; Pseudomonadaceae; Pseudomonas | PAS domain S-box protein | 6 |
|  | **WP_002718258.1** | 5.2E-07 | Qa.III | IDEITQTIGRVNEIASAIASAVEEQGAATREIANNVSQAAE------------ | *Afipia felis* | Bacteria; Proteobacteria; Alphaproteobacteria; Rhizobiales; Bradyrhizobiaceae; Afipia | methyl-accepting chemotaxis protein | 6 |
|  | **WP_002727409.1** | 5.2E-07 | Qa.IV | -----QTIRDLNELSSAIADAIEQQSAATAEISRNVEEASTNTTQT------- | *Phaeospirillum molischianum* | Bacteria; Proteobacteria; Alphaproteobacteria; Rhodospirillales; Rhodospirillaceae; Phaeospirillum | methyl-accepting chemotaxis protein | 6 |
|  | **WP_027539079.1** | 5.2E-07 | Qa.IV | -KEISGTIERLSEISSTIAAAVEEQGAATQEISRNVQQAAQ------------ | *Bradyrhizobium sp. URHA0002* | Bacteria; Proteobacteria; Alphaproteobacteria; Rhizobiales; Bradyrhizobiaceae; Bradyrhizobium | methyl-accepting chemotaxis protein | 6 |
|  | **WP_045008314.1** | 5.2E-07 | Qa.IV | IRDISATIERLSEVSSTIAAAVEEQGAATQEIARNVQQAA------------- | *Bradyrhizobium* | Bacteria; Proteobacteria; Alphaproteobacteria; Rhizobiales; Bradyrhizobiaceae | MULTISPECIES: methyl-accepting chemotaxis protein | 6 |
|  | **WP_051957725.1** | 5.2E-07 | Qa.III | IKDISGIIQNVNEIVSGIATAVEEQSVTTGEIAENMNRTSLKMQEVN------ | *Desulfobacter vibrioformis* | Bacteria; Proteobacteria; Deltaproteobacteria; Desulfobacterales; Desulfobacteraceae; Desulfobacter | methyl-accepting chemotaxis protein | 6 |
|  | **WP_058545205.1** | 5.2E-07 | Qb.I | LHRTKQHLQETIEQSELSSEELKASNEEMQAVNEELRSATEELETS------- | *Pseudomonas fluorescens* | Bacteria; Proteobacteria; Gammaproteobacteria; Pseudomonadales; Pseudomonadaceae; Pseudomonas | PAS domain S-box protein | 6 |
|  | **WP_063702910.1** | 5.2E-07 | Qa.IV | IRDISATIERLSEVSSTIAAAVEEQGAATQEIARNVQQAA------------- | *Bradyrhizobium sp. BR 10245* | Bacteria; Proteobacteria; Alphaproteobacteria; Rhizobiales; Bradyrhizobiaceae; Bradyrhizobium | methyl-accepting chemotaxis protein | 6 |
|  | **WP_074123342.1** | 5.2E-07 | Qa.IV | IRDISATIERLSEVSSTIAAAVEEQGAATQEIARNVQQAA------------- | *Bradyrhizobium sp. AS23.2* | Bacteria; Proteobacteria; Alphaproteobacteria; Rhizobiales; Bradyrhizobiaceae; Bradyrhizobium | methyl-accepting chemotaxis protein | 6 |
|  | **WP_083843533.1** | 5.2E-07 | Qa.III | IEEISAVIADVNDIVSTIATAVEEQSSATREIAQNIAQASSGIEEVN------ | *Desulfobacter postgatei* | Bacteria; Proteobacteria; Deltaproteobacteria; Desulfobacterales; Desulfobacteraceae; Desulfobacter | methyl-accepting chemotaxis protein | 6 |
|  | **WP_100419358.1** | 5.2E-07 | Qa.IV | -KEISGTIERLSEISSTIAAAVEEQGAATQEISRNVQQAAQ------------ | *Bradyrhizobium lablabi* | Bacteria; Proteobacteria; Alphaproteobacteria; Rhizobiales; Bradyrhizobiaceae; Bradyrhizobium | methyl-accepting chemotaxis protein | 6 |
|  | **WP_103012541.1** | 5.2E-07 | Qa.IV | IREIGSTIERMSEISSSIASAVEQQGAATQEIARNVQEAA------------- | *Rhodopseudomonas palustris* | Bacteria; Proteobacteria; Alphaproteobacteria; Rhizobiales; Bradyrhizobiaceae; Rhodopseudomonas | methyl-accepting chemotaxis protein | 6 |
|  | **PIE65048.1** | 5.3E-07 | SNAP.b | LNNMLKRIDGDRERLRQTIVSLEENREKLIRTQQEMVRAEKLASV-------- | *Desulfobacterales bacterium* | Bacteria; Proteobacteria; Deltaproteobacteria; Desulfobacterales | hypothetical protein CSA26_05225 | 6 |
| | **OYX69250.1** | 5.4E-07 | Qc.III | IERVGAQIRKMSDVANEISAAVEEQGMATKEIVRNVDQAATGTNSVTSHISD- | *Rhizobiales bacterium 32-66-11* | Bacteria; Proteobacteria; Alphaproteobacteria; Rhizobiales | hypothetical protein B7Y95_18645 | 6 |
|  | **WP_027279520.1** | 5.4E-07 | Qa.IV | IKEISDTIAQMSEISSTIAAAVEEQGAATQEISRNVQQAA------------- | *Rhodopseudomonas palustris* | Bacteria; Proteobacteria; Alphaproteobacteria; Rhizobiales; Bradyrhizobiaceae; Rhodopseudomonas | methyl-accepting chemotaxis protein | 6 |
|  | **WP_054700599.1** | 5.4E-07 | Qa.III | IRMISAVIVEVNEIVTGIAAAVEEQSTATQEIASNVSQASQGIQE-------- | *Desulfosarcina cetonica* | Bacteria; Proteobacteria; Deltaproteobacteria; Desulfobacterales; Desulfobacteraceae; Desulfosarcina | hypothetical protein | 6 |
|  | **WP_081674462.1** | 5.4E-07 | Qb.I | ---ITNLIKDLAEKSQANMEAITNSSGAVSKAGDSFSKINQELNSTA------ | *Butyrivibrio sp. VCD2006* | Bacteria; Firmicutes; Clostridia; Clostridiales; Lachnospiraceae; Butyrivibrio | methyl-accepting chemotaxis protein | 6 |
|  | **WP_103011439.1** | 5.4E-07 | Qa.IV | IKEISDTIAQMSEISSTIAAAVEEQGAATQEISRNVQQAA------------- | *Rhodopseudomonas palustris* | Bacteria; Proteobacteria; Alphaproteobacteria; Rhizobiales; Bradyrhizobiaceae; Rhodopseudomonas | methyl-accepting chemotaxis protein | 6 |
|  | **WP_106944642.1** | 5.4E-07 | Qa.IV | IRDISATIERLSEVSSTIAAAVEEQGAATQEISRNVQQAA------------- | *Bradyrhizobium sp. MOS002* | Bacteria; Proteobacteria; Alphaproteobacteria; Rhizobiales; Bradyrhizobiaceae; Bradyrhizobium | methyl-accepting chemotaxis protein | 6 |
|  | **WP_110780922.1** | 5.4E-07 | Qa.IV | IKEISDTIAQMSEISSTIAAAVEEQGAATQEISRNVQQAA------------- | *Rhodopseudomonas faecalis* | Bacteria; Proteobacteria; Alphaproteobacteria; Rhizobiales; Bradyrhizobiaceae; Rhodopseudomonas | methyl-accepting chemotaxis protein | 6 |
|  | **WP_002714793.1** | 5.5E-07 | Qa.IV | IKEISSTINLMSEISSTIAAAVEEQGAATQEIARNVQQAAE------------ | *Afipia clevelandensis* | Bacteria; Proteobacteria; Alphaproteobacteria; Rhizobiales; Bradyrhizobiaceae; Afipia | methyl-accepting chemotaxis protein | 6 |
|  | **WP_008551941.1** | 5.5E-07 | Qa.III | ISAITRTMADVNDITATIAAAVEQQGAATTEISENVQQAAVGTQN-------- | *Pseudovibrio sp. JE062* | Bacteria; Proteobacteria; Alphaproteobacteria; Rhodobacterales; Rhodobacteraceae; Pseudovibrio | methyl-accepting chemotaxis protein | 6 |
|  | **WP_009031126.1** | 5.5E-07 | Qa.IV | IKEISSTIEKLSEISSTIAAAVEEQGAATQEISRNVQQAA------------- | *Bradyrhizobium sp. ORS 375* | Bacteria; Proteobacteria; Alphaproteobacteria; Rhizobiales; Bradyrhizobiaceae; Bradyrhizobium | HAMP domain-containing protein | 6 |
|  | **WP_009735415.1** | 5.5E-07 | Qa.IV | IKEISSTINLMSEISSTIAAAVEEQGAATQEIARNVQQAAE------------ | *Bradyrhizobiaceae bacterium SG-6C* | Bacteria; Proteobacteria; Alphaproteobacteria; Rhizobiales; Bradyrhizobiaceae | methyl-accepting chemotaxis protein | 6 |
|  | **WP_018647155.1** | 5.5E-07 | Qa.IV | IRDISATIERLSEVSSTIAAAVEEQGAATQEISRNVQQAA------------- | *Bradyrhizobium japonicum* | Bacteria; Proteobacteria; Alphaproteobacteria; Rhizobiales; Bradyrhizobiaceae; Bradyrhizobium | methyl-accepting chemotaxis protein | 6 |
|  | **WP_027549083.1** | 5.5E-07 | Qa.IV | IRDISATIERLSEVSSTIAAAVEEQGAATQEISRNVQQAA------------- | *Bradyrhizobium sp. WSM2254* | Bacteria; Proteobacteria; Alphaproteobacteria; Rhizobiales; Bradyrhizobiaceae; Bradyrhizobium | methyl-accepting chemotaxis protein | 6 |
|  | **WP_027554608.1** | 5.5E-07 | Qa.IV | IRDISATIERLSEVSSTIAAAVEEQGAATQEIARNVQQAA------------- | *Bradyrhizobium sp. Cp5.3* | Bacteria; Proteobacteria; Alphaproteobacteria; Rhizobiales; Bradyrhizobiaceae; Bradyrhizobium | methyl-accepting chemotaxis protein | 6 |
|  | **WP_027560013.1** | 5.5E-07 | Qa.IV | IRDISATIERLSEVSSTIAAAVEEQGAATQEISRNVQQAA------------- | *Bradyrhizobium* | Bacteria; Proteobacteria; Alphaproteobacteria; Rhizobiales; Bradyrhizobiaceae | MULTISPECIES: methyl-accepting chemotaxis protein | 6 |
|  | **WP_028350031.1** | 5.5E-07 | Qa.IV | IQAISSTIEKLSEISSTIAAAVEEQGAATQEISRNVQQAA------------- | *Bradyrhizobium elkanii* | Bacteria; Proteobacteria; Alphaproteobacteria; Rhizobiales; Bradyrhizobiaceae; Bradyrhizobium | HAMP domain-containing protein | 6 |
|  | **WP_057847170.1** | 5.5E-07 | Qa.IV | IQAISSTIEKLSEISSTIAAAVEEQGAATQEISRNVQQAA------------- | *Bradyrhizobium retamae* | Bacteria; Proteobacteria; Alphaproteobacteria; Rhizobiales; Bradyrhizobiaceae; Bradyrhizobium | HAMP domain-containing protein | 6 |
|  | **WP_057854098.1** | 5.5E-07 | Qa.IV | IQAISSTIEKLSEISSTIAAAVEEQGAATQEISRNVQQAA------------- | *Bradyrhizobium valentinum* | Bacteria; Proteobacteria; Alphaproteobacteria; Rhizobiales; Bradyrhizobiaceae; Bradyrhizobium | HAMP domain-containing protein | 6 |
|  | **WP_057863451.1** | 5.5E-07 | Qa.IV | IQAISSTIEKLSEISSTIAAAVEEQGAATQEISRNVQQAA------------- | *Bradyrhizobium lablabi* | Bacteria; Proteobacteria; Alphaproteobacteria; Rhizobiales; Bradyrhizobiaceae; Bradyrhizobium | HAMP domain-containing protein | 6 |
|  | **WP_057902535.1** | 5.5E-07 | Qa.IV | IQAISSTIEKLSEISSTIAAAVEEQGAATQEISRNVQQAA------------- | *Bradyrhizobium valentinum* | Bacteria; Proteobacteria; Alphaproteobacteria; Rhizobiales; Bradyrhizobiaceae; Bradyrhizobium | HAMP domain-containing protein | 6 |
|  | **WP_063675797.1** | 5.5E-07 | Qa.IV | IRDISATIERLSEVSSTIAAAVEEQGAATQEIARNVQQAA------------- | *Bradyrhizobium neotropicale* | Bacteria; Proteobacteria; Alphaproteobacteria; Rhizobiales; Bradyrhizobiaceae; Bradyrhizobium | methyl-accepting chemotaxis protein | 6 |
|  | **WP_063682649.1** | 5.5E-07 | Qa.IV | IRDISATIERLSEVSSTIAAAVEEQGAATQEISRNVQQAA------------- | *Bradyrhizobium stylosanthis* | Bacteria; Proteobacteria; Alphaproteobacteria; Rhizobiales; Bradyrhizobiaceae; Bradyrhizobium | methyl-accepting chemotaxis protein | 6 |
|  | **WP_065733043.1** | 5.5E-07 | Qa.IV | IQAISSTIEKLSEISSTIAAAVEEQGAATQEISRNVQQAA------------- | *Bradyrhizobium icense* | Bacteria; Proteobacteria; Alphaproteobacteria; Rhizobiales; Bradyrhizobiaceae; Bradyrhizobium | methyl-accepting chemotaxis protein | 6 |
|  | **WP_074280698.1** | 5.5E-07 | Qa.IV | -KEISGTIERLSEISSTIAAAVEEQGAATQEISRNVQQASQ------------ | *Bradyrhizobium erythrophlei* | Bacteria; Proteobacteria; Alphaproteobacteria; Rhizobiales; Bradyrhizobiaceae; Bradyrhizobium | methyl-accepting chemotaxis protein | 6 |
|  | **WP_075697255.1** | 5.5E-07 | Qa.III | ISAITRTMADVNDITATIAAAVEQQGAATTEISENVQQAAVGTQN-------- | *Pseudovibrio sp. Ab134* | Bacteria; Proteobacteria; Alphaproteobacteria; Rhodobacterales; Rhodobacteraceae; Pseudovibrio | methyl-accepting chemotaxis protein | 6 |
|  | **WP_080134358.1** | 5.5E-07 | Qa.IV | IRDISATIERLSEVSSTIAAAVEEQGAATQEISRNVQQAA------------- | *Bradyrhizobium sp. BR10280* | Bacteria; Proteobacteria; Alphaproteobacteria; Rhizobiales; Bradyrhizobiaceae; Bradyrhizobium | methyl-accepting chemotaxis protein | 6 |
|  | **WP_083416486.1** | 5.5E-07 | Qa.III | ISAITRTMADVNDITATIAAAVEQQGAATTEISENVQQAAVGTQN-------- | *Pseudovibrio denitrificans* | Bacteria; Proteobacteria; Alphaproteobacteria; Rhodobacterales; Rhodobacteraceae; Pseudovibrio | methyl-accepting chemotaxis protein | 6 |
|  | **WP_085590576.1** | 5.5E-07 | Qc.III | -SEIGKIIENINEIATTISAAVEEQSAATNEISRNVRGAAER---------- | *Thalassospira sp. MCCC 1A01428* | Bacteria; Proteobacteria; Alphaproteobacteria; Rhodospirillales; Rhodospirillaceae; Thalassospira | HAMP domain-containing protein | 6 |
|  | **WP_092213726.1** | 5.5E-07 | Qa.IV | IRDISATIERLSEVSSTIAAAVEEQGAATQEISRNVQQAA------------- | *Bradyrhizobium arachidis* | Bacteria; Proteobacteria; Alphaproteobacteria; Rhizobiales; Bradyrhizobiaceae; Bradyrhizobium | methyl-accepting chemotaxis protein | 6 |
|  | **RDD60364.1** | 5.6E-07 | Qa.III | IERIERSVKEIDEAASSIASAIEEQSSATREITRNVNEASTGTQEVNKN---- | *Rhodospirillaceae bacterium* | Bacteria; Proteobacteria; Alphaproteobacteria; Rhodospirillales; Rhodospirillaceae | HAMP domain-containing protein | 6 |
|  | **WP_019195975.1** | 5.6E-07 | Qa.IV | IKEISATINLMSEISSTIAAAVEEQGAATQEIARNVQQAAE------------ | *Afipia birgiae* | Bacteria; Proteobacteria; Alphaproteobacteria; Rhizobiales; Bradyrhizobiaceae; Afipia | methyl-accepting chemotaxis protein | 6 |
|  | **WP_027518664.1** | 5.6E-07 | Qa.IV | IRDISTTIERLSEVSSTIAAAVEEQGAATQEISRNVQQAA------------- | *Bradyrhizobium sp. WSM1417* | Bacteria; Proteobacteria; Alphaproteobacteria; Rhizobiales; Bradyrhizobiaceae; Bradyrhizobium | methyl-accepting chemotaxis protein | 6 |
|  | **WP_027568436.1** | 5.6E-07 | Qa.IV | IRDISTTIERLSEVSSTIAAAVEEQGAATQEISRNVQQAA------------- | *Bradyrhizobium sp. URHA0013* | Bacteria; Proteobacteria; Alphaproteobacteria; Rhizobiales; Bradyrhizobiaceae; Bradyrhizobium | methyl-accepting chemotaxis protein | 6 |
|  | **WP_079566119.1** | 5.6E-07 | Qa.IV | -KEISGTIERLSEISSTIAAAVEEQGAATQEISRNVQQAAQ------------ | *Bradyrhizobium erythrophlei* | Bacteria; Proteobacteria; Alphaproteobacteria; Rhizobiales; Bradyrhizobiaceae; Bradyrhizobium | methyl-accepting chemotaxis protein | 6 |
|  | **WP_084807233.1** | 5.7E-07 | Qa.IV | IRDISATIERLSEVSSTIAAAVEEQGAATQEIARNVQQAA------------- | *Bradyrhizobium sp. NAS80.1* | Bacteria; Proteobacteria; Alphaproteobacteria; Rhizobiales; Bradyrhizobiaceae; Bradyrhizobium | methyl-accepting chemotaxis protein | 6 |
|  | **WP_092291366.1** | 5.7E-07 | Qa.IV | IKAISETIERLSEISSTIAAAVEEQGAATQEIARNVQQAA------------- | *Bradyrhizobium sp. Ghvi* | Bacteria; Proteobacteria; Alphaproteobacteria; Rhizobiales; Bradyrhizobiaceae; Bradyrhizobium | methyl-accepting chemotaxis protein | 6 |
|  | **WP_092619778.1** | 5.7E-07 | Qc.III | IEDIAQVINRIDEIAQAISAAVEEQGAATREIADSTQRASGS----------- | *Roseospirillum parvum* | Bacteria; Proteobacteria; Alphaproteobacteria; Rhizobiales; Rhodobiaceae; Roseospirillum | methyl-accepting chemotaxis protein | 6 |
|  | **WP_036142990.1** | 5.8E-07 | Qb.I | LHRTKERMQATIEQSETSSEELKASNEELQAINEELRSATEELETS------- | *Lysobacter sp. URHA0019* | Bacteria; Proteobacteria; Gammaproteobacteria; Xanthomonadales; Xanthomonadaceae; Lysobacter | PAS domain S-box protein | 6 |
|  | **WP_057460932.1** | 5.8E-07 | Qa.III | ISAITKTMSDVNDITATIAAAVEQQGAATTEISENVQQAAVGTQN-------- | *Pseudovibrio sp. POLY-S9* | Bacteria; Proteobacteria; Alphaproteobacteria; Rhodobacterales; Rhodobacteraceae; Pseudovibrio | methyl-accepting chemotaxis protein | 6 |
|  | **WP_063288427.1** | 5.8E-07 | Qa.III | ISAITKTMSDVNDITATIAAAVEQQGAATTEISENVQQAAVGTQN-------- | *Pseudovibrio* | Bacteria; Proteobacteria; Alphaproteobacteria; Rhodobacterales; Rhodobacteraceae | MULTISPECIES: methyl-accepting chemotaxis protein | 6 |
|  | **WP_063304750.1** | 5.8E-07 | Qa.III | ISAITKTMSDVNDITATIAAAVEQQGAATTEISENVQQAAVGTQN-------- | *Pseudovibrio sp. W64* | Bacteria; Proteobacteria; Alphaproteobacteria; Rhodobacterales; Rhodobacteraceae; Pseudovibrio | methyl-accepting chemotaxis protein | 6 |
|  | **WP_063310829.1** | 5.8E-07 | Qa.III | ISAITKTMSDVNDITATIAAAVEQQGAATTEISENVQQAAVGTQN-------- | *Pseudovibrio sp. Ad26* | Bacteria; Proteobacteria; Alphaproteobacteria; Rhodobacterales; Rhodobacteraceae; Pseudovibrio | methyl-accepting chemotaxis protein | 6 |

| | | | | | | | | |
|---|---|---|---|---|---|---|---|---|
|  | **WP_108817492.1** | 5.8E-07 | Qa.III | ISAITKTMSDVNDITATIAAAVEQQGAATTEISENVQQAAVGTQN-------- | *Pseudovibrio sp. Alg231-02* | Bacteria; Proteobacteria; Alphaproteobacteria; Rhodobacterales; Rhodobacteraceae; Pseudovibrio | methyl-accepting chemotaxis protein | 6 |
|  | **WP_108942553.1** | 5.8E-07 | Qa.IV | IQQIGRTIVSVNDITGSIAATVVQQTAATNEIARNAAEAA------------ | *Methylobacterium sp. DM1* | Bacteria; Proteobacteria; Alphaproteobacteria; Rhizobiales; Methylobacteriaceae; Methylobacterium | HAMP domain-containing protein | 6 |
|  | **WP_027569737.1** | 5.9E-07 | Qa.IV | -KEISGTIERLSEISSTIAAAVEEQGAATQEISRNVQQAAQ----------- | *Bradyrhizobium sp. URHA0013* | Bacteria; Proteobacteria; Alphaproteobacteria; Rhizobiales; Bradyrhizobiaceae; Bradyrhizobium | methyl-accepting chemotaxis protein | 6 |
|  | **WP_080134429.1** | 5.9E-07 | Qa.III | IQSICTTIQQVDEISAAIAAAVGQQGTATQEIAQNVQQAA------------ | *Bradyrhizobium sp. BR10280* | Bacteria; Proteobacteria; Alphaproteobacteria; Rhizobiales; Bradyrhizobiaceae; Bradyrhizobium | methyl-accepting chemotaxis protein | 6 |
|  | **WP_092296123.1** | 5.9E-07 | Qa.IV | IKEISSTIERLSEISSAIAAAVEEQGAATQEIARNVQQAA------------ | *Bradyrhizobium sp. Ghvi* | Bacteria; Proteobacteria; Alphaproteobacteria; Rhizobiales; Bradyrhizobiaceae; Bradyrhizobium | methyl-accepting chemotaxis protein | 6 |
|  | **WP_026105367.1** | 6E-07 | Qa.IV | IQQIGRTIVSVNDITGSIAATVVQQTAATNEIARNAAEAA------------ | *Methylobacterium sp. MB200* | Bacteria; Proteobacteria; Alphaproteobacteria; Rhizobiales; Methylobacteriaceae; Methylobacterium | HAMP domain-containing protein | 6 |
|  | **WP_027528773.1** | 6E-07 | Qa.IV | IKEISSTIERLSEISSAIAAAVEEQGAATQEIARNVQQAA------------ | *Bradyrhizobium sp. WSM3983* | Bacteria; Proteobacteria; Alphaproteobacteria; Rhizobiales; Bradyrhizobiaceae; Bradyrhizobium | methyl-accepting chemotaxis protein | 6 |
|  | **WP_049840834.1** | 6E-07 | R.IV | IREINQAVNDLDRMVQQNAELVVESAAAAGALQTQAGDLAETAGHFR------ | *Klebsiella sp. RIT-PI-d* | Bacteria; Proteobacteria; Gammaproteobacteria; Enterobacterales; Enterobacteriaceae; Klebsiella | methyl-accepting chemotaxis protein | 6 |
|  | **WP_050626163.1** | 6E-07 | Qa.IV | IKEISSTIEKLSEISSTIAAAVEEQGAATQEISRNVQQAAQ----------- | *Bradyrhizobium viridifuturi* | Bacteria; Proteobacteria; Alphaproteobacteria; Rhizobiales; Bradyrhizobiaceae; Bradyrhizobium | methyl-accepting chemotaxis protein | 6 |
| | **WP_057863362.1** | 6E-07 | Qc.III | IKEISGTIERLAEIASTIAAAVEEQGAATQEISRNVQQAAEGTQQVSANIT-- | *Bradyrhizobium lablabi* | Bacteria; Proteobacteria; Alphaproteobacteria; Rhizobiales; Bradyrhizobiaceae; Bradyrhizobium | HAMP domain-containing protein | 6 |
|  | **WP_074447502.1** | 6E-07 | Qa.IV | IKEISSTIERLSEISSAIAAAVEEQGAATQEIARNVQQAA------------ | *Bradyrhizobium yuanmingense* | Bacteria; Proteobacteria; Alphaproteobacteria; Rhizobiales; Bradyrhizobiaceae; Bradyrhizobium | methyl-accepting chemotaxis protein | 6 |
|  | **WP_074447503.1** | 6E-07 | Qa.IV | IKEISSTIERLSEISSAIAAAVEEQGAATQEIARNVQQAA------------ | *Bradyrhizobium yuanmingense* | Bacteria; Proteobacteria; Alphaproteobacteria; Rhizobiales; Bradyrhizobiaceae; Bradyrhizobium | methyl-accepting chemotaxis protein | 6 |
|  | **WP_095733572.1** | 6E-07 | Qa.IV | IKEISSTIEKLSEISSTIAAAVEEQGAATQEISRNVQQAAQ----------- | *Bradyrhizobium sp. UFLA03-84* | Bacteria; Proteobacteria; Alphaproteobacteria; Rhizobiales; Bradyrhizobiaceae; Bradyrhizobium | methyl-accepting chemotaxis protein | 6 |
|  | **OOF17462.1** | 6.1E-07 | Qc.III | -DEISSNLANIKTLSGDMNEAIDQLGPIVSDLKDNVDSLSSAIKH-------- | *Salinivibrio sp. PR932* | Bacteria; Proteobacteria; Gammaproteobacteria; Vibrionales; Vibrionaceae; Salinivibrio | methyl-accepting chemotaxis protein partial | 6 |
| | **WP_101267984.1** | 6.1E-07 | Qc.III | IEMIGSIIENINEIATTISAAIEQQGSATNEITRNVRGAADRTRAVSSSINE- | *Thalassospira marina* | Bacteria; Proteobacteria; Alphaproteobacteria; Rhodospirillales; Rhodospirillaceae; Thalassospira | HAMP domain-containing protein | 6 |
| | **WP_101284380.1** | 6.1E-07 | Qc.III | IEMIGSIIENINEIATTISAAIEQQGSATNEITRNVRGAADRTRAVSSSINE- | *Thalassospira marina* | Bacteria; Proteobacteria; Alphaproteobacteria; Rhodospirillales; Rhodospirillaceae; Thalassospira | HAMP domain-containing protein | 6 |
|  | **WP_027537351.1** | 6.2E-07 | Qa.IV | IRSISSTISEINDVTTAIAAAVEEQGAATREIARNIQHAA------------ | *Bradyrhizobium sp. URHA0002* | Bacteria; Proteobacteria; Alphaproteobacteria; Rhizobiales; Bradyrhizobiaceae; Bradyrhizobium | HAMP domain-containing protein | 6 |
|  | **WP_074822843.1** | 6.2E-07 | Qa.IV | IRSISSTISEINDVTTAIAAAVEEQGAATREIARNIQHAA------------ | *Bradyrhizobium* | Bacteria; Proteobacteria; Alphaproteobacteria; Rhizobiales; Bradyrhizobiaceae | MULTISPECIES: HAMP domain-containing protein | 6 |
|  | **WP_079586314.1** | 6.2E-07 | Qa.IV | IRSISSTISEINDVTTAIAAAVEEQGAATREIARNIQHAA------------ | *Bradyrhizobium lablabi* | Bacteria; Proteobacteria; Alphaproteobacteria; Rhizobiales; Bradyrhizobiaceae; Bradyrhizobium | HAMP domain-containing protein | 6 |
|  | **PIW26965.1** | 6.3E-07 | Qa.III | IEGITRTINDINEIASSVAAAVEQQGVATQEIARNVEQAAAGTQEV------- | *Rhodospirillales bacterium CG15_BIG_FIL_POST_REV_8_21_14_020_66_15* | Bacteria; Proteobacteria; Alphaproteobacteria; Rhodospirillales; unclassified Rhodospirillales | hypothetical protein COW30_12255 partial | 6 |
|  | **WP_111419114.1** | 6.3E-07 | Qc.III | IKEIGGTIGRIAEIAGAIAAAVEQQGAATSEISRNIQKAASG----------- | *Rhodoplanes roseus* | Bacteria; Proteobacteria; Alphaproteobacteria; Rhizobiales; Hyphomicrobiaceae; Rhodoplanes | methyl-accepting chemotaxis protein | 6 |
|  | **WP_065935764.1** | 6.4E-07 | Qc.III | LDEMTDSVVRINDLSTQIATAAEEQSAVTEEINSNMVAVQ------------ | *Pseudomonas sp. 34 E 7* | Bacteria; Proteobacteria; Gammaproteobacteria; Pseudomonadales; Pseudomonadaceae; Pseudomonas | methyl-accepting chemotaxis protein | 6 |
|  | **WP_092024364.1** | 6.4E-07 | Qa.IV | IKEISSTIERLSEISSAIAAAVEEQGAATQEIARNVQQAAQ----------- | *Bradyrhizobium sp. OK095* | Bacteria; Proteobacteria; Alphaproteobacteria; Rhizobiales; Bradyrhizobiaceae; Bradyrhizobium | methyl-accepting chemotaxis protein | 6 |
|  | **WP_109866615.1** | 6.4E-07 | Qa.IV | IKEISSTIERLSEISSAIAAAVEEQGAATQEIARNVQQAAQ----------- | *Bradyrhizobium ottawaense* | Bacteria; Proteobacteria; Alphaproteobacteria; Rhizobiales; Bradyrhizobiaceae; Bradyrhizobium | methyl-accepting chemotaxis protein | 6 |
|  | **WP_017981124.1** | 6.5E-07 | Qa.IV | ---------EVNNAFLKIKEAIDKANETIVRIQESAKTSKENVETALRHIQ-- | *Methanocaldococcus villosus* | Archaea; Euryarchaeota; Methanococci; Methanococcales; Methanocaldococcaceae; Methanocaldococcus | HAMP domain-containing protein | 6 |
|  | **RCK47440.1** | 6.6E-07 | Qa.III | IQTIVASVNDVNGILKAIADAVLEQGTATGQISNSISEAAGYA---------- | *Thalassospira xiamenensis* | Bacteria; Proteobacteria; Alphaproteobacteria; Rhodospirillales; Rhodospirillaceae; Thalassospira | chemotaxis protein | 6 |
|  | **WP_071239531.1** | 6.6E-07 | Qa.III | IQTIVASVNDVNGILKAIADAVLEQGTATGQISNSISEAAGYA---------- | *Thalassospira sp. MIT1004* | Bacteria; Proteobacteria; Alphaproteobacteria; Rhodospirillales; Rhodospirillaceae; Thalassospira | methyl-accepting chemotaxis protein | 6 |
|  | **WP_085645625.1** | 6.6E-07 | Qa.III | IQTIVASVNDVNGILKAIADAVLEQGTATGQISNSISEAAGYA---------- | *Thalassospira sp. MCCC 1A03138* | Bacteria; Proteobacteria; Alphaproteobacteria; Rhodospirillales; Rhodospirillaceae; Thalassospira | methyl-accepting chemotaxis protein | 6 |
|  | **WP_114111453.1** | 6.6E-07 | Qa.III | IQTIVASVNDVNGILKAIADAVLEQGTATGQISNSISEAAGYA---------- | *Thalassospira xiamenensis* | Bacteria; Proteobacteria; Alphaproteobacteria; Rhodospirillales; Rhodospirillaceae; Thalassospira | HAMP domain-containing protein | 6 |
|  | **WP_114123283.1** | 6.6E-07 | Qa.III | IQTIVASVNDVNGILKAIADAVLEQGTATGQISNSISEAAGYA---------- | *Thalassospira xianhensis* | Bacteria; Proteobacteria; Alphaproteobacteria; Rhodospirillales; Rhodospirillaceae; Thalassospira | HAMP domain-containing protein | 6 |
|  | **PLX38548.1** | 6.7E-07 | Qa.III | IDAIKEIIAQISDISTSIASAIEEQSTVTREISENMQTAS------------ | *Rhizobiales bacterium* | Bacteria; Proteobacteria; Alphaproteobacteria; Rhizobiales | chemotaxis protein | 6 |
|  | **PRP55328.1** | 6.7E-07 | Qa.III | LDKIQDSAGEVTKATDDIARAVSEQTEATKSVTGYMEKTLDEVE--------- | *Bacillus halotolerans* | Bacteria; Firmicutes; Bacilli; Bacillales; Bacillaceae; Bacillus | methyl-accepting chemotaxis protein partial | 6 |
|  | **PXX35829.1** | 6.7E-07 | Qa.III | IQTIVASVNDVNAILKAIADAVLEQGTATGQISNSISEAAGYA---------- | *Thalassospira sp. 11-3* | Bacteria; Proteobacteria; Alphaproteobacteria; Rhodospirillales; Rhodospirillaceae; Thalassospira | methyl-accepting chemotaxis protein | 6 |
|  | **WP_007090953.1** | 6.7E-07 | Qa.III | IQTIVASVNDVNAILKAIADAVLEQGTATGQISNSISEAAGYA---------- | *Thalassospira xiamenensis* | Bacteria; Proteobacteria; Alphaproteobacteria; Rhodospirillales; Rhodospirillaceae; Thalassospira | methyl-accepting chemotaxis protein | 6 |
|  | **WP_037990873.1** | 6.7E-07 | Qa.III | IQTIVASVNDVNAILKAIADAVLEQGTATGQISNSISEAAGYA---------- | *Thalassospira permensis* | Bacteria; Proteobacteria; Alphaproteobacteria; Rhodospirillales; Rhodospirillaceae; Thalassospira | methyl-accepting chemotaxis protein | 6 |
|  | **WP_111358624.1** | 6.7E-07 | Qa.IV | IKEIGRTINRVSEIAGNIAAAVEQQGAATQEIARNVQQAA------------ | *Rhodoplanes elegans* | Bacteria; Proteobacteria; Alphaproteobacteria; Rhizobiales; Hyphomicrobiaceae; Rhodoplanes | methyl-accepting chemotaxis protein | 6 |
|  | **WP_114095063.1** | 6.7E-07 | Qa.III | IQTIVASVNDVNAILKAIADAVLEQGTATGQISNSISEAAGYA---------- | *Thalassospira xiamenensis* | Bacteria; Proteobacteria; Alphaproteobacteria; Rhodospirillales; Rhodospirillaceae; Thalassospira | HAMP domain-containing protein | 6 |
|  | **WP_036346129.1** | 6.8E-07 | Qa.IV | IQEIARTITEMSQISTSIAAAMEEQGAATAEIARNVQEAA------------ | *Microvirga sp. BSC39* | Bacteria; Proteobacteria; Alphaproteobacteria; Rhizobiales; Methylobacteriaceae; Microvirga | chemotaxis protein | 6 |
|  | **WP_036347106.1** | 6.8E-07 | Qa.IV | IQEIARTITEMSQISTSIAAAMEEQGAATAEIARNVQEAA------------ | *Microvirga sp. BSC39* | Bacteria; Proteobacteria; Alphaproteobacteria; Rhizobiales; Methylobacteriaceae; Microvirga | chemotaxis protein | 6 |
| | **WP_107991826.1** | 6.8E-07 | Qc.III | IEAIMSIIAQISDISTAISSGVEEQSAVTSEISRNMQSASEGVETLTSNMR-- | *Breoghania corrubedonensis* | Bacteria; Proteobacteria; Alphaproteobacteria; Rhizobiales; Cohaesibacteraceae; Breoghania | PAS domain-containing protein | 6 |
|  | **WP_011188517.1** | 6.9E-07 | Qa.III | IEQITRVIEDVNEIVATIAATVEEQSVTTREISANIVQASA----------- | *Desulfotalea psychrophila* | Bacteria; Proteobacteria; Deltaproteobacteria; Desulfobacterales; Desulfobulbaceae; Desulfotalea | HAMP domain-containing protein | 6 |
|  | **WP_047764450.1** | 6.9E-07 | Qa.III | IRSITGTISQVNDIAQTIGDSIEQQDLATREISQNVQETAINTTEASNE---- | *Kiloniella spongiae* | Bacteria; Proteobacteria; Alphaproteobacteria; Kiloniellales; Kiloniellaceae; Kiloniella | methyl-accepting chemotaxis protein | 6 |
|  | **WP_085581343.1** | 6.9E-07 | Qc.III.c | -SEIGKIIEHINEIATTISAAVEEQSAATNEISRNVRGAAER---------- | *Thalassospira mesophila* | Bacteria; Proteobacteria; Alphaproteobacteria; Rhodospirillales; Rhodospirillaceae; Thalassospira | HAMP domain-containing protein | 6 |
| | **WP_085590574.1** | 7E-07 | Qc.III | IEMIGSIIENINEIATTISAAIEQQGSATNEITRNVRGAADRTRAVSSSIT-- | *Thalassospira sp. MCCC 1A01428* | Bacteria; Proteobacteria; Alphaproteobacteria; Rhodospirillales; Rhodospirillaceae; Thalassospira | HAMP domain-containing protein | 6 |
|  | **WP_109317447.1** | 7E-07 | Qa.III | ISAITKTMSDVNDITATIAAAVEQQGAATTEISENVQQAAVGTQNVAES---- | *Pseudovibrio sp. AU243* | Bacteria; Proteobacteria; Alphaproteobacteria; Rhodobacterales; Rhodobacteraceae; Pseudovibrio | methyl-accepting chemotaxis protein | 6 |
|  | **KFG70522.1** | 7.1E-07 | Qa.IV | IQEIARTITEMSQISTSIAAAMEEQGAATAEIARNVQEAA------------ | *Microvirga sp. BSC39* | Bacteria; Proteobacteria; Alphaproteobacteria; Rhizobiales; Methylobacteriaceae; Microvirga | chemotaxis protein partial | 6 |
|  | **KFG70853.1** | 7.1E-07 | Qa.IV | IQEIARTITEMSQISTSIAAAMEEQGAATAEIARNVQEAA------------ | *Microvirga sp. BSC39* | Bacteria; Proteobacteria; Alphaproteobacteria; Rhizobiales; Methylobacteriaceae; Microvirga | chemotaxis protein partial | 6 |
|  | **OGR22404.1** | 7.1E-07 | Qa.III | IDAISKVITNINEIVASIASAVEEQSVVTKEISGNIAQAS------------ | *Desulfobacterales bacterium RIFOXYA12_FULL_46_15* | Bacteria; Proteobacteria; Deltaproteobacteria; Desulfobacterales | hypothetical protein A2277_10945 | 6 |
|  | **PID44315.1** | 7.1E-07 | Qa.III | MEQINAVINDVNTIVTTIAAAVEEQATSTDEIASNVAQAAQ----------- | *Proteobacteria bacterium* | Bacteria; Proteobacteria | hypothetical protein CR981_01480 | 6 |
| | **WP_028152152.1** | 7.1E-07 | Qc.III | IKDISDTIERLSEISSAIAAAVEEQGAATQEISRNVQQAAEGTHQVSSNITD- | *Bradyrhizobium japonicum* | Bacteria; Proteobacteria; Alphaproteobacteria; Rhizobiales; Bradyrhizobiaceae; Bradyrhizobium | methyl-accepting chemotaxis protein | 6 |
| | **WP_028158174.1** | 7.1E-07 | Qc.III | IKDISDTIERLSEISSAIAAAVEEQGAATQEISRNVQQAAEGTHQVSSNITD- | *Bradyrhizobium japonicum* | Bacteria; Proteobacteria; Alphaproteobacteria; Rhizobiales; Bradyrhizobiaceae; Bradyrhizobium | methyl-accepting chemotaxis protein | 6 |
| | **WP_038944557.1** | 7.1E-07 | Qc.III | IKDISDTIERLSEISSAIAAAVEEQGAATQEISRNVQQAAEGTHQVSSNITD- | *Bradyrhizobium japonicum* | Bacteria; Proteobacteria; Alphaproteobacteria; Rhizobiales; Bradyrhizobiaceae; Bradyrhizobium | methyl-accepting chemotaxis protein | 6 |
| | **WP_063985094.1** | 7.1E-07 | Qc.III | IKDISDTIERLSEISSAIAAAVEEQGAATQEISRNVQQAAEGTHQVSSNITD- | *Bradyrhizobium sp.* | Bacteria; Proteobacteria; Alphaproteobacteria; Rhizobiales; Bradyrhizobiaceae; Bradyrhizobium | methyl-accepting chemotaxis protein | 6 |
| | **WP_071916122.1** | 7.1E-07 | Qc.III | IKDISDTIERLSEISSAIAAAVEEQGAATQEISRNVQQAAEGTHQVSSNITD- | *Bradyrhizobium japonicum* | Bacteria; Proteobacteria; Alphaproteobacteria; Rhizobiales; Bradyrhizobiaceae; Bradyrhizobium | methyl-accepting chemotaxis protein | 6 |
| | **WP_085402566.1** | 7.1E-07 | Qc.III | IKDISDTIERLSEISSAIAAAVEEQGAATQEISRNVQQAAEGTHQVSSNITD- | *Bradyrhizobium japonicum* | Bacteria; Proteobacteria; Alphaproteobacteria; Rhizobiales; Bradyrhizobiaceae; Bradyrhizobium | methyl-accepting chemotaxis protein | 6 |
|  | **WP_108512846.1** | 7.1E-07 | Qc.III | IQEIGGTIGRIAEISQAIAAAVEEQGAATQEISRNVQQAA------------ | *Bradyrhizobium algeriense* | Bacteria; Proteobacteria; Alphaproteobacteria; Rhizobiales; Bradyrhizobiaceae; Bradyrhizobium | HAMP domain-containing protein | 6 |
| | **PTW55704.1** | 7.2E-07 | Qc.III | IEAIMSIIAQISDISTAISSGVEEQSAVTSEISRNMQSASEGVETLTSNMR-- | *Breoghania corrubedonensis* | Bacteria; Proteobacteria; Alphaproteobacteria; Rhizobiales; Cohaesibacteraceae; Breoghania | methyl-accepting chemotaxis sensory transducer with Pas/Pac sensor | 6 |
|  | **WP_092516016.1** | 7.2E-07 | Qa.IV | -KEISGTIERLSEISSTIAAAVEEQGAATQEISRNVQQAAQ----------- | *Afipia sp. GAS231* | Bacteria; Proteobacteria; Alphaproteobacteria; Rhizobiales; Bradyrhizobiaceae; Afipia | HAMP domain-containing protein | 6 |
|  | **WP_008965075.1** | 7.3E-07 | Qa.IV | IKEISSTIEKLSDISSTIAAAVEEQGAATQEISRNVQQAA------------ | *Bradyrhizobium sp. STM 3809* | Bacteria; Proteobacteria; Alphaproteobacteria; Rhizobiales; Bradyrhizobiaceae; Bradyrhizobium | methyl-accepting chemotaxis protein | 6 |
| | **WP_015666782.1** | 7.3E-07 | Qa.IV | IKEISTTIERLAEISSTIAAAVEQQGAATQEISRNIQEASRGTNEVNTNI--- | *Bradyrhizobium oligotrophicum* | Bacteria; Proteobacteria; Alphaproteobacteria; Rhizobiales; Bradyrhizobiaceae; Bradyrhizobium | PAS domain S-box protein | 6 |

| | | | | | | | |
|---|---|---|---|---|---|---|---|
| ANV98879.1 | 7.4E-07 | Qc.III | IQEIGGTIGRIAEISQAIAAAVEEQGAATQEISRNVQQAA------------- | *Bradyrhizobium icense* | Bacteria; Proteobacteria; Alphaproteobacteria; Rhizobiales; Bradyrhizobiaceae; Bradyrhizobium | chemotaxis protein | 6 |
| WP_067217888.1 | 7.4E-07 | Qa.IV | IRSITETIREMDIVTTAIASAVEQQGAATQEIARNVQSAA------------- | *Stappia indica* | Bacteria; Proteobacteria; Alphaproteobacteria; Rhodobacterales; Rhodobacteraceae; Stappia | methyl-accepting chemotaxis protein | 6 |
| WP_083219393.1 | 7.4E-07 | Qc.III | IQEIGGTIGRIAEISQAIAAAVEEQGAATQEISRNVQQAA------------- | *Bradyrhizobium icense* | Bacteria; Proteobacteria; Alphaproteobacteria; Rhizobiales; Bradyrhizobiaceae; Bradyrhizobium | HAMP domain-containing protein | 6 |
| PTL41910.1 | 7.5E-07 | Qa.III | IEEIRHVILKLNEVAQEIAAAVEEQSATTQEIANNVASANDSVQQV------- | *Blastopirellula marina* | Bacteria; Planctomycetes; Planctomycetia; Planctomycetales; Planctomycetaceae; Blastopirellula | HAMP domain-containing protein | 6 |
| WP_011085715.1 | 7.5E-07 | Qa.IV | IKEISSTIERLSEISSAIAAAVEEQGAATQEIARNVQQAAQ------------ | *Bradyrhizobium diazoefficiens* | Bacteria; Proteobacteria; Alphaproteobacteria; Rhizobiales; Bradyrhizobiaceae; Bradyrhizobium | methyl-accepting chemotaxis protein | 6 |
| WP_027544030.1 | 7.5E-07 | Qa.IV | IKEISSTIERLSEISSAIAAAVEEQGAATQEIARNVQQAAQ------------ | *Bradyrhizobium sp. WSM2254* | Bacteria; Proteobacteria; Alphaproteobacteria; Rhizobiales; Bradyrhizobiaceae; Bradyrhizobium | HAMP domain-containing protein | 6 |
| WP_080804399.1 | 7.5E-07 | Qc.III | -KAISQIINNINDISSTIAAAVEEQSASTSEITQNVQQAADGLNDVNENVA-- | *Desulfamplus magnetovallimortis* | Bacteria; Proteobacteria; Deltaproteobacteria; Desulfobacterales; Desulfobacteraceae; Desulfamplus | methyl-accepting chemotaxis protein | 6 |
| WP_094408167.1 | 7.5E-07 | Qa.IV | -ADITRTMREVNEVAAAIAAAVEQQGAATQEIARTVQEV----AQ-------- | *Elstera cyanobacteriorum* | Bacteria; Proteobacteria; Alphaproteobacteria; Rhodospirillales; Rhodospirillaceae; Elstera | HAMP domain-containing protein | 6 |
| AWL95360.1 | 7.6E-07 | Qa.IV | IRDISVTIERLSEVSSTIAAAVEEQGAATQEISRNVQQAA------------- | *Bradyrhizobium ottawaense* | Bacteria; Proteobacteria; Alphaproteobacteria; Rhizobiales; Bradyrhizobiaceae; Bradyrhizobium | methyl-accepting chemotaxis protein | 6 |
| EXJ48743.1 | 7.6E-07 | Qa.III | LDEINAAVALISDMATQIATAAEEQTHVTNEITQNVTSIK------------- | *Vibrio parahaemolyticus VPTS-2010* | Bacteria; Proteobacteria; Gammaproteobacteria; Vibrionales; Vibrionaceae; Vibrio | methyl-accepting chemotaxis (MCP) signaling domain protein | 6 |
| WP_009402188.1 | 7.6E-07 | Qb.I | LHRTKEQLQATVEQSETSTEELKASNEELQAINEELRSTTEELETSK------ | *Pseudomonas putida* | Bacteria; Proteobacteria; Gammaproteobacteria; Pseudomonadales; Pseudomonadaceae; Pseudomonas | PAS domain S-box protein | 6 |
| WP_028141088.1 | 7.6E-07 | Qa.IV | IRDISVTIERLSEVSSTIAAAVEEQGAATQEISRNVQQAA------------- | *Bradyrhizobium* | Bacteria; Proteobacteria; Alphaproteobacteria; Rhizobiales; Bradyrhizobiaceae | MULTISPECIES: methyl-accepting chemotaxis protein | 6 |
| WP_038950517.1 | 7.6E-07 | Qa.IV | IRDISVTIERLSEVSSTIAAAVEEQGAATQEISRNVQQAA------------- | *Bradyrhizobium sp. CCBAU 15544* | Bacteria; Proteobacteria; Alphaproteobacteria; Rhizobiales; Bradyrhizobiaceae; Bradyrhizobium | methyl-accepting chemotaxis protein | 6 |
| WP_038972367.1 | 7.6E-07 | Qa.IV | IRDISVTIERLSEVSSTIAAAVEEQGAATQEISRNVQQAA------------- | *Bradyrhizobium sp. CCBAU 15635* | Bacteria; Proteobacteria; Alphaproteobacteria; Rhizobiales; Bradyrhizobiaceae; Bradyrhizobium | methyl-accepting chemotaxis protein | 6 |
| WP_064304507.1 | 7.6E-07 | Qb.I | LHRTKEQLQATVEQSETSTEELKASNEELQAINEELRSTTEELETSK------ | *Pseudomonas putida* | Bacteria; Proteobacteria; Gammaproteobacteria; Pseudomonadales; Pseudomonadaceae; Pseudomonas | PAS domain S-box protein | 6 |
| WP_091957645.1 | 7.6E-07 | Qa.IV | IRDISVTIERLSEVSSTIAAAVEEQGAATQEISRNVQQAA------------- | *Bradyrhizobium shewense* | Bacteria; Proteobacteria; Alphaproteobacteria; Rhizobiales; Bradyrhizobiaceae; Bradyrhizobium | methyl-accepting chemotaxis protein | 6 |
| WP_106953190.1 | 7.6E-07 | Qa.IV | IRDISVTIERLSEVSSTIAAAVEEQGAATQEISRNVQQAA------------- | *Bradyrhizobium sp. MOS003* | Bacteria; Proteobacteria; Alphaproteobacteria; Rhizobiales; Bradyrhizobiaceae; Bradyrhizobium | methyl-accepting chemotaxis protein | 6 |
| WP_056156407.1 | 7.7E-07 | Qb.I | LHRTRAQLQDTIEHSETSNEELRASNEELQAMNEELRSATEELETS------- | *Duganella sp. Leaf126* | Bacteria; Proteobacteria; Betaproteobacteria; Burkholderiales; Oxalobacteraceae; Duganella | PAS domain S-box protein | 6 |
| WP_057863366.1 | 7.7E-07 | Qc.III | IKEISGTIERLAEISSAIAAAVEEQGAATQEISRNVQQAARGTQQVSSNITD- | *Bradyrhizobium lablabi* | Bacteria; Proteobacteria; Alphaproteobacteria; Rhizobiales; Bradyrhizobiaceae; Bradyrhizobium | methyl-accepting chemotaxis protein | 6 |
| ETP61373.1 | 7.8E-07 | Qa.III | ----VAAVERVNAIMADISTAVDEQSTGIEQVNVTVTQMEETMQ--------- | *Burkholderia dolosa PC543* | Bacteria; Proteobacteria; Betaproteobacteria; Burkholderiales; Burkholderiaceae; Burkholderia; Burkholderia cepacia complex | methyl-accepting chemotaxis protein | 6 |
| WP_036043199.1 | 7.8E-07 | Qa.IV | IKEISSTIERLSEISSAIAAAVEEQGAATQEIARNVQQAA------------- | *Bradyrhizobium yuanmingense* | Bacteria; Proteobacteria; Alphaproteobacteria; Rhizobiales; Bradyrhizobiaceae; Bradyrhizobium | methyl-accepting chemotaxis protein partial | 6 |
| WP_088874121.1 | 7.8E-07 | Qc.III | --AISGTIENINNIATAIASAVEEQGAATQEIARNVQQAATGTDSVTQNIQ-- | *Nitrospirillum amazonense* | Bacteria; Proteobacteria; Alphaproteobacteria; Rhodospirillales; Rhodospirillaceae; Nitrospirillum | methyl-accepting chemotaxis protein | 6 |
| WP_114088356.1 | 7.8E-07 | Qc.III | IQSISSTITEINEIAASIASAVEQQGAATEEITRNVREASSGTREVTSNIH-- | *Thalassospira profundimaris* | Bacteria; Proteobacteria; Alphaproteobacteria; Rhodospirillales; Rhodospirillaceae; Thalassospira | HAMP domain-containing protein | 6 |
| WP_027288172.1 | 8E-07 | Qa.IV | LQKIITRISEMHEITTTVASAVEEQESATGEIARNVEQAA------------- | *Rhodovibrio salinarum* | Bacteria; Proteobacteria; Alphaproteobacteria; Rhodospirillales; Rhodospirillaceae; Rhodovibrio | methyl-accepting chemotaxis protein | 6 |
| WP_006968205.1 | 8.1E-07 | Qa.III | ISQITGVINQVDEIVSTIAAAVEEQSVTTREIAGNVSQAARNM---------- | *Desulfotignum phosphitoxidans* | Bacteria; Proteobacteria; Deltaproteobacteria; Desulfobacterales; Desulfobacteraceae; Desulfotignum | methyl-accepting chemotaxis protein | 6 |
| WP_024335261.1 | 8.1E-07 | Qa.III | ISQITGVINQVDEIVSTIAAAVEEQSVTTREIAGNVSQAARNM---------- | *Desulfotignum balticum* | Bacteria; Proteobacteria; Deltaproteobacteria; Desulfobacterales; Desulfobacteraceae; Desulfotignum | methyl-accepting chemotaxis protein | 6 |
| WP_006767146.1 | 8.2E-07 | Qa.III | ----VAAVERVNAIMADISTAVDEQSTGIEQVNVTVTQMEETMQ--------- | *Burkholderia* | Bacteria; Proteobacteria; Betaproteobacteria; Burkholderiales; Burkholderiaceae | MULTISPECIES: HAMP domain-containing protein | 6 |
| CUU49089.1 | 8.3E-07 | Qb.I | ---INKLVSNTVKQSHDSAEYINESSYLISGTVDTFDNIFNKINSANDLIQE- | *Clostridium beijerinckii* | Bacteria; Firmicutes; Clostridia; Clostridiales; Clostridiaceae; Clostridium | Chemotaxis protein | 6 |
| PCI41461.1 | 8.4E-07 | Qa.IV | IQSIAGTISQIDEASSSIASAVEQQGAATQEIARNVEQAAN------------ | *Rhodospirillaceae bacterium* | Bacteria; Proteobacteria; Alphaproteobacteria; Rhodospirillales; Rhodospirillaceae | chemotaxis protein | 6 |
| WP_079568939.1 | 8.4E-07 | Qa.IV | IKEIGSTIAQISEISTAIAAAVEEQGAATQEIARNVQQAA------------- | *Bradyrhizobium erythrophlei* | Bacteria; Proteobacteria; Alphaproteobacteria; Rhizobiales; Bradyrhizobiaceae; Bradyrhizobium | HAMP domain-containing protein | 6 |
| WP_086508455.1 | 8.4E-07 | R.IV | -AQLNQAMDDMTRVTQDNQQLVILSTQSAHALREKAEHMQAHAGRFRITA--- | *Halomonas desiderata* | Bacteria; Proteobacteria; Gammaproteobacteria; Oceanospirillales; Halomonadaceae; Halomonas | HAMP domain-containing protein | 6 |
| EKD37939.1 | 8.5E-07 | Qa.III | IEKISKIIEEINEMINTMATAVEEQSAATAEIANNITQASQ------------ | *uncultured bacterium* | Bacteria; environmental samples | chemotaxis transducer partial | 6 |
| WP_093805199.1 | 8.5E-07 | Qc.III | IDSIMAVIKNVGDIAVSISSAVEEQTAVTNDISSNMQFAARGV---------- | *Stappia sp. ES.058* | Bacteria; Proteobacteria; Alphaproteobacteria; Rhodobacterales; Rhodobacteraceae; Stappia | PAS domain S-box protein | 6 |
| KJS35686.1 | 8.6E-07 | Qc.III | IQGIGSTIGQINEIAAAIAAAVEEQGAATREIARNVQEASTGTTEVTSNIHN- | *Rhodospirillaceae bacterium BRH_c57* | Bacteria; Proteobacteria; Alphaproteobacteria; Rhodospirillales; Rhodospirillaceae | chemotaxis protein | 6 |
| WP_026792753.1 | 8.6E-07 | Qa.IV | IRAIAATMEEVNRATLGIASAVEQQEASTNEISRNVNDASGNTV--------- | *Pleomorphomonas oryzae* | Bacteria; Proteobacteria; Alphaproteobacteria; Rhizobiales; Methylocystaceae; Pleomorphomonas | methyl-accepting chemotaxis protein | 6 |
| WP_049974371.1 | 8.6E-07 | Qa.III | ITSIASTIVRINEIASTIASAVEEQGAATREIASNVQQAA------------- | *Azospirillum sp. B4* | Bacteria; Proteobacteria; Alphaproteobacteria; Rhodospirillales; Rhodospirillaceae; Azospirillum | HAMP domain-containing protein partial | 6 |
| WP_057863365.1 | 8.6E-07 | Qc.III | IKEISGTIERLSEIASTIAAAVEEQGAATQEISRNVQQAAEGTQQVSSNITD- | *Bradyrhizobium lablabi* | Bacteria; Proteobacteria; Alphaproteobacteria; Rhizobiales; Bradyrhizobiaceae; Bradyrhizobium | chemotaxis protein | 6 |
| SDT36917.1 | 8.7E-07 | Qa.IV | ------TIAEINEISNAIAAAVEQQGAATREISGNVQQAANGTREVNENI--- | *Bradyrhizobium canariense* | Bacteria; Proteobacteria; Alphaproteobacteria; Rhizobiales; Bradyrhizobiaceae; Bradyrhizobium | methyl-accepting chemotaxis protein | 6 |
| WP_005505144.1 | 8.7E-07 | Qa.III | LEQITNMMADINDMNTQIATAATEQSSVTEEITENVTNIAR------------ | *Grimontia hollisae* | Bacteria; Proteobacteria; Gammaproteobacteria; Vibrionales; Vibrionaceae; Grimontia | methyl-accepting chemotaxis protein | 6 |
| WP_089229567.1 | 8.7E-07 | Qc.III | -RSIGEVIGQMSEIATSISAAVEEQGAATDEIAQTMSRAADGTQTVAGNI--- | *Tistlia consotensis* | Bacteria; Proteobacteria; Alphaproteobacteria; Rhodospirillales; Rhodospirillaceae; Tistlia | HAMP domain-containing protein | 6 |
| WP_089229568.1 | 8.7E-07 | Qc.III | -RSIGEVIGQMSEIATSISAAVEEQGAATDEIAQTMSRAADGTQTVAGNI--- | *Tistlia consotensis* | Bacteria; Proteobacteria; Alphaproteobacteria; Rhodospirillales; Rhodospirillaceae; Tistlia | methyl-accepting chemotaxis protein | 6 |
| WP_100385377.1 | 8.7E-07 | Qa.IV | ------TIAEINEISNAIAAAVEQQGAATREISGNVQQAANGTREVNENI--- | *Afipia broomeae* | Bacteria; Proteobacteria; Alphaproteobacteria; Rhizobiales; Bradyrhizobiaceae; Afipia | chemotaxis protein | 6 |
| PIE64159.1 | 8.8E-07 | Qa.III | MEQINAVINDVNTIVTTIAAAVEEQATSTDEIASNVAQAA------------- | *Desulfobacterales bacterium* | Bacteria; Proteobacteria; Deltaproteobacteria; Desulfobacterales | chemotaxis protein | 6 |
| WP_056563713.1 | 8.8E-07 | Qb.I | LQRTKGQLQATIEQAETSTEELKASNEELQAINEELRSTTEELETGK------ | *Achromobacter sp. Root565* | Bacteria; Proteobacteria; Betaproteobacteria; Burkholderiales; Alcaligenaceae; Achromobacter | response regulator | 6 |
| WP_080975006.1 | 8.8E-07 | Qa.III | LEQITASINQISDMATQISSAAEEQRAVTDEVSRNIQAVK------------- | *Aeromonas hydrophila* | Bacteria; Proteobacteria; Gammaproteobacteria; Aeromonadales; Aeromonadaceae; Aeromonas | methyl-accepting chemotaxis protein | 6 |
| WP_057745847.1 | 8.9E-07 | Qa.IV | IRDISATIERLSEVSSTIAAAVEEQGAATQEISRNVQQAA------------- | *Bradyrhizobium manausense* | Bacteria; Proteobacteria; Alphaproteobacteria; Rhizobiales; Bradyrhizobiaceae; Bradyrhizobium | methyl-accepting chemotaxis protein | 6 |
| WP_051375093.1 | 9E-07 | Qa.III | IDQISKVVNEVNEIVSTIATAVEEQSVTTREIAENVVQAS------------- | *Desulfosarcina sp. BuS5* | Bacteria; Proteobacteria; Deltaproteobacteria; Desulfobacterales; Desulfobacteraceae; Desulfosarcina | HAMP domain-containing protein | 6 |
| WP_077481082.1 | 9E-07 | Qc.III | -DEISSNLANIKTLSVDMNEAIDQLGPIVSDLKDNVDSLSSAIKH-------- | *Salinivibrio sp. PR919* | Bacteria; Proteobacteria; Gammaproteobacteria; Vibrionales; Vibrionaceae; Salinivibrio | methyl-accepting chemotaxis protein | 6 |
| WP_091129691.1 | 9E-07 | Qa.IV | IQEIARTINEMSQISVSIAAAMEEQGAATAEIARNVQEAAR------------ | *Microvirga guangxiensis* | Bacteria; Proteobacteria; Alphaproteobacteria; Rhizobiales; Methylobacteriaceae; Microvirga | methyl-accepting chemotaxis protein | 6 |
| OXC90405.1 | 9.1E-07 | Qb.I | LQRTKDQLQATIEQAETSTEELKASNEELQAINEELRSTTEELETGK------ | *Achromobacter sp. KAs 3-5* | Bacteria; Proteobacteria; Betaproteobacteria; Burkholderiales; Alcaligenaceae; Achromobacter | chemotaxis protein | 6 |
| WP_014744168.1 | 9.1E-07 | Qa.III | IEGIRKIIVEINDIATSIASAVEQQSAATQEIASNVQAAARGTEQV------- | *Tistrella mobilis* | Bacteria; Proteobacteria; Alphaproteobacteria; Rhodospirillales; Rhodospirillaceae; Tistrella | methyl-accepting chemotaxis protein | 6 |
| WP_029005712.1 | 9.1E-07 | Qc.III | IEGVSQQIRQMSDVATGISAAVEEQGIATREIVRNVDQAATGTNSVTSHITD- | *Azorhizobium doebereinerae* | Bacteria; Proteobacteria; Alphaproteobacteria; Rhizobiales; Xanthobacteraceae; Azorhizobium | methyl-accepting chemotaxis protein | 6 |
| WP_056323345.1 | 9.1E-07 | Qb.I | LQRTKDQLQATIEQAETSTEELKASNEELQAINEELRSTTEELETGK------ | *Achromobacter sp. Root170* | Bacteria; Proteobacteria; Betaproteobacteria; Burkholderiales; Alcaligenaceae; Achromobacter | response regulator | 6 |
| WP_062842126.1 | 9.1E-07 | Qb.I | LQRTKDQLQATIEQAETSTEELKASNEELQAINEELRSTTEELETGK------ | *Achromobacter xylosoxidans* | Bacteria; Proteobacteria; Betaproteobacteria; Burkholderiales; Alcaligenaceae; Achromobacter | response regulator | 6 |
| WP_063954671.1 | 9.1E-07 | Qb.I | LQRTKDQLQATIEQAETSTEELKASNEELQAINEELRSTTEELETGK------ | *Achromobacter xylosoxidans* | Bacteria; Proteobacteria; Betaproteobacteria; Burkholderiales; Alcaligenaceae; Achromobacter | response regulator | 6 |
| WP_105239835.1 | 9.1E-07 | Qb.I | LQRTKDQLQATIEQAETSTEELKASNEELQAINEELRSTTEELETGK------ | *Achromobacter spanius* | Bacteria; Proteobacteria; Betaproteobacteria; Burkholderiales; Alcaligenaceae; Achromobacter | response regulator | 6 |
| WP_108005449.1 | 9.1E-07 | Qb.I | LQRTKDQLQATIEQAETSTEELKASNEELQAINEELRSTTEELETGK------ | *Achromobacter mucicolens* | Bacteria; Proteobacteria; Betaproteobacteria; Burkholderiales; Alcaligenaceae; Achromobacter | response regulator | 6 |
| WP_074767759.1 | 9.3E-07 | Qc.III | LDGIGQTIRTLNELSTAIAGAVEQQSAATSEISRNVEEASANTSRTSGNI--- | *Phaeospirillum fulvum* | Bacteria; Proteobacteria; Alphaproteobacteria; Rhodospirillales; Rhodospirillaceae; Phaeospirillum | methyl-accepting chemotaxis protein | 6 |
| WP_075803584.1 | 9.3E-07 | Qb.I | LQRTKEQLQATVEQSETSTEELKASNEELQAINEELRSTTEELETSK------ | *Pseudomonas putida* | Bacteria; Proteobacteria; Gammaproteobacteria; Pseudomonadales; Pseudomonadaceae; Pseudomonas | PAS domain S-box protein | 6 |
| WP_020594493.1 | 9.4E-07 | Qa.III.b | --QIDAVIKEIDEIAAGIANAVQEQGSATREISQTVNQSASN----------- | *Kiloniella laminariae* | Bacteria; Proteobacteria; Alphaproteobacteria; Kiloniellales; Kiloniellaceae; Kiloniella | HAMP domain-containing protein | 6 |
| WP_108519852.1 | 9.6E-07 | Qa.IV | IQAISATIEKLSEISSTIAAAVEEQGAATQEISRNVQQAA------------- | *Bradyrhizobium algeriense* | Bacteria; Proteobacteria; Alphaproteobacteria; Rhizobiales; Bradyrhizobiaceae; Bradyrhizobium | HAMP domain-containing protein | 6 |

| | | | | | | | | |
|---|---|---|---|---|---|---|---|---|
| | **WP_111067744.1** | 9.6E-07 | Qa.III | IRGIADTVQQVNQIATSIASAVEEQTAATGEISRNVQQAAQGTAEVSRSI--- | *Azospirillum sp. M2T2B2* | Bacteria; Proteobacteria; Alphaproteobacteria; Rhodospirillales; Rhodospirillaceae; Azospirillum | methyl-accepting chemotaxis protein | 6 |
|  | **PWT85508.1** | 9.7E-07 | Qa.IV | IKEISDTIEKLSEISSTIAAAVEEQGAATQEISRNVQQAAE------------ | *Proteobacteria bacterium* | Bacteria; Proteobacteria | methyl-accepting chemotaxis protein | 6 |
|  | **WP_097658672.1** | 9.8E-07 | Qa.IV | IRDISVTIERLSEVSSTIAAAVEEQGAATQEISRNVQQAA------------- | *Bradyrhizobium sp. Y36* | Bacteria; Proteobacteria; Alphaproteobacteria; Rhizobiales; Bradyrhizobiaceae; Bradyrhizobium | methyl-accepting chemotaxis protein | 6 |
|  | **WP_027278069.1** | 9.9E-07 | Qa.IV | IREIGSTIERMSEISSSIASAVEEQGAATQEIARNVQEAA------------- | *Rhodopseudomonas palustris* | Bacteria; Proteobacteria; Alphaproteobacteria; Rhizobiales; Bradyrhizobiaceae; Rhodopseudomonas | methyl-accepting chemotaxis protein | 6 |
| | **WP_056490969.1** | 9.9E-07 | Qc.III | IERIAGIIGRIEESAISIGSAVEEQSATATEIRRSVEEASGRTETVSSSV--- | *Methylobacterium sp. Leaf111* | Bacteria; Proteobacteria; Alphaproteobacteria; Rhizobiales; Methylobacteriaceae; Methylobacterium | chemotaxis protein | 6 |
|  | **WP_110782365.1** | 9.9E-07 | Qa.IV | IREIGSTIERMSEISSSIASAVEEQGAATQEIARNVQEAA------------- | *Rhodopseudomonas faecalis* | Bacteria; Proteobacteria; Alphaproteobacteria; Rhizobiales; Bradyrhizobiaceae; Rhodopseudomonas | methyl-accepting chemotaxis protein | 6 |
|  | **KYG21476.1** | 1E-06 | Qa.IV | IKDISDTIEKLSEISSTIAAAVEEQGAATQEISRNVQQAAE------------ | *Bradyrhizobium sp. AT1* | Bacteria; Proteobacteria; Alphaproteobacteria; Rhizobiales; Bradyrhizobiaceae; Bradyrhizobium | chemotaxis protein | 6 |
|  | **PIW30604.1** | 1E-06 | Qa.III | IDGITRIISEVSNIATGIASAVEQQGAATREIASNVEEAAR------------ | *Rhodospirillales bacterium CG15_BIG_FIL_POST_REV_8_21_14_020_66_15* | Bacteria; Proteobacteria; Alphaproteobacteria; Rhodospirillales; unclassified Rhodospirillales | methyl-accepting chemotaxis protein | 6 |
|  | **WP_014102596.1** | 1E-06 | Qa.IV | ------TIRNIDEISTTVAAAVEEQSAATQQISRNVQEVSTATNEVNKNI--- | *Micavibrio aeruginosavorus* | Bacteria; Proteobacteria; Alphaproteobacteria; Micavibrio | hypothetical protein | 6 |
|  | **WP_027301062.1** | 1E-06 | Qa.IV | IRNISTTILEVNEISSSIAAAVEEQTAATQEITRNVQQAAN------------ | *Rhodospirillales bacterium URHD0088* | Bacteria; Proteobacteria; Alphaproteobacteria; Rhodospirillales; unclassified Rhodospirillales | HAMP domain-containing protein | 6 |
| | **WP_028350032.1** | 1E-06 | Qc.III | IKEISGTIERLAEISSAIAAAVEEQGAATQEISRNVQQAARGTHQVSANIT-- | *Bradyrhizobium elkanii* | Bacteria; Proteobacteria; Alphaproteobacteria; Rhizobiales; Bradyrhizobiaceae; Bradyrhizobium | methyl-accepting chemotaxis protein | 6 |
| | **WP_044830221.1** | 1E-06 | Qa.III | IDGITGIIANLNEIASEVAAAVEEQGQATTEISTSVQQAVQSTQEVSNHI--- | *Thalassospira sp. HJ* | Bacteria; Proteobacteria; Alphaproteobacteria; Rhodospirillales; Rhodospirillaceae; Thalassospira | methyl-accepting chemotaxis protein | 6 |
| | **WP_057847163.1** | 1E-06 | Qc.III | IKEISGTIERLAEISSAIAAAVEEQGAATQEISRNVQQAARGTHQVSTNIT-- | *Bradyrhizobium retamae* | Bacteria; Proteobacteria; Alphaproteobacteria; Rhizobiales; Bradyrhizobiaceae; Bradyrhizobium | chemotaxis protein | 6 |
|  | **WP_063196529.1** | 1E-06 | Qa.IV | IKDISDTIEKLSEISSTIAAAVEEQGAATQEISRNVQQAAE------------ | *Bradyrhizobium sp. AT1* | Bacteria; Proteobacteria; Alphaproteobacteria; Rhizobiales; Bradyrhizobiaceae; Bradyrhizobium | methyl-accepting chemotaxis protein | 6 |
|  | **WP_063291837.1** | 1E-06 | Qa.III | ISAITKTMSDVNDITATIAAAVEQQGAATTEISENVQQAAV------------ | *Pseudovibrio sp. Ad5* | Bacteria; Proteobacteria; Alphaproteobacteria; Rhodobacterales; Rhodobacteraceae; Pseudovibrio | methyl-accepting chemotaxis protein | 6 |
|  | **WP_063307234.1** | 1E-06 | Qa.III | ISAITKTMSDVNDITATIAAAVEQQGAATTEISENVQQAAV------------ | *Pseudovibrio sp. Ad46* | Bacteria; Proteobacteria; Alphaproteobacteria; Rhodobacterales; Rhodobacteraceae; Pseudovibrio | methyl-accepting chemotaxis protein | 6 |
|  | **WP_080801925.1** | 1E-06 | Qa.III | --EISKVIQEVNEIISTIATAVEEQSLTTREIVENIAQASTGVQEVNE----- | *Desulfamplus magnetovallimortis* | Bacteria; Proteobacteria; Deltaproteobacteria; Desulfobacterales; Desulfobacteraceae; Desulfamplus | methyl-accepting chemotaxis protein | 6 |
|  | **WP_103011284.1** | 1E-06 | Qa.IV | IKEISQTISRMSEIASTIASAVEEQGAATQEISRNVQEAA------------- | *Rhodopseudomonas palustris* | Bacteria; Proteobacteria; Alphaproteobacteria; Rhizobiales; Bradyrhizobiaceae; Rhodopseudomonas | methyl-accepting chemotaxis protein | 6 |
| | **WP_109451292.1** | 1E-06 | Qa.III | IRNIGRTITQVNEISNSIAAAVEEQTAATGEIGRNVQQAAEGTQEVSSSI--- | *Azospirillum sp. TSH64* | Bacteria; Proteobacteria; Alphaproteobacteria; Rhodospirillales; Rhodospirillaceae; Azospirillum | methyl-accepting chemotaxis protein | 6 |
|  | **WP_110353751.1** | 1E-06 | Qa.IV | -ISITDRIREISGLTTSIAAAVEEQGAATQEIVRNVAQAAQ------------ | *Methylobacterium sp. B4* | Bacteria; Proteobacteria; Alphaproteobacteria; Rhizobiales; Methylobacteriaceae; Methylobacterium | HAMP domain-containing protein | 6 |
|  | **ACK81664.1** | 1.1E-06 | Qa.IV | IRAIVSTITDLSEISMSVSSAVTEQAAVTQEIASNMQTAARSV---------- | *Methylorubrum extorquens CM4* | Bacteria; Proteobacteria; Alphaproteobacteria; Rhizobiales; Methylobacteriaceae; Methylorubrum | methyl-accepting chemotaxis sensory transducer with Pas/Pac sensor | 6 |
|  | **AND93918.1** | 1.1E-06 | Qa.IV | IKEISSTIERLSEISSAIAAAVEEQGAATQEIARNVQQAAQ------------ | *Bradyrhizobium diazoefficiens USDA 110* | Bacteria; Proteobacteria; Alphaproteobacteria; Rhizobiales; Bradyrhizobiaceae; Bradyrhizobium | chemotaxis protein | 6 |
|  | **BAR60108.1** | 1.1E-06 | Qa.IV | IKEISSTIERLSEISSAIAAAVEEQGAATQEIARNVQQAAQ------------ | *Bradyrhizobium diazoefficiens* | Bacteria; Proteobacteria; Alphaproteobacteria; Rhizobiales; Bradyrhizobiaceae; Bradyrhizobium | hypothetical protein NK6_6957 | 6 |
|  | **GAK84059.1** | 1.1E-06 | Qa.III | LEKITQSIQEISEMASHISNAAFEQRTVTEEVSKNIQQAN------------- | *Vibrio ponticus* | Bacteria; Proteobacteria; Gammaproteobacteria; Vibrionales; Vibrionaceae; Vibrio | methyl-accepting chemotaxis protein | 6 |
|  | **KGJ68477.1** | 1.1E-06 | Qa.IV | IKEISSTIERLSEISSAIAAAVEEQGAATQEIARNVQQAAQ------------ | *Bradyrhizobium diazoefficiens SEMIA 5080* | Bacteria; Proteobacteria; Alphaproteobacteria; Rhizobiales; Bradyrhizobiaceae; Bradyrhizobium | putative chemotaxis protein | 6 |
|  | **KOY10463.1** | 1.1E-06 | Qa.IV | IKEISSTIERLSEISSAIAAAVEEQGAATQEIARNVQQAAQ------------ | *Bradyrhizobium japonicum* | Bacteria; Proteobacteria; Alphaproteobacteria; Rhizobiales; Bradyrhizobiaceae; Bradyrhizobium | chemotaxis protein | 6 |
|  | **OEU46385.1** | 1.1E-06 | Qa.III | IEEISHVINNVNEMVSTIAAAVEQQSVTTKKIADNVAQAS------------- | *Desulfobacterales bacterium S3730MH5* | Bacteria; Proteobacteria; Deltaproteobacteria; Desulfobacterales | hypothetical protein BA861_11525 | 6 |
|  | **OJX78591.1** | 1.1E-06 | Qa.IV | INEILATIGEISEASTTIASAVEQQQAATNEIARNVEQAS------------- | *Magnetospirillum sp. 64-120* | Bacteria; Proteobacteria; Alphaproteobacteria; Rhodospirillales; Rhodospirillaceae; Magnetospirillum | hypothetical protein BGO92_01740 | 6 |
| | **OZA91769.1** | 1.1E-06 | Qc.III | IERVGTQIRKMSDVANEISAAVEEQGMATKEIVRNVDQAATGTNSVTSHISD- | *Rhizobiales bacterium 39-66-18* | Bacteria; Proteobacteria; Alphaproteobacteria; Rhizobiales | hypothetical protein B7X67_29870 partial | 6 |
| | **PIW26631.1** | 1.1E-06 | Qc.III | --EINRTIGRIEETSTAISAAVEEQDAATREIAVNVEQAAAGTAEVTRNVA-- | *Rhodospirillales bacterium CG15_BIG_FIL_POST_REV_8_21_14_020_66_15* | Bacteria; Proteobacteria; Alphaproteobacteria; Rhodospirillales; unclassified Rhodospirillales | chemotaxis protein | 6 |
|  | **SBW09516.1** | 1.1E-06 | Qa.IV | IQAIATTIGRISEISSAIASAVEEQGAATQEIARNVQQAA------------- | *uncultured Alphaproteobacteria bacterium* | Bacteria; Proteobacteria; Alphaproteobacteria; environmental samples | Putative Methyl-accepting chemotaxis protein (fragment) | 6 |
|  | **SCB55649.1** | 1.1E-06 | Qa.IV | -RDISDTIEKLSEISSTIAAAVEEQGAATQEISRNVQQAAE------------ | *Bradyrhizobium shewense* | Bacteria; Proteobacteria; Alphaproteobacteria; Rhizobiales; Bradyrhizobiaceae; Bradyrhizobium | Methyl-accepting chemotaxis protein | 6 |
|  | **SFI39098.1** | 1.1E-06 | Qa.IV | IKEISSTIERLSEISSAIAAAVEEQGAATQEIARNVQQAAQ------------ | *Bradyrhizobium sp. cf659* | Bacteria; Proteobacteria; Alphaproteobacteria; Rhizobiales; Bradyrhizobiaceae; Bradyrhizobium | methyl-accepting chemotaxis sensory transducer with Cache sensor | 6 |
|  | **WP_014285304.1** | 1.1E-06 | Qa.III | ISAITRTMANVNDITATIAAAVEQQGAATTEISENVQQAAVGTQN-------- | *Pseudovibrio sp. FO-BEG1* | Bacteria; Proteobacteria; Alphaproteobacteria; Rhodobacterales; Rhodobacteraceae; Pseudovibrio | methyl-accepting chemotaxis protein | 6 |
|  | **WP_057862430.1** | 1.1E-06 | Qa.IV | IRNISSTIAEINEVTTAIAAAVEEQGAATREIARNIQHAA------------- | *Bradyrhizobium lablabi* | Bacteria; Proteobacteria; Alphaproteobacteria; Rhizobiales; Bradyrhizobiaceae; Bradyrhizobium | HAMP domain-containing protein | 6 |
| | **WP_082733749.1** | 1.1E-06 | Qc.III | IESIASIIERISGIAQAISGAVEEQGAATSAITRNIDEAAKGTEAVSASI--- | *Pseudovibrio hongkongensis* | Bacteria; Proteobacteria; Alphaproteobacteria; Rhodobacterales; Rhodobacteraceae; Pseudovibrio | hypothetical protein | 6 |
|  | **WP_084301890.1** | 1.1E-06 | Qa.IV | IKDISDTIEKLSEISSTIAAAVEEQGAATQEISRNVQQAAE------------ | *Bradyrhizobium sp. WSM2254* | Bacteria; Proteobacteria; Alphaproteobacteria; Rhizobiales; Bradyrhizobiaceae; Bradyrhizobium | methyl-accepting chemotaxis protein | 6 |
|  | **WP_091967314.1** | 1.1E-06 | Qa.IV | -RDISDTIEKLSEISSTIAAAVEEQGAATQEISRNVQQAAE------------ | *Bradyrhizobium shewense* | Bacteria; Proteobacteria; Alphaproteobacteria; Rhizobiales; Bradyrhizobiaceae; Bradyrhizobium | HAMP domain-containing protein | 6 |
| | **WP_101246891.1** | 1.1E-06 | Qc.III | IEKVTQTISSISEIANGISSAVNEQNAAVNEIANSATSASAGSTEVVANIDE- | *Thalassospira lohafexi* | Bacteria; Proteobacteria; Alphaproteobacteria; Rhodospirillales; Rhodospirillaceae; Thalassospira | methyl-accepting chemotaxis protein | 6 |
|  | **WP_104462860.1** | 1.1E-06 | Qa.IV | -RDISDTIEKLSEISSTIAAAVEEQGAATQEISRNVQQAAE------------ | *Bradyrhizobium sp. AC87j1* | Bacteria; Proteobacteria; Alphaproteobacteria; Rhizobiales; Bradyrhizobiaceae; Bradyrhizobium | methyl-accepting chemotaxis protein | 6 |
|  | **WP_111418193.1** | 1.1E-06 | Qa.IV | IKEIGRTINRVSEIASNIAASVEQQGAATQEIARNVQQAA------------- | *Rhodoplanes roseus* | Bacteria; Proteobacteria; Alphaproteobacteria; Rhizobiales; Hyphomicrobiaceae; Rhodoplanes | methyl-accepting chemotaxis protein | 6 |
| | **WP_114112266.1** | 1.1E-06 | Qc.III | IEKVTQTISSISEIANGISSAVNEQNAAVNEIANSATSASAGSTEVVANIDE- | *Thalassospira profundimaris* | Bacteria; Proteobacteria; Alphaproteobacteria; Rhodospirillales; Rhodospirillaceae; Thalassospira | HAMP domain-containing protein | 6 |
| | **AFK54332.1** | 1.2E-06 | Qc.III | IEEIGATINRVNEISTAISAAMQEQGAATREISRNVQEASQGTVEVTGNI--- | *Tistrella mobilis KA081020-065* | Bacteria; Proteobacteria; Alphaproteobacteria; Rhodospirillales; Rhodospirillaceae; Tistrella | Methyl-accepting chemotaxis protein | 6 |
|  | **AWO89940.1** | 1.2E-06 | Qa.IV | IKEISSTIERLSEISSAIAAAVEEQGAATQEIARNVQQAAQ------------ | *Bradyrhizobium diazoefficiens* | Bacteria; Proteobacteria; Alphaproteobacteria; Rhizobiales; Bradyrhizobiaceae; Bradyrhizobium | methyl-accepting chemotaxis protein | 6 |
| | **EKD26980.1** | 1.2E-06 | Qa.III | --KISKIIEEVNNITHTIASAVEEQSATTSEIASNVENTSTAARESSKNI--- | *uncultured bacterium* | Bacteria; environmental samples | hypothetical protein ACD_79C00945G0001 partial | 6 |
| | **KKJ76948.1** | 1.2E-06 | Qc.III | LEAIGNSIREVNEIGEAISTAVEEQGAATQEIARNVEQASAGTREVSSNIQ-- | *Kiloniella litopenaei* | Bacteria; Proteobacteria; Alphaproteobacteria; Kiloniellales; Kiloniellaceae; Kiloniella | hypothetical protein WH95_10960 | 6 |
|  | **NP_769572.1** | 1.2E-06 | Qa.IV | IKEISSTIERLSEISSAIAAAVEEQGAATQEIARNVQQAAQ------------ | *Bradyrhizobium diazoefficiens USDA 110* | Bacteria; Proteobacteria; Alphaproteobacteria; Rhizobiales; Bradyrhizobiaceae; Bradyrhizobium | hypothetical protein blr2932 | 6 |
| | **PVE22810.1** | 1.2E-06 | Qa.IV | IQEIARTINEMSQISISIAAAMEQQGAATAEIARNVQEAARGTEQVTGNI--- | *Microvirga sp. KLBC 81* | Bacteria; Proteobacteria; Alphaproteobacteria; Rhizobiales; Methylobacteriaceae; Microvirga | methyl-accepting chemotaxis protein | 6 |
| | **WP_011925755.1** | 1.2E-06 | Qc.III | IKEIGSTIGRLSEISAAVSAAVEEQGAATQEISRNVQHAANGTQRVSHNIDD- | *Bradyrhizobium sp. ORS 278* | Bacteria; Proteobacteria; Alphaproteobacteria; Rhizobiales; Bradyrhizobiaceae; Bradyrhizobium | methyl-accepting chemotaxis protein | 6 |
|  | **WP_022665797.1** | 1.2E-06 | Qa.III | IEEITRAITDVNEMIDSVAAVVEEQSVTTKEIAENVSQAAR------------ | *Desulfospira joergensenii* | Bacteria; Proteobacteria; Deltaproteobacteria; Desulfobacterales; Desulfobacteraceae; Desulfospira | methyl-accepting chemotaxis protein | 6 |
|  | **WP_022853134.1** | 1.2E-06 | Qc.III.c | -ERISKVIERVNEVSNTIAAAVEEQTATVSGISENMNDVAS------------ | *Thermodesulfatator atlanticus* | Bacteria; Thermodesulfobacteria; Thermodesulfobacteriales; Thermodesulfobacteriaceae; Thermodesulfatator | methyl-accepting chemotaxis protein | 6 |
| | **WP_041604956.1** | 1.2E-06 | Qc.III | IEEIGATINRVNEISTAISAAMQEQGAATREISRNVQEASQGTVEVTGNI--- | *Tistrella mobilis* | Bacteria; Proteobacteria; Alphaproteobacteria; Rhodospirillales; Rhodospirillaceae; Tistrella | methyl-accepting chemotaxis protein | 6 |
|  | **WP_047138359.1** | 1.2E-06 | Qb.I | LQRTRDRLQATIEQSETSTEELKASNEELQAINEELRSATEELETS------- | *Luteimonas sp. FCS-9* | Bacteria; Proteobacteria; Gammaproteobacteria; Xanthomonadales; Xanthomonadaceae; Luteimonas | response regulator | 6 |
|  | **WP_051155374.1** | 1.2E-06 | Qb.I | LQRTKEQLQETVEHSEVSTEELKASNEELQAINEELRSATEELETS------- | *Noviherbaspirillum massiliense* | Bacteria; Proteobacteria; Betaproteobacteria; Burkholderiales; Oxalobacteraceae; Noviherbaspirillum | PAS domain S-box protein | 6 |
| | **WP_051357016.1** | 1.2E-06 | Qc.III | IEGVGGQIRRMSDVATSISAAVEQQGVATREIVRNVDQAATGTNTVTSHISD- | *Azorhizobium doebereinerae* | Bacteria; Proteobacteria; Alphaproteobacteria; Rhizobiales; Xanthobacteraceae; Azorhizobium | HAMP domain-containing protein | 6 |
|  | **WP_057195871.1** | 1.2E-06 | Qa.IV | IREISVTIERLSEISSAVAAAVEQQGAATQEISRNVQQAAD------------ | *Bradyrhizobium sp. Leaf396* | Bacteria; Proteobacteria; Alphaproteobacteria; Rhizobiales; Bradyrhizobiaceae; Bradyrhizobium | methyl-accepting chemotaxis protein | 6 |
| | **WP_072825830.1** | 1.2E-06 | Qc.III | -REISGTIEKLSEIASAIAAAVEEQGAATQEISRNVQRAAQGTMQVSSNIAD- | *Bradyrhizobium erythrophlei* | Bacteria; Proteobacteria; Alphaproteobacteria; Rhizobiales; Bradyrhizobiaceae; Bradyrhizobium | HAMP domain-containing protein | 6 |
|  | **WP_074077141.1** | 1.2E-06 | Qa.IV | IKEISSTIERLSEISSAIAAAVEEQGAATQEIARNVQQAAQ------------ | *Bradyrhizobium* | Bacteria; Proteobacteria; Alphaproteobacteria; Rhizobiales; Bradyrhizobiaceae | MULTISPECIES: methyl-accepting chemotaxis protein | 6 |
|  | **WP_074272832.1** | 1.2E-06 | Qa.III | IQAIGGTIAEIDEISNEIAAAVDQQGAATREIAGNVQQAANGTRDVNNN---- | *Bradyrhizobium erythrophlei* | Bacteria; Proteobacteria; Alphaproteobacteria; Rhizobiales; Bradyrhizobiaceae; Bradyrhizobium | chemotaxis protein | 6 |

| | | | | | | | | |
|---|---|---|---|---|---|---|---|---|
|  | WP_080135670.1 | 1.2E-06 | Qa.IV | IKDISDTIEKLSEISSTIAAAVEEQGAATQEISRNVQQAA------------- | *Bradyrhizobium sp. BR10280* | Bacteria; Proteobacteria; Alphaproteobacteria; Rhizobiales; Bradyrhizobiaceae; Bradyrhizobium | HAMP domain-containing protein | 6 |
| | WP_080138430.1 | 1.2E-06 | Qc.III | -RSIGATIQRVNEIAAAIAAAVEEQGAATREIARNVQQASQGTNEVSRHIS-- | *Bradyrhizobium sp. BR10280* | Bacteria; Proteobacteria; Alphaproteobacteria; Rhizobiales; Bradyrhizobiaceae; Bradyrhizobium | HAMP domain-containing protein | 6 |
|  | WP_082756239.1 | 1.2E-06 | Qa.IV | IKEISSTIERLSEISSAIAAAVEEQGAATQEIARNVQQAAQ------------ | *Bradyrhizobium diazoefficiens* | Bacteria; Proteobacteria; Alphaproteobacteria; Rhizobiales; Bradyrhizobiaceae; Bradyrhizobium | methyl-accepting chemotaxis protein | 6 |
|  | WP_083860045.1 | 1.2E-06 | Qa.IV | IKEISSTIERLSEISSAIAAAVEEQGAATQEIARNVQQAAQ------------ | *Bradyrhizobium diazoefficiens* | Bacteria; Proteobacteria; Alphaproteobacteria; Rhizobiales; Bradyrhizobiaceae; Bradyrhizobium | methyl-accepting chemotaxis protein | 6 |
|  | WP_092186099.1 | 1.2E-06 | Qa.IV | IKEISSTIERLSEISSAIAAAVEEQGAATQEIARNVQQAAQ------------ | *Bradyrhizobium sp. cf659* | Bacteria; Proteobacteria; Alphaproteobacteria; Rhizobiales; Bradyrhizobiaceae; Bradyrhizobium | methyl-accepting chemotaxis protein | 6 |
|  | WP_093269943.1 | 1.2E-06 | Qa.III | LDEINRAVAVISDMATQIATAAEEQSHVTDEITQNVTSIK------------- | *Vibrio xiamenensis* | Bacteria; Proteobacteria; Gammaproteobacteria; Vibrionales; Vibrionaceae; Vibrio | methyl-accepting chemotaxis protein | 6 |
|  | WP_094407443.1 | 1.2E-06 | Qa.IV | IRQIVLTIQQLDEISSVVATAVEQQGAATQEIARNVSEAATSTGEALE----- | *Elstera cyanobacteriorum* | Bacteria; Proteobacteria; Alphaproteobacteria; Rhodospirillales; Rhodospirillaceae; Elstera | methyl-accepting chemotaxis protein | 6 |
| | WP_106750644.1 | 1.2E-06 | SNAP.c | IDKVGQVIGEMRGIAASVAAAVEEQNAAVSTIAEAVNRASDETRDGAEAM--- | *Phreatobacter cathodiphilus* | Bacteria; Proteobacteria; Alphaproteobacteria; Phreatobacter | HAMP domain-containing protein | 6 |
| | WP_108519816.1 | 1.2E-06 | Qc.III | IKDISGTIERLAEISSAIAAAVEEQGAATQEISRNVQQAARGTQQVSSNISD- | *Bradyrhizobium algeriense* | Bacteria; Proteobacteria; Alphaproteobacteria; Rhizobiales; Bradyrhizobiaceae; Bradyrhizobium | methyl-accepting chemotaxis protein | 6 |
|  | ANK81589.1 | 1.3E-06 | Qa.IV | IQKVTGAVKELNGIASTIASAVEEQTAVTQDIAKNVQQAA------------- | *Rhizobiales bacterium NRL2* | Bacteria; Proteobacteria; Alphaproteobacteria; Rhizobiales | hypothetical protein TEF_12890 | 6 |
|  | AWL97789.1 | 1.3E-06 | Qa.IV | -RDISDTIEKLSEISSTIAAAVEEQGAATQEISRNVQQAAE------------ | *Bradyrhizobium ottawaense* | Bacteria; Proteobacteria; Alphaproteobacteria; Rhizobiales; Bradyrhizobiaceae; Bradyrhizobium | methyl-accepting chemotaxis protein | 6 |
|  | CCD92075.1 | 1.3E-06 | Qa.IV | IRAISTTIERLSEISSAIAAAVEEQGAATQEISRNVQQAS------------- | *Bradyrhizobium sp. ORS 375* | Bacteria; Proteobacteria; Alphaproteobacteria; Rhizobiales; Bradyrhizobiaceae; Bradyrhizobium | putative methyl-accepting chemotaxis receptor/sensory transducer precursor | 6 |
|  | RCK26994.1 | 1.3E-06 | Qa.III | IDGITSIISNLNEIATEVAAAVQEQGQATTEISNSVQQAVQSTQE-------- | *Thalassospira lucentensis MCCC 1A00383 = DSM 14000* | Bacteria; Proteobacteria; Alphaproteobacteria; Rhodospirillales; Rhodospirillaceae; Thalassospira | chemotaxis protein | 6 |
|  | SFH87482.1 | 1.3E-06 | Qa.III | IQSICVTIREVDEISAAIAAAVGQQGTATQEIAQNVQQAA------------- | *Bradyrhizobium sp. Gha* | Bacteria; Proteobacteria; Alphaproteobacteria; Rhizobiales; Bradyrhizobiaceae; Bradyrhizobium | methyl-accepting chemotaxis sensory transducer | 6 |
|  | SFN86938.1 | 1.3E-06 | Qa.IV | -RDISDTIEKLSEISSTIAAAVEEQGAATQEISRNVQQAAE------------ | *Bradyrhizobium sp. Rc3b* | Bacteria; Proteobacteria; Alphaproteobacteria; Rhizobiales; Bradyrhizobiaceae; Bradyrhizobium | methyl-accepting chemotaxis sensory transducer | 6 |
| | SHN66634.1 | 1.3E-06 | Qc.III | -REISGTIEKLSEIASAIAAAVEEQGAATQEISRNVQRAAQGTMQVSSNIAD- | *Bradyrhizobium erythrophlei* | Bacteria; Proteobacteria; Alphaproteobacteria; Rhizobiales; Bradyrhizobiaceae; Bradyrhizobium | methyl-accepting chemotaxis sensory transducer with Cache sensor | 6 |
| | SMH62626.1 | 1.3E-06 | Qa.III | IRGIAGTIADINEIAATIAAAVEQQGAATAEIARNVQQAARGTEEVARSIT-- | *Azospirillum lipoferum* | Bacteria; Proteobacteria; Alphaproteobacteria; Rhodospirillales; Rhodospirillaceae; Azospirillum | Methyl-accepting chemotaxis protein (MCP) signalling domain-containing protein partial | 6 |
|  | WP_006610690.1 | 1.3E-06 | Qa.IV | IRAISTTIERLSEISSAIAAAVEEQGAATQEISRNVQQAS------------- | *Bradyrhizobium sp. ORS 285* | Bacteria; Proteobacteria; Alphaproteobacteria; Rhizobiales; Bradyrhizobiaceae; Bradyrhizobium | HAMP domain-containing protein | 6 |
|  | WP_008945532.1 | 1.3E-06 | Qa.IV | IREIGQTIAEINDIAANIAAAVEEQGAATQEISRNVQEAA------------- | *Oceanibaculum indicum* | Bacteria; Proteobacteria; Alphaproteobacteria; Rhodospirillales; Rhodospirillaceae; Oceanibaculum | methyl-accepting chemotaxis protein | 6 |
|  | WP_011925590.1 | 1.3E-06 | Qc.III | IKEISGTIERLSEIASTIAAAVEEQGAATSEISRNVQHASQGTKDVS------ | *Bradyrhizobium sp. ORS 278* | Bacteria; Proteobacteria; Alphaproteobacteria; Rhizobiales; Bradyrhizobiaceae; Bradyrhizobium | HAMP domain-containing protein | 6 |
|  | WP_028141643.1 | 1.3E-06 | Qa.IV | -RDISDTIEKLSEISSTIAAAVEEQGAATQEISRNVQQAAE------------ | *Bradyrhizobium* | Bacteria; Proteobacteria; Alphaproteobacteria; Rhizobiales; Bradyrhizobiaceae | MULTISPECIES: methyl-accepting chemotaxis protein | 6 |
|  | WP_036347291.1 | 1.3E-06 | Qa.IV | IQEIARTITEMSQISTSIAAAMEEQGAATAEIARNVQEAA------------- | *Microvirga sp. BSC39* | Bacteria; Proteobacteria; Alphaproteobacteria; Rhizobiales; Methylobacteriaceae; Microvirga | methyl-accepting chemotaxis protein partial | 6 |
|  | WP_036347649.1 | 1.3E-06 | Qa.IV | IQEIARTITEMSQISTSIAAAMEEQGAATAEIARNVQEAA------------- | *Microvirga sp. BSC39* | Bacteria; Proteobacteria; Alphaproteobacteria; Rhizobiales; Methylobacteriaceae; Microvirga | methyl-accepting chemotaxis protein partial | 6 |
|  | WP_038948271.1 | 1.3E-06 | Qa.IV | -RDISDTIEKLSEISSTIAAAVEEQGAATQEISRNVQQAAE------------ | *Bradyrhizobium* | Bacteria; Proteobacteria; Alphaproteobacteria; Rhizobiales; Bradyrhizobiaceae | MULTISPECIES: HAMP domain-containing protein | 6 |
|  | WP_049823230.1 | 1.3E-06 | Qa.IV | IKAISETIERLSEISSTIAAAVEEQGAATQEIARNVQQAA------------- | *Bradyrhizobium sp. WSM2254* | Bacteria; Proteobacteria; Alphaproteobacteria; Rhizobiales; Bradyrhizobiaceae; Bradyrhizobium | methyl-accepting chemotaxis protein | 6 |
| | WP_051244921.1 | 1.3E-06 | Qa.III.b | IRMIAESTREVDEITTTIADSIDQQGQATREISESAQQAATSVERTVSTI--- | *Thalassobaculum salexigens* | Bacteria; Proteobacteria; Alphaproteobacteria; Rhodospirillales; Rhodospirillaceae; Thalassobaculum | HAMP domain-containing protein | 6 |
|  | WP_065730698.1 | 1.3E-06 | Qa.IV | IKEISSTIERLAEISSAIAAAVEEQGAATQEISRNVQQAA------------- | *Bradyrhizobium icense* | Bacteria; Proteobacteria; Alphaproteobacteria; Rhizobiales; Bradyrhizobiaceae; Bradyrhizobium | methyl-accepting chemotaxis protein | 6 |
|  | WP_076574895.1 | 1.3E-06 | Qa.III | LEQIAESIGQISDMATQISSAAEEQRAVTDEVSRNIQAVK------------- | *Aeromonas sp. RU39B* | Bacteria; Proteobacteria; Gammaproteobacteria; Aeromonadales; Aeromonadaceae; Aeromonas | methyl-accepting chemotaxis protein | 6 |
|  | WP_081928705.1 | 1.3E-06 | Qa.IV | IQEIARTITEMSQISTSIAAAMEEQGAATAEIARNVQEAA------------- | *Microvirga sp. BSC39* | Bacteria; Proteobacteria; Alphaproteobacteria; Rhizobiales; Methylobacteriaceae; Microvirga | methyl-accepting chemotaxis protein partial | 6 |
|  | WP_090272411.1 | 1.3E-06 | Qa.III | -TNIRRAIGQVSDMTGQIASAAEEQGAVVEEINHNITNIAA------------ | *Pseudomonas litoralis* | Bacteria; Proteobacteria; Gammaproteobacteria; Pseudomonadales; Pseudomonadaceae; Pseudomonas | PAS domain S-box protein | 6 |
|  | WP_092262613.1 | 1.3E-06 | Qa.IV | -RDISDTIEKLSEISSTIAAAVEEQGAATQEISRNVQQAAE------------ | *Bradyrhizobium sp. Rc3b* | Bacteria; Proteobacteria; Alphaproteobacteria; Rhizobiales; Bradyrhizobiaceae; Bradyrhizobium | methyl-accepting chemotaxis protein | 6 |
|  | WP_092512206.1 | 1.3E-06 | Qa.III | IQAIGGTIAEIDEISNEIAAAVDQQGAATREIAGNVQQAADGTREVN------ | *Afipia sp. GAS231* | Bacteria; Proteobacteria; Alphaproteobacteria; Rhizobiales; Bradyrhizobiaceae; Afipia | chemotaxis protein | 6 |
| | WP_093154164.1 | 1.3E-06 | Qa.III.b | IRMIAESTREVDEITTTIADSIDQQGQATREISESAQQAATSVERTVSTI--- | *Thalassobaculum litoreum* | Bacteria; Proteobacteria; Alphaproteobacteria; Rhodospirillales; Rhodospirillaceae; Thalassobaculum | HAMP domain-containing protein | 6 |
|  | WP_101301013.1 | 1.3E-06 | Qa.III | IDGITSIISNLNEIATEVAAAVQEQGQATTEISNSVQQAVQSTQE-------- | *Thalassospira lohafexi* | Bacteria; Proteobacteria; Alphaproteobacteria; Rhodospirillales; Rhodospirillaceae; Thalassospira | methyl-accepting chemotaxis protein | 6 |
|  | WP_102784531.1 | 1.3E-06 | Qa.III | IDGITSIISNLNEIATEVAAAVQEQGQATTEISNSVQQAVQSTQE-------- | *Thalassospira sp. GB04J01* | Bacteria; Proteobacteria; Alphaproteobacteria; Rhodospirillales; Rhodospirillaceae; Thalassospira | methyl-accepting chemotaxis protein | 6 |
| | WP_106950445.1 | 1.3E-06 | Qc.III | -REISGSIGRLSEIASAIAAAIEEQGAATQEIARNVQNAAQGTQQVSANIG-- | *Bradyrhizobium sp. MOS003* | Bacteria; Proteobacteria; Alphaproteobacteria; Rhizobiales; Bradyrhizobiaceae; Bradyrhizobium | HAMP domain-containing protein | 6 |
|  | WP_109794642.1 | 1.3E-06 | Qa.IV | IQKVTGAVKELNGIASTIASAVEEQTAVTQDIAKNVQQAA------------- | *Rhizobiales bacterium* | Bacteria; Proteobacteria; Alphaproteobacteria; Rhizobiales | HAMP domain-containing protein | 6 |
|  | AEH44199.1 | 1.4E-06 | Qc.III | IEEITEIISRINDISNMIASAVEEQTAMMGEISQAANMSAESVDQIKEK---- | *Thermodesulfatator indicus DSM 15286* | Bacteria; Thermodesulfobacteria; Thermodesulfobacteriales; Thermodesulfobacteriaceae; Thermodesulfatator | methyl-accepting chemotaxis sensory transducer | 6 |
| | KLN59676.1 | 1.4E-06 | Qc.III | LEAIGNSIREVNEIGEAISTAVEEQAAATQEIARNVEQASAGTREVSSNIQ-- | *Kiloniella spongiae* | Bacteria; Proteobacteria; Alphaproteobacteria; Kiloniellales; Kiloniellaceae; Kiloniella | hypothetical protein WH96_16645 | 6 |
|  | OJJ12709.1 | 1.4E-06 | Qa.III | --AITSTISKINEITTSISVAVEQQGAATNDIASNIQQAAAGTQQVSD----- | *Alphaproteobacteria bacterium AO1-B* | Bacteria; Proteobacteria; Alphaproteobacteria | chemotaxis protein | 6 |
| | PLX41388.1 | 1.4E-06 | Qa.IV | VRSITDQVKDISNVTSAIAAAIEQQQSATAEISRSVQMTSDETNQAAMSV--- | *Rhizobiales bacterium* | Bacteria; Proteobacteria; Alphaproteobacteria; Rhizobiales | hypothetical protein C0605_05705 | 6 |
| | RCK52479.1 | 1.4E-06 | Qa.IV | ---IAGTIRELDAATGRIAAAVEEQGAATQEIARNVDEAARGVARVNADIDAA | *Thalassospira profundimaris* | Bacteria; Proteobacteria; Alphaproteobacteria; Rhodospirillales; Rhodospirillaceae; Thalassospira | hypothetical protein TH25_05385 | 6 |
|  | SIO65949.1 | 1.4E-06 | Qa.IV | -KEISGTIERLAEISSTIAAAVEEQGAATQEISRNVQQAAQ------------ | *Bradyrhizobium erythrophlei* | Bacteria; Proteobacteria; Alphaproteobacteria; Rhizobiales; Bradyrhizobiaceae; Bradyrhizobium | methyl-accepting chemotaxis sensory transducer | 6 |
|  | SMX59631.1 | 1.4E-06 | Qa.IV | IKEISSTIEKLSEISSTIAAAVEEQGAATQEISRNVQQAA------------- | *Bradyrhizobium sp. ORS 285* | Bacteria; Proteobacteria; Alphaproteobacteria; Rhizobiales; Bradyrhizobiaceae; Bradyrhizobium | putative methyl-accepting chemotaxis receptor/sensory transducer (Chemoreceptor) | 6 |
|  | WP_008144692.1 | 1.4E-06 | Qa.IV | IKEISATIARISDISTTIAATIEEQGAATAEIARNVNEAAK------------ | *Bradyrhizobium sp. YR681* | Bacteria; Proteobacteria; Alphaproteobacteria; Rhizobiales; Bradyrhizobiaceae; Bradyrhizobium | methyl-accepting chemotaxis protein | 6 |
|  | WP_008963891.1 | 1.4E-06 | Qa.IV | IRAISTTIERLSEISSAIAAAVEEQGAATQEISRNVQQAS------------- | *Bradyrhizobium sp. STM 3809* | Bacteria; Proteobacteria; Alphaproteobacteria; Rhizobiales; Bradyrhizobiaceae; Bradyrhizobium | HAMP domain-containing protein | 6 |
|  | WP_017221403.1 | 1.4E-06 | Qc.III.c | LDSMTESVLQINDLGIQIATAAEEQSSVTEEINRNMTSIQTMVHQLTN----- | *Moritella dasanensis* | Bacteria; Proteobacteria; Gammaproteobacteria; Alteromonadales; Moritellaceae; Moritella | methyl-accepting chemotaxis protein | 6 |
|  | WP_018648774.1 | 1.4E-06 | Qa.IV | -RDISDTIEKLSEISSTIAAAVEEQGAATQEISRNVQQAAE------------ | *Bradyrhizobium japonicum* | Bacteria; Proteobacteria; Alphaproteobacteria; Rhizobiales; Bradyrhizobiaceae; Bradyrhizobium | methyl-accepting chemotaxis protein | 6 |
|  | WP_019644243.1 | 1.4E-06 | Qc.III.c | --AIGGTISRINAISSAIASAVEEQTAVTRDLSANMQSTSSGVGTIT------ | *Novispirillum itersonii* | Bacteria; Proteobacteria; Alphaproteobacteria; Rhodospirillales; Rhodospirillaceae; Novispirillum | methyl-accepting chemotaxis protein | 6 |
|  | WP_020084505.1 | 1.4E-06 | Qa.IV | INEIASTINRISAVSSNIAAAVEQQGATTQEISRNVMEAA------------- | *Hyphomicrobium zavarzinii* | Bacteria; Proteobacteria; Alphaproteobacteria; Rhizobiales; Hyphomicrobiaceae; Hyphomicrobium | methyl-accepting chemotaxis protein | 6 |
| | WP_028350028.1 | 1.4E-06 | Qc.III | IKEISGTIERLAEIASTIAAAVEEQGAATQEISRNVQQAADGTQQVSANIT-- | *Bradyrhizobium elkanii* | Bacteria; Proteobacteria; Alphaproteobacteria; Rhizobiales; Bradyrhizobiaceae; Bradyrhizobium | HAMP domain-containing protein | 6 |
|  | WP_035639630.1 | 1.4E-06 | Qa.IV | IRAISTTIERLSEISSAIAAAVEEQGAATQEISRNVQQAS------------- | *Bradyrhizobium sp. ORS 375* | Bacteria; Proteobacteria; Alphaproteobacteria; Rhizobiales; Bradyrhizobiaceae; Bradyrhizobium | HAMP domain-containing protein | 6 |
|  | WP_035643645.1 | 1.4E-06 | Qc.III | IKEISGTIERLSEIASTIAAAVEEQGAATQEISRNVQHASQGTRDVS------ | *Bradyrhizobium sp. ORS 375* | Bacteria; Proteobacteria; Alphaproteobacteria; Rhizobiales; Bradyrhizobiaceae; Bradyrhizobium | methyl-accepting chemotaxis protein | 6 |
|  | WP_035647811.1 | 1.4E-06 | Qa.IV | IKEISSTIEKLSEISSTIAAAVEEQGAATQEISRNVQQAA------------- | *Bradyrhizobium sp. ORS 285* | Bacteria; Proteobacteria; Alphaproteobacteria; Rhizobiales; Bradyrhizobiaceae; Bradyrhizobium | HAMP domain-containing protein | 6 |
|  | WP_042048745.1 | 1.4E-06 | Qa.III | LEQIAGSIGQISDMATQISSAAEEQRAVTDEVSRNIQAVK------------- | *Aeromonas simiae* | Bacteria; Proteobacteria; Gammaproteobacteria; Aeromonadales; Aeromonadaceae; Aeromonas | methyl-accepting chemotaxis protein | 6 |

| | | | | | | | | |
|---|---|---|---|---|---|---|---|---|
|  | **WP_044407419.1** | 1.4E-06 | Qa.IV | IKEISSTIGRMSEISSTIASAVEEQGAATQEISRNVQQAAQ------------ | *Rhodopseudomonas palustris* | Bacteria; Proteobacteria; Alphaproteobacteria; Rhizobiales; Bradyrhizobiaceae; Rhodopseudomonas | methyl-accepting chemotaxis protein | 6 |
|  | **WP_045485841.1** | 1.4E-06 | Qa.III | LDEINSAVAQISDMATQIATAAEEQTHVTNEITQNVTSIK------------- | *Vibrio harveyi* | Bacteria; Proteobacteria; Gammaproteobacteria; Vibrionales; Vibrionaceae; Vibrio | methyl-accepting chemotaxis protein | 6 |
|  | **WP_047193729.1** | 1.4E-06 | Qb.I | LKHTKQQLQNTIEQSETSTEELKASNEELQAINEELRSTTEELETSK------ | *[Polyangium] brachysporum* | Bacteria; Proteobacteria; Betaproteobacteria; Burkholderiales | PAS domain S-box protein | 6 |
|  | **WP_050922370.1** | 1.4E-06 | Qa.III | LDEINSAVAQISDMATQIATAAEEQTHVTNEITQNVTSIK------------- | *Vibrio harveyi* | Bacteria; Proteobacteria; Gammaproteobacteria; Vibrionales; Vibrionaceae; Vibrio | methyl-accepting chemotaxis protein | 6 |
|  | **WP_050938611.1** | 1.4E-06 | Qa.III | LDEINSAVAQISDMATQIATAAEEQTHVTNEITQNVTSIK------------- | *Vibrio harveyi* | Bacteria; Proteobacteria; Gammaproteobacteria; Vibrionales; Vibrionaceae; Vibrio | methyl-accepting chemotaxis protein | 6 |
|  | **WP_050940873.1** | 1.4E-06 | Qa.III | LDEINSAVAQISDMATQIATAAEEQTHVTNEITQNVTSIK------------- | *Vibrio harveyi* | Bacteria; Proteobacteria; Gammaproteobacteria; Vibrionales; Vibrionaceae; Vibrio | methyl-accepting chemotaxis protein | 6 |
|  | **WP_054163810.1** | 1.4E-06 | Qa.IV | IKEISLTISQMSEIASTIASAVEEQGAATQEISRNVQQAA------------- | *Rhodopseudomonas sp. AAP120* | Bacteria; Proteobacteria; Alphaproteobacteria; Rhizobiales; Bradyrhizobiaceae; Rhodopseudomonas | HAMP domain-containing protein | 6 |
| | **WP_057847160.1** | 1.4E-06 | Qc.III | IKEISGTIERLAEIASTIAAAVEEQGAATQEISRNVQQAADGTQQVSANIT-- | *Bradyrhizobium retamae* | Bacteria; Proteobacteria; Alphaproteobacteria; Rhizobiales; Bradyrhizobiaceae; Bradyrhizobium | methyl-accepting chemotaxis protein | 6 |
|  | **WP_071854123.1** | 1.4E-06 | Qb.I | LQRTKEQLQSTIEQSETSTEELKASNEELQAINEELRSATEELETS------- | *Burkholderia ubonensis* | Bacteria; Proteobacteria; Betaproteobacteria; Burkholderiales; Burkholderiaceae; Burkholderia; Burkholderia cepacia complex | PAS domain S-box protein partial | 6 |
|  | **WP_073616796.1** | 1.4E-06 | Qb.III.d | --QSERILGDIESLPAGSRQELHQYKEQIERINDELQRRQEQL---------- | *Desulfopila aestuarii* | Bacteria; Proteobacteria; Deltaproteobacteria; Desulfobacterales; Desulfobulbaceae; Desulfopila | PAS domain S-box protein | 6 |
|  | **WP_074281433.1** | 1.4E-06 | Qa.IV | -KEISGTIERLAEISSTIAAAVEEQGAATQEISRNVQQAAQ------------ | *Bradyrhizobium erythrophlei* | Bacteria; Proteobacteria; Alphaproteobacteria; Rhizobiales; Bradyrhizobiaceae; Bradyrhizobium | methyl-accepting chemotaxis protein | 6 |
|  | **WP_085615605.1** | 1.4E-06 | Qa.III | IEGITSIISNLNGIASEVAAAVQEQGHATSEISTSVQQAVQSTQE-------- | *Thalassospira alkalitolerans* | Bacteria; Proteobacteria; Alphaproteobacteria; Rhodospirillales; Rhodospirillaceae; Thalassospira | methyl-accepting chemotaxis protein | 6 |
|  | **WP_097660542.1** | 1.4E-06 | Qa.IV | IKEISATIARISDISTTIAATIEEQGAATAEIARNVNEAAK------------ | *Bradyrhizobium sp. Y36* | Bacteria; Proteobacteria; Alphaproteobacteria; Rhizobiales; Bradyrhizobiaceae; Bradyrhizobium | methyl-accepting chemotaxis protein | 6 |
|  | **WP_099473549.1** | 1.4E-06 | Qc.III.c | -DRIKRSIDKVQASSDVIREEVTRQRAATRDIADSADRAQASVQMVT------ | *Emcibacter congregatus* | Bacteria; Proteobacteria; Alphaproteobacteria; Emcibacterales; Emcibacteraceae; Emcibacter | hypothetical protein | 6 |
|  | **WP_100175319.1** | 1.4E-06 | Qa.IV | IKEISATIARISDISTTIAATIEEQGAATAEIARNVNEAAK------------ | *Bradyrhizobium sp. TSA1* | Bacteria; Proteobacteria; Alphaproteobacteria; Rhizobiales; Bradyrhizobiaceae; Bradyrhizobium | methyl-accepting chemotaxis protein | 6 |
| | **WP_109110705.1** | 1.4E-06 | Qa.III | IRGIADTVQQVNQIATSIASAVEEQTAATGEISRNVQQAAQGTAEVSRSI--- | *Azospirillum sp. TSO35-2* | Bacteria; Proteobacteria; Alphaproteobacteria; Rhodospirillales; Rhodospirillaceae; Azospirillum | HAMP domain-containing protein | 6 |
|  | **WP_110782636.1** | 1.4E-06 | Qa.IV | IKEIGQTISRMSEIASTIASAVEEQGAATQEISRNVQQAA------------- | *Rhodopseudomonas faecalis* | Bacteria; Proteobacteria; Alphaproteobacteria; Rhizobiales; Bradyrhizobiaceae; Rhodopseudomonas | methyl-accepting chemotaxis protein | 6 |
| | **WP_114087529.1** | 1.4E-06 | Qa.IV | ---IAGTIRELDAATGRIAAAVEEQGAATQEIARNVDEAARGVARVNADIDAA | *Thalassospira profundimaris* | Bacteria; Proteobacteria; Alphaproteobacteria; Rhodospirillales; Rhodospirillaceae; Thalassospira | HAMP domain-containing protein | 6 |
|  | **WP_114320389.1** | 1.4E-06 | Qa.III | LDEINSAVAQISDMATQIATAAEEQTHVTNEITQNVTSIK------------- | *Vibrio harveyi* | Bacteria; Proteobacteria; Gammaproteobacteria; Vibrionales; Vibrionaceae; Vibrio | HAMP domain-containing protein | 6 |
|  | **ACB78882.1** | 1.5E-06 | Qa.IV | IRAIVSTITDLSEISMSVSSAVTEQAAVTQEIASNMQTAARSV---------- | *Methylorubrum populi BJ001* | Bacteria; Proteobacteria; Alphaproteobacteria; Rhizobiales; Methylobacteriaceae; Methylorubrum | methyl-accepting chemotaxis sensory transducer with Pas/Pac sensor | 6 |
|  | **CCD97186.1** | 1.5E-06 | Qc.III | IKEISGTIERLSEIASTIAAAVEEQGAATQEISRNVQHASQGTRDVS------ | *Bradyrhizobium sp. ORS 375* | Bacteria; Proteobacteria; Alphaproteobacteria; Rhizobiales; Bradyrhizobiaceae; Bradyrhizobium | putative methyl-accepting chemotaxis protein; signal peptide | 6 |
|  | **EVU13461.1** | 1.5E-06 | Qa.III | LDEINAAVALISDMATQIATAAEEQTHVTNEITQNVTSIK------------- | *Vibrio parahaemolyticus V-223/04* | Bacteria; Proteobacteria; Gammaproteobacteria; Vibrionales; Vibrionaceae; Vibrio | methyl-accepting chemotaxis (MCP) signaling domain protein | 6 |
|  | **PPD16667.1** | 1.5E-06 | Qa.III | -----ADIAELAPICAGISSAVQEQRQTIDNLRQQMETAQGAVEGVVREVG-- | *Methylobacterium sp.* | Bacteria; Proteobacteria; Alphaproteobacteria; Rhizobiales; Methylobacteriaceae; Methylobacterium | hypothetical protein CTY25_01650 | 6 |
|  | **RCK28686.1** | 1.5E-06 | Qa.III | --DIASVITRITDMTNTISDAVQQQSEATDEIAQNVEQAAIGTQEVSS----- | *Thalassospira lucentensis MCCC 1A00383 = DSM 14000* | Bacteria; Proteobacteria; Alphaproteobacteria; Rhodospirillales; Rhodospirillaceae; Thalassospira | chemotaxis protein | 6 |
|  | **WP_012829728.1** | 1.5E-06 | Qb.I | LEETKAKLQATIEQMDANTEELQATNEELTAANEELQSTNEELH--------- | *Haliangium ochraceum* | Bacteria; Proteobacteria; Deltaproteobacteria; Myxococcales; Nannocystineae; Kofleriaceae; Haliangium | PAS domain S-box protein | 6 |
|  | **WP_015668556.1** | 1.5E-06 | Qa.IV | IRAISTTIERLSEISSAIAAAVEEQGAATQEISRNVQQAS------------- | *Bradyrhizobium oligotrophicum* | Bacteria; Proteobacteria; Alphaproteobacteria; Rhizobiales; Bradyrhizobiaceae; Bradyrhizobium | HAMP domain-containing protein | 6 |
|  | **WP_026230364.1** | 1.5E-06 | Qb.I | LQRTKEQLQATIEQSETSTEELKASNEELQAINEELRSATEELETS------- | *Paraburkholderia symbiotica* | Bacteria; Proteobacteria; Betaproteobacteria; Burkholderiales; Burkholderiaceae; Paraburkholderia | response regulator | 6 |
|  | **WP_036347245.1** | 1.5E-06 | Qa.IV | IQEIARTITEMSQISTSIAAAMEEQGAATAEISRNVQEAAR------------ | *Microvirga sp. BSC39* | Bacteria; Proteobacteria; Alphaproteobacteria; Rhizobiales; Methylobacteriaceae; Microvirga | HAMP domain-containing protein | 6 |
|  | **WP_036349971.1** | 1.5E-06 | Qa.IV | IQEIARTITEMSQISTSIAAAMEEQGAATAEIARNVQEAAR------------ | *Microvirga sp. BSC39* | Bacteria; Proteobacteria; Alphaproteobacteria; Rhizobiales; Methylobacteriaceae; Microvirga | methyl-accepting chemotaxis protein | 6 |
|  | **WP_074820127.1** | 1.5E-06 | Qa.III | IQAIGGTIAEIDEISNEIAAAVDQQGAATREIAGNVQQAADGTREVN------ | *Bradyrhizobium* | Bacteria; Proteobacteria; Alphaproteobacteria; Rhizobiales; Bradyrhizobiaceae | MULTISPECIES: chemotaxis protein | 6 |
| | **WP_079438362.1** | 1.5E-06 | Qb.I | ---INKLVSNTVKQSEESAGYINESSYLISGTVDTFDSIFDRINDTNNLI--- | *Clostridium chromiireducens* | Bacteria; Firmicutes; Clostridia; Clostridiales; Clostridiaceae; Clostridium | methyl-accepting chemotaxis protein | 6 |
|  | **WP_081422088.1** | 1.5E-06 | Qa.III | IKQIGNTIGEISGIASEIASAVEQQGAATREIAQNVQRVAH------------ | *Tardiphaga robiniae* | Bacteria; Proteobacteria; Alphaproteobacteria; Rhizobiales; Bradyrhizobiaceae; Tardiphaga | methyl-accepting chemotaxis protein | 6 |
|  | **WP_081715892.1** | 1.5E-06 | Qb.I | LQRTKEQLQSTVEQSETSTEELKASNEELQAINEELRSATEELETS------- | *Paraburkholderia mimosarum* | Bacteria; Proteobacteria; Betaproteobacteria; Burkholderiales; Burkholderiaceae; Paraburkholderia | PAS domain S-box protein | 6 |
|  | **WP_081723292.1** | 1.5E-06 | Qb.I | LQRTKEQLQSTVEQSETSTEELKASNEELQAINEELRSATEELETS------- | *Paraburkholderia mimosarum* | Bacteria; Proteobacteria; Betaproteobacteria; Burkholderiales; Burkholderiaceae; Paraburkholderia | PAS domain S-box protein | 6 |
|  | **WP_091129688.1** | 1.5E-06 | Qa.IV | IQDIARTITEMSQISTSIAAAMEEQGAATAEIARNVQEAAR------------ | *Microvirga guangxiensis* | Bacteria; Proteobacteria; Alphaproteobacteria; Rhizobiales; Methylobacteriaceae; Microvirga | methyl-accepting chemotaxis protein | 6 |
|  | **WP_092190828.1** | 1.5E-06 | Qa.III | IDRIVKVIGEVNSIVANIAAAIEEQSAVTREVASNIAQATEDVQDAN------ | *Desulfomicrobium norvegicum* | Bacteria; Proteobacteria; Deltaproteobacteria; Desulfovibrionales; Desulfomicrobiaceae; Desulfomicrobium | methyl-accepting chemotaxis protein | 6 |
|  | **WP_092227466.1** | 1.5E-06 | Qa.IV | IKDISDTIEKLSEISSTIAAAVEEQGAATQEISRNVQQAAQ------------ | *Bradyrhizobium sp. Gha* | Bacteria; Proteobacteria; Alphaproteobacteria; Rhizobiales; Bradyrhizobiaceae; Bradyrhizobium | methyl-accepting chemotaxis protein | 6 |
|  | **WP_092516017.1** | 1.5E-06 | Qa.IV | IKEISSTIEKLSEISSTIAAAVEEQGAATQEISRNVQQAAQ------------ | *Afipia sp. GAS231* | Bacteria; Proteobacteria; Alphaproteobacteria; Rhizobiales; Bradyrhizobiaceae; Afipia | HAMP domain-containing protein | 6 |
|  | **WP_100177237.1** | 1.5E-06 | Qa.IV | IRDISVTIERLSEVSSTIAAAVEEQGAATQEISRNVQQAA------------- | *Bradyrhizobium sp. TSA1* | Bacteria; Proteobacteria; Alphaproteobacteria; Rhizobiales; Bradyrhizobiaceae; Bradyrhizobium | methyl-accepting chemotaxis protein | 6 |
|  | **WP_101304631.1** | 1.5E-06 | Qa.III | --DIASVITRITDMTNTISDAVQQQSEATDEIAQNVEQAAIGTQEVSS----- | *Thalassospira lohafexi* | Bacteria; Proteobacteria; Alphaproteobacteria; Rhodospirillales; Rhodospirillaceae; Thalassospira | HAMP domain-containing protein | 6 |
|  | **WP_102784959.1** | 1.5E-06 | Qa.III | --DIASVITRITDMTNTISDAVQQQSEATDEIAQNVEQAAIGTQEVSS----- | *Thalassospira sp. GB04J01* | Bacteria; Proteobacteria; Alphaproteobacteria; Rhodospirillales; Rhodospirillaceae; Thalassospira | HAMP domain-containing protein | 6 |
|  | **WP_106751858.1** | 1.5E-06 | Qa.III | IQEIASTMNEVNQFTATIADAISQQGEATREITYNVQKAA------------- | *Pannonibacter carbonis* | Bacteria; Proteobacteria; Alphaproteobacteria; Rhodobacterales; Rhodobacteraceae; Pannonibacter | methyl-accepting chemotaxis protein | 6 |
|  | **WP_107149255.1** | 1.5E-06 | Qb.I | LQRTKEQLQATIEQSETSTEELKASNEELQAINEELRSATEELETS------- | *Paraburkholderia symbiotica* | Bacteria; Proteobacteria; Betaproteobacteria; Burkholderiales; Burkholderiaceae; Paraburkholderia | response regulator | 6 |
|  | **WP_108938772.1** | 1.5E-06 | Qa.IV | IRAIVSTITELSEISMSVSSAVTEQAAVTQEIATNMQTAARSV---------- | *Methylobacterium sp. DM1* | Bacteria; Proteobacteria; Alphaproteobacteria; Rhizobiales; Methylobacteriaceae; Methylobacterium | PAS domain S-box protein | 6 |
|  | **WP_111433457.1** | 1.5E-06 | Qa.IV | IQQIEATIQEMNQITTGIASAMEQQNAATEEIARNVEQAA------------- | *Rhodobium orientis* | Bacteria; Proteobacteria; Alphaproteobacteria; Rhizobiales; Rhodobiaceae; Rhodobium | methyl-accepting chemotaxis protein | 6 |
|  | **AAZ64263.1** | 1.6E-06 | Qb.I | LQRTKEQLQATIEQSETSTEELKASNEELQAINEELRSATEELETS------- | *Cupriavidus pinatubonensis JMP134* | Bacteria; Proteobacteria; Betaproteobacteria; Burkholderiales; Burkholderiaceae; Cupriavidus | PAS | 6 |
|  | **AND93872.1** | 1.6E-06 | Qa.IV | -RDISDTIEKLSEISSTIAAAVEEQGAATQEISRNVQQAAQ------------ | *Bradyrhizobium diazoefficiens USDA 110* | Bacteria; Proteobacteria; Alphaproteobacteria; Rhizobiales; Bradyrhizobiaceae; Bradyrhizobium | chemotaxis protein | 6 |
|  | **APO55491.1** | 1.6E-06 | Qa.IV | -RDISDTIEKLSEISSTIAAAVEEQGAATQEISRNVQQAAQ------------ | *Bradyrhizobium diazoefficiens* | Bacteria; Proteobacteria; Alphaproteobacteria; Rhizobiales; Bradyrhizobiaceae; Bradyrhizobium | chemotaxis protein | 6 |
|  | **ERB67204.1** | 1.6E-06 | Qc.III.c | ------SIVRINDLGLEIATAASQQNAVTEEVSRNMTTIQSMVDELTSNG--- | *Vibrio coralliilyticus OCN008* | Bacteria; Proteobacteria; Gammaproteobacteria; Vibrionales; Vibrionaceae; Vibrio | hypothetical protein N779_00175 | 6 |
|  | **KZD20836.1** | 1.6E-06 | Qa.III | IKQIGNTIGEISGIASEIASAVEQQGAATREIAQNVQRVAH------------ | *Tardiphaga robiniae* | Bacteria; Proteobacteria; Alphaproteobacteria; Rhizobiales; Bradyrhizobiaceae; Tardiphaga | chemotaxis protein | 6 |
|  | **OHE62881.1** | 1.6E-06 | Qa.III | -------IRQVNEIVTGIAAAIEEQSVATRDIASNVAQASQGVEDANRNI--- | *Treponema sp. GWC1_61_84* | Bacteria; Spirochaetes; Spirochaetales; Spirochaetaceae; Treponema | hypothetical protein A2001_00315 | 6 |
|  | **OHE72494.1** | 1.6E-06 | Qa.III.b | -EEITKIVYEINEIANTIAASVAEQSTTTGSISQNMNGVSEAT---------- | *Verrucomicrobia bacterium GWC2_42_7* | Bacteria; Verrucomicrobia; unclassified Verrucomicrobia | hypothetical protein A2007_01640 | 6 |
|  | **WP_003606904.1** | 1.6E-06 | Qa.IV | IRAIVSTITDLSEISMSVSSAVTEQAAVTQEIASNMQTAARSV---------- | *Methylorubrum extorquens* | Bacteria; Proteobacteria; Alphaproteobacteria; Rhizobiales; Methylobacteriaceae; Methylorubrum | PAS domain S-box protein | 6 |
|  | **WP_011665943.1** | 1.6E-06 | Qa.III | IKQIGDTIGQISSIASTIASAVTEQGSATREIAQNVQSVAR------------ | *Rhodopseudomonas palustris* | Bacteria; Proteobacteria; Alphaproteobacteria; Rhizobiales; Bradyrhizobiaceae; Rhodopseudomonas | methyl-accepting chemotaxis protein | 6 |
|  | **WP_012252469.1** | 1.6E-06 | Qa.IV | IRAIVSTITDLSEISMSVSSAVTEQAAVTQEIASNMQTAARSV---------- | *Methylobacteriaceae* | Bacteria; Proteobacteria; Alphaproteobacteria; Rhizobiales | MULTISPECIES: PAS domain S-box protein | 6 |
|  | **WP_012355628.1** | 1.6E-06 | Qb.I | LQRTKEQLQATIEQSETSTEELKASNEELQAINEELRSATEELETS------- | *Cupriavidus taiwanensis* | Bacteria; Proteobacteria; Betaproteobacteria; Burkholderiales; Burkholderiaceae; Cupriavidus | PAS domain S-box protein | 6 |
|  | **WP_015821223.1** | 1.6E-06 | Qa.IV | IRAIVSTITDLSEISMSVSSAVTEQAAVTQEIASNMQTAARSV---------- | *Methylorubrum* | Bacteria; Proteobacteria; Alphaproteobacteria; Rhizobiales; Methylobacteriaceae | MULTISPECIES: PAS domain S-box protein | 6 |
|  | **WP_018007395.1** | 1.6E-06 | Qb.I | LQRTKEQLQATIEQSETSTEELKASNEELQAINEELRSATEELETS------- | *Cupriavidus taiwanensis* | Bacteria; Proteobacteria; Betaproteobacteria; Burkholderiales; Burkholderiaceae; Cupriavidus | PAS domain S-box protein | 6 |
|  | **WP_025581717.1** | 1.6E-06 | Qb.I | LQRTKEQLQATIEQSETSTEELKASNEELQAINEELRSATEELETS------- | *Cupriavidus taiwanensis* | Bacteria; Proteobacteria; Betaproteobacteria; Burkholderiales; Burkholderiaceae; Cupriavidus | PAS domain S-box protein | 6 |
|  | **WP_027558191.1** | 1.6E-06 | Qa.IV | IKDISDTIEKLSEISSTIAAAVEEQGAATQEISRNVQQAAQ------------ | *Bradyrhizobium genosp. SA-4* | Bacteria; Proteobacteria; Alphaproteobacteria; Rhizobiales; Bradyrhizobiaceae; Bradyrhizobium | methyl-accepting chemotaxis protein | 6 |

| | | | | | | | | |
|---|---|---|---|---|---|---|---|---|
|  | **WP_038958329.1** | 1.6E-06 | Qa.IV | IKDISDTIEKLSEISSTIAAAVEEQGAATQEISRNVQQAAQ------------ | *Bradyrhizobium japonicum* | Bacteria; Proteobacteria; Alphaproteobacteria; Rhizobiales; Bradyrhizobiaceae; Bradyrhizobium | methyl-accepting chemotaxis protein | 6 |
|  | **WP_041680272.1** | 1.6E-06 | Qb.I | LQRTKEQLQATIEQSETSTEELKASNEELQAINEELRSATEELETS------- | *Cupriavidus pinatubonensis* | Bacteria; Proteobacteria; Betaproteobacteria; Burkholderiales; Burkholderiaceae; Cupriavidus | PAS domain S-box protein | 6 |
|  | **WP_056112558.1** | 1.6E-06 | Qa.IV | IRAIVSTITDLSEISMSVSSAVTEQAAVTQEIASNMQTAARSV---------- | *Methylobacterium sp. Leaf122* | Bacteria; Proteobacteria; Alphaproteobacteria; Rhizobiales; Methylobacteriaceae; Methylobacterium | PAS domain S-box protein | 6 |
|  | **WP_056456980.1** | 1.6E-06 | Qa.IV | IRAIVSTITDLSEISMSVSSAVTEQAAVTQEIASNMQTAARSV---------- | *Methylobacterium sp. Leaf90* | Bacteria; Proteobacteria; Alphaproteobacteria; Rhizobiales; Methylobacteriaceae; Methylobacterium | PAS domain S-box protein | 6 |
|  | **WP_056498418.1** | 1.6E-06 | Qa.IV | IRAIVSTITDLSEISMSVSSAVTEQAAVTQEIASNMQTAARSV---------- | *Methylobacterium sp. Leaf121* | Bacteria; Proteobacteria; Alphaproteobacteria; Rhizobiales; Methylobacteriaceae; Methylobacterium | PAS domain S-box protein | 6 |
|  | **WP_062803373.1** | 1.6E-06 | Qb.I | LQRTKEQLQATIEQSETSTEELKASNEELQAINEELRSATEELETS------- | *Cupriavidus nantongensis* | Bacteria; Proteobacteria; Betaproteobacteria; Burkholderiales; Burkholderiaceae; Cupriavidus | PAS domain S-box protein | 6 |
|  | **WP_076641086.1** | 1.6E-06 | Qa.IV | IRAIVSTITDLSEISMSVSSAVTEQAAVTQEIASNMQTAARSV---------- | *Methylorubrum extorquens* | Bacteria; Proteobacteria; Alphaproteobacteria; Rhizobiales; Methylobacteriaceae; Methylorubrum | PAS domain S-box protein | 6 |
|  | **WP_082291645.1** | 1.6E-06 | Qa.IV | IRAIVSTITDLSEISMSVSSAVTEQAAVTQEIASNMQTAARSV---------- | *Methylorubrum extorquens* | Bacteria; Proteobacteria; Alphaproteobacteria; Rhizobiales; Methylobacteriaceae; Methylorubrum | PAS domain S-box protein | 6 |
| | **WP_111418820.1** | 1.6E-06 | Qc.III | IKEIGGTIGRIAEIASTIAAAVEEQGAATGEIARNVQQAATGTTEVARNIT-- | *Rhodoplanes roseus* | Bacteria; Proteobacteria; Alphaproteobacteria; Rhizobiales; Hyphomicrobiaceae; Rhodoplanes | methyl-accepting chemotaxis protein | 6 |
|  | **AEG16281.1** | 1.7E-06 | Qc.I | -----RMARQLHDLYTGLEQKVQERTAQLQEANELLERQRQELENANRELT-- | *Desulfofundulus kuznetsovii DSM 6115* | Bacteria; Firmicutes; Clostridia; Clostridiales; Peptococcaceae; Desulfofundulus | integral membrane sensor signal transduction histidine kinase | 6 |
| | **GAJ33561.1** | 1.7E-06 | Qc.III | IKDISHTIERLSEISSTIAAAVEEQGAATQEISRNVQQAATGTHQVSSNITD- | *Bradyrhizobium sp. DOA9* | Bacteria; Proteobacteria; Alphaproteobacteria; Rhizobiales; Bradyrhizobiaceae; Bradyrhizobium | chemoreceptor protein A | 6 |
|  | **KGJ68024.1** | 1.7E-06 | Qa.IV | -RDISDTIEKLSEISSTIAAAVEEQGAATQEISRNVQQAAQ------------ | *Bradyrhizobium diazoefficiens SEMIA 5080* | Bacteria; Proteobacteria; Alphaproteobacteria; Rhizobiales; Bradyrhizobiaceae; Bradyrhizobium | putative chemotaxis protein | 6 |
|  | **OEU70277.1** | 1.7E-06 | Qa.III.b | IRQISKVINDVNDIVTTIASSVEEQSVTTKEIATNVAQAS------------- | *Desulfuromonadales bacterium C00003093* | Bacteria; Proteobacteria; Deltaproteobacteria; Desulfuromonadales | hypothetical protein BA864_14845 | 6 |
|  | **SMX59623.1** | 1.7E-06 | Qc.III | IKEISGTIERLSEIASTIAAAVEEQGAATQEISRNVQHASQGTKDV------- | *Bradyrhizobium sp. ORS 285* | Bacteria; Proteobacteria; Alphaproteobacteria; Rhizobiales; Bradyrhizobiaceae; Bradyrhizobium | putative methyl-accepting chemotaxis protein; signal peptide | 6 |
|  | **WP_009493622.1** | 1.7E-06 | Qa.IV | IQEIARTITEMSQISTSIAAAMEEQGAATAEIARNVQEAA------------- | *Microvirga lotononidis* | Bacteria; Proteobacteria; Alphaproteobacteria; Rhizobiales; Methylobacteriaceae; Microvirga | HAMP domain-containing protein | 6 |
|  | **WP_009493624.1** | 1.7E-06 | Qa.IV | IQEIARTITEMSQISVSIAAAMEEQGAATAEIARNVQEAAR------------ | *Microvirga lotononidis* | Bacteria; Proteobacteria; Alphaproteobacteria; Rhizobiales; Methylobacteriaceae; Microvirga | HAMP domain-containing protein | 6 |
|  | **WP_011085333.1** | 1.7E-06 | Qa.IV | -RDISDTIEKLSEISSTIAAAVEEQGAATQEISRNVQQAAQ------------ | *Bradyrhizobium* | Bacteria; Proteobacteria; Alphaproteobacteria; Rhizobiales; Bradyrhizobiaceae | MULTISPECIES: methyl-accepting chemotaxis protein | 6 |
| | **WP_020590736.1** | 1.7E-06 | Qc.III | --QISKTIGRINEFANAIAAAIEEQSAATQEIASSVQKAAAGTQEVSSNV--- | *Kiloniella laminariae* | Bacteria; Proteobacteria; Alphaproteobacteria; Kiloniellales; Kiloniellaceae; Kiloniella | methyl-accepting chemotaxis protein | 6 |
| | **WP_022730339.1** | 1.7E-06 | Qc.III | --DIGTIIASLNEIATTISAAVEEQGAATDEITRNVRGAADRTRSVSASI--- | *Thalassospira lucentensis* | Bacteria; Proteobacteria; Alphaproteobacteria; Rhodospirillales; Rhodospirillaceae; Thalassospira | HAMP domain-containing protein | 6 |
|  | **WP_027276275.1** | 1.7E-06 | Qa.IV | IKEIGQTISRMSEIASTIASAVEEQGAATQEISRNVQEAA------------- | *Rhodopseudomonas palustris* | Bacteria; Proteobacteria; Alphaproteobacteria; Rhizobiales; Bradyrhizobiaceae; Rhodopseudomonas | methyl-accepting chemotaxis protein | 6 |
|  | **WP_028688897.1** | 1.7E-06 | Qb.I | LHRTRLQLQETIEQSEFSSQELTASNEEMQAINEELRSASEELETSK------ | *Pseudomonas mosselii* | Bacteria; Proteobacteria; Gammaproteobacteria; Pseudomonadales; Pseudomonadaceae; Pseudomonas | PAS domain S-box protein | 6 |
|  | **WP_035647852.1** | 1.7E-06 | Qc.III | IKEISGTIERLSEIASTIAAAVEEQGAATQEISRNVQHASQGTKDV------- | *Bradyrhizobium sp. ORS 285* | Bacteria; Proteobacteria; Alphaproteobacteria; Rhizobiales; Bradyrhizobiaceae; Bradyrhizobium | methyl-accepting chemotaxis protein | 6 |
|  | **WP_038706119.1** | 1.7E-06 | Qb.I | LHRTRLQLQETIEQSEFSSQELTASNEEMQAINEELRSASEELETSK------ | *Pseudomonas mosselii* | Bacteria; Proteobacteria; Gammaproteobacteria; Pseudomonadales; Pseudomonadaceae; Pseudomonas | PAS domain S-box protein | 6 |
|  | **WP_041371032.1** | 1.7E-06 | Qa.IV | IRAIVSTITDLSEISMSVSSAVTEQAAVTQEIASNMQTAARSV---------- | *Methylorubrum populi* | Bacteria; Proteobacteria; Alphaproteobacteria; Rhizobiales; Methylobacteriaceae; Methylorubrum | PAS domain S-box protein | 6 |
|  | **WP_050406067.1** | 1.7E-06 | Qa.IV | -----TTIAEIDQVSGEIAAAVDQQGAATREISSNLQQAADRTRDVNK----- | *Bradyrhizobium embrapense* | Bacteria; Proteobacteria; Alphaproteobacteria; Rhizobiales; Bradyrhizobiaceae; Bradyrhizobium | chemotaxis protein | 6 |
|  | **WP_050705692.1** | 1.7E-06 | Qb.I | LHRTRLQLQETIEQSEFSSQELTASNEEMQAINEELRSASEELETSK------ | *Pseudomonas sp. 250J* | Bacteria; Proteobacteria; Gammaproteobacteria; Pseudomonadales; Pseudomonadaceae; Pseudomonas | PAS domain S-box protein | 6 |
|  | **WP_054167036.1** | 1.7E-06 | Qc.III.c | --EIGATVEQVNQISTSIASAVEEQNAATTEIARSARHASNGVETVT------ | *alpha proteobacterium AAP38* | Bacteria; Proteobacteria; Alphaproteobacteria | methyl-accepting chemotaxis protein | 6 |
|  | **WP_054894637.1** | 1.7E-06 | Qb.I | LHRTRLQLQETIEQSEFSSQELTASNEEMQAINEELRSASEELETSK------ | *Pseudomonas* | Bacteria; Proteobacteria; Gammaproteobacteria; Pseudomonadales; Pseudomonadaceae | MULTISPECIES: PAS domain S-box protein | 6 |
|  | **WP_060768515.1** | 1.7E-06 | Qa.IV | IRAIVSTITDLSEISMSVSSAVTEQAAVTQEIASNMQTAARSV---------- | *Methylobacterium sp. AMS5* | Bacteria; Proteobacteria; Alphaproteobacteria; Rhizobiales; Methylobacteriaceae; Methylobacterium | PAS domain S-box protein | 6 |
|  | **WP_062362439.1** | 1.7E-06 | Qb.I | LHRTRLQLQETIEQSEFSSQELTASNEEMQAINEELRSASEELETSK------ | *Pseudomonas mosselii* | Bacteria; Proteobacteria; Gammaproteobacteria; Pseudomonadales; Pseudomonadaceae; Pseudomonas | PAS domain S-box protein | 6 |
|  | **WP_062574102.1** | 1.7E-06 | Qb.I | LHRTRLQLQETIEQSEFSSQELTASNEEMQAINEELRSASEELETSK------ | *Pseudomonas putida* | Bacteria; Proteobacteria; Gammaproteobacteria; Pseudomonadales; Pseudomonadaceae; Pseudomonas | PAS domain S-box protein | 6 |
|  | **WP_063544624.1** | 1.7E-06 | Qb.I | LHRTRLQLQETIEQSEFSSQELTASNEEMQAINEELRSASEELETSK------ | *Pseudomonas putida* | Bacteria; Proteobacteria; Gammaproteobacteria; Pseudomonadales; Pseudomonadaceae; Pseudomonas | PAS domain S-box protein | 6 |
|  | **WP_063691911.1** | 1.7E-06 | Qa.IV | IKDISDTIEKLSEISSTIAAAVEEQGAATQEISRNVQQAA------------- | *Bradyrhizobium stylosanthis* | Bacteria; Proteobacteria; Alphaproteobacteria; Rhizobiales; Bradyrhizobiaceae; Bradyrhizobium | methyl-accepting chemotaxis protein | 6 |
|  | **WP_065863546.1** | 1.7E-06 | Qb.I | LHRTRLQLQETIEQSEFSSQELTASNEEMQAINEELRSASEELETSK------ | *Pseudomonas putida* | Bacteria; Proteobacteria; Gammaproteobacteria; Pseudomonadales; Pseudomonadaceae; Pseudomonas | PAS domain S-box protein | 6 |
|  | **WP_067555730.1** | 1.7E-06 | Qa.IV | --------AQINEISSTISAAVEEQNATTTEIARNVQEAASATEGVTGNIEK- | *Oceanibaculum pacificum* | Bacteria; Proteobacteria; Alphaproteobacteria; Rhodospirillales; Rhodospirillaceae; Oceanibaculum | methyl-accepting chemotaxis protein | 6 |
|  | **WP_068716178.1** | 1.7E-06 | Qa.III | LEQIKMSVSEISDMATQIATAAEEQTHVTGEITQNITAIK------------- | *Vibrio tritonius* | Bacteria; Proteobacteria; Gammaproteobacteria; Vibrionales; Vibrionaceae; Vibrio | methyl-accepting chemotaxis protein | 6 |
|  | **WP_069017602.1** | 1.7E-06 | Qb.I | LHRTRLQLQETIEQSEFSSQELTASNEEMQAINEELRSASEELETSK------ | *Pseudomonas mosselii* | Bacteria; Proteobacteria; Gammaproteobacteria; Pseudomonadales; Pseudomonadaceae; Pseudomonas | PAS domain S-box protein | 6 |
| | **WP_083222885.1** | 1.7E-06 | Qc.III | --DISRTIHEINEVSAAIAAAVEEQGAATSDISTKVSHVADQSTKVSDRIG-- | *Terasakiella sp. PR1* | Bacteria; Proteobacteria; Alphaproteobacteria; Rhizobiales; Methylocystaceae; Terasakiella | hypothetical protein | 6 |
|  | **WP_084942255.1** | 1.7E-06 | Qb.I | LHRTRLQLQETIEQSEFSSQELTASNEEMQAINEELRSASEELETSK------ | *Pseudomonas mosselii* | Bacteria; Proteobacteria; Gammaproteobacteria; Pseudomonadales; Pseudomonadaceae; Pseudomonas | PAS domain S-box protein | 6 |
|  | **WP_088852815.1** | 1.7E-06 | Qb.I | LHRTRLQLQETIEQSEFSSQELTASNEEMQAINEELRSASEELETSK------ | *Pseudomonas sp. DrBHI1* | Bacteria; Proteobacteria; Gammaproteobacteria; Pseudomonadales; Pseudomonadaceae; Pseudomonas | PAS domain S-box protein | 6 |
|  | **WP_092187664.1** | 1.7E-06 | Qa.IV | -RDISDTIEKLSEISSTIAAAVEEQGAATQEISRNVQQAAQ------------ | *Bradyrhizobium sp. cf659* | Bacteria; Proteobacteria; Alphaproteobacteria; Rhizobiales; Bradyrhizobiaceae; Bradyrhizobium | methyl-accepting chemotaxis protein | 6 |
|  | **WP_094009891.1** | 1.7E-06 | Qb.I | LHRTRLQLQETIEQSEFSSQELTASNEEMQAINEELRSASEELETSK------ | *Pseudomonas* | Bacteria; Proteobacteria; Gammaproteobacteria; Pseudomonadales; Pseudomonadaceae | MULTISPECIES: PAS domain S-box protein | 6 |
|  | **WP_096050099.1** | 1.7E-06 | Qb.I | LHRTRLQLQETIEQSEFSSQELTASNEEMQAINEELRSASEELETSK------ | *Pseudomonas mosselii* | Bacteria; Proteobacteria; Gammaproteobacteria; Pseudomonadales; Pseudomonadaceae; Pseudomonas | PAS domain S-box protein | 6 |
| | **WP_101304516.1** | 1.7E-06 | Qc.III | --DIGTIIASLNEIATTISAAVEEQGAATDEITRNVRGAADRTRSVSASI--- | *Thalassospira lohafexi* | Bacteria; Proteobacteria; Alphaproteobacteria; Rhodospirillales; Rhodospirillaceae; Thalassospira | HAMP domain-containing protein | 6 |
|  | **WP_102111701.1** | 1.7E-06 | Qc.III.c | --EIGATVEQVNQISTSIASAVEEQNAATTEIARSARHASNGVETVT------ | *Niveispirillum cyanobacteriorum* | Bacteria; Proteobacteria; Alphaproteobacteria; Rhodospirillales; Rhodospirillaceae; Niveispirillum | methyl-accepting chemotaxis protein | 6 |
|  | **WP_108914330.1** | 1.7E-06 | Qa.IV | IKDISDTIEKLSEISSTIAAAVEEQGAATQEISRNVQQAAQ------------ | *Bradyrhizobium arachidis* | Bacteria; Proteobacteria; Alphaproteobacteria; Rhizobiales; Bradyrhizobiaceae; Bradyrhizobium | methyl-accepting chemotaxis protein | 6 |
|  | **WP_110116006.1** | 1.7E-06 | Qa.IV | -RDISDTIEKLSEISSTIAAAVEEQGAATQEISRNVQQAAQ------------ | *Bradyrhizobium diazoefficiens* | Bacteria; Proteobacteria; Alphaproteobacteria; Rhizobiales; Bradyrhizobiaceae; Bradyrhizobium | methyl-accepting chemotaxis protein | 6 |
|  | **WP_110604718.1** | 1.7E-06 | Qb.I | LHRTRLQLQETIEQSEFSSQELTASNEEMQAINEELRSASEELETSK------ | *Pseudomonas soli* | Bacteria; Proteobacteria; Gammaproteobacteria; Pseudomonadales; Pseudomonadaceae; Pseudomonas | PAS domain S-box protein | 6 |
|  | **WP_110703291.1** | 1.7E-06 | Qb.I | LHRTRLQLQETIEQSEFSSQELTASNEEMQAINEELRSASEELETSK------ | *Pseudomonas soli* | Bacteria; Proteobacteria; Gammaproteobacteria; Pseudomonadales; Pseudomonadaceae; Pseudomonas | PAS domain S-box protein | 6 |
|  | **WP_111532877.1** | 1.7E-06 | Qb.I | LHRTRLQLQETIEQSEFSSQELTASNEEMQAINEELRSASEELETSK------ | *Pseudomonas sp. 2848* | Bacteria; Proteobacteria; Gammaproteobacteria; Pseudomonadales; Pseudomonadaceae; Pseudomonas | PAS domain S-box protein | 6 |
|  | **WP_112898063.1** | 1.7E-06 | Qb.I | LHRTRLQLQETIEQSEFSSQELTASNEEMQAINEELRSASEELETSK------ | *Pseudomonas putida* | Bacteria; Proteobacteria; Gammaproteobacteria; Pseudomonadales; Pseudomonadaceae; Pseudomonas | PAS domain S-box protein | 6 |
| | **AWL99845.1** | 1.8E-06 | Qc.III | IKDISHTIERLSEISSTIAAAVEEQGAATQEISRNVQQAAEGTHQVSSNITD- | *Bradyrhizobium sp. 2 39S1MB* | Bacteria; Proteobacteria; Alphaproteobacteria; Rhizobiales; Bradyrhizobiaceae; Bradyrhizobium | methyl-accepting chemotaxis protein | 6 |
|  | **BAM86516.1** | 1.8E-06 | Qa.IV | VKQVATAMRQMADVTTAIASAVEQQGAATREISHNVQMAA------------- | *Bradyrhizobium oligotrophicum S58* | Bacteria; Proteobacteria; Alphaproteobacteria; Rhizobiales; Bradyrhizobiaceae; Bradyrhizobium | methyl-accepting chemotaxis protein | 6 |
|  | **CEG06835.1** | 1.8E-06 | Qa.IV | IKEISTTISEISVITTTIAAAVEEQGVATQEIARNVQQAA------------- | *Afipia felis* | Bacteria; Proteobacteria; Alphaproteobacteria; Rhizobiales; Bradyrhizobiaceae; Afipia | Methyl-accepting chemotaxis protein 2 | 6 |
|  | **EFI52578.1** | 1.8E-06 | Qa.IV | IKEISTTISEISVITTTIAAAVEEQGVATQEIARNVQQAA------------- | *Afipia sp. 1NLS2* | Bacteria; Proteobacteria; Alphaproteobacteria; Rhizobiales; Bradyrhizobiaceae; Afipia | methyl-accepting chemotaxis sensory transducer | 6 |
|  | **PIT02460.1** | 1.8E-06 | Qa.III | IQSICATIQQVDEISAAIAAAVGQQGTATQEIAKNVQQAA------------- | *Bradyrhizobium sp. TSA1* | Bacteria; Proteobacteria; Alphaproteobacteria; Rhizobiales; Bradyrhizobiaceae; Bradyrhizobium | chemotaxis protein | 6 |
|  | **SFP41929.1** | 1.8E-06 | Qa.IV | IKDISDTIEKLSEISSTIAAAVEEQGAATQEISRNVQQAA------------- | *Bradyrhizobium sp. Ghvi* | Bacteria; Proteobacteria; Alphaproteobacteria; Rhizobiales; Bradyrhizobiaceae; Bradyrhizobium | Methyl-accepting chemotaxis protein | 6 |
| | **WP_007001909.1** | 1.8E-06 | Qc.III | --GIGSTVERTNEISTAIAAAVEQQGAATQEIARSASQVAEATGTVTQRIAR- | *Roseomonas cervicalis* | Bacteria; Proteobacteria; Alphaproteobacteria; Rhodospirillales; Acetobacteraceae; Roseomonas | HAMP domain-containing protein | 6 |
|  | **WP_011663483.1** | 1.8E-06 | Qa.IV | IKEIGATISQMAEIASTIASAVEQQGAATQEISRNVQQAA------------- | *Rhodopseudomonas palustris* | Bacteria; Proteobacteria; Alphaproteobacteria; Rhizobiales; Bradyrhizobiaceae; Rhodopseudomonas | chemotaxis protein | 6 |
| | **WP_022663879.1** | 1.8E-06 | Qc.III | ---IENQVGNLLDVSQNIQTAVGNQETMTSRIAENAQKTSADIRDVTGRISRV | *Desulfospira joergensenii* | Bacteria; Proteobacteria; Deltaproteobacteria; Desulfobacterales; Desulfobacteraceae; Desulfospira | HAMP domain-containing protein | 6 |

| | | | | | | | | |
|---|---|---|---|---|---|---|---|---|
|  | **WP_028335948.1** | 1.8E-06 | Qa.IV | -----TTIAEIDQVSGEIAAAVDQQGAATREISSNLQQAADRTRDVNK----- | *Bradyrhizobium elkanii* | Bacteria; Proteobacteria; Alphaproteobacteria; Rhizobiales; Bradyrhizobiaceae; Bradyrhizobium | chemotaxis protein | 6 |
|  | **WP_028482170.1** | 1.8E-06 | Qa.III.b | -GEIAKTMGDVNDITQSIAAAVEQQGAATTEISDNVQQAAQ------------ | *Nesiotobacter exalbescens* | Bacteria; Proteobacteria; Alphaproteobacteria; Rhodobacterales; Rhodobacteraceae; Nesiotobacter | methyl-accepting chemotaxis protein | 6 |
|  | **WP_035970310.1** | 1.8E-06 | Qa.IV | IKDISDTIEKLSEISSTIAAAVEEQGAATQEISRNVQQAA------------- | *Bradyrhizobium sp. WSM3983* | Bacteria; Proteobacteria; Alphaproteobacteria; Rhizobiales; Bradyrhizobiaceae; Bradyrhizobium | methyl-accepting chemotaxis protein | 6 |
|  | **WP_040297761.1** | 1.8E-06 | Qa.IV | IKEISTTISEISVITTTIAAAVEEQGVATQEIARNVQQAA------------- | *Afipia sp. 1NLS2* | Bacteria; Proteobacteria; Alphaproteobacteria; Rhizobiales; Bradyrhizobiaceae; Afipia | HAMP domain-containing protein | 6 |
|  | **WP_048755547.1** | 1.8E-06 | Qa.IV | IKEISTTISEISVITTTIAAAVEEQGVATQEIARNVQQAA------------- | *Afipia felis* | Bacteria; Proteobacteria; Alphaproteobacteria; Rhizobiales; Bradyrhizobiaceae; Afipia | HAMP domain-containing protein | 6 |
| | **WP_059398873.1** | 1.8E-06 | Qc.III | --AIGGIIGQINDISTAIAAAIEEQGAATNEISRNVQEAATGTHQVSGSIT-- | *Azospirillum* | Bacteria; Proteobacteria; Alphaproteobacteria; Rhodospirillales; Rhodospirillaceae | MULTISPECIES: HAMP domain-containing protein | 6 |
|  | **WP_074121594.1** | 1.8E-06 | Qa.IV | IKDISDTIEKLSEISSTIAAAVEEQGAATQEISRNVQQAA------------- | *Bradyrhizobium sp. AS23.2* | Bacteria; Proteobacteria; Alphaproteobacteria; Rhizobiales; Bradyrhizobiaceae; Bradyrhizobium | methyl-accepting chemotaxis protein | 6 |
|  | **WP_084807381.1** | 1.8E-06 | Qa.IV | IKDISDTIEKLSEISSTIAAAVEEQGAATQEISRNVQQAA------------- | *Bradyrhizobium sp. NAS80.1* | Bacteria; Proteobacteria; Alphaproteobacteria; Rhizobiales; Bradyrhizobiaceae; Bradyrhizobium | methyl-accepting chemotaxis protein | 6 |
|  | **WP_092078827.1** | 1.8E-06 | Qa.IV | -----TTIAEIDQVSGEIAAAVDQQGAATREISSNLQQAADRTRDVNK----- | *Bradyrhizobium sp. R5* | Bacteria; Proteobacteria; Alphaproteobacteria; Rhizobiales; Bradyrhizobiaceae; Bradyrhizobium | chemotaxis protein | 6 |
|  | **WP_092298463.1** | 1.8E-06 | Qa.IV | IKDISDTIEKLSEISSTIAAAVEEQGAATQEISRNVQQAA------------- | *Bradyrhizobium sp. Ghvi* | Bacteria; Proteobacteria; Alphaproteobacteria; Rhizobiales; Bradyrhizobiaceae; Bradyrhizobium | methyl-accepting chemotaxis protein | 6 |
|  | **WP_096628513.1** | 1.8E-06 | Qc.III | -DEISSNLANIKTLSGDMNEAIDQLGPIVSDLKDNVDSLSSAIKH-------- | *Salinivibrio sp. YCSC6* | Bacteria; Proteobacteria; Gammaproteobacteria; Vibrionales; Vibrionaceae; Salinivibrio | methyl-accepting chemotaxis protein | 6 |
|  | **AUT57643.1** | 1.9E-06 | Qb.I | LQRTKEQLQSTIEQSETSTEELKASNEELQAINEELRSATEELETS------- | *Paraburkholderia caribensis* | Bacteria; Proteobacteria; Betaproteobacteria; Burkholderiales; Burkholderiaceae; Paraburkholderia | PAS domain S-box protein | 6 |
| | **BAI72055.1** | 1.9E-06 | Qa.III | IRGIADTVRQVNQIATSIASAVEEQTAATGEISRNVQQAAQGTAEVSRSI--- | *Azospirillum sp. B510* | Bacteria; Proteobacteria; Alphaproteobacteria; Rhodospirillales; Rhodospirillaceae; Azospirillum | methyl-accepting chemotaxis protein | 6 |
|  | **CCE00404.1** | 1.9E-06 | Qa.IV | VKEVATAMRQMADVTTAIASAVEQQGAATREISHNVQMAA------------- | *Bradyrhizobium sp. STM 3809* | Bacteria; Proteobacteria; Alphaproteobacteria; Rhizobiales; Bradyrhizobiaceae; Bradyrhizobium | Methyl-accepting chemotaxis protein (MCP) | 6 |
|  | **EEA02744.1** | 1.9E-06 | Qb.I | LQRTKEQLQSTIEQSETSTEELKASNEELQAINEELRSATEELETS------- | *Burkholderia sp. H160* | Bacteria; Proteobacteria; Betaproteobacteria; Burkholderiales; Burkholderiaceae; Burkholderia | signal transduction histidine kinase with CheB and CheR activity | 6 |
|  | **EEE47967.1** | 1.9E-06 | Qc.III | IQSISQTIGRIKDISDTISGAVSEQGIATREIADSTQQAAS------------ | *Labrenzia alexandrii DFL-11* | Bacteria; Proteobacteria; Alphaproteobacteria; Rhodobacterales; Rhodobacteraceae; Labrenzia | Methyl-accepting chemotaxis protein signaling domain | 6 |
|  | **EKM33263.1** | 1.9E-06 | Qa.III | -SKIREAIQQISDMATQIASAAEEQSLVTAEINANTESVRE------------ | *Vibrio harveyi* | Bacteria; Proteobacteria; Gammaproteobacteria; Vibrionales; Vibrionaceae; Vibrio | methyl-accepting chemotaxis (MCP) signaling domain protein | 6 |
|  | **ERS05909.1** | 1.9E-06 | Qa.IV | IQEIASTMEEVNQFTAAIAAAISQQGDATNEISHNVQQAA------------- | *Labrenzia sp. C1B70* | Bacteria; Proteobacteria; Alphaproteobacteria; Rhodobacterales; Rhodobacteraceae; Labrenzia | hypothetical protein Q675_29765 | 6 |
|  | **KIZ33810.1** | 1.9E-06 | Qa.IV | ---ITTTIEEISAISATIAAAIEQQGAATSEIARNVTQTAQATQD-------- | *Rhodopseudomonas palustris* | Bacteria; Proteobacteria; Alphaproteobacteria; Rhizobiales; Bradyrhizobiaceae; Rhodopseudomonas | chemotaxis protein partial | 6 |
|  | **KVC58984.1** | 1.9E-06 | Qb.I | LQRTKEQLQSTIEQSETSTEELKASNEELQAINEELRSATEELETS------- | *Burkholderia stagnalis* | Bacteria; Proteobacteria; Betaproteobacteria; Burkholderiales; Burkholderiaceae; Burkholderia; Burkholderia cepacia complex | chemotaxis protein | 6 |
|  | **KVC69404.1** | 1.9E-06 | Qb.I | LQRTKEQLQSTIEQSETSTEELKASNEELQAINEELRSATEELETS------- | *Burkholderia ubonensis* | Bacteria; Proteobacteria; Betaproteobacteria; Burkholderiales; Burkholderiaceae; Burkholderia; Burkholderia cepacia complex | chemotaxis protein | 6 |
|  | **KVC73002.1** | 1.9E-06 | Qb.I | LQRTKEQLQSTIEQSETSTEELKASNEELQAINEELRSATEELETS------- | *Burkholderia ubonensis* | Bacteria; Proteobacteria; Betaproteobacteria; Burkholderiales; Burkholderiaceae; Burkholderia; Burkholderia cepacia complex | chemotaxis protein | 6 |
|  | **KVC88384.1** | 1.9E-06 | Qb.I | LQRTKEQLQSTIEQSETSTEELKASNEELQAINEELRSATEELETS------- | *Burkholderia ubonensis* | Bacteria; Proteobacteria; Betaproteobacteria; Burkholderiales; Burkholderiaceae; Burkholderia; Burkholderia cepacia complex | chemotaxis protein | 6 |
|  | **KVC97579.1** | 1.9E-06 | Qb.I | LQRTKEQLQSTIEQSETSTEELKASNEELQAINEELRSATEELETS------- | *Burkholderia ubonensis* | Bacteria; Proteobacteria; Betaproteobacteria; Burkholderiales; Burkholderiaceae; Burkholderia; Burkholderia cepacia complex | chemotaxis protein | 6 |
|  | **KVD25417.1** | 1.9E-06 | Qb.I | LQRTKEQLQSTIEQSETSTEELKASNEELQAINEELRSATEELETS------- | *Burkholderia ubonensis* | Bacteria; Proteobacteria; Betaproteobacteria; Burkholderiales; Burkholderiaceae; Burkholderia; Burkholderia cepacia complex | chemotaxis protein | 6 |
|  | **KVD31535.1** | 1.9E-06 | Qb.I | LQRTKEQLQSTIEQSETSTEELKASNEELQAINEELRSATEELETS------- | *Burkholderia ubonensis* | Bacteria; Proteobacteria; Betaproteobacteria; Burkholderiales; Burkholderiaceae; Burkholderia; Burkholderia cepacia complex | chemotaxis protein | 6 |
|  | **KVD73791.1** | 1.9E-06 | Qb.I | LQRTKEQLQSTIEQSETSTEELKASNEELQAINEELRSATEELETS------- | *Burkholderia ubonensis* | Bacteria; Proteobacteria; Betaproteobacteria; Burkholderiales; Burkholderiaceae; Burkholderia; Burkholderia cepacia complex | chemotaxis protein | 6 |
|  | **KVD81202.1** | 1.9E-06 | Qb.I | LQRTKEQLQSTIEQSETSTEELKASNEELQAINEELRSATEELETS------- | *Burkholderia ubonensis* | Bacteria; Proteobacteria; Betaproteobacteria; Burkholderiales; Burkholderiaceae; Burkholderia; Burkholderia cepacia complex | chemotaxis protein | 6 |
|  | **KVD87057.1** | 1.9E-06 | Qb.I | LQRTKEQLQSTIEQSETSTEELKASNEELQAINEELRSATEELETS------- | *Burkholderia ubonensis* | Bacteria; Proteobacteria; Betaproteobacteria; Burkholderiales; Burkholderiaceae; Burkholderia; Burkholderia cepacia complex | chemotaxis protein | 6 |
|  | **OAN11140.1** | 1.9E-06 | Qa.III | LEQINNSVALISDMATQIATAAEEQTHVTDEITQNTTTIK------------- | *Photobacterium jeanii* | Bacteria; Proteobacteria; Gammaproteobacteria; Vibrionales; Vibrionaceae; Photobacterium | chemotaxis protein | 6 |
|  | **OHC52984.1** | 1.9E-06 | Qc.III | --GISDAVSNIADMADQIAAAAEEQSSVADEINRNITNIAQLSD--------- | *Pseudomonadales bacterium RIFCSPLOWO2_02_FULL_63_210* | Bacteria; Proteobacteria; Gammaproteobacteria; Pseudomonadales | hypothetical protein A3J25_10995 | 6 |
| | **OOM40599.1** | 1.9E-06 | Qb.I | ---INKLVSDTVKQSHDSAEYINESSYLISGTADTFDNIFNKINSANDLIQE- | *Clostridium beijerinckii* | Bacteria; Firmicutes; Clostridia; Clostridiales; Clostridiaceae; Clostridium | methyl-accepting chemotaxis protein PctC | 6 |
|  | **PHQ89084.1** | 1.9E-06 | Qa.III | ------SVTTINDMSTQIATAATEQSSVTEEINRNVY----AIQEIVNELT-- | *Pseudoalteromonas sp.* | Bacteria; Proteobacteria; Gammaproteobacteria; Alteromonadales; Pseudoalteromonadaceae; Pseudoalteromonas | chemotaxis protein partial | 6 |
|  | **PIE60210.1** | 1.9E-06 | Qa.III | IDGIVKIINQINEIISTITTAVEEQSVSTREITANITQANE------------ | *Desulfobulbus propionicus* | Bacteria; Proteobacteria; Deltaproteobacteria; Desulfobacterales; Desulfobulbaceae; Desulfobulbus | hypothetical protein CSA32_00045 | 6 |
|  | **PST90659.1** | 1.9E-06 | Qa.III | LEQINNSVALISDMATQIATAAEEQTHVTDEITQNTTTIK------------- | *Photobacterium jeanii* | Bacteria; Proteobacteria; Gammaproteobacteria; Vibrionales; Vibrionaceae; Photobacterium | methyl-accepting chemotaxis protein | 6 |
| | **PWC32442.1** | 1.9E-06 | Qa.III | IRGIADTVQQVNQIATSIASAVEEQTAATGEISRNVQQAAQGTAEVSRSI--- | *Azospirillum sp. TSO35-2* | Bacteria; Proteobacteria; Alphaproteobacteria; Rhodospirillales; Rhodospirillaceae; Azospirillum | chemotaxis protein | 6 |
| | **RAI19905.1** | 1.9E-06 | Qa.III | IESITGTIKQIDEISSTIASSVEEQGSATGEIAQNVQQAAQGTQEVSTSIV-- | *Afifella marina DSM 2698* | Bacteria; Proteobacteria; Alphaproteobacteria; Rhizobiales; Rhodobiaceae; Afifella | methyl-accepting chemotaxis protein | 6 |
| | **SOD93684.1** | 1.9E-06 | Qc.III | -QGITGTIGRINEIAAAIAAAVEQQGAATQEIARNVQQASSGTSEVTSNIS-- | *Caenispirillum bisanense* | Bacteria; Proteobacteria; Alphaproteobacteria; Rhodospirillales; Rhodospirillaceae; Caenispirillum | Methyl-accepting chemotaxis protein | 6 |
| | **SQB01419.1** | 1.9E-06 | Qb.I | ---INKLVSDTVKQSHDSAEYINESSYLISGTADTFDNIFNKINSANDLIQE- | *Clostridium beijerinckii* | Bacteria; Firmicutes; Clostridia; Clostridiales; Clostridiaceae; Clostridium | methyl-accepting chemotaxis sensory transducer | 6 |
|  | **WP_010099354.1** | 1.9E-06 | Qb.I | LQRTKEQLQSTIEQSETSTEELKASNEELQAINEELRSATEELETS------- | *Burkholderia ubonensis* | Bacteria; Proteobacteria; Betaproteobacteria; Burkholderiales; Burkholderiaceae; Burkholderia; Burkholderia cepacia complex | PAS domain S-box protein | 6 |
| | **WP_011502286.1** | 1.9E-06 | Qc.III | IKEIGDTIGRLSEISSIIASAVEQQDAVTQEITHNVQDASNGTRQVTSHIGD- | *Rhodopseudomonas palustris* | Bacteria; Proteobacteria; Alphaproteobacteria; Rhizobiales; Bradyrhizobiaceae; Rhodopseudomonas | PAS domain S-box protein | 6 |
|  | **WP_012405588.1** | 1.9E-06 | Qb.I | LQRTKEQLQSTIEQSETSTEELKASNEELQAINEELRSATEELETS------- | *Paraburkholderia phymatum* | Bacteria; Proteobacteria; Betaproteobacteria; Burkholderiales; Burkholderiaceae; Paraburkholderia | PAS domain S-box protein | 6 |
|  | **WP_013094103.1** | 1.9E-06 | Qb.I | LQRTKEQLQSTIEQSETSTEELKASNEELQAINEELRSATEELETS------- | *Burkholderia sp. CCGE1002* | Bacteria; Proteobacteria; Betaproteobacteria; Burkholderiales; Burkholderiaceae; Burkholderia | PAS domain S-box protein | 6 |
| | **WP_017211047.1** | 1.9E-06 | Qb.I | ---INKLVSDTVKQSHDSAEYINESSYLISGTADTFDNIFNKINSANDLIQE- | *Clostridium beijerinckii* | Bacteria; Firmicutes; Clostridia; Clostridiales; Clostridiaceae; Clostridium | methyl-accepting chemotaxis protein | 6 |
|  | **WP_018422248.1** | 1.9E-06 | Qb.I | LQRTKEQLQSTIEQSETSTEELKASNEELQAINEELRSATEELETS------- | *Paraburkholderia tuberum* | Bacteria; Proteobacteria; Betaproteobacteria; Burkholderiales; Burkholderiaceae; Paraburkholderia | PAS domain S-box protein | 6 |
|  | **WP_018434944.1** | 1.9E-06 | Qb.I | LQRTKEQLQSTIEQSETSTEELKASNEELQAINEELRSATEELETS------- | *Burkholderia sp. JPY251* | Bacteria; Proteobacteria; Betaproteobacteria; Burkholderiales; Burkholderiaceae; Burkholderia | PAS domain S-box protein | 6 |
| | **WP_023976472.1** | 1.9E-06 | Qb.I | ---INKLVSDTVKQSHDSAEYINESSYLISGTADTFDNIFNKINSANDLIQE- | *Clostridium* | Bacteria; Firmicutes; Clostridia; Clostridiales; Clostridiaceae | MULTISPECIES: methyl-accepting chemotaxis protein | 6 |
|  | **WP_025482523.1** | 1.9E-06 | R.IV | IHLIQENLNQITDVVQNNVATAEASSAAAEELNHQAEALKQS----------- | *Clostridiales bacterium VE202-28* | Bacteria; Firmicutes; Clostridia; Clostridiales | methyl-accepting chemotaxis protein | 6 |
|  | **WP_027194809.1** | 1.9E-06 | Qb.I | LQRTKEQLQSTIEQSETSTEELKASNEELQAINEELRSATEELETS------- | *Paraburkholderia sprentiae* | Bacteria; Proteobacteria; Betaproteobacteria; Burkholderiales; Burkholderiaceae; Paraburkholderia | PAS domain S-box protein | 6 |
|  | **WP_027316793.1** | 1.9E-06 | Qa.IV | IQEIARTITEMSQISTSIAAAMEEQGAATAEIARNVQEAA------------- | *Microvirga flocculans* | Bacteria; Proteobacteria; Alphaproteobacteria; Rhizobiales; Methylobacteriaceae; Microvirga | HAMP domain-containing protein | 6 |
|  | **WP_028204293.1** | 1.9E-06 | Qb.I | LQRTKEQLQSTIEQSETSTEELKASNEELQAINEELRSATEELETS------- | *Paraburkholderia nodosa* | Bacteria; Proteobacteria; Betaproteobacteria; Burkholderiales; Burkholderiaceae; Paraburkholderia | PAS domain S-box protein | 6 |
|  | **WP_028344135.1** | 1.9E-06 | Qa.IV | -----TTIAEIDQVSGEIAAAVDQQGAATREISSNLQQAADRTRDVNK----- | *Bradyrhizobium elkanii* | Bacteria; Proteobacteria; Alphaproteobacteria; Rhizobiales; Bradyrhizobiaceae; Bradyrhizobium | chemotaxis protein | 6 |
|  | **WP_028364432.1** | 1.9E-06 | Qb.I | LQRTKEQLQSTIEQSETSTEELKASNEELQAINEELRSATEELETS------- | *Burkholderia sp. UYPR1.413* | Bacteria; Proteobacteria; Betaproteobacteria; Burkholderiales; Burkholderiaceae; Burkholderia | PAS domain S-box protein | 6 |
|  | **WP_035991715.1** | 1.9E-06 | Qb.I | LQRTKEQLQSTIEQSETSTEELKASNEELQAINEELRSATEELETS------- | *Paraburkholderia caribensis* | Bacteria; Proteobacteria; Betaproteobacteria; Burkholderiales; Burkholderiaceae; Paraburkholderia | PAS domain S-box protein | 6 |
|  | **WP_038385637.1** | 1.9E-06 | Qa.IV | -----TTIAEIDQVSGEIAAAVDQQGAATREISSNLQQAADRTRDVNK----- | *Bradyrhizobium elkanii* | Bacteria; Proteobacteria; Alphaproteobacteria; Rhizobiales; Bradyrhizobiaceae; Bradyrhizobium | chemotaxis protein | 6 |
| | **WP_041898066.1** | 1.9E-06 | Qb.I | ---INKLVSDTVKQSHDSAEYINESSYLISGTADTFDNIFNKINSANDLIQE- | *Clostridium beijerinckii* | Bacteria; Firmicutes; Clostridia; Clostridiales; Clostridiaceae; Clostridium | methyl-accepting chemotaxis protein | 6 |
|  | **WP_042338593.1** | 1.9E-06 | Qa.IV | VKQVATAMRQMADVTTAIASAVEQQGAATREISHNVQMAA------------- | *Bradyrhizobium oligotrophicum* | Bacteria; Proteobacteria; Alphaproteobacteria; Rhizobiales; Bradyrhizobiaceae; Bradyrhizobium | methyl-accepting chemotaxis protein | 6 |
|  | **WP_042340906.1** | 1.9E-06 | Qc.III | -REISGTIERLSEIASTIAAAVEEQGAATQEISRNVQHASQGTKDVS------ | *Bradyrhizobium oligotrophicum* | Bacteria; Proteobacteria; Alphaproteobacteria; Rhizobiales; Bradyrhizobiaceae; Bradyrhizobium | HAMP domain-containing protein | 6 |
|  | **WP_042587668.1** | 1.9E-06 | Qb.I | LQRTKEQLQSTIEQSETSTEELKASNEELQAINEELRSATEELETS------- | *Burkholderia sp. MSHR3999* | Bacteria; Proteobacteria; Betaproteobacteria; Burkholderiales; Burkholderiaceae; Burkholderia | PAS domain S-box protein | 6 |

| | | | | | | | | |
|---|---|---|---|---|---|---|---|---|
|  | **WP_044848673.1** | 1.9E-06 | Qb.I | LQRTKEQLQSTIEQSETSTEELKASNEELQAINEELRSATEELETS------- | *Burkholderia sp. USM B20* | Bacteria; Proteobacteria; Betaproteobacteria; Burkholderiales; Burkholderiaceae; Burkholderia | PAS domain S-box protein | 6 |
|  | **WP_045564790.1** | 1.9E-06 | Qb.I | LQRTKEQLQSTIEQSETSTEELKASNEELQAINEELRSATEELETS------- | *Burkholderia ubonensis* | Bacteria; Proteobacteria; Betaproteobacteria; Burkholderiales; Burkholderiaceae; Burkholderia; Burkholderia cepacia complex | PAS domain S-box protein | 6 |
|  | **WP_054931578.1** | 1.9E-06 | Qb.I | LQRTKEQLQSTIEQSETSTEELKASNEELQAINEELRSATEELETS------- | *Burkholderia sp. E7m48* | Bacteria; Proteobacteria; Betaproteobacteria; Burkholderiales; Burkholderiaceae; Burkholderia | PAS domain S-box protein | 6 |
|  | **WP_059586104.1** | 1.9E-06 | Qb.I | LQRTKEQLQSTIEQSETSTEELKASNEELQAINEELRSATEELETS------- | *Burkholderia ubonensis* | Bacteria; Proteobacteria; Betaproteobacteria; Burkholderiales; Burkholderiaceae; Burkholderia; Burkholderia cepacia complex | PAS domain S-box protein | 6 |
|  | **WP_059612396.1** | 1.9E-06 | Qb.I | LQRTKEQLQSTIEQSETSTEELKASNEELQAINEELRSATEELETS------- | *Burkholderia ubonensis* | Bacteria; Proteobacteria; Betaproteobacteria; Burkholderiales; Burkholderiaceae; Burkholderia; Burkholderia cepacia complex | PAS domain S-box protein | 6 |
|  | **WP_059629308.1** | 1.9E-06 | Qb.I | LQRTKEQLQSTIEQSETSTEELKASNEELQAINEELRSATEELETS------- | *Burkholderia ubonensis* | Bacteria; Proteobacteria; Betaproteobacteria; Burkholderiales; Burkholderiaceae; Burkholderia; Burkholderia cepacia complex | PAS domain S-box protein | 6 |
|  | **WP_059635950.1** | 1.9E-06 | Qb.I | LQRTKEQLQSTIEQSETSTEELKASNEELQAINEELRSATEELETS------- | *Burkholderia ubonensis* | Bacteria; Proteobacteria; Betaproteobacteria; Burkholderiales; Burkholderiaceae; Burkholderia; Burkholderia cepacia complex | PAS domain S-box protein | 6 |
|  | **WP_059640434.1** | 1.9E-06 | Qb.I | LQRTKEQLQSTIEQSETSTEELKASNEELQAINEELRSATEELETS------- | *Burkholderia ubonensis* | Bacteria; Proteobacteria; Betaproteobacteria; Burkholderiales; Burkholderiaceae; Burkholderia; Burkholderia cepacia complex | PAS domain S-box protein | 6 |
|  | **WP_059650181.1** | 1.9E-06 | Qb.I | LQRTKEQLQSTIEQSETSTEELKASNEELQAINEELRSATEELETS------- | *Burkholderia ubonensis* | Bacteria; Proteobacteria; Betaproteobacteria; Burkholderiales; Burkholderiaceae; Burkholderia; Burkholderia cepacia complex | PAS domain S-box protein | 6 |
|  | **WP_059654486.1** | 1.9E-06 | Qb.I | LQRTKEQLQSTIEQSETSTEELKASNEELQAINEELRSATEELETS------- | *Burkholderia ubonensis* | Bacteria; Proteobacteria; Betaproteobacteria; Burkholderiales; Burkholderiaceae; Burkholderia; Burkholderia cepacia complex | PAS domain S-box protein | 6 |
|  | **WP_059659041.1** | 1.9E-06 | Qb.I | LQRTKEQLQSTIEQSETSTEELKASNEELQAINEELRSATEELETS------- | *Burkholderia ubonensis* | Bacteria; Proteobacteria; Betaproteobacteria; Burkholderiales; Burkholderiaceae; Burkholderia; Burkholderia cepacia complex | PAS domain S-box protein | 6 |
|  | **WP_059660366.1** | 1.9E-06 | Qb.I | LQRTKEQLQSTIEQSETSTEELKASNEELQAINEELRSATEELETS------- | *Burkholderia ubonensis* | Bacteria; Proteobacteria; Betaproteobacteria; Burkholderiales; Burkholderiaceae; Burkholderia; Burkholderia cepacia complex | PAS domain S-box protein | 6 |
|  | **WP_059673887.1** | 1.9E-06 | Qb.I | LQRTKEQLQSTIEQSETSTEELKASNEELQAINEELRSATEELETS------- | *Burkholderia ubonensis* | Bacteria; Proteobacteria; Betaproteobacteria; Burkholderiales; Burkholderiaceae; Burkholderia; Burkholderia cepacia complex | PAS domain S-box protein | 6 |
|  | **WP_059711385.1** | 1.9E-06 | Qb.I | LQRTKEQLQSTIEQSETSTEELKASNEELQAINEELRSATEELETS------- | *Burkholderia ubonensis* | Bacteria; Proteobacteria; Betaproteobacteria; Burkholderiales; Burkholderiaceae; Burkholderia; Burkholderia cepacia complex | PAS domain S-box protein | 6 |
|  | **WP_059713347.1** | 1.9E-06 | Qb.I | LQRTKEQLQSTIEQSETSTEELKASNEELQAINEELRSATEELETS------- | *Burkholderia ubonensis* | Bacteria; Proteobacteria; Betaproteobacteria; Burkholderiales; Burkholderiaceae; Burkholderia; Burkholderia cepacia complex | PAS domain S-box protein | 6 |
|  | **WP_059728565.1** | 1.9E-06 | Qb.I | LQRTKEQLQSTIEQSETSTEELKASNEELQAINEELRSATEELETS------- | *Burkholderia ubonensis* | Bacteria; Proteobacteria; Betaproteobacteria; Burkholderiales; Burkholderiaceae; Burkholderia; Burkholderia cepacia complex | PAS domain S-box protein | 6 |
|  | **WP_059756884.1** | 1.9E-06 | Qb.I | LQRTKEQLQSTIEQSETSTEELKASNEELQAINEELRSATEELETS------- | *Burkholderia ubonensis* | Bacteria; Proteobacteria; Betaproteobacteria; Burkholderiales; Burkholderiaceae; Burkholderia; Burkholderia cepacia complex | PAS domain S-box protein | 6 |
|  | **WP_059768308.1** | 1.9E-06 | Qb.I | LQRTKEQLQSTIEQSETSTEELKASNEELQAINEELRSATEELETS------- | *Burkholderia ubonensis* | Bacteria; Proteobacteria; Betaproteobacteria; Burkholderiales; Burkholderiaceae; Burkholderia; Burkholderia cepacia complex | PAS domain S-box protein | 6 |
|  | **WP_059778110.1** | 1.9E-06 | Qb.I | LQRTKEQLQSTIEQSETSTEELKASNEELQAINEELRSATEELETS------- | *Burkholderia ubonensis* | Bacteria; Proteobacteria; Betaproteobacteria; Burkholderiales; Burkholderiaceae; Burkholderia; Burkholderia cepacia complex | PAS domain S-box protein | 6 |
|  | **WP_059786609.1** | 1.9E-06 | Qb.I | LQRTKEQLQSTIEQSETSTEELKASNEELQAINEELRSATEELETS------- | *Burkholderia ubonensis* | Bacteria; Proteobacteria; Betaproteobacteria; Burkholderiales; Burkholderiaceae; Burkholderia; Burkholderia cepacia complex | PAS domain S-box protein | 6 |
|  | **WP_059795304.1** | 1.9E-06 | Qb.I | LQRTKEQLQSTIEQSETSTEELKASNEELQAINEELRSATEELETS------- | *Burkholderia ubonensis* | Bacteria; Proteobacteria; Betaproteobacteria; Burkholderiales; Burkholderiaceae; Burkholderia; Burkholderia cepacia complex | PAS domain S-box protein | 6 |
|  | **WP_059808592.1** | 1.9E-06 | Qb.I | LQRTKEQLQSTIEQSETSTEELKASNEELQAINEELRSATEELETS------- | *Burkholderia ubonensis* | Bacteria; Proteobacteria; Betaproteobacteria; Burkholderiales; Burkholderiaceae; Burkholderia; Burkholderia cepacia complex | PAS domain S-box protein | 6 |
|  | **WP_059812663.1** | 1.9E-06 | Qb.I | LQRTKEQLQSTIEQSETSTEELKASNEELQAINEELRSATEELETS------- | *Burkholderia ubonensis* | Bacteria; Proteobacteria; Betaproteobacteria; Burkholderiales; Burkholderiaceae; Burkholderia; Burkholderia cepacia complex | PAS domain S-box protein | 6 |
|  | **WP_059821241.1** | 1.9E-06 | Qb.I | LQRTKEQLQSTIEQSETSTEELKASNEELQAINEELRSATEELETS------- | *Burkholderia ubonensis* | Bacteria; Proteobacteria; Betaproteobacteria; Burkholderiales; Burkholderiaceae; Burkholderia; Burkholderia cepacia complex | PAS domain S-box protein | 6 |
|  | **WP_059825327.1** | 1.9E-06 | Qb.I | LQRTKEQLQSTIEQSETSTEELKASNEELQAINEELRSATEELETS------- | *Burkholderia ubonensis* | Bacteria; Proteobacteria; Betaproteobacteria; Burkholderiales; Burkholderiaceae; Burkholderia; Burkholderia cepacia complex | PAS domain S-box protein | 6 |
|  | **WP_059838397.1** | 1.9E-06 | Qb.I | LQRTKEQLQSTIEQSETSTEELKASNEELQAINEELRSATEELETS------- | *Burkholderia ubonensis* | Bacteria; Proteobacteria; Betaproteobacteria; Burkholderiales; Burkholderiaceae; Burkholderia; Burkholderia cepacia complex | PAS domain S-box protein | 6 |
|  | **WP_059849367.1** | 1.9E-06 | Qb.I | LQRTKEQLQSTIEQSETSTEELKASNEELQAINEELRSATEELETS------- | *Burkholderia ubonensis* | Bacteria; Proteobacteria; Betaproteobacteria; Burkholderiales; Burkholderiaceae; Burkholderia; Burkholderia cepacia complex | PAS domain S-box protein | 6 |
|  | **WP_059854907.1** | 1.9E-06 | Qb.I | LQRTKEQLQSTIEQSETSTEELKASNEELQAINEELRSATEELETS------- | *Burkholderia ubonensis* | Bacteria; Proteobacteria; Betaproteobacteria; Burkholderiales; Burkholderiaceae; Burkholderia; Burkholderia cepacia complex | PAS domain S-box protein | 6 |
|  | **WP_059863178.1** | 1.9E-06 | Qb.I | LQRTKEQLQSTIEQSETSTEELKASNEELQAINEELRSATEELETS------- | *Burkholderia ubonensis* | Bacteria; Proteobacteria; Betaproteobacteria; Burkholderiales; Burkholderiaceae; Burkholderia; Burkholderia cepacia complex | PAS domain S-box protein | 6 |
|  | **WP_059866618.1** | 1.9E-06 | Qb.I | LQRTKEQLQSTIEQSETSTEELKASNEELQAINEELRSATEELETS------- | *Burkholderia ubonensis* | Bacteria; Proteobacteria; Betaproteobacteria; Burkholderiales; Burkholderiaceae; Burkholderia; Burkholderia cepacia complex | PAS domain S-box protein | 6 |
|  | **WP_059866787.1** | 1.9E-06 | Qb.I | LQRTKEQLQSTIEQSETSTEELKASNEELQAINEELRSATEELETS------- | *Burkholderia ubonensis* | Bacteria; Proteobacteria; Betaproteobacteria; Burkholderiales; Burkholderiaceae; Burkholderia; Burkholderia cepacia complex | PAS domain S-box protein | 6 |
|  | **WP_059872123.1** | 1.9E-06 | Qb.I | LQRTKEQLQSTIEQSETSTEELKASNEELQAINEELRSATEELETS------- | *Burkholderia ubonensis* | Bacteria; Proteobacteria; Betaproteobacteria; Burkholderiales; Burkholderiaceae; Burkholderia; Burkholderia cepacia complex | PAS domain S-box protein | 6 |
|  | **WP_059877111.1** | 1.9E-06 | Qb.I | LQRTKEQLQSTIEQSETSTEELKASNEELQAINEELRSATEELETS------- | *Burkholderia ubonensis* | Bacteria; Proteobacteria; Betaproteobacteria; Burkholderiales; Burkholderiaceae; Burkholderia; Burkholderia cepacia complex | PAS domain S-box protein | 6 |
|  | **WP_059880460.1** | 1.9E-06 | Qb.I | LQRTKEQLQSTIEQSETSTEELKASNEELQAINEELRSATEELETS------- | *Burkholderia stagnalis* | Bacteria; Proteobacteria; Betaproteobacteria; Burkholderiales; Burkholderiaceae; Burkholderia; Burkholderia cepacia complex | PAS domain S-box protein | 6 |
|  | **WP_059881183.1** | 1.9E-06 | Qb.I | LQRTKEQLQSTIEQSETSTEELKASNEELQAINEELRSATEELETS------- | *Burkholderia ubonensis* | Bacteria; Proteobacteria; Betaproteobacteria; Burkholderiales; Burkholderiaceae; Burkholderia; Burkholderia cepacia complex | PAS domain S-box protein | 6 |
|  | **WP_059886863.1** | 1.9E-06 | Qb.I | LQRTKEQLQSTIEQSETSTEELKASNEELQAINEELRSATEELETS------- | *Burkholderia ubonensis* | Bacteria; Proteobacteria; Betaproteobacteria; Burkholderiales; Burkholderiaceae; Burkholderia; Burkholderia cepacia complex | PAS domain S-box protein | 6 |
|  | **WP_059892770.1** | 1.9E-06 | Qb.I | LQRTKEQLQSTIEQSETSTEELKASNEELQAINEELRSATEELETS------- | *Burkholderia ubonensis* | Bacteria; Proteobacteria; Betaproteobacteria; Burkholderiales; Burkholderiaceae; Burkholderia; Burkholderia cepacia complex | PAS domain S-box protein | 6 |
|  | **WP_059899316.1** | 1.9E-06 | Qb.I | LQRTKEQLQSTIEQSETSTEELKASNEELQAINEELRSATEELETS------- | *Burkholderia ubonensis* | Bacteria; Proteobacteria; Betaproteobacteria; Burkholderiales; Burkholderiaceae; Burkholderia; Burkholderia cepacia complex | PAS domain S-box protein | 6 |
|  | **WP_059908218.1** | 1.9E-06 | Qb.I | LQRTKEQLQSTIEQSETSTEELKASNEELQAINEELRSATEELETS------- | *Burkholderia stagnalis* | Bacteria; Proteobacteria; Betaproteobacteria; Burkholderiales; Burkholderiaceae; Burkholderia; Burkholderia cepacia complex | PAS domain S-box protein | 6 |
|  | **WP_059911649.1** | 1.9E-06 | Qb.I | LQRTKEQLQSTIEQSETSTEELKASNEELQAINEELRSATEELETS------- | *Burkholderia ubonensis* | Bacteria; Proteobacteria; Betaproteobacteria; Burkholderiales; Burkholderiaceae; Burkholderia; Burkholderia cepacia complex | PAS domain S-box protein | 6 |
|  | **WP_059912179.1** | 1.9E-06 | Qb.I | LQRTKEQLQSTIEQSETSTEELKASNEELQAINEELRSATEELETS------- | *Burkholderia ubonensis* | Bacteria; Proteobacteria; Betaproteobacteria; Burkholderiales; Burkholderiaceae; Burkholderia; Burkholderia cepacia complex | PAS domain S-box protein | 6 |
|  | **WP_059919269.1** | 1.9E-06 | Qb.I | LQRTKEQLQSTIEQSETSTEELKASNEELQAINEELRSATEELETS------- | *Burkholderia ubonensis* | Bacteria; Proteobacteria; Betaproteobacteria; Burkholderiales; Burkholderiaceae; Burkholderia; Burkholderia cepacia complex | PAS domain S-box protein | 6 |
|  | **WP_059921814.1** | 1.9E-06 | Qb.I | LQRTKEQLQSTIEQSETSTEELKASNEELQAINEELRSATEELETS------- | *Burkholderia stagnalis* | Bacteria; Proteobacteria; Betaproteobacteria; Burkholderiales; Burkholderiaceae; Burkholderia; Burkholderia cepacia complex | PAS domain S-box protein | 6 |
|  | **WP_059922115.1** | 1.9E-06 | Qb.I | LQRTKEQLQSTIEQSETSTEELKASNEELQAINEELRSATEELETS------- | *Burkholderia ubonensis* | Bacteria; Proteobacteria; Betaproteobacteria; Burkholderiales; Burkholderiaceae; Burkholderia; Burkholderia cepacia complex | PAS domain S-box protein | 6 |
|  | **WP_059925157.1** | 1.9E-06 | Qb.I | LQRTKEQLQSTIEQSETSTEELKASNEELQAINEELRSATEELETS------- | *Burkholderia ubonensis* | Bacteria; Proteobacteria; Betaproteobacteria; Burkholderiales; Burkholderiaceae; Burkholderia; Burkholderia cepacia complex | PAS domain S-box protein | 6 |
|  | **WP_059936452.1** | 1.9E-06 | Qb.I | LQRTKEQLQSTIEQSETSTEELKASNEELQAINEELRSATEELETS------- | *Burkholderia ubonensis* | Bacteria; Proteobacteria; Betaproteobacteria; Burkholderiales; Burkholderiaceae; Burkholderia; Burkholderia cepacia complex | PAS domain S-box protein | 6 |
|  | **WP_059940018.1** | 1.9E-06 | Qb.I | LQRTKEQLQSTIEQSETSTEELKASNEELQAINEELRSATEELETS------- | *Burkholderia ubonensis* | Bacteria; Proteobacteria; Betaproteobacteria; Burkholderiales; Burkholderiaceae; Burkholderia; Burkholderia cepacia complex | PAS domain S-box protein | 6 |
|  | **WP_059946195.1** | 1.9E-06 | Qb.I | LQRTKEQLQSTIEQSETSTEELKASNEELQAINEELRSATEELETS------- | *Burkholderia ubonensis* | Bacteria; Proteobacteria; Betaproteobacteria; Burkholderiales; Burkholderiaceae; Burkholderia; Burkholderia cepacia complex | PAS domain S-box protein | 6 |
|  | **WP_059950405.1** | 1.9E-06 | Qb.I | LQRTKEQLQSTIEQSETSTEELKASNEELQAINEELRSATEELETS------- | *Burkholderia ubonensis* | Bacteria; Proteobacteria; Betaproteobacteria; Burkholderiales; Burkholderiaceae; Burkholderia; Burkholderia cepacia complex | PAS domain S-box protein | 6 |
|  | **WP_059954482.1** | 1.9E-06 | Qb.I | LQRTKEQLQSTIEQSETSTEELKASNEELQAINEELRSATEELETS------- | *Burkholderia ubonensis* | Bacteria; Proteobacteria; Betaproteobacteria; Burkholderiales; Burkholderiaceae; Burkholderia; Burkholderia cepacia complex | PAS domain S-box protein | 6 |
|  | **WP_059957269.1** | 1.9E-06 | Qb.I | LQRTKEQLQSTIEQSETSTEELKASNEELQAINEELRSATEELETS------- | *Burkholderia ubonensis* | Bacteria; Proteobacteria; Betaproteobacteria; Burkholderiales; Burkholderiaceae; Burkholderia; Burkholderia cepacia complex | PAS domain S-box protein | 6 |
|  | **WP_059959369.1** | 1.9E-06 | Qb.I | LQRTKEQLQSTIEQSETSTEELKASNEELQAINEELRSATEELETS------- | *Burkholderia stagnalis* | Bacteria; Proteobacteria; Betaproteobacteria; Burkholderiales; Burkholderiaceae; Burkholderia; Burkholderia cepacia complex | PAS domain S-box protein | 6 |
|  | **WP_059965063.1** | 1.9E-06 | Qb.I | LQRTKEQLQSTIEQSETSTEELKASNEELQAINEELRSATEELETS------- | *Burkholderia ubonensis* | Bacteria; Proteobacteria; Betaproteobacteria; Burkholderiales; Burkholderiaceae; Burkholderia; Burkholderia cepacia complex | PAS domain S-box protein | 6 |
|  | **WP_059969660.1** | 1.9E-06 | Qb.I | LQRTKEQLQSTIEQSETSTEELKASNEELQAINEELRSATEELETS------- | *Burkholderia ubonensis* | Bacteria; Proteobacteria; Betaproteobacteria; Burkholderiales; Burkholderiaceae; Burkholderia; Burkholderia cepacia complex | PAS domain S-box protein | 6 |
|  | **WP_059977325.1** | 1.9E-06 | Qb.I | LQRTKEQLQSTIEQSETSTEELKASNEELQAINEELRSATEELETS------- | *Burkholderia ubonensis* | Bacteria; Proteobacteria; Betaproteobacteria; Burkholderiales; Burkholderiaceae; Burkholderia; Burkholderia cepacia complex | PAS domain S-box protein | 6 |
|  | **WP_059979536.1** | 1.9E-06 | Qb.I | LQRTKEQLQSTIEQSETSTEELKASNEELQAINEELRSATEELETS------- | *Burkholderia ubonensis* | Bacteria; Proteobacteria; Betaproteobacteria; Burkholderiales; Burkholderiaceae; Burkholderia; Burkholderia cepacia complex | PAS domain S-box protein | 6 |
|  | **WP_059982296.1** | 1.9E-06 | Qb.I | LQRTKEQLQSTIEQSETSTEELKASNEELQAINEELRSATEELETS------- | *Burkholderia ubonensis* | Bacteria; Proteobacteria; Betaproteobacteria; Burkholderiales; Burkholderiaceae; Burkholderia; Burkholderia cepacia complex | PAS domain S-box protein | 6 |
|  | **WP_059988473.1** | 1.9E-06 | Qb.I | LQRTKEQLQSTIEQSETSTEELKASNEELQAINEELRSATEELETS------- | *Burkholderia stagnalis* | Bacteria; Proteobacteria; Betaproteobacteria; Burkholderiales; Burkholderiaceae; Burkholderia; Burkholderia cepacia complex | PAS domain S-box protein | 6 |
|  | **WP_059989740.1** | 1.9E-06 | Qb.I | LQRTKEQLQSTIEQSETSTEELKASNEELQAINEELRSATEELETS------- | *Burkholderia ubonensis* | Bacteria; Proteobacteria; Betaproteobacteria; Burkholderiales; Burkholderiaceae; Burkholderia; Burkholderia cepacia complex | PAS domain S-box protein | 6 |
|  | **WP_059995465.1** | 1.9E-06 | Qb.I | LQRTKEQLQSTIEQSETSTEELKASNEELQAINEELRSATEELETS------- | *Burkholderia ubonensis* | Bacteria; Proteobacteria; Betaproteobacteria; Burkholderiales; Burkholderiaceae; Burkholderia; Burkholderia cepacia complex | PAS domain S-box protein | 6 |
|  | **WP_059996946.1** | 1.9E-06 | Qb.I | LQRTKEQLQSTIEQSETSTEELKASNEELQAINEELRSATEELETS------- | *Burkholderia stagnalis* | Bacteria; Proteobacteria; Betaproteobacteria; Burkholderiales; Burkholderiaceae; Burkholderia; Burkholderia cepacia complex | PAS domain S-box protein | 6 |
|  | **WP_060000282.1** | 1.9E-06 | Qb.I | LQRTKEQLQSTIEQSETSTEELKASNEELQAINEELRSATEELETS------- | *Burkholderia ubonensis* | Bacteria; Proteobacteria; Betaproteobacteria; Burkholderiales; Burkholderiaceae; Burkholderia; Burkholderia cepacia complex | PAS domain S-box protein | 6 |

| | | | | | | | | |
|---|---|---|---|---|---|---|---|---|
|  | **WP_060006724.1** | 1.9E-06 | Qb.I | LQRTKEQLQSTIEQSETSTEELKASNEELQAINEELRSATEELETS------- | *Burkholderia ubonensis* | Bacteria; Proteobacteria; Betaproteobacteria; Burkholderiales; Burkholderiaceae; Burkholderia; Burkholderia cepacia complex | PAS domain S-box protein | 6 |
|  | **WP_060010610.1** | 1.9E-06 | Qb.I | LQRTKEQLQSTIEQSETSTEELKASNEELQAINEELRSATEELETS------- | *Burkholderia ubonensis* | Bacteria; Proteobacteria; Betaproteobacteria; Burkholderiales; Burkholderiaceae; Burkholderia; Burkholderia cepacia complex | PAS domain S-box protein | 6 |
|  | **WP_060014126.1** | 1.9E-06 | Qb.I | LQRTKEQLQSTIEQSETSTEELKASNEELQAINEELRSATEELETS------- | *Burkholderia ubonensis* | Bacteria; Proteobacteria; Betaproteobacteria; Burkholderiales; Burkholderiaceae; Burkholderia; Burkholderia cepacia complex | PAS domain S-box protein | 6 |
|  | **WP_060026404.1** | 1.9E-06 | Qb.I | LQRTKEQLQSTIEQSETSTEELKASNEELQAINEELRSATEELETS------- | *Burkholderia ubonensis* | Bacteria; Proteobacteria; Betaproteobacteria; Burkholderiales; Burkholderiaceae; Burkholderia; Burkholderia cepacia complex | PAS domain S-box protein | 6 |
|  | **WP_060027819.1** | 1.9E-06 | Qb.I | LQRTKEQLQSTIEQSETSTEELKASNEELQAINEELRSATEELETS------- | *Burkholderia stagnalis* | Bacteria; Proteobacteria; Betaproteobacteria; Burkholderiales; Burkholderiaceae; Burkholderia; Burkholderia cepacia complex | PAS domain S-box protein | 6 |
|  | **WP_060029755.1** | 1.9E-06 | Qb.I | LQRTKEQLQSTIEQSETSTEELKASNEELQAINEELRSATEELETS------- | *Burkholderia ubonensis* | Bacteria; Proteobacteria; Betaproteobacteria; Burkholderiales; Burkholderiaceae; Burkholderia; Burkholderia cepacia complex | PAS domain S-box protein | 6 |
|  | **WP_060032598.1** | 1.9E-06 | Qb.I | LQRTKEQLQSTIEQSETSTEELKASNEELQAINEELRSATEELETS------- | *Burkholderia ubonensis* | Bacteria; Proteobacteria; Betaproteobacteria; Burkholderiales; Burkholderiaceae; Burkholderia; Burkholderia cepacia complex | PAS domain S-box protein | 6 |
|  | **WP_060037111.1** | 1.9E-06 | Qb.I | LQRTKEQLQSTIEQSETSTEELKASNEELQAINEELRSATEELETS------- | *Burkholderia ubonensis* | Bacteria; Proteobacteria; Betaproteobacteria; Burkholderiales; Burkholderiaceae; Burkholderia; Burkholderia cepacia complex | PAS domain S-box protein | 6 |
|  | **WP_060053730.1** | 1.9E-06 | Qb.I | LQRTKEQLQSTIEQSETSTEELKASNEELQAINEELRSATEELETS------- | *Burkholderia ubonensis* | Bacteria; Proteobacteria; Betaproteobacteria; Burkholderiales; Burkholderiaceae; Burkholderia; Burkholderia cepacia complex | PAS domain S-box protein | 6 |
|  | **WP_060058866.1** | 1.9E-06 | Qb.I | LQRTKEQLQSTIEQSETSTEELKASNEELQAINEELRSATEELETS------- | *Burkholderia ubonensis* | Bacteria; Proteobacteria; Betaproteobacteria; Burkholderiales; Burkholderiaceae; Burkholderia; Burkholderia cepacia complex | PAS domain S-box protein | 6 |
|  | **WP_060062001.1** | 1.9E-06 | Qb.I | LQRTKEQLQSTIEQSETSTEELKASNEELQAINEELRSATEELETS------- | *Burkholderia ubonensis* | Bacteria; Proteobacteria; Betaproteobacteria; Burkholderiales; Burkholderiaceae; Burkholderia; Burkholderia cepacia complex | PAS domain S-box protein | 6 |
|  | **WP_060073907.1** | 1.9E-06 | Qb.I | LQRTKEQLQSTIEQSETSTEELKASNEELQAINEELRSATEELETS------- | *Burkholderia ubonensis* | Bacteria; Proteobacteria; Betaproteobacteria; Burkholderiales; Burkholderiaceae; Burkholderia; Burkholderia cepacia complex | PAS domain S-box protein | 6 |
|  | **WP_060076229.1** | 1.9E-06 | Qb.I | LQRTKEQLQSTIEQSETSTEELKASNEELQAINEELRSATEELETS------- | *Burkholderia ubonensis* | Bacteria; Proteobacteria; Betaproteobacteria; Burkholderiales; Burkholderiaceae; Burkholderia; Burkholderia cepacia complex | PAS domain S-box protein | 6 |
|  | **WP_060077236.1** | 1.9E-06 | Qb.I | LQRTKEQLQSTIEQSETSTEELKASNEELQAINEELRSATEELETS------- | *Burkholderia ubonensis* | Bacteria; Proteobacteria; Betaproteobacteria; Burkholderiales; Burkholderiaceae; Burkholderia; Burkholderia cepacia complex | PAS domain S-box protein | 6 |
|  | **WP_060080274.1** | 1.9E-06 | Qb.I | LQRTKEQLQSTIEQSETSTEELKASNEELQAINEELRSATEELETS------- | *Burkholderia ubonensis* | Bacteria; Proteobacteria; Betaproteobacteria; Burkholderiales; Burkholderiaceae; Burkholderia; Burkholderia cepacia complex | PAS domain S-box protein | 6 |
|  | **WP_060088018.1** | 1.9E-06 | Qb.I | LQRTKEQLQSTIEQSETSTEELKASNEELQAINEELRSATEELETS------- | *Burkholderia ubonensis* | Bacteria; Proteobacteria; Betaproteobacteria; Burkholderiales; Burkholderiaceae; Burkholderia; Burkholderia cepacia complex | PAS domain S-box protein | 6 |
|  | **WP_060090225.1** | 1.9E-06 | Qb.I | LQRTKEQLQSTIEQSETSTEELKASNEELQAINEELRSATEELETS------- | *Burkholderia ubonensis* | Bacteria; Proteobacteria; Betaproteobacteria; Burkholderiales; Burkholderiaceae; Burkholderia; Burkholderia cepacia complex | PAS domain S-box protein | 6 |
|  | **WP_060104332.1** | 1.9E-06 | Qb.I | LQRTKEQLQSTIEQSETSTEELKASNEELQAINEELRSATEELETS------- | *Burkholderia ubonensis* | Bacteria; Proteobacteria; Betaproteobacteria; Burkholderiales; Burkholderiaceae; Burkholderia; Burkholderia cepacia complex | PAS domain S-box protein | 6 |
|  | **WP_060112192.1** | 1.9E-06 | Qb.I | LQRTKEQLQSTIEQSETSTEELKASNEELQAINEELRSATEELETS------- | *Burkholderia ubonensis* | Bacteria; Proteobacteria; Betaproteobacteria; Burkholderiales; Burkholderiaceae; Burkholderia; Burkholderia cepacia complex | PAS domain S-box protein | 6 |
|  | **WP_060119404.1** | 1.9E-06 | Qb.I | LQRTKEQLQSTIEQSETSTEELKASNEELQAINEELRSATEELETS------- | *Burkholderia ubonensis* | Bacteria; Proteobacteria; Betaproteobacteria; Burkholderiales; Burkholderiaceae; Burkholderia; Burkholderia cepacia complex | PAS domain S-box protein | 6 |
|  | **WP_060130795.1** | 1.9E-06 | Qb.I | LQRTKEQLQSTIEQSETSTEELKASNEELQAINEELRSATEELETS------- | *Burkholderia ubonensis* | Bacteria; Proteobacteria; Betaproteobacteria; Burkholderiales; Burkholderiaceae; Burkholderia; Burkholderia cepacia complex | PAS domain S-box protein | 6 |
|  | **WP_060134513.1** | 1.9E-06 | Qb.I | LQRTKEQLQSTIEQSETSTEELKASNEELQAINEELRSATEELETS------- | *Burkholderia ubonensis* | Bacteria; Proteobacteria; Betaproteobacteria; Burkholderiales; Burkholderiaceae; Burkholderia; Burkholderia cepacia complex | PAS domain S-box protein | 6 |
|  | **WP_060137345.1** | 1.9E-06 | Qb.I | LQRTKEQLQSTIEQSETSTEELKASNEELQAINEELRSATEELETS------- | *Burkholderia ubonensis* | Bacteria; Proteobacteria; Betaproteobacteria; Burkholderiales; Burkholderiaceae; Burkholderia; Burkholderia cepacia complex | PAS domain S-box protein | 6 |
|  | **WP_060149216.1** | 1.9E-06 | Qb.I | LQRTKEQLQSTIEQSETSTEELKASNEELQAINEELRSATEELETS------- | *Burkholderia stagnalis* | Bacteria; Proteobacteria; Betaproteobacteria; Burkholderiales; Burkholderiaceae; Burkholderia; Burkholderia cepacia complex | PAS domain S-box protein | 6 |
|  | **WP_060156126.1** | 1.9E-06 | Qb.I | LQRTKEQLQSTIEQSETSTEELKASNEELQAINEELRSATEELETS------- | *Burkholderia ubonensis* | Bacteria; Proteobacteria; Betaproteobacteria; Burkholderiales; Burkholderiaceae; Burkholderia; Burkholderia cepacia complex | PAS domain S-box protein | 6 |
|  | **WP_060160485.1** | 1.9E-06 | Qb.I | LQRTKEQLQSTIEQSETSTEELKASNEELQAINEELRSATEELETS------- | *Burkholderia ubonensis* | Bacteria; Proteobacteria; Betaproteobacteria; Burkholderiales; Burkholderiaceae; Burkholderia; Burkholderia cepacia complex | PAS domain S-box protein | 6 |
|  | **WP_060161075.1** | 1.9E-06 | Qb.I | LQRTKEQLQSTIEQSETSTEELKASNEELQAINEELRSATEELETS------- | *Burkholderia stagnalis* | Bacteria; Proteobacteria; Betaproteobacteria; Burkholderiales; Burkholderiaceae; Burkholderia; Burkholderia cepacia complex | PAS domain S-box protein | 6 |
|  | **WP_060167041.1** | 1.9E-06 | Qb.I | LQRTKEQLQSTIEQSETSTEELKASNEELQAINEELRSATEELETS------- | *Burkholderia ubonensis* | Bacteria; Proteobacteria; Betaproteobacteria; Burkholderiales; Burkholderiaceae; Burkholderia; Burkholderia cepacia complex | PAS domain S-box protein | 6 |
|  | **WP_060197740.1** | 1.9E-06 | Qb.I | LQRTKEQLQSTIEQSETSTEELKASNEELQAINEELRSATEELETS------- | *Burkholderia ubonensis* | Bacteria; Proteobacteria; Betaproteobacteria; Burkholderiales; Burkholderiaceae; Burkholderia; Burkholderia cepacia complex | PAS domain S-box protein | 6 |
|  | **WP_060216026.1** | 1.9E-06 | Qb.I | LQRTKEQLQSTIEQSETSTEELKASNEELQAINEELRSATEELETS------- | *Burkholderia ubonensis* | Bacteria; Proteobacteria; Betaproteobacteria; Burkholderiales; Burkholderiaceae; Burkholderia; Burkholderia cepacia complex | PAS domain S-box protein | 6 |
|  | **WP_060226705.1** | 1.9E-06 | Qb.I | LQRTKEQLQSTIEQSETSTEELKASNEELQAINEELRSATEELETS------- | *Burkholderia ubonensis* | Bacteria; Proteobacteria; Betaproteobacteria; Burkholderiales; Burkholderiaceae; Burkholderia; Burkholderia cepacia complex | PAS domain S-box protein | 6 |
|  | **WP_060233220.1** | 1.9E-06 | Qb.I | LQRTKEQLQSTIEQSETSTEELKASNEELQAINEELRSATEELETS------- | *Burkholderia ubonensis* | Bacteria; Proteobacteria; Betaproteobacteria; Burkholderiales; Burkholderiaceae; Burkholderia; Burkholderia cepacia complex | PAS domain S-box protein | 6 |
|  | **WP_060244873.1** | 1.9E-06 | Qb.I | LQRTKEQLQSTIEQSETSTEELKASNEELQAINEELRSATEELETS------- | *Burkholderia ubonensis* | Bacteria; Proteobacteria; Betaproteobacteria; Burkholderiales; Burkholderiaceae; Burkholderia; Burkholderia cepacia complex | PAS domain S-box protein | 6 |
|  | **WP_060250033.1** | 1.9E-06 | Qb.I | LQRTKEQLQSTIEQSETSTEELKASNEELQAINEELRSATEELETS------- | *Burkholderia ubonensis* | Bacteria; Proteobacteria; Betaproteobacteria; Burkholderiales; Burkholderiaceae; Burkholderia; Burkholderia cepacia complex | PAS domain S-box protein | 6 |
|  | **WP_060257874.1** | 1.9E-06 | Qb.I | LQRTKEQLQSTIEQSETSTEELKASNEELQAINEELRSATEELETS------- | *Burkholderia ubonensis* | Bacteria; Proteobacteria; Betaproteobacteria; Burkholderiales; Burkholderiaceae; Burkholderia; Burkholderia cepacia complex | PAS domain S-box protein | 6 |
|  | **WP_060268749.1** | 1.9E-06 | Qb.I | LQRTKEQLQSTIEQSETSTEELKASNEELQAINEELRSATEELETS------- | *Burkholderia ubonensis* | Bacteria; Proteobacteria; Betaproteobacteria; Burkholderiales; Burkholderiaceae; Burkholderia; Burkholderia cepacia complex | PAS domain S-box protein | 6 |
|  | **WP_060281883.1** | 1.9E-06 | Qb.I | LQRTKEQLQSTIEQSETSTEELKASNEELQAINEELRSATEELETS------- | *Burkholderia ubonensis* | Bacteria; Proteobacteria; Betaproteobacteria; Burkholderiales; Burkholderiaceae; Burkholderia; Burkholderia cepacia complex | PAS domain S-box protein | 6 |
|  | **WP_060289884.1** | 1.9E-06 | Qb.I | LQRTKEQLQSTIEQSETSTEELKASNEELQAINEELRSATEELETS------- | *Burkholderia ubonensis* | Bacteria; Proteobacteria; Betaproteobacteria; Burkholderiales; Burkholderiaceae; Burkholderia; Burkholderia cepacia complex | PAS domain S-box protein | 6 |
|  | **WP_060291607.1** | 1.9E-06 | Qb.I | LQRTKEQLQSTIEQSETSTEELKASNEELQAINEELRSATEELETS------- | *Burkholderia stagnalis* | Bacteria; Proteobacteria; Betaproteobacteria; Burkholderiales; Burkholderiaceae; Burkholderia; Burkholderia cepacia complex | PAS domain S-box protein | 6 |
|  | **WP_060308360.1** | 1.9E-06 | Qb.I | LQRTKEQLQSTIEQSETSTEELKASNEELQAINEELRSATEELETS------- | *Burkholderia ubonensis* | Bacteria; Proteobacteria; Betaproteobacteria; Burkholderiales; Burkholderiaceae; Burkholderia; Burkholderia cepacia complex | PAS domain S-box protein | 6 |
|  | **WP_060337273.1** | 1.9E-06 | Qb.I | LQRTKEQLQSTIEQSETSTEELKASNEELQAINEELRSATEELETS------- | *Burkholderia ubonensis* | Bacteria; Proteobacteria; Betaproteobacteria; Burkholderiales; Burkholderiaceae; Burkholderia; Burkholderia cepacia complex | PAS domain S-box protein | 6 |
|  | **WP_060351271.1** | 1.9E-06 | Qb.I | LQRTKEQLQSTIEQSETSTEELKASNEELQAINEELRSATEELETS------- | *Burkholderia stagnalis* | Bacteria; Proteobacteria; Betaproteobacteria; Burkholderiales; Burkholderiaceae; Burkholderia; Burkholderia cepacia complex | PAS domain S-box protein | 6 |
|  | **WP_060371492.1** | 1.9E-06 | Qb.I | LQRTKEQLQSTIEQSETSTEELKASNEELQAINEELRSATEELETS------- | *Burkholderia ubonensis* | Bacteria; Proteobacteria; Betaproteobacteria; Burkholderiales; Burkholderiaceae; Burkholderia; Burkholderia cepacia complex | PAS domain S-box protein | 6 |
|  | **WP_060379808.1** | 1.9E-06 | Qb.I | LQRTKEQLQSTIEQSETSTEELKASNEELQAINEELRSATEELETS------- | *Burkholderia ubonensis* | Bacteria; Proteobacteria; Betaproteobacteria; Burkholderiales; Burkholderiaceae; Burkholderia; Burkholderia cepacia complex | PAS domain S-box protein | 6 |
|  | **WP_060401395.1** | 1.9E-06 | Qb.I | LQRTKEQLQSTIEQSETSTEELKASNEELQAINEELRSATEELETS------- | *Burkholderia ubonensis* | Bacteria; Proteobacteria; Betaproteobacteria; Burkholderiales; Burkholderiaceae; Burkholderia; Burkholderia cepacia complex | PAS domain S-box protein | 6 |
|  | **WP_060449538.1** | 1.9E-06 | Qb.I | LQRTKEQLQSTIEQSETSTEELKASNEELQAINEELRSATEELETS------- | *Burkholderia ubonensis* | Bacteria; Proteobacteria; Betaproteobacteria; Burkholderiales; Burkholderiaceae; Burkholderia; Burkholderia cepacia complex | PAS domain S-box protein | 6 |
|  | **WP_062918793.1** | 1.9E-06 | Qb.I | LQRTKEQLQSTIEQSETSTEELKASNEELQAINEELRSATEELETS------- | *Paraburkholderia caribensis* | Bacteria; Proteobacteria; Betaproteobacteria; Burkholderiales; Burkholderiaceae; Paraburkholderia | PAS domain S-box protein | 6 |
|  | **WP_065065780.1** | 1.9E-06 | Qb.I | LQRTKEQLQSTIEQSETSTEELKASNEELQAINEELRSATEELETS------- | *Paraburkholderia tropica* | Bacteria; Proteobacteria; Betaproteobacteria; Burkholderiales; Burkholderiaceae; Paraburkholderia | PAS domain S-box protein | 6 |
| | **WP_065416142.1** | 1.9E-06 | Qb.I | ---INKLVSDTVKQSHDSAEYINESSYLISGTADTFDNIFNKINSANDLIQE- | *Clostridium beijerinckii* | Bacteria; Firmicutes; Clostridia; Clostridiales; Clostridiaceae; Clostridium | methyl-accepting chemotaxis protein | 6 |
|  | **WP_069237679.1** | 1.9E-06 | Qb.I | LQRTKEQLQSTIEQSETSTEELKASNEELQAINEELRSATEELETS------- | *Burkholderia ubonensis* | Bacteria; Proteobacteria; Betaproteobacteria; Burkholderiales; Burkholderiaceae; Burkholderia; Burkholderia cepacia complex | PAS domain S-box protein | 6 |
|  | **WP_069262287.1** | 1.9E-06 | Qb.I | LQRTKEQLQSTIEQSETSTEELKASNEELQAINEELRSATEELETS------- | *Paraburkholderia nodosa* | Bacteria; Proteobacteria; Betaproteobacteria; Burkholderiales; Burkholderiaceae; Paraburkholderia | PAS domain S-box protein | 6 |
|  | **WP_069271361.1** | 1.9E-06 | Qb.I | LQRTKEQLQSTIEQSETSTEELKASNEELQAINEELRSATEELETS------- | *Burkholderia cepacia complex* | Bacteria; Proteobacteria; Betaproteobacteria; Burkholderiales; Burkholderiaceae; Burkholderia | MULTISPECIES: PAS domain S-box protein | 6 |
|  | **WP_069307526.1** | 1.9E-06 | Qa.III | IEAISNVIQELNGIATTIAGAVREQGSATQEIARNVQEAS------------ | *Methylobrevis pamukkalensis* | Bacteria; Proteobacteria; Alphaproteobacteria; Rhizobiales; Methylobrevis | HAMP domain-containing protein | 6 |
|  | **WP_071755127.1** | 1.9E-06 | Qb.I | LQRTKEQLQSTIEQSETSTEELKASNEELQAINEELRSATEELETS------- | *Burkholderia ubonensis* | Bacteria; Proteobacteria; Betaproteobacteria; Burkholderiales; Burkholderiaceae; Burkholderia; Burkholderia cepacia complex | PAS domain S-box protein | 6 |
|  | **WP_071774243.1** | 1.9E-06 | Qb.I | LQRTKEQLQSTIEQSETSTEELKASNEELQAINEELRSATEELETS------- | *Burkholderia ubonensis* | Bacteria; Proteobacteria; Betaproteobacteria; Burkholderiales; Burkholderiaceae; Burkholderia; Burkholderia cepacia complex | PAS domain S-box protein | 6 |
|  | **WP_071776636.1** | 1.9E-06 | Qb.I | LQRTKEQLQSTIEQSETSTEELKASNEELQAINEELRSATEELETS------- | *Burkholderia ubonensis* | Bacteria; Proteobacteria; Betaproteobacteria; Burkholderiales; Burkholderiaceae; Burkholderia; Burkholderia cepacia complex | PAS domain S-box protein | 6 |
|  | **WP_074987333.1** | 1.9E-06 | Qb.I | LQRTKEQLQSTIEQSETSTEELKASNEELQAINEELRSATEELETS------- | *Paraburkholderia tropica* | Bacteria; Proteobacteria; Betaproteobacteria; Burkholderiales; Burkholderiaceae; Paraburkholderia | PAS domain S-box protein | 6 |
|  | **WP_076474843.1** | 1.9E-06 | Qb.I | LQRTKEQLQSTIEQSETSTEELKASNEELQAINEELRSATEELETS------- | *Burkholderia ubonensis* | Bacteria; Proteobacteria; Betaproteobacteria; Burkholderiales; Burkholderiaceae; Burkholderia; Burkholderia cepacia complex | PAS domain S-box protein | 6 |
|  | **WP_076834912.1** | 1.9E-06 | Qa.IV | -----STIAEIDQVSGEIAAAVDQQGAATREISSNLQQAADRTRDVNK----- | *Bradyrhizobium sp. UFLA 03-321* | Bacteria; Proteobacteria; Alphaproteobacteria; Rhizobiales; Bradyrhizobiaceae; Bradyrhizobium | chemotaxis protein | 6 |
|  | **WP_076864537.1** | 1.9E-06 | Qa.IV | -----TTIAEIDQVSGEIAAAVDQQGAATREISSNLQQAADRTRDVNK----- | *Bradyrhizobium mercantei* | Bacteria; Proteobacteria; Alphaproteobacteria; Rhizobiales; Bradyrhizobiaceae; Bradyrhizobium | chemotaxis protein | 6 |
| | **WP_077856140.1** | 1.9E-06 | Qb.I | ---INKLVSDTVKQSHDSAEYINESSYLISGTADTFDNIFNKINSANDLIQE- | *Clostridium beijerinckii* | Bacteria; Firmicutes; Clostridia; Clostridiales; Clostridiaceae; Clostridium | methyl-accepting chemotaxis protein | 6 |
|  | **WP_083455897.1** | 1.9E-06 | Qa.III | IRMISAVIVEVNEIVTGIAAAVEEQSTATQEIASNVSQASQGIQE-------- | *Desulfosarcina cetonica* | Bacteria; Proteobacteria; Deltaproteobacteria; Desulfobacterales; Desulfobacteraceae; Desulfosarcina | HAMP domain-containing protein | 6 |

| | | | | | | | | |
|---|---|---|---|---|---|---|---|---|
|  | WP_089429316.1 | 1.9E-06 | Qb.I | LQRTKEQLQSTIEQSETSTEELKASNEELQAINEELRSATEELETS------- | *Burkholderia sp. HI2500* | Bacteria; Proteobacteria; Betaproteobacteria; Burkholderiales; Burkholderiaceae; Burkholderia | response regulator | 6 |
|  | WP_089463701.1 | 1.9E-06 | Qb.I | LQRTKEQLQSTIEQSETSTEELKASNEELQAINEELRSATEELETS------- | *Burkholderia* | Bacteria; Proteobacteria; Betaproteobacteria; Burkholderiales; Burkholderiaceae | MULTISPECIES: PAS domain S-box protein | 6 |
|  | WP_090865395.1 | 1.9E-06 | Qb.I | LQRTKEQLQSTIEQSETSTEELKASNEELQAINEELRSATEELETS------- | *Paraburkholderia diazotrophica* | Bacteria; Proteobacteria; Betaproteobacteria; Burkholderiales; Burkholderiaceae; Paraburkholderia | PAS domain S-box protein | 6 |
|  | WP_092006295.1 | 1.9E-06 | Qb.I | LQRTKEQLQSTIEQSETSTEELKASNEELQAINEELRSATEELETS------- | *Burkholderia sp. TNe-862* | Bacteria; Proteobacteria; Betaproteobacteria; Burkholderiales; Burkholderiaceae; Burkholderia | PAS domain S-box protein | 6 |
| | WP_092683361.1 | 1.9E-06 | Qc.III | IKEIGDTIGRLSEISSIIASAVEQQDAVTQEITHNVQDASNGTRQVTSHIGD- | *Rhodopseudomonas pseudopalustris* | Bacteria; Proteobacteria; Alphaproteobacteria; Rhizobiales; Bradyrhizobiaceae; Rhodopseudomonas | PAS domain S-box protein | 6 |
|  | WP_095413035.1 | 1.9E-06 | Qb.I | LQRTKEQLQSTIEQSETSTEELKASNEELQAINEELRSATEELETS------- | *Burkholderia ubonensis* | Bacteria; Proteobacteria; Betaproteobacteria; Burkholderiales; Burkholderiaceae; Burkholderia; Burkholderia cepacia complex | PAS domain S-box protein | 6 |
|  | WP_097674379.1 | 1.9E-06 | Qa.IV | -----TTIAEIDQVSGEIAAAVDQQGAATREISSNLQQAADRTRDVNK----- | *Bradyrhizobium sp. C9* | Bacteria; Proteobacteria; Alphaproteobacteria; Rhizobiales; Bradyrhizobiaceae; Bradyrhizobium | chemotaxis protein | 6 |
|  | WP_102817280.1 | 1.9E-06 | Qb.I | LQRTKEQLQSTIEQSETSTEELKASNEELQAINEELRSATEELETS------- | *Burkholderia ubonensis* | Bacteria; Proteobacteria; Betaproteobacteria; Burkholderiales; Burkholderiaceae; Burkholderia; Burkholderia cepacia complex | PAS domain S-box protein | 6 |
| | WP_105411095.1 | 1.9E-06 | Qb.I | ---INKLVSDTVKQSHDSAEYINESSYLISGTADTFDNIFNKINSANDLIQE- | *Clostridium* | Bacteria; Firmicutes; Clostridia; Clostridiales; Clostridiaceae | MULTISPECIES: methyl-accepting chemotaxis protein | 6 |
|  | WP_107203135.1 | 1.9E-06 | Qb.I | LQRTKEQLQSTIEQSETSTEELKASNEELQAINEELRSATEELETS------- | *Paraburkholderia caribensis* | Bacteria; Proteobacteria; Betaproteobacteria; Burkholderiales; Burkholderiaceae; Paraburkholderia | PAS domain S-box protein | 6 |
|  | WP_110329726.1 | 1.9E-06 | Qb.I | LQRTKEQLQSTIEQSETSTEELKASNEELQAINEELRSATEELETS------- | *Paraburkholderia tropica* | Bacteria; Proteobacteria; Betaproteobacteria; Burkholderiales; Burkholderiaceae; Paraburkholderia | PAS domain S-box protein | 6 |
|  | AMN54291.1 | 2E-06 | Qa.IV | IQEIASTMEEVNQFTAAIAAAISQQGDATNEISHNVQQAA------------- | *Labrenzia sp. CP4* | Bacteria; Proteobacteria; Alphaproteobacteria; Rhodobacterales; Rhodobacteraceae; Labrenzia | hypothetical protein ACP90_19820 | 6 |
|  | BAM91033.1 | 2E-06 | Qc.III | -REISGTIERLSEIASTIAAAVEEQGAATQEISRNVQHASQGTKDVS------ | *Bradyrhizobium oligotrophicum S58* | Bacteria; Proteobacteria; Alphaproteobacteria; Rhizobiales; Bradyrhizobiaceae; Bradyrhizobium | methyl-accepting chemotaxis protein signal peptide | 6 |
| | EME67552.1 | 2E-06 | Qc.III.c | ---IGGTIREINGISTAIASAVEEQSAATSEISRNVHEASKAVNDVNKNIA-- | *Magnetospirillum caucaseum* | Bacteria; Proteobacteria; Alphaproteobacteria; Rhodospirillales; Rhodospirillaceae; Magnetospirillum | methyl-accepting chemotaxis sensory transducer partial | 6 |
| | KZY95361.1 | 2E-06 | Qa.III | LDEIVAAIANIQDLSATIASATEEQVSVVNEVAQNINSIRELNETSNTIAQKT | *Oleiphilus sp. HI0073* | Bacteria; Proteobacteria; Gammaproteobacteria; Oceanospirillales; Oleiphilaceae; Oleiphilus | hypothetical protein A3744_34765 | 6 |
|  | OWW22954.1 | 2E-06 | Qb.I | LQRTKEQLQETIEQSEVSNEELRASNEELQAINEELRSATEELETS------- | *Noviherbaspirillum denitrificans* | Bacteria; Proteobacteria; Betaproteobacteria; Burkholderiales; Oxalobacteraceae; Noviherbaspirillum | chemotaxis protein | 6 |
|  | PWC82105.1 | 2E-06 | Qa.III | IRGIADTVQQVNQIATSIASAVEEQTAATGEISRNVQQAAQ------------ | *Azospirillum sp. TSH100* | Bacteria; Proteobacteria; Alphaproteobacteria; Rhodospirillales; Rhodospirillaceae; Azospirillum | chemotaxis protein | 6 |
|  | SDR60476.1 | 2E-06 | Qb.I | LQRTKEQLQSTIEQSETSTEELKASNEELQAINEELRSATEELETS------- | *Paraburkholderia tuberum* | Bacteria; Proteobacteria; Betaproteobacteria; Burkholderiales; Burkholderiaceae; Paraburkholderia | two-component system chemotaxis family CheB/ CheR fusion protein | 6 |
|  | WP_035656357.1 | 2E-06 | Qa.IV | VKEVATAMRQMADVTTAIASAVEQQGAATREISHNVQMAA------------- | *Bradyrhizobium sp. STM 3809* | Bacteria; Proteobacteria; Alphaproteobacteria; Rhizobiales; Bradyrhizobiaceae; Bradyrhizobium | methyl-accepting chemotaxis protein | 6 |
|  | WP_040451535.1 | 2E-06 | Qc.III | IQSISQTIGRIKDISDTISGAVSEQGIATREIADSTQQAAS------------ | *Labrenzia alexandrii* | Bacteria; Proteobacteria; Alphaproteobacteria; Rhodobacterales; Rhodobacteraceae; Labrenzia | methyl-accepting chemotaxis protein | 6 |
|  | WP_041633847.1 | 2E-06 | Qa.I | IEDIGRTIADISTVNTTIASAVEEQQAATREIARNVDEAA------------- | *Magnetospirillum gryphiswaldense* | Bacteria; Proteobacteria; Alphaproteobacteria; Rhodospirillales; Rhodospirillaceae; Magnetospirillum | methyl-accepting chemotaxis protein | 6 |
|  | WP_056913007.1 | 2E-06 | Qa.IV | INAIERTIREIGNISGAIAAAVTEQGAATSEIARSVETAALRT---------- | *Pseudolabrys sp. Root1462* | Bacteria; Proteobacteria; Alphaproteobacteria; Rhizobiales; Xanthobacteraceae; Pseudolabrys | HAMP domain-containing protein | 6 |
|  | WP_073014697.1 | 2E-06 | Qa.IV | -NEITETVAKINEITSTISAAVEQQGAATAEIARNVQEAAE------------ | *Labrenzia suaedae* | Bacteria; Proteobacteria; Alphaproteobacteria; Rhodobacterales; Rhodobacteraceae; Labrenzia | methyl-accepting chemotaxis protein | 6 |
| | WP_073954018.1 | 2E-06 | Qc.III | IETIGSIIENINEIATTISAAIEQQGSATNEITRNVRGAADRTRAVSSSINE- | *Thalassospira sp. TSL5-1* | Bacteria; Proteobacteria; Alphaproteobacteria; Rhodospirillales; Rhodospirillaceae; Thalassospira | HAMP domain-containing protein | 6 |
|  | WP_088708103.1 | 2E-06 | Qb.I | LQRTKEQLQETIEQSEVSNEELRASNEELQAINEELRSATEELETS------- | *Noviherbaspirillum denitrificans* | Bacteria; Proteobacteria; Betaproteobacteria; Burkholderiales; Oxalobacteraceae; Noviherbaspirillum | PAS domain S-box protein | 6 |
|  | WP_097669880.1 | 2E-06 | Qc.III | -RRIFDVINRVGETATSIVSILDEQNAAIGDISRSAESAAA------------ | *Bradyrhizobium ottawaense* | Bacteria; Proteobacteria; Alphaproteobacteria; Rhizobiales; Bradyrhizobiaceae; Bradyrhizobium | hypothetical protein | 6 |
|  | WP_099512823.1 | 2E-06 | Qa.IV | -ETIARTIGEMSQISVSISAAIEEQGAATSEISRNVQEAAR------------ | *Microvirga ossetica* | Bacteria; Proteobacteria; Alphaproteobacteria; Rhizobiales; Methylobacteriaceae; Microvirga | methyl-accepting chemotaxis protein | 6 |
|  | WP_106325515.1 | 2E-06 | Qc.III.c | --AISQVIVQISEISTVIASAVEEQAAATKEVTANINGTTKAIDDA------- | *Rhizobium tropici* | Bacteria; Proteobacteria; Alphaproteobacteria; Rhizobiales; Rhizobiaceae; Rhizobium/Agrobacterium group; Rhizobium | hypothetical protein | 6 |
|  | WP_109443714.1 | 2E-06 | Qa.III | IRGIADTVQQVNQIATSIASAVEEQTAATGEISRNVQQAAQ------------ | *Azospirillum sp. TSH100* | Bacteria; Proteobacteria; Alphaproteobacteria; Rhodospirillales; Rhodospirillaceae; Azospirillum | methyl-accepting chemotaxis protein | 6 |
| | WP_111420849.1 | 2E-06 | Qc.III | IKEIGGTIARIAEIAAAIAAAVEEQGAATSEIARNVQQAAAGTTEVARNISD- | *Rhodoplanes roseus* | Bacteria; Proteobacteria; Alphaproteobacteria; Rhizobiales; Hyphomicrobiaceae; Rhodoplanes | methyl-accepting chemotaxis protein | 6 |
|  | ADG13135.1 | 2.1E-06 | Qa.IV | --------DEVNNAFLKIKEAVDKAHEKIEGIKQAAQNVTENVEKALRDVQ-- | *Methanocaldococcus infernus ME* | Archaea; Euryarchaeota; Methanococci; Methanococcales; Methanocaldococcaceae; Methanocaldococcus | methyl-accepting chemotaxis sensory transducer | 6 |
|  | OSI82311.1 | 2.1E-06 | Qa.III | IRAITGTIGTINEIASSIASAVEEQSAATQEIARNVQ----HAAQGANE---- | *Bradyrhizobium canariense* | Bacteria; Proteobacteria; Alphaproteobacteria; Rhizobiales; Bradyrhizobiaceae; Bradyrhizobium | hypothetical protein BSZ21_00085 partial | 6 |
|  | WP_027533933.1 | 2.1E-06 | Qa.III | IQGIGATINEVNQIAASIAAAVEEQGTATGEIARNVQQAAQ------------ | *Bradyrhizobium sp. WSM3983* | Bacteria; Proteobacteria; Alphaproteobacteria; Rhizobiales; Bradyrhizobiaceae; Bradyrhizobium | methyl-accepting chemotaxis protein | 6 |
|  | WP_043461097.1 | 2.1E-06 | Qb.I | LRRVKEQLQGTIEQSETSNEELKASNEELQAINEELRSATEELETS------- | *Azohydromonas australica* | Bacteria; Proteobacteria; Betaproteobacteria; Burkholderiales; Alcaligenaceae; Azohydromonas | PAS domain S-box protein | 6 |
|  | WP_054030258.1 | 2.1E-06 | Qa.III | IENITRVVKSVNEVVTTIASAVEEQSTTTGEIAGNIAQAS------------- | *Desulfatitalea tepidiphila* | Bacteria; Proteobacteria; Deltaproteobacteria; Desulfobacterales; Desulfobacteraceae; Desulfatitalea | methyl-accepting chemotaxis protein | 6 |
| | WP_056183474.1 | 2.1E-06 | Qc.III | --GITEVIQRLNRISGDISAAVDQQDAATQEIARNTVRTSEGTRAVSGHIA-- | *Methylobacterium sp. Leaf113* | Bacteria; Proteobacteria; Alphaproteobacteria; Rhizobiales; Methylobacteriaceae; Methylobacterium | methyl-accepting chemotaxis protein | 6 |
|  | WP_081713261.1 | 2.1E-06 | Qa.IV | IQEIASTMEEVNQFTAAIAAAISQQGDATNEISHNVQQAA------------- | *Labrenzia* | Bacteria; Proteobacteria; Alphaproteobacteria; Rhodobacterales; Rhodobacteraceae | MULTISPECIES: methyl-accepting chemotaxis protein | 6 |
|  | WP_102648780.1 | 2.1E-06 | Qb.I | LQRTREQLQATIEHSETSTEELKASNEELQAINEELRSATEELETS------- | *Burkholderia dabaoshanensis* | Bacteria; Proteobacteria; Betaproteobacteria; Burkholderiales; Burkholderiaceae; Burkholderia | response regulator | 6 |
| | CCC98891.1 | 2.2E-06 | Qc.III | --AIGGIIGQINEISTTIAAAIEEQGAATNEISRNVQEAATGTHQVSGSIT-- | *Azospirillum brasilense Sp245* | Bacteria; Proteobacteria; Alphaproteobacteria; Rhodospirillales; Rhodospirillaceae; Azospirillum | putative methyl-accepting chemotaxis protein | 6 |
| | KIZ42319.1 | 2.2E-06 | Qc.III | IKEIGTTIGRMAEISATIASAVEQQGAATQEISRNVQQAAQGTHQVTANITD- | *Rhodopseudomonas palustris* | Bacteria; Proteobacteria; Alphaproteobacteria; Rhizobiales; Bradyrhizobiaceae; Rhodopseudomonas | chemotaxis protein partial | 6 |
|  | OYU89374.1 | 2.2E-06 | Qa.III | IKQIGNTIGEISGIASEIASAVEQQGAATREIAQNVQRVAH------------ | *Bradyrhizobiaceae bacterium PARB1* | Bacteria; Proteobacteria; Alphaproteobacteria; Rhizobiales; Bradyrhizobiaceae | methyl-accepting chemotaxis protein | 6 |
|  | PXW52606.1 | 2.2E-06 | Qa.IV | -ISITDRIREISGLTTSIAAAVEEQGAATQEIVRNVAQAAQ------------ | *Methylobacterium sp. B4* | Bacteria; Proteobacteria; Alphaproteobacteria; Rhizobiales; Methylobacteriaceae; Methylobacterium | methyl-accepting chemotaxis protein | 6 |
|  | RAI36322.1 | 2.2E-06 | Qc.III | IKEIGGTIGRIAEITATIAAAVEEQGAATNEIARNVQQAASGTTEVA------ | *Rhodoplanes roseus* | Bacteria; Proteobacteria; Alphaproteobacteria; Rhizobiales; Hyphomicrobiaceae; Rhodoplanes | methyl-accepting chemotaxis protein partial | 6 |
|  | WP_045013855.1 | 2.2E-06 | Qa.III | IQSICATIQQVDEISAAIAAAVGQQGAATQEIAQNVQQAA------------- | *Bradyrhizobium sp. LTSP849* | Bacteria; Proteobacteria; Alphaproteobacteria; Rhizobiales; Bradyrhizobiaceae; Bradyrhizobium | methyl-accepting chemotaxis protein | 6 |
|  | WP_054031145.1 | 2.2E-06 | Qa.III | ISNITEVVNEVNDIVSKIAAAVEEQSTTTKEIAGNVAQAS------------- | *Desulfatitalea tepidiphila* | Bacteria; Proteobacteria; Deltaproteobacteria; Desulfobacterales; Desulfobacteraceae; Desulfatitalea | methyl-accepting chemotaxis protein | 6 |
| | WP_073012393.1 | 2.2E-06 | Qa.III | ---ITQTISKVSEIATSIASAVEEQTAATTEISHNVQMAAHNTEEFVSEIVR- | *Labrenzia suaedae* | Bacteria; Proteobacteria; Alphaproteobacteria; Rhodobacterales; Rhodobacteraceae; Labrenzia | chemotaxis protein | 6 |
|  | WP_082809792.1 | 2.2E-06 | Qa.IV | IQEIASTMEEVNQFTAAIAAAISQQGDATNEISHNVQQAA------------- | *Labrenzia sp. CP4* | Bacteria; Proteobacteria; Alphaproteobacteria; Rhodobacterales; Rhodobacteraceae; Labrenzia | methyl-accepting chemotaxis protein | 6 |
|  | WP_109084413.1 | 2.2E-06 | Qa.III | IRNIGRTITQVNEISNSIAAAVEEQSAATGEIGRNVQQAAQGTQEVSSS---- | *Azospirillum sp. TSH100* | Bacteria; Proteobacteria; Alphaproteobacteria; Rhodospirillales; Rhodospirillaceae; Azospirillum | HAMP domain-containing protein | 6 |
| | WP_111360137.1 | 2.2E-06 | Qc.III | IKEIGGTIGRIAEIASTIAAAVEEQGAATSEIARNVQQAAAGTGEVARNV--- | *Rhodoplanes elegans* | Bacteria; Proteobacteria; Alphaproteobacteria; Rhizobiales; Hyphomicrobiaceae; Rhodoplanes | methyl-accepting chemotaxis protein | 6 |
| | WP_114103573.1 | 2.2E-06 | Qc.III | IESIGKTIGEISEIAATIASAVEEQGAATEEINRNVQEASTGTTDVSSNI--- | *Thalassospira profundimaris* | Bacteria; Proteobacteria; Alphaproteobacteria; Rhodospirillales; Rhodospirillaceae; Thalassospira | HAMP domain-containing protein | 6 |
|  | RAI17669.1 | 2.3E-06 | Qc.III.c | -EGMTRTIGRIHEMSGAIAAAVEEQGAATAEIAVHVDEASQR----------- | *Rhodoblastus acidophilus* | Bacteria; Proteobacteria; Alphaproteobacteria; Rhizobiales; Beijerinckiaceae; Rhodoblastus | methyl-accepting chemotaxis protein | 6 |
|  | WP_008196467.1 | 2.3E-06 | Qa.III | --AITSTIGKINEITTSISAAVEQQGAATNDIASNIQQAAAGTQQVSD----- | *Labrenzia alexandrii* | Bacteria; Proteobacteria; Alphaproteobacteria; Rhodobacterales; Rhodobacteraceae; Labrenzia | HAMP domain-containing protein | 6 |
|  | WP_050666614.1 | 2.3E-06 | Qa.III | LEQIVESIGQISDMATQISSAAEEQRAVTDEVSRNIQAVK------------- | *Aeromonas schubertii* | Bacteria; Proteobacteria; Gammaproteobacteria; Aeromonadales; Aeromonadaceae; Aeromonas | methyl-accepting chemotaxis protein | 6 |
|  | WP_055672372.1 | 2.3E-06 | Qa.III | --AITSTIGKINEITTSISAAVEQQGAATNDIASNIQQAAAGTQQVSD----- | *Labrenzia alexandrii* | Bacteria; Proteobacteria; Alphaproteobacteria; Rhodobacterales; Rhodobacteraceae; Labrenzia | HAMP domain-containing protein | 6 |
| | WP_056481546.1 | 2.3E-06 | Qc.III | --GITEVIQRLNRISGDISAAVDQQDAATQEIARNTVRTSEGTRAVSGHI--- | *Methylobacterium sp. Leaf117* | Bacteria; Proteobacteria; Alphaproteobacteria; Rhizobiales; Methylobacteriaceae; Methylobacterium | methyl-accepting chemotaxis protein | 6 |
| | WP_080672155.1 | 2.3E-06 | Qc.III | -RDIGGSIGRLSEIASAIAAAVEEQGAATQEIARNVQKAAQGTQQVSANV--- | *Bradyrhizobium japonicum* | Bacteria; Proteobacteria; Alphaproteobacteria; Rhizobiales; Bradyrhizobiaceae; Bradyrhizobium | HAMP domain-containing protein | 6 |
|  | WP_081486889.1 | 2.3E-06 | Qa.IV | IQAIAATIGDISEITTTIASAVEEQQNATTTEISRSVAQAA------------ | *Nitrospirillum amazonense* | Bacteria; Proteobacteria; Alphaproteobacteria; Rhodospirillales; Rhodospirillaceae; Nitrospirillum | HAMP domain-containing protein | 6 |
| | WP_082114107.1 | 2.3E-06 | Qc.III | LEAIGNSIREVNEIGEAISTAVEEQGAATQEIARNVEQASAGTREVSSNIQ-- | *Kiloniella litopenaei* | Bacteria; Proteobacteria; Alphaproteobacteria; Kiloniellales; Kiloniellaceae; Kiloniella | methyl-accepting chemotaxis protein | 6 |
| | WP_085899186.1 | 2.3E-06 | Qc.III | LEAIGNSIREVNEIGEAISTAVEEQGAATQEIARNVEQASAGTREVSSNIQ-- | *Kiloniella majae* | Bacteria; Proteobacteria; Alphaproteobacteria; Kiloniellales; Kiloniellaceae; Kiloniella | methyl-accepting chemotaxis protein | 6 |

| | | | | | | | | |
|---|---|---|---|---|---|---|---|---|
| | **WP_085905987.1** | 2.3E-06 | Qc.III | LEAIGNSIREVNEIGEAISTAVEEQGAATQEIARNVEQASAGTREVSSNIQ-- | *Kiloniella majae* | Bacteria; Proteobacteria; Alphaproteobacteria; Kiloniellales; Kiloniellaceae; Kiloniella | methyl-accepting chemotaxis protein | 6 |
| | **WP_085909961.1** | 2.3E-06 | Qc.III | LEAIGNSIREVNEIGEAISTAVEEQGAATQEIARNVEQASAGTREVSSNIQ-- | *Kiloniella majae* | Bacteria; Proteobacteria; Alphaproteobacteria; Kiloniellales; Kiloniellaceae; Kiloniella | methyl-accepting chemotaxis protein | 6 |
| | **WP_086087583.1** | 2.3E-06 | Qa.IV | ---IKNTIRDLGEISSAIAAAVTEQGAATQEIARSVETASRRSADTADQIGK- | *Pseudorhodoplanes sinuspersici* | Bacteria; Proteobacteria; Alphaproteobacteria; Rhizobiales; Pseudorhodoplanes | methyl-accepting chemotaxis protein | 6 |
|  | **WP_094457510.1** | 2.3E-06 | Qc.III.c | ---IGTIVEQVNRISIAIASAVEEQNAATTEIARSARHASGGVETVT------ | *Niveispirillum lacus* | Bacteria; Proteobacteria; Alphaproteobacteria; Rhodospirillales; Rhodospirillaceae; Niveispirillum | methyl-accepting chemotaxis protein | 6 |
| | **WP_097665434.1** | 2.3E-06 | Qc.III | -RDIGGSIGRLSEIASAIAAAVEEQGAATQEIARNVQKAAQGTQQVSANV--- | *Bradyrhizobium sp. Y36* | Bacteria; Proteobacteria; Alphaproteobacteria; Rhizobiales; Bradyrhizobiaceae; Bradyrhizobium | HAMP domain-containing protein | 6 |
|  | **WP_111421413.1** | 2.3E-06 | Qa.IV | IKEIADTIGRLSEIASTIAAAVEQQGAATQEISRNVQQAA------------ | *Rhodoplanes roseus* | Bacteria; Proteobacteria; Alphaproteobacteria; Rhizobiales; Hyphomicrobiaceae; Rhodoplanes | methyl-accepting chemotaxis protein | 6 |
|  | **CDL01374.1** | 2.4E-06 | Qa.I | IEDIGRTIADISTVNTTIASAVEEQQAATREIARNVDEAA------------ | *Magnetospirillum gryphiswaldense MSR-1 v2* | Bacteria; Proteobacteria; Alphaproteobacteria; Rhodospirillales; Rhodospirillaceae; Magnetospirillum | putative Methyl-accepting chemotaxis protein | 6 |
|  | **EJW09834.1** | 2.4E-06 | Qc.III | IKEIGETIGKIAEIAAAIAAAVEEQGAATSEISRNIQQAATGTTQVAAH---- | *Rhodovulum sp. PH10* | Bacteria; Proteobacteria; Alphaproteobacteria; Rhodobacterales; Rhodobacteraceae; Rhodovulum | Methyl-accepting chemotaxis receptor protein | 6 |
| | **PLX44104.1** | 2.4E-06 | Qc.III | IEAIMKIIQQISEISSGISSAVEEQSAVTQDISTHMQSASQGVELVTNNIQ-- | *Rhizobiales bacterium* | Bacteria; Proteobacteria; Alphaproteobacteria; Rhizobiales | chemotaxis protein | 6 |
| | **SFH98220.1** | 2.4E-06 | Qc.III | IQEISGTIERLSEVSAAIAAAVEEQGAATQEISRNVQQASVGTQQVSSNISD- | *Bradyrhizobium sp. Gha* | Bacteria; Proteobacteria; Alphaproteobacteria; Rhizobiales; Bradyrhizobiaceae; Bradyrhizobium | methyl-accepting chemotaxis sensory transducer | 6 |
|  | **SOD98204.1** | 2.4E-06 | Qc.III.c | --DMSGTIDLLNEISGNISAAVEEQSAVTRDISQNMQVAADGVHRI------- | *Caenispirillum bisanense* | Bacteria; Proteobacteria; Alphaproteobacteria; Rhodospirillales; Rhodospirillaceae; Caenispirillum | methyl-accepting chemotaxis sensory transducer with Pas/Pac sensor | 6 |
|  | **WP_005348206.1** | 2.4E-06 | Qa.III | LEQIVESIGQISDMATQISSAAEEQRAVTDEVSRNIQAVK------------ | *Aeromonas diversa* | Bacteria; Proteobacteria; Gammaproteobacteria; Aeromonadales; Aeromonadaceae; Aeromonas | methyl-accepting chemotaxis protein | 6 |
| | **WP_008889109.1** | 2.4E-06 | Qa.III | IDGITGIIANLNDIASEVAAAVEEQGQATTEISTSVQQAVQSTQEVSNHI--- | *Thalassospira profundimaris* | Bacteria; Proteobacteria; Alphaproteobacteria; Rhodospirillales; Rhodospirillaceae; Thalassospira | methyl-accepting chemotaxis protein | 6 |
| | **WP_008942673.1** | 2.4E-06 | Qc.III | IQSITQIIERMHEASTAIAAAIEEQTAATREIARNVEQAAAGTDEVTRSIAKV | *Oceanibaculum indicum* | Bacteria; Proteobacteria; Alphaproteobacteria; Rhodospirillales; Rhodospirillaceae; Oceanibaculum | HAMP domain-containing protein | 6 |
| | **WP_008960397.1** | 2.4E-06 | Qc.III | -REISGTIERLSEIASTIAAAVEEQGAATQEISRNVQQAAQGTQQVSSNVS-- | *Bradyrhizobium sp. STM 3809* | Bacteria; Proteobacteria; Alphaproteobacteria; Rhizobiales; Bradyrhizobiaceae; Bradyrhizobium | methyl-accepting chemotaxis protein | 6 |
| | **WP_025034554.1** | 2.4E-06 | Qc.III | IKDISHTIERLSEISSTIAAAVEEQGAATQEISRNVQQAATGTHQVSSNITD- | *Bradyrhizobium sp. DOA9* | Bacteria; Proteobacteria; Alphaproteobacteria; Rhizobiales; Bradyrhizobiaceae; Bradyrhizobium | HAMP domain-containing protein | 6 |
|  | **WP_040621756.1** | 2.4E-06 | Qc.III | IKEIGETIGKIAEIAAAIAAAVEEQGAATSEISRNIQQAATGTTQVAAH---- | *Rhodovulum sp. PH10* | Bacteria; Proteobacteria; Alphaproteobacteria; Rhodobacterales; Rhodobacteraceae; Rhodovulum | methyl-accepting chemotaxis protein | 6 |
| | **WP_041811170.1** | 2.4E-06 | Qc.III | --AIGGIIGQINEISTTIAAAIEEQGAATNEISRNVQEAATGTHQVSGSIT-- | *Azospirillum brasilense* | Bacteria; Proteobacteria; Alphaproteobacteria; Rhodospirillales; Rhodospirillaceae; Azospirillum | HAMP domain-containing protein | 6 |
| | **WP_063085683.1** | 2.4E-06 | Qa.III | IDGITGIIANLNDIASEVAAAVEEQGQATTEISTSVQQAVQSTQEVSNHI--- | *Thalassospira* | Bacteria; Proteobacteria; Alphaproteobacteria; Rhodospirillales; Rhodospirillaceae | MULTISPECIES: methyl-accepting chemotaxis protein | 6 |
| | **WP_064780427.1** | 2.4E-06 | Qa.III | IDGITGIIANLNDIASEVAAAVEEQGQATTEISTSVQQAVQSTQEVSNHI--- | *Thalassospira tepidiphila* | Bacteria; Proteobacteria; Alphaproteobacteria; Rhodospirillales; Rhodospirillaceae; Thalassospira | methyl-accepting chemotaxis protein | 6 |
| | **WP_064788185.1** | 2.4E-06 | Qa.III | IDGITGIIANLNDIASEVAAAVEEQGQATTEISTSVQQAVQSTQEVSNHI--- | *Thalassospira profundimaris* | Bacteria; Proteobacteria; Alphaproteobacteria; Rhodospirillales; Rhodospirillaceae; Thalassospira | methyl-accepting chemotaxis protein | 6 |
|  | **WP_083456147.1** | 2.4E-06 | Qa.III | INQIGKVIDTVNEIVATIATAVEEQSSATKEIAENIAQISE----------- | *Desulfosarcina cetonica* | Bacteria; Proteobacteria; Deltaproteobacteria; Desulfobacterales; Desulfobacteraceae; Desulfosarcina | methyl-accepting chemotaxis protein | 6 |
| | **WP_084195015.1** | 2.4E-06 | Qa.III.b | IRDISSVIGEVNALAQDIAGAVDEQGMATRQIASNVSEAARGTQDVSNSV--- | *Insolitispirillum peregrinum* | Bacteria; Proteobacteria; Alphaproteobacteria; Rhodospirillales; Rhodospirillaceae; Insolitispirillum | methyl-accepting chemotaxis protein | 6 |
| | **WP_092227467.1** | 2.4E-06 | Qc.III | IQEISGTIERLSEVSAAIAAAVEEQGAATQEISRNVQQASVGTQQVSSNISD- | *Bradyrhizobium sp. Gha* | Bacteria; Proteobacteria; Alphaproteobacteria; Rhizobiales; Bradyrhizobiaceae; Bradyrhizobium | methyl-accepting chemotaxis protein | 6 |
|  | **WP_103012421.1** | 2.4E-06 | Qa.IV | IKEIGQTITRMSEIASTIASAVEEQGAATQEISRNVQEAA------------ | *Rhodopseudomonas palustris* | Bacteria; Proteobacteria; Alphaproteobacteria; Rhizobiales; Bradyrhizobiaceae; Rhodopseudomonas | HAMP domain-containing protein | 6 |
|  | **AIB15151.1** | 2.5E-06 | Qa.III | IQGIAAVIGEVNEIATAIAAAVEEQGAATQEITRNVQQAAA----------- | *Azospirillum brasilense* | Bacteria; Proteobacteria; Alphaproteobacteria; Rhodospirillales; Rhodospirillaceae; Azospirillum | hypothetical protein ABAZ39_24980 (plasmid) | 6 |
|  | **OGB20357.1** | 2.5E-06 | Qb.I | LQRTKEQLQETIEHSETSTEELKASNEELQAINEELRSATEELETS------- | *Burkholderiales bacterium RIFCSPLOWO2_02_FULL_57_36* | Bacteria; Proteobacteria; Betaproteobacteria; Burkholderiales | chemotaxis protein | 6 |
|  | **PNQ97736.1** | 2.5E-06 | Qa.III | IQGIAAVIGEVNEIATAIAAAVEEQGAATQEITRNVQQAAA----------- | *Azospirillum brasilense* | Bacteria; Proteobacteria; Alphaproteobacteria; Rhodospirillales; Rhodospirillaceae; Azospirillum | methyl-accepting chemotaxis protein (plasmid) | 6 |
| | **SIT19155.1** | 2.5E-06 | Qa.III.b | IRDISSVIGEVNALAQDIAGAVDEQGMATRQIASNVSEAARGTQDVSNSV--- | *Insolitispirillum peregrinum* | Bacteria; Proteobacteria; Alphaproteobacteria; Rhodospirillales; Rhodospirillaceae; Insolitispirillum | methyl-accepting chemotaxis sensory transducer with Cache sensor | 6 |
|  | **WP_017487447.1** | 2.5E-06 | Qa.IV | IRAIVSTITDLSEISMSVSSAVTEQAAVTQEIATNMQTAARSV---------- | *Methylobacterium sp. MB200* | Bacteria; Proteobacteria; Alphaproteobacteria; Rhizobiales; Methylobacteriaceae; Methylobacterium | PAS domain S-box protein | 6 |
|  | **WP_029582404.1** | 2.5E-06 | Qa.IV | IRNIGQTISEINEVTAAIAAAVEQQGAATREIARNIQHAA------------ | *Bradyrhizobium sp. URHD0069* | Bacteria; Proteobacteria; Alphaproteobacteria; Rhizobiales; Bradyrhizobiaceae; Bradyrhizobium | HAMP domain-containing protein | 6 |
|  | **CCE04793.1** | 2.6E-06 | Qa.IV | IQAISSTIQRVNQIASAIAAAVEEQGAATREIARNVQQAS------------ | *Bradyrhizobium sp. STM 3843* | Bacteria; Proteobacteria; Alphaproteobacteria; Rhizobiales; Bradyrhizobiaceae; Bradyrhizobium | fragment of conserved hypothetical protein (part 2) | 6 |
|  | **EAR59773.1** | 2.6E-06 | Qa.IV | -KQIRNSISEINDMSTLIASSTEEQTAVTSNITQTIETTDSLTKN-------- | *Neptuniibacter caesariensis* | Bacteria; Proteobacteria; Gammaproteobacteria; Oceanospirillales; Neptuniibacter | chemotaxis sensory transducer | 6 |
|  | **EGH18032.1** | 2.6E-06 | Qa.III | ---ISQALRNLNSLNTSIASATLQQTHVVDDINQNVTQAAQLSQS-------- | *Pseudomonas savastanoi pv. glycinea str. race 4* | Bacteria; Proteobacteria; Gammaproteobacteria; Pseudomonadales; Pseudomonadaceae; Pseudomonas | methyl-accepting chemotaxis protein partial | 6 |
| | **OYY13241.1** | 2.6E-06 | Qc.III | LGAIHGTISNLNEVSTIISGAVEEQTAVTREVSANMHDAANGIDSVLVNMN-- | *Rhizobiales bacterium 35-68-8* | Bacteria; Proteobacteria; Alphaproteobacteria; Rhizobiales | hypothetical protein B7Y70_02785 | 6 |
|  | **WP_006959133.1** | 2.6E-06 | Qa.III | LEEINTAVQTISDMATQIATAAEEQTHVTNEITQNITSIK------------ | *Vibrio coralliilyticus* | Bacteria; Proteobacteria; Gammaproteobacteria; Vibrionales; Vibrionaceae; Vibrio | methyl-accepting chemotaxis protein | 6 |
|  | **WP_019276682.1** | 2.6E-06 | Qa.III | LEEINTAVQTISDMATQIATAAEEQTHVTNEITQNITSIK------------ | *Vibrio coralliilyticus* | Bacteria; Proteobacteria; Gammaproteobacteria; Vibrionales; Vibrionaceae; Vibrio | methyl-accepting chemotaxis protein | 6 |
|  | **WP_027276276.1** | 2.6E-06 | Qa.IV | IKEIGQTISRMSEIASTIASAVEEQGAATQEISRNVQQAA------------ | *Rhodopseudomonas palustris* | Bacteria; Proteobacteria; Alphaproteobacteria; Rhizobiales; Bradyrhizobiaceae; Rhodopseudomonas | methyl-accepting chemotaxis protein | 6 |
| | **WP_027300779.1** | 2.6E-06 | Qc.III | IQSIGGTIGAINEIATTIASAVEEQGAATQEIARNVQGAAQGTEQVSSNI--- | *Rhodospirillales bacterium URHD0088* | Bacteria; Proteobacteria; Alphaproteobacteria; Rhodospirillales; unclassified Rhodospirillales | methyl-accepting chemotaxis protein | 6 |
|  | **WP_038162201.1** | 2.6E-06 | Qa.III | LEEINTAVQTISDMATQIATAAEEQTHVTNEITQNITSIK------------ | *Vibrio sp. B183* | Bacteria; Proteobacteria; Gammaproteobacteria; Vibrionales; Vibrionaceae; Vibrio | methyl-accepting chemotaxis protein | 6 |
|  | **WP_042701040.1** | 2.6E-06 | Qa.III | IRNIGRTITQVNEISNSIAAAVEEQSAATGEIGRNVQQAAQGTQ--------- | *Azospirillum sp. B506* | Bacteria; Proteobacteria; Alphaproteobacteria; Rhodospirillales; Rhodospirillaceae; Azospirillum | HAMP domain-containing protein | 6 |
| | **WP_044563049.1** | 2.6E-06 | Qc.III | IQGIGGIINRINAISGAMAAAVEEQNAVARDISRNMQTAADGVSSISESMT-- | *Azospirillum sp. B4* | Bacteria; Proteobacteria; Alphaproteobacteria; Rhodospirillales; Rhodospirillaceae; Azospirillum | methyl-accepting chemotaxis protein | 6 |
| | **WP_053006066.1** | 2.6E-06 | Qc.III | LEAIGNSIREVNEIGEAISTAVEEQAAATQEIARNVEQASAGTREVSSNIQ-- | *Kiloniella spongiae* | Bacteria; Proteobacteria; Alphaproteobacteria; Kiloniellales; Kiloniellaceae; Kiloniella | HAMP domain-containing protein | 6 |
|  | **WP_099609794.1** | 2.6E-06 | Qa.III | LEEINTAVQTISDMATQIATAAEEQTHVTNEITQNITSIK------------ | *Vibrio coralliilyticus* | Bacteria; Proteobacteria; Gammaproteobacteria; Vibrionales; Vibrionaceae; Vibrio | methyl-accepting chemotaxis protein | 6 |
|  | **WP_103011283.1** | 2.6E-06 | Qa.IV | IKEIGQTISRMSEIASTIASAVEEQGAATQEISRNVQQAA------------ | *Rhodopseudomonas palustris* | Bacteria; Proteobacteria; Alphaproteobacteria; Rhizobiales; Bradyrhizobiaceae; Rhodopseudomonas | methyl-accepting chemotaxis protein | 6 |
|  | **EKD34872.1** | 2.7E-06 | Qa.III | IDKISKVIADINSVINGIATAVVEQSAATSEIATNISQAS------------ | *uncultured bacterium* | Bacteria; environmental samples | chemotaxis transducer | 6 |
|  | **KJE37440.1** | 2.7E-06 | Qa.III | -TEIVTSISDINAMVEQIAVAAEEQVNVTDEINQNISDIDNKSQ--------- | *Thalassomonas actiniarum* | Bacteria; Proteobacteria; Gammaproteobacteria; Alteromonadales; Colwelliaceae; Thalassomonas | hypothetical protein SG35_30640 | 6 |
|  | **OCR95364.1** | 2.7E-06 | Qa.IV | --EIEANVNILSQGINEMSQSINEQTEAINQINEAVATVDEQTKQ-NV----- | *Campylobacter fetus subsp. testudinum* | Bacteria; Proteobacteria; Epsilonproteobacteria; Campylobacterales; Campylobacteraceae; Campylobacter | chemotaxis protein partial | 6 |
|  | **SFK28642.1** | 2.7E-06 | Qa.IV | IQSIAGRIEEVSQIATLIASAVEEQGAATQEIARSVQQTVEA---------- | *Roseomonas stagni DSM 19981* | Bacteria; Proteobacteria; Alphaproteobacteria; Rhodospirillales; Acetobacteraceae; Roseomonas | methyl-accepting chemotaxis protein | 6 |
| | **WP_011389831.1** | 2.7E-06 | Qc.III | INEIATTIGEINEISANIAAAMEQQGAATQEIARNVEQAATGTDQVTRNIH-- | *Rhodospirillum rubrum* | Bacteria; Proteobacteria; Alphaproteobacteria; Rhodospirillales; Rhodospirillaceae; Rhodospirillum | HAMP domain-containing protein | 6 |
|  | **WP_022723137.1** | 2.7E-06 | Qa.III | IREISGTIEKLSEISSTIASAVEEQGTATQEISRNVQQASQGTQQ-------- | *Rhodopseudomonas sp. B29* | Bacteria; Proteobacteria; Alphaproteobacteria; Rhizobiales; Bradyrhizobiaceae; Rhodopseudomonas | HAMP domain-containing protein | 6 |
| | **WP_027533484.1** | 2.7E-06 | Qc.III | IQSIGTTIQRVNEIAAAIAAAVEEQGAATREIARNVQQASQGTNEVSRHIS-- | *Bradyrhizobium sp. WSM3983* | Bacteria; Proteobacteria; Alphaproteobacteria; Rhizobiales; Bradyrhizobiaceae; Bradyrhizobium | HAMP domain-containing protein | 6 |
|  | **WP_028241848.1** | 2.7E-06 | Qb.I | LARTKEQLQTTIEQSETSNEELKASNEELQAINEELRSATEELETS------- | *Pseudomonas azotifigens* | Bacteria; Proteobacteria; Gammaproteobacteria; Pseudomonadales; Pseudomonadaceae; Pseudomonas | PAS domain S-box protein | 6 |
| | **WP_045005086.1** | 2.7E-06 | Qc.III | IQSIGTTIQRVNEIAAAIAAAVEEQGAATREIARNVQQASQGTNEVSRHIS-- | *Bradyrhizobium sp. LTSP857* | Bacteria; Proteobacteria; Alphaproteobacteria; Rhizobiales; Bradyrhizobiaceae; Bradyrhizobium | HAMP domain-containing protein | 6 |
|  | **WP_045005207.1** | 2.7E-06 | Qa.IV | IREISSTIGRLSEISSTVAAAVEEQGAATQEISRNVQQAA------------ | *Bradyrhizobium sp. LTSP857* | Bacteria; Proteobacteria; Alphaproteobacteria; Rhizobiales; Bradyrhizobiaceae; Bradyrhizobium | methyl-accepting chemotaxis protein | 6 |
|  | **WP_045010599.1** | 2.7E-06 | Qa.IV | IREISSTIGRLSEISSTVAAAVEEQGAATQEISRNVQQAA------------ | *Bradyrhizobium sp. LTSP849* | Bacteria; Proteobacteria; Alphaproteobacteria; Rhizobiales; Bradyrhizobiaceae; Bradyrhizobium | methyl-accepting chemotaxis protein | 6 |
| | **WP_045012442.1** | 2.7E-06 | Qc.III | IQSIGTTIQRVNEIAAAIAAAVEEQGAATREIARNVQQASQGTNEVSRHIS-- | *Bradyrhizobium sp. LTSP849* | Bacteria; Proteobacteria; Alphaproteobacteria; Rhizobiales; Bradyrhizobiaceae; Bradyrhizobium | methyl-accepting chemotaxis protein | 6 |
|  | **WP_053239283.1** | 2.7E-06 | Qa.III | IKKIGEIIEEINNISGNISSTVRQQGSSTQEIANNIQEAAKSVQDV------- | *Rickettsiales bacterium Ac37b* | Bacteria; Proteobacteria; Alphaproteobacteria; Rickettsiales | methyl-accepting chemotaxis protein | 6 |
| | **WP_057743573.1** | 2.7E-06 | Qc.III | IQSIGTTIQRVNEIAAAIAAAVEEQGAATREIARNVQQASQGTNEVSRHIS-- | *Bradyrhizobium manausense* | Bacteria; Proteobacteria; Alphaproteobacteria; Rhizobiales; Bradyrhizobiaceae; Bradyrhizobium | HAMP domain-containing protein | 6 |

| | | | | | | | | |
|---|---|---|---|---|---|---|---|---|
|  | **WP_057854068.1** | 2.7E-06 | Qa.IV | -KEISGTIERLAEISSTIAAAVEEQGAATQEISRNVQQAAQ------------ | *Bradyrhizobium valentinum* | Bacteria; Proteobacteria; Alphaproteobacteria; Rhizobiales; Bradyrhizobiaceae; Bradyrhizobium | methyl-accepting chemotaxis protein | 6 |
| | **WP_090074047.1** | 2.7E-06 | Qc.III | IGEISATIEHMNQIAAAITEAVQQQGQATQEIARNIQEASAGTREVSSNIA-- | *Cohaesibacter marisflavi* | Bacteria; Proteobacteria; Alphaproteobacteria; Rhizobiales; Cohaesibacteraceae | methyl-accepting chemotaxis protein | 6 |
| | **WP_092231054.1** | 2.7E-06 | Qc.III | IQSIGTTIQRVNEIAAAIAAAVEEQGAATREIARNVQQASQGTNEVSRHIS-- | *Bradyrhizobium sp. Gha* | Bacteria; Proteobacteria; Alphaproteobacteria; Rhizobiales; Bradyrhizobiaceae; Bradyrhizobium | methyl-accepting chemotaxis protein | 6 |
|  | **WP_111384553.1** | 2.7E-06 | Qa.IV | IKEIGRTITRVSEIAADIATSVEQQGAATSEIARNVQQAA------------- | *Rhodoplanes piscinae* | Bacteria; Proteobacteria; Alphaproteobacteria; Rhizobiales; Hyphomicrobiaceae; Rhodoplanes | methyl-accepting chemotaxis protein | 6 |
|  | **CCG42370.1** | 2.8E-06 | Qa.IV | IVSIVERIEEINQIATTIASAVEEQSAATNEIARNVQQAA------------- | *Phaeospirillum molischianum DSM 120* | Bacteria; Proteobacteria; Alphaproteobacteria; Rhodospirillales; Rhodospirillaceae; Phaeospirillum | Putative methyl-accepting chemotaxis protein (fragment) | 6 |
|  | **OCR84520.1** | 2.8E-06 | Qa.IV | --EIEANVNILSQGINEMSQSINEQTEAINQINEAVATVDEQTKQ-NV----- | *Campylobacter fetus subsp. testudinum* | Bacteria; Proteobacteria; Epsilonproteobacteria; Campylobacterales; Campylobacteraceae; Campylobacter | chemotaxis protein partial | 6 |
| | **PCI85944.1** | 2.8E-06 | Qa.III | IEQVEEIINNLSEASTGVASAVEQQAAAIDEIAANVNTASTLSEQSSQSMKS- | *Rhizobiales bacterium* | Bacteria; Proteobacteria; Alphaproteobacteria; Rhizobiales | hypothetical protein COB24_11395 | 6 |
| | **PTB98200.1** | 2.8E-06 | Qc.III | IDSIGTTITEISEIASAIASAVEQQGAATDEINRNVQEASAGTGEVTANI--- | *Thalassospira xiamenensis* | Bacteria; Proteobacteria; Alphaproteobacteria; Rhodospirillales; Rhodospirillaceae; Thalassospira | methyl-accepting chemotaxis protein partial | 6 |
|  | **RAI34966.1** | 2.8E-06 | Qa.IV | IKEIGRTITRVSEIAADIATSVEQQGAATSEIARNVQQAA------------- | *Rhodoplanes piscinae* | Bacteria; Proteobacteria; Alphaproteobacteria; Rhizobiales; Hyphomicrobiaceae; Rhodoplanes | hypothetical protein CH340_07370 | 6 |
|  | **WP_005490198.1** | 2.8E-06 | Qa.I | ------QVEEVSSIIDTLAENINQVSYNTESMTDQSKNVMENIKDGNQALK-- | *Halanaerobium saccharolyticum* | Bacteria; Firmicutes; Clostridia; Halanaerobiales; Halanaerobiaceae; Halanaerobium | methyl-accepting chemotaxis protein | 6 |
|  | **WP_019439839.1** | 2.8E-06 | Qc.III | LDSMTESVLQINDLGIQIATAAEEQSSVTEEINRNMTSIQTMVYQLTSN---- | *Moritella marina* | Bacteria; Proteobacteria; Gammaproteobacteria; Alteromonadales; Moritellaceae; Moritella | methyl-accepting chemotaxis protein | 6 |
|  | **WP_049962271.1** | 2.8E-06 | Qb.III | -KDSARSAHEVEDFSAGVADKLTEQNEQMAKVQDAMKEIEDK----------- | *Oribacterium sp. FC2011* | Bacteria; Firmicutes; Clostridia; Clostridiales; Lachnospiraceae; Oribacterium | hypothetical protein | 6 |
|  | **WP_050627054.1** | 2.8E-06 | Qc.III.c | --AISSVIVQISEISTVIASAVEEQAAATKEVTANINGTTKAIDDA------- | *Bradyrhizobium viridifuturi* | Bacteria; Proteobacteria; Alphaproteobacteria; Rhizobiales; Bradyrhizobiaceae; Bradyrhizobium | hypothetical protein | 6 |
|  | **WP_082036457.1** | 2.8E-06 | Qa.IV | -RTIATVIEQVRQISSGIASAVEEQGAATQEIARNVQQAAQ------------ | *Magnetospirillum magnetotacticum* | Bacteria; Proteobacteria; Alphaproteobacteria; Rhodospirillales; Rhodospirillaceae; Magnetospirillum | HAMP domain-containing protein | 6 |
|  | **WP_084693035.1** | 2.8E-06 | Qa.III | -TEIVTSISDINAMVEQIAVAAEEQVNVTDEINQNISDIDNKSQ--------- | *Thalassomonas actiniarum* | Bacteria; Proteobacteria; Gammaproteobacteria; Alteromonadales; Colwelliaceae; Thalassomonas | HAMP domain-containing protein | 6 |
|  | **WP_089266922.1** | 2.8E-06 | Qa.III | IKQIGSTIGEISGIASEIASAVEQQGAATREIAQNVQRVAH------------ | *Tardiphaga sp. OK246* | Bacteria; Proteobacteria; Alphaproteobacteria; Rhizobiales; Bradyrhizobiaceae; Tardiphaga | methyl-accepting chemotaxis protein | 6 |
|  | **WP_089526797.1** | 2.8E-06 | Qc.III | --QVALSAGELATGANEISEEVGDMSESIREIENKVSSYADSSRQMNER---- | *Paenibacillus herberti* | Bacteria; Firmicutes; Bacilli; Bacillales; Paenibacillaceae; Paenibacillus | methyl-accepting chemotaxis protein | 6 |
|  | **WP_090816339.1** | 2.8E-06 | Qb.III | -KDSARSAHEVEDFSAGVADKLTEQNEQMAKVQDAMKEIEDK----------- | *Oribacterium sp. KHPX15* | Bacteria; Firmicutes; Clostridia; Clostridiales; Lachnospiraceae; Oribacterium | hypothetical protein | 6 |
|  | **WP_092141840.1** | 2.8E-06 | Qa.III | IKQIGSTIGEISGIASEIASAVEQQGAATREIAQNVQRVAH------------ | *Bradyrhizobium sp. NFR13* | Bacteria; Proteobacteria; Alphaproteobacteria; Rhizobiales; Bradyrhizobiaceae; Bradyrhizobium | methyl-accepting chemotaxis protein | 6 |
|  | **WP_093758241.1** | 2.8E-06 | Qa.III | IKQIGSTIGEISGIASEIASAVEQQGAATREIAQNVQRVAH------------ | *Tardiphaga sp. OK245* | Bacteria; Proteobacteria; Alphaproteobacteria; Rhizobiales; Bradyrhizobiaceae; Tardiphaga | methyl-accepting chemotaxis protein | 6 |
| | **WP_109517472.1** | 2.8E-06 | Qc.III | --GIGNTVERTNEISTAIAAAVEEQGAATQEIARSAAQVAEATGTVTQRIA-- | *Roseomonas aestuarii* | Bacteria; Proteobacteria; Alphaproteobacteria; Rhodospirillales; Acetobacteraceae; Roseomonas | HAMP domain-containing protein | 6 |
|  | **OEU74331.1** | 2.9E-06 | Qa.III.b | -----KVINDVNEIVSTIATAVEEQSVTTKEIAGNVAQASEGIKEVNE----- | *Desulfuromonadales bacterium C00003093* | Bacteria; Proteobacteria; Deltaproteobacteria; Desulfuromonadales | hypothetical protein BA864_08370 | 6 |
| | **OSJ26015.1** | 2.9E-06 | Qc.III | IQEIGGTIERLSEVAAAIAAAVEEQGAATQEISRNVQQASVGTQEVSSNIT-- | *Bradyrhizobium canariense* | Bacteria; Proteobacteria; Alphaproteobacteria; Rhizobiales; Bradyrhizobiaceae; Bradyrhizobium | methyl-accepting chemotaxis protein | 6 |
|  | **PCK06025.1** | 2.9E-06 | Qa.III.b | -GKINTVIGDVNQIVTTIAAAVEEQNVTTKSIAGNIGQASDGV---------- | *Alteromonadaceae bacterium* | Bacteria; Proteobacteria; Gammaproteobacteria; Alteromonadales; Alteromonadaceae | chemotaxis protein partial | 6 |
|  | **PIE71333.1** | 2.9E-06 | Qa.III | MEQINTVINDVNSIVTTIASAVEEQAVSTNEIASNVAQAA------------- | *Deltaproteobacteria bacterium* | Bacteria; Proteobacteria; Deltaproteobacteria | chemotaxis protein | 6 |
|  | **WP_027278318.1** | 2.9E-06 | Qa.IV | IKEIGQTITRMSEIASTIASAVEEQGAATQEISRNVQEAA------------- | *Rhodopseudomonas palustris* | Bacteria; Proteobacteria; Alphaproteobacteria; Rhizobiales; Bradyrhizobiaceae; Rhodopseudomonas | methyl-accepting chemotaxis protein | 6 |
| | **WP_027516464.1** | 2.9E-06 | Qc.III | IQEIGGTIERLSEVAAAIAAAVEEQGAATQEISRNVQQASVGTQEVSSNIT-- | *Bradyrhizobium sp. WSM1417* | Bacteria; Proteobacteria; Alphaproteobacteria; Rhizobiales; Bradyrhizobiaceae; Bradyrhizobium | methyl-accepting chemotaxis protein | 6 |
| | **WP_027568630.1** | 2.9E-06 | Qc.III | IQEIGGTIERLSEVAAAIAAAVEEQGAATQEISRNVQQASVGTQEVSSNIT-- | *Bradyrhizobium sp. URHA0013* | Bacteria; Proteobacteria; Alphaproteobacteria; Rhizobiales; Bradyrhizobiaceae; Bradyrhizobium | methyl-accepting chemotaxis protein | 6 |
|  | **WP_057863128.1** | 2.9E-06 | Qa.IV | IQSICTTIQEVDEISAAIAAAVGQQGSATQEIARNVQQAAARTG--------- | *Bradyrhizobium lablabi* | Bacteria; Proteobacteria; Alphaproteobacteria; Rhizobiales; Bradyrhizobiaceae; Bradyrhizobium | methyl-accepting chemotaxis protein | 6 |
| | **WP_063993453.1** | 2.9E-06 | Qc.III | IQEIGGTIERLSEVAAAIAAAVEEQGAATQEISRNVQQASVGTQEVSSNIT-- | *Bradyrhizobium sp.* | Bacteria; Proteobacteria; Alphaproteobacteria; Rhizobiales; Bradyrhizobiaceae; Bradyrhizobium | methyl-accepting chemotaxis protein | 6 |
| | **WP_081728431.1** | 2.9E-06 | Qc.III | IQGISETVERINEIAQSVASAVEEQGAATQEIARNVQEASQGTQEVARNIS-- | *Rhodovibrio salinarum* | Bacteria; Proteobacteria; Alphaproteobacteria; Rhodospirillales; Rhodospirillaceae; Rhodovibrio | methyl-accepting chemotaxis protein | 6 |
|  | **WP_085126970.1** | 2.9E-06 | Qa.III | IKGIGQTVETIDEIAASISAAVQEQGAATQEIARNVQQASESTRQVASS---- | *Tistlia consotensis* | Bacteria; Proteobacteria; Alphaproteobacteria; Rhodospirillales; Rhodospirillaceae; Tistlia | methyl-accepting chemotaxis protein | 6 |
| | **WP_085352599.1** | 2.9E-06 | Qc.III | IQEIGGTIERLSEVAAAIAAAVEEQGAATQEISRNVQQASVGTQEVSSNIT-- | *Bradyrhizobium canariense* | Bacteria; Proteobacteria; Alphaproteobacteria; Rhizobiales; Bradyrhizobiaceae; Bradyrhizobium | methyl-accepting chemotaxis protein | 6 |
| | **WP_085361345.1** | 2.9E-06 | Qc.III | IQEIGGTIERLSEVAAAIAAAVEEQGAATQEISRNVQQASVGTQEVSSNIT-- | *Bradyrhizobium canariense* | Bacteria; Proteobacteria; Alphaproteobacteria; Rhizobiales; Bradyrhizobiaceae; Bradyrhizobium | methyl-accepting chemotaxis protein | 6 |
| | **WP_088930801.1** | 2.9E-06 | Qc.III | IQEIGGTIERLSEVAAAIAAAVEEQGAATQEISRNVQQASVGTQEVSSNIT-- | *Bradyrhizobium sp. WSM4349* | Bacteria; Proteobacteria; Alphaproteobacteria; Rhizobiales; Bradyrhizobiaceae; Bradyrhizobium | methyl-accepting chemotaxis protein | 6 |
|  | **WP_111383698.1** | 2.9E-06 | Qa.IV | IKEIGSTIHRIAEIASSIAAAVEEQGAATNEIARNVQQAAQ------------ | *Rhodoplanes piscinae* | Bacteria; Proteobacteria; Alphaproteobacteria; Rhizobiales; Hyphomicrobiaceae; Rhodoplanes | methyl-accepting chemotaxis protein | 6 |
|  | **PLX60048.1** | 3E-06 | Qa.III | LDEINAAVALISDMATQIATAAEEQTHVTNEITQNVTSIK------------- | *Vibrio alginolyticus* | Bacteria; Proteobacteria; Gammaproteobacteria; Vibrionales; Vibrionaceae; Vibrio | chemotaxis protein partial | 6 |
|  | **PVX55459.1** | 3E-06 | Qa.III | IKQIGSTIGEISGIASEIASAVEQQGAATREIAQNVQRVAH------------ | *Tardiphaga sp. OV697* | Bacteria; Proteobacteria; Alphaproteobacteria; Rhizobiales; Bradyrhizobiaceae; Tardiphaga | methyl-accepting chemotaxis sensory transducer | 6 |
|  | **SEH86439.1** | 3E-06 | Qa.III | IKQIGSTIGEISGIASEIASAVEQQGAATREIAQNVQRVAH------------ | *Tardiphaga sp. OK245* | Bacteria; Proteobacteria; Alphaproteobacteria; Rhizobiales; Bradyrhizobiaceae; Tardiphaga | methyl-accepting chemotaxis sensory transducer | 6 |
|  | **SFL35948.1** | 3E-06 | Qa.III | IKQIGSTIGEISGIASEIASAVEQQGAATREIAQNVQRVAH------------ | *Bradyrhizobium sp. NFR13* | Bacteria; Proteobacteria; Alphaproteobacteria; Rhizobiales; Bradyrhizobiaceae; Bradyrhizobium | methyl-accepting chemotaxis sensory transducer | 6 |
|  | **SNT57288.1** | 3E-06 | Qa.III | IKQIGSTIGEISGIASEIASAVEQQGAATREIAQNVQRVAH------------ | *Tardiphaga sp. OK246* | Bacteria; Proteobacteria; Alphaproteobacteria; Rhizobiales; Bradyrhizobiaceae; Tardiphaga | methyl-accepting chemotaxis sensory transducer | 6 |
| | **WP_006611665.1** | 3E-06 | Qc.III | -REISDTIERLSEISATIAAAVEEQGAATQEISRNVQQAAQGTQQVSSNITD- | *Bradyrhizobium sp. ORS 285* | Bacteria; Proteobacteria; Alphaproteobacteria; Rhizobiales; Bradyrhizobiaceae; Bradyrhizobium | methyl-accepting chemotaxis protein | 6 |
| | **WP_011472306.1** | 3E-06 | Qc.III | INEISDTIGRMSEISSMIASAVEEQGAATQEISRNVQHAADGTSKVSSNLT-- | *Rhodopseudomonas palustris* | Bacteria; Proteobacteria; Alphaproteobacteria; Rhizobiales; Bradyrhizobiaceae; Rhodopseudomonas | methyl-accepting chemotaxis protein | 6 |
|  | **WP_029233070.1** | 3E-06 | Qb.I | ---ITNLIKDLAEKSQANMEAITNSRGAVSKAGDSFSKINQELNSTA------ | *Butyrivibrio sp. VCB2006* | Bacteria; Firmicutes; Clostridia; Clostridiales; Lachnospiraceae; Butyrivibrio | methyl-accepting chemotaxis protein | 6 |
|  | **WP_035672138.1** | 3E-06 | Qa.III | IVGIGHTIQDINQIAGGIAAGIGQQGSAIRSIVSNMDQAA------------- | *Azospirillum* | Bacteria; Proteobacteria; Alphaproteobacteria; Rhodospirillales; Rhodospirillaceae | MULTISPECIES: chemotaxis protein | 6 |
|  | **WP_035673232.1** | 3E-06 | Qa.III.b | ---VARVIEQVNGIATIIASAVEQQGAATQEISRNVQQAAEVTREVARD---- | *Azospirillum brasilense* | Bacteria; Proteobacteria; Alphaproteobacteria; Rhodospirillales; Rhodospirillaceae; Azospirillum | HAMP domain-containing protein | 6 |
|  | **WP_053955732.1** | 3E-06 | Qb.III.d | ------QIQDTNQIAYQTVENVNNGTEVIKNTKNQIDKIESSIHNANSVVKN- | *Inediibacterium massiliense* | Bacteria; Firmicutes; Clostridia; Clostridiales; Clostridiaceae; Inediibacterium | methyl-accepting chemotaxis protein | 6 |
| | **WP_092029122.1** | 3E-06 | Qc.III | IQEIGGTIERLSEVAAAIAAAVEEQGAATQEISRNVQQASVGTQEVSSNIT-- | *Bradyrhizobium sp. OK095* | Bacteria; Proteobacteria; Alphaproteobacteria; Rhizobiales; Bradyrhizobiaceae; Bradyrhizobium | methyl-accepting chemotaxis protein | 6 |
| | **WP_106945087.1** | 3E-06 | Qc.III | IQEIGGTIERLSEVAAAIAAAVEEQGAATQEISRNVQQASVGTQEVSSNIT-- | *Bradyrhizobium sp. MOS002* | Bacteria; Proteobacteria; Alphaproteobacteria; Rhizobiales; Bradyrhizobiaceae; Bradyrhizobium | methyl-accepting chemotaxis protein | 6 |
| | **WP_109331087.1** | 3E-06 | Qc.III | --GVGSIIGQINDIATAIAAAVEEQNAATLEIARNVQQAASGTQQVTSNID-- | *Azospirillum sp. CFH 70021* | Bacteria; Proteobacteria; Alphaproteobacteria; Rhodospirillales; Rhodospirillaceae; Azospirillum | methyl-accepting chemotaxis protein | 6 |
|  | **WP_111070341.1** | 3E-06 | Qa.III | IRSIAGTIREVNEIAATIAAAVEQQSSATREIARNVGEAADST---------- | *Azospirillum sp. M2T2B2* | Bacteria; Proteobacteria; Alphaproteobacteria; Rhodospirillales; Rhodospirillaceae; Azospirillum | HAMP domain-containing protein | 6 |
|  | **ALJ38336.1** | 3.1E-06 | Qa.III | IQDIAGVIGEVNEIATAIAAAVEEQGAATQEITRNVQQAAVGTQ--------- | *Azospirillum brasilense* | Bacteria; Proteobacteria; Alphaproteobacteria; Rhodospirillales; Rhodospirillaceae; Azospirillum | hypothetical protein AMK58_22810 (plasmid) | 6 |
|  | **OLE32715.1** | 3.1E-06 | Qa.IV | IKEISSTIEKLSEISSAIAAAVEEQGAATQEISRNVQQAAQ------------ | *Alphaproteobacteria bacterium 13_1_20CM_3_64_12* | Bacteria; Proteobacteria; Alphaproteobacteria | methyl-accepting chemotaxis protein | 6 |
|  | **WP_011661501.1** | 3.1E-06 | Qa.IV | -KQISATIGRMSEISSTIASAVEEQGAATQEISRNVQQAAQ------------ | *Rhodopseudomonas palustris* | Bacteria; Proteobacteria; Alphaproteobacteria; Rhizobiales; Bradyrhizobiaceae; Rhodopseudomonas | methyl-accepting chemotaxis protein | 6 |
| | **WP_018645912.1** | 3.1E-06 | Qc.III | -RSIGSTIQRVNEIAAAIAAAVEQQGAATREIARNVQQASQGTNEVSRHIS-- | *Bradyrhizobium japonicum* | Bacteria; Proteobacteria; Alphaproteobacteria; Rhizobiales; Bradyrhizobiaceae; Bradyrhizobium | HAMP domain-containing protein | 6 |
| | **WP_019646449.1** | 3.1E-06 | Qc.III | -DEVSGTIGAIEQIAATITESVEQQSAATREMVEHITVASQRTRTVVERLE-- | *Novispirillum itersonii* | Bacteria; Proteobacteria; Alphaproteobacteria; Rhodospirillales; Rhodospirillaceae; Novispirillum | methyl-accepting chemotaxis protein | 6 |
| | **WP_027546774.1** | 3.1E-06 | Qc.III | -RSIGSTIQRVNEIAAAIAAAVEQQGAATREIARNVQQASQGTNEVSRHIS-- | *Bradyrhizobium sp. WSM2254* | Bacteria; Proteobacteria; Alphaproteobacteria; Rhizobiales; Bradyrhizobiaceae; Bradyrhizobium | HAMP domain-containing protein | 6 |
|  | **WP_031337856.1** | 3.1E-06 | Qa.IV | -REIGATISRMSEIASTIASAVEEQGAATQEISRNVQQAAQ------------ | *Rhodopseudomonas sp. B29* | Bacteria; Proteobacteria; Alphaproteobacteria; Rhizobiales; Bradyrhizobiaceae; Rhodopseudomonas | HAMP domain-containing protein | 6 |
|  | **WP_051140681.1** | 3.1E-06 | Qa.III | IQDIAGVIGEVNEIATAIAAAVEEQGAATQEITRNVQQAAVGTQ--------- | *Azospirillum brasilense* | Bacteria; Proteobacteria; Alphaproteobacteria; Rhodospirillales; Rhodospirillaceae; Azospirillum | methyl-accepting chemotaxis protein | 6 |
|  | **WP_056641367.1** | 3.1E-06 | Qc.III | -QQVAASANELATGANEISEDVSEMSESITEIEEKVSDYASSTKEMNTR---- | *Paenibacillus sp. Soil522* | Bacteria; Firmicutes; Bacilli; Bacillales; Paenibacillaceae; Paenibacillus | methyl-accepting chemotaxis protein | 6 |

| | | | | | | | | |
|---|---|---|---|---|---|---|---|---|
| | **WP_092192042.1** | 3.1E-06 | Qc.III | -RSIGSTIQRVNEIAAAIAAAVEQQGAATREIARNVQQASQGTNEVSRHIS-- | *Bradyrhizobium sp. cf659* | Bacteria; Proteobacteria; Alphaproteobacteria; Rhizobiales; Bradyrhizobiaceae; Bradyrhizobium | HAMP domain-containing protein | 6 |
|  | **WP_102692293.1** | 3.1E-06 | Qa.IV | -IKVEKSITEVKENIEGIAKEIDKVTESINRVSENVEKTNDKI---------- | *Rummeliibacillus pycnus* | Bacteria; Firmicutes; Bacilli; Bacillales; Planococcaceae; Rummeliibacillus | PAS domain S-box protein | 6 |
|  | **WP_109583876.1** | 3.1E-06 | Qb.I | LQRTKEQLQATVEHSETSTEELKASNEELQAINEELRSATEELETS------- | *Cupriavidus plantarum* | Bacteria; Proteobacteria; Betaproteobacteria; Burkholderiales; Burkholderiaceae; Cupriavidus | PAS domain S-box protein | 6 |
|  | **WP_110769791.1** | 3.1E-06 | Qc.III | -QQVAASANELATGANEISEDVSEMSESITEIEEKVSDYASSTKEMNTR---- | *Paenibacillus sp. OV191* | Bacteria; Firmicutes; Bacilli; Bacillales; Paenibacillaceae; Paenibacillus | methyl-accepting chemotaxis protein | 6 |
| | **WP_111388223.1** | 3.1E-06 | Qc.III | IKEIGGTIGRIAEIASTIAAAVEEQGAATGEIARNVQQASQGTTEVASNI--- | *Rhodoplanes piscinae* | Bacteria; Proteobacteria; Alphaproteobacteria; Rhizobiales; Hyphomicrobiaceae; Rhodoplanes | methyl-accepting chemotaxis protein | 6 |
|  | **AWJ88502.1** | 3.2E-06 | Qa.III | IQGIGETIRSINEISTTIASAVQEQAAATQEISNNVAQAAR------------ | *Azospirillum brasilense Sp245* | Bacteria; Proteobacteria; Alphaproteobacteria; Rhodospirillales; Rhodospirillaceae; Azospirillum | chemotaxis protein | 6 |
|  | **OGR16624.1** | 3.2E-06 | Qa.III | IEKISKIIEEINEMINTMATAVEEQSAATAEIANNITQAS------------- | *Desulfobacterales bacterium GWB2_56_26* | Bacteria; Proteobacteria; Deltaproteobacteria; Desulfobacterales | hypothetical protein A2X81_01190 | 6 |
|  | **OPH18690.1** | 3.2E-06 | Qa.III | IQDIAGVIGEVNEIATAIAAAVEEQGAATQEITRNVQQAAVGTQ--------- | *Azospirillum brasilense* | Bacteria; Proteobacteria; Alphaproteobacteria; Rhodospirillales; Rhodospirillaceae; Azospirillum | hypothetical protein FE88_24320 | 6 |
|  | **OPH19875.1** | 3.2E-06 | Qa.III | IQGIGETIRSINEISTTIASAVQEQAAATQEISNNVAQAAR------------ | *Azospirillum brasilense* | Bacteria; Proteobacteria; Alphaproteobacteria; Rhodospirillales; Rhodospirillaceae; Azospirillum | hypothetical protein FE88_17340 | 6 |
|  | **PWC70856.1** | 3.2E-06 | Qa.III | IQGIGETIRSINEISTTIASAVQEQAAATQEISNNVAQAAR------------ | *Azospirillum sp. TSH58* | Bacteria; Proteobacteria; Alphaproteobacteria; Rhodospirillales; Rhodospirillaceae; Azospirillum | hypothetical protein TSH58_13195 | 6 |
|  | **RCK05218.1** | 3.2E-06 | Qa.III | IEKITETISNINEITSAIAAAVEQQGAATAEISGNVQQAA------------- | *Thalassospira xianhensis MCCC 1A02616* | Bacteria; Proteobacteria; Alphaproteobacteria; Rhodospirillales; Rhodospirillaceae; Thalassospira | hypothetical protein TH5_15220 | 6 |
|  | **WP_011991337.1** | 3.2E-06 | Qb.I | ---TKENLQATIEEMQAANEELKSSNEELQSTNEELQSTNEELETSK------ | *Methanoregula boonei* | Archaea; Euryarchaeota; Methanomicrobia; Methanomicrobiales; Methanoregulaceae; Methanoregula | PAS domain-containing protein | 6 |
|  | **WP_038531582.1** | 3.2E-06 | Qa.III | IQGIGETIRSINEISTTIASAVQEQAAATQEISNNVAQAAR------------ | *Azospirillum brasilense* | Bacteria; Proteobacteria; Alphaproteobacteria; Rhodospirillales; Rhodospirillaceae; Azospirillum | hypothetical protein | 6 |
|  | **WP_054884496.1** | 3.2E-06 | Qb.I | LQRTRQQLQETIEQSEVSSEALTASNEEMQAINEELRSASEELETSK------ | *Pseudomonas sp. NBRC 111130* | Bacteria; Proteobacteria; Gammaproteobacteria; Pseudomonadales; Pseudomonadaceae; Pseudomonas | response regulator | 6 |
|  | **WP_056194352.1** | 3.2E-06 | Qa.IV | IRAIVSTITDLSEISMSVSSAVTEQAAVTQEIASNMQTAA------------- | *Methylobacterium sp. Leaf123* | Bacteria; Proteobacteria; Alphaproteobacteria; Rhizobiales; Methylobacteriaceae; Methylobacterium | PAS domain S-box protein | 6 |
|  | **WP_062960932.1** | 3.2E-06 | Qa.III | IEKITETISNINEITSAIAAAVEQQGAATAEISGNVQQAA------------- | *Thalassospira xiamenensis* | Bacteria; Proteobacteria; Alphaproteobacteria; Rhodospirillales; Rhodospirillaceae; Thalassospira | methyl-accepting chemotaxis protein | 6 |
|  | **WP_075215249.1** | 3.2E-06 | Qa.III | -TEISGVISQLKDISTSIASAVEQQGAATREIAQNVNQASRGTED-------- | *Mongoliimonas terrestris* | Bacteria; Proteobacteria; Alphaproteobacteria; Rhizobiales; Mongoliimonas | hypothetical protein | 6 |
|  | **WP_075652594.1** | 3.2E-06 | Qa.III | LEKITVSIQEISDMALQISNAASEQRTVTDEVSKNIQQAN------------- | *Vibrio ponticus* | Bacteria; Proteobacteria; Gammaproteobacteria; Vibrionales; Vibrionaceae; Vibrio | methyl-accepting chemotaxis protein | 6 |
|  | **WP_084813698.1** | 3.2E-06 | Qa.III | -------IDEINEIVNGIATAVDEQSKATQEISANVSQASEGIQEVN------ | *Desulfobulbus japonicus* | Bacteria; Proteobacteria; Deltaproteobacteria; Desulfobacterales; Desulfobulbaceae; Desulfobulbus | methyl-accepting chemotaxis protein | 6 |
|  | **WP_097050912.1** | 3.2E-06 | Qa.III | IEKITETISNINEITSAIAAAVEQQGAATAEISGNVQQAA------------- | *Thalassospira xiamenensis* | Bacteria; Proteobacteria; Alphaproteobacteria; Rhodospirillales; Rhodospirillaceae; Thalassospira | methyl-accepting chemotaxis protein | 6 |
|  | **WP_099814954.1** | 3.2E-06 | Qb.I | LQRTRQQLQETIEQSEVSSEALTASNEEMQAINEELRSASEELETSK------ | *Pseudomonas sp. 382* | Bacteria; Proteobacteria; Gammaproteobacteria; Pseudomonadales; Pseudomonadaceae; Pseudomonas | response regulator | 6 |
|  | **WP_103039963.1** | 3.2E-06 | Qa.III | IQGIGETIRSINEISTTIASAVQEQAAATQEISNNVAQAAR------------ | *Azospirillum brasilense* | Bacteria; Proteobacteria; Alphaproteobacteria; Rhodospirillales; Rhodospirillaceae; Azospirillum | chemotaxis protein | 6 |
|  | **WP_104675366.1** | 3.2E-06 | Qa.III | IQGIGETIRSINEISTTIASAVQEQAAATQEISNNVAQAAR------------ | *Azospirillum* | Bacteria; Proteobacteria; Alphaproteobacteria; Rhodospirillales; Rhodospirillaceae | MULTISPECIES: chemotaxis protein | 6 |
|  | **WP_109069978.1** | 3.2E-06 | Qa.III | IQGIGETIRSINEISTTIASAVQEQAAATQEISNNVAQAAR------------ | *Azospirillum sp. TSH58* | Bacteria; Proteobacteria; Alphaproteobacteria; Rhodospirillales; Rhodospirillaceae; Azospirillum | chemotaxis protein | 6 |
|  | **WP_114122502.1** | 3.2E-06 | Qa.III | IEKITETISNINEITSAIAAAVEQQGAATAEISGNVQQAA------------- | *Thalassospira xianhensis* | Bacteria; Proteobacteria; Alphaproteobacteria; Rhodospirillales; Rhodospirillaceae; Thalassospira | HAMP domain-containing protein | 6 |
|  | **AWJ90417.1** | 3.3E-06 | Qa.III | IVGIGHTIQDINQIAGGIAAGIGQQGSAIRSIVSNMDQAA------------- | *Azospirillum brasilense Sp245* | Bacteria; Proteobacteria; Alphaproteobacteria; Rhodospirillales; Rhodospirillaceae; Azospirillum | chemotaxis protein | 6 |
|  | **OZG41492.1** | 3.3E-06 | Qa.III | LEQITASINQISDMATQISSAAEEQRAVTDEVSRNIQAVK------------- | *Aeromonas sp. A35_P* | Bacteria; Proteobacteria; Gammaproteobacteria; Aeromonadales; Aeromonadaceae; Aeromonas | hypothetical protein CAK78_13155 | 6 |
|  | **PNQ97872.1** | 3.3E-06 | Qa.III | IVGIGHTIQDINQIAGGIAAGIGQQGSAIRSIVSNMDQAA------------- | *Azospirillum brasilense* | Bacteria; Proteobacteria; Alphaproteobacteria; Rhodospirillales; Rhodospirillaceae; Azospirillum | chemotaxis protein | 6 |
|  | **WP_012497025.1** | 3.3E-06 | Qa.IV | IKEIGTTIGQMSEISSAIASAVEQQGSATQEISRNVQQAA------------- | *Rhodopseudomonas palustris* | Bacteria; Proteobacteria; Alphaproteobacteria; Rhizobiales; Bradyrhizobiaceae; Rhodopseudomonas | HAMP domain-containing protein | 6 |
|  | **WP_045003306.1** | 3.3E-06 | Qa.IV | IQEISTTINIVSEISSTIAAAVEEQGAATQEIARNVQQAAQ------------ | *Bradyrhizobium sp. LTSP857* | Bacteria; Proteobacteria; Alphaproteobacteria; Rhizobiales; Bradyrhizobiaceae; Bradyrhizobium | methyl-accepting chemotaxis protein | 6 |
|  | **WP_045013268.1** | 3.3E-06 | Qa.IV | IQEISTTINIVSEISSTIAAAVEEQGAATQEIARNVQQAAQ------------ | *Bradyrhizobium sp. LTSP849* | Bacteria; Proteobacteria; Alphaproteobacteria; Rhizobiales; Bradyrhizobiaceae; Bradyrhizobium | methyl-accepting chemotaxis protein | 6 |
| | **WP_049974314.1** | 3.3E-06 | Qc.III | IEGIAAVITDINQIATGISSAVEEQSAATREIARNVDQAAKGTQDVSRSI--- | *Azospirillum sp. B4* | Bacteria; Proteobacteria; Alphaproteobacteria; Rhodospirillales; Rhodospirillaceae; Azospirillum | methyl-accepting chemotaxis protein | 6 |
|  | **WP_079285577.1** | 3.3E-06 | Qa.III | IQDIAGVIGEVNEIATAIAAAVEEQGAATQEITRNVQQAAVGTQ--------- | *Azospirillum* | Bacteria; Proteobacteria; Alphaproteobacteria; Rhodospirillales; Rhodospirillaceae | MULTISPECIES: methyl-accepting chemotaxis protein | 6 |
| | **WP_086464893.1** | 3.3E-06 | Qc.III | IQSITKIIERMHEASTAIAAAIEEQTAATREIARNVEQAAAGTDEVTRSIAKV | *Oceanibaculum nanhaiense* | Bacteria; Proteobacteria; Alphaproteobacteria; Rhodospirillales; Rhodospirillaceae; Oceanibaculum | methyl-accepting chemotaxis protein | 6 |
|  | **WP_094301411.1** | 3.3E-06 | Qa.III | IQGIGETIRSINEISTTIASAVQEQAAATQEISNNVAQAAR------------ | *Azospirillum brasilense* | Bacteria; Proteobacteria; Alphaproteobacteria; Rhodospirillales; Rhodospirillaceae; Azospirillum | hypothetical protein | 6 |
|  | **WP_094304500.1** | 3.3E-06 | Qa.III | IQDIAAVIGEVNEIATAIAAAVEEQGAATQEITRNVQQAAVGTQ--------- | *Azospirillum brasilense* | Bacteria; Proteobacteria; Alphaproteobacteria; Rhodospirillales; Rhodospirillaceae; Azospirillum | methyl-accepting chemotaxis protein | 6 |
|  | **WP_096619409.1** | 3.3E-06 | Qa.III | LEKIAGMMTAINDMNTQIATAASQQSTVTEEITENVNNIA------------- | *bacterium symbiont of Cryptopsaras couesii* | Bacteria | methyl-accepting chemotaxis protein | 6 |
|  | **WP_098736108.1** | 3.3E-06 | Qa.III | IRNIGRTITQVNEISNSIAAAVEEQTAATGEIGRNVQQAAQGTQEVSSS---- | *Azospirillum sp. B2* | Bacteria; Proteobacteria; Alphaproteobacteria; Rhodospirillales; Rhodospirillaceae; Azospirillum | HAMP domain-containing protein | 6 |
|  | **WP_100181106.1** | 3.3E-06 | Qa.III | IHGITATIEEINEITSSIAAAVVEQDAATREIARNVQEAA------------- | *Bradyrhizobium sp. TSA1* | Bacteria; Proteobacteria; Alphaproteobacteria; Rhizobiales; Bradyrhizobiaceae; Bradyrhizobium | HAMP domain-containing protein | 6 |
|  | **WP_107345363.1** | 3.3E-06 | Qa.IV | IKEIGTTIGQMSEISSAIASAVEQQGSATQEISRNVQQAA------------- | *Rhodopseudomonas palustris* | Bacteria; Proteobacteria; Alphaproteobacteria; Rhizobiales; Bradyrhizobiaceae; Rhodopseudomonas | HAMP domain-containing protein | 6 |
|  | **WP_107356391.1** | 3.3E-06 | Qa.IV | IKEIGTTIGQMSEISSAIASAVEQQGSATQEISRNVQQAA------------- | *Rhodopseudomonas palustris* | Bacteria; Proteobacteria; Alphaproteobacteria; Rhizobiales; Bradyrhizobiaceae; Rhodopseudomonas | methyl-accepting chemotaxis protein | 6 |
|  | **WP_109050271.1** | 3.3E-06 | Qa.III | IRNIGRTITQVNEISNSIAAAVEEQTAATGEIGRNVQQAAQGTQEVSSS---- | *Azospirillum sp. TSA6c* | Bacteria; Proteobacteria; Alphaproteobacteria; Rhodospirillales; Rhodospirillaceae; Azospirillum | methyl-accepting chemotaxis protein | 6 |
|  | **WP_110788794.1** | 3.3E-06 | Qa.IV | IKEIGTTIGQMSEISSAIASAVEQQGSATQEISRNVQQAA------------- | *Rhodopseudomonas palustris* | Bacteria; Proteobacteria; Alphaproteobacteria; Rhizobiales; Bradyrhizobiaceae; Rhodopseudomonas | HAMP domain-containing protein | 6 |
|  | **WP_114359256.1** | 3.3E-06 | Qa.IV | IKEIGTTIGQMSEISSAIASAVEQQGSATQEISRNVQQAA------------- | *Rhodopseudomonas pentothenatexigens* | Bacteria; Proteobacteria; Alphaproteobacteria; Rhizobiales; Bradyrhizobiaceae; Rhodopseudomonas | HAMP domain-containing protein | 6 |
| | **WP_012172367.1** | 3.4E-06 | Qc.III | IESVGQQIRSMSDVATGISAAVEEQGIATREIVRNVDQAATGTNSVTGHIAD- | *Azorhizobium caulinodans* | Bacteria; Proteobacteria; Alphaproteobacteria; Rhizobiales; Xanthobacteraceae; Azorhizobium | methyl-accepting chemotaxis protein | 6 |
| | **WP_042340920.1** | 3.4E-06 | Qc.III | IQAISSTIERLAEIAAAIAAAVEEQGAATQEISRNVQQAAHGTQQVSSNI--- | *Bradyrhizobium oligotrophicum* | Bacteria; Proteobacteria; Alphaproteobacteria; Rhizobiales; Bradyrhizobiaceae; Bradyrhizobium | HAMP domain-containing protein | 6 |
| | **BAM91137.1** | 3.5E-06 | Qc.III | IQAISSTIERLAEIAAAIAAAVEEQGAATQEISRNVQQAAHGTQQVSSNI--- | *Bradyrhizobium oligotrophicum S58* | Bacteria; Proteobacteria; Alphaproteobacteria; Rhizobiales; Bradyrhizobiaceae; Bradyrhizobium | receptor/sensory transducer | 6 |
|  | **PWB60000.1** | 3.5E-06 | Qa.IV | IKEIKATIGRISDISSTIAAAVEEQGATTQEIARNLQQAA------------- | *Bradyrhizobiaceae bacterium* | Bacteria; Proteobacteria; Alphaproteobacteria; Rhizobiales; Bradyrhizobiaceae | hypothetical protein C3F17_15555 | 6 |
|  | **WP_047188771.1** | 3.5E-06 | Qa.IV | IQEIARTINEMSQISISIAAAMEEQGAATAEIARNVQEAA------------- | *Microvirga vignae* | Bacteria; Proteobacteria; Alphaproteobacteria; Rhizobiales; Methylobacteriaceae; Microvirga | methyl-accepting chemotaxis protein | 6 |
| | **WP_092065333.1** | 3.5E-06 | Qa.IV | IKDITQSISDIDSIISTIAAAVEEQSTTTRDIAENIGSVSQHIGEVNENVNQ- | *Desulfonauticus submarinus* | Bacteria; Proteobacteria; Deltaproteobacteria; Desulfovibrionales; Desulfohalobiaceae; Desulfonauticus | methyl-accepting chemotaxis protein | 6 |
|  | **WP_097670514.1** | 3.5E-06 | Qa.IV | -RAIGSTIERLSEISSAIAAAIEEQGAATQEISRNVQQAAQ----------- | *Bradyrhizobium sp. C9* | Bacteria; Proteobacteria; Alphaproteobacteria; Rhizobiales; Bradyrhizobiaceae; Bradyrhizobium | methyl-accepting chemotaxis protein | 6 |
|  | **CAE29039.1** | 3.6E-06 | Qa.IV | IKEIGTTIGQMSEISSAIASAVEQQGSATQEISRNVQQAA------------- | *Rhodopseudomonas palustris CGA009* | Bacteria; Proteobacteria; Alphaproteobacteria; Rhizobiales; Bradyrhizobiaceae; Rhodopseudomonas | methyl-accepting chemotaxis receptor/ sensory transducer | 6 |
|  | **OYW58975.1** | 3.6E-06 | Qa.III.b | IERIASSIEQLNQVSTAIATVVTEQGAATGDIRANVSQAA------------- | *Bosea sp. 12-68-7* | Bacteria; Proteobacteria; Alphaproteobacteria; Rhizobiales; Bradyrhizobiaceae; Bosea | hypothetical protein B7Z40_22270 | 6 |
| | **PWE82251.1** | 3.6E-06 | Qc.III | IQEIGGTIERLSEVSAAIAAAVEEQGAATQEISRNVQQASLGTQEVSSNIT-- | *Bradyrhizobium sp. SUTN9-2* | Bacteria; Proteobacteria; Alphaproteobacteria; Rhizobiales; Bradyrhizobiaceae; Bradyrhizobium | chemotaxis protein partial | 6 |
|  | **SOE01619.1** | 3.6E-06 | Qa.IV | -REIAATIERISDITEGIAEAVKQQGAATTEIARNIQEAARGTADA------- | *Caenispirillum bisanense* | Bacteria; Proteobacteria; Alphaproteobacteria; Rhodospirillales; Rhodospirillaceae; Caenispirillum | methyl-accepting chemotaxis protein | 6 |
|  | **WP_013501970.1** | 3.6E-06 | Qa.IV | IRDISSTIEKLSEIASTIASAVEEQGAATQEISRNVQQAAQ------------ | *Rhodopseudomonas palustris* | Bacteria; Proteobacteria; Alphaproteobacteria; Rhizobiales; Bradyrhizobiaceae; Rhodopseudomonas | HAMP domain-containing protein | 6 |
| | **WP_114096178.1** | 3.6E-06 | Qc.III | IEKVTQTISSISEIANGISSAVNEQNSAVNEIANSATSASSGSTEVVANIDE- | *Thalassospira profundimaris* | Bacteria; Proteobacteria; Alphaproteobacteria; Rhodospirillales; Rhodospirillaceae; Thalassospira | HAMP domain-containing protein | 6 |
|  | **KZK90546.1** | 3.7E-06 | Qa.IV | IQEITSSIRQVNEFSSNIAASVEQQGAATAEITENVHQAA------------- | *Pseudovibrio sp. Ad46* | Bacteria; Proteobacteria; Alphaproteobacteria; Rhodobacterales; Rhodobacteraceae; Pseudovibrio | Methyl-accepting chemotaxis protein PctC | 6 |
|  | **KZK92732.1** | 3.7E-06 | Qa.IV | IQEITSSIRQVNEFSSNIAASVEQQGAATAEITENVHQAA------------- | *Pseudovibrio sp. Ad5* | Bacteria; Proteobacteria; Alphaproteobacteria; Rhodobacterales; Rhodobacteraceae; Pseudovibrio | Methyl-accepting chemotaxis protein PctC | 6 |
|  | **KZL25799.1** | 3.7E-06 | Qa.IV | IQEITSSIRQVNEFSSNIAASVEQQGAATAEITENVHQAA------------- | *Pseudovibrio sp. Ad37* | Bacteria; Proteobacteria; Alphaproteobacteria; Rhodobacterales; Rhodobacteraceae; Pseudovibrio | Methyl-accepting chemotaxis protein PctC | 6 |
|  | **PIE61018.1** | 3.7E-06 | Qa.III | MENINTVINNVNTIVTTIASAVDEQATSTDKIASNVAQVA------------- | *Desulfobacterales bacterium* | Bacteria; Proteobacteria; Deltaproteobacteria; Desulfobacterales | chemotaxis protein partial | 6 |

| | | | | | | | | |
|---|---|---|---|---|---|---|---|---|
|  | **SDG67245.1** | 3.7E-06 | Qa.III | IQSICATIREVDEISAAIAAAVGQQGAATQEIAQNVQQAA------------- | *Bradyrhizobium sp. Rc2d* | Bacteria; Proteobacteria; Alphaproteobacteria; Rhizobiales; Bradyrhizobiaceae; Bradyrhizobium | methyl-accepting chemotaxis sensory transducer | 6 |
| | **WP_012172237.1** | 3.7E-06 | Qc.III | IESVGQQIRSMSDVATGISAAVEEQGIATREIVRNVDQAATGTNSVTGHITD- | *Azorhizobium caulinodans* | Bacteria; Proteobacteria; Alphaproteobacteria; Rhizobiales; Xanthobacteraceae; Azorhizobium | HAMP domain-containing protein | 6 |
| | **WP_047703544.1** | 3.7E-06 | Qa.III | --RICAEVAQISGMSLQMAAAIEQQGQVAEQINQQITEIAGFTERTSQQAGR- | *Pseudomonas mediterranea* | Bacteria; Proteobacteria; Gammaproteobacteria; Pseudomonadales; Pseudomonadaceae; Pseudomonas | PAS domain S-box protein | 6 |
| | **WP_061240771.1** | 3.7E-06 | Qa.III | --RICAEVAQISGMSLQMAAAIEQQGQVAEQINQQITEIAGFTERTSQQAGR- | *Pseudomonas composti* | Bacteria; Proteobacteria; Gammaproteobacteria; Pseudomonadales; Pseudomonadaceae; Pseudomonas | PAS domain S-box protein | 6 |
| | **EKQ57510.1** | 3.8E-06 | Qb.I | ---INKLVTGTVKQSHESAEYINESSYLISETVNTFDSIFGKINSTNDLIQS- | *Clostridium sp. Maddingley MBC34-26* | Bacteria; Firmicutes; Clostridia; Clostridiales; Clostridiaceae; Clostridium | methyl-accepting chemotaxis protein | 6 |
| | **GAP34295.1** | 3.8E-06 | R.IV | IDQVNTAVTQLDDATQQNASLVQQSSEAAASLQHQAQRLAEAIGVFKTAT--- | *Ideonella sakaiensis* | Bacteria; Proteobacteria; Betaproteobacteria; Burkholderiales; Ideonella | methyl-accepting chemotaxis protein I | 6 |
|  | **OJX79430.1** | 3.8E-06 | Qc.III | MDDIIKVVGDIGQMATEMTHAIEEQGNATREIAQGAEQASSATRTVT------ | *Magnetospirillum sp. 64-120* | Bacteria; Proteobacteria; Alphaproteobacteria; Rhodospirillales; Rhodospirillaceae; Magnetospirillum | hypothetical protein BGO92_13210 | 6 |
|  | **PVO74066.1** | 3.8E-06 | Qc.III | IQAIGGIIGEVNEVATAIAAAVQEQGAATQEITRSTQYAAQGTKNVS------ | *Salmonella enterica subsp. enterica serovar Anatum* | Bacteria; Proteobacteria; Gammaproteobacteria; Enterobacterales; Enterobacteriaceae; Salmonella | methyl-accepting chemotaxis protein partial | 6 |
|  | **WP_050626164.1** | 3.8E-06 | Qa.IV | -RAIGSTIERLSEISSAIAAAIEEQGAATQEISRNVQQAAQ----------- | *Bradyrhizobium* | Bacteria; Proteobacteria; Alphaproteobacteria; Rhizobiales; Bradyrhizobiaceae | MULTISPECIES: methyl-accepting chemotaxis protein | 6 |
|  | **WP_063292932.1** | 3.8E-06 | Qa.IV | IQEITSSIRQVNEFSSNIAASVEQQGAATAEITENVHQAA------------- | *Pseudovibrio* | Bacteria; Proteobacteria; Alphaproteobacteria; Rhodobacterales; Rhodobacteraceae | MULTISPECIES: methyl-accepting chemotaxis protein | 6 |
|  | **WP_063296103.1** | 3.8E-06 | Qa.IV | IQEITSSIRQVNEFSSNIAASVEQQGAATAEITENVHQAA------------- | *Pseudovibrio sp. Ad37* | Bacteria; Proteobacteria; Alphaproteobacteria; Rhodobacterales; Rhodobacteraceae; Pseudovibrio | methyl-accepting chemotaxis protein | 6 |
|  | **WP_063306789.1** | 3.8E-06 | Qa.IV | IQEITSSIRQVNEFSSNIAASVEQQGAATAEITENVHQAA------------- | *Pseudovibrio* | Bacteria; Proteobacteria; Alphaproteobacteria; Rhodobacterales; Rhodobacteraceae | MULTISPECIES: methyl-accepting chemotaxis protein | 6 |
|  | **WP_085997598.1** | 3.8E-06 | Qa.III | IQSIGATIAEINEIASSIASAVEEQSAATGEIARNVEQAAS------------ | *Polymorphum gilvum* | Bacteria; Proteobacteria; Alphaproteobacteria; Polymorphum | methyl-accepting chemotaxis protein | 6 |
| | **WP_109067753.1** | 3.8E-06 | Qc.III | --AIGGIISQINDISTAIAAAIEEQSAATNEISRNVQEAAAGTHQVSGSIT-- | *Azospirillum sp. TSH58* | Bacteria; Proteobacteria; Alphaproteobacteria; Rhodospirillales; Rhodospirillaceae; Azospirillum | methyl-accepting chemotaxis protein | 6 |
|  | **PWC80370.1** | 3.9E-06 | Qa.III | IRDIATVIGEVNEIATAIAAAVEEQGAATQEITRNVQQAAVGTQ--------- | *Azospirillum sp. TSH58* | Bacteria; Proteobacteria; Alphaproteobacteria; Rhodospirillales; Rhodospirillaceae; Azospirillum | hypothetical protein TSH58_02250 | 6 |
|  | **SFK21399.1** | 3.9E-06 | Qa.IV | ------TIVQISQISGEIAAAVDQQSAATREIAANLQQASDRTKDVNRS---- | *Bradyrhizobium sp. Gha* | Bacteria; Proteobacteria; Alphaproteobacteria; Rhizobiales; Bradyrhizobiaceae; Bradyrhizobium | methyl-accepting chemotaxis protein | 6 |
|  | **WP_073956493.1** | 3.9E-06 | Qa.III | IQGIGTTISELNEIAASIASAVEQQGAATGEITRNIQQAS------------- | *Thalassospira sp. TSL5-1* | Bacteria; Proteobacteria; Alphaproteobacteria; Rhodospirillales; Rhodospirillaceae; Thalassospira | methyl-accepting chemotaxis protein | 6 |
| | **WP_087456038.1** | 3.9E-06 | Qc.III | -DRVVEIAEQFSSQSQEVNAAVEEQTASMEEIHASAEALSEMSEKLQRLVTRF | *Tumebacillus avium* | Bacteria; Firmicutes; Bacilli; Bacillales; Alicyclobacillaceae; Tumebacillus | HAMP domain-containing protein | 6 |
|  | **WP_092811301.1** | 3.9E-06 | Qa.III | IESITGTIKQIDEISSTIASSVEEQGSATGEIAQNVQQAAQGTQE-------- | *Afifella marina* | Bacteria; Proteobacteria; Alphaproteobacteria; Rhizobiales; Rhodobiaceae; Afifella | methyl-accepting chemotaxis protein | 6 |
|  | **WP_101267985.1** | 3.9E-06 | Qc.III.c | -GEIGRIIEHINEIATTISAAVEQQGSATNEISRNVRGAAER----------- | *Thalassospira marina* | Bacteria; Proteobacteria; Alphaproteobacteria; Rhodospirillales; Rhodospirillaceae; Thalassospira | HAMP domain-containing protein | 6 |
|  | **WP_101284379.1** | 3.9E-06 | Qc.III.c | -GEIGRIIEHINEIATTISAAVEQQGSATNEISRNVRGAAER----------- | *Thalassospira marina* | Bacteria; Proteobacteria; Alphaproteobacteria; Rhodospirillales; Rhodospirillaceae; Thalassospira | HAMP domain-containing protein | 6 |
| | **WP_018460439.1** | 4E-06 | Qc.III | IKEISGTIGRISEISSAIAAAVEEQGMATQEISRNVQQAAQGTHEVASNIT-- | *Bradyrhizobium sp. WSM4349* | Bacteria; Proteobacteria; Alphaproteobacteria; Rhizobiales; Bradyrhizobiaceae; Bradyrhizobium | methyl-accepting chemotaxis protein | 6 |
|  | **WP_019613035.1** | 4E-06 | Qc.III.c | LDSMAVSVVKINDLGVQIATAAEQQSSATDEINRNMSAIQGMVTQL------- | *Psychromonas ossibalaenae* | Bacteria; Proteobacteria; Gammaproteobacteria; Alteromonadales; Psychromonadaceae; Psychromonas | methyl-accepting chemotaxis protein | 6 |
| | **WP_085350185.1** | 4E-06 | Qc.III | IKEISGTIGRISEISSAIAAAVEEQGMATQEISRNVQQAAQGTHEVASNIT-- | *Bradyrhizobium canariense* | Bacteria; Proteobacteria; Alphaproteobacteria; Rhizobiales; Bradyrhizobiaceae; Bradyrhizobium | methyl-accepting chemotaxis protein | 6 |
| | **WP_085360752.1** | 4E-06 | Qc.III | IKEISGTIGRISEISSAIAAAVEEQGMATQEISRNVQQAAQGTHEVASNIT-- | *Bradyrhizobium canariense* | Bacteria; Proteobacteria; Alphaproteobacteria; Rhizobiales; Bradyrhizobiaceae; Bradyrhizobium | methyl-accepting chemotaxis protein | 6 |
| | **WP_085383588.1** | 4E-06 | Qc.III | IKEISGTIGRISEISSAIAAAVEEQGMATQEISRNVQQAAQGTHEVASNIT-- | *Bradyrhizobium canariense* | Bacteria; Proteobacteria; Alphaproteobacteria; Rhizobiales; Bradyrhizobiaceae; Bradyrhizobium | methyl-accepting chemotaxis protein | 6 |
|  | **WP_106993209.1** | 4E-06 | R.IV | IQEINRAVTHLDSMVQQNAELVVESAAAASALQGQAGELAETAGHFR------ | *Enterobacter sp. FS01* | Bacteria; Proteobacteria; Gammaproteobacteria; Enterobacterales; Enterobacteriaceae; Enterobacter | methyl-accepting chemotaxis protein | 6 |
|  | **WP_107988083.1** | 4E-06 | Qa.IV | IQQIAETMEEVNSYTSAIAAAVEQQGSATTEISRSVQDATRDT---------- | *Breoghania corrubedonensis* | Bacteria; Proteobacteria; Alphaproteobacteria; Rhizobiales; Cohaesibacteraceae; Breoghania | methyl-accepting chemotaxis protein | 6 |
|  | **WP_109067977.1** | 4E-06 | Qa.III | IRDIATVIGEVNEIATAIAAAVEEQGAATQEITRNVQQAAVGTQ--------- | *Azospirillum sp. TSH58* | Bacteria; Proteobacteria; Alphaproteobacteria; Rhodospirillales; Rhodospirillaceae; Azospirillum | methyl-accepting chemotaxis protein | 6 |
|  | **OJW27437.1** | 4.1E-06 | Qa.III | IQGITGTIEEVSTIATTIASAVEEQGAATNEIARNVQQTAQ------------ | *Rhodospirillales bacterium 69-11* | Bacteria; Proteobacteria; Alphaproteobacteria; Rhodospirillales; unclassified Rhodospirillales | methyl-accepting chemotaxis protein | 6 |
|  | **WP_041756223.1** | 4.1E-06 | Qc.III | IKEISGTIERLAEIASTIAAAVEEQGAATQEISRNVQQAS------------- | *Bradyrhizobium sp. ORS 278* | Bacteria; Proteobacteria; Alphaproteobacteria; Rhizobiales; Bradyrhizobiaceae; Bradyrhizobium | methyl-accepting chemotaxis protein | 6 |
|  | **WP_077607872.1** | 4.1E-06 | Qa.III | -NAINEQIVQISDMNTQVATATEEQSTVVNDINQNIEEINHSSQQTA------ | *Salinivibrio sp. IB282* | Bacteria; Proteobacteria; Gammaproteobacteria; Vibrionales; Vibrionaceae; Salinivibrio | methyl-accepting chemotaxis protein | 6 |
| | **WP_079542961.1** | 4.1E-06 | Qc.III | IEEISGTIEKLSEISSTIAAAVEEQGAATQEISRNVQQAAQGTQRVSSNITD- | *Bradyrhizobium lablabi* | Bacteria; Proteobacteria; Alphaproteobacteria; Rhizobiales; Bradyrhizobiaceae; Bradyrhizobium | HAMP domain-containing protein | 6 |
|  | **CDF93959.1** | 4.2E-06 | Qb.I | LQRTKLHLQDTIEQSEVSNEELKASNEEMQAINEELRSATEELETS------- | *Pseudomonas sp. SHC52* | Bacteria; Proteobacteria; Gammaproteobacteria; Pseudomonadales; Pseudomonadaceae; Pseudomonas | Chemotaxis protein methyltransferase CheR | 6 |
|  | **KIZ45586.1** | 4.2E-06 | Qa.IV | IREITDTIRQMSEISATIAAAVEQQGAATAAISRDVQQVAGDTRN-------- | *Rhodopseudomonas palustris* | Bacteria; Proteobacteria; Alphaproteobacteria; Rhizobiales; Bradyrhizobiaceae; Rhodopseudomonas | chemotaxis protein | 6 |
|  | **OJU20488.1** | 4.2E-06 | Qa.IV | IQAIDGTIGEINQLSAEVAEAISQQGAATQDIASNLQQAAN------------ | *Afipia sp. 62-7* | Bacteria; Proteobacteria; Alphaproteobacteria; Rhizobiales; Bradyrhizobiaceae; Afipia | chemotaxis protein | 6 |
|  | **WP_008969108.1** | 4.2E-06 | Qa.IV | IRDISSTIEKLSEIASTIAAAVEEQGAATQEISRNVQQAA------------- | *Bradyrhizobium sp. STM 3843* | Bacteria; Proteobacteria; Alphaproteobacteria; Rhizobiales; Bradyrhizobiaceae; Bradyrhizobium | methyl-accepting chemotaxis protein | 6 |
|  | **WP_041021738.1** | 4.2E-06 | Qb.I | LQRTKLHLQDTIEQSEVSNEELKASNEEMQAINEELRSATEELETS------- | *Pseudomonas sp. SHC52* | Bacteria; Proteobacteria; Gammaproteobacteria; Pseudomonadales; Pseudomonadaceae; Pseudomonas | PAS domain S-box protein | 6 |
|  | **WP_057852575.1** | 4.2E-06 | Qa.III | IQAIGGTIAEIDEISNEVAAAVDQQGAATREIAGNVQQAAN------------ | *Bradyrhizobium valentinum* | Bacteria; Proteobacteria; Alphaproteobacteria; Rhizobiales; Bradyrhizobiaceae; Bradyrhizobium | chemotaxis protein | 6 |
| | **WP_076955655.1** | 4.2E-06 | Qc.III | --GIGSTVERTNEISTAIAAAVEQQGAATQEIARSASQVAEATGTVSQRIGR- | *Roseomonas deserti* | Bacteria; Proteobacteria; Alphaproteobacteria; Rhodospirillales; Acetobacteraceae; Roseomonas | methyl-accepting chemotaxis protein | 6 |
| | **WP_082130372.1** | 4.2E-06 | Qc.III | --GISKTIDSIHEVATAIASAVEEQGAATAEITKSVQQAANGTDQVSSNIT-- | *Kiloniella spongiae* | Bacteria; Proteobacteria; Alphaproteobacteria; Kiloniellales; Kiloniellaceae; Kiloniella | hypothetical protein | 6 |
|  | **WP_108512320.1** | 4.2E-06 | Qa.III | IQAIGGTIAEIDEISNEVAAAVDQQGAATREIAGNVQQAAN------------ | *Bradyrhizobium algeriense* | Bacteria; Proteobacteria; Alphaproteobacteria; Rhizobiales; Bradyrhizobiaceae; Bradyrhizobium | chemotaxis protein | 6 |
| | **WP_111357234.1** | 4.2E-06 | Qc.III | IKEIGGTIGRIAEIASSIAAAIEEQGAATQEIARNVQQAARGTEEVARNIG-- | *Rhodoplanes elegans* | Bacteria; Proteobacteria; Alphaproteobacteria; Rhizobiales; Hyphomicrobiaceae; Rhodoplanes | methyl-accepting chemotaxis protein | 6 |
|  | **WP_114123247.1** | 4.2E-06 | Qa.III | IDGITSIIANLNEIASEVAAAVQEQGKATSEISHSVQQAVQSTQ--------- | *Thalassospira xianhensis* | Bacteria; Proteobacteria; Alphaproteobacteria; Rhodospirillales; Rhodospirillaceae; Thalassospira | HAMP domain-containing protein | 6 |
|  | **CAL75451.1** | 4.3E-06 | Qc.III | IKEISGTIERLAEIASTIAAAVEEQGAATQEISRNVQQAS------------- | *Bradyrhizobium sp. ORS 278* | Bacteria; Proteobacteria; Alphaproteobacteria; Rhizobiales; Bradyrhizobiaceae; Bradyrhizobium | Methyl-accepting chemotaxis protein | 6 |
|  | **EXJ44942.1** | 4.3E-06 | Qa.III | LDEINAAVALISDMATQIATAAEEQTHVTNEITQNVTSIK------------- | *Vibrio parahaemolyticus VPTS-2010_2* | Bacteria; Proteobacteria; Gammaproteobacteria; Vibrionales; Vibrionaceae; Vibrio | methyl-accepting chemotaxis (MCP) signaling domain protein | 6 |
| | **PWB66502.1** | 4.3E-06 | Qc.III | --EIGGTITRISEIASAIAAAVEEQGAATQEIARNVGEAAKGTNQVAESI--- | *Bradyrhizobiaceae bacterium* | Bacteria; Proteobacteria; Alphaproteobacteria; Rhizobiales; Bradyrhizobiaceae | methyl-accepting chemotaxis protein | 6 |
| | **RCK33643.1** | 4.3E-06 | Qc.III | IESIGKTIGEISEIAATIASAVEEQGAATEEINRNVQEASTGTTDVSSNI--- | *Thalassospira profundimaris* | Bacteria; Proteobacteria; Alphaproteobacteria; Rhodospirillales; Rhodospirillaceae; Thalassospira | hypothetical protein TH19_17195 | 6 |
|  | **WP_026511823.1** | 4.3E-06 | SNAP.b | -NDLMKDAGESVQVVQLLIDNIEEQGKQLDTTRDSMDKMQQ------------ | *Butyrivibrio sp. LC3010* | Bacteria; Firmicutes; Clostridia; Clostridiales; Lachnospiraceae; Butyrivibrio | methyl-accepting chemotaxis protein | 6 |
|  | **WP_040567222.1** | 4.3E-06 | Qa.IV | IVSIVERIEEINQIATTIASAVEEQSAATNEIARNVQQAA------------- | *Phaeospirillum molischianum* | Bacteria; Proteobacteria; Alphaproteobacteria; Rhodospirillales; Rhodospirillaceae; Phaeospirillum | HAMP domain-containing protein | 6 |
|  | **WP_044408098.1** | 4.3E-06 | Qa.IV | IREITDTIRQMSEISATIAAAVEQQGAATAAISRDVQQVAGDTRN-------- | *Rhodopseudomonas palustris* | Bacteria; Proteobacteria; Alphaproteobacteria; Rhizobiales; Bradyrhizobiaceae; Rhodopseudomonas | methyl-accepting chemotaxis protein | 6 |
|  | **WP_101081039.1** | 4.3E-06 | Qc.III.c | LDSMTASVTEINDLGIQIATAAEEQSSVTEEINRNMTAIQDMVAQLT------ | *Psychromonas sp. Urea-02u-13* | Bacteria; Proteobacteria; Gammaproteobacteria; Alteromonadales; Psychromonadaceae; Psychromonas | methyl-accepting chemotaxis protein | 6 |
| | **KJS35697.1** | 4.4E-06 | Qc.III | --GISKIIGRISEISVSIASAIEEQSAATDEIARNVQQASAGSQEVSYNIQ-- | *Rhodospirillaceae bacterium BRH_c57* | Bacteria; Proteobacteria; Alphaproteobacteria; Rhodospirillales; Rhodospirillaceae | hypothetical protein VR70_15510 | 6 |
|  | **OCR89188.1** | 4.4E-06 | Qa.IV | --EIEANVNILSQGINEMSQSINEQTEAINQINEAVATVDEQTKQ-NV----- | *Campylobacter fetus subsp. testudinum* | Bacteria; Proteobacteria; Epsilonproteobacteria; Campylobacterales; Campylobacteraceae; Campylobacter | chemotaxis protein partial | 6 |
|  | **OCR89985.1** | 4.4E-06 | Qa.IV | --EIEANVNILSQGINEMSQSINEQTEAINQINEAVATVDEQTKQ-NV----- | *Campylobacter fetus subsp. testudinum* | Bacteria; Proteobacteria; Epsilonproteobacteria; Campylobacterales; Campylobacteraceae; Campylobacter | chemotaxis protein partial | 6 |
| | **PHS28120.1** | 4.4E-06 | Qc.III | IEGISKTISDINEFSTAISAAMEEQTAVTGEMSSNMQTAASGVDNITRGI--- | *Robiginitomaculum sp.* | Bacteria; Proteobacteria; Alphaproteobacteria; Rhodobacterales; Hyphomonadaceae; Robiginitomaculum | chemotaxis protein | 6 |
| | **WP_002975744.1** | 4.4E-06 | Qc.II | --DLFKTLKSIREITTDFNNQIESQDKSIHELTEFVSSFSESIQSSVDNIGK- | *Leptospira terpstrae* | Bacteria; Spirochaetes; Leptospirales; Leptospiraceae; Leptospira | methyl-accepting chemotaxis protein | 6 |
|  | **WP_024341498.1** | 4.4E-06 | Qa.IV | IREISVTIERLSEISSAVAAAVEQQGAATQEISRNVQRAAD------------ | *Bradyrhizobium japonicum* | Bacteria; Proteobacteria; Alphaproteobacteria; Rhizobiales; Bradyrhizobiaceae; Bradyrhizobium | methyl-accepting chemotaxis protein | 6 |
|  | **WP_038156125.1** | 4.4E-06 | Qc.III.c | ------SIVRINDLGLEIATAASQQNAVTEEVSRNMTTIQSMVDELTSNG--- | *Vibrio sp. B183* | Bacteria; Proteobacteria; Gammaproteobacteria; Vibrionales; Vibrionaceae; Vibrio | methyl-accepting chemotaxis protein | 6 |
|  | **WP_038176013.1** | 4.4E-06 | Qc.III.c | ------SIVRINDLGLEIATAASQQNAVTEEVSRNMTTIQSMVDELTSNG--- | *Vibrio coralliilyticus* | Bacteria; Proteobacteria; Gammaproteobacteria; Vibrionales; Vibrionaceae; Vibrio | methyl-accepting chemotaxis protein | 6 |

| | | | | | | | |
|---|---|---|---|---|---|---|---|
| WP_039952572.1 | 4.4E-06 | Qc.III.c | ------SIVRINDLGLEIATAASQQNAVTEEVSRNMTTIQSMVDELTSNG--- | *Vibrio coralliilyticus* | Bacteria; Proteobacteria; Gammaproteobacteria; Vibrionales; Vibrionaceae; Vibrio | methyl-accepting chemotaxis protein | 6 |
| WP_051100023.1 | 4.4E-06 | Qb.I | LQRTRLLLQETIEQSEVSSEELTASNEEMQAINEELRSASEELETSK------ | *Pseudomonas sp. 10-1B* | Bacteria; Proteobacteria; Gammaproteobacteria; Pseudomonadales; Pseudomonadaceae; Pseudomonas | response regulator | 6 |
| WP_098694408.1 | 4.4E-06 | Qa.IV | ----ETTITELRDVTTTFNDMVDRQREIVSGLVEQDQRAQRFVSDVAHEL--- | *Nocardia terpenica* | Bacteria; Actinobacteria; Corynebacteriales; Nocardiaceae; Nocardia | sensor histidine kinase | 6 |
| WP_099607491.1 | 4.4E-06 | Qc.III.c | ------SIVRINDLGLEIATAASQQNAVTEEVSRNMTTIQSMVDELTSNG--- | *Vibrio coralliilyticus* | Bacteria; Proteobacteria; Gammaproteobacteria; Vibrionales; Vibrionaceae; Vibrio | methyl-accepting chemotaxis protein | 6 |
| APP07696.1 | 4.5E-06 | Qa.III | -SKIREAIQQISDMATQIASAAEEQSLVTAEINANTESVRE------------ | *Vibrio harveyi* | Bacteria; Proteobacteria; Gammaproteobacteria; Vibrionales; Vibrionaceae; Vibrio | methyl-accepting chemotaxis protein | 6 |
| EEX31712.1 | 4.5E-06 | Qc.III.c | ------SIVRINDLGLEIATAASQQNAVTEEVSRNMTTIQSMVDELTSNG--- | *Vibrio coralliilyticus ATCC BAA-450* | Bacteria; Proteobacteria; Gammaproteobacteria; Vibrionales; Vibrionaceae; Vibrio | hypothetical methyl-accepting chemotaxis protein | 6 |
| EEZ86627.1 | 4.5E-06 | Qa.III | -SKIREAIQQISDMATQIASAAEEQSLVTAEINANTESVRE------------ | *Vibrio harveyi 1DA3* | Bacteria; Proteobacteria; Gammaproteobacteria; Vibrionales; Vibrionaceae; Vibrio | methyl-accepting chemotaxis protein | 6 |
| EKD25751.1 | 4.5E-06 | Qa.III | IEKITKIIDEVNNITQTIASSVEEQSATTNEIASNIETTSSAARESSK----- | *uncultured bacterium* | Bacteria; environmental samples | hypothetical protein ACD_79C01527G0009 | 6 |
| KFI13854.1 | 4.5E-06 | Qc.III.c | ------SIVRINDLGLEIATAASQQNAVTEEVSRNMTTIQSMVDELTSNG--- | *Vibrio sp. B183* | Bacteria; Proteobacteria; Gammaproteobacteria; Vibrionales; Vibrionaceae; Vibrio | chemotaxis protein | 6 |
| PIE58826.1 | 4.5E-06 | Qa.III | MEQINTVINNVNTIVTTIASAVEEQALSTDEIASNVAQAA------------- | *Desulfobulbus propionicus* | Bacteria; Proteobacteria; Deltaproteobacteria; Desulfobacterales; Desulfobulbaceae; Desulfobulbus | chemotaxis protein | 6 |
| PSW25642.1 | 4.5E-06 | Qa.III | LDQISGSVASISDMATQIATAAEEQTHVTDEITQNTTTIK------------- | *Photobacterium swingsii* | Bacteria; Proteobacteria; Gammaproteobacteria; Vibrionales; Vibrionaceae; Photobacterium | methyl-accepting chemotaxis protein | 6 |
| WP_019645511.1 | 4.5E-06 | Qa.III.b | IHEIADVITEINALAQDISSAVAQQGESTRAIAINVSEAAKGTQDVAQSVIS- | *Novispirillum itersonii* | Bacteria; Proteobacteria; Alphaproteobacteria; Rhodospirillales; Rhodospirillaceae; Novispirillum | methyl-accepting chemotaxis protein | 6 |
| WP_022694481.1 | 4.5E-06 | Qa.III | --AIERVIGQVNEVSVAISATVEEQATVTQEISGNMSSANTAV---------- | *Ponticaulis koreensis* | Bacteria; Proteobacteria; Alphaproteobacteria; Rhodobacterales; Hyphomonadaceae; Ponticaulis | methyl-accepting chemotaxis protein | 6 |
| WP_074121595.1 | 4.5E-06 | Qc.III | IKDISDTIERLSEISSAIAAAVEEQGAATQEISRNVAQASQGTQQVSSNITD- | *Bradyrhizobium sp. AS23.2* | Bacteria; Proteobacteria; Alphaproteobacteria; Rhizobiales; Bradyrhizobiaceae; Bradyrhizobium | HAMP domain-containing protein | 6 |
| WP_079324920.1 | 4.5E-06 | Qa.III | -SKIREAIQQISDMATQIASAAEEQSLVTAEINANTESVRE------------ | *Vibrio* | Bacteria; Proteobacteria; Gammaproteobacteria; Vibrionales; Vibrionaceae | MULTISPECIES: methyl-accepting chemotaxis protein | 6 |
| WP_082245653.1 | 4.5E-06 | Qa.III | -SKIREAIQQISDMATQIASAAEEQSLVTAEINANTESVRE------------ | *Vibrio harveyi* | Bacteria; Proteobacteria; Gammaproteobacteria; Vibrionales; Vibrionaceae; Vibrio | methyl-accepting chemotaxis protein | 6 |
| WP_084807382.1 | 4.5E-06 | Qc.III | IKDISDTIERLSEISSAIAAAVEEQGAATQEISRNVAQASQGTQQVSSNITD- | *Bradyrhizobium sp. NAS80.1* | Bacteria; Proteobacteria; Alphaproteobacteria; Rhizobiales; Bradyrhizobiaceae; Bradyrhizobium | HAMP domain-containing protein | 6 |
| WP_092029120.1 | 4.5E-06 | Qc.III | IKDISHTIERLSEISSAIAAAVEEQGAATQEISRNVAQAAEGTQQVSSNIT-- | *Bradyrhizobium sp. OK095* | Bacteria; Proteobacteria; Alphaproteobacteria; Rhizobiales; Bradyrhizobiaceae; Bradyrhizobium | methyl-accepting chemotaxis protein | 6 |
| WP_114322037.1 | 4.5E-06 | Qa.III | -SKIREAIQQISDMATQIASAAEEQSLVTAEINANTESVRE------------ | *Vibrio harveyi* | Bacteria; Proteobacteria; Gammaproteobacteria; Vibrionales; Vibrionaceae; Vibrio | HAMP domain-containing protein | 6 |
| AND93636.1 | 4.6E-06 | Qa.IV | IKEIGTTISRISDISTTIAATIEEQGAATAEIARNVSEAAK------------ | *Bradyrhizobium diazoefficiens USDA 110* | Bacteria; Proteobacteria; Alphaproteobacteria; Rhizobiales; Bradyrhizobiaceae; Bradyrhizobium | chemotaxis protein | 6 |
| BAR57062.1 | 4.6E-06 | Qa.IV | IKEIGTTISRISDISTTIAATIEEQGAATAEIARNVSEAAK------------ | *Bradyrhizobium diazoefficiens* | Bacteria; Proteobacteria; Alphaproteobacteria; Rhizobiales; Bradyrhizobiaceae; Bradyrhizobium | putative methyl accepting chemotaxis protein | 6 |
| KEX93810.1 | 4.6E-06 | Qb.I | LQRTKLHLQETIEQSEVSSEELKASNEEMQAINEELRSATEELETS------- | *Pseudomonas putida* | Bacteria; Proteobacteria; Gammaproteobacteria; Pseudomonadales; Pseudomonadaceae; Pseudomonas | hypothetical protein HA62_11210 partial | 6 |
| KGJ65012.1 | 4.6E-06 | Qa.IV | IKEIGTTISRISDISTTIAATIEEQGAATAEIARNVSEAAK------------ | *Bradyrhizobium diazoefficiens SEMIA 5080* | Bacteria; Proteobacteria; Alphaproteobacteria; Rhizobiales; Bradyrhizobiaceae; Bradyrhizobium | hypothetical protein BJA5080_01655 | 6 |
| OCR95425.1 | 4.6E-06 | Qa.IV | --EIEANVNILSQGINEMSQSINEQTEAINQINEAVATVDEQTKQ-NV----- | *Campylobacter fetus subsp. testudinum* | Bacteria; Proteobacteria; Epsilonproteobacteria; Campylobacterales; Campylobacteraceae; Campylobacter | chemotaxis protein partial | 6 |
| OCR95609.1 | 4.6E-06 | Qa.IV | --EIEANVNILSQGINEMSQSINEQTEAINQINEAVATVDEQTKQ-NV----- | *Campylobacter fetus subsp. testudinum* | Bacteria; Proteobacteria; Epsilonproteobacteria; Campylobacterales; Campylobacteraceae; Campylobacter | chemotaxis protein partial | 6 |
| OXE33453.1 | 4.6E-06 | Qa.III | LDEINAAVALISDMATQIATAAEEQTHVTNEITQNVTSIK------------- | *Vibrio parahaemolyticus* | Bacteria; Proteobacteria; Gammaproteobacteria; Vibrionales; Vibrionaceae; Vibrio | chemotaxis protein partial | 6 |
| PTC01217.1 | 4.6E-06 | Qa.III | IDGITSIIANLNEIASEVAAAVQEQGQATSEISHSVQQAVQSTQ--------- | *Thalassospira xiamenensis* | Bacteria; Proteobacteria; Alphaproteobacteria; Rhodospirillales; Rhodospirillaceae; Thalassospira | methyl-accepting chemotaxis protein | 6 |
| PXX27550.1 | 4.6E-06 | Qa.III | IDGITSIIANLNEIASEVAAAVQEQGQATSEISHSVQQAVQSTQ--------- | *Thalassospira sp. 11-3* | Bacteria; Proteobacteria; Alphaproteobacteria; Rhodospirillales; Rhodospirillaceae; Thalassospira | methyl-accepting chemotaxis protein | 6 |
| WP_007091851.1 | 4.6E-06 | Qa.III | IDGITSIIANLNEIASEVAAAVQEQGQATSEISHSVQQAVQSTQ--------- | *Thalassospira* | Bacteria; Proteobacteria; Alphaproteobacteria; Rhodospirillales; Rhodospirillaceae | MULTISPECIES: methyl-accepting chemotaxis protein | 6 |
| WP_011083405.1 | 4.6E-06 | Qa.IV | IKEIGTTISRISDISTTIAATIEEQGAATAEIARNVSEAAK------------ | *Bradyrhizobium diazoefficiens* | Bacteria; Proteobacteria; Alphaproteobacteria; Rhizobiales; Bradyrhizobiaceae; Bradyrhizobium | HAMP domain-containing protein | 6 |
| WP_037989418.1 | 4.6E-06 | Qa.III | IDGITSIIANLNEIASEVAAAVQEQGQATSEISHSVQQAVQSTQ--------- | *Thalassospira* | Bacteria; Proteobacteria; Alphaproteobacteria; Rhodospirillales; Rhodospirillaceae | MULTISPECIES: methyl-accepting chemotaxis protein | 6 |
| WP_051357368.1 | 4.6E-06 | Qc.III | IESVSQQIRSMSDVATGISAAVEEQGIATREIVRNVDQAATGTNSVTAHITD- | *Azorhizobium doebereinerae* | Bacteria; Proteobacteria; Alphaproteobacteria; Rhizobiales; Xanthobacteraceae; Azorhizobium | HAMP domain-containing protein | 6 |
| WP_051475087.1 | 4.6E-06 | Qa.IV | VKDIVATIGEINRLSGEVSEAVDQQNAATSEIARNVQQAA------------- | *Rhodospirillales bacterium URHD0088* | Bacteria; Proteobacteria; Alphaproteobacteria; Rhodospirillales; unclassified Rhodospirillales | methyl-accepting chemotaxis protein | 6 |
| WP_067743312.1 | 4.6E-06 | Qa.III | LQDIASGISQINDINTQVATATEEQTVVVEDINQNIDTISNNS---------- | *Alteromonadaceae bacterium XY-R5* | Bacteria; Proteobacteria; Gammaproteobacteria; Alteromonadales; Alteromonadaceae | methyl-accepting chemotaxis protein | 6 |
| WP_074076446.1 | 4.6E-06 | Qa.IV | IKEIGTTISRISDISTTIAATIEEQGAATAEIARNVSEAAK------------ | *Bradyrhizobium* | Bacteria; Proteobacteria; Alphaproteobacteria; Rhizobiales; Bradyrhizobiaceae | MULTISPECIES: methyl-accepting chemotaxis protein | 6 |
| WP_082755728.1 | 4.6E-06 | Qa.IV | IKEIGTTISRISDISTTIAATIEEQGAATAEIARNVSEAAK------------ | *Bradyrhizobium diazoefficiens* | Bacteria; Proteobacteria; Alphaproteobacteria; Rhizobiales; Bradyrhizobiaceae; Bradyrhizobium | methyl-accepting chemotaxis protein | 6 |
| WP_082793157.1 | 4.6E-06 | Qb.I | LAATREYLQSVIEQQDAANEELQSSSEEIQSANEELQSINEELETSK------ | *Ramlibacter tataouinensis* | Bacteria; Proteobacteria; Betaproteobacteria; Burkholderiales; Comamonadaceae; Ramlibacter | PAS domain S-box protein | 6 |
| WP_085646114.1 | 4.6E-06 | Qa.III | IDGITSIIANLNEIASEVAAAVQEQGQATSEISHSVQQAVQSTQ--------- | *Thalassospira sp. MCCC 1A03138* | Bacteria; Proteobacteria; Alphaproteobacteria; Rhodospirillales; Rhodospirillaceae; Thalassospira | methyl-accepting chemotaxis protein | 6 |
| WP_091881142.1 | 4.6E-06 | Qa.III | IESIRATIMEVNEISSSIAAAVEEQGAATQAIARNVQEAA------------- | *Bradyrhizobium sp. Rc2d* | Bacteria; Proteobacteria; Alphaproteobacteria; Rhizobiales; Bradyrhizobiaceae; Bradyrhizobium | methyl-accepting chemotaxis protein | 6 |
| WP_092189338.1 | 4.6E-06 | Qa.IV | IKEIGTTISRISDISTTIAATIEEQGAATAEIARNVSEAAK------------ | *Bradyrhizobium sp. cf659* | Bacteria; Proteobacteria; Alphaproteobacteria; Rhizobiales; Bradyrhizobiaceae; Bradyrhizobium | methyl-accepting chemotaxis protein | 6 |
| AKU95205.1 | 4.7E-06 | Qa.III | MDEIVGSVKRVTDIMTDIAGATREQSSGLDQVNTAIVQID------------- | *Labilithrix luteola* | Bacteria; Proteobacteria; Deltaproteobacteria; Myxococcales; Sorangiineae; Labilitrichaceae; Labilithrix | Methyl-accepting chemotaxis protein I (serine chemoreceptor protein) | 6 |
| KOY43264.1 | 4.7E-06 | Qa.III | LDEINAAVALISDMATQIATAAEEQTHVTNEITQNVTSIK------------- | *Vibrio parahaemolyticus* | Bacteria; Proteobacteria; Gammaproteobacteria; Vibrionales; Vibrionaceae; Vibrio | chemotaxis protein partial | 6 |
| WP_009868888.1 | 4.7E-06 | Qc.III | --GIATTISAINDISATISHAVAEQGQATQEIARSVDQAAQRTGEVSEKIE-- | *Magnetospirillum magnetotacticum* | Bacteria; Proteobacteria; Alphaproteobacteria; Rhodospirillales; Rhodospirillaceae; Magnetospirillum | methyl-accepting chemotaxis protein | 6 |
| WP_018315628.1 | 4.7E-06 | Qc.III | IGEISGTIARLSEIAAAIAAAVEQQGAATQEIARNVQQAAQRTQQVSSNVG-- | *Bradyrhizobium sp. WSM2793* | Bacteria; Proteobacteria; Alphaproteobacteria; Rhizobiales; Bradyrhizobiaceae; Bradyrhizobium | PAS domain S-box protein | 6 |
| WP_037293629.1 | 4.7E-06 | Qa.III | IQEIAQTAYQVSEAASEMKTEATAGGESIDRVGDQMSAIS------------- | *Saccharibacillus sacchari* | Bacteria; Firmicutes; Bacilli; Bacillales; Paenibacillaceae; Saccharibacillus | methyl-accepting chemotaxis protein | 6 |
| WP_046019042.1 | 4.7E-06 | Qa.IV | ---ITEQISQINTLSLQIATSTKEQTQMIDVILNNVEN----LNDGVHKTGD- | *Marinomonas sp. S3726* | Bacteria; Proteobacteria; Gammaproteobacteria; Oceanospirillales; Marinomonas | methyl-accepting chemotaxis protein | 6 |
| WP_063339687.1 | 4.7E-06 | Qa.IV | ---ITEQISQINTLSLQIATSTKEQTQMIDVILNNVEN----LNDGVHKTGD- | *Marinomonas* | Bacteria; Proteobacteria; Gammaproteobacteria; Oceanospirillales | MULTISPECIES: methyl-accepting chemotaxis protein | 6 |
| WP_082367929.1 | 4.7E-06 | R.IV | IDQVNTAVTQLDDATQQNASLVQQSSEAAASLQHQAQRLAEAIGVFKTAT--- | *Ideonella sakaiensis* | Bacteria; Proteobacteria; Betaproteobacteria; Burkholderiales; Ideonella | hypothetical protein | 6 |
| WP_109328712.1 | 4.7E-06 | Qc.III | IQGIGGTITRMNEIATAISGAIAEQNATTSEISRSVQHASRATQSVSAS---- | *Azospirillum sp. CFH 70021* | Bacteria; Proteobacteria; Alphaproteobacteria; Rhodospirillales; Rhodospirillaceae; Azospirillum | HAMP domain-containing protein | 6 |
| PWB61606.1 | 4.8E-06 | Qa.IV | IKEIGQTIGQIANISSTIAAAVEEQGAATQEIARNVQQAAE------------ | *Bradyrhizobiaceae bacterium* | Bacteria; Proteobacteria; Alphaproteobacteria; Rhizobiales; Bradyrhizobiaceae | methyl-accepting chemotaxis protein | 6 |
| WP_009540318.1 | 4.8E-06 | Qc.III | IQGIGGTIGRISEITGAIAAAVEEQSAATQEIARNVQEASSGTTEVTSNIS-- | *Caenispirillum salinarum* | Bacteria; Proteobacteria; Alphaproteobacteria; Rhodospirillales; Rhodospirillaceae; Caenispirillum | methyl-accepting chemotaxis protein | 6 |
| WP_011389829.1 | 4.8E-06 | Qc.III | INEIATTIGEINEISANIAAAMEQQGAATQEIARNVEQAATGTDQVTRNIH-- | *Rhodospirillum rubrum* | Bacteria; Proteobacteria; Alphaproteobacteria; Rhodospirillales; Rhodospirillaceae; Rhodospirillum | methyl-accepting chemotaxis protein | 6 |
| WP_011663877.1 | 4.8E-06 | Qa.IV | IKEIGSTIERMSDIASTIAAAVEQQGAATQEIARNVQQAS------------- | *Rhodopseudomonas palustris* | Bacteria; Proteobacteria; Alphaproteobacteria; Rhizobiales; Bradyrhizobiaceae; Rhodopseudomonas | methyl-accepting chemotaxis protein | 6 |
| WP_039469761.1 | 4.8E-06 | Qa.III | LEQINASVAMISDMATQIATAAEEQTHVTDEITQNTTTIK------------- | *Photobacterium gaetbulicola* | Bacteria; Proteobacteria; Gammaproteobacteria; Vibrionales; Vibrionaceae; Photobacterium | methyl-accepting chemotaxis protein | 6 |
| WP_044620771.1 | 4.8E-06 | Qa.III | LEQINASVAMISDMATQIATAAEEQTHVTDEITQNTTTIK------------- | *Photobacterium gaetbulicola* | Bacteria; Proteobacteria; Gammaproteobacteria; Vibrionales; Vibrionaceae; Photobacterium | methyl-accepting chemotaxis protein | 6 |
| WP_084327421.1 | 4.8E-06 | Qc.III | IESIGATIQRMHEISGTIASAAEEQQSATQEIASNVSDAANGTDAVN------ | *Salinarimonas rosea* | Bacteria; Proteobacteria; Alphaproteobacteria; Rhizobiales; Bradyrhizobiaceae; Salinarimonas | methyl-accepting chemotaxis protein | 6 |
| WP_084539766.1 | 4.8E-06 | Qa.III | ITGIASTIKQMNEISTTIASAVTQQSAATDEIANSIARASEGTRNASEN---- | *Azorhizobium doebereinerae* | Bacteria; Proteobacteria; Alphaproteobacteria; Rhizobiales; Xanthobacteraceae; Azorhizobium | methyl-accepting chemotaxis protein | 6 |
| WP_114087132.1 | 4.8E-06 | Qa.III | IQGIGTTIAELNEIAASIASAVEQQGAATGEITRNIQQAS------------- | *Thalassospira profundimaris* | Bacteria; Proteobacteria; Alphaproteobacteria; Rhodospirillales; Rhodospirillaceae; Thalassospira | HAMP domain-containing protein | 6 |
| APS54140.1 | 4.9E-06 | Qa.III | LDKIQDSAGEVTKATDDIARAVSEQTEATKSVTGYMEKTLDEVE--------- | *Piscirickettsia salmonis* | Bacteria; Proteobacteria; Gammaproteobacteria; Thiotrichales; Piscirickettsiaceae; Piscirickettsia | chemotaxis protein | 6 |
| APS84152.1 | 4.9E-06 | Qa.III | LDKIQDSAGEVTKATDDIARAVSEQTEATKSVTGYMEKTLDEVE--------- | *Piscirickettsia salmonis* | Bacteria; Proteobacteria; Gammaproteobacteria; Thiotrichales; Piscirickettsiaceae; Piscirickettsia | chemotaxis protein | 6 |

| | | | | | | | | |
|---|---|---|---|---|---|---|---|---|
|  | **PEQ17218.1** | 4.9E-06 | Qa.III | LDKIQDSAGEVTKATDDIARAVSEQTEATKSVTGYMEKTLDEVE--------- | *Piscirickettsia salmonis* | Bacteria; Proteobacteria; Gammaproteobacteria; Thiotrichales; Piscirickettsiaceae; Piscirickettsia | methyl-accepting chemotaxis protein | 6 |
|  | **PJJ31230.1** | 4.9E-06 | Qa.III | IENIGGTIGAINEITTAIASAVEEQGSATKEIARNVQEAA------------ | *Afipia broomeae* | Bacteria; Proteobacteria; Alphaproteobacteria; Rhizobiales; Bradyrhizobiaceae; Afipia | methyl-accepting chemotaxis sensory transducer with Cache sensor | 6 |
| | **PPR20095.1** | 4.9E-06 | Qc.III | IEGITKTITEINEIAGTIAAAVEEQGAATGEISRNAQEAATGTQEVNSNIAN- | *Alphaproteobacteria bacterium MarineAlpha10_Bin2* | Bacteria; Proteobacteria; Alphaproteobacteria | Methyl-accepting chemotaxis protein CtpH | 6 |
|  | **SDS73919.1** | 4.9E-06 | Qa.III | IENIGGTIGAINEITTAIASAVEEQGSATKEIARNVQEAA------------ | *Bradyrhizobium canariense* | Bacteria; Proteobacteria; Alphaproteobacteria; Rhizobiales; Bradyrhizobiaceae; Bradyrhizobium | methyl-accepting chemotaxis sensory transducer with Cache sensor | 6 |
| | **WP_027301976.1** | 4.9E-06 | Qc.III | -KGIGETISRVAEIATTIASAVEEQGAATQEIARNVQEAAAGTQEVSSHI--- | *Rhodospirillales bacterium URHD0088* | Bacteria; Proteobacteria; Alphaproteobacteria; Rhodospirillales; unclassified Rhodospirillales | HAMP domain-containing protein | 6 |
|  | **WP_035039702.1** | 4.9E-06 | Qb.I | ----KAKAESAVNMMEDSNKMVVEQSEKVGETSEKFDEISKAVENSKTIVS-- | *Catonella morbi* | Bacteria; Firmicutes; Clostridia; Clostridiales; Lachnospiraceae; Catonella | methyl-accepting chemotaxis protein | 6 |
|  | **WP_063308211.1** | 4.9E-06 | Qc.III.c | ---IGETIEQINALSTGINEAVGDQKQATEDISRDLNSAHQNVSQV------- | *Pseudovibrio sp. Ad46* | Bacteria; Proteobacteria; Alphaproteobacteria; Rhodobacterales; Rhodobacteraceae; Pseudovibrio | methyl-accepting chemotaxis protein | 6 |
|  | **WP_080133924.1** | 4.9E-06 | Qa.IV | IKEIGSTISRISDISTTIAATIEEQGAATAEIARNVSEAAK----------- | *Bradyrhizobium sp. BR10280* | Bacteria; Proteobacteria; Alphaproteobacteria; Rhizobiales; Bradyrhizobiaceae; Bradyrhizobium | HAMP domain-containing protein | 6 |
|  | **WP_099510139.1** | 4.9E-06 | Qa.IV | IQEIARTITEMAQISTSIAAAMEEQGAATAEIARNVQEAA------------ | *Microvirga ossetica* | Bacteria; Proteobacteria; Alphaproteobacteria; Rhizobiales; Methylobacteriaceae; Microvirga | HAMP domain-containing protein | 6 |
| | **KIZ36211.1** | 5E-06 | Qc.III | IKEISDTIGRMSEIASTIASAVEEQGAATQEIARNVQLAAQGTTQVSANI--- | *Rhodopseudomonas palustris* | Bacteria; Proteobacteria; Alphaproteobacteria; Rhizobiales; Bradyrhizobiaceae; Rhodopseudomonas | chemotaxis protein partial | 6 |
|  | **OOE71553.1** | 5E-06 | Qc.III | -DEISGNLSNIKNLSSDMNEAIEQLGPIVSNLKENVDSLSGAIRHL------- | *Salinivibrio kushneri* | Bacteria; Proteobacteria; Gammaproteobacteria; Vibrionales; Vibrionaceae; Salinivibrio | methyl-accepting chemotaxis protein | 6 |
| | **WP_063701014.1** | 5E-06 | Qc.III | IQSISETIQRVSQIAAAIAAAVEEQGAATREIARNVQQASEGTTEVSRHIA-- | *Bradyrhizobium sp. BR 10245* | Bacteria; Proteobacteria; Alphaproteobacteria; Rhizobiales; Bradyrhizobiaceae; Bradyrhizobium | HAMP domain-containing protein | 6 |
|  | **WP_064004073.1** | 5E-06 | Qa.III | LDKIQDSAGEVTKATDDIARAVSEQTEATKSVTGYMEKTLDEVE--------- | *Piscirickettsiaceae bacterium NZ-RLO1* | Bacteria; Proteobacteria; Gammaproteobacteria; Thiotrichales; Piscirickettsiaceae; unclassified Piscirickettsiaceae | methyl-accepting chemotaxis protein | 6 |
| | **WP_076865993.1** | 5E-06 | Qc.III | INEISATIKRLSEVSSAIAAAVEEQGAVTQEIARNVQRAALGTQEVSSSIS-- | *Bradyrhizobium mercantei* | Bacteria; Proteobacteria; Alphaproteobacteria; Rhizobiales; Bradyrhizobiaceae; Bradyrhizobium | methyl-accepting chemotaxis protein | 6 |
|  | **WP_094409854.1** | 5E-06 | Qa.III | IQGIAATILKINDISSGIAAAVEQQSAATREITRNVQEAARSTEDVS------ | *Elstera cyanobacteriorum* | Bacteria; Proteobacteria; Alphaproteobacteria; Rhodospirillales; Rhodospirillaceae; Elstera | HAMP domain-containing protein | 6 |
|  | **WP_101267278.1** | 5E-06 | Qa.III | IDGIARIINDLNAISAEVAAAVRQQGSATSEISHSVQQAVSSTQQ-------- | *Thalassospira marina* | Bacteria; Proteobacteria; Alphaproteobacteria; Rhodospirillales; Rhodospirillaceae; Thalassospira | methyl-accepting chemotaxis protein | 6 |
|  | **WP_110782638.1** | 5E-06 | Qa.IV | IKEISLTISRMSEIASTIASAVEEQGAATQEISRNVQEAA------------ | *Rhodopseudomonas faecalis* | Bacteria; Proteobacteria; Alphaproteobacteria; Rhizobiales; Bradyrhizobiaceae; Rhodopseudomonas | methyl-accepting chemotaxis protein | 6 |
|  | **OYU50299.1** | 5.1E-06 | Qc.III | IDAISQTISAINGISISISAAVEEQAAVTSDMSHNMQEAARG----------- | *Rhizobiales bacterium PAR1* | Bacteria; Proteobacteria; Alphaproteobacteria; Rhizobiales | chemotaxis protein | 6 |
| | **WP_011925586.1** | 5.1E-06 | Qc.III | ---ISGTIERLSEIASAIAAAVEEQGAATQEISRNVQQAAQGTQQVSSNIAD- | *Bradyrhizobium sp. ORS 278* | Bacteria; Proteobacteria; Alphaproteobacteria; Rhizobiales; Bradyrhizobiaceae; Bradyrhizobium | methyl-accepting chemotaxis protein | 6 |
|  | **WP_079599852.1** | 5.1E-06 | Qa.IV | IRNIGQTISEINEVTAAIAAAVEEQGAATREIARNIQHAA------------ | *Bradyrhizobium erythrophlei* | Bacteria; Proteobacteria; Alphaproteobacteria; Rhizobiales; Bradyrhizobiaceae; Bradyrhizobium | HAMP domain-containing protein | 6 |
|  | **WP_089264137.1** | 5.1E-06 | Qa.III | IQGITATIEEVSAIATTIGSAIEEQGTATAEIARNVTQTAQATQN-------- | *Tardiphaga sp. OK246* | Bacteria; Proteobacteria; Alphaproteobacteria; Rhizobiales; Bradyrhizobiaceae; Tardiphaga | methyl-accepting chemotaxis protein | 6 |
|  | **WP_092143469.1** | 5.1E-06 | Qa.III | IQGITATIEEVSAIATTIGSAIEEQGTATAEIARNVTQTAQATQN-------- | *Bradyrhizobium sp. NFR13* | Bacteria; Proteobacteria; Alphaproteobacteria; Rhizobiales; Bradyrhizobiaceae; Bradyrhizobium | HAMP domain-containing protein | 6 |
|  | **WP_093758414.1** | 5.1E-06 | Qa.III | IQGITATIEEVSAIATTIGSAIEEQGTATAEIARNVTQTAQATQN-------- | *Tardiphaga sp. OK245* | Bacteria; Proteobacteria; Alphaproteobacteria; Rhizobiales; Bradyrhizobiaceae; Tardiphaga | methyl-accepting chemotaxis protein | 6 |
|  | **WP_102709963.1** | 5.1E-06 | Qc.III | -QQVSASVNELATGANEISEDVSHMSESITEIEEKVSDYADSTKEMNTR---- | *Paenibacillus castaneae* | Bacteria; Firmicutes; Bacilli; Bacillales; Paenibacillaceae; Paenibacillus | methyl-accepting chemotaxis protein | 6 |
|  | **KJS34929.1** | 5.2E-06 | Qc.III | -GEIVHVVQRVNEVSAAIASAVEEQNAATQEISRNVQEAANG----------- | *Rhodospirillaceae bacterium BRH_c57* | Bacteria; Proteobacteria; Alphaproteobacteria; Rhodospirillales; Rhodospirillaceae | hypothetical protein VR70_16770 | 6 |
|  | **WP_066473160.1** | 5.2E-06 | Qa.III | MKDIVQSVGQVTGLLGEISHALQEQSEGI----DHVNTAVAHMDSTNQE---- | *Comamonas terrae* | Bacteria; Proteobacteria; Betaproteobacteria; Burkholderiales; Comamonadaceae; Comamonas | HAMP domain-containing protein | 6 |
|  | **WP_083204530.1** | 5.2E-06 | Qc.III | --DIGGVIDRMKEISTAISAAMEEQGAATNEIARNVQEAAKGTGEVT------ | *Salinarimonadaceae bacterium HL-109* | Bacteria; Proteobacteria; Alphaproteobacteria; Rhizobiales; Salinarimonadaceae | methyl-accepting chemotaxis protein | 6 |
| | **WP_085592826.1** | 5.2E-06 | Qa.III | IDGITRIINDLNSISAEVAAAVRQQGTATSEISHSVQQAVSSTQQVSTHI--- | *Thalassospira sp. MCCC 1A01428* | Bacteria; Proteobacteria; Alphaproteobacteria; Rhodospirillales; Rhodospirillaceae; Thalassospira | methyl-accepting chemotaxis protein | 6 |
|  | **WP_100380382.1** | 5.2E-06 | Qa.III | IENIGGTIGAINEITTAIASAVEEQGSATKEIARNVQEAA------------ | *Afipia broomeae* | Bacteria; Proteobacteria; Alphaproteobacteria; Rhizobiales; Bradyrhizobiaceae; Afipia | HAMP domain-containing protein | 6 |
| | **WP_108519813.1** | 5.2E-06 | Qc.III | IQAISGTIERLSEIASTIAAAVEEQGAATQEISRNVQQAAQGTQQVSSNITD- | *Bradyrhizobium algeriense* | Bacteria; Proteobacteria; Alphaproteobacteria; Rhizobiales; Bradyrhizobiaceae; Bradyrhizobium | methyl-accepting chemotaxis protein | 6 |
| | **WP_111419403.1** | 5.2E-06 | Qc.III | IEEVAATIRELENISTSISSAVEEQSAATSEIAQNVGQTASHVGRVSALMDD- | *Rhodoplanes roseus* | Bacteria; Proteobacteria; Alphaproteobacteria; Rhizobiales; Hyphomicrobiaceae; Rhodoplanes | hypothetical protein | 6 |
| | **AWM06051.1** | 5.3E-06 | Qc.III | -RDISHTIEKLSEISATIAAAVEEQGAATQEISRNVQQAAEGTHQVSSNITD- | *Bradyrhizobium sp. 3 85S1MB* | Bacteria; Proteobacteria; Alphaproteobacteria; Rhizobiales; Bradyrhizobiaceae; Bradyrhizobium | methyl-accepting chemotaxis protein | 6 |
|  | **ESL03325.1** | 5.3E-06 | Qb.I | ----KAKAESAVNMMEDSNKMVVEQSEKVGETSEKFDEISKAVENSKTIVS-- | *Catonella morbi ATCC 51271* | Bacteria; Firmicutes; Clostridia; Clostridiales; Lachnospiraceae; Catonella | methyl-accepting chemotaxis protein signaling domain protein | 6 |
| | **OSI68988.1** | 5.3E-06 | Qa.IV | -KEIGATIGQISSISTSIASAVEQQGAATQEIARSVQTVAQGTQTAATDI--- | *Bradyrhizobium canariense* | Bacteria; Proteobacteria; Alphaproteobacteria; Rhizobiales; Bradyrhizobiaceae; Bradyrhizobium | hypothetical protein BSZ15_00130 partial | 6 |
|  | **OYW52230.1** | 5.3E-06 | Qa.IV | -------ISDIRQVTAAIASAVEQQGAATREIAQNTQHAAHGTEAVNDNIQ-- | *Rhizobiales bacterium 12-68-15* | Bacteria; Proteobacteria; Alphaproteobacteria; Rhizobiales | hypothetical protein B7Z30_16110 | 6 |
| | **OYX13803.1** | 5.3E-06 | Qc.III | --GVCAQIQRMAEVASGISAAVEEQGVATGEIVRNVDQAAAGTRSVTAHIAD- | *Rhizobiales bacterium 32-66-8* | Bacteria; Proteobacteria; Alphaproteobacteria; Rhizobiales | hypothetical protein B7Z15_06060 | 6 |
| | **PIT06076.1** | 5.3E-06 | Qc.III | LHELRDVINEVNEVAVSISAAVEEQTAATSEIGRNIMEAAGRTEAVTRSI--- | *Bradyrhizobium sp. TSA1* | Bacteria; Proteobacteria; Alphaproteobacteria; Rhizobiales; Bradyrhizobiaceae; Bradyrhizobium | hypothetical protein TSA1_14840 | 6 |
|  | **PIW25982.1** | 5.3E-06 | Qc.III.c | ------VINRINEIAGAVAAAVEEQGAATSEIARNAEQAASGVATVTTNLG-- | *Rhodospirillales bacterium CG15_BIG_FIL_POST_REV_8_21_14_020_66_15* | Bacteria; Proteobacteria; Alphaproteobacteria; Rhodospirillales; unclassified Rhodospirillales | methyl-accepting chemotaxis protein | 6 |
| | **PVV22797.1** | 5.3E-06 | Qa.III | -SEINKAIETINEMNTQIATAAEEQCSVVEEINLNITNISENSKQTSQNAK-- | *gamma proteobacterium symbiont of Ctena orbiculata* | Bacteria; Proteobacteria; Gammaproteobacteria | hypothetical protein B6D78_04420 | 6 |
|  | **SCC81413.1** | 5.3E-06 | Qc.III | --DIGGVIDRMKEISTAISAAMEEQGAATNEIARNVQEAAKGTGEVT------ | *Salinarimonadaceae bacterium HL-109* | Bacteria; Proteobacteria; Alphaproteobacteria; Rhizobiales; Salinarimonadaceae | methyl-accepting chemotaxis sensory transducer with Cache sensor | 6 |
| | **WP_014497319.1** | 5.3E-06 | Qc.III | -RDISDTIEKLSEISSAIAAAVEEQGAATQEISRNVQQAAEGTHQVSSNITD- | *Bradyrhizobium japonicum* | Bacteria; Proteobacteria; Alphaproteobacteria; Rhizobiales; Bradyrhizobiaceae; Bradyrhizobium | methyl-accepting chemotaxis protein | 6 |
|  | **WP_018269047.1** | 5.3E-06 | Qa.IV | IRNISSTIGEINDVTTAIAAAVEQQGAATREIARNIQHAA------------ | *Bradyrhizobium elkanii* | Bacteria; Proteobacteria; Alphaproteobacteria; Rhizobiales; Bradyrhizobiaceae; Bradyrhizobium | HAMP domain-containing protein | 6 |
| | **WP_028152151.1** | 5.3E-06 | Qc.III | -RDISDTIEKLSEISSAIAAAVEEQGAATQEISRNVQQAAEGTHQVSSNITD- | *Bradyrhizobium japonicum* | Bacteria; Proteobacteria; Alphaproteobacteria; Rhizobiales; Bradyrhizobiaceae; Bradyrhizobium | methyl-accepting chemotaxis protein | 6 |
| | **WP_028158175.1** | 5.3E-06 | Qc.III | -RDISDTIEKLSEISSAIAAAVEEQGAATQEISRNVQQAAEGTHQVSSNITD- | *Bradyrhizobium japonicum* | Bacteria; Proteobacteria; Alphaproteobacteria; Rhizobiales; Bradyrhizobiaceae; Bradyrhizobium | methyl-accepting chemotaxis protein | 6 |
|  | **WP_028333723.1** | 5.3E-06 | Qa.IV | IRNISSTIGEINDVTTAIAAAVEQQGAATREIARNIQHAA------------ | *Bradyrhizobium elkanii* | Bacteria; Proteobacteria; Alphaproteobacteria; Rhizobiales; Bradyrhizobiaceae; Bradyrhizobium | HAMP domain-containing protein | 6 |
|  | **WP_028338943.1** | 5.3E-06 | Qa.IV | IRNISSTIGEINDVTTAIAAAVEQQGAATREIARNIQHAA------------ | *Bradyrhizobium elkanii* | Bacteria; Proteobacteria; Alphaproteobacteria; Rhizobiales; Bradyrhizobiaceae; Bradyrhizobium | HAMP domain-containing protein | 6 |
|  | **WP_038384425.1** | 5.3E-06 | Qa.IV | IRNISSTIGEINDVTTAIAAAVEQQGAATREIARNIQHAA------------ | *Bradyrhizobium elkanii* | Bacteria; Proteobacteria; Alphaproteobacteria; Rhizobiales; Bradyrhizobiaceae; Bradyrhizobium | HAMP domain-containing protein | 6 |
| | **WP_038935330.1** | 5.3E-06 | Qc.III | -RDISDTIEKLSEISSAIAAAVEEQGAATQEISRNVQQAAEGTHQVSSNITD- | *Bradyrhizobium japonicum* | Bacteria; Proteobacteria; Alphaproteobacteria; Rhizobiales; Bradyrhizobiaceae; Bradyrhizobium | methyl-accepting chemotaxis protein | 6 |
| | **WP_038944556.1** | 5.3E-06 | Qc.III | -RDISDTIEKLSEISSAIAAAVEEQGAATQEISRNVQQAAEGTHQVSSNITD- | *Bradyrhizobium japonicum* | Bacteria; Proteobacteria; Alphaproteobacteria; Rhizobiales; Bradyrhizobiaceae; Bradyrhizobium | methyl-accepting chemotaxis protein | 6 |
| | **WP_038956604.1** | 5.3E-06 | Qc.III | -RDISDTIEKLSEISSAIAAAVEEQGAATQEISRNVQQAAEGTHQVSSNITD- | *Bradyrhizobium japonicum* | Bacteria; Proteobacteria; Alphaproteobacteria; Rhizobiales; Bradyrhizobiaceae; Bradyrhizobium | methyl-accepting chemotaxis protein | 6 |
| | **WP_051664492.1** | 5.3E-06 | Qc.III | -RDISDTIEKLSEISSAIAAAVEEQGAATQEISRNVQQAAEGTHQVSSNITD- | *Bradyrhizobium japonicum* | Bacteria; Proteobacteria; Alphaproteobacteria; Rhizobiales; Bradyrhizobiaceae; Bradyrhizobium | methyl-accepting chemotaxis protein | 6 |
|  | **WP_057018145.1** | 5.3E-06 | Qa.IV | IRNISSTIGEINDVTTAIAAAVEQQGAATREIARNIQHAA------------ | *Bradyrhizobium* | Bacteria; Proteobacteria; Alphaproteobacteria; Rhizobiales; Bradyrhizobiaceae | MULTISPECIES: HAMP domain-containing protein | 6 |
|  | **WP_060512342.1** | 5.3E-06 | Qb.I | LQRTKTHLQETIEQSEVSSEELKASNEEMQAINEELRSATEELETS------- | *Pseudomonas sp. NBRC 111124* | Bacteria; Proteobacteria; Gammaproteobacteria; Pseudomonadales; Pseudomonadaceae; Pseudomonas | PAS domain S-box protein | 6 |
| | **WP_063986624.1** | 5.3E-06 | Qc.III | -RDISDTIEKLSEISSAIAAAVEEQGAATQEISRNVQQAAEGTHQVSSNITD- | *Bradyrhizobium sp.* | Bacteria; Proteobacteria; Alphaproteobacteria; Rhizobiales; Bradyrhizobiaceae; Bradyrhizobium | methyl-accepting chemotaxis protein | 6 |
|  | **WP_069280270.1** | 5.3E-06 | Qa.IV | IRNISSTIGEINDVTTAIAAAVEQQGAATREIARNIQHAA------------ | *Bradyrhizobium elkanii* | Bacteria; Proteobacteria; Alphaproteobacteria; Rhizobiales; Bradyrhizobiaceae; Bradyrhizobium | HAMP domain-containing protein | 6 |
| | **WP_071917655.1** | 5.3E-06 | Qc.III | -RDISDTIEKLSEISSAIAAAVEEQGAATQEISRNVQQAAEGTHQVSSNITD- | *Bradyrhizobium japonicum* | Bacteria; Proteobacteria; Alphaproteobacteria; Rhizobiales; Bradyrhizobiaceae; Bradyrhizobium | methyl-accepting chemotaxis protein | 6 |
|  | **WP_083763513.1** | 5.3E-06 | Qa.IV | ------TIREINGISTAIASAVEEQSAATSEISRNVHEASKAVNDVNKNI--- | *Magnetospirillum magneticum* | Bacteria; Proteobacteria; Alphaproteobacteria; Rhodospirillales; Rhodospirillaceae; Magnetospirillum | methyl-accepting chemotaxis protein | 6 |
| | **WP_085402595.1** | 5.3E-06 | Qc.III | -RDISDTIEKLSEISSAIAAAVEEQGAATQEISRNVQQAAEGTHQVSSNITD- | *Bradyrhizobium japonicum* | Bacteria; Proteobacteria; Alphaproteobacteria; Rhizobiales; Bradyrhizobiaceae; Bradyrhizobium | methyl-accepting chemotaxis protein | 6 |
|  | **WP_092080195.1** | 5.3E-06 | Qa.IV | IRNISSTIGEINDVTTAIAAAVEQQGAATREIARNIQHAA------------ | *Bradyrhizobium sp. R5* | Bacteria; Proteobacteria; Alphaproteobacteria; Rhizobiales; Bradyrhizobiaceae; Bradyrhizobium | HAMP domain-containing protein | 6 |

| | | | | | | | | |
|---|---|---|---|---|---|---|---|---|
| | **WP_099556408.1** | 5.3E-06 | Qc.III | INSIADAIEELNRIAAAISTGVDQQAVATDEISSSMQTVARNSNQVSENIN-- | *Hartmannibacter diazotrophicus* | Bacteria; Proteobacteria; Alphaproteobacteria; Rhizobiales; Hartmannibacter | methyl-accepting chemotaxis protein | 6 |
|  | **WP_107988082.1** | 5.3E-06 | Qa.IV | IKEITRTMDDVNSYTSAIAAAVEQQGSATTEISRNVQGAA------------ | *Breoghania corrubedonensis* | Bacteria; Proteobacteria; Alphaproteobacteria; Rhizobiales; Cohaesibacteraceae; Breoghania | methyl-accepting chemotaxis protein | 6 |
| | **WP_109332196.1** | 5.3E-06 | Qc.III | --GVGSIIGQINDIATTIAAAVEEQNAATLEIARNVQQAASGTQQVTSNID-- | *Azospirillum sp. CFH 70021* | Bacteria; Proteobacteria; Alphaproteobacteria; Rhodospirillales; Rhodospirillaceae; Azospirillum | methyl-accepting chemotaxis protein | 6 |
|  | **WP_111421952.1** | 5.3E-06 | Qc.III | IKEIGGTIGKIAEIAATIAAAVEEQGAATNEIARNVQQASSGTTEV------- | *Rhodoplanes roseus* | Bacteria; Proteobacteria; Alphaproteobacteria; Rhizobiales; Hyphomicrobiaceae; Rhodoplanes | methyl-accepting chemotaxis protein | 6 |
|  | **WP_114088037.1** | 5.3E-06 | Qa.III | IQNISRTINEVDAISTAIASAVEQQGAATREISSSVYQAVQSTESAQAN---- | *Thalassospira profundimaris* | Bacteria; Proteobacteria; Alphaproteobacteria; Rhodospirillales; Rhodospirillaceae; Thalassospira | chemotaxis protein | 6 |
| | **AJA64982.1** | 5.4E-06 | Qc.III | -RDISDTIEKLSEISSAIAAAVEEQGAATQEISRNVQQAAEGTHQVSSNITD- | *Bradyrhizobium japonicum* | Bacteria; Proteobacteria; Alphaproteobacteria; Rhizobiales; Bradyrhizobiaceae; Bradyrhizobium | chemotaxis protein | 6 |
| | **KGT73809.1** | 5.4E-06 | Qc.III | -RDISDTIEKLSEISSAIAAAVEEQGAATQEISRNVQQAAEGTHQVSSNITD- | *Bradyrhizobium japonicum* | Bacteria; Proteobacteria; Alphaproteobacteria; Rhizobiales; Bradyrhizobiaceae; Bradyrhizobium | chemotaxis protein | 6 |
|  | **OJJ12878.1** | 5.4E-06 | Qa.IV | --EIAETMTEVNSYTTAIASAVEQQGAATAEISQNVQRAAEATG--------- | *Alphaproteobacteria bacterium AO1-B* | Bacteria; Proteobacteria; Alphaproteobacteria | chemotaxis protein | 6 |
| | **WP_024507688.1** | 5.4E-06 | Qc.III | IKRIGGIIGEVNEVATAIAAAVQEQGAATQEITRSTQYAAQGTKNVSENMT-- | *Bradyrhizobium sp. ARR65* | Bacteria; Proteobacteria; Alphaproteobacteria; Rhizobiales; Bradyrhizobiaceae; Bradyrhizobium | methyl-accepting chemotaxis protein | 6 |
| | **WP_037546997.1** | 5.4E-06 | Qc.III | -RSIAETVERIHSISNTILSAVQKQMAATGEITQSVDFATQNTESVASSMT-- | *Stappia stellulata* | Bacteria; Proteobacteria; Alphaproteobacteria; Rhodobacterales; Rhodobacteraceae; Stappia | methyl-accepting chemotaxis protein | 6 |
|  | **WP_055670511.1** | 5.4E-06 | Qa.IV | --EIAETMTEVNSYTTAIASAVEQQGAATAEISQNVQRAAEATG--------- | *Labrenzia alexandrii* | Bacteria; Proteobacteria; Alphaproteobacteria; Rhodobacterales; Rhodobacteraceae; Labrenzia | HAMP domain-containing protein | 6 |
| | **WP_100177096.1** | 5.4E-06 | Qc.III | LHELRDVINEVNEVAVSISAAVEEQTAATSEIGRNIMEAAGRTEAVTRSI--- | *Bradyrhizobium sp. TSA1* | Bacteria; Proteobacteria; Alphaproteobacteria; Rhizobiales; Bradyrhizobiaceae; Bradyrhizobium | methyl-accepting chemotaxis protein | 6 |
|  | **CDE35798.1** | 5.5E-06 | Qc.I | -EEVTSHAQSVKTSAEQMEQVVDESSRAIQTVSEKLNTAKEQIDK-------- | *Eubacterium sp. CAG:38* | Bacteria; Firmicutes; Clostridia; Clostridiales; Eubacteriaceae; Eubacterium; environmental samples | methyl-accepting chemotaxis protein | 6 |
|  | **OOE70692.1** | 5.5E-06 | Qa.III | -NAINEQIVQISDMNTQVATATEEQSTVVNDINQNIEEINHSSQQTA------ | *Salinivibrio kushneri* | Bacteria; Proteobacteria; Gammaproteobacteria; Vibrionales; Vibrionaceae; Salinivibrio | methyl-accepting chemotaxis protein | 6 |
|  | **SIS38948.1** | 5.5E-06 | Qa.IV | IRDITRQITEINGISTAVAAAVEEQEAATAEISRNVQETS------------ | *Insolitispirillum peregrinum* | Bacteria; Proteobacteria; Alphaproteobacteria; Rhodospirillales; Rhodospirillaceae; Insolitispirillum | hemerythrin-like metal-binding domain protein | 6 |
| | **WP_011439929.1** | 5.5E-06 | Qc.III | -GSIGQTINRLSEIAAAIASAVEQQGVATREISRNVHHAAEGTHQVTVNIAE- | *Rhodopseudomonas palustris* | Bacteria; Proteobacteria; Alphaproteobacteria; Rhizobiales; Bradyrhizobiaceae; Rhodopseudomonas | methyl-accepting chemotaxis protein | 6 |
|  | **WP_025739406.1** | 5.5E-06 | Qa.III | -NAINEQIVQISDMNTQVATATEEQSTVVNDINQNIEEINHSSQQTA------ | *Salinivibrio socompensis* | Bacteria; Proteobacteria; Gammaproteobacteria; Vibrionales; Vibrionaceae; Salinivibrio | methyl-accepting chemotaxis protein | 6 |
|  | **WP_027299294.1** | 5.5E-06 | Qc.III | IQGIGGTIGSINEIATTIAAAVEEQGAATQEIARNVQEAAQGTGQVSSN---- | *Rhodospirillales bacterium URHD0088* | Bacteria; Proteobacteria; Alphaproteobacteria; Rhodospirillales; unclassified Rhodospirillales | HAMP domain-containing protein | 6 |
|  | **WP_045671158.1** | 5.5E-06 | Qc.III | -EQVAASAGELATGANEITNDVGEMTESIREIEEKVTGYVEASK--------- | *Paenibacillus beijingensis* | Bacteria; Firmicutes; Bacilli; Bacillales; Paenibacillaceae; Paenibacillus | methyl-accepting chemotaxis protein | 6 |
|  | **WP_046073991.1** | 5.5E-06 | Qa.III | -NAINEQIVQISDMNTQVATATEEQSTVVNDINQNIEEINHSSQQTA------ | *Salinivibrio* | Bacteria; Proteobacteria; Gammaproteobacteria; Vibrionales; Vibrionaceae | MULTISPECIES: methyl-accepting chemotaxis protein | 6 |
|  | **WP_074213459.1** | 5.5E-06 | Qa.III | -NAINEQIVQISDMNTQVATATEEQSTVVNDINQNIEEINHSSQQTA------ | *Salinivibrio sp. ES.052* | Bacteria; Proteobacteria; Gammaproteobacteria; Vibrionales; Vibrionaceae; Salinivibrio | methyl-accepting chemotaxis protein | 6 |
|  | **WP_075767383.1** | 5.5E-06 | Qc.III.c | LERITESIQHISDMATQISSAAEEQRAVTDEISRNTQAVN------------ | *Photobacterium proteolyticum* | Bacteria; Proteobacteria; Gammaproteobacteria; Vibrionales; Vibrionaceae; Photobacterium | methyl-accepting chemotaxis protein | 6 |
|  | **WP_077456503.1** | 5.5E-06 | Qa.III | -NAINEQIVQISDMNTQVATATEEQSTVVNDINQNIEEINHSSQQTA------ | *Salinivibrio kushneri* | Bacteria; Proteobacteria; Gammaproteobacteria; Vibrionales; Vibrionaceae; Salinivibrio | methyl-accepting chemotaxis protein | 6 |
|  | **WP_077484268.1** | 5.5E-06 | Qa.III | -NAINEQIVQISDMNTQVATATEEQSTVVNDINQNIEEINHSSQQTA------ | *Salinivibrio sp. ML323* | Bacteria; Proteobacteria; Gammaproteobacteria; Vibrionales; Vibrionaceae; Salinivibrio | methyl-accepting chemotaxis protein | 6 |
|  | **WP_077599646.1** | 5.5E-06 | Qa.III | -NAINEQIVQISDMNTQVATATEEQSTVVNDINQNIEEINHSSQQTA------ | *Salinivibrio kushneri* | Bacteria; Proteobacteria; Gammaproteobacteria; Vibrionales; Vibrionaceae; Salinivibrio | methyl-accepting chemotaxis protein | 6 |
|  | **WP_077641630.1** | 5.5E-06 | Qa.III | -NAINEQIVQISDMNTQVATATEEQSTVVNDINQNIEEINHSSQQTA------ | *Salinivibrio sp. IB870* | Bacteria; Proteobacteria; Gammaproteobacteria; Vibrionales; Vibrionaceae; Salinivibrio | methyl-accepting chemotaxis protein | 6 |
|  | **WP_077653512.1** | 5.5E-06 | Qa.III | -NAINEQIVQISDMNTQVATATEEQSTVVNDINQNIEEINHSSQQTA------ | *Salinivibrio kushneri* | Bacteria; Proteobacteria; Gammaproteobacteria; Vibrionales; Vibrionaceae; Salinivibrio | methyl-accepting chemotaxis protein | 6 |
|  | **WP_085580556.1** | 5.5E-06 | Qc.III | -RDISEIISRTRETSGSISSAVEQQNAATQEIARNISQASAGTNEV------- | *Thalassospira mesophila* | Bacteria; Proteobacteria; Alphaproteobacteria; Rhodospirillales; Rhodospirillaceae; Thalassospira | methyl-accepting chemotaxis protein | 6 |
| | **WP_092516018.1** | 5.5E-06 | Qc.III | -REISGTIERLSEISSTIAAAVEEQGAATQEISRNVQQAAQGTQQVSSNIAD- | *Afipia sp. GAS231* | Bacteria; Proteobacteria; Alphaproteobacteria; Rhizobiales; Bradyrhizobiaceae; Afipia | HAMP domain-containing protein | 6 |
|  | **WP_102505697.1** | 5.5E-06 | Qa.III | -NAINEQIVQISDMNTQVATATEEQSTVVNDINQNIEEINHSSQQTA------ | *Salinivibrio costicola* | Bacteria; Proteobacteria; Gammaproteobacteria; Vibrionales; Vibrionaceae; Salinivibrio | methyl-accepting chemotaxis protein | 6 |
|  | **EFH09912.1** | 5.6E-06 | Qa.III | IEAITRSIDQVSHVASAIAGAVEQQGQTTRQIAAHVQQAAESNRQ-------- | *Roseomonas cervicalis ATCC 49957* | Bacteria; Proteobacteria; Alphaproteobacteria; Rhodospirillales; Acetobacteraceae; Roseomonas | methyl-accepting chemotaxis protein signaling domain protein | 6 |
|  | **OXH83240.1** | 5.6E-06 | Qa.III | --EIVAAVERVNAIMSDISTAFDEQSSGIEQVNATVTQMEQTMQ--------- | *Burkholderia multivorans* | Bacteria; Proteobacteria; Betaproteobacteria; Burkholderiales; Burkholderiaceae; Burkholderia; Burkholderia cepacia complex | hypothetical protein CA830_35185 partial | 6 |
|  | **PSQ58555.1** | 5.6E-06 | Qb.I | LAETKRQLQDTIEESETTTEELESSNEELLSMNEELQSKNEEIETSK------ | *Bacteroidetes bacterium QH_10_64_19* | Bacteria; Bacteroidetes | hypothetical protein BRD27_06215 | 6 |
|  | **WP_008193551.1** | 5.6E-06 | Qa.IV | --EIAETMTEVNSYTTAIASAVEQQGAATAEISQNVQRAAEATG--------- | *Labrenzia alexandrii* | Bacteria; Proteobacteria; Alphaproteobacteria; Rhodobacterales; Rhodobacteraceae; Labrenzia | HAMP domain-containing protein | 6 |
| | **WP_068302577.1** | 5.6E-06 | Qc.III | --AIDETISKLNQVTLAISSAVEEQEAATSEISRSVQFASEQTDKVASEI--- | *Kordiimonas lacus* | Bacteria; Proteobacteria; Alphaproteobacteria; Kordiimonadales; Kordiimonadaceae; Kordiimonas | HAMP domain-containing protein | 6 |
|  | **WP_074130979.1** | 5.6E-06 | Qa.IV | ------TIAEIDHVSGQIAAAVEQQGAATREISSNLQQAADRTRDVNK----- | *Bradyrhizobium sp. NAS96.2* | Bacteria; Proteobacteria; Alphaproteobacteria; Rhizobiales; Bradyrhizobiaceae; Bradyrhizobium | chemotaxis protein | 6 |
|  | **WP_084618739.1** | 5.6E-06 | Qc.III | --SISETISRMEEMAQAIAGAVEQQNAATGEITQNVAQATQGTQEVTR----- | *Thalassobaculum* | Bacteria; Proteobacteria; Alphaproteobacteria; Rhodospirillales; Rhodospirillaceae | MULTISPECIES: hypothetical protein | 6 |
|  | **KIT38283.1** | 5.7E-06 | Qa.III | LDEINAAIALISDMATQIATAAEEQTHVTNEITQNVTSIK------------ | *Vibrio parahaemolyticus 3644* | Bacteria; Proteobacteria; Gammaproteobacteria; Vibrionales; Vibrionaceae; Vibrio | chemotaxis protein | 6 |
|  | **OUX63257.1** | 5.7E-06 | Qa.IV | IKEIGATINLISEISSSIAAAVEEQGAATQEIARNVQYAAQ----------- | *Afipia sp. TMED4* | Bacteria; Proteobacteria; Alphaproteobacteria; Rhizobiales; Bradyrhizobiaceae; Afipia | methyl-accepting chemotaxis protein partial | 6 |
| | **WP_009027178.1** | 5.7E-06 | Qc.III | IQAISSTIERLAEIGAAIAAAVEEQGAATQEISRNVQQAAHGTQQVSSNI--- | *Bradyrhizobium sp. ORS 375* | Bacteria; Proteobacteria; Alphaproteobacteria; Rhizobiales; Bradyrhizobiaceae; Bradyrhizobium | HAMP domain-containing protein | 6 |
|  | **WP_020591301.1** | 5.7E-06 | Qa.IV | IKTIDNVIQTINEISSSIASAVEQQGAATKEIARNIQEAA------------ | *Kiloniella laminariae* | Bacteria; Proteobacteria; Alphaproteobacteria; Kiloniellales; Kiloniellaceae; Kiloniella | hypothetical protein | 6 |
|  | **WP_092375052.1** | 5.7E-06 | Qa.III | IDRIVNVIGEVNSIVGNIAAAIEEQSAVTREVANNIAQATEDVQDAN------ | *Desulfomicrobium apsheronum* | Bacteria; Proteobacteria; Deltaproteobacteria; Desulfovibrionales; Desulfomicrobiaceae; Desulfomicrobium | methyl-accepting chemotaxis protein | 6 |
|  | **WP_107282121.1** | 5.7E-06 | Qa.III | LEQINNSVALISDMATQIATAAEEQTQVTDEITQNTTSIK------------ | *Photobacterium lipolyticum* | Bacteria; Proteobacteria; Gammaproteobacteria; Vibrionales; Vibrionaceae; Photobacterium | methyl-accepting chemotaxis protein | 6 |
|  | **BAU89355.1** | 5.8E-06 | Qa.IV | IRAIVSTITDLSEISMSVSSAVTEQAAVTQEIATNMQTAA------------ | *Methylorubrum populi* | Bacteria; Proteobacteria; Alphaproteobacteria; Rhizobiales; Methylobacteriaceae; Methylorubrum | methyl-accepting chemotaxis receptor/ sensory transducer | 6 |
|  | **KPQ09297.1** | 5.8E-06 | Qc.III | --DIGGVIDRMKEISTAISAAMEEQGAATNEIARNVQEAAKGTGEVT------ | *Salinarimonadaceae bacterium HL-109* | Bacteria; Proteobacteria; Alphaproteobacteria; Rhizobiales; Salinarimonadaceae | chemotaxis signal relay system methyl-accepting signal transducer | 6 |
|  | **PSQ73689.1** | 5.8E-06 | Qb.I | LAETKRQLQDTIEESETTTEELESSNEELLSMNEELQSKNEEIETSK------ | *Bacteroidetes bacterium QH_9_64_21* | Bacteria; Bacteroidetes | hypothetical protein BRD39_03690 | 6 |
|  | **WP_016523904.1** | 5.8E-06 | SNAP.c | LNQITGNIENVKDRVMVQHTEVNETAAKIDAITQTMSSLDFHIDNQAK----- | *Treponema medium* | Bacteria; Spirochaetes; Spirochaetales; Spirochaetaceae; Treponema | methyl-accepting chemotaxis protein | 6 |
|  | **WP_039030708.1** | 5.8E-06 | R.IV | IQEINRAVLNLDSMVQQNAELVVESAAAASALQGQAGDLAETAGHFR------ | *Leclercia adecarboxylata* | Bacteria; Proteobacteria; Gammaproteobacteria; Enterobacterales; Enterobacteriaceae; Leclercia | methyl-accepting chemotaxis protein | 6 |
|  | **WP_044828549.1** | 5.8E-06 | Qa.IV | IRTIVESVQGINEILKLTAESVMEQGTATGQISTSIKEAAGY---------- | *Thalassospira sp. HJ* | Bacteria; Proteobacteria; Alphaproteobacteria; Rhodospirillales; Rhodospirillaceae; Thalassospira | methyl-accepting chemotaxis protein | 6 |
|  | **WP_056913828.1** | 5.8E-06 | Qc.III | --SISQSIARIHTVARAITEAVSVQSEATQRIAQTVEGAAVRTRQV------- | *Pseudolabrys sp. Root1462* | Bacteria; Proteobacteria; Alphaproteobacteria; Rhizobiales; Xanthobacteraceae; Pseudolabrys | HAMP domain-containing protein | 6 |
|  | **WP_084194509.1** | 5.8E-06 | Qa.IV | IRDITRQITEINGISTAVAAAVEEQEAATAEISRNVQETS------------ | *Insolitispirillum peregrinum* | Bacteria; Proteobacteria; Alphaproteobacteria; Rhodospirillales; Rhodospirillaceae; Insolitispirillum | HAMP domain-containing protein | 6 |
|  | **WP_085905251.1** | 5.8E-06 | Qa.III.b | -----STINKVNDITKQIAGVAQEQEEATQEISRNIQQASQGTQEVSSSI--- | *Kiloniella majae* | Bacteria; Proteobacteria; Alphaproteobacteria; Kiloniellales; Kiloniellaceae; Kiloniella | methyl-accepting chemotaxis protein | 6 |
|  | **WP_103041624.1** | 5.8E-06 | Qa.III | IRAIAGTIAEINEIASAIAAAVEQQGAATREIARNVQEAA------------ | *Azospirillum brasilense* | Bacteria; Proteobacteria; Alphaproteobacteria; Rhodospirillales; Rhodospirillaceae; Azospirillum | urea ABC transporter substrate-binding protein partial | 6 |
|  | **OZA84465.1** | 5.9E-06 | Qa.IV | --KIVTIISDIRQATSAIASAIEEQGAATREIAYNTQSAAQGTQGVND----- | *Azorhizobium sp. 39-67-5* | Bacteria; Proteobacteria; Alphaproteobacteria; Rhizobiales; Xanthobacteraceae; Azorhizobium | hypothetical protein B7X76_06745 | 6 |
|  | **SCZ28747.1** | 5.9E-06 | Qa.III | IESITGTIKQIDEISSTIASSVEEQGSATGEIAQNVQQAAQGTQE-------- | *Afifella marina DSM 2698* | Bacteria; Proteobacteria; Alphaproteobacteria; Rhizobiales; Rhodobiaceae; Afifella | methyl-accepting chemotaxis sensory transducer | 6 |
|  | **WP_009539221.1** | 5.9E-06 | Qa.IV | IKSVTSTIDQINEIAGAIASAVEEQGAATQEIARNTQQAA------------ | *Caenispirillum salinarum* | Bacteria; Proteobacteria; Alphaproteobacteria; Rhodospirillales; Rhodospirillaceae; Caenispirillum | methyl-accepting chemotaxis protein | 6 |
|  | **WP_027549664.1** | 5.9E-06 | Qa.IV | IRDIGQTIGEINDVTTAIAAAVEQQGAATREIARNIQHAA------------ | *Bradyrhizobium sp. Cp5.3* | Bacteria; Proteobacteria; Alphaproteobacteria; Rhizobiales; Bradyrhizobiaceae; Bradyrhizobium | HAMP domain-containing protein | 6 |
|  | **WP_048747327.1** | 5.9E-06 | Qc.III | -EQVASSAGELATGANEISGEVGDMTDSIRDIETKVSAYAES---------- | *Paenibacillus* | Bacteria; Firmicutes; Bacilli; Bacillales; Paenibacillaceae | MULTISPECIES: methyl-accepting chemotaxis protein | 6 |
|  | **WP_063680077.1** | 5.9E-06 | Qa.IV | IRDIGQTIGEINDVTTAIAAAVEQQGAATREIARNIQHAA------------ | *Bradyrhizobium neotropicale* | Bacteria; Proteobacteria; Alphaproteobacteria; Rhizobiales; Bradyrhizobiaceae; Bradyrhizobium | HAMP domain-containing protein | 6 |

| | | | | | | | | |
|---|---|---|---|---|---|---|---|---|
|  | **ADZ69890.1** | 6E-06 | Qa.III | IQSIGATIAEINEIASSIASAVEEQSAATGEIARNVEQAAS------------ | *Polymorphum gilvum SL003B-26A1* | Bacteria; Proteobacteria; Alphaproteobacteria; Polymorphum | Methyl-accepting chemotaxis sensory transducer | 6 |
|  | **OCA61807.1** | 6E-06 | Qa.III | LEQITASINQISDMATQISSAAEEQRAVTDEVSRNIQAVK------------- | *Aeromonas piscicola* | Bacteria; Proteobacteria; Gammaproteobacteria; Aeromonadales; Aeromonadaceae; Aeromonas | chemotaxis protein | 6 |
|  | **OOD34295.1** | 6E-06 | Qa.III | LEQITASINQISDMATQISSAAEEQRAVTDEVSRNIQAVK------------- | *Aeromonas hydrophila* | Bacteria; Proteobacteria; Gammaproteobacteria; Aeromonadales; Aeromonadaceae; Aeromonas | methyl-accepting chemotaxis protein | 6 |
|  | **OSO91254.1** | 6E-06 | Qa.III | LEQITASINQISDMATQISSAAEEQRAVTDEVSRNIQAVK------------- | *Aeromonas hydrophila* | Bacteria; Proteobacteria; Gammaproteobacteria; Aeromonadales; Aeromonadaceae; Aeromonas | chemotaxis protein | 6 |
|  | **OXH83459.1** | 6E-06 | Qa.III | --EIVAAVERVNAIMSDISTAFDEQSSGIEQVNATVTQMEQTMQ--------- | *Burkholderia multivorans* | Bacteria; Proteobacteria; Betaproteobacteria; Burkholderiales; Burkholderiaceae; Burkholderia; Burkholderia cepacia complex | hypothetical protein CA831_33540 partial | 6 |
|  | **PCS23773.1** | 6E-06 | Qa.III | LEGIAGMMQEINDMNMQIATAASEQSSVTEEITENVNNIA------------- | *bacterium symbiont of Melanocetus johnsonii* | Bacteria | Methyl-accepting chemotaxis protein | 6 |
|  | **PKD23526.1** | 6E-06 | Qa.III | LEQITASINQISDMATQISSAAEEQRAVTDEVSRNIQAVK------------- | *Aeromonas hydrophila* | Bacteria; Proteobacteria; Gammaproteobacteria; Aeromonadales; Aeromonadaceae; Aeromonas | methyl-accepting chemotaxis protein | 6 |
|  | **PNO61289.1** | 6E-06 | Qa.III | LEQITASINQISDMATQISSAAEEQRAVTDEVSRNIQAVK------------- | *Aeromonas hydrophila* | Bacteria; Proteobacteria; Gammaproteobacteria; Aeromonadales; Aeromonadaceae; Aeromonas | methyl-accepting chemotaxis protein | 6 |
|  | **PPA30742.1** | 6E-06 | Qa.III | LEQITASINQISDMATQISSAAEEQRAVTDEVSRNIQAVK------------- | *Aeromonas jandaei* | Bacteria; Proteobacteria; Gammaproteobacteria; Aeromonadales; Aeromonadaceae; Aeromonas | methyl-accepting chemotaxis protein | 6 |
|  | **WP_005298343.1** | 6E-06 | Qa.III | LEQITASINQISDMATQISSAAEEQRAVTDEVSRNIQAVK------------- | *Aeromonas* | Bacteria; Proteobacteria; Gammaproteobacteria; Aeromonadales; Aeromonadaceae | MULTISPECIES: methyl-accepting chemotaxis protein | 6 |
|  | **WP_017764984.1** | 6E-06 | Qa.III | LEQITASINQISDMATQISSAAEEQRAVTDEVSRNIQAVK------------- | *Aeromonas* | Bacteria; Proteobacteria; Gammaproteobacteria; Aeromonadales; Aeromonadaceae | MULTISPECIES: methyl-accepting chemotaxis protein | 6 |
|  | **WP_017781600.1** | 6E-06 | Qa.III | LEQITASINQISDMATQISSAAEEQRAVTDEVSRNIQAVK------------- | *Aeromonas* | Bacteria; Proteobacteria; Gammaproteobacteria; Aeromonadales; Aeromonadaceae | MULTISPECIES: methyl-accepting chemotaxis protein | 6 |
|  | **WP_024944206.1** | 6E-06 | Qa.III | LEQITASINQISDMATQISSAAEEQRAVTDEVSRNIQAVK------------- | *Aeromonas hydrophila* | Bacteria; Proteobacteria; Gammaproteobacteria; Aeromonadales; Aeromonadaceae; Aeromonas | methyl-accepting chemotaxis protein | 6 |
|  | **WP_027694369.1** | 6E-06 | Qa.III | LEEILTSVGQISDMATQIATAAEEQTHVTGEITQNVTAIKD------------ | *Vibrio gazogenes* | Bacteria; Proteobacteria; Gammaproteobacteria; Vibrionales; Vibrionaceae; Vibrio | methyl-accepting chemotaxis protein | 6 |
|  | **WP_029299351.1** | 6E-06 | Qa.III | LEQITASINQISDMATQISSAAEEQRAVTDEVSRNIQAVK------------- | *Aeromonas* | Bacteria; Proteobacteria; Gammaproteobacteria; Aeromonadales; Aeromonadaceae | MULTISPECIES: methyl-accepting chemotaxis protein | 6 |
|  | **WP_029304221.1** | 6E-06 | Qa.III | LEQITASINQISDMATQISSAAEEQRAVTDEVSRNIQAVK------------- | *Aeromonas hydrophila* | Bacteria; Proteobacteria; Gammaproteobacteria; Aeromonadales; Aeromonadaceae; Aeromonas | methyl-accepting chemotaxis protein | 6 |
|  | **WP_033137549.1** | 6E-06 | Qa.III | LEQITASINQISDMATQISSAAEEQRAVTDEVSRNIQAVK------------- | *Aeromonas finlandiensis* | Bacteria; Proteobacteria; Gammaproteobacteria; Aeromonadales; Aeromonadaceae; Aeromonas | methyl-accepting chemotaxis protein | 6 |
|  | **WP_037264583.1** | 6E-06 | Qc.III | IQSIGGTIGSINEIATTIASAVEEQGAATQEIARNVQEAAQGTGQVSSN---- | *Rhodospirillales bacterium URHD0088* | Bacteria; Proteobacteria; Alphaproteobacteria; Rhodospirillales; unclassified Rhodospirillales | methyl-accepting chemotaxis protein | 6 |
|  | **WP_037338794.1** | 6E-06 | Qb.III.d | --SSEQLASSSEELSAvtdDSTQNLRRQNEEIEQAATAVNELTA------------ | *Saccharospirillum impatiens* | Bacteria; Proteobacteria; Gammaproteobacteria; Oceanospirillales; Saccharospirillaceae; Saccharospirillum | methyl-accepting chemotaxis protein | 6 |
|  | **WP_039215687.1** | 6E-06 | Qa.III | LEQITASINQISDMATQISSAAEEQRAVTDEVSRNIQAVK------------- | *Aeromonas hydrophila* | Bacteria; Proteobacteria; Gammaproteobacteria; Aeromonadales; Aeromonadaceae; Aeromonas | methyl-accepting chemotaxis protein | 6 |
|  | **WP_041210005.1** | 6E-06 | Qa.III | LEQITASINQISDMATQISSAAEEQRAVTDEVSRNIQAVK------------- | *Aeromonas* | Bacteria; Proteobacteria; Gammaproteobacteria; Aeromonadales; Aeromonadaceae | MULTISPECIES: methyl-accepting chemotaxis protein | 6 |
|  | **WP_041218279.1** | 6E-06 | Qa.III | LEQITASINQISDMATQISSAAEEQRAVTDEVSRNIQAVK------------- | *Aeromonas hydrophila* | Bacteria; Proteobacteria; Gammaproteobacteria; Aeromonadales; Aeromonadaceae; Aeromonas | methyl-accepting chemotaxis protein | 6 |
|  | **WP_041993033.1** | 6E-06 | Qa.III | LEQITASINQISDMATQISSAAEEQRAVTDEVSRNIQAVK------------- | *Aeromonas bivalvium* | Bacteria; Proteobacteria; Gammaproteobacteria; Aeromonadales; Aeromonadaceae; Aeromonas | methyl-accepting chemotaxis protein | 6 |
|  | **WP_042029782.1** | 6E-06 | Qa.III | LEQITASINQISDMATQISSAAEEQRAVTDEVSRNIQAVK------------- | *Aeromonas jandaei* | Bacteria; Proteobacteria; Gammaproteobacteria; Aeromonadales; Aeromonadaceae; Aeromonas | methyl-accepting chemotaxis protein | 6 |
|  | **WP_042034673.1** | 6E-06 | Qa.III | LEQITASINQISDMATQISSAAEEQRAVTDEVSRNIQAVK------------- | *Aeromonas popoffii* | Bacteria; Proteobacteria; Gammaproteobacteria; Aeromonadales; Aeromonadaceae; Aeromonas | methyl-accepting chemotaxis protein | 6 |
|  | **WP_042040875.1** | 6E-06 | Qa.III | LEQITASINQISDMATQISSAAEEQRAVTDEVSRNIQAVK------------- | *Aeromonas rivuli* | Bacteria; Proteobacteria; Gammaproteobacteria; Aeromonadales; Aeromonadaceae; Aeromonas | methyl-accepting chemotaxis protein | 6 |
|  | **WP_042065018.1** | 6E-06 | Qa.III | LEQITASINQISDMATQISSAAEEQRAVTDEVSRNIQAVK------------- | *Aeromonas hydrophila* | Bacteria; Proteobacteria; Gammaproteobacteria; Aeromonadales; Aeromonadaceae; Aeromonas | methyl-accepting chemotaxis protein | 6 |
|  | **WP_042867668.1** | 6E-06 | Qa.III | LEQITASINQISDMATQISSAAEEQRAVTDEVSRNIQAVK------------- | *Aeromonas piscicola* | Bacteria; Proteobacteria; Gammaproteobacteria; Aeromonadales; Aeromonadaceae; Aeromonas | methyl-accepting chemotaxis protein | 6 |
|  | **WP_043135693.1** | 6E-06 | Qa.III | LEQITASINQISDMATQISSAAEEQRAVTDEVSRNIQAVK------------- | *Aeromonas salmonicida* | Bacteria; Proteobacteria; Gammaproteobacteria; Aeromonadales; Aeromonadaceae; Aeromonas | methyl-accepting chemotaxis protein | 6 |
|  | **WP_043161447.1** | 6E-06 | Qa.III | LEQITASINQISDMATQISSAAEEQRAVTDEVSRNIQAVK------------- | *Aeromonas hydrophila* | Bacteria; Proteobacteria; Gammaproteobacteria; Aeromonadales; Aeromonadaceae; Aeromonas | methyl-accepting chemotaxis protein | 6 |
|  | **WP_043163647.1** | 6E-06 | Qa.III | LEQITASINQISDMATQISSAAEEQRAVTDEVSRNIQAVK------------- | *Aeromonas hydrophila* | Bacteria; Proteobacteria; Gammaproteobacteria; Aeromonadales; Aeromonadaceae; Aeromonas | methyl-accepting chemotaxis protein | 6 |
|  | **WP_043171198.1** | 6E-06 | Qa.III | LEQITASINQISDMATQISSAAEEQRAVTDEVSRNIQAVK------------- | *Aeromonas* | Bacteria; Proteobacteria; Gammaproteobacteria; Aeromonadales; Aeromonadaceae | MULTISPECIES: methyl-accepting chemotaxis protein | 6 |
|  | **WP_043555132.1** | 6E-06 | Qa.III | LEQITASINQISDMATQISSAAEEQRAVTDEVSRNIQAVK------------- | *Aeromonas bestiarum* | Bacteria; Proteobacteria; Gammaproteobacteria; Aeromonadales; Aeromonadaceae; Aeromonas | methyl-accepting chemotaxis protein | 6 |
|  | **WP_044800561.1** | 6E-06 | Qa.III | LEQITASINQISDMATQISSAAEEQRAVTDEVSRNIQAVK------------- | *Aeromonas hydrophila* | Bacteria; Proteobacteria; Gammaproteobacteria; Aeromonadales; Aeromonadaceae; Aeromonas | methyl-accepting chemotaxis protein | 6 |
|  | **WP_053372094.1** | 6E-06 | Qc.III | -QQVAASANELATGANEISEDVSEMSESITEIEEKVSDYATSTKEMNTR---- | *Bacillales* | Bacteria; Firmicutes; Bacilli | MULTISPECIES: methyl-accepting chemotaxis protein | 6 |
|  | **WP_072954518.1** | 6E-06 | Qa.III | LEEILTSVGQISDMATQIATAAEEQTHVTGEITQNVTAIKD------------ | *Vibrio gazogenes* | Bacteria; Proteobacteria; Gammaproteobacteria; Vibrionales; Vibrionaceae; Vibrio | methyl-accepting chemotaxis protein | 6 |
|  | **WP_076267493.1** | 6E-06 | Qc.III | -QQVAASANELATGANEISEDVSEMSESITEIEEKVSDYATSTKEMNTR---- | *Paenibacillus sp. FSL A5-0031* | Bacteria; Firmicutes; Bacilli; Bacillales; Paenibacillaceae; Paenibacillus | methyl-accepting chemotaxis protein | 6 |
|  | **WP_076360364.1** | 6E-06 | Qa.III | LEQITASINQISDMATQISSAAEEQRAVTDEVSRNIQAVK------------- | *Aeromonas* | Bacteria; Proteobacteria; Gammaproteobacteria; Aeromonadales; Aeromonadaceae | MULTISPECIES: methyl-accepting chemotaxis protein | 6 |
| | **WP_079589945.1** | 6E-06 | Qb.I | --NIKSLVDDTVENTQKSSENIKQSSNLVETASETFDKIYNQVNQSNVLV--- | *Acetoanaerobium noterae* | Bacteria; Firmicutes; Clostridia; Clostridiales; Peptostreptococcaceae; Acetoanaerobium | methyl-accepting chemotaxis protein | 6 |
| | **WP_082509100.1** | 6E-06 | Qc.III.c | --GISISIRSINETSTAIASAVEEQEAATREIVQAIAQASSGIGEVTANIT-- | *Methylobacterium sp. Leaf121* | Bacteria; Proteobacteria; Alphaproteobacteria; Rhizobiales; Methylobacteriaceae; Methylobacterium | hypothetical protein | 6 |
|  | **WP_098982958.1** | 6E-06 | Qa.III | LEQITASINQISDMATQISSAAEEQRAVTDEVSRNIQAVK------------- | *Aeromonas sp. CA23* | Bacteria; Proteobacteria; Gammaproteobacteria; Aeromonadales; Aeromonadaceae; Aeromonas | methyl-accepting chemotaxis protein | 6 |
|  | **WP_101614087.1** | 6E-06 | Qa.III | LEQITASINQISDMATQISSAAEEQRAVTDEVSRNIQAVK------------- | *Aeromonas hydrophila* | Bacteria; Proteobacteria; Gammaproteobacteria; Aeromonadales; Aeromonadaceae; Aeromonas | methyl-accepting chemotaxis protein | 6 |
|  | **WP_103470979.1** | 6E-06 | Qa.III | LEQITASINQISDMATQISSAAEEQRAVTDEVSRNIQAVK------------- | *Aeromonas bestiarum* | Bacteria; Proteobacteria; Gammaproteobacteria; Aeromonadales; Aeromonadaceae; Aeromonas | methyl-accepting chemotaxis protein | 6 |
|  | **WP_106553145.1** | 6E-06 | Qa.III | LEQITASINQISDMATQISSAAEEQRAVTDEVSRNIQAVK------------- | *Aeromonas hydrophila* | Bacteria; Proteobacteria; Gammaproteobacteria; Aeromonadales; Aeromonadaceae; Aeromonas | methyl-accepting chemotaxis protein | 6 |
|  | **WP_108570804.1** | 6E-06 | Qa.III | LEQITASINQISDMATQISSAAEEQRAVTDEVSRNIQAVK------------- | *Aeromonas sp. HMWF016* | Bacteria; Proteobacteria; Gammaproteobacteria; Aeromonadales; Aeromonadaceae; Aeromonas | methyl-accepting chemotaxis protein | 6 |
|  | **WP_111328835.1** | 6E-06 | Qa.III | IESITGTIKQIDEISSTIASSVEEQGSATGEIAQNVQQAAQGTQE-------- | *Afifella marina* | Bacteria; Proteobacteria; Alphaproteobacteria; Rhizobiales; Rhodobiaceae; Afifella | methyl-accepting chemotaxis protein | 6 |
|  | **WP_111874046.1** | 6E-06 | Qa.III | LEQITASINQISDMATQISSAAEEQRAVTDEVSRNIQAVK------------- | *Aeromonas* | Bacteria; Proteobacteria; Gammaproteobacteria; Aeromonadales; Aeromonadaceae | MULTISPECIES: HAMP domain-containing protein | 6 |
|  | **WP_113994187.1** | 6E-06 | Qa.III | LEQITASINQISDMATQISSAAEEQRAVTDEVSRNIQAVK------------- | *Aeromonas hydrophila* | Bacteria; Proteobacteria; Gammaproteobacteria; Aeromonadales; Aeromonadaceae; Aeromonas | HAMP domain-containing protein | 6 |
|  | **SHE39118.1** | 6.1E-06 | Qa.III | LEEILTSVGQISDMATQIATAAEEQTHVTGEITQNVTAIKD------------ | *Vibrio gazogenes DSM 21264 = NBRC 103151* | Bacteria; Proteobacteria; Gammaproteobacteria; Vibrionales; Vibrionaceae; Vibrio | methyl-accepting chemotaxis sensory transducer with Cache sensor | 6 |
|  | **WP_011925119.1** | 6.1E-06 | Qa.IV | IRAISTTIEKLSEISSAIAAAVEEQGAATQEISRNVQQAS------------- | *Bradyrhizobium sp. ORS 278* | Bacteria; Proteobacteria; Alphaproteobacteria; Rhizobiales; Bradyrhizobiaceae; Bradyrhizobium | HAMP domain-containing protein | 6 |
|  | **WP_114395831.1** | 6.1E-06 | Qa.III | IQGIAQVIGRVNDVATSIASAVEEQGAATREISRNVQQAAQ------------ | *Rhodospirillaceae bacterium NAU-10* | Bacteria; Proteobacteria; Alphaproteobacteria; Rhodospirillales; Rhodospirillaceae | HAMP domain-containing protein | 6 |
| | **WP_043949872.1** | 6.2E-06 | Qc.III | IESIAKTVDELNGISTSISSAVEEQSAATGEIARNVQEAAGGTDEVSRSLT-- | *Candidatus Phaeomarinobacter ectocarpi* | Bacteria; Proteobacteria; Alphaproteobacteria; Rhizobiales; Rhodobiaceae; Candidatus Phaeomarinobacter | HAMP domain-containing protein | 6 |
| | **WP_057757271.1** | 6.2E-06 | Qc.III | -KGISDTIEKLSDISSTIAAAVEEQGAATQEISRNVQKAAEGTHQVSANIT-- | *Bradyrhizobium manausense* | Bacteria; Proteobacteria; Alphaproteobacteria; Rhizobiales; Bradyrhizobiaceae; Bradyrhizobium | methyl-accepting chemotaxis protein | 6 |
|  | **WP_094302373.1** | 6.2E-06 | Qa.III | IRAIAGTIAEINEIASAIAAAVEQQGAATREIARNVQEAA------------- | *Azospirillum brasilense* | Bacteria; Proteobacteria; Alphaproteobacteria; Rhodospirillales; Rhodospirillaceae; Azospirillum | urea ABC transporter substrate-binding protein partial | 6 |
|  | **WP_096483891.1** | 6.2E-06 | Qa.IV | IRAIVSTITDLSEISMSVSSAVTEQAAVTQEIATNMQTAA------------- | *Methylorubrum populi* | Bacteria; Proteobacteria; Alphaproteobacteria; Rhizobiales; Methylobacteriaceae; Methylorubrum | PAS domain S-box protein | 6 |
|  | **WP_097355955.1** | 6.2E-06 | Qa.III | LEGIAGMMQEINDMNMQIATAASEQSSVTEEITENVNNIA------------- | *bacterium symbiont of Melanocetus johnsonii* | Bacteria | methyl-accepting chemotaxis protein | 6 |
|  | **WP_100416203.1** | 6.2E-06 | Qa.IV | IRNISTTITEINDVTAAIAAAVEEQGAATREIARNIQHAA------------- | *Bradyrhizobium lablabi* | Bacteria; Proteobacteria; Alphaproteobacteria; Rhizobiales; Bradyrhizobiaceae; Bradyrhizobium | HAMP domain-containing protein | 6 |

| | | | | | | | | |
|---|---|---|---|---|---|---|---|---|
| | WP_111417916.1 | 6.2E-06 | Qc.III | IKEIGGTIGQIAEIASAIAAAVEQQGAATGEIARNVQQAAKGTDEVARHITD- | *Rhodoplanes roseus* | Bacteria; Proteobacteria; Alphaproteobacteria; Rhizobiales; Hyphomicrobiaceae; Rhodoplanes | methyl-accepting chemotaxis protein | 6 |
|  | PIE61214.1 | 6.3E-06 | Qa.III | IKRIANVISNVNEIVDSIATAVNDQTNVTNDIAMNISQISE----------- | *Desulfobacterales bacterium* | Bacteria; Proteobacteria; Deltaproteobacteria; Desulfobacterales | methyl-accepting chemotaxis protein | 6 |
| | PIE67576.1 | 6.3E-06 | Qa.IV | --QINTVINDVNTIVTTIASAVKEQAASTDEIASNVAQAAQGIGEVNEHVAQ- | *Deltaproteobacteria bacterium* | Bacteria; Proteobacteria; Deltaproteobacteria | chemotaxis protein | 6 |
| | WP_011925489.1 | 6.3E-06 | Qc.III | IQAISATIERLAEIGAAIAAAVEEQGAATQEISRNVQQAAHGTQQVSSNI--- | *Bradyrhizobium sp. ORS 278* | Bacteria; Proteobacteria; Alphaproteobacteria; Rhizobiales; Bradyrhizobiaceae; Bradyrhizobium | HAMP domain-containing protein | 6 |
|  | WP_020527780.1 | 6.3E-06 | Qc.III | -------TVNLLQESQKMSEELQLQQEILRQNAEEMEATQEELKRTNHRLE-- | *Flexithrix dorotheae* | Bacteria; Bacteroidetes; Cytophagia; Cytophagales; Flammeovirgaceae; Flexithrix | PAS domain S-box protein | 6 |
|  | WP_027301045.1 | 6.3E-06 | Qa.IV | MKTIATTIATVSEVSSAIAAAVEQQGAAVREIARNTQEAA------------- | *Rhodospirillales bacterium URHD0088* | Bacteria; Proteobacteria; Alphaproteobacteria; Rhodospirillales; unclassified Rhodospirillales | HAMP domain-containing protein | 6 |
| | WP_029005916.1 | 6.3E-06 | Qc.III | IEGVGQQIRQMSDVASGISAAVEEQGIATREIVRNVDQAATGTNSVNGHIAE- | *Azorhizobium doebereinerae* | Bacteria; Proteobacteria; Alphaproteobacteria; Rhizobiales; Xanthobacteraceae; Azorhizobium | HAMP domain-containing protein | 6 |
|  | WP_049895880.1 | 6.3E-06 | Qb.III | -KESAKSAHDVEDFSAGVADKLTEQNEQMARVQDAMKEIEDK----------- | *Oribacterium sp. NK2B42* | Bacteria; Firmicutes; Clostridia; Clostridiales; Lachnospiraceae; Oribacterium | hypothetical protein | 6 |
|  | WP_068517790.1 | 6.3E-06 | Qa.III | IEGITSIIANLNEIATEVAAAVEEQGQATTEISNSVQQAVQSTQE-------- | *Thalassospira* | Bacteria; Proteobacteria; Alphaproteobacteria; Rhodospirillales; Rhodospirillaceae | MULTISPECIES: methyl-accepting chemotaxis protein | 6 |
| | RAI45253.1 | 6.4E-06 | Qc.III | IKEIGGTIGQIAEIASAIAAAVEQQGAATGEIARNVQQAAKGTDEVARHITD- | *Rhodoplanes roseus* | Bacteria; Proteobacteria; Alphaproteobacteria; Rhizobiales; Hyphomicrobiaceae; Rhodoplanes | hypothetical protein CH341_04895 | 6 |
|  | WP_063303586.1 | 6.4E-06 | Qc.III.c | ---IGETIEQINSLSTGINEAVGDQKQATEDISRDLNSAHQNVSQV------- | *Pseudovibrio sp. W64* | Bacteria; Proteobacteria; Alphaproteobacteria; Rhodobacterales; Rhodobacteraceae; Pseudovibrio | methyl-accepting chemotaxis protein | 6 |
|  | WP_068148975.1 | 6.4E-06 | SNAP.b | ----VQKMDEVYSDLVNSQTELENKNEQLKKAYEELDDAHKKLTLAQERL--- | *Kordiimonas lipolytica* | Bacteria; Proteobacteria; Alphaproteobacteria; Kordiimonadales; Kordiimonadaceae; Kordiimonas | sensor histidine kinase | 6 |
|  | WP_077646579.1 | 6.4E-06 | Qc.III | -DEISSNLSNIKTLSSDMHEAIDQLGPIVSDLKDNVDALNGA----------- | *Salinivibrio sp. IB643* | Bacteria; Proteobacteria; Gammaproteobacteria; Vibrionales; Vibrionaceae; Salinivibrio | methyl-accepting chemotaxis protein | 6 |
|  | WP_081863097.1 | 6.4E-06 | Qa.III | IRAIAGTIAEINEIASAIAAAVEQQGAATREIARNVQEAA------------- | *Azospirillum brasilense* | Bacteria; Proteobacteria; Alphaproteobacteria; Rhodospirillales; Rhodospirillaceae; Azospirillum | urea ABC transporter substrate-binding protein | 6 |
|  | WP_114097766.1 | 6.4E-06 | Qa.III.b | ----------ITEMANTISTAVGEQAEATDEIAQNVEQAAIGTQEVSSNIE-- | *Thalassospira profundimaris* | Bacteria; Proteobacteria; Alphaproteobacteria; Rhodospirillales; Rhodospirillaceae; Thalassospira | HAMP domain-containing protein | 6 |
| | WP_011385443.1 | 6.5E-06 | Qc.III | --GISHTISRINELATAIAGAVEEQGAATAEIARNVELANEGTSEVARNI--- | *Magnetospirillum magneticum* | Bacteria; Proteobacteria; Alphaproteobacteria; Rhodospirillales; Rhodospirillaceae; Magnetospirillum | methyl-accepting chemotaxis protein | 6 |
|  | WP_114096100.1 | 6.5E-06 | Qa.III | IDGITSIIANLNEIASEVAAAVEEQGQATTEISNSVQQAVQSTQE-------- | *Thalassospira profundimaris* | Bacteria; Proteobacteria; Alphaproteobacteria; Rhodospirillales; Rhodospirillaceae; Thalassospira | HAMP domain-containing protein | 6 |
|  | WP_114100701.1 | 6.5E-06 | Qa.III | IDGITSIIANLNEIASEVAAAVEEQGQATTEISNSVQQAVQSTQE-------- | *Thalassospira profundimaris* | Bacteria; Proteobacteria; Alphaproteobacteria; Rhodospirillales; Rhodospirillaceae; Thalassospira | HAMP domain-containing protein | 6 |
|  | AUD61363.1 | 6.6E-06 | Qa.III | LENILNAISTISRMSEQIARAVNEQTSVVNSVSESVLIIKSNAQ--------- | *Shewanella sp. Pdp11* | Bacteria; Proteobacteria; Gammaproteobacteria; Alteromonadales; Shewanellaceae; Shewanella | hypothetical protein AYJ58_18615 | 6 |
|  | OOF08383.1 | 6.6E-06 | Qc.III | -DEISSNLANIKTLSGDMNEAIDQLGPIVSXLKDNVDSLSSAIKH-------- | *Salinivibrio sp. PR5* | Bacteria; Proteobacteria; Gammaproteobacteria; Vibrionales; Vibrionaceae; Salinivibrio | methyl-accepting chemotaxis protein partial | 6 |
| | OUR79443.1 | 6.6E-06 | Qc.III | -EGIRKTISEMNDIAIMITSAVDEQGVATNEISRSVQEAASGTQNVSQSID-- | *Alphaproteobacteria bacterium 46_93_T64* | Bacteria; Proteobacteria; Alphaproteobacteria | hypothetical protein A9Q83_03290 | 6 |
|  | SNY91698.1 | 6.6E-06 | Qa.III | IQEIGTTIGDMNAIASSIASAVEQQSSATGEIAQNVEQAA------------- | *Cohaesibacter sp. ES.047* | Bacteria; Proteobacteria; Alphaproteobacteria; Rhizobiales; Cohaesibacteraceae | methyl-accepting chemotaxis sensory transducer with Cache sensor | 6 |
|  | WP_009027013.1 | 6.6E-06 | Qa.IV | -----GTIRQISTIASSIATAVEQQGGATQEIAKNVQSVAAGTQAAASDI--- | *Bradyrhizobium sp. ORS 375* | Bacteria; Proteobacteria; Alphaproteobacteria; Rhizobiales; Bradyrhizobiaceae; Bradyrhizobium | methyl-accepting chemotaxis protein | 6 |
|  | WP_015664643.1 | 6.6E-06 | Qa.IV | IREISETIERLSEISATIAAAVEQQGAATQEISRNVQQAAQ------------ | *Bradyrhizobium oligotrophicum* | Bacteria; Proteobacteria; Alphaproteobacteria; Rhizobiales; Bradyrhizobiaceae; Bradyrhizobium | methyl-accepting chemotaxis protein | 6 |
| | WP_028165071.1 | 6.6E-06 | Qc.III | -KGIGSIIGEVNEVATAIAAAVQEQGAATQEITRSTQFAAQGTKNVSENIT-- | *Bradyrhizobium elkanii* | Bacteria; Proteobacteria; Alphaproteobacteria; Rhizobiales; Bradyrhizobiaceae; Bradyrhizobium | HAMP domain-containing protein | 6 |
| | WP_079538820.1 | 6.6E-06 | Qc.III | -KGIGSIIGEVNEVATAIAAAVQEQGAATQEITRSTQFAAQGTKNVSENIT-- | *Bradyrhizobium lablabi* | Bacteria; Proteobacteria; Alphaproteobacteria; Rhizobiales; Bradyrhizobiaceae; Bradyrhizobium | methyl-accepting chemotaxis protein | 6 |
|  | OJI46220.1 | 6.7E-06 | Qc.III | LENIMASVQRINDMSTQIATAAEEQSLVTEEITRNTQAVNDVSN--------- | *Vibrio vulnificus* | Bacteria; Proteobacteria; Gammaproteobacteria; Vibrionales; Vibrionaceae; Vibrio | Methyl-accepting chemotaxis protein PctC | 6 |
|  | WP_017422381.1 | 6.7E-06 | Qc.III | LENIMASVQRINDMSTQIATAAEEQSLVTEEITRNTQAVNDVSN--------- | *Vibrio vulnificus* | Bacteria; Proteobacteria; Gammaproteobacteria; Vibrionales; Vibrionaceae; Vibrio | methyl-accepting chemotaxis protein | 6 |
|  | WP_024575067.1 | 6.7E-06 | Qc.III | -KAIGQTIEQINDIATSIAAAVEQQGAATEEIAKSVQAAASVTT--------- | *Afipia* | Bacteria; Proteobacteria; Alphaproteobacteria; Rhizobiales; Bradyrhizobiaceae | MULTISPECIES: methyl-accepting chemotaxis protein | 6 |
| | WP_035678192.1 | 6.7E-06 | Qc.III | IKDISHTIERLSEISSTIAAAVEEQGAATQEISRNVQQAAAGTHQVSSNI--- | *Bradyrhizobium* | Bacteria; Proteobacteria; Alphaproteobacteria; Rhizobiales; Bradyrhizobiaceae | MULTISPECIES: HAMP domain-containing protein | 6 |
|  | WP_038939999.1 | 6.7E-06 | Qc.III | LENIMASVQRINDMSTQIATAAEEQSLVTEEITRNTQAVNDVSN--------- | *Vibrio vulnificus* | Bacteria; Proteobacteria; Gammaproteobacteria; Vibrionales; Vibrionaceae; Vibrio | methyl-accepting chemotaxis protein | 6 |
|  | WP_038963819.1 | 6.7E-06 | Qc.III | LENIMASVQRINDMSTQIATAAEEQSLVTEEITRNTQAVNDVSN--------- | *Vibrio vulnificus* | Bacteria; Proteobacteria; Gammaproteobacteria; Vibrionales; Vibrionaceae; Vibrio | methyl-accepting chemotaxis protein | 6 |
|  | WP_045590345.1 | 6.7E-06 | Qc.III | LENIMASVQRINDMSTQIATAAEEQSLVTEEITRNTQAVNDVSN--------- | *Vibrio vulnificus* | Bacteria; Proteobacteria; Gammaproteobacteria; Vibrionales; Vibrionaceae; Vibrio | methyl-accepting chemotaxis protein | 6 |
|  | WP_045610497.1 | 6.7E-06 | Qc.III | LENIMASVQRINDMSTQIATAAEEQSLVTEEITRNTQAVNDVSN--------- | *Vibrio vulnificus* | Bacteria; Proteobacteria; Gammaproteobacteria; Vibrionales; Vibrionaceae; Vibrio | methyl-accepting chemotaxis protein | 6 |
|  | WP_046028671.1 | 6.7E-06 | Qc.III | LENIMASVQRINDMSTQIATAAEEQSLVTEEITRNTQAVNDVSN--------- | *Vibrio vulnificus* | Bacteria; Proteobacteria; Gammaproteobacteria; Vibrionales; Vibrionaceae; Vibrio | methyl-accepting chemotaxis protein | 6 |
|  | WP_072608970.1 | 6.7E-06 | Qc.III | LENIMASVQRINDMSTQIATAAEEQSLVTEEITRNTQAVNDVSN--------- | *Vibrio vulnificus* | Bacteria; Proteobacteria; Gammaproteobacteria; Vibrionales; Vibrionaceae; Vibrio | methyl-accepting chemotaxis protein | 6 |
|  | WP_080541305.1 | 6.7E-06 | Qc.III | LENIMASVQRINDMSTQIATAAEEQSLVTEEITRNTQAVNDVSN--------- | *Vibrio vulnificus* | Bacteria; Proteobacteria; Gammaproteobacteria; Vibrionales; Vibrionaceae; Vibrio | methyl-accepting chemotaxis protein | 6 |
|  | WP_086467944.1 | 6.7E-06 | Qc.III | LENIMASVQRINDMSTQIATAAEEQSLVTEEITRNTQAVNDVSN--------- | *Vibrio vulnificus* | Bacteria; Proteobacteria; Gammaproteobacteria; Vibrionales; Vibrionaceae; Vibrio | methyl-accepting chemotaxis protein | 6 |
|  | WP_103154648.1 | 6.7E-06 | Qc.III | LENIMASVQRINDMSTQIATAAEEQSLVTEEITRNTQAVNDVSN--------- | *Vibrio vulnificus* | Bacteria; Proteobacteria; Gammaproteobacteria; Vibrionales; Vibrionaceae; Vibrio | methyl-accepting chemotaxis protein | 6 |
|  | WP_103156398.1 | 6.7E-06 | Qc.III | LENIMASVQRINDMSTQIATAAEEQSLVTEEITRNTQAVNDVSN--------- | *Vibrio vulnificus* | Bacteria; Proteobacteria; Gammaproteobacteria; Vibrionales; Vibrionaceae; Vibrio | methyl-accepting chemotaxis protein | 6 |
|  | WP_103163937.1 | 6.7E-06 | Qc.III | LENIMASVQRINDMSTQIATAAEEQSLVTEEITRNTQAVNDVSN--------- | *Vibrio vulnificus* | Bacteria; Proteobacteria; Gammaproteobacteria; Vibrionales; Vibrionaceae; Vibrio | methyl-accepting chemotaxis protein | 6 |
|  | EWY40261.1 | 6.8E-06 | Qa.IV | IQAIARVIIEVNEISTTVAAAVEEQAAATQEIARNIQQAA------------- | *Skermanella stibiiresistens SB22* | Bacteria; Proteobacteria; Alphaproteobacteria; Rhodospirillales; Rhodospirillaceae; Skermanella | methyl-accepting chemotaxis protein | 6 |
| | KQQ06698.1 | 6.8E-06 | Qc.III.c | --GISISIRSINETSTAIASAVEEQEAATREIVQAIAQASSGIGEVTANIT-- | *Methylobacterium sp. Leaf121* | Bacteria; Proteobacteria; Alphaproteobacteria; Rhizobiales; Methylobacteriaceae; Methylobacterium | hypothetical protein ASF59_23720 | 6 |
|  | WP_007107027.1 | 6.8E-06 | Qc.I | -----------KNMNMAMSEQLRQEKRQLDEVNQELNAAYDKLNAINNTLEE- | *Paraglaciecola polaris* | Bacteria; Proteobacteria; Gammaproteobacteria; Alteromonadales; Alteromonadaceae; Paraglaciecola | EAL domain-containing protein | 6 |
| | WP_009539374.1 | 6.8E-06 | Qc.III | IDSINRVIERMAEISAAVSAAVEEQNAATAEIARSVDQASEGTREVSRNI--- | *Caenispirillum salinarum* | Bacteria; Proteobacteria; Alphaproteobacteria; Rhodospirillales; Rhodospirillaceae; Caenispirillum | HAMP domain-containing protein | 6 |
|  | WP_051512268.1 | 6.8E-06 | Qa.IV | IQAIARVIIEVNEISTTVAAAVEEQAAATQEIARNIQQAA------------- | *Skermanella stibiiresistens* | Bacteria; Proteobacteria; Alphaproteobacteria; Rhodospirillales; Rhodospirillaceae; Skermanella | methyl-accepting chemotaxis protein | 6 |
|  | WP_052657661.1 | 6.8E-06 | Qa.III | IEKVTETISNINEIATAIASAVEEQGAATAEISSNVQQAA------------- | *Thalassospira sp. HJ* | Bacteria; Proteobacteria; Alphaproteobacteria; Rhodospirillales; Rhodospirillaceae; Thalassospira | methyl-accepting chemotaxis protein | 6 |
|  | WP_064607315.1 | 6.8E-06 | Qa.III | IESIEQSVGQIADMSQQIASACSEQDSVTDELGRNVETINQSSK--------- | *Photobacterium sp. J15* | Bacteria; Proteobacteria; Gammaproteobacteria; Vibrionales; Vibrionaceae; Photobacterium | methyl-accepting chemotaxis protein | 6 |
|  | WP_081370264.1 | 6.8E-06 | Qc.III | LENIMASVQRINDMSTQIATAAEEQSLVTEEITRNTQAVNDVSN--------- | *Vibrio vulnificus* | Bacteria; Proteobacteria; Gammaproteobacteria; Vibrionales; Vibrionaceae; Vibrio | methyl-accepting chemotaxis protein | 6 |
|  | WP_085617000.1 | 6.8E-06 | Qa.IV | ---------EVSQLSSHIASSVQQQTAAISEISNSAQEVSTHMQGISGDIAQA | *Thalassospira alkalitolerans* | Bacteria; Proteobacteria; Alphaproteobacteria; Rhodospirillales; Rhodospirillaceae; Thalassospira | chemotaxis protein | 6 |
|  | WP_096173283.1 | 6.8E-06 | Qa.III | IQEIGTTIGDMNAIASSIASAVEQQSSATGEIAQNVEQAA------------- | *Cohaesibacter sp. ES.047* | Bacteria; Proteobacteria; Alphaproteobacteria; Rhizobiales; Cohaesibacteraceae | methyl-accepting chemotaxis protein | 6 |
|  | WP_044562036.1 | 6.9E-06 | Qa.IV | IITISKTIAEINDISDMVAGAVEEQDAATKEIARNVQQAS------------- | *Azospirillum sp. B4* | Bacteria; Proteobacteria; Alphaproteobacteria; Rhodospirillales; Rhodospirillaceae; Azospirillum | methyl-accepting chemotaxis protein | 6 |
|  | WP_053240013.1 | 6.9E-06 | Qa.III | IRAIAGTMEEVNRATLGIASAVEQQEASTNEISRNVSEASDNTV--------- | *Pleomorphomonas koreensis* | Bacteria; Proteobacteria; Alphaproteobacteria; Rhizobiales; Methylocystaceae; Pleomorphomonas | methyl-accepting chemotaxis protein | 6 |
|  | WP_055456134.1 | 6.9E-06 | Qa.IV | IQEIAETMSEVNQFTTTIADAISQQGEATREITVNVQRAAD------------ | *Pannonibacter indicus* | Bacteria; Proteobacteria; Alphaproteobacteria; Rhodobacterales; Rhodobacteraceae; Pannonibacter | methyl-accepting chemotaxis protein | 6 |
|  | KPL54199.1 | 7E-06 | Qa.IV | -EEIQSTIRTIDALTMQVAAAVEEQGASTNEIARNVSEAARAT---------- | *Prosthecomicrobium hirschii* | Bacteria; Proteobacteria; Alphaproteobacteria; Rhizobiales; Hyphomicrobiaceae; Prosthecomicrobium | hypothetical protein ABB55_19900 | 6 |
|  | KZK83910.1 | 7E-06 | Qc.III.c | ---IGETIEQINSLSTGINEAVGDQKQATEDISRDLNSAHQNVSQV------- | *Pseudovibrio sp. W64* | Bacteria; Proteobacteria; Alphaproteobacteria; Rhodobacterales; Rhodobacteraceae; Pseudovibrio | Methyl-accepting chemotaxis protein 4 | 6 |
|  | OQW58620.1 | 7E-06 | Qa.III.b | IQGICATILQVNEISSSIAESVGQQGSATQEIASSVQKAAAGTSQVSQ----- | *Proteobacteria bacterium SG_bin9* | Bacteria; Proteobacteria | chemotaxis protein | 6 |
|  | WP_057013266.1 | 7E-06 | Qb.I | LQRTKLHLQDTIEQSEVSSEELKASNEEMQAVNEELRSATEELETS------- | *Pseudomonas libanensis* | Bacteria; Proteobacteria; Gammaproteobacteria; Pseudomonadales; Pseudomonadaceae; Pseudomonas | PAS domain S-box protein | 6 |
|  | WP_063290281.1 | 7E-06 | Qc.III.c | ---IGETIEQINSLSTGINEAVGDQKQATEDISRDLNSAHQNVSQV------- | *Pseudovibrio* | Bacteria; Proteobacteria; Alphaproteobacteria; Rhodobacterales; Rhodobacteraceae | MULTISPECIES: methyl-accepting chemotaxis protein | 6 |
|  | CRM87106.1 | 7.1E-06 | Qb.I | LQRTKLHLQDTIEQAEVSSEELKASNEEIQAINEELRSATEELETS------- | *Pseudomonas sp. 22 E 5* | Bacteria; Proteobacteria; Gammaproteobacteria; Pseudomonadales; Pseudomonadaceae; Pseudomonas | Aerobic respiration control sensor protein ArcB | 6 |

| | | | | | | | |
|---|---|---|---|---|---|---|---|
| WP_017223047.1 | 7.1E-06 | Qa.III | LEEINRSVVLISDMATQIATAAEEQTQVTSEISQNVISIK------------- | *Moritella dasanensis* | Bacteria; Proteobacteria; Gammaproteobacteria; Alteromonadales; Moritellaceae; Moritella | methyl-accepting chemotaxis protein | 6 |
| WP_070881405.1 | 7.1E-06 | Qb.I | LQNTKEQLQATVEQAETSTEELKASNEELQAINEELRSTTEELETSK------ | *Pseudomonas punonensis* | Bacteria; Proteobacteria; Gammaproteobacteria; Pseudomonadales; Pseudomonadaceae; Pseudomonas | PAS domain S-box protein | 6 |
| WP_109107890.1 | 7.1E-06 | Qa.III | IVGIGRTIHAINEIAAGISAGIEQQGSAVREIVANMEQAAA------------ | *Azospirillum sp. TSO35-2* | Bacteria; Proteobacteria; Alphaproteobacteria; Rhodospirillales; Rhodospirillaceae; Azospirillum | chemotaxis protein | 6 |
| AWJ93595.1 | 7.2E-06 | Qc.III | -GSIGQTIGRMSEIATTIAAAIEEQAAATREIASSVSQAAQGTEEVSRNIA-- | *Azospirillum brasilense Sp245* | Bacteria; Proteobacteria; Alphaproteobacteria; Rhodospirillales; Rhodospirillaceae; Azospirillum | chemotaxis protein (plasmid) | 6 |
| ESQ89297.1 | 7.2E-06 | Qc.III.b | --DIIHAVQQLHDIAQTVSGRVDEQARTTSEMSHVVRRASSEAQT-------- | *Asticcacaulis sp. AC460* | Bacteria; Proteobacteria; Alphaproteobacteria; Caulobacterales; Caulobacteraceae; Asticcacaulis | hypothetical protein ABAC460_13465 | 6 |
| WP_008438668.1 | 7.2E-06 | Qb.I | LQRTKLHLQDTIEQSEVSSEELKASNEEMQAVNEELRSATEELETS------- | *Pseudomonas sp. Ag1* | Bacteria; Proteobacteria; Gammaproteobacteria; Pseudomonadales; Pseudomonadaceae; Pseudomonas | PAS domain S-box protein | 6 |
| WP_014746010.1 | 7.2E-06 | Qa.III | IERIVGMIGEISSMTTSIATAVEEQSAATGEITRNVQEAANGTEQIRQDVQ-- | *Tistrella mobilis* | Bacteria; Proteobacteria; Alphaproteobacteria; Rhodospirillales; Rhodospirillaceae; Tistrella | HAMP domain-containing protein | 6 |
| WP_027553769.1 | 7.2E-06 | Qc.III | IQEIGGTIERLSEVASAIAAAVEEQGAATQEISRNVQQASVGTQEVSSNIT-- | *Bradyrhizobium sp. Cp5.3* | Bacteria; Proteobacteria; Alphaproteobacteria; Rhizobiales; Bradyrhizobiaceae; Bradyrhizobium | HAMP domain-containing protein | 6 |
| WP_027616407.1 | 7.2E-06 | Qb.I | LQRTKLHLQDTIEQSEVSSEELKASNEEMQAVNEELRSATEELETS------- | *Pseudomonas sp. URHB0015* | Bacteria; Proteobacteria; Gammaproteobacteria; Pseudomonadales; Pseudomonadaceae; Pseudomonas | PAS domain S-box protein | 6 |
| WP_034103397.1 | 7.2E-06 | Qb.I | LQRTKLHLQDTIEQSEVSSEELKASNEEMQAVNEELRSATEELETS------- | *Pseudomonas fluorescens* | Bacteria; Proteobacteria; Gammaproteobacteria; Pseudomonadales; Pseudomonadaceae; Pseudomonas | PAS domain S-box protein | 6 |
| WP_040136010.1 | 7.2E-06 | Qc.III | -GSIGQTIGRMSEIATTIAAAIEEQAAATREIASSVSQAAQGTEEVSRNIA-- | *Azospirillum brasilense* | Bacteria; Proteobacteria; Alphaproteobacteria; Rhodospirillales; Rhodospirillaceae; Azospirillum | methyl-accepting chemotaxis protein | 6 |
| WP_043046424.1 | 7.2E-06 | Qb.I | LQRTKLHLQDTIEQSEVSSEELKASNEEMQAVNEELRSATEELETS------- | *Pseudomonas fluorescens* | Bacteria; Proteobacteria; Gammaproteobacteria; Pseudomonadales; Pseudomonadaceae; Pseudomonas | PAS domain S-box protein | 6 |
| WP_054068922.1 | 7.2E-06 | Qb.I | LQRTKLHLQDTIEQSEVSSEELKASNEEMQAVNEELRSATEELETS------- | *Pseudomonas syringae group genomosp. 3* | Bacteria; Proteobacteria; Gammaproteobacteria; Pseudomonadales; Pseudomonadaceae; Pseudomonas | PAS domain S-box protein | 6 |
| WP_059399560.1 | 7.2E-06 | Qc.III | -GSIGQTIGRMSEIATTIAAAIEEQAAATREIASSVSQAAQGTEEVSRNIA-- | *Azospirillum* | Bacteria; Proteobacteria; Alphaproteobacteria; Rhodospirillales; Rhodospirillaceae | MULTISPECIES: methyl-accepting chemotaxis protein | 6 |
| WP_060693664.1 | 7.2E-06 | Qb.I | LQRTKLHLQDTIEQSEVSSEELKASNEEMQAVNEELRSATEELETS------- | *Pseudomonas versuta* | Bacteria; Proteobacteria; Gammaproteobacteria; Pseudomonadales; Pseudomonadaceae; Pseudomonas | PAS domain S-box protein | 6 |
| WP_063678697.1 | 7.2E-06 | Qc.III | IQEIGGTIERLSEVASAIAAAVEEQGAATQEISRNVQQASVGTQEVSSNIT-- | *Bradyrhizobium neotropicale* | Bacteria; Proteobacteria; Alphaproteobacteria; Rhizobiales; Bradyrhizobiaceae; Bradyrhizobium | HAMP domain-containing protein | 6 |
| WP_065946801.1 | 7.2E-06 | Qb.I | LQRTKLHLQDTIEQSEVSSEELKASNEEMQAVNEELRSATEELETS------- | *Pseudomonas sp. 1 R 17* | Bacteria; Proteobacteria; Gammaproteobacteria; Pseudomonadales; Pseudomonadaceae; Pseudomonas | PAS domain S-box protein | 6 |
| WP_073510264.1 | 7.2E-06 | Qb.I | LQRTKLHLQDTIEQSEVSSEELKASNEEMQAVNEELRSATEELETS------- | *Pseudomonas versuta* | Bacteria; Proteobacteria; Gammaproteobacteria; Pseudomonadales; Pseudomonadaceae; Pseudomonas | PAS domain S-box protein | 6 |
| WP_073613474.1 | 7.2E-06 | Qa.III.b | IGNIAKVVGQVNELVATIAAAVEEQSVTTRDIAGNVAKATEGV---------- | *Desulfopila aestuarii* | Bacteria; Proteobacteria; Deltaproteobacteria; Desulfobacterales; Desulfobulbaceae; Desulfopila | methyl-accepting chemotaxis protein | 6 |
| WP_074846686.1 | 7.2E-06 | Qb.I | LQRTKLHLQDTIEQSEVSSEELKASNEEMQAVNEELRSATEELETS------- | *Pseudomonas marginalis* | Bacteria; Proteobacteria; Gammaproteobacteria; Pseudomonadales; Pseudomonadaceae; Pseudomonas | PAS domain S-box protein | 6 |
| WP_078738015.1 | 7.2E-06 | Qb.I | LQRTKLHLQDTIEQSEVSSEELKASNEEMQAVNEELRSATEELETS------- | *Pseudomonas fluorescens* | Bacteria; Proteobacteria; Gammaproteobacteria; Pseudomonadales; Pseudomonadaceae; Pseudomonas | PAS domain S-box protein | 6 |
| WP_078828212.1 | 7.2E-06 | Qb.I | LQRTKLHLQDTIEQSEVSSEELKASNEEMQAVNEELRSATEELETS------- | *Pseudomonas fluorescens* | Bacteria; Proteobacteria; Gammaproteobacteria; Pseudomonadales; Pseudomonadaceae; Pseudomonas | PAS domain S-box protein | 6 |
| WP_081716565.1 | 7.2E-06 | Qc.III.b | --DIIHAVQQLHDIAQTVSGRVDEQARTTSEMSHVVRRASSEAQT-------- | *Asticcacaulis sp. AC460* | Bacteria; Proteobacteria; Alphaproteobacteria; Caulobacterales; Caulobacteraceae; Asticcacaulis | methyl-accepting chemotaxis protein | 6 |
| WP_094303584.1 | 7.2E-06 | Qc.III | -GSIGQTIGRMSEIATTIAAAIEEQAAATREIASSVSQAAQGTEEVSRNIA-- | *Azospirillum brasilense* | Bacteria; Proteobacteria; Alphaproteobacteria; Rhodospirillales; Rhodospirillaceae; Azospirillum | methyl-accepting chemotaxis protein | 6 |
| WP_099583087.1 | 7.2E-06 | Qb.I | LQRTKLHLQDTIEQSEVSSEELKASNEEMQAVNEELRSATEELETS------- | *Pseudomonas sp. 2822-17* | Bacteria; Proteobacteria; Gammaproteobacteria; Pseudomonadales; Pseudomonadaceae; Pseudomonas | PAS domain S-box protein | 6 |
| WP_103040454.1 | 7.2E-06 | Qc.III | -GSIGQTIGRMSEIATTIAAAIEEQAAATREIASSVSQAAQGTEEVSRNIA-- | *Azospirillum brasilense* | Bacteria; Proteobacteria; Alphaproteobacteria; Rhodospirillales; Rhodospirillaceae; Azospirillum | methyl-accepting chemotaxis protein | 6 |
| WP_109068799.1 | 7.2E-06 | Qc.III | -GSIGQTIGRMSEIATTIAAAIEEQAAATREIASSVSQAAQGTEEVSRNIA-- | *Azospirillum sp. TSH58* | Bacteria; Proteobacteria; Alphaproteobacteria; Rhodospirillales; Rhodospirillaceae; Azospirillum | methyl-accepting chemotaxis protein | 6 |
| WP_111434241.1 | 7.2E-06 | Qa.III | --GITATMGEVNEYTNAIAAAVVEQGSATNEISNNIGNAASRTQSAV------ | *Rhodobium orientis* | Bacteria; Proteobacteria; Alphaproteobacteria; Rhizobiales; Rhodobiaceae; Rhodobium | methyl-accepting chemotaxis protein | 6 |
| WP_112195373.1 | 7.2E-06 | Qb.I | LQRTKLHLQDTIEQSEVSSEELKASNEEMQAVNEELRSATEELETS------- | *Pseudomonas sp. LG1E9* | Bacteria; Proteobacteria; Gammaproteobacteria; Pseudomonadales; Pseudomonadaceae; Pseudomonas | PAS domain S-box protein | 6 |
| AWM04541.1 | 7.3E-06 | Qc.III | IQEIGGTIERLSEVSAAIAAAVEEQGAATQEISRNVQQASVGTQEVSSNIT-- | *Bradyrhizobium sp. 2 39S1MB* | Bacteria; Proteobacteria; Alphaproteobacteria; Rhizobiales; Bradyrhizobiaceae; Bradyrhizobium | HAMP domain-containing protein | 6 |
| AWM06052.1 | 7.3E-06 | Qc.III | IQEIGGTIERLSEVSAAIAAAVEEQGAATQEISRNVQQASVGTQEVSSNIT-- | *Bradyrhizobium sp. 3 85S1MB* | Bacteria; Proteobacteria; Alphaproteobacteria; Rhizobiales; Bradyrhizobiaceae; Bradyrhizobium | methyl-accepting chemotaxis protein | 6 |
| SCM73103.1 | 7.3E-06 | Qa.IV | IRAIAATMEEVNRATLGIASAVEQQEASTNEISRNVSDASGNTV--------- | *uncultured Pleomorphomonas sp.* | Bacteria; Proteobacteria; Alphaproteobacteria; Rhizobiales; Methylocystaceae; Pleomorphomonas; environmental samples | putative Methyl-accepting chemotaxis protein | 6 |
| WP_025744322.1 | 7.3E-06 | Qa.III | -NAINEQIIQISDMNTQVATATEEQSTVVNDINQNIEEINHSSQQTA------ | *Salinivibrio* | Bacteria; Proteobacteria; Gammaproteobacteria; Vibrionales; Vibrionaceae | MULTISPECIES: methyl-accepting chemotaxis protein | 6 |
| WP_058821840.1 | 7.3E-06 | Qb.I | LQRTKLHLQDTIEQSEVSSEELKASNEEMQAVNEELRSATEELETS------- | *Pseudomonas syringae* | Bacteria; Proteobacteria; Gammaproteobacteria; Pseudomonadales; Pseudomonadaceae; Pseudomonas | PAS domain S-box protein | 6 |
| WP_077459846.1 | 7.3E-06 | Qa.III | -NAINEQIIQISDMNTQVATATEEQSTVVNDINQNIEEINHSSQQTA------ | *Salinivibrio sp. AR647* | Bacteria; Proteobacteria; Gammaproteobacteria; Vibrionales; Vibrionaceae; Salinivibrio | methyl-accepting chemotaxis protein | 6 |
| WP_077576191.1 | 7.3E-06 | Qa.III | -NAINEQIIQISDMNTQVATATEEQSTVVNDINQNIEEINHSSQQTA------ | *Salinivibrio sp. AR640* | Bacteria; Proteobacteria; Gammaproteobacteria; Vibrionales; Vibrionaceae; Salinivibrio | methyl-accepting chemotaxis protein | 6 |
| WP_077645797.1 | 7.3E-06 | Qa.III | -NAINEQIIQISDMNTQVATATEEQSTVVNDINQNIEEINHSSQQTA------ | *Salinivibrio sp. IB643* | Bacteria; Proteobacteria; Gammaproteobacteria; Vibrionales; Vibrionaceae; Salinivibrio | methyl-accepting chemotaxis protein | 6 |
| WP_077660108.1 | 7.3E-06 | Qa.III | -NAINEQIIQISDMNTQVATATEEQSTVVNDINQNIEEINHSSQQTA------ | *Salinivibrio sp. MA351* | Bacteria; Proteobacteria; Gammaproteobacteria; Vibrionales; Vibrionaceae; Salinivibrio | methyl-accepting chemotaxis protein | 6 |
| WP_077669152.1 | 7.3E-06 | Qa.III | -NAINEQIIQISDMNTQVATATEEQSTVVNDINQNIEEINHSSQQTA------ | *Salinivibrio costicola* | Bacteria; Proteobacteria; Gammaproteobacteria; Vibrionales; Vibrionaceae; Salinivibrio | methyl-accepting chemotaxis protein | 6 |
| WP_077676612.1 | 7.3E-06 | Qa.III | -NAINEQIIQISDMNTQVATATEEQSTVVNDINQNIEEINHSSQQTA------ | *Salinivibrio sp. MA607* | Bacteria; Proteobacteria; Gammaproteobacteria; Vibrionales; Vibrionaceae; Salinivibrio | methyl-accepting chemotaxis protein | 6 |
| WP_077678566.1 | 7.3E-06 | Qa.III | -NAINEQIIQISDMNTQVATATEEQSTVVNDINQNIEEINHSSQQTA------ | *Salinivibrio sp. MA440* | Bacteria; Proteobacteria; Gammaproteobacteria; Vibrionales; Vibrionaceae; Salinivibrio | methyl-accepting chemotaxis protein | 6 |
| WP_085126565.1 | 7.3E-06 | Qa.IV | IQAIVASIGEVGSVTGTIATAVQQQASATTEISRNVQAASSATGE-------- | *Tistlia consotensis* | Bacteria; Proteobacteria; Alphaproteobacteria; Rhodospirillales; Rhodospirillaceae; Tistlia | hypothetical protein | 6 |
| WP_100078860.1 | 7.3E-06 | Qa.IV | IRAIAATMEEVNRATLGIASAVEQQEASTNEISRNVSDASGNTV--------- | *Pleomorphomonas sp. SVCO-16* | Bacteria; Proteobacteria; Alphaproteobacteria; Rhizobiales; Methylocystaceae; Pleomorphomonas | methyl-accepting chemotaxis protein | 6 |
| WP_109070144.1 | 7.3E-06 | Qa.III | IRAIAGTIAEINEIASAIAAAVEQQGAATREIARNVQEAA------------- | *Azospirillum sp. TSH58* | Bacteria; Proteobacteria; Alphaproteobacteria; Rhodospirillales; Rhodospirillaceae; Azospirillum | urea ABC transporter substrate-binding protein | 6 |
| CBH21084.1 | 7.4E-06 | Qb.I | --NIKSLVDDTVENTQKSSENIKQSSNLVETASETFDKIYNQVNQSNIL---- | *[Clostridium] sticklandii* | Bacteria; Firmicutes; Clostridia; Clostridiales; Peptostreptococcaceae; Peptoclostridium | Histidine kinase HAMP region:chemotaxis sensory transducer precursor | 6 |
| OSQ36182.1 | 7.4E-06 | Qc.III.b | ---INGNIGEINANARHVSDTVNQQYEMIAEIGRRVDGAVEQMRK-------- | *Thalassospira mesophila* | Bacteria; Proteobacteria; Alphaproteobacteria; Rhodospirillales; Rhodospirillaceae; Thalassospira | hypothetical protein TMES_18830 | 6 |
| SFP41881.1 | 7.4E-06 | Qc.III | IQEIGGTIERLSEVSAAIAAAVEEQGAATQEISRNVQQASVGTQEVSSNIT-- | *Bradyrhizobium sp. Ghvi* | Bacteria; Proteobacteria; Alphaproteobacteria; Rhizobiales; Bradyrhizobiaceae; Bradyrhizobium | methyl-accepting chemotaxis protein | 6 |
| WP_011189139.1 | 7.4E-06 | Qa.III | IDEIRQVIADVDQIVSTIATAVEEQSVTTREIAINVSQAS------------- | *Desulfotalea psychrophila* | Bacteria; Proteobacteria; Deltaproteobacteria; Desulfobacterales; Desulfobulbaceae; Desulfotalea | methyl-accepting chemotaxis protein | 6 |
| WP_027558192.1 | 7.4E-06 | Qc.III | IQEIGGTIERLSEVSAAIAAAVEEQGAATQEISRNVQQASVGTQEVSSNIT-- | *Bradyrhizobium genosp. SA-4* | Bacteria; Proteobacteria; Alphaproteobacteria; Rhizobiales; Bradyrhizobiaceae; Bradyrhizobium | methyl-accepting chemotaxis protein | 6 |
| WP_045005143.1 | 7.4E-06 | Qc.III | IQEIGGTIERLSEVSAAIAAAVEEQGAATQEISRNVQQASVGTQEVSSNIT-- | *Bradyrhizobium sp. LTSP857* | Bacteria; Proteobacteria; Alphaproteobacteria; Rhizobiales; Bradyrhizobiaceae; Bradyrhizobium | methyl-accepting chemotaxis protein | 6 |
| WP_045014261.1 | 7.4E-06 | Qc.III | IQEIGGTIERLSEVSAAIAAAVEEQGAATQEISRNVQQASVGTQEVSSNIT-- | *Bradyrhizobium sp. LTSP849* | Bacteria; Proteobacteria; Alphaproteobacteria; Rhizobiales; Bradyrhizobiaceae; Bradyrhizobium | methyl-accepting chemotaxis protein | 6 |
| WP_084608394.1 | 7.4E-06 | Qc.III | ---ITQTIEQVDQIAAAISAAVEEQTAATQEIARNVEEASAGTNQVNSNISE- | *Nisaea denitrificans* | Bacteria; Proteobacteria; Alphaproteobacteria; Rhodospirillales; Rhodospirillaceae; Nisaea | methyl-accepting chemotaxis protein | 6 |
| WP_108914366.1 | 7.4E-06 | Qc.III | IQEIGGTIERLSEVSAAIAAAVEEQGAATQEISRNVQQASVGTQEVSSNIT-- | *Bradyrhizobium arachidis* | Bacteria; Proteobacteria; Alphaproteobacteria; Rhizobiales; Bradyrhizobiaceae; Bradyrhizobium | methyl-accepting chemotaxis protein | 6 |
| WP_111418810.1 | 7.4E-06 | Qc.III | IKEISGTIGKIAEISSAIAAAVEEQGAATSEISRNIQHAATGTT--------- | *Rhodoplanes roseus* | Bacteria; Proteobacteria; Alphaproteobacteria; Rhizobiales; Hyphomicrobiaceae; Rhodoplanes | methyl-accepting chemotaxis protein | 6 |
| WP_111433453.1 | 7.4E-06 | Qa.IV | IQQIGATIKDMNEIALGISSAVEEQNAATNEIARNVEQAA------------- | *Rhodobium orientis* | Bacteria; Proteobacteria; Alphaproteobacteria; Rhizobiales; Rhodobiaceae; Rhodobium | methyl-accepting chemotaxis protein | 6 |
| BAF87315.1 | 7.5E-06 | Qc.III | IENVSNQIRQMSDVATGISAAVEQQGVATKEIVRNVDQAATGTNTVTSHIAD- | *Azorhizobium caulinodans ORS 571* | Bacteria; Proteobacteria; Alphaproteobacteria; Rhizobiales; Xanthobacteraceae; Azorhizobium | methyl-accepting chemotaxis protein | 6 |
| OYU50507.1 | 7.5E-06 | Qa.III | --QIAGILSEMSTIATSVASAVEEQSSAIATISDNVNEAARSSAEGASAI--- | *Rhizobiales bacterium PAR1* | Bacteria; Proteobacteria; Alphaproteobacteria; Rhizobiales | hypothetical protein CFE31_02865 | 6 |
| SDE97338.1 | 7.5E-06 | Qa.III.b | IKGIVAVIGEINEITVTVSSAVEEQTAATGEISENAQHAAESTQNASDN---- | *Rhodospira trueperi* | Bacteria; Proteobacteria; Alphaproteobacteria; Rhodospirillales; Rhodospirillaceae; Rhodospira | methyl-accepting chemotaxis protein | 6 |
| SOD96516.1 | 7.5E-06 | Qa.IV | IQTIVRTIEEVNEVASAIAAAVEQQNAATHEISRNVQEAA------------- | *Caenispirillum bisanense* | Bacteria; Proteobacteria; Alphaproteobacteria; Rhodospirillales; Rhodospirillaceae; Caenispirillum | methyl-accepting chemotaxis sensory transducer with Cache sensor | 6 |

| | | | | | | | |
|---|---|---|---|---|---|---|---|
| **WP_014853076.1** | 7.5E-06 | Qc.III | LDQMADSVNRIHDLSAQIAAAAEEQSAVSGEINQNMVAVRD------------ | *Pseudomonas stutzeri* | Bacteria; Proteobacteria; Gammaproteobacteria; Pseudomonadales; Pseudomonadaceae; Pseudomonas | methyl-accepting chemotaxis protein | 6 |
| **WP_022664241.1** | 7.5E-06 | Qa.III | IQAISKDITSVNEMIDTVAAAVEEQSVTTREISANVTQAA------------- | *Desulfospira joergensenii* | Bacteria; Proteobacteria; Deltaproteobacteria; Desulfobacterales; Desulfobacteraceae; Desulfospira | methyl-accepting chemotaxis protein | 6 |
| **WP_027109916.1** | 7.5E-06 | Qb.I | ---IRKLIGDAVSQASESADNIKENSELIHVAVETFDQIYDIIQDTNSLIT-- | *Lachnospiraceae bacterium NC2008* | Bacteria; Firmicutes; Clostridia; Clostridiales; Lachnospiraceae | methyl-accepting chemotaxis protein | 6 |
| **WP_036177302.1** | 7.5E-06 | Qb.I | LQRTKEQLQETIEHSEISTEELRASNEELQAINEELRSATEELETS------- | *Massilia sp. 9096* | Bacteria; Proteobacteria; Betaproteobacteria; Burkholderiales; Oxalobacteraceae; Massilia | PAS domain S-box protein | 6 |
| **WP_052233788.1** | 7.5E-06 | Qb.I | LQRTKEQLQETIEHSEISTEELRASNEELQAINEELRSATEELETS------- | *Massilia* | Bacteria; Proteobacteria; Betaproteobacteria; Burkholderiales; Oxalobacteraceae | MULTISPECIES: PAS domain S-box protein | 6 |
| **WP_092217948.1** | 7.5E-06 | Qc.III | IQEIGGTIERLSEVSAAIAAAVEEQGAATQEISRNVQQASVGTQEVSSNIT-- | *Bradyrhizobium arachidis* | Bacteria; Proteobacteria; Alphaproteobacteria; Rhizobiales; Bradyrhizobiaceae; Bradyrhizobium | methyl-accepting chemotaxis protein | 6 |
| **WP_092298459.1** | 7.5E-06 | Qc.III | IQEIGGTIERLSEVSAAIAAAVEEQGAATQEISRNVQQASVGTQEVSSNIT-- | *Bradyrhizobium sp. Ghvi* | Bacteria; Proteobacteria; Alphaproteobacteria; Rhizobiales; Bradyrhizobiaceae; Bradyrhizobium | methyl-accepting chemotaxis protein | 6 |
| **WP_109634005.1** | 7.5E-06 | Qb.I | LQRTKLHLQDTIEQSEVSSEELKASNEEMQAINEELRSATEELETS------- | *Pseudomonas sp. 31-12* | Bacteria; Proteobacteria; Gammaproteobacteria; Pseudomonadales; Pseudomonadaceae; Pseudomonas | PAS domain S-box protein | 6 |
| **WP_020407345.1** | 7.6E-06 | Qa.III | ---------TANEMNAQIASAAREQHQVVEEINQNIASINELSHQSSDSAT-- | *Hahella ganghwensis* | Bacteria; Proteobacteria; Gammaproteobacteria; Oceanospirillales; Hahellaceae; Hahella | methyl-accepting chemotaxis protein | 6 |
| **WP_027516462.1** | 7.6E-06 | Qc.III | IKDISGTIMKLSEISSTIAAAVEEQGAATQEISRNVQQAAEGTHQVSSNITD- | *Bradyrhizobium sp. WSM1417* | Bacteria; Proteobacteria; Alphaproteobacteria; Rhizobiales; Bradyrhizobiaceae; Bradyrhizobium | methyl-accepting chemotaxis protein | 6 |
| **WP_083463780.1** | 7.6E-06 | Qa.IV | -EEIQSTIRTIDALTMQVAAAVEEQGASTNEIARNVSEAARAT---------- | *Prosthecomicrobium hirschii* | Bacteria; Proteobacteria; Alphaproteobacteria; Rhizobiales; Hyphomicrobiaceae; Prosthecomicrobium | methyl-accepting chemotaxis protein | 6 |
| **WP_092787931.1** | 7.6E-06 | Qa.III.b | IKGIVAVIGEINEITVTVSSAVEEQTAATGEISENAQHAAESTQNASDN---- | *Rhodospira trueperi* | Bacteria; Proteobacteria; Alphaproteobacteria; Rhodospirillales; Rhodospirillaceae; Rhodospira | methyl-accepting chemotaxis protein | 6 |
| **ETR68800.1** | 7.7E-06 | Qc.III.c | --EISQIIDRMNNAAKTILSAVEEQTRAADNLSHNANEADSR----------- | *Candidatus Magnetoglobus multicellularis str. Araruama* | Bacteria; Proteobacteria; Deltaproteobacteria; Desulfobacterales; Desulfobacteraceae; Candidatus Magnetoglobus | methyl-accepting chemotaxis protein | 6 |
| **OJY13089.1** | 7.7E-06 | Qa.IV | IKEIGSTISQISDIASTIATAVEQQSSATQEIANNVQN----VAQGTEE---- | *Rhizobiales bacterium 62-47* | Bacteria; Proteobacteria; Alphaproteobacteria; Rhizobiales | methyl-accepting chemotaxis protein | 6 |
| **RAI19903.1** | 7.7E-06 | Qa.III.b | -------IQQIDEIASTIASSVEEQGSATGEIAQNVQQAAQGTQEVSS----- | *Afifella marina DSM 2698* | Bacteria; Proteobacteria; Alphaproteobacteria; Rhizobiales; Rhodobiaceae; Afifella | methyl-accepting chemotaxis protein | 6 |
| **WP_090852539.1** | 7.7E-06 | Qa.III | -----KSMKETNEAFEHISTTTVDTTHHLDQLATDVQTFKEQMRKVVKE---- | *Paraliobacillus sp. PM-2* | Bacteria; Firmicutes; Bacilli; Bacillales; Bacillaceae; Paraliobacillus | methyl-accepting chemotaxis protein | 6 |
| **WP_092293350.1** | 7.7E-06 | Qc.III | IQSIGTTIQRVNEIAAAIAAAVEQQGAATREIARNVQQASQGTNEVSRHIS-- | *Bradyrhizobium sp. Ghvi* | Bacteria; Proteobacteria; Alphaproteobacteria; Rhizobiales; Bradyrhizobiaceae; Bradyrhizobium | HAMP domain-containing protein | 6 |
| **WP_097279738.1** | 7.7E-06 | Qa.IV | IQTIVRTIEEVNEVASAIAAAVEQQNAATHEISRNVQEAA------------- | *Caenispirillum bisanense* | Bacteria; Proteobacteria; Alphaproteobacteria; Rhodospirillales; Rhodospirillaceae; Caenispirillum | HAMP domain-containing protein | 6 |
| **AFT64122.1** | 7.8E-06 | SNAP.c | -GDVSDVIHEMSQIAAAVAASVEQQNAAIQSITENVANATQ------------ | *alpha proteobacterium D323* | Bacteria; Proteobacteria; Alphaproteobacteria | methyl-accepting chemotaxis sensory transducer | 6 |
| **PCJ50001.1** | 7.8E-06 | Qa.III | LREIVNAVQQVNSIIAEISSSAREQSDGIDQVNTAITQMDEATQ--------- | *Gammaproteobacteria bacterium* | Bacteria; Proteobacteria; Gammaproteobacteria | chemotaxis protein | 6 |
| **PKL94981.1** | 7.8E-06 | Qc.III | IQGIADTIGKVAEIAAAIASAVEEQSAATGEISANVTEAAASTQEVSSNIQ-- | *Gammaproteobacteria bacterium HGW-Gammaproteobacteria-8* | Bacteria; Proteobacteria; Gammaproteobacteria | methyl-accepting chemotaxis protein partial | 6 |
| **WP_009540649.1** | 7.8E-06 | Qc.III | IEGIVRTVEEVNEVASAIASAVEEQNAATSEISRNVQEAAAGSQEVSRNIS-- | *Caenispirillum salinarum* | Bacteria; Proteobacteria; Alphaproteobacteria; Rhodospirillales; Rhodospirillaceae; Caenispirillum | methyl-accepting chemotaxis protein | 6 |
| **WP_029004510.1** | 7.8E-06 | Qa.IV | --RIVNTISQIRNASSAIASAIEEQGAATQEIAVNTQQAARGTETVNQNI--- | *Azorhizobium doebereinerae* | Bacteria; Proteobacteria; Alphaproteobacteria; Rhizobiales; Xanthobacteraceae; Azorhizobium | methyl-accepting chemotaxis protein | 6 |
| **WP_057463479.1** | 7.8E-06 | SNAP.c | -GDVSDVIHEMSQIAAAVAASVEQQNAAIQSITENVANATQ------------ | *Pseudovibrio sp. POLY-S9* | Bacteria; Proteobacteria; Alphaproteobacteria; Rhodobacterales; Rhodobacteraceae; Pseudovibrio | HAMP domain-containing protein | 6 |
| **WP_063286050.1** | 7.8E-06 | SNAP.c | -GDVSDVIHEMSQIAAAVAASVEQQNAAIQSITENVANATQ------------ | *Pseudovibrio* | Bacteria; Proteobacteria; Alphaproteobacteria; Rhodobacterales; Rhodobacteraceae | MULTISPECIES: HAMP domain-containing protein | 6 |
| **WP_063291179.1** | 7.8E-06 | SNAP.c | -GDVSDVIHEMSQIAAAVAASVEQQNAAIQSITENVANATQ------------ | *Pseudovibrio sp. Ad5* | Bacteria; Proteobacteria; Alphaproteobacteria; Rhodobacterales; Rhodobacteraceae; Pseudovibrio | HAMP domain-containing protein | 6 |
| **WP_063294436.1** | 7.8E-06 | SNAP.c | -GDVSDVIHEMSQIAAAVAASVEQQNAAIQSITENVANATQ------------ | *Pseudovibrio sp. Ad37* | Bacteria; Proteobacteria; Alphaproteobacteria; Rhodobacterales; Rhodobacteraceae; Pseudovibrio | HAMP domain-containing protein | 6 |
| **WP_063301614.1** | 7.8E-06 | SNAP.c | -GDVSDVIHEMSQIAAAVAASVEQQNAAIQSITENVANATQ------------ | *Pseudovibrio sp. WM33* | Bacteria; Proteobacteria; Alphaproteobacteria; Rhodobacterales; Rhodobacteraceae; Pseudovibrio | HAMP domain-containing protein | 6 |
| **WP_063305055.1** | 7.8E-06 | SNAP.c | -GDVSDVIHEMSQIAAAVAASVEQQNAAIQSITENVANATQ------------ | *Pseudovibrio sp. W64* | Bacteria; Proteobacteria; Alphaproteobacteria; Rhodobacterales; Rhodobacteraceae; Pseudovibrio | HAMP domain-containing protein | 6 |
| **WP_063305912.1** | 7.8E-06 | SNAP.c | -GDVSDVIHEMSQIAAAVAASVEQQNAAIQSITENVANATQ------------ | *Pseudovibrio sp. Ad46* | Bacteria; Proteobacteria; Alphaproteobacteria; Rhodobacterales; Rhodobacteraceae; Pseudovibrio | HAMP domain-containing protein | 6 |
| **WP_063308799.1** | 7.8E-06 | SNAP.c | -GDVSDVIHEMSQIAAAVAASVEQQNAAIQSITENVANATQ------------ | *Pseudovibrio sp. Ad26* | Bacteria; Proteobacteria; Alphaproteobacteria; Rhodobacterales; Rhodobacteraceae; Pseudovibrio | HAMP domain-containing protein | 6 |
| **WP_063316968.1** | 7.8E-06 | SNAP.c | -GDVSDVIHEMSQIAAAVAASVEQQNAAIQSITENVANATQ------------ | *Pseudovibrio sp. Ad13* | Bacteria; Proteobacteria; Alphaproteobacteria; Rhodobacterales; Rhodobacteraceae; Pseudovibrio | HAMP domain-containing protein | 6 |
| **WP_073372822.1** | 7.8E-06 | Qa.III | IKNITEVINEINEIITTIATAVEQQSAATSEISENVAQAA------------- | *Desulfofustis glycolicus* | Bacteria; Proteobacteria; Deltaproteobacteria; Desulfobacterales; Desulfobulbaceae; Desulfofustis | methyl-accepting chemotaxis protein | 6 |
| **WP_085598972.1** | 7.8E-06 | Qb.I | LQRTRLQLQDTIEQSEVSSEELTASNEEMQAINEELRSASEELETSK------ | *Pseudomonas sp. B10(2017)* | Bacteria; Proteobacteria; Gammaproteobacteria; Pseudomonadales; Pseudomonadaceae; Pseudomonas | response regulator | 6 |
| **WP_093515864.1** | 7.8E-06 | SNAP.c | -GDVSDVIHEMSQIAAAVAASVEQQNAAIQSITENVANATQ------------ | *Pseudovibrio* | Bacteria; Proteobacteria; Alphaproteobacteria; Rhodobacterales; Rhodobacteraceae | MULTISPECIES: HAMP domain-containing protein | 6 |
| **WP_100386883.1** | 7.8E-06 | Qc.III | IKEIGATIGRISEISSTIASAVEEQGAATQEISRNVQRAAEG----------- | *Afipia broomeae* | Bacteria; Proteobacteria; Alphaproteobacteria; Rhizobiales; Bradyrhizobiaceae; Afipia | HAMP domain-containing protein | 6 |
| **WP_108818719.1** | 7.8E-06 | SNAP.c | -GDVSDVIHEMSQIAAAVAASVEQQNAAIQSITENVANATQ------------ | *Pseudovibrio sp. Alg231-02* | Bacteria; Proteobacteria; Alphaproteobacteria; Rhodobacterales; Rhodobacteraceae; Pseudovibrio | HAMP domain-containing protein | 6 |
| **EGY01659.1** | 7.9E-06 | Qc.III | --AISGTIENINSIATTIASAVEEQGAATQEIARNVQQAAAGADSVTHNIQ-- | *Nitrospirillum amazonense Y2* | Bacteria; Proteobacteria; Alphaproteobacteria; Rhodospirillales; Rhodospirillaceae; Nitrospirillum | Methyl-accepting chemotaxis receptor/ sensory transducer | 6 |
| **OJX68612.1** | 7.9E-06 | Qc.III | -SSISQTIENINELSSAISAAVEEQGAATQEIARSVGQAAEGTH--------- | *Magnetospirillum sp. 64-120* | Bacteria; Proteobacteria; Alphaproteobacteria; Rhodospirillales; Rhodospirillaceae; Magnetospirillum | chemotaxis protein | 6 |
| **OOE86053.1** | 7.9E-06 | Qc.III | -DEISGNLSNIKNLSSDMNEAIEQLGPIVSNLKENVDSLSGAIR--------- | *Salinivibrio siamensis* | Bacteria; Proteobacteria; Gammaproteobacteria; Vibrionales; Vibrionaceae; Salinivibrio | methyl-accepting chemotaxis protein | 6 |
| **PJJ33041.1** | 7.9E-06 | Qc.III | IKEIGATIGRISEISSTIASAVEEQGAATQEISRNVQRAAEG----------- | *Afipia broomeae* | Bacteria; Proteobacteria; Alphaproteobacteria; Rhizobiales; Bradyrhizobiaceae; Afipia | methyl-accepting chemotaxis sensory transducer with Cache sensor | 6 |
| **SDS17891.1** | 7.9E-06 | Qc.III | IKEIGATIGRISEISSTIASAVEEQGAATQEISRNVQRAAEG----------- | *Bradyrhizobium canariense* | Bacteria; Proteobacteria; Alphaproteobacteria; Rhizobiales; Bradyrhizobiaceae; Bradyrhizobium | methyl-accepting chemotaxis sensory transducer with Cache sensor | 6 |
| **SFV15920.1** | 7.9E-06 | Qc.III | IQEIGGTIERLSEVSAAIAAAVEEQGAATQEISRNVQQASVGTQEVSSNIT-- | *Bradyrhizobium arachidis* | Bacteria; Proteobacteria; Alphaproteobacteria; Rhizobiales; Bradyrhizobiaceae; Bradyrhizobium | methyl-accepting chemotaxis protein | 6 |
| **WP_011189138.1** | 7.9E-06 | Qa.III | IDEIRQVIADVDQIVSTIATAVEEQSVTTREIAINVSQAS------------- | *Desulfotalea psychrophila* | Bacteria; Proteobacteria; Deltaproteobacteria; Desulfobacterales; Desulfobulbaceae; Desulfotalea | methyl-accepting chemotaxis protein | 6 |
| **WP_054034307.1** | 7.9E-06 | Qa.III | -ETITRVISNVNELVATIASAVEEQSSATKEIAANIGQASV------------ | *Desulfatitalea tepidiphila* | Bacteria; Proteobacteria; Deltaproteobacteria; Desulfobacterales; Desulfobacteraceae; Desulfatitalea | methyl-accepting chemotaxis protein | 6 |
| **WP_066642731.1** | 7.9E-06 | Qc.III | -EQVNNGAKQIADGSQSLAQGATEQASSIEELSASISDVSDKIRQNTSQIS-- | *Clostridium sp. W14A* | Bacteria; Firmicutes; Clostridia; Clostridiales; Clostridiaceae; Clostridium | methyl-accepting chemotaxis protein | 6 |
| **WP_074213577.1** | 7.9E-06 | Qc.III | -DEISGNLSNIKNLSSDMNEAIEQLGPIVSNLKENVDSLSGAIR--------- | *Salinivibrio sp. ES.052* | Bacteria; Proteobacteria; Gammaproteobacteria; Vibrionales; Vibrionaceae; Salinivibrio | methyl-accepting chemotaxis protein | 6 |
| **WP_076400735.1** | 7.9E-06 | Qa.III | IRAISSTISQVNDISANIAAAVEEQSSVTRGIAASMQTAAESVDQVSRSLG-- | *Insolitispirillum peregrinum* | Bacteria; Proteobacteria; Alphaproteobacteria; Rhodospirillales; Rhodospirillaceae; Insolitispirillum | PAS domain S-box protein | 6 |
| **WP_077523055.1** | 7.9E-06 | Qc.III | -DEISGNLSNIKNLSSDMNEAIEQLGPIVSNLKENVDSLSGAIR--------- | *Salinivibrio kushneri* | Bacteria; Proteobacteria; Gammaproteobacteria; Vibrionales; Vibrionaceae; Salinivibrio | methyl-accepting chemotaxis protein | 6 |
| **WP_077599983.1** | 7.9E-06 | Qc.III | -DEISGNLSNIKNLSSDMNEAIEQLGPIVSNLKENVDSLSGAIR--------- | *Salinivibrio kushneri* | Bacteria; Proteobacteria; Gammaproteobacteria; Vibrionales; Vibrionaceae; Salinivibrio | methyl-accepting chemotaxis protein | 6 |
| **WP_077638489.1** | 7.9E-06 | Qc.III | -DEISGNLSNIKNLSSDMNEAIEQLGPIVSNLKENVDSLSGAIR--------- | *Salinivibrio sp. ML198* | Bacteria; Proteobacteria; Gammaproteobacteria; Vibrionales; Vibrionaceae; Salinivibrio | methyl-accepting chemotaxis protein | 6 |
| **WP_077653754.1** | 7.9E-06 | Qc.III | -DEISGNLSNIKNLSSDMNEAIEQLGPIVSNLKENVDSLSGAIR--------- | *Salinivibrio kushneri* | Bacteria; Proteobacteria; Gammaproteobacteria; Vibrionales; Vibrionaceae; Salinivibrio | methyl-accepting chemotaxis protein | 6 |
| **WP_077666261.1** | 7.9E-06 | Qc.III | -DEISGNLSNIKNLSSDMNEAIEQLGPIVSNLKENVDSLSGAIR--------- | *Salinivibrio sp. PR6* | Bacteria; Proteobacteria; Gammaproteobacteria; Vibrionales; Vibrionaceae; Salinivibrio | methyl-accepting chemotaxis protein | 6 |
| **SCY34513.1** | 8E-06 | Qa.III | -----KTIHEVNDITLTIATAIEEQSAATRDISGNVTQASERFQRVNAS---- | *Desulfoluna spongiiphila* | Bacteria; Proteobacteria; Deltaproteobacteria; Desulfobacterales; Desulfobacteraceae; Desulfoluna | methyl-accepting chemotaxis protein | 6 |
| **SNB70500.1** | 8E-06 | Qa.IV | IKDIGQTIEQVNGVTTTIASAVEEQDAATREIARNIEATA------------- | *Arboricoccus pini* | Bacteria; Proteobacteria; Alphaproteobacteria; Rhodospirillales; Geminicoccaceae; Arboricoccus | methyl-accepting chemotaxis protein | 6 |
| **WP_008454203.1** | 8E-06 | R.IV | -QEINHAVTHLDNMVQQNAELVVESAAAAGALQSQAGELAETAGHFR------ | *Enterobacter sp. Ag1* | Bacteria; Proteobacteria; Gammaproteobacteria; Enterobacterales; Enterobacteriaceae; Enterobacter | methyl-accepting chemotaxis protein | 6 |
| **WP_085540049.1** | 8E-06 | R.IV | -QEINHAVTHLDNMVQQNAELVVESAAAAGALQSQAGELAETAGHFR------ | *Cedecea sp. NFIX57* | Bacteria; Proteobacteria; Gammaproteobacteria; Enterobacterales; Enterobacteriaceae; Cedecea | methyl-accepting chemotaxis protein | 6 |
| **WP_092210799.1** | 8E-06 | Qa.III | -----KTIHEVNDITLTIATAIEEQSAATRDISGNVTQASERFQRVNAS---- | *Desulfoluna spongiiphila* | Bacteria; Proteobacteria; Deltaproteobacteria; Desulfobacterales; Desulfobacteraceae; Desulfoluna | methyl-accepting chemotaxis protein | 6 |
| **WP_109594966.1** | 8E-06 | Qa.IV | IQQIAQTIAEMSQISTSIAAAMEEQGAATQEISRNVQEAA------------- | *Microvirga sp. 17 mud 1-3* | Bacteria; Proteobacteria; Alphaproteobacteria; Rhizobiales; Methylobacteriaceae; Microvirga | HAMP domain-containing protein | 6 |

| | | | | | | | | |
|---|---|---|---|---|---|---|---|---|
| | **KGJ64754.1** | 8.1E-06 | Qa.IV | --EIGATIGQISSISTSIASAVEQQGAATQEIARSVQTVAQGTHAAATDI--- | *Bradyrhizobium diazoefficiens SEMIA 5080* | Bacteria; Proteobacteria; Alphaproteobacteria; Rhizobiales; Bradyrhizobiaceae; Bradyrhizobium | putative Methyl-accepting chemotaxis protein | 6 |
| | **OYW08423.1** | 8.1E-06 | Qc.III | --GIGATVERTSEIATAIAAAVEEQGAATQEIARSAAQVAEATQTVAQRI--- | *Rhodospirillales bacterium 12-71-4* | Bacteria; Proteobacteria; Alphaproteobacteria; Rhodospirillales; unclassified Rhodospirillales | hypothetical protein B7Z53_04755 | 6 |
| | **WP_008387844.1** | 8.1E-06 | Qc.III | IKEIGGTIGHIAEISAAIAAAVEEQGAATSEIARNVQQAALGTTQVASNITK- | *Rhodovulum sp. PH10* | Bacteria; Proteobacteria; Alphaproteobacteria; Rhodobacterales; Rhodobacteraceae; Rhodovulum | methyl-accepting chemotaxis protein | 6 |
|  | **WP_072325902.1** | 8.1E-06 | Qa.III | LEAIQRSVNDINDMNTQIASAAEEQSAVADEINRNFSSITEASEEAAQ----- | *Marinospirillum alkaliphilum* | Bacteria; Proteobacteria; Gammaproteobacteria; Oceanospirillales; Marinospirillum | methyl-accepting chemotaxis protein | 6 |
|  | **WP_085585450.1** | 8.1E-06 | Qc.III.b | ---INGNIGEINANARHVSDTVNQQYEMIAEIGRRVDGAVEQMRK-------- | *Thalassospira mesophila* | Bacteria; Proteobacteria; Alphaproteobacteria; Rhodospirillales; Rhodospirillaceae; Thalassospira | hypothetical protein | 6 |
|  | **WP_114010374.1** | 8.1E-06 | Qa.III.b | IRGIGDVIVEINEIASEIAASVDQQGGATREIAHNVNEAAKGTQEVS------ | *Cohaesibacter sp. YE-B6* | Bacteria; Proteobacteria; Alphaproteobacteria; Rhizobiales; Cohaesibacteraceae | HAMP domain-containing protein | 6 |
| | **EIM28120.1** | 8.2E-06 | Qc.III | IDNIGRIMEELSGIATAIAAAVEEQAAVTSEISVSVNQTTDHTHAVVRSIE-- | *Microvirga lotononidis* | Bacteria; Proteobacteria; Alphaproteobacteria; Rhizobiales; Methylobacteriaceae; Microvirga | methyl-accepting chemotaxis protein | 6 |
|  | **WP_073162609.1** | 8.2E-06 | Qc.I | ------MARQLHDLYTGLEQKVQERTAQLQAANELLERQRQELEKANQELS-- | *Desulfofundulus australicus* | Bacteria; Firmicutes; Clostridia; Clostridiales; Peptococcaceae; Desulfofundulus | DUF3365 domain-containing protein | 6 |
|  | **WP_103169265.1** | 8.2E-06 | Qa.IV | IRDVARTMTEIHQVTSSIAGAIEEQDAATREIAASAQMAA------------- | *bacterium YEK0313* | Bacteria | chemotaxis protein | 6 |
|  | **WP_111359126.1** | 8.2E-06 | Qc.III | IKEISGTIGKIAEISSAIAAAVEEQGAATSEISRNIQHAAAGT---------- | *Rhodoplanes elegans* | Bacteria; Proteobacteria; Alphaproteobacteria; Rhizobiales; Hyphomicrobiaceae; Rhodoplanes | methyl-accepting chemotaxis protein | 6 |
|  | **WP_010195422.1** | 8.3E-06 | Qa.III | ---ISQALRNLNSLNTSIASATLQQTHVVDDINQNVTQAAQLSQ--------- | *Pseudomonas amygdali* | Bacteria; Proteobacteria; Gammaproteobacteria; Pseudomonadales; Pseudomonadaceae; Pseudomonas | methyl-accepting chemotaxis protein partial | 6 |
|  | **WP_011322584.1** | 8.3E-06 | Qc.III.b | IESLSVAANRIADSAVEANEEIIEQNDSLRSINDEMQNLSATMEE-------- | *Natronomonas pharaonis* | Archaea; Euryarchaeota; Halobacteria; Halobacteriales; Haloarculaceae; Natronomonas | PAS domain-containing protein | 6 |
|  | **WP_083455964.1** | 8.3E-06 | Qa.III | IRQISSGIVDVNEIVGSIAAAVEEQSVSTGNIAGNVIQASQGIQEVSQ----- | *Desulfosarcina cetonica* | Bacteria; Proteobacteria; Deltaproteobacteria; Desulfobacterales; Desulfobacteraceae; Desulfosarcina | HAMP domain-containing protein | 6 |
|  | **WP_088561772.1** | 8.3E-06 | Qa.IV | IKDIGQTIEQVNGVTTTIASAVEEQDAATREIARNIEATA------------- | *Arboricoccus pini* | Bacteria; Proteobacteria; Alphaproteobacteria; Rhodospirillales; Geminicoccaceae; Arboricoccus | methyl-accepting chemotaxis protein | 6 |
|  | **EKD35812.1** | 8.4E-06 | Qa.III | IEKISKIIEEIHEMINTIATAVEEQSAATAEIADNITQAS------------- | *uncultured bacterium* | Bacteria; environmental samples | hypothetical protein ACD_75C01769G0001 | 6 |
|  | **OYX85085.1** | 8.4E-06 | Qa.IV | --KIATAIDEVQNISTTVASAVEEQGAATREIAENTHRAARRTE--------- | *Azorhizobium sp. 32-67-21* | Bacteria; Proteobacteria; Alphaproteobacteria; Rhizobiales; Xanthobacteraceae; Azorhizobium | hypothetical protein B7Y84_15970 | 6 |
|  | **OZA01244.1** | 8.4E-06 | Qa.IV | --KIATAIDEVQNISTTVASAVEEQGAATREIAENTHRAARRTE--------- | *Rhizobiales bacterium 17-65-6* | Bacteria; Proteobacteria; Alphaproteobacteria; Rhizobiales | hypothetical protein B7X99_01395 | 6 |
|  | **RAU37215.1** | 8.4E-06 | Qb.I | LQRTKEQLQLTIEQSETSTEELKASNEELQAINEELRSATEELETS------- | *Pseudomonas sp. RIT 411* | Bacteria; Proteobacteria; Gammaproteobacteria; Pseudomonadales; Pseudomonadaceae; Pseudomonas | PAS domain S-box protein | 6 |
|  | **WP_007163109.1** | 8.4E-06 | Qb.I | LQRTKEQLQLTIEQSETSTEELKASNEELQAINEELRSATEELETS------- | *Pseudomonas psychrotolerans* | Bacteria; Proteobacteria; Gammaproteobacteria; Pseudomonadales; Pseudomonadaceae; Pseudomonas | response regulator | 6 |
|  | **WP_024576123.1** | 8.4E-06 | Qa.IV | IKEIGSTINLISEISSSIAAAVEEQGAATQEIARNVQYAA------------- | *Afipia* | Bacteria; Proteobacteria; Alphaproteobacteria; Rhizobiales; Bradyrhizobiaceae | MULTISPECIES: methyl-accepting chemotaxis protein | 6 |
|  | **WP_027600073.1** | 8.4E-06 | Qb.I | LQRTKEQLQLTIEQSETSTEELKASNEELQAINEELRSATEELETS------- | *Pseudomonas sp. MOIL14HWK12:I2* | Bacteria; Proteobacteria; Gammaproteobacteria; Pseudomonadales; Pseudomonadaceae; Pseudomonas | PAS domain S-box protein | 6 |
|  | **WP_027600928.1** | 8.4E-06 | Qb.I | LQRTKEQLQLTIEQSETSTEELKASNEELQAINEELRSATEELETS------- | *Pseudomonas* | Bacteria; Proteobacteria; Gammaproteobacteria; Pseudomonadales; Pseudomonadaceae | MULTISPECIES: PAS domain S-box protein | 6 |
|  | **WP_042138583.1** | 8.4E-06 | Qb.I | LQRTKEQLQLTIEQSETSTEELKASNEELQAINEELRSATEELETS------- | *Pseudomonas oryzihabitans* | Bacteria; Proteobacteria; Gammaproteobacteria; Pseudomonadales; Pseudomonadaceae; Pseudomonas | response regulator | 6 |
|  | **WP_044341651.1** | 8.4E-06 | Qb.I | LQRTKEQLQLTIEQSETSTEELKASNEELQAINEELRSATEELETS------- | *Pseudomonas oryzihabitans* | Bacteria; Proteobacteria; Gammaproteobacteria; Pseudomonadales; Pseudomonadaceae; Pseudomonas | PAS domain S-box protein | 6 |
|  | **WP_058790375.1** | 8.4E-06 | Qb.I | LQRTKEQLQLTIEQSETSTEELKASNEELQAINEELRSATEELETS------- | *Pseudomonas psychrotolerans* | Bacteria; Proteobacteria; Gammaproteobacteria; Pseudomonadales; Pseudomonadaceae; Pseudomonas | PAS domain S-box protein | 6 |
|  | **WP_059230918.1** | 8.4E-06 | Qb.I | LQRTKEQLQLTIEQSETSTEELKASNEELQAINEELRSATEELETS------- | *Pseudomonas sp. EpS/L25* | Bacteria; Proteobacteria; Gammaproteobacteria; Pseudomonadales; Pseudomonadaceae; Pseudomonas | PAS domain S-box protein | 6 |
|  | **WP_067046440.1** | 8.4E-06 | Qc.III.c | LDSMTESVLQINDLGIQIATAAEEQSSVTEEINRNMTAIQHMVQQLTS----- | *Moritella sp. JT01* | Bacteria; Proteobacteria; Gammaproteobacteria; Alteromonadales; Moritellaceae; Moritella | methyl-accepting chemotaxis protein | 6 |
|  | **WP_074585089.1** | 8.4E-06 | Qb.I | LQRTKEQLQLTIEQSETSTEELKASNEELQAINEELRSATEELETS------- | *Pseudomonas psychrotolerans* | Bacteria; Proteobacteria; Gammaproteobacteria; Pseudomonadales; Pseudomonadaceae; Pseudomonas | PAS domain S-box protein | 6 |
|  | **WP_077171517.1** | 8.4E-06 | Qb.I | LQRTKEQLQLTIEQSETSTEELKASNEELQAINEELRSATEELETS------- | *Pseudomonas psychrotolerans* | Bacteria; Proteobacteria; Gammaproteobacteria; Pseudomonadales; Pseudomonadaceae; Pseudomonas | PAS domain S-box protein | 6 |
| | **WP_082700711.1** | 8.4E-06 | Qc.III | --GISGTISRINELATIIAGAVEEQGAATAEIARNVDQANEGTREVASNIS-- | *Magnetospirillum sp. XM-1* | Bacteria; Proteobacteria; Alphaproteobacteria; Rhodospirillales; Rhodospirillaceae; Magnetospirillum | methyl-accepting chemotaxis protein | 6 |
| | **WP_084681161.1** | 8.4E-06 | Qc.III | -GKISGTVKDLNEIVTTIAAAVEEQSIATREIARNVSQAAEGINNTAENV--- | *Desulfospira joergensenii* | Bacteria; Proteobacteria; Deltaproteobacteria; Desulfobacterales; Desulfobacteraceae; Desulfospira | methyl-accepting chemotaxis protein | 6 |
| | **CUW39229.1** | 8.5E-06 | Qc.III | --GISGTISRINELATIIAGAVEEQGAATAEIARNVDQANEGTREVASNIS-- | *Magnetospirillum sp. XM-1* | Bacteria; Proteobacteria; Alphaproteobacteria; Rhodospirillales; Rhodospirillaceae; Magnetospirillum | putative Methyl-accepting chemotaxis protein | 6 |
| | **OSQ26706.1** | 8.5E-06 | Qc.III | -RRISETIAKVADSSTAIAAAVEEQHAAIGEIARNVEEAANGTREVSERIE-- | *Thalassospira sp. MCCC 1A03138* | Bacteria; Proteobacteria; Alphaproteobacteria; Rhodospirillales; Rhodospirillaceae; Thalassospira | chemotaxis protein partial | 6 |
| | **WP_063691176.1** | 8.5E-06 | Qc.III | -SEISGTIARLSEISSAIAAAVEEQGAATQEIARNVQQAAQGTQQVSSKVGD- | *Bradyrhizobium stylosanthis* | Bacteria; Proteobacteria; Alphaproteobacteria; Rhizobiales; Bradyrhizobiaceae; Bradyrhizobium | PAS domain S-box protein | 6 |
| | **WP_083223032.1** | 8.5E-06 | Qc.III | ---TGRTISQIDEIATAIASAVEEQGASTENIASNMQEVAENSEMVRSRIVK- | *Terasakiella sp. PR1* | Bacteria; Proteobacteria; Alphaproteobacteria; Rhizobiales; Methylocystaceae; Terasakiella | hypothetical protein | 6 |
| | **WP_097278773.1** | 8.5E-06 | Qc.III | IEGISGTIVEINEIAAAIAAAVEEQGAATGEISRNVQEAARGSQEVTESI--- | *Caenispirillum bisanense* | Bacteria; Proteobacteria; Alphaproteobacteria; Rhodospirillales; Rhodospirillaceae; Caenispirillum | HAMP domain-containing protein | 6 |
|  | **WP_106651308.1** | 8.5E-06 | Qa.III | LENILNAISTISRMSEQIARAVNEQTSVVNSVSESVLIIKSNAQ--------- | *Shewanella sp. Pdp11* | Bacteria; Proteobacteria; Gammaproteobacteria; Alteromonadales; Shewanellaceae; Shewanella | hypothetical protein | 6 |
|  | **WP_114103332.1** | 8.5E-06 | Qa.III | --AITSSINELDDITTALSSAVEQQASATHEITQNAQAANDTTRRAAD----- | *Thalassospira profundimaris* | Bacteria; Proteobacteria; Alphaproteobacteria; Rhodospirillales; Rhodospirillaceae; Thalassospira | HAMP domain-containing protein | 6 |
|  | **OCC14532.1** | 8.6E-06 | Qa.III | -NQIAYSVEEVTNLANTIASATEEQTATINEITENVSNVNS------------ | *Dissulfuribacter thermophilus* | Bacteria; Proteobacteria; Deltaproteobacteria; Dissulfuribacter | Methyl-accepting chemotaxis protein | 6 |
|  | **PWB64157.1** | 8.6E-06 | Qa.IV | --NIQSTIREVGEITSAIAAAVTEQGAATREIARSAETASHRTRE-------- | *Bradyrhizobiaceae bacterium* | Bacteria; Proteobacteria; Alphaproteobacteria; Rhizobiales; Bradyrhizobiaceae | methyl-accepting chemotaxis protein | 6 |
| | **WP_019644397.1** | 8.6E-06 | Qa.III.b | IRSIGSVIGTINGLSGDIAGAVDDQGQSTREIAQNVAEAAKGTQEVSRSVIG- | *Novispirillum itersonii* | Bacteria; Proteobacteria; Alphaproteobacteria; Rhodospirillales; Rhodospirillaceae; Novispirillum | methyl-accepting chemotaxis protein | 6 |
|  | **WP_092027789.1** | 8.6E-06 | Qa.IV | IKEIGATISRISDISTTIAATIEEQGAATAEIARNVSEAAK------------ | *Bradyrhizobium sp. OK095* | Bacteria; Proteobacteria; Alphaproteobacteria; Rhizobiales; Bradyrhizobiaceae; Bradyrhizobium | HAMP domain-containing protein | 6 |
|  | **BAS01002.1** | 8.7E-06 | Qa.III.b | --QITEIIRQLNEIATTIASAVEEQGAATTEIARNVQQASSGTAEVSSN---- | *Blastochloris viridis* | Bacteria; Proteobacteria; Alphaproteobacteria; Rhizobiales; Hyphomicrobiaceae; Blastochloris | methyl-accepting chemotaxis protein | 6 |
|  | **KJE42679.1** | 8.7E-06 | Qa.III | -TEIVTSISDINAMVEQIAVAAEEQVNVTNEINQNISDIDHKSQ--------- | *Thalassomonas viridans* | Bacteria; Proteobacteria; Gammaproteobacteria; Alteromonadales; Colwelliaceae; Thalassomonas | hypothetical protein SG34_27595 | 6 |
| | **PDT62821.1** | 8.7E-06 | Qa.IV | --EIGATIGQISSISTSIASAVEQQGAATQEIARSVQTVAQGTQTAATDI--- | *Bradyrhizobium ottawaense* | Bacteria; Proteobacteria; Alphaproteobacteria; Rhizobiales; Bradyrhizobiaceae; Bradyrhizobium | methyl-accepting chemotaxis protein partial | 6 |
|  | **SHE32308.1** | 8.7E-06 | Qa.III | LDKIQQSVATINDRAAQIATATEEQTSVTEEVNQNTT----DIQRGSKELV-- | *Marinomonas polaris DSM 16579* | Bacteria; Proteobacteria; Gammaproteobacteria; Oceanospirillales; Marinomonas | Methyl-accepting chemotaxis protein (MCP) signalling domain-containing protein | 6 |
|  | **WP_047763914.1** | 8.7E-06 | Qa.III | --QIDNVIKEINDIAGGISTAVEEQGGATREISQTVNQSANN----------- | *Kiloniella spongiae* | Bacteria; Proteobacteria; Alphaproteobacteria; Kiloniellales; Kiloniellaceae; Kiloniella | HAMP domain-containing protein | 6 |
|  | **WP_052716189.1** | 8.7E-06 | Qa.III | IETIARVIADIAEISTNIAAAVEQQGVATQEIARNVDEASD------------ | *Magnetospira sp. QH-2* | Bacteria; Proteobacteria; Alphaproteobacteria; Rhodospirillales; Rhodospirillaceae; Magnetospira | HAMP domain-containing protein | 6 |
|  | **WP_083186757.1** | 8.7E-06 | Qa.III | -NQIAYSVEEVTNLANTIASATEEQTATINEITENVSNVNS------------ | *Dissulfuribacter thermophilus* | Bacteria; Proteobacteria; Deltaproteobacteria; Dissulfuribacter | bacteriohemerythrin | 6 |
| | **WP_084175090.1** | 8.7E-06 | Qc.III | IEAISQTISRMSEIATAIASAVEEQGAATREIARNVQEAATGTSEVSSNI--- | *Afifella pfennigii* | Bacteria; Proteobacteria; Alphaproteobacteria; Rhizobiales; Rhodobiaceae; Afifella | methyl-accepting chemotaxis protein | 6 |
|  | **WP_085620566.1** | 8.7E-06 | Qa.I | IDSIGTTIAEISEIASTIASAVEQQGAATEEINRNVQEAS------------- | *Thalassospira alkalitolerans* | Bacteria; Proteobacteria; Alphaproteobacteria; Rhodospirillales; Rhodospirillaceae; Thalassospira | methyl-accepting chemotaxis protein | 6 |
|  | **WP_108548374.1** | 8.7E-06 | Qa.IV | IQEVSATVTRMNEITATIAAAIEQQGAATREIADNINNAAE------------ | *Azospirillum humicireducens* | Bacteria; Proteobacteria; Alphaproteobacteria; Rhodospirillales; Rhodospirillaceae; Azospirillum | hypothetical protein | 6 |
| | **WP_049973819.1** | 8.8E-06 | Qc.III | --AISGTIGNINNIATTIASAVEEQGAATQEIARNVQQAAAGADSVTQNI--- | *Azospirillum sp. B4* | Bacteria; Proteobacteria; Alphaproteobacteria; Rhodospirillales; Rhodospirillaceae; Azospirillum | methyl-accepting chemotaxis protein | 6 |
|  | **WP_054696677.1** | 8.8E-06 | Qa.III | IDEIAKVIGNINDIVGTMAAAVDEQSAATREIANNIAQASQGIQEV------- | *Desulfosarcina cetonica* | Bacteria; Proteobacteria; Deltaproteobacteria; Desulfobacterales; Desulfobacteraceae; Desulfosarcina | methyl-accepting chemotaxis protein | 6 |
|  | **WP_082416782.1** | 8.8E-06 | Qa.III.b | --QITEIIRQLNEIATTIASAVEEQGAATTEIARNVQQASSGTAEVSSN---- | *Blastochloris viridis* | Bacteria; Proteobacteria; Alphaproteobacteria; Rhizobiales; Hyphomicrobiaceae; Blastochloris | HAMP domain-containing protein | 6 |
|  | **WP_086058523.1** | 8.8E-06 | Qb.I | LQRTKEQLQLTMEQSETSTEELKASNEELQAINEELRSTTEELETGK------ | *Bordetella genomosp. 9* | Bacteria; Proteobacteria; Betaproteobacteria; Burkholderiales; Alcaligenaceae; Bordetella | PAS domain S-box protein | 6 |
|  | **WP_086073085.1** | 8.8E-06 | Qb.I | LQRTKEQLQLTMEQSETSTEELKASNEELQAINEELRSTTEELETGK------ | *Bordetella genomosp. 9* | Bacteria; Proteobacteria; Betaproteobacteria; Burkholderiales; Alcaligenaceae; Bordetella | PAS domain S-box protein | 6 |
| | **WP_103969270.1** | 8.8E-06 | R.IV | -AQLNQAMDDMTRVTLDNQQLVILSTQSAHALREKAEHMQAHAGRFRITA--- | *Halomonas daqingensis* | Bacteria; Proteobacteria; Gammaproteobacteria; Oceanospirillales; Halomonadaceae; Halomonas | HAMP domain-containing protein | 6 |
|  | **OYU90115.1** | 8.9E-06 | Qa.IV | IQEIGTTIGQMSEIASTIASAVEQQGAATQEIARNVQQAA------------- | *Bradyrhizobiaceae bacterium PARB1* | Bacteria; Proteobacteria; Alphaproteobacteria; Rhizobiales; Bradyrhizobiaceae | methyl-accepting chemotaxis protein | 6 |
|  | **WP_084724148.1** | 8.9E-06 | Qa.III | -TEIVTSISDINAMVEQIAVAAEEQVNVTNEINQNISDIDHKSQ--------- | *Thalassomonas viridans* | Bacteria; Proteobacteria; Gammaproteobacteria; Alteromonadales; Colwelliaceae; Thalassomonas | HAMP domain-containing protein | 6 |

| | | | | | | | | |
|---|---|---|---|---|---|---|---|---|
|  | WP_092304996.1 | 8.9E-06 | Qb.I | LQRTKLHLQDTIEQSEVSSEELKASNEEMQALNEELRSATEELETS------- | *Pseudomonas sp. NFIX28* | Bacteria; Proteobacteria; Gammaproteobacteria; Pseudomonadales; Pseudomonadaceae; Pseudomonas | PAS domain S-box protein | 6 |
| | KYG25509.1 | 9E-06 | Qa.IV | --EIGATIGQISSISTSIASAVEQQGAATQEIARSVQTVAQGTQTAATDI--- | *Bradyrhizobium sp. AT1* | Bacteria; Proteobacteria; Alphaproteobacteria; Rhizobiales; Bradyrhizobiaceae; Bradyrhizobium | chemotaxis protein partial | 6 |
| | WP_029584445.1 | 9E-06 | Qc.III | -REIGATIGQIDEIATTIAAAVEEQGGSTNEIARSVQQAAKGTHEVMKNIA-- | *Bradyrhizobium sp. URHD0069* | Bacteria; Proteobacteria; Alphaproteobacteria; Rhizobiales; Bradyrhizobiaceae; Bradyrhizobium | HAMP domain-containing protein | 6 |
|  | WP_100233297.1 | 9E-06 | Qa.IV | IKEIGATIARISDISTTIAATIEQQGAATAEIARNVSEAAK------------ | *Bradyrhizobium sp. INPA54B* | Bacteria; Proteobacteria; Alphaproteobacteria; Rhizobiales; Bradyrhizobiaceae; Bradyrhizobium | chemotaxis protein | 6 |
|  | BAR53379.1 | 9.1E-06 | Qa.IV | IRDISATIERLSEVSSTIPAAVEEQGAATQEISRNVQQAA------------- | *Bradyrhizobium diazoefficiens* | Bacteria; Proteobacteria; Alphaproteobacteria; Rhizobiales; Bradyrhizobiaceae; Bradyrhizobium | hypothetical protein NK6_191 | 6 |
|  | EFO47568.1 | 9.1E-06 | Qa.III | LDEINAAVALISDMATQIATAAEEQTHVTNEITQNVTSIK------------- | *Vibrio parahaemolyticus AQ4037* | Bacteria; Proteobacteria; Gammaproteobacteria; Vibrionales; Vibrionaceae; Vibrio | histidine kinase | 6 |
|  | OAZ11460.1 | 9.1E-06 | Qa.IV | IRTIVESVQGINEILRLTAESVMEQGNATGQISTSITEAAGY----------- | *Thalassospira profundimaris* | Bacteria; Proteobacteria; Alphaproteobacteria; Rhodospirillales; Rhodospirillaceae; Thalassospira | chemotaxis protein | 6 |
| | SCY64237.1 | 9.1E-06 | Qa.IV | INEINTSIHNVNEMVGAIAAAVEEQSATTREIAENVNQAAQGVDDVNTNIA-- | *Desulfoluna spongiiphila* | Bacteria; Proteobacteria; Deltaproteobacteria; Desulfobacterales; Desulfobacteraceae; Desulfoluna | Methyl-accepting chemotaxis protein | 6 |
| | SEG02704.1 | 9.1E-06 | R.IV | -AQLNQAMDDMTRVTLDNQQLVILSTQSAHALREKAEHMQAHAGRFRITA--- | *Halomonas daqingensis* | Bacteria; Proteobacteria; Gammaproteobacteria; Oceanospirillales; Halomonadaceae; Halomonas | methyl-accepting chemotaxis sensory transducer with TarH sensor | 6 |
|  | WP_066344963.1 | 9.1E-06 | Qb.I | LQRTKEQLQLTMEQSETSTEELKASNEELQAINEELRSTTEELETGK------ | *Bordetella bronchialis* | Bacteria; Proteobacteria; Betaproteobacteria; Burkholderiales; Alcaligenaceae; Bordetella | PAS domain S-box protein | 6 |
|  | WP_066668370.1 | 9.1E-06 | Qb.I | LQRTKEQLQLTMEQSETSTEELKASNEELQAINEELRSTTEELETGK------ | *Bordetella bronchialis* | Bacteria; Proteobacteria; Betaproteobacteria; Burkholderiales; Alcaligenaceae; Bordetella | PAS domain S-box protein | 6 |
|  | WP_068004194.1 | 9.1E-06 | Qc.III.c | ---IGETIGQINTLSTGINEAVGEQKQATDEISLNLNHAHQNVSQV------- | *Pseudovibrio axinellae* | Bacteria; Proteobacteria; Alphaproteobacteria; Rhodobacterales; Rhodobacteraceae; Pseudovibrio | methyl-accepting chemotaxis protein | 6 |
|  | WP_073954610.1 | 9.1E-06 | Qa.IV | ------TIRELDTSTGRIAAAVEEQGAATQEIARNVDEAARGVGRVNSDIDAA | *Thalassospira sp. TSL5-1* | Bacteria; Proteobacteria; Alphaproteobacteria; Rhodospirillales; Rhodospirillaceae; Thalassospira | methyl-accepting chemotaxis protein | 6 |
|  | OQX57763.1 | 9.2E-06 | Qa.III | ---------ELSRITSEISSMAQEGQEAIRQISDTMASVKSSMEEVSRK---- | *Thermotoga sp. 4484_232* | Bacteria; Thermotogae; Thermotogales; Thermotogaceae; Thermotoga | hypothetical protein B5M49_03105 | 6 |
|  | WP_004274109.1 | 9.2E-06 | Qa.IV | IITISRTIAEINDISDMVAGAVEEQDAATKEIARNVQQAS------------- | *Nitrospirillum amazonense* | Bacteria; Proteobacteria; Alphaproteobacteria; Rhodospirillales; Rhodospirillaceae; Nitrospirillum | HAMP domain-containing protein | 6 |
|  | WP_039588581.1 | 9.2E-06 | Qb.I | LQRTKSHLQDTIEQAEVSSEELKASNEEMQAINEELRSATEELETS------- | *Pseudomonas frederiksbergensis* | Bacteria; Proteobacteria; Gammaproteobacteria; Pseudomonadales; Pseudomonadaceae; Pseudomonas | PAS domain S-box protein | 6 |
|  | WP_045155960.1 | 9.2E-06 | Qb.I | LQRTKSHLQDTIEQAEVSSEELKASNEEMQAINEELRSATEELETS------- | *Pseudomonas fluorescens* | Bacteria; Proteobacteria; Gammaproteobacteria; Pseudomonadales; Pseudomonadaceae; Pseudomonas | PAS domain S-box protein | 6 |
|  | WP_085899250.1 | 9.2E-06 | Qa.IV | ---ITGTIEQINEITSSVAAAVEEQGAATLEISRNVQEAAAAT---------- | *Kiloniella majae* | Bacteria; Proteobacteria; Alphaproteobacteria; Kiloniellales; Kiloniellaceae; Kiloniella | methyl-accepting chemotaxis protein | 6 |
|  | WP_085908912.1 | 9.2E-06 | Qa.IV | ---ITGTIEQINEITSSVAAAVEEQGAATLEISRNVQEAAAAT---------- | *Kiloniella majae* | Bacteria; Proteobacteria; Alphaproteobacteria; Kiloniellales; Kiloniellaceae; Kiloniella | methyl-accepting chemotaxis protein | 6 |
|  | WP_088870889.1 | 9.2E-06 | Qa.IV | IITISRTIAEINDISDMVAGAVEEQDAATKEIARNVQQAS------------- | *Nitrospirillum amazonense* | Bacteria; Proteobacteria; Alphaproteobacteria; Rhodospirillales; Rhodospirillaceae; Nitrospirillum | methyl-accepting chemotaxis protein | 6 |
|  | WP_094855171.1 | 9.2E-06 | Qb.I | LQRTKEQLQLTMEQSETSTEELKASNEELQAINEELRSTTEELETGK------ | *Bordetella genomosp. 10* | Bacteria; Proteobacteria; Betaproteobacteria; Burkholderiales; Alcaligenaceae; Bordetella | PAS domain S-box protein | 6 |
|  | WP_109070227.1 | 9.2E-06 | Qa.III.b | ---VARVIEQVNGIATIIASAVEEQGAATQAISRNVQQAAEVTREVARD---- | *Azospirillum sp. TSH58* | Bacteria; Proteobacteria; Alphaproteobacteria; Rhodospirillales; Rhodospirillaceae; Azospirillum | HAMP domain-containing protein | 6 |
|  | CUU20990.1 | 9.3E-06 | Qc.III | IQEIGGTIERLSEVAAAIAAAVEEQGAATQEISRNVQQASVGTQ--------- | *Bradyrhizobium sp.* | Bacteria; Proteobacteria; Alphaproteobacteria; Rhizobiales; Bradyrhizobiaceae; Bradyrhizobium | blr2548 putative chemotaxis protein CDS | 6 |
|  | OSJ29416.1 | 9.3E-06 | Qc.III | IQEIGGTIERLSEVAAAIAAAVEEQGAATQEISRNVQQASVGTQ--------- | *Bradyrhizobium japonicum* | Bacteria; Proteobacteria; Alphaproteobacteria; Rhizobiales; Bradyrhizobiaceae; Bradyrhizobium | methyl-accepting chemotaxis protein | 6 |
|  | PLX37393.1 | 9.3E-06 | Qa.III | IRNIAQSVGEVSTVTTTIASAVEEQEAATREISNSIQSAAN------------ | *Rhizobiales bacterium* | Bacteria; Proteobacteria; Alphaproteobacteria; Rhizobiales | hypothetical protein C0605_07450 | 6 |
|  | WP_028152153.1 | 9.3E-06 | Qc.III | IQEIGGTIERLSEVAAAIAAAVEEQGAATQEISRNVQQASVGTQ--------- | *Bradyrhizobium japonicum* | Bacteria; Proteobacteria; Alphaproteobacteria; Rhizobiales; Bradyrhizobiaceae; Bradyrhizobium | methyl-accepting chemotaxis protein | 6 |
| | WP_046504352.1 | 9.3E-06 | Qc.III | ---ISSTIERMNEFANAIAAAIEEQGAATQEISNSVQKAAAGTQEVSSNV--- | *Kiloniella litopenaei* | Bacteria; Proteobacteria; Alphaproteobacteria; Kiloniellales; Kiloniellaceae; Kiloniella | methyl-accepting chemotaxis protein | 6 |
| | WP_047762640.1 | 9.3E-06 | Qc.III | ---ISSTIERMNEFANAIAAAIEEQGAATQEISNSVQKAAAGTQEVSSNV--- | *Kiloniella spongiae* | Bacteria; Proteobacteria; Alphaproteobacteria; Kiloniellales; Kiloniellaceae; Kiloniella | methyl-accepting chemotaxis protein | 6 |
|  | WP_063985092.1 | 9.3E-06 | Qc.III | IQEIGGTIERLSEVAAAIAAAVEEQGAATQEISRNVQQASVGTQ--------- | *Bradyrhizobium sp.* | Bacteria; Proteobacteria; Alphaproteobacteria; Rhizobiales; Bradyrhizobiaceae; Bradyrhizobium | methyl-accepting chemotaxis protein | 6 |
| | WP_069956697.1 | 9.3E-06 | Qc.III | -GSIGGIINHINEITSAIAAAVEEQGAATSEIARNVEQASSGTQEVSSNIS-- | *Magnetovibrio blakemorei* | Bacteria; Proteobacteria; Alphaproteobacteria; Rhodospirillales; Rhodospirillaceae; Magnetovibrio | hypothetical protein | 6 |
|  | WP_071916120.1 | 9.3E-06 | Qc.III | IQEIGGTIERLSEVAAAIAAAVEEQGAATQEISRNVQQASVGTQ--------- | *Bradyrhizobium japonicum* | Bacteria; Proteobacteria; Alphaproteobacteria; Rhizobiales; Bradyrhizobiaceae; Bradyrhizobium | methyl-accepting chemotaxis protein | 6 |
|  | WP_081655470.1 | 9.3E-06 | SNAP.c | ------------AQARTHARALEEQNETLRRTERRLEEANTQLEASNERLEQF | *Halopiger goleimassiliensis* | Archaea; Euryarchaeota; Halobacteria; Natrialbales; Natrialbaceae; Halopiger | histidine kinase | 6 |
|  | WP_085965529.1 | 9.3E-06 | Qc.III | IQEIGGTIERLSEVAAAIAAAVEEQGAATQEISRNVQQASVGTQ--------- | *Bradyrhizobium japonicum* | Bacteria; Proteobacteria; Alphaproteobacteria; Rhizobiales; Bradyrhizobiaceae; Bradyrhizobium | methyl-accepting chemotaxis protein | 6 |
| | WP_092212600.1 | 9.3E-06 | Qa.IV | INEINTSIHNVNEMVGAIAAAVEEQSATTREIAENVNQAAQGVDDVNTNIA-- | *Desulfoluna spongiiphila* | Bacteria; Proteobacteria; Deltaproteobacteria; Desulfobacterales; Desulfobacteraceae; Desulfoluna | methyl-accepting chemotaxis protein | 6 |
|  | WP_094184203.1 | 9.3E-06 | Qc.III | IQEIGGTIERLSEVAAAIAAAVEEQGAATQEISRNVQQASVGTQ--------- | *Bradyrhizobium japonicum* | Bacteria; Proteobacteria; Alphaproteobacteria; Rhizobiales; Bradyrhizobiaceae; Bradyrhizobium | methyl-accepting chemotaxis protein | 6 |
| | WP_100085412.1 | 9.3E-06 | Qc.III | --AISGTIENINSIATTIASAVEEQGAATQEIARNVQQAAAGADSVTHNIQ-- | *Nitrospirillum amazonense* | Bacteria; Proteobacteria; Alphaproteobacteria; Rhodospirillales; Rhodospirillaceae; Nitrospirillum | chemotaxis protein partial | 6 |
| | WP_106951122.1 | 9.3E-06 | Qc.III | IKDISHTIEKLSEISSTIAAAVEEQGAATQEISRNVQQAAEGTHQVSSNITD- | *Bradyrhizobium sp. MOS003* | Bacteria; Proteobacteria; Alphaproteobacteria; Rhizobiales; Bradyrhizobiaceae; Bradyrhizobium | methyl-accepting chemotaxis protein | 6 |
|  | PUB85602.1 | 9.4E-06 | Qa.III | --EISNAIDTINNMNTQIATAAEQQCSVVDEINRNIASIVENSRQTSQ----- | *gamma proteobacterium symbiont of Ctena orbiculata* | Bacteria; Proteobacteria; Gammaproteobacteria | chemotaxis protein partial | 6 |
|  | SER23734.1 | 9.4E-06 | Qc.III.c | ---IGETIGQINTLSTGINEAVGEQKQATDEISLNLNHAHQNVSQV------- | *Pseudovibrio axinellae* | Bacteria; Proteobacteria; Alphaproteobacteria; Rhodobacterales; Rhodobacteraceae; Pseudovibrio | methyl-accepting chemotaxis protein | 6 |
| | WP_085300483.1 | 9.4E-06 | R.IV | --QATDQVATAINEMSTSIDEVTNRTAEASNAADNSKQLVNTSSAAVNQT--- | *Colwellia mytili* | Bacteria; Proteobacteria; Gammaproteobacteria; Alteromonadales; Colwelliaceae; Colwellia | methyl-accepting chemotaxis protein | 6 |
| | GAJ33562.1 | 9.5E-06 | Qc.III | IQEIGGTIERLSEVSAAIAAAVEEQGAATQEISRNVQQASLGTQEVSSNIT-- | *Bradyrhizobium sp. DOA9* | Bacteria; Proteobacteria; Alphaproteobacteria; Rhizobiales; Bradyrhizobiaceae; Bradyrhizobium | protein pilJ | 6 |
| | WP_025034555.1 | 9.5E-06 | Qc.III | IQEIGGTIERLSEVSAAIAAAVEEQGAATQEISRNVQQASLGTQEVSSNIT-- | *Bradyrhizobium sp. DOA9* | Bacteria; Proteobacteria; Alphaproteobacteria; Rhizobiales; Bradyrhizobiaceae; Bradyrhizobium | HAMP domain-containing protein | 6 |
|  | WP_028164403.1 | 9.5E-06 | Qa.IV | IRNISTTIGEINDVTTAIAAAVEEQGAATREIARNIQHAA------------- | *Bradyrhizobium elkanii* | Bacteria; Proteobacteria; Alphaproteobacteria; Rhizobiales; Bradyrhizobiaceae; Bradyrhizobium | HAMP domain-containing protein | 6 |
| | WP_036038279.1 | 9.5E-06 | Qc.III | IQEIGGTIERLSEVSAAIAAAVEEQGAATQEISRNVQQASLGTQEVSSNIT-- | *Bradyrhizobium yuanmingense* | Bacteria; Proteobacteria; Alphaproteobacteria; Rhizobiales; Bradyrhizobiaceae; Bradyrhizobium | methyl-accepting chemotaxis protein | 6 |
| | WP_050733983.1 | 9.5E-06 | Qc.III | --SIQDTIVRLNEVATAISSAVEQQSAVTREMSASMQTAAQGVGAITTNL--- | *Methylobacterium sp. ARG-1* | Bacteria; Proteobacteria; Alphaproteobacteria; Rhizobiales; Methylobacteriaceae; Methylobacterium | methyl-accepting chemotaxis protein | 6 |
| | WP_065731903.1 | 9.5E-06 | Qc.III | IKEIGDTIGRMSEIAATIAAAVEQQGAATQEISRNVQQAANGTQQVSSNIAD- | *Bradyrhizobium icense* | Bacteria; Proteobacteria; Alphaproteobacteria; Rhizobiales; Bradyrhizobiaceae; Bradyrhizobium | methyl-accepting chemotaxis protein | 6 |
|  | WP_085906239.1 | 9.5E-06 | Qa.III.b | -EQIDSVIKEINDIAGGISTAVEEQGGATREISQTVNQSANN----------- | *Kiloniella majae* | Bacteria; Proteobacteria; Alphaproteobacteria; Kiloniellales; Kiloniellaceae; Kiloniella | HAMP domain-containing protein | 6 |
|  | WP_085907855.1 | 9.5E-06 | Qa.III.b | -EQIDSVIKEINDIAGGISTAVEEQGGATREISQTVNQSANN----------- | *Kiloniella majae* | Bacteria; Proteobacteria; Alphaproteobacteria; Kiloniellales; Kiloniellaceae; Kiloniella | HAMP domain-containing protein | 6 |
| | WP_085968859.1 | 9.5E-06 | Qc.III | IQEIGGTIERLSEVSAAIAAAVEEQGAATQEISRNVQQASLGTQEVSSNIT-- | *Bradyrhizobium yuanmingense* | Bacteria; Proteobacteria; Alphaproteobacteria; Rhizobiales; Bradyrhizobiaceae; Bradyrhizobium | methyl-accepting chemotaxis protein | 6 |
| | WP_085972346.1 | 9.5E-06 | Qc.III | IQEIGGTIERLSEVSAAIAAAVEEQGAATQEISRNVQQASLGTQEVSSNIT-- | *Bradyrhizobium yuanmingense* | Bacteria; Proteobacteria; Alphaproteobacteria; Rhizobiales; Bradyrhizobiaceae; Bradyrhizobium | methyl-accepting chemotaxis protein | 6 |
| | WP_085974497.1 | 9.5E-06 | Qc.III | IQEIGGTIERLSEVSAAIAAAVEEQGAATQEISRNVQQASLGTQEVSSNIT-- | *Bradyrhizobium yuanmingense* | Bacteria; Proteobacteria; Alphaproteobacteria; Rhizobiales; Bradyrhizobiaceae; Bradyrhizobium | methyl-accepting chemotaxis protein | 6 |
|  | BAI73053.1 | 9.6E-06 | Qa.III | IQGISNIIEQIKGISTTISSAVEEQGSATREIARNVQQAAQGTHQ-------- | *Azospirillum sp. B510* | Bacteria; Proteobacteria; Alphaproteobacteria; Rhodospirillales; Rhodospirillaceae; Azospirillum | methyl-accepting chemotaxis protein | 6 |
|  | PWC69140.1 | 9.6E-06 | Qc.III | -RDIADTVRRVNDIAGSIAQAVERQSAATGAISRNVQQAAEG----------- | *Azospirillum sp. TSH58* | Bacteria; Proteobacteria; Alphaproteobacteria; Rhodospirillales; Rhodospirillaceae; Azospirillum | chemotaxis protein partial | 6 |
|  | WP_008071810.1 | 9.6E-06 | Qb.I | LQRTKLHLQDTIEQSEVSSEELKASNEEMQAINEELRSATEELETS------- | *Pseudomonas sp. GM79* | Bacteria; Proteobacteria; Gammaproteobacteria; Pseudomonadales; Pseudomonadaceae; Pseudomonas | PAS domain S-box protein | 6 |
|  | WP_008147940.1 | 9.6E-06 | Qb.I | LQRTKLHLQDTIEQSEVSSEELKASNEEMQAINEELRSATEELETS------- | *Pseudomonas sp. GM41(2012)* | Bacteria; Proteobacteria; Gammaproteobacteria; Pseudomonadales; Pseudomonadaceae; Pseudomonas | PAS domain S-box protein | 6 |
|  | WP_020005102.1 | 9.6E-06 | Qb.I | ----KNMMSESMEYIEGAANIIAQTTSNIEDVHQASDKIKDITK--------- | *Brachyspira innocens* | Bacteria; Spirochaetes; Brachyspirales; Brachyspiraceae; Brachyspira | methyl-accepting chemotaxis protein | 6 |
| | WP_027530187.1 | 9.6E-06 | Qc.III | -QAIGGIIGEVNEVATAIAAAVQEQGAATQEITRSTQYAAQGTKNVSENIT-- | *Bradyrhizobium sp. WSM3983* | Bacteria; Proteobacteria; Alphaproteobacteria; Rhizobiales; Bradyrhizobiaceae; Bradyrhizobium | methyl-accepting chemotaxis protein | 6 |
|  | WP_027923944.1 | 9.6E-06 | Qb.I | LQRTKLHLQDTIEQSEVSSEELKASNEEMQAINEELRSATEELETS------- | *Pseudomonas sp. URMO17WK12:I12* | Bacteria; Proteobacteria; Gammaproteobacteria; Pseudomonadales; Pseudomonadaceae; Pseudomonas | PAS domain S-box protein | 6 |
|  | WP_033055227.1 | 9.6E-06 | Qb.I | LQRTKLHLQDTIEQSEVSSEELKASNEEMQAINEELRSATEELETS------- | *Pseudomonas mandelii* | Bacteria; Proteobacteria; Gammaproteobacteria; Pseudomonadales; Pseudomonadaceae; Pseudomonas | PAS domain S-box protein | 6 |
|  | WP_046046524.1 | 9.6E-06 | Qb.I | LQRTKLHLQDTIEQSEVSSEELKASNEEMQAINEELRSATEELETS------- | *Pseudomonas fluorescens* | Bacteria; Proteobacteria; Gammaproteobacteria; Pseudomonadales; Pseudomonadaceae; Pseudomonas | PAS domain S-box protein | 6 |
|  | WP_046052821.1 | 9.6E-06 | Qb.I | LQRTKLHLQDTIEQSEVSSEELKASNEEMQAINEELRSATEELETS------- | *Pseudomonas fluorescens* | Bacteria; Proteobacteria; Gammaproteobacteria; Pseudomonadales; Pseudomonadaceae; Pseudomonas | PAS domain S-box protein | 6 |

| | | | | | | | | |
|---|---|---|---|---|---|---|---|---|
|  | **WP_052293666.1** | 9.6E-06 | Qa.III | IQGISNIIEQIKGISTTISSAVEEQGSATREIARNVQQAAQGTHQ-------- | *Azospirillum lipoferum* | Bacteria; Proteobacteria; Alphaproteobacteria; Rhodospirillales; Rhodospirillaceae; Azospirillum | methyl-accepting chemotaxis protein | 6 |
|  | **WP_059182307.1** | 9.6E-06 | Qb.I | LQRTKLHLQDTIEQSEVSSEELKASNEEMQAINEELRSATEELETS------- | *Pseudomonas sp. URMO17WK12:I11* | Bacteria; Proteobacteria; Gammaproteobacteria; Pseudomonadales; Pseudomonadaceae; Pseudomonas | PAS domain S-box protein | 6 |
|  | **WP_063342838.1** | 9.6E-06 | Qb.I | LQRTKLHLQDTIEQSEVSSEELKASNEEMQAINEELRSATEELETS------- | *Pseudomonas fluorescens* | Bacteria; Proteobacteria; Gammaproteobacteria; Pseudomonadales; Pseudomonadaceae; Pseudomonas | PAS domain S-box protein | 6 |
|  | **WP_074879192.1** | 9.6E-06 | Qb.I | LQRTKLHLQDTIEQSEVSSEELKASNEEMQAINEELRSATEELETS------- | *Pseudomonas frederiksbergensis* | Bacteria; Proteobacteria; Gammaproteobacteria; Pseudomonadales; Pseudomonadaceae; Pseudomonas | PAS domain S-box protein | 6 |
|  | **WP_076028306.1** | 9.6E-06 | Qb.I | LQRTKLHLQDTIEQSEVSSEELKASNEEMQAINEELRSATEELETS------- | *Pseudomonas frederiksbergensis* | Bacteria; Proteobacteria; Gammaproteobacteria; Pseudomonadales; Pseudomonadaceae; Pseudomonas | PAS domain S-box protein | 6 |
|  | **WP_090187036.1** | 9.6E-06 | Qb.I | LQRTKLHLQDTIEQSEVSSEELKASNEEMQAINEELRSATEELETS------- | *Pseudomonas arsenicoxydans* | Bacteria; Proteobacteria; Gammaproteobacteria; Pseudomonadales; Pseudomonadaceae; Pseudomonas | PAS domain S-box protein | 6 |
|  | **WP_090288613.1** | 9.6E-06 | Qb.I | LQRTKLHLQDTIEQSEVSSEELKASNEEMQAINEELRSATEELETS------- | *Pseudomonas granadensis* | Bacteria; Proteobacteria; Gammaproteobacteria; Pseudomonadales; Pseudomonadaceae; Pseudomonas | PAS domain S-box protein | 6 |
|  | **WP_093106542.1** | 9.6E-06 | Qb.I | LQRTKLHLQDTIEQSEVSSEELKASNEEMQAINEELRSATEELETS------- | *Pseudomonas sp. Z003-0.4C(8344-21)* | Bacteria; Proteobacteria; Gammaproteobacteria; Pseudomonadales; Pseudomonadaceae; Pseudomonas | PAS domain S-box protein | 6 |
|  | **WP_102616616.1** | 9.6E-06 | Qb.I | LQRTKLHLQDTIEQSEVSSEELKASNEEMQAINEELRSATEELETS------- | *Pseudomonas* | Bacteria; Proteobacteria; Gammaproteobacteria; Pseudomonadales; Pseudomonadaceae | MULTISPECIES: PAS domain S-box protein | 6 |
|  | **WP_111451677.1** | 9.6E-06 | Qb.I | LQRTKLHLQDTIEQSEVSSEELKASNEEMQAINEELRSATEELETS------- | *Pseudomonas sp. 478* | Bacteria; Proteobacteria; Gammaproteobacteria; Pseudomonadales; Pseudomonadaceae; Pseudomonas | PAS domain S-box protein | 6 |
|  | **WP_114395913.1** | 9.6E-06 | Qa.IV | IQDIATIIEQVREISTGISSAVEQQGAATAEIARNVQEAAD------------ | *Rhodospirillaceae bacterium NAU-10* | Bacteria; Proteobacteria; Alphaproteobacteria; Rhodospirillales; Rhodospirillaceae | HAMP domain-containing protein | 6 |
|  | **SDS52141.1** | 9.7E-06 | Qb.I | LQRTKLHLQDTIEQSEVSSEELKASNEEMQAINEELRSATEELETS------- | *Pseudomonas sp. Z003-0.4C(8344-21)* | Bacteria; Proteobacteria; Gammaproteobacteria; Pseudomonadales; Pseudomonadaceae; Pseudomonas | two-component system chemotaxis family CheB/CheR fusion protein | 6 |
|  | **SDT41756.1** | 9.7E-06 | Qb.I | LQRTKLHLQDTIEQSEVSSEELKASNEEMQAINEELRSATEELETS------- | *Pseudomonas granadensis* | Bacteria; Proteobacteria; Gammaproteobacteria; Pseudomonadales; Pseudomonadaceae; Pseudomonas | two-component system chemotaxis family CheB/CheR fusion protein | 6 |
| | **WP_026312971.1** | 9.7E-06 | Qc.III | IQEIGGTIERLSEVSAAIAAAVEEQGAATQEISRNVQQASLGTQEVSSNIT-- | *Bradyrhizobium japonicum* | Bacteria; Proteobacteria; Alphaproteobacteria; Rhizobiales; Bradyrhizobiaceae; Bradyrhizobium | methyl-accepting chemotaxis protein | 6 |
|  | **WP_028864202.1** | 9.7E-06 | Qc.III.c | LDSMTVSVVKINDLGVQIATAAEQQSSATDEINRNMSAIQGMVTQL------- | *Psychromonas aquimarina* | Bacteria; Proteobacteria; Gammaproteobacteria; Alteromonadales; Psychromonadaceae; Psychromonas | methyl-accepting chemotaxis protein | 6 |
|  | **WP_047187879.1** | 9.7E-06 | Qa.IV | IQNIARTIAEMSQISVSISAAMEEQGAATAEISRNVQEAAR------------ | *Microvirga vignae* | Bacteria; Proteobacteria; Alphaproteobacteria; Rhizobiales; Methylobacteriaceae; Microvirga | methyl-accepting chemotaxis protein | 6 |
|  | **WP_047458404.1** | 9.7E-06 | Qa.III | IKRIQGSIMQINDMNNQIAAAAEEQSSVAEEINNNTYRIKD------------ | *Vibrio furnissii* | Bacteria; Proteobacteria; Gammaproteobacteria; Vibrionales; Vibrionaceae; Vibrio | methyl-accepting chemotaxis protein | 6 |
|  | **ALV26193.1** | 9.8E-06 | Qc.III | -SRIVRTIGHIQEVAQAISAAVEEQGAATSEIAANTQRAATG----------- | *Pannonibacter phragmitetus* | Bacteria; Proteobacteria; Alphaproteobacteria; Rhodobacterales; Rhodobacteraceae; Pannonibacter | hypothetical protein APZ00_03150 | 6 |
|  | **PVE24679.1** | 9.8E-06 | Qa.IV | IQSIARTIAEMSQISVSISAAMEEQGAATAEISRNVQEAAR------------ | *Microvirga sp. KLBC 81* | Bacteria; Proteobacteria; Alphaproteobacteria; Rhizobiales; Methylobacteriaceae; Microvirga | methyl-accepting chemotaxis protein | 6 |
|  | **WP_069312574.1** | 9.8E-06 | Qa.III | LDKIQDSAGEVTKATDDIARAVSEQTEATKSVTGYMEKTLGEVE--------- | *Piscirickettsia litoralis* | Bacteria; Proteobacteria; Gammaproteobacteria; Thiotrichales; Piscirickettsiaceae; Piscirickettsia | methyl-accepting chemotaxis protein | 6 |
|  | **KJB97071.1** | 9.9E-06 | Qa.IV | IKSVLSGVSDMNEICTLIAAAVEEQAAATREIARNVDQAA------------- | *Skermanella aerolata KACC 11604* | Bacteria; Proteobacteria; Alphaproteobacteria; Rhodospirillales; Rhodospirillaceae; Skermanella | methyl-accepting chemotaxis protein | 6 |
|  | **OCS05964.1** | 9.9E-06 | Qa.IV | --EIEANVNILSQGINEMSQSINEQTEAINQINEAVATVDEQTKQ-NV----- | *Campylobacter fetus subsp. testudinum* | Bacteria; Proteobacteria; Epsilonproteobacteria; Campylobacterales; Campylobacteraceae; Campylobacter | chemotaxis protein partial | 6 |
| | **SFV18060.1** | 9.9E-06 | Qa.IV | --EIGATIGQISSISTSIASAVEQQGAATQEIARSVQTVAQGTQTAATDI--- | *Bradyrhizobium arachidis* | Bacteria; Proteobacteria; Alphaproteobacteria; Rhizobiales; Bradyrhizobiaceae; Bradyrhizobium | Methyl-accepting chemotaxis protein (MCP) signalling domain-containing protein partial | 6 |
|  | **WP_005347961.1** | 9.9E-06 | Qa.III | LEQITSSINQISDMATQISSAAEEQRAVTDEVSRNIQAVK------------- | *Aeromonas veronii* | Bacteria; Proteobacteria; Gammaproteobacteria; Aeromonadales; Aeromonadaceae; Aeromonas | methyl-accepting chemotaxis protein | 6 |
|  | **WP_011664925.1** | 9.9E-06 | Qc.III | IQEISDTIGRMSEIASTIASAVEEQGAATQEISRNVQQAAQGTQRVSAN---- | *Rhodopseudomonas palustris* | Bacteria; Proteobacteria; Alphaproteobacteria; Rhizobiales; Bradyrhizobiaceae; Rhodopseudomonas | HAMP domain-containing protein | 6 |
| | **WP_090074979.1** | 9.9E-06 | Qa.III | IEKIAGTISHINEISSSIATAVEQQASAAKEIADNVTQASGGAEQINAAL--- | *Cohaesibacter marisflavi* | Bacteria; Proteobacteria; Alphaproteobacteria; Rhizobiales; Cohaesibacteraceae | hypothetical protein | 6 |
|  | **AHZ72111.1** | 1E-05 | Qb.I | LQRTKLHLQDTIEQSEVSSEELKASNEEMQAINEELRSATEELETS------- | *Pseudomonas mandelii JR-1* | Bacteria; Proteobacteria; Gammaproteobacteria; Pseudomonadales; Pseudomonadaceae; Pseudomonas | signal transduction histidine kinase with CheB and CheR activity | 6 |
| | **KZZ70007.1** | 1E-05 | Qa.III | LDEIVAAIANIQDLSATIASATEEQVSVVNEVAQNINSIRELNETSNTIAQKT | *Oleiphilus sp. HI0130* | Bacteria; Proteobacteria; Gammaproteobacteria; Oceanospirillales; Oleiphilaceae; Oleiphilus | hypothetical protein A3765_03320 | 6 |
| | **OKO67271.1** | 1E-05 | Qa.IV | -KEIGATIGQISTISTSIASAVEQQGAATQEIARSVQTVAQGTQAAATDI--- | *Bradyrhizobium sp. NAS80.1* | Bacteria; Proteobacteria; Alphaproteobacteria; Rhizobiales; Bradyrhizobiaceae; Bradyrhizobium | chemotaxis protein partial | 6 |
|  | **OYX88347.1** | 1E-05 | Qa.IV | --KIVTIISDIRQATSAIASAIEEQGAATREIAYNTQSAAQGTQGVND----- | *Azorhizobium sp. 35-67-5* | Bacteria; Proteobacteria; Alphaproteobacteria; Rhizobiales; Xanthobacteraceae; Azorhizobium | hypothetical protein B7Y75_00755 | 6 |
|  | **SPX65612.1** | 1E-05 | R.IV | IQEINRAVTHLDSMVQQNAELVVQSAAAASALQGQAGELAETAGHFR------ | *Leclercia adecarboxylata* | Bacteria; Proteobacteria; Gammaproteobacteria; Enterobacterales; Enterobacteriaceae; Leclercia | Serine chemoreceptor protein | 6 |
|  | **WP_019244199.1** | 1E-05 | Qa.III.b | TKNIQKAIRNLNEAFTGINEYTDNQAAAIQQIAATIQEVSKNAQ--------- | *Bacillus massilioanorexius* | Bacteria; Firmicutes; Bacilli; Bacillales; Bacillaceae; Bacillus | hypothetical protein | 6 |
|  | **WP_021076529.1** | 1E-05 | Qa.IV | IRNISSTIGEINDVTTAIAAAVEEQGAATREIARNIQHAA------------- | *Bradyrhizobium sp. MOS004* | Bacteria; Proteobacteria; Alphaproteobacteria; Rhizobiales; Bradyrhizobiaceae; Bradyrhizobium | HAMP domain-containing protein | 6 |
|  | **WP_021133549.1** | 1E-05 | Qa.IV | -----QTIRTLNELSTAIAGAVEQQSAATAEISRNVEEASANTSR-------- | *Phaeospirillum fulvum* | Bacteria; Proteobacteria; Alphaproteobacteria; Rhodospirillales; Rhodospirillaceae; Phaeospirillum | methyl-accepting chemotaxis protein | 6 |
|  | **WP_024579017.1** | 1E-05 | Qa.IV | IRNISSTIGEINDVTTAIAAAVEEQGAATREIARNIQHAA------------- | *Bradyrhizobium* | Bacteria; Proteobacteria; Alphaproteobacteria; Rhizobiales; Bradyrhizobiaceae | MULTISPECIES: HAMP domain-containing protein | 6 |
|  | **WP_044542967.1** | 1E-05 | Qa.IV | IRNISSTIGEINDVTTAIAAAVEEQGAATREIARNIQHAA------------- | *Bradyrhizobium sp. LTSP885* | Bacteria; Proteobacteria; Alphaproteobacteria; Rhizobiales; Bradyrhizobiaceae; Bradyrhizobium | HAMP domain-containing protein | 6 |
|  | **WP_044589627.1** | 1E-05 | Qa.IV | IRNISSTIGEINDVTTAIAAAVEEQGAATREIARNIQHAA------------- | *Bradyrhizobium sp. LTSPM299* | Bacteria; Proteobacteria; Alphaproteobacteria; Rhizobiales; Bradyrhizobiaceae; Bradyrhizobium | HAMP domain-containing protein | 6 |
|  | **WP_045585159.1** | 1E-05 | Qa.IV | IRSIAVTIREVNEISATIAAAVEQQSAATREIARNVGEAA------------- | *Azospirillum thiophilum* | Bacteria; Proteobacteria; Alphaproteobacteria; Rhodospirillales; Rhodospirillaceae; Azospirillum | HAMP domain-containing protein | 6 |
|  | **WP_050403352.1** | 1E-05 | Qa.IV | IRNISSTIGEINDVTTAIAAAVEEQGAATREIARNIQHAA------------- | *Bradyrhizobium embrapense* | Bacteria; Proteobacteria; Alphaproteobacteria; Rhizobiales; Bradyrhizobiaceae; Bradyrhizobium | HAMP domain-containing protein | 6 |
|  | **WP_050630079.1** | 1E-05 | Qa.IV | IRNISSTIGEINDVTTAIAAAVEEQGAATREIARNIQHAA------------- | *Bradyrhizobium viridifuturi* | Bacteria; Proteobacteria; Alphaproteobacteria; Rhizobiales; Bradyrhizobiaceae; Bradyrhizobium | HAMP domain-containing protein | 6 |
|  | **WP_060845346.1** | 1E-05 | Qc.III | -DGIATTVGTIADIGTSIASAVEEQNAATGEITQMMQTVANRTQT-------- | *Methylobacterium aquaticum* | Bacteria; Proteobacteria; Alphaproteobacteria; Rhizobiales; Methylobacteriaceae; Methylobacterium | HAMP domain-containing protein | 6 |
|  | **WP_062175102.1** | 1E-05 | Qb.I | LRTTRTQLQASLEHADASTEELKASNEELQAINEELRSTTEELETSK------ | *Burkholderia sp. PAMC 26561* | Bacteria; Proteobacteria; Betaproteobacteria; Burkholderiales; Burkholderiaceae; Burkholderia | PAS domain S-box protein | 6 |
|  | **WP_066513345.1** | 1E-05 | Qa.IV | IRNISSTIGEINDVTTAIAAAVEEQGAATREIARNIQHAA------------- | *Bradyrhizobium sp. BR 10303* | Bacteria; Proteobacteria; Alphaproteobacteria; Rhizobiales; Bradyrhizobiaceae; Bradyrhizobium | methyl-accepting chemotaxis protein | 6 |
| | **WP_068452686.1** | 1E-05 | Qa.III | LDEIVAAIANIQDLSATIASATEEQVSVVNEVAQNINSIRELNETSNTIAQKT | *Oleiphilus* | Bacteria; Proteobacteria; Gammaproteobacteria; Oceanospirillales; Oleiphilaceae | MULTISPECIES: methyl-accepting chemotaxis protein | 6 |
| | **WP_072817535.1** | 1E-05 | Qc.III | -KAIGGIIGEVNEVATAIAAAVEEQGAATQEISRNTQYAAQGTRNVSENIA-- | *Bradyrhizobium erythrophlei* | Bacteria; Proteobacteria; Alphaproteobacteria; Rhizobiales; Bradyrhizobiaceae; Bradyrhizobium | methyl-accepting chemotaxis protein | 6 |
|  | **WP_074129623.1** | 1E-05 | Qa.IV | IRNISSTIGEINDVTTAIAAAVEEQGAATREIARNIQHAA------------- | *Bradyrhizobium sp. NAS96.2* | Bacteria; Proteobacteria; Alphaproteobacteria; Rhizobiales; Bradyrhizobiaceae; Bradyrhizobium | HAMP domain-containing protein | 6 |
|  | **WP_076864336.1** | 1E-05 | Qa.IV | IRNISSTIGEINDVTTAIAAAVEEQGAATREIARNIQHAA------------- | *Bradyrhizobium mercantei* | Bacteria; Proteobacteria; Alphaproteobacteria; Rhizobiales; Bradyrhizobiaceae; Bradyrhizobium | HAMP domain-containing protein | 6 |
|  | **WP_083724179.1** | 1E-05 | Qa.III | LEQIAQSMQQLVALNASIASATTQQSTVVEDVNRNVTEAAEL----------- | *Pseudomonas pachastrellae* | Bacteria; Proteobacteria; Gammaproteobacteria; Pseudomonadales; Pseudomonadaceae; Pseudomonas | methyl-accepting chemotaxis protein | 6 |
|  | **WP_084720479.1** | 1E-05 | Qa.IV | IKSVLSGVSDMNEICTLIAAAVEEQAAATREIARNVDQAA------------- | *Skermanella aerolata* | Bacteria; Proteobacteria; Alphaproteobacteria; Rhodospirillales; Rhodospirillaceae; Skermanella | hypothetical protein | 6 |
|  | **WP_088274026.1** | 1E-05 | Qa.III | LEQIAQSMQQLVALNASIASATTQQSTVVEDVNRNVTEAAEL----------- | *Pseudomonas aestusnigri* | Bacteria; Proteobacteria; Gammaproteobacteria; Pseudomonadales; Pseudomonadaceae; Pseudomonas | methyl-accepting chemotaxis protein | 6 |
|  | **WP_091750550.1** | 1E-05 | Qc.III | -DGIATTVGTIADIGTSIASAVEEQNAATGEITQMMQTVANRTQT-------- | *Methylobacterium sp. ap11* | Bacteria; Proteobacteria; Alphaproteobacteria; Rhizobiales; Methylobacteriaceae; Methylobacterium | HAMP domain-containing protein | 6 |
|  | **WP_091882986.1** | 1E-05 | Qc.III | -DGIATTVGTIADIGTSIASAVEEQNAATGEITQMMQTVANRTQT-------- | *Methylobacterium sp. yr596* | Bacteria; Proteobacteria; Alphaproteobacteria; Rhizobiales; Methylobacteriaceae; Methylobacterium | HAMP domain-containing protein | 6 |
|  | **WP_092123108.1** | 1E-05 | Qa.IV | IRNISSTIGEINDVTTAIAAAVEEQGAATREIARNIQHAA------------- | *Bradyrhizobium erythrophlei* | Bacteria; Proteobacteria; Alphaproteobacteria; Rhizobiales; Bradyrhizobiaceae; Bradyrhizobium | HAMP domain-containing protein | 6 |
|  | **WP_095731117.1** | 1E-05 | Qa.IV | IRNISSTIGEINDVTTAIAAAVEEQGAATREIARNIQHAA------------- | *Bradyrhizobium sp. UFLA03-84* | Bacteria; Proteobacteria; Alphaproteobacteria; Rhizobiales; Bradyrhizobiaceae; Bradyrhizobium | HAMP domain-containing protein | 6 |
|  | **WP_096173630.1** | 1E-05 | Qa.III | INEIRQVIESVNQIAQSIAAAVEEQSAVTRDISDNMQQAS------------- | *Cohaesibacter sp. ES.047* | Bacteria; Proteobacteria; Alphaproteobacteria; Rhizobiales; Cohaesibacteraceae | methyl-accepting chemotaxis protein | 6 |
|  | **WP_097674075.1** | 1E-05 | Qa.IV | IRNISSTIGEINDVTTAIAAAVEEQGAATREIARNIQHAA------------- | *Bradyrhizobium sp. C9* | Bacteria; Proteobacteria; Alphaproteobacteria; Rhizobiales; Bradyrhizobiaceae; Bradyrhizobium | HAMP domain-containing protein | 6 |
|  | **WP_100294045.1** | 1E-05 | Qc.III | LEQIAESINQISDMATQISSAAEEQRAVTSEVSRNIQAVNDVSDE-------- | *Aeromonas cavernicola* | Bacteria; Proteobacteria; Gammaproteobacteria; Aeromonadales; Aeromonadaceae; Aeromonas | methyl-accepting chemotaxis protein | 6 |
|  | **WP_100951841.1** | 1E-05 | Qa.IV | IRNISSTIGEINDVTTAIAAAVEEQGAATREIARNIQHAA------------- | *Bradyrhizobium sp. SK17* | Bacteria; Proteobacteria; Alphaproteobacteria; Rhizobiales; Bradyrhizobiaceae; Bradyrhizobium | HAMP domain-containing protein | 6 |
|  | **WP_101287388.1** | 1E-05 | Qa.IV | IRAIAATMEEVNRATLGIASAVEQQEASTNEISRNVSEASGNTV--------- | *Pleomorphomonas diazotrophica* | Bacteria; Proteobacteria; Alphaproteobacteria; Rhizobiales; Methylocystaceae; Pleomorphomonas | methyl-accepting chemotaxis protein | 6 |
|  | **WP_106324086.1** | 1E-05 | Qa.IV | IRNISSTIGEINDVTTAIAAAVEEQGAATREIARNIQHAA------------- | *Rhizobium tropici* | Bacteria; Proteobacteria; Alphaproteobacteria; Rhizobiales; Rhizobiaceae; Rhizobium/Agrobacterium group; Rhizobium | HAMP domain-containing protein | 6 |

| | | | | | | | | |
|---|---|---|---|---|---|---|---|---|
|  | **ACJ00923.1** | 1.1E-05 | Qa.IV | IITISRTISEINEISDQVSHSVEEQDAATKEIARNVQQAS------------- | *Rhodospirillum centenum SW* | Bacteria; Proteobacteria; Alphaproteobacteria; Rhodospirillales; Rhodospirillaceae; Rhodospirillum | methyl-accepting chemotaxis protein putative | 6 |
|  | **ANJ54693.1** | 1.1E-05 | Qb.I | LQRTKLHLQDTIEQSEVSSEELKASNEEMQAINEELRSATEELETS------- | *Pseudomonas silesiensis* | Bacteria; Proteobacteria; Gammaproteobacteria; Pseudomonadales; Pseudomonadaceae; Pseudomonas | chemotaxis protein | 6 |
|  | **EJW13570.1** | 1.1E-05 | Qc.III | IKEIGGTIGRISEIAAAIAAAVEQQGAATSEVARNIQHASQGTAR-------- | *Rhodovulum sp. PH10* | Bacteria; Proteobacteria; Alphaproteobacteria; Rhodobacterales; Rhodobacteraceae; Rhodovulum | putative methyl-accepting chemotaxis receptor/sensory transducer | 6 |
|  | **KPW11271.1** | 1.1E-05 | Qa.III | ---ISQALRNLNSLNTSIASATLQQTHVVDDINQNVTQAAQLSQ--------- | *Pseudomonas amygdali pv. aesculi* | Bacteria; Proteobacteria; Gammaproteobacteria; Pseudomonadales; Pseudomonadaceae; Pseudomonas; Pseudomonas amygdali | Methyl-accepting chemotaxis protein | 6 |
| | **KZD24784.1** | 1.1E-05 | Qc.III | IKEIGDTIGRMSEIASTIASAVEEQGAATQEISRNVQQAAQGTMQVSSNIT-- | *Tardiphaga robiniae* | Bacteria; Proteobacteria; Alphaproteobacteria; Rhizobiales; Bradyrhizobiaceae; Tardiphaga | chemotaxis protein | 6 |
| | **OKO67282.1** | 1.1E-05 | Qa.IV | -KEIGATIGQISTISTSIASAVEQQGAATQEIARSVQTVAQGTQAAATDI--- | *Bradyrhizobium sp. NAS80.1* | Bacteria; Proteobacteria; Alphaproteobacteria; Rhizobiales; Bradyrhizobiaceae; Bradyrhizobium | chemotaxis protein partial | 6 |
|  | **OPK11111.1** | 1.1E-05 | Qb.I | LQRTKLHLQDTIEQSEVSSEELKASNEEMQAINEELRSATEELETS------- | *Pseudomonas sp. VI4.1* | Bacteria; Proteobacteria; Gammaproteobacteria; Pseudomonadales; Pseudomonadaceae; Pseudomonas | chemotaxis protein | 6 |
| | **OYU87016.1** | 1.1E-05 | Qc.III | IKEIGDTIGRMSEIASTIASAVEEQGAATQEISRNVQQAAQGTMQVSSNIT-- | *Bradyrhizobiaceae bacterium PARB1* | Bacteria; Proteobacteria; Alphaproteobacteria; Rhizobiales; Bradyrhizobiaceae | methyl-accepting chemotaxis protein | 6 |
| | **OYU90114.1** | 1.1E-05 | Qc.III | IKEIGDTIGRMSEIASTIASAVEEQGAATQEISRNVQQAAQGTMQVSSNIT-- | *Bradyrhizobiaceae bacterium PARB1* | Bacteria; Proteobacteria; Alphaproteobacteria; Rhizobiales; Bradyrhizobiaceae | methyl-accepting chemotaxis protein | 6 |
|  | **PIR01551.1** | 1.1E-05 | Qc.III | IDEISQIINEINDISNTIASAVEEQTATTSEMGRNVTQAARGSN--------- | *Nitrospinae bacterium CG11_big_fil_rev_8_21_14_0_20_45_15* | Bacteria; Nitrospinae/Tectomicrobia group; Nitrospinae | hypothetical protein COV66_00975 | 6 |
| | **PVX48603.1** | 1.1E-05 | Qc.III | IKEIGDTIGRMSEIASTIASAVEEQGAATQEISRNVQQAAQGTMQVSSNIT-- | *Tardiphaga sp. OV697* | Bacteria; Proteobacteria; Alphaproteobacteria; Rhizobiales; Bradyrhizobiaceae; Tardiphaga | methyl-accepting chemotaxis protein | 6 |
| | **PVX53825.1** | 1.1E-05 | Qc.III | IKEIGDTIGRMSEIASTIASAVEEQGAATQEISRNVQQAAQGTMQVSSNIT-- | *Tardiphaga sp. OV697* | Bacteria; Proteobacteria; Alphaproteobacteria; Rhizobiales; Bradyrhizobiaceae; Tardiphaga | methyl-accepting chemotaxis sensory transducer | 6 |
|  | **PWC35792.1** | 1.1E-05 | Qa.III | IEDIGVVIGQVSGIADDIAAAVEQQGEATREIARNVQQAA------------- | *Azospirillum sp. TSO35-2* | Bacteria; Proteobacteria; Alphaproteobacteria; Rhodospirillales; Rhodospirillaceae; Azospirillum | hypothetical protein TSO352_11195 | 6 |
| | **PWC39491.1** | 1.1E-05 | Qa.III | IKDVARVITRINEVATSISSAVEQQGAATREISRNVQQAAQGTQETSRSIVG- | *Azospirillum sp. TSO35-2* | Bacteria; Proteobacteria; Alphaproteobacteria; Rhodospirillales; Rhodospirillaceae; Azospirillum | hypothetical protein TSO352_04920 | 6 |
|  | **PWC87378.1** | 1.1E-05 | Qa.III | IVGIGRTIQSINQIASGISAGIEQQGSAVREIVVNMEQAA------------- | *Azospirillum sp. TSO5* | Bacteria; Proteobacteria; Alphaproteobacteria; Rhodospirillales; Rhodospirillaceae; Azospirillum | chemotaxis protein | 6 |
|  | **SEQ16439.1** | 1.1E-05 | SNAP.c | -GDVSDVINEMNQIASAVAASVEQQNAAIQAITENVTNATQ------------ | *Pseudovibrio axinellae* | Bacteria; Proteobacteria; Alphaproteobacteria; Rhodobacterales; Rhodobacteraceae; Pseudovibrio | methyl-accepting chemotaxis protein | 6 |
| | **SOD99820.1** | 1.1E-05 | Qc.III | --RIRGTIQELDEIAQSIAAAVEQQNAATMDISRGVQRAADGVREAARQVD-- | *Caenispirillum bisanense* | Bacteria; Proteobacteria; Alphaproteobacteria; Rhodospirillales; Rhodospirillaceae; Caenispirillum | methyl-accepting chemotaxis protein | 6 |
|  | **WP_008033069.1** | 1.1E-05 | Qb.I | LQRTKLHLQDTIEQSEVSSEELKASNEEMQAINEELRSATEELETS------- | *Pseudomonas sp. GM60* | Bacteria; Proteobacteria; Gammaproteobacteria; Pseudomonadales; Pseudomonadaceae; Pseudomonas | PAS domain S-box protein | 6 |
|  | **WP_008038448.1** | 1.1E-05 | Qb.I | LQRTKLHLQDTIEQSEVSSEELKASNEEMQAINEELRSATEELETS------- | *Pseudomonas sp. GM67* | Bacteria; Proteobacteria; Gammaproteobacteria; Pseudomonadales; Pseudomonadaceae; Pseudomonas | PAS domain S-box protein | 6 |
|  | **WP_009027584.1** | 1.1E-05 | Qc.III | -KAIGSTIERINEIAASISAAVEQQRAATQNITQSVRSAASG----------- | *Bradyrhizobium sp. ORS 375* | Bacteria; Proteobacteria; Alphaproteobacteria; Rhizobiales; Bradyrhizobiaceae; Bradyrhizobium | HAMP domain-containing protein | 6 |
| | **WP_009031103.1** | 1.1E-05 | Qc.III | IKEISGTIERLSEISSTIAAAVEEQGAATQEISRNIQQAAQGTHEVSSNITD- | *Bradyrhizobium sp. ORS 375* | Bacteria; Proteobacteria; Alphaproteobacteria; Rhizobiales; Bradyrhizobiaceae; Bradyrhizobium | methyl-accepting chemotaxis protein | 6 |
|  | **WP_010457177.1** | 1.1E-05 | Qb.I | LQRTKLHLQDTIEQSEVSSEELKASNEEMQAINEELRSATEELETS------- | *Pseudomonas mandelii* | Bacteria; Proteobacteria; Gammaproteobacteria; Pseudomonadales; Pseudomonadaceae; Pseudomonas | PAS domain S-box protein | 6 |
|  | **WP_016773601.1** | 1.1E-05 | Qb.I | LQRTKLHLQDTIEQSEISSEELKASNEEMQAINEELRSATEELETS------- | *Pseudomonas sp. R62* | Bacteria; Proteobacteria; Gammaproteobacteria; Pseudomonadales; Pseudomonadaceae; Pseudomonas | response regulator | 6 |
|  | **WP_017340209.1** | 1.1E-05 | Qb.I | LQRTKLHLQDTIEQSEVSSEELKASNEEMQAINEELRSATEELETS------- | *Pseudomonas fluorescens* | Bacteria; Proteobacteria; Gammaproteobacteria; Pseudomonadales; Pseudomonadaceae; Pseudomonas | PAS domain S-box protein | 6 |
|  | **WP_018926330.1** | 1.1E-05 | Qb.I | LQRTKLHLQDTIEQSEISSEELKASNEEMQAINEELRSATEELETS------- | *Pseudomonas* | Bacteria; Proteobacteria; Gammaproteobacteria; Pseudomonadales; Pseudomonadaceae | MULTISPECIES: PAS domain S-box protein | 6 |
|  | **WP_019582898.1** | 1.1E-05 | Qb.I | LQRTKLHLQDTIEQSEISSEELKASNEEMQAINEELRSATEELETS------- | *Pseudomonas* | Bacteria; Proteobacteria; Gammaproteobacteria; Pseudomonadales; Pseudomonadaceae | MULTISPECIES: PAS domain S-box protein | 6 |
|  | **WP_019650358.1** | 1.1E-05 | Qb.I | LQRTKLHLQDTIEQSEISSEELKASNEEMQAINEELRSATEELETS------- | *Pseudomonas sp. 45MFCol3.1* | Bacteria; Proteobacteria; Gammaproteobacteria; Pseudomonadales; Pseudomonadaceae; Pseudomonas | PAS domain S-box protein | 6 |
|  | **WP_024013991.1** | 1.1E-05 | Qb.I | LQRTKLHLQDTIEQSEISSEELKASNEEMQAINEELRSATEELETS------- | *Pseudomonas moraviensis* | Bacteria; Proteobacteria; Gammaproteobacteria; Pseudomonadales; Pseudomonadaceae; Pseudomonas | response regulator | 6 |
| | **WP_028135260.1** | 1.1E-05 | Qc.III | IGEISGTIGRLSEIASAIAAAVQQQGAATQEIARNVQQAAQGTQQVSSNVG-- | *Bradyrhizobium japonicum* | Bacteria; Proteobacteria; Alphaproteobacteria; Rhizobiales; Bradyrhizobiaceae; Bradyrhizobium | PAS domain S-box protein | 6 |
|  | **WP_028145863.1** | 1.1E-05 | Qa.IV | IKEIGATIARISDISTTIAATIEQQGAATAEIARNVSEAA------------- | *Bradyrhizobium* | Bacteria; Proteobacteria; Alphaproteobacteria; Rhizobiales; Bradyrhizobiaceae | MULTISPECIES: chemotaxis protein | 6 |
|  | **WP_039757865.1** | 1.1E-05 | Qb.I | LQRTKLHLQDTIEQSEISSEELKASNEEMQAINEELRSATEELETS------- | *Pseudomonas fluorescens* | Bacteria; Proteobacteria; Gammaproteobacteria; Pseudomonadales; Pseudomonadaceae; Pseudomonas | PAS domain S-box protein | 6 |
|  | **WP_041756538.1** | 1.1E-05 | Qc.III | -KAIGSTIERINEIAASISAAVEQQRAATQNITQSVRSAASG----------- | *Bradyrhizobium sp. ORS 278* | Bacteria; Proteobacteria; Alphaproteobacteria; Rhizobiales; Bradyrhizobiaceae; Bradyrhizobium | HAMP domain-containing protein | 6 |
| | **WP_045013238.1** | 1.1E-05 | Qc.III | IKEISGTIGRISEISSTIAAAVEEQGAATQEISRNVQQAAQGTAQVSSNI--- | *Bradyrhizobium sp. LTSP849* | Bacteria; Proteobacteria; Alphaproteobacteria; Rhizobiales; Bradyrhizobiaceae; Bradyrhizobium | methyl-accepting chemotaxis protein | 6 |
| | **WP_046510234.1** | 1.1E-05 | Qc.III | --GISKTIDSIHEVATAIASAVEEQGAATAEITKSVQQAANGTDQVSSNIT-- | *Kiloniella litopenaei* | Bacteria; Proteobacteria; Alphaproteobacteria; Kiloniellales; Kiloniellaceae; Kiloniella | HAMP domain-containing protein partial | 6 |
|  | **WP_051644849.1** | 1.1E-05 | Qa.III | ------AISDLNEIAASIAAAVEEQSAATSEIAGNVQQAAVASQEVSDSI--- | *Labrenzia sp. DG1229* | Bacteria; Proteobacteria; Alphaproteobacteria; Rhodobacterales; Rhodobacteraceae; Labrenzia | methyl-accepting chemotaxis protein | 6 |
|  | **WP_054300313.1** | 1.1E-05 | Qa.III | -EEIEIAVKKINEMSTDISNAVQEQNTVLKEMNDKTSQIRAASELGD------ | *Piscirickettsia salmonis* | Bacteria; Proteobacteria; Gammaproteobacteria; Thiotrichales; Piscirickettsiaceae; Piscirickettsia | methyl-accepting chemotaxis protein | 6 |
|  | **WP_055110953.1** | 1.1E-05 | Qa.III | ------AISDLNEIAASIAAAVEEQSAATSEIAGNVQQAAVASQEVSDSI--- | *Labrenzia* | Bacteria; Proteobacteria; Alphaproteobacteria; Rhodobacterales; Rhodobacteraceae | MULTISPECIES: methyl-accepting chemotaxis protein | 6 |
|  | **WP_061272465.1** | 1.1E-05 | R.IV | -QEINHAVTHLDNMVQQNAELVVESAAAAGALQSQAGDLAETAGHFR------ | *Cedecea neteri* | Bacteria; Proteobacteria; Gammaproteobacteria; Enterobacterales; Enterobacteriaceae; Cedecea | methyl-accepting chemotaxis protein | 6 |
|  | **WP_064616472.1** | 1.1E-05 | Qb.I | LQRTKLHLQDTIEQSEISSEELKASNEEMQAINEELRSATEELETS------- | *Pseudomonas sp. GR 6-02* | Bacteria; Proteobacteria; Gammaproteobacteria; Pseudomonadales; Pseudomonadaceae; Pseudomonas | response regulator | 6 |
| | **WP_068728912.1** | 1.1E-05 | Qc.III | IKEIGDTIGRMSEIASTIASAVEEQGAATQEISRNVQQAAQGTMQVSSNIT-- | *Tardiphaga robiniae* | Bacteria; Proteobacteria; Alphaproteobacteria; Rhizobiales; Bradyrhizobiaceae; Tardiphaga | HAMP domain-containing protein | 6 |
|  | **WP_070425341.1** | 1.1E-05 | Qa.III | MDEIVVSVKRVSDLMADISTAAQQQSEGIEQINQVVEHIDESTQ--------- | *Stenotrophomonas sp. LM091* | Bacteria; Proteobacteria; Gammaproteobacteria; Xanthomonadales; Xanthomonadaceae; Stenotrophomonas | PAS domain S-box protein | 6 |
|  | **WP_070471844.1** | 1.1E-05 | Qa.III | MDEIVVSVKRVSDLMADISTAAQQQSEGIEQINQVVEHIDESTQ--------- | *Stenotrophomonas sp. HMSC10F06* | Bacteria; Proteobacteria; Gammaproteobacteria; Xanthomonadales; Xanthomonadaceae; Stenotrophomonas | PAS domain S-box protein | 6 |
|  | **WP_072819429.1** | 1.1E-05 | Qa.IV | ----------INQITTAIASSVEQQGAATQEIARNIQEAAQGTEKVTQNI--- | *Bradyrhizobium erythrophlei* | Bacteria; Proteobacteria; Alphaproteobacteria; Rhizobiales; Bradyrhizobiaceae; Bradyrhizobium | methyl-accepting chemotaxis protein | 6 |
|  | **WP_074444747.1** | 1.1E-05 | Qa.IV | IQQITERMREINSATVTIASAMTQQGAATSEISRNVQHAAK------------ | *Salinarimonadaceae bacterium HL-109* | Bacteria; Proteobacteria; Alphaproteobacteria; Rhizobiales; Salinarimonadaceae | methyl-accepting chemotaxis protein | 6 |
|  | **WP_075219407.1** | 1.1E-05 | Qa.IV | -------ITDIRETFDQVAAAVHEQQSATRSISSSAQDAANGVSNAATTL--- | *Acuticoccus yangtzensis* | Bacteria; Proteobacteria; Alphaproteobacteria; Rhodobacterales; Rhodobacteraceae; Acuticoccus | hypothetical protein | 6 |
|  | **WP_075274652.1** | 1.1E-05 | Qa.III | -EEIEIAVKKINEMSTDISNAVQEQNTVLKEMNDKTSQIRAASELGD------ | *Piscirickettsia salmonis* | Bacteria; Proteobacteria; Gammaproteobacteria; Thiotrichales; Piscirickettsiaceae; Piscirickettsia | methyl-accepting chemotaxis protein | 6 |
|  | **WP_076563865.1** | 1.1E-05 | Qb.I | LQRTKLHLQDTIEQSEISSEELKASNEEMQAINEELRSATEELETS------- | *Pseudomonas sp. B10* | Bacteria; Proteobacteria; Gammaproteobacteria; Pseudomonadales; Pseudomonadaceae; Pseudomonas | response regulator | 6 |
|  | **WP_077749186.1** | 1.1E-05 | Qb.I | LQRTKLHLQDTIEQSEVSSEELKASNEEMQAINEELRSATEELETS------- | *Pseudomonas sp. FSL W5-0299* | Bacteria; Proteobacteria; Gammaproteobacteria; Pseudomonadales; Pseudomonadaceae; Pseudomonas | PAS domain S-box protein | 6 |
|  | **WP_079538073.1** | 1.1E-05 | Qa.IV | IKEIGATIARISEISATIASAVEEQGAATQEISRNVQRAAE------------ | *Bradyrhizobium lablabi* | Bacteria; Proteobacteria; Alphaproteobacteria; Rhizobiales; Bradyrhizobiaceae; Bradyrhizobium | HAMP domain-containing protein | 6 |
|  | **WP_082012933.1** | 1.1E-05 | Qa.III | IQGIASTIGEVNQIAGAIAAAVEEQGSATQEIARNVQQAA------------- | *Belnapia sp. F-4-1* | Bacteria; Proteobacteria; Alphaproteobacteria; Rhodospirillales; Acetobacteraceae; Belnapia | methyl-accepting chemotaxis protein | 6 |
|  | **WP_082012981.1** | 1.1E-05 | Qa.III | IQGIASTIGEVNQIAGAIAAAVEEQGSATQEIARNVQQAA------------- | *Belnapia sp. F-4-1* | Bacteria; Proteobacteria; Alphaproteobacteria; Rhodospirillales; Acetobacteraceae; Belnapia | methyl-accepting chemotaxis protein | 6 |
|  | **WP_082700526.1** | 1.1E-05 | Qa.IV | ------TIREINGISTAIASAVEEQSAATSEISRNVHEASKAVNDVNKNI--- | *Magnetospirillum sp. XM-1* | Bacteria; Proteobacteria; Alphaproteobacteria; Rhodospirillales; Rhodospirillaceae; Magnetospirillum | HAMP domain-containing protein | 6 |
|  | **WP_083375937.1** | 1.1E-05 | Qb.I | LQRTKLHLQDTIEQSEVSSEELKASNEEMQAINEELRSATEELETS------- | *Pseudomonas mandelii* | Bacteria; Proteobacteria; Gammaproteobacteria; Pseudomonadales; Pseudomonadaceae; Pseudomonas | PAS domain S-box protein | 6 |
|  | **WP_083850241.1** | 1.1E-05 | Qc.III | IKEIGGTIGRISEIAAAIAAAVEQQGAATSEVARNIQHASQGTAR-------- | *Rhodovulum sp. PH10* | Bacteria; Proteobacteria; Alphaproteobacteria; Rhodobacterales; Rhodobacteraceae; Rhodovulum | methyl-accepting chemotaxis protein | 6 |
|  | **WP_084321954.1** | 1.1E-05 | Qb.I | LQRTKLHLQDTIEQSEVSSEELKASNEEMQAINEELRSATEELETS------- | *Pseudomonas migulae* | Bacteria; Proteobacteria; Gammaproteobacteria; Pseudomonadales; Pseudomonadaceae; Pseudomonas | PAS domain S-box protein | 6 |
| | **WP_085375383.1** | 1.1E-05 | Qc.III.c | ---IGGTIREINGISTAIASAVEEQSAATSEISRNVHEASKAVNDVNKNIA-- | *Magnetospirillum sp. ME-1* | Bacteria; Proteobacteria; Alphaproteobacteria; Rhodospirillales; Rhodospirillaceae; Magnetospirillum | HAMP domain-containing protein | 6 |
|  | **WP_085901720.1** | 1.1E-05 | Qa.III.b | -EQIDNVIKEINDIAGGISTAVEEQGGATREISQTVNQSANN----------- | *Kiloniella majae* | Bacteria; Proteobacteria; Alphaproteobacteria; Kiloniellales; Kiloniellaceae; Kiloniella | HAMP domain-containing protein | 6 |
|  | **WP_086944555.1** | 1.1E-05 | Qb.I | LQRTKLHLQDTIEQSEISSEELKASNEEMQAINEELRSATEELETS------- | *Pseudomonas* | Bacteria; Proteobacteria; Gammaproteobacteria; Pseudomonadales; Pseudomonadaceae | MULTISPECIES: response regulator | 6 |
| | **WP_089265831.1** | 1.1E-05 | Qc.III | IKEIGDTIGRMSEIASTIASAVEEQGAATQEISRNVQQAAQGTMQVSSNIT-- | *Tardiphaga sp. OK246* | Bacteria; Proteobacteria; Alphaproteobacteria; Rhizobiales; Bradyrhizobiaceae; Tardiphaga | HAMP domain-containing protein | 6 |
| | **WP_089267187.1** | 1.1E-05 | Qc.III | IKEIGDTIGRMSEIASTIASAVEEQGAATQEISRNVQQAAQGTMQVSSNIT-- | *Tardiphaga sp. OK246* | Bacteria; Proteobacteria; Alphaproteobacteria; Rhizobiales; Bradyrhizobiaceae; Tardiphaga | methyl-accepting chemotaxis protein | 6 |

| | | | | | | | | |
|---|---|---|---|---|---|---|---|---|
| | **WP_092145860.1** | 1.1E-05 | Qc.III | IKEIGDTIGRMSEIASTIASAVEEQGAATQEISRNVQQAAQGTMQVSSNIT-- | *Bradyrhizobium sp. NFR13* | Bacteria; Proteobacteria; Alphaproteobacteria; Rhizobiales; Bradyrhizobiaceae; Bradyrhizobium | methyl-accepting chemotaxis protein | 6 |
| | **WP_092151354.1** | 1.1E-05 | Qc.III | IKEIGDTIGRMSEIASTIASAVEEQGAATQEISRNVQQAAQGTMQVSSNIT-- | *Bradyrhizobium sp. NFR13* | Bacteria; Proteobacteria; Alphaproteobacteria; Rhizobiales; Bradyrhizobiaceae; Bradyrhizobium | methyl-accepting chemotaxis protein | 6 |
|  | **WP_092616373.1** | 1.1E-05 | Qa.IV | IQAIAKVVAEINEITSTISAATEEQTATTQEISRNAQHAA------------- | *Roseospirillum parvum* | Bacteria; Proteobacteria; Alphaproteobacteria; Rhizobiales; Rhodobiaceae; Roseospirillum | HAMP domain-containing protein | 6 |
| | **WP_093757184.1** | 1.1E-05 | Qc.III | IKEIGDTIGRMSEIASTIASAVEEQGAATQEISRNVQQAAQGTMQVSSNIT-- | *Tardiphaga sp. OK245* | Bacteria; Proteobacteria; Alphaproteobacteria; Rhizobiales; Bradyrhizobiaceae; Tardiphaga | methyl-accepting chemotaxis protein | 6 |
| | **WP_093758888.1** | 1.1E-05 | Qc.III | IKEIGDTIGRMSEIASTIASAVEEQGAATQEISRNVQQAAQGTMQVSSNIT-- | *Tardiphaga sp. OK245* | Bacteria; Proteobacteria; Alphaproteobacteria; Rhizobiales; Bradyrhizobiaceae; Tardiphaga | methyl-accepting chemotaxis protein | 6 |
|  | **WP_094471120.1** | 1.1E-05 | Qb.I | LQRTKLHLQDTIEQSEVSSEELKASNEEMQAINEELRSATEELETS------- | *Pseudomonas mandelii* | Bacteria; Proteobacteria; Gammaproteobacteria; Pseudomonadales; Pseudomonadaceae; Pseudomonas | PAS domain S-box protein | 6 |
|  | **WP_094958585.1** | 1.1E-05 | Qa.III | LDQISSSVALISDMATQIATAAEEQTHVTDEITQNTTTIK------------- | *Photobacterium sanguinicancri* | Bacteria; Proteobacteria; Gammaproteobacteria; Vibrionales; Vibrionaceae; Photobacterium | methyl-accepting chemotaxis protein | 6 |
|  | **WP_095634347.1** | 1.1E-05 | Qb.I | LQRTKLHLQDTIEQSEISSEELKASNEEMQAINEELRSATEELETS------- | *Pseudomonas sp. PICF141* | Bacteria; Proteobacteria; Gammaproteobacteria; Pseudomonadales; Pseudomonadaceae; Pseudomonas | PAS domain S-box protein | 6 |
|  | **WP_096174255.1** | 1.1E-05 | Qc.III.c | -DEISATIERVSEISAAIAAAVEEQSSATREIANNVEEAAN------------ | *Cohaesibacter sp. ES.047* | Bacteria; Proteobacteria; Alphaproteobacteria; Rhizobiales; Cohaesibacteraceae | methyl-accepting chemotaxis protein | 6 |
|  | **WP_096480929.1** | 1.1E-05 | Qb.I | LQRTKLHLQDTIEQSEISSEELKASNEEMQAINEELRSATEELETS------- | *Pseudomonas frederiksbergensis* | Bacteria; Proteobacteria; Gammaproteobacteria; Pseudomonadales; Pseudomonadaceae; Pseudomonas | response regulator | 6 |
|  | **WP_096702157.1** | 1.1E-05 | Qa.IV | ------TIREINGISTAIASAVEEQSAATSEISRNVHEASKAVNDVNKNI--- | *Magnetospirillum sp. 15-1* | Bacteria; Proteobacteria; Alphaproteobacteria; Rhodospirillales; Rhodospirillaceae; Magnetospirillum | HAMP domain-containing protein | 6 |
|  | **WP_101063546.1** | 1.1E-05 | Qa.III | LEEINKSVVLISDMATQIATAAEEQTQVTGEISQNVIAIK------------- | *Moritella sp. Urea-trap-13* | Bacteria; Proteobacteria; Gammaproteobacteria; Alteromonadales; Moritellaceae; Moritella | methyl-accepting chemotaxis protein | 6 |
|  | **WP_101156028.1** | 1.1E-05 | Qb.I | LQRTKLHLQDTIEQSEISSEELKASNEEMQAINEELRSATEELETS------- | *Pseudomonas sp. Choline-02u-1* | Bacteria; Proteobacteria; Gammaproteobacteria; Pseudomonadales; Pseudomonadaceae; Pseudomonas | response regulator | 6 |
|  | **WP_102595131.1** | 1.1E-05 | Qb.I | LQRTKLHLQDTIEQSEVSSEELKASNEEMQAINEELRSATEELETS------- | *Pseudomonas* | Bacteria; Proteobacteria; Gammaproteobacteria; Pseudomonadales; Pseudomonadaceae | MULTISPECIES: PAS domain S-box protein | 6 |
|  | **WP_102789470.1** | 1.1E-05 | Qa.III | MDEIVVSVKRVSDLMADISTAAQQQSEGIEQINQVVEHIDESTQ--------- | *Stenotrophomonas bentonitica* | Bacteria; Proteobacteria; Gammaproteobacteria; Xanthomonadales; Xanthomonadaceae; Stenotrophomonas | PAS domain S-box protein | 6 |
|  | **WP_104701247.1** | 1.1E-05 | Qa.III | ------AISDLNEIAASIAAAVEEQSAATSEIAGNVQQAAVASQEVSDSI--- | *Labrenzia sp. EL143* | Bacteria; Proteobacteria; Alphaproteobacteria; Rhodobacterales; Rhodobacteraceae; Labrenzia | methyl-accepting chemotaxis protein | 6 |
|  | **WP_105346329.1** | 1.1E-05 | Qb.I | LQRTKLHLQDTIEQSEISSEELKASNEEMQAINEELRSATEELETS------- | *Pseudomonas frederiksbergensis* | Bacteria; Proteobacteria; Gammaproteobacteria; Pseudomonadales; Pseudomonadaceae; Pseudomonas | PAS domain S-box protein | 6 |
|  | **WP_109107465.1** | 1.1E-05 | Qa.III | IEDIGVVIGQVSGIADDIAAAVEQQGEATREIARNVQQAA------------- | *Azospirillum sp. TSO35-2* | Bacteria; Proteobacteria; Alphaproteobacteria; Rhodospirillales; Rhodospirillaceae; Azospirillum | methyl-accepting chemotaxis protein | 6 |
| | **WP_109108900.1** | 1.1E-05 | Qc.III | -GSIGQTIGRMSEIATAIAAAIEEQAAATRDIAASVTQAAQGTGEVSSNI--- | *Azospirillum sp. TSO35-2* | Bacteria; Proteobacteria; Alphaproteobacteria; Rhodospirillales; Rhodospirillaceae; Azospirillum | methyl-accepting chemotaxis protein | 6 |
|  | **WP_109156027.1** | 1.1E-05 | Qa.III | IVGIGRTIQSINQIASGISAGIEQQGSAVREIVVNMEQAA------------- | *Azospirillum* | Bacteria; Proteobacteria; Alphaproteobacteria; Rhodospirillales; Rhodospirillaceae | MULTISPECIES: chemotaxis protein | 6 |
| | **WP_111420144.1** | 1.1E-05 | Qc.III | IKEIGGTIGRVAEIASTIAAAVEEQGAATGEIARNVQQASAGTTEVARNIAD- | *Rhodoplanes roseus* | Bacteria; Proteobacteria; Alphaproteobacteria; Rhizobiales; Hyphomicrobiaceae; Rhodoplanes | methyl-accepting chemotaxis protein | 6 |
|  | **WP_111455181.1** | 1.1E-05 | Qb.I | LQRTKLHLQDTIEQSEISSEELKASNEEMQAINEELRSATEELETS------- | *Pseudomonas sp. URMO17WK12:I6* | Bacteria; Proteobacteria; Gammaproteobacteria; Pseudomonadales; Pseudomonadaceae; Pseudomonas | response regulator | 6 |
|  | **AMN39323.1** | 1.2E-05 | Qa.IV | -----RTIHELTSVSTSIAAAVDEQAATTREIATSIQTAAGNTARASVEIK-- | *Rhodoplanes sp. Z2-YC6860* | Bacteria; Proteobacteria; Alphaproteobacteria; Rhizobiales; Hyphomicrobiaceae; Rhodoplanes | methyl-accepting chemotaxis protein McpB | 6 |
| | **BAF86724.1** | 1.2E-05 | Qc.III | IEGVGSQIRKMSDVATSISAAVEEQGVATREIVRNVDQAATGTNSVTAHISE- | *Azorhizobium caulinodans ORS 571* | Bacteria; Proteobacteria; Alphaproteobacteria; Rhizobiales; Xanthobacteraceae; Azorhizobium | histidine kinase | 6 |
|  | **BAR97939.1** | 1.2E-05 | Qa.III | IEAITETIRKMNEIATTIASAVEEQGAATTEIARNVQQAS------------- | *Blastochloris viridis* | Bacteria; Proteobacteria; Alphaproteobacteria; Rhizobiales; Hyphomicrobiaceae; Blastochloris | methyl-accepting chemotaxis protein | 6 |
|  | **CAL76835.1** | 1.2E-05 | Qa.IV | IQAISSTIQRVNQIAAAIAAAVEQQGAATREIARNVQQAS------------- | *Bradyrhizobium sp. ORS 278* | Bacteria; Proteobacteria; Alphaproteobacteria; Rhizobiales; Bradyrhizobiaceae; Bradyrhizobium | conserved hypothetical protein putative methyl-accepting chemotaxis receptor/sensory transducer | 6 |
|  | **CAL77022.1** | 1.2E-05 | Qc.III | -KAIGSTIERINEIAASISAAVEQQRAATQNITQSVRSAASG----------- | *Bradyrhizobium sp. ORS 278* | Bacteria; Proteobacteria; Alphaproteobacteria; Rhizobiales; Bradyrhizobiaceae; Bradyrhizobium | putative methyl-accepting chemotaxis protein | 6 |
|  | **CUU41425.1** | 1.2E-05 | Qa.III | IEAITETIRKMNEIATTIASAVEEQGAATTEIARNVQQAS------------- | *Blastochloris viridis* | Bacteria; Proteobacteria; Alphaproteobacteria; Rhizobiales; Hyphomicrobiaceae; Blastochloris | Methyl-accepting chemotaxis protein 4 | 6 |
|  | **EGH92764.1** | 1.2E-05 | Qa.III | ---ISQALRNLNSLNTSIASATLQQTHVVDDINQNVTQAAQLSQ--------- | *Pseudomonas amygdali pv. tabaci str. ATCC 11528* | Bacteria; Proteobacteria; Gammaproteobacteria; Pseudomonadales; Pseudomonadaceae; Pseudomonas; Pseudomonas amygdali | methyl-accepting chemotaxis protein partial | 6 |
|  | **OGR18174.1** | 1.2E-05 | Qa.III | IDQISEVISGVNEIVTTIAAAVEEQTAATREIADNIAQAS------------- | *Desulfobacterales bacterium GWB2_56_26* | Bacteria; Proteobacteria; Deltaproteobacteria; Desulfobacterales | hypothetical protein A2X81_09775 | 6 |
|  | **OGR21159.1** | 1.2E-05 | Qa.III.b | IKEISAIIAEVNEIVAGIAASVQEQSATTGEIAENMNKTSSKM---------- | *Desulfobacula sp. RIFOXYA12_FULL_46_16* | Bacteria; Proteobacteria; Deltaproteobacteria; Desulfobacterales; Desulfobacteraceae; Desulfobacula | hypothetical protein A3J85_01245 | 6 |
|  | **OJU14831.1** | 1.2E-05 | Qa.IV | IKEIGKTITLISEISSAIAAAVEEQGAATQEISRSVQQAAQ------------ | *Afipia sp. 62-7* | Bacteria; Proteobacteria; Alphaproteobacteria; Rhizobiales; Bradyrhizobiaceae; Afipia | methyl-accepting chemotaxis protein | 6 |
|  | **OKP42638.1** | 1.2E-05 | Qa.III | LEQITTSINQISDMATQISSAAEEQRAVTDEVSRNIQAVK------------- | *Aeromonas allosaccharophila* | Bacteria; Proteobacteria; Gammaproteobacteria; Aeromonadales; Aeromonadaceae; Aeromonas | chemotaxis protein partial | 6 |
|  | **OMF22222.1** | 1.2E-05 | Qc.III | -EQVAASATELATGANEISEDVSEMSESITEIEEKVSDYANSTKEMNTR---- | *Paenibacillus sp. FSL H8-0548* | Bacteria; Firmicutes; Bacilli; Bacillales; Paenibacillaceae; Paenibacillus | methyl-accepting chemotaxis protein | 6 |
|  | **OOE90173.1** | 1.2E-05 | Qa.III | -NAINEQIVQISDMNTQVATATEEQSTVVNDINQNIEEINQSSQQTA------ | *Salinivibrio sharmensis* | Bacteria; Proteobacteria; Gammaproteobacteria; Vibrionales; Vibrionaceae; Salinivibrio | methyl-accepting chemotaxis protein | 6 |
|  | **OYX67744.1** | 1.2E-05 | Qa.III.b | ------VIGQINEIAASIASAVDQQGVATREIAHNVARASQGTQQVSENV--- | *Rhizobiales bacterium 32-66-11* | Bacteria; Proteobacteria; Alphaproteobacteria; Rhizobiales | methyl-accepting chemotaxis protein | 6 |
|  | **OYZ98636.1** | 1.2E-05 | Qa.IV | -------ISDIRQVTAAIASAVEQQGAATREIAQNTQHAAHGTEAVNDNIQ-- | *Rhizobiales bacterium 17-65-6* | Bacteria; Proteobacteria; Alphaproteobacteria; Rhizobiales | hypothetical protein B7X99_10825 | 6 |
| | **PEQ15229.1** | 1.2E-05 | Qa.III | -EEIEIAVKKINEMSTDISNAVQEQNTVLKEMNDKTSQIRTASELGDASVE-- | *Piscirickettsia salmonis* | Bacteria; Proteobacteria; Gammaproteobacteria; Thiotrichales; Piscirickettsiaceae; Piscirickettsia | methyl-accepting chemotaxis protein | 6 |
| | **PIT03837.1** | 1.2E-05 | Qc.III | -RDIGGSIGQLSEIASAIAAAVEEQGAATQEIARNVQKAAQGTQQVSANIG-- | *Bradyrhizobium sp. TSA1* | Bacteria; Proteobacteria; Alphaproteobacteria; Rhizobiales; Bradyrhizobiaceae; Bradyrhizobium | chemotaxis protein | 6 |
| | **PIU06208.1** | 1.2E-05 | Qa.III | IEAIARSIRSLSEISLSVSSAVAEQSAVTRDMSVNMQTAAESVREVRSHM--- | *Methylobacterium sp. CG09_land_8_20_14_0_10_71_15* | Bacteria; Proteobacteria; Alphaproteobacteria; Rhizobiales; Methylobacteriaceae; Methylobacterium | histidine kinase | 6 |
| | **PWT85507.1** | 1.2E-05 | Qc.III | IQEIGGTIERLSEVSAAIAAAIEEQGAATQEISRNVQQASVGTQEVSSNIT-- | *Proteobacteria bacterium* | Bacteria; Proteobacteria | methyl-accepting chemotaxis protein | 6 |
| | **RAI20176.1** | 1.2E-05 | Qc.III.c | -EAISQIINQVKEISSGISAAVEQQASVTREISSNMHTAAQGVSEITHSMS-- | *Afifella marina DSM 2698* | Bacteria; Proteobacteria; Alphaproteobacteria; Rhizobiales; Rhodobiaceae; Afifella | chemotaxis protein | 6 |
|  | **WP_007004176.1** | 1.2E-05 | Qa.IV | IQAIAATIEEVSAITVTIAAAVEEQSAATGEIARTVQSTAQATDE-------- | *Roseomonas cervicalis* | Bacteria; Proteobacteria; Alphaproteobacteria; Rhodospirillales; Acetobacteraceae; Roseomonas | HAMP domain-containing protein | 6 |
| | **WP_010627906.1** | 1.2E-05 | R.IV | -AQLNQAMDDMTRVTQDNQQLVAHSSQSAHALREKAEHMQAHAGRFRITA--- | *Halomonas sp. KM-1* | Bacteria; Proteobacteria; Gammaproteobacteria; Oceanospirillales; Halomonadaceae; Halomonas | HAMP domain-containing protein | 6 |
|  | **WP_011473290.1** | 1.2E-05 | Qa.IV | ---ITATIEEVSAIATTIASAIEEQGAATAEIARNVTQTAQATQA-------- | *Rhodopseudomonas palustris* | Bacteria; Proteobacteria; Alphaproteobacteria; Rhizobiales; Bradyrhizobiaceae; Rhodopseudomonas | HAMP domain-containing protein | 6 |
| | **WP_012169255.1** | 1.2E-05 | Qc.III | IEGVGSQIRKMSDVATSISAAVEEQGVATREIVRNVDQAATGTNSVTAHISE- | *Azorhizobium caulinodans* | Bacteria; Proteobacteria; Alphaproteobacteria; Rhizobiales; Xanthobacteraceae; Azorhizobium | methyl-accepting chemotaxis protein | 6 |
| | **WP_012169256.1** | 1.2E-05 | Qc.III | IEGVGSQIRKMSDVATSISAAVEEQGVATREIVRNVDQAATGTNSVTAHISE- | *Azorhizobium caulinodans* | Bacteria; Proteobacteria; Alphaproteobacteria; Rhizobiales; Xanthobacteraceae; Azorhizobium | methyl-accepting chemotaxis protein | 6 |
|  | **WP_013972622.1** | 1.2E-05 | Qb.I | LQRTRLQLQETIEQSEVSSEELTASNEEMQAINEELRSASEELETSK------ | *Pseudomonas putida* | Bacteria; Proteobacteria; Gammaproteobacteria; Pseudomonadales; Pseudomonadaceae; Pseudomonas | response regulator | 6 |
|  | **WP_015270333.1** | 1.2E-05 | Qb.I | LQRTRLQLQETIEQSEVSSEELTASNEEMQAINEELRSASEELETSK------ | *Pseudomonas* | Bacteria; Proteobacteria; Gammaproteobacteria; Pseudomonadales; Pseudomonadaceae | MULTISPECIES: response regulator | 6 |
|  | **WP_023661128.1** | 1.2E-05 | Qb.I | LQRTRLQLQETIEQSEVSSEELTASNEEMQAINEELRSASEELETSK------ | *Pseudomonas plecoglossicida* | Bacteria; Proteobacteria; Gammaproteobacteria; Pseudomonadales; Pseudomonadaceae; Pseudomonas | response regulator | 6 |
|  | **WP_024087132.1** | 1.2E-05 | Qb.I | LQRTRLQLQETIEQSEVSSEELTASNEEMQAINEELRSASEELETSK------ | *Pseudomonas putida group* | Bacteria; Proteobacteria; Gammaproteobacteria; Pseudomonadales; Pseudomonadaceae; Pseudomonas | MULTISPECIES: response regulator | 6 |
|  | **WP_025339100.1** | 1.2E-05 | Qb.I | LQRTRLQLQETIEQSEVSSEELTASNEEMQAINEELRSASEELETSK------ | *Pseudomonas sp. FGI182* | Bacteria; Proteobacteria; Gammaproteobacteria; Pseudomonadales; Pseudomonadaceae; Pseudomonas | response regulator | 6 |
|  | **WP_026312827.1** | 1.2E-05 | Qa.III.b | -EAIAQTIAEINGITASIATSIEQQGFATREIARNVQEASA------------ | *Bradyrhizobium japonicum* | Bacteria; Proteobacteria; Alphaproteobacteria; Rhizobiales; Bradyrhizobiaceae; Bradyrhizobium | chemotaxis protein | 6 |
|  | **WP_026355456.1** | 1.2E-05 | Qc.III | -SRIVRTIGHIQEVAQAISAAVEEQGAATSEIAANTQRAATG----------- | *Pannonibacter phragmitetus* | Bacteria; Proteobacteria; Alphaproteobacteria; Rhodobacterales; Rhodobacteraceae; Pannonibacter | methyl-accepting chemotaxis protein | 6 |
| | **WP_028134921.1** | 1.2E-05 | Qc.III | IQEIGGTIERLSEVASAIAAAVEEQGAATQEISRNVQQAALGTQEVSSNI--- | *Bradyrhizobium japonicum* | Bacteria; Proteobacteria; Alphaproteobacteria; Rhizobiales; Bradyrhizobiaceae; Bradyrhizobium | HAMP domain-containing protein | 6 |
|  | **WP_028583759.1** | 1.2E-05 | Qa.III | IEQISAVITDVNDIVTTIATAVEEQTAATQEIADNIAQASVGIQEVN------ | *Desulfobulbus mediterraneus* | Bacteria; Proteobacteria; Deltaproteobacteria; Desulfobacterales; Desulfobulbaceae; Desulfobulbus | methyl-accepting chemotaxis protein | 6 |
|  | **WP_029858276.1** | 1.2E-05 | Qa.III | LDEINAAVALISDMATQIATAAEEQTHVTNEITQNVTSIK------------- | *Vibrio parahaemolyticus* | Bacteria; Proteobacteria; Gammaproteobacteria; Vibrionales; Vibrionaceae; Vibrio | methyl-accepting chemotaxis protein partial | 6 |
|  | **WP_034328367.1** | 1.2E-05 | Qc.I | -DEFAQMVRDIKENIKDVTTGLEQDQKLVEQSLQVIERAKQ------------ | *Helicobacter sp. MIT 03-1616* | Bacteria; Proteobacteria; Epsilonproteobacteria; Campylobacterales; Helicobacteraceae; Helicobacter | chemotaxis protein partial | 6 |
| | **WP_037000656.1** | 1.2E-05 | Qa.III | --RICAEVAQISGMSLQMAAAIEQQGQVAEQINQQITEIAGFTERSSQQAGR- | *Pseudomonas pseudoalcaligenes* | Bacteria; Proteobacteria; Gammaproteobacteria; Pseudomonadales; Pseudomonadaceae; Pseudomonas; Pseudomonas oleovorans/ pseudoalcaligenes group | PAS domain S-box protein | 6 |

| | | | | | | | | |
|---|---|---|---|---|---|---|---|---|
|  | **WP_038409645.1** | 1.2E-05 | Qb.I | LQRTRLQLQETIEQSEVSSEELTASNEEMQAINEELRSASEELETSK------ | *Pseudomonas putida* | Bacteria; Proteobacteria; Gammaproteobacteria; Pseudomonadales; Pseudomonadaceae; Pseudomonas | response regulator | 6 |
|  | **WP_039613390.1** | 1.2E-05 | Qb.I | LQRTRLQLQETIEQSEVSSEELTASNEEMQAINEELRSASEELETSK------ | *Pseudomonas sp. C5pp* | Bacteria; Proteobacteria; Gammaproteobacteria; Pseudomonadales; Pseudomonadaceae; Pseudomonas | response regulator | 6 |
|  | **WP_046614161.1** | 1.2E-05 | Qb.I | LQRTRLQLQETIEQSEVSSEELTASNEEMQAINEELRSASEELETSK------ | *Pseudomonas putida* | Bacteria; Proteobacteria; Gammaproteobacteria; Pseudomonadales; Pseudomonadaceae; Pseudomonas | response regulator | 6 |
| | **WP_050991793.1** | 1.2E-05 | Qc.III | IKDISHTIEKLSEISSTIAAAVEEQGAATQEISRNVQQAATGTHQVSSNITD- | *Bradyrhizobium yuanmingense* | Bacteria; Proteobacteria; Alphaproteobacteria; Rhizobiales; Bradyrhizobiaceae; Bradyrhizobium | methyl-accepting chemotaxis protein | 6 |
| | **WP_050996044.1** | 1.2E-05 | Qc.III | IKDISHTIEKLSEISSTIAAAVEEQGAATQEISRNVQQAATGTHQVSSNITD- | *Bradyrhizobium yuanmingense* | Bacteria; Proteobacteria; Alphaproteobacteria; Rhizobiales; Bradyrhizobiaceae; Bradyrhizobium | methyl-accepting chemotaxis protein | 6 |
| | **WP_051056864.1** | 1.2E-05 | Qc.III | IKDISHTIEKLSEISSTIAAAVEEQGAATQEISRNVQQAATGTHQVSSNITD- | *Bradyrhizobium yuanmingense* | Bacteria; Proteobacteria; Alphaproteobacteria; Rhizobiales; Bradyrhizobiaceae; Bradyrhizobium | methyl-accepting chemotaxis protein | 6 |
|  | **WP_051305399.1** | 1.2E-05 | Qa.III | IEEISAVIIEVNEVIATIASAVEEQTAATREISDNISQAS------------ | *Desulfobulbus mediterraneus* | Bacteria; Proteobacteria; Deltaproteobacteria; Desulfobacterales; Desulfobulbaceae; Desulfobulbus | methyl-accepting chemotaxis protein | 6 |
|  | **WP_054054551.1** | 1.2E-05 | Qb.I | LQRTKLQLQDTIEQSEVSSEELKASNEEMQAINEELRSATEELETS------- | *Pseudomonas sp. 655* | Bacteria; Proteobacteria; Gammaproteobacteria; Pseudomonadales; Pseudomonadaceae; Pseudomonas | response regulator | 6 |
|  | **WP_054573300.1** | 1.2E-05 | Qb.I | LQRTRLQLQETIEQSEVSSEELTASNEEMQAINEELRSASEELETSK------ | *Pseudomonas putida* | Bacteria; Proteobacteria; Gammaproteobacteria; Pseudomonadales; Pseudomonadaceae; Pseudomonas | response regulator | 6 |
|  | **WP_054614203.1** | 1.2E-05 | Qb.I | LQRTKLQLQDTIEQSEVSSEELKASNEEMQAINEELRSATEELETS------- | *Pseudomonas sp. In5* | Bacteria; Proteobacteria; Gammaproteobacteria; Pseudomonadales; Pseudomonadaceae; Pseudomonas | response regulator | 6 |
|  | **WP_056797076.1** | 1.2E-05 | Qb.I | LQRTRLQLQETIEQSEVSSEELTASNEEMQAINEELRSASEELETSK------ | *Pseudomonas sp. Leaf58* | Bacteria; Proteobacteria; Gammaproteobacteria; Pseudomonadales; Pseudomonadaceae; Pseudomonas | response regulator | 6 |
| | **WP_057027010.1** | 1.2E-05 | Qc.III | IKDISHTIEKLSEISSTIAAAVEEQGAATQEISRNVQQAATGTHQVSSNITD- | *Bradyrhizobium yuanmingense* | Bacteria; Proteobacteria; Alphaproteobacteria; Rhizobiales; Bradyrhizobiaceae; Bradyrhizobium | methyl-accepting chemotaxis protein | 6 |
|  | **WP_057259997.1** | 1.2E-05 | Qa.III | MHDIVKSVQNVASLMQEIASASQEQSLGIDQVNQAVTQMDNTTQQ-------- | *Duganella sp. Root1480D1* | Bacteria; Proteobacteria; Betaproteobacteria; Burkholderiales; Oxalobacteraceae; Duganella | methyl-accepting chemotaxis protein | 6 |
|  | **WP_059404881.1** | 1.2E-05 | Qb.I | LQRTKLHLQDTIEQSEISSEELKASNEEMQAINEELRSATEELETS------- | *Pseudomonas sp. RIT-PI-q* | Bacteria; Proteobacteria; Gammaproteobacteria; Pseudomonadales; Pseudomonadaceae; Pseudomonas | PAS domain S-box protein | 6 |
|  | **WP_060507047.1** | 1.2E-05 | Qb.I | LQRTRLQLQETIEQSEVSSEELTASNEEMQAINEELRSASEELETSK------ | *Pseudomonas sp. NBRC 111124* | Bacteria; Proteobacteria; Gammaproteobacteria; Pseudomonadales; Pseudomonadaceae; Pseudomonas | response regulator | 6 |
|  | **WP_061551220.1** | 1.2E-05 | Qb.I | LQRTRLQLQETIEQSEVSSEELTASNEEMQAINEELRSASEELETSK------ | *Pseudomonas sp. ABFPK* | Bacteria; Proteobacteria; Gammaproteobacteria; Pseudomonadales; Pseudomonadaceae; Pseudomonas | response regulator | 6 |
|  | **WP_068309225.1** | 1.2E-05 | Qa.IV | -------VREIAELVNAVSAAVEQQQAATSEIARNIQEAAKGTREVNENI--- | *Kordiimonas lacus* | Bacteria; Proteobacteria; Alphaproteobacteria; Kordiimonadales; Kordiimonadaceae; Kordiimonas | hypothetical protein | 6 |
| | **WP_068812421.1** | 1.2E-05 | Qa.III | --EINDSVVSIADMTVHVAAAVEEQSVVIEEINQNLASISEQSVTTSDSATK- | *Pseudohongiella nitratireducens* | Bacteria; Proteobacteria; Gammaproteobacteria; Pseudohongiella | methyl-accepting chemotaxis protein | 6 |
|  | **WP_075045228.1** | 1.2E-05 | Qb.I | LQRTRLQLQETIEQSEVSSEELTASNEEMQAINEELRSASEELETSK------ | *Pseudomonas putida* | Bacteria; Proteobacteria; Gammaproteobacteria; Pseudomonadales; Pseudomonadaceae; Pseudomonas | response regulator | 6 |
|  | **WP_079541968.1** | 1.2E-05 | Qa.IV | IRNIGQTIGEINDVTTAIAAAVEEQGAATREIARNIQHAA------------ | *Bradyrhizobium lablabi* | Bacteria; Proteobacteria; Alphaproteobacteria; Rhizobiales; Bradyrhizobiaceae; Bradyrhizobium | HAMP domain-containing protein | 6 |
|  | **WP_080804051.1** | 1.2E-05 | Qa.III.b | IVKIISVISNVNEIVTTIASAVEEQTVTTQEITGNLSQASAGIEEVNSN---- | *Desulfamplus magnetovallimortis* | Bacteria; Proteobacteria; Deltaproteobacteria; Desulfobacterales; Desulfobacteraceae; Desulfamplus | methyl-accepting chemotaxis protein | 6 |
|  | **WP_082675860.1** | 1.2E-05 | Qc.III | -SRIVRTIGHIQEVAQAISAAVEEQGAATSEIAANTQRAATG----------- | *Pannonibacter phragmitetus* | Bacteria; Proteobacteria; Alphaproteobacteria; Rhodobacterales; Rhodobacteraceae; Pannonibacter | methyl-accepting chemotaxis protein | 6 |
|  | **WP_083676533.1** | 1.2E-05 | Qc.III | -EQVAASATELATGANEISEDVSEMSESITEIEEKVSDYANSTKEMNTR---- | *Paenibacillus sp. FSL H8-0548* | Bacteria; Firmicutes; Bacilli; Bacillales; Paenibacillaceae; Paenibacillus | methyl-accepting chemotaxis protein | 6 |
|  | **WP_084438569.1** | 1.2E-05 | Qc.III | --GIAGTIRTINETATAIAGAVEEQTAATQEIARNVDEAARGTDDASR----- | *Niveispirillum irakense* | Bacteria; Proteobacteria; Alphaproteobacteria; Rhodospirillales; Rhodospirillaceae; Niveispirillum | methyl-accepting chemotaxis protein | 6 |
|  | **WP_085589675.1** | 1.2E-05 | Qb.I | LQRTRLQLQETIEQSEVSSEELTASNEEMQAINEELRSASEELETSK------ | *Pseudomonas sp. B14(2017)* | Bacteria; Proteobacteria; Gammaproteobacteria; Pseudomonadales; Pseudomonadaceae; Pseudomonas | response regulator | 6 |
|  | **WP_085615736.1** | 1.2E-05 | Qb.I | LQRTRLQLQETIEQSEVSSEELTASNEEMQAINEELRSASEELETSK------ | *Pseudomonas* | Bacteria; Proteobacteria; Gammaproteobacteria; Pseudomonadales; Pseudomonadaceae | MULTISPECIES: response regulator | 6 |
|  | **WP_085625069.1** | 1.2E-05 | Qb.I | LQRTRLQLQETIEQSEVSSEELTASNEEMQAINEELRSASEELETSK------ | *Pseudomonas* | Bacteria; Proteobacteria; Gammaproteobacteria; Pseudomonadales; Pseudomonadaceae | MULTISPECIES: response regulator | 6 |
|  | **WP_085664889.1** | 1.2E-05 | Qb.I | LQRTRLQLQETIEQSEVSSEELTASNEEMQAINEELRSASEELETSK------ | *Pseudomonas sp. B5(2017)* | Bacteria; Proteobacteria; Gammaproteobacteria; Pseudomonadales; Pseudomonadaceae; Pseudomonas | response regulator | 6 |
|  | **WP_085692593.1** | 1.2E-05 | Qb.I | LQRTRLQLQETIEQSEVSSEELTASNEEMQAINEELRSASEELETSK------ | *Pseudomonas* | Bacteria; Proteobacteria; Gammaproteobacteria; Pseudomonadales; Pseudomonadaceae | MULTISPECIES: response regulator | 6 |
|  | **WP_085720169.1** | 1.2E-05 | Qb.I | LQRTRLQLQETIEQSEVSSEELTASNEEMQAINEELRSASEELETSK------ | *Pseudomonas sp. B22(2017)* | Bacteria; Proteobacteria; Gammaproteobacteria; Pseudomonadales; Pseudomonadaceae; Pseudomonas | response regulator | 6 |
|  | **WP_092191296.1** | 1.2E-05 | Qa.III | IQSIVQVIQDVNDIVTTIAAAIEEQAMATREVATNIAQATTGVQDAN------ | *Desulfomicrobium norvegicum* | Bacteria; Proteobacteria; Deltaproteobacteria; Desulfovibrionales; Desulfomicrobiaceae; Desulfomicrobium | methyl-accepting chemotaxis protein | 6 |
|  | **WP_092385976.1** | 1.2E-05 | Qa.III | -DGISQAIEQINDMTNQIASAAEEQSAVAEEINRNITTIAD------------ | *Pseudomonas salegens* | Bacteria; Proteobacteria; Gammaproteobacteria; Pseudomonadales; Pseudomonadaceae; Pseudomonas | PAS domain S-box protein | 6 |
| | **WP_094407839.1** | 1.2E-05 | Qc.III | --GISGTIEQISEIATSVAAAVEEQGAATDEIARNVQQAASGTHQVTQHVGS- | *Elstera cyanobacteriorum* | Bacteria; Proteobacteria; Alphaproteobacteria; Rhodospirillales; Rhodospirillaceae; Elstera | methyl-accepting chemotaxis protein | 6 |
|  | **WP_094409886.1** | 1.2E-05 | Qa.IV | IQQVTATMLQVNSTSGIIATAVVSQGAATEEIARNAQEVAQRT---------- | *Elstera cyanobacteriorum* | Bacteria; Proteobacteria; Alphaproteobacteria; Rhodospirillales; Rhodospirillaceae; Elstera | methyl-accepting chemotaxis protein | 6 |
|  | **WP_095942703.1** | 1.2E-05 | Qb.I | LQRTKLHLQDTIEQSEISSEELKASNEEMQAINEELRSATEELETS------- | *Pseudomonas sp. ACN8* | Bacteria; Proteobacteria; Gammaproteobacteria; Pseudomonadales; Pseudomonadaceae; Pseudomonas | response regulator | 6 |
| | **WP_100178965.1** | 1.2E-05 | Qc.III | -RDIGGSIGQLSEIASAIAAAVEEQGAATQEIARNVQKAAQGTQQVSANIG-- | *Bradyrhizobium sp. TSA1* | Bacteria; Proteobacteria; Alphaproteobacteria; Rhizobiales; Bradyrhizobiaceae; Bradyrhizobium | HAMP domain-containing protein | 6 |
|  | **WP_100413617.1** | 1.2E-05 | Qb.I | LQRTRLQLQETIEQSEVSSEELTASNEEMQAINEELRSASEELETSK------ | *Pseudomonas sp. MR 02* | Bacteria; Proteobacteria; Gammaproteobacteria; Pseudomonadales; Pseudomonadaceae; Pseudomonas | response regulator | 6 |
|  | **WP_102084010.1** | 1.2E-05 | Qb.I | LQRTRLQLQETIEQSEVSSEELTASNEEMQAINEELRSASEELETSK------ | *Pseudomonas plecoglossicida* | Bacteria; Proteobacteria; Gammaproteobacteria; Pseudomonadales; Pseudomonadaceae; Pseudomonas | response regulator | 6 |
| | **WP_106738700.1** | 1.2E-05 | Qa.III | --RICAEVAQISGMSLQMAAAIEQQGQVAEQINQQITEIAGFTERTSQQAGR- | *Pseudomonas mendocina* | Bacteria; Proteobacteria; Gammaproteobacteria; Pseudomonadales; Pseudomonadaceae; Pseudomonas | PAS domain S-box protein | 6 |
|  | **WP_108127530.1** | 1.2E-05 | Qa.III | -RKIQAFMQEINDISTQIATATEEQSQVVAGINQNLQSIADYS---------- | *Saccharospirillum mangrovi* | Bacteria; Proteobacteria; Gammaproteobacteria; Oceanospirillales; Saccharospirillaceae; Saccharospirillum | methyl-accepting chemotaxis protein | 6 |
|  | **WP_108215250.1** | 1.2E-05 | Qb.I | LQRTKLHLQDTIEQSEISSEELKASNEEMQAINEELRSATEELETS------- | *Pseudomonas sp. GV085* | Bacteria; Proteobacteria; Gammaproteobacteria; Pseudomonadales; Pseudomonadaceae; Pseudomonas | PAS domain S-box protein | 6 |
|  | **WP_108480981.1** | 1.2E-05 | Qb.I | LQRTRLQLQETIEQSEVSSEELTASNEEMQAINEELRSASEELETSK------ | *Pseudomonas plecoglossicida* | Bacteria; Proteobacteria; Gammaproteobacteria; Pseudomonadales; Pseudomonadaceae; Pseudomonas | response regulator | 6 |
| | **WP_109106221.1** | 1.2E-05 | Qa.III | IKDVARVITRINEVATSISSAVEQQGAATREISRNVQQAAQGTQETSRSIVG- | *Azospirillum sp. TSO35-2* | Bacteria; Proteobacteria; Alphaproteobacteria; Rhodospirillales; Rhodospirillaceae; Azospirillum | methyl-accepting chemotaxis protein | 6 |
| | **WP_109145903.1** | 1.2E-05 | Qc.III | IKDISHTIEKLSEISSTIAAAVEEQGAATQEISRNVQQAATGTHQVSSNITD- | *Bradyrhizobium sp. SUTN9-2* | Bacteria; Proteobacteria; Alphaproteobacteria; Rhizobiales; Bradyrhizobiaceae; Bradyrhizobium | methyl-accepting chemotaxis protein | 6 |
|  | **WP_109154063.1** | 1.2E-05 | Qa.III | IVGIGRTIQSINQIASGISAGIEQQGSAVREIVVNMEQAA------------ | *Azospirillum sp. TSO5* | Bacteria; Proteobacteria; Alphaproteobacteria; Rhodospirillales; Rhodospirillaceae; Azospirillum | chemotaxis protein | 6 |
|  | **WP_110745388.1** | 1.2E-05 | Qb.I | LQRTRLQLQETIEQSEVSSEELTASNEEMQAINEELRSASEELETSK------ | *Pseudomonas sp. LB-090624* | Bacteria; Proteobacteria; Gammaproteobacteria; Pseudomonadales; Pseudomonadaceae; Pseudomonas | response regulator | 6 |
|  | **WP_111958146.1** | 1.2E-05 | Qa.III | IQDITDVINQVNDIVSTIAMAVEEQTGTIAEIDENVSQGAHALQEVNAN---- | *Desulfobacter hydrogenophilus* | Bacteria; Proteobacteria; Deltaproteobacteria; Desulfobacterales; Desulfobacteraceae; Desulfobacter | methyl-accepting chemotaxis protein | 6 |
| | **WP_112211008.1** | 1.2E-05 | Qa.III | --RICAEVAQISGMSLQMAAAIEQQGQVAEQINQQITEIAGFTERTSQQAGR- | *Pseudomonas* | Bacteria; Proteobacteria; Gammaproteobacteria; Pseudomonadales; Pseudomonadaceae | MULTISPECIES: PAS domain S-box protein | 6 |
|  | **WP_114010731.1** | 1.2E-05 | Qa.IV | IRAIATKMTEISEYTTAISAAVQQQNASTNEISHNIQQAAE----------- | *Cohaesibacter sp. YE-B6* | Bacteria; Proteobacteria; Alphaproteobacteria; Rhizobiales; Cohaesibacteraceae | HAMP domain-containing protein | 6 |
|  | **AWZ02129.1** | 1.3E-05 | Qa.IV | IREVTSTVGEISEISGAIASSIEQQQEATVEISSNVQEAA------------ | *Rhodobiaceae bacterium* | Bacteria; Proteobacteria; Alphaproteobacteria; Rhizobiales; Rhodobiaceae; unclassified Rhodobiaceae | methyl-accepting chemotaxis protein McpB | 6 |
|  | **CDE55863.1** | 1.3E-05 | Qb.I | --ETKNSVNGVVEKSEHMTTLMEHSSEMVSDIRKRLDTQESLLQ--------- | *Roseburia sp. CAG:303* | Bacteria; Firmicutes; Clostridia; Clostridiales; Lachnospiraceae; Roseburia; environmental samples | methyl-accepting chemotaxis sensory transducer | 6 |
|  | **EGH25779.1** | 1.3E-05 | Qa.III | ---ISQALRNLNSLNTSIASATLQQTHVVDDINQNVTQAAQLSQ--------- | *Pseudomonas amygdali pv. mori str. 301020* | Bacteria; Proteobacteria; Gammaproteobacteria; Pseudomonadales; Pseudomonadaceae; Pseudomonas; Pseudomonas amygdali | methyl-accepting chemotaxis protein partial | 6 |
|  | **OCR92162.1** | 1.3E-05 | Qa.IV | --EIEANVNILSQGINEMSQSINEQTEAINQINEAVATVDEQTKQ-NV----- | *Campylobacter fetus subsp. testudinum* | Bacteria; Proteobacteria; Epsilonproteobacteria; Campylobacterales; Campylobacteraceae; Campylobacter | chemotaxis protein partial | 6 |
|  | **ODM81340.1** | 1.3E-05 | Qc.III.c | --AISQVIVQISEISTVIASAVEEQAAATKEVTANINGTTKAIDDA------- | *Bradyrhizobium elkanii* | Bacteria; Proteobacteria; Alphaproteobacteria; Rhizobiales; Bradyrhizobiaceae; Bradyrhizobium | hypothetical protein A6452_21550 | 6 |
|  | **OIN99761.1** | 1.3E-05 | SNAP.b | --EAMRTLHTMSEKLRESMQRLGKQSEAIGQVMGVINDIADQT---------- | *Desulfovibrionaceae bacterium CG1_02_65_16* | Bacteria; Proteobacteria; Deltaproteobacteria; Desulfovibrionales; Desulfovibrionaceae | chemotaxis protein | 6 |
|  | **OIQ02941.1** | 1.3E-05 | Qc.III | ---VGDVIGKMNEINRMISAAADEQNQATREIAQSAQYASEATHNVTE----- | *Zetaproteobacteria bacterium CG2_30_59_37* | Bacteria; Proteobacteria; Zetaproteobacteria | hypothetical protein AUK36_06100 | 6 |
| | **PIT01033.1** | 1.3E-05 | Qc.III | IKDISHTIEKLSEISSTIAAAVEEQGAATQEISRNVQQAAAGTHQVSSNITD- | *Bradyrhizobium sp. TSA1* | Bacteria; Proteobacteria; Alphaproteobacteria; Rhizobiales; Bradyrhizobiaceae; Bradyrhizobium | chemotaxis protein | 6 |
|  | **PJJ13551.1** | 1.3E-05 | Qa.III | IREIRTTIGSMSEIASDIASAVETQGVMTREIATNVGQAA------------ | *Bradyrhizobium lablabi* | Bacteria; Proteobacteria; Alphaproteobacteria; Rhizobiales; Bradyrhizobiaceae; Bradyrhizobium | methyl-accepting chemotaxis protein | 6 |
|  | **PKF62306.1** | 1.3E-05 | Qa.III | LNEIVQNIQQISDLNTQVATATEEQSVVINEINTHVHSISDSSEESANS---- | *Psychromonas sp. psych-6C06* | Bacteria; Proteobacteria; Gammaproteobacteria; Alteromonadales; Psychromonadaceae; Psychromonas | methyl-accepting chemotaxis protein | 6 |
|  | **PTY38509.1** | 1.3E-05 | Qa.III | -QKIQTYMQEINDISTQIATATEEQSQVVSGINENLQSIADYS---------- | *Saccharospirillum sp. MSK14-1* | Bacteria; Proteobacteria; Gammaproteobacteria; Oceanospirillales; Saccharospirillaceae; Saccharospirillum | hypothetical protein BGP77_12445 | 6 |
|  | **PZP40535.1** | 1.3E-05 | Qa.IV | IEEITRTIGEVSEATVTIAAAVEEQTAATAEIARTVQQTA------------ | *Azospirillum brasilense* | Bacteria; Proteobacteria; Alphaproteobacteria; Rhodospirillales; Rhodospirillaceae; Azospirillum | hypothetical protein DI601_23690 | 6 |

| | | | | | | | | |
|---|---|---|---|---|---|---|---|---|
|  | **RAI28462.1** | 1.3E-05 | Qa.IV | IQQIGSTIKDMNQITVGIASAMEEQNAATDEIARNVEQAA------------ | *Rhodobium orientis* | Bacteria; Proteobacteria; Alphaproteobacteria; Rhizobiales; Rhodobiaceae; Rhodobium | hypothetical protein CH339_06115 | 6 |
| | **WP_008944934.1** | 1.3E-05 | Qc.III | --EIASIIGEINEIATVIASAVEEQNAATQEIARNVQQAASGTQEVSSNI--- | *Oceanibaculum indicum* | Bacteria; Proteobacteria; Alphaproteobacteria; Rhodospirillales; Rhodospirillaceae; Oceanibaculum | HAMP domain-containing protein | 6 |
|  | **WP_008973265.1** | 1.3E-05 | Qc.III | -RDISGTIERLAEIASAIAAAVEEQGAATQEISRNIQQASVGTQQV------- | *Bradyrhizobium sp. STM 3843* | Bacteria; Proteobacteria; Alphaproteobacteria; Rhizobiales; Bradyrhizobiaceae; Bradyrhizobium | HAMP domain-containing protein | 6 |
|  | **WP_009031132.1** | 1.3E-05 | Qa.IV | IREISTTIERLSEISATVAAAVEEQGAATSEIARNIQHAA------------ | *Bradyrhizobium sp. ORS 375* | Bacteria; Proteobacteria; Alphaproteobacteria; Rhizobiales; Bradyrhizobiaceae; Bradyrhizobium | PAS domain S-box protein | 6 |
| | **WP_013717281.1** | 1.3E-05 | Qa.III | --RICAEVAQISGMSLQMAAAIEQQGQVAEQINQQISEIAGFTERTSQQAGR- | *Pseudomonas mendocina* | Bacteria; Proteobacteria; Gammaproteobacteria; Pseudomonadales; Pseudomonadaceae; Pseudomonas | PAS domain S-box protein | 6 |
| | **WP_017362066.1** | 1.3E-05 | Qa.III | --RICAEVAQISGMSLQMAAAIEQQGQVAEQINQQISEIAGFTERTSQQAGR- | *Pseudomonas mendocina* | Bacteria; Proteobacteria; Gammaproteobacteria; Pseudomonadales; Pseudomonadaceae; Pseudomonas | PAS domain S-box protein | 6 |
|  | **WP_020589137.1** | 1.3E-05 | Qa.III | ITEISKIISEVNDIVSSIAAAIEEQSSATREIAENILQVSSGIQEVNK----- | *Desulfobacter curvatus* | Bacteria; Proteobacteria; Deltaproteobacteria; Desulfobacterales; Desulfobacteraceae; Desulfobacter | hypothetical protein | 6 |
|  | **WP_021081828.1** | 1.3E-05 | Qc.III.c | --AISQVIVQISEISTVIASAVEEQAAATKEVTANINGTTKAIDDA------- | *Bradyrhizobium sp. MOS004* | Bacteria; Proteobacteria; Alphaproteobacteria; Rhizobiales; Bradyrhizobiaceae; Bradyrhizobium | PAS domain-containing protein | 6 |
| | **WP_024341539.1** | 1.3E-05 | Qc.III | IKDISHTIEKLSEISSTIAAAVEEQGAATQEISRNVQQAAAGTHQVSSNITD- | *Bradyrhizobium japonicum* | Bacteria; Proteobacteria; Alphaproteobacteria; Rhizobiales; Bradyrhizobiaceae; Bradyrhizobium | methyl-accepting chemotaxis protein | 6 |
|  | **WP_027281646.1** | 1.3E-05 | Qa.IV | IEEITRTIGEVSEATVTIAAAVEEQTAATAEIARTVQQTA------------ | *Roseomonas gilardii* | Bacteria; Proteobacteria; Alphaproteobacteria; Rhodospirillales; Acetobacteraceae; Roseomonas | HAMP domain-containing protein | 6 |
| | **WP_027671846.1** | 1.3E-05 | Qa.III | --NIRDSVRTITSLTEQMSAAVEQQGTVAEHINQQIVEIADSTSATHTSANS- | *Rheinheimera baltica* | Bacteria; Proteobacteria; Gammaproteobacteria; Chromatiales; Chromatiaceae; Rheinheimera | PAS domain S-box protein | 6 |
|  | **WP_028579965.1** | 1.3E-05 | Qa.III | IDQISEVINKINEIVAIISTAVGEQSAATQEIADNIAQAS------------ | *Desulfobulbus japonicus* | Bacteria; Proteobacteria; Deltaproteobacteria; Desulfobacterales; Desulfobulbaceae; Desulfobulbus | methyl-accepting chemotaxis protein | 6 |
|  | **WP_029832524.1** | 1.3E-05 | Qa.III | LDEINAAVALISDMATQIATAAEEQTHVTNEITQNVTSIK------------ | *Vibrio parahaemolyticus* | Bacteria; Proteobacteria; Gammaproteobacteria; Vibrionales; Vibrionaceae; Vibrio | methyl-accepting chemotaxis protein partial | 6 |
|  | **WP_029847035.1** | 1.3E-05 | Qa.III | LDEINAAVALISDMATQIATAAEEQTHVTNEITQNVTSIK------------ | *Vibrio parahaemolyticus* | Bacteria; Proteobacteria; Gammaproteobacteria; Vibrionales; Vibrionaceae; Vibrio | methyl-accepting chemotaxis protein partial | 6 |
|  | **WP_035633959.1** | 1.3E-05 | Qc.III.c | --AISQVIVQISEISTVIASAVEEQAAATKEVTANINGTTKAIDDA------- | *Bradyrhizobium elkanii* | Bacteria; Proteobacteria; Alphaproteobacteria; Rhizobiales; Bradyrhizobiaceae; Bradyrhizobium | PAS domain-containing protein | 6 |
|  | **WP_038376907.1** | 1.3E-05 | Qc.III.c | --AISQVIVQISEISTVIASAVEEQAAATKEVTANINGTTKAIDDA------- | *Bradyrhizobium elkanii* | Bacteria; Proteobacteria; Alphaproteobacteria; Rhizobiales; Bradyrhizobiaceae; Bradyrhizobium | PAS domain-containing protein | 6 |
|  | **WP_038377474.1** | 1.3E-05 | Qc.III.c | --AISQVIVQISEISTVIASAVEEQAAATKEVTANINGTTKAIDDA------- | *Bradyrhizobium* | Bacteria; Proteobacteria; Alphaproteobacteria; Rhizobiales; Bradyrhizobiaceae | MULTISPECIES: PAS domain-containing protein | 6 |
|  | **WP_038379838.1** | 1.3E-05 | Qc.III.c | --AISQVIVQISEISTVIASAVEEQAAATKEVTANINGTTKAIDDA------- | *Bradyrhizobium elkanii* | Bacteria; Proteobacteria; Alphaproteobacteria; Rhizobiales; Bradyrhizobiaceae; Bradyrhizobium | PAS domain-containing protein | 6 |
| | **WP_041980470.1** | 1.3E-05 | Qa.III | --RICAEVAQISGMSLQMAAAIEQQGQVAEQINQQISEIAGFTERTSQQAGR- | *Pseudomonas mendocina* | Bacteria; Proteobacteria; Gammaproteobacteria; Pseudomonadales; Pseudomonadaceae; Pseudomonas | PAS domain S-box protein | 6 |
|  | **WP_042955604.1** | 1.3E-05 | Qb.I | LQRTKLHLQETIEQSEVSSEELKASNEEMQAINEELRSATEELETS------- | *Pseudomonas sp. G5(2012)* | Bacteria; Proteobacteria; Gammaproteobacteria; Pseudomonadales; Pseudomonadaceae; Pseudomonas | PAS domain S-box protein partial | 6 |
| | **WP_049820014.1** | 1.3E-05 | Qc.III | IKDISHTIEKLSEISSTIAAAVEEQGAATQEISRNVQQAAAGTHQVSSNITD- | *Bradyrhizobium japonicum* | Bacteria; Proteobacteria; Alphaproteobacteria; Rhizobiales; Bradyrhizobiaceae; Bradyrhizobium | methyl-accepting chemotaxis protein | 6 |
|  | **WP_050735583.1** | 1.3E-05 | Qa.IV | MQQIARTIASVSEISGSIASTVVEQTAATSEISRNAGQAA------------ | *Methylobacterium sp. ARG-1* | Bacteria; Proteobacteria; Alphaproteobacteria; Rhizobiales; Methylobacteriaceae; Methylobacterium | methyl-accepting chemotaxis protein | 6 |
|  | **WP_051002922.1** | 1.3E-05 | Qc.III.c | --AISQVIVQISEISTVIASAVEEQAAATKEVTANINGTTKAIDDA------- | *Bradyrhizobium elkanii* | Bacteria; Proteobacteria; Alphaproteobacteria; Rhizobiales; Bradyrhizobiaceae; Bradyrhizobium | PAS domain-containing protein | 6 |
| | **WP_051284999.1** | 1.3E-05 | Qc.III.c | -EALAGTVNDIDQVAGTIAAAAEEQSAATNEIARSVESTSANVQDVTVNIGE- | *Nisaea denitrificans* | Bacteria; Proteobacteria; Alphaproteobacteria; Rhodospirillales; Rhodospirillaceae; Nisaea | HAMP domain-containing protein | 6 |
|  | **WP_051402751.1** | 1.3E-05 | Qa.IV | IREVTSTVGEISEISGAIASSIEQQQEATVEISSNVQEAA------------ | *alpha proteobacterium Mf 1.05b.01* | Bacteria; Proteobacteria; Alphaproteobacteria | methyl-accepting chemotaxis protein | 6 |
|  | **WP_054026866.1** | 1.3E-05 | Qc.III | -QQVAASANELATGANEISEDVSEMSESITEIEVKVSDYATSTKEMNTR---- | *Bacillus sp. FJAT-28004* | Bacteria; Firmicutes; Bacilli; Bacillales; Bacillaceae; Bacillus | methyl-accepting chemotaxis protein | 6 |
| | **WP_057195848.1** | 1.3E-05 | Qc.III | IKDISHTIEKLSEISSTIAAAVEEQGAATQEISRNVQQAAAGTHQVSSNITD- | *Bradyrhizobium sp. Leaf396* | Bacteria; Proteobacteria; Alphaproteobacteria; Rhizobiales; Bradyrhizobiaceae; Bradyrhizobium | methyl-accepting chemotaxis protein | 6 |
|  | **WP_063691902.1** | 1.3E-05 | Qc.III | IQEIGGTIERLSEVSAAIAAAVEEQGAATQEISRNVQQASVGTQEVSSN---- | *Bradyrhizobium stylosanthis* | Bacteria; Proteobacteria; Alphaproteobacteria; Rhizobiales; Bradyrhizobiaceae; Bradyrhizobium | methyl-accepting chemotaxis protein | 6 |
|  | **WP_063790157.1** | 1.3E-05 | Qc.III.c | --AISQVIVQISEISTVIASAVEEQAAATKEVTANINGTTKAIDDA------- | *Bradyrhizobium* | Bacteria; Proteobacteria; Alphaproteobacteria; Rhizobiales; Bradyrhizobiaceae | MULTISPECIES: PAS domain-containing protein | 6 |
|  | **WP_069680634.1** | 1.3E-05 | Qc.III | -EKVNDIAKQNSELTQQISSQLDEQSNSLDKFNDTLNNLSKDI---------- | *Clostridium taeniosporum* | Bacteria; Firmicutes; Clostridia; Clostridiales; Clostridiaceae; Clostridium | chemotaxis protein | 6 |
|  | **WP_069937493.1** | 1.3E-05 | Qb.I | ----NTIVAQTVEQMNLVQDSTTESSEVVNALGQKSKEIGQIVE--------- | *Domibacillus iocasae* | Bacteria; Firmicutes; Bacilli; Bacillales; Bacillaceae; Domibacillus | methyl-accepting chemotaxis protein | 6 |
|  | **WP_075798831.1** | 1.3E-05 | Qa.IV | IEEITRTIGEVSEATVTIAAAVEEQTAATAEIARTVQQTA------------ | *Roseomonas gilardii* | Bacteria; Proteobacteria; Alphaproteobacteria; Rhodospirillales; Acetobacteraceae; Roseomonas | HAMP domain-containing protein | 6 |
| | **WP_081753032.1** | 1.3E-05 | Qc.III | -KGIDGTIGRISEISTTIAAAIEEQGAATGEITRSTQETARGTEAVSRNIA-- | *Rhodospirillales bacterium URHD0088* | Bacteria; Proteobacteria; Alphaproteobacteria; Rhodospirillales; unclassified Rhodospirillales | methyl-accepting chemotaxis protein | 6 |
|  | **WP_082416710.1** | 1.3E-05 | Qa.III | IEAITETIRKMNEIATTIASAVEEQGAATTEIARNVQQAS------------ | *Blastochloris viridis* | Bacteria; Proteobacteria; Alphaproteobacteria; Rhizobiales; Hyphomicrobiaceae; Blastochloris | hypothetical protein | 6 |
| | **WP_084020828.1** | 1.3E-05 | Qc.III | IDHIGRIMEELSTIATAIAAAVEEQAAVTSEIAVNVNQTTDHTHAVVRSIE-- | *Microvirga flocculans* | Bacteria; Proteobacteria; Alphaproteobacteria; Rhizobiales; Methylobacteriaceae; Microvirga | chemotaxis protein | 6 |
| | **WP_085617850.1** | 1.3E-05 | Qc.III | -RRISETIARVAESSTAISAAVEEQHAAIGEISRNVTQAASGTEEVSQRIE-- | *Thalassospira alkalitolerans* | Bacteria; Proteobacteria; Alphaproteobacteria; Rhodospirillales; Rhodospirillaceae; Thalassospira | HAMP domain-containing protein | 6 |
|  | **WP_086465944.1** | 1.3E-05 | Qc.III | -REIGDTISEINDIAANIAAAVEEQGAATQEISRNVQEAAKGTQEV------- | *Oceanibaculum nanhaiense* | Bacteria; Proteobacteria; Alphaproteobacteria; Rhodospirillales; Rhodospirillaceae; Oceanibaculum | methyl-accepting chemotaxis protein | 6 |
|  | **WP_092076722.1** | 1.3E-05 | Qc.III | -DQISAGADQISDTSQALSQAATEQASSLEEINSSMTEMASRTKQ-------- | *Desulfuromonas thiophila* | Bacteria; Proteobacteria; Deltaproteobacteria; Desulfuromonadales; Desulfuromonadaceae; Desulfuromonas | DUF3365 domain-containing protein | 6 |
|  | **WP_092085573.1** | 1.3E-05 | Qc.III.c | --AISQVIVQISEISTVIASAVEEQAAATKEVTANINGTTKAIDDA------- | *Bradyrhizobium sp. R5* | Bacteria; Proteobacteria; Alphaproteobacteria; Rhizobiales; Bradyrhizobiaceae; Bradyrhizobium | PAS domain-containing protein | 6 |
|  | **WP_092280328.1** | 1.3E-05 | Qb.I | LQRTKLQLQDTIEQSEVSSEELKASNEEMQAINEELRSATEELETS------- | *Pseudomonas prosekii* | Bacteria; Proteobacteria; Gammaproteobacteria; Pseudomonadales; Pseudomonadaceae; Pseudomonas | PAS domain S-box protein | 6 |
| | **WP_095744056.1** | 1.3E-05 | R.IV | IEQINQAIAQMDQVTQQNAALVEEASATAQSLQEQAEGLVRAAGVFRTGAG-- | *Variovorax boronicumulans* | Bacteria; Proteobacteria; Betaproteobacteria; Burkholderiales; Comamonadaceae; Variovorax | PAS domain S-box protein | 6 |
| | **WP_095946606.1** | 1.3E-05 | R.IV | IEQINQAIAQMDQVTQQNAALVEEASATAQSLQEQAEGLVRAAGVFRTGAG-- | *Variovorax boronicumulans* | Bacteria; Proteobacteria; Betaproteobacteria; Burkholderiales; Comamonadaceae; Variovorax | PAS domain S-box protein | 6 |
| | **WP_097280859.1** | 1.3E-05 | Qc.III.c | --RIRGTIQELDEIAQSIAAAVEQQNAATMDISRGVQRAADGVREAARQVD-- | *Caenispirillum bisanense* | Bacteria; Proteobacteria; Alphaproteobacteria; Rhodospirillales; Rhodospirillaceae; Caenispirillum | methyl-accepting chemotaxis protein | 6 |
|  | **WP_097672462.1** | 1.3E-05 | Qc.III.c | --AISQVIVQISEISTVIASAVEEQAAATKEVTANINGTTKAIDDA------- | *Bradyrhizobium sp. C9* | Bacteria; Proteobacteria; Alphaproteobacteria; Rhizobiales; Bradyrhizobiaceae; Bradyrhizobium | PAS domain-containing protein | 6 |
| | **WP_099792029.1** | 1.3E-05 | R.IV | IEQINQAIAQMDQVTQQNAALVEEASATAQSLQEQAEGLVRAAGVFRTGAG-- | *Variovorax sp. 54* | Bacteria; Proteobacteria; Betaproteobacteria; Burkholderiales; Comamonadaceae; Variovorax | PAS domain S-box protein | 6 |
| | **WP_100236281.1** | 1.3E-05 | Qc.III | IKDISHTIEKLSEISSTIAAAVEEQGAATQEISRNVQQAAAGTHQVSSNITD- | *Bradyrhizobium sp. INPA54B* | Bacteria; Proteobacteria; Alphaproteobacteria; Rhizobiales; Bradyrhizobiaceae; Bradyrhizobium | methyl-accepting chemotaxis protein | 6 |
|  | **WP_100417720.1** | 1.3E-05 | Qa.III | IREIRTTIGSMSEIASDIASAVETQGVMTREIATNVGQAA------------ | *Bradyrhizobium lablabi* | Bacteria; Proteobacteria; Alphaproteobacteria; Rhizobiales; Bradyrhizobiaceae; Bradyrhizobium | methyl-accepting chemotaxis protein | 6 |
|  | **WP_101209013.1** | 1.3E-05 | Qb.I | LQRTKLQLQDTIEQSEVSSEELKASNEEMQAINEELRSATEELETS------- | *Pseudomonas sp. 43NM1* | Bacteria; Proteobacteria; Gammaproteobacteria; Pseudomonadales; Pseudomonadaceae; Pseudomonas | PAS domain S-box protein | 6 |
| | **WP_101489174.1** | 1.3E-05 | R.IV | IEQINQAIAQMDQVTQQNAALVEEASATAQSLQEQAEGLVRAAGVFRTGAG-- | *Variovorax sp. RO1* | Bacteria; Proteobacteria; Betaproteobacteria; Burkholderiales; Comamonadaceae; Variovorax | PAS domain S-box protein | 6 |
|  | **WP_108163374.1** | 1.3E-05 | Qa.III | -QKIQTYMQEINDISTQIATATEEQSQVVSGINENLQSIADYS---------- | *Saccharospirillum sp. MSK14-1* | Bacteria; Proteobacteria; Gammaproteobacteria; Oceanospirillales; Saccharospirillaceae; Saccharospirillum | methyl-accepting chemotaxis protein | 6 |
|  | **WP_109503748.1** | 1.3E-05 | Qb.I | LQRTKLQLQDTIEQSEVSSEELKASNEEMQAINEELRSATEELETS------- | *Pseudomonas prosekii* | Bacteria; Proteobacteria; Gammaproteobacteria; Pseudomonadales; Pseudomonadaceae; Pseudomonas | PAS domain S-box protein | 6 |
|  | **WP_109522414.1** | 1.3E-05 | Qb.I | LQRTKLQLQDTIEQSEVSSEELKASNEEMQAINEELRSATEELETS------- | *Pseudomonas prosekii* | Bacteria; Proteobacteria; Gammaproteobacteria; Pseudomonadales; Pseudomonadaceae; Pseudomonas | PAS domain S-box protein | 6 |
|  | **WP_111433456.1** | 1.3E-05 | Qa.IV | IQQIGSTIKDMNQITVGIASAMEEQNAATDEIARNVEQAA------------ | *Rhodobium orientis* | Bacteria; Proteobacteria; Alphaproteobacteria; Rhizobiales; Rhodobiaceae; Rhodobium | methyl-accepting chemotaxis protein | 6 |
|  | **WP_112251775.1** | 1.3E-05 | Qb.I | LLRTRLLLQETIEQSEVSSEELTASNEEMQAINEELRSASEELETSK------ | *Pseudomonas putida* | Bacteria; Proteobacteria; Gammaproteobacteria; Pseudomonadales; Pseudomonadaceae; Pseudomonas | response regulator | 6 |
|  | **AUW39007.1** | 1.4E-05 | Qa.III | LDEINAAVALISDMATQIATAAEEQTHVTNEITQNVTSIK------------ | *Vibrio parahaemolyticus* | Bacteria; Proteobacteria; Gammaproteobacteria; Vibrionales; Vibrionaceae; Vibrio | methyl-accepting chemotaxis protein | 6 |
|  | **AVF63103.1** | 1.4E-05 | Qa.III | LDEINAAVALISDMATQIATAAEEQTHVTNEITQNVTSIK------------ | *Vibrio alginolyticus* | Bacteria; Proteobacteria; Gammaproteobacteria; Vibrionales; Vibrionaceae; Vibrio | chemotaxis protein | 6 |
|  | **AVF73153.1** | 1.4E-05 | Qa.III | LDEINAAVALISDMATQIATAAEEQTHVTNEITQNVTSIK------------ | *Vibrio alginolyticus* | Bacteria; Proteobacteria; Gammaproteobacteria; Vibrionales; Vibrionaceae; Vibrio | methyl-accepting chemotaxis protein | 6 |
|  | **AVW97761.1** | 1.4E-05 | Qa.III | LDEINAAVALISDMATQIATAAEEQTHVTNEITQNVTSIK------------ | *Vibrio parahaemolyticus* | Bacteria; Proteobacteria; Gammaproteobacteria; Vibrionales; Vibrionaceae; Vibrio | methyl-accepting chemotaxis protein | 6 |
|  | **AWA92224.1** | 1.4E-05 | Qa.III | LDEINAAVALISDMATQIATAAEEQTHVTNEITQNVTSIK------------ | *Vibrio parahaemolyticus* | Bacteria; Proteobacteria; Gammaproteobacteria; Vibrionales; Vibrionaceae; Vibrio | methyl-accepting chemotaxis protein | 6 |
|  | **AWG87114.1** | 1.4E-05 | Qa.III | LDEINAAVALISDMATQIATAAEEQTHVTNEITQNVTSIK------------ | *Vibrio parahaemolyticus* | cellular organisms; Bacteria; Proteobacteria; Gammaproteobacteria; Vibrionales; Vibrionaceae; Vibrio; Vibrio harveyi group | chemotaxis protein | 6 |

| | | | | | | | | |
|---|---|---|---|---|---|---|---|---|
|  | CRH04773.1 | 1.4E-05 | Qc.III | --KISEMVNTIHGNNQNILNEVSEQDRMLGDVGSAMNQVTQQ----------- | *magneto-ovoid bacterium MO-1* | Bacteria | Putative methyl-accepting chemotaxis sensory transducer | 6 |
|  | EGH17618.1 | 1.4E-05 | Qa.III | ---ISQALRNLNSLNTSIASATLQQTHVVDDINQNVTQAAQLSQ--------- | *Pseudomonas savastanoi pv. glycinea str. race 4* | Bacteria; Proteobacteria; Gammaproteobacteria; Pseudomonadales; Pseudomonadaceae; Pseudomonas | methyl-accepting chemotaxis protein partial | 6 |
| | EME70892.1 | 1.4E-05 | Qc.III | ---ISGTISRINELATIIAGAVEEQGAATAEIARNVDQANEGTREVASNIS-- | *Magnetospirillum caucaseum* | Bacteria; Proteobacteria; Alphaproteobacteria; Rhodospirillales; Rhodospirillaceae; Magnetospirillum | methyl-accepting chemotaxis protein | 6 |
|  | GAJ71869.1 | 1.4E-05 | Qa.III | LDEINAAVALISDMATQIATAAEEQTHVTNEITQNVTSIK------------- | *Vibrio sp. JCM 18904* | Bacteria; Proteobacteria; Gammaproteobacteria; Vibrionales; Vibrionaceae; Vibrio | N-acetylglucosamine regulated methyl-accepting chemotaxis protein | 6 |
|  | GAK17058.1 | 1.4E-05 | Qa.III | LDEINAAVALISDMATQIATAAEEQTHVTNEITQNVTSIK------------- | *Vibrio sp. JCM 19053* | Bacteria; Proteobacteria; Gammaproteobacteria; Vibrionales; Vibrionaceae; Vibrio | LOW QUALITY PROTEIN: N-acetylglucosamine regulated methyl-accepting chemotaxis protein | 6 |
| | KJS36172.1 | 1.4E-05 | Qc.III | IEEIVHAVGKMNEVSGSIASAVEEQNAATQEIARNTQQAAAGTQEVSSNIQ-- | *Rhodospirillaceae bacterium BRH_c57* | Bacteria; Proteobacteria; Alphaproteobacteria; Rhodospirillales; Rhodospirillaceae | hypothetical protein VR70_14350 | 6 |
|  | KOE86426.1 | 1.4E-05 | Qa.III | LDEINAAVALISDMATQIATAAEEQTHVTNEITQNVTSIK------------- | *Vibrio parahaemolyticus* | Bacteria; Proteobacteria; Gammaproteobacteria; Vibrionales; Vibrionaceae; Vibrio | chemotaxis protein | 6 |
|  | KOY35435.1 | 1.4E-05 | Qa.III | LDEINAAVALISDMATQIATAAEEQTHVTNEITQNVTSIK------------- | *Vibrio parahaemolyticus* | Bacteria; Proteobacteria; Gammaproteobacteria; Vibrionales; Vibrionaceae; Vibrio | chemotaxis protein | 6 |
|  | KOY45348.1 | 1.4E-05 | Qa.III | LDEINAAVALISDMATQIATAAEEQTHVTNEITQNVTSIK------------- | *Vibrio parahaemolyticus* | Bacteria; Proteobacteria; Gammaproteobacteria; Vibrionales; Vibrionaceae; Vibrio | chemotaxis protein | 6 |
|  | OCS02383.1 | 1.4E-05 | Qa.IV | --EIEANVNILSQGINEMSQSINEQTEAINQINEAVATVDEQTKQ-NV----- | *Campylobacter fetus subsp. testudinum* | Bacteria; Proteobacteria; Epsilonproteobacteria; Campylobacterales; Campylobacteraceae; Campylobacter | chemotaxis protein partial | 6 |
|  | OTW24054.1 | 1.4E-05 | Qa.III | LDEINAAVALISDMATQIATAAEEQTHVTNEITQNVTSIK------------- | *Vibrio parahaemolyticus* | Bacteria; Proteobacteria; Gammaproteobacteria; Vibrionales; Vibrionaceae; Vibrio | chemotaxis protein | 6 |
|  | OUD75410.1 | 1.4E-05 | Qa.III | LDEINAAVALISDMATQIATAAEEQTHVTNEITQNVTSIK------------- | *Vibrio parahaemolyticus* | Bacteria; Proteobacteria; Gammaproteobacteria; Vibrionales; Vibrionaceae; Vibrio | chemotaxis protein | 6 |
|  | OUJ23541.1 | 1.4E-05 | Qa.III | LDEINAAVALISDMATQIATAAEEQTHVTNEITQNVTSIK------------- | *Vibrio parahaemolyticus* | Bacteria; Proteobacteria; Gammaproteobacteria; Vibrionales; Vibrionaceae; Vibrio | chemotaxis protein | 6 |
|  | OUJ32622.1 | 1.4E-05 | Qa.III | LDEINAAVALISDMATQIATAAEEQTHVTNEITQNVTSIK------------- | *Vibrio parahaemolyticus* | Bacteria; Proteobacteria; Gammaproteobacteria; Vibrionales; Vibrionaceae; Vibrio | chemotaxis protein | 6 |
|  | OUR77762.1 | 1.4E-05 | Qa.III | -EGIRKTIAEMNEIATMIASAVEEQGAATSEISRNIQEAATGTQS-------- | *Alphaproteobacteria bacterium 46_93_T64* | Bacteria; Proteobacteria; Alphaproteobacteria | hypothetical protein A9Q83_09595 | 6 |
|  | OXD28369.1 | 1.4E-05 | Qa.III | LDEINAAVALISDMATQIATAAEEQTHVTNEITQNVTSIK------------- | *Vibrio parahaemolyticus* | Bacteria; Proteobacteria; Gammaproteobacteria; Vibrionales; Vibrionaceae; Vibrio | methyl-accepting chemotaxis protein | 6 |
|  | OXD38341.1 | 1.4E-05 | Qa.III | LDEINAAVALISDMATQIATAAEEQTHVTNEITQNVTSIK------------- | *Vibrio parahaemolyticus* | Bacteria; Proteobacteria; Gammaproteobacteria; Vibrionales; Vibrionaceae; Vibrio | methyl-accepting chemotaxis protein | 6 |
|  | OXD51482.1 | 1.4E-05 | Qa.III | LDEINAAVALISDMATQIATAAEEQTHVTNEITQNVTSIK------------- | *Vibrio parahaemolyticus* | Bacteria; Proteobacteria; Gammaproteobacteria; Vibrionales; Vibrionaceae; Vibrio | methyl-accepting chemotaxis protein | 6 |
| | OYU86373.1 | 1.4E-05 | Qa.IV | IKEIGATITLISEVSSAIAAAVEEQGAATQEIARNVQQSAQLSSQVAHEV--- | *Bradyrhizobiaceae bacterium PARB1* | Bacteria; Proteobacteria; Alphaproteobacteria; Rhizobiales; Bradyrhizobiaceae | methyl-accepting chemotaxis protein | 6 |
|  | OYU91474.1 | 1.4E-05 | Qc.III | IKMIGGIIGEVNEVATAIAAAVEEQGAATQEITRSTQQAAEGTRNVS------ | *Bradyrhizobiaceae bacterium PARB1* | Bacteria; Proteobacteria; Alphaproteobacteria; Rhizobiales; Bradyrhizobiaceae | methyl-accepting chemotaxis protein | 6 |
|  | PID71773.1 | 1.4E-05 | Qa.IV | -------IVDMDQIVETIATTVEEQTATTREIAENVQQASTGISEVNENVA-- | *Desulfobulbus propionicus* | Bacteria; Proteobacteria; Deltaproteobacteria; Desulfobacterales; Desulfobulbaceae; Desulfobulbus | chemotaxis protein | 6 |
|  | PIO90621.1 | 1.4E-05 | Qc.III | ---VGDVIGKMNEINRMISAAADEQNQATREIAQSAQYASEATHNVTE----- | *Zetaproteobacteria bacterium CG23_combo_of_CG06-09_8_20_14_all_59_86* | Bacteria; Proteobacteria; Zetaproteobacteria | hypothetical protein COX56_02410 | 6 |
| | PIU05770.1 | 1.4E-05 | Qc.III | ---VVSVIQRLARIGGDIAAAVEEQGAATGEIARTTARTSEGTRAVSSHIA-- | *Methylobacterium sp. CG09_land_8_20_14_0_10_71_15* | Bacteria; Proteobacteria; Alphaproteobacteria; Rhizobiales; Methylobacteriaceae; Methylobacterium | chemotaxis protein | 6 |
|  | PLX38652.1 | 1.4E-05 | Qc.III | -----EAVAQLRDSVQGVRREVDHQTKMAQEATRTAEHSNSQVRSLM------ | *Rhizobiales bacterium* | Bacteria; Proteobacteria; Alphaproteobacteria; Rhizobiales | hypothetical protein C0606_10750 | 6 |
|  | PMT82540.1 | 1.4E-05 | Qa.III | LDEINAAVALISDMATQIATAAEEQTHVTNEITQNVTSIK------------- | *Vibrio parahaemolyticus* | Bacteria; Proteobacteria; Gammaproteobacteria; Vibrionales; Vibrionaceae; Vibrio | methyl-accepting chemotaxis protein | 6 |
|  | PNM80051.1 | 1.4E-05 | Qa.III | LDEINAAVALISDMATQIATAAEEQTHVTNEITQNVTSIK------------- | *Vibrio parahaemolyticus* | Bacteria; Proteobacteria; Gammaproteobacteria; Vibrionales; Vibrionaceae; Vibrio | chemotaxis protein | 6 |
|  | PNP28290.1 | 1.4E-05 | Qa.III | LDEINAAVALISDMATQIATAAEEQTHVTNEITQNVTSIK------------- | *Vibrio alginolyticus* | Bacteria; Proteobacteria; Gammaproteobacteria; Vibrionales; Vibrionaceae; Vibrio | methyl-accepting chemotaxis protein | 6 |
|  | PXB18904.1 | 1.4E-05 | Qa.III | LDEINAAVALISDMATQIATAAEEQTHVTNEITQNVTSIK------------- | *Vibrio parahaemolyticus* | Bacteria; Proteobacteria; Gammaproteobacteria; Vibrionales; Vibrionaceae; Vibrio | methyl-accepting chemotaxis protein | 6 |
|  | PXC15205.1 | 1.4E-05 | Qa.III | LDEINAAVALISDMATQIATAAEEQTHVTNEITQNVTSIK------------- | *Vibrio parahaemolyticus* | Bacteria; Proteobacteria; Gammaproteobacteria; Vibrionales; Vibrionaceae; Vibrio | methyl-accepting chemotaxis protein | 6 |
|  | PZN41190.1 | 1.4E-05 | Qa.III | IDSITETIRQIDEVSTTIASAVDQQGATTNEIARNVSQAAQGTQ--------- | *Proteobacteria bacterium* | Bacteria; Proteobacteria | chemotaxis protein | 6 |
|  | PZR15135.1 | 1.4E-05 | Qa.IV | IEEITRTIGEVSEATVTIAAAVEEQTAATAEIARTVQQTA------------- | *Azospirillum brasilense* | Bacteria; Proteobacteria; Alphaproteobacteria; Rhodospirillales; Rhodospirillaceae; Azospirillum | hypothetical protein DI532_07065 | 6 |
| | RCK48451.1 | 1.4E-05 | Qc.III | IDSIGTTITEISEIASAIASAVEQQGAATDEINRNVQEASAGTGEVTVNIH-- | *Thalassospira xiamenensis* | Bacteria; Proteobacteria; Alphaproteobacteria; Rhodospirillales; Rhodospirillaceae; Thalassospira | chemotaxis protein | 6 |
|  | SDD93252.1 | 1.4E-05 | Qa.III | IQAIVRVIAEINEITTGVSAAVEQQAAATGEISQNAQQAAHSTQDASQN---- | *Rhodospira trueperi* | Bacteria; Proteobacteria; Alphaproteobacteria; Rhodospirillales; Rhodospirillaceae; Rhodospira | Methyl-accepting chemotaxis protein | 6 |
|  | SHO46430.1 | 1.4E-05 | Qa.III | IEKISTVIQEINEMIATIATAVEEQSATTVEIAGNISQAA------------- | *Desulfopila aestuarii DSM 18488* | Bacteria; Proteobacteria; Deltaproteobacteria; Desulfobacterales; Desulfobulbaceae; Desulfopila | methyl-accepting chemotaxis protein | 6 |
|  | SQA41875.1 | 1.4E-05 | Qa.III | LDEINAAVALISDMATQIATAAEEQTHVTNEITQNVTSIK------------- | *Vibrio alginolyticus* | Bacteria; Proteobacteria; Gammaproteobacteria; Vibrionales; Vibrionaceae; Vibrio | putative methyl-accepting chemotaxis protein | 6 |
|  | WP_005374718.1 | 1.4E-05 | Qa.III | LDEINAAVALISDMATQIATAAEEQTHVTNEITQNVTSIK------------- | *Vibrio alginolyticus* | Bacteria; Proteobacteria; Gammaproteobacteria; Vibrionales; Vibrionaceae; Vibrio | methyl-accepting chemotaxis protein | 6 |
|  | WP_005383451.1 | 1.4E-05 | Qa.III | LDEINAAVALISDMATQIATAAEEQTHVTNEITQNVTSIK------------- | *Vibrio* | Bacteria; Proteobacteria; Gammaproteobacteria; Vibrionales; Vibrionaceae | MULTISPECIES: methyl-accepting chemotaxis protein | 6 |
|  | WP_005395590.1 | 1.4E-05 | Qa.III | LDEINAAVALISDMATQIATAAEEQTHVTNEITQNVTSIK------------- | *Vibrio alginolyticus* | Bacteria; Proteobacteria; Gammaproteobacteria; Vibrionales; Vibrionaceae; Vibrio | methyl-accepting chemotaxis protein | 6 |
|  | WP_005500014.1 | 1.4E-05 | Qa.III | LDEINAAVALISDMATQIATAAEEQTHVTNEITQNVTSIK------------- | *Vibrio parahaemolyticus* | Bacteria; Proteobacteria; Gammaproteobacteria; Vibrionales; Vibrionaceae; Vibrio | methyl-accepting chemotaxis protein | 6 |
|  | WP_009194687.1 | 1.4E-05 | Qc.III | -------THRLLDESQKMSRELQMQQEVLRQNAEEMEATQEELKRTNHRLE-- | *Cesiribacter andamanensis* | Bacteria; Bacteroidetes; Cytophagia; Cytophagales; Flammeovirgaceae; Cesiribacter | PAS domain S-box protein | 6 |
|  | WP_010648050.1 | 1.4E-05 | Qb.I | ---------QSVLLIGETNDVIQTQAKSVNDTEQAFTDIQSAIHTNNQLIIE- | *Oceanobacillus massiliensis* | Bacteria; Firmicutes; Bacilli; Bacillales; Bacillaceae; Oceanobacillus | methyl-accepting chemotaxis protein | 6 |
|  | WP_011217234.1 | 1.4E-05 | Qc.III | LDKIVEQVQKINDMNFHIASAAEQQTAVADEMNQNLTNVR------------- | *Photobacterium profundum* | Bacteria; Proteobacteria; Gammaproteobacteria; Vibrionales; Vibrionaceae; Photobacterium | methyl-accepting chemotaxis protein | 6 |
|  | WP_012030405.1 | 1.4E-05 | Qa.IV | -----GTIRQISTIASSIATAVEQQGGATQEIARNVQSVAAGTQAAASDI--- | *Bradyrhizobium sp. ORS 278* | Bacteria; Proteobacteria; Alphaproteobacteria; Rhizobiales; Bradyrhizobiaceae; Bradyrhizobium | methyl-accepting chemotaxis protein | 6 |
|  | WP_021131581.1 | 1.4E-05 | Qa.I | IRSISGTIENISELTSAIAAAVEEQGAATSEIARSVEEAA------------- | *Phaeospirillum fulvum* | Bacteria; Proteobacteria; Alphaproteobacteria; Rhodospirillales; Rhodospirillaceae; Phaeospirillum | PAS domain S-box protein | 6 |
|  | WP_023584474.1 | 1.4E-05 | Qa.III | LDEINAAVALISDMATQIATAAEEQTHVTNEITQNVTSIK------------- | *Vibrio parahaemolyticus* | Bacteria; Proteobacteria; Gammaproteobacteria; Vibrionales; Vibrionaceae; Vibrio | methyl-accepting chemotaxis protein | 6 |
|  | WP_025498764.1 | 1.4E-05 | Qa.III | LDEINAAVALISDMATQIATAAEEQTHVTNEITQNVTSIK------------- | *Vibrio parahaemolyticus* | Bacteria; Proteobacteria; Gammaproteobacteria; Vibrionales; Vibrionaceae; Vibrio | methyl-accepting chemotaxis protein | 6 |
|  | WP_025505249.1 | 1.4E-05 | Qa.III | LDEINAAVALISDMATQIATAAEEQTHVTNEITQNVTSIK------------- | *Vibrio parahaemolyticus* | Bacteria; Proteobacteria; Gammaproteobacteria; Vibrionales; Vibrionaceae; Vibrio | methyl-accepting chemotaxis protein | 6 |
|  | WP_025508004.1 | 1.4E-05 | Qa.III | LDEINAAVALISDMATQIATAAEEQTHVTNEITQNVTSIK------------- | *Vibrio parahaemolyticus* | Bacteria; Proteobacteria; Gammaproteobacteria; Vibrionales; Vibrionaceae; Vibrio | methyl-accepting chemotaxis protein | 6 |
|  | WP_025515589.1 | 1.4E-05 | Qa.III | LDEINAAVALISDMATQIATAAEEQTHVTNEITQNVTSIK------------- | *Vibrio parahaemolyticus* | Bacteria; Proteobacteria; Gammaproteobacteria; Vibrionales; Vibrionaceae; Vibrio | methyl-accepting chemotaxis protein | 6 |
|  | WP_025521234.1 | 1.4E-05 | Qa.III | LDEINAAVALISDMATQIATAAEEQTHVTNEITQNVTSIK------------- | *Vibrio parahaemolyticus* | Bacteria; Proteobacteria; Gammaproteobacteria; Vibrionales; Vibrionaceae; Vibrio | methyl-accepting chemotaxis protein | 6 |
|  | WP_025524283.1 | 1.4E-05 | Qa.III | LDEINAAVALISDMATQIATAAEEQTHVTNEITQNVTSIK------------- | *Vibrio parahaemolyticus* | Bacteria; Proteobacteria; Gammaproteobacteria; Vibrionales; Vibrionaceae; Vibrio | methyl-accepting chemotaxis protein | 6 |
|  | WP_025532891.1 | 1.4E-05 | Qa.III | LDEINAAVALISDMATQIATAAEEQTHVTNEITQNVTSIK------------- | *Vibrio parahaemolyticus* | Bacteria; Proteobacteria; Gammaproteobacteria; Vibrionales; Vibrionaceae; Vibrio | methyl-accepting chemotaxis protein | 6 |
|  | WP_025559484.1 | 1.4E-05 | Qa.III | LDEINAAVALISDMATQIATAAEEQTHVTNEITQNVTSIK------------- | *Vibrio parahaemolyticus* | Bacteria; Proteobacteria; Gammaproteobacteria; Vibrionales; Vibrionaceae; Vibrio | methyl-accepting chemotaxis protein | 6 |
|  | WP_025623954.1 | 1.4E-05 | Qa.III | LDEINAAVALISDMATQIATAAEEQTHVTNEITQNVTSIK------------- | *Vibrio parahaemolyticus* | Bacteria; Proteobacteria; Gammaproteobacteria; Vibrionales; Vibrionaceae; Vibrio | methyl-accepting chemotaxis protein | 6 |
|  | WP_025627722.1 | 1.4E-05 | Qa.III | LDEINAAVALISDMATQIATAAEEQTHVTNEITQNVTSIK------------- | *Vibrio parahaemolyticus* | Bacteria; Proteobacteria; Gammaproteobacteria; Vibrionales; Vibrionaceae; Vibrio | methyl-accepting chemotaxis protein | 6 |
|  | WP_025635522.1 | 1.4E-05 | Qa.III | LDEINAAVALISDMATQIATAAEEQTHVTNEITQNVTSIK------------- | *Vibrio parahaemolyticus* | Bacteria; Proteobacteria; Gammaproteobacteria; Vibrionales; Vibrionaceae; Vibrio | methyl-accepting chemotaxis protein | 6 |
|  | WP_025638667.1 | 1.4E-05 | Qa.III | LDEINAAVALISDMATQIATAAEEQTHVTNEITQNVTSIK------------- | *Vibrio parahaemolyticus* | Bacteria; Proteobacteria; Gammaproteobacteria; Vibrionales; Vibrionaceae; Vibrio | methyl-accepting chemotaxis protein | 6 |
|  | WP_025768674.1 | 1.4E-05 | Qa.III | LDEINAAVALISDMATQIATAAEEQTHVTNEITQNVTSIK------------- | *Vibrio harveyi group* | Bacteria; Proteobacteria; Gammaproteobacteria; Vibrionales; Vibrionaceae; Vibrio | MULTISPECIES: methyl-accepting chemotaxis protein | 6 |

| | | | | | | | | |
|---|---|---|---|---|---|---|---|---|
|  | WP_025793649.1 | 1.4E-05 | Qa.III | LDEINAAVALISDMATQIATAAEEQTHVTNEITQNVTSIK------------- | *Vibrio parahaemolyticus* | Bacteria; Proteobacteria; Gammaproteobacteria; Vibrionales; Vibrionaceae; Vibrio | methyl-accepting chemotaxis protein | 6 |
|  | WP_025794015.1 | 1.4E-05 | Qa.III | LDEINAAVALISDMATQIATAAEEQTHVTNEITQNVTSIK------------- | *Vibrio parahaemolyticus* | Bacteria; Proteobacteria; Gammaproteobacteria; Vibrionales; Vibrionaceae; Vibrio | methyl-accepting chemotaxis protein | 6 |
|  | WP_027917371.1 | 1.4E-05 | Qb.I | LQRTRLQLQETIEQSEISSEELTASNEEMQAINEELRSASEELETSK------ | *Pseudomonas* | Bacteria; Proteobacteria; Gammaproteobacteria; Pseudomonadales; Pseudomonadaceae | MULTISPECIES: response regulator | 6 |
| | WP_028134450.1 | 1.4E-05 | Qc.III | --TIGGTIERLSEISSTIAAAVEEQGAATSEISRNVQNAAAGTTQVSANI--- | *Bradyrhizobium japonicum* | Bacteria; Proteobacteria; Alphaproteobacteria; Rhizobiales; Bradyrhizobiaceae; Bradyrhizobium | methyl-accepting chemotaxis protein | 6 |
|  | WP_028318941.1 | 1.4E-05 | Qa.III | IKHISQVIHNINEIVTVISTAVDEQSSATQEIANNIAQAS------------- | *Desulfobulbus elongatus* | Bacteria; Proteobacteria; Deltaproteobacteria; Desulfobacterales; Desulfobulbaceae; Desulfobulbus | methyl-accepting chemotaxis protein | 6 |
|  | WP_029826758.1 | 1.4E-05 | Qa.III | LDEINAAVALISDMATQIATAAEEQTHVTNEITQNVTSIK------------- | *Vibrio parahaemolyticus* | Bacteria; Proteobacteria; Gammaproteobacteria; Vibrionales; Vibrionaceae; Vibrio | methyl-accepting chemotaxis protein | 6 |
|  | WP_029833637.1 | 1.4E-05 | Qa.III | LDEINAAVALISDMATQIATAAEEQTHVTNEITQNVTSIK------------- | *Vibrio parahaemolyticus* | Bacteria; Proteobacteria; Gammaproteobacteria; Vibrionales; Vibrionaceae; Vibrio | methyl-accepting chemotaxis protein | 6 |
|  | WP_029841129.1 | 1.4E-05 | Qa.III | LDEINAAVALISDMATQIATAAEEQTHVTNEITQNVTSIK------------- | *Vibrio parahaemolyticus* | Bacteria; Proteobacteria; Gammaproteobacteria; Vibrionales; Vibrionaceae; Vibrio | methyl-accepting chemotaxis protein | 6 |
|  | WP_029852974.1 | 1.4E-05 | Qa.III | LDEINAAVALISDMATQIATAAEEQTHVTNEITQNVTSIK------------- | *Vibrio parahaemolyticus* | Bacteria; Proteobacteria; Gammaproteobacteria; Vibrionales; Vibrionaceae; Vibrio | methyl-accepting chemotaxis protein | 6 |
|  | WP_029853996.1 | 1.4E-05 | Qa.III | LDEINAAVALISDMATQIATAAEEQTHVTNEITQNVTSIK------------- | *Vibrio parahaemolyticus* | Bacteria; Proteobacteria; Gammaproteobacteria; Vibrionales; Vibrionaceae; Vibrio | methyl-accepting chemotaxis protein | 6 |
|  | WP_029862580.1 | 1.4E-05 | Qa.III | LDEINAAVALISDMATQIATAAEEQTHVTNEITQNVTSIK------------- | *Vibrio parahaemolyticus* | Bacteria; Proteobacteria; Gammaproteobacteria; Vibrionales; Vibrionaceae; Vibrio | methyl-accepting chemotaxis protein | 6 |
|  | WP_029864441.1 | 1.4E-05 | Qa.III | LDEINAAVALISDMATQIATAAEEQTHVTNEITQNVTSIK------------- | *Vibrio parahaemolyticus* | Bacteria; Proteobacteria; Gammaproteobacteria; Vibrionales; Vibrionaceae; Vibrio | methyl-accepting chemotaxis protein | 6 |
|  | WP_031812644.1 | 1.4E-05 | Qa.III | LDEINAAVALISDMATQIATAAEEQTHVTNEITQNVTSIK------------- | *Vibrio parahaemolyticus* | Bacteria; Proteobacteria; Gammaproteobacteria; Vibrionales; Vibrionaceae; Vibrio | methyl-accepting chemotaxis protein | 6 |
|  | WP_032611325.1 | 1.4E-05 | R.IV | IQEINRAVTHLDSMVQQNAELVVQSAAAASALQGQAGELAETAGHFR------ | *Leclercia adecarboxylata* | Bacteria; Proteobacteria; Gammaproteobacteria; Enterobacterales; Enterobacteriaceae; Leclercia | methyl-accepting chemotaxis protein | 6 |
|  | WP_042764455.1 | 1.4E-05 | Qa.III | LDEINAAVALISDMATQIATAAEEQTHVTNEITQNVTSIK------------- | *Vibrio parahaemolyticus* | Bacteria; Proteobacteria; Gammaproteobacteria; Vibrionales; Vibrionaceae; Vibrio | methyl-accepting chemotaxis protein | 6 |
|  | WP_042779726.1 | 1.4E-05 | Qa.III | LDEINAAVALISDMATQIATAAEEQTHVTNEITQNVTSIK------------- | *Vibrio parahaemolyticus* | Bacteria; Proteobacteria; Gammaproteobacteria; Vibrionales; Vibrionaceae; Vibrio | methyl-accepting chemotaxis protein | 6 |
|  | WP_043042543.1 | 1.4E-05 | Qa.III | LDEINAAVALISDMATQIATAAEEQTHVTNEITQNVTSIK------------- | *Vibrio parahaemolyticus* | Bacteria; Proteobacteria; Gammaproteobacteria; Vibrionales; Vibrionaceae; Vibrio | methyl-accepting chemotaxis protein | 6 |
|  | WP_045545569.1 | 1.4E-05 | Qa.III | LDEINAAVALISDMATQIATAAEEQTHVTNEITQNVTSIK------------- | *Vibrio* | Bacteria; Proteobacteria; Gammaproteobacteria; Vibrionales; Vibrionaceae | MULTISPECIES: methyl-accepting chemotaxis protein | 6 |
|  | WP_045607726.1 | 1.4E-05 | Qa.III | LDEINAAVALISDMATQIATAAEEQTHVTNEITQNVTSIK------------- | *Vibrio parahaemolyticus* | Bacteria; Proteobacteria; Gammaproteobacteria; Vibrionales; Vibrionaceae; Vibrio | methyl-accepting chemotaxis protein | 6 |
|  | WP_045627212.1 | 1.4E-05 | Qa.III | LDEINAAVALISDMATQIATAAEEQTHVTNEITQNVTSIK------------- | *Vibrio parahaemolyticus* | Bacteria; Proteobacteria; Gammaproteobacteria; Vibrionales; Vibrionaceae; Vibrio | methyl-accepting chemotaxis protein | 6 |
|  | WP_046875685.1 | 1.4E-05 | Qa.III | LDEINAAVALISDMATQIATAAEEQTHVTNEITQNVTSIK------------- | *Vibrio alginolyticus* | Bacteria; Proteobacteria; Gammaproteobacteria; Vibrionales; Vibrionaceae; Vibrio | methyl-accepting chemotaxis protein | 6 |
|  | WP_047009659.1 | 1.4E-05 | Qa.III | LDEINAAVALISDMATQIATAAEEQTHVTNEITQNVTSIK------------- | *Vibrio alginolyticus* | Bacteria; Proteobacteria; Gammaproteobacteria; Vibrionales; Vibrionaceae; Vibrio | methyl-accepting chemotaxis protein | 6 |
|  | WP_047706277.1 | 1.4E-05 | Qa.III | LDEINAAVALISDMATQIATAAEEQTHVTNEITQNVTSIK------------- | *Vibrio parahaemolyticus* | Bacteria; Proteobacteria; Gammaproteobacteria; Vibrionales; Vibrionaceae; Vibrio | methyl-accepting chemotaxis protein | 6 |
|  | WP_047731556.1 | 1.4E-05 | Qa.III | LDEINAAVALISDMATQIATAAEEQTHVTNEITQNVTSIK------------- | *Vibrio parahaemolyticus* | Bacteria; Proteobacteria; Gammaproteobacteria; Vibrionales; Vibrionaceae; Vibrio | methyl-accepting chemotaxis protein | 6 |
|  | WP_048625453.1 | 1.4E-05 | Qa.III | LDEINAAVALISDMATQIATAAEEQTHVTNEITQNVTSIK------------- | *Vibrio diabolicus* | Bacteria; Proteobacteria; Gammaproteobacteria; Vibrionales; Vibrionaceae; Vibrio | methyl-accepting chemotaxis protein | 6 |
|  | WP_053048381.1 | 1.4E-05 | Qa.III | LDEINAAVALISDMATQIATAAEEQTHVTNEITQNVTSIK------------- | *Vibrio parahaemolyticus* | Bacteria; Proteobacteria; Gammaproteobacteria; Vibrionales; Vibrionaceae; Vibrio | methyl-accepting chemotaxis protein | 6 |
|  | WP_053302365.1 | 1.4E-05 | Qa.III | LDEINAAVALISDMATQIATAAEEQTHVTNEITQNVTSIK------------- | *Vibrio parahaemolyticus* | Bacteria; Proteobacteria; Gammaproteobacteria; Vibrionales; Vibrionaceae; Vibrio | methyl-accepting chemotaxis protein | 6 |
|  | WP_053316119.1 | 1.4E-05 | Qa.III | LDEINAAVALISDMATQIATAAEEQTHVTNEITQNVTSIK------------- | *Vibrio parahaemolyticus* | Bacteria; Proteobacteria; Gammaproteobacteria; Vibrionales; Vibrionaceae; Vibrio | methyl-accepting chemotaxis protein | 6 |
|  | WP_053318114.1 | 1.4E-05 | Qa.III | LDEINAAVALISDMATQIATAAEEQTHVTNEITQNVTSIK------------- | *Vibrio parahaemolyticus* | Bacteria; Proteobacteria; Gammaproteobacteria; Vibrionales; Vibrionaceae; Vibrio | methyl-accepting chemotaxis protein | 6 |
|  | WP_053340306.1 | 1.4E-05 | Qa.III | LDEINAAVALISDMATQIATAAEEQTHVTNEITQNVTSIK------------- | *Vibrio* | Bacteria; Proteobacteria; Gammaproteobacteria; Vibrionales; Vibrionaceae | MULTISPECIES: methyl-accepting chemotaxis protein | 6 |
|  | WP_053343482.1 | 1.4E-05 | Qa.III | LDEINAAVALISDMATQIATAAEEQTHVTNEITQNVTSIK------------- | *Vibrio parahaemolyticus* | Bacteria; Proteobacteria; Gammaproteobacteria; Vibrionales; Vibrionaceae; Vibrio | methyl-accepting chemotaxis protein | 6 |
|  | WP_053350167.1 | 1.4E-05 | Qa.III | LDEINAAVALISDMATQIATAAEEQTHVTNEITQNVTSIK------------- | *Vibrio alginolyticus* | Bacteria; Proteobacteria; Gammaproteobacteria; Vibrionales; Vibrionaceae; Vibrio | methyl-accepting chemotaxis protein | 6 |
|  | WP_053815227.1 | 1.4E-05 | Qa.III | LDEINAAVALISDMATQIATAAEEQTHVTNEITQNVTSIK------------- | *Vibrio parahaemolyticus* | Bacteria; Proteobacteria; Gammaproteobacteria; Vibrionales; Vibrionaceae; Vibrio | methyl-accepting chemotaxis protein | 6 |
|  | WP_054030448.1 | 1.4E-05 | Qa.III | IDKISKVIHSVNDLVSTIATAVEEQSVTTKEIAGNVAQASQGIQEVN------ | *Desulfatitalea tepidiphila* | Bacteria; Proteobacteria; Deltaproteobacteria; Desulfobacterales; Desulfobacteraceae; Desulfatitalea | methyl-accepting chemotaxis protein | 6 |
|  | WP_054575394.1 | 1.4E-05 | Qa.III | LDEINAAVALISDMATQIATAAEEQTHVTNEITQNVTSIK------------- | *Vibrio alginolyticus* | Bacteria; Proteobacteria; Gammaproteobacteria; Vibrionales; Vibrionaceae; Vibrio | methyl-accepting chemotaxis protein | 6 |
|  | WP_054576558.1 | 1.4E-05 | Qa.III | LDEINAAVALISDMATQIATAAEEQTHVTNEITQNVTSIK------------- | *Vibrio harveyi group* | Bacteria; Proteobacteria; Gammaproteobacteria; Vibrionales; Vibrionaceae; Vibrio | MULTISPECIES: methyl-accepting chemotaxis protein | 6 |
|  | WP_054578607.1 | 1.4E-05 | Qa.III | LDEINAAVALISDMATQIATAAEEQTHVTNEITQNVTSIK------------- | *Vibrio alginolyticus* | Bacteria; Proteobacteria; Gammaproteobacteria; Vibrionales; Vibrionaceae; Vibrio | methyl-accepting chemotaxis protein | 6 |
| | WP_056214488.1 | 1.4E-05 | Qc.III | IERIAGIIGRIDESAISIGSAVEEQSATATEIRRSVEEASGRTETVSSSV--- | *Methylobacterium sp. Leaf94* | Bacteria; Proteobacteria; Alphaproteobacteria; Rhizobiales; Methylobacteriaceae; Methylobacterium | chemotaxis protein | 6 |
| | WP_056275190.1 | 1.4E-05 | Qc.III | IERIAGIIGRIDESAISIGSAVEEQSATATEIRRSVEEASGRTETVSSSV--- | *Methylobacterium sp. Leaf88* | Bacteria; Proteobacteria; Alphaproteobacteria; Rhizobiales; Methylobacteriaceae; Methylobacterium | chemotaxis protein | 6 |
| | WP_056352389.1 | 1.4E-05 | Qc.III | IERIAGIIGRIDESAISIGSAVEEQSATATEIRRSVEEASGRTETVSSSV--- | *Methylobacterium sp. Leaf89* | Bacteria; Proteobacteria; Alphaproteobacteria; Rhizobiales; Methylobacteriaceae; Methylobacterium | chemotaxis protein | 6 |
| | WP_056381203.1 | 1.4E-05 | Qc.III | IERIAGIIGRIDESAISIGSAVEEQSATATEIRRSVEEASGRTETVSSSV--- | *Methylobacterium sp. Leaf465* | Bacteria; Proteobacteria; Alphaproteobacteria; Rhizobiales; Methylobacteriaceae; Methylobacterium | chemotaxis protein | 6 |
|  | WP_059307692.1 | 1.4E-05 | R.IV | IQEINRAVTHLDSMVQQNAELVVQSAAAASALQGQAGELAETAGHFR------ | *Leclercia adecarboxylata* | Bacteria; Proteobacteria; Gammaproteobacteria; Enterobacterales; Enterobacteriaceae; Leclercia | methyl-accepting chemotaxis protein | 6 |
|  | WP_060485470.1 | 1.4E-05 | Qb.I | LQRTRLQLQETIEQSEISSEELTASNEEMQAINEELRSASEELETSK------ | *Pseudomonas sp. NBRC 111123* | Bacteria; Proteobacteria; Gammaproteobacteria; Pseudomonadales; Pseudomonadaceae; Pseudomonas | response regulator | 6 |
|  | WP_062491430.1 | 1.4E-05 | Qc.III | -EQVAASAGELATGANEISRDVGDMTESIREIEEQVASYADSTREMNRR---- | *Paenibacillus sp. 32O-W* | Bacteria; Firmicutes; Bacilli; Bacillales; Paenibacillaceae; Paenibacillus | methyl-accepting chemotaxis protein | 6 |
|  | WP_062855397.1 | 1.4E-05 | Qa.III | LDEINAAVALISDMATQIATAAEEQTHVTNEITQNVTSIK------------- | *Vibrio parahaemolyticus* | Bacteria; Proteobacteria; Gammaproteobacteria; Vibrionales; Vibrionaceae; Vibrio | methyl-accepting chemotaxis protein | 6 |
|  | WP_062907449.1 | 1.4E-05 | Qa.III | LDEINAAVALISDMATQIATAAEEQTHVTNEITQNVTSIK------------- | *Vibrio parahaemolyticus* | Bacteria; Proteobacteria; Gammaproteobacteria; Vibrionales; Vibrionaceae; Vibrio | methyl-accepting chemotaxis protein | 6 |
|  | WP_064354987.1 | 1.4E-05 | Qa.III | LDEINAAVALISDMATQIATAAEEQTHVTNEITQNVTSIK------------- | *Vibrio alginolyticus* | Bacteria; Proteobacteria; Gammaproteobacteria; Vibrionales; Vibrionaceae; Vibrio | methyl-accepting chemotaxis protein | 6 |
|  | WP_065646140.1 | 1.4E-05 | Qa.III | LDEINAAVALISDMATQIATAAEEQTHVTNEITQNVTSIK------------- | *Vibrio alginolyticus* | Bacteria; Proteobacteria; Gammaproteobacteria; Vibrionales; Vibrionaceae; Vibrio | methyl-accepting chemotaxis protein | 6 |
|  | WP_065729702.1 | 1.4E-05 | Qa.III | IEAIGKTIGEISEIATTIASAIEEQGAATQEIARNVQQAA------------- | *Bradyrhizobium icense* | Bacteria; Proteobacteria; Alphaproteobacteria; Rhizobiales; Bradyrhizobiaceae; Bradyrhizobium | methyl-accepting chemotaxis protein | 6 |
|  | WP_065764635.1 | 1.4E-05 | Qa.III | LDEINAAVALISDMATQIATAAEEQTHVTNEITQNVTSIK------------- | *Vibrio parahaemolyticus* | Bacteria; Proteobacteria; Gammaproteobacteria; Vibrionales; Vibrionaceae; Vibrio | methyl-accepting chemotaxis protein | 6 |
|  | WP_065769536.1 | 1.4E-05 | Qa.III | LDEINAAVALISDMATQIATAAEEQTHVTNEITQNVTSIK------------- | *Vibrio parahaemolyticus* | Bacteria; Proteobacteria; Gammaproteobacteria; Vibrionales; Vibrionaceae; Vibrio | methyl-accepting chemotaxis protein | 6 |
|  | WP_068865210.1 | 1.4E-05 | Qa.III | LDEINAAVALISDMATQIATAAEEQTHVTNEITQNVTSIK------------- | *Vibrio parahaemolyticus* | Bacteria; Proteobacteria; Gammaproteobacteria; Vibrionales; Vibrionaceae; Vibrio | methyl-accepting chemotaxis protein | 6 |
|  | WP_069487925.1 | 1.4E-05 | Qa.III | LDEINAAVALISDMATQIATAAEEQTHVTNEITQNVTSIK------------- | *Vibrio parahaemolyticus* | Bacteria; Proteobacteria; Gammaproteobacteria; Vibrionales; Vibrionaceae; Vibrio | methyl-accepting chemotaxis protein | 6 |
|  | WP_069489707.1 | 1.4E-05 | Qa.III | LDEINAAVALISDMATQIATAAEEQTHVTNEITQNVTSIK------------- | *Vibrio parahaemolyticus* | Bacteria; Proteobacteria; Gammaproteobacteria; Vibrionales; Vibrionaceae; Vibrio | methyl-accepting chemotaxis protein | 6 |
|  | WP_069493264.1 | 1.4E-05 | Qa.III | LDEINAAVALISDMATQIATAAEEQTHVTNEITQNVTSIK------------- | *Vibrio parahaemolyticus* | Bacteria; Proteobacteria; Gammaproteobacteria; Vibrionales; Vibrionaceae; Vibrio | methyl-accepting chemotaxis protein | 6 |
|  | WP_069496551.1 | 1.4E-05 | Qa.III | LDEINAAVALISDMATQIATAAEEQTHVTNEITQNVTSIK------------- | *Vibrio parahaemolyticus* | Bacteria; Proteobacteria; Gammaproteobacteria; Vibrionales; Vibrionaceae; Vibrio | methyl-accepting chemotaxis protein | 6 |
|  | WP_069498168.1 | 1.4E-05 | Qa.III | LDEINAAVALISDMATQIATAAEEQTHVTNEITQNVTSIK------------- | *Vibrio parahaemolyticus* | Bacteria; Proteobacteria; Gammaproteobacteria; Vibrionales; Vibrionaceae; Vibrio | methyl-accepting chemotaxis protein | 6 |
|  | WP_069502658.1 | 1.4E-05 | Qa.III | LDEINAAVALISDMATQIATAAEEQTHVTNEITQNVTSIK------------- | *Vibrio parahaemolyticus* | Bacteria; Proteobacteria; Gammaproteobacteria; Vibrionales; Vibrionaceae; Vibrio | methyl-accepting chemotaxis protein | 6 |
|  | WP_069505528.1 | 1.4E-05 | Qa.III | LDEINAAVALISDMATQIATAAEEQTHVTNEITQNVTSIK------------- | *Vibrio parahaemolyticus* | Bacteria; Proteobacteria; Gammaproteobacteria; Vibrionales; Vibrionaceae; Vibrio | methyl-accepting chemotaxis protein | 6 |
|  | WP_069515325.1 | 1.4E-05 | Qa.III | LDEINAAVALISDMATQIATAAEEQTHVTNEITQNVTSIK------------- | *Vibrio parahaemolyticus* | Bacteria; Proteobacteria; Gammaproteobacteria; Vibrionales; Vibrionaceae; Vibrio | methyl-accepting chemotaxis protein | 6 |
|  | WP_069534251.1 | 1.4E-05 | Qa.III | LDEINAAVALISDMATQIATAAEEQTHVTNEITQNVTSIK------------- | *Vibrio parahaemolyticus* | Bacteria; Proteobacteria; Gammaproteobacteria; Vibrionales; Vibrionaceae; Vibrio | methyl-accepting chemotaxis protein | 6 |

| | | | | | | | |
|---|---|---|---|---|---|---|---|
| WP_069536927.1 | 1.4E-05 | Qa.III | LDEINAAVALISDMATQIATAAEEQTHVTNEITQNVTSIK------------- | *Vibrio parahaemolyticus* | Bacteria; Proteobacteria; Gammaproteobacteria; Vibrionales; Vibrionaceae; Vibrio | methyl-accepting chemotaxis protein | 6 |
| WP_069539342.1 | 1.4E-05 | Qa.III | LDEINAAVALISDMATQIATAAEEQTHVTNEITQNVTSIK------------- | *Vibrio parahaemolyticus* | Bacteria; Proteobacteria; Gammaproteobacteria; Vibrionales; Vibrionaceae; Vibrio | methyl-accepting chemotaxis protein | 6 |
| WP_069543878.1 | 1.4E-05 | Qa.III | LDEINAAVALISDMATQIATAAEEQTHVTNEITQNVTSIK------------- | *Vibrio parahaemolyticus* | Bacteria; Proteobacteria; Gammaproteobacteria; Vibrionales; Vibrionaceae; Vibrio | methyl-accepting chemotaxis protein | 6 |
| WP_069549233.1 | 1.4E-05 | Qa.III | LDEINAAVALISDMATQIATAAEEQTHVTNEITQNVTSIK------------- | *Vibrio parahaemolyticus* | Bacteria; Proteobacteria; Gammaproteobacteria; Vibrionales; Vibrionaceae; Vibrio | methyl-accepting chemotaxis protein | 6 |
| WP_071653048.1 | 1.4E-05 | Qa.III | LDEINAAVALISDMATQIATAAEEQTHVTNEITQNVTSIK------------- | *Vibrio parahaemolyticus* | Bacteria; Proteobacteria; Gammaproteobacteria; Vibrionales; Vibrionaceae; Vibrio | methyl-accepting chemotaxis protein | 6 |
| WP_077227096.1 | 1.4E-05 | R.IV | IQEINRAVTHLDSMVQQNAELVVQSAAAASALQGQAGELAETAGHFR------ | *Leclercia adecarboxylata* | Bacteria; Proteobacteria; Gammaproteobacteria; Enterobacterales; Enterobacteriaceae; Leclercia | methyl-accepting chemotaxis protein | 6 |
| WP_077283808.1 | 1.4E-05 | Qa.III | LDEINAAVALISDMATQIATAAEEQTHVTNEITQNVTSIK------------- | *Vibrio sp. OULL4* | Bacteria; Proteobacteria; Gammaproteobacteria; Vibrionales; Vibrionaceae; Vibrio | methyl-accepting chemotaxis protein | 6 |
| WP_080607911.1 | 1.4E-05 | Qa.III | LDEINAAVALISDMATQIATAAEEQTHVTNEITQNVTSIK------------- | *Vibrio parahaemolyticus* | Bacteria; Proteobacteria; Gammaproteobacteria; Vibrionales; Vibrionaceae; Vibrio | methyl-accepting chemotaxis protein | 6 |
| WP_083119908.1 | 1.4E-05 | Qa.III | LDEINAAVALISDMATQIATAAEEQTHVTNEITQNVTSIK------------- | *Vibrio parahaemolyticus* | Bacteria; Proteobacteria; Gammaproteobacteria; Vibrionales; Vibrionaceae; Vibrio | methyl-accepting chemotaxis protein | 6 |
| WP_083859889.1 | 1.4E-05 | Qa.III.b | IKGIGATIAQINEIATTIASAVEEQGAATAEISRNVQEAAK------------ | *Oceanibaculum indicum* | Bacteria; Proteobacteria; Alphaproteobacteria; Rhodospirillales; Rhodospirillaceae; Oceanibaculum | methyl-accepting chemotaxis protein | 6 |
| WP_084244305.1 | 1.4E-05 | Qa.IV | -----RTIHELTSVSTSIAAAVDEQAATTREIATSIQTAAGNTARASVEIK-- | *Rhodoplanes sp. Z2-YC6860* | Bacteria; Proteobacteria; Alphaproteobacteria; Rhizobiales; Hyphomicrobiaceae; Rhodoplanes | hypothetical protein | 6 |
| WP_084553679.1 | 1.4E-05 | Qa.III | IEKISTVIQEINEMIATIATAVEEQSATTVEIAGNISQAA------------- | *Desulfopila aestuarii* | Bacteria; Proteobacteria; Deltaproteobacteria; Desulfobacterales; Desulfobulbaceae; Desulfopila | methyl-accepting chemotaxis protein | 6 |
| WP_085569429.1 | 1.4E-05 | Qa.III | LDEINAAVALISDMATQIATAAEEQTHVTNEITQNVTSIK------------- | *Vibrio alginolyticus* | Bacteria; Proteobacteria; Gammaproteobacteria; Vibrionales; Vibrionaceae; Vibrio | methyl-accepting chemotaxis protein | 6 |
| WP_085588861.1 | 1.4E-05 | Qa.III | IQSIGGTIAQINEIASNIASAVEQQGAATQEITRNVQQAS------------- | *Thalassospira sp. MCCC 1A01428* | Bacteria; Proteobacteria; Alphaproteobacteria; Rhodospirillales; Rhodospirillaceae; Thalassospira | methyl-accepting chemotaxis protein | 6 |
| WP_085644911.1 | 1.4E-05 | Qc.III | IDSIGTTITEISEIASAIASAVEQQGAATDEINRNVQEASAGTGEVTVNIH-- | *Thalassospira sp. MCCC 1A03138* | Bacteria; Proteobacteria; Alphaproteobacteria; Rhodospirillales; Rhodospirillaceae; Thalassospira | methyl-accepting chemotaxis protein | 6 |
| WP_085677503.1 | 1.4E-05 | Qb.I | LQRTRLQLQETIEQSEISSEELTASNEEMQAINEELRSASEELETSK------ | *Pseudomonas* | Bacteria; Proteobacteria; Gammaproteobacteria; Pseudomonadales; Pseudomonadaceae | MULTISPECIES: response regulator | 6 |
| WP_086465226.1 | 1.4E-05 | Qa.IV | --------AQVNEISATISAAVEEQNATTAEIARNVQQAASATEGVTGNIG-- | *Oceanibaculum nanhaiense* | Bacteria; Proteobacteria; Alphaproteobacteria; Rhodospirillales; Rhodospirillaceae; Oceanibaculum | methyl-accepting chemotaxis protein | 6 |
| WP_086583474.1 | 1.4E-05 | Qa.III | LDEINAAVALISDMATQIATAAEEQTHVTNEITQNVTSIK------------- | *Vibrio parahaemolyticus* | Bacteria; Proteobacteria; Gammaproteobacteria; Vibrionales; Vibrionaceae; Vibrio | methyl-accepting chemotaxis protein | 6 |
| WP_086585752.1 | 1.4E-05 | Qa.III | LDEINAAVALISDMATQIATAAEEQTHVTNEITQNVTSIK------------- | *Vibrio parahaemolyticus* | Bacteria; Proteobacteria; Gammaproteobacteria; Vibrionales; Vibrionaceae; Vibrio | methyl-accepting chemotaxis protein | 6 |
| WP_092303541.1 | 1.4E-05 | Qc.III | -KAISGTIERLSEISSTIAAAVEEQGAATQEISRNVQQAAQGTQQVSSNIND- | *Bradyrhizobium sp. Ghvi* | Bacteria; Proteobacteria; Alphaproteobacteria; Rhizobiales; Bradyrhizobiaceae; Bradyrhizobium | methyl-accepting chemotaxis protein | 6 |
| WP_094455213.1 | 1.4E-05 | Qa.IV | IRSITGTIARVNEIATAIAAAVEEQGAATQEIARNVQQAA------------- | *Niveispirillum lacus* | Bacteria; Proteobacteria; Alphaproteobacteria; Rhodospirillales; Rhodospirillaceae; Niveispirillum | HAMP domain-containing protein | 6 |
| WP_096700624.1 | 1.4E-05 | Qc.III | ---ISGTISRINELATIIAGAVEEQGAATAEIARNVDQANEGTREVASNIS-- | *Magnetospirillum sp. 15-1* | Bacteria; Proteobacteria; Alphaproteobacteria; Rhodospirillales; Rhodospirillaceae; Magnetospirillum | methyl-accepting chemotaxis protein | 6 |
| WP_100176251.1 | 1.4E-05 | Qc.III | IKDISHTIEKLSEISSTIAAAVEEQGAATQEISRNVQQAAAGTHQVSSNITD- | *Bradyrhizobium sp. TSA1* | Bacteria; Proteobacteria; Alphaproteobacteria; Rhizobiales; Bradyrhizobiaceae; Bradyrhizobium | methyl-accepting chemotaxis protein | 6 |
| WP_100417594.1 | 1.4E-05 | Qc.III | -KAIGQTIERISDIATSISAAVEQQSAATKNIAQSVRAAAS------------ | *Bradyrhizobium lablabi* | Bacteria; Proteobacteria; Alphaproteobacteria; Rhizobiales; Bradyrhizobiaceae; Bradyrhizobium | methyl-accepting chemotaxis protein | 6 |
| WP_103792034.1 | 1.4E-05 | R.IV | IQEINRAVTHLDSMVQQNAELVVQSAAAASALQGQAGELAETAGHFR------ | *Leclercia* | Bacteria; Proteobacteria; Gammaproteobacteria; Enterobacterales; Enterobacteriaceae | MULTISPECIES: methyl-accepting chemotaxis protein | 6 |
| WP_104972973.1 | 1.4E-05 | Qa.III | LDEINAAVALISDMATQIATAAEEQTHVTNEITQNVTSIK------------- | *Vibrio diabolicus* | Bacteria; Proteobacteria; Gammaproteobacteria; Vibrionales; Vibrionaceae; Vibrio | methyl-accepting chemotaxis protein | 6 |
| WP_109108981.1 | 1.4E-05 | Qa.III | IRNIGQTIGRLNDIASSIAAAVEEQGAATSEIARNVQQAAGGTAQASQNIGQ- | *Azospirillum sp. TSO35-2* | Bacteria; Proteobacteria; Alphaproteobacteria; Rhodospirillales; Rhodospirillaceae; Azospirillum | HAMP domain-containing protein | 6 |
| WP_109885313.1 | 1.4E-05 | Qa.III | LDEINAAVALISDMATQIATAAEEQTHVTNEITQNVTSIK------------- | *Vibrio parahaemolyticus* | Bacteria; Proteobacteria; Gammaproteobacteria; Vibrionales; Vibrionaceae; Vibrio | methyl-accepting chemotaxis protein | 6 |
| WP_110637645.1 | 1.4E-05 | Qb.I | LQRTRLQLQETIEQSEISSEELTASNEEMQAINEELRSASEELETSK------ | *Pseudomonas sp. CC120222-01b* | Bacteria; Proteobacteria; Gammaproteobacteria; Pseudomonadales; Pseudomonadaceae; Pseudomonas | response regulator | 6 |
| WP_114089712.1 | 1.4E-05 | Qa.IV | -ETITRTISELSTLTASIAAAIEQQGSATAEISQSANSAAN------------ | *Thalassospira profundimaris* | Bacteria; Proteobacteria; Alphaproteobacteria; Rhodospirillales; Rhodospirillaceae; Thalassospira | HAMP domain-containing protein | 6 |
| WP_114097331.1 | 1.4E-05 | Qa.III | --AITTSINELDDITTALSSAVEQQASATHEITQNAQAANDTTRRAAD----- | *Thalassospira profundimaris* | Bacteria; Proteobacteria; Alphaproteobacteria; Rhodospirillales; Rhodospirillaceae; Thalassospira | HAMP domain-containing protein | 6 |
| WP_114139544.1 | 1.4E-05 | Qa.III | LDEINAAVALISDMATQIATAAEEQTHVTNEITQNVTSIK------------- | *Vibrio alginolyticus* | Bacteria; Proteobacteria; Gammaproteobacteria; Vibrionales; Vibrionaceae; Vibrio | HAMP domain-containing protein | 6 |
| WP_114322725.1 | 1.4E-05 | Qa.III | LDEINAAVALISDMATQIATAAEEQTHVTNEITQNVTSIK------------- | *Vibrio parahaemolyticus* | Bacteria; Proteobacteria; Gammaproteobacteria; Vibrionales; Vibrionaceae; Vibrio | HAMP domain-containing protein | 6 |
| ALG54817.1 | 1.5E-05 | Qa.III | LDEINAAVALISDMATQIATAAEEQTHVTNEITQNVTSIK------------- | *Vibrio parahaemolyticus* | Bacteria; Proteobacteria; Gammaproteobacteria; Vibrionales; Vibrionaceae; Vibrio | N-acetylglucosamine regulated methyl-accepting chemotaxis protein | 6 |
| ALM69338.1 | 1.5E-05 | Qa.III | LDEINAAVALISDMATQIATAAEEQTHVTNEITQNVTSIK------------- | *Vibrio parahaemolyticus* | Bacteria; Proteobacteria; Gammaproteobacteria; Vibrionales; Vibrionaceae; Vibrio | N-acetylglucosamine regulated methyl-accepting chemotaxis protein | 6 |
| APE87028.1 | 1.5E-05 | Qa.III | LDEINAAVALISDMATQIATAAEEQTHVTNEITQNVTSIK------------- | *Vibrio parahaemolyticus* | Bacteria; Proteobacteria; Gammaproteobacteria; Vibrionales; Vibrionaceae; Vibrio | N-acetylglucosamine regulated methyl-accepting chemotaxis protein | 6 |
| EHN68826.1 | 1.5E-05 | Qc.III | LEEISYSIQQISDMAMQISNAAAEQRTVTDEVNQNIQSASDVSDQ-------- | *Aliivibrio fischeri SR5* | Bacteria; Proteobacteria; Gammaproteobacteria; Vibrionales; Vibrionaceae; Aliivibrio | methyl-accepting chemotaxis protein | 6 |
| GAJ74489.1 | 1.5E-05 | Qa.III | LDEINAAVALISDMATQIATAAEEQTHVTNEITQNVTSIK------------- | *Vibrio sp. JCM 18905* | Bacteria; Proteobacteria; Gammaproteobacteria; Vibrionales; Vibrionaceae; Vibrio | LOW QUALITY PROTEIN: N-acetylglucosamine regulated methyl-accepting chemotaxis protein | 6 |
| KES23052.1 | 1.5E-05 | Qc.III.c | LQEIAASVSMVEQMSQQIAAAAEQQSAVADQVGESMERVQ------------- | *Pseudomonas sp. AAC* | Bacteria; Proteobacteria; Gammaproteobacteria; Pseudomonadales; Pseudomonadaceae; Pseudomonas | chemotaxis protein | 6 |
| KIT34201.1 | 1.5E-05 | Qa.III | LDEINAAVALISDMATQIATAAEEQTHVTNEITQNVTSIK------------- | *Vibrio parahaemolyticus 49* | Bacteria; Proteobacteria; Gammaproteobacteria; Vibrionales; Vibrionaceae; Vibrio | chemotaxis protein | 6 |
| KIT58424.1 | 1.5E-05 | Qa.III | LDEINAAVALISDMATQIATAAEEQTHVTNEITQNVTSIK------------- | *Vibrio parahaemolyticus EN9701072* | Bacteria; Proteobacteria; Gammaproteobacteria; Vibrionales; Vibrionaceae; Vibrio | chemotaxis protein | 6 |
| KQU27437.1 | 1.5E-05 | Qc.III | IERIAGIIGRIDESAISIGSAVEEQSATATEIRRSVEEASGRTETVSSSV--- | *Methylobacterium sp. Leaf94* | Bacteria; Proteobacteria; Alphaproteobacteria; Rhizobiales; Methylobacteriaceae; Methylobacterium | chemotaxis protein | 6 |
| OGQ93856.1 | 1.5E-05 | Qa.IV | -------INDINNIMTNVATAIEEQSATTREISKNVNQAFSGVTEANEN---- | *Deltaproteobacteria bacterium RIFOXYC2_FULL_48_10* | Bacteria; Proteobacteria; Deltaproteobacteria | hypothetical protein A2464_11975 | 6 |
| OJU24149.1 | 1.5E-05 | Qc.III | IKEIGGTIERLSEIASIIAAAVEEQGAATQEISRNVQQASQGTQQVSSNIAD- | *Nitrobacter sp. 62-13* | Bacteria; Proteobacteria; Alphaproteobacteria; Rhizobiales; Bradyrhizobiaceae; Nitrobacter | methyl-accepting chemotaxis protein | 6 |
| PIB13538.1 | 1.5E-05 | Qa.III | LDEINAAVALISDMATQIATAAEEQTHVTNEITQNVTTIK------------- | *Vibrio rotiferianus CAIM 577 = LMG 21460* | Bacteria; Proteobacteria; Gammaproteobacteria; Vibrionales; Vibrionaceae; Vibrio | histidine kinase | 6 |
| PID77982.1 | 1.5E-05 | Qa.III | ISKISNVISNVNDIVNTIAASVEEQSVTTQEISNNVQQAT------------- | *Deltaproteobacteria bacterium* | Bacteria; Proteobacteria; Deltaproteobacteria | hypothetical protein CSB21_02500 | 6 |
| PTM51354.1 | 1.5E-05 | SNAP.c | -EKVNQVIGEMRGIAASVAAAVEEQNAAVATIAEAVNRASDETRG-------- | *Phreatobacter oligotrophus* | Bacteria; Proteobacteria; Alphaproteobacteria; Phreatobacter | methyl-accepting chemotaxis protein | 6 |
| PYS80908.1 | 1.5E-05 | SNAP.b | -----------NRDLVQVLGELEEKREQLEQVNRRLEESNRELNEANAKLREL | *Acidobacteria bacterium* | Bacteria; Acidobacteria | hypothetical protein DMF67_18775 | 6 |
| WP_007560259.1 | 1.5E-05 | Qc.III | --SIQGTIVRLNEVATAISSAVEQQSAVTREMSASMQTAAQGVGAITTNL--- | *Methylobacterium sp. GXF4* | Bacteria; Proteobacteria; Alphaproteobacteria; Rhizobiales; Methylobacteriaceae; Methylobacterium | methyl-accepting chemotaxis protein | 6 |
| WP_008055658.1 | 1.5E-05 | Qb.I | LQRTKLHLQETIEQSEVSSEELKASNEEMQAINEELRSATEELETS------- | *Pseudomonas sp. GM78* | Bacteria; Proteobacteria; Gammaproteobacteria; Pseudomonadales; Pseudomonadaceae; Pseudomonas | PAS domain S-box protein | 6 |
| WP_009540380.1 | 1.5E-05 | Qa.III | IEAISGTIAEINEITSAIAAAVEEQGAATSEISRNVQEAAQGSQE-------- | *Caenispirillum salinarum* | Bacteria; Proteobacteria; Alphaproteobacteria; Rhodospirillales; Rhodospirillaceae; Caenispirillum | methyl-accepting chemotaxis protein | 6 |
| WP_010453828.1 | 1.5E-05 | Qa.III | LDEINAAVALISDMATQIATAAEEQTHVTNEITQNVTTIK------------- | *Vibrio rotiferianus* | Bacteria; Proteobacteria; Gammaproteobacteria; Vibrionales; Vibrionaceae; Vibrio | methyl-accepting chemotaxis protein | 6 |
| WP_015668126.1 | 1.5E-05 | Qc.III | ---ISGTIERLSEIASAIAAAVEEQGAATQEISRNVQQAAQGTQQVSANIA-- | *Bradyrhizobium oligotrophicum* | Bacteria; Proteobacteria; Alphaproteobacteria; Rhizobiales; Bradyrhizobiaceae; Bradyrhizobium | methyl-accepting chemotaxis protein | 6 |
| WP_019024576.1 | 1.5E-05 | Qc.III | --SITQSVTRISDMNTQIASAAEEQSTVAEEINRNVANVTEVLDQT------- | *Thioalkalivibrio* | Bacteria; Proteobacteria; Gammaproteobacteria; Chromatiales; Ectothiorhodospiraceae | MULTISPECIES: methyl-accepting chemotaxis protein | 6 |
| WP_023450339.1 | 1.5E-05 | Qc.III.b | --DIIQAVQHLHDIAQTVSGRVDEQSRTTSEMSHVVRRAANEAQT-------- | *Asticcacaulis sp. AC402* | Bacteria; Proteobacteria; Alphaproteobacteria; Caulobacterales; Caulobacteraceae; Asticcacaulis | methyl-accepting chemotaxis protein | 6 |
| WP_024921919.1 | 1.5E-05 | Qa.IV | IKEIGATINLISEISSSIAAAVEEQGAATQEIARNVQYAAQ------------ | *Afipia* | Bacteria; Proteobacteria; Alphaproteobacteria; Rhizobiales; Bradyrhizobiaceae | MULTISPECIES: methyl-accepting chemotaxis protein | 6 |
| WP_027299293.1 | 1.5E-05 | Qc.III | -GSIGGTIGTINEIATTIAAAVEEQGAATQEIARNVHGAAEGTHQVS------ | *Rhodospirillales bacterium URHD0088* | Bacteria; Proteobacteria; Alphaproteobacteria; Rhodospirillales; unclassified Rhodospirillales | HAMP domain-containing protein | 6 |

| | | | | | | | | |
|---|---|---|---|---|---|---|---|---|
|  | **WP_027568629.1** | 1.5E-05 | Qa.IV | IKEIGNTIARMSEISSAIAAAVEEQGAATQEISRNIQQAAN------------ | *Bradyrhizobium sp. URHA0013* | Bacteria; Proteobacteria; Alphaproteobacteria; Rhizobiales; Bradyrhizobiaceae; Bradyrhizobium | methyl-accepting chemotaxis protein | 6 |
|  | **WP_029863444.1** | 1.5E-05 | Qa.III | LDEINAAVALISDMATQIATAAEEQTHVTNEITQNVTSIK------------- | *Vibrio parahaemolyticus* | Bacteria; Proteobacteria; Gammaproteobacteria; Vibrionales; Vibrionaceae; Vibrio | methyl-accepting chemotaxis protein partial | 6 |
|  | **WP_035238995.1** | 1.5E-05 | Qa.III | IKEIVEIIGDINSIVTSVASAIEEQSATTQEISNNVSQAA------------- | *Desulfobacter vibrioformis* | Bacteria; Proteobacteria; Deltaproteobacteria; Desulfobacterales; Desulfobacteraceae; Desulfobacter | methyl-accepting chemotaxis protein | 6 |
| | **WP_040475802.1** | 1.5E-05 | Qc.III | ---ISGTISRINELATIIAGAVEEQGAATAEIARNVDQANEGTREVASNIS-- | *Magnetospirillum caucaseum* | Bacteria; Proteobacteria; Alphaproteobacteria; Rhodospirillales; Rhodospirillaceae; Magnetospirillum | methyl-accepting chemotaxis protein | 6 |
|  | **WP_043271486.1** | 1.5E-05 | Qc.III.c | LQEIAASVSMVEQMSQQIAAAAEQQSAVADQVGESMERVQ------------- | *Pseudomonas sp. AAC* | Bacteria; Proteobacteria; Gammaproteobacteria; Pseudomonadales; Pseudomonadaceae; Pseudomonas | methyl-accepting chemotaxis protein | 6 |
|  | **WP_051548046.1** | 1.5E-05 | Qa.IV | ---INSSIEDISNAVAMIASAVEEQGSATHKISQSVQSASQNTQT-------- | *Sneathiella glossodoripedis* | Bacteria; Proteobacteria; Alphaproteobacteria; Sneathiellales; Sneathiellaceae; Sneathiella | hypothetical protein | 6 |
|  | **WP_051957915.1** | 1.5E-05 | Qa.III | IESIVGIINEINSMVTSIAAAIEEQSTTTQEISSNVSQAGQGVQA-------- | *Desulfobacter vibrioformis* | Bacteria; Proteobacteria; Deltaproteobacteria; Desulfobacterales; Desulfobacteraceae; Desulfobacter | methyl-accepting chemotaxis protein | 6 |
|  | **WP_056852781.1** | 1.5E-05 | Qb.I | LQRTKLHLQETIEQSEVSSEELKASNEEMQAINEELRSATEELETS------- | *Pseudomonas sp. Root562* | Bacteria; Proteobacteria; Gammaproteobacteria; Pseudomonadales; Pseudomonadaceae; Pseudomonas | PAS domain S-box protein | 6 |
|  | **WP_067046444.1** | 1.5E-05 | Qc.III | LDSMTDSVMKINDLGIQIATAAEEQSSVTEEINRNMTMIQSMVNQLTEN---- | *Moritella sp. JT01* | Bacteria; Proteobacteria; Gammaproteobacteria; Alteromonadales; Moritellaceae; Moritella | methyl-accepting chemotaxis protein | 6 |
| | **WP_068502704.1** | 1.5E-05 | Qc.III | --GIASTIETINELSAAISAAVEEQGAATQEIARSVSQAAEGTHQAAENV--- | *Magnetospirillum moscoviense* | Bacteria; Proteobacteria; Alphaproteobacteria; Rhodospirillales; Rhodospirillaceae; Magnetospirillum | PAS domain S-box protein | 6 |
|  | **WP_074121597.1** | 1.5E-05 | Qc.III | IQEIGGTIERLSEVSAAIAAAVEEQGAATQEISRNVQQASVGTQEVS------ | *Bradyrhizobium sp. AS23.2* | Bacteria; Proteobacteria; Alphaproteobacteria; Rhizobiales; Bradyrhizobiaceae; Bradyrhizobium | methyl-accepting chemotaxis protein | 6 |
|  | **WP_075701588.1** | 1.5E-05 | SNAP.c | -GDVSDVIHEMSQIAAAVAVSVEQQNAAIQSITENVANATQ------------ | *Pseudovibrio sp. Ab134* | Bacteria; Proteobacteria; Alphaproteobacteria; Rhodobacterales; Rhodobacteraceae; Pseudovibrio | HAMP domain-containing protein | 6 |
|  | **WP_080562785.1** | 1.5E-05 | Qc.III | LEEISYSIQQISDMAMQISNAAAEQRTVTDEVNQNIQSASDVSDQ-------- | *Aliivibrio fischeri* | Bacteria; Proteobacteria; Gammaproteobacteria; Vibrionales; Vibrionaceae; Aliivibrio | methyl-accepting chemotaxis protein | 6 |
|  | **WP_080951991.1** | 1.5E-05 | Qc.III | LEEISYSIQQISDMAMQISNAAAEQRTVTDEVNQNIQSASDVSDQ-------- | *Aliivibrio fischeri* | Bacteria; Proteobacteria; Gammaproteobacteria; Vibrionales; Vibrionaceae; Aliivibrio | methyl-accepting chemotaxis protein | 6 |
|  | **WP_081249009.1** | 1.5E-05 | Qc.III | LEEISYSIQQISDMAMQISNAAAEQRTVTDEVNQNIQSASDVSDQ-------- | *Aliivibrio fischeri* | Bacteria; Proteobacteria; Gammaproteobacteria; Vibrionales; Vibrionaceae; Aliivibrio | methyl-accepting chemotaxis protein | 6 |
|  | **WP_081249112.1** | 1.5E-05 | Qc.III | LEEISYSIQQISDMAMQISNAAAEQRTVTDEVNQNIQSASDVSDQ-------- | *Aliivibrio fischeri* | Bacteria; Proteobacteria; Gammaproteobacteria; Vibrionales; Vibrionaceae; Aliivibrio | methyl-accepting chemotaxis protein | 6 |
|  | **WP_081249457.1** | 1.5E-05 | Qc.III | LEEISYSIQQISDMAMQISNAAAEQRTVTDEVNQNIQSASDVSDQ-------- | *Aliivibrio fischeri* | Bacteria; Proteobacteria; Gammaproteobacteria; Vibrionales; Vibrionaceae; Aliivibrio | methyl-accepting chemotaxis protein | 6 |
|  | **WP_081249698.1** | 1.5E-05 | Qc.III | LEEISYSIQQISDMAMQISNAAAEQRTVTDEVNQNIQSASDVSDQ-------- | *Aliivibrio fischeri* | Bacteria; Proteobacteria; Gammaproteobacteria; Vibrionales; Vibrionaceae; Aliivibrio | methyl-accepting chemotaxis protein | 6 |
|  | **WP_081250081.1** | 1.5E-05 | Qc.III | LEEISYSIQQISDMAMQISNAAAEQRTVTDEVNQNIQSASDVSDQ-------- | *Aliivibrio fischeri* | Bacteria; Proteobacteria; Gammaproteobacteria; Vibrionales; Vibrionaceae; Aliivibrio | methyl-accepting chemotaxis protein | 6 |
|  | **WP_081300254.1** | 1.5E-05 | Qc.III | LEEISYSIQQISDMAMQISNAAAEQRTVTDEVNQNIQSASDVSDQ-------- | *Aliivibrio fischeri* | Bacteria; Proteobacteria; Gammaproteobacteria; Vibrionales; Vibrionaceae; Aliivibrio | methyl-accepting chemotaxis protein | 6 |
|  | **WP_081302368.1** | 1.5E-05 | Qc.III | LEEISYSIQQISDMAMQISNAAAEQRTVTDEVNQNIQSASDVSDQ-------- | *Aliivibrio fischeri* | Bacteria; Proteobacteria; Gammaproteobacteria; Vibrionales; Vibrionaceae; Aliivibrio | methyl-accepting chemotaxis protein | 6 |
|  | **WP_081302772.1** | 1.5E-05 | Qc.III | LEEISYSIQQISDMAMQISNAAAEQRTVTDEVNQNIQSASDVSDQ-------- | *Aliivibrio fischeri* | Bacteria; Proteobacteria; Gammaproteobacteria; Vibrionales; Vibrionaceae; Aliivibrio | methyl-accepting chemotaxis protein | 6 |
|  | **WP_081304269.1** | 1.5E-05 | Qc.III | LEEISYSIQQISDMAMQISNAAAEQRTVTDEVNQNIQSASDVSDQ-------- | *Aliivibrio fischeri* | Bacteria; Proteobacteria; Gammaproteobacteria; Vibrionales; Vibrionaceae; Aliivibrio | methyl-accepting chemotaxis protein | 6 |
|  | **WP_081329338.1** | 1.5E-05 | Qc.III | LEEISYSIQQISDMAMQISNAAAEQRTVTDEVNQNIQSASDVSDQ-------- | *Aliivibrio fischeri* | Bacteria; Proteobacteria; Gammaproteobacteria; Vibrionales; Vibrionaceae; Aliivibrio | methyl-accepting chemotaxis protein | 6 |
| | **WP_081433888.1** | 1.5E-05 | Qc.III | IEGVGSQIRKMSDVATSISAAVEEQGVATREIVRNVDQAATGTNSVTAHISE- | *Azorhizobium caulinodans* | Bacteria; Proteobacteria; Alphaproteobacteria; Rhizobiales; Xanthobacteraceae; Azorhizobium | methyl-accepting chemotaxis protein | 6 |
|  | **WP_083754912.1** | 1.5E-05 | Qc.III | IKEIGGTIGRISDIAATIASAVEQQSAATQEIARSVQNVAEG----------- | *Bradyrhizobium elkanii* | Bacteria; Proteobacteria; Alphaproteobacteria; Rhizobiales; Bradyrhizobiaceae; Bradyrhizobium | methyl-accepting chemotaxis protein | 6 |
|  | **WP_084618349.1** | 1.5E-05 | Qc.III.c | -EELAGVAARRKELATHFEEEVLRHAQQLSSVSHDLRDVTSTLTTDT------ | *Jonesia quinghaiensis* | Bacteria; Actinobacteria; Micrococcales; Jonesiaceae; Jonesia | hypothetical protein | 6 |
|  | **WP_086028509.1** | 1.5E-05 | Qc.III | LEEISYSIQQISDMAMQISNAAAEQRTVTDEVNQNIQSASDVSDQ-------- | *Aliivibrio fischeri* | Bacteria; Proteobacteria; Gammaproteobacteria; Vibrionales; Vibrionaceae; Aliivibrio | methyl-accepting chemotaxis protein | 6 |
| | **WP_091863981.1** | 1.5E-05 | Qc.III | --SIQGTIVRLNEVATAISSAVEQQSAVTREMSASMQTAAQGVGAITTNL--- | *Methylobacterium brachiatum* | Bacteria; Proteobacteria; Alphaproteobacteria; Rhizobiales; Methylobacteriaceae; Methylobacterium | methyl-accepting chemotaxis protein | 6 |
|  | **WP_092782468.1** | 1.5E-05 | Qa.III | IQAIVRVIAEINEITTGVSAAVEQQAAATGEISQNAQQAAHSTQDASQN---- | *Rhodospira trueperi* | Bacteria; Proteobacteria; Alphaproteobacteria; Rhodospirillales; Rhodospirillaceae; Rhodospira | HAMP domain-containing protein | 6 |
|  | **WP_092787363.1** | 1.5E-05 | Qc.III | ----ADTVGRIGEISTTVAAAVEEQNAAIGEVSRNTAEAASGTQEVSHSI--- | *Rhodospira trueperi* | Bacteria; Proteobacteria; Alphaproteobacteria; Rhodospirillales; Rhodospirillaceae; Rhodospira | methyl-accepting chemotaxis protein | 6 |
|  | **WP_108179046.1** | 1.5E-05 | SNAP.c | -EKVNQVIGEMRGIAASVAAAVEEQNAAVATIAEAVNRASDETRG-------- | *Phreatobacter oligotrophus* | Bacteria; Proteobacteria; Alphaproteobacteria; Phreatobacter | methyl-accepting chemotaxis protein | 6 |
|  | **WP_108652222.1** | 1.5E-05 | Qa.III | LEQITESINQITDMATQISSAAEEQRAVTDEVSRNLQAVK------------- | *Gammaproteobacteria bacterium DM2* | Bacteria; Proteobacteria; Gammaproteobacteria | methyl-accepting chemotaxis protein | 6 |
|  | **WP_108652223.1** | 1.5E-05 | Qa.III | LEQITESINQITDMATQISSAAEEQRAVTDEVSRNLQAVK------------- | *Gammaproteobacteria bacterium DM2* | Bacteria; Proteobacteria; Gammaproteobacteria | methyl-accepting chemotaxis protein | 6 |
|  | **WP_109120332.1** | 1.5E-05 | Qc.III | ---IGRTIQSINSIAGGIADAIEHQGGAIREIVHNMEHAAAGTR--------- | *Azospirillum sp. TSO22-1* | Bacteria; Proteobacteria; Alphaproteobacteria; Rhodospirillales; Rhodospirillaceae; Azospirillum | chemotaxis protein | 6 |
|  | **WP_111357237.1** | 1.5E-05 | Qc.III | IKEIGSTIGRIAEIAASIAAAIEQQGAATGEIARNVQQASQGTVAVARS---- | *Rhodoplanes elegans* | Bacteria; Proteobacteria; Alphaproteobacteria; Rhizobiales; Hyphomicrobiaceae; Rhodoplanes | methyl-accepting chemotaxis protein | 6 |
|  | **WP_114010221.1** | 1.5E-05 | Qa.III.b | IQGIAETMETVNEYTRTISLAVDEQGSATFEISQNVAKAA------------- | *Cohaesibacter sp. YE-B6* | Bacteria; Proteobacteria; Alphaproteobacteria; Rhizobiales; Cohaesibacteraceae | HAMP domain-containing protein | 6 |
|  | **ACH64207.1** | 1.6E-05 | Qc.III | LEEISYSIQQISDMAMQISNAAAEQRTVTDEVNQNIQSASDVSDQ-------- | *Aliivibrio fischeri MJ11* | Bacteria; Proteobacteria; Gammaproteobacteria; Vibrionales; Vibrionaceae; Aliivibrio | methyl-accepting chemotaxis protein | 6 |
| | **AMA58845.1** | 1.6E-05 | Qc.III | IKQIGDTISQISGIATTIAAAVEQQGAATTEIARNVQKASDGTREVSGSITK- | *Bradyrhizobium sp. CCGE-LA001* | Bacteria; Proteobacteria; Alphaproteobacteria; Rhizobiales; Bradyrhizobiaceae; Bradyrhizobium | chemotaxis protein | 6 |
|  | **CDD74021.1** | 1.6E-05 | Qa.IV | ---INQNIMEISAAMEETGANIDSQTTSIQNIDKTCKDTTHAV---------- | *Clostridium sp. CAG:62* | Bacteria; Firmicutes; Clostridia; Clostridiales; Clostridiaceae; Clostridium; environmental samples | methyl-accepting chemotaxis sensory transducer with Cache sensor | 6 |
|  | **EGA67825.1** | 1.6E-05 | Qc.III.c | LDVMVGSIARINDLGLEIATAASQQSAVTDEVSRNMTTIQTMVEELTAN---- | *Vibrio sinaloensis DSM 21326* | Bacteria; Proteobacteria; Gammaproteobacteria; Vibrionales; Vibrionaceae; Vibrio; Vibrio oreintalis group | methyl-accepting chemotaxis protein | 6 |
|  | **KJR42451.1** | 1.6E-05 | Qc.III | -------INEINDISAAIAAAVEEQSAATNEITRNVREASEGMGEVNENI--- | *Candidatus Magnetoovum chiemensis* | Bacteria; Nitrospirae; Nitrospirales; Nitrospiraceae; Candidatus Magnetoovum | methyl-accepting chemotaxis sensory transducer partial | 6 |
| | **KYG98956.1** | 1.6E-05 | Qc.III | IKQIGDTISQISGIATTIAAAVEQQGAATTEIARNVQKASDGTREVSGSITK- | *Bradyrhizobium sp. DOA1* | Bacteria; Proteobacteria; Alphaproteobacteria; Rhizobiales; Bradyrhizobiaceae; Bradyrhizobium | chemotaxis protein | 6 |
|  | **OCH04516.1** | 1.6E-05 | Qc.III | LEEISYSIQQISDMAMQISNAAAEQRTVTDEVNQNIQSASDVSDQ-------- | *Aliivibrio fischeri* | Bacteria; Proteobacteria; Gammaproteobacteria; Vibrionales; Vibrionaceae; Aliivibrio | chemotaxis protein | 6 |
|  | **OCH48942.1** | 1.6E-05 | Qc.III | LEEISYSIQQISDMAMQISNAAAEQRTVTDEVNQNIQSASDVSDQ-------- | *Aliivibrio fischeri* | Bacteria; Proteobacteria; Gammaproteobacteria; Vibrionales; Vibrionaceae; Aliivibrio | chemotaxis protein | 6 |
|  | **OED51138.1** | 1.6E-05 | Qc.III | LEEISYSIQQISDMAMQISNAAAEQRTVTDEVNQNIQSASDVSDQ-------- | *Aliivibrio fischeri* | Bacteria; Proteobacteria; Gammaproteobacteria; Vibrionales; Vibrionaceae; Aliivibrio | chemotaxis protein | 6 |
|  | **OED54896.1** | 1.6E-05 | Qc.III | LEEISYSIQQISDMAMQISNAAAEQRTVTDEVNQNIQSASDVSDQ-------- | *Aliivibrio fischeri* | Bacteria; Proteobacteria; Gammaproteobacteria; Vibrionales; Vibrionaceae; Aliivibrio | chemotaxis protein | 6 |
|  | **OEE09433.1** | 1.6E-05 | Qc.III | LEEISYSIQQISDMAMQISNAAAEQRTVTDEVNQNIQSASDVSDQ-------- | *Aliivibrio fischeri ZF-211* | Bacteria; Proteobacteria; Gammaproteobacteria; Vibrionales; Vibrionaceae; Aliivibrio | chemotaxis protein | 6 |
|  | **OFI50210.1** | 1.6E-05 | Qa.IV | --EIEANVNILSQGINEMSQSINEQTEAINQINEAVATVDEQTKQ-NV----- | *Campylobacter fetus subsp. fetus* | Bacteria; Proteobacteria; Epsilonproteobacteria; Campylobacterales; Campylobacteraceae; Campylobacter | hypothetical protein CFF27374_01010 | 6 |
|  | **OLA02277.1** | 1.6E-05 | Qa.IV | ---INQNIMEISAAMEETGANIDSQTTSIQNIDKTCKDTTHAV---------- | *Clostridium sp. CAG:62_40_43* | Bacteria; Firmicutes; Clostridia; Clostridiales; Clostridiaceae; Clostridium | hypothetical protein BHW11_05645 | 6 |
| | **PCI42315.1** | 1.6E-05 | Qc.III | -GSIGGTINELNEISSAIAAAVEEQGAATQEIARNVEQAANGTTEVSSNI--- | *Rhodospirillaceae bacterium* | Bacteria; Proteobacteria; Alphaproteobacteria; Rhodospirillales; Rhodospirillaceae | methyl-accepting chemotaxis protein | 6 |
| | **PHS79572.1** | 1.6E-05 | Qc.III | -GSIGGTINELNEISSAIAAAVEEQGAATQEIARNVEQAANGTTEVSSNI--- | *Rhodospirillaceae bacterium* | Bacteria; Proteobacteria; Alphaproteobacteria; Rhodospirillales; Rhodospirillaceae | methyl-accepting chemotaxis protein | 6 |
| | **PXX36214.1** | 1.6E-05 | Qc.III | IDSIGTTITEISEIASAIASAVEQQGAATDEINRNVQEASAGTGEVTANI--- | *Thalassospira sp. 11-3* | Bacteria; Proteobacteria; Alphaproteobacteria; Rhodospirillales; Rhodospirillaceae; Thalassospira | methyl-accepting chemotaxis protein | 6 |
|  | **PYM42448.1** | 1.6E-05 | Qb.I | LAATREYLQSVIEQQEVANEELQSSNEEVQSANEELQSVNEELETSK------ | *Candidatus Rokubacteria bacterium* | Bacteria; Candidatus Rokubacteria | hypothetical protein DME16_23815 partial | 6 |
|  | **PYM69383.1** | 1.6E-05 | Qb.I | LAATREYLQSVIEQQEVANEELQSSNEEVQSANEELQSVNEELETSK------ | *Candidatus Rokubacteria bacterium* | Bacteria; Candidatus Rokubacteria | hypothetical protein DME10_23915 partial | 6 |
| | **SFQ26388.1** | 1.6E-05 | Qc.III | -KAISGTIERLSEISSTIAAAVEEQGAATQEISRNVQQAAQGTQQVSSNIND- | *Bradyrhizobium sp. Ghvi* | Bacteria; Proteobacteria; Alphaproteobacteria; Rhizobiales; Bradyrhizobiaceae; Bradyrhizobium | methyl-accepting chemotaxis sensory transducer | 6 |
| | **SIS98180.1** | 1.6E-05 | Qc.III | IDSIGTTITEISEIASAIASAVEQQGAATDEINRNVQEASAGTGEVTANI--- | *Thalassospira xiamenensis M-5 = DSM 17429* | Bacteria; Proteobacteria; Alphaproteobacteria; Rhodospirillales; Rhodospirillaceae; Thalassospira | methyl-accepting chemotaxis protein | 6 |
| | **WP_007089901.1** | 1.6E-05 | Qc.III | IDSIGTTITEISEIASAIASAVEQQGAATDEINRNVQEASAGTGEVTANI--- | *Thalassospira xiamenensis* | Bacteria; Proteobacteria; Alphaproteobacteria; Rhodospirillales; Rhodospirillaceae; Thalassospira | methyl-accepting chemotaxis protein | 6 |
|  | **WP_011643976.1** | 1.6E-05 | Qc.III | --RIGSVIGQIEEIASGISAAVEEQVAVTQDISRNMGEAATDVSHV------- | *Maricaulis maris* | Bacteria; Proteobacteria; Alphaproteobacteria; Rhodobacterales; Hyphomonadaceae; Maricaulis | methyl-accepting chemotaxis protein | 6 |
| | **WP_019962750.1** | 1.6E-05 | Qc.III | -GSILATIESIRNISSTIAAAVEEQTAVTRDISHNMQAASTGVQSITSSMG-- | *Pannonibacter phragmitetus* | Bacteria; Proteobacteria; Alphaproteobacteria; Rhodobacterales; Rhodobacteraceae; Pannonibacter | PAS domain S-box protein | 6 |

| | | | | | | | | |
|---|---|---|---|---|---|---|---|---|
|  | **WP_027298731.1** | 1.6E-05 | Qa.III | IQGIGTTIAEVNSIASSIAAAVEQQGAATQEIARNVQQAAQGT---------- | *Rhodospirillales bacterium URHD0088* | Bacteria; Proteobacteria; Alphaproteobacteria; Rhodospirillales; unclassified Rhodospirillales | methyl-accepting chemotaxis protein | 6 |
| | **WP_037987557.1** | 1.6E-05 | Qc.III | IDSIGTTITEISEIASAIASAVEQQGAATDEINRNVQEASAGTGEVTANI--- | *Thalassospira permensis* | Bacteria; Proteobacteria; Alphaproteobacteria; Rhodospirillales; Rhodospirillaceae; Thalassospira | methyl-accepting chemotaxis protein | 6 |
|  | **WP_041805099.1** | 1.6E-05 | Qa.III | IKQIGNIIGQMSEISSTIASAVEEQGAATKEISANIQQAA------------- | *Rhodopseudomonas palustris* | Bacteria; Proteobacteria; Alphaproteobacteria; Rhizobiales; Bradyrhizobiaceae; Rhodopseudomonas | HAMP domain-containing protein | 6 |
|  | **WP_044916282.1** | 1.6E-05 | SNAP.b | -NDLMKDAGESVQVVQLLIDNIEEQGKQLDTTRDSMDRMQQ------------ | *Butyrivibrio sp. WCE2006* | Bacteria; Firmicutes; Clostridia; Clostridiales; Lachnospiraceae; Butyrivibrio | methyl-accepting chemotaxis protein | 6 |
|  | **WP_045774323.1** | 1.6E-05 | Qa.III | -------IGQINDITTTIAAAIEQQGAATGEISRNVQQAASGTEEAS------ | *Elstera litoralis* | Bacteria; Proteobacteria; Alphaproteobacteria; Rhodospirillales; Rhodospirillaceae; Elstera | HAMP domain-containing protein | 6 |
|  | **WP_050626811.1** | 1.6E-05 | Qa.IV | ------TIAEIDHVSGEIAAAVDQQGAATREISSNLQQAADRTRDVNK----- | *Bradyrhizobium viridifuturi* | Bacteria; Proteobacteria; Alphaproteobacteria; Rhizobiales; Bradyrhizobiaceae; Bradyrhizobium | chemotaxis protein | 6 |
|  | **WP_055254777.1** | 1.6E-05 | SNAP.b | -QNLSSAAFQINEVMAQSNNDLRSQSEQIDQSSESISQMNIAIDEVSHN---- | *Pseudoalteromonas sp. P1-11* | Bacteria; Proteobacteria; Gammaproteobacteria; Alteromonadales; Pseudoalteromonadaceae; Pseudoalteromonas | methyl-accepting chemotaxis protein | 6 |
|  | **WP_056248306.1** | 1.6E-05 | Qa.III | IRAITATIGNLSEISLSVSSAVTEQAAVTHEISNNMQVAARNVDAVR------ | *Methylobacterium sp. Leaf456* | Bacteria; Proteobacteria; Alphaproteobacteria; Rhizobiales; Methylobacteriaceae; Methylobacterium | methyl-accepting chemotaxis protein | 6 |
| | **WP_063092957.1** | 1.6E-05 | Qc.III | IDSIGTTITEISEIASAIASAVEQQGAATDEINRNVQEASAGTGEVTANI--- | *Thalassospira* | Bacteria; Proteobacteria; Alphaproteobacteria; Rhodospirillales; Rhodospirillaceae | MULTISPECIES: methyl-accepting chemotaxis protein | 6 |
| | **WP_063197165.1** | 1.6E-05 | Qc.III | --TIGGTIERLSEISSTIAAAVEEQGAATSEISRNVQNAAAGTTQVSANI--- | *Bradyrhizobium sp. AT1* | Bacteria; Proteobacteria; Alphaproteobacteria; Rhizobiales; Bradyrhizobiaceae; Bradyrhizobium | HAMP domain-containing protein | 6 |
| | **WP_071240166.1** | 1.6E-05 | Qc.III | IDSIGTTITEISEIASAIASAVEQQGAATDEINRNVQEASAGTGEVTANI--- | *Thalassospira sp. MIT1004* | Bacteria; Proteobacteria; Alphaproteobacteria; Rhodospirillales; Rhodospirillaceae; Thalassospira | methyl-accepting chemotaxis protein | 6 |
| | **WP_082813707.1** | 1.6E-05 | Qc.III | IKQIGDTISQISGIATTIAAAVEQQGAATTEIARNVQKASDGTREVSGSITK- | *Bradyrhizobium sp. DOA1* | Bacteria; Proteobacteria; Alphaproteobacteria; Rhizobiales; Bradyrhizobiaceae; Bradyrhizobium | HAMP domain-containing protein | 6 |
| | **WP_083543376.1** | 1.6E-05 | Qc.III | IKQIGDTISQISGIATTIAAAVEQQGAATTEIARNVQKASDGTREVSGSITK- | *Bradyrhizobium sp. CCGE-LA001* | Bacteria; Proteobacteria; Alphaproteobacteria; Rhizobiales; Bradyrhizobiaceae; Bradyrhizobium | HAMP domain-containing protein | 6 |
|  | **WP_084536898.1** | 1.6E-05 | Qc.III | -RDIGKTVSDVNEIAGTIAAAVEEQAAATQEIGRNVQQAAR------------ | *Azospirillum halopraeferens* | Bacteria; Proteobacteria; Alphaproteobacteria; Rhodospirillales; Rhodospirillaceae; Azospirillum | hypothetical protein | 6 |
|  | **WP_085690556.1** | 1.6E-05 | Qb.I | LQRTKLHLQETIEQSEVSSEELKASNEEMQAINEELRSATEELETS------- | *Pseudomonas* | Bacteria; Proteobacteria; Gammaproteobacteria; Pseudomonadales; Pseudomonadaceae | MULTISPECIES: PAS domain S-box protein | 6 |
| | **WP_092191981.1** | 1.6E-05 | Qa.III | IQQITNVINDVNSIVTAIAAAVEEQSVTTREITQNLAQASSGIQEINENVTQ- | *Desulfomicrobium norvegicum* | Bacteria; Proteobacteria; Deltaproteobacteria; Desulfovibrionales; Desulfomicrobiaceae; Desulfomicrobium | methyl-accepting chemotaxis protein | 6 |
| | **WP_092259481.1** | 1.6E-05 | Qc.III | IGEISGTIARLSEIAAAIAAAVEQQGAATQEIARNVQQAAQGTQQVSSNVG-- | *Bradyrhizobium sp. Rc3b* | Bacteria; Proteobacteria; Alphaproteobacteria; Rhizobiales; Bradyrhizobiaceae; Bradyrhizobium | PAS domain S-box protein | 6 |
|  | **WP_092685778.1** | 1.6E-05 | Qa.III | IKQIGNIIGQMSEISSTIASAVEEQGAATKEISANIQQAA------------- | *Rhodopseudomonas pseudopalustris* | Bacteria; Proteobacteria; Alphaproteobacteria; Rhizobiales; Bradyrhizobiaceae; Rhodopseudomonas | HAMP domain-containing protein | 6 |
|  | **WP_095732344.1** | 1.6E-05 | Qa.IV | ------TIAEIDHVSGEIAAAVDQQGAATREISSNLQQAADRTRDVNK----- | *Bradyrhizobium sp. UFLA03-84* | Bacteria; Proteobacteria; Alphaproteobacteria; Rhizobiales; Bradyrhizobiaceae; Bradyrhizobium | chemotaxis protein | 6 |
| | **WP_097657613.1** | 1.6E-05 | Qc.III | --TIGGTIERLSEISSTIAAAVEEQGAATSEISRNVQNAAAGTTQVSANI--- | *Bradyrhizobium sp. Y36* | Bacteria; Proteobacteria; Alphaproteobacteria; Rhizobiales; Bradyrhizobiaceae; Bradyrhizobium | methyl-accepting chemotaxis protein | 6 |
| | **WP_106949979.1** | 1.6E-05 | Qc.III | IGEISGTIARLSEIAAAIAAAVEQQGAATQEIARNVQQAAQGTQQVSSNVG-- | *Bradyrhizobium sp. MOS003* | Bacteria; Proteobacteria; Alphaproteobacteria; Rhizobiales; Bradyrhizobiaceae; Bradyrhizobium | PAS domain S-box protein | 6 |
|  | **WP_109068617.1** | 1.6E-05 | Qa.III | IVGIGHTIQDINHIAGGIAAGIGQQGSAIRSIVSNMDQAA------------- | *Azospirillum sp. TSH58* | Bacteria; Proteobacteria; Alphaproteobacteria; Rhodospirillales; Rhodospirillaceae; Azospirillum | chemotaxis protein | 6 |
|  | **WP_111434517.1** | 1.6E-05 | Qc.III | IKEIGATIGRLSDISAGIAAAIEEQGAATQEITRNVTEAATG----------- | *Rhodobium orientis* | Bacteria; Proteobacteria; Alphaproteobacteria; Rhizobiales; Rhodobiaceae; Rhodobium | methyl-accepting chemotaxis protein | 6 |
| | **WP_114094443.1** | 1.6E-05 | Qc.III | IDSIGTTITEISEIASAIASAVEQQGAATDEINRNVQEASAGTGEVTANI--- | *Thalassospira xiamenensis* | Bacteria; Proteobacteria; Alphaproteobacteria; Rhodospirillales; Rhodospirillaceae; Thalassospira | HAMP domain-containing protein | 6 |
|  | **AND88043.1** | 1.7E-05 | Qc.III | IQEIGGTIERLSEVSAAIAAAVEEQGAATQEISRNVQQASVGTQEV------- | *Bradyrhizobium diazoefficiens USDA 110* | Bacteria; Proteobacteria; Alphaproteobacteria; Rhizobiales; Bradyrhizobiaceae; Bradyrhizobium | chemotaxis protein | 6 |
|  | **ATY78516.1** | 1.7E-05 | Qa.III | LEQITTSINQISDMATQISSAAEEQRAVTDEVSRNIQAVK------------- | *Aeromonas veronii* | Bacteria; Proteobacteria; Gammaproteobacteria; Aeromonadales; Aeromonadaceae; Aeromonas | methyl-accepting chemotaxis protein | 6 |
|  | **AXB07713.1** | 1.7E-05 | Qa.III | LEQITTSINQISDMATQISSAAEEQRAVTDEVSRNIQAVK------------- | *Aeromonas caviae* | Bacteria; Proteobacteria; Gammaproteobacteria; Aeromonadales; Aeromonadaceae; Aeromonas | methyl-accepting chemotaxis protein | 6 |
|  | **BAK98943.1** | 1.7E-05 | Qb.III | IDATAQNTNDASRLADDSSAELKNENEQMQQMITAMDEISS------------ | *Oscillibacter valericigenes Sjm18-20* | Bacteria; Firmicutes; Clostridia; Clostridiales; Oscillospiraceae; Oscillibacter | methyl-accepting chemotaxis protein | 6 |
| | **BAM86552.1** | 1.7E-05 | Qa.III | IKQIGGTIGQISAIASSIATAVEQQGGATQEIARNVQSVAAGTQEAASDLTQ- | *Bradyrhizobium oligotrophicum S58* | Bacteria; Proteobacteria; Alphaproteobacteria; Rhizobiales; Bradyrhizobiaceae; Bradyrhizobium | methyl-accepting chemotaxis sensory transducer | 6 |
|  | **EDM59323.1** | 1.7E-05 | Qa.III | LDEINAAVALISDMATQIATAAEEQTHVTNEITQNVTSIK------------- | *Vibrio parahaemolyticus AQ3810* | Bacteria; Proteobacteria; Gammaproteobacteria; Vibrionales; Vibrionaceae; Vibrio | methyl-accepting chemotaxis protein | 6 |
|  | **EJN05932.1** | 1.7E-05 | Qc.III | IQEIGGTIERLSEVSAAIAAAVEEQGAATQEISRNVQQASVGTQEV------- | *Bradyrhizobium sp. YR681* | Bacteria; Proteobacteria; Alphaproteobacteria; Rhizobiales; Bradyrhizobiaceae; Bradyrhizobium | methyl-accepting chemotaxis protein | 6 |
|  | **EKF07914.1** | 1.7E-05 | Qc.III | INEITDIVGELNESASAIAAAVEEQGAATQEISGSVENAS------------- | *Thalassospira profundimaris WP0211* | Bacteria; Proteobacteria; Alphaproteobacteria; Rhodospirillales; Rhodospirillaceae; Thalassospira | methyl-accepting chemotaxis protein | 6 |
|  | **ESQ87857.1** | 1.7E-05 | Qa.III | IKGIGETIKHMNEIASSIADSVHQQGDATNEIARNVNEASD------------ | *Asticcacaulis benevestitus DSM 16100 = ATCC BAA-896* | Bacteria; Proteobacteria; Alphaproteobacteria; Caulobacterales; Caulobacteraceae; Asticcacaulis | methyl-accepting chemotaxis protein | 6 |
|  | **KEP89506.1** | 1.7E-05 | Qa.III | LEQITTSINQISDMATQISSAAEEQRAVTDEVSRNIQAVK------------- | *Aeromonas caviae* | Bacteria; Proteobacteria; Gammaproteobacteria; Aeromonadales; Aeromonadaceae; Aeromonas | chemotaxis protein | 6 |
|  | **KOY08007.1** | 1.7E-05 | Qc.III | IQEIGGTIERLSEVSAAIAAAVEEQGAATQEISRNVQQASVGTQEV------- | *Bradyrhizobium japonicum* | Bacteria; Proteobacteria; Alphaproteobacteria; Rhizobiales; Bradyrhizobiaceae; Bradyrhizobium | chemotaxis protein | 6 |
|  | **KZC98166.1** | 1.7E-05 | Qc.III | INEITDIVGELNESASAIAAAVEEQGAATQEISGSVENAS------------- | *Thalassospira sp. MCCC 1A02898* | Bacteria; Proteobacteria; Alphaproteobacteria; Rhodospirillales; Rhodospirillaceae; Thalassospira | hypothetical protein AUQ41_18205 | 6 |
|  | **OCQ42930.1** | 1.7E-05 | Qa.III | LEQITTSINQISDMATQISSAAEEQRAVTDEVSRNIQAVK------------- | *Aeromonas veronii* | Bacteria; Proteobacteria; Gammaproteobacteria; Aeromonadales; Aeromonadaceae; Aeromonas | chemotaxis protein | 6 |
|  | **OCR98885.1** | 1.7E-05 | Qa.IV | --EIEANVNILSQGINEMSQSINEQTEAINQINEAVATVDEQTKQ-NV----- | *Campylobacter fetus subsp. testudinum* | Bacteria; Proteobacteria; Epsilonproteobacteria; Campylobacterales; Campylobacteraceae; Campylobacter | chemotaxis protein partial | 6 |
|  | **OEU75746.1** | 1.7E-05 | Qc.III | -EQISGGADQISDTSQQLSQAATEQASSLEEITSSMAEMGSRTKQ-------- | *Desulfuromonadales bacterium C00003068* | Bacteria; Proteobacteria; Deltaproteobacteria; Desulfuromonadales | hypothetical protein BA874_04350 | 6 |
| | **PCI40422.1** | 1.7E-05 | Qc.III | -GSIGGTINELNEISSAIAAAVEEQGAATQEIARNVEQAANGTTEVSSNI--- | *Rhodospirillaceae bacterium* | Bacteria; Proteobacteria; Alphaproteobacteria; Rhodospirillales; Rhodospirillaceae | methyl-accepting chemotaxis protein | 6 |
|  | **PMU05442.1** | 1.7E-05 | Qa.III | LEQITTSINQISDMATQISSAAEEQRAVTDEVSRNIQAVK------------- | *Aeromonas salmonicida* | Bacteria; Proteobacteria; Gammaproteobacteria; Aeromonadales; Aeromonadaceae; Aeromonas | methyl-accepting chemotaxis protein | 6 |
|  | **PNO62055.1** | 1.7E-05 | Qa.III | LEQITTSINQISDMATQISSAAEEQRAVTDEVSRNIQAVK------------- | *Aeromonas caviae* | Bacteria; Proteobacteria; Gammaproteobacteria; Aeromonadales; Aeromonadaceae; Aeromonas | methyl-accepting chemotaxis protein | 6 |
|  | **POG19239.1** | 1.7E-05 | Qa.III | LEQITTSINQISDMATQISSAAEEQRAVTDEVSRNIQAVK------------- | *Aeromonas veronii* | Bacteria; Proteobacteria; Gammaproteobacteria; Aeromonadales; Aeromonadaceae; Aeromonas | methyl-accepting chemotaxis protein | 6 |
|  | **POV91625.1** | 1.7E-05 | Qa.III | LEQITTSINQISDMATQISSAAEEQRAVTDEVSRNIQAVK------------- | *Aeromonas sp. ASNIH8* | Bacteria; Proteobacteria; Gammaproteobacteria; Aeromonadales; Aeromonadaceae; Aeromonas | methyl-accepting chemotaxis protein | 6 |
|  | **PXV58009.1** | 1.7E-05 | Qa.III | LEQITTSINQISDMATQISSAAEEQRAVTDEVSRNIQAVK------------- | *Aeromonas diversa* | Bacteria; Proteobacteria; Gammaproteobacteria; Aeromonadales; Aeromonadaceae; Aeromonas | UNVERIFIED_ORG: methyl-accepting chemotaxis sensory transducer with Cache sensor | 6 |
|  | **PYP37962.1** | 1.7E-05 | Qb.I | ---TKDYLQSVIEQQEAANEELQSSNEEVQSANEELQSINEEIETSK------ | *Gemmatimonadetes bacterium* | Bacteria; Gemmatimonadetes | hypothetical protein DMD48_10385 | 6 |
|  | **RAI25614.1** | 1.7E-05 | Qc.III | IKEISGTIGQIAEVASAIAAAVEEQGAATAEISRNIQQAASGT---------- | *Rhodoplanes elegans* | Bacteria; Proteobacteria; Alphaproteobacteria; Rhizobiales; Hyphomicrobiaceae; Rhodoplanes | chemotaxis protein partial | 6 |
|  | **RAJ09351.1** | 1.7E-05 | Qa.III | LEQITTSINQISDMATQISSAAEEQRAVTDEVSRNIQAVK------------- | *Aeromonas salmonicida* | Bacteria; Proteobacteria; Gammaproteobacteria; Aeromonadales; Aeromonadaceae; Aeromonas | methyl-accepting chemotaxis sensory transducer with Cache sensor | 6 |
|  | **RDE64331.1** | 1.7E-05 | Qa.III | LEQITTSINQISDMATQISSAAEEQRAVTDEVSRNIQAVK------------- | *Aeromonas veronii* | Bacteria; Proteobacteria; Gammaproteobacteria; Aeromonadales; Aeromonadaceae; Aeromonas | methyl-accepting chemotaxis protein | 6 |
|  | **SDG99699.1** | 1.7E-05 | Qa.IV | IKEIGATIARISDISTTIAATIEEQGAATSEIARNVSEAAK------------ | *Bradyrhizobium sp. Rc2d* | Bacteria; Proteobacteria; Alphaproteobacteria; Rhizobiales; Bradyrhizobiaceae; Bradyrhizobium | methyl-accepting chemotaxis protein | 6 |
|  | **SES39387.1** | 1.7E-05 | SNAP.b | --------TEMTSQVENNATKLSEQNEQFKNVFGEMREMTKMLNESVE----- | *Butyrivibrio fibrisolvens* | Bacteria; Firmicutes; Clostridia; Clostridiales; Lachnospiraceae; Butyrivibrio | Methyl-accepting chemotaxis protein | 6 |
|  | **SFJ09905.1** | 1.7E-05 | Qc.III | IQEIGGTIERLSEVSAAIAAAVEEQGAATQEISRNVQQASVGTQEV------- | *Bradyrhizobium sp. cf659* | Bacteria; Proteobacteria; Alphaproteobacteria; Rhizobiales; Bradyrhizobiaceae; Bradyrhizobium | methyl-accepting chemotaxis protein | 6 |
|  | **SHJ02051.1** | 1.7E-05 | SNAP.b | --------TEMTSQVENNATKLSEQNEQFKNVFGEMREMTKMLNESVE----- | *Butyrivibrio fibrisolvens DSM 3071* | Bacteria; Firmicutes; Clostridia; Clostridiales; Lachnospiraceae; Butyrivibrio | Methyl-accepting chemotaxis protein | 6 |
| | **SMX61296.1** | 1.7E-05 | Qc.III | -REISGTIEKLAEISSTIAAAVEEQGAATQEISRNIQQAAQGTHEVSSNITD- | *Bradyrhizobium sp. ORS 285* | Bacteria; Proteobacteria; Alphaproteobacteria; Rhizobiales; Bradyrhizobiaceae; Bradyrhizobium | putative methyl-accepting chemotaxis protein | 6 |
|  | **WP_005317198.1** | 1.7E-05 | Qa.III | LEQITTSINQISDMATQISSAAEEQRAVTDEVSRNIQAVK------------- | *Aeromonas salmonicida* | Bacteria; Proteobacteria; Gammaproteobacteria; Aeromonadales; Aeromonadaceae; Aeromonas | methyl-accepting chemotaxis protein | 6 |
|  | **WP_005329900.1** | 1.7E-05 | Qa.III | LEQITTSINQISDMATQISSAAEEQRAVTDEVSRNIQAVK------------- | *Aeromonas media* | Bacteria; Proteobacteria; Gammaproteobacteria; Aeromonadales; Aeromonadaceae; Aeromonas | methyl-accepting chemotaxis protein | 6 |
|  | **WP_005335052.1** | 1.7E-05 | Qa.III | LEQITTSINQISDMATQISSAAEEQRAVTDEVSRNIQAVK------------- | *Aeromonas* | Bacteria; Proteobacteria; Gammaproteobacteria; Aeromonadales; Aeromonadaceae | MULTISPECIES: methyl-accepting chemotaxis protein | 6 |
|  | **WP_005354019.1** | 1.7E-05 | Qa.III | LEQITTSINQISDMATQISSAAEEQRAVTDEVSRNIQAVK------------- | *Aeromonas* | Bacteria; Proteobacteria; Gammaproteobacteria; Aeromonadales; Aeromonadaceae | MULTISPECIES: methyl-accepting chemotaxis protein | 6 |

| | | | | | | | | |
|---|---|---|---|---|---|---|---|---|
|  | **WP_005477512.1** | 1.7E-05 | Qa.III | LDEINAAVALISDMATQIATAAEEQTHVTNEITQNVTSIK------------- | *Vibrio parahaemolyticus* | Bacteria; Proteobacteria; Gammaproteobacteria; Vibrionales; Vibrionaceae; Vibrio | methyl-accepting chemotaxis protein | 6 |
| | **WP_008961865.1** | 1.7E-05 | Qc.III | -REISGTIEKLAEISSTIAAAVEEQGAATQEISRNIQQAAQGTHEVSSNITD- | *Bradyrhizobium sp. STM 3809* | Bacteria; Proteobacteria; Alphaproteobacteria; Rhizobiales; Bradyrhizobiaceae; Bradyrhizobium | methyl-accepting chemotaxis protein | 6 |
| | **WP_009031104.1** | 1.7E-05 | Qc.III | -REISGTIEKLAEISSTIAAAVEEQGAATQEISRNIQQAAQGTHEVSSNITD- | *Bradyrhizobium sp. ORS 375* | Bacteria; Proteobacteria; Alphaproteobacteria; Rhizobiales; Bradyrhizobiaceae; Bradyrhizobium | methyl-accepting chemotaxis protein | 6 |
|  | **WP_011085335.1** | 1.7E-05 | Qc.III | IQEIGGTIERLSEVSAAIAAAVEEQGAATQEISRNVQQASVGTQEV------- | *Bradyrhizobium diazoefficiens* | Bacteria; Proteobacteria; Alphaproteobacteria; Rhizobiales; Bradyrhizobiaceae; Bradyrhizobium | methyl-accepting chemotaxis protein | 6 |
|  | **WP_011188222.1** | 1.7E-05 | Qa.III | IDGISQVINEVNEIVSTIVAAVEEQAVTTNEIAANVSQAS------------- | *Desulfotalea psychrophila* | Bacteria; Proteobacteria; Deltaproteobacteria; Desulfobacterales; Desulfobulbaceae; Desulfotalea | methyl-accepting chemotaxis protein | 6 |
|  | **WP_011470548.1** | 1.7E-05 | Qa.IV | IKEIGATITRIADISSTIAATVEEQGAATAEIARNVSEAAK------------ | *Rhodopseudomonas palustris* | Bacteria; Proteobacteria; Alphaproteobacteria; Rhizobiales; Bradyrhizobiaceae; Rhodopseudomonas | chemotaxis protein | 6 |
| | **WP_012030049.1** | 1.7E-05 | Qc.III | -REISGTIEKLAEISSTIAAAVEEQGAATQEISRNIQQAAQGTHEVSSNITD- | *Bradyrhizobium sp. ORS 278* | Bacteria; Proteobacteria; Alphaproteobacteria; Rhizobiales; Bradyrhizobiaceae; Bradyrhizobium | methyl-accepting chemotaxis protein | 6 |
|  | **WP_013113426.1** | 1.7E-05 | Qb.I | ----KNMMSESMDYIEGAANIIAQTTSNIEDVHQASDKIKDITK--------- | *Brachyspira murdochii* | Bacteria; Spirochaetes; Brachyspirales; Brachyspiraceae; Brachyspira | HAMP domain-containing protein | 6 |
|  | **WP_017412438.1** | 1.7E-05 | Qa.III | LEQITTSINQISDMATQISSAAEEQRAVTDEVSRNIQAVK------------- | *Aeromonas salmonicida* | Bacteria; Proteobacteria; Gammaproteobacteria; Aeromonadales; Aeromonadaceae; Aeromonas | methyl-accepting chemotaxis protein | 6 |
|  | **WP_025326132.1** | 1.7E-05 | Qa.III | LEQITTSINQISDMATQISSAAEEQRAVTDEVSRNIQAVK------------- | *Aeromonas hydrophila* | Bacteria; Proteobacteria; Gammaproteobacteria; Aeromonadales; Aeromonadaceae; Aeromonas | methyl-accepting chemotaxis protein | 6 |
| | **WP_026345235.1** | 1.7E-05 | Qc.III | LGEISSTIVQMNEISTTIASAVEEQGAATQEIARNAEQAATGTEQVTNSV--- | *Novispirillum itersonii* | Bacteria; Proteobacteria; Alphaproteobacteria; Rhodospirillales; Rhodospirillaceae; Novispirillum | methyl-accepting chemotaxis protein | 6 |
|  | **WP_027591486.1** | 1.7E-05 | Qa.III | LRQIVESLRSISGLAASIASATLEQSHVVDEINGNVTQTAS------------ | *Pseudomonas sp. RL* | Bacteria; Proteobacteria; Gammaproteobacteria; Pseudomonadales; Pseudomonadaceae; Pseudomonas | methyl-accepting chemotaxis protein | 6 |
|  | **WP_031842957.1** | 1.7E-05 | Qa.III | LDEINAAVALISDMATQIATAAEEQTHVTNEITQNVTSIK------------- | *Vibrio parahaemolyticus* | Bacteria; Proteobacteria; Gammaproteobacteria; Vibrionales; Vibrionaceae; Vibrio | methyl-accepting chemotaxis protein | 6 |
|  | **WP_039562705.1** | 1.7E-05 | Qa.III | LRQIVESLRSISGLAASIASATLEQSHVVDEINGNVTQTAS------------ | *Pseudomonas flexibilis* | Bacteria; Proteobacteria; Gammaproteobacteria; Pseudomonadales; Pseudomonadaceae; Pseudomonas | methyl-accepting chemotaxis protein | 6 |
|  | **WP_039605845.1** | 1.7E-05 | Qa.III | LRQIVESLRSISGLAASIASATLEQSHVVDEINGNVTQTAS------------ | *Pseudomonas flexibilis* | Bacteria; Proteobacteria; Gammaproteobacteria; Pseudomonadales; Pseudomonadaceae; Pseudomonas | methyl-accepting chemotaxis protein | 6 |
|  | **WP_040098086.1** | 1.7E-05 | Qa.III | LEQITTSINQISDMATQISSAAEEQRAVTDEVSRNIQAVK------------- | *Aeromonas australiensis* | Bacteria; Proteobacteria; Gammaproteobacteria; Aeromonadales; Aeromonadaceae; Aeromonas | methyl-accepting chemotaxis protein | 6 |
|  | **WP_041212095.1** | 1.7E-05 | Qa.III | LEQITTSINQISDMATQISSAAEEQRAVTDEVSRNIQAVK------------- | *Aeromonas allosaccharophila* | Bacteria; Proteobacteria; Gammaproteobacteria; Aeromonadales; Aeromonadaceae; Aeromonas | methyl-accepting chemotaxis protein | 6 |
|  | **WP_042013836.1** | 1.7E-05 | Qa.III | LEQITTSINQISDMATQISSAAEEQRAVTDEVSRNIQAVK------------- | *Aeromonas fluvialis* | Bacteria; Proteobacteria; Gammaproteobacteria; Aeromonadales; Aeromonadaceae; Aeromonas | methyl-accepting chemotaxis protein | 6 |
|  | **WP_042022906.1** | 1.7E-05 | Qa.III | LEQITTSINQISDMATQISSAAEEQRAVTDEVSRNIQAVK------------- | *Aeromonas sobria* | Bacteria; Proteobacteria; Gammaproteobacteria; Aeromonadales; Aeromonadaceae; Aeromonas | methyl-accepting chemotaxis protein | 6 |
|  | **WP_042063910.1** | 1.7E-05 | Qa.III | LEQITTSINQISDMATQISSAAEEQRAVTDEVSRNIQAVK------------- | *Aeromonas allosaccharophila* | Bacteria; Proteobacteria; Gammaproteobacteria; Aeromonadales; Aeromonadaceae; Aeromonas | methyl-accepting chemotaxis protein | 6 |
|  | **WP_042078946.1** | 1.7E-05 | Qa.III | LEQITTSINQISDMATQISSAAEEQRAVTDEVSRNIQAVK------------- | *Aeromonas sanarellii* | Bacteria; Proteobacteria; Gammaproteobacteria; Aeromonadales; Aeromonadaceae; Aeromonas | methyl-accepting chemotaxis protein | 6 |
|  | **WP_042083552.1** | 1.7E-05 | Qa.III | LEQITTSINQISDMATQISSAAEEQRAVTDEVSRNIQAVK------------- | *Aeromonas* | Bacteria; Proteobacteria; Gammaproteobacteria; Aeromonadales; Aeromonadaceae | MULTISPECIES: methyl-accepting chemotaxis protein | 6 |
|  | **WP_042643993.1** | 1.7E-05 | Qa.III | LEQITTSINQISDMATQISSAAEEQRAVTDEVSRNIQAVK------------- | *Aeromonas eucrenophila* | Bacteria; Proteobacteria; Gammaproteobacteria; Aeromonadales; Aeromonadaceae; Aeromonas | methyl-accepting chemotaxis protein | 6 |
|  | **WP_042652716.1** | 1.7E-05 | Qa.III | LEQITTSINQISDMATQISSAAEEQRAVTDEVSRNIQAVK------------- | *Aeromonas encheleia* | Bacteria; Proteobacteria; Gammaproteobacteria; Aeromonadales; Aeromonadaceae; Aeromonas | methyl-accepting chemotaxis protein | 6 |
|  | **WP_042656009.1** | 1.7E-05 | Qa.III | LEQITTSINQISDMATQISSAAEEQRAVTDEVSRNIQAVK------------- | *Aeromonas allosaccharophila* | Bacteria; Proteobacteria; Gammaproteobacteria; Aeromonadales; Aeromonadaceae; Aeromonas | methyl-accepting chemotaxis protein | 6 |
|  | **WP_042658602.1** | 1.7E-05 | Qa.III | LEQITTSINQISDMATQISSAAEEQRAVTDEVSRNIQAVK------------- | *Aeromonas allosaccharophila* | Bacteria; Proteobacteria; Gammaproteobacteria; Aeromonadales; Aeromonadaceae; Aeromonas | methyl-accepting chemotaxis protein | 6 |
|  | **WP_042880252.1** | 1.7E-05 | Qa.III | LEQITTSINQISDMATQISSAAEEQRAVTDEVSRNIQAVK------------- | *Aeromonas sp. ZOR0002* | Bacteria; Proteobacteria; Gammaproteobacteria; Aeromonadales; Aeromonadaceae; Aeromonas | methyl-accepting chemotaxis protein | 6 |
|  | **WP_043762143.1** | 1.7E-05 | Qa.III | LEQITTSINQISDMATQISSAAEEQRAVTDEVSRNIQAVK------------- | *Aeromonas taiwanensis* | Bacteria; Proteobacteria; Gammaproteobacteria; Aeromonadales; Aeromonadaceae; Aeromonas | methyl-accepting chemotaxis protein | 6 |
|  | **WP_044303633.1** | 1.7E-05 | Qa.III | LEQITTSINQISDMATQISSAAEEQRAVTDEVSRNIQAVK------------- | *Aeromonas salmonicida* | Bacteria; Proteobacteria; Gammaproteobacteria; Aeromonadales; Aeromonadaceae; Aeromonas | methyl-accepting chemotaxis protein | 6 |
|  | **WP_047437966.1** | 1.7E-05 | Qa.III | LEQITTSINQISDMATQISSAAEEQRAVTDEVSRNIQAVK------------- | *Aeromonas sp. ZOR0001* | Bacteria; Proteobacteria; Gammaproteobacteria; Aeromonadales; Aeromonadaceae; Aeromonas | methyl-accepting chemotaxis protein | 6 |
|  | **WP_048207133.1** | 1.7E-05 | Qa.III | LEQITTSINQISDMATQISSAAEEQRAVTDEVSRNIQAVK------------- | *Aeromonas caviae* | Bacteria; Proteobacteria; Gammaproteobacteria; Aeromonadales; Aeromonadaceae; Aeromonas | methyl-accepting chemotaxis protein | 6 |
| | **WP_051243787.1** | 1.7E-05 | Qc.III | IQEIVKVIGEINQIATVIAAAVEEQTAATQEIARNVTEASRGTTEVSSNI--- | *Thalassobaculum salexigens* | Bacteria; Proteobacteria; Alphaproteobacteria; Rhodospirillales; Rhodospirillaceae; Thalassobaculum | methyl-accepting chemotaxis protein | 6 |
|  | **WP_053288723.1** | 1.7E-05 | Qa.III | LEQITTSINQISDMATQISSAAEEQRAVTDEVSRNIQAVK------------- | *Aeromonas caviae* | Bacteria; Proteobacteria; Gammaproteobacteria; Aeromonadales; Aeromonadaceae; Aeromonas | methyl-accepting chemotaxis protein | 6 |
| | **WP_056431246.1** | 1.7E-05 | Qc.III | --SIQDTIMTLNEVASAISAAVEEQSAVTREMSASMQTASNGVGQITSSLS-- | *Methylobacterium* | Bacteria; Proteobacteria; Alphaproteobacteria; Rhizobiales; Methylobacteriaceae | MULTISPECIES: methyl-accepting chemotaxis protein | 6 |
|  | **WP_058054257.1** | 1.7E-05 | Qa.III | LEQITTSINQISDMATQISSAAEEQRAVTDEVSRNIQAVK------------- | *Aeromonas allosaccharophila* | Bacteria; Proteobacteria; Gammaproteobacteria; Aeromonadales; Aeromonadaceae; Aeromonas | methyl-accepting chemotaxis protein | 6 |
|  | **WP_058055633.1** | 1.7E-05 | Qa.III | LEQITTSINQISDMATQISSAAEEQRAVTDEVSRNIQAVK------------- | *Aeromonas veronii* | Bacteria; Proteobacteria; Gammaproteobacteria; Aeromonadales; Aeromonadaceae; Aeromonas | methyl-accepting chemotaxis protein | 6 |
|  | **WP_058059222.1** | 1.7E-05 | Qa.III | LEQITTSINQISDMATQISSAAEEQRAVTDEVSRNIQAVK------------- | *Aeromonas veronii* | Bacteria; Proteobacteria; Gammaproteobacteria; Aeromonadales; Aeromonadaceae; Aeromonas | methyl-accepting chemotaxis protein | 6 |
|  | **WP_059111473.1** | 1.7E-05 | Qa.III | LEQITTSINQISDMATQISSAAEEQRAVTDEVSRNIQAVK------------- | *Aeromonas salmonicida* | Bacteria; Proteobacteria; Gammaproteobacteria; Aeromonadales; Aeromonadaceae; Aeromonas | methyl-accepting chemotaxis protein | 6 |
|  | **WP_059168381.1** | 1.7E-05 | Qa.III | LEQITTSINQISDMATQISSAAEEQRAVTDEVSRNIQAVK------------- | *Aeromonas salmonicida* | Bacteria; Proteobacteria; Gammaproteobacteria; Aeromonadales; Aeromonadaceae; Aeromonas | methyl-accepting chemotaxis protein | 6 |
|  | **WP_060736721.1** | 1.7E-05 | Qc.III | IQEIGGTIERLSEVSAAIAAAVEEQGAATQEISRNVQQASVGTQEV------- | *Bradyrhizobium sp. CCGE-LA001* | Bacteria; Proteobacteria; Alphaproteobacteria; Rhizobiales; Bradyrhizobiaceae; Bradyrhizobium | HAMP domain-containing protein | 6 |
|  | **WP_061847049.1** | 1.7E-05 | Qc.III | IQEIGGTIERLSEVSAAIAAAVEEQGAATQEISRNVQQASVGTQEV------- | *Bradyrhizobium sp. DOA1* | Bacteria; Proteobacteria; Alphaproteobacteria; Rhizobiales; Bradyrhizobiaceae; Bradyrhizobium | HAMP domain-containing protein | 6 |
|  | **WP_063518374.1** | 1.7E-05 | Qa.III | LDEINAAVALISDMATQIATAAEEQTHVTNEITQNVTSIK------------- | *Vibrio parahaemolyticus* | Bacteria; Proteobacteria; Gammaproteobacteria; Vibrionales; Vibrionaceae; Vibrio | methyl-accepting chemotaxis protein | 6 |
|  | **WP_065765871.1** | 1.7E-05 | Qa.III | LDEINAAVALISDMATQIATAAEEQTHVTNEITQNVTSIK------------- | *Vibrio parahaemolyticus* | Bacteria; Proteobacteria; Gammaproteobacteria; Vibrionales; Vibrionaceae; Vibrio | methyl-accepting chemotaxis protein | 6 |
|  | **WP_068978034.1** | 1.7E-05 | Qa.III | LEQITTSINQISDMATQISSAAEEQRAVTDEVSRNIQAVK------------- | *Aeromonas sp. EERV15* | Bacteria; Proteobacteria; Gammaproteobacteria; Aeromonadales; Aeromonadaceae; Aeromonas | methyl-accepting chemotaxis protein | 6 |
|  | **WP_069652132.1** | 1.7E-05 | Qa.III | LEQITTSINQISDMATQISSAAEEQRAVTDEVSRNIQAVK------------- | *Aeromonas caviae* | Bacteria; Proteobacteria; Gammaproteobacteria; Aeromonadales; Aeromonadaceae; Aeromonas | methyl-accepting chemotaxis protein | 6 |
|  | **WP_071463015.1** | 1.7E-05 | Qc.II | -----------RTIAHRASDEARESSTVIEQMTQAMDRINQAVTQASQRLT-- | *Idiomarina sp. MD25a* | Bacteria; Proteobacteria; Gammaproteobacteria; Alteromonadales; Idiomarinaceae; Idiomarina | methyl-accepting chemotaxis protein | 6 |
|  | **WP_074077225.1** | 1.7E-05 | Qc.III | IQEIGGTIERLSEVSAAIAAAVEEQGAATQEISRNVQQASVGTQEV------- | *Bradyrhizobium* | Bacteria; Proteobacteria; Alphaproteobacteria; Rhizobiales; Bradyrhizobiaceae | MULTISPECIES: methyl-accepting chemotaxis protein | 6 |
|  | **WP_075114169.1** | 1.7E-05 | Qa.III | LEQITTSINQISDMATQISSAAEEQRAVTDEVSRNIQAVK------------- | *Gammaproteobacteria* | Bacteria; Proteobacteria | MULTISPECIES: methyl-accepting chemotaxis protein | 6 |
|  | **WP_077280614.1** | 1.7E-05 | Qa.III | LDEINAAVALISDMATQIATAAEEQTHVTNEITQNVTSIK------------- | *Vibrio sp. SALL6* | Bacteria; Proteobacteria; Gammaproteobacteria; Vibrionales; Vibrionaceae; Vibrio | methyl-accepting chemotaxis protein | 6 |
|  | **WP_082756087.1** | 1.7E-05 | Qc.III | IQEIGGTIERLSEVSAAIAAAVEEQGAATQEISRNVQQASVGTQEV------- | *Bradyrhizobium diazoefficiens* | Bacteria; Proteobacteria; Alphaproteobacteria; Rhizobiales; Bradyrhizobiaceae; Bradyrhizobium | methyl-accepting chemotaxis protein | 6 |
| | **WP_083938782.1** | 1.7E-05 | Qa.III | IKQIGGTIGQISAIASSIATAVEQQGGATQEIARNVQSVAAGTQEAASDLTQ- | *Bradyrhizobium oligotrophicum* | Bacteria; Proteobacteria; Alphaproteobacteria; Rhizobiales; Bradyrhizobiaceae; Bradyrhizobium | methyl-accepting chemotaxis protein | 6 |
|  | **WP_083991915.1** | 1.7E-05 | Qc.III.c | --AISQVIVQISEISTVIASAVEEQAAATKEVTANINGTTKAIDDA------- | *Bradyrhizobium embrapense* | Bacteria; Proteobacteria; Alphaproteobacteria; Rhizobiales; Bradyrhizobiaceae; Bradyrhizobium | PAS domain-containing protein | 6 |
|  | **WP_084301888.1** | 1.7E-05 | Qc.III | IQEIGGTIERLSEVSAAIAAAVEEQGAATQEISRNVQQASVGTQEV------- | *Bradyrhizobium sp. WSM2254* | Bacteria; Proteobacteria; Alphaproteobacteria; Rhizobiales; Bradyrhizobiaceae; Bradyrhizobium | methyl-accepting chemotaxis protein | 6 |
|  | **WP_084807383.1** | 1.7E-05 | Qc.III | IQEIGGTIERLSEVSAAIAAAVEEQGAATQEISRNVQQASVGTQEV------- | *Bradyrhizobium sp. NAS80.1* | Bacteria; Proteobacteria; Alphaproteobacteria; Rhizobiales; Bradyrhizobiaceae; Bradyrhizobium | HAMP domain-containing protein | 6 |
|  | **WP_085969993.1** | 1.7E-05 | Qc.III | IQEIGGTIERLSEVSAAIAAAVEEQGAATQEISRNVQQASVGTQEV------- | *Bradyrhizobium sp. YR681* | Bacteria; Proteobacteria; Alphaproteobacteria; Rhizobiales; Bradyrhizobiaceae; Bradyrhizobium | HAMP domain-containing protein | 6 |
|  | **WP_085985429.1** | 1.7E-05 | Qc.III.c | -DGISASVRAINETATAIASAIEEQGAATREIVQAIAQASS------------ | *Methylorubrum populi* | Bacteria; Proteobacteria; Alphaproteobacteria; Rhizobiales; Methylobacteriaceae; Methylorubrum | methyl-accepting chemotaxis protein | 6 |
|  | **WP_091882755.1** | 1.7E-05 | Qa.IV | IKEIGATIARISDISTTIAATIEEQGAATSEIARNVSEAAK------------ | *Bradyrhizobium sp. Rc2d* | Bacteria; Proteobacteria; Alphaproteobacteria; Rhizobiales; Bradyrhizobiaceae; Bradyrhizobium | HAMP domain-containing protein | 6 |
|  | **WP_092187662.1** | 1.7E-05 | Qc.III | IQEIGGTIERLSEVSAAIAAAVEEQGAATQEISRNVQQASVGTQEV------- | *Bradyrhizobium sp. cf659* | Bacteria; Proteobacteria; Alphaproteobacteria; Rhizobiales; Bradyrhizobiaceae; Bradyrhizobium | methyl-accepting chemotaxis protein | 6 |
|  | **WP_092363871.1** | 1.7E-05 | Qb.I | LQSTKEQLQATVEQAETSTEELKASNEELQAINEELRSTTEELETSK------ | *Pseudomonas seleniipraecipitans* | Bacteria; Proteobacteria; Gammaproteobacteria; Pseudomonadales; Pseudomonadaceae; Pseudomonas | PAS domain S-box protein | 6 |
|  | **WP_096511787.1** | 1.7E-05 | Qb.I | LQRTKLHLQETIEQSEVSSEELKASNEEMQAINEELRSATEELETS------- | *Pseudomonas sp. LAB-08* | Bacteria; Proteobacteria; Gammaproteobacteria; Pseudomonadales; Pseudomonadaceae; Pseudomonas | PAS domain S-box protein | 6 |

| | | | | | | | | |
|---|---|---|---|---|---|---|---|---|
|  | **WP_097659748.1** | 1.7E-05 | Qc.III | IQEIGGTIERLSEVSAAIAAAVEEQGAATQEISRNVQQASVGTQEV------- | *Bradyrhizobium sp. Y36* | Bacteria; Proteobacteria; Alphaproteobacteria; Rhizobiales; Bradyrhizobiaceae; Bradyrhizobium | methyl-accepting chemotaxis protein | 6 |
|  | **WP_098180699.1** | 1.7E-05 | Qa.III | -EAITRRIQQVAALVRDMSAATGEQSAGIEQINQAVTQLEE------------ | *Halomonas salina* | Bacteria; Proteobacteria; Gammaproteobacteria; Oceanospirillales; Halomonadaceae; Halomonas | methyl-accepting chemotaxis protein | 6 |
|  | **WP_098970253.1** | 1.7E-05 | Qa.III | LEQITTSINQISDMATQISSAAEEQRAVTDEVSRNIQAVK------------- | *Aeromonas sp. CU5* | Bacteria; Proteobacteria; Gammaproteobacteria; Aeromonadales; Aeromonadaceae; Aeromonas | methyl-accepting chemotaxis protein | 6 |
|  | **WP_101321019.1** | 1.7E-05 | Qa.III | LEQITTSINQISDMATQISSAAEEQRAVTDEVSRNIQAVK------------- | *Aeromonas sobria* | Bacteria; Proteobacteria; Gammaproteobacteria; Aeromonadales; Aeromonadaceae; Aeromonas | methyl-accepting chemotaxis protein | 6 |
|  | **WP_101325531.1** | 1.7E-05 | Qa.III | LEQITTSINQISDMATQISSAAEEQRAVTDEVSRNIQAVK------------- | *Aeromonas sobria* | Bacteria; Proteobacteria; Gammaproteobacteria; Aeromonadales; Aeromonadaceae; Aeromonas | methyl-accepting chemotaxis protein | 6 |
|  | **WP_101347795.1** | 1.7E-05 | Qa.III | LEQITTSINQISDMATQISSAAEEQRAVTDEVSRNIQAVK------------- | *Aeromonas sobria* | Bacteria; Proteobacteria; Gammaproteobacteria; Aeromonadales; Aeromonadaceae; Aeromonas | methyl-accepting chemotaxis protein | 6 |
|  | **WP_104618009.1** | 1.7E-05 | Qb.I | ----KNMMSESMDYIEGAANIIAQTTSNIEDVHQASDKIKDITK--------- | *Brachyspira murdochii* | Bacteria; Spirochaetes; Brachyspirales; Brachyspiraceae; Brachyspira | HAMP domain-containing protein | 6 |
|  | **WP_106887561.1** | 1.7E-05 | Qa.III | LEQITTSINQISDMATQISSAAEEQRAVTDEVSRNIQAVK------------- | *Aeromonas rivipollensis* | Bacteria; Proteobacteria; Gammaproteobacteria; Aeromonadales; Aeromonadaceae; Aeromonas | methyl-accepting chemotaxis protein | 6 |
|  | **WP_109422503.1** | 1.7E-05 | Qa.III | LEQITTSINQISDMATQISSAAEEQRAVTDEVSRNIQAVK------------- | *Aeromonas caviae* | Bacteria; Proteobacteria; Gammaproteobacteria; Aeromonadales; Aeromonadaceae; Aeromonas | methyl-accepting chemotaxis protein | 6 |
| | **WP_111355205.1** | 1.7E-05 | Qc.III | IKEIGGTIGRISEISSSIAAAIEEQGAATSEIARNVQQASTGTNHVASNIS-- | *Rhodoplanes elegans* | Bacteria; Proteobacteria; Alphaproteobacteria; Rhizobiales; Hyphomicrobiaceae; Rhodoplanes | methyl-accepting chemotaxis protein | 6 |
| | **WP_111358981.1** | 1.7E-05 | Qc.III | IKEIGATIAHIAEIAAAIAAAVEEQGAATGEIARNVQQAAKGTDAVARHITE- | *Rhodoplanes elegans* | Bacteria; Proteobacteria; Alphaproteobacteria; Rhizobiales; Hyphomicrobiaceae; Rhodoplanes | HAMP domain-containing protein | 6 |
|  | **WP_111805342.1** | 1.7E-05 | Qa.III | LEQITTSINQISDMATQISSAAEEQRAVTDEVSRNIQAVK------------- | *Aeromonas allosaccharophila* | Bacteria; Proteobacteria; Gammaproteobacteria; Aeromonadales; Aeromonadaceae; Aeromonas | HAMP domain-containing protein | 6 |
|  | **WP_111910343.1** | 1.7E-05 | Qa.III | LEQITTSINQISDMATQISSAAEEQRAVTDEVSRNIQAVK------------- | *Aeromonas media* | Bacteria; Proteobacteria; Gammaproteobacteria; Aeromonadales; Aeromonadaceae; Aeromonas | HAMP domain-containing protein | 6 |
|  | **WP_113068839.1** | 1.7E-05 | Qa.III | LEQITTSINQISDMATQISSAAEEQRAVTDEVSRNIQAVK------------- | *Aeromonas caviae* | Bacteria; Proteobacteria; Gammaproteobacteria; Aeromonadales; Aeromonadaceae; Aeromonas | methyl-accepting chemotaxis protein | 6 |
|  | **WP_113725006.1** | 1.7E-05 | Qa.III | LEQITTSINQISDMATQISSAAEEQRAVTDEVSRNIQAVK------------- | *Aeromonas* | Bacteria; Proteobacteria; Gammaproteobacteria; Aeromonadales; Aeromonadaceae | MULTISPECIES: HAMP domain-containing protein | 6 |
|  | **WP_113739701.1** | 1.7E-05 | Qa.III | LEQITTSINQISDMATQISSAAEEQRAVTDEVSRNIQAVK------------- | *Aeromonas veronii* | Bacteria; Proteobacteria; Gammaproteobacteria; Aeromonadales; Aeromonadaceae; Aeromonas | HAMP domain-containing protein | 6 |
|  | **AEF82555.1** | 1.8E-05 | SNAP.c | INEITANIQSIKGRVINQSASVTETNATMEQITVNIDKLNGHVDKQT------ | *Treponema azotonutricium ZAS-9* | Bacteria; Spirochaetes; Spirochaetales; Spirochaetaceae; Treponema | methyl-accepting chemotaxis protein | 6 |
|  | **CCD87468.1** | 1.8E-05 | Qa.IV | IKAISTTIEKLSEIASAIAAAVEQQGAATQEISRNVQQAA------------- | *Bradyrhizobium sp. ORS 285* | Bacteria; Proteobacteria; Alphaproteobacteria; Rhizobiales; Bradyrhizobiaceae; Bradyrhizobium | Methyl-accepting chemotaxis protein | 6 |
|  | **EKE67458.1** | 1.8E-05 | Qa.III.b | IKGIGATIAQINEIATTIASAVEEQGAATAEISRNVQEAAK------------ | *Oceanibaculum indicum P24* | Bacteria; Proteobacteria; Alphaproteobacteria; Rhodospirillales; Rhodospirillaceae; Oceanibaculum | methyl-accepting chemotaxis sensory transducer | 6 |
| | **KZM49176.1** | 1.8E-05 | Qc.III | IQSISEIIENMNSVAATMAASVEEQTAATSEIASNVDNASRGTNVVSENISR- | *Labrenzia sp. OB1* | Bacteria; Proteobacteria; Alphaproteobacteria; Rhodobacterales; Rhodobacteraceae; Labrenzia | hypothetical protein OA90_16635 | 6 |
|  | **OLS59240.1** | 1.8E-05 | Qb.I | LQRTKQHLQDVIEQSEVSGEELKASNEEMQAVNEELRSATEELETS------- | *Pseudomonas putida* | Bacteria; Proteobacteria; Gammaproteobacteria; Pseudomonadales; Pseudomonadaceae; Pseudomonas | Chemotaxis protein methyltransferase | 6 |
| | **OYX84968.1** | 1.8E-05 | Qc.III | LGAIHGTISNLNEVSTIISGAVEEQTAVTREVSANMHDAANGIDSVLVNMN-- | *Azorhizobium sp. 32-67-21* | Bacteria; Proteobacteria; Alphaproteobacteria; Rhizobiales; Xanthobacteraceae; Azorhizobium | chemotaxis protein | 6 |
|  | **PIE63548.1** | 1.8E-05 | Qa.III | IDEIVKIINQVNNIIKAITTAVEEQSTTTQEITTNITQAS------------- | *Desulfobacterales bacterium* | Bacteria; Proteobacteria; Deltaproteobacteria; Desulfobacterales | hypothetical protein CSA26_12475 | 6 |
|  | **PUY42940.1** | 1.8E-05 | R.IV | IDQISKAVSELDSTTQQNAALVQESSAAAASLEQQAGELSQLISRF------- | *Cronobacter sakazakii* | Bacteria; Proteobacteria; Gammaproteobacteria; Enterobacterales; Enterobacteriaceae; Cronobacter | chemotaxis protein partial | 6 |
|  | **PUY80548.1** | 1.8E-05 | R.IV | IDQISKAVSELDSTTQQNAALVQESSAAAASLEQQAGELSQLISRF------- | *Cronobacter sakazakii* | Bacteria; Proteobacteria; Gammaproteobacteria; Enterobacterales; Enterobacteriaceae; Cronobacter | chemotaxis protein partial | 6 |
|  | **SMX56391.1** | 1.8E-05 | Qa.IV | IKAISTTIEKLSEIASAIAAAVEQQGAATQEISRNVQQAA------------- | *Bradyrhizobium sp. ORS 285* | Bacteria; Proteobacteria; Alphaproteobacteria; Rhizobiales; Bradyrhizobiaceae; Bradyrhizobium | Putative Methyl-accepting chemotaxis protein | 6 |
|  | **WP_008129829.1** | 1.8E-05 | Qa.I | IRGISGTIEEVSSISNVIATAVQAQGVATAEIARNVQQTSQSVK--------- | *Bradyrhizobium sp. YR681* | Bacteria; Proteobacteria; Alphaproteobacteria; Rhizobiales; Bradyrhizobiaceae; Bradyrhizobium | methyl-accepting chemotaxis protein | 6 |
|  | **WP_010795838.1** | 1.8E-05 | Qb.I | LHRTKEQLQLTIEHSETSTEELKASNEELQAINEELRSATEELETS------- | *Pseudomonas sp. HPB0071* | Bacteria; Proteobacteria; Gammaproteobacteria; Pseudomonadales; Pseudomonadaceae; Pseudomonas | PAS domain S-box protein | 6 |
|  | **WP_011662749.1** | 1.8E-05 | Qa.IV | IKEISDTIARMSEIASTIASAVEEQGAATQEISRNVQQAA------------- | *Rhodopseudomonas palustris* | Bacteria; Proteobacteria; Alphaproteobacteria; Rhizobiales; Bradyrhizobiaceae; Rhodopseudomonas | HAMP domain-containing protein | 6 |
|  | **WP_018644608.1** | 1.8E-05 | Qa.I | IRGISGTIEEVSSISSVIATAVQAQGVATAEIARNVQQTSQSVK--------- | *Bradyrhizobium japonicum* | Bacteria; Proteobacteria; Alphaproteobacteria; Rhizobiales; Bradyrhizobiaceae; Bradyrhizobium | methyl-accepting chemotaxis protein | 6 |
|  | **WP_019367349.1** | 1.8E-05 | Qb.I | LHRTKEQLQLTIEHSETSTEELKASNEELQAINEELRSATEELETS------- | *Pseudomonas luteola* | Bacteria; Proteobacteria; Gammaproteobacteria; Pseudomonadales; Pseudomonadaceae; Pseudomonas | PAS domain S-box protein | 6 |
| | **WP_022724015.1** | 1.8E-05 | Qc.III | -REISGTIGRMSEISSTIASAVEEQGAATQEISRNVQQAAQGTMRVSSNIAD- | *Rhodopseudomonas sp. B29* | Bacteria; Proteobacteria; Alphaproteobacteria; Rhizobiales; Bradyrhizobiaceae; Rhodopseudomonas | methyl-accepting chemotaxis protein | 6 |
|  | **WP_023353963.1** | 1.8E-05 | Qb.I | -----------VDMMEDSNKMVIEQSKKVAETSEKFEEISTAVENSKVIVKEI | *Catonella morbi* | Bacteria; Firmicutes; Clostridia; Clostridiales; Lachnospiraceae; Catonella | methyl-accepting chemotaxis protein | 6 |
| | **WP_024341540.1** | 1.8E-05 | Qc.III | IQEIGGTIERLSEVSSAIAAAVEEQGAATQEISRNVQQASVGTQEVSSNIT-- | *Bradyrhizobium japonicum* | Bacteria; Proteobacteria; Alphaproteobacteria; Rhizobiales; Bradyrhizobiaceae; Bradyrhizobium | HAMP domain-containing protein | 6 |
| | **WP_028880815.1** | 1.8E-05 | Qc.III.c | --EISEVIEEINQLSAMIAAAVEQQSAATVEISNTVNNAANQVDVITSRI--- | *Terasakiella pusilla* | Bacteria; Proteobacteria; Alphaproteobacteria; Rhizobiales; Methylocystaceae; Terasakiella | methyl-accepting chemotaxis protein | 6 |
|  | **WP_039959139.1** | 1.8E-05 | Qc.III.c | LDVMVGSIARINDLGLEIATAASQQSAVTDEVSRNMTTIQTMVEELTAN---- | *Vibrio sinaloensis* | Bacteria; Proteobacteria; Gammaproteobacteria; Vibrionales; Vibrionaceae; Vibrio; Vibrio oreintalis group | HAMP domain-containing protein | 6 |
|  | **WP_040823799.1** | 1.8E-05 | Qc.III | INEITDIVGELNESASAIAAAVEEQGAATQEISGSVENAS------------- | *Thalassospira profundimaris* | Bacteria; Proteobacteria; Alphaproteobacteria; Rhodospirillales; Rhodospirillaceae; Thalassospira | methyl-accepting chemotaxis protein | 6 |
|  | **WP_041798698.1** | 1.8E-05 | Qa.III | IRDISGTIEKLSEIASTIASAVEEQGAATQEISRNVQQASQGTQQ-------- | *Rhodopseudomonas palustris* | Bacteria; Proteobacteria; Alphaproteobacteria; Rhizobiales; Bradyrhizobiaceae; Rhodopseudomonas | chemotaxis protein | 6 |
| | **WP_044407782.1** | 1.8E-05 | Qc.III | -REIGETIGRMSEISATIASAVEEQGAATQEISRNVQLAAQGTQQVSSNITD- | *Rhodopseudomonas palustris* | Bacteria; Proteobacteria; Alphaproteobacteria; Rhizobiales; Bradyrhizobiaceae; Rhodopseudomonas | methyl-accepting chemotaxis protein | 6 |
|  | **WP_051689758.1** | 1.8E-05 | Qb.I | -------LQATIERLESANEELKSSNEELMSMNEEFQSTNEELETSK------ | *Pelobacter seleniigenes* | Bacteria; Proteobacteria; Deltaproteobacteria; Desulfuromonadales; Desulfuromonadaceae; Pelobacter | PAS domain S-box protein | 6 |
|  | **WP_054203649.1** | 1.8E-05 | Qa.III | -DGILSMISDINDMNNQVAVAVEEQSTVCEDINQNITGISESSDASSQS---- | *Pseudoalteromonas* | Bacteria; Proteobacteria; Gammaproteobacteria; Alteromonadales; Pseudoalteromonadaceae | MULTISPECIES: methyl-accepting chemotaxis protein | 6 |
| | **WP_057195847.1** | 1.8E-05 | Qc.III | IQEIGGTIERLSEVSSAIAAAVEEQGAATQEISRNVQQASVGTQEVSSNIT-- | *Bradyrhizobium sp. Leaf396* | Bacteria; Proteobacteria; Alphaproteobacteria; Rhizobiales; Bradyrhizobiaceae; Bradyrhizobium | HAMP domain-containing protein | 6 |
|  | **WP_059398748.1** | 1.8E-05 | Qa.III.b | ---VARVIEQVNGIATIIASAVEQQGAATQAISRNVQQAAEVTREVARD---- | *Azospirillum* | Bacteria; Proteobacteria; Alphaproteobacteria; Rhodospirillales; Rhodospirillaceae | MULTISPECIES: HAMP domain-containing protein | 6 |
|  | **WP_063198648.1** | 1.8E-05 | Qa.IV | IKEIGATIARISDISTTIAATIEEQGAATAEIARNVSEAAK------------ | *Bradyrhizobium sp. AT1* | Bacteria; Proteobacteria; Alphaproteobacteria; Rhizobiales; Bradyrhizobiaceae; Bradyrhizobium | chemotaxis protein | 6 |
| | **WP_063682014.1** | 1.8E-05 | Qa.IV | -KEIGATIGQISTISTSIASAVEQQGAATQEIARSVQNVAQGTQAAATDIG-- | *Bradyrhizobium neotropicale* | Bacteria; Proteobacteria; Alphaproteobacteria; Rhizobiales; Bradyrhizobiaceae; Bradyrhizobium | methyl-accepting chemotaxis protein partial | 6 |
| | **WP_063682041.1** | 1.8E-05 | Qa.IV | -KEIGATIGQISTISTSIASAVEQQGAATQEIARSVQNVAQGTQAAATDIG-- | *Bradyrhizobium neotropicale* | Bacteria; Proteobacteria; Alphaproteobacteria; Rhizobiales; Bradyrhizobiaceae; Bradyrhizobium | methyl-accepting chemotaxis protein partial | 6 |
| | **WP_063699082.1** | 1.8E-05 | Qa.IV | -KEIGATIGQISTISTSIASAVEQQGAATQEIARSVQNVAQGTQAAATDIG-- | *Bradyrhizobium sp. BR 10245* | Bacteria; Proteobacteria; Alphaproteobacteria; Rhizobiales; Bradyrhizobiaceae; Bradyrhizobium | methyl-accepting chemotaxis protein | 6 |
|  | **WP_064780737.1** | 1.8E-05 | Qc.III | INEITDIVGELNESASAIAAAVEEQGAATQEISGSVENAS------------- | *Thalassospira tepidiphila* | Bacteria; Proteobacteria; Alphaproteobacteria; Rhodospirillales; Rhodospirillaceae; Thalassospira | methyl-accepting chemotaxis protein | 6 |
|  | **WP_064789488.1** | 1.8E-05 | Qc.III | INEITDIVGELNESASAIAAAVEEQGAATQEISGSVENAS------------- | *Thalassospira profundimaris* | Bacteria; Proteobacteria; Alphaproteobacteria; Rhodospirillales; Rhodospirillaceae; Thalassospira | methyl-accepting chemotaxis protein | 6 |
|  | **WP_068731097.1** | 1.8E-05 | Qa.IV | IKEIGSTITLISEVSSAIAAAVEEQGAATQEIARNVQQSAQLS---------- | *Tardiphaga robiniae* | Bacteria; Proteobacteria; Alphaproteobacteria; Rhizobiales; Bradyrhizobiaceae; Tardiphaga | methyl-accepting chemotaxis protein | 6 |
|  | **WP_073450407.1** | 1.8E-05 | Qb.I | LHRTKEQLQLTIEHSETSTEELKASNEELQAINEELRSATEELETS------- | *Pseudomonas* | Bacteria; Proteobacteria; Gammaproteobacteria; Pseudomonadales; Pseudomonadaceae | MULTISPECIES: PAS domain S-box protein | 6 |
|  | **WP_074769782.1** | 1.8E-05 | Qa.IV | ------TIETISELTSAIAAAVEEQGAATSEIARSVQQAAQGTAAAAENVE-- | *Phaeospirillum fulvum* | Bacteria; Proteobacteria; Alphaproteobacteria; Rhodospirillales; Rhodospirillaceae; Phaeospirillum | PAS domain S-box protein | 6 |
|  | **WP_074821994.1** | 1.8E-05 | Qb.I | LHRTKEQLQLTIEHSETSTEELKASNEELQAINEELRSATEELETS------- | *Pseudomonas* | Bacteria; Proteobacteria; Gammaproteobacteria; Pseudomonadales; Pseudomonadaceae | MULTISPECIES: PAS domain S-box protein | 6 |
|  | **WP_080137015.1** | 1.8E-05 | Qa.I | IRGISGTIEEVSSISNVIATAVQAQGVATAEIARNVQQTSQSVK--------- | *Bradyrhizobium sp. BR10280* | Bacteria; Proteobacteria; Alphaproteobacteria; Rhizobiales; Bradyrhizobiaceae; Bradyrhizobium | methyl-accepting chemotaxis protein | 6 |
|  | **WP_080652630.1** | 1.8E-05 | SNAP.b | --------TEMTSQVENNATKLSEQNEQFKNVFGEMREMTKMLNESVE----- | *Butyrivibrio fibrisolvens* | Bacteria; Firmicutes; Clostridia; Clostridiales; Lachnospiraceae; Butyrivibrio | hypothetical protein | 6 |
|  | **WP_080659121.1** | 1.8E-05 | SNAP.b | --------TEMTSQVENNATKLSEQNEQFKNVFGEMREMTKMLNESVE----- | *Butyrivibrio fibrisolvens* | Bacteria; Firmicutes; Clostridia; Clostridiales; Lachnospiraceae; Butyrivibrio | hypothetical protein | 6 |
|  | **WP_080666209.1** | 1.8E-05 | SNAP.b | --------TEMTSQVENNATKLSEQNEQFKNVFGEMREMTKMLNESVE----- | *Butyrivibrio fibrisolvens* | Bacteria; Firmicutes; Clostridia; Clostridiales; Lachnospiraceae; Butyrivibrio | hypothetical protein | 6 |
|  | **WP_081357228.1** | 1.8E-05 | SNAP.b | --------TEMTSQVENNATKLSEQNEQFKNVFGEMREMTKMLNESVE----- | *Butyrivibrio fibrisolvens* | Bacteria; Firmicutes; Clostridia; Clostridiales; Lachnospiraceae; Butyrivibrio | hypothetical protein | 6 |
|  | **WP_081373989.1** | 1.8E-05 | SNAP.b | --------TEMTSQVENNATKLSEQNEQFKNVFGEMREMTKMLNESVE----- | *Butyrivibrio fibrisolvens* | Bacteria; Firmicutes; Clostridia; Clostridiales; Lachnospiraceae; Butyrivibrio | hypothetical protein | 6 |
|  | **WP_083952753.1** | 1.8E-05 | Qc.III | INEITDIVGELNESASAIAAAVEEQGAATQEISGSVENAS------------- | *Thalassospira* | Bacteria; Proteobacteria; Alphaproteobacteria; Rhodospirillales; Rhodospirillaceae | MULTISPECIES: methyl-accepting chemotaxis protein | 6 |
|  | **WP_085643909.1** | 1.8E-05 | Qa.III | ISSIAETIEKINEISSSIAAAVEQQQVATGEIARNVDEAAK------------ | *Thalassospira sp. MCCC 1A03138* | Bacteria; Proteobacteria; Alphaproteobacteria; Rhodospirillales; Rhodospirillaceae; Thalassospira | methyl-accepting chemotaxis protein | 6 |

| | | | | | | | |
|---|---|---|---|---|---|---|---|
| **WP_089264646.1** | 1.8E-05 | Qa.IV | IKEIGSTITLISEVSSAIAAAVEEQGAATQEIARNVQQSAQLS---------- | *Tardiphaga sp. OK246* | Bacteria; Proteobacteria; Alphaproteobacteria; Rhizobiales; Bradyrhizobiaceae; Tardiphaga | methyl-accepting chemotaxis protein | 6 |
| **WP_090645848.1** | 1.8E-05 | Qa.III | IKGIGETIKHMNEIASSIADSVHQQGDATNEIARNVNEAS------------- | *Asticcacaulis taihuensis* | Bacteria; Proteobacteria; Alphaproteobacteria; Caulobacterales; Caulobacteraceae; Asticcacaulis | methyl-accepting chemotaxis protein | 6 |
| **WP_092032204.1** | 1.8E-05 | Qc.III | -QTIGGIIGEVNEVATAIAAAVQEQGAATQEITRSTQFAAQGTKNVTD----- | *Bradyrhizobium sp. OK095* | Bacteria; Proteobacteria; Alphaproteobacteria; Rhizobiales; Bradyrhizobiaceae; Bradyrhizobium | methyl-accepting chemotaxis protein | 6 |
| **WP_092144596.1** | 1.8E-05 | Qa.IV | IKEIGSTITLISEVSSAIAAAVEEQGAATQEIARNVQQSAQLS---------- | *Bradyrhizobium sp. NFR13* | Bacteria; Proteobacteria; Alphaproteobacteria; Rhizobiales; Bradyrhizobiaceae; Bradyrhizobium | methyl-accepting chemotaxis protein | 6 |
| **WP_093759058.1** | 1.8E-05 | Qa.IV | IKEIGSTITLISEVSSAIAAAVEEQGAATQEIARNVQQSAQLS---------- | *Tardiphaga sp. OK245* | Bacteria; Proteobacteria; Alphaproteobacteria; Rhizobiales; Bradyrhizobiaceae; Tardiphaga | methyl-accepting chemotaxis protein | 6 |
| **WP_106951444.1** | 1.8E-05 | Qc.III | IQEIGGTIERLSEVSSAIAAAVEEQGAATQEISRNVQQASVGTQEVSSNIT-- | *Bradyrhizobium sp. MOS003* | Bacteria; Proteobacteria; Alphaproteobacteria; Rhizobiales; Bradyrhizobiaceae; Bradyrhizobium | HAMP domain-containing protein | 6 |
| **WP_111068655.1** | 1.8E-05 | Qc.III | IEGIGQTITTINEIAASISAAIEEQNATTGEITRNVQQAAQGTEEVSG----- | *Azospirillum sp. M2T2B2* | Bacteria; Proteobacteria; Alphaproteobacteria; Rhodospirillales; Rhodospirillaceae; Azospirillum | methyl-accepting chemotaxis protein | 6 |
| **ALA16729.1** | 1.9E-05 | Qa.III | IDSITETIRQIDEVSTTIASAVDQQGATTNEIARNVSQAAQGTQEV------- | *Chelatococcus sp. CO-6* | Bacteria; Proteobacteria; Alphaproteobacteria; Rhizobiales; Chelatococcaceae; Chelatococcus | hypothetical protein AL346_04000 | 6 |
| **APF38437.1** | 1.9E-05 | Qa.III | IDSITETIRQIDEVSTTIASAVDQQGATTNEIARNVSQAAQGTQEV------- | *Chelatococcus daeguensis* | Bacteria; Proteobacteria; Alphaproteobacteria; Rhizobiales; Chelatococcaceae; Chelatococcus | hypothetical protein BOQ54_14845 | 6 |
| **CCJ72211.1** | 1.9E-05 | R.IV | IDQISKAVSELDSTTQQNAALVQESSAAAASLEQQAGELSQLISLFK------ | *Cronobacter condimenti 1330* | Bacteria; Proteobacteria; Gammaproteobacteria; Enterobacterales; Enterobacteriaceae; Cronobacter | Methyl-accepting chemotaxis protein I (serine chemoreceptor protein) | 6 |
| **EGP06262.1** | 1.9E-05 | Qc.III | IEAISQTISRINEISAAIAASVEEQASVTQSISSNMQVAAKGVSDINSSMT-- | *Bradyrhizobiaceae bacterium SG-6C* | Bacteria; Proteobacteria; Alphaproteobacteria; Rhizobiales; Bradyrhizobiaceae | putative PAS/PAC sensor protein | 6 |
| **KUJ95765.1** | 1.9E-05 | Qa.III | -ESITQKVKEVNDLANTIASATEEQTVTMADITTNVRRTTEAAQKVK------ | *Desulfonauticus sp. 38_4375* | Bacteria; Proteobacteria; Deltaproteobacteria; Desulfovibrionales; Desulfohalobiaceae; Desulfonauticus | Methyl-accepting chemotaxis sensory transducer | 6 |
| **KZE34272.1** | 1.9E-05 | Qa.III | IDSITETIRQIDEVSTTIASAVDQQGATTNEIARNVSQAAQGTQEV------- | *Chelatococcus daeguensis* | Bacteria; Proteobacteria; Alphaproteobacteria; Rhizobiales; Chelatococcaceae; Chelatococcus | hypothetical protein AVW15_17305 | 6 |
| **ODU66786.1** | 1.9E-05 | Qb.I | LRRTREQLQMTVEQSETSGEELKASNEELQSINEELRSATEELETS------- | *Rhodanobacter sp. SCN 65-17* | Bacteria; Proteobacteria; Gammaproteobacteria; Xanthomonadales; Rhodanobacteraceae; Rhodanobacter | chemotaxis protein | 6 |
| **PSV60228.1** | 1.9E-05 | Qc.III | LDKIVEQVQKINDMNFHIASAAEQQTAVADEMNQNLTNVR------------- | *Photobacterium profundum* | Bacteria; Proteobacteria; Gammaproteobacteria; Vibrionales; Vibrionaceae; Photobacterium | methyl-accepting chemotaxis protein | 6 |
| **PYM55089.1** | 1.9E-05 | Qb.I | LAATREYLQSVIEQQEVANEELQSSNEEVQSANEELQSVNEELETSK------ | *Candidatus Rokubacteria bacterium* | Bacteria; Candidatus Rokubacteria | hypothetical protein DMD77_20955 partial | 6 |
| **SCC81049.1** | 1.9E-05 | Qa.IV | IQDITTTIDEMSELASSIASAVTQQSAATQEIASNVQEAAN------------ | *Salinarimonadaceae bacterium HL-109* | Bacteria; Proteobacteria; Alphaproteobacteria; Rhizobiales; Salinarimonadaceae | methyl-accepting chemotaxis protein | 6 |
| **SDO96914.1** | 1.9E-05 | Qc.III | -REISGTIERLSEISSTIAAAVEEQGAATQEISRNVQQAAQGTQQVSANIT-- | *Afipia sp. GAS231* | Bacteria; Proteobacteria; Alphaproteobacteria; Rhizobiales; Bradyrhizobiaceae; Afipia | methyl-accepting chemotaxis sensory transducer | 6 |
| **SMX61719.1** | 1.9E-05 | Qa.IV | -----GTIRQISTIASSIATAVEQQGGATQEIARNVQSVSAGTQAAASDI--- | *Bradyrhizobium sp. ORS 285* | Bacteria; Proteobacteria; Alphaproteobacteria; Rhizobiales; Bradyrhizobiaceae; Bradyrhizobium | putative methyl-accepting chemotaxis protein | 6 |
| **WP_002713101.1** | 1.9E-05 | Qc.III | IEAISQTISRINEISAAIAASVEEQASVTQSISSNMQVAAKGVSDINSSMT-- | *Afipia clevelandensis* | Bacteria; Proteobacteria; Alphaproteobacteria; Rhizobiales; Bradyrhizobiaceae; Afipia | methyl-accepting chemotaxis protein | 6 |
| **WP_005345507.1** | 1.9E-05 | Qc.III | LDGMADAVVRINDLSSQIATAAEEQSRVTEEINRNMVSIRDMLHQL------- | *Aeromonas diversa* | Bacteria; Proteobacteria; Gammaproteobacteria; Aeromonadales; Aeromonadaceae; Aeromonas | methyl-accepting chemotaxis protein | 6 |
| **WP_008962171.1** | 1.9E-05 | Qa.IV | -----GTIRQISTIASSIATAVEQQGGATQEIAKNVQGVAAGTQAAASDI--- | *Bradyrhizobium sp. STM 3809* | Bacteria; Proteobacteria; Alphaproteobacteria; Rhizobiales; Bradyrhizobiaceae; Bradyrhizobium | methyl-accepting chemotaxis protein | 6 |
| **WP_011500586.1** | 1.9E-05 | Qc.III | IKEISGTIERLSEISATIASAVEEQGAATLEISRNVQQAAQGTQRVSSNIAD- | *Rhodopseudomonas palustris* | Bacteria; Proteobacteria; Alphaproteobacteria; Rhizobiales; Bradyrhizobiaceae; Rhodopseudomonas | methyl-accepting chemotaxis protein | 6 |
| **WP_015664186.1** | 1.9E-05 | Qa.III | IKEISGTIEKLSEIASTIAAAVEEQGAATQEISRNVQQAA------------- | *Bradyrhizobium oligotrophicum* | Bacteria; Proteobacteria; Alphaproteobacteria; Rhizobiales; Bradyrhizobiaceae; Bradyrhizobium | HAMP domain-containing protein | 6 |
| **WP_022694480.1** | 1.9E-05 | Qc.III.c | --AISKIIEDVNAISVSISSAVDQQAAVTQRISAHMTEATNAVTSVNSGMG-- | *Ponticaulis koreensis* | Bacteria; Proteobacteria; Alphaproteobacteria; Rhodobacterales; Hyphomonadaceae; Ponticaulis | PAS domain S-box protein | 6 |
| **WP_028346433.1** | 1.9E-05 | Qa.III | IEAIGKTIGEISEIATTIASAIEQQGAATQEIARNVQQAA------------- | *Bradyrhizobium elkanii* | Bacteria; Proteobacteria; Alphaproteobacteria; Rhizobiales; Bradyrhizobiaceae; Bradyrhizobium | HAMP domain-containing protein | 6 |
| **WP_028794058.1** | 1.9E-05 | Qa.III | IQTIVSSIAEVSSVTGSIAAAVEQQSSATNEISRNVQEAS------------- | *Thalassobaculum salexigens* | Bacteria; Proteobacteria; Alphaproteobacteria; Rhodospirillales; Rhodospirillaceae; Thalassobaculum | hypothetical protein | 6 |
| **WP_043924117.1** | 1.9E-05 | Qc.III | IEAISQTISRINEISAAIAASVEEQASVTQSISSNMQVAAKGVSDINSSMT-- | *Bradyrhizobiaceae bacterium SG-6C* | Bacteria; Proteobacteria; Alphaproteobacteria; Rhizobiales; Bradyrhizobiaceae | methyl-accepting chemotaxis protein | 6 |
| **WP_054164102.1** | 1.9E-05 | Qc.III | IKEIGGTIGRMSEISSTIASAVEEQGAATQEISRNVSQAAQGTQQVSSNIT-- | *Rhodopseudomonas sp. AAP120* | Bacteria; Proteobacteria; Alphaproteobacteria; Rhizobiales; Bradyrhizobiaceae; Rhodopseudomonas | methyl-accepting chemotaxis protein | 6 |
| **WP_054164853.1** | 1.9E-05 | Qa.III | IREISGTIEKLSEIASTIASAVEEQGAATQEISRNVQQAAQGTQQ-------- | *Rhodopseudomonas sp. AAP120* | Bacteria; Proteobacteria; Alphaproteobacteria; Rhizobiales; Bradyrhizobiaceae; Rhodopseudomonas | chemotaxis protein | 6 |
| **WP_069649851.1** | 1.9E-05 | Qb.III | ---IENLALDMEERGKTIKNVLDKQRDMIKEIIDTVANIDNYMK--------- | *Caloranaerobacter ferrireducens* | Bacteria; Firmicutes; Clostridia; Clostridiales; Clostridiaceae; Caloranaerobacter | hypothetical protein | 6 |
| **WP_073611979.1** | 1.9E-05 | Qa.III | IREISAVVNEVNDIVSTIASAVEEQSATTNEIAGNVSQAS------------- | *Desulfopila aestuarii* | Bacteria; Proteobacteria; Deltaproteobacteria; Desulfobacterales; Desulfobulbaceae; Desulfopila | methyl-accepting chemotaxis protein | 6 |
| **WP_083204486.1** | 1.9E-05 | Qa.IV | IQDITTTIDEMSELASSIASAVTQQSAATQEIASNVQEAAN------------ | *Salinarimonadaceae bacterium HL-109* | Bacteria; Proteobacteria; Alphaproteobacteria; Rhizobiales; Salinarimonadaceae | hypothetical protein | 6 |
| **WP_083228332.1** | 1.9E-05 | Qa.III | -----SSISKVNEMNLEVASAATQQSSVVEEMNVNVNRINEKAEASSEQ---- | *Enterovibrio pacificus* | Bacteria; Proteobacteria; Gammaproteobacteria; Vibrionales; Vibrionaceae; Enterovibrio | methyl-accepting chemotaxis protein | 6 |
| **WP_085552326.1** | 1.9E-05 | Qc.III | --SIGQTIARMSEIATAIASAVEEQGAATRDIANSVTQAAQGTDEVSSNI--- | *Azospirillum lipoferum* | Bacteria; Proteobacteria; Alphaproteobacteria; Rhodospirillales; Rhodospirillaceae; Azospirillum | methyl-accepting chemotaxis protein | 6 |
| **WP_089265832.1** | 1.9E-05 | Qc.III | IQEIGDTIGRMSEIASTIASAVEEQGAATQEISRNVQQAAQGTMQVSSNI--- | *Tardiphaga sp. OK246* | Bacteria; Proteobacteria; Alphaproteobacteria; Rhizobiales; Bradyrhizobiaceae; Tardiphaga | methyl-accepting chemotaxis protein | 6 |
| **WP_091727685.1** | 1.9E-05 | Qa.IV | -NDITLQINEIANFINDISKNIDEQNRATESLAKNVENAASFIGD-------- | *Proteiniborus ethanoligenes* | Bacteria; Firmicutes; Clostridia; Clostridiales; Proteiniborus | hypothetical protein | 6 |
| **WP_092040604.1** | 1.9E-05 | Qc.III | --SIARIVGEIATSSTAIAAAIEEQGAATQEIARNVQQASSGTDRVTGNI--- | *Methylobacterium pseudosasicola* | Bacteria; Proteobacteria; Alphaproteobacteria; Rhizobiales; Methylobacteriaceae; Methylobacterium | HAMP domain-containing protein | 6 |
| **WP_092518642.1** | 1.9E-05 | Qc.III | -REISGTIERLSEISSTIAAAVEEQGAATQEISRNVQQAAQGTQQVSANIT-- | *Afipia sp. GAS231* | Bacteria; Proteobacteria; Alphaproteobacteria; Rhizobiales; Bradyrhizobiaceae; Afipia | methyl-accepting chemotaxis protein | 6 |
| **WP_092681494.1** | 1.9E-05 | Qc.III | IKEISGTIERLSEISATIASAVEEQGAATLEISRNVQQAAQGTQRVSSNIAD- | *Rhodopseudomonas pseudopalustris* | Bacteria; Proteobacteria; Alphaproteobacteria; Rhizobiales; Bradyrhizobiaceae; Rhodopseudomonas | methyl-accepting chemotaxis protein | 6 |
| **WP_093150820.1** | 1.9E-05 | Qa.III | IQTIVSSIAEVSSVTGSIAAAVEQQSSATNEISRNVQEAS------------- | *Thalassobaculum litoreum* | Bacteria; Proteobacteria; Alphaproteobacteria; Rhodospirillales; Rhodospirillaceae; Thalassobaculum | hypothetical protein | 6 |
| **WP_107243201.1** | 1.9E-05 | Qc.III | LDKIVEQVQKINDMNFHIASAAEQQTAVADEMNQNLTNVR------------- | *Photobacterium frigidiphilum* | Bacteria; Proteobacteria; Gammaproteobacteria; Vibrionales; Vibrionaceae; Photobacterium | methyl-accepting chemotaxis protein | 6 |
| **WP_107255508.1** | 1.9E-05 | Qc.III | LDKIVEQVQKINDMNFHIASAAEQQTAVADEMNQNLTNVR------------- | *Photobacterium indicum* | Bacteria; Proteobacteria; Gammaproteobacteria; Vibrionales; Vibrionaceae; Photobacterium | methyl-accepting chemotaxis protein | 6 |
| **WP_109794640.1** | 1.9E-05 | Qa.IV | IRKVTGAVKELNGIASTIASAVEEQTAVTRDIARNVQQAA------------- | *Rhizobiales bacterium* | Bacteria; Proteobacteria; Alphaproteobacteria; Rhizobiales | HAMP domain-containing protein | 6 |
| **WP_110163001.1** | 1.9E-05 | R.IV | IDQIGKAVTELDSTTQQNAALVQESSAAASSLEQQADQLSQLISVFR------ | *Pectobacterium carotovorum* | Bacteria; Proteobacteria; Gammaproteobacteria; Enterobacterales; Pectobacteriaceae; Pectobacterium | HAMP domain-containing protein | 6 |
| **ANK81587.1** | 2E-05 | Qa.IV | IRKVTGAVKELNGIASTIASAVEEQTAVTRDIARNVQQAA------------- | *Rhizobiales bacterium NRL2* | Bacteria; Proteobacteria; Alphaproteobacteria; Rhizobiales | hypothetical protein TEF_12880 | 6 |
| **SDS63217.1** | 2E-05 | Qc.III | IKEISGTIEKLSEISSTIAAAVEEQGAATQEISRNVQQAAQGTQQVSSNITD- | *Bradyrhizobium canariense* | Bacteria; Proteobacteria; Alphaproteobacteria; Rhizobiales; Bradyrhizobiaceae; Bradyrhizobium | methyl-accepting chemotaxis sensory transducer | 6 |
| **SHO44039.1** | 2E-05 | Qa.III | IREISAVVNEVNDIVSTIASAVEEQSATTNEIAGNVSQAS------------- | *Desulfopila aestuarii DSM 18488* | Bacteria; Proteobacteria; Deltaproteobacteria; Desulfobacterales; Desulfobulbaceae; Desulfopila | methyl-accepting chemotaxis sensory transducer with Cache sensor | 6 |
| **WP_005434270.1** | 2E-05 | Qa.III.b | LDKINSAVAMISDMATQIATAAEEQTHVTNEITQNVTSIK------------- | *Vibrio harveyi* | Bacteria; Proteobacteria; Gammaproteobacteria; Vibrionales; Vibrionaceae; Vibrio | methyl-accepting chemotaxis protein | 6 |
| **WP_012978285.1** | 2E-05 | Qa.III | IRSIAGTIREVNEIAATIAAAVEQQSAATREIARNVGEAANST---------- | *Azospirillum lipoferum* | Bacteria; Proteobacteria; Alphaproteobacteria; Rhodospirillales; Rhodospirillaceae; Azospirillum | HAMP domain-containing protein | 6 |
| **WP_013651549.1** | 2E-05 | Qa.IV | IRAIAATMAEVNEYTGAIAASVTEQGAATNEISANIQSAAQRSET-------- | *Polymorphum gilvum* | Bacteria; Proteobacteria; Alphaproteobacteria; Polymorphum | HAMP domain-containing protein | 6 |
| **WP_020592698.1** | 2E-05 | Qc.III | IEGISKIINQVNEIASGISAAVEEQGAATQEIARNTQEAARGTQEVSSNI--- | *Kiloniella laminariae* | Bacteria; Proteobacteria; Alphaproteobacteria; Kiloniellales; Kiloniellaceae; Kiloniella | methyl-accepting chemotaxis protein | 6 |
| **WP_023456604.1** | 2E-05 | Qa.III | IRGIGETIKHMNEIASSIADSVHQQGDATNEIARNVNEAS------------- | *Asticcacaulis sp. AC466* | Bacteria; Proteobacteria; Alphaproteobacteria; Caulobacterales; Caulobacteraceae; Asticcacaulis | methyl-accepting chemotaxis protein | 6 |
| **WP_035016319.1** | 2E-05 | Qb.I | --------ATAIEQQSTASQSISRSTQALSSTSEELKNYSDTAGETFALLHK- | *Catenovulum agarivorans* | Bacteria; Proteobacteria; Gammaproteobacteria; Alteromonadales; Alteromonadaceae; Catenovulum | chemotaxis protein | 6 |
| **WP_035962311.1** | 2E-05 | Qc.III | IKSISGTIERLSEISSTIAAAVEEQGAATQEIARNVQQASQGTQQVSSNI--- | *Bradyrhizobium sp. URHA0013* | Bacteria; Proteobacteria; Alphaproteobacteria; Rhizobiales; Bradyrhizobiaceae; Bradyrhizobium | methyl-accepting chemotaxis protein | 6 |
| **WP_044536660.1** | 2E-05 | Qa.III | IKEIGNTIAKMSEIASTIASAVEEQGAATREISRNVQQAAHGTQQVS------ | *Bradyrhizobium sp. LTSP885* | Bacteria; Proteobacteria; Alphaproteobacteria; Rhizobiales; Bradyrhizobiaceae; Bradyrhizobium | HAMP domain-containing protein | 6 |
| **WP_044540097.1** | 2E-05 | Qc.III | IKEIGDTIGRMSEISSTIAAAVEEQGAATQEISRNIQNAAQGTVQVSGNIA-- | *Bradyrhizobium sp. LTSP885* | Bacteria; Proteobacteria; Alphaproteobacteria; Rhizobiales; Bradyrhizobiaceae; Bradyrhizobium | HAMP domain-containing protein | 6 |
| **WP_045782829.1** | 2E-05 | R.IV | -QEINHAVNHLDSMVQQNAELVVESAAAAGALQSQAGDLAETAGHFR------ | *Klebsiella michiganensis* | Bacteria; Proteobacteria; Gammaproteobacteria; Enterobacterales; Enterobacteriaceae; Klebsiella | methyl-accepting chemotaxis protein | 6 |
| **WP_046868825.1** | 2E-05 | Qa.IV | IQQIAATIAEMSQISISIAAAMEEQGAATSEIARNVQEAAKGTEQVTG----- | *Microvirga massiliensis* | Bacteria; Proteobacteria; Alphaproteobacteria; Rhizobiales; Methylobacteriaceae; Microvirga | HAMP domain-containing protein | 6 |

| | | | | | | | | |
|---|---|---|---|---|---|---|---|---|
| | **WP_069958264.1** | 2E-05 | Qc.III | INSIGGIINQMNEIASTIAAAVEEQGAATQEIARNVEQAAAGTGEVSSNI--- | *Magnetovibrio blakemorei* | Bacteria; Proteobacteria; Alphaproteobacteria; Rhodospirillales; Rhodospirillaceae; Magnetovibrio | HAMP domain-containing protein | 6 |
| | **WP_074826578.1** | 2E-05 | Qc.III | IKEISGTIEKLSEISSTIAAAVEEQGAATQEISRNVQQAAQGTQQVSSNITD- | *Bradyrhizobium* | Bacteria; Proteobacteria; Alphaproteobacteria; Rhizobiales; Bradyrhizobiaceae | MULTISPECIES: methyl-accepting chemotaxis protein | 6 |
| | **WP_079567872.1** | 2E-05 | Qc.III | IKEISGTIEKLSEISSTIAAAVEEQGAATQEISRNVQQAAQGTQQVSSNITD- | *Bradyrhizobium erythrophlei* | Bacteria; Proteobacteria; Alphaproteobacteria; Rhizobiales; Bradyrhizobiaceae; Bradyrhizobium | methyl-accepting chemotaxis protein | 6 |
| | **WP_079586711.1** | 2E-05 | Qc.III | IKEISGTIEKLSEISSTIAAAVEEQGAATQEISRNVQQAAQGTQQVSSNITD- | *Bradyrhizobium lablabi* | Bacteria; Proteobacteria; Alphaproteobacteria; Rhizobiales; Bradyrhizobiaceae; Bradyrhizobium | methyl-accepting chemotaxis protein | 6 |
| | **WP_099954445.1** | 2E-05 | Qa.III | LRSIERSVAEAQSMVNGIASAAEEQSAVTQEISSAMQCASRGLTEVNQSLV-- | *Methylobacterium sp. PR1016A* | Bacteria; Proteobacteria; Alphaproteobacteria; Rhizobiales; Methylobacteriaceae; Methylobacterium | methyl-accepting chemotaxis protein | 6 |
| | **WP_100380697.1** | 2E-05 | Qc.III | IKEISGTIEKLSEISSTIAAAVEEQGAATQEISRNVQQAAQGTQQVSSNITD- | *Afipia broomeae* | Bacteria; Proteobacteria; Alphaproteobacteria; Rhizobiales; Bradyrhizobiaceae; Afipia | methyl-accepting chemotaxis protein | 6 |
|  | **WP_100861287.1** | 2E-05 | Qa.III | LVQITTSINQISDMATQISSAAEEQRAVTDEVSRNIQAVK------------ | *Aeromonas lusitana* | Bacteria; Proteobacteria; Gammaproteobacteria; Aeromonadales; Aeromonadaceae; Aeromonas | methyl-accepting chemotaxis protein | 6 |
|  | **WP_101263799.1** | 2E-05 | Qc.III | -RDISDVISRTRETASSISSAVDQQNSATQEIARNVSQASAGTQEVS------ | *Thalassospira marina* | Bacteria; Proteobacteria; Alphaproteobacteria; Rhodospirillales; Rhodospirillaceae; Thalassospira | methyl-accepting chemotaxis protein | 6 |
|  | **WP_101283730.1** | 2E-05 | Qc.III | -RDISDVISRTRETASSISSAVDQQNSATQEIARNVSQASAGTQEVS------ | *Thalassospira marina* | Bacteria; Proteobacteria; Alphaproteobacteria; Rhodospirillales; Rhodospirillaceae; Thalassospira | methyl-accepting chemotaxis protein | 6 |
| | **WP_108174472.1** | 2E-05 | Qc.III | -GDISGTIMRISEISTAIATAVEEQSAVTAEMSSNMRVAAEGVQAISHSMKE- | *Phreatobacter oligotrophus* | Bacteria; Proteobacteria; Alphaproteobacteria; Phreatobacter | PAS domain S-box protein | 6 |
|  | **WP_109118793.1** | 2E-05 | Qa.IV | IQAITGTIGDISQVSAAIASAVEQQGASTSEIARNVEQAA------------ | *Azospirillum sp. TSO22-1* | Bacteria; Proteobacteria; Alphaproteobacteria; Rhodospirillales; Rhodospirillaceae; Azospirillum | methyl-accepting chemotaxis protein | 6 |
|  | **WP_109516551.1** | 2E-05 | Qa.III | IQGIAGTIGEVNQIAGAIAAAVEEQGVATREIARNVAQAA------------ | *Roseomonas aestuarii* | Bacteria; Proteobacteria; Alphaproteobacteria; Rhodospirillales; Acetobacteraceae; Roseomonas | methyl-accepting chemotaxis protein | 6 |
|  | **BAM87517.1** | 2.1E-05 | Qa.I | VEAIARTIAEIDEIASTISAAVEQQGAATKNIAHNVQEAS------------ | *Bradyrhizobium oligotrophicum S58* | Bacteria; Proteobacteria; Alphaproteobacteria; Rhizobiales; Bradyrhizobiaceae; Bradyrhizobium | hypothetical protein S58_15090 | 6 |
|  | **GAJ37937.1** | 2.1E-05 | Qa.IV | IQSISQTIQRVSQIASVIAAAVEEQGAATREIARNVQQTS------------ | *Bradyrhizobium sp. DOA9* | Bacteria; Proteobacteria; Alphaproteobacteria; Rhizobiales; Bradyrhizobiaceae; Bradyrhizobium | probable chemoreceptor Y4FA | 6 |
|  | **KFG69471.1** | 2.1E-05 | Qa.IV | IQEIARTISEMSQISTSISAAMEQQGSATAEIARSVQEAAR----------- | *Microvirga sp. BSC39* | Bacteria; Proteobacteria; Alphaproteobacteria; Rhizobiales; Methylobacteriaceae; Microvirga | hypothetical protein JH26_10150 | 6 |
|  | **KXJ56645.1** | 2.1E-05 | Qa.III | LHTIHNEVTSISDATSQIAAAVEQQSVVTEQVSQNIERINELT---------- | *Colwellia sp. Phe_37* | Bacteria; Proteobacteria; Gammaproteobacteria; Alteromonadales; Colwelliaceae; Colwellia | chemotaxis protein | 6 |
| | **KZD25048.1** | 2.1E-05 | Qc.III | IKEIGDTIGRMSEIASTIASAVEEQGAATQEISRNVQQAAQGTMQVSSNI--- | *Tardiphaga robiniae* | Bacteria; Proteobacteria; Alphaproteobacteria; Rhizobiales; Bradyrhizobiaceae; Tardiphaga | chemotaxis protein | 6 |
|  | **OUX99907.1** | 2.1E-05 | Qa.III | ISAIAKVIEDVSSISVSISGAVEEQAVVTRNISANMSEATNAVT--------- | *Hyphomonadaceae bacterium TMED5* | Bacteria; Proteobacteria; Alphaproteobacteria; Rhodobacterales; Hyphomonadaceae | chemotaxis protein | 6 |
|  | **PVV07881.1** | 2.1E-05 | Qa.III | --EISNAIDTINNMNTQIATAAEQQCSVVDEINRNIASIVENSRQTSQ----- | *gamma proteobacterium symbiont of Ctena orbiculata* | Bacteria; Proteobacteria; Gammaproteobacteria | hypothetical protein B6D72_17625 | 6 |
| | **PVX48604.1** | 2.1E-05 | Qc.III | IKEIGDTIGRMSEIASTIASAVEEQGAATQEISRNVQQAAQGTMQVSSNI--- | *Tardiphaga sp. OV697* | Bacteria; Proteobacteria; Alphaproteobacteria; Rhizobiales; Bradyrhizobiaceae; Tardiphaga | methyl-accepting chemotaxis sensory transducer | 6 |
|  | **RAI25823.1** | 2.1E-05 | Qc.III | IKEISGTIGQIAEVASAIAAAVEEQGAATAEISRNIQQAASGT---------- | *Rhodoplanes elegans* | Bacteria; Proteobacteria; Alphaproteobacteria; Rhizobiales; Hyphomicrobiaceae; Rhodoplanes | chemotaxis protein partial | 6 |
| | **SDT60787.1** | 2.1E-05 | Qc.III | IKEISGTIEKLSEISSTIAAAVEEQGAATQEISRNVQQAAQGTQQVSSNITD- | *Bradyrhizobium canariense* | Bacteria; Proteobacteria; Alphaproteobacteria; Rhizobiales; Bradyrhizobiaceae; Bradyrhizobium | methyl-accepting chemotaxis sensory transducer | 6 |
|  | **SEE39299.1** | 2.1E-05 | Qa.IV | -RAIGSTIERLSEISSAIAAGIEEQGAATQEISRNVQQAAQ----------- | *Bradyrhizobium erythrophlei* | Bacteria; Proteobacteria; Alphaproteobacteria; Rhizobiales; Bradyrhizobiaceae; Bradyrhizobium | methyl-accepting chemotaxis sensory transducer | 6 |
|  | **SMX55864.1** | 2.1E-05 | Qc.III | IQRIGGIIGEVNEVATAIAAAVQEQGAATQEISRSTQFAADGTKNVS------ | *Bradyrhizobium sp. ORS 285* | Bacteria; Proteobacteria; Alphaproteobacteria; Rhizobiales; Bradyrhizobiaceae; Bradyrhizobium | putative methyl-accepting chemotaxis receptor/sensory transducer | 6 |
| | **SMX57839.1** | 2.1E-05 | Qc.III | IQAISSTIHRVNQIAAAIAAAVEQQGAATREIARNVQQASEGTNAVSHNIS-- | *Bradyrhizobium sp. ORS 285* | Bacteria; Proteobacteria; Alphaproteobacteria; Rhizobiales; Bradyrhizobiaceae; Bradyrhizobium | putative methyl-accepting chemotaxis receptor/sensory transducer | 6 |
| | **WP_008135826.1** | 2.1E-05 | Qc.III | ---IGGTIERLSEISSTIAAAVEEQGAATSEISRNVQNAAAGTTQVSANI--- | *Bradyrhizobium sp. YR681* | Bacteria; Proteobacteria; Alphaproteobacteria; Rhizobiales; Bradyrhizobiaceae; Bradyrhizobium | methyl-accepting chemotaxis protein | 6 |
| | **WP_008965076.1** | 2.1E-05 | Qc.III | --AISGTIERLSEISSTIAAAVEEQGAATQEISRNVQQAAQGTQQVSSNIAE- | *Bradyrhizobium sp. STM 3809* | Bacteria; Proteobacteria; Alphaproteobacteria; Rhizobiales; Bradyrhizobiaceae; Bradyrhizobium | HAMP domain-containing protein | 6 |
|  | **WP_011924218.1** | 2.1E-05 | Qc.III | IQRIGGIIGEVNEVATAIAAAVQEQGAATQEISRSTQFAADGTKNVS------ | *Bradyrhizobium sp. ORS 278* | Bacteria; Proteobacteria; Alphaproteobacteria; Rhizobiales; Bradyrhizobiaceae; Bradyrhizobium | methyl-accepting chemotaxis protein | 6 |
| | **WP_021133299.1** | 2.1E-05 | Qc.III | --RIISVIGEINEMSSAIAGAVEEQGAAMQEVVRNVEEAAKGTREVARSI--- | *Phaeospirillum fulvum* | Bacteria; Proteobacteria; Alphaproteobacteria; Rhodospirillales; Rhodospirillaceae; Phaeospirillum | HAMP domain-containing protein | 6 |
|  | **WP_027317021.1** | 2.1E-05 | Qc.III | IENIAATITRINEISAAIAAAVEQQAAVTQKMSANMQAAAEG----------- | *Microvirga flocculans* | Bacteria; Proteobacteria; Alphaproteobacteria; Rhizobiales; Methylobacteriaceae; Microvirga | methyl-accepting chemotaxis protein | 6 |
| | **WP_028144446.1** | 2.1E-05 | Qc.III | IKEISGTIRRISEISSAIAAAVEEQGMATQEIARNVQQAAQGTHEVASNIT-- | *Bradyrhizobium* | Bacteria; Proteobacteria; Alphaproteobacteria; Rhizobiales; Bradyrhizobiaceae | MULTISPECIES: methyl-accepting chemotaxis protein | 6 |
|  | **WP_042338923.1** | 2.1E-05 | Qa.I | VEAIARTIAEIDEIASTISAAVEQQGAATKNIAHNVQEAS------------ | *Bradyrhizobium oligotrophicum* | Bacteria; Proteobacteria; Alphaproteobacteria; Rhizobiales; Bradyrhizobiaceae; Bradyrhizobium | chemotaxis protein | 6 |
|  | **WP_051284069.1** | 2.1E-05 | Qa.III | IESISKVIHDVNDIVATIATAVEEQSATTKEIAGNIIQVSKE---------- | *Desulforegula conservatrix* | Bacteria; Proteobacteria; Deltaproteobacteria; Desulfobacterales; Desulfobacteraceae; Desulforegula | methyl-accepting chemotaxis protein | 6 |
|  | **WP_058076441.1** | 2.1E-05 | Qc.III.c | LREIAASVAVVGQMSQQIAAAAEEQSAVAEQVGQSMARVS------------ | *Pseudomonas sp. TTU2014-066ASC* | Bacteria; Proteobacteria; Gammaproteobacteria; Pseudomonadales; Pseudomonadaceae; Pseudomonas | methyl-accepting chemotaxis protein | 6 |
| | **WP_074116913.1** | 2.1E-05 | Qa.IV | -KEIGATIGQISTISTSIASAVEQQGAATQEIARSVQNVAQGTQAAASDI--- | *Bradyrhizobium sp. AS23.2* | Bacteria; Proteobacteria; Alphaproteobacteria; Rhizobiales; Bradyrhizobiaceae; Bradyrhizobium | methyl-accepting chemotaxis protein | 6 |
|  | **WP_080134976.1** | 2.1E-05 | Qc.III | -QAIGGIIGEVNEVATAIAAAVQEQGAATQEITRSTQFAAQGTKNVS------ | *Bradyrhizobium sp. BR10280* | Bacteria; Proteobacteria; Alphaproteobacteria; Rhizobiales; Bradyrhizobiaceae; Bradyrhizobium | methyl-accepting chemotaxis protein | 6 |
|  | **WP_081928884.1** | 2.1E-05 | Qa.IV | IQEIARTISEMSQISTSISAAMEQQGSATAEIARSVQEAAR----------- | *Microvirga sp. BSC39* | Bacteria; Proteobacteria; Alphaproteobacteria; Rhizobiales; Methylobacteriaceae; Microvirga | methyl-accepting chemotaxis protein | 6 |
|  | **WP_082312407.1** | 2.1E-05 | Qa.III | IDSITETIRQIDEVSTTIASAVDQQGATTNEIARNVSQAAQGTQEV------- | *Chelatococcus* | Bacteria; Proteobacteria; Alphaproteobacteria; Rhizobiales; Chelatococcaceae | MULTISPECIES: methyl-accepting chemotaxis protein | 6 |
|  | **WP_082464674.1** | 2.1E-05 | Qa.III | IDSITETIRQIDEVSTTIASAVDQQGATTNEIARNVSQAAQGTQEV------- | *Chelatococcus* | Bacteria; Proteobacteria; Alphaproteobacteria; Rhizobiales; Chelatococcaceae | MULTISPECIES: methyl-accepting chemotaxis protein | 6 |
|  | **WP_082831613.1** | 2.1E-05 | Qa.III | IDSITETIRQIDEVSTTIASAVDQQGATTNEIARNVSQAAQGTQEV------- | *Chelatococcus daeguensis* | Bacteria; Proteobacteria; Alphaproteobacteria; Rhizobiales; Chelatococcaceae; Chelatococcus | methyl-accepting chemotaxis protein | 6 |
| | **WP_083921939.1** | 2.1E-05 | Qc.III | LEAIGNSIREVSTISNAISIAVEEQAAATQEIARNVEQASAGTQQVSRSIED- | *Kiloniella laminariae* | Bacteria; Proteobacteria; Alphaproteobacteria; Kiloniellales; Kiloniellaceae; Kiloniella | methyl-accepting chemotaxis protein | 6 |
|  | **WP_085589171.1** | 2.1E-05 | Qa.III | IQGIGATIDEINEIAANIASAVEQQGAATGEITRNIQQAS------------ | *Thalassospira sp. MCCC 1A01428* | Bacteria; Proteobacteria; Alphaproteobacteria; Rhodospirillales; Rhodospirillaceae; Thalassospira | methyl-accepting chemotaxis protein | 6 |
| | **WP_085902146.1** | 2.1E-05 | Qc.III | --GISKTIDSIHEVATAIASAVEEQGAATAEITKSVQQAANGTDQVSSNIS-- | *Kiloniella majae* | Bacteria; Proteobacteria; Alphaproteobacteria; Kiloniellales; Kiloniellaceae; Kiloniella | methyl-accepting chemotaxis protein | 6 |
| | **WP_085902147.1** | 2.1E-05 | Qc.III | --GISKTIDSIHEVATAIASAVEEQGAATAEITKSVQQAANGTDQVSSNIS-- | *Kiloniella majae* | Bacteria; Proteobacteria; Alphaproteobacteria; Kiloniellales; Kiloniellaceae; Kiloniella | methyl-accepting chemotaxis protein | 6 |
| | **WP_085904858.1** | 2.1E-05 | Qc.III | --GISKTIDSIHEVATAIASAVEEQGAATAEITKSVQQAANGTDQVSSNIS-- | *Kiloniella majae* | Bacteria; Proteobacteria; Alphaproteobacteria; Kiloniellales; Kiloniellaceae; Kiloniella | methyl-accepting chemotaxis protein | 6 |
| | **WP_085908091.1** | 2.1E-05 | Qc.III | --GISKTIDSIHEVATAIASAVEEQGAATAEITKSVQQAANGTDQVSSNIS-- | *Kiloniella majae* | Bacteria; Proteobacteria; Alphaproteobacteria; Kiloniellales; Kiloniellaceae; Kiloniella | methyl-accepting chemotaxis protein | 6 |
| | **WP_085908092.1** | 2.1E-05 | Qc.III | --GISKTIDSIHEVATAIASAVEEQGAATAEITKSVQQAANGTDQVSSNIS-- | *Kiloniella majae* | Bacteria; Proteobacteria; Alphaproteobacteria; Kiloniellales; Kiloniellaceae; Kiloniella | methyl-accepting chemotaxis protein | 6 |
|  | **WP_090272429.1** | 2.1E-05 | Qa.III | LEQIATSMQQLVSLNASIASATTQQSTVVEDVNRNVTEAAEL---------- | *Pseudomonas litoralis* | Bacteria; Proteobacteria; Gammaproteobacteria; Pseudomonadales; Pseudomonadaceae; Pseudomonas | methyl-accepting chemotaxis protein | 6 |
|  | **WP_091944611.1** | 2.1E-05 | Qa.III | IRAITATIGNLSEISLSVSSAVTEQAAVTHEISNNMQVAARNVDA-------- | *Methylorubrum salsuginis* | Bacteria; Proteobacteria; Alphaproteobacteria; Rhizobiales; Methylobacteriaceae; Methylorubrum | methyl-accepting chemotaxis protein | 6 |
|  | **WP_092037178.1** | 2.1E-05 | Qc.III | IDSIQGTIVKLNEVATAISSAVEQQSAVTREMSASMQTAAQG---------- | *Methylobacterium pseudosasicola* | Bacteria; Proteobacteria; Alphaproteobacteria; Rhizobiales; Methylobacteriaceae; Methylobacterium | methyl-accepting chemotaxis protein | 6 |
|  | **WP_092125056.1** | 2.1E-05 | Qa.IV | -RAIGSTIERLSEISSAIAAGIEEQGAATQEISRNVQQAAQ----------- | *Bradyrhizobium erythrophlei* | Bacteria; Proteobacteria; Alphaproteobacteria; Rhizobiales; Bradyrhizobiaceae; Bradyrhizobium | methyl-accepting chemotaxis protein | 6 |
| | **WP_093757182.1** | 2.1E-05 | Qc.III | IKEIGDTIGRMSEIASTIASAVEEQGAATQEISRNVQQAAQGTMQVSSNI--- | *Tardiphaga sp. OK245* | Bacteria; Proteobacteria; Alphaproteobacteria; Rhizobiales; Bradyrhizobiaceae; Tardiphaga | methyl-accepting chemotaxis protein | 6 |
| | **WP_093757183.1** | 2.1E-05 | Qc.III | IKEIGDTIGRMSEIASTIASAVEEQGAATQEISRNVQQAAQGTMQVSSNI--- | *Tardiphaga sp. OK245* | Bacteria; Proteobacteria; Alphaproteobacteria; Rhizobiales; Bradyrhizobiaceae; Tardiphaga | methyl-accepting chemotaxis protein | 6 |
| | **WP_100383017.1** | 2.1E-05 | Qc.III | IKEISGTIEKLSEISSTIAAAVEEQGAATQEISRNVQQAAQGTQQVSSNITD- | *Afipia broomeae* | Bacteria; Proteobacteria; Alphaproteobacteria; Rhizobiales; Bradyrhizobiaceae; Afipia | methyl-accepting chemotaxis protein | 6 |
|  | **WP_103179942.1** | 2.1E-05 | R.IV | IQEINRAVLNLDSMVQQNAELVVQSAAAASALQGQAGDLAETAGHFR------ | *Leclercia* | Bacteria; Proteobacteria; Gammaproteobacteria; Enterobacterales; Enterobacteriaceae | MULTISPECIES: methyl-accepting chemotaxis protein | 6 |
|  | **WP_103822283.1** | 2.1E-05 | R.IV | IQEINRAVLNLDSMVQQNAELVVQSAAAASALQGQAGDLAETAGHFR------ | *Leclercia* | Bacteria; Proteobacteria; Gammaproteobacteria; Enterobacterales; Enterobacteriaceae | MULTISPECIES: methyl-accepting chemotaxis protein | 6 |
|  | **WP_108660384.1** | 2.1E-05 | Qc.III | IEKIGATIDSINEITTAISAAVEEQSAATSEISRNVTQAAT----------- | *Acuticoccus sp. J103* | Bacteria; Proteobacteria; Alphaproteobacteria; Rhodobacterales; Rhodobacteraceae; Acuticoccus | HAMP domain-containing protein | 6 |

| | | | | | | | | |
|---|---|---|---|---|---|---|---|---|
| | WP_109442871.1 | 2.1E-05 | Qc.III | -REIGSTIASINAVSAEIAATVDRQNEVAQTIAESVRAAAQESDRVNANVAD- | *Azospirillum sp. TSH100* | Bacteria; Proteobacteria; Alphaproteobacteria; Rhodospirillales; Rhodospirillaceae; Azospirillum | methyl-accepting chemotaxis protein | 6 |
|  | WP_109596109.1 | 2.1E-05 | Qc.III | -ETITATVMRINAISMAISQAVGEQSAVTQAMSTTMRTASQ------------ | *Microvirga sp. 17 mud 1-3* | Bacteria; Proteobacteria; Alphaproteobacteria; Rhizobiales; Methylobacteriaceae; Microvirga | methyl-accepting chemotaxis protein | 6 |
|  | WP_111360019.1 | 2.1E-05 | Qc.III | IKEIGGTIGRIAEITATIAAAVEEQGAATSEIARNVQQAASGTTEV------- | *Rhodoplanes elegans* | Bacteria; Proteobacteria; Alphaproteobacteria; Rhizobiales; Hyphomicrobiaceae; Rhodoplanes | methyl-accepting chemotaxis protein partial | 6 |
|  | CBS85984.1 | 2.2E-05 | Qa.IV | IQSIATTIGEVNDIVGTIAQAVDGQNAATMEIARNVQQAA------------- | *Azospirillum lipoferum 4B* | Bacteria; Proteobacteria; Alphaproteobacteria; Rhodospirillales; Rhodospirillaceae; Azospirillum | methyl-accepting chemotaxis protein | 6 |
|  | EGF91253.1 | 2.2E-05 | Qa.III | IRGIGETIKHMNEIASSIADSVHQQGDATNEIARNVNEAS------------- | *Asticcacaulis biprosthecum C19* | Bacteria; Proteobacteria; Alphaproteobacteria; Caulobacterales; Caulobacteraceae; Asticcacaulis | methyl-accepting chemotaxis protein MCP signaling domain protein | 6 |
|  | EKM29670.1 | 2.2E-05 | Qa.III | LDEINSAVALISDMATQIATAAEEQTHVTNEITQNVTSIK------------- | *Vibrio harveyi* | Bacteria; Proteobacteria; Gammaproteobacteria; Vibrionales; Vibrionaceae; Vibrio | HAMP domain protein | 6 |
|  | EQI16701.1 | 2.2E-05 | Qc.I | -----DMAGELRGMYEHLESQVQERTVDLREANALLERQRDKLKQLNAD---- | *Clostridioides difficile F501* | Bacteria; Firmicutes; Clostridia; Clostridiales; Peptostreptococcaceae; Clostridioides | UNVERIFIED_ORG: HAMP domain protein | 6 |
|  | KFG69743.1 | 2.2E-05 | Qa.IV | IQEIGRTIAEMSQISVSIAAAMEQQGAATTEIARNVQQAA------------- | *Microvirga sp. BSC39* | Bacteria; Proteobacteria; Alphaproteobacteria; Rhizobiales; Methylobacteriaceae; Microvirga | hypothetical protein JH26_08765 | 6 |
| | KZD25863.1 | 2.2E-05 | Qc.III | IKEIGDTIGRMSEIASTIASAVEQQGAATQEISRNVQQAAQGTMQVTSNITD- | *Tardiphaga robiniae* | Bacteria; Proteobacteria; Alphaproteobacteria; Rhizobiales; Bradyrhizobiaceae; Tardiphaga | chemotaxis protein | 6 |
|  | OQX26920.1 | 2.2E-05 | Qb.I | ---TKENLHATIEEMQAANEELKSGNEELQSTNEELQSTNEEMETSK------ | *Desulfobacteraceae bacterium IS3* | Bacteria; Proteobacteria; Deltaproteobacteria; Desulfobacterales; Desulfobacteraceae | SAM-dependent methyltransferase | 6 |
|  | PHS78355.1 | 2.2E-05 | Qc.III | -GSIGGTINELNEISAAIAAAVEEQGAATQEISRNVEQAAQ------------ | *Rhodospirillaceae bacterium* | Bacteria; Proteobacteria; Alphaproteobacteria; Rhodospirillales; Rhodospirillaceae | hypothetical protein COB59_07530 | 6 |
|  | PNB53033.1 | 2.2E-05 | Qc.III | IKEIGATIGKISSISTSIASAVEEQGAATQEIARSVQNVA------------- | *Pseudomonas sp. GW531-E2* | Bacteria; Proteobacteria; Gammaproteobacteria; Pseudomonadales; Pseudomonadaceae; Pseudomonas | methyl-accepting chemotaxis protein partial | 6 |
|  | PWC65304.1 | 2.2E-05 | Qc.III | --GIADTVLQVNQIAATITSAVEEQTAATGEISRNVQQAAQGTSEVSR----- | *Azospirillum sp. TSH20* | Bacteria; Proteobacteria; Alphaproteobacteria; Rhodospirillales; Rhodospirillaceae; Azospirillum | hypothetical protein TSH20_16705 | 6 |
| | RAI20059.1 | 2.2E-05 | Qa.III | --SIAHTIQNLSEVSSGIAAAVEQQTAVTREIAQNMQVASTGVEQISQSIS-- | *Rhodoblastus acidophilus* | Bacteria; Proteobacteria; Alphaproteobacteria; Rhizobiales; Beijerinckiaceae; Rhodoblastus | chemotaxis protein | 6 |
| | RAI34639.1 | 2.2E-05 | Qc.III | IKEIGATIAHIAEIAAAIAAAVEEQGAATGEIARNVQQAAKGTDAVARHITE- | *Rhodoplanes elegans* | Bacteria; Proteobacteria; Alphaproteobacteria; Rhizobiales; Hyphomicrobiaceae; Rhodoplanes | hypothetical protein CH338_20575 | 6 |
|  | RCK17700.1 | 2.2E-05 | Qa.III | LDEINSAVALISDMATQIATAAEEQTHVTNEITQNVTSIK------------- | *Vibrio harveyi* | Bacteria; Proteobacteria; Gammaproteobacteria; Vibrionales; Vibrionaceae; Vibrio | methyl-accepting chemotaxis protein | 6 |
| | SOE00788.1 | 2.2E-05 | Qc.III | IQSINKVIDRMSEISVAVSAAVEEQNAATAEIARSVDQTSQGTKEVSRSI--- | *Caenispirillum bisanense* | Bacteria; Proteobacteria; Alphaproteobacteria; Rhodospirillales; Rhodospirillaceae; Caenispirillum | methyl-accepting chemotaxis sensory transducer with Cache sensor | 6 |
|  | WP_003235671.1 | 2.2E-05 | Qb.I | LQRTKLHLQDTIEQAEVSSEELKASNEEMQAVNEELRSATEELETS------- | *Pseudomonas fluorescens* | Bacteria; Proteobacteria; Gammaproteobacteria; Pseudomonadales; Pseudomonadaceae; Pseudomonas | PAS domain S-box protein | 6 |
|  | WP_005449641.1 | 2.2E-05 | Qa.III | LDEINSAVALISDMATQIATAAEEQTHVTNEITQNVTSIK------------- | *Vibrio harveyi* | Bacteria; Proteobacteria; Gammaproteobacteria; Vibrionales; Vibrionaceae; Vibrio | methyl-accepting chemotaxis protein | 6 |
|  | WP_006610421.1 | 2.2E-05 | Qa.III.b | IQGIGGTIGQISEIAKTIATAVDQQGHATQEIARSVSQAAS------------ | *Bradyrhizobium sp. ORS 285* | Bacteria; Proteobacteria; Alphaproteobacteria; Rhizobiales; Bradyrhizobiaceae; Bradyrhizobium | PAS domain S-box protein | 6 |
|  | WP_008962152.1 | 2.2E-05 | Qa.III.b | IQGIGGTIGQISEIAKTIATAVDQQGHATQEIARSVSQAAS------------ | *Bradyrhizobium sp. STM 3809* | Bacteria; Proteobacteria; Alphaproteobacteria; Rhizobiales; Bradyrhizobiaceae; Bradyrhizobium | PAS domain S-box protein | 6 |
|  | WP_009026655.1 | 2.2E-05 | Qa.III.b | IQGIGGTIGQISEIAKTIATAVDQQGHATQEIARSVSQAAS------------ | *Bradyrhizobium sp. ORS 375* | Bacteria; Proteobacteria; Alphaproteobacteria; Rhizobiales; Bradyrhizobiaceae; Bradyrhizobium | methyl-accepting chemotaxis protein | 6 |
| | WP_011500764.1 | 2.2E-05 | Qc.III | IKEIGQTIGRMSEIASTIASAVEEQGAATQEISRNVQQAAQGTQQVSANIT-- | *Rhodopseudomonas* | Bacteria; Proteobacteria; Alphaproteobacteria; Rhizobiales; Bradyrhizobiaceae | MULTISPECIES: methyl-accepting chemotaxis protein | 6 |
|  | WP_011926775.1 | 2.2E-05 | Qa.III.b | IQGIGGTIGQISEIAKTIATAVDQQGHATQEIARSVSQAAS------------ | *Bradyrhizobium sp. ORS 278* | Bacteria; Proteobacteria; Alphaproteobacteria; Rhizobiales; Bradyrhizobiaceae; Bradyrhizobium | PAS domain S-box protein | 6 |
|  | WP_015371798.1 | 2.2E-05 | Qb.I | LQRTKLHLQDTIEQAEVSSEELKASNEEMQAVNEELRSATEELETS------- | *Pseudomonas poae* | Bacteria; Proteobacteria; Gammaproteobacteria; Pseudomonadales; Pseudomonadaceae; Pseudomonas | PAS domain S-box protein | 6 |
|  | WP_017189861.1 | 2.2E-05 | Qa.III | LDEINSAVALISDMATQIATAAEEQTHVTNEITQNVTSIK------------- | *Vibrio harveyi* | Bacteria; Proteobacteria; Gammaproteobacteria; Vibrionales; Vibrionaceae; Vibrio | methyl-accepting chemotaxis protein | 6 |
|  | WP_023450447.1 | 2.2E-05 | Qa.III | IRGIGETIKHMNEIASSIADSVHQQGDATNEIARNVNEAS------------- | *Asticcacaulis sp. AC402* | Bacteria; Proteobacteria; Alphaproteobacteria; Caulobacterales; Caulobacteraceae; Asticcacaulis | methyl-accepting chemotaxis protein | 6 |
|  | WP_023454302.1 | 2.2E-05 | Qa.III | IRGIGETIKHMNEIASSIADSVHQQGDATNEIARNVNEAS------------- | *Asticcacaulis sp. AC460* | Bacteria; Proteobacteria; Alphaproteobacteria; Caulobacterales; Caulobacteraceae; Asticcacaulis | methyl-accepting chemotaxis protein | 6 |
|  | WP_042337061.1 | 2.2E-05 | Qa.IV | IQSISQTIQRVSQIASVIAAAVEEQGAATREIARNVQQTS------------- | *Bradyrhizobium sp. DOA9* | Bacteria; Proteobacteria; Alphaproteobacteria; Rhizobiales; Bradyrhizobiaceae; Bradyrhizobium | methyl-accepting chemotaxis protein | 6 |
|  | WP_045446543.1 | 2.2E-05 | Qa.IV | IREVTVTVSEMREIAQTIASSVEQQQSATVEISRNVQEAAQ------------ | *Tepidicaulis marinus* | Bacteria; Proteobacteria; Alphaproteobacteria; Rhizobiales; Rhodobiaceae; Tepidicaulis | methyl-accepting chemotaxis protein | 6 |
|  | WP_047514733.1 | 2.2E-05 | Qa.III | LDEINSAVALISDMATQIATAAEEQTHVTNEITQNVTSIK------------- | *Vibrio harveyi* | Bacteria; Proteobacteria; Gammaproteobacteria; Vibrionales; Vibrionaceae; Vibrio | methyl-accepting chemotaxis protein | 6 |
|  | WP_065869816.1 | 2.2E-05 | Qb.I | LQRTKLHLQDTIEQAEVSSEELKASNEEMQAVNEELRSATEELETS------- | *Pseudomonas sp. 25 E 4* | Bacteria; Proteobacteria; Gammaproteobacteria; Pseudomonadales; Pseudomonadaceae; Pseudomonas | PAS domain S-box protein | 6 |
|  | WP_069217894.1 | 2.2E-05 | Qa.III | LDEINSAVALISDMATQIATAAEEQTHVTNEITQNVTSIK------------- | *Vibrio harveyi* | Bacteria; Proteobacteria; Gammaproteobacteria; Vibrionales; Vibrionaceae; Vibrio | methyl-accepting chemotaxis protein | 6 |
|  | WP_076954919.1 | 2.2E-05 | Qb.I | LQRTKLHLQDTIEQAEVSSEELKASNEEMQAVNEELRSATEELETS------- | *Pseudomonas cedrina* | Bacteria; Proteobacteria; Gammaproteobacteria; Pseudomonadales; Pseudomonadaceae; Pseudomonas | PAS domain S-box protein | 6 |
|  | WP_085319691.1 | 2.2E-05 | Qa.III | LDEINSAVALISDMATQIATAAEEQTHVTNEITQNVTSIK------------- | *Vibrio harveyi* | Bacteria; Proteobacteria; Gammaproteobacteria; Vibrionales; Vibrionaceae; Vibrio | methyl-accepting chemotaxis protein | 6 |
| | WP_089266235.1 | 2.2E-05 | Qc.III | IKEIGDTIGRMSEIASTIASAVEQQGAATQEISRNVQQAAQGTMQVTSNITD- | *Tardiphaga sp. OK246* | Bacteria; Proteobacteria; Alphaproteobacteria; Rhizobiales; Bradyrhizobiaceae; Tardiphaga | methyl-accepting chemotaxis protein | 6 |
| | WP_092146204.1 | 2.2E-05 | Qc.III | IKEIGDTIGRMSEIASTIASAVEQQGAATQEISRNVQQAAQGTMQVTSNITD- | *Bradyrhizobium sp. NFR13* | Bacteria; Proteobacteria; Alphaproteobacteria; Rhizobiales; Bradyrhizobiaceae; Bradyrhizobium | methyl-accepting chemotaxis protein | 6 |
| | WP_092478076.1 | 2.2E-05 | Qc.III | -DRLVVSASSTASKAQVLNENANLQTEKLSAVDESFSRVSSSVDDVTNSVKE- | *[Clostridium] polysaccharolyticum* | Bacteria; Firmicutes; Clostridia; Clostridiales; Lachnospiraceae | hypothetical protein | 6 |
| | WP_093757370.1 | 2.2E-05 | Qc.III | IKEIGDTIGRMSEIASTIASAVEQQGAATQEISRNVQQAAQGTMQVTSNITD- | *Tardiphaga sp. OK245* | Bacteria; Proteobacteria; Alphaproteobacteria; Rhizobiales; Bradyrhizobiaceae; Tardiphaga | methyl-accepting chemotaxis protein | 6 |
|  | WP_109153225.1 | 2.2E-05 | Qa.III | IRNISRTITQVNEISSTIAAAVEEQSAATKEIGRNVQQAAQGTQEVST----- | *Azospirillum sp. TSO5* | Bacteria; Proteobacteria; Alphaproteobacteria; Rhodospirillales; Rhodospirillaceae; Azospirillum | methyl-accepting chemotaxis protein | 6 |
|  | WP_111066825.1 | 2.2E-05 | Qa.III | IEAIAKTIGQMNEIATGIASAIEEQNAATQNIAGNITQAA------------- | *Azospirillum sp. M2T2B2* | Bacteria; Proteobacteria; Alphaproteobacteria; Rhodospirillales; Rhodospirillaceae; Azospirillum | HAMP domain-containing protein | 6 |
|  | WP_111571813.1 | 2.2E-05 | Qa.I | ------QVEEVSSIIDTLAEDITKVSNNTESMAGQSKNVMKNIKEGNQSLK-- | *Halanaerobium saccharolyticum* | Bacteria; Firmicutes; Clostridia; Halanaerobiales; Halanaerobiaceae; Halanaerobium | methyl-accepting chemotaxis protein | 6 |
|  | AHY49192.1 | 2.3E-05 | Qc.III | IQEIGGTIERLSEVSAAIAAAVEEQGAATQEISRNVQQASVGTQ--------- | *Bradyrhizobium japonicum SEMIA 5079* | Bacteria; Proteobacteria; Alphaproteobacteria; Rhizobiales; Bradyrhizobiaceae; Bradyrhizobium | chemotaxis protein | 6 |
| | AMW33868.1 | 2.3E-05 | Qc.III | ---VIETIGRLNQISAAIASAMEEQDAATREIARNVEEASNGTGEVTRNVR-- | *Haematospirillum jordaniae* | Bacteria; Proteobacteria; Alphaproteobacteria; Rhodospirillales; Rhodospirillaceae; Haematospirillum | hypothetical protein AY555_00320 | 6 |
| | KEO53376.1 | 2.3E-05 | Qc.III | -RRISETIAKVADSSTAIAAAVEEQHAAIGEIARNVEEAANGTREVSERIE-- | *Thalassospira permensis NBRC 106175* | Bacteria; Proteobacteria; Alphaproteobacteria; Rhodospirillales; Rhodospirillaceae; Thalassospira | hypothetical protein SMB34_21220 partial | 6 |
|  | KGH32042.1 | 2.3E-05 | Qa.III | MKEIMQSVGQVTGLLSEISGALHQQSEGI----AHVNTAVAHMDSTNQE---- | *Comamonas testosteroni* | Bacteria; Proteobacteria; Betaproteobacteria; Burkholderiales; Comamonadaceae; Comamonas | methyl-accepting chemotaxis protein | 6 |
| | OSI57798.1 | 2.3E-05 | Qa.IV | ---IGATIGQISSISTSIASAVEQQGAATQEIARSVQTVAQGTQTAATDI--- | *Bradyrhizobium canariense* | Bacteria; Proteobacteria; Alphaproteobacteria; Rhizobiales; Bradyrhizobiaceae; Bradyrhizobium | methyl-accepting chemotaxis protein partial | 6 |
| | OSI58814.1 | 2.3E-05 | Qa.IV | ---IGATIGQISSISTSIASAVEQQGAATQEIARSVQTVAQGTQTAATDI--- | *Bradyrhizobium canariense* | Bacteria; Proteobacteria; Alphaproteobacteria; Rhizobiales; Bradyrhizobiaceae; Bradyrhizobium | methyl-accepting chemotaxis protein partial | 6 |
| | OSI62809.1 | 2.3E-05 | Qa.IV | ---IGATIGQISSISTSIASAVEQQGAATQEIARSVQTVAQGTQTAATDI--- | *Bradyrhizobium canariense* | Bacteria; Proteobacteria; Alphaproteobacteria; Rhizobiales; Bradyrhizobiaceae; Bradyrhizobium | methyl-accepting chemotaxis protein partial | 6 |
|  | PRW78745.1 | 2.3E-05 | Qb.I | LQRTKLHLQDTIEQAEVSSEELKASNEEMQAVNEELRSATEELETS------- | *Pseudomonas fluorescens* | Bacteria; Proteobacteria; Gammaproteobacteria; Pseudomonadales; Pseudomonadaceae; Pseudomonas | PAS domain S-box protein | 6 |
| | PWC84292.1 | 2.3E-05 | Qc.III | IEEISRIITEISQISATIASAVEQQSAATQEISRNVQEAAQGTQQVTSNI--- | *Azospirillum sp. TSH100* | Bacteria; Proteobacteria; Alphaproteobacteria; Rhodospirillales; Rhodospirillaceae; Azospirillum | chemotaxis protein partial | 6 |
|  | RAK09784.1 | 2.3E-05 | Qa.I | ------QVEEVSSIIDTLAEDITKVSNNTESMAGQSKNVMKNIKEGNQSLK-- | *Halanaerobium saccharolyticum* | Bacteria; Firmicutes; Clostridia; Halanaerobiales; Halanaerobiaceae; Halanaerobium | methyl-accepting chemotaxis protein | 6 |
|  | SDI66988.1 | 2.3E-05 | Qc.III | IQEIGGTIERLSEVSAAIAAAVEEQGAATQEISRNVQQASVGTQ--------- | *Bradyrhizobium sp. Rc2d* | Bacteria; Proteobacteria; Alphaproteobacteria; Rhizobiales; Bradyrhizobiaceae; Bradyrhizobium | methyl-accepting chemotaxis protein | 6 |
| | SEH57234.1 | 2.3E-05 | Qc.III | IKEIGDTIGRMSEIASTIASAVEQQGAATQEISRNVQQAAQGTMQVTSNITD- | *Tardiphaga sp. OK245* | Bacteria; Proteobacteria; Alphaproteobacteria; Rhizobiales; Bradyrhizobiaceae; Tardiphaga | methyl-accepting chemotaxis sensory transducer | 6 |
| | SFL81492.1 | 2.3E-05 | Qc.III | IKEIGDTIGRMSEIASTIASAVEQQGAATQEISRNVQQAAQGTMQVTSNITD- | *Bradyrhizobium sp. NFR13* | Bacteria; Proteobacteria; Alphaproteobacteria; Rhizobiales; Bradyrhizobiaceae; Bradyrhizobium | methyl-accepting chemotaxis sensory transducer | 6 |
| | SFU08402.1 | 2.3E-05 | Qa.IV | ---ITAAVKEISELTSSISDAVEQQQRATAEIAEAVQAASDGTAKAASNV--- | *Pseudovibrio denitrificans* | Bacteria; Proteobacteria; Alphaproteobacteria; Rhodobacterales; Rhodobacteraceae; Pseudovibrio | methyl-accepting chemotaxis protein | 6 |
| | SNT49876.1 | 2.3E-05 | Qc.III | IKEIGDTIGRMSEIASTIASAVEQQGAATQEISRNVQQAAQGTMQVTSNITD- | *Tardiphaga sp. OK246* | Bacteria; Proteobacteria; Alphaproteobacteria; Rhizobiales; Bradyrhizobiaceae; Tardiphaga | methyl-accepting chemotaxis sensory transducer | 6 |
|  | WP_003190419.1 | 2.3E-05 | Qb.I | LQRTKLHLQDTIEQAEVSSEELKASNEEMQAVNEELRSATEELETS------- | *Pseudomonas fluorescens* | Bacteria; Proteobacteria; Gammaproteobacteria; Pseudomonadales; Pseudomonadaceae; Pseudomonas | PAS domain S-box protein | 6 |

| | | | | | | | | |
|---|---|---|---|---|---|---|---|---|
| | **WP_008970722.1** | 2.3E-05 | Qc.III | -REISGTIERLSEISSTIAAAVEQQGAATQEISRNVQQAARGTHEVSSNIT-- | *Bradyrhizobium sp. STM 3843* | Bacteria; Proteobacteria; Alphaproteobacteria; Rhizobiales; Bradyrhizobiaceae; Bradyrhizobium | HAMP domain-containing protein | 6 |
|  | **WP_014497317.1** | 2.3E-05 | Qc.III | IQEIGGTIERLSEVSAAIAAAVEEQGAATQEISRNVQQASVGTQ--------- | *Bradyrhizobium japonicum* | Bacteria; Proteobacteria; Alphaproteobacteria; Rhizobiales; Bradyrhizobiaceae; Bradyrhizobium | methyl-accepting chemotaxis protein | 6 |
|  | **WP_014718014.1** | 2.3E-05 | Qb.I | LQRTKLHLQDTIEQAEVSSEELKASNEEMQAVNEELRSATEELETS------- | *Pseudomonas fluorescens* | Bacteria; Proteobacteria; Gammaproteobacteria; Pseudomonadales; Pseudomonadaceae; Pseudomonas | PAS domain S-box protein | 6 |
|  | **WP_019821469.1** | 2.3E-05 | Qb.I | LQRTKLHLQDTIEQAEVSSEELKASNEEMQAVNEELRSATEELETS------- | *Pseudomonas* | Bacteria; Proteobacteria; Gammaproteobacteria; Pseudomonadales; Pseudomonadaceae | MULTISPECIES: PAS domain S-box protein | 6 |
|  | **WP_020303206.1** | 2.3E-05 | Qb.I | LQRTKLHLQDTIEQAEVSSEELKASNEEMQAVNEELRSATEELETS------- | *Pseudomonas* | Bacteria; Proteobacteria; Gammaproteobacteria; Pseudomonadales; Pseudomonadaceae | MULTISPECIES: PAS domain S-box protein | 6 |
|  | **WP_023536165.1** | 2.3E-05 | Qb.I | LQRTKLHLQETIEQSEVSSEELKASNEEMQAINQELRSATEELETS------- | *Pseudomonas putida* | Bacteria; Proteobacteria; Gammaproteobacteria; Pseudomonadales; Pseudomonadaceae; Pseudomonas | PAS domain S-box protein | 6 |
|  | **WP_028158171.1** | 2.3E-05 | Qc.III | IQEIGGTIERLSEVSAAIAAAVEEQGAATQEISRNVQQASVGTQ--------- | *Bradyrhizobium japonicum* | Bacteria; Proteobacteria; Alphaproteobacteria; Rhizobiales; Bradyrhizobiaceae; Bradyrhizobium | methyl-accepting chemotaxis protein | 6 |
|  | **WP_028170144.1** | 2.3E-05 | Qc.III | IQEIGGTIERLSEVSAAIAAAVEEQGAATQEISRNVQQASVGTQ--------- | *Bradyrhizobium japonicum* | Bacteria; Proteobacteria; Alphaproteobacteria; Rhizobiales; Bradyrhizobiaceae; Bradyrhizobium | methyl-accepting chemotaxis protein | 6 |
|  | **WP_032903489.1** | 2.3E-05 | Qb.I | LQRTKLHLQDTIEQAEVSSEELKASNEEMQAVNEELRSATEELETS------- | *Pseudomonas fluorescens* | Bacteria; Proteobacteria; Gammaproteobacteria; Pseudomonadales; Pseudomonadaceae; Pseudomonas | PAS domain S-box protein | 6 |
|  | **WP_034127092.1** | 2.3E-05 | Qb.I | LQRTKLHLQDTIEQAEVSSEELKASNEEMQAVNEELRSATEELETS------- | *Pseudomonas fluorescens* | Bacteria; Proteobacteria; Gammaproteobacteria; Pseudomonadales; Pseudomonadaceae; Pseudomonas | PAS domain S-box protein | 6 |
|  | **WP_034135305.1** | 2.3E-05 | Qb.I | LQRTKLHLQDTIEQAEVSSEELKASNEEMQAVNEELRSATEELETS------- | *Pseudomonas* | Bacteria; Proteobacteria; Gammaproteobacteria; Pseudomonadales; Pseudomonadaceae | MULTISPECIES: PAS domain S-box protein | 6 |
|  | **WP_038449742.1** | 2.3E-05 | Qb.I | LQRTKLHLQDTIEQAEVSSEELKASNEEMQAVNEELRSATEELETS------- | *Pseudomonas* | Bacteria; Proteobacteria; Gammaproteobacteria; Pseudomonadales; Pseudomonadaceae | MULTISPECIES: PAS domain S-box protein | 6 |
|  | **WP_039146588.1** | 2.3E-05 | Qc.III | IQEIGGTIERLSEVSAAIAAAVEEQGAATQEISRNVQQASVGTQ--------- | *Bradyrhizobium japonicum* | Bacteria; Proteobacteria; Alphaproteobacteria; Rhizobiales; Bradyrhizobiaceae; Bradyrhizobium | methyl-accepting chemotaxis protein | 6 |
|  | **WP_044273255.1** | 2.3E-05 | Qb.I | LQRTKLHLQDTIEQAEVSSEELKASNEEMQAVNEELRSATEELETS------- | *Pseudomonas fluorescens* | Bacteria; Proteobacteria; Gammaproteobacteria; Pseudomonadales; Pseudomonadaceae; Pseudomonas | PAS domain S-box protein | 6 |
|  | **WP_047710562.1** | 2.3E-05 | Qb.I | LQRTKLHLQDTIEQAEVSSEELKASNEEMQAVNEELRSATEELETS------- | *Pseudomonas fluorescens* | Bacteria; Proteobacteria; Gammaproteobacteria; Pseudomonadales; Pseudomonadaceae; Pseudomonas | PAS domain S-box protein | 6 |
| | **WP_051356551.1** | 2.3E-05 | Qc.III | -EGVSVQIRQMSDVATGISAAVEQQGAATKEIVRNMDQAAAGTHTVTSHIAD- | *Azorhizobium doebereinerae* | Bacteria; Proteobacteria; Alphaproteobacteria; Rhizobiales; Xanthobacteraceae; Azorhizobium | methyl-accepting chemotaxis protein | 6 |
|  | **WP_054895716.1** | 2.3E-05 | Qb.I | LQRTKLHLQDTIEQAEVSSEELKASNEEMQAVNEELRSATEELETS------- | *Pseudomonas* | Bacteria; Proteobacteria; Gammaproteobacteria; Pseudomonadales; Pseudomonadaceae | MULTISPECIES: PAS domain S-box protein | 6 |
|  | **WP_054921170.1** | 2.3E-05 | Qb.I | LQRTKLHLQDTIEQAEVSSEELKASNEEMQAVNEELRSATEELETS------- | *Pseudomonas sp. NBRC 111138* | Bacteria; Proteobacteria; Gammaproteobacteria; Pseudomonadales; Pseudomonadaceae; Pseudomonas | PAS domain S-box protein | 6 |
|  | **WP_056787536.1** | 2.3E-05 | Qb.I | LQRTKLHLQDTIEQAEVSSEELKASNEEMQAVNEELRSATEELETS------- | *Pseudomonas* | Bacteria; Proteobacteria; Gammaproteobacteria; Pseudomonadales; Pseudomonadaceae | MULTISPECIES: PAS domain S-box protein | 6 |
|  | **WP_057721925.1** | 2.3E-05 | Qb.I | LQRTKLHLQDTIEQAEVSSEELKASNEEMQAVNEELRSATEELETS------- | *Pseudomonas orientalis* | Bacteria; Proteobacteria; Gammaproteobacteria; Pseudomonadales; Pseudomonadaceae; Pseudomonas | PAS domain S-box protein | 6 |
|  | **WP_057978085.1** | 2.3E-05 | Qb.I | LQRTKLHLQDTIEQAEVSSEELKASNEEMQAVNEELRSATEELETS------- | *Pseudomonas* | Bacteria; Proteobacteria; Gammaproteobacteria; Pseudomonadales; Pseudomonadaceae | MULTISPECIES: PAS domain S-box protein | 6 |
|  | **WP_058410186.1** | 2.3E-05 | Qb.I | LQRTKLHLQDTIEQAEVSSEELKASNEEMQAVNEELRSATEELETS------- | *Pseudomonas sp. ICMP 19500* | Bacteria; Proteobacteria; Gammaproteobacteria; Pseudomonadales; Pseudomonadaceae; Pseudomonas | PAS domain S-box protein | 6 |
|  | **WP_060767482.1** | 2.3E-05 | Qb.I | LQRTKLHLQDTIEQAEVSSEELKASNEEMQAVNEELRSATEELETS------- | *Pseudomonas fluorescens* | Bacteria; Proteobacteria; Gammaproteobacteria; Pseudomonadales; Pseudomonadaceae; Pseudomonas | PAS domain S-box protein | 6 |
|  | **WP_060772977.1** | 2.3E-05 | Qb.I | LQRTKLHLQDTIEQAEVSSEELKASNEEMQAVNEELRSATEELETS------- | *Pseudomonas fluorescens* | Bacteria; Proteobacteria; Gammaproteobacteria; Pseudomonadales; Pseudomonadaceae; Pseudomonas | PAS domain S-box protein | 6 |
| | **WP_061878469.1** | 2.3E-05 | Qc.III | IQEIGGTIERLSEVSSAIAAAVEEQGAATQEISRNVQQASLGTQEVSSNIT-- | *Bradyrhizobium liaoningense* | Bacteria; Proteobacteria; Alphaproteobacteria; Rhizobiales; Bradyrhizobiaceae; Bradyrhizobium | HAMP domain-containing protein | 6 |
|  | **WP_068513197.1** | 2.3E-05 | Qa.III | -DEISEIINKINDVATTIASAVEEQSIAIRSIAEQIEQTRDT----------- | *Caldimicrobium thiodismutans* | Bacteria; Thermodesulfobacteria; Thermodesulfobacteriales; Thermodesulfobacteriaceae; Caldimicrobium | methyl-accepting chemotaxis protein | 6 |
|  | **WP_069558961.1** | 2.3E-05 | Qb.I | LQRTKLHLQDTIEQAEVSSEELKASNEEMQAVNEELRSATEELETS------- | *Pseudomonas sp. AP42* | Bacteria; Proteobacteria; Gammaproteobacteria; Pseudomonadales; Pseudomonadaceae; Pseudomonas | PAS domain S-box protein | 6 |
|  | **WP_072435636.1** | 2.3E-05 | Qb.I | LQRTKLHLQDTIEQAEVSSEELKASNEEMQAVNEELRSATEELETS------- | *Pseudomonas sp. NFR02* | Bacteria; Proteobacteria; Gammaproteobacteria; Pseudomonadales; Pseudomonadaceae; Pseudomonas | PAS domain S-box protein | 6 |
|  | **WP_072448498.1** | 2.3E-05 | Qb.I | LQRTKLHLQDTIEQAEVSSEELKASNEEMQAVNEELRSATEELETS------- | *Pseudomonas sp. NFPP02* | Bacteria; Proteobacteria; Gammaproteobacteria; Pseudomonadales; Pseudomonadaceae; Pseudomonas | PAS domain S-box protein | 6 |
|  | **WP_073453996.1** | 2.3E-05 | Qb.I | LQRTKLHLQDTIEQAEVSSEELKASNEEMQAVNEELRSATEELETS------- | *Pseudomonas sp. FSL W5-0203* | Bacteria; Proteobacteria; Gammaproteobacteria; Pseudomonadales; Pseudomonadaceae; Pseudomonas | PAS domain S-box protein | 6 |
|  | **WP_073525630.1** | 2.3E-05 | Qb.I | LQRTKLHLQDTIEQAEVSSEELKASNEEMQAVNEELRSATEELETS------- | *Pseudomonas fluorescens* | Bacteria; Proteobacteria; Gammaproteobacteria; Pseudomonadales; Pseudomonadaceae; Pseudomonas | PAS domain S-box protein | 6 |
|  | **WP_073842819.1** | 2.3E-05 | Qb.I | LQRTKLHLQDTIEQAEVSSEELKASNEEMQAVNEELRSATEELETS------- | *Pseudomonas sp. BTN1* | Bacteria; Proteobacteria; Gammaproteobacteria; Pseudomonadales; Pseudomonadaceae; Pseudomonas | PAS domain S-box protein | 6 |
|  | **WP_073954410.1** | 2.3E-05 | Qa.IV | -ETITRTISELSTLTASISAAIEQQGSATAEISQSANSAAN------------ | *Thalassospira sp. TSL5-1* | Bacteria; Proteobacteria; Alphaproteobacteria; Rhodospirillales; Rhodospirillaceae; Thalassospira | HAMP domain-containing protein | 6 |
|  | **WP_078467737.1** | 2.3E-05 | Qb.I | LQRTKLHLQDTIEQAEVSSEELKASNEEMQAVNEELRSATEELETS------- | *Pseudomonas sp. MF6394* | Bacteria; Proteobacteria; Gammaproteobacteria; Pseudomonadales; Pseudomonadaceae; Pseudomonas | PAS domain S-box protein | 6 |
|  | **WP_078802886.1** | 2.3E-05 | Qb.I | LQRTKLHLQDTIEQAEVSSEELKASNEEMQAVNEELRSATEELETS------- | *Pseudomonas fluorescens* | Bacteria; Proteobacteria; Gammaproteobacteria; Pseudomonadales; Pseudomonadaceae; Pseudomonas | PAS domain S-box protein | 6 |
|  | **WP_082637273.1** | 2.3E-05 | Qc.III | IKEIGGTIGRISDIAATIASAVEQQSAATQEIARSVQNVAQG----------- | *Bradyrhizobium retamae* | Bacteria; Proteobacteria; Alphaproteobacteria; Rhizobiales; Bradyrhizobiaceae; Bradyrhizobium | methyl-accepting chemotaxis protein | 6 |
| | **WP_082811775.1** | 2.3E-05 | Qc.III | ---VIETIGRLNQISAAIASAMEEQDAATREIARNVEEASNGTGEVTRNVR-- | *Haematospirillum jordaniae* | Bacteria; Proteobacteria; Alphaproteobacteria; Rhodospirillales; Rhodospirillaceae; Haematospirillum | methyl-accepting chemotaxis protein | 6 |
|  | **WP_087611285.1** | 2.3E-05 | R.IV | --EIARNVNNIKDATDQNVTEVNKNVQSATDLSGQAYTLAQAINRF------- | *Marinomonas sp. QM202* | Bacteria; Proteobacteria; Gammaproteobacteria; Oceanospirillales; Oceanospirillaceae; Marinomonas | methyl-accepting chemotaxis protein | 6 |
|  | **WP_089042636.1** | 2.3E-05 | Qb.I | LQRTKLHLQDTIEQAEVSSEELKASNEEMQAVNEELRSATEELETS------- | *Pseudomonas fluorescens* | Bacteria; Proteobacteria; Gammaproteobacteria; Pseudomonadales; Pseudomonadaceae; Pseudomonas | PAS domain S-box protein | 6 |
|  | **WP_091892592.1** | 2.3E-05 | Qc.III | IQEIGGTIERLSEVSAAIAAAVEEQGAATQEISRNVQQASVGTQ--------- | *Bradyrhizobium sp. Rc2d* | Bacteria; Proteobacteria; Alphaproteobacteria; Rhizobiales; Bradyrhizobiaceae; Bradyrhizobium | methyl-accepting chemotaxis protein | 6 |
|  | **WP_098952002.1** | 2.3E-05 | Qb.I | LQRTKLHLQDTIEQAEVSSEELKASNEEMQAVNEELRSATEELETS------- | *Pseudomonas sp. FDAARGOS_380* | Bacteria; Proteobacteria; Gammaproteobacteria; Pseudomonadales; Pseudomonadaceae; Pseudomonas | PAS domain S-box protein | 6 |
| | **WP_100236282.1** | 2.3E-05 | Qc.III | IQEIGGTIERLSEVSSAIAAAVEEQGAATQEISRNVQQASLGTQEVSSNIT-- | *Bradyrhizobium sp. INPA54B* | Bacteria; Proteobacteria; Alphaproteobacteria; Rhizobiales; Bradyrhizobiaceae; Bradyrhizobium | HAMP domain-containing protein | 6 |
|  | **WP_102666062.1** | 2.3E-05 | Qb.I | LQRTKLHLQDTIEQAEVSSEELKASNEEMQAVNEELRSATEELETS------- | *Pseudomonas* | Bacteria; Proteobacteria; Gammaproteobacteria; Pseudomonadales; Pseudomonadaceae | MULTISPECIES: PAS domain S-box protein | 6 |
|  | **WP_106111626.1** | 2.3E-05 | Qb.I | LQRTKLHLQDTIEQAEVSSEELKASNEEMQAVNEELRSATEELETS------- | *Pseudomonas fluorescens* | Bacteria; Proteobacteria; Gammaproteobacteria; Pseudomonadales; Pseudomonadaceae; Pseudomonas | PAS domain S-box protein | 6 |
|  | **WP_106118382.1** | 2.3E-05 | Qb.I | LQRTKLHLQDTIEQAEVSSEELKASNEEMQAVNEELRSATEELETS------- | *Pseudomonas fluorescens* | Bacteria; Proteobacteria; Gammaproteobacteria; Pseudomonadales; Pseudomonadaceae; Pseudomonas | PAS domain S-box protein | 6 |
|  | **WP_106754291.1** | 2.3E-05 | Qc.III | -RSIAGTIDRISDISNTIRASIQQQAEATDAIGRSVSVATS------------ | *Pannonibacter carbonis* | Bacteria; Proteobacteria; Alphaproteobacteria; Rhodobacterales; Rhodobacteraceae; Pannonibacter | methyl-accepting chemotaxis protein | 6 |
|  | **WP_108538450.1** | 2.3E-05 | Qb.I | LQRTKLHLQDTIEQAEVSSEELKASNEEMQAVNEELRSATEELETS------- | *Pseudomonas sp. HMWF034* | Bacteria; Proteobacteria; Gammaproteobacteria; Pseudomonadales; Pseudomonadaceae; Pseudomonas | PAS domain S-box protein | 6 |
|  | **WP_109075853.1** | 2.3E-05 | Qc.III | --GIADTVLQVNQIAATITSAVEEQTAATGEISRNVQQAAQGTSEVSR----- | *Azospirillum* | Bacteria; Proteobacteria; Alphaproteobacteria; Rhodospirillales; Rhodospirillaceae | MULTISPECIES: HAMP domain-containing protein | 6 |
|  | **WP_109095736.1** | 2.3E-05 | Qa.III | IKGIGGVIGEISEIASAIAAAVEEQGTATQEISRNVQQAAIGTQEV------- | *Azospirillum sp. TSH64* | Bacteria; Proteobacteria; Alphaproteobacteria; Rhodospirillales; Rhodospirillaceae; Azospirillum | HAMP domain-containing protein | 6 |
|  | **WP_109902387.1** | 2.3E-05 | Qc.III | -GRIDEMIRAMAEASTGIAAAVEEQIASIDEIARNIDRTSDD----------- | *Zavarzinia sp. HR-AS* | Bacteria; Proteobacteria; Alphaproteobacteria; Rhodospirillales; Acetobacteraceae; Zavarzinia | methyl-accepting chemotaxis protein | 6 |
| | **WP_110784509.1** | 2.3E-05 | Qc.III | IKEIGDIIGRMSEIASTIASAVEEQGAATQEIARNVQEAARGTQQVSSNI--- | *Rhodopseudomonas palustris* | Bacteria; Proteobacteria; Alphaproteobacteria; Rhizobiales; Bradyrhizobiaceae; Rhodopseudomonas | methyl-accepting chemotaxis protein | 6 |
| | **GBF58040.1** | 2.4E-05 | Qc.III.c | ---ITQVIGSINEVSAAIAAAVEEQSALTDAMSSNMAVASQSVQAISSRM--- | *Candidatus Phycosocius bacilliformis* | Bacteria; Proteobacteria; Alphaproteobacteria; Candidatus Phycosocius | biofilm dispersion protein BdlA | 6 |
|  | **OJJ11821.1** | 2.4E-05 | Qa.IV | -QSIAEVVKQLDEMTSSIAAAVEEQGAATQEISHSVQTVAN------------ | *Alphaproteobacteria bacterium AO1-B* | Bacteria; Proteobacteria; Alphaproteobacteria | chemotaxis protein | 6 |
|  | **PWC82881.1** | 2.4E-05 | Qa.III | IQSISNIIEQIQGISTTISSAVEEQGSATREISRNVQQAAQGTQQ-------- | *Azospirillum sp. TSH100* | Bacteria; Proteobacteria; Alphaproteobacteria; Rhodospirillales; Rhodospirillaceae; Azospirillum | hypothetical protein TSH100_22550 | 6 |
| | **SCB55651.1** | 2.4E-05 | Qc.III | IQEIGGTIERLSEVSSAIAAAVEEQGAATQEISRNVQQASLGTQEVSSNIT-- | *Bradyrhizobium shewense* | Bacteria; Proteobacteria; Alphaproteobacteria; Rhizobiales; Bradyrhizobiaceae; Bradyrhizobium | methyl-accepting chemotaxis protein | 6 |
|  | **WP_007042691.1** | 2.4E-05 | Qb.I | LKSTREDLRSTIEQMETSNEELKASNEEVMSMNEELQSTNEELETSK------ | *Thiorhodococcus drewsii* | Bacteria; Proteobacteria; Gammaproteobacteria; Chromatiales; Chromatiaceae; Thiorhodococcus | PAS domain S-box protein | 6 |
| | **WP_007607191.1** | 2.4E-05 | Qc.III | --AISNTIEKLSEISSTIAAAVEEQGAATQEISRNVQQAAEGTHQVSSNITD- | *Bradyrhizobium* | Bacteria; Proteobacteria; Alphaproteobacteria; Rhizobiales; Bradyrhizobiaceae | MULTISPECIES: methyl-accepting chemotaxis protein | 6 |
|  | **WP_008192796.1** | 2.4E-05 | Qa.IV | -QSIAEVVKQLDEMTSSIAAAVEEQGAATQEISHSVQTVAN------------ | *Labrenzia alexandrii* | Bacteria; Proteobacteria; Alphaproteobacteria; Rhodobacterales; Rhodobacteraceae; Labrenzia | methyl-accepting chemotaxis protein | 6 |
|  | **WP_008961155.1** | 2.4E-05 | Qc.III | -KAIGSTIERINEIAASISAAVEQQRAATQNITQSVRSAAN------------ | *Bradyrhizobium sp. STM 3809* | Bacteria; Proteobacteria; Alphaproteobacteria; Rhizobiales; Bradyrhizobiaceae; Bradyrhizobium | HAMP domain-containing protein | 6 |
|  | **WP_015408511.1** | 2.4E-05 | Qc.III.c | ---LEAVVEEVVDSAEEIAAHVDEQNGMLEDAVSEMQSFSARMEEV------- | *Natronomonas moolapensis* | Archaea; Euryarchaeota; Halobacteria; Halobacteriales; Haloarculaceae; Natronomonas | PAS domain-containing protein | 6 |
|  | **WP_019645342.1** | 2.4E-05 | Qc.III | LGEISGTIVQMNEISTTIASAIEEQGAATQEIARNAEQAATGTEQVTAS---- | *Novispirillum itersonii* | Bacteria; Proteobacteria; Alphaproteobacteria; Rhodospirillales; Rhodospirillaceae; Novispirillum | methyl-accepting chemotaxis protein | 6 |

| | | | | | | | | |
|---|---|---|---|---|---|---|---|---|
|  | **WP_022850495.1** | 2.4E-05 | SNAP.b | TREMMQRIASTVTEAEQTVRELGKSSEQIGEIIKVIDEIADQT---------- | *Geovibrio sp. L21-Ace-BES* | Bacteria; Deferribacteres; Deferribacterales; Deferribacteraceae; Geovibrio | methyl-accepting chemotaxis protein | 6 |
|  | **WP_024517586.1** | 2.4E-05 | Qc.III | IKRIGSIIGEVNEVATAIAAAVQEQGAATQEITRSTQFAAQGTKNV------- | *Bradyrhizobium sp. Tv2a-2* | Bacteria; Proteobacteria; Alphaproteobacteria; Rhizobiales; Bradyrhizobiaceae; Bradyrhizobium | methyl-accepting chemotaxis protein | 6 |
| | **WP_027516461.1** | 2.4E-05 | Qc.III | --AISNTIEKLSEISSTIAAAVEEQGAATQEISRNVQQAAEGTHQVSSNITD- | *Bradyrhizobium sp. WSM1417* | Bacteria; Proteobacteria; Alphaproteobacteria; Rhizobiales; Bradyrhizobiaceae; Bradyrhizobium | methyl-accepting chemotaxis protein | 6 |
| | **WP_041041610.1** | 2.4E-05 | Qc.III | IQGIGGIIERINQVSAAIAAAVEEQEATTRDIARNVAEAATGTREVSRHVTD- | *Magnetospirillum magnetotacticum* | Bacteria; Proteobacteria; Alphaproteobacteria; Rhodospirillales; Rhodospirillaceae; Magnetospirillum | HAMP domain-containing protein | 6 |
|  | **WP_055670142.1** | 2.4E-05 | Qa.IV | -QSIAEVVKQLDEMTSSIAAAVEEQGAATQEISHSVQTVAN------------ | *Labrenzia alexandrii* | Bacteria; Proteobacteria; Alphaproteobacteria; Rhodobacterales; Rhodobacteraceae; Labrenzia | chemotaxis protein | 6 |
| | **WP_056295865.1** | 2.4E-05 | Qc.III | IKEIGSTIGQISTIAGTITNAIEQQSAATQEIAHSIQNVAAGTQQVAGNI--- | *Afipia sp. Root123D2* | Bacteria; Proteobacteria; Alphaproteobacteria; Rhizobiales; Bradyrhizobiaceae; Afipia | methyl-accepting chemotaxis protein | 6 |
| | **WP_063706282.1** | 2.4E-05 | Qc.III | IQEIGGTIERLSEVSSAIAAAVEEQGAATQEISRNVQQASLGTQEVSSNIT-- | *Bradyrhizobium sp. BR 10245* | Bacteria; Proteobacteria; Alphaproteobacteria; Rhizobiales; Bradyrhizobiaceae; Bradyrhizobium | methyl-accepting chemotaxis protein | 6 |
|  | **WP_072242580.1** | 2.4E-05 | Qa.III | INAISETVREVDNMTATIAAAVEQQGAATREISSNIQSAA------------- | *Pannonibacter indicus* | Bacteria; Proteobacteria; Alphaproteobacteria; Rhodobacterales; Rhodobacteraceae; Pannonibacter | chemotaxis protein | 6 |
|  | **WP_073953451.1** | 2.4E-05 | Qc.III | -RDISEVISRTRETASSISSAVEEQNTATQEIAQNVAQASAGTQEV------- | *Thalassospira sp. TSL5-1* | Bacteria; Proteobacteria; Alphaproteobacteria; Rhodospirillales; Rhodospirillaceae; Thalassospira | methyl-accepting chemotaxis protein | 6 |
|  | **WP_077524479.1** | 2.4E-05 | Qa.III | LEQIARSLRNLTGLNASIASATLEQSQVVEDINQNVTQAASLAEE-------- | *Pseudomonas resinovorans* | Bacteria; Proteobacteria; Gammaproteobacteria; Pseudomonadales; Pseudomonadaceae; Pseudomonas | methyl-accepting chemotaxis protein | 6 |
| | **WP_081478484.1** | 2.4E-05 | Qb.II | --DTSSAITEIIANIESVHKQIRLQNESVESSSENVTGISDSISVLDHLIE-- | *Treponema saccharophilum* | Bacteria; Spirochaetes; Spirochaetales; Spirochaetaceae; Treponema | methyl-accepting chemotaxis protein | 6 |
|  | **WP_081928851.1** | 2.4E-05 | Qa.IV | IQEIGRTIAEMSQISVSIAAAMEQQGAATTEIARNVQQAA------------- | *Microvirga sp. BSC39* | Bacteria; Proteobacteria; Alphaproteobacteria; Rhizobiales; Methylobacteriaceae; Microvirga | methyl-accepting chemotaxis protein | 6 |
| | **WP_085964302.1** | 2.4E-05 | Qc.III | IQEIGGTIERLSEVSSAIAAAVEEQGAATQEISRNVQQASLGTQEVSSNIT-- | *Bradyrhizobium sp. WSM2793* | Bacteria; Proteobacteria; Alphaproteobacteria; Rhizobiales; Bradyrhizobiaceae; Bradyrhizobium | methyl-accepting chemotaxis protein | 6 |
| | **WP_091967318.1** | 2.4E-05 | Qc.III | IQEIGGTIERLSEVSSAIAAAVEEQGAATQEISRNVQQASLGTQEVSSNIT-- | *Bradyrhizobium shewense* | Bacteria; Proteobacteria; Alphaproteobacteria; Rhizobiales; Bradyrhizobiaceae; Bradyrhizobium | methyl-accepting chemotaxis protein | 6 |
| | **WP_104462862.1** | 2.4E-05 | Qc.III | IQEIGGTIERLSEVSSAIAAAVEEQGAATQEISRNVQQASLGTQEVSSNIT-- | *Bradyrhizobium sp. AC87j1* | Bacteria; Proteobacteria; Alphaproteobacteria; Rhizobiales; Bradyrhizobiaceae; Bradyrhizobium | methyl-accepting chemotaxis protein | 6 |
|  | **WP_108224725.1** | 2.4E-05 | Qb.I | LQRTKEHLQETIEHSETSSEELKASNEEMHAINEELRSATEELETS------- | *Pseudomonas sp. GV091* | Bacteria; Proteobacteria; Gammaproteobacteria; Pseudomonadales; Pseudomonadaceae; Pseudomonas | PAS domain S-box protein | 6 |
|  | **WP_111388716.1** | 2.4E-05 | Qc.III | IKEIGGTIGRIAEIAATIAAAVEEQGAATAEIARNVQQASQG----------- | *Rhodoplanes piscinae* | Bacteria; Proteobacteria; Alphaproteobacteria; Rhizobiales; Hyphomicrobiaceae; Rhodoplanes | methyl-accepting chemotaxis protein | 6 |
|  | **WP_114087727.1** | 2.4E-05 | Qc.III | -RDISEVISRTRETASSISSAVEEQNTATQEIAQNVAQASAGTQEV------- | *Thalassospira profundimaris* | Bacteria; Proteobacteria; Alphaproteobacteria; Rhodospirillales; Rhodospirillaceae; Thalassospira | HAMP domain-containing protein | 6 |
| | **BAI75326.1** | 2.5E-05 | Qc.III | -KGISATITQMSGIATAIASAVEEQNAATGEISRNVTEAASGTREVTVNV--- | *Azospirillum sp. B510* | Bacteria; Proteobacteria; Alphaproteobacteria; Rhodospirillales; Rhodospirillaceae; Azospirillum | methyl-accepting chemotaxis protein (plasmid) | 6 |
|  | **EGU39713.1** | 2.5E-05 | Qa.III | LEEINTSVALISDMATQIATAAEEQTHVTGEITQNITTIK------------- | *Vibrio splendidus ATCC 33789* | Bacteria; Proteobacteria; Gammaproteobacteria; Vibrionales; Vibrionaceae; Vibrio | methyl-accepting chemotaxis protein | 6 |
|  | **EKD36142.1** | 2.5E-05 | Qa.III | IDKISTIIADINNVINGIATAVVEQSAATSEIAGNISQAS------------- | *uncultured bacterium* | Bacteria; environmental samples | hypothetical protein ACD_75C01617G0001 | 6 |
|  | **KPP98581.1** | 2.5E-05 | Qa.III | IRGITDVIGEVGELVNGIATAIEEQSVVTRDVATNIAQATAGVQDANQ----- | *Bacteroidetes bacterium HLUCCA01* | Bacteria; Bacteroidetes | chemotaxis signal relay system methyl-accepting signal transducer | 6 |
|  | **KRR30460.1** | 2.5E-05 | Qc.III | IKEIGGTIGRISDIAATIASAVEQQSAATQEIARSVQNVAQG----------- | *Bradyrhizobium retamae* | Bacteria; Proteobacteria; Alphaproteobacteria; Rhizobiales; Bradyrhizobiaceae; Bradyrhizobium | chemotaxis protein | 6 |
|  | **OOF10036.1** | 2.5E-05 | Qc.III | -DEISSNLSNIKTLSSDMHEAIDQLGPIVSDLKDNVNALNG------------ | *Salinivibrio sp. MA427* | Bacteria; Proteobacteria; Gammaproteobacteria; Vibrionales; Vibrionaceae; Salinivibrio | methyl-accepting chemotaxis protein partial | 6 |
|  | **OYU88695.1** | 2.5E-05 | Qa.III | IRAITGTIGTINEIASSIASAVEEQSAATQEIARNVQ----HAAQGANE---- | *Bradyrhizobiaceae bacterium PARB1* | Bacteria; Proteobacteria; Alphaproteobacteria; Rhizobiales; Bradyrhizobiaceae | chemotaxis protein | 6 |
|  | **PMP70845.1** | 2.5E-05 | SNAP.b | TKEMMEKIAETVSHSANTVRALGESSEQIGQIIQVIDDIADQT---------- | *Calditerrivibrio nitroreducens* | Bacteria; Deferribacteres; Deferribacterales; Deferribacteraceae | chemotaxis protein | 6 |
|  | **PWC79828.1** | 2.5E-05 | Qa.III | IRSIAGTIREVNEIAATIAAAVEQQSAATREIARNVGEAAD------------ | *Azospirillum sp. TSH64* | Bacteria; Proteobacteria; Alphaproteobacteria; Rhodospirillales; Rhodospirillaceae; Azospirillum | chemotaxis protein | 6 |
| | **RCK26908.1** | 2.5E-05 | Qc.III | IEKVTQTISSISEIANGIASAVSEQNSAVNEIANSATSASSGSNEVVANIDE- | *Thalassospira lucentensis MCCC 1A00383 = DSM 14000* | Bacteria; Proteobacteria; Alphaproteobacteria; Rhodospirillales; Rhodospirillaceae; Thalassospira | chemotaxis protein | 6 |
| | **SDF05823.1** | 2.5E-05 | Qc.III | IQEIVKVIGEINQIATVIAAAVEEQTAATQEIARNVNEASRGTTEVSSNI--- | *Thalassobaculum litoreum DSM 18839* | Bacteria; Proteobacteria; Alphaproteobacteria; Rhodospirillales; Rhodospirillaceae; Thalassobaculum | Methyl-accepting chemotaxis protein | 6 |
|  | **SLM28001.1** | 2.5E-05 | Qa.III | LKSIVEIIDEINTIVTSVATAIEEQSATTQEISNNVSQAATGVQEVNEN---- | *Desulfamplus magnetovallimortis* | Bacteria; Proteobacteria; Deltaproteobacteria; Desulfobacterales; Desulfobacteraceae; Desulfamplus | hypothetical protein MTBBW1_1210041 | 6 |
|  | **WP_004579854.1** | 2.5E-05 | Qa.III | -KRILESVESISSINEQIAAATYEQSTTFNSVTQHITDIHNNAEQVS------ | *Marinobacter nanhaiticus* | Bacteria; Proteobacteria; Gammaproteobacteria; Alteromonadales; Alteromonadaceae; Marinobacter | methyl-accepting chemotaxis protein | 6 |
| | **WP_006611935.1** | 2.5E-05 | Qc.III | -QAISTTIERLAEIGAAIAAAVEEQGAATQEISRNVQQAAHGTQQVSSNI--- | *Bradyrhizobium sp. ORS 285* | Bacteria; Proteobacteria; Alphaproteobacteria; Rhizobiales; Bradyrhizobiaceae; Bradyrhizobium | HAMP domain-containing protein | 6 |
| | **WP_007566248.1** | 2.5E-05 | Qc.III | --SIARIVGEIATSSTAIAAAIEEQGAATQEIARNVQQAASGTDRVTGNI--- | *Methylobacterium sp. GXF4* | Bacteria; Proteobacteria; Alphaproteobacteria; Rhizobiales; Methylobacteriaceae; Methylobacterium | HAMP domain-containing protein | 6 |
|  | **WP_008382574.1** | 2.5E-05 | Qc.III | --EISGTIGKIAEIASAIAAAVEQQGAATGEISRNIQRAAAGTTEVS------ | *Rhodovulum sp. PH10* | Bacteria; Proteobacteria; Alphaproteobacteria; Rhodobacterales; Rhodobacteraceae; Rhodovulum | methyl-accepting chemotaxis protein | 6 |
|  | **WP_008960436.1** | 2.5E-05 | Qc.III | IQRIGGIIGEVNEVATAIAAAVQEQGAATQEISRSTQFAADGTRNVS------ | *Bradyrhizobium sp. STM 3809* | Bacteria; Proteobacteria; Alphaproteobacteria; Rhizobiales; Bradyrhizobiaceae; Bradyrhizobium | methyl-accepting chemotaxis protein | 6 |
|  | **WP_010438865.1** | 2.5E-05 | Qa.III | LEEINAAVVLISDMATQIATAAEEQTHVTNEVTQNVTAIK------------- | *Vibrio cyclitrophicus* | Bacteria; Proteobacteria; Gammaproteobacteria; Vibrionales; Vibrionaceae; Vibrio | methyl-accepting chemotaxis protein | 6 |
|  | **WP_012982359.1** | 2.5E-05 | Qc.III | ---ITQSVTRISDMNTQIASAAEEQSTVAEEINRNVANVTEVLDQT------- | *Thioalkalivibrio sp. K90mix* | Bacteria; Proteobacteria; Gammaproteobacteria; Chromatiales; Ectothiorhodospiraceae; Thioalkalivibrio | methyl-accepting chemotaxis protein | 6 |
|  | **WP_013450563.1** | 2.5E-05 | SNAP.b | TKEMMEKIAETVSHSANTVRALGESSEQIGQIIQVIDDIADQT---------- | *Calditerrivibrio nitroreducens* | Bacteria; Deferribacteres; Deferribacterales; Deferribacteraceae | methyl-accepting chemotaxis protein | 6 |
|  | **WP_018139124.1** | 2.5E-05 | Qc.III | ---ITQSVTRISDMNTQIASAAEEQSTVAEEINRNVANVTEVLDQT------- | *Thioalkalivibrio* | Bacteria; Proteobacteria; Gammaproteobacteria; Chromatiales; Ectothiorhodospiraceae | MULTISPECIES: methyl-accepting chemotaxis protein | 6 |
|  | **WP_018174488.1** | 2.5E-05 | Qc.III | ---ITQSVTRISDMNTQIASAAEEQSTVAEEINRNVANVTEVLDQT------- | *Thioalkalivibrio* | Bacteria; Proteobacteria; Gammaproteobacteria; Chromatiales; Ectothiorhodospiraceae | MULTISPECIES: methyl-accepting chemotaxis protein | 6 |
| | **WP_018458836.1** | 2.5E-05 | Qc.III | --AISNTIEKLSEISSTIAAAVEEQGAATQEISRNVQQAAEGTHQVSSNITD- | *Bradyrhizobium sp. WSM4349* | Bacteria; Proteobacteria; Alphaproteobacteria; Rhizobiales; Bradyrhizobiaceae; Bradyrhizobium | methyl-accepting chemotaxis protein | 6 |
|  | **WP_018861659.1** | 2.5E-05 | Qc.III | ---ITQSVTRISDMNTQIASAAEEQSTVAEEINRNVANVTEVLDQT------- | *Thioalkalivibrio sp. ALJ3* | Bacteria; Proteobacteria; Gammaproteobacteria; Chromatiales; Ectothiorhodospiraceae; Thioalkalivibrio | methyl-accepting chemotaxis protein | 6 |
|  | **WP_018864097.1** | 2.5E-05 | Qc.III | ---ITQSVTRISDMNTQIASAAEEQSTVAEEINRNVANVTEVLDQT------- | *Thioalkalivibrio sp. ARh3* | Bacteria; Proteobacteria; Gammaproteobacteria; Chromatiales; Ectothiorhodospiraceae; Thioalkalivibrio | methyl-accepting chemotaxis protein | 6 |
|  | **WP_018868569.1** | 2.5E-05 | Qc.III | ---ITQSVTRISDMNTQIASAAEEQSTVAEEINRNVANVTEVLDQT------- | *Thioalkalivibrio* | Bacteria; Proteobacteria; Gammaproteobacteria; Chromatiales; Ectothiorhodospiraceae | MULTISPECIES: methyl-accepting chemotaxis protein | 6 |
|  | **WP_018940472.1** | 2.5E-05 | Qc.III | ---ITQSVTRISDMNTQIASAAEEQSTVAEEINRNVANVTEVLDQT------- | *Thioalkalivibrio sp. AKL11* | Bacteria; Proteobacteria; Gammaproteobacteria; Chromatiales; Ectothiorhodospiraceae; Thioalkalivibrio | methyl-accepting chemotaxis protein | 6 |
|  | **WP_019623449.1** | 2.5E-05 | Qc.III | ---ITQSVTRISDMNTQIASAAEEQSTVAEEINRNVANVTEVLDQT------- | *Thioalkalivibrio* | Bacteria; Proteobacteria; Gammaproteobacteria; Chromatiales; Ectothiorhodospiraceae | MULTISPECIES: methyl-accepting chemotaxis protein | 6 |
|  | **WP_020146193.1** | 2.5E-05 | Qc.III | ---ITQSVTRISDMNTQIASAAEEQSTVAEEINRNVANVTEVLDQT------- | *Thioalkalivibrio sp. ALJ15* | Bacteria; Proteobacteria; Gammaproteobacteria; Chromatiales; Ectothiorhodospiraceae; Thioalkalivibrio | methyl-accepting chemotaxis protein | 6 |
|  | **WP_026148104.1** | 2.5E-05 | Qc.III | ---ITQSVTRISDMNTQIASAAEEQSTVAEEINRNVANVTEVLDQT------- | *Thioalkalivibrio sp. HL-Eb18* | Bacteria; Proteobacteria; Gammaproteobacteria; Chromatiales; Ectothiorhodospiraceae; Thioalkalivibrio | methyl-accepting chemotaxis protein | 6 |
|  | **WP_026202956.1** | 2.5E-05 | Qa.IV | IKEIGATIARISDISSTIAATIEEQGAATAEIARNVSEAAK------------ | *Bradyrhizobium sp. WSM2793* | Bacteria; Proteobacteria; Alphaproteobacteria; Rhizobiales; Bradyrhizobiaceae; Bradyrhizobium | chemotaxis protein | 6 |
| | **WP_027556453.1** | 2.5E-05 | Qc.III | IKDISGTIERLSEISSTIAAAVEEQGAATQEISRNVQQAAQGTMQVSSNI--- | *Bradyrhizobium sp. Cp5.3* | Bacteria; Proteobacteria; Alphaproteobacteria; Rhizobiales; Bradyrhizobiaceae; Bradyrhizobium | HAMP domain-containing protein | 6 |
| | **WP_035961728.1** | 2.5E-05 | Qc.III | --AISNTIEKLSEISSTIAAAVEEQGAATQEISRNVQQAAEGTHQVSSNITD- | *Bradyrhizobium sp. URHA0013* | Bacteria; Proteobacteria; Alphaproteobacteria; Rhizobiales; Bradyrhizobiaceae; Bradyrhizobium | methyl-accepting chemotaxis protein | 6 |
|  | **WP_042352603.1** | 2.5E-05 | Qa.III.b | -RRIQSSIQNLNDAFSGINQLSDNQAAAIEEISSTINEISRNAQ--------- | *Bacillus massiliogorillae* | Bacteria; Firmicutes; Bacilli; Bacillales; Bacillaceae; Bacillus | chemotaxis protein | 6 |
|  | **WP_063312872.1** | 2.5E-05 | Qc.III.c | ---IGETIEQINSLSTGINEAVGDQKQATEDISRDLNSAHLNVSQ-------- | *Pseudovibrio sp. Ad26* | Bacteria; Proteobacteria; Alphaproteobacteria; Rhodobacterales; Rhodobacteraceae; Pseudovibrio | methyl-accepting chemotaxis protein | 6 |
|  | **WP_084408135.1** | 2.5E-05 | Qc.III.c | ---IGETIEQINSLSTGINEAVGDQKQATEDISRDLNSAHLNVSQ-------- | *Pseudovibrio sp. Ad37* | Bacteria; Proteobacteria; Alphaproteobacteria; Rhodobacterales; Rhodobacteraceae; Pseudovibrio | methyl-accepting chemotaxis protein | 6 |
| | **WP_085352602.1** | 2.5E-05 | Qc.III | --AISNTIEKLSEISSTIAAAVEEQGAATQEISRNVQQAAEGTHQVSSNITD- | *Bradyrhizobium canariense* | Bacteria; Proteobacteria; Alphaproteobacteria; Rhizobiales; Bradyrhizobiaceae; Bradyrhizobium | methyl-accepting chemotaxis protein | 6 |
| | **WP_085354599.1** | 2.5E-05 | Qc.III | --AISNTIEKLSEISSTIAAAVEEQGAATQEISRNVQQAAEGTHQVSSNITD- | *Bradyrhizobium canariense* | Bacteria; Proteobacteria; Alphaproteobacteria; Rhizobiales; Bradyrhizobiaceae; Bradyrhizobium | methyl-accepting chemotaxis protein | 6 |
| | **WP_085361343.1** | 2.5E-05 | Qc.III | --AISNTIEKLSEISSTIAAAVEEQGAATQEISRNVQQAAEGTHQVSSNITD- | *Bradyrhizobium canariense* | Bacteria; Proteobacteria; Alphaproteobacteria; Rhizobiales; Bradyrhizobiaceae; Bradyrhizobium | methyl-accepting chemotaxis protein | 6 |
| | **WP_091863847.1** | 2.5E-05 | Qc.III | --SIARIVGEIATSSTAIAAAIEEQGAATQEIARNVQQAASGTDRVTGNI--- | *Methylobacterium brachiatum* | Bacteria; Proteobacteria; Alphaproteobacteria; Rhizobiales; Methylobacteriaceae; Methylobacterium | HAMP domain-containing protein | 6 |
| | **WP_092227261.1** | 2.5E-05 | Qc.III | -NGIAGTIGKMSEIAIAISSAVEEQGAATQEIARNIQSAAQGSSEITSHI--- | *Bradyrhizobium sp. Gha* | Bacteria; Proteobacteria; Alphaproteobacteria; Rhizobiales; Bradyrhizobiaceae; Bradyrhizobium | HAMP domain-containing protein | 6 |
|  | **WP_092251279.1** | 2.5E-05 | Qa.IV | IKEIGATIARISDISSTIAATIEEQGAATAEIARNVSEAAK------------ | *Bradyrhizobium sp. Rc3b* | Bacteria; Proteobacteria; Alphaproteobacteria; Rhizobiales; Bradyrhizobiaceae; Bradyrhizobium | chemotaxis protein | 6 |

| | | | | | | | |
|---|---|---|---|---|---|---|---|
| **WP_092293446.1** | 2.5E-05 | Qc.III | -NGIAGTIGKMSEIAIAISSAVEEQGAATQEIARNIQSAAQGSSEITSHI--- | *Bradyrhizobium sp. Ghvi* | Bacteria; Proteobacteria; Alphaproteobacteria; Rhizobiales; Bradyrhizobiaceae; Bradyrhizobium | HAMP domain-containing protein | 6 |
| **WP_093147349.1** | 2.5E-05 | Qc.III | IQEIVKVIGEINQIATVIAAAVEEQTAATQEIARNVNEASRGTTEVSSNI--- | *Thalassobaculum litoreum* | Bacteria; Proteobacteria; Alphaproteobacteria; Rhodospirillales; Rhodospirillaceae; Thalassobaculum | methyl-accepting chemotaxis protein | 6 |
| **WP_093520006.1** | 2.5E-05 | Qc.III.c | ---IGETIEQINSLSTGINEAVGDQKQATEDISRDLNSAHLNVSQ-------- | *Pseudovibrio* | Bacteria; Proteobacteria; Alphaproteobacteria; Rhodobacterales; Rhodobacteraceae | MULTISPECIES: methyl-accepting chemotaxis protein | 6 |
| **WP_098735520.1** | 2.5E-05 | Qa.III | IRSIAGTIREVNEIAATIAAAVEQQSAATREIARNVGEAAD------------ | *Azospirillum sp. B2* | Bacteria; Proteobacteria; Alphaproteobacteria; Rhodospirillales; Rhodospirillaceae; Azospirillum | HAMP domain-containing protein | 6 |
| **WP_101301137.1** | 2.5E-05 | Qc.III | IEKVTQTISSISEIANGIASAVSEQNSAVNEIANSATSASSGSNEVVANIDE- | *Thalassospira lohafexi* | Bacteria; Proteobacteria; Alphaproteobacteria; Rhodospirillales; Rhodospirillaceae; Thalassospira | methyl-accepting chemotaxis protein | 6 |
| **WP_106945090.1** | 2.5E-05 | Qc.III | --AISNTIEKLSEISSTIAAAVEEQGAATQEISRNVQQAAEGTHQVSSNITD- | *Bradyrhizobium sp. MOS002* | Bacteria; Proteobacteria; Alphaproteobacteria; Rhizobiales; Bradyrhizobiaceae; Bradyrhizobium | methyl-accepting chemotaxis protein | 6 |
| **WP_109050660.1** | 2.5E-05 | Qa.III | IRSIAGTIREVNEIAATIAAAVEQQSAATREIARNVGEAAD------------ | *Azospirillum sp. TSA6c* | Bacteria; Proteobacteria; Alphaproteobacteria; Rhodospirillales; Rhodospirillaceae; Azospirillum | HAMP domain-containing protein | 6 |
| **WP_109073598.1** | 2.5E-05 | Qa.III | IRSIAGTIREVNEIAATIAAAVEQQSAATREIARNVGEAAD------------ | *Azospirillum* | Bacteria; Proteobacteria; Alphaproteobacteria; Rhodospirillales; Rhodospirillaceae | MULTISPECIES: HAMP domain-containing protein | 6 |
| **WP_109449006.1** | 2.5E-05 | Qa.III | IRSIAGTIREVNEIAATIAAAVEQQSAATREIARNVGEAAD------------ | *Azospirillum sp. TSH64* | Bacteria; Proteobacteria; Alphaproteobacteria; Rhodospirillales; Rhodospirillaceae; Azospirillum | HAMP domain-containing protein | 6 |
| **AWL98015.1** | 2.6E-05 | Qa.IV | IKEIGATIARISDISSTIAATIEEQGAATAEIARNVSEAAK------------ | *Bradyrhizobium ottawaense* | Bacteria; Proteobacteria; Alphaproteobacteria; Rhizobiales; Bradyrhizobiaceae; Bradyrhizobium | methyl-accepting chemotaxis protein | 6 |
| **KIN11198.1** | 2.6E-05 | Qa.III | LDEINSAVALISDMATQIATAAEEQTHVTNEITQNVTSIK------------- | *Vibrio mytili* | Bacteria; Proteobacteria; Gammaproteobacteria; Vibrionales; Vibrionaceae; Vibrio | chemotaxis protein | 6 |
| **KMQ51874.1** | 2.6E-05 | Qa.I | --KINSVISEVGEIIYTIATAIEEQATVTRDIAENISGASDDVREANE----- | *Chitinispirillum alkaliphilum* | Bacteria; Fibrobacteres; Chitinispirillia; Chitinispirillales; Chitinispirillaceae; Chitinispirillum | Methyl-accepting chemotaxis protein | 6 |
| **KPW64140.1** | 2.6E-05 | Qa.III | ---ISQALRNLNSLNTSIASATLQQTHVVDDINQNVTQAAQLSQ--------- | *Pseudomonas amygdali pv. ciccaronei* | Bacteria; Proteobacteria; Gammaproteobacteria; Pseudomonadales; Pseudomonadaceae; Pseudomonas; Pseudomonas amygdali | Methyl-accepting chemotaxis protein | 6 |
| **PHS21443.1** | 2.6E-05 | Qc.III | IEGISKTIGDINEFSAAISAAMEEQTAVTGEMSSNMQTAASGVNNIT------ | *Robiginitomaculum sp.* | Bacteria; Proteobacteria; Alphaproteobacteria; Rhodobacterales; Hyphomonadaceae; Robiginitomaculum | chemotaxis protein | 6 |
| **PZR13770.1** | 2.6E-05 | Qa.IV | IQNIAATIEEVSAITVTIASAVEEQSAATGEISRTVNQTALATD--------- | *Azospirillum brasilense* | Bacteria; Proteobacteria; Alphaproteobacteria; Rhodospirillales; Rhodospirillaceae; Azospirillum | methyl-accepting chemotaxis protein | 6 |
| **SFE31099.1** | 2.6E-05 | Qa.IV | -------IREISGVATSIAAAVEEQGAATQEIVRNVAQASAETNEVTGNIA-- | *Methylobacterium sp. 13MFTsu3.1M2* | Bacteria; Proteobacteria; Alphaproteobacteria; Rhizobiales; Methylobacteriaceae; Methylobacterium | Methyl-accepting chemotaxis protein (MCP) signalling domain-containing protein | 6 |
| **STA92916.1** | 2.6E-05 | Qb.I | ----------AVEKMEQVNKISAAQSESVTRTEEKFEGIANSLEVTKNVVKKL | *Clostridium cochlearium* | Bacteria; Firmicutes; Clostridia; Clostridiales; Clostridiaceae; Clostridium | methyl-accepting chemotaxis protein | 6 |
| **WP_011665318.1** | 2.6E-05 | Qa.IV | -KEISDTISKMSEIASTIASAVEEQGAATQEISRNVQQAAQ------------ | *Rhodopseudomonas palustris* | Bacteria; Proteobacteria; Alphaproteobacteria; Rhizobiales; Bradyrhizobiaceae; Rhodopseudomonas | methyl-accepting chemotaxis protein | 6 |
| **WP_013006923.1** | 2.6E-05 | SNAP.c | -DEVIEVAERVKNIAVDTANNVENTNRIIATLSQLAEVLNDAVNR-------- | *Deferribacter desulfuricans* | Bacteria; Deferribacteres; Deferribacterales; Deferribacteraceae; Deferribacter | methyl-accepting chemotaxis protein | 6 |
| **WP_028138800.1** | 2.6E-05 | Qc.III | IKEISGTIEKLSEISATIASAVEEQGAATQEISRNVQQAAQGTQEVSSNIT-- | *Bradyrhizobium japonicum* | Bacteria; Proteobacteria; Alphaproteobacteria; Rhizobiales; Bradyrhizobiaceae; Bradyrhizobium | chemotaxis protein | 6 |
| **WP_047309193.1** | 2.6E-05 | Qa.III | IRDISGTIEKLSEIASTIASAVEEQGAATQEISRNVQQAAQGTQQ-------- | *Rhodopseudomonas palustris* | Bacteria; Proteobacteria; Alphaproteobacteria; Rhizobiales; Bradyrhizobiaceae; Rhodopseudomonas | HAMP domain-containing protein | 6 |
| **WP_049801921.1** | 2.6E-05 | Qa.IV | IKEIGATIARISDISSTIAATIEEQGAATAEIARNVSEAAK------------ | *Bradyrhizobium* | Bacteria; Proteobacteria; Alphaproteobacteria; Rhizobiales; Bradyrhizobiaceae | MULTISPECIES: methyl-accepting chemotaxis protein | 6 |
| **WP_050995718.1** | 2.6E-05 | Qa.IV | IKEIGATIARISDISSTIAATIEEQGAATAEIARNVSEAAK------------ | *Bradyrhizobium sp. CCBAU 15635* | Bacteria; Proteobacteria; Alphaproteobacteria; Rhizobiales; Bradyrhizobiaceae; Bradyrhizobium | methyl-accepting chemotaxis protein | 6 |
| **WP_051609788.1** | 2.6E-05 | Qc.III | -SSIGETITEIDQIANSIAAAVEEQTAATSEIARNVEQASQGTNEVTINIQS- | *Terasakiella pusilla* | Bacteria; Proteobacteria; Alphaproteobacteria; Rhizobiales; Methylocystaceae; Terasakiella | HAMP domain-containing protein | 6 |
| **WP_057846462.1** | 2.6E-05 | Qa.III | IEAIGKTIGEISEIATTIASAIEQQGAATQEIARNVQQAA------------- | *Bradyrhizobium retamae* | Bacteria; Proteobacteria; Alphaproteobacteria; Rhizobiales; Bradyrhizobiaceae; Bradyrhizobium | HAMP domain-containing protein | 6 |
| **WP_057851416.1** | 2.6E-05 | Qa.III | IEAIGKTIGEISEIATTIASAIEQQGAATQEIARNVQQAA------------- | *Bradyrhizobium valentinum* | Bacteria; Proteobacteria; Alphaproteobacteria; Rhizobiales; Bradyrhizobiaceae; Bradyrhizobium | HAMP domain-containing protein | 6 |
| **WP_061031531.1** | 2.6E-05 | Qa.III | LEEINTSVALISDMATQIATAAEEQTHVTGEITQNITTIK------------- | *Vibrio splendidus* | Bacteria; Proteobacteria; Gammaproteobacteria; Vibrionales; Vibrionaceae; Vibrio | methyl-accepting chemotaxis protein | 6 |
| **WP_063524340.1** | 2.6E-05 | Qa.III | LEEINTSVALISDMATQIATAAEEQTHVTGEITQNITTIK------------- | *Vibrio sp. HI00D65* | Bacteria; Proteobacteria; Gammaproteobacteria; Vibrionales; Vibrionaceae; Vibrio | methyl-accepting chemotaxis protein | 6 |
| **WP_067746289.1** | 2.6E-05 | Qa.III | LQDIATGISQINDVNTQVATATEEQTVVVEDINQNIENIARN----------- | *Alteromonadaceae bacterium XY-R5* | Bacteria; Proteobacteria; Gammaproteobacteria; Alteromonadales; Alteromonadaceae | methyl-accepting chemotaxis protein | 6 |
| **WP_069957631.1** | 2.6E-05 | Qc.III | -GSIGGTISKMNEIASAIAAAVEEQGAATQEIARNVEQAAGGTTEVTT----- | *Magnetovibrio blakemorei* | Bacteria; Proteobacteria; Alphaproteobacteria; Rhodospirillales; Rhodospirillaceae; Magnetovibrio | hypothetical protein | 6 |
| **WP_071989243.1** | 2.6E-05 | R.IV | IDQISKAVSELDSTTQQNAALVQESSAAAASLEQQAGELSQLISLFK------ | *Cronobacter condimenti* | Bacteria; Proteobacteria; Gammaproteobacteria; Enterobacterales; Enterobacteriaceae; Cronobacter | chemotaxis protein | 6 |
| **WP_075797238.1** | 2.6E-05 | Qa.IV | IQNIAATIEEVSAITVTIASAVEEQSAATGEISRTVNQTALATD--------- | *Roseomonas gilardii* | Bacteria; Proteobacteria; Alphaproteobacteria; Rhodospirillales; Acetobacteraceae; Roseomonas | methyl-accepting chemotaxis protein | 6 |
| **WP_076573605.1** | 2.6E-05 | Qa.III | LEQIVESIDRISQMATQISSAAEEQRSVTDEVSRNIQAVK------------- | *Aeromonas sp. RU39B* | Bacteria; Proteobacteria; Gammaproteobacteria; Aeromonadales; Aeromonadaceae; Aeromonas | methyl-accepting chemotaxis protein | 6 |
| **WP_080136710.1** | 2.6E-05 | Qa.III.b | -EAIAQTIAEINGITSSIATSIEQQGLATREIARNVQEASA------------ | *Bradyrhizobium sp. BR10280* | Bacteria; Proteobacteria; Alphaproteobacteria; Rhizobiales; Bradyrhizobiaceae; Bradyrhizobium | methyl-accepting chemotaxis protein | 6 |
| **WP_089202243.1** | 2.6E-05 | Qa.III | LDEINAXVALISDMATQIATAAEEQTHVTNEITQNVTSIK------------- | *Vibrio parahaemolyticus* | Bacteria; Proteobacteria; Gammaproteobacteria; Vibrionales; Vibrionaceae; Vibrio | methyl-accepting chemotaxis protein | 6 |
| **WP_090070847.1** | 2.6E-05 | Qa.III | IHAIKEVIQQVSTISSSIATAIEQQSAVTRDISGNMQTAS------------- | *Cohaesibacter marisflavi* | Bacteria; Proteobacteria; Alphaproteobacteria; Rhizobiales; Cohaesibacteraceae | PAS domain S-box protein | 6 |
| **WP_097281368.1** | 2.6E-05 | Qc.III | IQSINKVIDRMSEISVAVSAAVEEQNAATAEIARSVDQTSQGTKEVSRSI--- | *Caenispirillum bisanense* | Bacteria; Proteobacteria; Alphaproteobacteria; Rhodospirillales; Rhodospirillaceae; Caenispirillum | HAMP domain-containing protein | 6 |
| **WP_097668298.1** | 2.6E-05 | Qa.IV | IKEIGATIARISDISSTIAATIEEQGAATAEIARNVSEAAK------------ | *Bradyrhizobium ottawaense* | Bacteria; Proteobacteria; Alphaproteobacteria; Rhizobiales; Bradyrhizobiaceae; Bradyrhizobium | methyl-accepting chemotaxis protein | 6 |
| **WP_105056983.1** | 2.6E-05 | Qa.III | LEEINTSVALISDMATQIATAAEEQTHVTGEITQNITTIK------------- | *Vibrio splendidus* | Bacteria; Proteobacteria; Gammaproteobacteria; Vibrionales; Vibrionaceae; Vibrio | methyl-accepting chemotaxis protein | 6 |
| **WP_109331602.1** | 2.6E-05 | Qc.III | ------TIARINEISAQISAAVGQQSAAVDEIASSITHTAAGTRDVST----- | *Azospirillum sp. CFH 70021* | Bacteria; Proteobacteria; Alphaproteobacteria; Rhodospirillales; Rhodospirillaceae; Azospirillum | HAMP domain-containing protein | 6 |
| **WP_109444105.1** | 2.6E-05 | Qa.III | IRSIAGTIREVNEIAATIAAAVEQQSAATREIARNVGEAAD------------ | *Azospirillum sp. TSH100* | Bacteria; Proteobacteria; Alphaproteobacteria; Rhodospirillales; Rhodospirillaceae; Azospirillum | HAMP domain-containing protein | 6 |
| **WP_111921720.1** | 2.6E-05 | Qb.I | ----------AVEKMEQVNKISAAQSESVTRTEEKFEGIANSLEVTKNVVKKL | *Clostridium cochlearium* | Bacteria; Firmicutes; Clostridia; Clostridiales; Clostridiaceae; Clostridium | methyl-accepting chemotaxis protein | 6 |
| **WP_112712551.1** | 2.6E-05 | Qc.III | LDSMTDSVMQINDLGIQIATAAEEQSSVTEEINRNMTMIQ------------- | *Moritella yayanosii* | Bacteria; Proteobacteria; Gammaproteobacteria; Alteromonadales; Moritellaceae; Moritella | methyl-accepting chemotaxis protein | 6 |
| **WP_114373777.1** | 2.6E-05 | Qa.III | IDAIREVIEKLNSISQSVAAAVEEQSATTQEIANNVARTNEAVQQVSQS---- | *Blastopirellula cremea* | Bacteria; Planctomycetes; Planctomycetia; Planctomycetales; Planctomycetaceae; Blastopirellula | HAMP domain-containing protein | 6 |
| **ACB82311.1** | 2.7E-05 | Qc.III.c | -DGISASVRAINETATAIASAIEEQGAATREIVQAIAQASS------------ | *Methylorubrum populi BJ001* | Bacteria; Proteobacteria; Alphaproteobacteria; Rhizobiales; Methylobacteriaceae; Methylorubrum | methyl-accepting chemotaxis sensory transducer | 6 |
| **PWS55586.1** | 2.7E-05 | Qa.III | ------SVTTINDMSTQIATAATEQSSVTEEINRNVY----AIQEIVNELT-- | *Pseudoalteromonas sp. meg-B1* | Bacteria; Proteobacteria; Gammaproteobacteria; Alteromonadales; Pseudoalteromonadaceae; Pseudoalteromonas | methyl-accepting chemotaxis protein | 6 |
| **WP_007091957.1** | 2.7E-05 | Qc.III | ---ISETIAKVADSSTAIAAAVEEQNAAIGEISRNVQQASLGTNEVSERIG-- | *Thalassospira xiamenensis* | Bacteria; Proteobacteria; Alphaproteobacteria; Rhodospirillales; Rhodospirillaceae; Thalassospira | methyl-accepting chemotaxis protein | 6 |
| **WP_008111551.1** | 2.7E-05 | Qa.III | ------SVTTINDMSTQIATAATEQSSVTEEINRNVY----AIQEIVNELT-- | *Alteromonadales* | Bacteria; Proteobacteria; Gammaproteobacteria | MULTISPECIES: methyl-accepting chemotaxis protein | 6 |
| **WP_014250130.1** | 2.7E-05 | Qa.III | IRSIAGTIREVNEIAATIAAAVEQQSAATREIARNVGEAAD------------ | *Azospirillum lipoferum* | Bacteria; Proteobacteria; Alphaproteobacteria; Rhodospirillales; Rhodospirillaceae; Azospirillum | HAMP domain-containing protein | 6 |
| **WP_024602336.1** | 2.7E-05 | Qa.III | ------SVTTINDMSTQIATAATEQSSVTEEINRNVY----AIQEIVNELT-- | *Pseudoalteromonas* | Bacteria; Proteobacteria; Gammaproteobacteria; Alteromonadales; Pseudoalteromonadaceae | MULTISPECIES: methyl-accepting chemotaxis protein | 6 |
| **WP_028833944.1** | 2.7E-05 | Qa.III | ------SVTTINDMSTQIATAATEQSSVTEEINRNVY----AIQEIVNELT-- | *Pseudoalteromonas* | Bacteria; Proteobacteria; Gammaproteobacteria; Alteromonadales; Pseudoalteromonadaceae | MULTISPECIES: methyl-accepting chemotaxis protein | 6 |
| **WP_029291090.1** | 2.7E-05 | Qa.III | --RICEEVGQISDMSQQMAAAIEQQGQAAEQINQKINEIAGFTERSSQQA--- | *Pseudomonas fluorescens* | Bacteria; Proteobacteria; Gammaproteobacteria; Pseudomonadales; Pseudomonadaceae; Pseudomonas | PAS domain-containing protein | 6 |
| **WP_036981615.1** | 2.7E-05 | Qa.III | ------SVTTINDMSTQIATAATEQSSVTEEINRNVY----AIQEIVNELT-- | *Pseudoalteromonas sp. SCSIO_11900* | Bacteria; Proteobacteria; Gammaproteobacteria; Alteromonadales; Pseudoalteromonadaceae; Pseudoalteromonas | methyl-accepting chemotaxis protein | 6 |
| **WP_055013046.1** | 2.7E-05 | Qa.III | ------SVTTINDMSTQIATAATEQSSVTEEINRNVY----AIQEIVNELT-- | *Pseudoalteromonas sp. P1-16-1b* | Bacteria; Proteobacteria; Gammaproteobacteria; Alteromonadales; Pseudoalteromonadaceae; Pseudoalteromonas | methyl-accepting chemotaxis protein | 6 |
| **WP_063635939.1** | 2.7E-05 | Qa.III | IRSIAGTIREVNEIAATIAAAVEQQSAATREIARNVGEAAD------------ | *Azospirillum humicireducens* | Bacteria; Proteobacteria; Alphaproteobacteria; Rhodospirillales; Rhodospirillaceae; Azospirillum | HAMP domain-containing protein | 6 |
| **WP_069440027.1** | 2.7E-05 | Qa.III | ------SVTTINDMSTQIATAATEQSSVTEEINRNVY----AIQEIVNELT-- | *Pseudoalteromonas tetraodonis* | Bacteria; Proteobacteria; Gammaproteobacteria; Alteromonadales; Pseudoalteromonadaceae; Pseudoalteromonas | methyl-accepting chemotaxis protein | 6 |
| **WP_075169718.1** | 2.7E-05 | Qa.III | ------SVTTINDMSTQIATAATEQSSVTEEINRNVY----AIQEIVNELT-- | *Pseudoalteromonas haloplanktis* | Bacteria; Proteobacteria; Gammaproteobacteria; Alteromonadales; Pseudoalteromonadaceae; Pseudoalteromonas | methyl-accepting chemotaxis protein | 6 |
| **WP_076918321.1** | 2.7E-05 | Qa.III | ------SVTTINDMSTQIATAATEQSSVTEEINRNVY----AIQEIVNELT-- | *Pseudoalteromonas sp. SK18* | Bacteria; Proteobacteria; Gammaproteobacteria; Alteromonadales; Pseudoalteromonadaceae; Pseudoalteromonas | methyl-accepting chemotaxis protein | 6 |
| **WP_076920662.1** | 2.7E-05 | Qa.III | ------SVTTINDMSTQIATAATEQSSVTEEINRNVY----AIQEIVNELT-- | *Pseudoalteromonas sp. SK20* | Bacteria; Proteobacteria; Gammaproteobacteria; Alteromonadales; Pseudoalteromonadaceae; Pseudoalteromonas | methyl-accepting chemotaxis protein | 6 |

| | | | | | | | | |
|---|---|---|---|---|---|---|---|---|
|  | **WP_083223127.1** | 2.7E-05 | Qa.IV | IQQIGETIKDINEVSTAVAAGVEEQGNATNEIANSAEQAA------------ | *Terasakiella sp. PR1* | Bacteria; Proteobacteria; Alphaproteobacteria; Rhizobiales; Methylocystaceae; Terasakiella | HAMP domain-containing protein | 6 |
|  | **WP_085579416.1** | 2.7E-05 | Qa.IV | ---------EVSRLSNDISASVEQQTAAINEISGSAQEVSTHMKGIADNI--- | *Thalassospira mesophila* | Bacteria; Proteobacteria; Alphaproteobacteria; Rhodospirillales; Rhodospirillaceae; Thalassospira | chemotaxis protein | 6 |
|  | **WP_090495329.1** | 2.7E-05 | Qa.III | ------SVTTINDMSTQIATAATEQSSVTEEINRNVY----AIQEIVNELT-- | *Pseudoalteromonas* | Bacteria; Proteobacteria; Gammaproteobacteria; Alteromonadales; Pseudoalteromonadaceae | MULTISPECIES: methyl-accepting chemotaxis protein | 6 |
| | **WP_104912555.1** | 2.7E-05 | Qa.III | --RICEEVGQISDMSQQMAAAIEQQGQAAEQINQKINEIAGFTERSSQQA--- | *Pseudomonas sp. LG1D9* | Bacteria; Proteobacteria; Gammaproteobacteria; Pseudomonadales; Pseudomonadaceae; Pseudomonas | PAS domain-containing protein | 6 |
| | **AWL97435.1** | 2.8E-05 | Qc.III | IGEISGTIARLSEIAATIAAAVEQQGAATQEIARNVQQAAQGTQQVSSNVG-- | *Bradyrhizobium ottawaense* | Bacteria; Proteobacteria; Alphaproteobacteria; Rhizobiales; Bradyrhizobiaceae; Bradyrhizobium | PAS domain S-box protein | 6 |
|  | **GBE44239.1** | 2.8E-05 | Qa.III | IRGIAETMGEVSAATESIAAAVEQQGACTSEISNNVQQAA------------ | *bacterium BMS3Bbin10* | Bacteria | methyl-accepting chemotaxis protein 4 | 6 |
|  | **KRQ02621.1** | 2.8E-05 | Qa.IV | -KEIGTTIGRISDISTTIAATIEQQGAATAEIARNVSEAAK----------- | *Bradyrhizobium yuanmingense* | Bacteria; Proteobacteria; Alphaproteobacteria; Rhizobiales; Bradyrhizobiaceae; Bradyrhizobium | chemotaxis protein | 6 |
| | **PHS23021.1** | 2.8E-05 | Qc.III | -GEIRSTIDEMNEVSVAINAAVEEQAAATQEIARNTQEAATGTQDVSRNI--- | *Robiginitomaculum sp.* | Bacteria; Proteobacteria; Alphaproteobacteria; Rhodobacterales; Hyphomonadaceae; Robiginitomaculum | chemotaxis protein | 6 |
| | **PHS79319.1** | 2.8E-05 | Qc.III | -KGIGSTISRIDEASTAIASAVEEQGAATQEIARNVEQAANGTNEVTSNITE- | *Rhodospirillaceae bacterium* | Bacteria; Proteobacteria; Alphaproteobacteria; Rhodospirillales; Rhodospirillaceae | chemotaxis protein | 6 |
|  | **PSQ27044.1** | 2.8E-05 | SNAP.c | ------VASEVKELAEDTKDAAEEIEERLERIQAQTERTADEVQKTSTQV--- | *Halobacteriales archaeon QS_9_68_17* | Archaea; Euryarchaeota; Halobacteria; Halobacteriales | chemotaxis protein | 6 |
|  | **PTL46714.1** | 2.8E-05 | Qa.III | IDAIRRVVGNLSEISRTIASSIEEQSTATSEIANNVAQSSSAARQ-------- | *Blastopirellula marina* | Bacteria; Planctomycetes; Planctomycetia; Planctomycetales; Planctomycetaceae; Blastopirellula | methyl-accepting chemotaxis protein | 6 |
|  | **SCB16746.1** | 2.8E-05 | Qa.IV | -KEIGTTIGRISDISTTIAATIEQQGAATAEIARNVSEAAK----------- | *Bradyrhizobium yuanmingense* | Bacteria; Proteobacteria; Alphaproteobacteria; Rhizobiales; Bradyrhizobiaceae; Bradyrhizobium | methyl-accepting chemotaxis protein | 6 |
|  | **WP_011088255.1** | 2.8E-05 | Qa.IV | IRDISATIERLSEVSSTIPAAVEEQGAATQEISRNVQQAA------------ | *Bradyrhizobium diazoefficiens* | Bacteria; Proteobacteria; Alphaproteobacteria; Rhizobiales; Bradyrhizobiaceae; Bradyrhizobium | hypothetical protein | 6 |
| | **WP_028143382.1** | 2.8E-05 | Qc.III | IGEISGTIARLSEIAATIAAAVEQQGAATQEIARNVQQAAQGTQQVSSNVG-- | *Bradyrhizobium* | Bacteria; Proteobacteria; Alphaproteobacteria; Rhizobiales; Bradyrhizobiaceae | MULTISPECIES: PAS domain S-box protein | 6 |
|  | **WP_034469452.1** | 2.8E-05 | Qa.IV | IREIGKTIATISEISSTIAAAVEEQGAATKEIARNMQQASQ----------- | *Afipia sp. P52-10* | Bacteria; Proteobacteria; Alphaproteobacteria; Rhizobiales; Bradyrhizobiaceae; Afipia | methyl-accepting chemotaxis protein | 6 |
|  | **WP_036002948.1** | 2.8E-05 | Qa.IV | -KEIGTTIGRISDISTTIAATIEQQGAATAEIARNVSEAAK----------- | *Bradyrhizobium yuanmingense* | Bacteria; Proteobacteria; Alphaproteobacteria; Rhizobiales; Bradyrhizobiaceae; Bradyrhizobium | chemotaxis protein | 6 |
|  | **WP_036024236.1** | 2.8E-05 | Qa.IV | -KEIGTTIGRISDISTTIAATIEQQGAATAEIARNVSEAAK----------- | *Bradyrhizobium yuanmingense* | Bacteria; Proteobacteria; Alphaproteobacteria; Rhizobiales; Bradyrhizobiaceae; Bradyrhizobium | chemotaxis protein | 6 |
|  | **WP_036038846.1** | 2.8E-05 | Qa.IV | -KEIGTTIGRISDISTTIAATIEQQGAATAEIARNVSEAAK----------- | *Bradyrhizobium yuanmingense* | Bacteria; Proteobacteria; Alphaproteobacteria; Rhizobiales; Bradyrhizobiaceae; Bradyrhizobium | chemotaxis protein | 6 |
| | **WP_038947730.1** | 2.8E-05 | Qc.III | IGEISGTIARLSEIAATIAAAVEQQGAATQEIARNVQQAAQGTQQVSSNVG-- | *Bradyrhizobium* | Bacteria; Proteobacteria; Alphaproteobacteria; Rhizobiales; Bradyrhizobiaceae | MULTISPECIES: PAS domain S-box protein | 6 |
| | **WP_038973707.1** | 2.8E-05 | Qc.III | IGEISGTIARLSEIAATIAAAVEQQGAATQEIARNVQQAAQGTQQVSSNVG-- | *Bradyrhizobium sp. CCBAU 15635* | Bacteria; Proteobacteria; Alphaproteobacteria; Rhizobiales; Bradyrhizobiaceae; Bradyrhizobium | PAS domain S-box protein | 6 |
| | **WP_041278040.1** | 2.8E-05 | Qb.I | ---VAKMISEDVGHVHQAAEEIADSSSDVKRNAEELSDLSTQLK---KMV--- | *Desulfotalea psychrophila* | Bacteria; Proteobacteria; Deltaproteobacteria; Desulfobacterales; Desulfobulbaceae; Desulfotalea | methyl-accepting chemotaxis protein | 6 |
|  | **WP_042693753.1** | 2.8E-05 | Qa.III | ----TRTIERINQIAGQISAGVSQQSTAVDEIASNINDVAANTRD-------- | *Azospirillum sp. B506* | Bacteria; Proteobacteria; Alphaproteobacteria; Rhodospirillales; Rhodospirillaceae; Azospirillum | HAMP domain-containing protein | 6 |
|  | **WP_058191872.1** | 2.8E-05 | Qc.III | --SIQGTIVKLNEVATAISSAVEEQTAVTREMSASMQTAAQG----------- | *Methylobacterium sp. GXS13* | Bacteria; Proteobacteria; Alphaproteobacteria; Rhizobiales; Methylobacteriaceae; Methylobacterium | methyl-accepting chemotaxis protein | 6 |
|  | **WP_058760714.1** | 2.8E-05 | Qb.I | LQRTKEQLQLTIEQSETSTEELKASNEELQAINEELRSATEELETSK------ | *Pseudomonas psychrotolerans* | Bacteria; Proteobacteria; Gammaproteobacteria; Pseudomonadales; Pseudomonadaceae; Pseudomonas | PAS domain S-box protein | 6 |
|  | **WP_058766308.1** | 2.8E-05 | Qb.I | LQRTKEQLQLTIEQSETSTEELKASNEELQAINEELRSATEELETSK------ | *Pseudomonas psychrotolerans* | Bacteria; Proteobacteria; Gammaproteobacteria; Pseudomonadales; Pseudomonadaceae; Pseudomonas | PAS domain S-box protein | 6 |
|  | **WP_058772750.1** | 2.8E-05 | Qb.I | LQRTKEQLQLTIEQSETSTEELKASNEELQAINEELRSATEELETSK------ | *Pseudomonas psychrotolerans* | Bacteria; Proteobacteria; Gammaproteobacteria; Pseudomonadales; Pseudomonadaceae; Pseudomonas | PAS domain S-box protein | 6 |
|  | **WP_058783224.1** | 2.8E-05 | Qb.I | LQRTKEQLQLTIEQSETSTEELKASNEELQAINEELRSATEELETSK------ | *Pseudomonas psychrotolerans* | Bacteria; Proteobacteria; Gammaproteobacteria; Pseudomonadales; Pseudomonadaceae; Pseudomonas | PAS domain S-box protein | 6 |
|  | **WP_058787802.1** | 2.8E-05 | Qb.I | LQRTKEQLQLTIEQSETSTEELKASNEELQAINEELRSATEELETSK------ | *Pseudomonas psychrotolerans* | Bacteria; Proteobacteria; Gammaproteobacteria; Pseudomonadales; Pseudomonadaceae; Pseudomonas | PAS domain S-box protein | 6 |
|  | **WP_058789125.1** | 2.8E-05 | Qb.I | LQRTKEQLQLTIEQSETSTEELKASNEELQAINEELRSATEELETSK------ | *Pseudomonas psychrotolerans* | Bacteria; Proteobacteria; Gammaproteobacteria; Pseudomonadales; Pseudomonadaceae; Pseudomonas | PAS domain S-box protein | 6 |
|  | **WP_074528587.1** | 2.8E-05 | Qb.I | LQRTKEQLQLTIEQSETSTEELKASNEELQAINEELRSATEELETSK------ | *Pseudomonas psychrotolerans* | Bacteria; Proteobacteria; Gammaproteobacteria; Pseudomonadales; Pseudomonadaceae; Pseudomonas | PAS domain S-box protein | 6 |
| | **WP_080135668.1** | 2.8E-05 | Qc.III | IQEIGGTIERLSEVSAAIAAAVEQQGAATQEISRNVQQASLGTQEVSSNIT-- | *Bradyrhizobium sp. BR10280* | Bacteria; Proteobacteria; Alphaproteobacteria; Rhizobiales; Bradyrhizobiaceae; Bradyrhizobium | methyl-accepting chemotaxis protein | 6 |
|  | **WP_081830016.1** | 2.8E-05 | Qb.I | ----RDLVADAVAQANNSAKNIDESSASIKGSVDTFDEIYNNIQETNVLIK-- | *Butyrivibrio sp. NC2002* | Bacteria; Firmicutes; Clostridia; Clostridiales; Lachnospiraceae; Butyrivibrio | methyl-accepting chemotaxis protein | 6 |
| | **WP_082837097.1** | 2.8E-05 | Qc.III | IQSISEIIENMNSVAATMAASVEEQTAATSEIASNVDNASRGTNVVSENISR- | *Labrenzia sp. OB1* | Bacteria; Proteobacteria; Alphaproteobacteria; Rhodobacterales; Rhodobacteraceae; Labrenzia | methyl-accepting chemotaxis protein | 6 |
|  | **WP_092029580.1** | 2.8E-05 | Qc.III | --TIGGTIERLSEISSTIAAAVEEQGAATSEISRNVQNAAAGTTQVS------ | *Bradyrhizobium sp. OK095* | Bacteria; Proteobacteria; Alphaproteobacteria; Rhizobiales; Bradyrhizobiaceae; Bradyrhizobium | methyl-accepting chemotaxis protein | 6 |
|  | **WP_103674679.1** | 2.8E-05 | R.IV | --EINHAVNDLDRMVQQNAELVVESAAAAGALQGQAGDLAETAGHFR------ | *unclassified Enterobacteriaceae (miscellaneous)* | Bacteria; Proteobacteria; Gammaproteobacteria; Enterobacterales; Enterobacteriaceae | MULTISPECIES: methyl-accepting chemotaxis protein | 6 |
|  | **WP_109141992.1** | 2.8E-05 | Qa.IV | -KEIGTTIGRISDISTTIAATIEQQGAATAEIARNVSEAAK----------- | *Bradyrhizobium sp. SUTN9-2* | Bacteria; Proteobacteria; Alphaproteobacteria; Rhizobiales; Bradyrhizobiaceae; Bradyrhizobium | chemotaxis protein | 6 |
|  | **WP_109443154.1** | 2.8E-05 | Qa.III | IQSISNIIEQIQGISTTISSAVEEQGSATREISRNVQQAAQGTQQ-------- | *Azospirillum sp. TSH100* | Bacteria; Proteobacteria; Alphaproteobacteria; Rhodospirillales; Rhodospirillaceae; Azospirillum | methyl-accepting chemotaxis protein | 6 |
|  | **WP_110116426.1** | 2.8E-05 | Qa.IV | IRDISATIERLSEVSSTIPAAVEEQGAATQEISRNVQQAA------------ | *Bradyrhizobium diazoefficiens* | Bacteria; Proteobacteria; Alphaproteobacteria; Rhizobiales; Bradyrhizobiaceae; Bradyrhizobium | hypothetical protein | 6 |
|  | **WP_111070476.1** | 2.8E-05 | Qa.IV | ----------LNDIAASIATAVEQQGAATSEIARNVQQAAGGTAQASQNI--- | *Azospirillum sp. M2T2B2* | Bacteria; Proteobacteria; Alphaproteobacteria; Rhodospirillales; Rhodospirillaceae; Azospirillum | HAMP domain-containing protein | 6 |
| | **AWL93874.1** | 2.9E-05 | Qc.III | --TIGGTIERLSEISSTIAAAVEEQGAATGEISRNVQNAAAGTTQVSANI--- | *Bradyrhizobium ottawaense* | Bacteria; Proteobacteria; Alphaproteobacteria; Rhizobiales; Bradyrhizobiaceae; Bradyrhizobium | methyl-accepting chemotaxis protein | 6 |
| | **AWL99357.1** | 2.9E-05 | Qc.III | --TIGGTIERLSEISSTIAAAVEEQGAATGEISRNVQNAAAGTTQVSANI--- | *Bradyrhizobium sp. 2 39S1MB* | Bacteria; Proteobacteria; Alphaproteobacteria; Rhizobiales; Bradyrhizobiaceae; Bradyrhizobium | methyl-accepting chemotaxis protein | 6 |
|  | **AXA59410.1** | 2.9E-05 | Qb.I | LQRTKLHLQDTIEQAEVSSEELKASNEEMQAINEELRSATEELETS------- | *Pseudomonas thivervalensis* | Bacteria; Proteobacteria; Gammaproteobacteria; Pseudomonadales; Pseudomonadaceae; Pseudomonas | chemotaxis protein | 6 |
|  | **KEX94916.1** | 2.9E-05 | Qb.I | LQRTKMHLQETIEQSEVSSEELKASNEEMQAINEELRSATEELETS------- | *Pseudomonas putida* | Bacteria; Proteobacteria; Gammaproteobacteria; Pseudomonadales; Pseudomonadaceae; Pseudomonas | hypothetical protein HA62_03715 partial | 6 |
| | **KJC51439.1** | 2.9E-05 | Qa.IV | ---IGATIGQISSISTSIASAVEQQGAATQEIARSVQTVAQGTQTAATDI--- | *Bradyrhizobium sp. LTSP857* | Bacteria; Proteobacteria; Alphaproteobacteria; Rhizobiales; Bradyrhizobiaceae; Bradyrhizobium | chemotaxis protein partial | 6 |
|  | **OLF73962.1** | 2.9E-05 | Qc.III | --RIGSVIGQIEEIASGISAAVEEQVAVTQDISRNMGEAASD----------- | *Maricaulis sp. W15* | Bacteria; Proteobacteria; Alphaproteobacteria; Rhodobacterales; Hyphomonadaceae; Maricaulis | hypothetical protein AWH62_07375 | 6 |
|  | **OYX87840.1** | 2.9E-05 | Qa.IV | ---------DIRQVTAAIASAVEQQGAATREIAQNTQHAAHGTEAVNDNIQ-- | *Azorhizobium sp. 32-67-21* | Bacteria; Proteobacteria; Alphaproteobacteria; Rhizobiales; Xanthobacteraceae; Azorhizobium | hypothetical protein B7Y84_10390 | 6 |
|  | **OYY09574.1** | 2.9E-05 | Qa.IV | ---------DIRQVTAAIASAVEQQGAATREIAQNTQHAAHGTEAVNDNIQ-- | *Rhizobiales bacterium 35-68-8* | Bacteria; Proteobacteria; Alphaproteobacteria; Rhizobiales | hypothetical protein B7Y70_09895 | 6 |
| | **SFO62322.1** | 2.9E-05 | Qc.III | -EQLTKIASETVTYAEAVEAKVTDQNSLISETNQTADFLNQQVNQLNKLIDDL | *Halolactibacillus alkaliphilus* | Bacteria; Firmicutes; Bacilli; Bacillales; Bacillaceae; Halolactibacillus | methyl-accepting chemotaxis protein | 6 |
|  | **WP_003174485.1** | 2.9E-05 | Qb.I | LQRTKLHLQDTIEQAEVSSEELKASNEEMQAINEELRSATEELETS------- | *Pseudomonas fluorescens* | Bacteria; Proteobacteria; Gammaproteobacteria; Pseudomonadales; Pseudomonadaceae; Pseudomonas | PAS domain S-box protein | 6 |
|  | **WP_017529043.1** | 2.9E-05 | Qb.I | LQRTKLHLQDTIEQAEVSSEELKASNEEMQAINEELRSATEELETS------- | *Pseudomonas fluorescens* | Bacteria; Proteobacteria; Gammaproteobacteria; Pseudomonadales; Pseudomonadaceae; Pseudomonas | PAS domain S-box protein | 6 |
| | **WP_018318277.1** | 2.9E-05 | Qc.III | --TIGGTIERLSEISSTIAAAVEEQGAATGEISRNVQNAAAGTTQVSANI--- | *Bradyrhizobium sp. WSM2793* | Bacteria; Proteobacteria; Alphaproteobacteria; Rhizobiales; Bradyrhizobiaceae; Bradyrhizobium | methyl-accepting chemotaxis protein | 6 |
| | **WP_028140512.1** | 2.9E-05 | Qc.III | --TIGGTIERLSEISSTIAAAVEEQGAATGEISRNVQNAAAGTTQVSANI--- | *Bradyrhizobium* | Bacteria; Proteobacteria; Alphaproteobacteria; Rhizobiales; Bradyrhizobiaceae | MULTISPECIES: methyl-accepting chemotaxis protein | 6 |
|  | **WP_028634006.1** | 2.9E-05 | Qb.I | LQRTRLQLQETIEQSEVSSEELTASNEEMQAINEELRSASEELETSK------ | *Pseudomonas parafulva* | Bacteria; Proteobacteria; Gammaproteobacteria; Pseudomonadales; Pseudomonadaceae; Pseudomonas | response regulator | 6 |
| | **WP_038948126.1** | 2.9E-05 | Qc.III | --TIGGTIERLSEISSTIAAAVEEQGAATGEISRNVQNAAAGTTQVSANI--- | *Bradyrhizobium sp. CCBAU 15544* | Bacteria; Proteobacteria; Alphaproteobacteria; Rhizobiales; Bradyrhizobiaceae; Bradyrhizobium | methyl-accepting chemotaxis protein | 6 |
| | **WP_038971112.1** | 2.9E-05 | Qc.III | --TIGGTIERLSEISSTIAAAVEEQGAATGEISRNVQNAAAGTTQVSANI--- | *Bradyrhizobium* | Bacteria; Proteobacteria; Alphaproteobacteria; Rhizobiales; Bradyrhizobiaceae | MULTISPECIES: methyl-accepting chemotaxis protein | 6 |
|  | **WP_053188106.1** | 2.9E-05 | Qb.I | LQRTKLHLQDTIEQAEVSSEELKASNEEMQAINEELRSATEELETS------- | *Pseudomonas thivervalensis* | Bacteria; Proteobacteria; Gammaproteobacteria; Pseudomonadales; Pseudomonadaceae; Pseudomonas | PAS domain S-box protein | 6 |
|  | **WP_057849261.1** | 2.9E-05 | Qc.III | IKEISGTIGRVSEIAAAIAAAVEEQGAATQEIARNVQQAAVGS---------- | *Bradyrhizobium valentinum* | Bacteria; Proteobacteria; Alphaproteobacteria; Rhizobiales; Bradyrhizobiaceae; Bradyrhizobium | methyl-accepting chemotaxis protein | 6 |
|  | **WP_060483396.1** | 2.9E-05 | Qb.I | LQRTRLQLQETIEQSEVSSEELTASNEEMQAINEELRSASEELETSK------ | *Pseudomonas sp. NBRC 111119* | Bacteria; Proteobacteria; Gammaproteobacteria; Pseudomonadales; Pseudomonadaceae; Pseudomonas | response regulator | 6 |
|  | **WP_065879133.1** | 2.9E-05 | Qb.I | LQRTKLHLQDTIEQAEVSSEELKASNEEMQAINEELRSATEELETS------- | *Pseudomonas* | Bacteria; Proteobacteria; Gammaproteobacteria; Pseudomonadales; Pseudomonadaceae | MULTISPECIES: PAS domain S-box protein | 6 |
|  | **WP_078732627.1** | 2.9E-05 | Qb.I | LQRTKLHLQDTIEQAEVSSEELKASNEEMQAINEELRSATEELETS------- | *Pseudomonas fluorescens* | Bacteria; Proteobacteria; Gammaproteobacteria; Pseudomonadales; Pseudomonadaceae; Pseudomonas | PAS domain S-box protein | 6 |

| | | | | | | | | |
|---|---|---|---|---|---|---|---|---|
|  | WP_078824068.1 | 2.9E-05 | Qb.I | LQRTKLHLQDTIEQAEVSSEELKASNEEMQAINEELRSATEELETS------- | *Pseudomonas fluorescens* | Bacteria; Proteobacteria; Gammaproteobacteria; Pseudomonadales; Pseudomonadaceae; Pseudomonas | PAS domain S-box protein | 6 |
|  | WP_083635133.1 | 2.9E-05 | Qc.III | --RIGSVIGQIEEIASGISAAVEEQVAVTQDISRNMGEAASD----------- | *Maricaulis sp. W15* | Bacteria; Proteobacteria; Alphaproteobacteria; Rhodobacterales; Hyphomonadaceae; Maricaulis | methyl-accepting chemotaxis protein | 6 |
|  | WP_083901334.1 | 2.9E-05 | Qc.III | --AIADTIGEINRISAAISAAVDQQSAAANDIAGSVTKAAAGTNEVS------ | *Azospirillum sp. B4* | Bacteria; Proteobacteria; Alphaproteobacteria; Rhodospirillales; Rhodospirillaceae; Azospirillum | methyl-accepting chemotaxis protein | 6 |
| | WP_090018132.1 | 2.9E-05 | Qa.III | -EAVADNMREIDEAASSIASAVEEQTSSTREISRNVEEASTGVQDVSTQL--- | *Limimonas halophila* | Bacteria; Proteobacteria; Alphaproteobacteria; Rhodospirillales; Rhodospirillaceae; Limimonas | HAMP domain-containing protein | 6 |
|  | WP_092199839.1 | 2.9E-05 | Qb.I | LQRTKLHLQDTIEQAEVSSEELKASNEEMQAINEELRSATEELETS------- | *Pseudomonas* | Bacteria; Proteobacteria; Gammaproteobacteria; Pseudomonadales; Pseudomonadaceae | MULTISPECIES: PAS domain S-box protein | 6 |
|  | WP_092253552.1 | 2.9E-05 | Qb.I | LQRTKLHLQDTIEQAEVSSEELKASNEEMQAINEELRSATEELETS------- | *Pseudomonas sp. NFACC13-1* | Bacteria; Proteobacteria; Gammaproteobacteria; Pseudomonadales; Pseudomonadaceae; Pseudomonas | PAS domain S-box protein | 6 |
| | WP_092254987.1 | 2.9E-05 | Qc.III | --TIGGTIERLSEISSTIAAAVEEQGAATGEISRNVQNAAAGTTQVSANI--- | *Bradyrhizobium sp. Rc3b* | Bacteria; Proteobacteria; Alphaproteobacteria; Rhizobiales; Bradyrhizobiaceae; Bradyrhizobium | methyl-accepting chemotaxis protein | 6 |
|  | WP_109169271.1 | 2.9E-05 | Qa.III | LDEINAAVALISDMATQIATAAEEQTHVTNEITQNVTSIK------------ | *Vibrio sp. T9* | Bacteria; Proteobacteria; Gammaproteobacteria; Vibrionales; Vibrionaceae; Vibrio | methyl-accepting chemotaxis protein | 6 |
|  | AWL91825.1 | 3E-05 | Qc.III | -QTIGGIIGEVNEVATAIAAAVQEQGAATQEITRSTQFAAQGTKNVS------ | *Bradyrhizobium ottawaense* | Bacteria; Proteobacteria; Alphaproteobacteria; Rhizobiales; Bradyrhizobiaceae; Bradyrhizobium | methyl-accepting chemotaxis protein | 6 |
|  | AWM03451.1 | 3E-05 | Qc.III | -QTIGGIIGEVNEVATAIAAAVQEQGAATQEITRSTQFAAQGTKNVS------ | *Bradyrhizobium sp. 2 39S1MB* | Bacteria; Proteobacteria; Alphaproteobacteria; Rhizobiales; Bradyrhizobiaceae; Bradyrhizobium | methyl-accepting chemotaxis protein | 6 |
|  | EWY38200.1 | 3E-05 | Qa.IV | IQNITETIKRMSEISSGIAAAVQEQAAATQEIARNVNHAA------------ | *Skermanella stibiiresistens SB22* | Bacteria; Proteobacteria; Alphaproteobacteria; Rhodospirillales; Rhodospirillaceae; Skermanella | methyl-accepting chemotaxis protein | 6 |
|  | KPZ01119.1 | 3E-05 | Qa.III | -------IEEINGMNQSVAAATEEQTSVVESINVDITHINTLNQQGVDNLRQT | *Pseudomonas syringae pv. spinaceae* | Bacteria; Proteobacteria; Gammaproteobacteria; Pseudomonadales; Pseudomonadaceae; Pseudomonas; Pseudomonas syringae | Methyl-accepting chemotaxis protein partial | 6 |
|  | KZD22396.1 | 3E-05 | Qc.III | -KSIGGIIGEVNEVATAIAAAVEEQGAATQEITRSTQQAADGTRNVS------ | *Tardiphaga robiniae* | Bacteria; Proteobacteria; Alphaproteobacteria; Rhizobiales; Bradyrhizobiaceae; Tardiphaga | chemotaxis protein | 6 |
|  | OEU71906.1 | 3E-05 | Qc.III | --QISSGADQISDTSQQLSQAATEQASSLEEINSSMTEMASRTKQ-------- | *Desulfuromonadales bacterium C00003068* | Bacteria; Proteobacteria; Deltaproteobacteria; Desulfuromonadales | hypothetical protein BA874_00105 | 6 |
|  | PIQ95941.1 | 3E-05 | Qa.III | IEEITHIINEINDISSTIASAVEEQTATVAEIGRNITEAA------------ | *Nitrospinae bacterium CG11_big_fil_rev_8_21_14_0_20_5 6_8* | Bacteria; Nitrospinae/Tectomicrobia group; Nitrospinae | hypothetical protein COV67_12235 | 6 |
|  | SDT41027.1 | 3E-05 | Qc.III | -KGIGSIIGEVNEVATAIAAAVQEQGAATQEISRSTQHAAQGTKNVS------ | *Bradyrhizobium canariense* | Bacteria; Proteobacteria; Alphaproteobacteria; Rhizobiales; Bradyrhizobiaceae; Bradyrhizobium | Methyl-accepting chemotaxis protein | 6 |
| | SFN86979.1 | 3E-05 | Qc.III | IQEIGGTIERLSEVSSAIAAAVEEQGAATQEISRNVQQAALGTQEVSSNI--- | *Bradyrhizobium sp. Rc3b* | Bacteria; Proteobacteria; Alphaproteobacteria; Rhizobiales; Bradyrhizobiaceae; Bradyrhizobium | methyl-accepting chemotaxis protein | 6 |
|  | SNS58246.1 | 3E-05 | Qa.IV | VQEIGRTIEEMDHISGTIAAAIEEQGAATQEISRNVLQAAD----------- | *Azospirillum sp. RU38E* | Bacteria; Proteobacteria; Alphaproteobacteria; Rhodospirillales; Rhodospirillaceae; Azospirillum | methyl-accepting chemotaxis sensory transducer with TarH sensor | 6 |
|  | WP_008129796.1 | 3E-05 | Qc.III | -QTIGGIIGEVNEVATAIAAAVQEQGAATQEITRSTQFAAQGTKNVS------ | *Bradyrhizobium sp. YR681* | Bacteria; Proteobacteria; Alphaproteobacteria; Rhizobiales; Bradyrhizobiaceae; Bradyrhizobium | methyl-accepting chemotaxis protein | 6 |
|  | WP_008964505.1 | 3E-05 | Qa.IV | IKEIGTTITQISDIAATIAATVEQQGAATREIARNVGEAA------------ | *Bradyrhizobium sp. STM 3809* | Bacteria; Proteobacteria; Alphaproteobacteria; Rhizobiales; Bradyrhizobiaceae; Bradyrhizobium | methyl-accepting chemotaxis protein | 6 |
|  | WP_018318753.1 | 3E-05 | Qc.III | -QTIGGIIGEVNEVATAIAAAVQEQGAATQEITRSTQFAAQGTKNVS------ | *Bradyrhizobium sp. WSM2793* | Bacteria; Proteobacteria; Alphaproteobacteria; Rhizobiales; Bradyrhizobiaceae; Bradyrhizobium | methyl-accepting chemotaxis protein | 6 |
|  | WP_018646635.1 | 3E-05 | Qc.III | -QTIGGIIGEVNEVATAIAAAVQEQGAATQEITRSTQFAAQGTKNVS------ | *Bradyrhizobium japonicum* | Bacteria; Proteobacteria; Alphaproteobacteria; Rhizobiales; Bradyrhizobiaceae; Bradyrhizobium | methyl-accepting chemotaxis protein | 6 |
|  | WP_024872950.1 | 3E-05 | Qa.III | LEEITRAVSSISDMANQIATAAEEQSHVTGEITTNVTAVK------------ | *Tolumonas lignilytica* | Bacteria; Proteobacteria; Gammaproteobacteria; Aeromonadales; Aeromonadaceae; Tolumonas | methyl-accepting chemotaxis protein | 6 |
|  | WP_027515466.1 | 3E-05 | Qc.III | -QTIGGIIGEVNEVATAIAAAVQEQGAATQEITRSTQFAAQGTKNVS------ | *Bradyrhizobium sp. WSM1417* | Bacteria; Proteobacteria; Alphaproteobacteria; Rhizobiales; Bradyrhizobiaceae; Bradyrhizobium | methyl-accepting chemotaxis protein | 6 |
| | WP_027533099.1 | 3E-05 | Qc.III | IKEISGTIGRISEISSAIASAVEEQGAATQEIARNIQQAAQGTDQVAANI--- | *Bradyrhizobium sp. WSM3983* | Bacteria; Proteobacteria; Alphaproteobacteria; Rhizobiales; Bradyrhizobiaceae; Bradyrhizobium | methyl-accepting chemotaxis protein | 6 |
|  | WP_027561173.1 | 3E-05 | Qc.III | -QTIGGIIGEVNEVATAIAAAVQEQGAATQEITRSTQFAAQGTKNVS------ | *Bradyrhizobium arachidis* | Bacteria; Proteobacteria; Alphaproteobacteria; Rhizobiales; Bradyrhizobiaceae; Bradyrhizobium | methyl-accepting chemotaxis protein | 6 |
|  | WP_027570226.1 | 3E-05 | Qc.III | -QTIGGIIGEVNEVATAIAAAVQEQGAATQEITRSTQFAAQGTKNVS------ | *Bradyrhizobium sp. URHA0013* | Bacteria; Proteobacteria; Alphaproteobacteria; Rhizobiales; Bradyrhizobiaceae; Bradyrhizobium | methyl-accepting chemotaxis protein | 6 |
| | WP_028141641.1 | 3E-05 | Qc.III | IQEIGGTIERLSEVSSAIAAAVEEQGAATQEISRNVQQAALGTQEVSSNI--- | *Bradyrhizobium japonicum* | Bacteria; Proteobacteria; Alphaproteobacteria; Rhizobiales; Bradyrhizobiaceae; Bradyrhizobium | methyl-accepting chemotaxis protein | 6 |
|  | WP_028146462.1 | 3E-05 | Qc.III | -QTIGGIIGEVNEVATAIAAAVQEQGAATQEITRSTQFAAQGTKNVS------ | *Bradyrhizobium genosp. SA-4* | Bacteria; Proteobacteria; Alphaproteobacteria; Rhizobiales; Bradyrhizobiaceae; Bradyrhizobium | methyl-accepting chemotaxis protein | 6 |
|  | WP_038971357.1 | 3E-05 | Qc.III | -QTIGGIIGEVNEVATAIAAAVQEQGAATQEITRSTQFAAQGTKNVS------ | *Bradyrhizobium sp. CCBAU 15635* | Bacteria; Proteobacteria; Alphaproteobacteria; Rhizobiales; Bradyrhizobiaceae; Bradyrhizobium | methyl-accepting chemotaxis protein | 6 |
|  | WP_049973701.1 | 3E-05 | Qa.IV | ------TIVQLSEVSSTIAAAVEEQNAATAEISRSISEAAQDVEEVNT----- | *Azospirillum sp. B4* | Bacteria; Proteobacteria; Alphaproteobacteria; Rhodospirillales; Rhodospirillaceae; Azospirillum | HAMP domain-containing protein | 6 |
|  | WP_056743335.1 | 3E-05 | Qb.I | LQRTKLHLQDTIEQSEVSSEELKASNEEMQAINEELRSATEELETSK------ | *Pseudomonas sp. Root329* | Bacteria; Proteobacteria; Gammaproteobacteria; Pseudomonadales; Pseudomonadaceae; Pseudomonas | PAS domain S-box protein | 6 |
| | WP_061029419.1 | 3E-05 | Qc.III | IQEIGGTIERLSEVSSAIAAAVEEQGAATQEISRNVQQAALGTQEVSSNI--- | *Bradyrhizobium sp. CCH5-F6* | Bacteria; Proteobacteria; Alphaproteobacteria; Rhizobiales; Bradyrhizobiaceae; Bradyrhizobium | HAMP domain-containing protein | 6 |
|  | WP_080134975.1 | 3E-05 | Qc.III | -QTIGGIIGEVNEVATAIAAAVQEQGAATQEITRSTQFAAQGTKNVS------ | *Bradyrhizobium sp. BR10280* | Bacteria; Proteobacteria; Alphaproteobacteria; Rhizobiales; Bradyrhizobiaceae; Bradyrhizobium | methyl-accepting chemotaxis protein | 6 |
|  | WP_084134834.1 | 3E-05 | Qc.III | -QQVAASATELATGANEISEDVSEMSESINEIEIKVSDYASSTKEMNER---- | *Paenibacillus harenae* | Bacteria; Firmicutes; Bacilli; Bacillales; Paenibacillaceae; Paenibacillus | methyl-accepting chemotaxis protein | 6 |
|  | WP_085351743.1 | 3E-05 | Qc.III | -QTIGGIIGEVNEVATAIAAAVQEQGAATQEITRSTQFAAQGTKNVS------ | *Bradyrhizobium canariense* | Bacteria; Proteobacteria; Alphaproteobacteria; Rhizobiales; Bradyrhizobiaceae; Bradyrhizobium | methyl-accepting chemotaxis protein | 6 |
| | WP_085967692.1 | 3E-05 | Qc.III | IQEIGGTIERLSEVSSAIAAAVEEQGAATQEISRNVQQAALGTQEVSSNI--- | *Bradyrhizobium* | Bacteria; Proteobacteria; Alphaproteobacteria; Rhizobiales; Bradyrhizobiaceae | MULTISPECIES: methyl-accepting chemotaxis protein | 6 |
| | WP_085968430.1 | 3E-05 | Qc.III | IQEIGGTIERLSEVSSAIAAAVEEQGAATQEISRNVQQAALGTQEVSSNI--- | *Bradyrhizobium sp. CCBAU 15635* | Bacteria; Proteobacteria; Alphaproteobacteria; Rhizobiales; Bradyrhizobiaceae; Bradyrhizobium | methyl-accepting chemotaxis protein | 6 |
| | WP_085971727.1 | 3E-05 | Qc.III | IQEIGGTIERLSEVSSAIAAAVEEQGAATQEISRNVQQAALGTQEVSSNI--- | *Bradyrhizobium sp. CCBAU 15544* | Bacteria; Proteobacteria; Alphaproteobacteria; Rhizobiales; Bradyrhizobiaceae; Bradyrhizobium | methyl-accepting chemotaxis protein | 6 |
|  | WP_089265427.1 | 3E-05 | Qc.III | -KSIGGIIGEVNEVATAIAAAVEEQGAATQEITRSTQQAADGTRNVS------ | *Tardiphaga sp. OK246* | Bacteria; Proteobacteria; Alphaproteobacteria; Rhizobiales; Bradyrhizobiaceae; Tardiphaga | methyl-accepting chemotaxis protein | 6 |
|  | WP_089288632.1 | 3E-05 | Qa.IV | VQEIGRTIEEMDHISGTIAAAIEEQGAATQEISRNVLQAAD----------- | *Azospirillum* | Bacteria; Proteobacteria; Alphaproteobacteria; Rhodospirillales; Rhodospirillaceae | MULTISPECIES: HAMP domain-containing protein | 6 |
|  | WP_089290597.1 | 3E-05 | Qa.III | --EITRTIGEINEIATTIAAAVEEQDAATREIARNVQQAAVGTQE-------- | *Azospirillum* | Bacteria; Proteobacteria; Alphaproteobacteria; Rhodospirillales; Rhodospirillaceae | MULTISPECIES: methyl-accepting chemotaxis protein | 6 |
| | WP_089799193.1 | 3E-05 | Qc.III | -EQLTKIASETVTYAEAVEAKVTDQNSLISETNQTADFLNQQVNQLNKLIDDL | *Halolactibacillus alkaliphilus* | Bacteria; Firmicutes; Bacilli; Bacillales; Bacillaceae; Halolactibacillus | methyl-accepting chemotaxis protein | 6 |
| | WP_091885095.1 | 3E-05 | Qc.III | IKEISGTIGRISEISSAIASAVEEQGAATQEIARNIQQAAQGTDQVAANI--- | *Bradyrhizobium sp. Rc2d* | Bacteria; Proteobacteria; Alphaproteobacteria; Rhizobiales; Bradyrhizobiaceae; Bradyrhizobium | methyl-accepting chemotaxis protein | 6 |
|  | WP_091955962.1 | 3E-05 | Qc.III | -QTIGGIIGEVNEVATAIAAAVQEQGAATQEITRSTQFAAQGTKNVS------ | *Bradyrhizobium shewense* | Bacteria; Proteobacteria; Alphaproteobacteria; Rhizobiales; Bradyrhizobiaceae; Bradyrhizobium | methyl-accepting chemotaxis protein | 6 |
|  | WP_092144944.1 | 3E-05 | Qc.III | -KSIGGIIGEVNEVATAIAAAVEEQGAATQEITRSTQQAADGTRNVS------ | *Bradyrhizobiaceae* | Bacteria; Proteobacteria; Alphaproteobacteria; Rhizobiales | methyl-accepting chemotaxis protein | 6 |
|  | WP_092216580.1 | 3E-05 | Qc.III | -QTIGGIIGEVNEVATAIAAAVQEQGAATQEITRSTQFAAQGTKNVS------ | *Bradyrhizobium arachidis* | Bacteria; Proteobacteria; Alphaproteobacteria; Rhizobiales; Bradyrhizobiaceae; Bradyrhizobium | methyl-accepting chemotaxis protein | 6 |
|  | WP_092254902.1 | 3E-05 | Qc.III | -QTIGGIIGEVNEVATAIAAAVQEQGAATQEITRSTQFAAQGTKNVS------ | *Bradyrhizobium sp. Rc3b* | Bacteria; Proteobacteria; Alphaproteobacteria; Rhizobiales; Bradyrhizobiaceae; Bradyrhizobium | methyl-accepting chemotaxis protein | 6 |
| | WP_092262450.1 | 3E-05 | Qc.III | IQEIGGTIERLSEVSSAIAAAVEEQGAATQEISRNVQQAALGTQEVSSNI--- | *Bradyrhizobium sp. Rc3b* | Bacteria; Proteobacteria; Alphaproteobacteria; Rhizobiales; Bradyrhizobiaceae; Bradyrhizobium | methyl-accepting chemotaxis protein | 6 |
|  | WP_093756821.1 | 3E-05 | Qc.III | -KSIGGIIGEVNEVATAIAAAVEEQGAATQEITRSTQQAADGTRNVS------ | *Tardiphaga sp. OK245* | Bacteria; Proteobacteria; Alphaproteobacteria; Rhizobiales; Bradyrhizobiaceae; Tardiphaga | methyl-accepting chemotaxis protein | 6 |
| | WP_100234696.1 | 3E-05 | Qc.III | IKEISGTIGRISEISSAIASAVEEQGAATQEIARNIQQAAQGTDQVAANI--- | *Bradyrhizobium sp. INPA54B* | Bacteria; Proteobacteria; Alphaproteobacteria; Rhizobiales; Bradyrhizobiaceae; Bradyrhizobium | methyl-accepting chemotaxis protein | 6 |
|  | WP_104462059.1 | 3E-05 | Qc.III | -QTIGGIIGEVNEVATAIAAAVQEQGAATQEITRSTQFAAQGTKNVS------ | *Bradyrhizobium sp. AC87j1* | Bacteria; Proteobacteria; Alphaproteobacteria; Rhizobiales; Bradyrhizobiaceae; Bradyrhizobium | methyl-accepting chemotaxis protein | 6 |
|  | WP_104512139.1 | 3E-05 | Qa.IV | IRDISTTIEKLSEISSTIASAVEEQGAATLEISRNVQQAAQ------------ | *Rhodopseudomonas palustris* | Bacteria; Proteobacteria; Alphaproteobacteria; Rhizobiales; Bradyrhizobiaceae; Rhodopseudomonas | methyl-accepting chemotaxis protein | 6 |
|  | WP_106941859.1 | 3E-05 | Qc.III | -QTIGGIIGEVNEVATAIAAAVQEQGAATQEITRSTQFAAQGTKNVS------ | *Bradyrhizobium sp. MOS002* | Bacteria; Proteobacteria; Alphaproteobacteria; Rhizobiales; Bradyrhizobiaceae; Bradyrhizobium | methyl-accepting chemotaxis protein | 6 |
|  | WP_106948049.1 | 3E-05 | Qc.III | -QTIGGIIGEVNEVATAIAAAVQEQGAATQEITRSTQFAAQGTKNVS------ | *Bradyrhizobium sp. MOS003* | Bacteria; Proteobacteria; Alphaproteobacteria; Rhizobiales; Bradyrhizobiaceae; Bradyrhizobium | methyl-accepting chemotaxis protein | 6 |
| | AFK55596.1 | 3.1E-05 | Qc.III | --QIRATIGRIDEISTAIAAAVEEQSAATSDISTNAQQAARGVDEVSRSV--- | *Tistrella mobilis KA081020-065* | Bacteria; Proteobacteria; Alphaproteobacteria; Rhodospirillales; Rhodospirillaceae; Tistrella | methyl-accepting chemotaxis protein (plasmid) | 6 |
|  | BAR55884.1 | 3.1E-05 | Qc.III | -QTIGGIIGEVNEVATAIAAAVQEQGAATQEITRSTQFAAQGTRNVS------ | *Bradyrhizobium diazoefficiens* | Bacteria; Proteobacteria; Alphaproteobacteria; Rhizobiales; Bradyrhizobiaceae; Bradyrhizobium | putative methyl accepting chemotaxis protein | 6 |
| | KJS40805.1 | 3.1E-05 | Qa.III | IKGITDVIEQIDEIATSIATAVEEQSAATREISRNVEEAARGTEEVSRNI--- | *Rhodospirillaceae bacterium BRH_c57* | Bacteria; Proteobacteria; Alphaproteobacteria; Rhodospirillales; Rhodospirillaceae | hypothetical protein VR70_05505 | 6 |
| | KKM09550.1 | 3.1E-05 | Qc.I | --QLQEMARQLKELYENLEEKVEQRTLALREANRELQAQREALHRANEEL--- | *Clostridiales bacterium PH28_bin88* | Bacteria; Firmicutes; Clostridia; Clostridiales | histidine kinase | 6 |

| | | | | | | | | |
|---|---|---|---|---|---|---|---|---|
|  | **PVE23224.1** | 3.1E-05 | Qa.IV | IEQIARTIAEMSQISISIATAMEEQGAATAEIARKVQEAARGT---------- | *Microvirga sp. KLBC 81* | Bacteria; Proteobacteria; Alphaproteobacteria; Rhizobiales; Methylobacteriaceae; Microvirga | hypothetical protein DC522_16955 | 6 |
|  | **SMX60916.1** | 3.1E-05 | Qc.III | -REVSGTIERLSHISATIAAAIEQQGAATQEISRNVQHTAQGTQRV------- | *Bradyrhizobium sp. ORS 285* | Bacteria; Proteobacteria; Alphaproteobacteria; Rhizobiales; Bradyrhizobiaceae; Bradyrhizobium | putative methyl-accepting chemotaxis protein | 6 |
|  | **WP_008960762.1** | 3.1E-05 | Qc.III | -REVSGTIERLSHISATIAAAIEQQGAATQEISRNVQHTAQGTQRV------- | *Bradyrhizobium sp. STM 3809* | Bacteria; Proteobacteria; Alphaproteobacteria; Rhizobiales; Bradyrhizobiaceae; Bradyrhizobium | methyl-accepting chemotaxis protein | 6 |
|  | **WP_011500763.1** | 3.1E-05 | Qa.IV | IKEIGQTIDRMSEIASTIASAVEEQGAATQEISRNVQQAAQ------------ | *Rhodopseudomonas palustris* | Bacteria; Proteobacteria; Alphaproteobacteria; Rhizobiales; Bradyrhizobiaceae; Rhodopseudomonas | methyl-accepting chemotaxis protein | 6 |
|  | **WP_034364595.1** | 3.1E-05 | Qa.III | MKEIMQSVGQVTGLLSEISGALHQQSEGI----AHVNTAVAHMDSTNQE---- | *Comamonas testosteroni* | Bacteria; Proteobacteria; Betaproteobacteria; Burkholderiales; Comamonadaceae; Comamonas | methyl-accepting chemotaxis protein | 6 |
|  | **WP_041757802.1** | 3.1E-05 | Qc.III | -REVSGTIERLSHISATIAAAIEQQGAATQEISRNVQHTAQGTQRV------- | *Bradyrhizobium sp. ORS 278* | Bacteria; Proteobacteria; Alphaproteobacteria; Rhizobiales; Bradyrhizobiaceae; Bradyrhizobium | HAMP domain-containing protein | 6 |
| | **WP_068435556.1** | 3.1E-05 | Qc.III | IESIGAIIERINQVASAIAAAVEEQDATTRDIARNVAEAAGGTREVSRHVTD- | *Magnetospirillum sp. XM-1* | Bacteria; Proteobacteria; Alphaproteobacteria; Rhodospirillales; Rhodospirillaceae; Magnetospirillum | HAMP domain-containing protein | 6 |
|  | **WP_082641264.1** | 3.1E-05 | Qa.IV | -KEIGTTIGRISDISTTIAATIEQQGAATAEIARNVSEAAK------------ | *Bradyrhizobium yuanmingense* | Bacteria; Proteobacteria; Alphaproteobacteria; Rhizobiales; Bradyrhizobiaceae; Bradyrhizobium | methyl-accepting chemotaxis protein | 6 |
|  | **WP_083235087.1** | 3.1E-05 | Qa.IV | -KEIGTTIGRISDISTTIAATIEQQGAATAEIARNVSEAAK------------ | *Bradyrhizobium yuanmingense* | Bacteria; Proteobacteria; Alphaproteobacteria; Rhizobiales; Bradyrhizobiaceae; Bradyrhizobium | chemotaxis protein | 6 |
|  | **WP_084165010.1** | 3.1E-05 | Qa.IV | IQNITETIKRMSEISSGIAAAVQEQAAATQEIARNVNHAA------------- | *Skermanella stibiiresistens* | Bacteria; Proteobacteria; Alphaproteobacteria; Rhodospirillales; Rhodospirillaceae; Skermanella | methyl-accepting chemotaxis protein | 6 |
|  | **WP_084302068.1** | 3.1E-05 | Qc.III | -QTIGGIIGEVNEVATAIAAAVQEQGAATQEITRSTQFAAQGTKNVS------ | *Bradyrhizobium sp. WSM2254* | Bacteria; Proteobacteria; Alphaproteobacteria; Rhizobiales; Bradyrhizobiaceae; Bradyrhizobium | methyl-accepting chemotaxis protein | 6 |
|  | **WP_092686354.1** | 3.1E-05 | Qa.IV | IKEIGQTIDRMSEIASTIASAVEEQGAATQEISRNVQQAAQ------------ | *Rhodopseudomonas pseudopalustris* | Bacteria; Proteobacteria; Alphaproteobacteria; Rhizobiales; Bradyrhizobiaceae; Rhodopseudomonas | methyl-accepting chemotaxis protein | 6 |
|  | **WP_094983928.1** | 3.1E-05 | Qb.I | --RGAQILKENLNNSEKISSDIKRSSKLIEDLNQQSGEISKIVT--------- | *Cellvibrio mixtus* | Bacteria; Proteobacteria; Gammaproteobacteria; Cellvibrionales; Cellvibrionaceae; Cellvibrio | methyl-accepting chemotaxis protein | 6 |
| | **WP_098738077.1** | 3.1E-05 | Qa.III.b | IKGIGGVIGEISEIAAAIAAAVEEQGTATQEISRNVQQAAVGTQEVSSHIVS- | *Azospirillum sp. B2* | Bacteria; Proteobacteria; Alphaproteobacteria; Rhodospirillales; Rhodospirillaceae; Azospirillum | methyl-accepting chemotaxis protein | 6 |
| | **WP_109049528.1** | 3.1E-05 | Qa.III.b | IKGIGGVIGEISEIAAAIAAAVEEQGTATQEISRNVQQAAVGTQEVSSHIVS- | *Azospirillum sp. TSA6c* | Bacteria; Proteobacteria; Alphaproteobacteria; Rhodospirillales; Rhodospirillaceae; Azospirillum | HAMP domain-containing protein | 6 |
|  | **WP_111355664.1** | 3.1E-05 | Qc.III | IKEIGGTIGRIAEISAAIAAAVEEQGAATHEIARNVQQAAAGT---------- | *Rhodoplanes elegans* | Bacteria; Proteobacteria; Alphaproteobacteria; Rhizobiales; Hyphomicrobiaceae; Rhodoplanes | methyl-accepting chemotaxis protein | 6 |
|  | **WP_111359642.1** | 3.1E-05 | Qc.III | IKEIGGTIGQIAEIASAIAAAVEEQEAATAEISRNIQHASSGTTEVSA----- | *Rhodoplanes elegans* | Bacteria; Proteobacteria; Alphaproteobacteria; Rhizobiales; Hyphomicrobiaceae; Rhodoplanes | HAMP domain-containing protein | 6 |
|  | **CAL79861.1** | 3.2E-05 | Qc.III | -REVSGTIERLSHISATIAAAIEQQGAATQEISRNVQHTAQGTQRV------- | *Bradyrhizobium sp. ORS 278* | Bacteria; Proteobacteria; Alphaproteobacteria; Rhizobiales; Bradyrhizobiaceae; Bradyrhizobium | Putative methyl-accepting chemotaxis protein | 6 |
| | **GAJ31143.1** | 3.2E-05 | Qc.III | --TIGGTIERLSEISSTIAAAVEEQGAATSEISRNVQSAAAGTTQVSANI--- | *Bradyrhizobium sp. DOA9* | Bacteria; Proteobacteria; Alphaproteobacteria; Rhizobiales; Bradyrhizobiaceae; Bradyrhizobium | chemoreceptor protein A | 6 |
|  | **KZD22397.1** | 3.2E-05 | Qc.III | -KSIGGIIGEVNEVATAIAAAVEEQGAATQEITRSTQQAADGTRNVS------ | *Tardiphaga robiniae* | Bacteria; Proteobacteria; Alphaproteobacteria; Rhizobiales; Bradyrhizobiaceae; Tardiphaga | chemotaxis protein | 6 |
|  | **PPY04797.1** | 3.2E-05 | R.IV | IDQISKAVSELDSTTQQNAALVQESSAAAASLEQQAGELSQLISRF------- | *Cronobacter sakazakii* | Bacteria; Proteobacteria; Gammaproteobacteria; Enterobacterales; Enterobacteriaceae; Cronobacter | methyl-accepting chemotaxis protein partial | 6 |
|  | **PVX49580.1** | 3.2E-05 | Qc.III | -KSIGGIIGEVNEVATAIAAAVEEQGAATQEITRSTQQAADGTRNVS------ | *Tardiphaga sp. OV697* | Bacteria; Proteobacteria; Alphaproteobacteria; Rhizobiales; Bradyrhizobiaceae; Tardiphaga | methyl-accepting chemotaxis protein | 6 |
|  | **PWT93794.1** | 3.2E-05 | Qc.III | IKEIGTTITRISEIALVIATAVEEQSATTGEIARNVDNAARSSTQV------- | *Proteobacteria bacterium* | Bacteria; Proteobacteria | methyl-accepting chemotaxis protein partial | 6 |
|  | **RCK19986.1** | 3.2E-05 | Qa.III | IDEIVRMISDISERASSVAAAVEQQTVATSEISQNVEQAASATREIS------ | *Thalassospira lucentensis MCCC 1A00383 = DSM 14000* | Bacteria; Proteobacteria; Alphaproteobacteria; Rhodospirillales; Rhodospirillaceae; Thalassospira | hypothetical protein TH1_20390 | 6 |
|  | **SFK70759.1** | 3.2E-05 | Qa.III.b | IRGIAGVIEQVDQIAATIAAAVEQQGAATREIARNVAETAVGTQEVS------ | *Roseomonas stagni DSM 19981* | Bacteria; Proteobacteria; Alphaproteobacteria; Rhodospirillales; Acetobacteraceae; Roseomonas | Methyl-accepting chemotaxis protein | 6 |
|  | **WP_003249938.1** | 3.2E-05 | Qc.III.c | LSVIMSVTEKINENSTNIANAVDQQASAIEQLNRAIESIAS------------ | *Bacillaceae* | Bacteria; Firmicutes; Bacilli; Bacillales | MULTISPECIES: methyl-accepting chemotaxis protein | 6 |
|  | **WP_013400616.1** | 3.2E-05 | Qc.III.c | LSVIMSVTEKINENSTNIANAVDQQASAIEQLNRAIESIAS------------ | *Bacillaceae* | Bacteria; Firmicutes; Bacilli; Bacillales | MULTISPECIES: methyl-accepting chemotaxis protein | 6 |
| | **WP_037492635.1** | 3.2E-05 | Qc.III | -SEIFETISEINEISTTIASAVEEQGSATSEISRNAQEASRSSEEVTRNIDT- | *Sneathiella glossodoripedis* | Bacteria; Proteobacteria; Alphaproteobacteria; Sneathiellales; Sneathiellaceae; Sneathiella | HAMP domain-containing protein | 6 |
|  | **WP_089265722.1** | 3.2E-05 | Qc.III | -KSIGGIIGEVNEVATAIAAAVEEQGAATQEITRSTQQAADGTRNVS------ | *Tardiphaga sp. OK246* | Bacteria; Proteobacteria; Alphaproteobacteria; Rhizobiales; Bradyrhizobiaceae; Tardiphaga | methyl-accepting chemotaxis protein | 6 |
|  | **WP_092146073.1** | 3.2E-05 | Qc.III | -KSIGGIIGEVNEVATAIAAAVEEQGAATQEITRSTQQAADGTRNVS------ | *Bradyrhizobiaceae* | Bacteria; Proteobacteria; Alphaproteobacteria; Rhizobiales | methyl-accepting chemotaxis protein | 6 |
|  | **WP_093757327.1** | 3.2E-05 | Qc.III | -KSIGGIIGEVNEVATAIAAAVEEQGAATQEITRSTQQAADGTRNVS------ | *Tardiphaga sp. OK245* | Bacteria; Proteobacteria; Alphaproteobacteria; Rhizobiales; Bradyrhizobiaceae; Tardiphaga | methyl-accepting chemotaxis protein | 6 |
|  | **WP_101299629.1** | 3.2E-05 | Qa.III | IDEIVRMISDISERASSVAAAVEQQTVATSEISQNVEQAASATREIS------ | *Thalassospira lohafexi* | Bacteria; Proteobacteria; Alphaproteobacteria; Rhodospirillales; Rhodospirillaceae; Thalassospira | HAMP domain-containing protein | 6 |
|  | **WP_102783835.1** | 3.2E-05 | Qa.III | IDEIVRMISDISERASSVAAAVEQQTVATSEISQNVEQAASATREIS------ | *Thalassospira sp. GB04J01* | Bacteria; Proteobacteria; Alphaproteobacteria; Rhodospirillales; Rhodospirillaceae; Thalassospira | HAMP domain-containing protein | 6 |
| | **WP_107343793.1** | 3.2E-05 | Qc.III | -REIGGTIERMSEIASTIASAVEEQGAATQEISRNVQQAARGTHEVSSNISD- | *Rhodopseudomonas palustris* | Bacteria; Proteobacteria; Alphaproteobacteria; Rhizobiales; Bradyrhizobiaceae; Rhodopseudomonas | methyl-accepting chemotaxis protein | 6 |
| | **EJW12630.1** | 3.3E-05 | Qc.III | IKEIGGTIGQIAEISAAIAAAVEEQGAATHEIARNVQQAAKGTDEVARHITD- | *Rhodovulum sp. PH10* | Bacteria; Proteobacteria; Alphaproteobacteria; Rhodobacterales; Rhodobacteraceae; Rhodovulum | methyl-accepting chemotaxis sensory transducer | 6 |
|  | **OCR86518.1** | 3.3E-05 | Qa.IV | --EIEANVNILSQGINEMSQSINEQTEAINQINEAVATVDEQTKQ-NV----- | *Campylobacter fetus subsp. testudinum* | Bacteria; Proteobacteria; Epsilonproteobacteria; Campylobacterales; Campylobacteraceae; Campylobacter | chemotaxis protein partial | 6 |
| | **OYU89305.1** | 3.3E-05 | Qc.III | IKDIGDTIGRMSEIASTIASAVEEQGAATQEISRNVQQAAVGTTQVSSNIT-- | *Bradyrhizobiaceae bacterium PARB1* | Bacteria; Proteobacteria; Alphaproteobacteria; Rhizobiales; Bradyrhizobiaceae | methyl-accepting chemotaxis protein | 6 |
|  | **WP_009686883.1** | 3.3E-05 | Qb.I | LQRTRLQLQETIEQSQVSSEELTASNEEMQAINEELRSASEELETSK------ | *Pseudomonas* | Bacteria; Proteobacteria; Gammaproteobacteria; Pseudomonadales; Pseudomonadaceae | MULTISPECIES: response regulator | 6 |
| | **WP_024508706.1** | 3.3E-05 | Qc.III | IKEISGTIEKLSEISSTIAAAVEEQGAATQEIARNVQQAAQGTQQVSSNIAD- | *Bradyrhizobium sp. ARR65* | Bacteria; Proteobacteria; Alphaproteobacteria; Rhizobiales; Bradyrhizobiaceae; Bradyrhizobium | methyl-accepting chemotaxis protein | 6 |
|  | **WP_029886816.1** | 3.3E-05 | Qb.I | LQRTRLQLQETIEQSQVSSEELTASNEEMQAINEELRSASEELETSK------ | *Pseudomonas putida* | Bacteria; Proteobacteria; Gammaproteobacteria; Pseudomonadales; Pseudomonadaceae; Pseudomonas | response regulator | 6 |
|  | **WP_044016336.1** | 3.3E-05 | SNAP.c | LNEITGNIESVKDSVMVQHTEVTETAAKIDSIAQTMSSLDFHIDNQAK----- | *Treponema sp. OMZ 838* | Bacteria; Spirochaetes; Spirochaetales; Spirochaetaceae; Treponema | methyl-accepting chemotaxis protein | 6 |
| | **WP_052026860.1** | 3.3E-05 | Qc.III | IKEIGGTIGQIAEISAAIAAAVEEQGAATHEIARNVQQAAKGTDEVARHITD- | *Rhodovulum sp. PH10* | Bacteria; Proteobacteria; Alphaproteobacteria; Rhodobacterales; Rhodobacteraceae; Rhodovulum | methyl-accepting chemotaxis protein | 6 |
|  | **WP_054881611.1** | 3.3E-05 | Qb.I | LQRTRLQLQETIEQSQVSSEELTASNEEMQAINEELRSASEELETSK------ | *Pseudomonas sp. NBRC 111128* | Bacteria; Proteobacteria; Gammaproteobacteria; Pseudomonadales; Pseudomonadaceae; Pseudomonas | response regulator | 6 |
|  | **WP_054889890.1** | 3.3E-05 | Qb.I | LQRTRLQLQETIEQSQVSSEELTASNEEMQAINEELRSASEELETSK------ | *Pseudomonas sp. NBRC 111129* | Bacteria; Proteobacteria; Gammaproteobacteria; Pseudomonadales; Pseudomonadaceae; Pseudomonas | response regulator | 6 |
|  | **WP_054904689.1** | 3.3E-05 | Qb.I | LQRTRLQLQETIEQSQVSSEELTASNEEMQAINEELRSASEELETSK------ | *Pseudomonas sp. NBRC 111144* | Bacteria; Proteobacteria; Gammaproteobacteria; Pseudomonadales; Pseudomonadaceae; Pseudomonas | response regulator | 6 |
|  | **WP_054913139.1** | 3.3E-05 | Qb.I | LQRTRLQLQETIEQSQVSSEELTASNEEMQAINEELRSASEELETSK------ | *Pseudomonas sp. NBRC 111127* | Bacteria; Proteobacteria; Gammaproteobacteria; Pseudomonadales; Pseudomonadaceae; Pseudomonas | response regulator | 6 |
| | **WP_061029421.1** | 3.3E-05 | Qc.III | IKDISHTIEKLSEISSTIAAAVEEQGAATQEISRNVQQAAAGTHQVSSNI--- | *Bradyrhizobium sp. CCH5-F6* | Bacteria; Proteobacteria; Alphaproteobacteria; Rhizobiales; Bradyrhizobiaceae; Bradyrhizobium | methyl-accepting chemotaxis protein | 6 |
|  | **WP_064314799.1** | 3.3E-05 | Qb.I | LQRTRLQLQETIEQSQVSSEELTASNEEMQAINEELRSASEELETSK------ | *Pseudomonas* | Bacteria; Proteobacteria; Gammaproteobacteria; Pseudomonadales; Pseudomonadaceae | MULTISPECIES: response regulator | 6 |
|  | **WP_085552860.1** | 3.3E-05 | Qa.IV | -----ATIARLNDIATGIAAAVEEQGAATSEIARNVQQAAGGTAQASEN---- | *Azospirillum lipoferum* | Bacteria; Proteobacteria; Alphaproteobacteria; Rhodospirillales; Rhodospirillaceae; Azospirillum | HAMP domain-containing protein | 6 |
|  | **WP_088871323.1** | 3.3E-05 | Qa.IV | IQDITETITTMNATTQSIAAAVGAQGEATREIAENVAQAAQ------------ | *Nitrospirillum amazonense* | Bacteria; Proteobacteria; Alphaproteobacteria; Rhodospirillales; Rhodospirillaceae; Nitrospirillum | chemotaxis protein | 6 |
|  | **WP_089126589.1** | 3.3E-05 | Qc.I | -----QMAGQLKESFETLEQRVSDRTAALAESNHQLEHAKEKAEVAN------ | *Tolypothrix sp. NIES-4075* | Bacteria; Cyanobacteria; Nostocales; Tolypothrichaceae; Tolypothrix | hybrid sensor histidine kinase/response regulator | 6 |
| | **WP_092619023.1** | 3.3E-05 | Qa.III | IQGIVKVIGEINEVTTAVSAAVEEQTAATGEISHNAQQAAQSTQQAAQSIQ-- | *Roseospirillum parvum* | Bacteria; Proteobacteria; Alphaproteobacteria; Rhizobiales; Rhodobiaceae; Roseospirillum | HAMP domain-containing protein | 6 |
|  | **WP_092960892.1** | 3.3E-05 | Qa.III.b | IRGIAGVIEQVDQIAATIAAAVEQQGAATREIARNVAETAVGTQEVS------ | *Roseomonas stagni* | Bacteria; Proteobacteria; Alphaproteobacteria; Rhodospirillales; Acetobacteraceae; Roseomonas | methyl-accepting chemotaxis protein | 6 |
|  | **WP_109087202.1** | 3.3E-05 | Qa.III | IVGIGRTIQSINQIAAGISAGIEQQGSAVREIVVNMEQAA------------- | *Azospirillum sp. TSH100* | Bacteria; Proteobacteria; Alphaproteobacteria; Rhodospirillales; Rhodospirillaceae; Azospirillum | chemotaxis protein | 6 |
|  | **WP_110676350.1** | 3.3E-05 | Qb.I | LQRTRLQLQETIEQSQVSSEELTASNEEMQAINEELRSASEELETSK------ | *Pseudomonas sp. MB-090624* | Bacteria; Proteobacteria; Gammaproteobacteria; Pseudomonadales; Pseudomonadaceae; Pseudomonas | response regulator | 6 |
| | **ACF00548.1** | 3.4E-05 | Qc.III | -REIGGTIERMSEIASTIASAVEEQGAATQEISRNVQQAARGTHEVSSNIAD- | *Rhodopseudomonas palustris TIE-1* | Bacteria; Proteobacteria; Alphaproteobacteria; Rhizobiales; Bradyrhizobiaceae; Rhodopseudomonas | methyl-accepting chemotaxis sensory transducer | 6 |
|  | **ADU12355.1** | 3.4E-05 | Qa.III | IRGIGATIKHMNEIASSIADSVHQQGDATNEIARNVNEAS------------- | *Asticcacaulis excentricus CB 48* | Bacteria; Proteobacteria; Alphaproteobacteria; Caulobacterales; Caulobacteraceae; Asticcacaulis | methyl-accepting chemotaxis sensory transducer | 6 |
|  | **BAL13616.1** | 3.4E-05 | Qc.III | -QTIGGIIGEVNEVATAIAAAVQEQGAATQEITRSTQFAAQGTRNVS------ | *Bradyrhizobium japonicum USDA 6* | Bacteria; Proteobacteria; Alphaproteobacteria; Rhizobiales; Bradyrhizobiaceae; Bradyrhizobium | methyl accepting chemotaxis protein | 6 |

| | | | | | | | | |
|---|---|---|---|---|---|---|---|---|
|  | **BBF81565.1** | 3.4E-05 | Qa.III | IRGIGATIKHMNEIASSIADSVHQQGDATNEIARNVNEAS------------ | *Asticcacaulis excentricus* | Bacteria; Proteobacteria; Alphaproteobacteria; Caulobacterales; Caulobacteraceae; Asticcacaulis | methyl-accepting chemotaxis protein | 6 |
|  | **OGP49776.1** | 3.4E-05 | Qb.I | ---TKENLQATIEEFQAANEELKSTNEELQSTNEELQSTNEELETSK------ | *Deltaproteobacteria bacterium RBG_13_43_22* | Bacteria; Proteobacteria; Deltaproteobacteria | SAM-dependent methyltransferase | 6 |
| | **OHC76848.1** | 3.4E-05 | Qc.III | -KGIVKTISDIDEIAAAISAAVEEQGAATQEIARNVEQAAAGTNEVSSNIA-- | *Rhodospirillales bacterium RIFCSPLOWO2_12_FULL_58_28* | Bacteria; Proteobacteria; Alphaproteobacteria; Rhodospirillales; unclassified Rhodospirillales | hypothetical protein A3G18_03600 | 6 |
|  | **SMH60225.1** | 3.4E-05 | Qa.III | -------IETVNNVASSISAAIEEQGAATRNISANVNEAAQGVAQVSSSL--- | *Azospirillum lipoferum* | Bacteria; Proteobacteria; Alphaproteobacteria; Rhodospirillales; Rhodospirillaceae; Azospirillum | methyl-accepting chemotaxis sensory transducer with Cache sensor | 6 |
| | **WP_018458835.1** | 3.4E-05 | Qc.III | IKDISGTIEKLSEISSTIAAAVEEQGAATQEISRNVQQAAEGTHQVSANIT-- | *Bradyrhizobium sp. WSM4349* | Bacteria; Proteobacteria; Alphaproteobacteria; Rhizobiales; Bradyrhizobiaceae; Bradyrhizobium | HAMP domain-containing protein | 6 |
|  | **WP_025565284.1** | 3.4E-05 | Qa.III | -----MSITSINDMSTQIATATTEQSSVTEEITRNVY----EIQNIVNSL--- | *Psychromonas sp. SP041* | Bacteria; Proteobacteria; Gammaproteobacteria; Alteromonadales; Psychromonadaceae; Psychromonas | methyl-accepting chemotaxis protein | 6 |
|  | **WP_028694172.1** | 3.4E-05 | Qb.I | LQRTRLQLQETIEQSEISSEELTASNEEMQAINEELRSASEELETSK------ | *Pseudomonas cremoricolorata* | Bacteria; Proteobacteria; Gammaproteobacteria; Pseudomonadales; Pseudomonadaceae; Pseudomonas | response regulator | 6 |
|  | **WP_043213340.1** | 3.4E-05 | Qb.I | LQRTRLQLQETIEQSEISSEELTASNEEMQAINEELRSASEELETSK------ | *Pseudomonas putida* | Bacteria; Proteobacteria; Gammaproteobacteria; Pseudomonadales; Pseudomonadaceae; Pseudomonas | response regulator | 6 |
| | **WP_051609585.1** | 3.4E-05 | Qc.III | ---IYRTIEEINEVSAAIAAAVEEQGAATNDISDKVGVVATQSSKVSDRI--- | *Terasakiella pusilla* | Bacteria; Proteobacteria; Alphaproteobacteria; Rhizobiales; Methylocystaceae; Terasakiella | hypothetical protein | 6 |
|  | **WP_060477383.1** | 3.4E-05 | Qb.I | LLRTRLQLQETIEQSEVSSEELTASNEEMQAINEELRSASEELETSK------ | *Pseudomonas monteilii* | Bacteria; Proteobacteria; Gammaproteobacteria; Pseudomonadales; Pseudomonadaceae; Pseudomonas | response regulator | 6 |
| | **WP_063993455.1** | 3.4E-05 | Qc.III | IKDISGTIEKLSEISSTIAAAVEEQGAATQEISRNVQQAAEGTHQVSANIT-- | *Bradyrhizobium sp.* | Bacteria; Proteobacteria; Alphaproteobacteria; Rhizobiales; Bradyrhizobiaceae; Bradyrhizobium | methyl-accepting chemotaxis protein | 6 |
|  | **WP_070090917.1** | 3.4E-05 | Qb.I | LQRTRLQLQETIEQSEISSEELTASNEEMQAINEELRSASEELETSK------ | *Pseudomonas monteilii* | Bacteria; Proteobacteria; Gammaproteobacteria; Pseudomonadales; Pseudomonadaceae; Pseudomonas | response regulator | 6 |
|  | **WP_070573821.1** | 3.4E-05 | Qb.I | LQRTRLQLQETIEQSEISSEELTASNEEMQAINEELRSASEELETSK------ | *Pseudomonas sp. HMSC08G10* | Bacteria; Proteobacteria; Gammaproteobacteria; Pseudomonadales; Pseudomonadaceae; Pseudomonas | response regulator | 6 |
| | **WP_079544751.1** | 3.4E-05 | Qc.III | IKEISGTIEKLSEISSTIAAAVEEQGAATQEISRNVQQAAQGTMQVSSNIT-- | *Bradyrhizobium lablabi* | Bacteria; Proteobacteria; Alphaproteobacteria; Rhizobiales; Bradyrhizobiaceae; Bradyrhizobium | methyl-accepting chemotaxis protein | 6 |
|  | **WP_082125632.1** | 3.4E-05 | Qa.III.b | -----EVINQVNDISNTIASAVEEQTATTNEIGRNVTEAARGTSEIAQ----- | *Silvibacterium bohemicum* | Bacteria; Acidobacteria; Acidobacteriales; Acidobacteriaceae; Silvibacterium | methyl-accepting chemotaxis protein | 6 |
|  | **WP_084857798.1** | 3.4E-05 | Qb.I | LQRTRLQLQETIEQSEISSEELTASNEEMQAINEELRSASEELETSK------ | *Pseudomonas putida* | Bacteria; Proteobacteria; Gammaproteobacteria; Pseudomonadales; Pseudomonadaceae; Pseudomonas | response regulator | 6 |
| | **WP_085352601.1** | 3.4E-05 | Qc.III | IKDISGTIEKLSEISSTIAAAVEEQGAATQEISRNVQQAAEGTHQVSANIT-- | *Bradyrhizobium canariense* | Bacteria; Proteobacteria; Alphaproteobacteria; Rhizobiales; Bradyrhizobiaceae; Bradyrhizobium | methyl-accepting chemotaxis protein | 6 |
| | **WP_085361344.1** | 3.4E-05 | Qc.III | IKDISGTIEKLSEISSTIAAAVEEQGAATQEISRNVQQAAEGTHQVSANIT-- | *Bradyrhizobium canariense* | Bacteria; Proteobacteria; Alphaproteobacteria; Rhizobiales; Bradyrhizobiaceae; Bradyrhizobium | methyl-accepting chemotaxis protein | 6 |
|  | **WP_085556392.1** | 3.4E-05 | Qa.IV | IQSIAATIGEVNDIVGTIATAVDGQNAATMEIARNVQQAA------------ | *Azospirillum lipoferum* | Bacteria; Proteobacteria; Alphaproteobacteria; Rhodospirillales; Rhodospirillaceae; Azospirillum | methyl-accepting chemotaxis protein | 6 |
| | **WP_085939971.1** | 3.4E-05 | Qc.III | --AISGTIENINSIATTIASAVEEQGAATQEIARNVQQAAAGADSVTQNI--- | *Azospirillum sp. B4* | Bacteria; Proteobacteria; Alphaproteobacteria; Rhodospirillales; Rhodospirillaceae; Azospirillum | methyl-accepting chemotaxis protein | 6 |
|  | **WP_090075755.1** | 3.4E-05 | Qc.III | --RISETVGEISTITESITSAVEEQEQSTQKIAGSIQKASSEVDAARKN---- | *Cohaesibacter marisflavi* | Bacteria; Proteobacteria; Alphaproteobacteria; Rhizobiales; Cohaesibacteraceae | HAMP domain-containing protein | 6 |
|  | **WP_090342667.1** | 3.4E-05 | Qb.I | LQRTRLQLQETIEQSEISSEELTASNEEMQAINEELRSASEELETSK------ | *Pseudomonas guariconensis* | Bacteria; Proteobacteria; Gammaproteobacteria; Pseudomonadales; Pseudomonadaceae; Pseudomonas | response regulator | 6 |
| | **WP_092616636.1** | 3.4E-05 | Qc.III | -KSIGEVVRQVSEISQSIASAVEEQNAATGEIARNVEQASAGTGEVTGNIH-- | *Roseospirillum parvum* | Bacteria; Proteobacteria; Alphaproteobacteria; Rhizobiales; Rhodobiaceae; Roseospirillum | methyl-accepting chemotaxis protein | 6 |
|  | **WP_099593237.1** | 3.4E-05 | Qb.I | LQRTRLQLQETIEQSEISSEELTASNEEMQAINEELRSASEELETSK------ | *Pseudomonas putida* | Bacteria; Proteobacteria; Gammaproteobacteria; Pseudomonadales; Pseudomonadaceae; Pseudomonas | response regulator | 6 |
|  | **WP_102081700.1** | 3.4E-05 | Qb.I | LQRTRLQLQETIEQSEISSEELTASNEEMQAINEELRSASEELETSK------ | *Pseudomonas guariconensis* | Bacteria; Proteobacteria; Gammaproteobacteria; Pseudomonadales; Pseudomonadaceae; Pseudomonas | response regulator | 6 |
| | **WP_104512245.1** | 3.4E-05 | Qc.III | -REIGGTIERMSEIASTIASAVEEQGAATQEISRNVQQAARGTHEVSSNIAD- | *Rhodopseudomonas palustris* | Bacteria; Proteobacteria; Alphaproteobacteria; Rhizobiales; Bradyrhizobiaceae; Rhodopseudomonas | methyl-accepting chemotaxis protein | 6 |
| | **WP_107354980.1** | 3.4E-05 | Qc.III | -REIGGTIERMSEIASTIASAVEEQGAATQEISRNVQQAARGTHEVSSNIAD- | *Rhodopseudomonas palustris* | Bacteria; Proteobacteria; Alphaproteobacteria; Rhizobiales; Bradyrhizobiaceae; Rhodopseudomonas | methyl-accepting chemotaxis protein | 6 |
|  | **WP_109120760.1** | 3.4E-05 | Qa.III | IREIARTVGEVHETSSSIAAAIHEQHAAIGEISRNVAQAAH----------- | *Azospirillum sp. TSO22-1* | Bacteria; Proteobacteria; Alphaproteobacteria; Rhodospirillales; Rhodospirillaceae; Azospirillum | HAMP domain-containing protein | 6 |
| | **WP_110784437.1** | 3.4E-05 | Qc.III | -REIGGTIERMSEIASTIASAVEEQGAATQEISRNVQQAARGTHEVSSNIAD- | *Rhodopseudomonas palustris* | Bacteria; Proteobacteria; Alphaproteobacteria; Rhizobiales; Bradyrhizobiaceae; Rhodopseudomonas | methyl-accepting chemotaxis protein | 6 |
|  | **ALJ34176.1** | 3.5E-05 | Qa.III | IKTIGGTIGEINGIATTIAAAVEEQGAATGEIARNVQQAA------------ | *Azospirillum brasilense* | Bacteria; Proteobacteria; Alphaproteobacteria; Rhodospirillales; Rhodospirillaceae; Azospirillum | hypothetical protein AMK58_01380 | 6 |
|  | **AND87131.1** | 3.5E-05 | Qc.III | -QTIGGIIGEVNEVATAIAAAVQEQGAATQEITRSTQFAAQGTRNVS------ | *Bradyrhizobium diazoefficiens USDA 110* | Bacteria; Proteobacteria; Alphaproteobacteria; Rhizobiales; Bradyrhizobiaceae; Bradyrhizobium | chemotaxis protein | 6 |
| | **EKV27726.1** | 3.5E-05 | Qc.III | IDSISETIKELKEISSNIASAVEQQGAATHEIARNVEQAAVGTQEVTSNI--- | *Caenispirillum salinarum AK4* | Bacteria; Proteobacteria; Alphaproteobacteria; Rhodospirillales; Rhodospirillaceae; Caenispirillum | Methyl-accepting chemotaxis protein | 6 |
|  | **KGJ65937.1** | 3.5E-05 | Qc.III | -QTIGGIIGEVNEVATAIAAAVQEQGAATQEITRSTQFAAQGTRNVS------ | *Bradyrhizobium diazoefficiens SEMIA 5080* | Bacteria; Proteobacteria; Alphaproteobacteria; Rhizobiales; Bradyrhizobiaceae; Bradyrhizobium | putative chemoreceptor mcpA | 6 |
| | **KJS34338.1** | 3.5E-05 | Qc.III | -SRIGSVIGELEGIATSIAGAVEEQTAATSEIARNIEQAASGTQEVTSHIGD- | *Rhodospirillaceae bacterium BRH_c57* | Bacteria; Proteobacteria; Alphaproteobacteria; Rhodospirillales; Rhodospirillaceae | chemotaxis protein partial | 6 |
|  | **OCS05850.1** | 3.5E-05 | Qa.IV | --EIEANVNILSQGINEMSQSINEQTEAINQINEAVATVDEQTKQ-NV----- | *Campylobacter fetus subsp. testudinum* | Bacteria; Proteobacteria; Epsilonproteobacteria; Campylobacterales; Campylobacteraceae; Campylobacter | chemotaxis protein partial | 6 |
|  | **OJW63196.1** | 3.5E-05 | Qc.III | -REISGTIGRISEISSAIASAVEEQGAATQEISRNIMQSAEGTHQV------- | *Afipia sp. 64-13* | Bacteria; Proteobacteria; Alphaproteobacteria; Rhizobiales; Bradyrhizobiaceae; Afipia | methyl-accepting chemotaxis protein | 6 |
|  | **OYU90805.1** | 3.5E-05 | Qa.IV | ---ITSTIEEVSAIATTIGSAIEEQGAATAEIARNVTQTAQATQD-------- | *Bradyrhizobiaceae bacterium PARB1* | Bacteria; Proteobacteria; Alphaproteobacteria; Rhizobiales; Bradyrhizobiaceae | chemotaxis protein | 6 |
|  | **SNY45890.1** | 3.5E-05 | Qa.III | --AIVQEVADIESAMGTLSGIADDQHEIVERLNATVDATMASI---------- | *Actinoplanes atraurantiacus* | Bacteria; Actinobacteria; Micromonosporales; Micromonosporaceae; Actinoplanes | methyl-accepting chemotaxis protein | 6 |
|  | **WP_002716533.1** | 3.5E-05 | Qa.IV | IKEISATIDQISEISSAIAAAVEEQGAATQEIARNVRQAAE----------- | *Afipia felis* | Bacteria; Proteobacteria; Alphaproteobacteria; Rhizobiales; Bradyrhizobiaceae; Afipia | methyl-accepting chemotaxis protein | 6 |
|  | **WP_003181783.1** | 3.5E-05 | Qb.I | LQRTKLQLQDTIEQAEVSSEELKASNEEMQAINEELRSATEELETS------- | *Pseudomonas fluorescens* | Bacteria; Proteobacteria; Gammaproteobacteria; Pseudomonadales; Pseudomonadaceae; Pseudomonas | PAS domain S-box protein | 6 |
|  | **WP_009340445.1** | 3.5E-05 | Qa.IV | IKEISATIDQISEISSAIAAAVEEQGAATQEIARNVRQAAE----------- | *Afipia* | Bacteria; Proteobacteria; Alphaproteobacteria; Rhizobiales; Bradyrhizobiaceae | MULTISPECIES: methyl-accepting chemotaxis protein | 6 |
| | **WP_019645340.1** | 3.5E-05 | Qc.III | LGEISSTIAQMNEISTTIASAVEEQGAATQEIARNAEQAATGTEQVTNSV--- | *Novispirillum itersonii* | Bacteria; Proteobacteria; Alphaproteobacteria; Rhodospirillales; Rhodospirillaceae; Novispirillum | methyl-accepting chemotaxis protein | 6 |
|  | **WP_024574841.1** | 3.5E-05 | Qc.III | --EISGTISSISQIAGAIFEAVDQQNAATREVTSGIQVAATGS---------- | *Afipia* | Bacteria; Proteobacteria; Alphaproteobacteria; Rhizobiales; Bradyrhizobiaceae | MULTISPECIES: chemotaxis protein | 6 |
|  | **WP_027536589.1** | 3.5E-05 | Qa.III | IQGITATIEEVSTIATSIGSAIEEQGAATAEIARNVTQTAEATK--------- | *Bradyrhizobium sp. URHA0002* | Bacteria; Proteobacteria; Alphaproteobacteria; Rhizobiales; Bradyrhizobiaceae; Bradyrhizobium | methyl-accepting chemotaxis protein | 6 |
|  | **WP_028148393.1** | 3.5E-05 | Qc.III | -QTIGGIIGEVNEVATAIAAAVQEQGAATQEITRSTQFAAQGTRNVS------ | *Bradyrhizobium japonicum* | Bacteria; Proteobacteria; Alphaproteobacteria; Rhizobiales; Bradyrhizobiaceae; Bradyrhizobium | methyl-accepting chemotaxis protein | 6 |
|  | **WP_028157414.1** | 3.5E-05 | Qc.III | -QTIGGIIGEVNEVATAIAAAVQEQGAATQEITRSTQFAAQGTRNVS------ | *Bradyrhizobium japonicum* | Bacteria; Proteobacteria; Alphaproteobacteria; Rhizobiales; Bradyrhizobiaceae; Bradyrhizobium | methyl-accepting chemotaxis protein | 6 |
|  | **WP_028170316.1** | 3.5E-05 | Qc.III | -QTIGGIIGEVNEVATAIAAAVQEQGAATQEITRSTQFAAQGTRNVS------ | *Bradyrhizobium japonicum* | Bacteria; Proteobacteria; Alphaproteobacteria; Rhizobiales; Bradyrhizobiaceae; Bradyrhizobium | methyl-accepting chemotaxis protein | 6 |
|  | **WP_038959390.1** | 3.5E-05 | Qc.III | -QTIGGIIGEVNEVATAIAAAVQEQGAATQEITRSTQFAAQGTRNVS------ | *Bradyrhizobium japonicum* | Bacteria; Proteobacteria; Alphaproteobacteria; Rhizobiales; Bradyrhizobiaceae; Bradyrhizobium | methyl-accepting chemotaxis protein | 6 |
|  | **WP_039156242.1** | 3.5E-05 | Qc.III | -QTIGGIIGEVNEVATAIAAAVQEQGAATQEITRSTQFAAQGTRNVS------ | *Bradyrhizobium japonicum* | Bacteria; Proteobacteria; Alphaproteobacteria; Rhizobiales; Bradyrhizobiaceae; Bradyrhizobium | methyl-accepting chemotaxis protein | 6 |
|  | **WP_041956209.1** | 3.5E-05 | Qc.III | -QTIGGIIGEVNEVATAIAAAVQEQGAATQEITRSTQFAAQGTRNVS------ | *Bradyrhizobium japonicum* | Bacteria; Proteobacteria; Alphaproteobacteria; Rhizobiales; Bradyrhizobiaceae; Bradyrhizobium | methyl-accepting chemotaxis protein | 6 |
|  | **WP_063980987.1** | 3.5E-05 | Qc.III | -QTIGGIIGEVNEVATAIAAAVQEQGAATQEITRSTQFAAQGTRNVS------ | *Bradyrhizobium sp.* | Bacteria; Proteobacteria; Alphaproteobacteria; Rhizobiales; Bradyrhizobiaceae; Bradyrhizobium | methyl-accepting chemotaxis protein | 6 |
|  | **WP_063994689.1** | 3.5E-05 | Qc.III | -QTIGGIIGEVNEVATAIAAAVQEQGAATQEITRSTQFAAQGTRNVS------ | *Bradyrhizobium sp.* | Bacteria; Proteobacteria; Alphaproteobacteria; Rhizobiales; Bradyrhizobiaceae; Bradyrhizobium | methyl-accepting chemotaxis protein | 6 |
|  | **WP_069586621.1** | 3.5E-05 | Qa.III | ---INEQIVQISDMNTQVATATEEQSTVVNDINQNIEEINHSSQQTA------ | *Salinivibrio sp. DV* | Bacteria; Proteobacteria; Gammaproteobacteria; Vibrionales; Vibrionaceae; Salinivibrio | methyl-accepting chemotaxis protein | 6 |
|  | **WP_071916684.1** | 3.5E-05 | Qc.III | -QTIGGIIGEVNEVATAIAAAVQEQGAATQEITRSTQFAAQGTRNVS------ | *Bradyrhizobium japonicum* | Bacteria; Proteobacteria; Alphaproteobacteria; Rhizobiales; Bradyrhizobiaceae; Bradyrhizobium | methyl-accepting chemotaxis protein | 6 |
|  | **WP_074829078.1** | 3.5E-05 | Qa.III | IQGITATIEEVSTIATSIGSAIEEQGAATAEIARNVTQTAEATK--------- | *Bradyrhizobium* | Bacteria; Proteobacteria; Alphaproteobacteria; Rhizobiales; Bradyrhizobiaceae | MULTISPECIES: methyl-accepting chemotaxis protein | 6 |
| | **WP_076960122.1** | 3.5E-05 | Qc.III | IQGIGGTIGEISNIAGSIATAVDEQGQATQEIARSVQKTAVSTREVTQNI--- | *Roseomonas deserti* | Bacteria; Proteobacteria; Alphaproteobacteria; Rhodospirillales; Acetobacteraceae; Roseomonas | methyl-accepting chemotaxis protein | 6 |
|  | **WP_077478959.1** | 3.5E-05 | Qa.III | ---INEQIVQISDMNTQVATATEEQSTVVNDINQNIEEINHSSQQTA------ | *Salinivibrio* | Bacteria; Proteobacteria; Gammaproteobacteria; Vibrionales; Vibrionaceae | MULTISPECIES: methyl-accepting chemotaxis protein | 6 |
|  | **WP_077639566.1** | 3.5E-05 | Qa.III | ---INEQIVQISDMNTQVATATEEQSTVVNDINQNIEEINHSSQQTA------ | *Salinivibrio* | Bacteria; Proteobacteria; Gammaproteobacteria; Vibrionales; Vibrionaceae | MULTISPECIES: methyl-accepting chemotaxis protein | 6 |
|  | **WP_077764979.1** | 3.5E-05 | Qa.III | ---INEQIVQISDMNTQVATATEEQSTVVNDINQNIEEINHSSQQTA------ | *Salinivibrio sp. PR5* | Bacteria; Proteobacteria; Gammaproteobacteria; Vibrionales; Vibrionaceae; Salinivibrio | methyl-accepting chemotaxis protein | 6 |
|  | **WP_079587049.1** | 3.5E-05 | Qa.III | IQGITATIEEVSTIATSIGSAIEEQGAATAEIARNVTQTAEATK--------- | *Bradyrhizobium lablabi* | Bacteria; Proteobacteria; Alphaproteobacteria; Rhizobiales; Bradyrhizobiaceae; Bradyrhizobium | methyl-accepting chemotaxis protein | 6 |

| | | | | | | | |
|---|---|---|---|---|---|---|---|
| **WP_081837180.1** | 3.5E-05 | Qc.III.b | ---LGSAIERVTSISEEIGAMINEQTQAIEEITRSVTVASERSQDVSGNIS-- | *alpha proteobacterium Q-1* | Bacteria; Proteobacteria; Alphaproteobacteria | hypothetical protein | 6 |
| **WP_085401867.1** | 3.5E-05 | Qc.III | -QTIGGIIGEVNEVATAIAAAVQEQGAATQEITRSTQFAAQGTRNVS------ | *Bradyrhizobium japonicum* | Bacteria; Proteobacteria; Alphaproteobacteria; Rhizobiales; Bradyrhizobiaceae; Bradyrhizobium | methyl-accepting chemotaxis protein | 6 |
| **WP_092516896.1** | 3.5E-05 | Qa.III | IQGITATIEEVSTIATSIGSAIEEQGAATAEIARNVTQTAEATK--------- | *Afipia sp. GAS231* | Bacteria; Proteobacteria; Alphaproteobacteria; Rhizobiales; Bradyrhizobiaceae; Afipia | methyl-accepting chemotaxis protein | 6 |
| **WP_110749963.1** | 3.5E-05 | Qb.I | LQRTKLQLQETVEQSDVSSEELKASNEEMHAINEELRSATEELETS------- | *Pseudomonas sp. OV529* | Bacteria; Proteobacteria; Gammaproteobacteria; Pseudomonadales; Pseudomonadaceae; Pseudomonas | PAS domain S-box protein | 6 |
| **WP_114360897.1** | 3.5E-05 | Qc.III | IKEIGDTIGKMSEIASTIASAVEQQGAATQEISRNVQQAASGTQQVSSNITD- | *Rhodopseudomonas pentothenatexigens* | Bacteria; Proteobacteria; Alphaproteobacteria; Rhizobiales; Bradyrhizobiaceae; Rhodopseudomonas | HAMP domain-containing protein | 6 |
| **CDA24737.1** | 3.6E-05 | Qc.III | --QVSAGADQLAQSAQGLAEGATDQAAALEELTSSVNEVAENVAENTKSTD-- | *Roseburia sp. CAG:197* | Bacteria; Firmicutes; Clostridia; Clostridiales; Lachnospiraceae; Roseburia; environmental samples | methyl-accepting chemotaxis sensory transducer | 6 |
| **KJE35257.1** | 3.6E-05 | Qc.III | INEITDIVGELNESASAIAAAVEEQGAATQEISGSVENAS------------ | *Thalassospira sp. HJ* | Bacteria; Proteobacteria; Alphaproteobacteria; Rhodospirillales; Rhodospirillaceae; Thalassospira | hypothetical protein UF64_11495 | 6 |
| **KPU83444.1** | 3.6E-05 | Qc.III.c | -DTMSGSILEINDLGVQIATAAEEQSSVTEDISRQMTAIREMVNTLS------ | *Psychromonas sp. PRT-SC03* | Bacteria; Proteobacteria; Gammaproteobacteria; Alteromonadales; Psychromonadaceae; Psychromonas | chemotaxis protein | 6 |
| **OLA72313.1** | 3.6E-05 | Qc.III | --QVSAGADQLAQSAQGLAEGATDQAAALEELTSSVNEVAENVAENTKSTD-- | *Roseburia sp. CAG:197_41_10* | Bacteria; Firmicutes; Clostridia; Clostridiales; Lachnospiraceae; Roseburia | hypothetical protein BHW45_07780 | 6 |
| **PMU94787.1** | 3.6E-05 | Qa.III | -------IGEIDGMNQSVATATEEQTSVVESINMDITEINTLNQEGVENLQST | *Pseudomonas sp. MPBD4-3* | Bacteria; Proteobacteria; Gammaproteobacteria; Pseudomonadales; Pseudomonadaceae; Pseudomonas | chemotaxis protein partial | 6 |
| **SHJ75666.1** | 3.6E-05 | Qc.III | IKEISGTIEKLSEISSTIAAAVEEQGAATQEISRNVQQAAQGTMQVSSNIT-- | *Bradyrhizobium lablabi* | Bacteria; Proteobacteria; Alphaproteobacteria; Rhizobiales; Bradyrhizobiaceae; Bradyrhizobium | methyl-accepting chemotaxis sensory transducer | 6 |
| **SMX59626.1** | 3.6E-05 | Qc.III | ---ISGTIERLSEIASAIAAAVEEQGAATQEISRNVQQAARGTQEVSANI--- | *Bradyrhizobium sp. ORS 285* | Bacteria; Proteobacteria; Alphaproteobacteria; Rhizobiales; Bradyrhizobiaceae; Bradyrhizobium | putative sensory transducer protein; signal peptide | 6 |
| **WP_009030106.1** | 3.6E-05 | Qa.III | IRDIGTTIGQMNEITETIASAVEEQGVATKEIARSVQQAA------------ | *Bradyrhizobium sp. ORS 375* | Bacteria; Proteobacteria; Alphaproteobacteria; Rhizobiales; Bradyrhizobiaceae; Bradyrhizobium | HAMP domain-containing protein | 6 |
| **WP_013502732.1** | 3.6E-05 | Qa.III | IREIGATISKMSEIASTIASAVEEQGAATQEISRNVQQAAQGTQQVS------ | *Rhodopseudomonas palustris* | Bacteria; Proteobacteria; Alphaproteobacteria; Rhizobiales; Bradyrhizobiaceae; Rhodopseudomonas | HAMP domain-containing protein | 6 |
| **WP_022722817.1** | 3.6E-05 | Qc.III | -REIGHTIGRMSEIASTIASAVEEQGAATQEISRNVQQAAGGTQQVSANIA-- | *Rhodopseudomonas sp. B29* | Bacteria; Proteobacteria; Alphaproteobacteria; Rhizobiales; Bradyrhizobiaceae; Rhodopseudomonas | methyl-accepting chemotaxis protein partial | 6 |
| **WP_023461811.1** | 3.6E-05 | Qa.III | IRGIGATIKHMNEIASSIADSVHQQGDATNEIARNVNEAS------------ | *Asticcacaulis sp. YBE204* | Bacteria; Proteobacteria; Alphaproteobacteria; Caulobacterales; Caulobacteraceae; Asticcacaulis | methyl-accepting chemotaxis protein | 6 |
| **WP_029009125.1** | 3.6E-05 | Qa.IV | IREIAGTIRSINEIATTIAAAVEQQNTATREIARNVGEAA------------ | *Azospirillum halopraeferens* | Bacteria; Proteobacteria; Alphaproteobacteria; Rhodospirillales; Rhodospirillaceae; Azospirillum | HAMP domain-containing protein | 6 |
| **WP_051243461.1** | 3.6E-05 | Qc.III | --HIAQTIGRMNEIAAAVAAAVEQQGAATSEIARNVEEASSGTRDVSSNIA-- | *Thalassobaculum salexigens* | Bacteria; Proteobacteria; Alphaproteobacteria; Rhodospirillales; Rhodospirillaceae; Thalassobaculum | hypothetical protein | 6 |
| **WP_051382625.1** | 3.6E-05 | Qa.IV | IKEIGTTIARISEISAAIASAVEEQGAATQEISRNVQRAAE----------- | *Bradyrhizobium sp. Tv2a-2* | Bacteria; Proteobacteria; Alphaproteobacteria; Rhizobiales; Bradyrhizobiaceae; Bradyrhizobium | HAMP domain-containing protein | 6 |
| **WP_052741743.1** | 3.6E-05 | Qc.III | -SEIAAIVSNVKEVSTAIASAVEQQGAATQEIARNIEQASGGTAEVNNNIQ-- | *Kiloniella litopenaei* | Bacteria; Proteobacteria; Alphaproteobacteria; Kiloniellales; Kiloniellaceae; Kiloniella | methyl-accepting chemotaxis protein | 6 |
| **WP_082736991.1** | 3.6E-05 | Qa.III | -EAITETIDNVNGIFAVISTAVEEQSAVISDMAMNMTEAANS---------- | *Rhizobiales bacterium CCH3-A5* | Bacteria; Proteobacteria; Alphaproteobacteria; Rhizobiales | PAS domain S-box protein | 6 |
| **WP_085126365.1** | 3.6E-05 | Qc.III | -GSIGATITRINEIAATMAATVDQQGSATREIARSVQEAATGTRSATSNVE-- | *Tistlia consotensis* | Bacteria; Proteobacteria; Alphaproteobacteria; Rhodospirillales; Rhodospirillaceae; Tistlia | methyl-accepting chemotaxis protein | 6 |
| **WP_085902384.1** | 3.6E-05 | Qc.III | ---ISTTIERMNEFANAIAAAIEEQGAATQEISNSVQKAAAGTQEVSSNV--- | *Kiloniella majae* | Bacteria; Proteobacteria; Alphaproteobacteria; Kiloniellales; Kiloniellaceae; Kiloniella | methyl-accepting chemotaxis protein | 6 |
| **WP_085906598.1** | 3.6E-05 | Qc.III | ---ISTTIERMNEFANAIAAAIEEQGAATQEISNSVQKAAAGTQEVSSNV--- | *Kiloniella majae* | Bacteria; Proteobacteria; Alphaproteobacteria; Kiloniellales; Kiloniellaceae; Kiloniella | methyl-accepting chemotaxis protein | 6 |
| **WP_093150027.1** | 3.6E-05 | Qc.III | --HIAQTIGRMNEIAAAVAAAVEQQGAATSEIARNVEEASSGTRDVSSNIA-- | *Thalassobaculum litoreum* | Bacteria; Proteobacteria; Alphaproteobacteria; Rhodospirillales; Rhodospirillaceae; Thalassobaculum | hypothetical protein | 6 |
| **WP_094332797.1** | 3.6E-05 | Qc.I | -----QMAGQLKESFETLEQRVSDRTAALAESNQQLEHAKEKAEVA------- | *Nostoc* | Bacteria; Cyanobacteria; Nostocales; Nostocaceae | MULTISPECIES: response regulator | 6 |
| **WP_107344408.1** | 3.6E-05 | Qa.III | IREIGATISKMSEIASTIASAVEEQGAATQEISRNVQQAAQGTQQVS------ | *Rhodopseudomonas palustris* | Bacteria; Proteobacteria; Alphaproteobacteria; Rhizobiales; Bradyrhizobiaceae; Rhodopseudomonas | methyl-accepting chemotaxis protein | 6 |
| **WP_107877598.1** | 3.6E-05 | Qa.III | IRGIGATIKHMNEIASSIADSVHQQGDATNEIARNVNEAS------------ | *Asticcacaulis sp. CF398* | Bacteria; Proteobacteria; Alphaproteobacteria; Caulobacterales; Caulobacteraceae; Asticcacaulis | methyl-accepting chemotaxis protein | 6 |
| **WP_114359851.1** | 3.6E-05 | Qa.III | IREIGATISKMSEIASTIASAVEEQGAATQEISRNVQQAAQGTQQVS------ | *Rhodopseudomonas pentothenatexigens* | Bacteria; Proteobacteria; Alphaproteobacteria; Rhizobiales; Bradyrhizobiaceae; Rhodopseudomonas | HAMP domain-containing protein | 6 |
| **WP_114368185.1** | 3.6E-05 | Qa.III | --AITQVIEQIREASLSIASAVEEQSVTTKQISTNINQTAESA---------- | *Blastopirellula cremea* | Bacteria; Planctomycetes; Planctomycetia; Planctomycetales; Planctomycetaceae; Blastopirellula | HAMP domain-containing protein | 6 |
| **ODN72260.1** | 3.7E-05 | Qc.III | -SMISTTIGDINEISSSIASAVEEQNAATGEIARNVEEASR----------- | *Methylobrevis pamukkalensis* | Bacteria; Proteobacteria; Alphaproteobacteria; Rhizobiales; Methylobrevis | putative methyl-accepting chemotaxis protein YoaH | 6 |
| **OQW79666.1** | 3.7E-05 | Qa.IV | --EITSTISDISEISNAIASAVDEQNSATSEISRSIQTVTTQT---------- | *Proteobacteria bacterium ST_bin15* | Bacteria; Proteobacteria | hypothetical protein BVN31_14000 | 6 |
| **OYU80737.1** | 3.7E-05 | Qc.I | ---------RLKNYASHLEELVEERTKTLNNTIRALEKSKGKVSAALEK---- | *Flavobacterium sp. BFFFF1* | Bacteria; Bacteroidetes; Flavobacteriia; Flavobacteriales; Flavobacteriaceae; Flavobacterium | PAS domain-containing sensor histidine kinase | 6 |
| **PID28190.1** | 3.7E-05 | Qc.III | IEDVSEIIGKINEININIAQTVDEQTRTIGEIAKITARSSEKSEE-------- | *Candidatus Cloacimonetes bacterium* | Bacteria; Candidatus Cloacimonetes | hypothetical protein CSB55_05890 | 6 |
| **SCY05949.1** | 3.7E-05 | Qa.III | IRAITDVIVEVNDVVSTIAAAVEEQAVTTREIAANVAQAA------------ | *Desulfoluna spongiiphila* | Bacteria; Proteobacteria; Deltaproteobacteria; Desulfobacterales; Desulfobacteraceae; Desulfoluna | methyl-accepting chemotaxis sensory transducer with Cache sensor | 6 |
| **WP_009540922.1** | 3.7E-05 | Qc.III | ---IGKTVNEISEVASAIASAVEEQNAATSEIARNTNEASAGTHDVSSNI--- | *Caenispirillum salinarum* | Bacteria; Proteobacteria; Alphaproteobacteria; Rhodospirillales; Rhodospirillaceae; Caenispirillum | methyl-accepting chemotaxis protein | 6 |
| **WP_011084287.1** | 3.7E-05 | Qc.III | -QTIGGIIGEVNEVATAIAAAVQEQGAATQEITRSTQFAAQGTRNVS------ | *Bradyrhizobium diazoefficiens* | Bacteria; Proteobacteria; Alphaproteobacteria; Rhizobiales; Bradyrhizobiaceae; Bradyrhizobium | methyl-accepting chemotaxis protein | 6 |
| **WP_027538285.1** | 3.7E-05 | Qc.III | IKRIGGIIGEVNEVATAIAAAVQEQGAATQEITRSTQYAAQGTKNV------- | *Bradyrhizobium sp. URHA0002* | Bacteria; Proteobacteria; Alphaproteobacteria; Rhizobiales; Bradyrhizobiaceae; Bradyrhizobium | methyl-accepting chemotaxis protein | 6 |
| **WP_045156199.1** | 3.7E-05 | Qb.I | LQRTKLQLQDVIEQSEVSSEELKASNEEMQAINEELRSASEELETSK------ | *Pseudomonas fluorescens* | Bacteria; Proteobacteria; Gammaproteobacteria; Pseudomonadales; Pseudomonadaceae; Pseudomonas | PAS domain S-box protein | 6 |
| **WP_046503117.1** | 3.7E-05 | Qa.IV | --SITEVITEVNDISQTISDSIEQQNLATQEISQNVQETAINTTEASDEV--- | *Kiloniella litopenaei* | Bacteria; Proteobacteria; Alphaproteobacteria; Kiloniellales; Kiloniellaceae; Kiloniella | methyl-accepting chemotaxis protein | 6 |
| **WP_054701312.1** | 3.7E-05 | Qa.III | ISEITEVIRDVNDVVATIATAVEEQSAATNDIATNVSQASQGIQEV------- | *Desulfosarcina cetonica* | Bacteria; Proteobacteria; Deltaproteobacteria; Desulfobacterales; Desulfobacteraceae; Desulfosarcina | methyl-accepting chemotaxis protein | 6 |
| **WP_079564123.1** | 3.7E-05 | Qc.III | IKEISGTIERLSEISSTIAAAVEEQGAATQEISRNVQQAAHGTQQVSS----- | *Bradyrhizobium erythrophlei* | Bacteria; Proteobacteria; Alphaproteobacteria; Rhizobiales; Bradyrhizobiaceae; Bradyrhizobium | methyl-accepting chemotaxis protein | 6 |
| **WP_080668138.1** | 3.7E-05 | Qc.III | -QTIGGIIGEVNEVATAIAAAVQEQGAATQEITRSTQFAAQGTRNVS------ | *Bradyrhizobium* | Bacteria; Proteobacteria; Alphaproteobacteria; Rhizobiales; Bradyrhizobiaceae | MULTISPECIES: methyl-accepting chemotaxis protein | 6 |
| **WP_082049715.1** | 3.7E-05 | Qc.III | INEITDIVGELNESASAIAAAVEEQGAATQEISGSVENAS------------ | *Thalassospira sp. HJ* | Bacteria; Proteobacteria; Alphaproteobacteria; Rhodospirillales; Rhodospirillaceae; Thalassospira | methyl-accepting chemotaxis protein | 6 |
| **WP_084176963.1** | 3.7E-05 | Qc.III | IDEVGNIISRINETSAEISATMEQQDAATREIAVNVTEASRGTQ--------- | *Labrenzia sp. DG1229* | Bacteria; Proteobacteria; Alphaproteobacteria; Rhodobacterales; Rhodobacteraceae; Labrenzia | HAMP domain-containing protein | 6 |
| **WP_085643911.1** | 3.7E-05 | Qc.III | --SIAEVIEKINEISSSIAAAVEEQQVATGEIARNVDEAAKGTEDVNENIHE- | *Thalassospira sp. MCCC 1A03138* | Bacteria; Proteobacteria; Alphaproteobacteria; Rhodospirillales; Rhodospirillaceae; Thalassospira | methyl-accepting chemotaxis protein | 6 |
| **WP_088873681.1** | 3.7E-05 | Qc.III | --AISGTIENINSIATTIASAVEEQGAATQEIARNVQQAAAGADSVTHNI--- | *Nitrospirillum amazonense* | Bacteria; Proteobacteria; Alphaproteobacteria; Rhodospirillales; Rhodospirillaceae; Nitrospirillum | methyl-accepting chemotaxis protein | 6 |
| **WP_092209426.1** | 3.7E-05 | Qa.III | IRAITDVIVEVNDVVSTIAAAVEEQAVTTREIAANVAQAA------------ | *Desulfoluna spongiiphila* | Bacteria; Proteobacteria; Deltaproteobacteria; Desulfobacterales; Desulfobacteraceae; Desulfoluna | methyl-accepting chemotaxis protein | 6 |
| **WP_092511713.1** | 3.7E-05 | Qc.III | IKRIGGIIGEVNEVATAIAAAVQEQGAATQEITRSTQYAAQGTKNV------- | *Afipia sp. GAS231* | Bacteria; Proteobacteria; Alphaproteobacteria; Rhizobiales; Bradyrhizobiaceae; Afipia | methyl-accepting chemotaxis protein | 6 |
| **AAU92470.1** | 3.8E-05 | Qb.I | --ATREDLRQTIEKSRHVNDELRASNEEIMAMNEELQSANEELESSR------ | *Methylococcus capsulatus str. Bath* | Bacteria; Proteobacteria; Gammaproteobacteria; Methylococcales; Methylococcaceae; Methylococcus | putative methyltransferase CheR | 6 |
| **CCC96375.1** | 3.8E-05 | Qa.III | IKTIGGTIGEINGIATTIAAAVEEQGAATGEIARNVQQAA------------ | *Azospirillum brasilense Sp245* | Bacteria; Proteobacteria; Alphaproteobacteria; Rhodospirillales; Rhodospirillaceae; Azospirillum | putative methyl-accepting chemotaxis protein | 6 |
| **OJW63195.1** | 3.8E-05 | Qc.III | -REISGTISRISEISSAIASAVEEQGAATQEISRNIMQSAEGTHQV------- | *Afipia sp. 64-13* | Bacteria; Proteobacteria; Alphaproteobacteria; Rhizobiales; Bradyrhizobiaceae; Afipia | chemotaxis protein | 6 |
| **OPH20622.1** | 3.8E-05 | Qa.III | IKTIGGTIGEINGIATTIAAAVEEQGAATGEIARNVQQAA------------ | *Azospirillum brasilense* | Bacteria; Proteobacteria; Alphaproteobacteria; Rhodospirillales; Rhodospirillaceae; Azospirillum | hypothetical protein FE88_13220 | 6 |
| **PLX67633.1** | 3.8E-05 | SNAP.b | TRNMMERIAATVSESAQTVRKLGESSEQIGQIIQVIDDIADQT---------- | *Denitrovibrio sp.* | Bacteria; Deferribacteres; Deferribacterales; Deferribacteraceae; Denitrovibrio | chemotaxis protein | 6 |
| **PWC63511.1** | 3.8E-05 | Qa.III | IKTIGGTIGEINGIATTIAAAVEEQGAATGEIARNVQQAA------------ | *Azospirillum sp. TSH58* | Bacteria; Proteobacteria; Alphaproteobacteria; Rhodospirillales; Rhodospirillaceae; Azospirillum | hypothetical protein TSH58_23400 | 6 |
| **WP_008970145.1** | 3.8E-05 | Qc.III | -KGIGGIIGEVNEVATAIAAAVQEQGAATQEISRSTQFAAQGTKNVS------ | *Bradyrhizobium sp. STM 3843* | Bacteria; Proteobacteria; Alphaproteobacteria; Rhizobiales; Bradyrhizobiaceae; Bradyrhizobium | methyl-accepting chemotaxis protein | 6 |
| **WP_009030537.1** | 3.8E-05 | Qa.IV | IKEIGSTITQISDIAATIAATVEQQGAATREIARNVGEAA------------ | *Bradyrhizobium sp. ORS 375* | Bacteria; Proteobacteria; Alphaproteobacteria; Rhizobiales; Bradyrhizobiaceae; Bradyrhizobium | methyl-accepting chemotaxis protein | 6 |
| **WP_011662970.1** | 3.8E-05 | Qa.IV | IKEIGNTIERMSDIAATIAAAVEQQGAATQEISRNVQQAA------------ | *Rhodopseudomonas palustris* | Bacteria; Proteobacteria; Alphaproteobacteria; Rhizobiales; Bradyrhizobiaceae; Rhodopseudomonas | methyl-accepting chemotaxis protein | 6 |
| **WP_012495464.1** | 3.8E-05 | Qc.III | -SEISGTIEKLSEISAHIASAVEEQGAATQEISRNIQQTAQGTERV------- | *Rhodopseudomonas palustris* | Bacteria; Proteobacteria; Alphaproteobacteria; Rhizobiales; Bradyrhizobiaceae; Rhodopseudomonas | PAS domain S-box protein | 6 |

| | | | | | | | | |
|---|---|---|---|---|---|---|---|---|
| | **WP_015683012.1** | 3.8E-05 | Qc.II | --ELFKTLKSIREITTDFNVQIESQDKSIHELTEFVSSFSESIQTSVDNIGK- | *Leptospira wolbachii* | Bacteria; Spirochaetes; Leptospirales; Leptospiraceae; Leptospira | methyl-accepting chemotaxis protein | 6 |
|  | **WP_019644840.1** | 3.8E-05 | Qa.IV | ----SRTIADISAISSAILETVEQQAAATTEIARQAEHAAGNTDDVA------ | *Novispirillum itersonii* | Bacteria; Proteobacteria; Alphaproteobacteria; Rhodospirillales; Rhodospirillaceae; Novispirillum | hypothetical protein | 6 |
| | **WP_024517292.1** | 3.8E-05 | Qc.III | IKEISGTIEKLSEISSTIAAAVEEQGAATQEIARNVQQAAQGTQQVSSNITD- | *Bradyrhizobium sp. Tv2a-2* | Bacteria; Proteobacteria; Alphaproteobacteria; Rhizobiales; Bradyrhizobiaceae; Bradyrhizobium | HAMP domain-containing protein | 6 |
|  | **WP_036246533.1** | 3.8E-05 | Qb.I | --ATREDLRQTIEKSRHVNDELRASNEEIMAMNEELQSANEELESSR------ | *Methylococcus capsulatus* | Bacteria; Proteobacteria; Gammaproteobacteria; Methylococcales; Methylococcaceae; Methylococcus | EAL domain-containing protein | 6 |
|  | **WP_038971356.1** | 3.8E-05 | Qc.III | -QAIGGIIGEVNEVATAIAAAVQEQGAATQEITRSTQYAAQGTKNVS------ | *Bradyrhizobium sp. CCBAU 15635* | Bacteria; Proteobacteria; Alphaproteobacteria; Rhizobiales; Bradyrhizobiaceae; Bradyrhizobium | methyl-accepting chemotaxis protein | 6 |
|  | **WP_041360956.1** | 3.8E-05 | Qb.I | --ATREDLRQTIEKSRHVNDELRASNEEIMAMNEELQSANEELESSR------ | *Methylococcus capsulatus* | Bacteria; Proteobacteria; Gammaproteobacteria; Methylococcales; Methylococcaceae; Methylococcus | EAL domain-containing protein | 6 |
|  | **WP_042440966.1** | 3.8E-05 | Qc.III | -SEISGTIEKLSEISAHIASAVEEQGAATQEISRNIQQTAQGTERV------- | *Rhodopseudomonas palustris* | Bacteria; Proteobacteria; Alphaproteobacteria; Rhizobiales; Bradyrhizobiaceae; Rhodopseudomonas | chemotaxis protein | 6 |
| | **WP_050471385.1** | 3.8E-05 | Qc.III | -GSILATIESIRTISSTIAAAVEEQTAVTRDISHNMQAASTGVQSITSSMG-- | *Pannonibacter phragmitetus* | Bacteria; Proteobacteria; Alphaproteobacteria; Rhodobacterales; Rhodobacteraceae; Pannonibacter | PAS domain S-box protein | 6 |
|  | **WP_056820583.1** | 3.8E-05 | Qb.I | LVRLKAHLRDTVEQYEASTEELKASNEELQAMNEELRSASEELETS------- | *Rhizobacter* | Bacteria; Proteobacteria; Betaproteobacteria; Burkholderiales | MULTISPECIES: PAS domain S-box protein | 6 |
| | **WP_058900902.1** | 3.8E-05 | Qc.III | -GSILATIESIRTISSTIAAAVEEQTAVTRDISHNMQAASTGVQSITSSMG-- | *Pannonibacter phragmitetus* | Bacteria; Proteobacteria; Alphaproteobacteria; Rhodobacterales; Rhodobacteraceae; Pannonibacter | PAS domain S-box protein | 6 |
| | **WP_062469262.1** | 3.8E-05 | R.IV | IEQINQAIAQMDQVTQQNAALVEEASATAQSLQEQAQGLVRAAGVFRTGAG-- | *Variovorax boronicumulans* | Bacteria; Proteobacteria; Betaproteobacteria; Burkholderiales; Comamonadaceae; Variovorax | PAS domain S-box protein | 6 |
|  | **WP_068030054.1** | 3.8E-05 | Qc.III | IKEIGSTIGRISDIATSIAAAIEQQGATTSQIADGVESAAR------------ | *Rhodoplanes sp. Z2-YC6860* | Bacteria; Proteobacteria; Alphaproteobacteria; Rhizobiales; Hyphomicrobiaceae; Rhodoplanes | methyl-accepting chemotaxis protein | 6 |
|  | **WP_068585143.1** | 3.8E-05 | Qa.III | LQEIHASMDEISERVSTIATATNEQSVGIDSVSQSVSEMENSTQQ-------- | *Thiomicrospira sp. WB1* | Bacteria; Proteobacteria; Gammaproteobacteria; Thiotrichales; Piscirickettsiaceae; Thiomicrospira | HAMP domain-containing protein | 6 |
| | **WP_070061535.1** | 3.8E-05 | R.IV | IEQINQAIAQMDQVTQQNAALVEEASATAQSLQEQAQGLVRAAGVFRTGAG-- | *Variovorax boronicumulans* | Bacteria; Proteobacteria; Betaproteobacteria; Burkholderiales; Comamonadaceae; Variovorax | PAS domain S-box protein | 6 |
| | **WP_072821678.1** | 3.8E-05 | Qc.III | IKDISGTIEKLSEISSTIAAAVEEQGAATQEISRNVQQAAQGTQQVSSNITD- | *Bradyrhizobium erythrophlei* | Bacteria; Proteobacteria; Alphaproteobacteria; Rhizobiales; Bradyrhizobiaceae; Bradyrhizobium | methyl-accepting chemotaxis protein | 6 |
|  | **WP_083255415.1** | 3.8E-05 | Qc.III | -SMISTTIGDINEISSSIASAVEEQNAATGEIARNVEEASR------------ | *Methylobrevis pamukkalensis* | Bacteria; Proteobacteria; Alphaproteobacteria; Rhizobiales; Methylobrevis | hypothetical protein | 6 |
|  | **WP_094456326.1** | 3.8E-05 | Qa.IV | VQEISLTITEMDQISQAIASAIEEQGAATQEISRNVNQAAE------------ | *Niveispirillum lacus* | Bacteria; Proteobacteria; Alphaproteobacteria; Rhodospirillales; Rhodospirillaceae; Niveispirillum | methyl-accepting chemotaxis protein | 6 |
|  | **WP_104512275.1** | 3.8E-05 | Qc.III | -SEISGTIEKLSEISAHIASAVEEQGAATQEISRNIQQTAQGTERV------- | *Rhodopseudomonas palustris* | Bacteria; Proteobacteria; Alphaproteobacteria; Rhizobiales; Bradyrhizobiaceae; Rhodopseudomonas | chemotaxis protein | 6 |
|  | **WP_107343896.1** | 3.8E-05 | Qc.III | -SEISGTIEKLSEISAHIASAVEEQGAATQEISRNIQQTAQGTERV------- | *Rhodopseudomonas palustris* | Bacteria; Proteobacteria; Alphaproteobacteria; Rhizobiales; Bradyrhizobiaceae; Rhodopseudomonas | chemotaxis protein | 6 |
|  | **WP_109051611.1** | 3.8E-05 | Qa.III.b | --EIGRVIAEINHISTTIAAAVEEQGTATKEISRNVQQAARGTQQ-------- | *Azospirillum sp. TSA6c* | Bacteria; Proteobacteria; Alphaproteobacteria; Rhodospirillales; Rhodospirillaceae; Azospirillum | methyl-accepting chemotaxis protein | 6 |
|  | **WP_110785354.1** | 3.8E-05 | Qc.III | -SEISGTIEKLSEISAHIASAVEEQGAATQEISRNIQQTAQGTERV------- | *Rhodopseudomonas palustris* | Bacteria; Proteobacteria; Alphaproteobacteria; Rhizobiales; Bradyrhizobiaceae; Rhodopseudomonas | chemotaxis protein | 6 |
|  | **ADU12618.1** | 3.9E-05 | Qc.III | IERISSSMSEVEQIATAIAATAEEQGAATQEIARSVMRTSSG----------- | *Asticcacaulis excentricus CB 48* | Bacteria; Proteobacteria; Alphaproteobacteria; Caulobacterales; Caulobacteraceae; Asticcacaulis | methyl-accepting chemotaxis sensory transducer | 6 |
|  | **CAE27372.1** | 3.9E-05 | Qc.III | -SEISGTIEKLSEISAHIASAVEEQGAATQEISRNIQQTAQGTERV------- | *Rhodopseudomonas palustris CGA009* | Bacteria; Proteobacteria; Alphaproteobacteria; Rhizobiales; Bradyrhizobiaceae; Rhodopseudomonas | methyl-accepting chemotaxis receptor/ sensory transducer | 6 |
|  | **EEA94892.1** | 3.9E-05 | Qa.IV | IQEITSSIRQVNEFASNIAASVEQQGAATAEITENVHQAA------------- | *Pseudovibrio sp. JE062* | Bacteria; Proteobacteria; Alphaproteobacteria; Rhodobacterales; Rhodobacteraceae; Pseudovibrio | methyl-accepting chemotaxis sensory transducer | 6 |
|  | **EGH99314.1** | 3.9E-05 | Qa.III | ---ISQALRNLNSLNASIASATLQQTHVVDDINQNVTQAAQLSQ--------- | *Pseudomonas amygdali pv. lachrymans str. M302278* | Bacteria; Proteobacteria; Gammaproteobacteria; Pseudomonadales; Pseudomonadaceae; Pseudomonas; Pseudomonas amygdali | methyl-accepting chemotaxis protein | 6 |
| | **EKD38242.1** | 3.9E-05 | Qc.III | IEEITTVINNVNEIVATITTAVGEQSKATEEIASNINQAAGGLGEVNENVS-- | *uncultured bacterium* | Bacteria; environmental samples | hypothetical protein ACD_75C00809G0002 partial | 6 |
|  | **EPM82459.1** | 3.9E-05 | Qa.III | ---ISQALRNLNSLNASIASATLQQTHVVDDINQNVTQAAQLSQ--------- | *Pseudomonas syringae pv. actinidiae ICMP 18804* | Bacteria; Proteobacteria; Gammaproteobacteria; Pseudomonadales; Pseudomonadaceae; Pseudomonas; Pseudomonas syringae | methyl-accepting chemotaxis protein | 6 |
|  | **KJV10174.1** | 3.9E-05 | Qa.IV | INSILTRITDINEIAATVAAAVEQQSAATAEIARNVQQAAE------------ | *Elstera litoralis* | Bacteria; Proteobacteria; Alphaproteobacteria; Rhodospirillales; Rhodospirillaceae; Elstera | chemotaxis protein partial | 6 |
|  | **OFX07341.1** | 3.9E-05 | Qa.III | IKAINGTISRINEIAANVASAVEEQSSATQEIARNVQEAASGTQQVNN----- | *Alphaproteobacteria bacterium RIFOXYD12_FULL_60_8* | Bacteria; Proteobacteria; Alphaproteobacteria | hypothetical protein A2516_01755 | 6 |
| | **OUT41416.1** | 3.9E-05 | Qc.III | --EISKRIEEIDQIATSIAAAVEEQSAATSEIARSVSYSSDNMRRVTESLD-- | *Micavibrio sp. TMED2* | Bacteria; Proteobacteria; Alphaproteobacteria; Micavibrio | hypothetical protein CBB62_03470 | 6 |
|  | **SBV98160.1** | 3.9E-05 | Qc.III.c | -ETISTTIGRVSEISAAISAAVEEQGAATREIARNVEEAAQ------------ | *uncultured Alphaproteobacteria bacterium* | Bacteria; Proteobacteria; Alphaproteobacteria; environmental samples | Methyl-accepting chemotaxis sensory transducer | 6 |
|  | **SPJ13710.1** | 3.9E-05 | Qb.I | ---TRENLQATIEGQEAANEELKSTNEELQSTNEELQSMNEEMETAK------ | *Syntrophobacter sp. SbD2* | Bacteria; Proteobacteria; Deltaproteobacteria; Syntrophobacterales; Syntrophobacteraceae; Syntrophobacter | MCP methyltransferase/ methylesterase | 6 |
|  | **WP_006034449.1** | 3.9E-05 | Qc.III | LDSMTESVMQINDLGIQIATAAEEQSSVTEEINRNMTMIQGMVNQLTE----- | *Moritella sp. PE36* | Bacteria; Proteobacteria; Gammaproteobacteria; Alteromonadales; Moritellaceae; Moritella | methyl-accepting chemotaxis protein partial | 6 |
| | **WP_025032190.1** | 3.9E-05 | Qc.III | --TIGGTIERLSEISSTIAAAVEEQGAATSEISRNVQSAAAGTTQVSANI--- | *Bradyrhizobium sp. DOA9* | Bacteria; Proteobacteria; Alphaproteobacteria; Rhizobiales; Bradyrhizobiaceae; Bradyrhizobium | methyl-accepting chemotaxis protein | 6 |
|  | **WP_035646875.1** | 3.9E-05 | Qc.III | -KAIGSTIERINEIAASISGAVELQRAATQNITQSVRSAASG----------- | *Bradyrhizobium sp. ORS 285* | Bacteria; Proteobacteria; Alphaproteobacteria; Rhizobiales; Bradyrhizobiaceae; Bradyrhizobium | HAMP domain-containing protein | 6 |
| | **WP_036009519.1** | 3.9E-05 | Qc.III | --TIGGTIERLSEISSTIAAAVEEQGAATSEISRNVQSAAAGTTQVSANI--- | *Bradyrhizobium yuanmingense* | Bacteria; Proteobacteria; Alphaproteobacteria; Rhizobiales; Bradyrhizobiaceae; Bradyrhizobium | methyl-accepting chemotaxis protein | 6 |
| | **WP_036030946.1** | 3.9E-05 | Qc.III | --TIGGTIERLSEISSTIAAAVEEQGAATSEISRNVQSAAAGTTQVSANI--- | *Bradyrhizobium yuanmingense* | Bacteria; Proteobacteria; Alphaproteobacteria; Rhizobiales; Bradyrhizobiaceae; Bradyrhizobium | methyl-accepting chemotaxis protein | 6 |
| | **WP_036040118.1** | 3.9E-05 | Qc.III | --TIGGTIERLSEISSTIAAAVEEQGAATSEISRNVQSAAAGTTQVSANI--- | *Bradyrhizobium yuanmingense* | Bacteria; Proteobacteria; Alphaproteobacteria; Rhizobiales; Bradyrhizobiaceae; Bradyrhizobium | methyl-accepting chemotaxis protein | 6 |
|  | **WP_054695695.1** | 3.9E-05 | Qa.III | ISEITEVIRDVNDVVATIATAVEEQSAATNDIATNVSQASQGIQEV------- | *Desulfosarcina cetonica* | Bacteria; Proteobacteria; Deltaproteobacteria; Desulfobacterales; Desulfobacteraceae; Desulfosarcina | methyl-accepting chemotaxis protein | 6 |
| | **WP_057027708.1** | 3.9E-05 | Qc.III | --TIGGTIERLSEISSTIAAAVEEQGAATSEISRNVQSAAAGTTQVSANI--- | *Bradyrhizobium yuanmingense* | Bacteria; Proteobacteria; Alphaproteobacteria; Rhizobiales; Bradyrhizobiaceae; Bradyrhizobium | HAMP domain-containing protein | 6 |
|  | **WP_057508007.1** | 3.9E-05 | Qa.III | MRDIVQSVQQVNQIMAEIAAASQEQSAGIEQVSQSITQMDANTRQ-------- | *Stenotrophomonas chelatiphaga* | Bacteria; Proteobacteria; Gammaproteobacteria; Xanthomonadales; Xanthomonadaceae; Stenotrophomonas | methyl-accepting chemotaxis protein | 6 |
|  | **WP_063699027.1** | 3.9E-05 | Qc.III | -QAIGGIIGEVNEVATAIAAAVQEQGAATQEITRSTQYAAQGTKNVS------ | *Bradyrhizobium sp. BR 10245* | Bacteria; Proteobacteria; Alphaproteobacteria; Rhizobiales; Bradyrhizobiaceae; Bradyrhizobium | methyl-accepting chemotaxis protein | 6 |
|  | **WP_079285293.1** | 3.9E-05 | Qa.III | IKTIGGTIGEINGIATTIAAAVEEQGAATGEIARNVQQAA------------- | *Azospirillum* | Bacteria; Proteobacteria; Alphaproteobacteria; Rhodospirillales; Rhodospirillaceae | MULTISPECIES: HAMP domain-containing protein | 6 |
| | **WP_083876006.1** | 3.9E-05 | Qc.III | IDSISETIKELKEISSNIASAVEQQGAATHEIARNVEQAAVGTQEVTSNI--- | *Caenispirillum salinarum* | Bacteria; Proteobacteria; Alphaproteobacteria; Rhodospirillales; Rhodospirillaceae; Caenispirillum | HAMP domain-containing protein | 6 |
|  | **WP_088286006.1** | 3.9E-05 | Qc.III | IEQIGSIIGTIHEKQTTIAAAVEEQSAVTQEIARTVSEVTGGAGQVNE----- | *Kineosporia sp. A_224* | Bacteria; Actinobacteria; Kineosporiales; Kineosporiaceae; Kineosporia | methyl-accepting chemotaxis protein | 6 |
| | **WP_109145287.1** | 3.9E-05 | Qc.III | --TIGGTIERLSEISSTIAAAVEEQGAATSEISRNVQSAAAGTTQVSANI--- | *Bradyrhizobium sp. SUTN9-2* | Bacteria; Proteobacteria; Alphaproteobacteria; Rhizobiales; Bradyrhizobiaceae; Bradyrhizobium | methyl-accepting chemotaxis protein | 6 |
|  | **ADG12901.1** | 4E-05 | Qa.IV | --------DEVNNAFLKIKEAVDKAHEKMEVIKQAVQNVTENVEKALRDVQ-- | *Methanocaldococcus infernus ME* | Archaea; Euryarchaeota; Methanococci; Methanococcales; Methanocaldococcaceae; Methanocaldococcus | methyl-accepting chemotaxis sensory transducer | 6 |
|  | **AWJ88656.1** | 4E-05 | Qa.III | IKTIGGTIGEINGIATTIAAAVEEQGAATGEIARNVQQAA------------- | *Azospirillum brasilense Sp245* | Bacteria; Proteobacteria; Alphaproteobacteria; Rhodospirillales; Rhodospirillaceae; Azospirillum | chemotaxis protein | 6 |
| | **CAL78252.1** | 4E-05 | Qc.III | IKQIGGTIQQISEIAAGIAAAVEQQGAATSQISRNVQDVAGGTARVVGNI--- | *Bradyrhizobium sp. ORS 278* | Bacteria; Proteobacteria; Alphaproteobacteria; Rhizobiales; Bradyrhizobiaceae; Bradyrhizobium | putative bacterial chemotaxis sensory transducer | 6 |
|  | **CCD87243.1** | 4E-05 | Qc.III | -KAIGSTIERINEIAASISGAVELQRAATQNITQSVRSAASG----------- | *Bradyrhizobium sp. ORS 285* | Bacteria; Proteobacteria; Alphaproteobacteria; Rhizobiales; Bradyrhizobiaceae; Bradyrhizobium | putative methyl-accepting chemotaxis protein | 6 |
| | **KJS39395.1** | 4E-05 | Qc.III | IEEIVAAINEVNDVASTIASAVEEQNAATQEIARNVEQAAAGSQEVSSNI--- | *Rhodospirillaceae bacterium BRH_c57* | Bacteria; Proteobacteria; Alphaproteobacteria; Rhodospirillales; Rhodospirillaceae | hypothetical protein VR70_08135 | 6 |
| | **KLK93108.1** | 4E-05 | Qa.IV | IEQIARTIAEMSQISISIAAAMEEQGAATAEIARNVQEAARGTEQVTGNI--- | *Microvirga vignae* | Bacteria; Proteobacteria; Alphaproteobacteria; Rhizobiales; Methylobacteriaceae; Microvirga | hypothetical protein AA309_11160 | 6 |
|  | **KPZ09231.1** | 4E-05 | Qa.III | -------IGEIDDMNQSVAAATEEQTSVVESLNFDISQINDLNQAGVQNLQST | *Pseudomonas syringae pv. spinaceae* | Bacteria; Proteobacteria; Gammaproteobacteria; Pseudomonadales; Pseudomonadaceae; Pseudomonas; Pseudomonas syringae | Chemotaxis protein | 6 |
|  | **KYD16054.1** | 4E-05 | Qc.III.c | LSVIMSVTEKINENSTNIANAVDQQASAIEQLNRAIESIAS------------ | *Anoxybacillus flavithermus* | Bacteria; Firmicutes; Bacilli; Bacillales; Bacillaceae; Anoxybacillus | hypothetical protein B4168_2730 | 6 |
|  | **PGH53627.1** | 4E-05 | Qc.III | --GIGGTIGSINDITTAIAAAIEEQSAATNEISRNVQQAAAGTQEVTGS---- | *Azospirillum sp. B2* | Bacteria; Proteobacteria; Alphaproteobacteria; Rhodospirillales; Rhodospirillaceae; Azospirillum | chemotaxis protein partial | 6 |
|  | **PKN22665.1** | 4E-05 | Qb.I | LRGARQQLQAAIEQFETANEELKSANEELQANNEELQSTNEELESSK------ | *Deltaproteobacteria bacterium HGW-Deltaproteobacteria-21* | Bacteria; Proteobacteria; Deltaproteobacteria | hypothetical protein CVU64_24670 | 6 |
|  | **WP_007679529.1** | 4E-05 | R.IV | -QEINHAVTQLDQMVQQNAELVVQSAAAAGALQGQAGELAQTAGHFR------ | *Cronobacter condimenti* | Bacteria; Proteobacteria; Gammaproteobacteria; Enterobacterales; Enterobacteriaceae; Cronobacter | methyl-accepting chemotaxis protein | 6 |
| | **WP_015669618.1** | 4E-05 | Qc.III | IQRIGGIIGEVNEVATAIAAAVQQQGAATQEISRSTQFAADGTKHVSENI--- | *Bradyrhizobium oligotrophicum* | Bacteria; Proteobacteria; Alphaproteobacteria; Rhizobiales; Bradyrhizobiaceae; Bradyrhizobium | methyl-accepting chemotaxis protein | 6 |
| | **WP_018458444.1** | 4E-05 | Qc.III | IEGVSGTISSINEIAIAIASAVEQQGVATREIARSIQKVAQGTEDVTRNI--- | *Bradyrhizobium sp. WSM4349* | Bacteria; Proteobacteria; Alphaproteobacteria; Rhizobiales; Bradyrhizobiaceae; Bradyrhizobium | HAMP domain-containing protein | 6 |

| | | | | | | | |
|---|---|---|---|---|---|---|---|
| **WP_028345470.1** | 4E-05 | Qc.III | -KAIGQTIERISDIATSISAAVEQQRGATQNIAQSVRAAAS------------ | *Bradyrhizobium elkanii* | Bacteria; Proteobacteria; Alphaproteobacteria; Rhizobiales; Bradyrhizobiaceae; Bradyrhizobium | methyl-accepting chemotaxis protein | 6 |
| **WP_035709178.1** | 4E-05 | Qc.III | IGEISGTIARLSEIASAIAAAVEQQGAATQEIARNVQQAAQGTQQVSSNVG-- | *Bradyrhizobium sp. CCBAU 43298* | Bacteria; Proteobacteria; Alphaproteobacteria; Rhizobiales; Bradyrhizobiaceae; Bradyrhizobium | PAS domain S-box protein | 6 |
| **WP_038149291.1** | 4E-05 | Qa.III | LEEITRAVSSISDMANQIATAAEEQSHVTGEITTNVTAIK------------- | *Tolumonas lignilytica* | Bacteria; Proteobacteria; Gammaproteobacteria; Aeromonadales; Aeromonadaceae; Tolumonas | methyl-accepting chemotaxis protein | 6 |
| **WP_045013287.1** | 4E-05 | Qc.III | -KAIGGIIGEVNEVATAIAAAVEEQGAATQEITRSTQQAAEGTKNV------- | *Bradyrhizobium sp. LTSP849* | Bacteria; Proteobacteria; Alphaproteobacteria; Rhizobiales; Bradyrhizobiaceae; Bradyrhizobium | methyl-accepting chemotaxis protein | 6 |
| **WP_052741922.1** | 4E-05 | Qa.IV | --KITGTIEQINEITASVAAAVEEQGAATLEISRNVQEAATAT---------- | *Kiloniella litopenaei* | Bacteria; Proteobacteria; Alphaproteobacteria; Kiloniellales; Kiloniellaceae; Kiloniella | methyl-accepting chemotaxis protein | 6 |
| **WP_053531902.1** | 4E-05 | R.IV | -QEINHAVTQLDQMVQQNAELVVQSAAAAGALQGQAGELAQTAGHFR------ | *Cronobacter condimenti* | Bacteria; Proteobacteria; Gammaproteobacteria; Enterobacterales; Enterobacteriaceae; Cronobacter | methyl-accepting chemotaxis protein | 6 |
| **WP_057843664.1** | 4E-05 | Qc.III | -KAIGQTIERISDIATSISAAVEQQRGATQNIAQSVRAAAS------------ | *Bradyrhizobium retamae* | Bacteria; Proteobacteria; Alphaproteobacteria; Rhizobiales; Bradyrhizobiaceae; Bradyrhizobium | methyl-accepting chemotaxis protein | 6 |
| **WP_057855510.1** | 4E-05 | Qc.III | -KAIGQTIERISDIATSISAAVEQQRGATQNIAQSVRAAAS------------ | *Bradyrhizobium valentinum* | Bacteria; Proteobacteria; Alphaproteobacteria; Rhizobiales; Bradyrhizobiaceae; Bradyrhizobium | methyl-accepting chemotaxis protein | 6 |
| **WP_057858101.1** | 4E-05 | Qc.III | -KAIGQTIERISDIATSISAAVEQQRGATQNIAQSVRAAAS------------ | *Bradyrhizobium lablabi* | Bacteria; Proteobacteria; Alphaproteobacteria; Rhizobiales; Bradyrhizobiaceae; Bradyrhizobium | methyl-accepting chemotaxis protein | 6 |
| **WP_061878143.1** | 4E-05 | Qc.III | IGEISGTIARLSEIASAIAAAVEQQGAATQEIARNVQEAARGTQQVSSNVG-- | *Bradyrhizobium liaoningense* | Bacteria; Proteobacteria; Alphaproteobacteria; Rhizobiales; Bradyrhizobiaceae; Bradyrhizobium | PAS domain S-box protein | 6 |
| **WP_065728901.1** | 4E-05 | Qc.III | -KAIGQTIERISDIATSISAAVEQQRGATQNIAQSVRAAAS------------ | *Bradyrhizobium icense* | Bacteria; Proteobacteria; Alphaproteobacteria; Rhizobiales; Bradyrhizobiaceae; Bradyrhizobium | methyl-accepting chemotaxis protein | 6 |
| **WP_092024363.1** | 4E-05 | Qc.III | IGEISGTIARLSEIASAIAAAVEQQGAATQEIARNVQQAAQGTQQVSSNVG-- | *Bradyrhizobium sp. OK095* | Bacteria; Proteobacteria; Alphaproteobacteria; Rhizobiales; Bradyrhizobiaceae; Bradyrhizobium | PAS domain S-box protein | 6 |
| **WP_092322475.1** | 4E-05 | Qb.I | ----TKLIKDLAEKSQANMNAIGNSSEAVTKAGESFNKINQELNTN------- | *Butyrivibrio sp. YAB3001* | Bacteria; Firmicutes; Clostridia; Clostridiales; Lachnospiraceae; Butyrivibrio | methyl-accepting chemotaxis protein | 6 |
| **WP_092787006.1** | 4E-05 | Qa.III.b | IEDIARVIGEINDITQTVSAAVEQQTGATQEISQNAQQASHSTQESSDNIG-- | *Rhodospira trueperi* | Bacteria; Proteobacteria; Alphaproteobacteria; Rhodospirillales; Rhodospirillaceae; Rhodospira | methyl-accepting chemotaxis protein | 6 |
| **WP_100178966.1** | 4E-05 | Qc.III | IGEISGTIARLSEIASAIAAAVEQQGAATQEIARNVQQAAQGTQQVSSNVG-- | *Bradyrhizobium sp. TSA1* | Bacteria; Proteobacteria; Alphaproteobacteria; Rhizobiales; Bradyrhizobiaceae; Bradyrhizobium | PAS domain S-box protein | 6 |
| **WP_101264373.1** | 4E-05 | Qc.III | IQSIGGTIAQINEIAANIASAVEQQGAATQEITRNVQQASAGTSEVTQNI--- | *Thalassospira marina* | Bacteria; Proteobacteria; Alphaproteobacteria; Rhodospirillales; Rhodospirillaceae; Thalassospira | methyl-accepting chemotaxis protein | 6 |
| **WP_101284926.1** | 4E-05 | Qc.III | IQSIGGTIAQINEIAANIASAVEQQGAATQEITRNVQQASAGTSEVTQNI--- | *Thalassospira marina* | Bacteria; Proteobacteria; Alphaproteobacteria; Rhodospirillales; Rhodospirillaceae; Thalassospira | HAMP domain-containing protein | 6 |
| **WP_104738665.1** | 4E-05 | Qa.III | LEQIAQSMEQLVALNASIASATTQQSTVVEDVNRNVTEAAEL----------- | *Pseudomonas oceani* | Bacteria; Proteobacteria; Gammaproteobacteria; Pseudomonadales; Pseudomonadaceae; Pseudomonas | methyl-accepting chemotaxis protein | 6 |
| **WP_108518291.1** | 4E-05 | Qc.III | -KAIGQTIERISDIATSISAAVEQQRGATQNIAQSVRAAAS------------ | *Bradyrhizobium algeriense* | Bacteria; Proteobacteria; Alphaproteobacteria; Rhizobiales; Bradyrhizobiaceae; Bradyrhizobium | methyl-accepting chemotaxis protein | 6 |
| **AWL91826.1** | 4.1E-05 | Qc.III | -QAIGGIIGEVNEVATAIAAAVQEQGAATQEITRSTQYAAQGTKNVS------ | *Bradyrhizobium ottawaense* | Bacteria; Proteobacteria; Alphaproteobacteria; Rhizobiales; Bradyrhizobiaceae; Bradyrhizobium | methyl-accepting chemotaxis protein | 6 |
| **AWM03452.1** | 4.1E-05 | Qc.III | -QAIGGIIGEVNEVATAIAAAVQEQGAATQEITRSTQYAAQGTKNVS------ | *Bradyrhizobium sp. 2 39S1MB* | Bacteria; Proteobacteria; Alphaproteobacteria; Rhizobiales; Bradyrhizobiaceae; Bradyrhizobium | methyl-accepting chemotaxis protein | 6 |
| **GBE44000.1** | 4.1E-05 | Qc.III | IDDIGKVIQQIEDSSTAIAAAMEEQGATTQEVARNVQEAASGTKEVSANI--- | *bacterium BMS3Bbin10* | Bacteria | putative methyl-accepting chemotaxis protein YoaH | 6 |
| **OGR18232.1** | 4.1E-05 | Qc.III | IESISSIINNINEIIATIATAIEEQSAATNEISHSVTQTSEGIGEVNRNIS-- | *Desulfobacterales bacterium GWB2_56_26* | Bacteria; Proteobacteria; Deltaproteobacteria; Desulfobacterales | hypothetical protein A2X81_20495 | 6 |
| **OQY19015.1** | 4.1E-05 | Qc.III | -EQISGGADQISDTSQALSQAATEQASSLEEITSSMADMSSR----------- | *Desulfobacteraceae bacterium 4572_35.1* | Bacteria; Proteobacteria; Deltaproteobacteria; Desulfobacterales; Desulfobacteraceae | hypothetical protein B6I36_05470 partial | 6 |
| **PWC53183.1** | 4.1E-05 | Qa.III | IKGVAQVIGRVNDVATSIASAVEQQGAATREISRNIQQAAQ------------ | *Azospirillum sp. TSH7* | Bacteria; Proteobacteria; Alphaproteobacteria; Rhodospirillales; Rhodospirillaceae; Azospirillum | chemotaxis protein partial | 6 |
| **WP_002714302.1** | 4.1E-05 | Qa.III | IRNIGATITEINEVTTAIAAAVEEQGAATQEIARNIQQAA------------- | *Afipia clevelandensis* | Bacteria; Proteobacteria; Alphaproteobacteria; Rhizobiales; Bradyrhizobiaceae; Afipia | HAMP domain-containing protein | 6 |
| **WP_009735036.1** | 4.1E-05 | Qa.III | IRNIGATITEINEVTTAIAAAVEEQGAATQEIARNIQQAA------------- | *Bradyrhizobiaceae bacterium SG-6C* | Bacteria; Proteobacteria; Alphaproteobacteria; Rhizobiales; Bradyrhizobiaceae | HAMP domain-containing protein | 6 |
| **WP_011084286.1** | 4.1E-05 | Qc.III | -QAIGGIIGEVNEVATAIAAAVQEQGAATQEITRSTQYAAQGTKNVS------ | *Bradyrhizobium diazoefficiens* | Bacteria; Proteobacteria; Alphaproteobacteria; Rhizobiales; Bradyrhizobiaceae; Bradyrhizobium | methyl-accepting chemotaxis protein | 6 |
| **WP_017477087.1** | 4.1E-05 | Qb.III | -----SAADEMNSITAQSSDGLGRQNQEIEQAATAVNEMTAAVDEVAR----- | *Pseudomonas sp. PAMC 26793* | Bacteria; Proteobacteria; Gammaproteobacteria; Pseudomonadales; Pseudomonadaceae; Pseudomonas | methyl-accepting chemotaxis protein | 6 |
| **WP_018646634.1** | 4.1E-05 | Qc.III | -QAIGGIIGEVNEVATAIAAAVQEQGAATQEITRSTQYAAQGTKNVS------ | *Bradyrhizobium japonicum* | Bacteria; Proteobacteria; Alphaproteobacteria; Rhizobiales; Bradyrhizobiaceae; Bradyrhizobium | methyl-accepting chemotaxis protein | 6 |
| **WP_027553926.1** | 4.1E-05 | Qc.III | -QAIGGIIGEVNEVATAIAAAVQEQGAATQEITRSTQYAAQGTKNVS------ | *Bradyrhizobium sp. Cp5.3* | Bacteria; Proteobacteria; Alphaproteobacteria; Rhizobiales; Bradyrhizobiaceae; Bradyrhizobium | methyl-accepting chemotaxis protein | 6 |
| **WP_027561174.1** | 4.1E-05 | Qc.III | -QAIGGIIGEVNEVATAIAAAVQEQGAATQEITRSTQYAAQGTKNVS------ | *Bradyrhizobium* | Bacteria; Proteobacteria; Alphaproteobacteria; Rhizobiales; Bradyrhizobiaceae | MULTISPECIES: methyl-accepting chemotaxis protein | 6 |
| **WP_028142073.1** | 4.1E-05 | Qc.III | -QAIGGIIGEVNEVATAIAAAVQEQGAATQEITRSTQYAAQGTKNVS------ | *Bradyrhizobium* | Bacteria; Proteobacteria; Alphaproteobacteria; Rhizobiales; Bradyrhizobiaceae | MULTISPECIES: methyl-accepting chemotaxis protein | 6 |
| **WP_028173481.1** | 4.1E-05 | Qc.III | -QAIGGIIGEVNEVATAIAAAVQEQGAATQEITRSTQYAAQGTKNVS------ | *Bradyrhizobium* | Bacteria; Proteobacteria; Alphaproteobacteria; Rhizobiales; Bradyrhizobiaceae | MULTISPECIES: methyl-accepting chemotaxis protein | 6 |
| **WP_060909170.1** | 4.1E-05 | Qc.III | -QAIGGIIGEVNEVATAIAAAVQEQGAATQEITRSTQYAAQGTKNVS------ | *Bradyrhizobium diazoefficiens* | Bacteria; Proteobacteria; Alphaproteobacteria; Rhizobiales; Bradyrhizobiaceae; Bradyrhizobium | methyl-accepting chemotaxis protein | 6 |
| **WP_061846232.1** | 4.1E-05 | Qc.III | -QAIGGIIGEVNEVATAIAAAVQEQGAATQEITRSTQYAAQGTKNVS------ | *Bradyrhizobium sp. DOA1* | Bacteria; Proteobacteria; Alphaproteobacteria; Rhizobiales; Bradyrhizobiaceae; Bradyrhizobium | methyl-accepting chemotaxis protein | 6 |
| **WP_063200812.1** | 4.1E-05 | Qc.III | -QAIGGIIGEVNEVATAIAAAVQEQGAATQEITRSTQYAAQGTKNVS------ | *Bradyrhizobium sp. AT1* | Bacteria; Proteobacteria; Alphaproteobacteria; Rhizobiales; Bradyrhizobiaceae; Bradyrhizobium | methyl-accepting chemotaxis protein | 6 |
| **WP_063677266.1** | 4.1E-05 | Qc.III | -QAIGGIIGEVNEVATAIAAAVQEQGAATQEITRSTQYAAQGTKNVS------ | *Bradyrhizobium neotropicale* | Bacteria; Proteobacteria; Alphaproteobacteria; Rhizobiales; Bradyrhizobiaceae; Bradyrhizobium | methyl-accepting chemotaxis protein | 6 |
| **WP_063694222.1** | 4.1E-05 | Qc.III | -QAIGGIIGEVNEVATAIAAAVQEQGAATQEITRSTQYAAQGTKNVS------ | *Bradyrhizobium stylosanthis* | Bacteria; Proteobacteria; Alphaproteobacteria; Rhizobiales; Bradyrhizobiaceae; Bradyrhizobium | methyl-accepting chemotaxis protein | 6 |
| **WP_063699028.1** | 4.1E-05 | Qc.III | -QAIGGIIGEVNEVATAIAAAVQEQGAATQEITRSTQYAAQGTKNVS------ | *Bradyrhizobium sp. BR 10245* | Bacteria; Proteobacteria; Alphaproteobacteria; Rhizobiales; Bradyrhizobiaceae; Bradyrhizobium | methyl-accepting chemotaxis protein | 6 |
| **WP_074120267.1** | 4.1E-05 | Qc.III | -QAIGGIIGEVNEVATAIAAAVQEQGAATQEITRSTQYAAQGTKNVS------ | *Bradyrhizobium sp. AS23.2* | Bacteria; Proteobacteria; Alphaproteobacteria; Rhizobiales; Bradyrhizobiaceae; Bradyrhizobium | methyl-accepting chemotaxis protein | 6 |
| **WP_085556863.1** | 4.1E-05 | Qa.III | -------IETVNNVASSISAAIEEQGAATRNISANVNEAAQGVAQVSSSL--- | *Azospirillum lipoferum* | Bacteria; Proteobacteria; Alphaproteobacteria; Rhodospirillales; Rhodospirillaceae; Azospirillum | methyl-accepting chemotaxis protein | 6 |
| **WP_091881293.1** | 4.1E-05 | Qc.III | -QAIGGIIGEVNEVATAIAAAVQEQGAATQEITRSTQYAAQGTKNVS------ | *Bradyrhizobium sp. Rc2d* | Bacteria; Proteobacteria; Alphaproteobacteria; Rhizobiales; Bradyrhizobiaceae; Bradyrhizobium | methyl-accepting chemotaxis protein | 6 |
| **WP_092032199.1** | 4.1E-05 | Qc.III | -QAIGGIIGEVNEVATAIAAAVQEQGAATQEITRSTQYAAQGTKNVS------ | *Bradyrhizobium sp. OK095* | Bacteria; Proteobacteria; Alphaproteobacteria; Rhizobiales; Bradyrhizobiaceae; Bradyrhizobium | methyl-accepting chemotaxis protein | 6 |
| **WP_092186722.1** | 4.1E-05 | Qc.III | -QAIGGIIGEVNEVATAIAAAVQEQGAATQEITRSTQYAAQGTKNVS------ | *Bradyrhizobium sp. cf659* | Bacteria; Proteobacteria; Alphaproteobacteria; Rhizobiales; Bradyrhizobiaceae; Bradyrhizobium | methyl-accepting chemotaxis protein | 6 |
| **WP_092227296.1** | 4.1E-05 | Qc.III | -QAIGGIIGEVNEVATAIAAAVQEQGAATQEITRSTQYAAQGTKNVS------ | *Bradyrhizobium sp. Gha* | Bacteria; Proteobacteria; Alphaproteobacteria; Rhizobiales; Bradyrhizobiaceae; Bradyrhizobium | methyl-accepting chemotaxis protein | 6 |
| **WP_097321478.1** | 4.1E-05 | Qa.III | --AIVQEVADIESAMGTLSGIADDQHEIVERLNATVDATMASI---------- | *Actinoplanes atraurantiacus* | Bacteria; Actinobacteria; Micromonosporales; Micromonosporaceae; Actinoplanes | chemotaxis protein | 6 |
| **WP_097659953.1** | 4.1E-05 | Qc.III | -QAIGGIIGEVNEVATAIAAAVQEQGAATQEITRSTQYAAQGTKNVS------ | *Bradyrhizobium sp. Y36* | Bacteria; Proteobacteria; Alphaproteobacteria; Rhizobiales; Bradyrhizobiaceae; Bradyrhizobium | methyl-accepting chemotaxis protein | 6 |
| **WP_097668323.1** | 4.1E-05 | Qc.III | -QAIGGIIGEVNEVATAIAAAVQEQGAATQEITRSTQYAAQGTKNVS------ | *Bradyrhizobium ottawaense* | Bacteria; Proteobacteria; Alphaproteobacteria; Rhizobiales; Bradyrhizobiaceae; Bradyrhizobium | methyl-accepting chemotaxis protein | 6 |
| **WP_100235106.1** | 4.1E-05 | Qc.III | -QAIGGIIGEVNEVATAIAAAVQEQGAATQEITRSTQYAAQGTKNVS------ | *Bradyrhizobium sp. INPA54B* | Bacteria; Proteobacteria; Alphaproteobacteria; Rhizobiales; Bradyrhizobiaceae; Bradyrhizobium | methyl-accepting chemotaxis protein | 6 |
| **WP_107991411.1** | 4.1E-05 | Qa.IV | ---ITKTIDQVSEIATAIASAVEEQTIATSEISVNVQTAAKNTAD-------- | *Breoghania corrubedonensis* | Bacteria; Proteobacteria; Alphaproteobacteria; Rhizobiales; Cohaesibacteraceae; Breoghania | chemotaxis protein | 6 |
| **WP_109071830.1** | 4.1E-05 | Qa.III | IKTIGGTIGEINGIATTIAAAVEEQGAATGEIARNVQQAA------------- | *Azospirillum sp. TSH58* | Bacteria; Proteobacteria; Alphaproteobacteria; Rhodospirillales; Rhodospirillaceae; Azospirillum | HAMP domain-containing protein | 6 |
| **WP_109329592.1** | 4.1E-05 | Qa.III | IEEIAGTVQEVHQTSSSIAAAIQEQHAAISEISRNVQQAAAGTEEIN------ | *Azospirillum sp. CFH 70021* | Bacteria; Proteobacteria; Alphaproteobacteria; Rhodospirillales; Rhodospirillaceae; Azospirillum | methyl-accepting chemotaxis protein | 6 |
| **ANC92792.1** | 4.2E-05 | Qa.III | -QEVVRVIGHINEVATSIASAVEEQGAATREISRNIQQAAQGTQTV------- | *Azospirillum humicireducens* | Bacteria; Proteobacteria; Alphaproteobacteria; Rhodospirillales; Rhodospirillaceae; Azospirillum | HAMP domain-containing protein | 6 |
| **EJO59758.1** | 4.2E-05 | Qa.III | --EIVAAVERVNAIMSDISTAFDEQSSGIEQVNATVTQMEQTMQ--------- | *Burkholderia multivorans ATCC BAA-247* | Bacteria; Proteobacteria; Betaproteobacteria; Burkholderiales; Burkholderiaceae; Burkholderia; Burkholderia cepacia complex | methyl-accepting chemotaxis protein signaling domain protein partial | 6 |
| **EPN67892.1** | 4.2E-05 | Qa.III | ---ISQALRNLNSLNASIASATLQQTHVVDDINQNVTQAAQLSQ--------- | *Pseudomonas syringae pv. actinidiae ICMP 19079* | Bacteria; Proteobacteria; Gammaproteobacteria; Pseudomonadales; Pseudomonadaceae; Pseudomonas; Pseudomonas syringae | methyl-accepting chemotaxis protein | 6 |
| **OCS02507.1** | 4.2E-05 | Qa.IV | --EIEANVNILSQGINEMSQSINEQTEAINQINEAVATVDEQTKQ-NV----- | *Campylobacter fetus subsp. testudinum* | Bacteria; Proteobacteria; Epsilonproteobacteria; Campylobacterales; Campylobacteraceae; Campylobacter | chemotaxis protein partial | 6 |
| **OJX78327.1** | 4.2E-05 | Qc.III | -GDIGRTINRMNEIAGSIAAAVEEQGAATNEIARSVAEAAA------------ | *Magnetospirillum sp. 64-120* | Bacteria; Proteobacteria; Alphaproteobacteria; Rhodospirillales; Rhodospirillaceae; Magnetospirillum | chemotaxis protein | 6 |
| **OUR75690.1** | 4.2E-05 | Qa.III | IQGIAGTIQKVDEIAAAIASAVEEQGAATQEISSSVQKAAA------------ | *Alphaproteobacteria bacterium 46_93_T64* | Bacteria; Proteobacteria; Alphaproteobacteria | hypothetical protein A9Q83_16960 | 6 |
| **OUR79103.1** | 4.2E-05 | Qa.IV | IKDVTATVDQLSVISTSLSDTINEQTAATAEISQNAQNAAQGTSRVNDNIS-- | *Alphaproteobacteria bacterium 46_93_T64* | Bacteria; Proteobacteria; Alphaproteobacteria | hypothetical protein A9Q83_05845 | 6 |

| | | | | | | | | |
|---|---|---|---|---|---|---|---|---|
|  | **OYD86295.1** | 4.2E-05 | Qa.III | IKTIGGTIGEINGIATTIAAAVEEQGAATGEIARNVQQAA------------- | *Azospirillum brasilense* | Bacteria; Proteobacteria; Alphaproteobacteria; Rhodospirillales; Rhodospirillaceae; Azospirillum | hypothetical protein CHT98_01650 | 6 |
|  | **PID30142.1** | 4.2E-05 | Qa.III | ------IISELNEINTTIATSVEEQSATVNEIAGSIAEAAQNSEQVNR----- | *Candidatus Cloacimonetes bacterium* | Bacteria; Candidatus Cloacimonetes | hypothetical protein CSB55_00005 | 6 |
|  | **WP_007090325.1** | 4.2E-05 | Qa.III.b | IDSIGGTIAEISEIAATIASAVEQQGAATEEITRNVQQAS------------- | *Thalassospira xiamenensis* | Bacteria; Proteobacteria; Alphaproteobacteria; Rhodospirillales; Rhodospirillaceae; Thalassospira | methyl-accepting chemotaxis protein | 6 |
|  | **WP_011502875.1** | 4.2E-05 | Qa.III | IREIGATISKMSEIASTIASAVEEQGAATQEISRNVQQAAQGTQQVSSS---- | *Rhodopseudomonas palustris* | Bacteria; Proteobacteria; Alphaproteobacteria; Rhizobiales; Bradyrhizobiaceae; Rhodopseudomonas | HAMP domain-containing protein | 6 |
|  | **WP_012978433.1** | 4.2E-05 | Qa.III | -QEVVRVIGHINEVATSIASAVEEQGAATREISRNIQQAAQGTQTV------- | *Azospirillum lipoferum* | Bacteria; Proteobacteria; Alphaproteobacteria; Rhodospirillales; Rhodospirillaceae; Azospirillum | HAMP domain-containing protein | 6 |
|  | **WP_014250007.1** | 4.2E-05 | Qa.III | -QEVVRVIGHINEVATSIASAVEEQGAATREISRNIQQAAQGTQTV------- | *Azospirillum lipoferum* | Bacteria; Proteobacteria; Alphaproteobacteria; Rhodospirillales; Rhodospirillaceae; Azospirillum | HAMP domain-containing protein | 6 |
|  | **WP_046273930.1** | 4.2E-05 | Qa.III | MDEIVVSVKRVSDLMADISAAAQQQSEGIEQINQVVEHIDESTQ--------- | *Stenotrophomonas maltophilia* | Bacteria; Proteobacteria; Gammaproteobacteria; Xanthomonadales; Xanthomonadaceae; Stenotrophomonas; Stenotrophomonas maltophilia group | methyl-accepting chemotaxis protein | 6 |
|  | **WP_080114889.1** | 4.2E-05 | Qa.III | MDEIVVSVKRVSDLMADISAAAQQQSEGIEQINQVVEHIDESTQ--------- | *Stenotrophomonas maltophilia* | Bacteria; Proteobacteria; Gammaproteobacteria; Xanthomonadales; Xanthomonadaceae; Stenotrophomonas; Stenotrophomonas maltophilia group | methyl-accepting chemotaxis protein | 6 |
|  | **WP_080223331.1** | 4.2E-05 | Qa.III | ---ISQALRNLNSLNASIASATLQQTHVVDDINQNVTQAAQLSQ--------- | *Pseudomonas syringae* | Bacteria; Proteobacteria; Gammaproteobacteria; Pseudomonadales; Pseudomonadaceae; Pseudomonas | methyl-accepting chemotaxis protein | 6 |
|  | **WP_082822720.1** | 4.2E-05 | Qa.III.b | IDSIGGTIAEISEIAATIASAVEQQGAATEEITRNVQQAS------------- | *Thalassospira* | Bacteria; Proteobacteria; Alphaproteobacteria; Rhodospirillales; Rhodospirillaceae | MULTISPECIES: methyl-accepting chemotaxis protein | 6 |
| | **WP_084005685.1** | 4.2E-05 | Qc.III | IQSIGGTINTMNEITSAIAAAVEEQGAATQEIARNVEQAATGTNEVSSNIA-- | *Magnetovibrio blakemorei* | Bacteria; Proteobacteria; Alphaproteobacteria; Rhodospirillales; Rhodospirillaceae; Magnetovibrio | methyl-accepting chemotaxis protein | 6 |
|  | **WP_085556602.1** | 4.2E-05 | Qc.III | -RSISGTISRINEISTSIATAMEEQGAATTEIARNVQQASV------------ | *Azospirillum lipoferum* | Bacteria; Proteobacteria; Alphaproteobacteria; Rhodospirillales; Rhodospirillaceae; Azospirillum | chemotaxis protein | 6 |
|  | **WP_088435410.1** | 4.2E-05 | Qa.III | MDEIVVSVKRVSDLMADISAAAQQQSEGIEQINQVVEHIDESTQ--------- | *Stenotrophomonas maltophilia* | Bacteria; Proteobacteria; Gammaproteobacteria; Xanthomonadales; Xanthomonadaceae; Stenotrophomonas; Stenotrophomonas maltophilia group | methyl-accepting chemotaxis protein | 6 |
|  | **WP_090068248.1** | 4.2E-05 | Qa.III | IQGIAETMEQVNQYTASISQAVEEQGSATYEISENVAQAAS------------ | *Cohaesibacter marisflavi* | Bacteria; Proteobacteria; Alphaproteobacteria; Rhizobiales; Cohaesibacteraceae | methyl-accepting chemotaxis protein | 6 |
|  | **WP_092685709.1** | 4.2E-05 | Qa.III | IREIGATISKMSEIASTIASAVEEQGAATQEISRNVQQAAQGTQQVSSS---- | *Rhodopseudomonas pseudopalustris* | Bacteria; Proteobacteria; Alphaproteobacteria; Rhizobiales; Bradyrhizobiaceae; Rhodopseudomonas | HAMP domain-containing protein | 6 |
|  | **WP_094303053.1** | 4.2E-05 | Qa.III | IRGIGATISSINDISSSIAAAIEEQGAATQEISRNVQQAAQSTQE-------- | *Azospirillum brasilense* | Bacteria; Proteobacteria; Alphaproteobacteria; Rhodospirillales; Rhodospirillaceae; Azospirillum | methyl-accepting chemotaxis protein | 6 |
|  | **WP_109073437.1** | 4.2E-05 | Qa.III | -QEVVRVIGHINEVATSIASAVEEQGAATREISRNIQQAAQGTQTV------- | *Azospirillum* | Bacteria; Proteobacteria; Alphaproteobacteria; Rhodospirillales; Rhodospirillaceae | MULTISPECIES: HAMP domain-containing protein | 6 |
|  | **WP_109332668.1** | 4.2E-05 | Qa.III | IKGVAQVISRVNDVATSIASAVEEQGAATREISRNIQQAA------------- | *Azospirillum sp. CFH 70021* | Bacteria; Proteobacteria; Alphaproteobacteria; Rhodospirillales; Rhodospirillaceae; Azospirillum | HAMP domain-containing protein | 6 |
| | **WP_109516314.1** | 4.2E-05 | Qc.III | --GIGSTVERTNGIATAIAAAVEEQGAATQEIARAAAQVAEATGTVSSRI--- | *Roseomonas aestuarii* | Bacteria; Proteobacteria; Alphaproteobacteria; Rhodospirillales; Acetobacteraceae; Roseomonas | methyl-accepting chemotaxis protein | 6 |
|  | **EGH29184.1** | 4.3E-05 | Qa.III | ---ISQALRNLNSLNASIASATLQQTHVVDDINQNVTQAAQLSQ--------- | *Pseudomonas syringae pv. japonica str. M301072* | Bacteria; Proteobacteria; Gammaproteobacteria; Pseudomonadales; Pseudomonadaceae; Pseudomonas; Pseudomonas syringae | histidine kinase HAMP region: chemotaxis sensory transducer | 6 |
|  | **OJX68467.1** | 4.3E-05 | Qa.IV | IRSITDSIAGINEISSAVAAAVEQQEAATAEISRNVAETSA------------ | *Magnetospirillum sp. 64-120* | Bacteria; Proteobacteria; Alphaproteobacteria; Rhodospirillales; Rhodospirillaceae; Magnetospirillum | hypothetical protein BGO92_18740 | 6 |
| | **OLS28528.1** | 4.3E-05 | Qc.III | -EEINASTEEVAQTSQSMAQGASSQSAMISLVVDKLNFTSDTLEKVLKRIQ-- | *Candidatus Heimdallarchaeota archaeon LC_2* | Archaea; Asgard group; Candidatus Heimdallarchaeota | putative methyl-accepting chemotaxis protein YoaH | 6 |
|  | **PWC13817.1** | 4.3E-05 | Qa.III | INDIVQEVRQVSTMINEISHAAGEQTSALGLINTSIAQIEE------------ | *Brenneria sp. CFCC 11842* | Bacteria; Proteobacteria; Gammaproteobacteria; Enterobacterales; Pectobacteriaceae; Brenneria | hypothetical protein DDT56_14680 | 6 |
|  | **PYS77146.1** | 4.3E-05 | SNAP.b | -----------NRDLVQVLGELEEKRERLEQVNRRLEESNRELNEANAKLREL | *Acidobacteria bacterium* | Bacteria; Acidobacteria | hypothetical protein DMF66_10890 | 6 |
|  | **SIT23595.1** | 4.3E-05 | Qa.III.b | IDSIGGTIAEISEIAATIASAVEQQGAATEEITRNVQQAS------------- | *Thalassospira xiamenensis M-5 = DSM 17429* | Bacteria; Proteobacteria; Alphaproteobacteria; Rhodospirillales; Rhodospirillaceae; Thalassospira | methyl-accepting chemotaxis protein | 6 |
|  | **WP_013899745.1** | 4.3E-05 | Qb.I | LERLKSHLRDTVEQYEASTEELKASNEELQAMNEELRSATEELETS------- | *Ramlibacter tataouinensis* | Bacteria; Proteobacteria; Betaproteobacteria; Burkholderiales; Comamonadaceae; Ramlibacter | PAS domain S-box protein | 6 |
| | **WP_020399623.1** | 4.3E-05 | Qc.III.b | LASINGSVTNLHASVSEIAGSIDEQSGVTREISNNMLTAAEAVGEINRNLKDI | *Kordiimonas gwangyangensis* | Bacteria; Proteobacteria; Alphaproteobacteria; Kordiimonadales; Kordiimonadaceae; Kordiimonas | methyl-accepting chemotaxis protein | 6 |
| | **WP_057863364.1** | 4.3E-05 | Qc.III | IQAISGTIEKLSEISSTIAAAVEEQGAATQEISRNVQQAATGTQQVSSNITD- | *Bradyrhizobium lablabi* | Bacteria; Proteobacteria; Alphaproteobacteria; Rhizobiales; Bradyrhizobiaceae; Bradyrhizobium | HAMP domain-containing protein | 6 |
|  | **WP_063633981.1** | 4.3E-05 | Qa.IV | IQSIAATIGEVNDIVGTIAQAVDGQNAATMEIARNVQQAA------------- | *Azospirillum humicireducens* | Bacteria; Proteobacteria; Alphaproteobacteria; Rhodospirillales; Rhodospirillaceae; Azospirillum | HAMP domain-containing protein | 6 |
| | **WP_074116914.1** | 4.3E-05 | Qa.IV | -KEIGATIGQISTISTSIASAVEQQGAATQEIARSVQTVAQGTQTAATDI--- | *Bradyrhizobium sp. AS23.2* | Bacteria; Proteobacteria; Alphaproteobacteria; Rhizobiales; Bradyrhizobiaceae; Bradyrhizobium | methyl-accepting chemotaxis protein | 6 |
|  | **WP_077772310.1** | 4.3E-05 | Qc.III | -DEISGNLSNIKNLSSDMTEAIEQLGPIVSNLKENVDSLSGAIR--------- | *Salinivibrio sharmensis* | Bacteria; Proteobacteria; Gammaproteobacteria; Vibrionales; Vibrionaceae; Salinivibrio | methyl-accepting chemotaxis protein | 6 |
| | **WP_085909318.1** | 4.3E-05 | Qc.III | -SEIAVTVSNVKEVSAAIASAVEEQGAATQEIARNIEQASGGTSEVNSNIQQ- | *Kiloniella majae* | Bacteria; Proteobacteria; Alphaproteobacteria; Kiloniellales; Kiloniellaceae; Kiloniella | methyl-accepting chemotaxis protein | 6 |
|  | **WP_094301548.1** | 4.3E-05 | Qa.III | IKTIGGTIGEINGIATTIAAAVEEQGAATGEIARNVQQAA------------- | *Azospirillum brasilense* | Bacteria; Proteobacteria; Alphaproteobacteria; Rhodospirillales; Rhodospirillaceae; Azospirillum | HAMP domain-containing protein | 6 |
|  | **WP_101533980.1** | 4.3E-05 | Qa.III.b | IRGIGDVIIEINEISAEIAASVDQQGSATQEIAHNVSEAAKGTQEVS------ | *Cohaesibacter sp. H1304* | Bacteria; Proteobacteria; Alphaproteobacteria; Rhizobiales; Cohaesibacteraceae | methyl-accepting chemotaxis protein | 6 |
|  | **ACS85868.1** | 4.4E-05 | R.IV | --QIAQAVTELDSTTQQNAALVEESSTAANQLEEQAQRLEQSISAF------- | *Dickeya paradisiaca Ech703* | Bacteria; Proteobacteria; Gammaproteobacteria; Enterobacterales; Pectobacteriaceae; Dickeya | methyl-accepting chemotaxis sensory transducer | 6 |
|  | **AMB44846.1** | 4.4E-05 | Qc.III.c | -DGISASIRSINETATAIASAIEEQGAATREIVQAIAQASS------------ | *Methylobacterium sp. AMS5* | Bacteria; Proteobacteria; Alphaproteobacteria; Rhizobiales; Methylobacteriaceae; Methylobacterium | chemotaxis sensory transducer | 6 |
| | **KIZ34570.1** | 4.4E-05 | Qc.III | IKEIGVTIGRMSEISSTIASAVEEQGAATQEISRNVQQAAQGTTQVSSNISD- | *Rhodopseudomonas palustris* | Bacteria; Proteobacteria; Alphaproteobacteria; Rhizobiales; Bradyrhizobiaceae; Rhodopseudomonas | chemotaxis protein partial | 6 |
|  | **KQN57975.1** | 4.4E-05 | Qa.III | IREMVGNAGDVERLINDIARAAHEQSDGIRQINQAIGQIDTTTQ--------- | *Erwinia sp. Leaf53* | Bacteria; Proteobacteria; Gammaproteobacteria; Enterobacterales; Erwiniaceae; Erwinia | hypothetical protein ASF13_04045 | 6 |
|  | **OQX26458.1** | 4.4E-05 | Qa.IV | -------ISEINTIMNTIASAVEEQTASTNEISKSIAETASNSNSVSKNVQE- | *Desulfobacteraceae bacterium IS3* | Bacteria; Proteobacteria; Deltaproteobacteria; Desulfobacterales; Desulfobacteraceae | hypothetical protein BWK80_10300 | 6 |
| | **PCI41041.1** | 4.4E-05 | Qc.III | -KGIGSTISRIDEASTAIASAVEQQGAATQEIARNVEQAATGTNEVTSNITE- | *Rhodospirillaceae bacterium* | Bacteria; Proteobacteria; Alphaproteobacteria; Rhodospirillales; Rhodospirillaceae | chemotaxis protein | 6 |
|  | **RDD61115.1** | 4.4E-05 | Qc.III | IDGISNTVKHINESVAAISSAVQEQEATTGEIARNIEDVTKASETVA------ | *Rhodospirillaceae bacterium* | Bacteria; Proteobacteria; Alphaproteobacteria; Rhodospirillales; Rhodospirillaceae | methyl-accepting chemotaxis protein | 6 |
| | **SBV99334.1** | 4.4E-05 | Qc.III | -QGITDVIGRIRENSSGIASAVEEQGAATQEIARNVEQAAAGTQQVSGNI--- | *uncultured Alphaproteobacteria bacterium* | Bacteria; Proteobacteria; Alphaproteobacteria; environmental samples | Chemotaxis sensory transducer | 6 |
|  | **WP_038210123.1** | 4.4E-05 | Qc.III.c | ------SIVRINDLGLEIATAAEQQHSVTDEVNRNMTTIQEMVNDLT------ | *Vibrio tubiashii* | Bacteria; Proteobacteria; Gammaproteobacteria; Vibrionales; Vibrionaceae; Vibrio; Vibrio oreintalis group | HAMP domain-containing protein | 6 |
|  | **WP_058076774.1** | 4.4E-05 | Qc.III | LDQMADSTNRIHDLSAQIAAAAEEQSAVSGEINQNMVAVRD------------ | *Pseudomonas sp. TTU2014-066ASC* | Bacteria; Proteobacteria; Gammaproteobacteria; Pseudomonadales; Pseudomonadaceae; Pseudomonas | methyl-accepting chemotaxis protein | 6 |
|  | **WP_058192798.1** | 4.4E-05 | Qa.IV | IEQIAKTIAAVNEISGSIASTVVQQTAATSEISRNAGEAA------------- | *Methylobacterium sp. GXS13* | Bacteria; Proteobacteria; Alphaproteobacteria; Rhizobiales; Methylobacteriaceae; Methylobacterium | methyl-accepting chemotaxis protein | 6 |
|  | **WP_060770332.1** | 4.4E-05 | Qc.III.c | -DGISASIRSINETATAIASAIEEQGAATREIVQAIAQASS------------ | *Methylobacterium sp. AMS5* | Bacteria; Proteobacteria; Alphaproteobacteria; Rhizobiales; Methylobacteriaceae; Methylobacterium | methyl-accepting chemotaxis protein | 6 |
|  | **WP_065985099.1** | 4.4E-05 | Qc.III | LDQMADSTNRIHDLSAQIAAAAEEQSAVSGEINQNMVAVRD------------ | *Pseudomonas xanthomarina* | Bacteria; Proteobacteria; Gammaproteobacteria; Pseudomonadales; Pseudomonadaceae; Pseudomonas | methyl-accepting chemotaxis protein | 6 |
|  | **WP_109154959.1** | 4.4E-05 | Qc.III | IQKVSAVIERLSTIATTISAAVDQQNAATAEISRSVQQAASGTGEVSS----- | *Azospirillum sp. TSO5* | Bacteria; Proteobacteria; Alphaproteobacteria; Rhodospirillales; Rhodospirillaceae; Azospirillum | methyl-accepting chemotaxis protein | 6 |
| | **OAZ08819.1** | 4.5E-05 | Qc.III | IERVTETISNINEIATAIASAVEEQGAATAEISSNVQQAAAGTNEVSSSI--- | *Thalassospira tepidiphila MCCC 1A03514* | Bacteria; Proteobacteria; Alphaproteobacteria; Rhodospirillales; Rhodospirillaceae; Thalassospira | hypothetical protein TH4_15735 | 6 |
| | **ONH86977.1** | 4.5E-05 | Qc.III | IERVTETISNINEIATAIASAVEEQGAATAEISSNVQQAAAGTNEVSSSI--- | *Thalassospira sp. MCCC 1A02803* | Bacteria; Proteobacteria; Alphaproteobacteria; Rhodospirillales; Rhodospirillaceae; Thalassospira | methyl-accepting chemotaxis protein | 6 |
|  | **PHJ03294.1** | 4.5E-05 | Qa.IV | --EIEANVNILSQGINEMSQSINEQTEAINQINEAVANVDEQTKQ-------- | *Campylobacter fetus subsp. venerealis* | Bacteria; Proteobacteria; Epsilonproteobacteria; Campylobacterales; Campylobacteraceae; Campylobacter | chemotaxis protein partial | 6 |
| | **WP_011662748.1** | 4.5E-05 | Qc.III | IKEISDTIGKMSEIASTIASAVEEQGAATQEISRNVQQAAAGTQQVSANIA-- | *Rhodopseudomonas palustris* | Bacteria; Proteobacteria; Alphaproteobacteria; Rhizobiales; Bradyrhizobiaceae; Rhodopseudomonas | methyl-accepting chemotaxis protein | 6 |
| | **WP_013653786.1** | 4.5E-05 | Qc.III | --SVNETIEKMNAISATISAAVEQQEAATGEISRNVTEASTGTTFVTRSVG-- | *Polymorphum gilvum* | Bacteria; Proteobacteria; Alphaproteobacteria; Polymorphum | methyl-accepting chemotaxis sensory transducer | 6 |
|  | **WP_028350919.1** | 4.5E-05 | Qc.III | IKEISGTIGRVSEIAAAIAAAIEEQGAATQEIARNVQQAAVGS---------- | *Bradyrhizobium elkanii* | Bacteria; Proteobacteria; Alphaproteobacteria; Rhizobiales; Bradyrhizobiaceae; Bradyrhizobium | methyl-accepting chemotaxis protein | 6 |
| | **WP_035673283.1** | 4.5E-05 | Qc.III | IGEISGTIARLSEIASAIAAAVQQQGAATQEIARNVQEAARGTRQVSSNVG-- | *Bradyrhizobium liaoningense* | Bacteria; Proteobacteria; Alphaproteobacteria; Rhizobiales; Bradyrhizobiaceae; Bradyrhizobium | PAS domain S-box protein | 6 |
|  | **WP_048493160.1** | 4.5E-05 | Qa.III | IEGITGVIDKINDISSTMASAVEEQSATTNEIGRNINDAA------------- | *Nitrospina sp. SCGC_AAA799_C22* | Bacteria; Nitrospinae/Tectomicrobia group; Nitrospinae; Nitrospinia; Nitrospinales; Nitrospinaceae; Nitrospina | chemotaxis protein partial | 6 |
|  | **WP_051357078.1** | 4.5E-05 | Qa.IV | ---IVSTISDIRTSSSAIASAVEEQGAATREIAVNTQQAAQGTGTVNHN---- | *Azorhizobium doebereinerae* | Bacteria; Proteobacteria; Alphaproteobacteria; Rhizobiales; Xanthobacteraceae; Azorhizobium | methyl-accepting chemotaxis protein | 6 |
|  | **WP_057848028.1** | 4.5E-05 | Qc.III | IKEISGTIGRVSEIAAAIAAAIEEQGAATQEIARNVQQAAVGS---------- | *Bradyrhizobium retamae* | Bacteria; Proteobacteria; Alphaproteobacteria; Rhizobiales; Bradyrhizobiaceae; Bradyrhizobium | methyl-accepting chemotaxis protein | 6 |

| | | | | | | | |
|---|---|---|---|---|---|---|---|
| **WP_063087763.1** | 4.5E-05 | Qc.III | IERVTETISNINEIATAIASAVEEQGAATAEISSNVQQAAAGTNEVSSSI--- | *Thalassospira* | Bacteria; Proteobacteria; Alphaproteobacteria; Rhodospirillales; Rhodospirillaceae | MULTISPECIES: methyl-accepting chemotaxis protein | 6 |
| **WP_065730656.1** | 4.5E-05 | Qc.III | IKEISGTIGRVSEIAAAIAAAIEEQGAATQEIARNVQQAAVGS---------- | *Bradyrhizobium icense* | Bacteria; Proteobacteria; Alphaproteobacteria; Rhizobiales; Bradyrhizobiaceae; Bradyrhizobium | methyl-accepting chemotaxis protein | 6 |
| **WP_074129900.1** | 4.5E-05 | Qc.III.c | --AISQVIIQISEISTVIASAVEEQAAATKEVTANINGTTKAIDDA------- | *Bradyrhizobium sp. NAS96.2* | Bacteria; Proteobacteria; Alphaproteobacteria; Rhizobiales; Bradyrhizobiaceae; Bradyrhizobium | PAS domain-containing protein | 6 |
| **WP_084302312.1** | 4.5E-05 | Qc.III | --DIAGIIDKLQEISSSISAAIEEQSIVTNDISRSITNISSDMREASR----- | *Bradyrhizobium sp. WSM2254* | Bacteria; Proteobacteria; Alphaproteobacteria; Rhizobiales; Bradyrhizobiaceae; Bradyrhizobium | PAS domain S-box protein | 6 |
| **WP_085900173.1** | 4.5E-05 | Qc.III | IDSISETIQQIDQIAASVAAAVEEQTAATQEIARNVDQAAAGTQEVSSNIS-- | *Kiloniella majae* | Bacteria; Proteobacteria; Alphaproteobacteria; Kiloniellales; Kiloniellaceae; Kiloniella | methyl-accepting chemotaxis protein | 6 |
| **WP_094248197.1** | 4.5E-05 | Qc.III | -EQVASSAGELATGANEISGEVGDMTDSIRDIELKVSAYAESSRLMNERL--- | *Paenibacillus sp. RUD330* | Bacteria; Firmicutes; Bacilli; Bacillales; Paenibacillaceae; Paenibacillus | methyl-accepting chemotaxis protein | 6 |
| **WP_108522166.1** | 4.5E-05 | Qc.III | IKEIGDTIGRMSEIASTIAAAVEEQGAATQEISRNVQQAAHGTQQVSSNI--- | *Bradyrhizobium algeriense* | Bacteria; Proteobacteria; Alphaproteobacteria; Rhizobiales; Bradyrhizobiaceae; Bradyrhizobium | HAMP domain-containing protein | 6 |
| **EGH75136.1** | 4.6E-05 | Qa.III | ---ISQALRNLNSLNASIASATLQQTHVVDDINQNVTQAAQLSQ--------- | *Pseudomonas syringae pv. aptata str. DSM 50252* | Bacteria; Proteobacteria; Gammaproteobacteria; Pseudomonadales; Pseudomonadaceae; Pseudomonas; Pseudomonas syringae | histidine kinase HAMP region: chemotaxis sensory transducer partial | 6 |
| **KEO58507.1** | 4.6E-05 | Qa.III.b | IDSIGGTIAEISEIAATIASAVEQQGAATEEITRNVQQAS------------- | *Thalassospira permensis NBRC 106175* | Bacteria; Proteobacteria; Alphaproteobacteria; Rhodospirillales; Rhodospirillaceae; Thalassospira | methyl-accepting chemotaxis protein | 6 |
| **OEU67079.1** | 4.6E-05 | Qa.III.b | -------------LYNALSSMVTSLVELIGTAEEKTQEAKKQTEAANIALEEA | *Desulfovibrio sp. S3730MH75* | Bacteria; Proteobacteria; Deltaproteobacteria; Desulfovibrionales; Desulfovibrionaceae; Desulfovibrio | chemotaxis protein | 6 |
| **PID72283.1** | 4.6E-05 | SNAP.c | -GEISDIIGQVNDIVITITTAVEEQSAATNEIAGNIAQASQGIQDVNDAISR- | *Desulfobulbus propionicus* | Bacteria; Proteobacteria; Deltaproteobacteria; Desulfobacterales; Desulfobulbaceae; Desulfobulbus | chemotaxis protein | 6 |
| **PWC83583.1** | 4.6E-05 | Qc.III | IQKVSAVIERLSTIATTISAAVDQQNAATAEISRSVQQAASGTGEVSS----- | *Azospirillum sp. TSO5* | Bacteria; Proteobacteria; Alphaproteobacteria; Rhodospirillales; Rhodospirillaceae; Azospirillum | chemotaxis protein | 6 |
| **WP_008143582.1** | 4.6E-05 | Qc.III | -SEISGTIARLSEISSAIAAAVEQQGAATQEIARNVQQAASGTQQVSSNVG-- | *Bradyrhizobium sp. YR681* | Bacteria; Proteobacteria; Alphaproteobacteria; Rhizobiales; Bradyrhizobiaceae; Bradyrhizobium | PAS domain S-box protein | 6 |
| **WP_024511737.1** | 4.6E-05 | Qa.IV | ---IGATIAQISTIASSIAGAVEQQGAATQEIARNVQNVAAGTQDAAANIV-- | *Bradyrhizobium sp. ARR65* | Bacteria; Proteobacteria; Alphaproteobacteria; Rhizobiales; Bradyrhizobiaceae; Bradyrhizobium | HAMP domain-containing protein | 6 |
| **WP_025675281.1** | 4.6E-05 | Qa.III.b | -VDIANMVTEINQMTSHIATATQQQTQVTTEISENINDISQ------------ | *Salinivibrio socompensis* | Bacteria; Proteobacteria; Gammaproteobacteria; Vibrionales; Vibrionaceae; Salinivibrio | methyl-accepting chemotaxis protein | 6 |
| **WP_027714053.1** | 4.6E-05 | Qa.IV | IRSIVKSIEEVKSMTDDMVTATTRQRQNTQSIEAAVQSVSDMVGQ-------- | *Desulfuromonas sp. TF* | Bacteria; Proteobacteria; Deltaproteobacteria; Desulfuromonadales; Desulfuromonadaceae; Desulfuromonas | GAF domain-containing protein | 6 |
| **WP_079564124.1** | 4.6E-05 | Qc.III | IKEISGTIERLSEISSTIAAAVEEQGAATQEISRNVQQAAHGTQQVSS----- | *Bradyrhizobium erythrophlei* | Bacteria; Proteobacteria; Alphaproteobacteria; Rhizobiales; Bradyrhizobiaceae; Bradyrhizobium | methyl-accepting chemotaxis protein | 6 |
| **WP_082013011.1** | 4.6E-05 | Qa.III | IQGIASTIGEVNQIAGAIAAAVEEQGSATQEIARNVQQAAAGTQEVSVSV--- | *Belnapia sp. F-4-1* | Bacteria; Proteobacteria; Alphaproteobacteria; Rhodospirillales; Acetobacteraceae; Belnapia | methyl-accepting chemotaxis protein | 6 |
| **WP_082502331.1** | 4.6E-05 | Qa.III | IREMVGNAGDVERLINDIARAAHEQSDGIRQINQAIGQIDTTTQ--------- | *Erwinia* | Bacteria; Proteobacteria; Gammaproteobacteria; Enterobacterales; Erwiniaceae | MULTISPECIES: hypothetical protein | 6 |
| **WP_082832738.1** | 4.6E-05 | Qa.III.b | IDSIGSTIAEISEIAATIASAVEQQGAATEEITRNVQQAS------------- | *Thalassospira xiamenensis* | Bacteria; Proteobacteria; Alphaproteobacteria; Rhodospirillales; Rhodospirillaceae; Thalassospira | methyl-accepting chemotaxis protein | 6 |
| **WP_083997131.1** | 4.6E-05 | Qc.III | IERVTETISNINEIATAIASAVEEQGAATAEISSNVQQAAAGTNEVSSSI--- | *Thalassospira tepidiphila* | Bacteria; Proteobacteria; Alphaproteobacteria; Rhodospirillales; Rhodospirillaceae; Thalassospira | methyl-accepting chemotaxis protein | 6 |
| **WP_101300454.1** | 4.6E-05 | Qc.III | IDSISATIGRINEFCNAIAAAVEQQGAATQEIADSVSRAASGTQEVSQN---- | *Thalassospira lohafexi* | Bacteria; Proteobacteria; Alphaproteobacteria; Rhodospirillales; Rhodospirillaceae; Thalassospira | methyl-accepting chemotaxis protein | 6 |
| **WP_109106428.1** | 4.6E-05 | Qa.III | -TGIGGTIGNINDITTAIAAAIEEQGAATNEISRNVQQAAA------------ | *Azospirillum sp. TSO35-2* | Bacteria; Proteobacteria; Alphaproteobacteria; Rhodospirillales; Rhodospirillaceae; Azospirillum | methyl-accepting chemotaxis protein | 6 |
| **WP_109156759.1** | 4.6E-05 | Qc.III | IQKVSAVIERLSTIATTISAAVDQQNAATAEISRSVQQAASGTGEVSS----- | *Azospirillum* | Bacteria; Proteobacteria; Alphaproteobacteria; Rhodospirillales; Rhodospirillaceae | MULTISPECIES: methyl-accepting chemotaxis protein | 6 |
| **WP_114110853.1** | 4.6E-05 | Qa.III.b | IDSIGSTIAEISEIAATIASAVEQQGAATEEITRNVQQAS------------- | *Thalassospira xiamenensis* | Bacteria; Proteobacteria; Alphaproteobacteria; Rhodospirillales; Rhodospirillaceae; Thalassospira | HAMP domain-containing protein | 6 |
| **WP_114121208.1** | 4.6E-05 | Qa.III.b | IDSIGSTIAEISEIAATIASAVEQQGAATEEITRNVQQAS------------- | *Thalassospira xianhensis* | Bacteria; Proteobacteria; Alphaproteobacteria; Rhodospirillales; Rhodospirillaceae; Thalassospira | HAMP domain-containing protein | 6 |
| **EIC02347.1** | 4.7E-05 | Qb.II | --DTSSAITEIIANIESVHKQIRLQNESVESSSENVTGISDSISVLDHLIE-- | *Treponema saccharophilum DSM 2985* | Bacteria; Spirochaetes; Spirochaetales; Spirochaetaceae; Treponema | methyl-accepting chemotaxis sensory transducer | 6 |
| **OJX73815.1** | 4.7E-05 | Qc.III | IEGMGRTIGDINQIASAIAAAVEEQNAATQEISRNVQLAAEGTNGV------- | *Magnetospirillum sp. 64-120* | Bacteria; Proteobacteria; Alphaproteobacteria; Rhodospirillales; Rhodospirillaceae; Magnetospirillum | hypothetical protein BGO92_04680 | 6 |
| **OUR77905.1** | 4.7E-05 | Qc.III | --NIGNVIKTIREATVGISSAVEEQSAATNEISRNVQEASSGTNEVSSKI--- | *Alphaproteobacteria bacterium 46_93_T64* | Bacteria; Proteobacteria; Alphaproteobacteria | hypothetical protein A9Q83_09810 | 6 |
| **OUX61713.1** | 4.7E-05 | Qa.III.b | IDGITRTIGKVNEIASAIASAVEEQGAATREIAHNVSQAA------------- | *Afipia sp. TMED4* | Bacteria; Proteobacteria; Alphaproteobacteria; Rhizobiales; Bradyrhizobiaceae; Afipia | methyl-accepting chemotaxis protein | 6 |
| **PLX68495.1** | 4.7E-05 | SNAP.b | TRNMMEKIAESVSESASTVRKLGESSEQIGQIISVIDDIADQT---------- | *Denitrovibrio sp.* | Bacteria; Deferribacteres; Deferribacterales; Deferribacteraceae; Denitrovibrio | chemotaxis protein | 6 |
| **WP_006021090.1** | 4.7E-05 | Qa.III.b | IDGITRTIGKVNEIASAIASAVEEQGAATREIAHNVSQAA------------- | *Afipia broomeae* | Bacteria; Proteobacteria; Alphaproteobacteria; Rhizobiales; Bradyrhizobiaceae; Afipia | methyl-accepting chemotaxis protein | 6 |
| **WP_007434995.1** | 4.7E-05 | Qa.IV | ---IATTIDEVNSITGSIAAAVEEQGAATGEIARTVQQTARATEA-------- | *Acetobacteraceae bacterium AT-5844* | Bacteria; Proteobacteria; Alphaproteobacteria; Rhodospirillales; Acetobacteraceae; unclassified Acetobacteraceae | HAMP domain-containing protein | 6 |
| **WP_011503800.1** | 4.7E-05 | Qa.III | -EGITTIIGRVNEIATAIASAVREQGTATREIAHNVAEAARST---------- | *Rhodopseudomonas palustris* | Bacteria; Proteobacteria; Alphaproteobacteria; Rhizobiales; Bradyrhizobiaceae; Rhodopseudomonas | HAMP domain-containing protein | 6 |
| **WP_052741756.1** | 4.7E-05 | Qc.III | IEGISKIINQVNEIASGISAAVEEQGAATQEIARNTQEAARGTQEVS------ | *Kiloniella litopenaei* | Bacteria; Proteobacteria; Alphaproteobacteria; Kiloniellales; Kiloniellaceae; Kiloniella | methyl-accepting chemotaxis protein | 6 |
| **WP_059717740.1** | 4.7E-05 | Qa.III | --ELNAAVGNVAQITTEISAAVEEQSRAIDEVNQAVASMDQNTQQ-------- | *Burkholderia vietnamiensis* | Bacteria; Proteobacteria; Betaproteobacteria; Burkholderiales; Burkholderiaceae; Burkholderia; Burkholderia cepacia complex | methyl-accepting chemotaxis protein | 6 |
| **WP_083530807.1** | 4.7E-05 | Qc.III.c | -DGISASIRSINETATAIASAIEEQGAATREIVQAIAQASS------------ | *Methylobacterium sp. AMS5* | Bacteria; Proteobacteria; Alphaproteobacteria; Rhizobiales; Methylobacteriaceae; Methylobacterium | methyl-accepting chemotaxis protein | 6 |
| **WP_085899877.1** | 4.7E-05 | Qc.III | IEGISKIINQVNEIASGISAAVEEQGAATQEIARNTQEAARGTQEVS------ | *Kiloniella majae* | Bacteria; Proteobacteria; Alphaproteobacteria; Kiloniellales; Kiloniellaceae; Kiloniella | methyl-accepting chemotaxis protein | 6 |
| **WP_085905290.1** | 4.7E-05 | Qc.III | IEGISKIINQVNEIASGISAAVEEQGAATQEIARNTQEAARGTQEVS------ | *Kiloniella majae* | Bacteria; Proteobacteria; Alphaproteobacteria; Kiloniellales; Kiloniellaceae; Kiloniella | methyl-accepting chemotaxis protein | 6 |
| **WP_085909409.1** | 4.7E-05 | Qc.III | IEGISKIINQVNEIASGISAAVEEQGAATQEIARNTQEAARGTQEVS------ | *Kiloniella majae* | Bacteria; Proteobacteria; Alphaproteobacteria; Kiloniellales; Kiloniellaceae; Kiloniella | methyl-accepting chemotaxis protein | 6 |
| **WP_092682730.1** | 4.7E-05 | Qa.III | -EGITTIIGRVNEIATAIASAVREQGTATREIAHNVAEAARST---------- | *Rhodopseudomonas pseudopalustris* | Bacteria; Proteobacteria; Alphaproteobacteria; Rhizobiales; Bradyrhizobiaceae; Rhodopseudomonas | methyl-accepting chemotaxis protein | 6 |
| **WP_108125053.1** | 4.7E-05 | Qa.III.a | MREVVTAIKQVNDIMTEIASASAEQSTGLDEIDKAVSQMDEMT---------- | *Saccharospirillum mangrovi* | Bacteria; Proteobacteria; Gammaproteobacteria; Oceanospirillales; Saccharospirillaceae; Saccharospirillum | methyl-accepting chemotaxis protein | 6 |
| **WP_110350961.1** | 4.7E-05 | Qa.III | IRAIVSTITNLSEISMSVSSAVTEQAAVTQEISNNMQTAARSVEA-------- | *Methylobacterium sp. B4* | Bacteria; Proteobacteria; Alphaproteobacteria; Rhizobiales; Methylobacteriaceae; Methylobacterium | PAS domain S-box protein | 6 |
| **EPM99645.1** | 4.8E-05 | Qa.III | ---ISQALRNLNSLNASIASATLQQTHVVDDINQNVTQAAQLSQ--------- | *Pseudomonas syringae pv. actinidiae ICMP 19070* | Bacteria; Proteobacteria; Gammaproteobacteria; Pseudomonadales; Pseudomonadaceae; Pseudomonas; Pseudomonas syringae | methyl-accepting chemotaxis protein partial | 6 |
| **OHE65118.1** | 4.8E-05 | SNAP.c | IAQISANIRSVKRQTVSQAASVEETNLAMERITRRIKELDVHIEAQSESLDR- | *Treponema sp. GWC1_61_84* | Bacteria; Spirochaetes; Spirochaetales; Spirochaetaceae; Treponema | hypothetical protein A2001_17000 | 6 |
| **PIQ01429.1** | 4.8E-05 | Qa.III | --SINQSVEQITQLTLDMSAAVEEQGAVAEHINQQIIDIADSAEITKIS---- | *Shewanella sp. CG18_big_fil_WC_8_21_14_2_50_42_11* | Bacteria; Proteobacteria; Gammaproteobacteria; Alteromonadales; Shewanellaceae; Shewanella | chemotaxis protein | 6 |
| **RCK40030.1** | 4.8E-05 | Qa.III | IEKITETISNINEITGAIAAAVEQQGAATAEISGNVQQAA------------- | *Thalassospira xiamenensis* | Bacteria; Proteobacteria; Alphaproteobacteria; Rhodospirillales; Rhodospirillaceae; Thalassospira | hypothetical protein TH24_11745 | 6 |
| **WP_002717709.1** | 4.8E-05 | Qc.III | ---IGTTIERLSEISSAIAAAVEQQGAATQEIARNVQHAAEGTQQVSSNITE- | *Afipia felis* | Bacteria; Proteobacteria; Alphaproteobacteria; Rhizobiales; Bradyrhizobiaceae; Afipia | HAMP domain-containing protein | 6 |
| **WP_027553925.1** | 4.8E-05 | Qc.III | -QAIGGIIGEVNEVATAIAAAVQEQGAATQEITRSTQYAAQGTRNVS------ | *Bradyrhizobium sp. Cp5.3* | Bacteria; Proteobacteria; Alphaproteobacteria; Rhizobiales; Bradyrhizobiaceae; Bradyrhizobium | methyl-accepting chemotaxis protein | 6 |
| **WP_039689049.1** | 4.8E-05 | Qc.I | ------MAERLSSMYSNLESQVRDRTTQLSAANEELERQRQHVERVNDRLK-- | *Coriobacteriaceae bacterium 68-1-3* | Bacteria; Actinobacteria; Coriobacteriia; Coriobacteriales; Coriobacteriaceae | DUF3365 domain-containing protein | 6 |
| **WP_051228415.1** | 4.8E-05 | Qa.III | IVGITSIIEQLNAIASSIAAAVDQQGAATNEIARNVSQASSGTHQVSQNIV-- | *Pleomorphomonas oryzae* | Bacteria; Proteobacteria; Alphaproteobacteria; Rhizobiales; Methylocystaceae; Pleomorphomonas | methyl-accepting chemotaxis protein | 6 |
| **WP_056582350.1** | 4.8E-05 | R.IV | IEQINQAIAQMDQVTQQNAALVEEASAAAQSLQEQAEGLVRAAQVFRTGAG-- | *Variovorax sp. Root473* | Bacteria; Proteobacteria; Betaproteobacteria; Burkholderiales; Comamonadaceae; Variovorax | PAS domain S-box protein | 6 |
| **WP_063300994.1** | 4.8E-05 | Qc.III.c | ---IGETIEQINSLSTGINEAVGDQKQATENISRDLNSAHLNVSQ-------- | *Pseudovibrio sp. WM33* | Bacteria; Proteobacteria; Alphaproteobacteria; Rhodobacterales; Rhodobacteraceae; Pseudovibrio | methyl-accepting chemotaxis protein | 6 |
| **WP_063677265.1** | 4.8E-05 | Qc.III | -QAIGGIIGEVNEVATAIAAAVQEQGAATQEITRSTQYAAQGTRNVS------ | *Bradyrhizobium neotropicale* | Bacteria; Proteobacteria; Alphaproteobacteria; Rhizobiales; Bradyrhizobiaceae; Bradyrhizobium | methyl-accepting chemotaxis protein | 6 |
| **WP_089288586.1** | 4.8E-05 | Qa.IV | IRTITGTINDINEAMATIAAAVEQQNATTLEISRNVQQAA------------- | *Azospirillum* | Bacteria; Proteobacteria; Alphaproteobacteria; Rhodospirillales; Rhodospirillaceae | MULTISPECIES: methyl-accepting chemotaxis protein | 6 |
| **WP_101287884.1** | 4.8E-05 | Qa.III | IVGITSIIEQLNAIASSIAAAVDQQGAATNEIARNVSQASSGTHQVSQNIV-- | *Pleomorphomonas diazotrophica* | Bacteria; Proteobacteria; Alphaproteobacteria; Rhizobiales; Methylocystaceae; Pleomorphomonas | methyl-accepting chemotaxis protein | 6 |
| **KZD04520.1** | 4.9E-05 | Qa.III | IEKITETISNINEITGAIAAAVEQQGAATAEISGNVQQAA------------- | *Thalassospira xiamenensis* | Bacteria; Proteobacteria; Alphaproteobacteria; Rhodospirillales; Rhodospirillaceae; Thalassospira | hypothetical protein AUP40_14915 | 6 |
| **OHY98020.1** | 4.9E-05 | Qa.III | IEKITETISNINEITGAIAAAVEQQGAATAEISGNVQQAA------------- | *Thalassospira sp. MIT1004* | Bacteria; Proteobacteria; Alphaproteobacteria; Rhodospirillales; Rhodospirillaceae; Thalassospira | hypothetical protein BC440_16550 | 6 |
| **OUR79660.1** | 4.9E-05 | Qc.III | -SKINAVISEIRQITVTISSAVEEQSAATGNISRNVQEASAGTKDVAGKIDRV | *Alphaproteobacteria bacterium 46_93_T64* | Bacteria; Proteobacteria; Alphaproteobacteria | hypothetical protein A9Q83_04520 | 6 |

| | | | | | | | | |
|---|---|---|---|---|---|---|---|---|
|  | **PHR99624.1** | 4.9E-05 | SNAP.b | TQKSISSLSEISEQVAGTIRAISEQTERISQAAQVIEDIAEQT---------- | *Oceanobacter sp.* | Bacteria; Proteobacteria; Gammaproteobacteria; Oceanospirillales; Oceanobacter | chemotaxis protein | 6 |
|  | **PIT01031.1** | 4.9E-05 | Qc.III | IQEIGGTIERLSEVSAAIAAAVEQQGAATQEISRNVQQASVGTQEV------- | *Bradyrhizobium sp. TSA1* | Bacteria; Proteobacteria; Alphaproteobacteria; Rhizobiales; Bradyrhizobiaceae; Bradyrhizobium | chemotaxis protein | 6 |
|  | **PXX32862.1** | 4.9E-05 | Qa.III | IEKITETISNINEITGAIAAAVEQQGAATAEISGNVQQAA------------ | *Thalassospira sp. 11-3* | Bacteria; Proteobacteria; Alphaproteobacteria; Rhodospirillales; Rhodospirillaceae; Thalassospira | methyl-accepting chemotaxis sensory transducer with Cache sensor | 6 |
|  | **WP_007090447.1** | 4.9E-05 | Qa.III | IEKITETISNINEITGAIAAAVEQQGAATAEISGNVQQAA------------ | *Thalassospira xiamenensis* | Bacteria; Proteobacteria; Alphaproteobacteria; Rhodospirillales; Rhodospirillaceae; Thalassospira | methyl-accepting chemotaxis protein | 6 |
|  | **WP_007977488.1** | 4.9E-05 | Qc.II | --ELAESVDQASGSAHTIEERVDEQTSLLSTVATEMENFSASM---------- | *Haladaptatus paucihalophilus* | Archaea; Euryarchaeota; Halobacteria; Halobacteriales; Halobacteriaceae; Haladaptatus | PAS domain-containing protein | 6 |
|  | **WP_011158730.1** | 4.9E-05 | Qa.IV | IRDISATIEKLSEISSTIASAVEEQGAATLEISRNVQQAAQ----------- | *Rhodopseudomonas palustris* | Bacteria; Proteobacteria; Alphaproteobacteria; Rhizobiales; Bradyrhizobiaceae; Rhodopseudomonas | methyl-accepting chemotaxis protein | 6 |
|  | **WP_011383629.1** | 4.9E-05 | Qc.III | --SIAATIGRINEIAGAIAAAVEEQGAATAEIARNVQQASAG----------- | *Magnetospirillum magneticum* | Bacteria; Proteobacteria; Alphaproteobacteria; Rhodospirillales; Rhodospirillaceae; Magnetospirillum | methyl-accepting chemotaxis protein | 6 |
|  | **WP_021023273.1** | 4.9E-05 | Qc.III | -DEISSNLSNIKTLSSDMHEAIDQLGPIVSDLKDNVNALNG----------- | *Salinivibrio costicola* | Bacteria; Proteobacteria; Gammaproteobacteria; Vibrionales; Vibrionaceae; Salinivibrio | methyl-accepting chemotaxis protein | 6 |
|  | **WP_036105284.1** | 4.9E-05 | R.IV | IQEINQAVTRLDQMVQQNAELVVESAAAADALQMQAGELAETASLFR------ | *Mangrovibacter* | Bacteria; Proteobacteria; Gammaproteobacteria; Enterobacterales; Enterobacteriaceae | MULTISPECIES: methyl-accepting chemotaxis protein | 6 |
|  | **WP_051682374.1** | 4.9E-05 | Qa.III | IEKITETISNINEITGAIAAAVEQQGAATAEISGNVQQAA------------ | *Thalassospira* | Bacteria; Proteobacteria; Alphaproteobacteria; Rhodospirillales; Rhodospirillaceae | MULTISPECIES: methyl-accepting chemotaxis protein | 6 |
| | **WP_057855460.1** | 4.9E-05 | Qc.III | IKEIGDTIGRMSEIASTIAAAVEEQGAATQEISRNVQQAAHGTQQVSSNI--- | *Bradyrhizobium valentinum* | Bacteria; Proteobacteria; Alphaproteobacteria; Rhizobiales; Bradyrhizobiaceae; Bradyrhizobium | HAMP domain-containing protein | 6 |
|  | **WP_064593986.1** | 4.9E-05 | R.IV | IQEINQAVTRLDQMVQQNAELVVESAAAADALQMQAGELAETASLFR------ | *Mangrovibacter phragmitis* | Bacteria; Proteobacteria; Gammaproteobacteria; Enterobacterales; Enterobacteriaceae; Mangrovibacter | methyl-accepting chemotaxis protein | 6 |
| | **WP_069957826.1** | 4.9E-05 | Qc.III | -GSIGGTIAKMNEIASAIAAAVEEQGAATQEIARNVEQAAAGTNEVSTNID-- | *Magnetovibrio blakemorei* | Bacteria; Proteobacteria; Alphaproteobacteria; Rhodospirillales; Rhodospirillaceae; Magnetovibrio | methyl-accepting chemotaxis protein | 6 |
|  | **WP_072824078.1** | 4.9E-05 | Qa.IV | IKAIGQTIERISEITGAISAAVEEQGAATQNIAQGVQAAA------------ | *Bradyrhizobium erythrophlei* | Bacteria; Proteobacteria; Alphaproteobacteria; Rhizobiales; Bradyrhizobiaceae; Bradyrhizobium | methyl-accepting chemotaxis protein | 6 |
|  | **WP_085553856.1** | 4.9E-05 | Qa.III | MQGIGETIAQVNRITATIAAAVEEQGAATEEIVRNIQQAA------------ | *Azospirillum lipoferum* | Bacteria; Proteobacteria; Alphaproteobacteria; Rhodospirillales; Rhodospirillaceae; Azospirillum | methyl-accepting chemotaxis protein | 6 |
|  | **WP_085641885.1** | 4.9E-05 | Qa.III | IEKITETISNINEITGAIAAAVEQQGAATAEISGNVQQAA------------ | *Thalassospira sp. MCCC 1A03138* | Bacteria; Proteobacteria; Alphaproteobacteria; Rhodospirillales; Rhodospirillaceae; Thalassospira | methyl-accepting chemotaxis protein | 6 |
|  | **WP_100176249.1** | 4.9E-05 | Qc.III | IQEIGGTIERLSEVSAAIAAAVEQQGAATQEISRNVQQASVGTQEV------- | *Bradyrhizobium sp. TSA1* | Bacteria; Proteobacteria; Alphaproteobacteria; Rhizobiales; Bradyrhizobiaceae; Bradyrhizobium | HAMP domain-containing protein | 6 |
|  | **WP_107345032.1** | 4.9E-05 | Qa.IV | IRDISATIEKLSEISSTIASAVEEQGAATLEISRNVQQAAQ----------- | *Rhodopseudomonas palustris* | Bacteria; Proteobacteria; Alphaproteobacteria; Rhizobiales; Bradyrhizobiaceae; Rhodopseudomonas | HAMP domain-containing protein | 6 |
|  | **WP_107346727.1** | 4.9E-05 | Qa.III | IKGIGGTIEKMSEIASTIASAVEEQGAATREISRNIQQAAE----------- | *Rhodopseudomonas palustris* | Bacteria; Proteobacteria; Alphaproteobacteria; Rhizobiales; Bradyrhizobiaceae; Rhodopseudomonas | methyl-accepting chemotaxis protein | 6 |
|  | **WP_107356092.1** | 4.9E-05 | Qa.IV | IRDISATIEKLSEISSTIASAVEEQGAATLEISRNVQQAAQ----------- | *Rhodopseudomonas palustris* | Bacteria; Proteobacteria; Alphaproteobacteria; Rhizobiales; Bradyrhizobiaceae; Rhodopseudomonas | HAMP domain-containing protein | 6 |
|  | **WP_107357611.1** | 4.9E-05 | Qa.III | IKGIGGTIEKMSEIASTIASAVEEQGAATREISRNIQQAAE----------- | *Rhodopseudomonas palustris* | Bacteria; Proteobacteria; Alphaproteobacteria; Rhizobiales; Bradyrhizobiaceae; Rhodopseudomonas | methyl-accepting chemotaxis protein | 6 |
|  | **WP_114094582.1** | 4.9E-05 | Qa.III | IEKITETISNINEITGAIAAAVEQQGAATAEISGNVQQAA------------ | *Thalassospira xiamenensis* | Bacteria; Proteobacteria; Alphaproteobacteria; Rhodospirillales; Rhodospirillaceae; Thalassospira | HAMP domain-containing protein | 6 |
|  | **BAO30980.1** | 5E-05 | Qb.I | --RALAVMRETVNNMNSIASLIRESGSSVEQLDESSQKIGGIVQ--------- | *Sulfuritalea hydrogenivorans sk43H* | Bacteria; Proteobacteria; Betaproteobacteria; Nitrosomonadales; Sterolibacteriaceae; Sulfuritalea | methyl-accepting chemotaxis sensory transducer | 6 |
| | **OEJ68377.1** | 5E-05 | Qc.III | -GSIGGTIGKMSEIASAIAAAVEEQGAATQEIARNVEQASAGTGEVTSNIA-- | *Magnetovibrio blakemorei* | Bacteria; Proteobacteria; Alphaproteobacteria; Rhodospirillales; Rhodospirillaceae; Magnetovibrio | hypothetical protein BEN30_06350 | 6 |
| | **PLX70003.1** | 5E-05 | R.IV | TQSATSEISEVVDGMHKDIQTLLENTNSVqanmGAVSDNTVELKDKIQTFSKNAN[illegible] | *Denitrovibrio sp.* | Bacteria; Deferribacteres; Deferribacterales; Deferribacteraceae; Denitrovibrio | hypothetical protein C0603_03435 | 6 |
|  | **PSQ80633.1** | 5E-05 | Qb.I | LAETKRQLQNTIEESETTTEELESSNEELLSMNEELQSKNEEIETSK------ | *Bacteroidetes bacterium QS_1_63_11* | Bacteria; Bacteroidetes | hypothetical protein BRD41_04690 | 6 |
| | **WP_011474810.1** | 5E-05 | Qc.III | IKEIGDTIGKMSEIAATIASAVEEQGAATQEISRNVQQAALGTQQVSSNIT-- | *Rhodopseudomonas palustris* | Bacteria; Proteobacteria; Alphaproteobacteria; Rhizobiales; Bradyrhizobiaceae; Rhodopseudomonas | methyl-accepting chemotaxis protein | 6 |
|  | **WP_028880316.1** | 5E-05 | Qa.IV | ---IHSSITRVQETATTISAAIEQQRAATENIARNATQTADDV---------- | *Terasakiella pusilla* | Bacteria; Proteobacteria; Alphaproteobacteria; Rhizobiales; Methylocystaceae; Terasakiella | hypothetical protein | 6 |
|  | **WP_041100644.1** | 5E-05 | Qb.I | --RALAVMRETVNNMNSIASLIRESGSSVEQLDESSQKIGGIVQ--------- | *Sulfuritalea hydrogenivorans* | Bacteria; Proteobacteria; Betaproteobacteria; Nitrosomonadales; Sterolibacteriaceae; Sulfuritalea | HAMP domain-containing protein | 6 |
|  | **WP_054161958.1** | 5E-05 | Qa.III | IEGITTIISRVNEIATAIASAVREQGTATREIAHNVAEAAR----------- | *Rhodopseudomonas sp. AAP120* | Bacteria; Proteobacteria; Alphaproteobacteria; Rhizobiales; Bradyrhizobiaceae; Rhodopseudomonas | methyl-accepting chemotaxis protein | 6 |
| | **WP_074792184.1** | 5E-05 | Qa.IV | --EITTTVGEVNDGIEEVARATDEQAELTTDIASFVDDATNRVDEVTKEID-- | *Haloferax larsenii* | Archaea; Euryarchaeota; Halobacteria; Haloferacales; Haloferacaceae; Haloferax | globin-coupled sensor protein | 6 |
|  | **WP_085587050.1** | 5E-05 | Qa.IV | ---------EVSRLSSDISVSVEQQTAAINEISGSAQEVSTHMAGIAEDIG-- | *Thalassospira sp. MCCC 1A01428* | Bacteria; Proteobacteria; Alphaproteobacteria; Rhodospirillales; Rhodospirillaceae; Thalassospira | chemotaxis protein | 6 |
|  | **WP_097052390.1** | 5E-05 | Qa.III.b | IDSIGSTIAEISEIAATIASAVEQQGAATEEITRNVQQAS------------ | *Thalassospira xiamenensis* | Bacteria; Proteobacteria; Alphaproteobacteria; Rhodospirillales; Rhodospirillaceae; Thalassospira | methyl-accepting chemotaxis protein | 6 |
|  | **WP_107355580.1** | 5E-05 | Qa.III | IREIGATISKMSEIASTIASAVEEQGAATQEISRNVQQAAQGTQQVS------ | *Rhodopseudomonas palustris* | Bacteria; Proteobacteria; Alphaproteobacteria; Rhizobiales; Bradyrhizobiaceae; Rhodopseudomonas | methyl-accepting chemotaxis protein | 6 |
|  | **WP_109865066.1** | 5E-05 | Qa.III | -QEVVRVIGHINEVATSIASAVEEQGAATREISRNIQQAAQGTQ--------- | *Azospirillum sp. TSH100* | Bacteria; Proteobacteria; Alphaproteobacteria; Rhodospirillales; Rhodospirillaceae; Azospirillum | HAMP domain-containing protein | 6 |
|  | **AWJ88859.1** | 5.1E-05 | Qa.III | IRGIGATISSINDISSSIAAAIEEQGAATQEISRNVQQAAQSTQE-------- | *Azospirillum brasilense Sp245* | Bacteria; Proteobacteria; Alphaproteobacteria; Rhodospirillales; Rhodospirillaceae; Azospirillum | methyl-accepting chemotaxis protein | 6 |
|  | **EGH55533.1** | 5.1E-05 | Qa.III | ---ISQALRNLNSLNASIASATLQQTHVVDDINQNVTQAAQLSQ--------- | *Pseudomonas syringae Cit 7* | Bacteria; Proteobacteria; Gammaproteobacteria; Pseudomonadales; Pseudomonadaceae; Pseudomonas; Pseudomonas syringae | histidine kinase HAMP region: chemotaxis sensory transducer partial | 6 |
|  | **EPM70572.1** | 5.1E-05 | Qa.III | ---ISQALRNLNSLNASIASATLQQTHVVDDINQNVTQAAQLSQ--------- | *Pseudomonas syringae pv. actinidiae ICMP 18886* | Bacteria; Proteobacteria; Gammaproteobacteria; Pseudomonadales; Pseudomonadaceae; Pseudomonas; Pseudomonas syringae | methyl-accepting chemotaxis protein partial | 6 |
|  | **PIM53848.1** | 5.1E-05 | R.IV | -QQLRGTFTDVERMVQQNAALVEEAAASAASLNDQAERMNELASAFR------ | *Mitsuaria sp. HWN-4* | Bacteria; Proteobacteria; Betaproteobacteria; Burkholderiales; Mitsuaria | hypothetical protein CS062_07895 partial | 6 |
|  | **PWC34976.1** | 5.1E-05 | Qa.IV | VQEIARTIAEIDSISSMIAAAVEEQAAATHEISRNVNQTA------------ | *Azospirillum sp. TSO22-1* | Bacteria; Proteobacteria; Alphaproteobacteria; Rhodospirillales; Rhodospirillaceae; Azospirillum | chemotaxis protein partial | 6 |
|  | **RCK53315.1** | 5.1E-05 | Qa.III.b | IEGIGGAIDEVSQLSSDIASSVQQQTAAISEISTSAQEVSTHMQGISSD---- | *Thalassospira xiamenensis* | Bacteria; Proteobacteria; Alphaproteobacteria; Rhodospirillales; Rhodospirillaceae; Thalassospira | chemotaxis protein | 6 |
|  | **SDS29559.1** | 5.1E-05 | Qa.IV | ---ITATIEEVSTIATTIGSAIEEQGAATAEIARNVTQTAQATQD-------- | *Bradyrhizobium canariense* | Bacteria; Proteobacteria; Alphaproteobacteria; Rhizobiales; Bradyrhizobiaceae; Bradyrhizobium | methyl-accepting chemotaxis sensory transducer with Cache sensor | 6 |
|  | **WP_007092358.1** | 5.1E-05 | Qa.III.b | IEGIGGAIDEVSQLSSDIASSVQQQTAAISEISTSAQEVSTHMQGISSD---- | *Thalassospira xiamenensis* | Bacteria; Proteobacteria; Alphaproteobacteria; Rhodospirillales; Rhodospirillaceae; Thalassospira | methyl-accepting chemotaxis protein | 6 |
| | **WP_007539399.1** | 5.1E-05 | Qa.IV | --EITTTVGEVNDGIEEVARATDEQAELTTDIASFVDDATNRVDEVTKEID-- | *Haloferax larsenii* | Archaea; Euryarchaeota; Halobacteria; Haloferacales; Haloferacaceae; Haloferax | globin-coupled sensor protein | 6 |
|  | **WP_035675811.1** | 5.1E-05 | Qa.III | IRGIGATISSINDISSSIAAAIEEQGAATQEISRNVQQAAQSTQE-------- | *Azospirillum* | Bacteria; Proteobacteria; Alphaproteobacteria; Rhodospirillales; Rhodospirillaceae | MULTISPECIES: methyl-accepting chemotaxis protein | 6 |
|  | **WP_037987127.1** | 5.1E-05 | Qa.III.b | IEGIGGAIDEVSQLSSDIASSVQQQTAAISEISTSAQEVSTHMQGISSD---- | *Thalassospira permensis* | Bacteria; Proteobacteria; Alphaproteobacteria; Rhodospirillales; Rhodospirillaceae; Thalassospira | chemotaxis protein | 6 |
|  | **WP_038529592.1** | 5.1E-05 | Qa.III | IRGIGATISSINDISSSIAAAIEEQGAATQEISRNVQQAAQSTQE-------- | *Azospirillum brasilense* | Bacteria; Proteobacteria; Alphaproteobacteria; Rhodospirillales; Rhodospirillaceae; Azospirillum | methyl-accepting chemotaxis protein | 6 |
|  | **WP_063092064.1** | 5.1E-05 | Qa.III.b | IEGIGGAIDEVSQLSSDIASSVQQQTAAISEISTSAQEVSTHMQGISSD---- | *Thalassospira xiamenensis* | Bacteria; Proteobacteria; Alphaproteobacteria; Rhodospirillales; Rhodospirillaceae; Thalassospira | chemotaxis protein | 6 |
|  | **WP_065701838.1** | 5.1E-05 | Qa.III.b | IEGIGGAIDEVSQLSSDIASSVQQQTAAISEISTSAQEVSTHMQGISSD---- | *Thalassospira* | Bacteria; Proteobacteria; Alphaproteobacteria; Rhodospirillales; Rhodospirillaceae | MULTISPECIES: chemotaxis protein | 6 |
|  | **WP_069519740.1** | 5.1E-05 | Qa.III | LEQIAQSLRNLTGLNASIASATLQQSHVVEDINQNVTQAHGLARQS------- | *Pseudomonas* | Bacteria; Proteobacteria; Gammaproteobacteria; Pseudomonadales; Pseudomonadaceae | MULTISPECIES: methyl-accepting chemotaxis protein | 6 |
|  | **WP_076400607.1** | 5.1E-05 | SNAP.c | -GTIINDISTLNGIAQDIANAVDQQSSAVTEIERSSQVATSAVQQ-------- | *Insolitispirillum peregrinum* | Bacteria; Proteobacteria; Alphaproteobacteria; Rhodospirillales; Rhodospirillaceae; Insolitispirillum | hypothetical protein | 6 |
|  | **WP_079565038.1** | 5.1E-05 | Qc.III | -KGIGSIIGEVNEVATAIAAAVQEQGAATQEITRSTQYAAQGTKNVS------ | *Bradyrhizobium erythrophlei* | Bacteria; Proteobacteria; Alphaproteobacteria; Rhizobiales; Bradyrhizobiaceae; Bradyrhizobium | methyl-accepting chemotaxis protein | 6 |
| | **WP_080802465.1** | 5.1E-05 | Qc.III | -ETVESVVREMNSITLSIHDLMEEQTRMMVSMNDHMNEANDNSEQITRRMDE- | *Desulfamplus magnetovallimortis* | Bacteria; Proteobacteria; Deltaproteobacteria; Desulfobacterales; Desulfobacteraceae; Desulfamplus | hypothetical protein | 6 |
|  | **WP_085645485.1** | 5.1E-05 | Qa.III.b | IEGIGGAIDEVSQLSSDIASSVQQQTAAISEISTSAQEVSTHMQGISSD---- | *Thalassospira sp. MCCC 1A03138* | Bacteria; Proteobacteria; Alphaproteobacteria; Rhodospirillales; Rhodospirillaceae; Thalassospira | chemotaxis protein | 6 |
|  | **WP_097052142.1** | 5.1E-05 | Qa.III.b | IEGIGGAIDEVSQLSSDIASSVQQQTAAISEISTSAQEVSTHMQGISSD---- | *Thalassospira xiamenensis* | Bacteria; Proteobacteria; Alphaproteobacteria; Rhodospirillales; Rhodospirillaceae; Thalassospira | chemotaxis protein | 6 |
| | **WP_097659749.1** | 5.1E-05 | Qc.III | -RDISGTIEKLSEISSTIAAAVEEQGAATQEISRNVQQAAKGTQQVSSNITD- | *Bradyrhizobium sp. Y36* | Bacteria; Proteobacteria; Alphaproteobacteria; Rhizobiales; Bradyrhizobiaceae; Bradyrhizobium | methyl-accepting chemotaxis protein | 6 |
|  | **WP_102115242.1** | 5.1E-05 | Qa.IV | --AIDRTIREMDQISGTIAAAMEQQGAATKEIARNVQMAAMAT---------- | *Niveispirillum cyanobacteriorum* | Bacteria; Proteobacteria; Alphaproteobacteria; Rhodospirillales; Rhodospirillaceae; Niveispirillum | chemotaxis protein | 6 |

| | | | | | | | |
|---|---|---|---|---|---|---|---|
| WP_103041117.1 | 5.1E-05 | Qa.III | IRGIGATISSINDISSSIAAAIEEQGAATQEISRNVQQAAQSTQE-------- | *Azospirillum brasilense* | Bacteria; Proteobacteria; Alphaproteobacteria; Rhodospirillales; Rhodospirillaceae; Azospirillum | methyl-accepting chemotaxis protein | 6 |
| WP_109072264.1 | 5.1E-05 | Qa.III | IRGIGATISSINDISSSIAAAIEEQGAATQEISRNVQQAAQSTQE-------- | *Azospirillum sp. TSH58* | Bacteria; Proteobacteria; Alphaproteobacteria; Rhodospirillales; Rhodospirillaceae; Azospirillum | methyl-accepting chemotaxis protein | 6 |
| WP_110252496.1 | 5.1E-05 | Qa.III.b | IEGIGGAIDEVSQLSSDIASSVQQQTAAISEISTSAQEVSTHMQGISSD---- | *Thalassospira sp. 11-3* | Bacteria; Proteobacteria; Alphaproteobacteria; Rhodospirillales; Rhodospirillaceae; Thalassospira | chemotaxis protein | 6 |
| WP_111420931.1 | 5.1E-05 | Qc.III | IKEISGTIGRIAEISSAIAAAVEEQGAATHEIARNVQQAAAGTT--------- | *Rhodoplanes roseus* | Bacteria; Proteobacteria; Alphaproteobacteria; Rhizobiales; Hyphomicrobiaceae; Rhodoplanes | HAMP domain-containing protein | 6 |
| WP_114093539.1 | 5.1E-05 | Qa.III.b | IEGIGGAIDEVSQLSSDIASSVQQQTAAISEISTSAQEVSTHMQGISSD---- | *Thalassospira xiamenensis* | Bacteria; Proteobacteria; Alphaproteobacteria; Rhodospirillales; Rhodospirillaceae; Thalassospira | chemotaxis protein | 6 |
| WP_114109545.1 | 5.1E-05 | Qa.III.b | IEGIGGAIDEVSQLSSDIASSVQQQTAAISEISTSAQEVSTHMQGISSD---- | *Thalassospira xiamenensis* | Bacteria; Proteobacteria; Alphaproteobacteria; Rhodospirillales; Rhodospirillaceae; Thalassospira | chemotaxis protein | 6 |
| WP_114123197.1 | 5.1E-05 | Qa.III.b | IEGIGGAIDEVSQLSSDIASSVQQQTAAISEISTSAQEVSTHMQGISSD---- | *Thalassospira xianhensis* | Bacteria; Proteobacteria; Alphaproteobacteria; Rhodospirillales; Rhodospirillaceae; Thalassospira | chemotaxis protein | 6 |
| KRQ89819.1 | 5.2E-05 | Qa.IV | ---IERTIGLIGDTTALINTTITQQASATDEIAQNVSQAAAHTE--------- | *Bradyrhizobium valentinum* | Bacteria; Proteobacteria; Alphaproteobacteria; Rhizobiales; Bradyrhizobiaceae; Bradyrhizobium | hypothetical protein CP49_36525 | 6 |
| SDK23665.1 | 5.2E-05 | Qb.I | LQRSKEKLQETIEHAEVSNEELRASNEELQAINEELRSATEELETS------- | *Duganella sp. OV510* | Bacteria; Proteobacteria; Betaproteobacteria; Burkholderiales; Oxalobacteraceae; Duganella | two-component system chemotaxis family CheB/ CheR fusion protein | 6 |
| WP_002712326.1 | 5.2E-05 | Qa.III.b | IDGITRTISKVNEIASAIASAVEEQGAATREIAHNVSQAA------------- | *Afipia clevelandensis* | Bacteria; Proteobacteria; Alphaproteobacteria; Rhizobiales; Bradyrhizobiaceae; Afipia | methyl-accepting chemotaxis protein | 6 |
| WP_006034618.1 | 5.2E-05 | Qc.III | LDSMTNSVMQINDLGIQIATAAEEQSSVTEEINRNMTMIQNMVSQLTEN---- | *Moritella sp. PE36* | Bacteria; Proteobacteria; Gammaproteobacteria; Alteromonadales; Moritellaceae; Moritella | methyl-accepting chemotaxis protein partial | 6 |
| WP_007899145.1 | 5.2E-05 | Qb.I | LQRTKVHLQDTIEQSEISSEELKASNEEMQAINEELRSATEELETS------- | *Pseudomonas sp. GM102* | Bacteria; Proteobacteria; Gammaproteobacteria; Pseudomonadales; Pseudomonadaceae; Pseudomonas | PAS domain S-box protein | 6 |
| WP_008960738.1 | 5.2E-05 | Qc.III | IKEISGTIEKLSEISSTIAAAVEEQGAATQEISRNIQQAAQGTHEVSSNITD- | *Bradyrhizobium sp. STM 3809* | Bacteria; Proteobacteria; Alphaproteobacteria; Rhizobiales; Bradyrhizobiaceae; Bradyrhizobium | methyl-accepting chemotaxis protein | 6 |
| WP_009736783.1 | 5.2E-05 | Qa.III.b | IDGITRTISKVNEIASAIASAVEEQGAATREIAHNVSQAA------------- | *Bradyrhizobiaceae bacterium SG-6C* | Bacteria; Proteobacteria; Alphaproteobacteria; Rhizobiales; Bradyrhizobiaceae | methyl-accepting chemotaxis protein | 6 |
| WP_011382692.1 | 5.2E-05 | Qc.III | --GIAQTIARINEMSASIAGAVEQQGAAASEISRNVEQAAAGSREVVHNI--- | *Magnetospirillum magneticum* | Bacteria; Proteobacteria; Alphaproteobacteria; Rhodospirillales; Rhodospirillaceae; Magnetospirillum | methyl-accepting chemotaxis protein | 6 |
| WP_024574985.1 | 5.2E-05 | Qa.III.b | IDGITRTISKVNEIASAIASAVEEQGAATREIAHNVSQAA------------- | *Afipia* | Bacteria; Proteobacteria; Alphaproteobacteria; Rhizobiales; Bradyrhizobiaceae | MULTISPECIES: methyl-accepting chemotaxis protein | 6 |
| WP_035647814.1 | 5.2E-05 | Qc.III | ---ISGSIERLSEVSSTIAAAVEEQNAATKEISRNVQQAAQGTQQVSVNIT-- | *Bradyrhizobium sp. ORS 285* | Bacteria; Proteobacteria; Alphaproteobacteria; Rhizobiales; Bradyrhizobiaceae; Bradyrhizobium | HAMP domain-containing protein | 6 |
| WP_045110617.1 | 5.2E-05 | Qc.III.c | LDSMTESVLQINDLGIQIATAAEEQSSVTEEINRNMTTIQHMVQQL------- | *Moritella viscosa* | Bacteria; Proteobacteria; Gammaproteobacteria; Alteromonadales; Moritellaceae; Moritella | methyl-accepting chemotaxis protein | 6 |
| WP_049921902.1 | 5.2E-05 | SNAP.c | ----GAVAKQVKELAADTKSTAEDIEQRLERIDEQTERTATEVQRTADRIS-- | *Halopiger djelfimassiliensis* | Archaea; Euryarchaeota; Halobacteria; Natrialbales; Natrialbaceae; Halopiger | HAMP domain-containing protein | 6 |
| WP_075473123.1 | 5.2E-05 | Qc.III.c | LDSMTESVLQINDLGIQIATAAEEQSSVTEEINRNMTTIQHMVQQL------- | *Moritella viscosa* | Bacteria; Proteobacteria; Gammaproteobacteria; Alteromonadales; Moritellaceae; Moritella | methyl-accepting chemotaxis protein | 6 |
| WP_077599268.1 | 5.2E-05 | SNAP.c | ------GSEEVASLAASTARAVEEQNRTLDSVASAVEELSSAVRDV------- | *Salinivibrio kushneri* | Bacteria; Proteobacteria; Gammaproteobacteria; Vibrionales; Vibrionaceae; Salinivibrio | methyl-accepting chemotaxis protein | 6 |
| WP_092145858.1 | 5.2E-05 | Qc.III | IKEIGDTIGRMSEISSTIASAVEEQGAATQEISRNVQQAAQGTMQVSSNI--- | *Bradyrhizobiaceae* | Bacteria; Proteobacteria; Alphaproteobacteria; Rhizobiales | methyl-accepting chemotaxis protein | 6 |
| EJN07451.1 | 5.3E-05 | Qa.III.b | -EAIAQTIAEINGITASIATSIEQQGLATREIARNVQEASA------------ | *Bradyrhizobium sp. YR681* | Bacteria; Proteobacteria; Alphaproteobacteria; Rhizobiales; Bradyrhizobiaceae; Bradyrhizobium | methyl-accepting chemotaxis protein | 6 |
| OGS36165.1 | 5.3E-05 | Qa.III | --SITQVIEEVDAISQTIAAAVEEQSTTIREIAGNVGNASRSAEDI------- | *Elusimicrobia bacterium RIFOXYB2_FULL_49_7* | Bacteria; Elusimicrobia | hypothetical protein A2293_04595 | 6 |
| PHR89511.1 | 5.3E-05 | Qc.III | LDSMTESVMQINDLGIQIATAAEEQSSVTEEINRNMTMIQGMVNQLTE----- | *Moritella sp.* | Bacteria; Proteobacteria; Gammaproteobacteria; Alteromonadales; Moritellaceae; Moritella | methyl-accepting chemotaxis protein | 6 |
| PXX31994.1 | 5.3E-05 | Qa.III.b | IDSIGGTIAEISEIAATIASAVEQQGAATEEITRNVQQAS------------- | *Thalassospira sp. 11-3* | Bacteria; Proteobacteria; Alphaproteobacteria; Rhodospirillales; Rhodospirillaceae; Thalassospira | methyl-accepting chemotaxis protein | 6 |
| PZV00142.1 | 5.3E-05 | Qc.I | ------MVNQLRQTLDTLEQRVEERTAALATANQALQQQTHSLEIALQKLR-- | *Leptolyngbya sp.* | Bacteria; Cyanobacteria; Synechococcales; Leptolyngbyaceae; Leptolyngbya | GGDEF domain-containing protein | 6 |
| SEM54028.1 | 5.3E-05 | Qa.III.b | -EAIAQTIAEINGITASIATSIEQQGLATREIARNVQEASA------------ | *Bradyrhizobium sp. OK095* | Bacteria; Proteobacteria; Alphaproteobacteria; Rhizobiales; Bradyrhizobiaceae; Bradyrhizobium | methyl-accepting chemotaxis sensory transducer | 6 |
| SMX59629.1 | 5.3E-05 | Qc.III | ---ISGSIERLSEVSSTIAAAVEEQNAATKEISRNVQQAAQGTQQVSVNIT-- | *Bradyrhizobium sp. ORS 285* | Bacteria; Proteobacteria; Alphaproteobacteria; Rhizobiales; Bradyrhizobiaceae; Bradyrhizobium | putative methyl-accepting chemotaxis receptor/sensory transducer | 6 |
| WP_011160171.1 | 5.3E-05 | Qc.III | IKEIGQTIGRMSEIASTIASAVEEQGAATQEISRNVQQAAHGTQQVSSNI--- | *Rhodopseudomonas palustris* | Bacteria; Proteobacteria; Alphaproteobacteria; Rhizobiales; Bradyrhizobiaceae; Rhodopseudomonas | methyl-accepting chemotaxis protein | 6 |
| WP_011441869.1 | 5.3E-05 | Qa.III | IREIGATIGKMSEIASTIASAVEEQGAATQEISRNVQQAAQGTQQVSSS---- | *Rhodopseudomonas palustris* | Bacteria; Proteobacteria; Alphaproteobacteria; Rhizobiales; Bradyrhizobiaceae; Rhodopseudomonas | HAMP domain-containing protein | 6 |
| WP_012497777.1 | 5.3E-05 | Qc.III | IKEIGQTIGRMSEIASTIASAVEEQGAATQEISRNVQQAAHGTQQVSSNI--- | *Rhodopseudomonas palustris* | Bacteria; Proteobacteria; Alphaproteobacteria; Rhizobiales; Bradyrhizobiaceae; Rhodopseudomonas | methyl-accepting chemotaxis protein | 6 |
| WP_024339677.1 | 5.3E-05 | Qc.III | IGEISGTIARLSEIASAIAAAVEQQGAATQEIARNVQQAANGTQQVSSNVG-- | *Bradyrhizobium* | Bacteria; Proteobacteria; Alphaproteobacteria; Rhizobiales; Bradyrhizobiaceae | MULTISPECIES: PAS domain S-box protein | 6 |
| WP_037987997.1 | 5.3E-05 | Qa.III.b | IDSIGGTIAEISEIAATIASAVEQQGAATEEITRNVQQAS------------- | *Thalassospira* | Bacteria; Proteobacteria; Alphaproteobacteria; Rhodospirillales; Rhodospirillaceae | MULTISPECIES: methyl-accepting chemotaxis protein | 6 |
| WP_047308410.1 | 5.3E-05 | Qc.III | IKEIGQTIGRMSEIASTIASAVEEQGAATQEISRNVQQAAHGTQQVSSNI--- | *Rhodopseudomonas palustris* | Bacteria; Proteobacteria; Alphaproteobacteria; Rhizobiales; Bradyrhizobiaceae; Rhodopseudomonas | methyl-accepting chemotaxis protein | 6 |
| WP_071241166.1 | 5.3E-05 | Qa.III.b | IDSIGGTIAEISEIAATIASAVEQQGAATEEITRNVQQAS------------- | *Thalassospira sp. MIT1004* | Bacteria; Proteobacteria; Alphaproteobacteria; Rhodospirillales; Rhodospirillaceae; Thalassospira | methyl-accepting chemotaxis protein | 6 |
| WP_082822681.1 | 5.3E-05 | Qc.III.b | -------------MATTISAAVEEQGAATDEITRNVRSAADRTRTVSSSINDV | *Thalassospira xiamenensis* | Bacteria; Proteobacteria; Alphaproteobacteria; Rhodospirillales; Rhodospirillaceae; Thalassospira | chemotaxis protein | 6 |
| WP_084005030.1 | 5.3E-05 | Qc.III | -GSIGGTIGKMSEIASAIAAAVEEQGAATQEIARNVEQASAGTGEVTSNIA-- | *Magnetovibrio blakemorei* | Bacteria; Proteobacteria; Alphaproteobacteria; Rhodospirillales; Rhodospirillaceae; Magnetovibrio | methyl-accepting chemotaxis protein | 6 |
| WP_085645760.1 | 5.3E-05 | Qc.III.b | -------------MATTISAAVEEQGAATDEITRNVRSAADRTRTVSSSINDV | *Thalassospira sp. MCCC 1A03138* | Bacteria; Proteobacteria; Alphaproteobacteria; Rhodospirillales; Rhodospirillaceae; Thalassospira | chemotaxis protein | 6 |
| WP_097112669.1 | 5.3E-05 | Qa.III | --DIMQSVSTISDMTLQIASATEEQSSVVKEMDQNLLRVNSLTVETK------ | *Rheinheimera tuosuensis* | Bacteria; Proteobacteria; Gammaproteobacteria; Chromatiales; Chromatiaceae; Rheinheimera | HAMP domain-containing protein | 6 |
| WP_104513052.1 | 5.3E-05 | Qc.III | IKEIGQTIGRMSEIASTIASAVEEQGAATQEISRNVQQAAHGTQQVSSNI--- | *Rhodopseudomonas palustris* | Bacteria; Proteobacteria; Alphaproteobacteria; Rhizobiales; Bradyrhizobiaceae; Rhodopseudomonas | methyl-accepting chemotaxis protein | 6 |
| WP_107346750.1 | 5.3E-05 | Qc.III | IKEIGQTIGRMSEIASTIASAVEEQGAATQEISRNVQQAAHGTQQVSSNI--- | *Rhodopseudomonas palustris* | Bacteria; Proteobacteria; Alphaproteobacteria; Rhizobiales; Bradyrhizobiaceae; Rhodopseudomonas | HAMP domain-containing protein | 6 |
| WP_107357635.1 | 5.3E-05 | Qc.III | IKEIGQTIGRMSEIASTIASAVEEQGAATQEISRNVQQAAHGTQQVSSNI--- | *Rhodopseudomonas palustris* | Bacteria; Proteobacteria; Alphaproteobacteria; Rhizobiales; Bradyrhizobiaceae; Rhodopseudomonas | methyl-accepting chemotaxis protein | 6 |
| WP_109116988.1 | 5.3E-05 | Qc.III | IQGISGTIVTINEISTAIAAAVEEQAAATHEISRSVQQASAGTQEVTGNIAR- | *Azospirillum sp. TSO22-1* | Bacteria; Proteobacteria; Alphaproteobacteria; Rhodospirillales; Rhodospirillaceae; Azospirillum | methyl-accepting chemotaxis protein | 6 |
| WP_114356848.1 | 5.3E-05 | Qc.III | IKEIGQTIGRMSEIASTIASAVEEQGAATQEISRNVQQAAHGTQQVSSNI--- | *Rhodopseudomonas pentothenatexigens* | Bacteria; Proteobacteria; Alphaproteobacteria; Rhizobiales; Bradyrhizobiaceae; Rhodopseudomonas | HAMP domain-containing protein | 6 |
| AWJ88392.1 | 5.4E-05 | Qa.III.b | -----QVILRVNEIATSIASAVEEQGAATREISRNVQEAAGGTQEVS------ | *Azospirillum brasilense Sp245* | Bacteria; Proteobacteria; Alphaproteobacteria; Rhodospirillales; Rhodospirillaceae; Azospirillum | chemotaxis protein | 6 |
| OQW55537.1 | 5.4E-05 | Qc.III | -KDIGATITLISEISAAIATAVERQNVSTQEITRNVQHAANRSRSVAENIG-- | *Proteobacteria bacterium SG_bin9* | Bacteria; Proteobacteria | chemotaxis protein | 6 |
| PPD14678.1 | 5.4E-05 | Qa.III | -----------SSIATSVAAAVEEQSSALATISDNVNEAARSSAEGATAIK-- | *Methylobacterium sp.* | Bacteria; Proteobacteria; Alphaproteobacteria; Rhizobiales; Methylobacteriaceae; Methylobacterium | hypothetical protein CTY25_10105 | 6 |
| SHK45030.1 | 5.4E-05 | Qc.II | --ELAESVDQASGSAHTIEERVDEQTSLLSTVATEMENFSASM---------- | *Haladaptatus paucihalophilus DX253* | Archaea; Euryarchaeota; Halobacteria; Halobacteriales; Halobacteriaceae; Haladaptatus | methyl-accepting chemotaxis sensory transducer with Pas/Pac sensor | 6 |
| WP_021297147.1 | 5.4E-05 | Qa.I | -EETNKSAEHVSLSIHAMSDNIAIQARNVSESLTAVTGISEDIKQ-------- | *Alicyclobacillus acidoterrestris* | Bacteria; Firmicutes; Bacilli; Bacillales; Alicyclobacillaceae; Alicyclobacillus | methyl-accepting chemotaxis protein | 6 |
| WP_028577334.1 | 5.4E-05 | Qa.III | --QIVHVIGEVNTIVTAIAAAIEEQSAVTREVAENISHATHDVQDAN------ | *Desulfomicrobium escambiense* | Bacteria; Proteobacteria; Deltaproteobacteria; Desulfovibrionales; Desulfomicrobiaceae; Desulfomicrobium | methyl-accepting chemotaxis protein | 6 |
| WP_057427772.1 | 5.4E-05 | Qb.I | LQRTKSQLQDVIEQAEVSSEELKASNEEMQAINEELRSASEELETSK------ | *Pseudomonas syringae* | Bacteria; Proteobacteria; Gammaproteobacteria; Pseudomonadales; Pseudomonadaceae; Pseudomonas | PAS domain S-box protein partial | 6 |
| WP_085969952.1 | 5.4E-05 | Qa.III.b | -EAIAQTIAEINGITASIATSIEQQGLATREIARNVQEASA------------ | *Bradyrhizobium sp. YR681* | Bacteria; Proteobacteria; Alphaproteobacteria; Rhizobiales; Bradyrhizobiaceae; Bradyrhizobium | methyl-accepting chemotaxis protein | 6 |
| WP_092024316.1 | 5.4E-05 | Qa.III.b | -EAIAQTIAEINGITASIATSIEQQGLATREIARNVQEASA------------ | *Bradyrhizobium sp. OK095* | Bacteria; Proteobacteria; Alphaproteobacteria; Rhizobiales; Bradyrhizobiaceae; Bradyrhizobium | methyl-accepting chemotaxis protein | 6 |
| WP_094301320.1 | 5.4E-05 | Qa.III.b | -----QVILRVNEIATSIASAVEEQGAATREISRNVQEAAGGTQEVS------ | *Azospirillum brasilense* | Bacteria; Proteobacteria; Alphaproteobacteria; Rhodospirillales; Rhodospirillaceae; Azospirillum | HAMP domain-containing protein | 6 |
| WP_109069880.1 | 5.4E-05 | Qa.III.b | -----QVILRVNEIATSIASAVEEQGAATREISRNVQEAAGGTQEVS------ | *Azospirillum sp. TSH58* | Bacteria; Proteobacteria; Alphaproteobacteria; Rhodospirillales; Rhodospirillaceae; Azospirillum | HAMP domain-containing protein | 6 |
| WP_011663318.1 | 5.5E-05 | Qa.III | IKEIGETIAKMSEISSTIASAVEQQGTATQEISRNVQQAAHGTQQVS------ | *Rhodopseudomonas palustris* | Bacteria; Proteobacteria; Alphaproteobacteria; Rhizobiales; Bradyrhizobiaceae; Rhodopseudomonas | methyl-accepting chemotaxis protein | 6 |
| WP_012564033.1 | 5.5E-05 | Qa.IV | IKEISATIDQISEISSAIAAAVEEQGAATQEIARNVRQAAE------------ | *Oligotropha carboxidovorans* | Bacteria; Proteobacteria; Alphaproteobacteria; Rhizobiales; Bradyrhizobiaceae; Oligotropha | methyl-accepting chemotaxis protein | 6 |
| WP_045585265.1 | 5.5E-05 | Qa.III | IRGIGQTIGRLNDIAASIAAAVEQQGAATSEIARNVQQAAGGTAQASQDIGQ- | *Azospirillum thiophilum* | Bacteria; Proteobacteria; Alphaproteobacteria; Rhodospirillales; Rhodospirillaceae; Azospirillum | HAMP domain-containing protein | 6 |

| | | | | | | | |
|---|---|---|---|---|---|---|---|
| **WP_063085882.1** | 5.5E-05 | Qc.III | IEKVTHTISSISEIAAGISSAVSEQNAAVNEIANSATSASSGSSEVVANIN-- | *Thalassospira* | Bacteria; Proteobacteria; Alphaproteobacteria; Rhodospirillales; Rhodospirillaceae | MULTISPECIES: methyl-accepting chemotaxis protein | 6 |
| **SFN49883.1** | 5.6E-05 | Qc.III | -DDISKTIERVSEISTAIAAAVEQQSASTREISSNIEEAAV------------ | *Cohaesibacter marisflavi* | Bacteria; Proteobacteria; Alphaproteobacteria; Rhizobiales; Cohaesibacteraceae | Methyl-accepting chemotaxis protein | 6 |
| **WP_052711948.1** | 5.6E-05 | Qc.III | --DIHQTIGTIADISANVAAAVEEQTAATREISRSVDEASRGTRDA------- | *Elstera litoralis* | Bacteria; Proteobacteria; Alphaproteobacteria; Rhodospirillales; Rhodospirillaceae; Elstera | HAMP domain-containing protein | 6 |
| **WP_058122067.1** | 5.6E-05 | Qc.I | -DEFAQMVRDIKENIKDVTTGLEQDQKLVKQSLQVIERAKQ------------ | *Helicobacter typhlonius* | Bacteria; Proteobacteria; Epsilonproteobacteria; Campylobacterales; Helicobacteraceae; Helicobacter | chemotaxis protein | 6 |
| **WP_085553163.1** | 5.6E-05 | Qa.III | IRRIVGSIATINDISTGIAAAVDEQDAGTREIARNVEQAA------------- | *Azospirillum lipoferum* | Bacteria; Proteobacteria; Alphaproteobacteria; Rhodospirillales; Rhodospirillaceae; Azospirillum | HAMP domain-containing protein | 6 |
| **WP_109966992.1** | 5.6E-05 | Qb.I | ---TKENLQATIEEMQAANEELKSTNEELQSTNEELQSTNEELETSK------ | *Methanospirillum lacunae* | Archaea; Euryarchaeota; Methanomicrobia; Methanomicrobiales; Methanospirillaceae; Methanospirillum | SAM-dependent methyltransferase | 6 |
| **OHE62884.1** | 5.7E-05 | Qa.III | -------IRQVNEIVTGIAAAIEEQTVATRDIASSVAQASQGVEDANRNI--- | *Treponema sp. GWC1_61_84* | Bacteria; Spirochaetes; Spirochaetales; Spirochaetaceae; Treponema | hypothetical protein A2001_00330 | 6 |
| **PIP89537.1** | 5.7E-05 | Qc.III | --GITQIIEKLKEVNISVASAVEEQNAAISEISRAAKEASKGTSEVNTNIKL- | *Bdellovibrionales bacterium CG22_combo_CG10-13_8_21_14_all_38_13* | Bacteria; Proteobacteria; Oligoflexia; Bdellovibrionales | hypothetical protein COW79_11230 | 6 |
| **SFI66346.1** | 5.7E-05 | Qc.III | -QTIGGIIGEVNEVATAIAAAVQEQGAATQEITRSTQYAAQGTKNVS------ | *Bradyrhizobium sp. cf659* | Bacteria; Proteobacteria; Alphaproteobacteria; Rhizobiales; Bradyrhizobiaceae; Bradyrhizobium | Methyl-accepting chemotaxis protein | 6 |
| **WP_029010202.1** | 5.7E-05 | Qa.III | IKDVTRVINQVNEVATSIASAVEEQAAATREISRNVQQAAR------------ | *Azospirillum halopraeferens* | Bacteria; Proteobacteria; Alphaproteobacteria; Rhodospirillales; Rhodospirillaceae; Azospirillum | HAMP domain-containing protein | 6 |
| **WP_038958947.1** | 5.7E-05 | Qa.IV | IRDISATIERLSEVSSTIAAAVEEQGAATLEISRNVQQAA------------- | *Bradyrhizobium japonicum* | Bacteria; Proteobacteria; Alphaproteobacteria; Rhizobiales; Bradyrhizobiaceae; Bradyrhizobium | methyl-accepting chemotaxis protein | 6 |
| **WP_045002532.1** | 5.7E-05 | Qc.III | -QTIGGIIGEVNEVATAIAAAVQEQGAATQEITRSTQYAAQGTKNVS------ | *Bradyrhizobium sp. LTSP857* | Bacteria; Proteobacteria; Alphaproteobacteria; Rhizobiales; Bradyrhizobiaceae; Bradyrhizobium | methyl-accepting chemotaxis protein | 6 |
| **WP_045009707.1** | 5.7E-05 | Qc.III | -QTIGGIIGEVNEVATAIAAAVQEQGAATQEITRSTQYAAQGTKNVS------ | *Bradyrhizobium sp. LTSP849* | Bacteria; Proteobacteria; Alphaproteobacteria; Rhizobiales; Bradyrhizobiaceae; Bradyrhizobium | methyl-accepting chemotaxis protein | 6 |
| **WP_063203805.1** | 5.7E-05 | Qc.III | -QTIGGIIGEVNEVATAIAAAVQEQGAATQEITRSTQYAAQGTKNVS------ | *Bradyrhizobium sp. AT1* | Bacteria; Proteobacteria; Alphaproteobacteria; Rhizobiales; Bradyrhizobiaceae; Bradyrhizobium | methyl-accepting chemotaxis protein | 6 |
| **WP_063694218.1** | 5.7E-05 | Qc.III | -QTIGGIIGEVNEVATAIAAAVQEQGAATQEITRSTQYAAQGTKNVS------ | *Bradyrhizobium stylosanthis* | Bacteria; Proteobacteria; Alphaproteobacteria; Rhizobiales; Bradyrhizobiaceae; Bradyrhizobium | methyl-accepting chemotaxis protein | 6 |
| **WP_090067776.1** | 5.7E-05 | Qc.III | -DDISKTIERVSEISTAIAAAVEQQSASTREISSNIEEAAV------------ | *Cohaesibacter marisflavi* | Bacteria; Proteobacteria; Alphaproteobacteria; Rhizobiales; Cohaesibacteraceae | methyl-accepting chemotaxis protein | 6 |
| **WP_097659952.1** | 5.7E-05 | Qc.III | -QTIGGIIGEVNEVATAIAAAVQEQGAATQEITRSTQYAAQGTKNVS------ | *Bradyrhizobium sp. Y36* | Bacteria; Proteobacteria; Alphaproteobacteria; Rhizobiales; Bradyrhizobiaceae; Bradyrhizobium | methyl-accepting chemotaxis protein | 6 |
| **WP_100176119.1** | 5.7E-05 | Qc.III | -QTIGGIIGEVNEVATAIAAAVQEQGAATQEITRSTQYAAQGTKNVS------ | *Bradyrhizobium sp. TSA1* | Bacteria; Proteobacteria; Alphaproteobacteria; Rhizobiales; Bradyrhizobiaceae; Bradyrhizobium | methyl-accepting chemotaxis protein | 6 |
| **WP_110973329.1** | 5.7E-05 | Qc.III.c | LQGIREAVERITGMSQQMAAAAEEQAHVAEDIARQINNVAGTVERTA------ | *Pseudomonas sp. WCHPs060044* | Bacteria; Proteobacteria; Gammaproteobacteria; Pseudomonadales; Pseudomonadaceae; Pseudomonas | PAS domain S-box protein | 6 |
| **WP_114356784.1** | 5.7E-05 | Qa.IV | IREIGNTIARMSEISATIASAVEEQGAATQEISRNVHHAA------------- | *Rhodopseudomonas pentothenatexigens* | Bacteria; Proteobacteria; Alphaproteobacteria; Rhizobiales; Bradyrhizobiaceae; Rhodopseudomonas | HAMP domain-containing protein | 6 |
| **RDC06286.1** | 5.8E-05 | Qc.I | -----DMAGELRGMYEHLESQVQERTVDLREANALLERQRDKLEQLNA----- | *Eggerthella lenta* | Bacteria; Actinobacteria; Coriobacteriia; Eggerthellales; Eggerthellaceae; Eggerthella | histidine kinase | 6 |
| **RDC18950.1** | 5.8E-05 | Qc.I | -----DMAGELRGMYEHLESQVQERTVDLREANALLERQRDKLEQLNA----- | *Eggerthella lenta* | Bacteria; Actinobacteria; Coriobacteriia; Eggerthellales; Eggerthellaceae; Eggerthella | histidine kinase | 6 |
| **SFR76044.1** | 5.8E-05 | Qc.I | ----------LRGHASTLEDQVAQRTAQLTEANAELTQALANLKTAQREL--- | *Mitsuaria sp. PDC51* | Bacteria; Proteobacteria; Betaproteobacteria; Burkholderiales; Mitsuaria | His Kinase A (phospho-acceptor) domain-containing protein | 6 |
| **SOC23725.1** | 5.8E-05 | Qa.III.b | IDSIGSTIAEISEIAATIASAVEQQGAATEEITRNVQQAS------------- | *Thalassospira xiamenensis* | Bacteria; Proteobacteria; Alphaproteobacteria; Rhodospirillales; Rhodospirillaceae; Thalassospira | methyl-accepting chemotaxis protein | 6 |
| **WP_009305407.1** | 5.8E-05 | Qc.I | -----DMAGELRGMYEHLESQVQERTVDLREANALLERQRDKLEQLNA----- | *Eggerthella* | Bacteria; Actinobacteria; Coriobacteriia; Eggerthellales; Eggerthellaceae | DUF3365 domain-containing protein | 6 |
| **WP_009608487.1** | 5.8E-05 | Qc.I | -----DMAGELRGMYEHLESQVQERTVDLREANALLERQRDKLEQLNA----- | *Eggerthella* | Bacteria; Actinobacteria; Coriobacteriia; Eggerthellales; Eggerthellaceae | MULTISPECIES: DUF3365 domain-containing protein | 6 |
| **WP_025896695.1** | 5.8E-05 | Qa.III | MKSIRRGITTIAEMASGVASAVEEQRAATDEIARNVQEAAT------------ | *Kordiimonas gwangyangensis* | Bacteria; Proteobacteria; Alphaproteobacteria; Kordiimonadales; Kordiimonadaceae; Kordiimonas | methyl-accepting chemotaxis protein | 6 |
| **WP_038358605.1** | 5.8E-05 | Qa.IV | ---------SLNQISGAVAIVVEQQNVATAGIAENIRQAADETVTAAGHID-- | *Bosea sp. UNC402CLCol* | Bacteria; Proteobacteria; Alphaproteobacteria; Rhizobiales; Bradyrhizobiaceae; Bosea | hypothetical protein | 6 |
| **WP_039444850.1** | 5.8E-05 | Qa.III | LEKINAAVNLISDMATQIATAAEEQTHVTSEITQNITSIK------------- | *Vibrio navarrensis* | Bacteria; Proteobacteria; Gammaproteobacteria; Vibrionales; Vibrionaceae; Vibrio | methyl-accepting chemotaxis protein | 6 |
| **WP_047765001.1** | 5.8E-05 | Qa.IV | --KITGTIEQINEITASVAAAVEEQGAATLEISRNVQEAATA----------- | *Kiloniella spongiae* | Bacteria; Proteobacteria; Alphaproteobacteria; Kiloniellales; Kiloniellaceae; Kiloniella | methyl-accepting chemotaxis protein | 6 |
| **WP_052711521.1** | 5.8E-05 | Qa.III | IQGIAATILRINDISSGIAAAVEQQSAATREITRNVQEAARSTEDVS------ | *Elstera litoralis* | Bacteria; Proteobacteria; Alphaproteobacteria; Rhodospirillales; Rhodospirillaceae; Elstera | HAMP domain-containing protein | 6 |
| **WP_074118545.1** | 5.8E-05 | Qc.III | IKEIGATIGRISEISAAISASVEQQGSATQEISRNVQRAASGTSQV------- | *Bradyrhizobium sp. AS23.2* | Bacteria; Proteobacteria; Alphaproteobacteria; Rhizobiales; Bradyrhizobiaceae; Bradyrhizobium | HAMP domain-containing protein | 6 |
| **WP_084809452.1** | 5.8E-05 | Qc.III | IKEIGATIGRISEISAAISASVEQQGSATQEISRNVQRAASGTSQV------- | *Bradyrhizobium sp. NAS80.1* | Bacteria; Proteobacteria; Alphaproteobacteria; Rhizobiales; Bradyrhizobiaceae; Bradyrhizobium | HAMP domain-containing protein | 6 |
| **WP_085901969.1** | 5.8E-05 | Qa.IV | --KITGTIEQINEITASVAAAVEEQGAATLEISRNVQEAATA----------- | *Kiloniella majae* | Bacteria; Proteobacteria; Alphaproteobacteria; Kiloniellales; Kiloniellaceae; Kiloniella | methyl-accepting chemotaxis protein | 6 |
| **WP_085906747.1** | 5.8E-05 | Qa.IV | --KITGTIEQINEITASVAAAVEEQGAATLEISRNVQEAATA----------- | *Kiloniella majae* | Bacteria; Proteobacteria; Alphaproteobacteria; Kiloniellales; Kiloniellaceae; Kiloniella | HAMP domain-containing protein | 6 |
| **WP_085909894.1** | 5.8E-05 | Qa.IV | --KITGTIEQINEITASVAAAVEEQGAATLEISRNVQEAATA----------- | *Kiloniella majae* | Bacteria; Proteobacteria; Alphaproteobacteria; Kiloniellales; Kiloniellaceae; Kiloniella | methyl-accepting chemotaxis protein | 6 |
| **WP_090073073.1** | 5.8E-05 | R.III | -----REVSDTVGHLAQSVEEIASSADNASEISLKAANLSSNAND-------- | *Cohaesibacter marisflavi* | Bacteria; Proteobacteria; Alphaproteobacteria; Rhizobiales; Cohaesibacteraceae | ammonium transporter | 6 |
| **WP_092518255.1** | 5.8E-05 | Qc.III | IKEIGGTIGKISDIATTISDAVQQQSTATQEIARSVQNVAQG----------- | *Afipia sp. GAS231* | Bacteria; Proteobacteria; Alphaproteobacteria; Rhizobiales; Bradyrhizobiaceae; Afipia | methyl-accepting chemotaxis protein | 6 |
| **WP_109050807.1** | 5.8E-05 | Qa.III | IRSIAGTIAEINEIAAAIAAAVEQQGAATGEIARNVQEAA------------- | *Azospirillum sp. TSA6c* | Bacteria; Proteobacteria; Alphaproteobacteria; Rhodospirillales; Rhodospirillaceae; Azospirillum | urea ABC transporter substrate-binding protein | 6 |
| **WP_109095588.1** | 5.8E-05 | Qa.III | IRSIAGTIAEINEIAAAIAAAVEQQGAATGEIARNVQEAA------------- | *Azospirillum sp. TSH64* | Bacteria; Proteobacteria; Alphaproteobacteria; Rhodospirillales; Rhodospirillaceae; Azospirillum | urea ABC transporter substrate-binding protein | 6 |
| **WP_112764514.1** | 5.8E-05 | Qa.IV | ---------SLNQISGAVAIVVEQQNVATAGIAENIRQAADETVTAAGHID-- | *Rhizobiales bacterium* | Bacteria; Proteobacteria; Alphaproteobacteria; Rhizobiales | chemotaxis protein | 6 |
| **WP_113254721.1** | 5.8E-05 | Qa.IV | ---------SLNQISGAVAIVVEQQNVATAGIAENIRQAADETVTAAGHID-- | *Rhizobiales bacterium* | Bacteria; Proteobacteria; Alphaproteobacteria; Rhizobiales | chemotaxis protein | 6 |
| **AWJ90916.1** | 5.9E-05 | Qa.III | IEGIGRTIAAINEISTSIAAAIEEQSAATNEISRNVQQAA------------- | *Azospirillum brasilense Sp245* | Bacteria; Proteobacteria; Alphaproteobacteria; Rhodospirillales; Rhodospirillaceae; Azospirillum | methyl-accepting chemotaxis protein | 6 |
| **OGR05117.1** | 5.9E-05 | Qa.III | IRDIGTTIGEIDEISASIAAAVEEQGVVTLDIAQNTQQAA------------- | *Deltaproteobacteria bacterium RIFOXYD12_FULL_50_9* | Bacteria; Proteobacteria; Deltaproteobacteria | hypothetical protein A2511_13595 | 6 |
| **RCK06737.1** | 5.9E-05 | Qa.III.b | IDSIGSTIAEISEIAATIASAVEQQGAATEEITRNVQQAS------------- | *Thalassospira xianhensis MCCC 1A02616* | Bacteria; Proteobacteria; Alphaproteobacteria; Rhodospirillales; Rhodospirillaceae; Thalassospira | chemotaxis protein | 6 |
| **RCK32599.1** | 5.9E-05 | Qa.III.b | IDSIGSTIAEISEIAATIASAVEQQGAATEEITRNVQQAS------------- | *Thalassospira xiamenensis* | Bacteria; Proteobacteria; Alphaproteobacteria; Rhodospirillales; Rhodospirillaceae; Thalassospira | chemotaxis protein | 6 |
| **RCK50972.1** | 5.9E-05 | Qa.III.b | IDSIGSTIAEISEIAATIASAVEQQGAATEEITRNVQQAS------------- | *Thalassospira xiamenensis* | Bacteria; Proteobacteria; Alphaproteobacteria; Rhodospirillales; Rhodospirillaceae; Thalassospira | chemotaxis protein | 6 |
| **SON57049.1** | 5.9E-05 | Qa.III | INSISETVRNINEVSTSIALAVEQQRAATDEIARNVDEAAR------------ | *Hartmannibacter diazotrophicus* | Bacteria; Proteobacteria; Alphaproteobacteria; Rhizobiales; Hartmannibacter | Methyl-accepting chemotaxis protein 4 | 6 |
| **WP_067743901.1** | 5.9E-05 | Qc.III | LDNISQAVSQISDMNISIASAVEEQSAAANEISQNVVRVTG------------ | *Alteromonadaceae bacterium XY-R5* | Bacteria; Proteobacteria; Gammaproteobacteria; Alteromonadales; Alteromonadaceae | methyl-accepting chemotaxis protein | 6 |
| **WP_068499280.1** | 5.9E-05 | Qa.III | IDEIVGTIRQMDEITATIAQAVQKQEAATRDISANIEDVAQKAEE-------- | *Magnetospirillum moscoviense* | Bacteria; Proteobacteria; Alphaproteobacteria; Rhodospirillales; Rhodospirillaceae; Magnetospirillum | chemotaxis protein | 6 |
| **WP_077460595.1** | 5.9E-05 | Qc.III | -DEISSNLSNIKTLSSDMHEAIDQLGPIVSDLKDNVNALNG------------ | *Salinivibrio sp. AR647* | Bacteria; Proteobacteria; Gammaproteobacteria; Vibrionales; Vibrionaceae; Salinivibrio | methyl-accepting chemotaxis protein | 6 |
| **WP_077577201.1** | 5.9E-05 | Qc.III | -DEISSNLSNIKTLSSDMHEAIDQLGPIVSDLKDNVNALNG------------ | *Salinivibrio sp. AR640* | Bacteria; Proteobacteria; Gammaproteobacteria; Vibrionales; Vibrionaceae; Salinivibrio | methyl-accepting chemotaxis protein | 6 |
| **WP_077669100.1** | 5.9E-05 | Qc.III | -DEISSNLSNIKTLSSDMHEAIDQLGPIVSDLKDNVNALNG------------ | *Salinivibrio costicola* | Bacteria; Proteobacteria; Gammaproteobacteria; Vibrionales; Vibrionaceae; Salinivibrio | methyl-accepting chemotaxis protein | 6 |
| **WP_077676919.1** | 5.9E-05 | Qc.III | -DEISSNLSNIKTLSSDMHEAIDQLGPIVSDLKDNVNALNG------------ | *Salinivibrio sp. MA607* | Bacteria; Proteobacteria; Gammaproteobacteria; Vibrionales; Vibrionaceae; Salinivibrio | methyl-accepting chemotaxis protein | 6 |
| **WP_079864908.1** | 5.9E-05 | Qa.III | ---------EIDSMQQSVATATEEQTAVVDSLNMDITEINTLNQEGVENLQ-- | *Pseudomonas aeruginosa* | Bacteria; Proteobacteria; Gammaproteobacteria; Pseudomonadales; Pseudomonadaceae; Pseudomonas | hypothetical protein partial | 6 |
| **WP_082012422.1** | 5.9E-05 | Qc.III | -QGIAGTVGEASQIAGAIAAAVEEQGAATREIARSAQRAASGTQEVNRNIG-- | *Belnapia sp. F-4-1* | Bacteria; Proteobacteria; Alphaproteobacteria; Rhodospirillales; Acetobacteraceae; Belnapia | methyl-accepting chemotaxis protein | 6 |
| **WP_105629776.1** | 5.9E-05 | R.IV | IDQISKAVSELDSTTQQNAALVQESSAAAASLEQQAGELSQLISRF------- | *Cronobacter malonaticus* | Bacteria; Proteobacteria; Gammaproteobacteria; Enterobacterales; Enterobacteriaceae; Cronobacter | chemotaxis protein | 6 |
| **BAF69483.1** | 6E-05 | Qc.III | -------VKKIVANSNNISDEAEDVNKLLDDVVVKLDQLLTLIHKSN------ | *Nitratiruptor sp. SB155-2* | Bacteria; Proteobacteria; Epsilonproteobacteria; Nitratiruptor | methyl-accepting chemotaxis protein | 6 |
| **KLN61329.1** | 6E-05 | Qc.III | --GISKTIDSIHEVATAIASAVEEQGAATAEITKSVQQAANGTDQVSANI--- | *Kiloniella spongiae* | Bacteria; Proteobacteria; Alphaproteobacteria; Kiloniellales; Kiloniellaceae; Kiloniella | hypothetical protein WH96_06685 | 6 |
| **SDO33880.1** | 6E-05 | Qc.III | IKEIGGTIGKISDIATTISDAVQQQSTATQEIARSVQNVAQG----------- | *Afipia sp. GAS231* | Bacteria; Proteobacteria; Alphaproteobacteria; Rhizobiales; Bradyrhizobiaceae; Afipia | methyl-accepting chemotaxis sensory transducer | 6 |
| **WP_022665483.1** | 6E-05 | Qa.III | ITNITRVVNEVNDIVSTIAAAVTQQSATTGEIAENVSQAS------------- | *Desulfospira joergensenii* | Bacteria; Proteobacteria; Deltaproteobacteria; Desulfobacterales; Desulfobacteraceae; Desulfospira | methyl-accepting chemotaxis protein | 6 |

| | | | | | | | |
|---|---|---|---|---|---|---|---|
| **WP_051487599.1** | 6E-05 | Qa.III.b | LRQIANSVTEISGLVSEIAGSAQQQSLNLDEINGSVTQLDQSTQQVAARL--- | *Roseivivax marinus* | Bacteria; Proteobacteria; Alphaproteobacteria; Rhodobacterales; Rhodobacteraceae; Roseivivax | HAMP domain-containing protein | 6 |
| **WP_073629555.1** | 6E-05 | Qa.III | IDGIRVTIEKMNAVANEIALAVDEQHSATGEIVQNVDQAA------------ | *Pseudoxanthobacter soli* | Bacteria; Proteobacteria; Alphaproteobacteria; Rhizobiales; Xanthobacteraceae; Pseudoxanthobacter | methyl-accepting chemotaxis protein | 6 |
| **WP_082130371.1** | 6E-05 | Qc.III | --GISKTIDSIHEVATAIASAVEEQGAATAEITKSVQQAANGTDQVSANI--- | *Kiloniella spongiae* | Bacteria; Proteobacteria; Alphaproteobacteria; Kiloniellales; Kiloniellaceae; Kiloniella | HAMP domain-containing protein | 6 |
| **WP_084292247.1** | 6E-05 | Qc.III | -QTIGGIIGEVNEVATAIAAAVQEQGAATQEITRSTQYAAQGTKNVS------ | *Bradyrhizobium sp. WSM3983* | Bacteria; Proteobacteria; Alphaproteobacteria; Rhizobiales; Bradyrhizobiaceae; Bradyrhizobium | methyl-accepting chemotaxis protein | 6 |
| **WP_092186723.1** | 6E-05 | Qc.III | -QTIGGIIGEVNEVATAIAAAVQEQGAATQEITRSTQYAAQGTKNVS------ | *Bradyrhizobium sp. cf659* | Bacteria; Proteobacteria; Alphaproteobacteria; Rhizobiales; Bradyrhizobiaceae; Bradyrhizobium | methyl-accepting chemotaxis protein | 6 |
| **WP_092239546.1** | 6E-05 | Qc.III | IKEISGTIDRISEISSAIASAVEEQGAATQEIARNIQQAAQGTDQVAANI--- | *Bradyrhizobium sp. Gha* | Bacteria; Proteobacteria; Alphaproteobacteria; Rhizobiales; Bradyrhizobiaceae; Bradyrhizobium | methyl-accepting chemotaxis protein | 6 |
| **WP_092810919.1** | 6E-05 | Qa.III.b | LRQIANSVTEISGLVSEIAGSAQQQSLNLDEINGSVTQLDQSTQQVAARL--- | *Roseivivax marinus* | Bacteria; Proteobacteria; Alphaproteobacteria; Rhodobacterales; Rhodobacteraceae; Roseivivax | HAMP domain-containing protein | 6 |
| **KPP82248.1** | 6.1E-05 | Qc.III | IDEISRVIEQVDEISASIAAAVEEQSAVTEDISRNMQTAAAGVSQVSGAIKD- | *Oceanicaulis sp. HLUCCA04* | Bacteria; Proteobacteria; Alphaproteobacteria; Rhodobacterales; Hyphomonadaceae; Oceanicaulis | chemotaxis signal relay system methyl-accepting signal transducer | 6 |
| **WP_036510253.1** | 6.1E-05 | Qc.III | IDEISRVIEQVDEISASIAAAVEEQSAVTEDISRNMQTAAAGVSQVSGAIKD- | *Oceanicaulis sp. HL-87* | Bacteria; Proteobacteria; Alphaproteobacteria; Rhodobacterales; Hyphomonadaceae; Oceanicaulis | methyl-accepting chemotaxis protein | 6 |
| **WP_041756414.1** | 6.1E-05 | Qc.III | ---ISGSIERLSEVSSTIAAAVEEQNAATREISRNVQQAAQGTQQVSANIT-- | *Bradyrhizobium sp. ORS 278* | Bacteria; Proteobacteria; Alphaproteobacteria; Rhizobiales; Bradyrhizobiaceae; Bradyrhizobium | HAMP domain-containing protein | 6 |
| **WP_073955527.1** | 6.1E-05 | Qc.III.c | -EGIGGSIDEVNRLSIDISDAIQHQTTAIEDISASAKEVSSHMQ--------- | *Thalassospira sp. TSL5-1* | Bacteria; Proteobacteria; Alphaproteobacteria; Rhodospirillales; Rhodospirillaceae; Thalassospira | chemotaxis protein | 6 |
| **WP_103447404.1** | 6.1E-05 | Qb.I | LQRTRVQLQETIEQSEISSEELTASNEEMQAINEELRSASEELETSK------ | *Pseudomonas putida* | Bacteria; Proteobacteria; Gammaproteobacteria; Pseudomonadales; Pseudomonadaceae; Pseudomonas | response regulator | 6 |
| **AHK79125.1** | 6.2E-05 | Qc.III | LEDITQSIGTINDMNTQIASAAEEQSAVAEDVHRSLTRISTGV---------- | *Halorhodospira halochloris str. A* | Bacteria; Proteobacteria; Gammaproteobacteria; Chromatiales; Ectothiorhodospiraceae; Halorhodospira | hypothetical protein M911_08120 | 6 |
| **ALG71873.1** | 6.2E-05 | Qa.III | IVGIGRTIHSINQIAAGISAGIEQQGSAVREIVVNMEQAAA------------ | *Azospirillum thiophilum* | Bacteria; Proteobacteria; Alphaproteobacteria; Rhodospirillales; Rhodospirillaceae; Azospirillum | chemotaxis protein | 6 |
| **KFL46091.1** | 6.2E-05 | Qb.I | ----RERLQDALEETQHQASELQAQSEELRVSNEELEEQGAALRESQARL--- | *Sphingobium sp. ba1* | Bacteria; Proteobacteria; Alphaproteobacteria; Sphingomonadales; Sphingomonadaceae; Sphingobium | signal transduction histidine kinase | 6 |
| **KIU37350.1** | 6.2E-05 | Qa.III | IDAIQGTIRTLNAVSSTIAAAVEEQSAVMREMSGSMQTAAEDV---------- | *Methylobacterium radiotolerans* | Bacteria; Proteobacteria; Alphaproteobacteria; Rhizobiales; Methylobacteriaceae; Methylobacterium | chemotaxis protein | 6 |
| **KZB98570.1** | 6.2E-05 | Qa.III | IDAIQGTIRTLNAVSSTIAAAVEEQSAVMREMSGSMQTAAEDV---------- | *Methylobacterium radiotolerans* | Bacteria; Proteobacteria; Alphaproteobacteria; Rhizobiales; Methylobacteriaceae; Methylobacterium | Biofilm dispersion protein BdlA | 6 |
| **OUR77568.1** | 6.2E-05 | Qc.III | IKEISSTISKVDEIAASIASAVEEQGAATQEISSNAQRAASGTQEVTSSID-- | *Alphaproteobacteria bacterium 46_93_T64* | Bacteria; Proteobacteria; Alphaproteobacteria | hypothetical protein A9Q83_10540 | 6 |
| **PXB99245.1** | 6.2E-05 | Qa.III | -VGISEAVSNITEMTTQIAAAAEEQSAVAEEINRNISTIAALAEQTSDE---- | *Pseudomonas aeruginosa* | Bacteria; Proteobacteria; Gammaproteobacteria; Pseudomonadales; Pseudomonadaceae; Pseudomonas | chemotaxis protein partial | 6 |
| **SFK12081.1** | 6.2E-05 | Qc.III | IKEISGTIDRISEISSAIASAVEEQGAATQEIARNIQQAAQGTDQVAANI--- | *Bradyrhizobium sp. Gha* | Bacteria; Proteobacteria; Alphaproteobacteria; Rhizobiales; Bradyrhizobiaceae; Bradyrhizobium | methyl-accepting chemotaxis protein | 6 |
| **WP_011474508.1** | 6.2E-05 | Qc.III | IKQIGDTIGRMSEIASTIAAAVEQQGAATQEISRNIQHAAQGTQAVSSNI--- | *Rhodopseudomonas palustris* | Bacteria; Proteobacteria; Alphaproteobacteria; Rhizobiales; Bradyrhizobiaceae; Rhodopseudomonas | methyl-accepting chemotaxis protein | 6 |
| **WP_012317222.1** | 6.2E-05 | Qa.III | IDAIQGTIRTLNAVSSTIAAAVEEQSAVMREMSGSMQTAAEDV---------- | *Methylobacterium radiotolerans* | Bacteria; Proteobacteria; Alphaproteobacteria; Rhizobiales; Methylobacteriaceae; Methylobacterium | PAS domain S-box protein | 6 |
| **WP_015902923.1** | 6.2E-05 | Qa.IV | ---------NVNEMVTTIAAAVEEQSATTNEIASNVLQASKGIQEVNQNV--- | *Desulfobacterium autotrophicum* | Bacteria; Proteobacteria; Deltaproteobacteria; Desulfobacterales; Desulfobacteraceae; Desulfobacterium | methyl-accepting chemotaxis protein | 6 |
| **WP_018873534.1** | 6.2E-05 | Qa.III | LQQIRQSVTTIVEMTDQIAVAARQQSTVVEDINRNIVQISDEA---------- | *Thioalkalivibrio sp. ALJ16* | Bacteria; Proteobacteria; Gammaproteobacteria; Chromatiales; Ectothiorhodospiraceae; Thioalkalivibrio | methyl-accepting chemotaxis protein | 6 |
| **WP_059408242.1** | 6.2E-05 | Qa.III | IDAIQGTIRTLNAVSSTIAAAVEEQSAVMREMSGSMQTAAEDV---------- | *Methylobacterium radiotolerans* | Bacteria; Proteobacteria; Alphaproteobacteria; Rhizobiales; Methylobacteriaceae; Methylobacterium | PAS domain S-box protein | 6 |
| **WP_070998310.1** | 6.2E-05 | Qa.III | IDAIQGTIRTLNAVSSTIAAAVEEQSAVMREMSGSMQTAAEDV---------- | *Methylobacterium sp. C1* | Bacteria; Proteobacteria; Alphaproteobacteria; Rhizobiales; Methylobacteriaceae; Methylobacterium | PAS domain S-box protein | 6 |
| **WP_076729178.1** | 6.2E-05 | Qa.III | IDAIQGTIRTLNAVSSTIAAAVEEQSAVMREMSGSMQTAAEDV---------- | *Methylobacterium radiotolerans* | Bacteria; Proteobacteria; Alphaproteobacteria; Rhizobiales; Methylobacteriaceae; Methylobacterium | PAS domain S-box protein | 6 |
| **WP_080138448.1** | 6.2E-05 | Qa.IV | VKEVATAMRRIDEVTAAIASAVEQQGAATREISQNVQMAA------------- | *Bradyrhizobium sp. BR10280* | Bacteria; Proteobacteria; Alphaproteobacteria; Rhizobiales; Bradyrhizobiaceae; Bradyrhizobium | methyl-accepting chemotaxis protein | 6 |
| **WP_091723535.1** | 6.2E-05 | Qc.I | ----------LRGHASTLEDQVAQRTAQLTEANAELTQALANLKTAQREL--- | *Mitsuaria sp. PDC51* | Bacteria; Proteobacteria; Betaproteobacteria; Burkholderiales; Mitsuaria | HAMP domain-containing protein | 6 |
| **WP_092151755.1** | 6.2E-05 | Qa.IV | -----STIDEVDKISSAIAAAIEEQGAATQEIARNVNQAAARTDD-------- | *Bradyrhizobiaceae* | Bacteria; Proteobacteria; Alphaproteobacteria; Rhizobiales | MULTISPECIES: methyl-accepting chemotaxis protein | 6 |
| **WP_099557359.1** | 6.2E-05 | Qa.III | INSISETVRNINEVSTSIALAVEQQRAATDEIARNVDEAAR------------ | *Hartmannibacter diazotrophicus* | Bacteria; Proteobacteria; Alphaproteobacteria; Rhizobiales; Hartmannibacter | HAMP domain-containing protein | 6 |
| **WP_105329382.1** | 6.2E-05 | Qa.III | IDAIRAVIVNLNEVAQEIAAAVEEQSATTQEIAQSVANANCAVQQVSQS---- | *Blastopirellula* | Bacteria; Planctomycetes; Planctomycetia; Planctomycetales; Planctomycetaceae | MULTISPECIES: methyl-accepting chemotaxis protein | 6 |
| **CRH05505.1** | 6.3E-05 | Qc.III | --QVDEVISRLAETNQAITEAVDDQAHGVQEIQTSMESVTRASDEVTRN---- | *magneto-ovoid bacterium MO-1* | Bacteria | putative Methyl-accepting chemotaxis protein | 6 |
| **GAJ31678.1** | 6.3E-05 | Qa.IV | -----ATIGQISSISTSIASAVEQQGAATQEIARSVQTVAQGTQTAATDI--- | *Bradyrhizobium sp. DOA9* | Bacteria; Proteobacteria; Alphaproteobacteria; Rhizobiales; Bradyrhizobiaceae; Bradyrhizobium | probable chemoreceptor Y4FA | 6 |
| **PKF81266.1** | 6.3E-05 | Qa.III | LDQITNMLTELNDMNLQIATAANEQSSVTDEISSSITSIADASMQ-------- | *Vibrio sp. vnigr-6D03* | Bacteria; Proteobacteria; Gammaproteobacteria; Vibrionales; Vibrionaceae; Vibrio | methyl-accepting chemotaxis protein | 6 |
| **WP_011471720.1** | 6.3E-05 | Qc.III | IEEISATIGKMSEIASTIASAVEEQGAATREISRNVQQAAQGTSQVSSNISD- | *Rhodopseudomonas palustris* | Bacteria; Proteobacteria; Alphaproteobacteria; Rhizobiales; Bradyrhizobiaceae; Rhodopseudomonas | methyl-accepting chemotaxis protein | 6 |
| **WP_045449660.1** | 6.3E-05 | Qc.III | IQAISSTIGSIHEISSTISAAVEEQSTVTKEMSENMQSSAGSVNSIS------ | *Tepidicaulis marinus* | Bacteria; Proteobacteria; Alphaproteobacteria; Rhizobiales; Rhodobiaceae; Tepidicaulis | methyl-accepting chemotaxis protein | 6 |
| **WP_107989163.1** | 6.3E-05 | Qa.IV | INAITETIRQISEISTAISGAVEEQSAATQEIARNVQQAS------------- | *Breoghania corrubedonensis* | Bacteria; Proteobacteria; Alphaproteobacteria; Rhizobiales; Cohaesibacteraceae; Breoghania | chemotaxis protein | 6 |
| **WP_108549063.1** | 6.3E-05 | Qc.III | -KGIGATITQMSGIATAIASAVEEQNAATSEISRNVTQAASGTREVS------ | *Azospirillum humicireducens* | Bacteria; Proteobacteria; Alphaproteobacteria; Rhodospirillales; Rhodospirillaceae; Azospirillum | HAMP domain-containing protein | 6 |
| **WP_109332974.1** | 6.3E-05 | Qa.IV | IRSIGRTITQVNEISSTIAAAVEEQSAATAEIGRNVQN----VAEGAQN---- | *Azospirillum sp. CFH 70021* | Bacteria; Proteobacteria; Alphaproteobacteria; Rhodospirillales; Rhodospirillaceae; Azospirillum | methyl-accepting chemotaxis protein | 6 |
| **WP_112008837.1** | 6.3E-05 | Qb.I | LRRTREMLQSSVEQSNVSTEELKVSNEELQAINEELRSATEELETS------- | *Burkholderia sp. yr520* | Bacteria; Proteobacteria; Betaproteobacteria; Burkholderiales; Burkholderiaceae; Burkholderia | PAS domain S-box protein | 6 |
| **PCI01158.1** | 6.4E-05 | Qc.III | --HIGETIQKMNDINQAISAATDEQNEATREISQSVQYASDATRRVTE----- | *Zetaproteobacteria bacterium* | Bacteria; Proteobacteria; Zetaproteobacteria | hypothetical protein COB79_04610 | 6 |
| **PCJ41358.1** | 6.4E-05 | Qc.III.c | IDKIKKSIDEVQESSDVIRDELARQSDATREITTNVTGARNSVQTVT------ | *Alphaproteobacteria bacterium* | Bacteria; Proteobacteria; Alphaproteobacteria | hypothetical protein COA81_07195 | 6 |
| **PVX50390.1** | 6.4E-05 | Qa.IV | ---ITATIEEVSAIATTIGSAIEEQGAATAEIARNVTQTAQATQD-------- | *Tardiphaga sp. OV697* | Bacteria; Proteobacteria; Alphaproteobacteria; Rhizobiales; Bradyrhizobiaceae; Tardiphaga | methyl-accepting chemotaxis sensory transducer with Cache sensor | 6 |
| **PVX50391.1** | 6.4E-05 | Qa.IV | ---ITATIEEVSAIATTIGSAIEEQGAATAEIARNVTQTAQATQD-------- | *Tardiphaga sp. OV697* | Bacteria; Proteobacteria; Alphaproteobacteria; Rhizobiales; Bradyrhizobiaceae; Tardiphaga | methyl-accepting chemotaxis sensory transducer with Cache sensor partial | 6 |
| **WP_009870940.1** | 6.4E-05 | Qc.III | --GIAQTIAHINEMSASIAGAVEQQGAAASEISRNVEHAAAGTREVVHNI--- | *Magnetospirillum magnetotacticum* | Bacteria; Proteobacteria; Alphaproteobacteria; Rhodospirillales; Rhodospirillaceae; Magnetospirillum | methyl-accepting chemotaxis protein | 6 |
| **WP_052285936.1** | 6.4E-05 | Qc.III | ------QIRKMSDVAASISAAVEEQGVATREIVRNVDQAATGTNTVTSHIAD- | *Azorhizobium caulinodans* | Bacteria; Proteobacteria; Alphaproteobacteria; Rhizobiales; Xanthobacteraceae; Azorhizobium | methyl-accepting chemotaxis protein | 6 |
| **WP_063198350.1** | 6.4E-05 | Qa.IV | VKEVATAMRRIDEVTAAIASAVEQQGAATREISQNVQMAA------------- | *Bradyrhizobium sp. AT1* | Bacteria; Proteobacteria; Alphaproteobacteria; Rhizobiales; Bradyrhizobiaceae; Bradyrhizobium | methyl-accepting chemotaxis protein | 6 |
| **WP_068736633.1** | 6.4E-05 | Qa.IV | ---ITATIEEVSAIATTIGSAIEEQGAATAEIARNVTQTAQATQD-------- | *Tardiphaga robiniae* | Bacteria; Proteobacteria; Alphaproteobacteria; Rhizobiales; Bradyrhizobiaceae; Tardiphaga | HAMP domain-containing protein | 6 |
| **WP_078930352.1** | 6.4E-05 | SNAP.c | INQISSNINNIQTQTLNQAAGVEESSTTINRMIERIDHLNDSISSQK------ | *Treponema berlinense* | Bacteria; Spirochaetes; Spirochaetales; Spirochaetaceae; Treponema | hypothetical protein | 6 |
| **WP_081912816.1** | 6.4E-05 | Qb.I | ----RERLQDALEETQHQASELQAQSEELRVSNEELEEQGAALRESQARL--- | *Sphingobium sp. ba1* | Bacteria; Proteobacteria; Alphaproteobacteria; Sphingomonadales; Sphingomonadaceae; Sphingobium | response regulator | 6 |
| **WP_089264138.1** | 6.4E-05 | Qa.IV | ---ITATIEEVSAIATTIGSAIEEQGAATAEIARNVTQTAQATQD-------- | *Tardiphaga sp. OK246* | Bacteria; Proteobacteria; Alphaproteobacteria; Rhizobiales; Bradyrhizobiaceae; Tardiphaga | HAMP domain-containing protein | 6 |
| **WP_092143463.1** | 6.4E-05 | Qa.IV | ---ITATIEEVSAIATTIGSAIEEQGAATAEIARNVTQTAQATQD-------- | *Bradyrhizobiaceae* | Bacteria; Proteobacteria; Alphaproteobacteria; Rhizobiales | HAMP domain-containing protein | 6 |
| **WP_092192016.1** | 6.4E-05 | Qa.IV | VKEVATAMRRIDEVTAAIASAVEQQGAATREISQNVQMAA------------- | *Bradyrhizobium sp. cf659* | Bacteria; Proteobacteria; Alphaproteobacteria; Rhizobiales; Bradyrhizobiaceae; Bradyrhizobium | methyl-accepting chemotaxis protein | 6 |
| **WP_093758416.1** | 6.4E-05 | Qa.IV | ---ITATIEEVSAIATTIGSAIEEQGAATAEIARNVTQTAQATQD-------- | *Tardiphaga sp. OK245* | Bacteria; Proteobacteria; Alphaproteobacteria; Rhizobiales; Bradyrhizobiaceae; Tardiphaga | HAMP domain-containing protein | 6 |
| **WP_111387029.1** | 6.4E-05 | Qc.III | IKEIGAVIGRIDEIAAAIAAAVQQQGGATREISGNVTRAADGTRQV------- | *Rhodoplanes piscinae* | Bacteria; Proteobacteria; Alphaproteobacteria; Rhizobiales; Hyphomicrobiaceae; Rhodoplanes | chemotaxis protein | 6 |
| **ANC92748.1** | 6.5E-05 | Qa.IV | IRSIAGTIREINEIATTVAAAVEQQSAATREIARNVGEAA------------- | *Azospirillum humicireducens* | Bacteria; Proteobacteria; Alphaproteobacteria; Rhodospirillales; Rhodospirillaceae; Azospirillum | methyl-accepting chemotaxis protein | 6 |
| **AWL93007.1** | 6.5E-05 | Qc.III | -RQISGTIERLSEISSTIAAAIEEQGAATQEISRNVQQAAQGTHGVST----- | *Bradyrhizobium ottawaense* | Bacteria; Proteobacteria; Alphaproteobacteria; Rhizobiales; Bradyrhizobiaceae; Bradyrhizobium | methyl-accepting chemotaxis protein | 6 |
| **CAL76377.1** | 6.5E-05 | Qc.III | ---ISGSIERLSEVSSTIAAAVEEQNAATREISRNVQQAAQGTQQVSANIT-- | *Bradyrhizobium sp. ORS 278* | Bacteria; Proteobacteria; Alphaproteobacteria; Rhizobiales; Bradyrhizobiaceae; Bradyrhizobium | putative methyl-accepting chemotaxis receptor/sensory transducer | 6 |

| | | | | | | | |
|---|---|---|---|---|---|---|---|
| **CCK05163.1** | 6.5E-05 | R.IV | IDQISKAVSELDSTTQQNAALVQESSAAAASLEQQAGELSQLISRF------- | *Cronobacter sakazakii 701* | Bacteria; Proteobacteria; Gammaproteobacteria; Enterobacterales; Enterobacteriaceae; Cronobacter | Methyl-accepting chemotaxis protein I (serine chemoreceptor protein) | 6 |
| **PTW63500.1** | 6.5E-05 | Qa.IV | INAITETIRQISEISTAISGAVEEQSAATQEIARNVQQAS------------- | *Breoghania corrubedonensis* | Bacteria; Proteobacteria; Alphaproteobacteria; Rhizobiales; Cohaesibacteraceae; Breoghania | methyl-accepting chemotaxis protein | 6 |
| **WP_013502962.1** | 6.5E-05 | Qc.III | IQEIGATISRMSEIAATIASAVEEQGAATQEISRNVQQAAQGTQEV------- | *Rhodopseudomonas palustris* | Bacteria; Proteobacteria; Alphaproteobacteria; Rhizobiales; Bradyrhizobiaceae; Rhodopseudomonas | PAS domain S-box protein | 6 |
| **WP_027915586.1** | 6.5E-05 | Qb.I | LQRTRLQLQETIEQSEISSEELTASNEEMQTINEELRSASEELETSK------ | *Pseudomonas* | Bacteria; Proteobacteria; Gammaproteobacteria; Pseudomonadales; Pseudomonadaceae | MULTISPECIES: response regulator | 6 |
| **WP_028687534.1** | 6.5E-05 | Qb.I | LQRTRLQLQETIEQSEISSEELTASNEEMQTINEELRSASEELETSK------ | *Pseudomonas fulva* | Bacteria; Proteobacteria; Gammaproteobacteria; Pseudomonadales; Pseudomonadaceae; Pseudomonas | response regulator | 6 |
| **WP_049695953.1** | 6.5E-05 | Qb.I | LQRTRLQLQETIEQSEISSEELTASNEEMQTINEELRSASEELETSK------ | *Pseudomonas* | Bacteria; Proteobacteria; Gammaproteobacteria; Pseudomonadales; Pseudomonadaceae | MULTISPECIES: response regulator | 6 |
| **WP_059184001.1** | 6.5E-05 | Qb.I | LQRTRLQLQETIEQSEISSEELTASNEEMQTINEELRSASEELETSK------ | *Pseudomonas sp. URMO17WK12:I11* | Bacteria; Proteobacteria; Gammaproteobacteria; Pseudomonadales; Pseudomonadaceae; Pseudomonas | response regulator | 6 |
| **WP_072324657.1** | 6.5E-05 | Qc.III | LDEVASAIHHIADMNTQIASAAEQQAAVADEINRNVVNINELSDQ-------- | *Marinospirillum alkaliphilum* | Bacteria; Proteobacteria; Gammaproteobacteria; Oceanospirillales; Marinospirillum | methyl-accepting chemotaxis protein | 6 |
| **WP_083514155.1** | 6.5E-05 | Qc.III | IKQIGDTIAQISGIATTIAAAVEQQGAATTEIARNVQRASDGTREVSGSITK- | *Bradyrhizobium manausense* | Bacteria; Proteobacteria; Alphaproteobacteria; Rhizobiales; Bradyrhizobiaceae; Bradyrhizobium | methyl-accepting chemotaxis protein | 6 |
| **WP_088523641.1** | 6.5E-05 | Qb.I | LQRTRLQLQETIEQSEISSEELTASNEEMQTINEELRSASEELETSK------ | *Pseudomonas sp. LAIL14HWK12:I4* | Bacteria; Proteobacteria; Gammaproteobacteria; Pseudomonadales; Pseudomonadaceae; Pseudomonas | response regulator | 6 |
| **WP_092954032.1** | 6.5E-05 | Qc.III | ---IRDTIVKISDIAVAIAAAVEQQGAATQDIARSVQAAASGTRDVATRIE-- | *Roseomonas stagni* | Bacteria; Proteobacteria; Alphaproteobacteria; Rhodospirillales; Acetobacteraceae; Roseomonas | HAMP domain-containing protein | 6 |
| **WP_100230111.1** | 6.5E-05 | Qc.III | IGEISGTIARLSEIASAIAAAVEQQGAATQEIARNVQQAARGTQQVSSNVG-- | *Bradyrhizobium sp. INPA54B* | Bacteria; Proteobacteria; Alphaproteobacteria; Rhizobiales; Bradyrhizobiaceae; Bradyrhizobium | PAS domain S-box protein | 6 |
| **WP_104926212.1** | 6.5E-05 | Qb.I | LQRTRLQLQETIEQSEISSEELTASNEEMQTINEELRSASEELETSK------ | *Pseudomonas fulva* | Bacteria; Proteobacteria; Gammaproteobacteria; Pseudomonadales; Pseudomonadaceae; Pseudomonas | response regulator | 6 |
| **WP_110606507.1** | 6.5E-05 | Qb.I | LQRTRLQLQETIEQSEISSEELTASNEEMQTINEELRSASEELETSK------ | *Pseudomonas fulva* | Bacteria; Proteobacteria; Gammaproteobacteria; Pseudomonadales; Pseudomonadaceae; Pseudomonas | response regulator | 6 |
| **AHF04631.1** | 6.6E-05 | Qa.III | -ETISASVQRISDMTNQIASAAEQQSTVTEEINRNITNIADSIE--------- | *Marichromatium purpuratum 984* | Bacteria; Proteobacteria; Gammaproteobacteria; Chromatiales; Chromatiaceae; Marichromatium | methyl-accepting chemotaxis protein | 6 |
| **PSW20118.1** | 6.6E-05 | Qc.III | LEAITASIQHISDMATQISSAAEEQRAVAEEISRNTQAVNDVSDQ-------- | *Photobacterium sanctipauli* | Bacteria; Proteobacteria; Gammaproteobacteria; Vibrionales; Vibrionaceae; Photobacterium | methyl-accepting chemotaxis protein | 6 |
| **PWC90839.1** | 6.6E-05 | Qa.III | IQQISRIIAVIDEAATSIAATVEEQGSATREISRNVQQAAQGTQEV------- | *Azospirillum sp. TSH100* | Bacteria; Proteobacteria; Alphaproteobacteria; Rhodospirillales; Rhodospirillaceae; Azospirillum | hypothetical protein TSH100_02110 | 6 |
| **SMX61913.1** | 6.6E-05 | Qa.IV | IKEIGATITQISDIAATIAATVEQQGAATREIARNVGEAA------------- | *Bradyrhizobium sp. ORS 285* | Bacteria; Proteobacteria; Alphaproteobacteria; Rhizobiales; Bradyrhizobiaceae; Bradyrhizobium | putative methyl-accepting chemotaxis receptor/sensory transducer | 6 |
| **WP_002726012.1** | 6.6E-05 | Qa.IV | ----------ISELSSAIAAAVEEQGAATSEIARSVQQAAQGTAAAAENVEQ- | *Phaeospirillum molischianum* | Bacteria; Proteobacteria; Alphaproteobacteria; Rhodospirillales; Rhodospirillaceae; Phaeospirillum | PAS domain S-box protein | 6 |
| **WP_050471018.1** | 6.6E-05 | Qa.III.b | IQEIAETMGEVNQFTTTIADAISQQGEATREITVNVQRAA------------- | *Pannonibacter phragmitetus* | Bacteria; Proteobacteria; Alphaproteobacteria; Rhodobacterales; Rhodobacteraceae; Pannonibacter | methyl-accepting chemotaxis protein | 6 |
| **WP_052709990.1** | 6.6E-05 | Qa.III | IVGIGRTIHSINQIAAGISAGIEQQGSAVREIVVNMEQAAA------------ | *Azospirillum thiophilum* | Bacteria; Proteobacteria; Alphaproteobacteria; Rhodospirillales; Rhodospirillaceae; Azospirillum | chemotaxis protein | 6 |
| **WP_058900436.1** | 6.6E-05 | Qa.III.b | IQEIAETMGEVNQFTTTIADAISQQGEATREITVNVQRAA------------- | *Pannonibacter phragmitetus* | Bacteria; Proteobacteria; Alphaproteobacteria; Rhodobacterales; Rhodobacteraceae; Pannonibacter | methyl-accepting chemotaxis protein | 6 |
| **WP_084306806.1** | 6.6E-05 | Qc.III | LEDITQSIGTINDMNTQIASAAEEQSAVAEDVHRSLTRISTGV---------- | *Ectothiorhodospira haloalkaliphila* | Bacteria; Proteobacteria; Gammaproteobacteria; Chromatiales; Ectothiorhodospiraceae; Ectothiorhodospira | methyl-accepting chemotaxis protein | 6 |
| **WP_094462170.1** | 6.6E-05 | Qa.III.b | IQEIAETMGEVNQFTTTIADAISQQGEATREITVNVQRAA------------- | *Pannonibacter phragmitetus* | Bacteria; Proteobacteria; Alphaproteobacteria; Rhodobacterales; Rhodobacteraceae; Pannonibacter | methyl-accepting chemotaxis protein | 6 |
| **WP_109110807.1** | 6.6E-05 | Qa.III | -QQVVRVIGHINEVATSIASAVEEQGAATREISRNVQQAAQ------------ | *Azospirillum sp. TSO35-2* | Bacteria; Proteobacteria; Alphaproteobacteria; Rhodospirillales; Rhodospirillaceae; Azospirillum | HAMP domain-containing protein | 6 |
| **ALV76085.1** | 6.7E-05 | Qc.III | LESITQAVSVINDMNTQIASAAEEQSAVAEDINRNVANANQ------------ | *Pseudomonas aeruginosa* | Bacteria; Proteobacteria; Gammaproteobacteria; Pseudomonadales; Pseudomonadaceae; Pseudomonas | Methyl-accepting chemotaxis protein PctC | 6 |
| **BAF88710.1** | 6.7E-05 | Qc.III | ------QIRKMSDVAASISAAVEEQGVATREIVRNVDQAATGTNTVTSHIAD- | *Azorhizobium caulinodans ORS 571* | Bacteria; Proteobacteria; Alphaproteobacteria; Rhizobiales; Xanthobacteraceae; Azorhizobium | histidine kinase | 6 |
| **CCK09698.1** | 6.7E-05 | R.IV | IDQISKAVSELDSTTQQNAALVQESSAAAASLEQQAGELSQLISRF------- | *Cronobacter sakazakii 696* | Bacteria; Proteobacteria; Gammaproteobacteria; Enterobacterales; Enterobacteriaceae; Cronobacter | Methyl-accepting chemotaxis protein I (serine chemoreceptor protein) | 6 |
| **KZD06403.1** | 6.7E-05 | Qc.III | -SDIAQVISEINEIATVIASAVEEQNAATQEIARNVQQAASGTQEV------- | *Oceanibaculum pacificum* | Bacteria; Proteobacteria; Alphaproteobacteria; Rhodospirillales; Rhodospirillaceae; Oceanibaculum | hypothetical protein AUP43_10795 | 6 |
| **PCI41549.1** | 6.7E-05 | Qc.III | IQGIGSTIGKIDEASTAIASAVEEQGAATQEIARNVEQAANGTNEVTSNITE- | *Rhodospirillaceae bacterium* | Bacteria; Proteobacteria; Alphaproteobacteria; Rhodospirillales; Rhodospirillaceae | chemotaxis protein | 6 |
| **SCM74749.1** | 6.7E-05 | Qa.III | IVGITSIIEQLNAIASSIAVAVDQQGAATNEIARNVAQASSGTHQVSQNIV-- | *uncultured Pleomorphomonas sp.* | Bacteria; Proteobacteria; Alphaproteobacteria; Rhizobiales; Methylocystaceae; Pleomorphomonas; environmental samples | putative Chemotaxis sensory transducer | 6 |
| **WP_011662981.1** | 6.7E-05 | Qc.III | IKEISDTIGRMSEIASTIASAVEEQGAATQEISRNVQQAAH------------ | *Rhodopseudomonas palustris* | Bacteria; Proteobacteria; Alphaproteobacteria; Rhizobiales; Bradyrhizobiaceae; Rhodopseudomonas | methyl-accepting chemotaxis protein | 6 |
| **WP_012170941.1** | 6.7E-05 | Qc.III | -KEIGDTINRIAGIAQAIAAAVEQQGAATLEISRNIQQAAAGTEQVSAN---- | *Azorhizobium caulinodans* | Bacteria; Proteobacteria; Alphaproteobacteria; Rhizobiales; Xanthobacteraceae; Azorhizobium | methyl-accepting chemotaxis protein | 6 |
| **WP_013657865.1** | 6.7E-05 | SNAP.c | ------VADEIRKLATNTANAVTEVNTLIDNIQGALKNVNTHLESSNKQIE-- | *Cellulosilyticum lentocellum* | Bacteria; Firmicutes; Clostridia; Clostridiales; Lachnospiraceae; Cellulosilyticum | methyl-accepting chemotaxis sensory transducer | 6 |
| **WP_021130422.1** | 6.7E-05 | Qa.III | ---IVERIEEINQIATTIASAVEEQSAATNEIARNVQQAAIGTQEVSNTI--- | *Phaeospirillum fulvum* | Bacteria; Proteobacteria; Alphaproteobacteria; Rhodospirillales; Rhodospirillaceae; Phaeospirillum | methyl-accepting chemotaxis protein | 6 |
| **WP_034463411.1** | 6.7E-05 | Qa.IV | -EAITQTISRINEISAAIAASVEEQSSVTQNISENMQTAAR------------ | *Afipia sp. P52-10* | Bacteria; Proteobacteria; Alphaproteobacteria; Rhizobiales; Bradyrhizobiaceae; Afipia | methyl-accepting chemotaxis protein | 6 |
| **WP_053006072.1** | 6.7E-05 | Qc.III | IEGISNIINQVNEIASGISAAVEEQGAATQEIARNTQEAARGTQEVS------ | *Kiloniella spongiae* | Bacteria; Proteobacteria; Alphaproteobacteria; Kiloniellales; Kiloniellaceae; Kiloniella | methyl-accepting chemotaxis protein | 6 |
| **WP_054085688.1** | 6.7E-05 | Qa.III | --GISEAVANITDMTTQIATATEEQSSVAEEINRNISTIAELADKTADEA--- | *Pseudomonas syringae group* | Bacteria; Proteobacteria; Gammaproteobacteria; Pseudomonadales; Pseudomonadaceae; Pseudomonas | MULTISPECIES: PAS domain S-box protein | 6 |
| **WP_060737165.1** | 6.7E-05 | Qc.III | -QTIGGIIGEVNEVATAIAAAVQEQGAATQEITRSTQYAAQGTRNVS------ | *Bradyrhizobium sp. CCGE-LA001* | Bacteria; Proteobacteria; Alphaproteobacteria; Rhizobiales; Bradyrhizobiaceae; Bradyrhizobium | methyl-accepting chemotaxis protein | 6 |
| **WP_072671445.1** | 6.7E-05 | Qa.III | LDGIVQQVKLINEMNLQIATAAGQQSSVAVEINQNITNVRELVEASV------ | *Vibrio sp. M12-1144* | Bacteria; Proteobacteria; Gammaproteobacteria; Vibrionales; Vibrionaceae; Vibrio | methyl-accepting chemotaxis protein | 6 |
| **WP_072817911.1** | 6.7E-05 | Qa.III.b | IKGIESIIMEINQIATTTASAVEEQGAATAEIARNVQQAA------------- | *Bradyrhizobium erythrophlei* | Bacteria; Proteobacteria; Alphaproteobacteria; Rhizobiales; Bradyrhizobiaceae; Bradyrhizobium | HAMP domain-containing protein | 6 |
| **WP_085339864.1** | 6.7E-05 | Qc.III | ---ISQVIGSISEVSTAIAAAIEEQSALTESMSANMGVASGSVQAISARM--- | *Aquidulcibacter paucihalophilus* | Bacteria; Proteobacteria; Alphaproteobacteria; Caulobacterales; Caulobacteraceae; Aquidulcibacter | methyl-accepting chemotaxis protein | 6 |
| **WP_093811060.1** | 6.7E-05 | Qc.III.c | -DNIMSIINQMSEIATSISAAVEEQSVVSSEISANMQSVAEGVGTVTSTIE-- | *Stappia sp. ES.058* | Bacteria; Proteobacteria; Alphaproteobacteria; Rhodobacterales; Rhodobacteraceae; Stappia | methyl-accepting chemotaxis protein | 6 |
| **WP_100079043.1** | 6.7E-05 | Qa.III | IVGITSIIEQLNAIASSIAVAVDQQGAATNEIARNVAQASSGTHQVSQNIV-- | *Pleomorphomonas sp. SVCO-16* | Bacteria; Proteobacteria; Alphaproteobacteria; Rhizobiales; Methylocystaceae; Pleomorphomonas | methyl-accepting chemotaxis protein | 6 |
| **WP_101923117.1** | 6.7E-05 | Qc.III.c | ---IAASIGQVNDQARGIASAATQQGEVTHDIAERMSRTAGRVGQSN------ | *Tabrizicola sp. TH137* | Bacteria; Proteobacteria; Alphaproteobacteria; Rhodobacterales; Rhodobacteraceae; Tabrizicola | methyl-accepting chemotaxis protein | 6 |
| **WP_110167810.1** | 6.7E-05 | Qa.III | LDGIVQQVKLINEMNLQIATAAGQQSSVAVEINQNITNVRELVEASV------ | *Vibrio sp. 11986-1-5* | Bacteria; Proteobacteria; Gammaproteobacteria; Vibrionales; Vibrionaceae; Vibrio | methyl-accepting chemotaxis protein | 6 |
| **WP_110786269.1** | 6.7E-05 | Qa.III | IREIGATISKMSEIASTIASAVEEQGAATQEISRNVQQAAQGTQQVS------ | *Rhodopseudomonas palustris* | Bacteria; Proteobacteria; Alphaproteobacteria; Rhizobiales; Bradyrhizobiaceae; Rhodopseudomonas | HAMP domain-containing protein | 6 |
| **CBL46702.1** | 6.8E-05 | SNAP.c | ------VADEVRKLATETQETLKRTNHAVERIIVNINETASKTDSNAQRVQ-- | *gamma proteobacterium HdN1* | Bacteria; Proteobacteria; Gammaproteobacteria | hypothetical protein HDN1F_31190 | 6 |
| **KPW74903.1** | 6.8E-05 | Qa.III | --GISEAVANITDMTTQIATATEEQSSVAEEINRNISTIAELADKTADEA--- | *Pseudomonas syringae pv. coriandricola* | Bacteria; Proteobacteria; Gammaproteobacteria; Pseudomonadales; Pseudomonadaceae; Pseudomonas | Chemotaxis protein | 6 |
| **OQW49686.1** | 6.8E-05 | Qa.IV | --EITATISDISEISNAIASAVDEQNSATSEISRSIQTVTTQT---------- | *Proteobacteria bacterium SG_bin8* | Bacteria; Proteobacteria | hypothetical protein A4S15_02955 | 6 |
| **WP_009027341.1** | 6.8E-05 | Qc.III | IQRIGGIIGEVNEVATAIAAAVQQQGAATQEISRSTQFAADGTRNVS------ | *Bradyrhizobium sp. ORS 375* | Bacteria; Proteobacteria; Alphaproteobacteria; Rhizobiales; Bradyrhizobiaceae; Bradyrhizobium | methyl-accepting chemotaxis protein | 6 |
| **WP_035691627.1** | 6.8E-05 | Qc.III | --GVGATINHINEIATTIAAAVEQQAAATREIARNVQQAAEGTQSVT------ | *Azospirillum halopraeferens* | Bacteria; Proteobacteria; Alphaproteobacteria; Rhodospirillales; Rhodospirillaceae; Azospirillum | chemotaxis protein | 6 |
| **WP_083763622.1** | 6.8E-05 | Qc.III | -------IEEINHIATAISAAVEEQSAATAEIARNVDQASQGTRQVSANV--- | *Magnetospirillum magneticum* | Bacteria; Proteobacteria; Alphaproteobacteria; Rhodospirillales; Rhodospirillaceae; Magnetospirillum | HAMP domain-containing protein | 6 |
| **WP_084291943.1** | 6.8E-05 | Qc.III | IQEIGGTIERLSEVSSAIAAAVEEQGAATQEISRNVQQASL------------ | *Bradyrhizobium sp. WSM3983* | Bacteria; Proteobacteria; Alphaproteobacteria; Rhizobiales; Bradyrhizobiaceae; Bradyrhizobium | methyl-accepting chemotaxis protein | 6 |
| **WP_106749575.1** | 6.8E-05 | Qc.III | IKEIGSTIDHLAGIAAAIAAAVEEQGAATQEISRNVQEAAQ------------ | *Phreatobacter cathodiphilus* | Bacteria; Proteobacteria; Alphaproteobacteria; Phreatobacter | methyl-accepting chemotaxis protein | 6 |
| **WP_114010222.1** | 6.8E-05 | Qa.III | IQGIAETMEKVNEYTKTISLAVDEQGSATYEISQNVARAAS------------ | *Cohaesibacter sp. YE-B6* | Bacteria; Proteobacteria; Alphaproteobacteria; Rhizobiales; Cohaesibacteraceae | HAMP domain-containing protein | 6 |
| **BAF87223.1** | 6.9E-05 | Qc.III | --SIAGTIGRMNDISGSIASAVTQQQAATSEIAGSIARASEGTREANENMA-- | *Azorhizobium caulinodans ORS 571* | Bacteria; Proteobacteria; Alphaproteobacteria; Rhizobiales; Xanthobacteraceae; Azorhizobium | methyl-accepting chemotaxis sensory transducer | 6 |
| **EKD39041.1** | 6.9E-05 | Qa.III | IEEITTVINNVNEIVATITTAVGEQSKATEEIARNINQAAE------------ | *uncultured bacterium* | Bacteria; environmental samples | hypothetical protein ACD_75C00522G0002 | 6 |
| **WP_013503254.1** | 6.9E-05 | Qa.III | ISEISGTIEKLSEISAHIASAVEEQGAATQEISRNIQQTAH------------ | *Rhodopseudomonas palustris* | Bacteria; Proteobacteria; Alphaproteobacteria; Rhizobiales; Bradyrhizobiaceae; Rhodopseudomonas | chemotaxis protein | 6 |

| | | | | | | | | |
|---|---|---|---|---|---|---|---|---|
|  | WP_046074139.1 | 6.9E-05 | SNAP.c | ------GSEEVASLAASTARAVEEQNRTLDSVASAVEELSTAVRDV------- | *Salinivibrio sp. KP-1* | Bacteria; Proteobacteria; Gammaproteobacteria; Vibrionales; Vibrionaceae; Salinivibrio | methyl-accepting chemotaxis protein | 6 |
| | WP_054360059.1 | 6.9E-05 | Qc.III | --SIADVIGKISEISSGIAAAVEEQTAVTRDMSENMQTASKGVEVISRNMN-- | *Prosthecomicrobium hirschii* | Bacteria; Proteobacteria; Alphaproteobacteria; Rhizobiales; Hyphomicrobiaceae; Prosthecomicrobium | methyl-accepting chemotaxis protein | 6 |
|  | WP_077578906.1 | 6.9E-05 | SNAP.c | ------GSEEVASLAASTARAVEEQNRTLDSVASAVEELSTAVRDV------- | *Salinivibrio* | Bacteria; Proteobacteria; Gammaproteobacteria; Vibrionales; Vibrionaceae | MULTISPECIES: methyl-accepting chemotaxis protein | 6 |
|  | WP_077607300.1 | 6.9E-05 | SNAP.c | ------GSEEVASLAASTARAVEEQNRTLDSVASAVEELSTAVRDV------- | *Salinivibrio sp. ML290* | Bacteria; Proteobacteria; Gammaproteobacteria; Vibrionales; Vibrionaceae; Salinivibrio | methyl-accepting chemotaxis protein | 6 |
|  | WP_077666540.1 | 6.9E-05 | SNAP.c | ------GSEEVASLAASTARAVEEQNRTLDSVASAVEELSTAVRDV------- | *Salinivibrio sp. PR6* | Bacteria; Proteobacteria; Gammaproteobacteria; Vibrionales; Vibrionaceae; Salinivibrio | methyl-accepting chemotaxis protein | 6 |
|  | WP_077667305.1 | 6.9E-05 | SNAP.c | ------GSEEVASLAASTARAVEEQNRTLDSVASAVEELSTAVRDV------- | *Salinivibrio siamensis* | Bacteria; Proteobacteria; Gammaproteobacteria; Vibrionales; Vibrionaceae; Salinivibrio | methyl-accepting chemotaxis protein | 6 |
|  | WP_077771301.1 | 6.9E-05 | SNAP.c | ------GSEEVASLAASTARAVEEQNRTLDSVASAVEELSTAVRDV------- | *Salinivibrio sharmensis* | Bacteria; Proteobacteria; Gammaproteobacteria; Vibrionales; Vibrionaceae; Salinivibrio | methyl-accepting chemotaxis protein | 6 |
|  | WP_080670211.1 | 6.9E-05 | Qc.III | -RDIGGTIARLSEIAAAIAAAVEEQGAATQEITRNVQQAAKG----------- | *Bradyrhizobium* | Bacteria; Proteobacteria; Alphaproteobacteria; Rhizobiales; Bradyrhizobiaceae | MULTISPECIES: HAMP domain-containing protein | 6 |
|  | WP_081494429.1 | 6.9E-05 | Qc.III | -RDIGGTIARLSEIAAAIAAAVEEQGAATQEITRNVQQAAKG----------- | *Bradyrhizobium* | Bacteria; Proteobacteria; Alphaproteobacteria; Rhizobiales; Bradyrhizobiaceae | MULTISPECIES: HAMP domain-containing protein | 6 |
|  | WP_083859283.1 | 6.9E-05 | Qc.III | -RDIGGTIARLSEIAAAIAAAVEEQGAATQEITRNVQQAAKG----------- | *Bradyrhizobium sp. CCBAU 15544* | Bacteria; Proteobacteria; Alphaproteobacteria; Rhizobiales; Bradyrhizobiaceae; Bradyrhizobium | HAMP domain-containing protein | 6 |
|  | WP_084462781.1 | 6.9E-05 | Qc.III | -SDIAQVISEINEIATVIASAVEEQNAATQEIARNVQQAASGTQEV------- | *Oceanibaculum pacificum* | Bacteria; Proteobacteria; Alphaproteobacteria; Rhodospirillales; Rhodospirillaceae; Oceanibaculum | HAMP domain-containing protein | 6 |
|  | WP_097669843.1 | 6.9E-05 | Qc.III | -RDIGGTIARLSEIAAAIAAAVEEQGAATQEITRNVQQAAKG----------- | *Bradyrhizobium ottawaense* | Bacteria; Proteobacteria; Alphaproteobacteria; Rhizobiales; Bradyrhizobiaceae; Bradyrhizobium | HAMP domain-containing protein | 6 |
| | WP_109327184.1 | 6.9E-05 | Qc.III | ---ISGTITNINDISTTIAAAIEEQGAATNEIARSVQQAAGGTQEVSNNI--- | *Azospirillum sp. CFH 70021* | Bacteria; Proteobacteria; Alphaproteobacteria; Rhodospirillales; Rhodospirillaceae; Azospirillum | methyl-accepting chemotaxis protein | 6 |
|  | PZQ17097.1 | 7E-05 | Qa.III | --AIARTISEVSEIASSIAAAVGQQAAVTKEIAATAQSVARNVDSVR------ | *Starkeya novella* | Bacteria; Proteobacteria; Alphaproteobacteria; Rhizobiales; Xanthobacteraceae; Starkeya | chemotaxis protein | 6 |
|  | SKB31854.1 | 7E-05 | Qa.IV | ---------SLNHISGMVAVVIEQQNAATAGIAENIRQAADETIAAASHI--- | *Bosea thiooxidans* | Bacteria; Proteobacteria; Alphaproteobacteria; Rhizobiales; Bradyrhizobiaceae; Bosea | methyl-accepting chemotaxis protein | 6 |
| | WP_027276781.1 | 7E-05 | Qc.III | IKEIGDTIGKMSEIASTIASAVEQQGAATQEISRNVQQAAGGTHQVSIHITD- | *Rhodopseudomonas palustris* | Bacteria; Proteobacteria; Alphaproteobacteria; Rhizobiales; Bradyrhizobiaceae; Rhodopseudomonas | HAMP domain-containing protein | 6 |
|  | WP_028167779.1 | 7E-05 | Qa.III | IEGIGGTIGGINEITTVIAAAVEQQGSATKEIARNVQQAA------------- | *Bradyrhizobium elkanii* | Bacteria; Proteobacteria; Alphaproteobacteria; Rhizobiales; Bradyrhizobiaceae; Bradyrhizobium | methyl-accepting chemotaxis protein | 6 |
|  | WP_036489416.1 | 7E-05 | Qa.III.b | IRDIGGVIGEINAISTEISAAVDQQGGATQEIAHNVSEAAKGTQEVS------ | *Nesiotobacter exalbescens* | Bacteria; Proteobacteria; Alphaproteobacteria; Rhodobacterales; Rhodobacteraceae; Nesiotobacter | methyl-accepting chemotaxis protein | 6 |
|  | WP_045003286.1 | 7E-05 | Qc.III | -KAIGGIIGEVNEVATAIAAAVEEQGAATQEITRSTQQAAQGTKNV------- | *Bradyrhizobium sp. LTSP857* | Bacteria; Proteobacteria; Alphaproteobacteria; Rhizobiales; Bradyrhizobiaceae; Bradyrhizobium | methyl-accepting chemotaxis protein | 6 |
|  | WP_057862572.1 | 7E-05 | Qc.III | -KGIGSIIGEVNEVATAIAAAVQQQGAATQEITRSTQFAAQGTKNVS------ | *Bradyrhizobium lablabi* | Bacteria; Proteobacteria; Alphaproteobacteria; Rhizobiales; Bradyrhizobiaceae; Bradyrhizobium | methyl-accepting chemotaxis protein | 6 |
| | WP_100176881.1 | 7E-05 | Qc.III | -LSISKTIQRLSEIASVIAAAVEEQSAATLEISRNVQQAAEGTRQVSSNVA-- | *Bradyrhizobium sp. TSA1* | Bacteria; Proteobacteria; Alphaproteobacteria; Rhizobiales; Bradyrhizobiaceae; Bradyrhizobium | methyl-accepting chemotaxis protein | 6 |
|  | WP_111462178.1 | 7E-05 | Qa.III | --QIAQSLRNLTGLNASIASATLQQSHVVDDINQNVTQTAALAHSTA------ | *Pseudomonas sp. URMO17WK12:I2* | Bacteria; Proteobacteria; Gammaproteobacteria; Pseudomonadales; Pseudomonadaceae; Pseudomonas | methyl-accepting chemotaxis protein | 6 |
|  | GAJ34363.1 | 7.1E-05 | Qa.IV | IKTIAGTIGEISSISGSIASAVDQQTATTQEIARNAQHAA------------- | *Bradyrhizobium sp. DOA9* | Bacteria; Proteobacteria; Alphaproteobacteria; Rhizobiales; Bradyrhizobiaceae; Bradyrhizobium | protein pilJ | 6 |
|  | KRQ16862.1 | 7.1E-05 | Qa.IV | -----SNISRISQIASTIAAAIEEQGTATRHIAENIQHAAEMSSQVATNI--- | *Bradyrhizobium manausense* | Bacteria; Proteobacteria; Alphaproteobacteria; Rhizobiales; Bradyrhizobiaceae; Bradyrhizobium | hypothetical protein AOQ71_04340 | 6 |
| | PWC81506.1 | 7.1E-05 | Qc.III | IDDISGIITGINETATTIASAVEEQGAATREIARNVQQAANGAQEVVSRV--- | *Azospirillum sp. TSH100* | Bacteria; Proteobacteria; Alphaproteobacteria; Rhodospirillales; Rhodospirillaceae; Azospirillum | chemotaxis protein partial | 6 |
| | SDR32018.1 | 7.1E-05 | Qc.III.c | --EIGLTIDQINSLSTGINEAVGEQKQATEDISRSLTDAHQNVSQVASNM--- | *Pseudovibrio sp. Tun.PSC04-5.I4* | Bacteria; Proteobacteria; Alphaproteobacteria; Rhodobacterales; Rhodobacteraceae; Pseudovibrio | methyl-accepting chemotaxis protein | 6 |
|  | WP_008974633.1 | 7.1E-05 | Qc.III | -KAIGRTIERISEIAASISSAVEQQRAATENITQSVRFAASGTAEVVS----- | *Bradyrhizobium sp. STM 3843* | Bacteria; Proteobacteria; Alphaproteobacteria; Rhizobiales; Bradyrhizobiaceae; Bradyrhizobium | methyl-accepting chemotaxis protein | 6 |
|  | WP_055727976.1 | 7.1E-05 | Qa.IV | ---------SLNHISGMVAVVIEQQNAATAGIAENIRQAADETIAAASHI--- | *Bosea thiooxidans* | Bacteria; Proteobacteria; Alphaproteobacteria; Rhizobiales; Bradyrhizobiaceae; Bosea | hypothetical protein | 6 |
|  | WP_078482880.1 | 7.1E-05 | Qa.III | -DTITQAVSNINDMNAQIANAANEQSTVVEEINEHVNTISEVNELT------- | *Solemya velesiana gill symbiont* | Bacteria; Proteobacteria; Gammaproteobacteria; sulfur-oxidizing symbionts | PAS domain S-box protein | 6 |
| | WP_084798448.1 | 7.1E-05 | Qa.III | IVGITSIIEQLNAIASSIAAAVDQQGAATNEIARNVSQASAGTHQVSQNIV-- | *Pleomorphomonas koreensis* | Bacteria; Proteobacteria; Alphaproteobacteria; Rhizobiales; Methylocystaceae; Pleomorphomonas | methyl-accepting chemotaxis protein | 6 |
|  | WP_090022033.1 | 7.1E-05 | Qa.III.b | --DISNVISEINEIATTMSSSVEEQIASASEISQNISTAAENAERVAE----- | *Limimonas halophila* | Bacteria; Proteobacteria; Alphaproteobacteria; Rhodospirillales; Rhodospirillaceae; Limimonas | methyl-accepting chemotaxis protein | 6 |
|  | WP_092781445.1 | 7.1E-05 | Qa.IV | -------IEQLNETTTAISTAVEQQSAATKEISQNAQQAAHNTQDASENID-- | *Rhodospira trueperi* | Bacteria; Proteobacteria; Alphaproteobacteria; Rhodospirillales; Rhodospirillaceae; Rhodospira | methyl-accepting chemotaxis protein | 6 |
|  | WP_106751869.1 | 7.1E-05 | Qc.III | IQGIGEIIRRINSASTAISAAVEEQGAATQEIARNVNQAAYGTQQ-------- | *Pannonibacter carbonis* | Bacteria; Proteobacteria; Alphaproteobacteria; Rhodobacterales; Rhodobacteraceae; Pannonibacter | HAMP domain-containing protein | 6 |
|  | WP_111955467.1 | 7.1E-05 | Qa.III | IKEIVKIIDDVNSIVSSVAAAIEEQSATTQEISNNVSQAATGVQEVNE----- | *Desulfobacter hydrogenophilus* | Bacteria; Proteobacteria; Deltaproteobacteria; Desulfobacterales; Desulfobacteraceae; Desulfobacter | methyl-accepting chemotaxis protein | 6 |
|  | BAR60156.1 | 7.2E-05 | Qa.III.b | -EAIAQTIAEIDGITASIATAIEQQGLATREIAHNVQEASA------------ | *Bradyrhizobium diazoefficiens* | Bacteria; Proteobacteria; Alphaproteobacteria; Rhizobiales; Bradyrhizobiaceae; Bradyrhizobium | putative methyl accepting chemotaxis protein | 6 |
|  | OUM94022.1 | 7.2E-05 | Qc.III | ------SAGELATGANQISEDVSQMSESISEIESKVAAYADSTKQMNER---- | *Thermobacillus sp. ZCTH02-B1* | Bacteria; Firmicutes; Bacilli; Bacillales; Paenibacillaceae; Thermobacillus | chemotaxis protein | 6 |
|  | PSQ48867.1 | 7.2E-05 | Qc.II | -DALEDSVEQATESAEVIDERVEEQTDQLSEVATQMENFSASM---------- | *Halobacteriales archaeon SW_6_65_15* | Archaea; Euryarchaeota; Halobacteria; Halobacteriales | chemotaxis protein | 6 |
|  | WP_019645338.1 | 7.2E-05 | Qc.III | ---ISGTIGRISQISTTIASAVEEQSAATGEIARNAEQAAVGTERVTV----- | *Novispirillum itersonii* | Bacteria; Proteobacteria; Alphaproteobacteria; Rhodospirillales; Rhodospirillaceae; Novispirillum | methyl-accepting chemotaxis protein | 6 |
| | WP_022665203.1 | 7.2E-05 | Qc.III | IEKISLVINDVNDKVVNITQAIEEQSKITNQIADNVSNASQGIQEVTENI--- | *Desulfospira joergensenii* | Bacteria; Proteobacteria; Deltaproteobacteria; Desulfobacterales; Desulfobacteraceae; Desulfospira | methyl-accepting chemotaxis protein | 6 |
|  | WP_024512187.1 | 7.2E-05 | Qa.III | IQGIGGVVAQINEITSAIAAAVEEQGAATSDIAANIQQAA------------- | *Bradyrhizobium sp. ARR65* | Bacteria; Proteobacteria; Alphaproteobacteria; Rhizobiales; Bradyrhizobiaceae; Bradyrhizobium | methyl-accepting chemotaxis protein | 6 |
|  | WP_045586080.1 | 7.2E-05 | Qc.III | --EIARVVHDLSEISASIAAAVEEQNAATGEIGRGVTEAAQGVR--------- | *Azospirillum thiophilum* | Bacteria; Proteobacteria; Alphaproteobacteria; Rhodospirillales; Rhodospirillaceae; Azospirillum | HAMP domain-containing protein | 6 |
|  | WP_051283887.1 | 7.2E-05 | Qa.IV | -------ISETSDIVSTIASAIEEQSTATREIASNINNVAKNVYDANTS---- | *Desulforegula conservatrix* | Bacteria; Proteobacteria; Deltaproteobacteria; Desulfobacterales; Desulfobacteraceae; Desulforegula | methyl-accepting chemotaxis protein | 6 |
|  | WP_056474708.1 | 7.2E-05 | Qc.III | --GIMEVIQRLGRISGDISAAVDQQESATQEIARNTVLTSEGTRAVSGH---- | *Methylobacterium sp. Leaf104* | Bacteria; Proteobacteria; Alphaproteobacteria; Rhizobiales; Methylobacteriaceae; Methylobacterium | methyl-accepting chemotaxis protein | 6 |
|  | WP_085372001.1 | 7.2E-05 | Qc.III | -GSISSTIQQINQLTTTIAGAVEEQGAATGEIARNVEEASKGTREVATN---- | *Magnetospirillum sp. ME-1* | Bacteria; Proteobacteria; Alphaproteobacteria; Rhodospirillales; Rhodospirillaceae; Magnetospirillum | methyl-accepting chemotaxis protein | 6 |
|  | WP_092207233.1 | 7.2E-05 | Qa.III | MQEITTVIGEVNDIVSTIAGAVEEQSAATSEIAENVSQAS------------- | *Desulfoluna spongiiphila* | Bacteria; Proteobacteria; Deltaproteobacteria; Desulfobacterales; Desulfobacteraceae; Desulfoluna | methyl-accepting chemotaxis protein | 6 |
|  | PVX50393.1 | 7.3E-05 | Qa.IV | ---ITATIEEVSAIATTIGSAIEEQGAATAEIARNVTQTAQATQN-------- | *Tardiphaga sp. OV697* | Bacteria; Proteobacteria; Alphaproteobacteria; Rhizobiales; Bradyrhizobiaceae; Tardiphaga | methyl-accepting chemotaxis sensory transducer with Cache sensor | 6 |
| | WP_006033830.1 | 7.3E-05 | SNAP.c | IDEINLSVNQIKEIAKDTSAASFQQTNTLDEIQTNVNDVNQVTEENTTRA--- | *Moritella sp. PE36* | Bacteria; Proteobacteria; Gammaproteobacteria; Alteromonadales; Moritellaceae; Moritella | HAMP domain-containing protein | 6 |
|  | WP_092252107.1 | 7.3E-05 | Qa.III | --EITGTIRKLSEISSTIAAAVEEQGAATQEIARNVQQAASGTQ--------- | *Bradyrhizobium sp. Rc3b* | Bacteria; Proteobacteria; Alphaproteobacteria; Rhizobiales; Bradyrhizobiaceae; Bradyrhizobium | chemotaxis protein | 6 |
|  | WP_109084164.1 | 7.3E-05 | Qa.III | IQQISRIIAVIDEAATSIAATVEEQGSATREISRNVQQAAQGTQEV------- | *Azospirillum sp. TSH100* | Bacteria; Proteobacteria; Alphaproteobacteria; Rhodospirillales; Rhodospirillaceae; Azospirillum | methyl-accepting chemotaxis protein | 6 |
|  | WP_008550363.1 | 7.4E-05 | SNAP.c | -GDVSDVIHEMSQIASAVAASVEQQNAAIQAITENVANATQ------------ | *Pseudovibrio sp. JE062* | Bacteria; Proteobacteria; Alphaproteobacteria; Rhodobacterales; Rhodobacteraceae; Pseudovibrio | HAMP domain-containing protein | 6 |
|  | WP_014283016.1 | 7.4E-05 | SNAP.c | -GDVSDVIHEMSQIASAVAASVEQQNAAIQAITENVANATQ------------ | *Pseudovibrio sp. FO-BEG1* | Bacteria; Proteobacteria; Alphaproteobacteria; Rhodobacterales; Rhodobacteraceae; Pseudovibrio | HAMP domain-containing protein | 6 |
|  | WP_029583486.1 | 7.4E-05 | Qc.III | IKMISSTIGKISEITGAISAAIEEQGAATQEISGNIQRTAD------------ | *Bradyrhizobium sp. URHD0069* | Bacteria; Proteobacteria; Alphaproteobacteria; Rhizobiales; Bradyrhizobiaceae; Bradyrhizobium | hypothetical protein | 6 |
|  | WP_032552634.1 | 7.4E-05 | Qa.III | LEQIREAIQVISDMATQIASAAEEQSLVTAEINTNTESVRE------------ | *Vibrio fortis* | Bacteria; Proteobacteria; Gammaproteobacteria; Vibrionales; Vibrionaceae; Vibrio | methyl-accepting chemotaxis protein | 6 |
| | WP_040426652.1 | 7.4E-05 | Qc.III | --GISETIDRISSIAAAIAAAVEQQGAATNEIANNVQQAARGTARVTSNIA-- | *Bradyrhizobiaceae* | Bacteria; Proteobacteria; Alphaproteobacteria; Rhizobiales | MULTISPECIES: PAS domain-containing protein | 6 |
|  | WP_057461825.1 | 7.4E-05 | Qc.III.c | ---IGETIEQINFLSTGINEAVGDQKQATEDISRDLNSAHQNVSQV------- | *Pseudovibrio sp. POLY-S9* | Bacteria; Proteobacteria; Alphaproteobacteria; Rhodobacterales; Rhodobacteraceae; Pseudovibrio | methyl-accepting chemotaxis protein | 6 |
|  | WP_079446886.1 | 7.4E-05 | Qa.III | IREIGGTIEKLSEIASIIAAAVEEQGAATQEISRNVQQASQGTQQ-------- | *Nitrobacter vulgaris* | Bacteria; Proteobacteria; Alphaproteobacteria; Rhizobiales; Bradyrhizobiaceae; Nitrobacter | HAMP domain-containing protein | 6 |
|  | WP_082885148.1 | 7.4E-05 | Qc.III | IKEIGATIGRISEIAAAISTSVEQQGSATQEISRNVQRAATGTSQ-------- | *Bradyrhizobium stylosanthis* | Bacteria; Proteobacteria; Alphaproteobacteria; Rhizobiales; Bradyrhizobiaceae; Bradyrhizobium | HAMP domain-containing protein | 6 |
|  | WP_083416765.1 | 7.4E-05 | SNAP.c | -GDVSDVIHEMSQIASAVAASVEQQNAAIQAITENVANATQ------------ | *Pseudovibrio denitrificans* | Bacteria; Proteobacteria; Alphaproteobacteria; Rhodobacterales; Rhodobacteraceae; Pseudovibrio | HAMP domain-containing protein | 6 |
|  | WP_085940714.1 | 7.4E-05 | Qa.IV | ----------LNDIAASIAAAVEEQGAATSEIARNVQQAAGGTAQASQNI--- | *Azospirillum sp. B506* | Bacteria; Proteobacteria; Alphaproteobacteria; Rhodospirillales; Rhodospirillaceae; Azospirillum | HAMP domain-containing protein | 6 |
|  | WP_109155812.1 | 7.4E-05 | Qc.III | --GIGGTIGRINDITTGIAAAIEEQSAATNEISRNVQQAAAGTQEVTGS---- | *Azospirillum* | Bacteria; Proteobacteria; Alphaproteobacteria; Rhodospirillales; Rhodospirillaceae | MULTISPECIES: methyl-accepting chemotaxis protein | 6 |

| | | | | | | | | |
|---|---|---|---|---|---|---|---|---|
|  | WP_109865129.1 | 7.4E-05 | Qa.IV | ----------LNDIAASIAAAVEEQGAATSEIARNVQQAAGGTAQASQNI--- | *Azospirillum sp. TSH100* | Bacteria; Proteobacteria; Alphaproteobacteria; Rhodospirillales; Rhodospirillaceae; Azospirillum | HAMP domain-containing protein | 6 |
|  | AGA57031.1 | 7.5E-05 | Qc.III | ------SAGELATGANQISEDVSQMSESISEIESKVANYADSTRQMNER---- | *Thermobacillus composti KWC4* | Bacteria; Firmicutes; Bacilli; Bacillales; Paenibacillaceae; Thermobacillus | methyl-accepting chemotaxis protein | 6 |
|  | EGH72626.1 | 7.5E-05 | Qa.III | ---ISQALRNLNSLNASIASATLQQTHVVDDINQNVTQAAQLSQ--------- | *Pseudomonas syringae pv. aceris str. M302273* | Bacteria; Proteobacteria; Gammaproteobacteria; Pseudomonadales; Pseudomonadaceae; Pseudomonas; Pseudomonas syringae | histidine kinase HAMP region: chemotaxis sensory transducer partial | 6 |
|  | KGG88092.1 | 7.5E-05 | Qa.III | MKEIMQSVGQVTGLLSEISGALQQQSEGI----AHVNTAVAHMDSTNQE---- | *Comamonas testosteroni* | Bacteria; Proteobacteria; Betaproteobacteria; Burkholderiales; Comamonadaceae; Comamonas | methyl-accepting chemotaxis protein | 6 |
|  | KGH11332.1 | 7.5E-05 | Qa.III | MKEIMQSVGQVTGLLSEISGALQQQSEGI----AHVNTAVAHMDSTNQE---- | *Comamonas testosteroni* | Bacteria; Proteobacteria; Betaproteobacteria; Burkholderiales; Comamonadaceae; Comamonas | methyl-accepting chemotaxis protein | 6 |
|  | KGH29848.1 | 7.5E-05 | Qa.III | MKEIMQSVGQVTGLLSEISGALQQQSEGI----AHVNTAVAHMDSTNQE---- | *Comamonas testosteroni* | Bacteria; Proteobacteria; Betaproteobacteria; Burkholderiales; Comamonadaceae; Comamonas | methyl-accepting chemotaxis protein | 6 |
|  | OGB79284.1 | 7.5E-05 | SNAP.b | ---SLKKIQEIVTNSSAMVRNLSSQSEQIVGVVDTIKNISEQT---------- | *candidate division CPR3 bacterium RIFOXYB2_FULL_35_8* | Bacteria; candidate division CPR3 | hypothetical protein A2296_03910 | 6 |
|  | PWC80299.1 | 7.5E-05 | Qa.IV | ----------LNDIAASIAAAVEEQGAATSEIARNVQQAAGGTAQASQNI--- | *Azospirillum sp. TSH100* | Bacteria; Proteobacteria; Alphaproteobacteria; Rhodospirillales; Rhodospirillaceae; Azospirillum | chemotaxis protein | 6 |
|  | PYL45500.1 | 7.5E-05 | Qb.I | ----KQHLNATIEQYEAANEELKASNEELQAMNEEMRSTTEELETSK------ | *Verrucomicrobia bacterium* | Bacteria; Verrucomicrobia; unclassified Verrucomicrobia | hypothetical protein DMF40_15355 | 6 |
| | WP_009031130.1 | 7.5E-05 | Qc.III | ---ISGSIERLSEVSSTIAAAVEEQNAATREISRNVQQAAQGTQQVSANIT-- | *Bradyrhizobium sp. ORS 375* | Bacteria; Proteobacteria; Alphaproteobacteria; Rhizobiales; Bradyrhizobiaceae; Bradyrhizobium | methyl-accepting chemotaxis protein | 6 |
| | WP_011661961.1 | 7.5E-05 | Qc.III | IKEIGATITRISEIAGVIAATVEEQGAATSEIARNVGEAAKGTAEVASKI--- | *Rhodopseudomonas palustris* | Bacteria; Proteobacteria; Alphaproteobacteria; Rhizobiales; Bradyrhizobiaceae; Rhodopseudomonas | chemotaxis protein | 6 |
|  | WP_014414012.1 | 7.5E-05 | Qa.III | -SGIVGRIVEINEIATAIASAVEEQSAATAEIARNVQQTSEA----------- | *Pararhodospirillum photometricum* | Bacteria; Proteobacteria; Alphaproteobacteria; Rhodospirillales; Rhodospirillaceae; Pararhodospirillum | HAMP domain-containing protein | 6 |
|  | WP_028349299.1 | 7.5E-05 | Qc.III | -KGIGSIIGEVNEVATAIAAAVQQQGAATQEITRSTQFAAQGTKNVS------ | *Bradyrhizobium elkanii* | Bacteria; Proteobacteria; Alphaproteobacteria; Rhizobiales; Bradyrhizobiaceae; Bradyrhizobium | methyl-accepting chemotaxis protein | 6 |
|  | WP_073014508.1 | 7.5E-05 | Qa.III | -------INEMNDIARSISSAVEEQGTATQEIAINATEASRSTVEVTTSI--- | *Labrenzia suaedae* | Bacteria; Proteobacteria; Alphaproteobacteria; Rhodobacterales; Rhodobacteraceae; Labrenzia | methyl-accepting chemotaxis protein | 6 |
|  | WP_074819632.1 | 7.5E-05 | Qc.III | -KGIGSIIGEVNEVATAIAAAVQEQGAATQEITRSTQYAAQGTKNV------- | *Bradyrhizobium* | Bacteria; Proteobacteria; Alphaproteobacteria; Rhizobiales; Bradyrhizobiaceae | MULTISPECIES: methyl-accepting chemotaxis protein | 6 |
|  | WP_080138389.1 | 7.5E-05 | Qa.IV | -----ATIGQISSISTSIASAVEQQGAATQEIARSVQTVAQGTQTAATDI--- | *Bradyrhizobium sp. BR10280* | Bacteria; Proteobacteria; Alphaproteobacteria; Rhizobiales; Bradyrhizobiaceae; Bradyrhizobium | methyl-accepting chemotaxis protein partial | 6 |
|  | WP_080138922.1 | 7.5E-05 | Qa.IV | -----ATIGQISSISTSIASAVEQQGAATQEIARSVQTVAQGTQTAATDI--- | *Bradyrhizobium sp. BR10280* | Bacteria; Proteobacteria; Alphaproteobacteria; Rhizobiales; Bradyrhizobiaceae; Bradyrhizobium | methyl-accepting chemotaxis protein partial | 6 |
|  | WP_083222820.1 | 7.5E-05 | R.II | IQQISSDVQQTTSLLSNSVEKADVANEKIGHLANASDRISE------------ | *Terasakiella sp. PR1* | Bacteria; Proteobacteria; Alphaproteobacteria; Rhizobiales; Methylocystaceae; Terasakiella | methyl-accepting chemotaxis protein | 6 |
|  | WP_106409328.1 | 7.5E-05 | Qa.IV | ------TIAEVSSTTVSIASAVEEQTAATAEIARTVQATAEATEQVTVNI--- | *Roseomonas deserti* | Bacteria; Proteobacteria; Alphaproteobacteria; Rhodospirillales; Acetobacteraceae; Roseomonas | methyl-accepting chemotaxis protein | 6 |
|  | WP_109096252.1 | 7.5E-05 | Qa.IV | ----------LNDIAASIAAAVEEQGAATSEIARNVQQAAGGTAQASQNI--- | *Azospirillum sp. TSH64* | Bacteria; Proteobacteria; Alphaproteobacteria; Rhodospirillales; Rhodospirillaceae; Azospirillum | HAMP domain-containing protein | 6 |
|  | WP_111417882.1 | 7.5E-05 | Qc.III | IKEISGTIRQIAEVASAIAAAVEEQGAATSEISRNIQQAAA------------ | *Rhodoplanes roseus* | Bacteria; Proteobacteria; Alphaproteobacteria; Rhizobiales; Hyphomicrobiaceae; Rhodoplanes | methyl-accepting chemotaxis protein | 6 |
|  | WP_114354410.1 | 7.5E-05 | Qc.III | ----SLSINEVKDNVDTIIDKIDLATKSIETIGEKIDTSAEQINKTVEKFDE- | *Saliterribacillus persicus* | Bacteria; Firmicutes; Bacilli; Bacillales; Bacillaceae; Saliterribacillus | PAS domain S-box protein | 6 |
| | PVX48411.1 | 7.6E-05 | Qc.III | IKEIGSTIGRMSEIASTIASAVEEQGAATQEISRNVQQAARGTTDVSSNI--- | *Tardiphaga sp. OV697* | Bacteria; Proteobacteria; Alphaproteobacteria; Rhizobiales; Bradyrhizobiaceae; Tardiphaga | methyl-accepting chemotaxis protein | 6 |
|  | WP_018707133.1 | 7.6E-05 | Qb.III.d | IASAMKEITEKIKANMDILQTLNVQTEAIRSIVETIGNIS------------- | *Bacillus fordii* | Bacteria; Firmicutes; Bacilli; Bacillales; Bacillaceae; Bacillus | PAS domain S-box protein | 6 |
|  | WP_083742408.1 | 7.6E-05 | Qa.III | IMEITATIEKLSEVSSTIAAAVEEQGTATLEISGNVQQASHGTQQ-------- | *Bradyrhizobium mercantei* | Bacteria; Proteobacteria; Alphaproteobacteria; Rhizobiales; Bradyrhizobiaceae; Bradyrhizobium | methyl-accepting chemotaxis protein | 6 |
|  | WP_099043995.1 | 7.6E-05 | Qa.IV | --EIEANVNILSQGINEMSQSINEQTEAINQINEAVATVDEQTKQ-N------ | *Campylobacter fetus* | Bacteria; Proteobacteria; Epsilonproteobacteria; Campylobacterales; Campylobacteraceae; Campylobacter | chemotaxis protein partial | 6 |
|  | WP_111359421.1 | 7.6E-05 | Qc.III | IKEIGSTIGRIATIASTIAATVEAQGAVTSEIGREVQEAS------------- | *Rhodoplanes elegans* | Bacteria; Proteobacteria; Alphaproteobacteria; Rhizobiales; Hyphomicrobiaceae; Rhodoplanes | methyl-accepting chemotaxis protein | 6 |
| | AWM03360.1 | 7.7E-05 | Qc.III | IKMISSTIKSLSEISSTIAAAVEEQGAATQEIARNVQQAAQGTHQVSSNISD- | *Bradyrhizobium sp. 2 39S1MB* | Bacteria; Proteobacteria; Alphaproteobacteria; Rhizobiales; Bradyrhizobiaceae; Bradyrhizobium | PAS domain S-box protein | 6 |
| | AWM09586.1 | 7.7E-05 | Qc.III | IKMISSTIKSLSEISSTIAAAVEEQGAATQEIARNVQQAAQGTHQVSSNISD- | *Bradyrhizobium sp. 3 85S1MB* | Bacteria; Proteobacteria; Alphaproteobacteria; Rhizobiales; Bradyrhizobiaceae; Bradyrhizobium | PAS domain S-box protein | 6 |
|  | CCK00690.1 | 7.7E-05 | R.IV | IDQISKAVSELDSTTQQNAALVQESSAAAASLEQQAGELSQLISRF------- | *Cronobacter malonaticus 507* | Bacteria; Proteobacteria; Gammaproteobacteria; Enterobacterales; Enterobacteriaceae; Cronobacter | Methyl-accepting chemotaxis protein I (serine chemoreceptor protein) | 6 |
|  | CUT09432.1 | 7.7E-05 | Qa.IV | -----ATIGQISSISTSIASAVEQQGAATQEIARSVQTVAQGTQTAATDI--- | *Bradyrhizobium sp.* | Bacteria; Proteobacteria; Alphaproteobacteria; Rhizobiales; Bradyrhizobiaceae; Bradyrhizobium | Methylaccepting chemotaxis protein | 6 |
| | ERF85014.1 | 7.7E-05 | Qa.III | ISDIAGIIQRIAQIASDVADAIDAQGSATRSIAENVQVASERTSQVAVSIGQ- | *Bradyrhizobium sp. DFCI-1* | Bacteria; Proteobacteria; Alphaproteobacteria; Rhizobiales; Bradyrhizobiaceae; Bradyrhizobium | hypothetical protein C207_01594 | 6 |
|  | KPY04879.1 | 7.7E-05 | Qa.III | ---ISQALRNLNSLNASIASATLQQTHVVDDINQNVTQAAQLSQ--------- | *Pseudomonas coronafaciens pv. oryzae* | Bacteria; Proteobacteria; Gammaproteobacteria; Pseudomonadales; Pseudomonadaceae; Pseudomonas; Pseudomonas coronafaciens | Histidine kinase HAMP region: chemotaxis sensory transducer | 6 |
|  | PHR89510.1 | 7.7E-05 | Qc.III | LDSMTNSVMQINDLGIQIATAAEEQSSVTEEINRNMTMIQNMVSQLTE----- | *Moritella sp.* | Bacteria; Proteobacteria; Gammaproteobacteria; Alteromonadales; Moritellaceae; Moritella | methyl-accepting chemotaxis protein | 6 |
|  | WP_039799816.1 | 7.7E-05 | Qa.IV | -----ATIGQISSISTSIASAVEQQGAATQEIARSVQTVAQGTQTAATDI--- | *Bradyrhizobium sp. WSM4349* | Bacteria; Proteobacteria; Alphaproteobacteria; Rhizobiales; Bradyrhizobiaceae; Bradyrhizobium | methyl-accepting chemotaxis protein partial | 6 |
|  | WP_045006636.1 | 7.7E-05 | Qa.IV | -----ATIGQISSISTSIASAVEQQGAATQEIARSVQTVAQGTQTAATDI--- | *Bradyrhizobium sp. LTSP857* | Bacteria; Proteobacteria; Alphaproteobacteria; Rhizobiales; Bradyrhizobiaceae; Bradyrhizobium | methyl-accepting chemotaxis protein partial | 6 |
|  | WP_045006637.1 | 7.7E-05 | Qa.IV | -----ATIGQISSISTSIASAVEQQGAATQEIARSVQTVAQGTQTAATDI--- | *Bradyrhizobium sp. LTSP857* | Bacteria; Proteobacteria; Alphaproteobacteria; Rhizobiales; Bradyrhizobiaceae; Bradyrhizobium | methyl-accepting chemotaxis protein partial | 6 |
|  | WP_049662530.1 | 7.7E-05 | Qb.III.d | IQNSEDLLETVKEVSQESAdnvrnlAQLQKQAESIKGIVKTIQEIA--------- | *Lysinibacillus xylanilyticus* | Bacteria; Firmicutes; Bacilli; Bacillales; Bacillaceae; Lysinibacillus | PAS domain S-box protein | 6 |
|  | WP_053482978.1 | 7.7E-05 | Qb.III.d | IQNSEDLLETVKEVSQESAdnvrnlAQLQKQAESIKGIVKTIQEIA--------- | *Lysinibacillus sp. FJAT-14745* | Bacteria; Firmicutes; Bacilli; Bacillales; Bacillaceae; Lysinibacillus | PAS domain S-box protein | 6 |
|  | WP_079564070.1 | 7.7E-05 | Qb.III.d | IQNSEDLLETVKEVSQESAdnvrnlAQLQKQAESIKGIVKTIQEIA--------- | *Lysinibacillus* | Bacteria; Firmicutes; Bacilli; Bacillales; Bacillaceae | MULTISPECIES: PAS domain S-box protein | 6 |
|  | WP_081434035.1 | 7.7E-05 | Qa.III | -SQIQRVVTDIKQISSSIAGAVEEQGAATGEIAQNCQQAATGTQ--------- | *Azorhizobium caulinodans* | Bacteria; Proteobacteria; Alphaproteobacteria; Rhizobiales; Xanthobacteraceae; Azorhizobium | methyl-accepting chemotaxis protein | 6 |
|  | WP_082646499.1 | 7.7E-05 | Qa.IV | ---IERTIGLIGDTTALINTTITQQASATDEIAQNVSQAAAHTE--------- | *Bradyrhizobium valentinum* | Bacteria; Proteobacteria; Alphaproteobacteria; Rhizobiales; Bradyrhizobiaceae; Bradyrhizobium | hypothetical protein | 6 |
|  | WP_105535826.1 | 7.7E-05 | R.IV | IDQISKAVSELDSTTQQNAALVQESSAAAASLEQQAGELSQLISRF------- | *Cronobacter sakazakii* | Bacteria; Proteobacteria; Gammaproteobacteria; Enterobacterales; Enterobacteriaceae; Cronobacter | chemotaxis protein | 6 |
|  | OJY09515.1 | 7.8E-05 | Qc.III | --IISSTIGQISEITTAISAAIEQQGAATQEISSNIQRTAAG----------- | *Rhizobiales bacterium 62-47* | Bacteria; Proteobacteria; Alphaproteobacteria; Rhizobiales | hypothetical protein BGP05_17065 | 6 |
|  | OYY09573.1 | 7.8E-05 | Qa.IV | -------ISEIRQSTTAIATAIEEQGAATREIAQNTQHAANGTQGVNDNIQ-- | *Rhizobiales bacterium 35-68-8* | Bacteria; Proteobacteria; Alphaproteobacteria; Rhizobiales | hypothetical protein B7Y70_09890 | 6 |
|  | WP_020400251.1 | 7.8E-05 | Qa.III | MKSIRRGITTIAEMASGVASAVEEQRAATDEIARNVQEAAT------------ | *Kordiimonas gwangyangensis* | Bacteria; Proteobacteria; Alphaproteobacteria; Kordiimonadales; Kordiimonadaceae; Kordiimonas | methyl-accepting chemotaxis protein | 6 |
|  | WP_027535720.1 | 7.8E-05 | Qc.III | IGEISGTIEKLSEISSTIAAAVEEQGAATQEISRNVQQTAEG----------- | *Bradyrhizobium sp. WSM3983* | Bacteria; Proteobacteria; Alphaproteobacteria; Rhizobiales; Bradyrhizobiaceae; Bradyrhizobium | chemotaxis protein | 6 |
|  | WP_030140614.1 | 7.8E-05 | Qb.I | LQRTKLHLQDNIEQAEVSSEELKASNEEMQAINEELRSATEELETS------- | *Pseudomonas fluorescens* | Bacteria; Proteobacteria; Gammaproteobacteria; Pseudomonadales; Pseudomonadaceae; Pseudomonas | PAS domain S-box protein | 6 |
|  | WP_052832043.1 | 7.8E-05 | Qa.IV | ---------RMSEVSSGIASAVEQQDATTRDISANLQQAGIFTSGVNANL--- | *Skermanella aerolata* | Bacteria; Proteobacteria; Alphaproteobacteria; Rhodospirillales; Rhodospirillaceae; Skermanella | methyl-accepting chemotaxis protein | 6 |
|  | WP_055026142.1 | 7.8E-05 | Qc.I | -------AEQLKRSYEQLEDKVEKRTKAIAEINDNLQRQIMQRKKIEA----- | *Shewanella sp. P1-14-1* | Bacteria; Proteobacteria; Gammaproteobacteria; Alteromonadales; Shewanellaceae; Shewanella | diguanylate cyclase | 6 |
|  | WP_065108845.1 | 7.8E-05 | Qc.I | -------AEQLKRSYEQLEDKVEKRTKAIAEINDNLQRQIMQRKKIEA----- | *Shewanella sp. UCD-FRSSP16_17* | Bacteria; Proteobacteria; Gammaproteobacteria; Alteromonadales; Shewanellaceae; Shewanella | diguanylate cyclase | 6 |
| | WP_076400140.1 | 7.8E-05 | Qc.III | IDEIVGIIEDINQVASSIATAVEQQNAATREIARNVEQAAAGTGEVSSNIQ-- | *Insolitispirillum peregrinum* | Bacteria; Proteobacteria; Alphaproteobacteria; Rhodospirillales; Rhodospirillaceae; Insolitispirillum | HAMP domain-containing protein | 6 |
|  | WP_080917001.1 | 7.8E-05 | Qc.I | -------AEQLKRSYEQLEDKVEKRTKAIAEINDNLQRQIMQRKKIEA----- | *Shewanella japonica* | Bacteria; Proteobacteria; Gammaproteobacteria; Alteromonadales; Shewanellaceae; Shewanella | diguanylate cyclase | 6 |
| | WP_106940543.1 | 7.8E-05 | Qa.III | ISDIAGIIQRIAQIASDVADAIDAQGSATRSIAENVQVASERTSQVAVSIGQ- | *Bradyrhizobium sp. MOS004* | Bacteria; Proteobacteria; Alphaproteobacteria; Rhizobiales; Bradyrhizobiaceae; Bradyrhizobium | HAMP domain-containing protein | 6 |
| | WP_106945089.1 | 7.8E-05 | Qc.III | --AISNTIEKLSEISSTIAAAVEEQGAATQEISRNVQQAAEGTHQVSANIT-- | *Bradyrhizobium sp. MOS002* | Bacteria; Proteobacteria; Alphaproteobacteria; Rhizobiales; Bradyrhizobiaceae; Bradyrhizobium | methyl-accepting chemotaxis protein | 6 |
|  | AWL98693.1 | 7.9E-05 | Qa.IV | -----ATIGQISSISTSIASAVEQQGAATQEIARSVQTVAQGTQTAATDI--- | *Bradyrhizobium sp. 2 39S1MB* | Bacteria; Proteobacteria; Alphaproteobacteria; Rhizobiales; Bradyrhizobiaceae; Bradyrhizobium | methyl-accepting chemotaxis protein | 6 |
|  | AWL98694.1 | 7.9E-05 | Qa.IV | -----ATIGQISSISTSIASAVEQQGAATQEIARSVQTVAQGTQTAATDI--- | *Bradyrhizobium sp. 2 39S1MB* | Bacteria; Proteobacteria; Alphaproteobacteria; Rhizobiales; Bradyrhizobiaceae; Bradyrhizobium | methyl-accepting chemotaxis protein | 6 |
|  | AWM10751.1 | 7.9E-05 | Qa.IV | -----ATIGQISSISTSIASAVEQQGAATQEIARSVQTVAQGTQTAATDI--- | *Bradyrhizobium sp. 3 85S1MB* | Bacteria; Proteobacteria; Alphaproteobacteria; Rhizobiales; Bradyrhizobiaceae; Bradyrhizobium | methyl-accepting chemotaxis protein | 6 |

| | | | | | | | | |
|---|---|---|---|---|---|---|---|---|
|  | AWM10752.1 | 7.9E-05 | Qa.IV | -----ATIGQISSISTSIASAVEQQGAATQEIARSVQTVAQGTQTAATDI--- | *Bradyrhizobium sp. 3 85S1MB* | Bacteria; Proteobacteria; Alphaproteobacteria; Rhizobiales; Bradyrhizobiaceae; Bradyrhizobium | methyl-accepting chemotaxis protein | 6 |
|  | BAF89479.1 | 7.9E-05 | Qa.III | -SQIQRVVTDIKQISSSIAGAVEEQGAATGEIAQNCQQAATGTQ--------- | *Azorhizobium caulinodans ORS 571* | Bacteria; Proteobacteria; Alphaproteobacteria; Rhizobiales; Xanthobacteraceae; Azorhizobium | IMP dehydrogenase | 6 |
|  | WP_008137663.1 | 7.9E-05 | Qa.IV | -----ATIGQISSISTSIASAVEQQGAATQEIARSVQTVAQGTQTAATDI--- | *Bradyrhizobium sp. YR681* | Bacteria; Proteobacteria; Alphaproteobacteria; Rhizobiales; Bradyrhizobiaceae; Bradyrhizobium | methyl-accepting chemotaxis protein | 6 |
|  | WP_018646254.1 | 7.9E-05 | Qa.IV | -----ATIGQISSISTSIASAVEQQGAATQEIARSVQTVAQGTQTAATDI--- | *Bradyrhizobium japonicum* | Bacteria; Proteobacteria; Alphaproteobacteria; Rhizobiales; Bradyrhizobiaceae; Bradyrhizobium | methyl-accepting chemotaxis protein | 6 |
|  | WP_025032731.1 | 7.9E-05 | Qa.IV | -----ATIGQISSISTSIASAVEQQGAATQEIARSVQTVAQGTQTAATDI--- | *Bradyrhizobium sp. DOA9* | Bacteria; Proteobacteria; Alphaproteobacteria; Rhizobiales; Bradyrhizobiaceae; Bradyrhizobium | methyl-accepting chemotaxis protein | 6 |
|  | WP_026312500.1 | 7.9E-05 | Qa.IV | -----ATIGQISSISTSIASAVEQQGAATQEIARSVQTVAQGTQTAATDI--- | *Bradyrhizobium japonicum* | Bacteria; Proteobacteria; Alphaproteobacteria; Rhizobiales; Bradyrhizobiaceae; Bradyrhizobium | methyl-accepting chemotaxis protein | 6 |
|  | WP_027515100.1 | 7.9E-05 | Qa.IV | -----ATIGQISSISTSIASAVEQQGAATQEIARSVQTVAQGTQTAATDI--- | *Bradyrhizobium sp. WSM1417* | Bacteria; Proteobacteria; Alphaproteobacteria; Rhizobiales; Bradyrhizobiaceae; Bradyrhizobium | methyl-accepting chemotaxis protein | 6 |
|  | WP_027515101.1 | 7.9E-05 | Qa.IV | -----ATIGQISSISTSIASAVEQQGAATQEIARSVQTVAQGTQTAATDI--- | *Bradyrhizobium sp. WSM1417* | Bacteria; Proteobacteria; Alphaproteobacteria; Rhizobiales; Bradyrhizobiaceae; Bradyrhizobium | methyl-accepting chemotaxis protein | 6 |
|  | WP_027533453.1 | 7.9E-05 | Qa.IV | -----ATIGQISSISTSIASAVEQQGAATQEIARSVQTVAQGTQTAATDI--- | *Bradyrhizobium sp. WSM3983* | Bacteria; Proteobacteria; Alphaproteobacteria; Rhizobiales; Bradyrhizobiaceae; Bradyrhizobium | methyl-accepting chemotaxis protein | 6 |
|  | WP_027546739.1 | 7.9E-05 | Qa.IV | -----ATIGQISSISTSIASAVEQQGAATQEIARSVQTVAQGTQTAATDI--- | *Bradyrhizobium sp. WSM2254* | Bacteria; Proteobacteria; Alphaproteobacteria; Rhizobiales; Bradyrhizobiaceae; Bradyrhizobium | methyl-accepting chemotaxis protein | 6 |
|  | WP_027563541.1 | 7.9E-05 | Qa.IV | -----ATIGQISSISTSIASAVEQQGAATQEIARSVQTVAQGTQTAATDI--- | *Bradyrhizobium genosp. SA-4* | Bacteria; Proteobacteria; Alphaproteobacteria; Rhizobiales; Bradyrhizobiaceae; Bradyrhizobium | methyl-accepting chemotaxis protein | 6 |
|  | WP_027563542.1 | 7.9E-05 | Qa.IV | -----ATIGQISSISTSIASAVEQQGAATQEIARSVQTVAQGTQTAATDI--- | *Bradyrhizobium* | Bacteria; Proteobacteria; Alphaproteobacteria; Rhizobiales; Bradyrhizobiaceae | MULTISPECIES: methyl-accepting chemotaxis protein | 6 |
|  | WP_027568324.1 | 7.9E-05 | Qa.IV | -----ATIGQISSISTSIASAVEQQGAATQEIARSVQTVAQGTQTAATDI--- | *Bradyrhizobium sp. URHA0013* | Bacteria; Proteobacteria; Alphaproteobacteria; Rhizobiales; Bradyrhizobiaceae; Bradyrhizobium | methyl-accepting chemotaxis protein | 6 |
|  | WP_028096867.1 | 7.9E-05 | Qc.III | -------IGRQLENSTKIASAATEQAKSTNKLVESLSAATEKIGTVVRLIND- | *Dongia sp. URHE0060* | Bacteria; Proteobacteria; Alphaproteobacteria; Rhodospirillales; Rhodospirillaceae; Dongia | methyl-accepting chemotaxis protein | 6 |
|  | WP_028133156.1 | 7.9E-05 | Qa.IV | -----ATIGQISSISTSIASAVEQQGAATQEIARSVQTVAQGTQTAATDI--- | *Bradyrhizobium japonicum* | Bacteria; Proteobacteria; Alphaproteobacteria; Rhizobiales; Bradyrhizobiaceae; Bradyrhizobium | methyl-accepting chemotaxis protein | 6 |
|  | WP_028149279.1 | 7.9E-05 | Qa.IV | -----ATIGQISSISTSIASAVEQQGAATQEIARSVQTVAQGTQTAATDI--- | *Bradyrhizobium japonicum* | Bacteria; Proteobacteria; Alphaproteobacteria; Rhizobiales; Bradyrhizobiaceae; Bradyrhizobium | methyl-accepting chemotaxis protein | 6 |
|  | WP_028159623.1 | 7.9E-05 | Qa.IV | -----ATIGQISSISTSIASAVEQQGAATQEIARSVQTVAQGTQTAATDI--- | *Bradyrhizobium japonicum* | Bacteria; Proteobacteria; Alphaproteobacteria; Rhizobiales; Bradyrhizobiaceae; Bradyrhizobium | methyl-accepting chemotaxis protein | 6 |
|  | WP_038952154.1 | 7.9E-05 | Qa.IV | -----ATIGQISSISTSIASAVEQQGAATQEIARSVQTVAQGTQTAATDI--- | *Bradyrhizobium japonicum* | Bacteria; Proteobacteria; Alphaproteobacteria; Rhizobiales; Bradyrhizobiaceae; Bradyrhizobium | methyl-accepting chemotaxis protein | 6 |
|  | WP_038959064.1 | 7.9E-05 | Qa.IV | -----ATIGQISSISTSIASAVEQQGAATQEIARSVQTVAQGTQTAATDI--- | *Bradyrhizobium japonicum* | Bacteria; Proteobacteria; Alphaproteobacteria; Rhizobiales; Bradyrhizobiaceae; Bradyrhizobium | methyl-accepting chemotaxis protein | 6 |
|  | WP_039155437.1 | 7.9E-05 | Qa.IV | -----ATIGQISSISTSIASAVEQQGAATQEIARSVQTVAQGTQTAATDI--- | *Bradyrhizobium japonicum* | Bacteria; Proteobacteria; Alphaproteobacteria; Rhizobiales; Bradyrhizobiaceae; Bradyrhizobium | methyl-accepting chemotaxis protein | 6 |
|  | WP_045011999.1 | 7.9E-05 | Qa.IV | -----ATIGQISSISTSIASAVEQQGAATQEIARSVQTVAQGTQTAATDI--- | *Bradyrhizobium sp. LTSP849* | Bacteria; Proteobacteria; Alphaproteobacteria; Rhizobiales; Bradyrhizobiaceae; Bradyrhizobium | methyl-accepting chemotaxis protein | 6 |
|  | WP_054700054.1 | 7.9E-05 | Qa.III | ------MIAEINEIVANIASAMEEQSTSTREVSSNVNQAAQGIQEANV----- | *Desulfosarcina cetonica* | Bacteria; Proteobacteria; Deltaproteobacteria; Desulfobacterales; Desulfobacteraceae; Desulfosarcina | hypothetical protein | 6 |
|  | WP_057027412.1 | 7.9E-05 | Qa.IV | -----ATIGQISSISTSIASAVEQQGAATQEIARSVQTVAQGTQTAATDI--- | *Bradyrhizobium yuanmingense* | Bacteria; Proteobacteria; Alphaproteobacteria; Rhizobiales; Bradyrhizobiaceae; Bradyrhizobium | methyl-accepting chemotaxis protein partial | 6 |
|  | WP_063692751.1 | 7.9E-05 | Qa.IV | -----ATIGQISSISTSIASAVEQQGAATQEIARSVQTVAQGTQTAATDI--- | *Bradyrhizobium stylosanthis* | Bacteria; Proteobacteria; Alphaproteobacteria; Rhizobiales; Bradyrhizobiaceae; Bradyrhizobium | methyl-accepting chemotaxis protein | 6 |
|  | WP_063692756.1 | 7.9E-05 | Qa.IV | -----ATIGQISSISTSIASAVEQQGAATQEIARSVQTVAQGTQTAATDI--- | *Bradyrhizobium stylosanthis* | Bacteria; Proteobacteria; Alphaproteobacteria; Rhizobiales; Bradyrhizobiaceae; Bradyrhizobium | methyl-accepting chemotaxis protein | 6 |
|  | WP_063980506.1 | 7.9E-05 | Qa.IV | -----ATIGQISSISTSIASAVEQQGAATQEIARSVQTVAQGTQTAATDI--- | *Bradyrhizobium sp.* | Bacteria; Proteobacteria; Alphaproteobacteria; Rhizobiales; Bradyrhizobiaceae; Bradyrhizobium | methyl-accepting chemotaxis protein | 6 |
|  | WP_063992420.1 | 7.9E-05 | Qa.IV | -----ATIGQISSISTSIASAVEQQGAATQEIARSVQTVAQGTQTAATDI--- | *Bradyrhizobium sp.* | Bacteria; Proteobacteria; Alphaproteobacteria; Rhizobiales; Bradyrhizobiaceae; Bradyrhizobium | methyl-accepting chemotaxis protein | 6 |
|  | WP_063992421.1 | 7.9E-05 | Qa.IV | -----ATIGQISSISTSIASAVEQQGAATQEIARSVQTVAQGTQTAATDI--- | *Bradyrhizobium sp.* | Bacteria; Proteobacteria; Alphaproteobacteria; Rhizobiales; Bradyrhizobiaceae; Bradyrhizobium | methyl-accepting chemotaxis protein | 6 |
|  | WP_071908553.1 | 7.9E-05 | Qa.IV | -----ATIGQISSISTSIASAVEQQGAATQEIARSVQTVAQGTQTAATDI--- | *Bradyrhizobium japonicum* | Bacteria; Proteobacteria; Alphaproteobacteria; Rhizobiales; Bradyrhizobiaceae; Bradyrhizobium | methyl-accepting chemotaxis protein | 6 |
|  | WP_085348892.1 | 7.9E-05 | Qa.IV | -----ATIGQISSISTSIASAVEQQGAATQEIARSVQTVAQGTQTAATDI--- | *Bradyrhizobium canariense* | Bacteria; Proteobacteria; Alphaproteobacteria; Rhizobiales; Bradyrhizobiaceae; Bradyrhizobium | methyl-accepting chemotaxis protein | 6 |
|  | WP_085354318.1 | 7.9E-05 | Qa.IV | -----ATIGQISSISTSIASAVEQQGAATQEIARSVQTVAQGTQTAATDI--- | *Bradyrhizobium canariense* | Bacteria; Proteobacteria; Alphaproteobacteria; Rhizobiales; Bradyrhizobiaceae; Bradyrhizobium | methyl-accepting chemotaxis protein | 6 |
|  | WP_085354319.1 | 7.9E-05 | Qa.IV | -----ATIGQISSISTSIASAVEQQGAATQEIARSVQTVAQGTQTAATDI--- | *Bradyrhizobium canariense* | Bacteria; Proteobacteria; Alphaproteobacteria; Rhizobiales; Bradyrhizobiaceae; Bradyrhizobium | methyl-accepting chemotaxis protein | 6 |
|  | WP_085358521.1 | 7.9E-05 | Qa.IV | -----ATIGQISSISTSIASAVEQQGAATQEIARSVQTVAQGTQTAATDI--- | *Bradyrhizobium canariense* | Bacteria; Proteobacteria; Alphaproteobacteria; Rhizobiales; Bradyrhizobiaceae; Bradyrhizobium | methyl-accepting chemotaxis protein | 6 |
|  | WP_085403640.1 | 7.9E-05 | Qa.IV | -----ATIGQISSISTSIASAVEQQGAATQEIARSVQTVAQGTQTAATDI--- | *Bradyrhizobium japonicum* | Bacteria; Proteobacteria; Alphaproteobacteria; Rhizobiales; Bradyrhizobiaceae; Bradyrhizobium | methyl-accepting chemotaxis protein | 6 |
|  | WP_092030663.1 | 7.9E-05 | Qa.IV | -----ATIGQISSISTSIASAVEQQGAATQEIARSVQTVAQGTQTAATDI--- | *Bradyrhizobium sp. OK095* | Bacteria; Proteobacteria; Alphaproteobacteria; Rhizobiales; Bradyrhizobiaceae; Bradyrhizobium | methyl-accepting chemotaxis protein | 6 |
|  | WP_092293284.1 | 7.9E-05 | Qa.IV | -----ATIGQISSISTSIASAVEQQGAATQEIARSVQTVAQGTQTAATDI--- | *Bradyrhizobium sp. Ghvi* | Bacteria; Proteobacteria; Alphaproteobacteria; Rhizobiales; Bradyrhizobiaceae; Bradyrhizobium | methyl-accepting chemotaxis protein | 6 |
|  | WP_110747865.1 | 7.9E-05 | Qb.I | LQRTKMHLQETIEQSEVSSEELKASNEEMQAINEELRSATEELETS------- | *Pseudomonas sp. OV529* | Bacteria; Proteobacteria; Gammaproteobacteria; Pseudomonadales; Pseudomonadaceae; Pseudomonas | PAS domain S-box protein partial | 6 |
|  | GAJ31679.1 | 8E-05 | Qa.IV | -----ATIGQISSISTSIASAVEQQGAATQEIARSVQTVAQGTQTAATDI--- | *Bradyrhizobium sp. DOA9* | Bacteria; Proteobacteria; Alphaproteobacteria; Rhizobiales; Bradyrhizobiaceae; Bradyrhizobium | probable chemoreceptor Y4FA | 6 |
|  | PID76801.1 | 8E-05 | Qa.III.b | -------ISDVNDLVTTIAAAVEEQSATTREISGNVQQTSIKMDEVGSNVE-- | *Deltaproteobacteria bacterium* | Bacteria; Proteobacteria; Deltaproteobacteria | hypothetical protein CSB24_04725 partial | 6 |
|  | SIT13621.1 | 8E-05 | Qa.IV | IQEISDTISSVNEISTSIASAIQEQTAATREISRNVSQAAD------------ | *Thalassospira xiamenensis M-5 = DSM 17429* | Bacteria; Proteobacteria; Alphaproteobacteria; Rhodospirillales; Rhodospirillaceae; Thalassospira | methyl-accepting chemotaxis sensory transducer with Cache sensor | 6 |
|  | WP_024081945.1 | 8E-05 | Qc.III | ---ITGTIENINELSAAISAAVEEQGAATMEIARSVSQAAEGTH--------- | *Magnetospirillum gryphiswaldense* | Bacteria; Proteobacteria; Alphaproteobacteria; Rhodospirillales; Rhodospirillaceae; Magnetospirillum | PAS domain S-box protein | 6 |
| | WP_094410311.1 | 8E-05 | Qc.III | IDDIAKTILSLNEISTSIAAAVEEQSSATNDISRAVQEAARGTERLRENVQ-- | *Elstera cyanobacteriorum* | Bacteria; Proteobacteria; Alphaproteobacteria; Rhodospirillales; Rhodospirillaceae; Elstera | HAMP domain-containing protein | 6 |
|  | WP_095734104.1 | 8E-05 | Qc.III.c | --AISSVIVQISEISTVIASAVEEQAAATKEVTANINGTTKAIDDA------- | *Bradyrhizobium sp. UFLA03-84* | Bacteria; Proteobacteria; Alphaproteobacteria; Rhizobiales; Bradyrhizobiaceae; Bradyrhizobium | PAS domain-containing protein | 6 |
|  | WP_106002748.1 | 8E-05 | Qc.III | ---ITGTIENINELSAAISAAVEEQGAATMEIARSVSQAAEGTH--------- | *Magnetospirillum gryphiswaldense* | Bacteria; Proteobacteria; Alphaproteobacteria; Rhodospirillales; Rhodospirillaceae; Magnetospirillum | PAS domain S-box protein | 6 |
|  | WP_114357476.1 | 8E-05 | Qc.III | -SEISGTIEKLSEISAHIAAAVEQQGAATQEISRNIQQTAQGTERV------- | *Rhodopseudomonas pentothenatexigens* | Bacteria; Proteobacteria; Alphaproteobacteria; Rhizobiales; Bradyrhizobiaceae; Rhodopseudomonas | chemotaxis protein | 6 |
|  | BAR57410.1 | 8.1E-05 | Qa.IV | -----ATIGQISSISTSIASAVEQQGAATQEIARSVQTVAQGTQTAATDI--- | *Bradyrhizobium diazoefficiens* | Bacteria; Proteobacteria; Alphaproteobacteria; Rhizobiales; Bradyrhizobiaceae; Bradyrhizobium | putative methyl accepting chemotaxis protein | 6 |
|  | CUT10636.1 | 8.1E-05 | Qa.III.b | -EAIAQTITEINGITASIATSIEQQGLATREIARNVQEASS------------ | *Bradyrhizobium sp.* | Bacteria; Proteobacteria; Alphaproteobacteria; Rhizobiales; Bradyrhizobiaceae; Bradyrhizobium | Methylaccepting chemotaxis protein | 6 |
|  | KGJ64753.1 | 8.1E-05 | Qa.IV | -----ATIGQISSISTSIASAVEQQGAATQEIARSVQTVAQGTQTAATDI--- | *Bradyrhizobium diazoefficiens SEMIA 5080* | Bacteria; Proteobacteria; Alphaproteobacteria; Rhizobiales; Bradyrhizobiaceae; Bradyrhizobium | putative methyl-accepting chemotaxis protein | 6 |
| | OYX14508.1 | 8.1E-05 | Qc.III | IDRIVGTISQIRTASSAISAAIEEQGAATQEIASNTQKAARGTEVVNENIS-- | *Rhizobiales bacterium 32-66-8* | Bacteria; Proteobacteria; Alphaproteobacteria; Rhizobiales | hypothetical protein B7Z15_03725 | 6 |
|  | SFJ97897.1 | 8.1E-05 | Qa.IV | -----ATIGQISSISTSIASAVEQQGAATQEIARSVQTVAQGTQTAATDI--- | *Bradyrhizobium sp. cf659* | Bacteria; Proteobacteria; Alphaproteobacteria; Rhizobiales; Bradyrhizobiaceae; Bradyrhizobium | Methyl-accepting chemotaxis protein | 6 |
| | WP_023002861.1 | 8.1E-05 | Qc.III.c | -ESILAVIKNVGDIAASIASSVEEQSAVTNDISSNMHSAAQGVERITSNM--- | *Labrenzia* | Bacteria; Proteobacteria; Alphaproteobacteria; Rhodobacterales; Rhodobacteraceae | MULTISPECIES: methyl-accepting chemotaxis protein | 6 |
|  | WP_042955342.1 | 8.1E-05 | Qb.I | LQRTKMHLQETIEQSEVSSEELKASNEEMQAINEELRSATEELETS------- | *Pseudomonas sp. G5(2012)* | Bacteria; Proteobacteria; Gammaproteobacteria; Pseudomonadales; Pseudomonadaceae; Pseudomonas | PAS domain S-box protein partial | 6 |
|  | WP_060734554.1 | 8.1E-05 | Qa.IV | -----ATIGQISSISTSIASAVEQQGAATQEIARSVQTVAQGTQTAATDI--- | *Bradyrhizobium sp. CCGE-LA001* | Bacteria; Proteobacteria; Alphaproteobacteria; Rhizobiales; Bradyrhizobiaceae; Bradyrhizobium | methyl-accepting chemotaxis protein | 6 |
|  | WP_082847785.1 | 8.1E-05 | Qa.IV | -----ATIGQISSISTSIASAVEQQGAATQEIARSVQTVAQGTQTAATDI--- | *Bradyrhizobium sp. DOA9* | Bacteria; Proteobacteria; Alphaproteobacteria; Rhizobiales; Bradyrhizobiaceae; Bradyrhizobium | methyl-accepting chemotaxis protein | 6 |
| | WP_101264715.1 | 8.1E-05 | Qc.III | --DIVETIERVNSIASNIAAAIEEQQAATGEIARNVEQASRGTQNVNANIFD- | *Thalassospira marina* | Bacteria; Proteobacteria; Alphaproteobacteria; Rhodospirillales; Rhodospirillaceae; Thalassospira | methyl-accepting chemotaxis protein | 6 |
|  | WP_109469601.1 | 8.1E-05 | Qc.III | -RDIADTVRRVNDIAGSIAQAVERQSAATGAISRNVQQAAE------------ | *Azospirillum sp. TSH58* | Bacteria; Proteobacteria; Alphaproteobacteria; Rhodospirillales; Rhodospirillaceae; Azospirillum | HAMP domain-containing protein | 6 |
|  | OUR79887.1 | 8.2E-05 | Qa.III | -EEIAKTIEQIDEATSSIANAVETQASVIGNISSNAQSAAQ------------ | *Alphaproteobacteria bacterium 46_93_T64* | Bacteria; Proteobacteria; Alphaproteobacteria | hypothetical protein A9Q83_02765 | 6 |
|  | SCB55112.1 | 8.2E-05 | Qa.IV | -----ATIGQISSISTSIASAVEQQGAATQEIARSVQTVAQGTQTAATDI--- | *Bradyrhizobium shewense* | Bacteria; Proteobacteria; Alphaproteobacteria; Rhizobiales; Bradyrhizobiaceae; Bradyrhizobium | Methyl-accepting chemotaxis protein | 6 |
|  | WP_010683794.1 | 8.2E-05 | Qa.III | IDGIGGTIEQLSEISLSVSSAVTEQAAVTRDMTENMQSAAQSVEAAR------ | *Methylobacterium mesophilicum* | Bacteria; Proteobacteria; Alphaproteobacteria; Rhizobiales; Methylobacteriaceae; Methylobacterium | methyl-accepting chemotaxis protein | 6 |
|  | WP_022666794.1 | 8.2E-05 | Qa.III | IETIVSVINEVDEIVTSVAVAIEEQSVSTKEISNNVTQAAVGVQEVN------ | *Desulfospira joergensenii* | Bacteria; Proteobacteria; Deltaproteobacteria; Desulfobacterales; Desulfobacteraceae; Desulfospira | HAMP domain-containing protein | 6 |

| | | | | | | | | |
|---|---|---|---|---|---|---|---|---|
|  | **WP_035694661.1** | 8.2E-05 | Qa.IV | -----ATIGQISSISTSIASAVEQQGAATQEIARSVQTVAQGTQTAATDI--- | *Bradyrhizobium liaoningense* | Bacteria; Proteobacteria; Alphaproteobacteria; Rhizobiales; Bradyrhizobiaceae; Bradyrhizobium | methyl-accepting chemotaxis protein | 6 |
|  | **WP_038974743.1** | 8.2E-05 | Qa.IV | -----ATIGQISSISTSIASAVEQQGAATQEIARSVQTVAQGTQTAATDI--- | *Bradyrhizobium* | Bacteria; Proteobacteria; Alphaproteobacteria; Rhizobiales; Bradyrhizobiaceae | MULTISPECIES: methyl-accepting chemotaxis protein | 6 |
|  | **WP_060734553.1** | 8.2E-05 | Qa.IV | -----ATIGQISSISTSIASAVEQQGAATQEIARSVQTVAQGTQTAATDI--- | *Bradyrhizobium sp. CCGE-LA001* | Bacteria; Proteobacteria; Alphaproteobacteria; Rhizobiales; Bradyrhizobiaceae; Bradyrhizobium | methyl-accepting chemotaxis protein | 6 |
| | **WP_068493451.1** | 8.2E-05 | Qc.III | --SVSSTIGRIDEIASAIASAVEEQSAATQEIARNVEEAARGTQEVSSNI--- | *Magnetospirillum marisnigri* | Bacteria; Proteobacteria; Alphaproteobacteria; Rhodospirillales; Rhodospirillaceae; Magnetospirillum | HAMP domain-containing protein | 6 |
|  | **WP_074448075.1** | 8.2E-05 | Qa.IV | -----ATIGQISSISTSIASAVEQQGAATQEIARSVQTVAQGTQTAATDI--- | *Bradyrhizobium yuanmingense* | Bacteria; Proteobacteria; Alphaproteobacteria; Rhizobiales; Bradyrhizobiaceae; Bradyrhizobium | methyl-accepting chemotaxis protein | 6 |
|  | **WP_084292884.1** | 8.2E-05 | Qa.IV | -----ATIGQISSISTSIASAVEQQGAATQEIARSVQTVAQGTQTAATDI--- | *Bradyrhizobium sp. WSM3983* | Bacteria; Proteobacteria; Alphaproteobacteria; Rhizobiales; Bradyrhizobiaceae; Bradyrhizobium | methyl-accepting chemotaxis protein | 6 |
|  | **WP_091966444.1** | 8.2E-05 | Qa.IV | -----ATIGQISSISTSIASAVEQQGAATQEIARSVQTVAQGTQTAATDI--- | *Bradyrhizobium shewense* | Bacteria; Proteobacteria; Alphaproteobacteria; Rhizobiales; Bradyrhizobiaceae; Bradyrhizobium | methyl-accepting chemotaxis protein | 6 |
|  | **WP_100236546.1** | 8.2E-05 | Qa.IV | -----ATIGQISSISTSIASAVEQQGAATQEIARSVQTVAQGTQTAATDI--- | *Bradyrhizobium sp. INPA54B* | Bacteria; Proteobacteria; Alphaproteobacteria; Rhizobiales; Bradyrhizobiaceae; Bradyrhizobium | methyl-accepting chemotaxis protein | 6 |
|  | **WP_104464289.1** | 8.2E-05 | Qa.IV | -----ATIGQISSISTSIASAVEQQGAATQEIARSVQTVAQGTQTAATDI--- | *Bradyrhizobium sp. AC87j1* | Bacteria; Proteobacteria; Alphaproteobacteria; Rhizobiales; Bradyrhizobiaceae; Bradyrhizobium | methyl-accepting chemotaxis protein | 6 |
|  | **WP_105554824.1** | 8.2E-05 | R.IV | IDQISKAVSELDSTTQQNAALVQESSAAAASLEQQAGELSQLISRF------- | *Cronobacter sakazakii* | Bacteria; Proteobacteria; Gammaproteobacteria; Enterobacterales; Enterobacteriaceae; Cronobacter | chemotaxis protein | 6 |
|  | **WP_106942156.1** | 8.2E-05 | Qa.IV | -----ATIGQISSISTSIASAVEQQGAATQEIARSVQTVAQGTQTAATDI--- | *Bradyrhizobium sp. MOS002* | Bacteria; Proteobacteria; Alphaproteobacteria; Rhizobiales; Bradyrhizobiaceae; Bradyrhizobium | methyl-accepting chemotaxis protein | 6 |
|  | **WP_108914658.1** | 8.2E-05 | Qa.IV | -----ATIGQISSISTSIASAVEQQGAATQEIARSVQTVAQGTQTAATDI--- | *Bradyrhizobium arachidis* | Bacteria; Proteobacteria; Alphaproteobacteria; Rhizobiales; Bradyrhizobiaceae; Bradyrhizobium | methyl-accepting chemotaxis protein | 6 |
| | **OJW24429.1** | 8.3E-05 | Qc.III | -RSITATIQRMSEIAAGISGAVDRQSASTQEIAGAVQRAAAGTNEMNGNI--- | *Rhodospirillales bacterium 69-11* | Bacteria; Proteobacteria; Alphaproteobacteria; Rhodospirillales; unclassified Rhodospirillales | hypothetical protein BGO51_02840 | 6 |
|  | **OYV39626.1** | 8.3E-05 | Qc.III | -GEIVATIDRISGIASAIAAAVEEQGAATNEIASSVSHAAQGTRQV------- | *Rhodospirillales bacterium 20-64-7* | Bacteria; Proteobacteria; Alphaproteobacteria; Rhodospirillales; unclassified Rhodospirillales | hypothetical protein B7Z80_06740 | 6 |
|  | **WP_008076413.1** | 8.3E-05 | Qa.III | ---ISESVSSIHDMSTQIATAAEQQHLVAEEINRNIQQVHEDTQK-------- | *Vibrio sinaloensis* | Bacteria; Proteobacteria; Gammaproteobacteria; Vibrionales; Vibrionaceae; Vibrio; Vibrio oreintalis group | methyl-accepting chemotaxis protein | 6 |
|  | **WP_011443427.1** | 8.3E-05 | Qa.IV | IRNISQTITSINEVTTAITAAVEEQGAATREIARNIQQAA------------- | *Rhodopseudomonas palustris* | Bacteria; Proteobacteria; Alphaproteobacteria; Rhizobiales; Bradyrhizobiaceae; Rhodopseudomonas | HAMP domain-containing protein | 6 |
|  | **WP_011501263.1** | 8.3E-05 | Qa.IV | IRNISQTITSINEVTTAITAAVEEQGAATREIARNIQQAA------------- | *Rhodopseudomonas palustris* | Bacteria; Proteobacteria; Alphaproteobacteria; Rhizobiales; Bradyrhizobiaceae; Rhodopseudomonas | HAMP domain-containing protein | 6 |
|  | **WP_092684409.1** | 8.3E-05 | Qa.IV | IRNISQTITSINEVTTAITAAVEEQGAATREIARNIQQAA------------- | *Rhodopseudomonas pseudopalustris* | Bacteria; Proteobacteria; Alphaproteobacteria; Rhizobiales; Bradyrhizobiaceae; Rhodopseudomonas | HAMP domain-containing protein | 6 |
|  | **WP_109075901.1** | 8.3E-05 | Qa.III | IKGVAQVIGRVNDVATSIASAVEQQGAATREISRNIQQAAQ------------ | *Azospirillum sp. TSH20* | Bacteria; Proteobacteria; Alphaproteobacteria; Rhodospirillales; Rhodospirillaceae; Azospirillum | methyl-accepting chemotaxis protein | 6 |
|  | **WP_110780295.1** | 8.3E-05 | Qa.III | IKEIGDTIGQLSQISSTIASAVEQQGAATREISQNVHQAA------------- | *Rhodopseudomonas faecalis* | Bacteria; Proteobacteria; Alphaproteobacteria; Rhizobiales; Bradyrhizobiaceae; Rhodopseudomonas | PAS domain S-box protein | 6 |
|  | **ALV65607.1** | 8.4E-05 | Qa.IV | --EIEANVNILSQGINEMSQSINEQTEAINQINEAVATVDEQTKQ-NV----- | *Campylobacter fetus subsp. testudinum Sp3* | Bacteria; Proteobacteria; Epsilonproteobacteria; Campylobacterales; Campylobacteraceae; Campylobacter | MCP-domain signal transduction protein | 6 |
|  | **ANJ55539.1** | 8.4E-05 | Qb.I | LQSTRLQLQDTIEQSEISSEELKTSNEEMQSINEELRSATEELETS------- | *Pseudomonas silesiensis* | Bacteria; Proteobacteria; Gammaproteobacteria; Pseudomonadales; Pseudomonadaceae; Pseudomonas | chemotaxis protein | 6 |
| | **AUC96401.1** | 8.4E-05 | Qa.III.b | --DIAGIIQRISQIASHVAGAIEEQGRATRSIAENVQVASERTSQVAVSIGQ- | *Bradyrhizobium sp. SK17* | Bacteria; Proteobacteria; Alphaproteobacteria; Rhizobiales; Bradyrhizobiaceae; Bradyrhizobium | methyl-accepting chemotaxis protein | 6 |
|  | **ETX59254.1** | 8.4E-05 | Qc.III.c | IDQIEKGADNIAEMSIQIASACSEQSSVTEELHRNVEHIN------------- | *Vibrio parahaemolyticus SBR10290* | Bacteria; Proteobacteria; Gammaproteobacteria; Vibrionales; Vibrionaceae; Vibrio | methyl-accepting chemotaxis (MCP) signaling domain protein | 6 |
| | **SNT53146.1** | 8.4E-05 | Qc.III | IKEISATIGRISEIASTIAAAVEQQGAATCEISRNVQQAAAGTTKVSHSIF-- | *Tardiphaga sp. OK246* | Bacteria; Proteobacteria; Alphaproteobacteria; Rhizobiales; Bradyrhizobiaceae; Tardiphaga | Methyl-accepting chemotaxis protein (MCP) signalling domain-containing protein | 6 |
|  | **WP_011474980.1** | 8.4E-05 | Qa.III | IKEIGETIGQMAEIASTIASAVEQQGAATQEISRNVQQAARGTEQV------- | *Rhodopseudomonas palustris* | Bacteria; Proteobacteria; Alphaproteobacteria; Rhizobiales; Bradyrhizobiaceae; Rhodopseudomonas | methyl-accepting chemotaxis protein | 6 |
|  | **AWL98040.1** | 8.5E-05 | Qa.IV | -----ATIGQISSISTSIASAVEQQGAATQEIARSVQTVAQGTQTAATDI--- | *Bradyrhizobium ottawaense* | Bacteria; Proteobacteria; Alphaproteobacteria; Rhizobiales; Bradyrhizobiaceae; Bradyrhizobium | methyl-accepting chemotaxis protein | 6 |
| | **BAI74079.1** | 8.5E-05 | Qc.III | IEGIARTLGQVGEISTAIAGAVSQQNAATHEIASSVQNAASGSNEVTKAI--- | *Azospirillum sp. B510* | Bacteria; Proteobacteria; Alphaproteobacteria; Rhodospirillales; Rhodospirillaceae; Azospirillum | methyl-accepting chemotaxis protein (plasmid) | 6 |
|  | **WP_011083158.1** | 8.5E-05 | Qa.IV | -----ATIGQISSISTSIASAVEQQGAATQEIARSVQTVAQGTQTAATDI--- | *Bradyrhizobium diazoefficiens* | Bacteria; Proteobacteria; Alphaproteobacteria; Rhizobiales; Bradyrhizobiaceae; Bradyrhizobium | methyl-accepting chemotaxis protein | 6 |
| | **WP_020593757.1** | 8.5E-05 | Qc.III | --SISTTIDSIHEVATAIASAVEEQGAATAEITKSVQQAANGTDQVTSNI--- | *Kiloniella laminariae* | Bacteria; Proteobacteria; Alphaproteobacteria; Kiloniellales; Kiloniellaceae; Kiloniella | methyl-accepting chemotaxis protein | 6 |
|  | **WP_029059408.1** | 8.5E-05 | Qc.III | --GVSSTIQNMNDISTAIAAAVEEQGAATGEISRNISQASDGTGRVT------ | *Stappia stellulata* | Bacteria; Proteobacteria; Alphaproteobacteria; Rhodobacterales; Rhodobacteraceae; Stappia | hypothetical protein | 6 |
|  | **WP_082755779.1** | 8.5E-05 | Qa.IV | -----ATIGQISSISTSIASAVEQQGAATQEIARSVQTVAQGTQTAATDI--- | *Bradyrhizobium diazoefficiens* | Bacteria; Proteobacteria; Alphaproteobacteria; Rhizobiales; Bradyrhizobiaceae; Bradyrhizobium | methyl-accepting chemotaxis protein | 6 |
|  | **WP_083513925.1** | 8.5E-05 | Qa.IV | -----SNISRISQIASTIAAAIEEQGTATRHIAENIQHAAEMSSQVATNI--- | *Bradyrhizobium manausense* | Bacteria; Proteobacteria; Alphaproteobacteria; Rhizobiales; Bradyrhizobiaceae; Bradyrhizobium | methyl-accepting chemotaxis protein | 6 |
|  | **WP_085967937.1** | 8.5E-05 | Qa.IV | -----ATIGQISSISTSIASAVEQQGAATQEIARSVQTVAQGTQTAATDI--- | *Bradyrhizobium sp. CCBAU 15615* | Bacteria; Proteobacteria; Alphaproteobacteria; Rhizobiales; Bradyrhizobiaceae; Bradyrhizobium | methyl-accepting chemotaxis protein | 6 |
|  | **WP_092192113.1** | 8.5E-05 | Qa.IV | -----ATIGQISSISTSIASAVEQQGAATQEIARSVQTVAQGTQTAATDI--- | *Bradyrhizobium sp. cf659* | Bacteria; Proteobacteria; Alphaproteobacteria; Rhizobiales; Bradyrhizobiaceae; Bradyrhizobium | methyl-accepting chemotaxis protein | 6 |
|  | **WP_114357405.1** | 8.5E-05 | Qa.III | IREIGSTIAKMSEIASTIASAVEEQGAATQEISRNVQQAAQGTQQVS------ | *Rhodopseudomonas pentothenatexigens* | Bacteria; Proteobacteria; Alphaproteobacteria; Rhizobiales; Bradyrhizobiaceae; Rhodopseudomonas | HAMP domain-containing protein | 6 |
| | **KRQ14503.1** | 8.6E-05 | Qc.III | IKQIGDTIAQISGIATTIAAAVEQQGAATTEIARNVQRASDGTREVSGSITK- | *Bradyrhizobium manausense* | Bacteria; Proteobacteria; Alphaproteobacteria; Rhizobiales; Bradyrhizobiaceae; Bradyrhizobium | chemotaxis protein | 6 |
|  | **PCJ68318.1** | 8.6E-05 | Qc.III | --EVTDMIGKLNEISGAIAAAVEEQGAATTEIARSMGEASKGV---------- | *Rhodobiaceae bacterium* | Bacteria; Proteobacteria; Alphaproteobacteria; Rhizobiales; Rhodobiaceae; unclassified Rhodobiaceae | chemotaxis protein | 6 |
|  | **PLX94012.1** | 8.6E-05 | SNAP.b | -------FNEMASSLSASRDELTYNNEALQKANDDLKTMQEHLIRSERLA--- | *Desulfuromonas sp.* | Bacteria; Proteobacteria; Deltaproteobacteria; Desulfuromonadales; Desulfuromonadaceae; Desulfuromonas | hypothetical protein C0621_06135 | 6 |
|  | **WP_023386083.1** | 8.6E-05 | Qa.IV | --EIEANVNILSQGINEMSQSINEQTEAINQINEAVATVDEQTKQ-NV----- | *Campylobacter fetus* | Bacteria; Proteobacteria; Epsilonproteobacteria; Campylobacterales; Campylobacteraceae; Campylobacter | MCP-domain signal transduction protein | 6 |
| | **WP_079538339.1** | 8.6E-05 | Qc.III | IKEIGGTIGRMSEIASAIASAIEEQGAATQEISRNVQQAAHGTQQVSSNI--- | *Bradyrhizobium lablabi* | Bacteria; Proteobacteria; Alphaproteobacteria; Rhizobiales; Bradyrhizobiaceae; Bradyrhizobium | methyl-accepting chemotaxis protein | 6 |
|  | **WP_104464299.1** | 8.6E-05 | Qa.IV | -----ATIGQISSISTSIASAVEQQGAATQEIARSVQTVAQGTQTAATDI--- | *Bradyrhizobium sp. AC87j1* | Bacteria; Proteobacteria; Alphaproteobacteria; Rhizobiales; Bradyrhizobiaceae; Bradyrhizobium | methyl-accepting chemotaxis protein | 6 |
|  | **WP_105459573.1** | 8.6E-05 | Qa.IV | --EIEANVNILSQGINEMSQSINEQTEAINQINEAVATVDEQTKQ-NV----- | *Campylobacter fetus* | Bacteria; Proteobacteria; Epsilonproteobacteria; Campylobacterales; Campylobacteraceae; Campylobacter | chemotaxis protein | 6 |
| | **EGH61535.1** | 8.7E-05 | Qa.III | --GISEAVANITDMTTQIATATEEQSAVAEEINRNISTIAELADKTADEAR-- | *Pseudomonas syringae pv. maculicola str. ES4326* | Bacteria; Proteobacteria; Gammaproteobacteria; Pseudomonadales; Pseudomonadaceae; Pseudomonas | PAS protein partial | 6 |
|  | **WP_006615070.1** | 8.7E-05 | Qa.III | IREIGTTIGQMNEITETIAAAVEEQGVATKEIARSVQQAA------------- | *Bradyrhizobium sp. ORS 285* | Bacteria; Proteobacteria; Alphaproteobacteria; Rhizobiales; Bradyrhizobiaceae; Bradyrhizobium | HAMP domain-containing protein | 6 |
|  | **WP_041805531.1** | 8.7E-05 | Qc.III | -REIGGTIERMSEIASTIASAVEEQGAATQEISRNVQQAAQGTQQVSSN---- | *Rhodopseudomonas palustris* | Bacteria; Proteobacteria; Alphaproteobacteria; Rhizobiales; Bradyrhizobiaceae; Rhodopseudomonas | methyl-accepting chemotaxis protein | 6 |
|  | **WP_092681719.1** | 8.7E-05 | Qc.III | -REIGGTIERMSEIASTIASAVEEQGAATQEISRNVQQAAQGTQQVSSN---- | *Rhodopseudomonas pseudopalustris* | Bacteria; Proteobacteria; Alphaproteobacteria; Rhizobiales; Bradyrhizobiaceae; Rhodopseudomonas | methyl-accepting chemotaxis protein | 6 |
|  | **WP_105770241.1** | 8.7E-05 | Qa.III | --EIVAAVERVNAIMNDISTAFDEQSSGIEQVNATVTQMEQTMQ--------- | *Burkholderia multivorans* | Bacteria; Proteobacteria; Betaproteobacteria; Burkholderiales; Burkholderiaceae; Burkholderia; Burkholderia cepacia complex | HAMP domain-containing protein | 6 |
|  | **EPN41424.1** | 8.8E-05 | Qa.III | ---ISQALRNLNSLNASIASATLQQTHVVDDINQNVTQAAQLSQ--------- | *Pseudomonas syringae pv. actinidiae ICMP 18807* | Bacteria; Proteobacteria; Gammaproteobacteria; Pseudomonadales; Pseudomonadaceae; Pseudomonas; Pseudomonas syringae | methyl-accepting chemotaxis protein partial | 6 |
| | **PIE74531.1** | 8.8E-05 | Qa.IV | IKSVNKVIKDINTIIFTIATAIEQQNSATQEIASNISQVSQGTQDVNENIAQ- | *Deltaproteobacteria bacterium* | Bacteria; Proteobacteria; Deltaproteobacteria | methyl-accepting chemotaxis protein | 6 |
|  | **SEO13872.1** | 8.8E-05 | Qc.III | -REIGGTIERMSEIASTIASAVEEQGAATQEISRNVQQAAQGTQQVSSN---- | *Rhodopseudomonas pseudopalustris* | Bacteria; Proteobacteria; Alphaproteobacteria; Rhizobiales; Bradyrhizobiaceae; Rhodopseudomonas | Methyl-accepting chemotaxis protein | 6 |
| | **WP_012497776.1** | 8.8E-05 | Qc.III | -REIGQTIGRMSEISSAIAAAVEEQGAATQEISRNVQQAAHGTQQVSSNI--- | *Rhodopseudomonas palustris* | Bacteria; Proteobacteria; Alphaproteobacteria; Rhizobiales; Bradyrhizobiaceae; Rhodopseudomonas | methyl-accepting chemotaxis protein | 6 |
|  | **WP_042695312.1** | 8.8E-05 | Qa.III | ITEIARVIGEVNEVSSTIANAIGEQRAATDEISGSIQRAA------------- | *Azospirillum sp. B506* | Bacteria; Proteobacteria; Alphaproteobacteria; Rhodospirillales; Rhodospirillaceae; Azospirillum | methyl-accepting chemotaxis protein | 6 |
|  | **WP_045583775.1** | 8.8E-05 | Qc.III | --GIGRTIGRMSEIATAIAAAIEEQAAATREIANSVTQAAQG----------- | *Azospirillum thiophilum* | Bacteria; Proteobacteria; Alphaproteobacteria; Rhodospirillales; Rhodospirillaceae; Azospirillum | methyl-accepting chemotaxis protein | 6 |
|  | **WP_063993129.1** | 8.8E-05 | Qa.III.b | -EAIAQTITEINGITASIATSIEQQGLATREIARNVQEASS------------ | *Bradyrhizobium sp.* | Bacteria; Proteobacteria; Alphaproteobacteria; Rhizobiales; Bradyrhizobiaceae; Bradyrhizobium | chemotaxis protein | 6 |
|  | **WP_072189619.1** | 8.8E-05 | R.IV | IDQISKAVSELDSTTQQNAALVQESSAAAASLEQQAGELSQLISRF------- | *Cronobacter sakazakii* | Bacteria; Proteobacteria; Gammaproteobacteria; Enterobacterales; Enterobacteriaceae; Cronobacter | chemotaxis protein | 6 |
| | **WP_085552088.1** | 8.8E-05 | Qc.III | --GIGRTIGTIREVAASIAAAVEEQGAATGEIARNVQEAASGTQAVTSNI--- | *Azospirillum lipoferum* | Bacteria; Proteobacteria; Alphaproteobacteria; Rhodospirillales; Rhodospirillaceae; Azospirillum | HAMP domain-containing protein | 6 |
|  | **WP_094305922.1** | 8.8E-05 | Qa.III | IVGIGHTIQDINQIAGGIAAGIGQQGSAIRAIVSNMDQAA------------- | *Azospirillum brasilense* | Bacteria; Proteobacteria; Alphaproteobacteria; Rhodospirillales; Rhodospirillaceae; Azospirillum | chemotaxis protein | 6 |
|  | **WP_101723648.1** | 8.8E-05 | Qc.I | -----DMAAELRGMYEHLESQVQERTVDLREANALLERQRDKLEQLNA----- | *Eggerthella timonensis* | Bacteria; Actinobacteria; Coriobacteriia; Eggerthellales; Eggerthellaceae; Eggerthella | DUF3365 domain-containing protein | 6 |
| | **WP_104513051.1** | 8.8E-05 | Qc.III | -REIGQTIGRMSEISSAIAAAVEEQGAATQEISRNVQQAAHGTQQVSSNI--- | *Rhodopseudomonas palustris* | Bacteria; Proteobacteria; Alphaproteobacteria; Rhizobiales; Bradyrhizobiaceae; Rhodopseudomonas | methyl-accepting chemotaxis protein | 6 |

| | | | | | | | | |
|---|---|---|---|---|---|---|---|---|
| | **WP_107357634.1** | 8.8E-05 | Qc.III | -REIGQTIGRMSEISSAIAAAVEEQGAATQEISRNVQQAAHGTQQVSSNI--- | *Rhodopseudomonas palustris* | Bacteria; Proteobacteria; Alphaproteobacteria; Rhizobiales; Bradyrhizobiaceae; Rhodopseudomonas | methyl-accepting chemotaxis protein | 6 |
|  | **OHB42053.1** | 8.9E-05 | SNAP.b | -------LKASYDEKEKIMAQLRENVNNLSNANKELVKINEQLNEAQERL--- | *Planctomycetes bacterium GWB2_41_19* | Bacteria; Planctomycetes | hypothetical protein A2069_01995 | 6 |
|  | **OHB46627.1** | 8.9E-05 | SNAP.b | -------LKASYDEKEKIMAQLRENVNNLSNANKELVKINEQLNEAQERL--- | *Planctomycetes bacterium GWE2_41_14* | Bacteria; Planctomycetes | hypothetical protein A2094_00165 | 6 |
|  | **OHC06440.1** | 8.9E-05 | SNAP.b | -------LKASYDEKEKIMAQLRENVNNLSNANKELVKINEQLNEAQERL--- | *Planctomycetes bacterium RIFOXYC2_FULL_41_27* | Bacteria; Planctomycetes | hypothetical protein A3J92_06140 | 6 |
|  | **OHC08523.1** | 8.9E-05 | SNAP.b | -------LKASYDEKEKIMAQLRENVNNLSNANKELVKINEQLNEAQERL--- | *Planctomycetes bacterium RIFOXYD2_FULL_41_16* | Bacteria; Planctomycetes | hypothetical protein A2545_03065 | 6 |
| | **SIS90811.1** | 8.9E-05 | Qa.III | IRAIAGTIEDISHIAAGIAAAVEEQGAATQEIARNVQEAAHGTDEVNRNI--- | *Insolitispirillum peregrinum* | Bacteria; Proteobacteria; Alphaproteobacteria; Rhodospirillales; Rhodospirillaceae; Insolitispirillum | methyl-accepting chemotaxis sensory transducer with Cache sensor | 6 |
|  | **WP_008972179.1** | 8.9E-05 | Qa.IV | ---ITSTIEEVSAITTTIGSAIEEQGAATAEIACNVTQTAEAT---------- | *Bradyrhizobium sp. STM 3843* | Bacteria; Proteobacteria; Alphaproteobacteria; Rhizobiales; Bradyrhizobiaceae; Bradyrhizobium | HAMP domain-containing protein | 6 |
|  | **WP_045143946.1** | 8.9E-05 | Qc.III | -----GGAEQVAETSQTISEGATEQASAIEELTASLGEVNEKVQNSVKHAK-- | *Clostridium butyricum* | Bacteria; Firmicutes; Clostridia; Clostridiales; Clostridiaceae; Clostridium | methyl-accepting chemotaxis protein partial | 6 |
|  | **WP_082938271.1** | 8.9E-05 | Qc.I | ----------LRGHASTLEDQVTQRTAQLTESNAELTQALANLKTAQREL--- | *Mitsuaria sp. 7* | Bacteria; Proteobacteria; Betaproteobacteria; Burkholderiales; Mitsuaria | HAMP domain-containing protein | 6 |
| | **WP_084194843.1** | 8.9E-05 | Qa.III | IRAIAGTIEDISHIAAGIAAAVEEQGAATQEIARNVQEAAHGTDEVNRNI--- | *Insolitispirillum peregrinum* | Bacteria; Proteobacteria; Alphaproteobacteria; Rhodospirillales; Rhodospirillaceae; Insolitispirillum | HAMP domain-containing protein | 6 |
| | **WP_089266123.1** | 8.9E-05 | Qc.III | IKEISATIGRISEIASTIAAAVEQQGAATCEISRNVQQAAAGTTKVSHSIF-- | *Tardiphaga sp. OK246* | Bacteria; Proteobacteria; Alphaproteobacteria; Rhizobiales; Bradyrhizobiaceae; Tardiphaga | hypothetical protein | 6 |
|  | **WP_104402452.1** | 8.9E-05 | Qa.III | LDQITNMLTELNDMNLQIATAANEQSSVTDEISSSITSIADASMQ-------- | *Vibrio penaeicida* | Bacteria; Proteobacteria; Gammaproteobacteria; Vibrionales; Vibrionaceae; Vibrio | methyl-accepting chemotaxis protein | 6 |
|  | **AND93414.1** | 9.1E-05 | Qa.IV | -----ATIGQISSISTSIASAVEQQGAATQEIARSVQTVAQGTHAAATDI--- | *Bradyrhizobium diazoefficiens USDA 110* | Bacteria; Proteobacteria; Alphaproteobacteria; Rhizobiales; Bradyrhizobiaceae; Bradyrhizobium | chemotaxis protein | 6 |
|  | **KZD23464.1** | 9.1E-05 | Qa.IV | IRNIGQTIAEINHVSAAIATAVEQQGAATREIARNIQHAA------------- | *Tardiphaga robiniae* | Bacteria; Proteobacteria; Alphaproteobacteria; Rhizobiales; Bradyrhizobiaceae; Tardiphaga | chemotaxis protein | 6 |
|  | **OUR61943.1** | 9.1E-05 | Qc.I | -SQMGQMAKQMHSNYQEFTDALKQfRKQALDDFAGRLDQAKQESQS-------- | *Bermanella sp. 47_1433_sub80_T6* | Bacteria; Proteobacteria; Gammaproteobacteria; Oceanospirillales; Bermanella | hypothetical protein A9Q73_10635 | 6 |
|  | **PVX53297.1** | 9.1E-05 | Qa.IV | IRNIGQTIAEINHVSAAIATAVEQQGAATREIARNIQHAA------------- | *Tardiphaga sp. OV697* | Bacteria; Proteobacteria; Alphaproteobacteria; Rhizobiales; Bradyrhizobiaceae; Tardiphaga | methyl-accepting chemotaxis protein | 6 |
| | **SMX59627.1** | 9.1E-05 | Qc.III | --AISGTIEKLSEISSAIAAAVEEQGAATQEIARNVQQAAAGTEQVSSNITD- | *Bradyrhizobium sp. ORS 285* | Bacteria; Proteobacteria; Alphaproteobacteria; Rhizobiales; Bradyrhizobiaceae; Bradyrhizobium | Methyl-accepting chemotaxis sensory transducer | 6 |
|  | **WP_003241834.1** | 9.1E-05 | Qb.I | --DAVQRIAAALGQFEQTMQAFKNATADIARTRQEMTDQQGEI---------- | *Pseudomonas mendocina* | Bacteria; Proteobacteria; Gammaproteobacteria; Pseudomonadales; Pseudomonadaceae; Pseudomonas | methyl-accepting chemotaxis protein | 6 |
|  | **WP_023471609.1** | 9.1E-05 | Qc.III | IDSISGIINKINDISAVIASAVEEQAATANEIGRTVSDAAVGSDE-------- | *Betaproteobacteria bacterium MOLA814* | Bacteria; Proteobacteria; Betaproteobacteria | methyl-accepting chemotaxis protein | 6 |
| | **WP_035647850.1** | 9.1E-05 | Qc.III | --AISGTIEKLSEISSAIAAAVEEQGAATQEIARNVQQAAAGTEQVSSNITD- | *Bradyrhizobium sp. ORS 285* | Bacteria; Proteobacteria; Alphaproteobacteria; Rhizobiales; Bradyrhizobiaceae; Bradyrhizobium | chemotaxis protein | 6 |
|  | **WP_045012000.1** | 9.1E-05 | Qa.IV | -----ATIGQISSISTSIASAVEEQGAATQEIARSVQNVAQGTQTAATDI--- | *Bradyrhizobium sp. LTSP849* | Bacteria; Proteobacteria; Alphaproteobacteria; Rhizobiales; Bradyrhizobiaceae; Bradyrhizobium | methyl-accepting chemotaxis protein | 6 |
|  | **WP_089266849.1** | 9.1E-05 | Qa.IV | IRNIGQTIAEINHVSAAIATAVEQQGAATREIARNIQHAA------------- | *Tardiphaga sp. OK246* | Bacteria; Proteobacteria; Alphaproteobacteria; Rhizobiales; Bradyrhizobiaceae; Tardiphaga | HAMP domain-containing protein | 6 |
|  | **WP_089976704.1** | 9.1E-05 | Qb.I | LRRTKEQLQATIEESETSSEELKASNEELQSINEELRSATEELETS------- | *Luteibacter sp. UNCMF331Sha3.1* | Bacteria; Proteobacteria; Gammaproteobacteria; Xanthomonadales; Rhodanobacteraceae; Luteibacter | PAS domain S-box protein | 6 |
|  | **WP_091896535.1** | 9.1E-05 | Qa.IV | -----ATIGQISSISTSIASAVEQQGAATQEIARSVQTVAQGTQTAATDI--- | *Bradyrhizobium sp. Rc2d* | Bacteria; Proteobacteria; Alphaproteobacteria; Rhizobiales; Bradyrhizobiaceae; Bradyrhizobium | methyl-accepting chemotaxis protein | 6 |
|  | **WP_092140117.1** | 9.1E-05 | Qa.IV | IRNIGQTIAEINHVSAAIATAVEQQGAATREIARNIQHAA------------- | *Bradyrhizobiaceae* | Bacteria; Proteobacteria; Alphaproteobacteria; Rhizobiales | HAMP domain-containing protein | 6 |
|  | **WP_092261363.1** | 9.1E-05 | Qa.IV | -----ATIGQISSISTSIASAVEQQGAATQEIARSVQTVAQGTQTAATDI--- | *Bradyrhizobium sp. Rc3b* | Bacteria; Proteobacteria; Alphaproteobacteria; Rhizobiales; Bradyrhizobiaceae; Bradyrhizobium | methyl-accepting chemotaxis protein | 6 |
|  | **WP_093761638.1** | 9.1E-05 | Qa.IV | IRNIGQTIAEINHVSAAIATAVEQQGAATREIARNIQHAA------------- | *Tardiphaga sp. OK245* | Bacteria; Proteobacteria; Alphaproteobacteria; Rhizobiales; Bradyrhizobiaceae; Tardiphaga | HAMP domain-containing protein | 6 |
|  | **WP_110798670.1** | 9.1E-05 | R.IV | IDQISKAVSELDSTTQQNAALVQESSAAAASLEQQAGELSQLISRF------- | *Cronobacter sakazakii* | Bacteria; Proteobacteria; Gammaproteobacteria; Enterobacterales; Enterobacteriaceae; Cronobacter | chemotaxis protein | 6 |
| | **WP_114394222.1** | 9.1E-05 | Qc.III | --TIGQTIGSISEVSAAIAAAIEEQNATTSEITRSVQQAAEGTQMVSRNVS-- | *Rhodospirillaceae bacterium NAU-10* | Bacteria; Proteobacteria; Alphaproteobacteria; Rhodospirillales; Rhodospirillaceae | HAMP domain-containing protein | 6 |
|  | **CCY77420.1** | 9.2E-05 | Qc.III | -NGINDSVNIITNGSVNLSDRANSQASSLEELASSVEFVLSSLRETAK----- | *Brachyspira sp. CAG:700* | Bacteria; Spirochaetes; Brachyspirales; Brachyspiraceae; Brachyspira; environmental samples | methyl-accepting chemotaxis sensory transducer with Cache sensor | 6 |
|  | **ETX01807.1** | 9.2E-05 | Qc.I | -DSFNKMAGQLKESFEMLETRVEERTSELAESNKQLQTAKQQAETAN------ | *Candidatus Entotheonella factor* | Bacteria; Nitrospinae/Tectomicrobia group; Candidatus Tectomicrobia; Candidatus Entotheonella | hypothetical protein ETSY1_05970 partial | 6 |
|  | **PIQ39033.1** | 9.2E-05 | Qa.IV | ------SLLSISDRILQVASAIEEQTSVINNINENLSNAKHLSDQGM------ | *Thalassolituus sp. CG17_big_fil_post_rev_8_21_14_2_50_53_8* | Bacteria; Proteobacteria; Gammaproteobacteria; Oceanospirillales; Thalassolituus | hypothetical protein COW58_14000 | 6 |
|  | **SFN52050.1** | 9.2E-05 | Qa.IV | -----ATIGQISSISTSIASAVEQQGAATQEIARSVQTVAQGTQTAATDI--- | *Bradyrhizobium sp. Rc3b* | Bacteria; Proteobacteria; Alphaproteobacteria; Rhizobiales; Bradyrhizobiaceae; Bradyrhizobium | methyl-accepting chemotaxis sensory transducer | 6 |
|  | **WP_027316407.1** | 9.2E-05 | Qa.III | IHEIAQTITEMSQISTGIAAAIEEQGAATREISRNVQEAA------------- | *Microvirga flocculans* | Bacteria; Proteobacteria; Alphaproteobacteria; Rhizobiales; Methylobacteriaceae; Microvirga | methyl-accepting chemotaxis protein | 6 |
|  | **WP_035675816.1** | 9.2E-05 | Qa.III | IEGIGRTIAAINEISTSIAAAIEEQSAATNEISRNVQQAAI------------ | *Azospirillum* | Bacteria; Proteobacteria; Alphaproteobacteria; Rhodospirillales; Rhodospirillaceae | MULTISPECIES: methyl-accepting chemotaxis protein | 6 |
|  | **WP_042691540.1** | 9.2E-05 | Qa.III | IKGVAQVIGRVNDVATSIASAVEQQGAATREISRNIQQAAQ------------ | *Azospirillum sp. B506* | Bacteria; Proteobacteria; Alphaproteobacteria; Rhodospirillales; Rhodospirillaceae; Azospirillum | methyl-accepting chemotaxis protein | 6 |
|  | **WP_045585204.1** | 9.2E-05 | Qa.III | IQGVARVIGRVNDVATSIASAVEQQGAATREISRNIQQAAQ------------ | *Azospirillum thiophilum* | Bacteria; Proteobacteria; Alphaproteobacteria; Rhodospirillales; Rhodospirillaceae; Azospirillum | methyl-accepting chemotaxis protein | 6 |
|  | **WP_092040030.1** | 9.2E-05 | Qa.III | IDTITGRIREINTVATTIAAAVEQQGVATQEIVRNVTQAS------------- | *Methylobacterium pseudosasicola* | Bacteria; Proteobacteria; Alphaproteobacteria; Rhizobiales; Methylobacteriaceae; Methylobacterium | methyl-accepting chemotaxis protein | 6 |
|  | **SCB55810.1** | 9.3E-05 | Qa.IV | VKEVATAMRRIDEVTAAIASAVEEQGAATREISQNVQMAA------------- | *Bradyrhizobium shewense* | Bacteria; Proteobacteria; Alphaproteobacteria; Rhizobiales; Bradyrhizobiaceae; Bradyrhizobium | methyl-accepting chemotaxis protein | 6 |
| | **WP_006613796.1** | 9.3E-05 | Qc.III | -REISTTIERLSEIAAAVAAAVEEQGAATSEISRNIQHAATGTQQVSVNIT-- | *Bradyrhizobium sp. ORS 285* | Bacteria; Proteobacteria; Alphaproteobacteria; Rhizobiales; Bradyrhizobiaceae; Bradyrhizobium | PAS domain S-box protein | 6 |
|  | **WP_013721588.1** | 9.3E-05 | Qa.III | MQEIVHSVGQVSSLLGDISHALSEQSEGI----AHVNQAVAHMDGA------- | *Alicycliphilus* | Bacteria; Proteobacteria; Betaproteobacteria; Burkholderiales; Comamonadaceae | MULTISPECIES: HAMP domain-containing protein | 6 |
|  | **WP_037053929.1** | 9.3E-05 | Qb.I | --RYRQRLQEVLGQTQQLNEELQTQQEELRAANEELEQQANVLKESQ------ | *Pseudomonas oleovorans* | Bacteria; Proteobacteria; Gammaproteobacteria; Pseudomonadales; Pseudomonadaceae; Pseudomonas; Pseudomonas oleovorans/ pseudoalcaligenes group | response regulator | 6 |
|  | **WP_083929280.1** | 9.3E-05 | Qc.III | ---VVETIGKISDISNAISAAVAEQATATQDISNNVMEAAKGTSEVT------ | *Hyphomicrobium zavarzinii* | Bacteria; Proteobacteria; Alphaproteobacteria; Rhizobiales; Hyphomicrobiaceae; Hyphomicrobium | methyl-accepting chemotaxis protein | 6 |
|  | **WP_104213982.1** | 9.3E-05 | Qa.IV | VQELTTDVRQASQVVGQLDTNTQSAGSILDSISAIAEQTN------------- | *Vibrio cyclitrophicus* | Bacteria; Proteobacteria; Gammaproteobacteria; Vibrionales; Vibrionaceae; Vibrio | methyl-accepting chemotaxis protein partial | 6 |
| | **WP_114357361.1** | 9.3E-05 | Qc.III | -REIGGTIERMSEIASTIASAVEEQGAATQEISRNVQQAARGTHDVSSNIA-- | *Rhodopseudomonas pentothenatexigens* | Bacteria; Proteobacteria; Alphaproteobacteria; Rhizobiales; Bradyrhizobiaceae; Rhodopseudomonas | HAMP domain-containing protein | 6 |
| | **CRH04773.1** | 9.4E-05 | Qc.II | ---LSQSANTMKQVSGDLDTTASETLSRVDGMSQVMETVNSSMETTSQAVK-- | *magneto-ovoid bacterium MO-1* | Bacteria | Putative methyl-accepting chemotaxis sensory transducer | 6 |
|  | **EED27869.1** | 9.4E-05 | Qc.III | -DEISGNLSNIKNLSSDMNEAINQLEPIVLDLKRNVDDLNGAIH--------- | *Vibrio sp. 16* | Bacteria; Proteobacteria; Gammaproteobacteria; Vibrionales; Vibrionaceae; Vibrio | methyl-accepting chemotaxis protein | 6 |
| | **EWY36023.1** | 9.4E-05 | Qc.III | IDGIGTIIGQMNEITTAIAAAIEEQSAATRDIATNVSQASRGTDQVSSNIT-- | *Skermanella stibiiresistens SB22* | Bacteria; Proteobacteria; Alphaproteobacteria; Rhodospirillales; Rhodospirillaceae; Skermanella | hypothetical protein N825_31875 | 6 |
|  | **ONG44470.1** | 9.4E-05 | Qa.IV | ------TIAEVSSTTVSIASAVEEQTAATAEIARTVQATAEATEQVTVNI--- | *Roseomonas deserti* | Bacteria; Proteobacteria; Alphaproteobacteria; Rhodospirillales; Acetobacteraceae; Roseomonas | hypothetical protein BKE38_28230 partial | 6 |
|  | **OUX61397.1** | 9.4E-05 | Qa.III | IRNIGSTIGEINEVTTAIAAAVEEQGAATREIARNIQHAASGT---------- | *Afipia sp. TMED4* | Bacteria; Proteobacteria; Alphaproteobacteria; Rhizobiales; Bradyrhizobiaceae; Afipia | methyl-accepting chemotaxis protein | 6 |
|  | **PXZ80675.1** | 9.4E-05 | Qa.III | -VGISEAVSNITEMTTQIAAAAEEQSAVAEEINRNISTIAALAEQTSDE---- | *Pseudomonas aeruginosa* | Bacteria; Proteobacteria; Gammaproteobacteria; Pseudomonadales; Pseudomonadaceae; Pseudomonas | chemotaxis protein partial | 6 |
|  | **WP_016288419.1** | 9.4E-05 | R.IV | -HQLTAGMEQIAGVVQTNASTAEKSAASAKELQTQSEELKV------------ | *Lachnospiraceae bacterium 3-1* | Bacteria; Firmicutes; Clostridia; Clostridiales; Lachnospiraceae | methyl-accepting chemotaxis protein | 6 |
|  | **WP_020086559.1** | 9.4E-05 | Qa.III | IQEITGVINSINDISNGIAAAIEEQGAVTQEISANMLTASNGV---------- | *Hyphomicrobium zavarzinii* | Bacteria; Proteobacteria; Alphaproteobacteria; Rhizobiales; Hyphomicrobiaceae; Hyphomicrobium | methyl-accepting chemotaxis protein | 6 |
|  | **WP_024024713.1** | 9.4E-05 | Qb.I | --NGAKILRQTVETSNQIVSEIENSTHLIEQLNKQSENIAKIVT--------- | *Marinomonas profundimaris* | Bacteria; Proteobacteria; Gammaproteobacteria; Oceanospirillales; Marinomonas | PAS domain S-box protein | 6 |
| | **WP_044830158.1** | 9.4E-05 | Qc.III | IEKVTQTIASINEIATGISSAVSQQSSAVTEIANSATSASSGSNEVVSHVNE- | *Thalassospira sp. HJ* | Bacteria; Proteobacteria; Alphaproteobacteria; Rhodospirillales; Rhodospirillaceae; Thalassospira | methyl-accepting chemotaxis protein | 6 |
|  | **WP_051283787.1** | 9.4E-05 | Qa.I | IDQISKVIREIENIVSTIASSIEEQSATTKEIATNVAQAS------------- | *Desulforegula conservatrix* | Bacteria; Proteobacteria; Deltaproteobacteria; Desulfobacterales; Desulfobacteraceae; Desulforegula | methyl-accepting chemotaxis protein | 6 |
| | **WP_062958303.1** | 9.4E-05 | Qc.III.b | -DDISDVISRITNITNTISDAVREQADATGEIAQNVEQAAIGTQEVSSNIE-- | *Thalassospira xiamenensis* | Bacteria; Proteobacteria; Alphaproteobacteria; Rhodospirillales; Rhodospirillaceae; Thalassospira | HAMP domain-containing protein | 6 |

| | | | | | | | | |
|---|---|---|---|---|---|---|---|---|
|  | WP_111355392.1 | 9.4E-05 | Qa.IV | --EVAATIRELEGISSSISAAVEEQSAATSDIAQNVGQTATHVG--------- | *Rhodoplanes elegans* | Bacteria; Proteobacteria; Alphaproteobacteria; Rhizobiales; Hyphomicrobiaceae; Rhodoplanes | hypothetical protein | 6 |
|  | WP_114089184.1 | 9.4E-05 | Qa.I | ---IGGSIDEVSRLSQDISDAIQHQTSAIEDISASAKEVSSHM---------- | *Thalassospira profundimaris* | Bacteria; Proteobacteria; Alphaproteobacteria; Rhodospirillales; Rhodospirillaceae; Thalassospira | chemotaxis protein | 6 |
| | WP_114122164.1 | 9.4E-05 | Qc.III.b | -DDISDVISRITNITNTISDAVREQADATGEIAQNVEQAAIGTQEVSSNIE-- | *Thalassospira xianhensis* | Bacteria; Proteobacteria; Alphaproteobacteria; Rhodospirillales; Rhodospirillaceae; Thalassospira | HAMP domain-containing protein | 6 |
|  | AAZ37152.1 | 9.5E-05 | Qa.III | ---ISQALRNLNSLNTSIASATLQQTHVVDDINQNVTQAAQLSQ--------- | *Pseudomonas savastanoi pv. phaseolicola 1448A* | Bacteria; Proteobacteria; Gammaproteobacteria; Pseudomonadales; Pseudomonadaceae; Pseudomonas | methyl-accepting chemotaxis protein | 6 |
|  | ARD14671.1 | 9.5E-05 | Qa.III | ---ISQALRNLNSLNTSIASATLQQTHVVDDINQNVTQAAQLSQ--------- | *Pseudomonas savastanoi pv. savastanoi NCPPB 3335* | Bacteria; Proteobacteria; Gammaproteobacteria; Pseudomonadales; Pseudomonadaceae; Pseudomonas | chemotaxis protein | 6 |
|  | EGH03054.1 | 9.5E-05 | Qa.III | ---ISQALRNLNSLNTSIASATLQQTHVVDDINQNVTQAAQLSQ--------- | *Pseudomonas amygdali pv. aesculi str. 0893_23* | Bacteria; Proteobacteria; Gammaproteobacteria; Pseudomonadales; Pseudomonadaceae; Pseudomonas; Pseudomonas amygdali | methyl-accepting chemotaxis protein | 6 |
|  | KIY17847.1 | 9.5E-05 | Qa.III | ---ISQALRNLNSLNTSIASATLQQTHVVDDINQNVTQAAQLSQ--------- | *Pseudomonas amygdali pv. tabaci* | Bacteria; Proteobacteria; Gammaproteobacteria; Pseudomonadales; Pseudomonadaceae; Pseudomonas; Pseudomonas amygdali | chemotaxis protein | 6 |
|  | KKY49714.1 | 9.5E-05 | Qa.III | ---ISQALRNLNSLNTSIASATLQQTHVVDDINQNVTQAAQLSQ--------- | *Pseudomonas amygdali pv. tabaci str. ATCC 11528* | Bacteria; Proteobacteria; Gammaproteobacteria; Pseudomonadales; Pseudomonadaceae; Pseudomonas; Pseudomonas amygdali | chemotaxis protein | 6 |
|  | KKY59389.1 | 9.5E-05 | Qa.III | ---ISQALRNLNSLNTSIASATLQQTHVVDDINQNVTQAAQLSQ--------- | *Pseudomonas amygdali pv. lachrymans* | Bacteria; Proteobacteria; Gammaproteobacteria; Pseudomonadales; Pseudomonadaceae; Pseudomonas; Pseudomonas amygdali | chemotaxis protein | 6 |
|  | KPB69791.1 | 9.5E-05 | Qa.III | ---ISQALRNLNSLNTSIASATLQQTHVVDDINQNVTQAAQLSQ--------- | *Pseudomonas amygdali pv. mellea* | Bacteria; Proteobacteria; Gammaproteobacteria; Pseudomonadales; Pseudomonadaceae; Pseudomonas; Pseudomonas amygdali | Methyl-accepting chemotaxis protein | 6 |
|  | KPX80263.1 | 9.5E-05 | Qa.III | ---ISQALRNLNSLNTSIASATLQQTHVVDDINQNVTQAAQLSQ--------- | *Pseudomonas amygdali pv. photiniae* | Bacteria; Proteobacteria; Gammaproteobacteria; Pseudomonadales; Pseudomonadaceae; Pseudomonas; Pseudomonas amygdali | Methyl-accepting chemotaxis protein | 6 |
|  | KWS38879.1 | 9.5E-05 | Qa.III | ---ISQALRNLNSLNTSIASATLQQTHVVDDINQNVTQAAQLSQ--------- | *Pseudomonas syringae pv. rhaphiolepidis* | Bacteria; Proteobacteria; Gammaproteobacteria; Pseudomonadales; Pseudomonadaceae; Pseudomonas; Pseudomonas syringae | chemotaxis protein | 6 |
|  | KWS50141.1 | 9.5E-05 | Qa.III | ---ISQALRNLNSLNTSIASATLQQTHVVDDINQNVTQAAQLSQ--------- | *Pseudomonas amygdali pv. myricae* | Bacteria; Proteobacteria; Gammaproteobacteria; Pseudomonadales; Pseudomonadaceae; Pseudomonas; Pseudomonas amygdali | chemotaxis protein | 6 |
|  | KWS84660.1 | 9.5E-05 | Qa.III | ---ISQALRNLNSLNTSIASATLQQTHVVDDINQNVTQAAQLSQ--------- | *Pseudomonas amygdali pv. dendropanacis* | Bacteria; Proteobacteria; Gammaproteobacteria; Pseudomonadales; Pseudomonadaceae; Pseudomonas; Pseudomonas amygdali | chemotaxis protein | 6 |
|  | KWS97977.1 | 9.5E-05 | Qa.III | ---ISQALRNLNSLNTSIASATLQQTHVVDDINQNVTQAAQLSQ--------- | *Pseudomonas syringae pv. daphniphylli* | Bacteria; Proteobacteria; Gammaproteobacteria; Pseudomonadales; Pseudomonadaceae; Pseudomonas; Pseudomonas syringae | chemotaxis protein | 6 |
|  | KWT10528.1 | 9.5E-05 | Qa.III | ---ISQALRNLNSLNTSIASATLQQTHVVDDINQNVTQAAQLSQ--------- | *Pseudomonas syringae pv. broussonetiae* | Bacteria; Proteobacteria; Gammaproteobacteria; Pseudomonadales; Pseudomonadaceae; Pseudomonas; Pseudomonas syringae | chemotaxis protein | 6 |
|  | KWT38534.1 | 9.5E-05 | Qa.III | ---ISQALRNLNSLNTSIASATLQQTHVVDDINQNVTQAAQLSQ--------- | *Pseudomonas amygdali pv. aesculi* | Bacteria; Proteobacteria; Gammaproteobacteria; Pseudomonadales; Pseudomonadaceae; Pseudomonas; Pseudomonas amygdali | chemotaxis protein | 6 |
|  | ODS47502.1 | 9.5E-05 | Qa.III | ---ISQALRNLNSLNTSIASATLQQTHVVDDINQNVTQAAQLSQ--------- | *Pseudomonas sp. BDAL1* | Bacteria; Proteobacteria; Gammaproteobacteria; Pseudomonadales; Pseudomonadaceae; Pseudomonas | chemotaxis protein | 6 |
|  | PHN83063.1 | 9.5E-05 | Qa.III | ---ISQALRNLNSLNTSIASATLQQTHVVDDINQNVTQAAQLSQ--------- | *Pseudomonas syringae pv. cerasicola* | Bacteria; Proteobacteria; Gammaproteobacteria; Pseudomonadales; Pseudomonadaceae; Pseudomonas; Pseudomonas syringae | chemotaxis protein | 6 |
|  | POD50340.1 | 9.5E-05 | Qa.III | ---ISQALRNLNSLNTSIASATLQQTHVVDDINQNVTQAAQLSQ--------- | *Pseudomonas amygdali pv. morsprunorum* | Bacteria; Proteobacteria; Gammaproteobacteria; Pseudomonadales; Pseudomonadaceae; Pseudomonas; Pseudomonas amygdali | chemotaxis protein | 6 |
|  | PPS32877.1 | 9.5E-05 | Qa.III | ---ISQALRNLNSLNTSIASATLQQTHVVDDINQNVTQAAQLSQ--------- | *Pseudomonas amygdali pv. morsprunorum* | Bacteria; Proteobacteria; Gammaproteobacteria; Pseudomonadales; Pseudomonadaceae; Pseudomonas; Pseudomonas amygdali | chemotaxis protein | 6 |
| | RCK05547.1 | 9.5E-05 | Qc.III.b | -DDISDVISRITNITNTISDAVREQADATGEIAQNVEQAAIGTQEVSSNIE-- | *Thalassospira xianhensis MCCC 1A02616* | Bacteria; Proteobacteria; Alphaproteobacteria; Rhodospirillales; Rhodospirillaceae; Thalassospira | chemotaxis protein | 6 |
| | RCK45163.1 | 9.5E-05 | Qc.III.b | -DDISDVISRITNITNTISDAVREQADATGEIAQNVEQAAIGTQEVSSNIE-- | *Thalassospira xiamenensis* | Bacteria; Proteobacteria; Alphaproteobacteria; Rhodospirillales; Rhodospirillaceae; Thalassospira | chemotaxis protein | 6 |
|  | SIS90787.1 | 9.5E-05 | Qa.III | IHAIVGTIEQISHIAAGIAAAVEEQGAATQEISRNVQEAA------------- | *Insolitispirillum peregrinum* | Bacteria; Proteobacteria; Alphaproteobacteria; Rhodospirillales; Rhodospirillaceae; Insolitispirillum | methyl-accepting chemotaxis sensory transducer with Cache sensor | 6 |
|  | SPD81415.1 | 9.5E-05 | Qa.III | ---ISQALRNLNSLNTSIASATLQQTHVVDDINQNVTQAAQLSQ--------- | *Pseudomonas syringae* | Bacteria; Proteobacteria; Gammaproteobacteria; Pseudomonadales; Pseudomonadaceae; Pseudomonas | methyl-accepting chemotaxis protein | 6 |
|  | SPF16462.1 | 9.5E-05 | Qa.III | ---ISQALRNLNSLNTSIASATLQQTHVVDDINQNVTQAAQLSQ--------- | *Pseudomonas syringae pv. cerasicola* | Bacteria; Proteobacteria; Gammaproteobacteria; Pseudomonadales; Pseudomonadaceae; Pseudomonas; Pseudomonas syringae | methyl-accepting chemotaxis protein | 6 |
|  | WP_002552764.1 | 9.5E-05 | Qa.III | ---ISQALRNLNSLNTSIASATLQQTHVVDDINQNVTQAAQLSQ--------- | *Pseudomonas syringae group genomosp. 2* | Bacteria; Proteobacteria; Gammaproteobacteria; Pseudomonadales; Pseudomonadaceae; Pseudomonas | MULTISPECIES: methyl-accepting chemotaxis protein | 6 |
|  | WP_029241052.1 | 9.5E-05 | Qa.III | ---ISQALRNLNSLNTSIASATLQQTHVVDDINQNVTQAAQLSQ--------- | *Pseudomonas amygdali* | Bacteria; Proteobacteria; Gammaproteobacteria; Pseudomonadales; Pseudomonadaceae; Pseudomonas | methyl-accepting chemotaxis protein | 6 |
|  | WP_038138736.1 | 9.5E-05 | Qc.III | -DEISGNLSNIKNLSSDMNEAINQLEPIVLDLKRNVDDLNGAIH--------- | *Vibrio caribbeanicus* | Bacteria; Proteobacteria; Gammaproteobacteria; Vibrionales; Vibrionaceae; Vibrio | methyl-accepting chemotaxis protein | 6 |
|  | WP_038214779.1 | 9.5E-05 | Qc.III | -DEISGNLSNIKNLSSDMNEAINQLEPIVLDLKRNVDDLNGAIH--------- | *Vibrio variabilis* | Bacteria; Proteobacteria; Gammaproteobacteria; Vibrionales; Vibrionaceae; Vibrio | methyl-accepting chemotaxis protein | 6 |
|  | WP_043886650.1 | 9.5E-05 | Qc.III | -DEISGNLSNIKNLSSDMNEAINQLEPIVLDLKRNVDDLNGAIH--------- | *Vibrio sp. 16* | Bacteria; Proteobacteria; Gammaproteobacteria; Vibrionales; Vibrionaceae; Vibrio | methyl-accepting chemotaxis protein | 6 |
|  | WP_076400830.1 | 9.5E-05 | Qa.III | IHAIVGTIEQISHIAAGIAAAVEEQGAATQEISRNVQEAA------------- | *Insolitispirillum peregrinum* | Bacteria; Proteobacteria; Alphaproteobacteria; Rhodospirillales; Rhodospirillaceae; Insolitispirillum | HAMP domain-containing protein | 6 |
|  | WP_080164716.1 | 9.5E-05 | Qa.III | ---ISQALRNLNSLNTSIASATLQQTHVVDDINQNVTQAAQLSQ--------- | *Pseudomonas amygdali* | Bacteria; Proteobacteria; Gammaproteobacteria; Pseudomonadales; Pseudomonadaceae; Pseudomonas | methyl-accepting chemotaxis protein | 6 |
|  | WP_080392406.1 | 9.5E-05 | Qa.III | ---ISQALRNLNSLNTSIASATLQQTHVVDDINQNVTQAAQLSQ--------- | *Pseudomonas syringae* | Bacteria; Proteobacteria; Gammaproteobacteria; Pseudomonadales; Pseudomonadaceae; Pseudomonas | methyl-accepting chemotaxis protein | 6 |
|  | WP_080438307.1 | 9.5E-05 | Qa.III | ---ISQALRNLNSLNTSIASATLQQTHVVDDINQNVTQAAQLSQ--------- | *Pseudomonas syringae group* | Bacteria; Proteobacteria; Gammaproteobacteria; Pseudomonadales; Pseudomonadaceae; Pseudomonas | MULTISPECIES: methyl-accepting chemotaxis protein | 6 |
|  | WP_080438637.1 | 9.5E-05 | Qa.III | ---ISQALRNLNSLNTSIASATLQQTHVVDDINQNVTQAAQLSQ--------- | *Pseudomonas syringae group* | Bacteria; Proteobacteria; Gammaproteobacteria; Pseudomonadales; Pseudomonadaceae; Pseudomonas | MULTISPECIES: methyl-accepting chemotaxis protein | 6 |
|  | WP_080544363.1 | 9.5E-05 | Qa.III | ---ISQALRNLNSLNTSIASATLQQTHVVDDINQNVTQAAQLSQ--------- | *Pseudomonas amygdali* | Bacteria; Proteobacteria; Gammaproteobacteria; Pseudomonadales; Pseudomonadaceae; Pseudomonas | methyl-accepting chemotaxis protein | 6 |
|  | WP_080572400.1 | 9.5E-05 | Qa.III | ---ISQALRNLNSLNTSIASATLQQTHVVDDINQNVTQAAQLSQ--------- | *Pseudomonas savastanoi* | Bacteria; Proteobacteria; Gammaproteobacteria; Pseudomonadales; Pseudomonadaceae; Pseudomonas | methyl-accepting chemotaxis protein | 6 |
|  | WP_080900034.1 | 9.5E-05 | Qa.III | ---ISQALRNLNSLNTSIASATLQQTHVVDDINQNVTQAAQLSQ--------- | *Pseudomonas amygdali* | Bacteria; Proteobacteria; Gammaproteobacteria; Pseudomonadales; Pseudomonadaceae; Pseudomonas | methyl-accepting chemotaxis protein | 6 |
|  | WP_080953453.1 | 9.5E-05 | Qa.III | ---ISQALRNLNSLNTSIASATLQQTHVVDDINQNVTQAAQLSQ--------- | *Pseudomonas syringae group* | Bacteria; Proteobacteria; Gammaproteobacteria; Pseudomonadales; Pseudomonadaceae; Pseudomonas | MULTISPECIES: methyl-accepting chemotaxis protein | 6 |
|  | WP_081002624.1 | 9.5E-05 | Qa.III | ---ISQALRNLNSLNTSIASATLQQTHVVDDINQNVTQAAQLSQ--------- | *Pseudomonas savastanoi* | Bacteria; Proteobacteria; Gammaproteobacteria; Pseudomonadales; Pseudomonadaceae; Pseudomonas | methyl-accepting chemotaxis protein | 6 |
|  | WP_081004480.1 | 9.5E-05 | Qa.III | ---ISQALRNLNSLNTSIASATLQQTHVVDDINQNVTQAAQLSQ--------- | *Pseudomonas amygdali* | Bacteria; Proteobacteria; Gammaproteobacteria; Pseudomonadales; Pseudomonadaceae; Pseudomonas | methyl-accepting chemotaxis protein | 6 |
|  | WP_081007527.1 | 9.5E-05 | Qa.III | ---ISQALRNLNSLNTSIASATLQQTHVVDDINQNVTQAAQLSQ--------- | *Pseudomonas amygdali* | Bacteria; Proteobacteria; Gammaproteobacteria; Pseudomonadales; Pseudomonadaceae; Pseudomonas | methyl-accepting chemotaxis protein | 6 |
|  | WP_081025044.1 | 9.5E-05 | Qa.III | ---ISQALRNLNSLNTSIASATLQQTHVVDDINQNVTQAAQLSQ--------- | *Pseudomonas amygdali* | Bacteria; Proteobacteria; Gammaproteobacteria; Pseudomonadales; Pseudomonadaceae; Pseudomonas | methyl-accepting chemotaxis protein | 6 |
|  | WP_082303429.1 | 9.5E-05 | Qa.III | ---ISQALRNLNSLNTSIASATLQQTHVVDDINQNVTQAAQLSQ--------- | *Pseudomonas meliae* | Bacteria; Proteobacteria; Gammaproteobacteria; Pseudomonadales; Pseudomonadaceae; Pseudomonas | methyl-accepting chemotaxis protein | 6 |
|  | WP_082333329.1 | 9.5E-05 | Qa.III | ---ISQALRNLNSLNTSIASATLQQTHVVDDINQNVTQAAQLSQ--------- | *Pseudomonas syringae* | Bacteria; Proteobacteria; Gammaproteobacteria; Pseudomonadales; Pseudomonadaceae; Pseudomonas | methyl-accepting chemotaxis protein | 6 |
| | WP_084165333.1 | 9.5E-05 | Qc.III | IDGIGTIIGQMNEITTAIAAAIEEQSAATRDIATNVSQASRGTDQVSSNIT-- | *Skermanella stibiiresistens* | Bacteria; Proteobacteria; Alphaproteobacteria; Rhodospirillales; Rhodospirillaceae; Skermanella | methyl-accepting chemotaxis protein | 6 |
|  | WP_088871051.1 | 9.5E-05 | Qc.III.c | -DHINGTIAQLSEIAVVIASAVEEQTAATSEIARNVNEAARGV---------- | *Nitrospirillum amazonense* | Bacteria; Proteobacteria; Alphaproteobacteria; Rhodospirillales; Rhodospirillaceae; Nitrospirillum | methyl-accepting chemotaxis protein | 6 |
|  | WP_108514278.1 | 9.5E-05 | Qc.III | --NIGGIIGEVNEVATAIAAAVQQQGAATQEITRSTQFAAQGTKNVS------ | *Bradyrhizobium algeriense* | Bacteria; Proteobacteria; Alphaproteobacteria; Rhizobiales; Bradyrhizobiaceae; Bradyrhizobium | methyl-accepting chemotaxis protein | 6 |
|  | KPB19119.1 | 9.6E-05 | Qa.III | ---ISQALRNLNSLNTSIASATLQQTHVVDDINQNVTQAAQLSQ--------- | *Pseudomonas amygdali pv. sesami* | Bacteria; Proteobacteria; Gammaproteobacteria; Pseudomonadales; Pseudomonadaceae; Pseudomonas; Pseudomonas amygdali | Methyl-accepting chemotaxis protein | 6 |
|  | KPB39424.1 | 9.6E-05 | Qa.III | ---ISQALRNLNSLNTSIASATLQQTHVVDDINQNVTQAAQLSQ--------- | *Pseudomonas savastanoi pv. phaseolicola* | Bacteria; Proteobacteria; Gammaproteobacteria; Pseudomonadales; Pseudomonadaceae; Pseudomonas | Methyl-accepting chemotaxis protein | 6 |
|  | KPB59542.1 | 9.6E-05 | Qa.III | ---ISQALRNLNSLNTSIASATLQQTHVVDDINQNVTQAAQLSQ--------- | *Pseudomonas savastanoi pv. phaseolicola* | Bacteria; Proteobacteria; Gammaproteobacteria; Pseudomonadales; Pseudomonadaceae; Pseudomonas | Methyl-accepting chemotaxis protein | 6 |
|  | KPB97099.1 | 9.6E-05 | Qa.III | ---ISQALRNLNSLNTSIASATLQQTHVVDDINQNVTQAAQLSQ--------- | *Pseudomonas amygdali pv. lachrymans* | Bacteria; Proteobacteria; Gammaproteobacteria; Pseudomonadales; Pseudomonadaceae; Pseudomonas; Pseudomonas amygdali | Methyl-accepting chemotaxis protein | 6 |
|  | KPC41061.1 | 9.6E-05 | Qa.III | ---ISQALRNLNSLNTSIASATLQQTHVVDDINQNVTQAAQLSQ--------- | *Pseudomonas savastanoi pv. glycinea* | Bacteria; Proteobacteria; Gammaproteobacteria; Pseudomonadales; Pseudomonadaceae; Pseudomonas | Methyl-accepting chemotaxis protein | 6 |
|  | KPX04698.1 | 9.6E-05 | Qa.III | ---ISQALRNLNSLNTSIASATLQQTHVVDDINQNVTQAAQLSQ--------- | *Pseudomonas syringae pv. cunninghamiae* | Bacteria; Proteobacteria; Gammaproteobacteria; Pseudomonadales; Pseudomonadaceae; Pseudomonas; Pseudomonas syringae | Methyl-accepting chemotaxis protein | 6 |
|  | KPX24415.1 | 9.6E-05 | Qa.III | ---ISQALRNLNSLNTSIASATLQQTHVVDDINQNVTQAAQLSQ--------- | *Pseudomonas amygdali pv. dendropanacis* | Bacteria; Proteobacteria; Gammaproteobacteria; Pseudomonadales; Pseudomonadaceae; Pseudomonas; Pseudomonas amygdali | Methyl-accepting chemotaxis protein | 6 |
|  | KPX77863.1 | 9.6E-05 | Qa.III | ---ISQALRNLNSLNTSIASATLQQTHVVDDINQNVTQAAQLSQ--------- | *Pseudomonas amygdali pv. mellea* | Bacteria; Proteobacteria; Gammaproteobacteria; Pseudomonadales; Pseudomonadaceae; Pseudomonas; Pseudomonas amygdali | Methyl-accepting chemotaxis protein | 6 |
|  | KPX92562.1 | 9.6E-05 | Qa.III | ---ISQALRNLNSLNTSIASATLQQTHVVDDINQNVTQAAQLSQ--------- | *Pseudomonas amygdali pv. myricae* | Bacteria; Proteobacteria; Gammaproteobacteria; Pseudomonadales; Pseudomonadaceae; Pseudomonas; Pseudomonas amygdali | Methyl-accepting chemotaxis protein | 6 |
|  | ODS00173.1 | 9.6E-05 | Qa.IV | MQSIVTTMGDVNQNTQEIAGAVQQQSAATSEISNSVRQAA------------- | *Methyloceanibacter methanicus* | Bacteria; Proteobacteria; Alphaproteobacteria; Rhizobiales; Methyloceanibacter | hypothetical protein AUC68_03440 | 6 |
|  | PIZ30735.1 | 9.6E-05 | Qc.III.c | ------AIHRIRDNATTVSAAVEEQSGVVKTIAGSVRDAASRVNDVDANM--- | *Alphaproteobacteria bacterium CG_4_10_14_0_8_um_filter_53_9* | Bacteria; Proteobacteria; Alphaproteobacteria | hypothetical protein COY40_03790 | 6 |

| | | | | | | | |
|---|---|---|---|---|---|---|---|
| WP_052063587.1 | 9.6E-05 | Qc.III | --EIVNSVSVISDMNTQIATAAEEQSQVAQDMDHNINKVAEQARRTN------ | *Nitrincola sp. A-D6* | Bacteria; Proteobacteria; Gammaproteobacteria; Oceanospirillales; Nitrincola | hypothetical protein | 6 |
| WP_074119753.1 | 9.6E-05 | Qa.III.b | -EAIAQTITEINGITASIATSIEQQGLATREIARNVQEASA------------ | *Bradyrhizobium sp. AS23.2* | Bacteria; Proteobacteria; Alphaproteobacteria; Rhizobiales; Bradyrhizobiaceae; Bradyrhizobium | methyl-accepting chemotaxis protein | 6 |
| WP_075766186.1 | 9.6E-05 | Qa.III | LEQINDSVAMISDMATQIATAAEEQTHVTDEITQNTTTIK------------- | *Photobacterium proteolyticum* | Bacteria; Proteobacteria; Gammaproteobacteria; Vibrionales; Vibrionaceae; Photobacterium | methyl-accepting chemotaxis protein | 6 |
| WP_084807637.1 | 9.6E-05 | Qa.III.b | -EAIAQTITEINGITASIATSIEQQGLATREIARNVQEASA------------ | *Bradyrhizobium sp. NAS80.1* | Bacteria; Proteobacteria; Alphaproteobacteria; Rhizobiales; Bradyrhizobiaceae; Bradyrhizobium | methyl-accepting chemotaxis protein | 6 |
| WP_092331739.1 | 9.6E-05 | Qa.IV | -QNIENSMNDVSNAMDDVAATVDESARGIGNVASAVNDVTDNMKE-------- | *Butyrivibrio sp. YAB3001* | Bacteria; Firmicutes; Clostridia; Clostridiales; Lachnospiraceae; Butyrivibrio | methyl-accepting chemotaxis protein | 6 |
| WP_109513538.1 | 9.6E-05 | Qb.I | LQRTKLQLQETIEHSEVSAEELKASNEEMQAINEELRSATEELETSK------ | *Pseudomonas sp. F51* | Bacteria; Proteobacteria; Gammaproteobacteria; Pseudomonadales; Pseudomonadaceae; Pseudomonas | PAS domain S-box protein | 6 |
| ACY31580.1 | 9.7E-05 | Qa.III | MKEIMQSVGQVTGLLSEISGALQQQSEGI----AHVNTAVAHMDSTNQE---- | *Comamonas testosteroni CNB-2* | Bacteria; Proteobacteria; Betaproteobacteria; Burkholderiales; Comamonadaceae; Comamonas | methyl-accepting chemotaxis sensory transducer | 6 |
| EFW81253.1 | 9.7E-05 | Qa.III | ---ISQALRNLNSLNTSIASATLQQTHVVDDINQNVTQAAQLSQ--------- | *Pseudomonas savastanoi pv. glycinea str. B076* | Bacteria; Proteobacteria; Gammaproteobacteria; Pseudomonadales; Pseudomonadaceae; Pseudomonas | methyl-accepting chemotaxis protein | 6 |
| GAW77303.1 | 9.7E-05 | Qa.III | MKEIMQSVGQVTGLLSEISGALQQQSEGI----AHVNTAVAHMDSTNQE---- | *Comamonas testosteroni* | Bacteria; Proteobacteria; Betaproteobacteria; Burkholderiales; Comamonadaceae; Comamonas | methyl-accepting chemotaxis protein I | 6 |
| KPC14328.1 | 9.7E-05 | Qa.III | ---ISQALRNLNSLNTSIASATLQQTHVVDDINQNVTQAAQLSQ--------- | *Pseudomonas amygdali pv. lachrymans* | Bacteria; Proteobacteria; Gammaproteobacteria; Pseudomonadales; Pseudomonadaceae; Pseudomonas; Pseudomonas amygdali | Methyl-accepting chemotaxis protein | 6 |
| KPX78518.1 | 9.7E-05 | Qa.III | ---ISQALRNLNSLNTSIASATLQQTHVVDDINQNVTQAAQLSQ--------- | *Pseudomonas meliae* | Bacteria; Proteobacteria; Gammaproteobacteria; Pseudomonadales; Pseudomonadaceae; Pseudomonas | Methyl-accepting chemotaxis protein | 6 |
| KPY46108.1 | 9.7E-05 | Qa.III | ---ISQALRNLNSLNTSIASATLQQTHVVDDINQNVTQAAQLSQ--------- | *Pseudomonas syringae pv. rhaphiolepidis* | Bacteria; Proteobacteria; Gammaproteobacteria; Pseudomonadales; Pseudomonadaceae; Pseudomonas; Pseudomonas syringae | Methyl-accepting chemotaxis protein | 6 |
| WP_002712221.1 | 9.7E-05 | Qc.III | -KAIGHTIEQISDIATSIAAAVEQQGAATQEIARSVQAAASV----------- | *Afipia clevelandensis* | Bacteria; Proteobacteria; Alphaproteobacteria; Rhizobiales; Bradyrhizobiaceae; Afipia | methyl-accepting chemotaxis protein | 6 |
| WP_008965077.1 | 9.7E-05 | Qc.III | ---ISGTIERLSEISSTIAAAVEEQGAATQEISRNVQQAARGTQQVSANIA-- | *Bradyrhizobium sp. STM 3809* | Bacteria; Proteobacteria; Alphaproteobacteria; Rhizobiales; Bradyrhizobiaceae; Bradyrhizobium | methyl-accepting chemotaxis protein | 6 |
| WP_009736711.1 | 9.7E-05 | Qc.III | -KAIGHTIEQISDIATSIAAAVEQQGAATQEIARSVQAAASV----------- | *Bradyrhizobiaceae bacterium SG-6C* | Bacteria; Proteobacteria; Alphaproteobacteria; Rhizobiales; Bradyrhizobiaceae | HAMP domain-containing protein | 6 |
| WP_077949307.1 | 9.7E-05 | R.IV | -TQIAQAMTEMDTTTQQNAALVEESSAAASSLEEQALKLEEAISVFH------ | *Salmonella enterica* | Bacteria; Proteobacteria; Gammaproteobacteria; Enterobacterales; Enterobacteriaceae; Salmonella | HAMP domain-containing protein | 6 |
| WP_079226202.1 | 9.7E-05 | R.IV | -TQIAQAMTEMDTTTQQNAALVEESSAAASSLEEQALKLEEAISVFH------ | *Citrobacter werkmanii* | Bacteria; Proteobacteria; Gammaproteobacteria; Enterobacterales; Enterobacteriaceae; Citrobacter; Citrobacter freundii complex | HAMP domain-containing protein | 6 |
| WP_080669823.1 | 9.7E-05 | Qa.IV | --EIEANVNILSQGINEMSQSINEQTEAINQINEAVATVDEQTKQ-NV----- | *Campylobacter fetus* | Bacteria; Proteobacteria; Epsilonproteobacteria; Campylobacterales; Campylobacteraceae; Campylobacter | methyl-accepting chemotaxis protein | 6 |
| WP_111355211.1 | 9.7E-05 | Qc.III | -RQIGDTIGRLSEISSSIAAAVEEQGAATREISRNVQQAAQGTVEVSSH---- | *Rhodoplanes elegans* | Bacteria; Proteobacteria; Alphaproteobacteria; Rhizobiales; Hyphomicrobiaceae; Rhodoplanes | PAS domain S-box protein | 6 |
| ABP85892.1 | 9.8E-05 | Qb.I | --DAVQRISAALGQFEQTMQAFKNATADIARTRQEMTDQQGEI---------- | *Pseudomonas mendocina ymp* | Bacteria; Proteobacteria; Gammaproteobacteria; Pseudomonadales; Pseudomonadaceae; Pseudomonas | methyl-accepting chemotaxis sensory transducer | 6 |
| BAR57409.1 | 9.8E-05 | Qa.IV | -----ATIGQISSISTSIASAVEQQGAATQEIARSVQTVAQGTHAAATD---- | *Bradyrhizobium diazoefficiens* | Bacteria; Proteobacteria; Alphaproteobacteria; Rhizobiales; Bradyrhizobiaceae; Bradyrhizobium | putative methyl accepting chemotaxis protein | 6 |
| KRQ15896.1 | 9.8E-05 | Qc.III | --DIAGIIDKMREISRSISAEIEAQSIVTNDISKRITLISSDM---------- | *Bradyrhizobium manausense* | Bacteria; Proteobacteria; Alphaproteobacteria; Rhizobiales; Bradyrhizobiaceae; Bradyrhizobium | hypothetical protein AOQ71_07060 | 6 |
| PKM33934.1 | 9.8E-05 | Qb.I | --DAVQRISAALGQFEQTMQAFKNATADIARTRQEMTDQQGEI---------- | *Gammaproteobacteria bacterium HGW-Gammaproteobacteria-12* | Bacteria; Proteobacteria; Gammaproteobacteria | methyl-accepting chemotaxis protein | 6 |
| SEL45287.1 | 9.8E-05 | Qc.III | LEDITRSIGTINDMNTQIASAAEEQSAVAEDVNRSLTRISSGVEQ-------- | *Ectothiorhodospira marina* | Bacteria; Proteobacteria; Gammaproteobacteria; Chromatiales; Ectothiorhodospiraceae; Ectothiorhodospira | methyl-accepting chemotaxis protein | 6 |
| SMH58956.1 | 9.8E-05 | Qc.III | -QSVASVIDRLSGIATAISAAVEEQNAATAEISRSVQQAAAGT---------- | *Azospirillum lipoferum* | Bacteria; Proteobacteria; Alphaproteobacteria; Rhodospirillales; Rhodospirillaceae; Azospirillum | methyl-accepting chemotaxis sensory transducer with Cache sensor | 6 |
| WP_074076408.1 | 9.8E-05 | Qa.IV | -----ATIGQISSISTSIASAVEQQGAATQEIARSVQTVAQGTHAAATD---- | *Bradyrhizobium diazoefficiens* | Bacteria; Proteobacteria; Alphaproteobacteria; Rhizobiales; Bradyrhizobiaceae; Bradyrhizobium | methyl-accepting chemotaxis protein | 6 |
| WP_083901624.1 | 9.8E-05 | Qa.IV | IQAIAQRIQDISGIASAIAAAVEEQEASTHEIARNVQEAA------------- | *Azospirillum sp. B4* | Bacteria; Proteobacteria; Alphaproteobacteria; Rhodospirillales; Rhodospirillaceae; Azospirillum | hypothetical protein | 6 |
| WP_092192111.1 | 9.8E-05 | Qa.IV | -----ATIGQISSISTSIASAVEQQGAATQEIARSVQTVAQGTHAAATD---- | *Bradyrhizobium sp. cf659* | Bacteria; Proteobacteria; Alphaproteobacteria; Rhizobiales; Bradyrhizobiaceae; Bradyrhizobium | methyl-accepting chemotaxis protein | 6 |
| WP_092686057.1 | 9.8E-05 | Qa.III.b | --AISRRISQINEVSSTIAAAVEQQGAATREIAMNVQQASAGT---------- | *Rhodopseudomonas pseudopalustris* | Bacteria; Proteobacteria; Alphaproteobacteria; Rhizobiales; Bradyrhizobiaceae; Rhodopseudomonas | methyl-accepting chemotaxis protein | 6 |
| WP_096827057.1 | 9.8E-05 | Qb.I | --DAVQRISAALGQFEQTMQAFKNATADIARTRQEMTDQQGEI---------- | *Pseudomonas mendocina* | Bacteria; Proteobacteria; Gammaproteobacteria; Pseudomonadales; Pseudomonadaceae; Pseudomonas | methyl-accepting chemotaxis protein | 6 |
| WP_098736920.1 | 9.8E-05 | Qa.III | IRDVAQVIGRVNDVATSIASAVEQQGAATREISRNIQQAA------------- | *Azospirillum sp. B2* | Bacteria; Proteobacteria; Alphaproteobacteria; Rhodospirillales; Rhodospirillaceae; Azospirillum | methyl-accepting chemotaxis protein | 6 |
| WP_109050339.1 | 9.8E-05 | Qa.III | IRDVAQVIGRVNDVATSIASAVEQQGAATREISRNIQQAA------------- | *Azospirillum sp. TSA6c* | Bacteria; Proteobacteria; Alphaproteobacteria; Rhodospirillales; Rhodospirillaceae; Azospirillum | HAMP domain-containing protein | 6 |
| CAE27920.1 | 9.9E-05 | Qa.III | IREIGSTISKMSEIASTIASAVEEQGAATQEISRNVQQAAMGTQ--------- | *Rhodopseudomonas palustris CGA009* | Bacteria; Proteobacteria; Alphaproteobacteria; Rhizobiales; Bradyrhizobiaceae; Rhodopseudomonas | methyl-accepting chemotaxis receptor/ sensory transducer | 6 |
| KPX12410.1 | 9.9E-05 | Qa.III | --GISEAVANITDMTTQIATATEEQSSVAEEINRNIATIASLADQT------- | *Pseudomonas amygdali pv. dendropanacis* | Bacteria; Proteobacteria; Gammaproteobacteria; Pseudomonadales; Pseudomonadaceae; Pseudomonas; Pseudomonas amygdali | Aerotaxis receptor Aer | 6 |
| PWC83652.1 | 9.9E-05 | Qa.IV | IRAIVNTIRQVDESLSTVAAAVEEQDAATRDISERSQRAATDTVAVLQE---- | *Azospirillum sp. Sp 7* | Bacteria; Proteobacteria; Alphaproteobacteria; Rhodospirillales; Rhodospirillaceae; Azospirillum | chemotaxis protein partial | 6 |
| RDD63480.1 | 9.9E-05 | Qc.III | IERISDTIRQMDEISTTISSAVEEQGAATREITRNVREAATGTRDVNN----- | *Rhodospirillaceae bacterium* | Bacteria; Proteobacteria; Alphaproteobacteria; Rhodospirillales; Rhodospirillaceae | chemotaxis protein | 6 |
| WP_011323021.1 | 9.9E-05 | Qc.III.b | ---LEASIDQLGDASDRIDRTVEAQNDSLAEIGTEMENVSATMEEVA------ | *Natronomonas pharaonis* | Archaea; Euryarchaeota; Halobacteria; Halobacteriales; Haloarculaceae; Natronomonas | methyl-accepting chemotaxis protein | 6 |
| WP_027516791.1 | 9.9E-05 | Qc.III | IGEISGTIARLSEISSAIAAAVEEQGAATQEIARNVQQAAQGTQQVSANV--- | *Bradyrhizobium sp. WSM1417* | Bacteria; Proteobacteria; Alphaproteobacteria; Rhizobiales; Bradyrhizobiaceae; Bradyrhizobium | PAS domain S-box protein | 6 |
| WP_031481266.1 | 9.9E-05 | Qa.III.b | -------------LYNALSSMVTSLVELIGTAEEKTQEAKRQTEAANIALEEA | *Desulfovibrio frigidus* | Bacteria; Proteobacteria; Deltaproteobacteria; Desulfovibrionales; Desulfovibrionaceae; Desulfovibrio | methyl-accepting chemotaxis protein | 6 |
| WP_068502777.1 | 9.9E-05 | Qc.III | --GIARTIGEINDIAGSIAAAVEEQGAATQEIARNVSQAAQGTQEVT------ | *Magnetospirillum moscoviense* | Bacteria; Proteobacteria; Alphaproteobacteria; Rhodospirillales; Rhodospirillaceae; Magnetospirillum | chemotaxis protein | 6 |
| WP_090074276.1 | 9.9E-05 | Qa.III | ISKISETMAEVNEYTNSIAAAVEQQGSATMEISSNVQQAS------------- | *Cohaesibacter marisflavi* | Bacteria; Proteobacteria; Alphaproteobacteria; Rhizobiales; Cohaesibacteraceae | methyl-accepting chemotaxis protein | 6 |
| WP_109086756.1 | 9.9E-05 | Qa.IV | IQAIVRTVTAIDEISGSIAAAVEQQGAATQEIARSVADAA------------- | *Azospirillum sp. TSH100* | Bacteria; Proteobacteria; Alphaproteobacteria; Rhodospirillales; Rhodospirillaceae; Azospirillum | HAMP domain-containing protein | 6 |
| GAS75438.1 | 0.0001 | R.IV | -TQIAQAMTEMDTTTQQNAALVEESSAAASSLEEQALKLEEAISVFH------ | *Salmonella enterica* | Bacteria; Proteobacteria; Gammaproteobacteria; Enterobacterales; Enterobacteriaceae; Salmonella | methyl-accepting chemotaxis protein III | 6 |
| KPY04898.1 | 0.0001 | Qa.III | ---ISQALRNLNSLNTSIASATLQQTHVVDDINQNVTQAAQLSQ--------- | *Pseudomonas amygdali pv. mori* | Bacteria; Proteobacteria; Gammaproteobacteria; Pseudomonadales; Pseudomonadaceae; Pseudomonas; Pseudomonas amygdali | Methyl-accepting chemotaxis protein | 6 |
| KPY60756.1 | 0.0001 | Qa.III | ---ISQALRNLNSLNTSIASATLQQTHVVDDINQNVTQAAQLSQ--------- | *Pseudomonas amygdali pv. sesami* | Bacteria; Proteobacteria; Gammaproteobacteria; Pseudomonadales; Pseudomonadaceae; Pseudomonas; Pseudomonas amygdali | Histidine kinase HAMP region: chemotaxis sensory transducer | 6 |
| KZD25862.1 | 0.0001 | Qc.III | IKEIGDTIGRMSEIASTIASAVEQQGAATQEISRNVQQAAQGTMQVSSNI--- | *Tardiphaga robiniae* | Bacteria; Proteobacteria; Alphaproteobacteria; Rhizobiales; Bradyrhizobiaceae; Tardiphaga | chemotaxis protein | 6 |
| NP_766967.1 | 0.0001 | Qa.IV | -----ATIGQISSISTSIASAVEQQGAATQEIARSVQTVAQGTHAAATDI--- | *Bradyrhizobium diazoefficiens USDA 110* | Bacteria; Proteobacteria; Alphaproteobacteria; Rhizobiales; Bradyrhizobiaceae; Bradyrhizobium | methyl-accepting chemotaxis protein | 6 |
| OEE02788.1 | 0.0001 | Qa.IV | VQELTTDVRQASQVVGQLDTNTQSAGSILDSISAIAEQTN------------- | *Vibrio cyclitrophicus ZF270* | Bacteria; Proteobacteria; Gammaproteobacteria; Vibrionales; Vibrionaceae; Vibrio | chemotaxis protein partial | 6 |
| SEH57214.1 | 0.0001 | Qc.III | IKEIGDTIGRMSEIASTIASAVEQQGAATQEISRNVQQAAQGTMQVSSNI--- | *Tardiphaga sp. OK245* | Bacteria; Proteobacteria; Alphaproteobacteria; Rhizobiales; Bradyrhizobiaceae; Tardiphaga | Methyl-accepting chemotaxis protein | 6 |
| SFL81478.1 | 0.0001 | Qc.III | IKEIGDTIGRMSEIASTIASAVEQQGAATQEISRNVQQAAQGTMQVSSNI--- | *Bradyrhizobium sp. NFR13* | Bacteria; Proteobacteria; Alphaproteobacteria; Rhizobiales; Bradyrhizobiaceae; Bradyrhizobium | Methyl-accepting chemotaxis protein | 6 |
| SNT49886.1 | 0.0001 | Qc.III | IKEIGDTIGRMSEIASTIASAVEQQGAATQEISRNVQQAAQGTMQVSSNI--- | *Tardiphaga sp. OK246* | Bacteria; Proteobacteria; Alphaproteobacteria; Rhizobiales; Bradyrhizobiaceae; Tardiphaga | Methyl-accepting chemotaxis protein | 6 |
| WP_002715667.1 | 0.0001 | Qc.III | INEISNSIERLSEISAAIASAIEEQGATTQEVARNVQHAAQGAQQVSSNV--- | *Afipia felis* | Bacteria; Proteobacteria; Alphaproteobacteria; Rhizobiales; Bradyrhizobiaceae; Afipia | PAS domain S-box protein | 6 |
| WP_003058313.1 | 0.0001 | Qa.III | MKEIMQSVGQVTGLLSEISGALQQQSEGI----AHVNTAVAHMDSTNQE---- | *Comamonas testosteroni* | Bacteria; Proteobacteria; Betaproteobacteria; Burkholderiales; Comamonadaceae; Comamonas | methyl-accepting chemotaxis protein | 6 |
| WP_003066090.1 | 0.0001 | Qa.III | MKEIMQSVGQVTGLLSEISGALQQQSEGI----AHVNTAVAHMDSTNQE---- | *Comamonas testosteroni* | Bacteria; Proteobacteria; Betaproteobacteria; Burkholderiales; Comamonadaceae; Comamonas | methyl-accepting chemotaxis protein | 6 |
| WP_003073891.1 | 0.0001 | Qa.III | MKEIMQSVGQVTGLLSEISGALQQQSEGI----AHVNTAVAHMDSTNQE---- | *Comamonas testosteroni* | Bacteria; Proteobacteria; Betaproteobacteria; Burkholderiales; Comamonadaceae; Comamonas | methyl-accepting chemotaxis protein | 6 |
| WP_011663369.1 | 0.0001 | Qc.III | --EISDTIGRMSEISSLIAAAVEQQGAATQEISRNVEHAAHGTSRV------- | *Rhodopseudomonas palustris* | Bacteria; Proteobacteria; Alphaproteobacteria; Rhizobiales; Bradyrhizobiaceae; Rhodopseudomonas | methyl-accepting chemotaxis protein | 6 |
| WP_012495956.1 | 0.0001 | Qa.III | IREIGSTISKMSEIASTIASAVEEQGAATQEISRNVQQAAM------------ | *Rhodopseudomonas palustris* | Bacteria; Proteobacteria; Alphaproteobacteria; Rhizobiales; Bradyrhizobiaceae; Rhodopseudomonas | methyl-accepting chemotaxis protein | 6 |
| WP_012526682.1 | 0.0001 | Qa.III | IHSLAGAIAELAETSRHMSATATQQAQGLGQINQAMRDVKQTTDQTVQAIRQT | *Anaeromyxobacter sp. K* | Bacteria; Proteobacteria; Deltaproteobacteria; Myxococcales; Cystobacterineae; Anaeromyxobacteraceae; Anaeromyxobacter | methyl-accepting chemotaxis protein | 6 |
| WP_018044919.1 | 0.0001 | Qc.III | --SIQDTIMTLNEVASAISAAVEEQSAVTREMSASMQTASNG----------- | *Methylobacterium sp. 88A* | Bacteria; Proteobacteria; Alphaproteobacteria; Rhizobiales; Methylobacteriaceae; Methylobacterium | methyl-accepting chemotaxis protein | 6 |
| WP_025034937.1 | 0.0001 | Qc.III | IGEISGTIARLSEIASAIAAAVQQQGAATQEIARNVQQAAQGTQQVSSNVG-- | *Bradyrhizobium sp. DOA9* | Bacteria; Proteobacteria; Alphaproteobacteria; Rhizobiales; Bradyrhizobiaceae; Bradyrhizobium | PAS domain S-box protein | 6 |

| | | | | | | | | |
|---|---|---|---|---|---|---|---|---|
| | **WP_028178347.1** | 0.0001 | Qc.III | IGEISGTIARLSEIASAIAAAVQQQGAATQEIARNVQEAARGTQQVSSNVG-- | *Bradyrhizobium* | Bacteria; Proteobacteria; Alphaproteobacteria; Rhizobiales; Bradyrhizobiaceae | MULTISPECIES: PAS domain S-box protein | 6 |
|  | **WP_029010153.1** | 0.0001 | Qa.III | IQEITRVIGRVNEVSTSIASAVEEQGAATREISRSVQQAAQGTREV------- | *Azospirillum halopraeferens* | Bacteria; Proteobacteria; Alphaproteobacteria; Rhodospirillales; Rhodospirillaceae; Azospirillum | HAMP domain-containing protein | 6 |
|  | **WP_034357654.1** | 0.0001 | Qa.III | MKEIMQSVGQVTGLLSEISGALQQQSEGI----AHVNTAVAHMDSTNQE---- | *Comamonas* | Bacteria; Proteobacteria; Betaproteobacteria; Burkholderiales; Comamonadaceae | MULTISPECIES: methyl-accepting chemotaxis protein | 6 |
|  | **WP_034381733.1** | 0.0001 | Qa.III | MKEIMQSVGQVTGLLSEISGALQQQSEGI----AHVNTAVAHMDSTNQE---- | *Comamonas* | Bacteria; Proteobacteria; Betaproteobacteria; Burkholderiales; Comamonadaceae | MULTISPECIES: methyl-accepting chemotaxis protein | 6 |
|  | **WP_034389287.1** | 0.0001 | Qa.III | MKEIMQSVGQVTGLLSEISGALQQQSEGI----AHVNTAVAHMDSTNQE---- | *Comamonas testosteroni* | Bacteria; Proteobacteria; Betaproteobacteria; Burkholderiales; Comamonadaceae; Comamonas | methyl-accepting chemotaxis protein | 6 |
| | **WP_036044698.1** | 0.0001 | Qc.III | IGEISGTIARLSEIASAIAAAVQQQGAATQEIARNVQQAAQGTQQVSSNVG-- | *Bradyrhizobium yuanmingense* | Bacteria; Proteobacteria; Alphaproteobacteria; Rhizobiales; Bradyrhizobiaceae; Bradyrhizobium | PAS domain S-box protein | 6 |
|  | **WP_039049647.1** | 0.0001 | Qa.III | MKEIMQSVGQVTGLLSEISGALQQQSEGI----AHVNTAVAHMDSTNQE---- | *Comamonas testosteroni* | Bacteria; Proteobacteria; Betaproteobacteria; Burkholderiales; Comamonadaceae; Comamonas | methyl-accepting chemotaxis protein | 6 |
|  | **WP_041743913.1** | 0.0001 | Qa.III | MKEIMQSVGQVTGLLSEISGALQQQSEGI----AHVNTAVAHMDSTNQE---- | *Comamonas testosteroni* | Bacteria; Proteobacteria; Betaproteobacteria; Burkholderiales; Comamonadaceae; Comamonas | methyl-accepting chemotaxis protein | 6 |
|  | **WP_043003434.1** | 0.0001 | Qa.III | MKEIMQSVGQVTGLLSEISGALQQQSEGI----AHVNTAVAHMDSTNQE---- | *Comamonas testosteroni* | Bacteria; Proteobacteria; Betaproteobacteria; Burkholderiales; Comamonadaceae; Comamonas | methyl-accepting chemotaxis protein | 6 |
|  | **WP_056147997.1** | 0.0001 | Qc.III | --SIQDTIMTLNEVASAISAAVEEQSAVTREMSASMQTASNG----------- | *Methylobacterium sp. Leaf85* | Bacteria; Proteobacteria; Alphaproteobacteria; Rhizobiales; Methylobacteriaceae; Methylobacterium | methyl-accepting chemotaxis protein | 6 |
|  | **WP_056161870.1** | 0.0001 | Qc.III | --SIQDTIMTLNEVASAISAAVEEQSAVTREMSASMQTASNG----------- | *Methylobacterium sp. Leaf106* | Bacteria; Proteobacteria; Alphaproteobacteria; Rhizobiales; Methylobacteriaceae; Methylobacterium | methyl-accepting chemotaxis protein | 6 |
|  | **WP_056257335.1** | 0.0001 | Qc.III | --SIQDTIMTLNEVASAISAAVEEQSAVTREMSASMQTASNG----------- | *Methylobacterium sp. Leaf93* | Bacteria; Proteobacteria; Alphaproteobacteria; Rhizobiales; Methylobacteriaceae; Methylobacterium | methyl-accepting chemotaxis protein | 6 |
|  | **WP_057849514.1** | 0.0001 | Qc.III | IQEISGTIGKLSEISSAIAAAVEEQGAATQEISRNVQQAA------------- | *Bradyrhizobium valentinum* | Bacteria; Proteobacteria; Alphaproteobacteria; Rhizobiales; Bradyrhizobiaceae; Bradyrhizobium | methyl-accepting chemotaxis protein | 6 |
|  | **WP_057901223.1** | 0.0001 | Qc.III | IQEISGTIGKLSEISSAIAAAVEEQGAATQEISRNVQQAA------------- | *Bradyrhizobium valentinum* | Bacteria; Proteobacteria; Alphaproteobacteria; Rhizobiales; Bradyrhizobiaceae; Bradyrhizobium | methyl-accepting chemotaxis protein | 6 |
| | **WP_068313591.1** | 0.0001 | Qa.III | IRSIRGTIDQINELAGNIDRAVSDQRFATDGISDSLESAHSSAQRVAADIE-- | *Pseudovibrio hongkongensis* | Bacteria; Proteobacteria; Alphaproteobacteria; Rhodobacterales; Rhodobacteraceae; Pseudovibrio | methyl-accepting chemotaxis protein | 6 |
|  | **WP_081459207.1** | 0.0001 | SNAP.c | -AQIDQDFQMMNTMAAFFGSAKDIQNEQYFNILQRVDDAND------------ | *Faecalitalea cylindroides* | Bacteria; Firmicutes; Erysipelotrichia; Erysipelotrichales; Erysipelotrichaceae; Faecalitalea | EAL domain-containing protein partial | 6 |
|  | **WP_083860149.1** | 0.0001 | Qa.IV | -----ATIGQISSISTSIASAVEQQGAATQEIARSVQTVAQGTHAAATDI--- | *Bradyrhizobium diazoefficiens* | Bacteria; Proteobacteria; Alphaproteobacteria; Rhizobiales; Bradyrhizobiaceae; Bradyrhizobium | methyl-accepting chemotaxis protein | 6 |
|  | **WP_084326277.1** | 0.0001 | SNAP.c | --QVADASQSLASGATEQAATVEELTAAIQSVTEKAEKNTVSVRKAT------ | *[Clostridium] aerotolerans* | Bacteria; Firmicutes; Clostridia; Clostridiales; Lachnospiraceae | methyl-accepting chemotaxis protein | 6 |
|  | **WP_085556335.1** | 0.0001 | Qc.III | -QSVASVIDRLSGIATAISAAVEEQNAATAEISRSVQQAAAGT---------- | *Azospirillum lipoferum* | Bacteria; Proteobacteria; Alphaproteobacteria; Rhodospirillales; Rhodospirillaceae; Azospirillum | HAMP domain-containing protein | 6 |
|  | **WP_087084941.1** | 0.0001 | Qa.III | MKEIMQSVGQVTGLLSEISGALQQQSEGI----AHVNTAVAHMDSTNQE---- | *Comamonas testosteroni* | Bacteria; Proteobacteria; Betaproteobacteria; Burkholderiales; Comamonadaceae; Comamonas | methyl-accepting chemotaxis protein | 6 |
| | **WP_089266236.1** | 0.0001 | Qc.III | IKEIGDTIGRMSEIASTIASAVEQQGAATQEISRNVQQAAQGTMQVSSNI--- | *Tardiphaga sp. OK246* | Bacteria; Proteobacteria; Alphaproteobacteria; Rhizobiales; Bradyrhizobiaceae; Tardiphaga | methyl-accepting chemotaxis protein | 6 |
|  | **WP_090255030.1** | 0.0001 | Qc.III | LEDITRSIGTINDMNTQIASAAEEQSAVAEDVNRSLTRISSGVEQ-------- | *Ectothiorhodospira marina* | Bacteria; Proteobacteria; Gammaproteobacteria; Chromatiales; Ectothiorhodospiraceae; Ectothiorhodospira | methyl-accepting chemotaxis protein | 6 |
|  | **WP_092143466.1** | 0.0001 | Qa.III | IQGITATIEEVSAIATTIGSAIEEQGAATAEIARNVTQTAQAT---------- | *Bradyrhizobium sp. NFR13* | Bacteria; Proteobacteria; Alphaproteobacteria; Rhizobiales; Bradyrhizobiaceae; Bradyrhizobium | HAMP domain-containing protein | 6 |
| | **WP_092146202.1** | 0.0001 | Qc.III | IKEIGDTIGRMSEIASTIASAVEQQGAATQEISRNVQQAAQGTMQVSSNI--- | *Bradyrhizobiaceae* | Bacteria; Proteobacteria; Alphaproteobacteria; Rhizobiales | methyl-accepting chemotaxis protein | 6 |
| | **WP_093757369.1** | 0.0001 | Qc.III | IKEIGDTIGRMSEIASTIASAVEQQGAATQEISRNVQQAAQGTMQVSSNI--- | *Tardiphaga sp. OK245* | Bacteria; Proteobacteria; Alphaproteobacteria; Rhizobiales; Bradyrhizobiaceae; Tardiphaga | methyl-accepting chemotaxis protein | 6 |
|  | **WP_108518545.1** | 0.0001 | Qc.III | IQEISGTIGKLSEISSAIAAAVEEQGAATQEISRNVQQAA------------- | *Bradyrhizobium algeriense* | Bacteria; Proteobacteria; Alphaproteobacteria; Rhizobiales; Bradyrhizobiaceae; Bradyrhizobium | methyl-accepting chemotaxis protein | 6 |
|  | **WP_110686386.1** | 0.0001 | R.IV | IQQINLAVGEMDSVTQQNAGLVEESTTVTSHLQTQAESLA------------- | *Salinicola sp. CPA62* | Bacteria; Proteobacteria; Gammaproteobacteria; Oceanospirillales; Halomonadaceae; Salinicola | HAMP domain-containing protein | 6 |
|  | **WP_110686387.1** | 0.0001 | R.IV | IQQINLAVGEMDSVTQQNAGLVEESTTVTSHLQTQAESLA------------- | *Salinicola sp. CPA62* | Bacteria; Proteobacteria; Gammaproteobacteria; Oceanospirillales; Halomonadaceae; Salinicola | HAMP domain-containing protein | 6 |
| **▼ 7** | | | | | | | | |
| | **PPD46731.1** | 1.6E-08 | Qc.III | --ILMKQASEMETLGIQLKQKYDEQNQRLESAHPDMVATNAQIRRVNNKLAT- | *Methylobacter sp.* | Bacteria; Proteobacteria; Gammaproteobacteria; Methylococcales; Methylococcaceae; Methylobacter | capsular biosynthesis protein | 7 |
| | **WP_013818087.1** | 2.2E-08 | Qc.III | ---LLKQAAEMETLGIQLRQKYEEQNQRLEAAHPDMKATNAQIRRVNRKLA-- | *Methylomonas methanica* | Bacteria; Proteobacteria; Gammaproteobacteria; Methylococcales; Methylococcaceae; Methylomonas | capsular polysaccharide biosynthesis protein | 7 |
| | **WP_036243086.1** | 6.3E-08 | Qc.III | ---LLKQAAEMETLGIQLKQKYDEQSQRLESAHPDMQATNAQIKRVNSKLSKL | *Methylobacter luteus* | Bacteria; Proteobacteria; Gammaproteobacteria; Methylococcales; Methylococcaceae; Methylobacter | capsular biosynthesis protein | 7 |
| | **WP_019867119.1** | 1.1E-07 | Qc.III | --ILMKQASEMETLGIQLKQKYDEQSQRLESAHPDMVATNAQIRRVNNKLAT- | *Methylovulum miyakonense* | Bacteria; Proteobacteria; Gammaproteobacteria; Methylococcales; Methylococcaceae; Methylovulum | capsular polysaccharide biosynthesis protein | 7 |
| | **WP_024296755.1** | 3E-05 | Qc.III | ---LLKQAAEMETLGIQLKQKYDEQSQRLESAHPDMIATNSQIKHVNGKLA-- | *Methylosarcina lacus* | Bacteria; Proteobacteria; Gammaproteobacteria; Methylococcales; Methylococcaceae; Methylosarcina | capsular biosynthesis protein | 7 |
|  | **WP_077731768.1** | 6.6E-05 | SNAP.b | ----LKQAAEMETLGIQLRQKYEEQTQRLTDAHPDMMATKAQIGRVERKLADL | *Methylocaldum sp. 14B* | Bacteria; Proteobacteria; Gammaproteobacteria; Methylococcales; Methylococcaceae; Methylocaldum | capsular biosynthesis protein | 7 |
|  | **WP_086133669.1** | 6.6E-05 | SNAP.b | ----LKQAAEMETLGIQLRQKYEEQTQRLTDAHPDMMATKAQIGRVERKLADL | *Methylocaldum sp. SAD2* | Bacteria; Proteobacteria; Gammaproteobacteria; Methylococcales; Methylococcaceae; Methylocaldum | capsular biosynthesis protein | 7 |
| **▼ 8** | | | | | | | | |
|  | **SEJ95423.1** | 2.2E-08 | SNAP.c | -------LGQIHGLVQALKDGQDQQNRRMDSFDTRFDALDGRLRSVEQR---- | *Variovorax sp. OK202* | Bacteria; Proteobacteria; Betaproteobacteria; Burkholderiales; Comamonadaceae; Variovorax | hypothetical protein SAMN05518853_105110 | 8 |
|  | **PBI84102.1** | 4.1E-08 | SNAP.c | -------LGQIHGLVQALKEGQDQQNRRMDSFDARFDALDGRLRAVEQRA--- | *Variovorax boronicumulans* | Bacteria; Proteobacteria; Betaproteobacteria; Burkholderiales; Comamonadaceae; Variovorax | hypothetical protein BKP43_55320 | 8 |
|  | **SCX70901.1** | 5E-08 | SNAP.c | -------LGQIHGLVQALKDGQDRQNRRMDGFDTRFDALDGRLRSVEQRA--- | *Variovorax sp. EL159* | Bacteria; Proteobacteria; Betaproteobacteria; Burkholderiales; Comamonadaceae; Variovorax | hypothetical protein SAMN03159363_3693 | 8 |
|  | **OJZ14246.1** | 8.8E-08 | SNAP.c | -------LGQIHGLVQALKDGQDRQNRRMDGFDTRFDALDGRLRTVEQRA--- | *Variovorax sp. 67-131* | Bacteria; Proteobacteria; Betaproteobacteria; Burkholderiales; Comamonadaceae; Variovorax | hypothetical protein BGP22_06020 | 8 |
|  | **SFB93767.1** | 2.3E-06 | SNAP.c | -------LGQIHGLVQALKDGQDRQTRRMDGFDARFDALDGRLRTVEQRA--- | *Variovorax sp. NFACC26* | Bacteria; Proteobacteria; Betaproteobacteria; Burkholderiales; Comamonadaceae; Variovorax | hypothetical protein SAMN03159379_01087 | 8 |
|  | **WP_093343527.1** | 2.9E-06 | SNAP.c | -------LGQIHGLVSALKEGQDRQSRRMDAVDARFDSLDSRLRAVEQR---- | *Variovorax sp. PDC80* | Bacteria; Proteobacteria; Betaproteobacteria; Burkholderiales; Comamonadaceae; Variovorax | hypothetical protein | 8 |
| **▼ 9** | | | | | | | | |
|  | **KYP87036.1** | 1.3E-07 | Qb.I | ---IAQMVQDATQQAKQALDQLNQTKAAIQQAKDQFDNYKNLVTGNNKL---- | *bacteria symbiont BFo1 of Frankliniella occidentalis* | Bacteria | conjugal transfer protein | 9 |
|  | **ELO16036.1** | 6E-06 | Qb.I | ---IAQLVSNAQQQAQQALDQLNETKAAIQQAKSQYDHYKNLVTGNNKL---- | *Salmonella enterica subsp. enterica serovar Enteritidis str. 648900 1-16* | Bacteria; Proteobacteria; Gammaproteobacteria; Enterobacterales; Enterobacteriaceae; Salmonella | conjugative transfer protein partial | 9 |
|  | **WP_023180318.1** | 2.8E-05 | Qb.I | ---IAQLVSNAQQQAQQALDQLNETKAAIQQAKSQYDHYKNLVTGNNKL---- | *Salmonella enterica* | Bacteria; Proteobacteria; Gammaproteobacteria; Enterobacterales; Enterobacteriaceae; Salmonella | P-type DNA transfer protein VirB5 | 9 |
|  | **WP_032679801.1** | 2.8E-05 | Qb.I | ---IAQLVSNAQQQAQQALDQLNETKAAIQQAKSQYDHYKNLVTGNNKL---- | *Enterobacter sp. BIDMC 26* | Bacteria; Proteobacteria; Gammaproteobacteria; Enterobacterales; Enterobacteriaceae; Enterobacter; Enterobacter cloacae complex | P-type DNA transfer protein VirB5 | 9 |
|  | **WP_047625766.1** | 2.8E-05 | Qb.I | ---IAQLVSNAQQQAQQALDQLNETKAAIQQAKSQYDHYKNLVTGNNKL---- | *Enterobacter sp. ZOR0014* | Bacteria; Proteobacteria; Gammaproteobacteria; Enterobacterales; Enterobacteriaceae; Enterobacter | P-type DNA transfer protein VirB5 | 9 |
|  | **WP_096879145.1** | 3.1E-05 | Qb.I | ---IAQLVSNAQQQAQQALDQLNETKAAIQQAKSQYDHYKNLVTGNNKL---- | *Citrobacter freundii* | Bacteria; Proteobacteria; Gammaproteobacteria; Enterobacterales; Enterobacteriaceae; Citrobacter; Citrobacter freundii complex | P-type DNA transfer protein VirB5 | 9 |
| **▼ 10** | | | | | | | | |
|  | **AMG33613.1** | 1.9E-07 | Qa.III | IKNLNTDISTVNSSMSDISVLVNNQKTEIDNLKTQIDTANTE----------- | *Staphylococcus saprophyticus* | Bacteria; Firmicutes; Bacilli; Bacillales; Staphylococcaceae; Staphylococcus | hypothetical protein AL494_07565 | 10 |
|  | **WP_107553006.1** | 6.9E-06 | Qa.III | IKNLNTDISIINSSMTDISVLVNNQKTEIDNLKTQIDTANTE----------- | *Staphylococcus succinus* | Bacteria; Firmicutes; Bacilli; Bacillales; Staphylococcaceae; Staphylococcus | hypothetical protein | 10 |
| **▼ 11** | | | | | | | | |
|  | **PXY54931.1** | 2.3E-07 | SNAP.b | -----KLVDDAIKTVSAARSHLGANQNRLDHTINNLDNAQENLTAAESRIR-- | *Virgibacillus profundi* | Bacteria; Firmicutes; Bacilli; Bacillales; Bacillaceae; Virgibacillus | hypothetical protein CIT14_03295 | 11 |
| | **WP_008374909.1** | 2E-06 | Qb.II | IDASISAIDAALSTINSARADLGAKQNRFASTISNLNNIVSNATASRSTIQ-- | *Pseudomonas sp. M47T1* | Bacteria; Proteobacteria; Gammaproteobacteria; Pseudomonadales; Pseudomonadaceae; Pseudomonas | flagellin | 11 |
| | **WP_027981681.1** | 3.5E-06 | Qb.II | -DASISAIDAALATINSARADLGAKQNRFSSTISNLNNIVSNATASRSTIQ-- | *gamma proteobacterium L18* | Bacteria; Proteobacteria; Gammaproteobacteria | flagellin | 11 |
|  | **WP_093194172.1** | 6.3E-06 | Qb.II | ---SQSGADDAISTINTAIETVSAQRANLGATQNRLEHTINNLGTSS------ | *Salimicrobium halophilum* | Bacteria; Firmicutes; Bacilli; Bacillales; Bacillaceae; Salimicrobium | flagellin | 11 |
|  | **PUU87115.1** | 1.4E-05 | SNAP.b | -------AQEALDTLDSAISEVSDQRSQLGAVQNRLDHTINNLNTAEENLT-- | *Halanaerobium sp.* | Bacteria; Firmicutes; Clostridia; Halanaerobiales; Halanaerobiaceae; Halanaerobium | flagellin | 11 |
|  | **OHD08308.1** | 1.8E-05 | Qb.III | ---ALATIDDAITIVAEERANLGAQQNRLTSAVDNLTSRVTNLDEAKSR---- | *Sphingopyxis sp. RIFCSPHIGHO2_12_FULL_65_19* | Bacteria; Proteobacteria; Alphaproteobacteria; Sphingomonadales; Sphingomonadaceae; Sphingopyxis | flagellin | 11 |
|  | **WP_058803468.1** | 1.8E-05 | Qb.III | ---ALATIDDAITIVAEERANLGAQQNRLTSAVDNLTSRVTNLDEAKSR---- | *Sphingopyxis sp. H115* | Bacteria; Proteobacteria; Alphaproteobacteria; Sphingomonadales; Sphingomonadaceae; Sphingopyxis | flagellin | 11 |
|  | **WP_056373231.1** | 2.3E-05 | Qb.III | ---ALAVLDTAIDAVASERANLGAQQNRLTSAVDNLTSRVTNLDESKSR---- | *Sphingopyxis* | Bacteria; Proteobacteria; Alphaproteobacteria; Sphingomonadales; Sphingomonadaceae | MULTISPECIES: flagellin | 11 |
| | **WP_089802119.1** | 3.7E-05 | SNAP.b | ---AIKTINDAIETVSAERSKLGANQNRLEHTIANLDNSSENLSAAESRIR-- | *Halolactibacillus alkaliphilus* | Bacteria; Firmicutes; Bacilli; Bacillales; Bacillaceae; Halolactibacillus | flagellin | 11 |
|  | **WP_038483738.1** | 4.1E-05 | Qb.III | ------------DTVSSERSKLGANQNRLEHTINNLDNASENLSAAESRIRD- | *Bacillus* | Bacteria; Firmicutes; Bacilli; Bacillales; Bacillaceae | MULTISPECIES: flagellin | 11 |
|  | **WP_058300365.1** | 4.4E-05 | SNAP.b | ------SVDDAIKAVSAARSKLGANQNRLEHTIKNLDNASENLTAAESRV--- | *Gorillibacterium timonense* | Bacteria; Firmicutes; Bacilli; Bacillales; Paenibacillaceae; Gorillibacterium | flagellin | 11 |

| | | | | | | | | |
|---|---|---|---|---|---|---|---|---|
|  | **WP_019440874.1** | 4.8E-05 | Qb.II | --GAIADLDSAIAQVTDIRADLGAKQNRFSSTIRNVTNIAENVSASK------ | *Moritella marina* | Bacteria; Proteobacteria; Gammaproteobacteria; Alteromonadales; Moritellaceae; Moritella | flagellin | 11 |
|  | **WP_053430275.1** | 5.2E-05 | SNAP.b | ------TINDAINTLSTSRAQLGAWQNRLEHTIKNLDNASENLQAAESRIR-- | *Bacillus okuhidensis* | Bacteria; Firmicutes; Bacilli; Bacillales; Bacillaceae; Bacillus | flagellin | 11 |
|  | **WP_078431752.1** | 5.3E-05 | Qb.II | ---TNAAASSSISQIDAAIDKVSKQRSVLGAVQNRLEHTINNLGASQ------ | *Bacillus halosaccharovorans* | Bacteria; Firmicutes; Bacilli; Bacillales; Bacillaceae; Bacillus | flagellin | 11 |
|  | **ATC98046.1** | 5.5E-05 | Qb.II | ---AIAALDDALSTVNAVRADLGAKQNRFSSTIRNLKSVAENVSASK------ | *Pseudoalteromonas spongiae UST010723-006* | Bacteria; Proteobacteria; Gammaproteobacteria; Alteromonadales; Pseudoalteromonadaceae; Pseudoalteromonas | flagellin | 11 |
| | **WP_093054350.1** | 5.6E-05 | Qb.III | ---AITTINEAIETVSAERSMLGANQNRLEHTISNLNNASENLSAAESRIRD- | *Salipaludibacillus aurantiacus* | Bacteria; Firmicutes; Bacilli; Bacillales; Bacillaceae | flagellin | 11 |
| | **WP_091732383.1** | 6.1E-05 | SNAP.b | --ATLGKVDEAIKTVSSVRSKLGANQNRLEYTIKNLDNAAENLQAAESRIR-- | *Proteiniborus ethanoligenes* | Bacteria; Firmicutes; Clostridia; Clostridiales; Proteiniborus | flagellin | 11 |
|  | **CDA54472.1** | 6.6E-05 | Qb.III | --ESIATVQNAIEQAALWRDTFGGQQEQLEHALRGTDNTSENTQRAE------ | *Roseburia intestinalis CAG:13* | Bacteria; Firmicutes; Clostridia; Clostridiales; Lachnospiraceae; Roseburia; environmental samples | flagellin | 11 |
|  | **WP_078579115.1** | 6.6E-05 | SNAP.b | --------DETLATVDAALQQVSDQRSALGSVQNRLEHTIRNLDNSSENL--- | *Salipaludibacillus agaradhaerens* | Bacteria; Firmicutes; Bacilli; Bacillales; Bacillaceae | flagellin | 11 |
|  | **WP_017220306.1** | 7.1E-05 | Qb.II | ------SLDTAIASVTDIRGDLGAKQNRFSSTIRNVTNIAENVSASK------ | *Moritella dasanensis* | Bacteria; Proteobacteria; Gammaproteobacteria; Alteromonadales; Moritellaceae; Moritella | flagellin | 11 |
|  | **WP_054982976.1** | 7.2E-05 | Qb.II | -QAAITAIDDALSTVNEVRADLGAKQNRFSSTIRNLENVAENVSASK------ | *Pseudoalteromonas sp. P1-30* | Bacteria; Proteobacteria; Gammaproteobacteria; Alteromonadales; Pseudoalteromonadaceae; Pseudoalteromonas | flagellin | 11 |
|  | **WP_079576914.1** | 9.6E-05 | Qb.III | ---LKGVQSNTIGLGLDTLKNLGANSLDITVARDWLDDIDNSLT--------- | *Arcobacter* | Bacteria; Proteobacteria; Epsilonproteobacteria; Campylobacterales; Campylobacteraceae | MULTISPECIES: flagellin | 11 |
|  | **WP_099344576.1** | 9.6E-05 | Qb.III | ---LKGVQSNTIGLGLDTLKNLGANSLDITVARDWLDDIDNSLT--------- | *Arcobacter marinus* | Bacteria; Proteobacteria; Epsilonproteobacteria; Campylobacterales; Campylobacteraceae; Arcobacter | flagellin | 11 |
| **▼ 12** | | | | | | | | |
|  | **WP_066377374.1** | 2.4E-07 | Qc.III.b | -----------KHGVDDVRFELKDQVRRFDALTERVTRSEERINQAHKRIDKL | *Bacillus sp. FJAT-27264* | Bacteria; Firmicutes; Bacilli; Bacillales; Bacillaceae; Bacillus | hypothetical protein | 12 |
|  | **WP_039291876.1** | 1.6E-05 | Qc.III.b | -----------KHGVDDVRFELKDQVRRFDALTERVTRVEESTKQAHKRIDKI | *Paenibacillus sp. IHB B 3415* | Bacteria; Firmicutes; Bacilli; Bacillales; Paenibacillaceae; Paenibacillus | hypothetical protein | 12 |
|  | **WP_025850202.1** | 3.3E-05 | Qc.III | ---------YIKRGVDDIRIEQRTQGKRIDDLAERVTRVEESSKQAHKRIDRL | *Paenibacillus ehimensis* | Bacteria; Firmicutes; Bacilli; Bacillales; Paenibacillaceae; Paenibacillus | hypothetical protein | 12 |
|  | **WP_025852983.1** | 3.4E-05 | Qc.III | ----------IKRGVDDIRIEQRTQGKRIDDLAERVTRVEESSKQAHKRIDRL | *Paenibacillus ehimensis* | Bacteria; Firmicutes; Bacilli; Bacillales; Paenibacillaceae; Paenibacillus | hypothetical protein | 12 |
|  | **WP_087916303.1** | 7.6E-05 | Qc.III | ----------IKRGVDDIRADVRQQGQRTDALAERVTRVEESAKQAHKRIDKI | *Paenibacillus donghaensis* | Bacteria; Firmicutes; Bacilli; Bacillales; Paenibacillaceae; Paenibacillus | hypothetical protein | 12 |
| **▼ 13** | | | | | | | | |
|  | **WP_045174574.1** | 2.7E-07 | SNAP.b | -----------NEILQAIVANLEKINERLDTIEKRLDKIEQRLDMVEQRLD-- | *Caldicellulosiruptor danielii* | Bacteria; Firmicutes; Clostridia; Thermoanaerobacterales; Thermoanaerobacterales Family III. Incertae Sedis; Caldicellulosiruptor | hypothetical protein | 13 |
|  | **WP_039765456.1** | 5.5E-07 | SNAP.b | -----------NDILQAIVANLEKINERLDTIEKRLDKIEQRLDMVEQRLD-- | *Caldicellulosiruptor sp. F32* | Bacteria; Firmicutes; Clostridia; Thermoanaerobacterales; Thermoanaerobacterales Family III. Incertae Sedis; Caldicellulosiruptor | hypothetical protein | 13 |
| | **WP_110939540.1** | 2.5E-06 | Qc.III | IDTLGKKIDRLEEKVDRIEERLDRIEERLDRVEERLDRVEERLDRVEERLDRV | *Geosporobacter subterraneus* | Bacteria; Firmicutes; Clostridia; Clostridiales; Clostridiaceae; Geosporobacter | DUF16 domain-containing protein | 13 |
| | **PDO09732.1** | 2.6E-06 | Qc.III | --QLIGIVKATNEKVSALTERVDRLEQKVDQLEARVDRLEQKIDQLEARVDRI | *Paenibacillaceae bacterium JTherm* | Bacteria; Firmicutes; Bacilli; Bacillales; Paenibacillaceae | hypothetical protein BLM47_10955 | 13 |
|  | **WP_079350681.1** | 3.6E-06 | SNAP.b | -----------DQVLSQILNQLQQMDKRFDNIDGRLEKIEDRLDTIEEQTKNI | *Paenibacillus sp. VT-16-81* | Bacteria; Firmicutes; Bacilli; Bacillales; Paenibacillaceae; Paenibacillus | hypothetical protein | 13 |
| | **WP_049975120.1** | 1.2E-05 | Qc.III | LDKLENKVDTLGNKVDTLGNKVDTLGNKVDTLENKVDTLENKVDTLENKVDT- | *Blautia schinkii* | Bacteria; Firmicutes; Clostridia; Clostridiales; Lachnospiraceae; Blautia | hypothetical protein | 13 |
|  | **WP_090903958.1** | 2.1E-05 | SNAP.b | -----------DQVLSQILNQLQQMDKRFDNIDGRLEKIEDRLDTVDGRLDTM | *Paenibacillus sp. OK076* | Bacteria; Firmicutes; Bacilli; Bacillales; Paenibacillaceae; Paenibacillus | hypothetical protein | 13 |
|  | **WP_003329566.1** | 4.2E-05 | SNAP.b | ------------KRLDAIEQRLDANDQRFDAIDKRLDGIDERLDGIDERLDGI | *Bacillus azotoformans* | Bacteria; Firmicutes; Bacilli; Bacillales; Bacillaceae; Bacillus | hypothetical protein | 13 |
|  | **WP_099083991.1** | 4.5E-05 | Qc.III | --DLIRIVGHTNSLVEELNTKVEKLNTKVEELDAKVDRLEARVERL------- | *Desulfotomaculum profundi* | Bacteria; Firmicutes; Clostridia; Clostridiales; Peptococcaceae; Desulfotomaculum | DUF4686 domain-containing protein | 13 |
|  | **AKB76684.1** | 5.7E-05 | Qc.III | ----------IREMNSNLCQKVDLLGGKIDVLGGKMDSLGGKMDSLGGKIDT- | *Methanosarcina horonobensis HB-1 = JCM 15518* | Archaea; Euryarchaeota; Methanomicrobia; Methanosarcinales; Methanosarcinaceae; Methanosarcina | Acylphosphatase | 13 |
|  | **WP_091261003.1** | 6.2E-05 | SNAP.b | ----------MEAVLNRILGELQHVNQKLENIDNRLDKVEQRMGKLEQRMD-- | *Aneurinibacillus thermoaerophilus* | Bacteria; Firmicutes; Bacilli; Bacillales; Paenibacillaceae; Aneurinibacillus group; Aneurinibacillus | hypothetical protein | 13 |
|  | **WP_048136754.1** | 7.2E-05 | Qc.III | ----------IREMNSNLCQKVDLLGGKIDVLGGKMDSLGGKMDSLGGKIDT- | *Methanosarcina horonobensis* | Archaea; Euryarchaeota; Methanomicrobia; Methanosarcinales; Methanosarcinaceae; Methanosarcina | hypothetical protein | 13 |
| **▼ 14** | | | | | | | | |
|  | **WP_105054878.1** | 2.8E-07 | Qc.III | -------VGALKSQNASLGMQVEEQNKQIDKLTNQVDLQITMLSRAK------ | *Aliivibrio sifiae* | Bacteria; Proteobacteria; Gammaproteobacteria; Vibrionales; Vibrionaceae; Aliivibrio | hypothetical protein | 14 |
|  | **WP_060991154.1** | 2.9E-07 | Qc.III | -------VGALKSQNASLGMQVEEQNKQIDKLTNQVDLQITMLSRAK------ | *Aliivibrio* | Bacteria; Proteobacteria; Gammaproteobacteria; Vibrionales; Vibrionaceae | MULTISPECIES: hypothetical protein | 14 |
|  | **WP_061011365.1** | 1.5E-05 | Qc.III | -------VGALKSQNASLGMQVEEQNKQIDKLMNQVDLQITMLSRAK------ | *Aliivibrio wodanis* | Bacteria; Proteobacteria; Gammaproteobacteria; Vibrionales; Vibrionaceae; Aliivibrio | hypothetical protein | 14 |
| **▼ 15** | | | | | | | | |
|  | **WP_096528190.1** | 3.3E-07 | SNAP.c | -----NAINSLNNQTHQLNQEIDKQNQAIDNLNQEIAGLNNEISAQNQAIKQ- | *Helicobacter pylori* | Bacteria; Proteobacteria; Epsilonproteobacteria; Campylobacterales; Helicobacteraceae; Helicobacter | UV radiation resistance protein | 15 |
|  | **WP_096529070.1** | 3.3E-07 | SNAP.c | -----NAINSLNNQTHQLNQEIDKQNQAIDNLNQEIAGLNNEISAQNQAIKQ- | *Helicobacter pylori* | Bacteria; Proteobacteria; Epsilonproteobacteria; Campylobacterales; Helicobacteraceae; Helicobacter | UV radiation resistance protein | 15 |
| **▼ 16** | | | | | | | | |
| | **PCS17992.1** | 3.4E-07 | Qa.III | --NVVRDMKELKDLVDQMSTTIATQAQMIDTRDQLLDEAYLKLESAETELI-- | *Lactococcus lactis subsp. lactis* | Bacteria; Firmicutes; Bacilli; Lactobacillales; Streptococcaceae; Lactococcus | hypothetical protein RU91_GL001756 | 16 |
| | **KSU27237.1** | 5.5E-06 | Qa.III | --NVVRDMKELKDLVDQMSATIATQAQMIDTRDQLLDEVYLKLESAETELI-- | *Lactococcus lactis subsp. lactis* | Bacteria; Firmicutes; Bacilli; Lactobacillales; Streptococcaceae; Lactococcus | hypothetical protein N42_1257 | 16 |
| | **ADZ64768.1** | 7.7E-06 | Qa.III | --NVVRDMTELKNLVDQMSSTIATQEQMIDTRDQLLDEVYLKLESAETELI-- | *Lactococcus lactis subsp. lactis CV56* | Bacteria; Firmicutes; Bacilli; Lactobacillales; Streptococcaceae; Lactococcus | phage protein | 16 |
| | **WP_023164645.1** | 3.7E-05 | Qa.III | --NVVRDMKELKDLVDQMTVTIATQAQMIDTRDQLLDEAFLKLESAETELI-- | *Lactococcus lactis* | Bacteria; Firmicutes; Bacilli; Lactobacillales; Streptococcaceae; Lactococcus | hypothetical protein | 16 |
| **▼ 17** | | | | | | | | |
|  | **WP_085164300.1** | 3.7E-07 | Qc.III | ----LAAVNALGANQREIGTTVRDHTARLDHIEGKVDLLATKSDDTNERVRS- | *Mycobacterium nebraskense* | Bacteria; Actinobacteria; Corynebacteriales; Mycobacteriaceae; Mycobacterium | hypothetical protein | 17 |
|  | **WP_067588660.1** | 4.7E-07 | SNAP.c | -----AVLSSMNALSQQTRERLDRQDARFDEVDRRFDHVDRRLDDTNARVR-- | *Nocardia terpenica* | Bacteria; Actinobacteria; Corynebacteriales; Nocardiaceae; Nocardia | hypothetical protein | 17 |
|  | **WP_015291713.1** | 6.4E-06 | Qc.III | -----AAVNALGANQREIATTVREHTGRLDRVETKVDQLAAKSDDTNARVRS- | *Mycobacterium canettii* | Bacteria; Actinobacteria; Corynebacteriales; Mycobacteriaceae; Mycobacterium; Mycobacterium tuberculosis complex | hypothetical protein | 17 |
|  | **WP_100470466.1** | 7.8E-06 | Qc.III | -----AAVNALGANQREIATTVREHTARLDRVETKVDQLAAKSDDTNARVRS- | *Mycobacteroides abscessus* | Bacteria; Actinobacteria; Corynebacteriales; Mycobacteriaceae | hypothetical protein | 17 |
|  | **WP_059020469.1** | 7.4E-05 | Qc.III | -----AAVNALGANQREMAGALAEHGVRLTSLESTLDDTNARVRS-------- | *Mycobacterium sp. M26* | Bacteria; Actinobacteria; Corynebacteriales; Mycobacteriaceae; Mycobacterium | hypothetical protein | 17 |
| **▼ 18** | | | | | | | | |
|  | **WP_015172395.1** | 4.2E-07 | SNAP.b | -------IRSAEERQRKAAAALEEQNQKLAQAQAEAERIKQQAQESAKKVRE- | *Geitlerinema sp. PCC 7407* | Bacteria; Cyanobacteria; Oscillatoriophycideae; Oscillatoriales; Coleofasciculaceae; Geitlerinema | F0F1 ATP synthase subunit B | 18 |
|  | **WP_023068158.1** | 1.1E-05 | Qb.III | ----EQAITDAEQRQKQAAETLAQQQQKLTQAQAEAERIRAEAQERAKALK-- | *Lyngbya aestuarii* | Bacteria; Cyanobacteria; Oscillatoriophycideae; Oscillatoriales; Oscillatoriaceae; Lyngbya | F0F1 ATP synthase subunit B | 18 |
| **▼ 19** | | | | | | | | |
|  | **WP_006826581.1** | 4.7E-07 | Qb.III | --------EATDESAEAVSDELATQSEQLRDAIDDLGIPDENVEEGRYRI--- | *Natrialba taiwanensis* | Archaea; Euryarchaeota; Halobacteria; Natrialbales; Natrialbaceae; Natrialba | DUF541 domain-containing protein | 19 |
|  | **WP_006110210.1** | 1.8E-06 | Qb.III | --------EATDESAEAVSDELATQSERLRDAIDELGIPDENVEEGRYRIR-- | *Natrialba asiatica* | Archaea; Euryarchaeota; Halobacteria; Natrialbales; Natrialbaceae; Natrialba | DUF541 domain-containing protein | 19 |
| **► 20** | | | | | | | | |
| **▼ 21** | | | | | | | | |
|  | **WP_054534729.1** | 5.9E-07 | Qc.I | ---------GLRERVQTLESSLNTRETNLDSLQKQIDSNLNSLLSIQSRLNEL | *Herpetosiphon geysericola* | Bacteria; Chloroflexi; Chloroflexia; Herpetosiphonales; Herpetosiphonaceae; Herpetosiphon | hypothetical protein | 21 |
|  | **WP_110515440.1** | 4.9E-06 | Qc.I | ---------GLRERVQTIESNLNTRETNLDSLQKQIDSNLNSLLSIQSRLDEL | *Herpetosiphon sp. CA052B* | Bacteria; Chloroflexi; Chloroflexia; Herpetosiphonales; Herpetosiphonaceae; Herpetosiphon | hypothetical protein | 21 |
| **▼ 22** | | | | | | | | |
|  | **WP_020027672.1** | 7.4E-07 | Qa.III | -VSLEEKIAYLQNMIDDLNMVVFRQGEKIEQLNSKLKETNEK----------- | *Candidatus Thioglobus singularis* | Bacteria; Proteobacteria; Gammaproteobacteria; Candidatus Thioglobus | hypothetical protein | 22 |
|  | **ALE01762.1** | 9E-07 | Qa.III | -VSLEEKIAYLQNMIDDLNMVVFRQGEKIEQLNSKLKETNEK----------- | *Candidatus Thioglobus singularis PS1* | Bacteria; Proteobacteria; Gammaproteobacteria; Candidatus Thioglobus | hypothetical protein W908_03730 | 22 |
| **► 23** | | | | | | | | |
| **▼ 24** | | | | | | | | |
|  | **CCJ34291.1** | 8E-07 | Qc.III | ----LKLLENLTEGVNGLREDLRETNRKLDRLEERMDKLESRMDKLESE---- | *Caloramator australicus RC3* | Bacteria; Firmicutes; Clostridia; Clostridiales; Clostridiaceae; Caloramator | hypothetical protein CAAU_2207 | 24 |
|  | **WP_051004417.1** | 8.1E-07 | Qc.III | ----LKLLENLTEGVNGLREDLRETNRKLDRLEERMDKLESRMDKLESE---- | *Caloramator australicus* | Bacteria; Firmicutes; Clostridia; Clostridiales; Clostridiaceae; Caloramator | hypothetical protein | 24 |
| | **EHQ91986.1** | 9.1E-06 | Qc.III | --ALSEKVDNQGNQIGALSEKVDNQGNQISALSEKVDNQGSQISALSEKVD-- | *Desulfosporosinus youngiae DSM 17734* | Bacteria; Firmicutes; Clostridia; Clostridiales; Peptococcaceae; Desulfosporosinus | hypothetical protein DesyoDRAFT_5052 | 24 |
| **▼ 25** | | | | | | | | |
| | **WP_020959621.1** | 1.2E-06 | Qc.III | IKELADQLQQTNGQVRQLGQQVQQMNEQVQQLSQHVQQTSEKVQHLNSQVQQL | *Geobacillus genomosp. 3* | Bacteria; Firmicutes; Bacilli; Bacillales; Bacillaceae; Geobacillus | hypothetical protein | 25 |
| | **KZE93102.1** | 8.2E-06 | Qc.III | -QQLGQQVQQTNEQVQQLGQQVQQTNEQVQQLGQQVQHLNSQVQQLNHQVQ-- | *Geobacillus stearothermophilus* | Bacteria; Firmicutes; Bacilli; Bacillales; Bacillaceae; Geobacillus | Chromosome partition protein Smc | 25 |
| | **ADU93899.1** | 9.9E-06 | Qc.III | -QQLGQQVQQTNEQVQQLGQQVQQTNEQVQQLGQQVQHLNSQVQQLNHQVQ-- | *Geobacillus sp. Y412MC52* | Bacteria; Firmicutes; Bacilli; Bacillales; Bacillaceae; Geobacillus | hypothetical protein GYMC52_1446 | 25 |
| | **WP_095859282.1** | 1E-05 | Qc.III | -QQLGQQVQQTNEQVQQLGQQVQQTNEQVQQLGQQVQHLNSQVQQLNHQVQ-- | *Geobacillus stearothermophilus* | Bacteria; Firmicutes; Bacilli; Bacillales; Bacillaceae; Geobacillus | hypothetical protein | 25 |
| | **WP_011230982.1** | 1.2E-05 | Qc.III | -QQLGQQVQQTNEQVQQLGQQVQQTNEQVQQLGQQVQHLNSQVQQLNHQVQ-- | *Geobacillus* | Bacteria; Firmicutes; Bacilli; Bacillales; Bacillaceae | MULTISPECIES: hypothetical protein | 25 |
| | **WP_020279830.1** | 1.2E-05 | Qc.III | -QQLGQQVQQTNEQVQQLGQQVQQTNEQVQQLGQQVQHLNSQVQQLNHQVQ-- | *Geobacillus kaustophilus* | Bacteria; Firmicutes; Bacilli; Bacillales; Bacillaceae; Geobacillus; Geobacillus thermoleovorans group | hypothetical protein | 25 |

| | | | | | | | | |
|---|---|---|---|---|---|---|---|---|
| | **WP_033843997.1** | 1.2E-05 | Qc.III | -ERLAKQVQQTNEQVEQLTKQVQQTNEQVERLAEQVQQTNEQVHRLSEQVSD- | *Geobacillus subterraneus* | Bacteria; Firmicutes; Bacilli; Bacillales; Bacillaceae; Geobacillus | hypothetical protein | 25 |
| | **PJW14603.1** | 1.3E-05 | Qc.III | -QQLGQQVQQTNEQVQQLGQQVQQTNEQVQQLGQQVQHLNSQVQQLNHQVQ-- | *Geobacillus sp. Manikaran-105* | Bacteria; Firmicutes; Bacilli; Bacillales; Bacillaceae; Geobacillus | hypothetical protein CV945_07745 | 25 |
| | **RAN23251.1** | 3.3E-05 | Qc.III | IKELADQLQQTNGQVQQLGEQVQQTNEKVQQLGQQVQHLNSQVQQLNHQVQ-- | *Geobacillus sp. A8* | Bacteria; Firmicutes; Bacilli; Bacillales; Bacillaceae; Geobacillus | hypothetical protein VC88_06955 | 25 |
| | **KPD01038.1** | 3.5E-05 | Qc.III | IKELADQLQQTNGQVQQLGEQVQQTNEKVQQLGQQVQHLNSQVQQLNHQVQ-- | *Geobacillus sp. BCO2* | Bacteria; Firmicutes; Bacilli; Bacillales; Bacillaceae; Geobacillus | Chromosome partition protein Smc | 25 |
| | **AMQ20382.1** | 4.4E-05 | Qc.III | -QQLGQQVQQTNEQVQQLGQQVQQTNEQVQQLGQQIQHLNSQVQQLNHQVQ-- | *Geobacillus sp. JS12* | Bacteria; Firmicutes; Bacilli; Bacillales; Bacillaceae; Geobacillus | hypothetical protein A0V43_04755 | 25 |
| | **WP_025950101.1** | 4.7E-05 | Qc.III | IKKLADQLQQTNGQVQQLGEQVQQTNEKVQQLGQQVQHLNSQVQQLNHQVQ-- | *Geobacillus thermocatenulatus* | Bacteria; Firmicutes; Bacilli; Bacillales; Bacillaceae; Geobacillus; Geobacillus thermoleovorans group | hypothetical protein | 25 |
| | **WP_051870828.1** | 5.9E-05 | Qc.III | -ERLAKQVQQTNEQVKQLTKQVQQTNEQVEQLTKQVQQTNEQVEQLTKQVQ-- | *Geobacillus vulcani* | Bacteria; Firmicutes; Bacilli; Bacillales; Bacillaceae; Geobacillus; Geobacillus thermoleovorans group | hypothetical protein | 25 |
| | **AKM18786.1** | 0.0001 | Qc.III | IKELADQLQQTNGQVQQLGQQVQQTNEQVQQLGQQVQHLNSQVQQLNHQVQ-- | *Geobacillus sp. 12AMOR1* | Bacteria; Firmicutes; Bacilli; Bacillales; Bacillaceae; Geobacillus | chromosome segregation protein | 25 |
| ▼ 26 | | | | | | | | |
|  | **PRC57448.1** | 1.4E-06 | Qa.III | ITQLVTDVANVNQLLQDLGAEIQMQQESVNKAILDVQTARDNASAAQLE---- | *Mycobacterium sp. ITM-2017-0098* | Bacteria; Actinobacteria; Corynebacteriales; Mycobacteriaceae; Mycobacterium | peptidase M23 partial | 26 |
|  | **PRC57449.1** | 1.6E-06 | Qa.III | ITQLVTDVANVNQLLQDLGAEIQMQQEGVNKAILDVQTARDNASAAQLE---- | *Mycobacterium sp. ITM-2017-0098* | Bacteria; Actinobacteria; Corynebacteriales; Mycobacteriaceae; Mycobacterium | peptidase M23 partial | 26 |
| ▼ 27 | | | | | | | | |
|  | **WP_026851021.1** | 1.5E-06 | Qa.III | --ELRESIRNFATLSADLARTVNAQSKNLDRISTNVQTGLTT---------- | *Gemmatimonas phototrophica* | Bacteria; Gemmatimonadetes; Gemmatimonadales; Gemmatimonadaceae; Gemmatimonas | MCE family protein | 27 |
| | **WP_096352890.1** | 1E-05 | Qa.III.b | ---IANSLQTLEGTTKKIDNLVGTQTVHINSIMSNTETVTSNLKTSSANL--- | *Mucilaginibacter gotjawali* | Bacteria; Bacteroidetes; Sphingobacteriia; Sphingobacteriales; Sphingobacteriaceae; Mucilaginibacter | MCE family protein | 27 |
| | **WP_008510398.1** | 2.1E-05 | Qa.IV | --SIANTLQSLEGTTQKVDALVGKQSVRINNILANVESISANLKSSNVYLT-- | *Mucilaginibacter paludis* | Bacteria; Bacteroidetes; Sphingobacteriia; Sphingobacteriales; Sphingobacteriaceae; Mucilaginibacter | MCE family protein | 27 |
|  | **WP_090647093.1** | 5.6E-05 | Qa.III | --SIANSLQTLEGVTKKIDNLVGTQTKHIDVILSNAEVVSSNLK--------- | *Mucilaginibacter sp. OK283* | Bacteria; Bacteroidetes; Sphingobacteriia; Sphingobacteriales; Sphingobacteriaceae; Mucilaginibacter | MCE family protein | 27 |
| ▼ 28 | | | | | | | | |
| | **PWI47941.1** | 1.6E-06 | Qc.III | -DQINGQMNQISSNLLTLTQRLEEQGVKIDRIDEKITDVQSKLQKIQSTLA-- | *Candidatus Heimdallarchaeota archaeon B3-JM-08* | Archaea; Asgard group; Candidatus Heimdallarchaeota | hypothetical protein CEE45_09200 | 28 |
| | **OLS22354.1** | 5.8E-05 | Qa.III | -NNLNAQISLIKSALDNMAGQLDEQGVVLVNIDEKIDVLDSKLDKAQEMLIK- | *Candidatus Heimdallarchaeota archaeon LC_2* | Archaea; Asgard group; Candidatus Heimdallarchaeota | hypothetical protein HeimC2_29900 | 28 |
| ▼ 29 | | | | | | | | |
| | **RAN66195.1** | 1.8E-06 | R.IV | ---LRNELETEKGKLNDAIGRIENNEGHIEALQDRANSLETRATNLEKQAN-- | *Dolosigranulum pigrum* | Bacteria; Firmicutes; Bacilli; Lactobacillales; Carnobacteriaceae; Dolosigranulum | hypothetical protein B8A45_00005 partial | 29 |
| | **RAN54946.1** | 6.6E-06 | R.IV | ---LRSDLEAEQTKLNDAINRIEKNEGHIEALQGKTDNLQNGITDLEKGAKEL | *Dolosigranulum pigrum* | Bacteria; Firmicutes; Bacilli; Lactobacillales; Carnobacteriaceae; Dolosigranulum | hypothetical protein B8A42_04700 partial | 29 |
| ▼ 30 | | | | | | | | |
|  | **WP_101562499.1** | 1.9E-06 | Qb.I | -----SKLSQEISKLKKTNKSLKQDNEKLKNSHEKLEKTAETLQS-------- | *Bacillus velezensis* | Bacteria; Firmicutes; Bacilli; Bacillales; Bacillaceae; Bacillus; Bacillus amyloliquefaciens group | hypothetical protein | 30 |
|  | **WP_039253607.1** | 5.9E-06 | Qb.I | -----SKLSQEISKLKKTNKSLKQDNEKLKNSHKKLEKTAETLQS-------- | *Bacillus* | Bacteria; Firmicutes; Bacilli; Bacillales; Bacillaceae | MULTISPECIES: membrane protein | 30 |
|  | **PRT02019.1** | 1.4E-05 | Qb.I | -----SKLSQEISKLKKTNKSLKQDNEKLKNSQKKLEKTAETLQS-------- | *Bacillus velezensis* | Bacteria; Firmicutes; Bacilli; Bacillales; Bacillaceae; Bacillus; Bacillus amyloliquefaciens group | hypothetical protein C6354_02680 | 30 |
|  | **WP_017418786.1** | 1.4E-05 | Qb.I | -----SKLSQEISKLKKTNKSLKQDNEKLKNSQKKLEKTAETLQS-------- | *Bacillus amyloliquefaciens group* | Bacteria; Firmicutes; Bacilli; Bacillales; Bacillaceae; Bacillus | MULTISPECIES: membrane protein | 30 |
|  | **WP_060673948.1** | 1.4E-05 | Qb.I | -----SKLSQEISKLKKTNKSLKQDNEKLKNSQKKLEKTAETLQS-------- | *Bacillus amyloliquefaciens* | Bacteria; Firmicutes; Bacilli; Bacillales; Bacillaceae; Bacillus; Bacillus amyloliquefaciens group | hypothetical protein | 30 |
|  | **WP_063636506.1** | 1.4E-05 | Qb.I | -----SKLSQEISKLKKTNKSLKQDNEKLKNSQKKLEKTAETLQS-------- | *Bacillus velezensis* | Bacteria; Firmicutes; Bacilli; Bacillales; Bacillaceae; Bacillus; Bacillus amyloliquefaciens group | hypothetical protein | 30 |
| ▼ 31 | | | | | | | | |
|  | **WP_089764467.1** | 2.1E-06 | Qc.III | -----AGVANLQEQADAIEDRLDETVDRLDDVDRDVDDVAAEVDA-------- | *Halobellus clavatus* | Archaea; Euryarchaeota; Halobacteria; Haloferacales; Haloferacaceae; Halobellus | hypothetical protein | 31 |
| | **WP_006629910.1** | 8.4E-06 | Qc.III.b | ---VDEAVESVRDAVSTVEAEVDDLGTDIDDLGTDVDRVEGEAEAVAESVDDL | *Halorubrum tebenquichense* | Archaea; Euryarchaeota; Halobacteria; Haloferacales; Halorubraceae; Halorubrum | hypothetical protein | 31 |
| ▼ 32 | | | | | | | | |
| | **WP_021620755.1** | 2.1E-06 | SNAP.c | LDLILEGIKELQQITRAIHDRQEETDAKLEALAMEVHKLHGEVASVKEDTKRI | *Aneurinibacillus aneurinilyticus* | Bacteria; Firmicutes; Bacilli; Bacillales; Paenibacillaceae; Aneurinibacillus group; Aneurinibacillus | hypothetical protein | 32 |
| | **WP_021622915.1** | 2.1E-06 | SNAP.c | LDLILEGIKELQQITRAIHDRQEETDAKLEALAMEVHKLHGEVASVKEDTKRI | *Aneurinibacillus aneurinilyticus* | Bacteria; Firmicutes; Bacilli; Bacillales; Paenibacillaceae; Aneurinibacillus group; Aneurinibacillus | hypothetical protein | 32 |
| | **WP_061566932.1** | 8.8E-05 | SNAP.c | ---MKSGQDELYQMVRAIHDRQEETDAKLDALSMDVHQAHGEIAGVKDAVKEL | *Bacillus coagulans* | Bacteria; Firmicutes; Bacilli; Bacillales; Bacillaceae; Bacillus | hypothetical protein | 32 |
| ▼ 33 | | | | | | | | |
|  | **WP_087440691.1** | 2.4E-06 | Qc.III | -QMLSGDLQAIARNANHMTEELQGQLKKIEPVADSVQNAGEALNELT------ | *Paenibacillus thiaminolyticus* | Bacteria; Firmicutes; Bacilli; Bacillales; Paenibacillaceae; Paenibacillus | DUF948 domain-containing protein | 33 |
|  | **WP_111156875.1** | 3.1E-06 | Qc.III | -QMLSGDLQAIARNANHMTEELQGQLKKIEPVADSVQNAGEALNELT------ | *Paenibacillus dendritiformis* | Bacteria; Firmicutes; Bacilli; Bacillales; Paenibacillaceae; Paenibacillus | DUF948 domain-containing protein | 33 |
|  | **WP_006677149.1** | 5.2E-06 | Qc.III | --MLSGDLQAIARNANHMTEELQGQLKKIEPVADSVQNAGEALNELT------ | *Paenibacillus dendritiformis* | Bacteria; Firmicutes; Bacilli; Bacillales; Paenibacillaceae; Paenibacillus | DUF948 domain-containing protein | 33 |
|  | **WP_006287073.1** | 4.7E-05 | Qc.III | --MLSGDLQAVARNANHMTEELQGQLKKIEPVADSVQNAGEALNELT------ | *Paenibacillus popilliae* | Bacteria; Firmicutes; Bacilli; Bacillales; Paenibacillaceae; Paenibacillus | DUF948 domain-containing protein | 33 |
|  | **WP_028544524.1** | 6.4E-05 | Qc.III | -EALSGDLQAVARNANEMTAGIQQQVKKIEPLVDSIENIGEALNEVT------ | *Paenibacillus taiwanensis* | Bacteria; Firmicutes; Bacilli; Bacillales; Paenibacillaceae; Paenibacillus | DUF948 domain-containing protein | 33 |
| ▼ 34 | | | | | | | | |
| | **WP_092625088.1** | 2.6E-06 | Qb.III | LEQAEAAVDAAEDQVDDAEAALDDQRQRLERAQEAVERLTEELAAARREL--- | *Jiangella sp. DSM 45060* | Bacteria; Actinobacteria; Jiangellales; Jiangellaceae; Jiangella | hypothetical protein | 34 |
| | **WP_069109736.1** | 2.9E-06 | Qb.III | LEQAEAAVDAAEEQVDDAEAALDDQRQRLERAQEAVERLTEELAAARREL--- | *Jiangella alba* | Bacteria; Actinobacteria; Jiangellales; Jiangellaceae; Jiangella | hypothetical protein | 34 |
| | **WP_111255308.1** | 7.2E-06 | Qb.III | LERAEADVDAAEDRVDDAEAALDDQRQRLERAQDAVERLTEELNAARREL--- | *Jiangella sp. GTF31* | Bacteria; Actinobacteria; Jiangellales; Jiangellaceae; Jiangella | hypothetical protein | 34 |
| | **WP_112247881.1** | 7.2E-06 | Qb.III.b | -EDAVAAAESTEAAVADLAGQLEAAREALADAQEAVHRIGDELDEARKAAR-- | *Kribbella sp. NEAU-SW521* | Bacteria; Actinobacteria; Propionibacteriales; Nocardioidaceae; Kribbella | hypothetical protein | 34 |
| | **WP_053204370.1** | 1.1E-05 | Qb.III | LEQAEAAVDAAEEQVDDAEAALDDQRRRLGRAQEAVERLTEELAAARREL--- | *Jiangella muralis* | Bacteria; Actinobacteria; Jiangellales; Jiangellaceae; Jiangella | hypothetical protein | 34 |
|  | **WP_107102838.1** | 2.1E-05 | SNAP.b | -DRARRQVSAAEQQLRKAREALARAQEQVGRAEEHAGHAEEDLDRAEK----- | *Streptomyces torulosus* | Bacteria; Actinobacteria; Streptomycetales; Streptomycetaceae; Streptomyces | hypothetical protein | 34 |
| | **SDU12491.1** | 7.5E-05 | Qb.III | LEQAESAVDAAEEQVDDAEAALDEQRQRLERAQQTAERLTEELAAARREL--- | *Jiangella alkaliphila* | Bacteria; Actinobacteria; Jiangellales; Jiangellaceae; Jiangella | hypothetical protein SAMN04488563_0195 | 34 |
| ▼ 35 | | | | | | | | |
|  | **PYN93259.1** | 2.8E-06 | Qc.I | ---------QLKTNFAAMEATVEADRAAVDNARAAIRANQAALDTVRLNLG-- | *Candidatus Rokubacteria bacterium* | Bacteria; Candidatus Rokubacteria | hypothetical protein DMD91_30520 partial | 35 |
|  | **WP_056527516.1** | 3.8E-06 | Qc.III | -----AQVERDQAQSGDIGATIDRQPALIEQAAAEVDAAQARLGF-------- | *Methylobacterium sp. Leaf361* | Bacteria; Proteobacteria; Alphaproteobacteria; Rhizobiales; Methylobacteriaceae; Methylobacterium | HlyD family secretion protein | 35 |
|  | **PYU71650.1** | 1.3E-05 | Qc.I | -----------KANADATASALKADQERLAQAERNVDVAKAQLGAAKARVE-- | *Acidobacteria bacterium* | Bacteria; Acidobacteria | HlyD family secretion protein | 35 |
|  | **OLB19831.1** | 1.9E-05 | Qc.I | -----------KANADATASALKADQEKLAQAQRNVDVTKAQLDAAKARVE-- | *Acidobacteria bacterium 13_2_20CM_57_7* | Bacteria; Acidobacteria | hypothetical protein AUH16_06355 | 35 |
|  | **PYT46827.1** | 2E-05 | Qc.I | -----------KANADATASALKADQEKLAQAQRNVDVTKAQLDAAKARVE-- | *Acidobacteria bacterium* | Bacteria; Acidobacteria | HlyD family secretion protein | 35 |
|  | **PYU48547.1** | 2E-05 | Qc.I | -----------KANADATASALKADQEKLAQAQRNVDVTKAQLDAAKARVE-- | *Acidobacteria bacterium* | Bacteria; Acidobacteria | HlyD family secretion protein | 35 |
|  | **OLE47468.1** | 2.3E-05 | Qc.I | -----------KANADATASALKADQERLAQAERNVDVAKAQLDAAKARVE-- | *Acidobacteria bacterium 13_1_20CM_3_58_11* | Bacteria; Acidobacteria | hypothetical protein AUG46_06400 | 35 |
|  | **PYU63255.1** | 2.4E-05 | Qc.I | -----------KANADATASALKADQERLAQAERNVDVAKAQLDAAKARVE-- | *Acidobacteria bacterium* | Bacteria; Acidobacteria | HlyD family secretion protein | 35 |
|  | **PYU64012.1** | 2.4E-05 | Qc.I | -----------KANADATASALKADQERLAQAERNVDVAKAQLDAAKARVE-- | *Acidobacteria bacterium* | Bacteria; Acidobacteria | HlyD family secretion protein | 35 |
|  | **PYU85121.1** | 2.4E-05 | Qc.I | -----------KANADATASALKADQERLAQAERNVDVAKAQLDAAKARVE-- | *Acidobacteria bacterium* | Bacteria; Acidobacteria | HlyD family secretion protein | 35 |
|  | **PYU69244.1** | 3.1E-05 | Qc.I | -----------KANADATASALKADQEKLAQAQRNVDVAKAQLDAAKARVE-- | *Acidobacteria bacterium* | Bacteria; Acidobacteria | HlyD family secretion protein | 35 |
|  | **WP_109502814.1** | 6.2E-05 | SNAP.b | ----------ARAQLQNARATLERQTSVIAQAQATVVSAKAEMDFAEHELN-- | *Pseudomonas prosekii* | Bacteria; Proteobacteria; Gammaproteobacteria; Pseudomonadales; Pseudomonadaceae; Pseudomonas | HlyD family secretion protein partial | 35 |
|  | **PYU05495.1** | 8E-05 | Qc.I | -----------KANADATGSALKADEERLAQAQHNVEITKAQSLAATARVEE- | *Acidobacteria bacterium* | Bacteria; Acidobacteria | HlyD family secretion protein | 35 |
|  | **OLC99654.1** | 0.0001 | Qc.I | -----------KANADATASTLKADQQRLAQAQRNVDVAKAQLDAAKARVE-- | *Acidobacteria bacterium 13_1_40CM_4_58_4* | Bacteria; Acidobacteria | hypothetical protein AUH86_02020 | 35 |
| ▼ 36 | | | | | | | | |
|  | **PKM77570.1** | 3.3E-06 | Qc.II | -EELFHRIKALRNYGSQIkyHNEVKGYNSRLDELQAAFLRVK------------ | *Firmicutes bacterium HGW-Firmicutes-15* | Bacteria; Firmicutes | erythromycin biosynthesis sensory transduction protein eryC1 | 36 |

| | | | | | | | | |
|---|---|---|---|---|---|---|---|---|
|  | **WP_107869754.1** | 3.5E-06 | Qc.II | -DALAEQLRMLRNYGSRIkyQNEVQGHNSRLDELQAAILRA------------- | *Massilia* | Bacteria; Proteobacteria; Betaproteobacteria; Burkholderiales; Oxalobacteraceae | MULTISPECIES: DegT/DnrJ/EryC1/StrS family aminotransferase | 36 |
|  | **WP_004514702.1** | 1.1E-05 | Qc.II | -DQLAKKIRVLRNYGSQIkyHNEIKGYNSRLDELQAAFLRVK------------ | *Geobacter metallireducens* | Bacteria; Proteobacteria; Deltaproteobacteria; Desulfuromonadales; Geobacteraceae; Geobacter | DegT/DnrJ/EryC1/StrS family aminotransferase | 36 |
|  | **WP_085811387.1** | 8.9E-05 | Qc.II | -DVLADKVAALRNYGSKVkyHNEVIGYNSRLDEIHAAFLRVK------------ | *Sphingomonas sp. TZW2008* | Bacteria; Proteobacteria; Alphaproteobacteria; Sphingomonadales; Sphingomonadaceae; Sphingomonas | DegT/DnrJ/EryC1/StrS family aminotransferase | 36 |
| ▼ **37** | | | | | | | | |
|  | **KRN30566.1** | 3.4E-06 | Qb.I | -------LAQEIERNHYSSKILAASFHTVQQVNSALENGAQAVTMGAD----- | *Lactobacillus selangorensis* | Bacteria; Firmicutes; Bacilli; Lactobacillales; Lactobacillaceae; Lactobacillus | fructose-6-phosphate aldolase | 37 |
|  | **WP_057770480.1** | 3.4E-06 | Qb.I | -------LAQEIERNHYSSKILAASFHTVQQVNSALENGAQAVTMGAD----- | *Lactobacillus selangorensis* | Bacteria; Firmicutes; Bacilli; Lactobacillales; Lactobacillaceae; Lactobacillus | fructose-6-phosphate aldolase | 37 |
|  | **WP_049486453.1** | 1.6E-05 | Qb.I | -----AQLADSIEKNQSSSKILAASFKNVGQVNRAFKEGAQAITAGA------ | *Streptococcus* | Bacteria; Firmicutes; Bacilli; Lactobacillales; Streptococcaceae | MULTISPECIES: fructose-6-phosphate aldolase | 37 |
|  | **PKZ96739.1** | 9.5E-05 | Qb.I | -----AQLAEAIEKNHSSSKILAASFKNVGQVNRAFKEGAQAITAGA------ | *Streptococcus parasanguinis* | Bacteria; Firmicutes; Bacilli; Lactobacillales; Streptococcaceae; Streptococcus | fructose-6-phosphate aldolase | 37 |
|  | **WP_003014188.1** | 9.5E-05 | Qb.I | -----AQLAEAIEKNHSSSKILAASFKNVGQVNRAFKEGAQAITAGA------ | *Streptococcus parasanguinis* | Bacteria; Firmicutes; Bacilli; Lactobacillales; Streptococcaceae; Streptococcus | fructose-6-phosphate aldolase | 37 |
|  | **WP_049518621.1** | 9.5E-05 | Qb.I | -----AQLAEAIEKNHSSSKILAASFKNVGQVNRAFKEGAQAITAGA------ | *Streptococcus parasanguinis* | Bacteria; Firmicutes; Bacilli; Lactobacillales; Streptococcaceae; Streptococcus | fructose-6-phosphate aldolase | 37 |
|  | **AFJ25377.1** | 9.7E-05 | Qb.I | -----AQLAEAIEKNHSSSKILAASFKNVGQVNRAFKEGAQAITAGA------ | *Streptococcus parasanguinis FW213* | Bacteria; Firmicutes; Bacilli; Lactobacillales; Streptococcaceae; Streptococcus | Transaldolase-like protein | 37 |
| ▼ **38** | | | | | | | | |
| | **WP_078434151.1** | 3.6E-06 | Qc.III | --RLNNEIKRLNEENRRMNREITRLNEQVNRVNNRVEQHTRRLNRLNQRLR-- | *Bacillus halosaccharovorans* | Bacteria; Firmicutes; Bacilli; Bacillales; Bacillaceae; Bacillus | hypothetical protein | 38 |
|  | **WP_110067799.1** | 2.6E-05 | Qc.III | --RLNRQLTNLSDQVRMLNRTVDRHTRRLNRLNQRLRAVENRLN--------- | *Bacillus oceanisediminis* | Bacteria; Firmicutes; Bacilli; Bacillales; Bacillaceae; Bacillus | hypothetical protein | 38 |
|  | **WP_026561684.1** | 5.5E-05 | Qc.III | -------IKRLNEENRRMNREISRLNEQVVRVNNRVEQHTRRLNRLNQRLR-- | *Bacillus sp. J37* | Bacteria; Firmicutes; Bacilli; Bacillales; Bacillaceae; Bacillus | hypothetical protein | 38 |
| ▼ **39** | | | | | | | | |
|  | **WP_089993184.1** | 3.6E-06 | Qc.I | -----------KSNAFAISEKLGLDKRIIERAAREMDETEGNFEDLLKDLE-- | *Lachnospiraceae bacterium G41* | Bacteria; Firmicutes; Clostridia; Clostridiales; Lachnospiraceae | endonuclease MutS2 | 39 |
|  | **OGN81777.1** | 2.6E-05 | Qb.III | IQSARGLLDPTNLQADDLLDEINQQRTIARKARGAADRARFEVDQIRKD---- | *Chloroflexi bacterium GWB2_54_36* | Bacteria; Chloroflexi | hypothetical protein A2X24_04495 | 39 |
| | **OEU77091.1** | 9E-05 | Qc.II | LAALAGQLTALRELSRRLLNSIGSRGELLDSASFELGDLRYRIRQTRTRIK-- | *Desulfuromonadales bacterium C00003093* | Bacteria; Proteobacteria; Deltaproteobacteria; Desulfuromonadales | hypothetical protein BA864_14750 | 39 |
| ▼ **40** | | | | | | | | |
|  | **WP_056244371.1** | 3.8E-06 | Qc.III | ----SDQLARIEDKAARIEDKYARSEALLSRVEDKVENASSRMNEAARQ---- | *Methylobacterium sp. Leaf456* | Bacteria; Proteobacteria; Alphaproteobacteria; Rhizobiales; Methylobacteriaceae; Methylobacterium | hypothetical protein | 40 |
|  | **WP_091950135.1** | 5.6E-06 | Qc.III | -----DQLARIEDKAARIEDKYARSEALLSRVEDKVENASSRMNEAARQ---- | *Methylorubrum salsuginis* | Bacteria; Proteobacteria; Alphaproteobacteria; Rhizobiales; Methylobacteriaceae; Methylorubrum | hypothetical protein | 40 |
|  | **SFL60932.1** | 7E-06 | Qc.III | -----DQLARIEDKAARIEDKYARSEALLSRVEDKVENASSRMNEAARQ---- | *Methylorubrum salsuginis* | Bacteria; Proteobacteria; Alphaproteobacteria; Rhizobiales; Methylobacteriaceae; Methylorubrum | hypothetical protein SAMN04488125_1199 | 40 |
|  | **WP_099956808.1** | 4E-05 | Qc.III | --QVADQLARIEDKASRIEDKYARSEALLTRVEDRVDAAGARMNEAARQ---- | *Methylobacterium sp. PR1016A* | Bacteria; Proteobacteria; Alphaproteobacteria; Rhizobiales; Methylobacteriaceae; Methylobacterium | hypothetical protein | 40 |
|  | **OAS17275.1** | 0.0001 | Qc.III | --QVADQLARIEDKASRIEDKYARSEALLGRVEDRVEAAGARMNEAARQ---- | *Methylobacterium platani* | Bacteria; Proteobacteria; Alphaproteobacteria; Rhizobiales; Methylobacteriaceae; Methylobacterium | hypothetical protein A5481_27510 | 40 |
|  | **WP_109962236.1** | 0.0001 | Qc.III | --QVADQLARIEDKASRIEDKYARSEALLGRVEDRVEAAGARMNEAARQ---- | *Methylobacterium sp. 17Sr1-28* | Bacteria; Proteobacteria; Alphaproteobacteria; Rhizobiales; Methylobacteriaceae; Methylobacterium | hypothetical protein | 40 |
| ▼ **41** | | | | | | | | |
|  | **AKT34085.1** | 3.9E-06 | Qc.III | ---------RIDEVAKSLSARIDETNKRIDEVNKRIDETNKRIDDLARRVDEL | *Pyrobaculum sp. WP30* | Archaea; Crenarchaeota; Thermoprotei; Thermoproteales; Thermoproteaceae; Pyrobaculum | hypothetical protein PYWP30_00429 | 41 |
|  | **KUO87578.1** | 4.5E-06 | Qc.III | ------------ELAKSLSARIDDTNKRIDELAKRMDDLAKRIDETNKRIDA- | *Thermoproteus sp. JCHS_4* | Archaea; Crenarchaeota; Thermoprotei; Thermoproteales; Thermoproteaceae; Thermoproteus | hypothetical protein AT715_05195 | 41 |
|  | **KUO82390.1** | 1.3E-05 | Qc.III | --------NRINAVNDALSRKIEETNKRIDEVNRRIDELSKGMNEMRESIN-- | *Vulcanisaeta sp. JCHS_4* | Archaea; Crenarchaeota; Thermoprotei; Thermoproteales; Thermoproteaceae; Vulcanisaeta | hypothetical protein AT718_08285 | 41 |
|  | **AEM39092.1** | 8.8E-05 | Qc.III | --------KRINELNERINERIDGLNKRIDEVNRRIDKVHDELSEKISR---- | *Pyrolobus fumarii 1A* | Archaea; Crenarchaeota; Thermoprotei; Desulfurococcales; Pyrodictiaceae; Pyrolobus | paREP15 putative coiled-coil protein | 41 |
| ▼ **42** | | | | | | | | |
|  | **PYR54580.1** | 4E-06 | Qc.III.c | -EELSAALNNVRAQAEMMRSEVKSQRERVETLTAHVNEAREAQ---------- | *Acidobacteria bacterium* | Bacteria; Acidobacteria | hypothetical protein DMF85_21900 | 42 |
|  | **PYR77467.1** | 4E-06 | Qc.III.c | -EELSAALNNVRAQAEMMRSEVKSQRERVETLTAHVNEAREAQ---------- | *Acidobacteria bacterium* | Bacteria; Acidobacteria | hypothetical protein DMF86_09090 | 42 |
| ▼ **43** | | | | | | | | |
|  | **KKU99810.1** | 4E-06 | Qb.III | LKQTIDNATATGKQVDELLKQLNTQKATIEDFVQKFTEINQSVGQSK------ | *Parcubacteria group bacterium GW2011_GWA2_48_9* | Bacteria; unclassified Parcubacteria group | hypothetical protein UY34_C0037G0005 | 43 |
|  | **OGY84025.1** | 4E-06 | Qb.III | LKQTIDNATATGKQVDELLKQLNTQKATIEDFVQKFTEINQSVGQSK------ | *Candidatus Kerfeldbacteria bacterium RIFCSPLOWO2_01_FULL_48_11* | Bacteria; Candidatus Kerfeldbacteria | hypothetical protein A2898_02025 | 43 |
| ▼ **44** | | | | | | | | |
|  | **KFF41983.1** | 4.3E-06 | Qa.IV | --EIEVLIKGLRKLPSQIEIILEKQGEQIQELSHNFCDTQDF----------- | *Candidatus Atelocyanobacterium thalassa isolate SIO64986* | Bacteria; Cyanobacteria; Oscillatoriophycideae; Chroococcales; Aphanothecaceae; Candidatus Atelocyanobacterium | glutamine--fructose-6-phosphate transaminase | 44 |
|  | **WP_012953769.1** | 4.5E-05 | Qa.IV | --KIEILIKGLRKLPNQIEIILEKQGEQIQELSHNFCDTQDF----------- | *Candidatus Atelocyanobacterium thalassa* | Bacteria; Cyanobacteria; Oscillatoriophycideae; Chroococcales; Aphanothecaceae; Candidatus Atelocyanobacterium | glutamine--fructose-6-phosphate transaminase (isomerizing) | 44 |
| ▼ **45** | | | | | | | | |
|  | **CUO65260.1** | 4.7E-06 | Qc.III.b | --------NRTVTLVNSINSTIGTHTTKIDELSGNITSVDTKVNSVQRDL--- | *Clostridium disporicum* | Bacteria; Firmicutes; Clostridia; Clostridiales; Clostridiaceae; Clostridium | phage minor structural protein | 45 |
|  | **OSB07979.1** | 8.8E-05 | Qc.III.b | --------NRTVQTVNSINSTLGSHTTQINEATGKITSVDTKVNSVQRDL--- | *Paraclostridium bifermentans* | Bacteria; Firmicutes; Clostridia; Clostridiales; Peptostreptococcaceae; Paraclostridium | hypothetical protein B2H97_16045 | 45 |
|  | **WP_085283750.1** | 8.8E-05 | Qc.III.b | --------NRTVQTVNSINSTLGSHTTQINEATGKITSVDTKVNSVQRDL--- | *Paraclostridium bifermentans* | Bacteria; Firmicutes; Clostridia; Clostridiales; Peptostreptococcaceae; Paraclostridium | hypothetical protein | 45 |
| ▼ **46** | | | | | | | | |
|  | **WP_004753016.1** | 4.9E-06 | Qc.I | ---------QLKQQVTHLEDHLKKTLEIMQDLVKKMHTQQEVLDRTSGKLQQ- | *Acinetobacter sp. ANC 3789* | Bacteria; Proteobacteria; Gammaproteobacteria; Pseudomonadales; Moraxellaceae; Acinetobacter | hypothetical protein | 46 |
|  | **WP_004903937.1** | 5.9E-06 | Qc.I | ---------QLKQQVTHLEDHLKKTLEIMQDLAKKMHIQQEVLDRTSGKLQQ- | *Acinetobacter brisouii* | Bacteria; Proteobacteria; Gammaproteobacteria; Pseudomonadales; Moraxellaceae; Acinetobacter | hypothetical protein | 46 |
|  | **WP_045793660.1** | 6.8E-06 | Qc.I | ---------QLKQQVTHLEDHLKKTLEIMQDLAKKMHIQQEVLDRTTGKLQQ- | *Acinetobacter brisouii* | Bacteria; Proteobacteria; Gammaproteobacteria; Pseudomonadales; Moraxellaceae; Acinetobacter | hypothetical protein | 46 |
| ▼ **47** | | | | | | | | |
| | **PSN10961.1** | 5.1E-06 | Qc.III | --RLERDMTRQEDLVNALQDELEAQKDLADDLKAQLERQQAETEQVMAQLD-- | *filamentous cyanobacterium CCT1* | Bacteria; Cyanobacteria | hypothetical protein C7293_26170 partial | 47 |
| | **PSN77808.1** | 6E-06 | Qc.III | --RLERDMTRQEDLVNALQDELEAQKDLADDLKAQLERQQAETEQVMAQLD-- | *filamentous cyanobacterium CCP4* | Bacteria; Cyanobacteria | hypothetical protein C8B47_20100 partial | 47 |
|  | **WP_073608118.1** | 6.9E-06 | Qc.III | -------LNRQQDLVNSLQDELRAQENLADDLKAQLERQQAETQAVREQLN-- | *Phormidium tenue* | Bacteria; Cyanobacteria; Oscillatoriophycideae; Oscillatoriales; Oscillatoriaceae; Phormidium | hypothetical protein | 47 |
| | **WP_106919665.1** | 7.7E-06 | Qc.III | --RLERDMTRQEDLVNALQDELEAQKDLADDLKAQLERQQAETEQVMAQLDN- | *filamentous cyanobacterium CCP3* | Bacteria; Cyanobacteria | hypothetical protein | 47 |
| | **PZO43921.1** | 6.3E-05 | Qc.III | -DRMERDLTRQQDLVTSLQDELSAQKDLADSLKTQLERQQTETDKVLEQLD-- | *Leptolyngbya antarctica* | Bacteria; Cyanobacteria; Synechococcales; Leptolyngbyaceae; Leptolyngbya | hypothetical protein DCF17_05130 | 47 |
|  | **PZV09497.1** | 7.1E-05 | Qc.III | -------LNRQQDLVSSLQDELRAQENLADDLKAQLERQQAETEAVLEQLD-- | *Leptolyngbya sp.* | Bacteria; Cyanobacteria; Synechococcales; Leptolyngbyaceae; Leptolyngbya | hypothetical protein DCF32_02285 | 47 |
| | **PZV20627.1** | 7.3E-05 | Qc.III | -DRMERDLTRQQDLVTSLQDELSAQKDLADSLKTQLERQQTETDKVLEQLD-- | *Leptolyngbya sp.* | Bacteria; Cyanobacteria; Synechococcales; Leptolyngbyaceae; Leptolyngbya | hypothetical protein DCF21_04470 | 47 |
| ▼ **48** | | | | | | | | |
|  | **WP_089347130.1** | 5.8E-06 | Qb.III.d | --------TMAEYLSPELAQEFKAEREAIESVDTEVMFIDAELVRADT----- | *Pseudoalteromonas espejiana* | Bacteria; Proteobacteria; Gammaproteobacteria; Alteromonadales; Pseudoalteromonadaceae; Pseudoalteromonas | Tim44 domain-containing protein | 48 |
|  | **WP_055014029.1** | 5.9E-06 | Qb.III.d | --------TMAEYLSPELAQEFKAEREAIESVDTEVMFIDAELVRADT----- | *Pseudoalteromonas* | Bacteria; Proteobacteria; Gammaproteobacteria; Alteromonadales; Pseudoalteromonadaceae | MULTISPECIES: Tim44 domain-containing protein | 48 |
|  | **WP_055019201.1** | 5.9E-06 | Qb.III.d | --------TMAEYLSPELAQEFKAEREAIESVDTEVMFIDAELVRADT----- | *Pseudoalteromonas sp. P1-7a* | Bacteria; Proteobacteria; Gammaproteobacteria; Alteromonadales; Pseudoalteromonadaceae; Pseudoalteromonas | Tim44 domain-containing protein | 48 |
|  | **WP_039609090.1** | 2.5E-05 | Qb.III.d | -------ATMSEYLSPELVEEFRQEREQQGDVETEVMFVDAELARADV----- | *Pseudoalteromonas luteoviolacea* | Bacteria; Proteobacteria; Gammaproteobacteria; Alteromonadales; Pseudoalteromonadaceae; Pseudoalteromonas | Tim44 domain-containing protein | 48 |
|  | **WP_063367376.1** | 2.5E-05 | Qb.III.d | -------ATMSEYLSPELVEEFRQEREQQGDVETEVMFVDAELARADV----- | *Pseudoalteromonas luteoviolacea* | Bacteria; Proteobacteria; Gammaproteobacteria; Alteromonadales; Pseudoalteromonadaceae; Pseudoalteromonas | Tim44 domain-containing protein | 48 |
|  | **WP_063378232.1** | 5.3E-05 | Qb.III.d | --------TMAEYLSPELVEEFRQEREQQGDVETEVMFVDAELARADVT---- | *Pseudoalteromonas luteoviolacea* | Bacteria; Proteobacteria; Gammaproteobacteria; Alteromonadales; Pseudoalteromonadaceae; Pseudoalteromonas | Tim44 domain-containing protein | 48 |
| ▼ **49** | | | | | | | | |
|  | **EKD81026.1** | 6.9E-06 | Qc.I | ------------QNIQSAGKRVEQDKARLKEAWAKMGKALEATKTETGKLKE- | *uncultured bacterium* | Bacteria; environmental samples | hypothetical protein ACD_39C02024G0001 partial | 49 |
|  | **OGK07990.1** | 2.9E-05 | Qc.I | ------------QNIQSAGKRVEQDKARLKEAWAKMGKALEATKTETGKLKE- | *Candidatus Riflebacteria bacterium GWC2_50_8* | Bacteria; Candidatus Riflebacteria | hypothetical protein A2W80_09220 | 49 |

| | | | | | | | | |
|---|---|---|---|---|---|---|---|---|
| ▼ 50 | | | | | | | | |
|  | BAI93715.1 | 7.3E-06 | Qc.I | -EELLDAANRLKDMMEQISNIIDQTSDTVvttkDTMVDAADTTSTGL--------- | *Arthrospira platensis NIES-39* | Bacteria; Cyanobacteria; Oscillatoriophycideae; Oscillatoriales; Microcoleaceae; Arthrospira | hypothetical protein NIES39_O04680 | 50 |
|  | KDR56878.1 | 7.3E-06 | Qc.I | -EELLDAANRLKDMMEQISNIIDQTSDTVvttkDTMVDAADTTSTGL--------- | *Arthrospira platensis str. Paraca* | Bacteria; Cyanobacteria; Oscillatoriophycideae; Oscillatoriales; Microcoleaceae; Arthrospira | hypothetical protein APPUASWS_014040 | 50 |
| ▼ 51 | | | | | | | | |
|  | KRN55034.1 | 7.6E-06 | Qc.III | LDQLKKMVEKLEYKVDEINERLNAHTVDLSKIDLRVTALENRM---------- | *Atopobium minutum* | Bacteria; Actinobacteria; Coriobacteriia; Coriobacteriales; Atopobiaceae; Atopobium | hypothetical protein IV72_GL000530 | 51 |
|  | SEB43353.1 | 1.4E-05 | Qc.III | LDQLKKMVEKLEYKVDEINERLNAHTVDLSKIDLRVTALENRM---------- | *Atopobium minutum* | Bacteria; Actinobacteria; Coriobacteriia; Coriobacteriales; Atopobiaceae; Atopobium | hypothetical protein SAMN04489746_0205 | 51 |
| ▼ 52 | | | | | | | | |
|  | WP_004248171.1 | 9.2E-06 | Qc.III | LDQISSDVNRLNTQVQQLSSDVQSANAQAKAAYDEAARANQRLDNQ------- | *Proteus* | Bacteria; Proteobacteria; Gammaproteobacteria; Enterobacterales; Morganellaceae | MULTISPECIES: hypothetical protein | 52 |
|  | WP_074924084.1 | 9.2E-06 | Qc.III | LDQISSDVNRLNTQVQQLSSDVQSANAQAKAAYDEAARANQRLDNQ------- | *Proteus mirabilis* | Bacteria; Proteobacteria; Gammaproteobacteria; Enterobacterales; Morganellaceae; Proteus | murein lipoprotein | 52 |
|  | EEI48914.1 | 1.5E-05 | Qc.III | LDQISSDVNRLNTQVQQLSSDVQSANAQAKAAYDEAARANQRLDNQ------- | *Proteus mirabilis ATCC 29906* | Bacteria; Proteobacteria; Gammaproteobacteria; Enterobacterales; Morganellaceae; Proteus | lipoprotein LPP repeat protein | 52 |
|  | WP_012988932.1 | 2.9E-05 | Qc.III.b | ------KVEQLATQVNTLGSKVDQLRNDLSAVHSDVQSAKEEAARANKRLD-- | *Xenorhabdus bovienii* | Bacteria; Proteobacteria; Gammaproteobacteria; Enterobacterales; Morganellaceae; Xenorhabdus | major outer membrane lipoprotein | 52 |
|  | WP_038190978.1 | 2.9E-05 | Qc.III.b | ------KVEQLATQVNTLGSKVDQLRNDLSAVHSDVQSAKEEAARANKRLD-- | *Xenorhabdus bovienii* | Bacteria; Proteobacteria; Gammaproteobacteria; Enterobacterales; Morganellaceae; Xenorhabdus | major outer membrane lipoprotein | 52 |
|  | WP_038233883.1 | 4.8E-05 | Qc.III.b | ------KVEQLASQVQTLGSKIDQLNNDLSSVRSDVQTAKDEAARANQRLN-- | *Xenorhabdus szentirmaii* | Bacteria; Proteobacteria; Gammaproteobacteria; Enterobacterales; Morganellaceae; Xenorhabdus | major outer membrane lipoprotein | 52 |
|  | WP_011411257.1 | 4.9E-05 | Qc.III | IDQLSSDVQTLNAKVDHISNDVNALSSDLQAAKDDAARANQRLDNQ------- | *Sodalis glossinidius* | Bacteria; Proteobacteria; Gammaproteobacteria; Enterobacterales; Pectobacteriaceae; Sodalis | major outer membrane lipoprotein | 52 |
|  | WP_047964342.1 | 5.2E-05 | Qc.III.b | ------KVEQLASQVQTLGSKVDQLSNDLSSIRSDVQSAKDEAARANQRLN-- | *Xenorhabdus* | Bacteria; Proteobacteria; Gammaproteobacteria; Enterobacterales; Morganellaceae | MULTISPECIES: major outer membrane lipoprotein | 52 |
| | WP_046594720.1 | 5.8E-05 | Qc.III | IDQLSSDVQTLSAKVDQLSNDVNAMRSDIQAAKDDAARANQRLDNKVSRVRK- | *Salmonella enterica* | Bacteria; Proteobacteria; Gammaproteobacteria; Enterobacterales; Enterobacteriaceae; Salmonella | major outer membrane lipoprotein | 52 |
|  | WP_103517453.1 | 8.6E-05 | Qc.III | IDQLSSDVQTLNAKADQLSNDVNAMRSDVQAAKDDAARANQRLDNQ------- | *Klebsiella pneumoniae* | Bacteria; Proteobacteria; Gammaproteobacteria; Enterobacterales; Enterobacteriaceae; Klebsiella | major outer membrane lipoprotein | 52 |
| | WP_001082322.1 | 9E-05 | Qc.III | IDQLSSDVQTLSAKVDQLSNDVNAMRSDIQAAKDDAARANQRLDNKVSRVRK- | *Salmonella enterica* | Bacteria; Proteobacteria; Gammaproteobacteria; Enterobacterales; Enterobacteriaceae; Salmonella | hypothetical protein | 52 |
| | WP_016506271.1 | 9E-05 | Qc.III | IDQLSSDVQTLSAKVDQLSNDVNAMRSDIQAAKDDAARANQRLDNKVSRVRK- | *Salmonella enterica* | Bacteria; Proteobacteria; Gammaproteobacteria; Enterobacterales; Enterobacteriaceae; Salmonella | major outer membrane lipoprotein | 52 |
| | WP_076735685.1 | 9E-05 | Qc.III | IDQLSSDVQTLSAKVDQLSNDVNAMRSDIQAAKDDAARANQRLDNKVSRVRK- | *Salmonella enterica* | Bacteria; Proteobacteria; Gammaproteobacteria; Enterobacterales; Enterobacteriaceae; Salmonella | major outer membrane lipoprotein | 52 |
| | WP_016506877.1 | 9.3E-05 | Qc.III | IDQLSSDVQTLNAKVDQLSNDVNAMRSDVQAAKDDAARANQRLDNKVSRVRK- | *Salmonella enterica* | Bacteria; Proteobacteria; Gammaproteobacteria; Enterobacterales; Enterobacteriaceae; Salmonella | major outer membrane lipoprotein | 52 |
|  | WP_010847307.1 | 9.7E-05 | Qc.III.b | ------KVEQLASQVQTLGSKVDQLSNDLSSVRSDVQSAKDEAARANQRLN-- | *Xenorhabdus nematophila* | Bacteria; Proteobacteria; Gammaproteobacteria; Enterobacterales; Morganellaceae; Xenorhabdus | major outer membrane lipoprotein | 52 |
| ▼ 53 | | | | | | | | |
| | KTC91028.1 | 1.1E-05 | Qc.III | --SMSQQVGNVNNNVNALNAQITKLNQMITDLNTQVVKQSEVINVLMERTK-- | *Fluoribacter dumoffii NY 23* | Bacteria; Proteobacteria; Gammaproteobacteria; Legionellales; Legionellaceae; Fluoribacter | Component of the Dot/Icm secretion system | 53 |
| | WP_058477170.1 | 1.4E-05 | Qc.III | --SMSQQVGNVNNNVNALNAQITKLNQMISELNTQVAKQSEVINILMERTK-- | *Legionella steigerwaltii* | Bacteria; Proteobacteria; Gammaproteobacteria; Legionellales; Legionellaceae; Legionella | hypothetical protein | 53 |
|  | WP_028380929.1 | 2.1E-05 | Qc.III | --SMSQQVGNVNNNVNALNAQITKLNQMITDLNNQVVKQSEVINVLIER---- | *Legionella cherrii* | Bacteria; Proteobacteria; Gammaproteobacteria; Legionellales; Legionellaceae; Legionella | hypothetical protein | 53 |
|  | WP_058387387.1 | 2.1E-05 | Qc.III | --SMSQQVGNVNNNVNALNAQITKLNQMITDLNNQVVKQSEVINVLIER---- | *Legionella cherrii* | Bacteria; Proteobacteria; Gammaproteobacteria; Legionellales; Legionellaceae; Legionella | hypothetical protein | 53 |
|  | SIQ65597.1 | 2.8E-05 | Qc.III | --SMNQQVGNVNNNVNALNAQITKLNQMISDLNNQVVKQSEVINVLIER---- | *Fluoribacter gormanii* | Bacteria; Proteobacteria; Gammaproteobacteria; Legionellales; Legionellaceae; Fluoribacter | intracellular multiplication protein IcmG | 53 |
|  | WP_058521434.1 | 3E-05 | Qc.III | --SMNQQVSNINNNVNSLNEQITKLNQVISDLNNQVEKQSEVINILIER---- | *Legionella tucsonensis* | Bacteria; Proteobacteria; Gammaproteobacteria; Legionellales; Legionellaceae; Legionella | hypothetical protein | 53 |
|  | WP_058458694.1 | 5.8E-05 | Qc.III | --SMNQQVSNINNNVNALNAQITKLNQVISDLNNQVVKQSEVINVLIER---- | *Fluoribacter bozemanae* | Bacteria; Proteobacteria; Gammaproteobacteria; Legionellales; Legionellaceae; Fluoribacter | hypothetical protein | 53 |
|  | WP_110141742.1 | 5.9E-05 | Qc.III | --SMNQQVGNVNNNVNALNAQITKLNQMISDLNNQVAKQSEVINVLIER---- | *Legionella sp. km488* | Bacteria; Proteobacteria; Gammaproteobacteria; Legionellales; Legionellaceae; Legionella | type IV secretion protein IcmG | 53 |
| ▼ 54 | | | | | | | | |
| | EKD56163.1 | 1.1E-05 | Qc.III | --QILATLRIIQQESNKMGEDLNVLNKHITNAKNTMDSVNNGYSRLSSKIDN- | *uncultured bacterium* | Bacteria; environmental samples | hypothetical protein ACD_58C00287G0012 | 54 |
|  | WP_039137987.1 | 4.1E-05 | Qc.I | ------------TEFSAFGDILDKTQKKLQEASNVIDKAGIRTRAIEKKLRK- | *Flavihumibacter solisilvae* | Bacteria; Bacteroidetes; Chitinophagia; Chitinophagales; Chitinophagaceae; Flavihumibacter | DNA recombination protein RmuC | 54 |
|  | WP_060544393.1 | 5.7E-05 | Qc.I | TESMSNLTAPLKQNIEQLQQAIQHTNSETAKSTASLSQQLREMS--------- | *Prevotella enoeca* | Bacteria; Bacteroidetes; Bacteroidia; Bacteroidales; Prevotellaceae; Prevotella | DNA recombination protein RmuC | 54 |
|  | PRR86643.1 | 5.9E-05 | Qc.I | -----------KTQFSKFGDILEKTQKKLQEASNTIETASRKSRTIERKLRS- | *Clostridium luticellarii* | Bacteria; Firmicutes; Clostridia; Clostridiales; Clostridiaceae; Clostridium | RmuC family protein | 54 |
|  | WP_106007945.1 | 6.1E-05 | Qc.I | -----------KTQFSKFGDILEKTQKKLQEASNTIETASRKSRTIERKLRS- | *Clostridium luticellarii* | Bacteria; Firmicutes; Clostridia; Clostridiales; Clostridiaceae; Clostridium | DNA recombination protein RmuC | 54 |
| | WP_056157554.1 | 6.9E-05 | Qc.III | --ALADQARDIHRLGRELHERLASMNGHLDQLGRSLNAAVGHYNQTVGSL--- | *Nocardioides* | Bacteria; Actinobacteria; Propionibacteriales; Nocardioidaceae | MULTISPECIES: DNA recombination protein RmuC | 54 |
| | WP_109509389.1 | 9.2E-05 | Qc.III | --ALADQAQEIHRLGVELHERLARMSTHLDQVGRSLNAAVGHYNQAMGSL--- | *Nocardioides sp. YIM ART13* | Bacteria; Actinobacteria; Propionibacteriales; Nocardioidaceae; Nocardioides | DNA recombination protein RmuC | 54 |
| ▼ 55 | | | | | | | | |
| | PAK20948.1 | 1.2E-05 | Qc.III | -KSLGSQINKINVQNVAINSKVETLATKVENLEIKFDNLETKVDNLEIKFDN- | *Mycoplasma agassizii* | Bacteria; Tenericutes; Mollicutes; Mycoplasmataceae; Mycoplasma | hypothetical protein CJJ23_04635 partial | 55 |
| | PAK21388.1 | 3.9E-05 | Qc.III | -KSLGSQINKINVQNVAINSKVETLATKVENIEIKFDNLETKVDNLETKVDNL | *Mycoplasma agassizii* | Bacteria; Tenericutes; Mollicutes; Mycoplasmataceae; Mycoplasma | hypothetical protein CJJ23_02500 partial | 55 |
| | SMC19600.1 | 5.8E-05 | SNAP.c | ---IGSQIDKINTQNVAINSKVDNLETKFDNLENKVDNLEVQVKEVKEQNKEL | *Mycoplasma agassizii* | Bacteria; Tenericutes; Mollicutes; Mycoplasmataceae; Mycoplasma | Uncharacterised protein family (UPF0184) partial | 55 |
| | WP_095279144.1 | 7.5E-05 | SNAP.c | ---IGSQIDKINTQNVAINSKVDNLETKFDNLENKVDNLEVQVKEVKEQNKEL | *Mycoplasma agassizii* | Bacteria; Tenericutes; Mollicutes; Mycoplasmataceae; Mycoplasma | hypothetical protein | 55 |
| ▼ 56 | | | | | | | | |
|  | WP_113896634.1 | 1.2E-05 | SNAP.c | ---ISTLTRDVHYLSQEMEDVLSNTNTLLDDINHKSEQLDPAVKA-------- | *Lactobacillus reuteri* | Bacteria; Firmicutes; Bacilli; Lactobacillales; Lactobacillaceae; Lactobacillus | DUF948 domain-containing protein | 56 |
|  | WP_076461248.1 | 1.8E-05 | SNAP.c | ---ISTLTRDVHYLSQEMEDVLSNTNTLLDDINHKSEQLDPAIK--------- | *Lactobacillus sp. Marseille-P3519* | Bacteria; Firmicutes; Bacilli; Lactobacillales; Lactobacillaceae; Lactobacillus | DUF948 domain-containing protein | 56 |
|  | AWD62013.1 | 2.2E-05 | SNAP.c | -----TLTRDVHYLSQEMEDVLSNTNTLLDDINHKSEQLDPAVKA-------- | *Lactobacillus reuteri* | Bacteria; Firmicutes; Bacilli; Lactobacillales; Lactobacillaceae; Lactobacillus | chemotaxis protein | 56 |
|  | PUH36169.1 | 2.2E-05 | SNAP.c | -----TLTRDVHYLSQEMEDVLSNTNTLLDDINHKSEQLDPAVKA-------- | *Lactobacillus reuteri* | Bacteria; Firmicutes; Bacilli; Lactobacillales; Lactobacillaceae; Lactobacillus | DUF948 domain-containing protein | 56 |
|  | PWT61843.1 | 2.2E-05 | SNAP.c | -----TLTRDVHYLSQEMEDVLSNTNTLLDDINHKSEQLDPAVKA-------- | *Lactobacillus reuteri* | Bacteria; Firmicutes; Bacilli; Lactobacillales; Lactobacillaceae; Lactobacillus | DUF948 domain-containing protein | 56 |
|  | WP_003666701.1 | 2.2E-05 | SNAP.c | -----TLTRDVHYLSQEMEDVLSNTNTLLDDINHKSEQLDPAVKA-------- | *Lactobacillus* | Bacteria; Firmicutes; Bacilli; Lactobacillales; Lactobacillaceae | MULTISPECIES: DUF948 domain-containing protein | 56 |
|  | WP_003671904.1 | 2.2E-05 | SNAP.c | -----TLTRDVHYLSQEMEDVLSNTNTLLDDINHKSEQLDPAVKA-------- | *Lactobacillus* | Bacteria; Firmicutes; Bacilli; Lactobacillales; Lactobacillaceae | MULTISPECIES: DUF948 domain-containing protein | 56 |
|  | WP_078009179.1 | 2.2E-05 | SNAP.c | -----TLTRDVHYLSQEMEDVLSNTNTLLDDINHKSEQLDPAVKA-------- | *Lactobacillus reuteri* | Bacteria; Firmicutes; Bacilli; Lactobacillales; Lactobacillaceae; Lactobacillus | DUF948 domain-containing protein | 56 |
|  | WP_085650402.1 | 2.2E-05 | SNAP.c | -----TLTRDVHYLSQEMEDVLSNTNTLLDDINHKSEQLDPAVKA-------- | *Lactobacillus reuteri* | Bacteria; Firmicutes; Bacilli; Lactobacillales; Lactobacillaceae; Lactobacillus | DUF948 domain-containing protein | 56 |
|  | WP_086142166.1 | 2.2E-05 | SNAP.c | -----TLTRDVHYLSQEMEDVLSNTNTLLDDINHKSEQLDPAVKA-------- | *Lactobacillus reuteri* | Bacteria; Firmicutes; Bacilli; Lactobacillales; Lactobacillaceae; Lactobacillus | DUF948 domain-containing protein | 56 |
|  | WP_094508208.1 | 2.2E-05 | SNAP.c | -----TLTRDVHYLSQEMEDVLSNTNTLLDDINHKSEQLDPAVKA-------- | *Lactobacillus reuteri* | Bacteria; Firmicutes; Bacilli; Lactobacillales; Lactobacillaceae; Lactobacillus | DUF948 domain-containing protein | 56 |
|  | OXE59513.1 | 2.5E-05 | SNAP.c | -----TLTRDIHYLSQEMEDVLSNTNTLLDDINHKSEQLDPAVKA-------- | *Lactobacillus reuteri* | Bacteria; Firmicutes; Bacilli; Lactobacillales; Lactobacillaceae; Lactobacillus | chemotaxis protein | 56 |
| ▼ 57 | | | | | | | | |
|  | OUT53718.1 | 1.3E-05 | SNAP.c | ------QLAQIHIQLRSLGNRLAATNRKLDRLIDLMEKNNDQTTRSREILK-- | *Muricauda sp. TMED12* | Bacteria; Bacteroidetes; Flavobacteriia; Flavobacteriales; Flavobacteriaceae; Muricauda | hypothetical protein CBB72_11515 | 57 |
|  | WP_072882363.1 | 1.6E-05 | SNAP.c | ------QLAQIHIQLRSLGNRLASTNRKLDRLIDLMEKNNDQTTRSREILK-- | *Muricauda antarctica* | Bacteria; Bacteroidetes; Flavobacteriia; Flavobacteriales; Flavobacteriaceae; Muricauda | hypothetical protein | 57 |
| ▼ 58 | | | | | | | | |
|  | WP_004437096.1 | 1.4E-05 | Qc.III | LDKMEEMLSTLISMVGNINDKQNQMVEKLDRVENRLDKVEERLDKL------- | *Bacillus methanolicus* | Bacteria; Firmicutes; Bacilli; Bacillales; Bacillaceae; Bacillus | hypothetical protein | 58 |
|  | AIE60429.1 | 4.9E-05 | Qc.III | ---LISMVGNINDKQNQMGEKLDRVENRLDKLEKRLDKLEDGQK--------- | *Bacillus methanolicus MGA3* | Bacteria; Firmicutes; Bacilli; Bacillales; Bacillaceae; Bacillus | hypothetical protein BMMGA3_10165 | 58 |
| ▼ 59 | | | | | | | | |
|  | OHC11588.1 | 1.4E-05 | SNAP.b | -NKMINDITLQKEEAEKIAVSLKGQDEQLRLVNKKLEEAKAKL---------- | *Planctomycetes bacterium RIFOXYB12_FULL_42_10* | Bacteria; Planctomycetes | hypothetical protein A3K25_01370 | 59 |

| | | | | | | | | |
|---|---|---|---|---|---|---|---|---|
|  | OHB40696.1 | 3.1E-05 | SNAP.b | --KMISDITRQKEDAEKVASSVREQGEQLKFVNKKLEDAKAKL---------- | *Planctomycetes bacterium GWB2_41_19* | Bacteria; Planctomycetes | hypothetical protein A2069_05495 | 59 |
|  | OHB46684.1 | 3.1E-05 | SNAP.b | --KMISDITRQKEDAEKVASSVREQGEQLKFVNKKLEDAKAKL---------- | *Planctomycetes bacterium GWE2_41_14* | Bacteria; Planctomycetes | hypothetical protein A2094_04105 | 59 |
|  | OHC07644.1 | 3.1E-05 | SNAP.b | --KMISDITRQKEDAEKVASSVREQGEQLKFVNKKLEDAKAKL---------- | *Planctomycetes bacterium RIFOXYD2_FULL_41_16* | Bacteria; Planctomycetes | hypothetical protein A2545_03330 | 59 |
|  | KXK25642.1 | 4.6E-05 | SNAP.b | LNRMVNDITRQKEEAERIAASFKEQGEQLKEVNNKIEDTKT------------ | *Candidatus Brocadia sinica* | Bacteria; Planctomycetes; Planctomycetia; Candidatus Brocadiales; Candidatus Brocadiaceae; Candidatus Brocadia | hypothetical protein UZ01_03226 | 59 |
|  | WP_052565382.1 | 4.6E-05 | SNAP.b | LNRMVNDITRQKEEAERIAASFKEQGEQLKEVNNKIEDTKT------------ | *Candidatus Brocadia sinica* | Bacteria; Planctomycetes; Planctomycetia; Candidatus Brocadiales; Candidatus Brocadiaceae; Candidatus Brocadia | hypothetical protein | 59 |
| ▼ 60 | | | | | | | | |
|  | WP_044873633.1 | 1.4E-05 | Qa.IV | IKQLQARVAELAKQQQNTNTQVGSQGKRVDQLAADVKTAQ------------- | *Pseudomonas sp. LFM046* | Bacteria; Proteobacteria; Gammaproteobacteria; Pseudomonadales; Pseudomonadaceae; Pseudomonas | ATPase | 60 |
|  | WP_028628704.1 | 0.0001 | Qa.IV | IKQLQAKVVELSKQQQSTNAQLGSQGKRVDQLAADVKSAQ------------- | *Pseudomonas resinovorans* | Bacteria; Proteobacteria; Gammaproteobacteria; Pseudomonadales; Pseudomonadaceae; Pseudomonas | ATPase | 60 |
| ▼ 61 | | | | | | | | |
|  | AGN00875.1 | 1.7E-05 | Qc.III | ---TAKKVQKLADTAETLYEQIKDLQRRMVNLEESVDESTARLDRL------- | *Salinarchaeum sp. Harcht-Bsk1* | Archaea; Euryarchaeota; Halobacteria; Natrialbales; Natrialbaceae; Salinarchaeum | hypothetical protein L593_04630 | 61 |
|  | ACV48116.1 | 7.5E-05 | Qc.III | --STAKKVQKLADLAEKMYKRINHMVEQLQDLRGTVDETGARVEEIER----- | *Halomicrobium mukohataei DSM 12286* | Archaea; Euryarchaeota; Halobacteria; Halobacteriales; Haloarculaceae; Halomicrobium | conserved hypothetical protein | 61 |
|  | WP_018259042.1 | 7.5E-05 | Qc.III | --STAKKVQKLADLAEKMYKRINHMVEQLQDLRGTVDETGARVEEIER----- | *Halomicrobium katesii* | Archaea; Euryarchaeota; Halobacteria; Halobacteriales; Haloarculaceae; Halomicrobium | hypothetical protein | 61 |
| ▼ 62 | | | | | | | | |
| | WP_043726358.1 | 1.7E-05 | Qc.I | --ELTLRARALEARATASEQALDEQGEVLNELAALTRKTLDGIGRIDRRAET- | *Nocardia asiatica* | Bacteria; Actinobacteria; Corynebacteriales; Nocardiaceae; Nocardia | hypothetical protein | 62 |
| | WP_039798387.1 | 3.8E-05 | Qc.I | --ELTLRARALEARATASEQALDEQGEVLNELAALTRKALDGIGRIDRRAET- | *Nocardia araoensis* | Bacteria; Actinobacteria; Corynebacteriales; Nocardiaceae; Nocardia | hypothetical protein | 62 |
| | WP_063018145.1 | 4.3E-05 | Qc.I | --ELTLRARALEARATASEQALDEQGEVLNELAALTRKALDGIGRIDRRAET- | *Nocardia niwae* | Bacteria; Actinobacteria; Corynebacteriales; Nocardiaceae; Nocardia | hypothetical protein | 62 |
| | WP_067807687.1 | 4.6E-05 | Qc.I | --ELTLRARALEARATASEQALDEQGEVLNELAALTRKALDGIGRIDRRAES- | *Nocardia beijingensis* | Bacteria; Actinobacteria; Corynebacteriales; Nocardiaceae; Nocardia | hypothetical protein | 62 |
| | WP_014986140.1 | 5E-05 | Qc.I | --ELTLRARALEARATASEQALDEQGDVLNELAALTRKALDGIHRIDERAER- | *Nocardia* | Bacteria; Actinobacteria; Corynebacteriales; Nocardiaceae | MULTISPECIES: hypothetical protein | 62 |
|  | WP_040867150.1 | 5.3E-05 | Qc.I | -------ARALEARATASEQALDEQGEVLNELAALTRKALDGIGRIDRRAET- | *Nocardia exalbida* | Bacteria; Actinobacteria; Corynebacteriales; Nocardiaceae; Nocardia | hypothetical protein | 62 |
|  | WP_062972838.1 | 5.9E-05 | Qc.I | -------ARALEARATASEQALDEQGEVLNELAALTRKALDGIGRIDRRAET- | *Nocardia gamkensis* | Bacteria; Actinobacteria; Corynebacteriales; Nocardiaceae; Nocardia | hypothetical protein | 62 |
| ▼ 63 | | | | | | | | |
| | WP_067515281.1 | 1.7E-05 | Qb.I | --DISNLAHSTLSGFEHTSEMMDGSHKMLDDMSEHLETINTRLDETNKLV--- | *Endozoicomonas ascidiicola* | Bacteria; Proteobacteria; Gammaproteobacteria; Oceanospirillales; Hahellaceae; Endozoicomonas | hypothetical protein | 63 |
|  | WP_101747893.1 | 4E-05 | SNAP.b | ---THERVDETYERLNETNERLGENSERLDEANERLDENNERL---------- | *Endozoicomonas acroporae* | Bacteria; Proteobacteria; Gammaproteobacteria; Oceanospirillales; Endozoicomonaceae; Endozoicomonas | hypothetical protein | 63 |
|  | WP_066013969.1 | 4.9E-05 | SNAP.b | ---THERVDETYERLNETNERLGENNERLDEANERLDENNERL---------- | *Endozoicomonas atrinae* | Bacteria; Proteobacteria; Gammaproteobacteria; Oceanospirillales; Hahellaceae; Endozoicomonas | hypothetical protein | 63 |
|  | WP_020581923.1 | 6.3E-05 | SNAP.b | ---THERVDETYERLNETNERLGENNERLDEANERLDENNERL---------- | *Endozoicomonas elysicola* | Bacteria; Proteobacteria; Gammaproteobacteria; Oceanospirillales; Hahellaceae; Endozoicomonas | hypothetical protein | 63 |
| ▼ 64 | | | | | | | | |
| | WP_037349910.1 | 1.8E-05 | SNAP.b | --RASQMLSRAKKKLARASQMLSRARKKLARASQMLSRAKKKLARASRMLS-- | *Saprospira grandis* | Bacteria; Bacteroidetes; Saprospiria; Saprospirales; Saprospiraceae; Saprospira | hypothetical protein | 64 |
| | WP_015692742.1 | 5.4E-05 | SNAP.b | --RARKELARANQTLSRARKKLARANQTLSRARKKLARANQTLSRARRILSR- | *Saprospira grandis* | Bacteria; Bacteroidetes; Saprospiria; Saprospirales; Saprospiraceae; Saprospira | hypothetical protein | 64 |
| | WP_015692742.1 | 5.6E-05 | SNAP.b | --RARKKLAQARRILSRARKELARANQTLSRARKKLARANQTLSRARKEL--- | *Saprospira grandis* | Bacteria; Bacteroidetes; Saprospiria; Saprospirales; Saprospiraceae; Saprospira | hypothetical protein | 64 |
| ▼ 65 | | | | | | | | |
|  | WP_067931873.1 | 1.8E-05 | Qc.III | --------NQLRSQADQVSSEIARQRAKANAIQSKIDDYQAHIDSMNQQI--- | *Alicyclobacillus kakegawensis* | Bacteria; Firmicutes; Bacilli; Bacillales; Alicyclobacillaceae; Alicyclobacillus | hypothetical protein | 65 |
|  | CRK80255.1 | 3.4E-05 | Qc.III | --KLDNQIIIGMDKAQKLNDEIKKQQGQIDKTKVEIDKAQKDF---------- | *Bacillus sp. LF1* | Bacteria; Firmicutes; Bacilli; Bacillales; Bacillaceae; Bacillus | NLP/P60 protein | 65 |
|  | WP_043434369.1 | 3.6E-05 | Qb.III | LQQLYRQAEQATETYDATAEELKRQRAKVTRLDGELSRARLALQ--------- | *Streptomyces pluripotens* | Bacteria; Actinobacteria; Streptomycetales; Streptomycetaceae; Streptomyces | hypothetical protein | 65 |
|  | WP_067924474.1 | 4.4E-05 | Qc.III | --------NKLQSQADQVSSEIARQRAKANAIQSKIDDYQAHIDSMNQQI--- | *Alicyclobacillus shizuokensis* | Bacteria; Firmicutes; Bacilli; Bacillales; Alicyclobacillaceae; Alicyclobacillus | hypothetical protein | 65 |
| ▼ 66 | | | | | | | | |
| | WP_017417019.1 | 2.1E-05 | Qc.III | --ELVSGVNQLADGSVALNDGLKELNGKIPELTDGTQKLNDGSKELAEKLN-- | *Clostridium tunisiense* | Bacteria; Firmicutes; Clostridia; Clostridiales; Clostridiaceae; Clostridium | YhgE/Pip domain-containing protein | 66 |
|  | EFV35642.1 | 7.3E-05 | Qc.II | INTLENDLTKVKEVNNKIYTNIDNYDEIADSLKNDFADVTKRI---------- | *Gemella morbillorum M424* | Bacteria; Firmicutes; Bacilli; Bacillales; Bacillales Family XI. Incertae Sedis; Gemella | YhgE/Pip domain-containing protein | 66 |
| ▼ 67 | | | | | | | | |
| | GAF23256.1 | 2.1E-05 | SNAP.b | ---LLRKLTEVEIALAKVTQQLESQTSVLEEVRKELSMTREKADAAEEKA--- | *Bacillus sp. JCM 19047* | Bacteria; Firmicutes; Bacilli; Bacillales; Bacillaceae; Bacillus | hypothetical protein JCM19047_3062 | 67 |
| | WP_055737291.1 | 4.3E-05 | SNAP.b | ---LLRKLTEVEIALAKVTQQLESQTSVLEEVRKELSMTREKADAAEEKA--- | *Bacillus* | Bacteria; Firmicutes; Bacilli; Bacillales; Bacillaceae | MULTISPECIES: hypothetical protein | 67 |
| ▼ 68 | | | | | | | | |
|  | WP_004415526.1 | 2.2E-05 | Qa.IV | ----------LLNLSTNLAKRVKGQQEAIDLVSEAIKRAKAKINDPNRPI--- | *Mycoplasma arginini* | Bacteria; Tenericutes; Mollicutes; Mycoplasmataceae; Mycoplasma | AAA family ATPase | 68 |
|  | WP_060823481.1 | 2.2E-05 | Qa.IV | ----------LLNLSTNLAKRVKGQQEAIDLVSEAIKRAKAKINDPNRPI--- | *Mycoplasma arginini* | Bacteria; Tenericutes; Mollicutes; Mycoplasmataceae; Mycoplasma | AAA family ATPase | 68 |
|  | WP_109246977.1 | 2.2E-05 | Qa.IV | ----------LLNLSTNLAKRVKGQQEAIDLVSEAIKRAKAKINDPNRPI--- | *Mycoplasma arginini* | Bacteria; Tenericutes; Mollicutes; Mycoplasmataceae; Mycoplasma | AAA family ATPase | 68 |
|  | PIR13100.1 | 8.9E-05 | Qa.IV | ----------LKNLESNLNKEIISQGQAIAKISSTIQRAKLGLRNANRPL--- | *Candidatus Falkowbacteria bacterium CG11_big_fil_rev_8_21_14_0_20_39_10* | Bacteria; Candidatus Falkowbacteria | hypothetical protein COV49_03300 | 68 |
| ▼ 69 | | | | | | | | |
|  | WP_090713500.1 | 2.3E-05 | Qc.I | -EEMLQLVSSIKQQSQNTKTALQEEIAVLREQVAGLEKTTMLL---------- | *Paenibacillus typhae* | Bacteria; Firmicutes; Bacilli; Bacillales; Paenibacillaceae; Paenibacillus | hypothetical protein | 69 |
|  | WP_082707847.1 | 3.3E-05 | Qc.I | -EEMLQLVSSIKQQSQSTKAALQEEIALLREQVAGLEKTTMLL---------- | *Paenibacillus sp. DMB5* | Bacteria; Firmicutes; Bacilli; Bacillales; Paenibacillaceae; Paenibacillus | hypothetical protein | 69 |
| ▼ 70 | | | | | | | | |
|  | KXI22960.1 | 2.4E-05 | Qc.II | -----QDIKKLKGIAEYIRIDAQVQKVLVDGYSDQLGKFSSKMRISRQRA--- | *Photobacterium sanguinicancri* | Bacteria; Proteobacteria; Gammaproteobacteria; Vibrionales; Vibrionaceae; Photobacterium | hypothetical protein AS132_10555 | 70 |
|  | PSW22395.1 | 2.4E-05 | Qc.II | -----QDINKLKGIAEYIRVDAQVQKVLVDGYSDQLGQFSSKMRVSRQRA--- | *Photobacterium swingsii* | Bacteria; Proteobacteria; Gammaproteobacteria; Vibrionales; Vibrionaceae; Photobacterium | OmpA family protein | 70 |
|  | WP_083540894.1 | 2.5E-05 | Qc.II | -----QDIKKLKGIAEYIRIDAQVQKVLVDGYSDQLGKFSSKMRISRQRA--- | *Photobacterium sanguinicancri* | Bacteria; Proteobacteria; Gammaproteobacteria; Vibrionales; Vibrionaceae; Photobacterium | OmpA family protein | 70 |
| ▼ 71 | | | | | | | | |
|  | SDB97410.1 | 2.6E-05 | SNAP.c | --------DPLKKKAIRMEKEISQTSDRIDTLRKKMKKNESKLKYHQKQ---- | *Melghirimyces thermohalophilus* | Bacteria; Firmicutes; Bacilli; Bacillales; Thermoactinomycetaceae; Melghirimyces | CHAP domain-containing protein | 71 |
|  | WP_091565665.1 | 3.5E-05 | SNAP.c | --------DPLKKKAIRMEKEISQTSDRIDTLRKKMKKNESKLKYHQKQ---- | *Melghirimyces thermohalophilus* | Bacteria; Firmicutes; Bacilli; Bacillales; Thermoactinomycetaceae; Melghirimyces | CHAP domain-containing protein | 71 |
| ▼ 72 | | | | | | | | |
|  | AFK62016.1 | 2.8E-05 | Qb.II | ------QLESVTQQRNALQSQLESQTETLNKAQGQIKDLTTNLALIKKVI--- | *Advenella kashmirensis WT001* | Bacteria; Proteobacteria; Betaproteobacteria; Burkholderiales; Alcaligenaceae | hypothetical protein TKWG_08105 | 72 |
|  | WP_041709116.1 | 3.3E-05 | Qb.II | ------QLESVTQQRNALQSQLESQTETLNKAQGQIKDLTTNLALIKKVI--- | *Advenella kashmirensis* | Bacteria; Proteobacteria; Betaproteobacteria; Burkholderiales; Alcaligenaceae | hypothetical protein | 72 |
| ▼ 73 | | | | | | | | |
|  | WP_042452523.1 | 2.9E-05 | Qb.III | -----ARIHELEQIAAELRSDLDSERRRADTMTTRTEELQDELTAARASLKRM | *Rhodococcus erythropolis* | Bacteria; Actinobacteria; Corynebacteriales; Nocardiaceae; Rhodococcus | hypothetical protein | 73 |
|  | WP_047269065.1 | 2.9E-05 | Qb.III | -----ARIHELEQIAAELRSDLDSERRRADTMTTRTEELQDELTAARASLKRM | *Rhodococcus* | Bacteria; Actinobacteria; Corynebacteriales; Nocardiaceae | MULTISPECIES: hypothetical protein | 73 |
|  | WP_073511952.1 | 2.9E-05 | Qb.III | -----ARIHELEQIAAELRSDLDSERRRADTMTTRTEELQDELTAARASLKRM | *Rhodococcus* | Bacteria; Actinobacteria; Corynebacteriales; Nocardiaceae | MULTISPECIES: hypothetical protein | 73 |
|  | WP_064445081.1 | 3E-05 | Qb.III | -----ARIHELEQIAAELRSDLDSERRRADTMTTRTEELQDELTAARASLKRM | *Rhodococcus* | Bacteria; Actinobacteria; Corynebacteriales; Nocardiaceae | MULTISPECIES: hypothetical protein | 73 |
| ▼ 74 | | | | | | | | |
|  | SDZ29583.1 | 2.9E-05 | Qb.I | -----------LNQFEQMKSELEESKNKLQAVKSECDSYKDELLQSKREIEK- | *Proteiniborus ethanoligenes* | Bacteria; Firmicutes; Clostridia; Clostridiales; Proteiniborus | cell division protein ZapA | 74 |
|  | WP_091732019.1 | 3.2E-05 | Qb.I | -----------LNQFEQMKSELEESKNKLQAVKSECDSYKDELLQSKREIEK- | *Proteiniborus ethanoligenes* | Bacteria; Firmicutes; Clostridia; Clostridiales; Proteiniborus | cell division protein ZapA | 74 |
| ▼ 75 | | | | | | | | |
| | PJY31902.1 | 2.9E-05 | Qc.III | --QIKANITNLNRATNTVEEELTRMTGLLEKTAEKVSHSQSSIRNVKSSV--- | *Escherichia coli* | Bacteria; Proteobacteria; Gammaproteobacteria; Enterobacterales; Enterobacteriaceae; Escherichia | methyl-accepting chemotaxis protein partial | 75 |
| | CSC39943.1 | 3.3E-05 | Qc.III | --QIKANITNLNRATNTVEEELTRMTGLLEKTAEKVSHSQSSIRNVKSSV--- | *Vibrio cholerae* | Bacteria; Proteobacteria; Gammaproteobacteria; Vibrionales; Vibrionaceae; Vibrio | methyl-accepting chemotaxis protein | 75 |
| | CSB43113.1 | 3.9E-05 | Qc.III | --QIKANITNLNRATNTVEEELTRMTGLLEKTAEKVSHSQSSIRNVKSSV--- | *Vibrio cholerae* | Bacteria; Proteobacteria; Gammaproteobacteria; Vibrionales; Vibrionaceae; Vibrio | methyl-accepting chemotaxis protein | 75 |
| ▼ 76 | | | | | | | | |
|  | WP_076387522.1 | 3E-05 | Qb.I | ----NQQIAKQITQSAAQIKQLEKSYSLLKDAQEKYQKVNGYIQQ-------- | *Chryseobacterium chaponense* | Bacteria; Bacteroidetes; Flavobacteriia; Flavobacteriales; Flavobacteriaceae; Chryseobacterium | hypothetical protein | 76 |

| | | | | | | | | |
|---|---|---|---|---|---|---|---|---|
|  | **SIS89960.1** | 5.2E-05 | Qb.I | ----NQQIAKQITQSAAQIKQLEKSYSLLKDAQEKYQKVNGYIQQ-------- | *Chryseobacterium chaponense* | Bacteria; Bacteroidetes; Flavobacteriia; Flavobacteriales; Flavobacteriaceae; Chryseobacterium | hypothetical protein SAMN05421789_11067 | 76 |
| ▼ 77 | | | | | | | | |
|  | **WP_051195220.1** | 3E-05 | SNAP.b | -------AVETAKQGADTAKnnALQYQNEQIPKAEAKADKIVQNATAS------- | *Pseudobutyrivibrio ruminis* | Bacteria; Firmicutes; Clostridia; Clostridiales; Lachnospiraceae; Pseudobutyrivibrio | FtsH protease activity modulator HflK | 77 |
|  | **WP_090544720.1** | 3E-05 | SNAP.b | -------AVETAKQGADTAKnnALQYQNEQIPKAEAKADKIVQNATAS------- | *Pseudobutyrivibrio sp. OR37* | Bacteria; Firmicutes; Clostridia; Clostridiales; Lachnospiraceae; Pseudobutyrivibrio | FtsH protease activity modulator HflK | 77 |
| ▼ 78 | | | | | | | | |
|  | **OHA10970.1** | 3.1E-05 | R.IV | LEKLKNEMADLKTTVRRSVADRIESAKSLGDLSENAEYQEAKD---------- | *Candidatus Sungbacteria bacterium RIFCSPLOWO2_02_FULL_48_13b* | Bacteria; Candidatus Sungbacteria | transcription elongation factor GreA | 78 |
|  | **PSO45776.1** | 3.7E-05 | R.IV | --ELKEELDHLKNTRRQEIAEEIERTKALGDLSENAEYQQARADQ-------- | *Parcubacteria group bacterium SW_6_46_9* | Bacteria; unclassified Parcubacteria group | transcription elongation factor GreA | 78 |
| ▼ 79 | | | | | | | | |
| | **OJW62167.1** | 3.3E-05 | Qc.I | ---IADLDRQMREEASKISRSLENDARIANARVDGLNASLDQLKRQATSSN-- | *Afipia sp. 64-13* | Bacteria; Proteobacteria; Alphaproteobacteria; Rhizobiales; Bradyrhizobiaceae; Afipia | lipopolysaccharide biosynthesis protein | 79 |
|  | **WP_034471277.1** | 7.8E-05 | Qc.I | --------RQLRDEAAKLSRSLENDARIAGARAESMGANLDQLKKQAVSTNT- | *Afipia sp. P52-10* | Bacteria; Proteobacteria; Alphaproteobacteria; Rhizobiales; Bradyrhizobiaceae; Afipia | LPS biosynthesis protein | 79 |
| ▼ 80 | | | | | | | | |
|  | **WP_054652631.1** | 3.7E-05 | SNAP.b | -----QLINQVKQTADEATQTLVDQGKAINEVKSTASQLRVDLQTA------- | *Lactobacillus equigenerosi* | Bacteria; Firmicutes; Bacilli; Lactobacillales; Lactobacillaceae; Lactobacillus | hypothetical protein | 80 |
|  | **KRL91844.1** | 4E-05 | SNAP.b | -----QLINQVKQTADEATQTLVDQGKAINEVKSTASQLRVDLQTA------- | *Lactobacillus equigenerosi DSM 18793 = JCM 14505* | Bacteria; Firmicutes; Bacilli; Lactobacillales; Lactobacillaceae; Lactobacillus | hypothetical protein FC21_GL000614 | 80 |
|  | **WP_061209664.1** | 7.2E-05 | SNAP.b | -----QLINQVKQTADEATQTLVDQGKAINEVKSTASQLRVDLQTA------- | *Lactobacillus equigenerosi* | Bacteria; Firmicutes; Bacilli; Lactobacillales; Lactobacillaceae; Lactobacillus | hypothetical protein | 80 |
| ▼ 81 | | | | | | | | |
|  | **WP_033020884.1** | 3.9E-05 | Qc.III | -DSLHQQVQTLNEQTNMMNKRMEDMERRMEEMDKRIGEMDKRIE--------- | *Geobacillus* | Bacteria; Firmicutes; Bacilli; Bacillales; Bacillaceae | MULTISPECIES: hypothetical protein | 81 |
|  | **RAN23443.1** | 4.5E-05 | Qc.III | -DSLQRQVQALSEQTNTMNKRIEDMDKRIEDMDKRMEGMNGRMK--------- | *Geobacillus sp. A8* | Bacteria; Firmicutes; Bacilli; Bacillales; Bacillaceae; Geobacillus | hypothetical protein VC88_06625 | 81 |
|  | **KYD29028.1** | 4.7E-05 | Qc.III | -DSLHQQVQTLNEQTNMMNKRMEDMERRMEEMDKRIGEMDKRIEE-------- | *Geobacillus sp. B4113_201601* | Bacteria; Firmicutes; Bacilli; Bacillales; Bacillaceae; Geobacillus | hypothetical protein B4113_2651 | 81 |
| ▼ 82 | | | | | | | | |
|  | **WP_113467932.1** | 4.5E-05 | Qc.I | ----------LKQGAQHYAAALERSGEALIRTaAQQLERQASDLERAGRNLAA- | *Rhizobiales bacterium* | Bacteria; Proteobacteria; Alphaproteobacteria; Rhizobiales | hypothetical protein partial | 82 |
|  | **WP_099782323.1** | 4.9E-05 | Qc.I | ----------LKQGAQHYAAVLERSGEALIRTaAQQLERQASDLERAGRNLSAH | *Roseomonas sp. FDAARGOS_362* | Bacteria; Proteobacteria; Alphaproteobacteria; Rhodospirillales; Acetobacteraceae; Roseomonas | hypothetical protein | 82 |
|  | **WP_094538648.1** | 6E-05 | Qc.I | ----------LKQGAQHYAAALERSGEALIRTaAQQLERQASDLERAGRNLAA- | *Ochrobactrum grignonense* | Bacteria; Proteobacteria; Alphaproteobacteria; Rhizobiales; Brucellaceae; Ochrobactrum | hypothetical protein | 82 |
| ▼ 83 | | | | | | | | |
|  | **PBV74855.1** | 4.7E-05 | Qc.III | --QVGEAVEAIAEHSRDLGEQLGEQKKQTQQLASQLDKVTKEL---------- | *Pseudomonas aeruginosa* | Bacteria; Proteobacteria; Gammaproteobacteria; Pseudomonadales; Pseudomonadaceae; Pseudomonas | phage capsid protein partial | 83 |
|  | **OPE06917.1** | 5.4E-05 | Qc.III | --QVGEAVEAIAEHSRDLGEQLGEQKKQTQQLASQLDKVTKEL---------- | *Pseudomonas aeruginosa* | Bacteria; Proteobacteria; Gammaproteobacteria; Pseudomonadales; Pseudomonadaceae; Pseudomonas | phage capsid protein partial | 83 |
|  | **WP_003098374.1** | 6.4E-05 | Qc.III | --QVGEAVEAIAEHSRDLGEQLGEQKKQTQQLASQVDKVTKEL---------- | *Pseudomonas aeruginosa* | Bacteria; Proteobacteria; Gammaproteobacteria; Pseudomonadales; Pseudomonadaceae; Pseudomonas | phage capsid scaffolding protein | 83 |
| ▼ 84 | | | | | | | | |
|  | **WP_050928686.1** | 4.8E-05 | Qb.I | -QDIRTMLQQLAQASKQANETLAQTNRLLRNANGLMDGQGKQL---------- | *Yersinia enterocolitica* | Bacteria; Proteobacteria; Gammaproteobacteria; Enterobacterales; Yersiniaceae; Yersinia | MCE family protein | 84 |
|  | **WP_076707491.1** | 4.8E-05 | Qb.I | -QDIRTMLQQLAQASKQANETLAQTNRLLRNANGLMDGQGKQL---------- | *Yersinia enterocolitica* | Bacteria; Proteobacteria; Gammaproteobacteria; Enterobacterales; Yersiniaceae; Yersinia | MCE family protein | 84 |
|  | **WP_095632731.1** | 6.7E-05 | Qb.I | --DIRQAMQQLASVGKQASATLEQTTALMRNANGLLNDQGKHMFGS------- | *Pseudomonas sp. PICF141* | Bacteria; Proteobacteria; Gammaproteobacteria; Pseudomonadales; Pseudomonadaceae; Pseudomonas | MCE family protein | 84 |
| ▼ 85 | | | | | | | | |
|  | **WP_074265673.1** | 5E-05 | Qb.III.b | ---------NALEQGDSLNKTLRDQRESISSLEGNIAEARTALKSAHRLLATM | *Paraburkholderia phenazinium* | Bacteria; Proteobacteria; Betaproteobacteria; Burkholderiales; Burkholderiaceae; Paraburkholderia | hypothetical protein | 85 |
|  | **SIO25896.1** | 5.7E-05 | Qb.III.b | ---------NALEQGDSLNKTLRDQRESISSLEGNIAEARTALKSAHRLLATM | *Paraburkholderia phenazinium* | Bacteria; Proteobacteria; Betaproteobacteria; Burkholderiales; Burkholderiaceae; Paraburkholderia | hypothetical protein SAMN05444168_3866 | 85 |
| ▼ 86 | | | | | | | | |
|  | **WP_094991345.1** | 5.4E-05 | Qc.III.b | LDRLAQQVEVLKRNAGatdvtALKNTLDSQARTLEKLDREVQ------------- | *Pseudomonas lundensis* | Bacteria; Proteobacteria; Gammaproteobacteria; Pseudomonadales; Pseudomonadaceae; Pseudomonas | hypothetical protein | 86 |
|  | **WP_052893284.1** | 6.4E-05 | Qc.III.b | LDKLAQQVEVLKRNAGatdvtALKNTLDSQARTLEKLDREVQ------------- | *Pseudomonas lundensis* | Bacteria; Proteobacteria; Gammaproteobacteria; Pseudomonadales; Pseudomonadaceae; Pseudomonas | hypothetical protein | 86 |
|  | **WP_052959408.1** | 6.4E-05 | Qc.III.b | LDKLAQQVEVLKRNAGatdvtALKNTLDSQARTLEKLDREVQ------------- | *Pseudomonas lundensis* | Bacteria; Proteobacteria; Gammaproteobacteria; Pseudomonadales; Pseudomonadaceae; Pseudomonas | hypothetical protein | 86 |
|  | **WP_053070052.1** | 6.4E-05 | Qc.III.b | LDKLAQQVEVLKRNAGatdvtALKNTLDSQARTLEKLDREVQ------------- | *Pseudomonas lundensis* | Bacteria; Proteobacteria; Gammaproteobacteria; Pseudomonadales; Pseudomonadaceae; Pseudomonas | hypothetical protein | 86 |
|  | **WP_070412149.1** | 6.4E-05 | Qc.III.b | LDKLAQQVEVLKRNAGatdvtALKNTLDSQARTLEKLDREVQ------------- | *Pseudomonas lundensis* | Bacteria; Proteobacteria; Gammaproteobacteria; Pseudomonadales; Pseudomonadaceae; Pseudomonas | hypothetical protein | 86 |
|  | **WP_095010215.1** | 6.4E-05 | Qc.III.b | LDKLAQQVEVLKRNAGatdvtALKNTLDSQARTLEKLDREVQ------------- | *Pseudomonas lundensis* | Bacteria; Proteobacteria; Gammaproteobacteria; Pseudomonadales; Pseudomonadaceae; Pseudomonas | hypothetical protein | 86 |
|  | **WP_052892180.1** | 9.8E-05 | Qc.III.b | LDKLAQQVEVLKRNAGatdvtALKNTLDSQGRTLEKLDREVQ------------- | *Pseudomonas* | Bacteria; Proteobacteria; Gammaproteobacteria; Pseudomonadales; Pseudomonadaceae | MULTISPECIES: hypothetical protein | 86 |
|  | **WP_052892955.1** | 9.8E-05 | Qc.III.b | LDKLAQQVEVLKRNAGatdvtALKNTLDSQGRTLEKLDREVQ------------- | *Pseudomonas lundensis* | Bacteria; Proteobacteria; Gammaproteobacteria; Pseudomonadales; Pseudomonadaceae; Pseudomonas | hypothetical protein | 86 |
|  | **WP_052959668.1** | 9.8E-05 | Qc.III.b | LDKLAQQVEVLKRNAGatdvtALKNTLDSQGRTLEKLDREVQ------------- | *Pseudomonas lundensis* | Bacteria; Proteobacteria; Gammaproteobacteria; Pseudomonadales; Pseudomonadaceae; Pseudomonas | hypothetical protein | 86 |
|  | **WP_094989907.1** | 9.8E-05 | Qc.III.b | LDKLAQQVEVLKRNAGatdvtALKNTLDSQGRTLEKLDREVQ------------- | *Pseudomonas lundensis* | Bacteria; Proteobacteria; Gammaproteobacteria; Pseudomonadales; Pseudomonadaceae; Pseudomonas | hypothetical protein | 86 |
|  | **WP_097191820.1** | 9.8E-05 | Qc.III.b | LDKLAQQVEVLKRNAGatdvtALKNTLDSQGRTLEKLDREVQ------------- | *Pseudomonas lundensis* | Bacteria; Proteobacteria; Gammaproteobacteria; Pseudomonadales; Pseudomonadaceae; Pseudomonas | hypothetical protein | 86 |
| ▼ 87 | | | | | | | | |
|  | **WP_042403592.1** | 5.7E-05 | Qc.III | -------VDQFKRITDATNRAAENQEKLLTRIADTTDSNTAAIRELTK----- | *Geomicrobium sp. JCM 19037* | Bacteria; Firmicutes; Bacilli; Bacillales; Geomicrobium | hypothetical protein | 87 |
|  | **GAK05654.1** | 6.6E-05 | Qc.III | -------VDQFKRITDATNRAAENQEKLLTRIADTTDSNTAAIRELTK----- | *Geomicrobium sp. JCM 19037* | Bacteria; Firmicutes; Bacilli; Bacillales; Geomicrobium | hypothetical protein JCM19037_4173 | 87 |
| ▼ 88 | | | | | | | | |
| | **PHS66078.1** | 5.8E-05 | Qb.III.d | --RTANLIATSSKVEAASAGDFRNQLEVIaQNITDTADKIRAHLSQARRLLG-- | *Thalassobium sp.* | Bacteria; Proteobacteria; Alphaproteobacteria; Rhodobacterales; Rhodobacteraceae; Thalassobium | hypothetical protein COB09_01015 | 88 |
| | **PIQ41161.1** | 5.8E-05 | Qb.III.d | --RTANLIATSSKVEAASAGDFRNQLEVIaQNITDTADKIRAHLSQARRLLG-- | *Thalassolituus sp. CG17_big_fil_post_rev_8_21_14_2_50_53_8* | Bacteria; Proteobacteria; Gammaproteobacteria; Oceanospirillales; Thalassolituus | hypothetical protein COW58_02595 | 88 |
| | **KZZ11945.1** | 8.1E-05 | Qb.III.d | --RTANLIATSSHVEAATAGDFREQLEVIaDNIRKTADRIAAHLNRANSLL--- | *Oleibacter sp. HI0075* | Bacteria; Proteobacteria; Gammaproteobacteria; Oceanospirillales; Oleibacter | hypothetical protein A3746_25960 partial | 88 |
| | **KZY96496.1** | 9.9E-05 | Qb.III.d | --RTANLIATSSHVEAATAGDFREQLEVIaDNIRKTADRIAAHLNRANSLL--- | *Oleibacter sp. HI0075* | Bacteria; Proteobacteria; Gammaproteobacteria; Oceanospirillales; Oleibacter | hypothetical protein A3746_10485 | 88 |
| | **OUX65553.1** | 9.9E-05 | Qb.III.d | --RTANLIATSSHVEAATAGDFREQLEVIaDNIRKTADRIAAHLNRANSLL--- | *Oceanospirillaceae bacterium TMED276* | Bacteria; Proteobacteria; Gammaproteobacteria; Oceanospirillales; unclassified Oceanospirillaceae | hypothetical protein CBE36_04870 | 88 |
| | **WP_076513501.1** | 9.9E-05 | Qb.III.d | --RTANLIATSSHVEAATAGDFREQLEVIaDNIRKTADRIAAHLNRANSLL--- | *Oleibacter marinus* | Bacteria; Proteobacteria; Gammaproteobacteria; Oceanospirillales; Oleibacter | hypothetical protein | 88 |
| ▼ 89 | | | | | | | | |
|  | **AKG12185.1** | 5.8E-05 | SNAP.c | --------EKSKAMGTKFANEIKQANAKIDEMSKKMQKTEQEVQKAKR----- | *Moraxella bovoculi* | Bacteria; Proteobacteria; Gammaproteobacteria; Pseudomonadales; Moraxellaceae; Moraxella | hypothetical protein AAX07_09610 | 89 |
|  | **AKG14155.1** | 5.8E-05 | SNAP.c | --------EKSKAMGTKFANEIKQANAKIDEMSKKMQKTEQEVQKAKR----- | *Moraxella bovoculi* | Bacteria; Proteobacteria; Gammaproteobacteria; Pseudomonadales; Moraxellaceae; Moraxella | hypothetical protein AAX11_09155 | 89 |
| ▼ 90 | | | | | | | | |
|  | **KKW08935.1** | 6.6E-05 | Qc.I | -----EIAAQLKEISQSINGHLQKNKEAIDSILANMDVTVKNMSSITKNVDE- | *Candidatus Kaiserbacteria bacterium GW2011_GWA2_49_19* | Bacteria; Candidatus Kaiserbacteria | Mammalian cell entry related domain protein | 90 |
|  | **OGX21724.1** | 6.6E-05 | Qc.I | -----EIAAQLKEISQSINGHLQKNKEAIDSILANMDVTVKNMSSITKNVDE- | *Omnitrophica WOR_2 bacterium GWC2_45_7* | Bacteria; Candidatus Omnitrophica | hypothetical protein A2Y04_06220 | 90 |
| ▼ 91 | | | | | | | | |
|  | **WP_010784602.1** | 6.7E-05 | Qa.IV | -------VSSLEDLTSELSTTVTESTEAFKQLASDLTNTVESVGTAKTEMKKA | *Enterococcus faecalis* | Bacteria; Firmicutes; Bacilli; Lactobacillales; Enterococcaceae; Enterococcus | phage minor structural protein | 91 |
|  | **WP_010784776.1** | 6.7E-05 | Qa.IV | -------VSSLEDLTSELSTTVTESTEAFKQLASDLTNTVESVGTAKTEMKKA | *Enterococcus faecalis* | Bacteria; Firmicutes; Bacilli; Lactobacillales; Enterococcaceae; Enterococcus | phage minor structural protein | 91 |
|  | **WP_033624668.1** | 6.7E-05 | Qa.IV | -------VSSLEDLTSELSTTVTESTEAFKQLASDLTNTVESVGTAKTEMKKA | *Enterococcus faecalis* | Bacteria; Firmicutes; Bacilli; Lactobacillales; Enterococcaceae; Enterococcus | peptidase M23 | 91 |
| ▼ 92 | | | | | | | | |
|  | **WP_084646689.1** | 7.8E-05 | SNAP.b | -------ARDASAQARAVADMMAEQGQALKQQRRKAGELALDLEVAQSQVKGL | *Mesorhizobium sp. WSM2561* | Bacteria; Proteobacteria; Alphaproteobacteria; Rhizobiales; Phyllobacteriaceae; Mesorhizobium | hypothetical protein | 92 |
|  | **WP_095495056.1** | 8.7E-05 | SNAP.b | -------ARDASAQARAVADTMAEQGQALNQQRRRAEELALDLEVAQREVKGL | *Mesorhizobium temperatum* | Bacteria; Proteobacteria; Alphaproteobacteria; Rhizobiales; Phyllobacteriaceae; Mesorhizobium | hypothetical protein | 92 |

| | | | | | | | | |
|---|---|---|---|---|---|---|---|---|
| ▼ 93 | | | | | | | | |
|  | **WP_051583229.1** | 7.9E-05 | SNAP.c | -------------QVILMQSRLDQLDGRISRMAGDIDGLTDALSTLRDRISRL | *Sphingomonas sp. RIT328* | Bacteria; Proteobacteria; Alphaproteobacteria; Sphingomonadales; Sphingomonadaceae; Sphingomonas | hypothetical protein | 93 |
|  | **EZP56152.1** | 8.5E-05 | SNAP.c | -------------QVILMQSRLDQLDGRISRMAGDIDGLTDALSTLRDRISRL | *Sphingomonas sp. RIT328* | Bacteria; Proteobacteria; Alphaproteobacteria; Sphingomonadales; Sphingomonadaceae; Sphingomonas | hypothetical protein BW41_00796 | 93 |
| ▼ 94 | | | | | | | | |
|  | **SFP99017.1** | 9E-05 | Qc.III.c | --ALGKLAKRFGDVAKTLGKTAGRHGKALRQIEGKVDDWNNK----------- | *Yuhushiella deserti* | Bacteria; Actinobacteria; Pseudonocardiales; Pseudonocardiaceae; Yuhushiella | hypothetical protein SAMN05421810_104123 | 94 |
|  | **WP_092530518.1** | 9E-05 | Qc.III.c | --ALGKLAKRFGDVAKTLGKTAGRHGKALRQIEGKVDDWNNK----------- | *Yuhushiella deserti* | Bacteria; Actinobacteria; Pseudonocardiales; Pseudonocardiaceae; Yuhushiella | hypothetical protein | 94 |
| ▼ 95 | | | | | | | | |
| | **KRM93311.1** | 9.4E-05 | Qc.III | LEGVASDTKNIGTNAKLLQTELEQMQTKLMAAGNGIDTVNDGLGQTNQYLKG- | *Lactobacillus senioris DSM 24302 = JCM 17472* | Bacteria; Firmicutes; Bacilli; Lactobacillales; Lactobacillaceae; Lactobacillus | hypothetical protein FC56_GL000977 | 95 |
| | **WP_082620611.1** | 9.7E-05 | Qc.III | LEGVASDTKNIGTNAKLLQTELEQMQTKLMAAGNGIDTVNDGLGQTNQYLKG- | *Lactobacillus senioris* | Bacteria; Firmicutes; Bacilli; Lactobacillales; Lactobacillaceae; Lactobacillus | MMPL family transporter | 95 |
| ▼ 96 | | | | | | | | |
|  | **GBE75732.1** | 0.0001 | Qa.III | -----------NLAVSDLKNTIQQQQDGITQIKERAISTVESLEKTNDKLR-- | *Microcystis aeruginosa NIES-87* | Bacteria; Cyanobacteria; Oscillatoriophycideae; Chroococcales; Microcystaceae; Microcystis | hypothetical protein myaer87_29590 | 96 |
|  | **GBL10172.1** | 0.0001 | Qa.III | -----------NLAVSDLKNTIQQQQDGITQIKERAISTVESLEKTNDKLR-- | *Microcystis aeruginosa Sj* | Bacteria; Cyanobacteria; Oscillatoriophycideae; Chroococcales; Microcystaceae; Microcystis | hypothetical protein MSj_01656 | 96 |
|  | **WP_002796633.1** | 0.0001 | Qa.III | -----------NLAVSDLKNTIQQQQDGITQIKERAISTVESLEKTNDKLR-- | *Microcystis aeruginosa* | Bacteria; Cyanobacteria; Oscillatoriophycideae; Chroococcales; Microcystaceae; Microcystis | hypothetical protein | 96 |
|  | **WP_002800886.1** | 0.0001 | Qa.III | -----------NLAVSDLKNTIQQQQDGITQIKERAISTVESLEKTNDKLR-- | *Microcystis aeruginosa* | Bacteria; Cyanobacteria; Oscillatoriophycideae; Chroococcales; Microcystaceae; Microcystis | hypothetical protein | 96 |
|  | **WP_008202910.1** | 0.0001 | Qa.III | -----------NLAVSDLKNTIQQQQDGITQIKERAISTVESLEKTNDKLR-- | *Microcystis sp. T1-4* | Bacteria; Cyanobacteria; Oscillatoriophycideae; Chroococcales; Microcystaceae; Microcystis | hypothetical protein | 96 |
|  | **WP_108936541.1** | 0.0001 | Qa.III | -----------NLAVSDLKNTIQQQQDGITQIKERAISTVESLEKTNDKLR-- | *Microcystis sp. 0824* | Bacteria; Cyanobacteria; Oscillatoriophycideae; Chroococcales; Microcystaceae; Microcystis | hypothetical protein | 96 |
